# A multi-scale *in silico* mouse model for insulin resistance and humanoid type 2 diabetes

**DOI:** 10.1101/2021.05.19.443124

**Authors:** Christian Simonsson, William Lövfors, Niclas Bergqvist, Elin Nyman, Peter Gennemark, Karin G Stenkula, Gunnar Cedersund

**Affiliations:** Department of Biomedical Engineering, Linköping University, Linköping, Sweden; Center for Medicine Imaging and Visualization Science (CMIV), Linköping University, Linköping, Sweden; Department of Mathematics, Linköping University, Linköping, Sweden; Drug Metabolism and Pharmacokinetics, Research and Early Development, Cardiovascular, Renal and Metabolism (CVRM), BioPharmaceuticals R&D, AstraZeneca, Gothenburg, Sweden; Department of Experimental Medical Science, Lund University, Lund, Sweden

## Abstract

Insulin resistance (IR) causes compensatory insulin production, which in humans eventually progresses to beta-cell failure and type 2 diabetes (T2D). This disease progression involves multi-scale processes, ranging from intracellular signaling to organ-organ and whole-body level regulations, on timescales from minutes to years. T2D progression is commonly studied using overfed and genetically modified rodents. However, rodents do not exhibit human T2D progression, with IR-driven beta-cell failure, and available multi-scale data is too complex to fully comprehend using traditional analysis. To help resolve these issues, we here present an *in silico* mouse model. This is the first mathematical model that simultaneously explains multi-scale mouse IR data on all three levels – cells, organs, body – ranging from minutes to months. The model correctly predicts new independent multi-scale validation data and provides insights into non-measured processes. Finally, we present a humanoid *in silico* mouse exhibiting disease progression from IR to IR-driven T2D.

## Introduction

The increasing prevalence of obesity is one of the major health risks in society today. Obesity with metabolic dysfunctions, such as decreased insulin sensitivity, may lead to onset of diabetes mellitus type 2 (T2D) and cardiovascular diseases. Therefore, it is of importance to understand the underlying mechanisms of how increased fat mass contributes to development of insulin resistance and T2D. This understanding is made more complicated by the fact that obesity and insulin resistance induce metabolic changes in different organs due to crosstalk signals, resulting in both short and long-term effects.

Murine animal models are commonly used to study obesity-related aspects of insulin resistance and T2D, through both genetic and diet-induced perturbations, such as high-fat diet (HFD)-feeding (1). These murine models allow for investigations of multi-scale mechanisms, i.e., mechanisms at several different timescales and body levels (Fig. 1A). The top body level is the *whole-body level* which is characterized by different biomarkers such as bodyweight and fat tissue mass. The next level, the *tissue and organ level*, covers the interplay between different tissues such as muscle and fat tissue, which primarily interact via the circulatory system, i.e., blood and lymph. One such interplay is the relationship between glucose and insulin levels. Finally, the *cellular level* includes cellular aspects, such as intracellular signaling, gene expression, and energy metabolism. The dynamics of these biochemical processes at the three levels occur on timescales which differ by orders of magnitude. The slower dynamics often occurs at the whole-body and tissue levels and can be monitored *in vivo*. More rapid dynamic events primarily occur at the cellular level, and can often only be captured through *in vitro* experiments, utilizing isolated tissue, at various time points.

**Figure 1:**
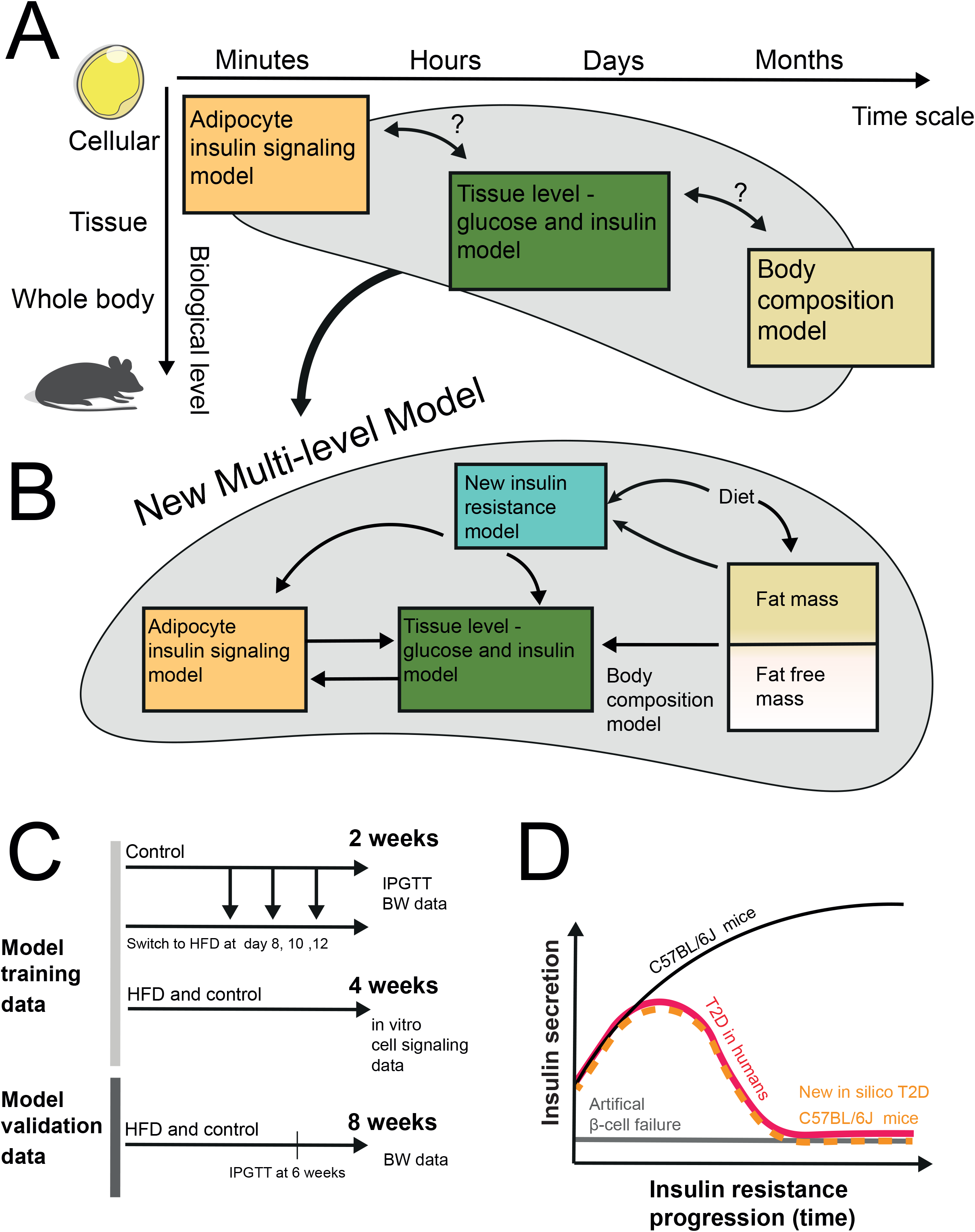
**A)** Systems operate over multiple body levels and timescales. Current mathematical descriptions lack connections between all three levels. **B)** A schematic over the multi-level and multi-scale model structure that would connect multiple body levels and timescales **C)** Diet schedules used to generate training and validation data in C57BL/6 mice. The training data set included body weight data as well as Intraperitoneal Glucose Tolerance Test (IPGTT) data from a two week dietary intervention, in which mice ate different durations of HFD (2). The training data set also included data from in vitro experiments on primary adipocytes taken from mice after a four-week HFD intervention (20). The validation data set includes new unpublished body weight data after an eight-week HFD intervention, with IPGTT data from the six-week mark. **D)** HFD induces insulin resistance in C57BL/6 mice causing an increase in insulin secretion (black line). In contrast to humans (red line), mice never develop diet induced T2D as the increased insulin production is not followed by a failure of the beta cells. Instead, beta-cell failure is artificially induced in T2D in vivo mouse models (grey line). The proposed humanoid in silico mouse develops diet induced T2D (orange line).

These three body levels have been experimentally examined using different murine models to examine the etiology of insulin resistance and T2D, where each model has its own strengths and weaknesses. While these murine models can describe different aspects of T2D progression, none of them captures the human T2D etiology. In short, the human etiology commonly starts with peripheral insulin resistance (in e.g., muscle and adipose tissue) which leads to compensatory hyperinsulinemia and an increased demand on the beta cells, which ultimately leads to pancreatic failure and manifested T2D. We have previously used C57BL/6J mice to study the short-term progression of insulin resistance using different feeding schedules combining chow diet and HFD (Fig. 1C) (2). In this mouse model, HFD-feeding induces insulin resistance, which in turn triggers a compensatory increase in insulin production (3)(Fig. 1D, black). A shortcoming of this mouse strain as a model for T2D is that the increased insulin production is not followed by beta-cell failure (4). In other words, these mice only become pre-diabetic, i.e. insulin resistant, but they do not develop T2D (Fig. 1D, red) (5, 6). Other mice models have been developed to study the diabetic hypo-insulinemic phenotype, either derived by genetic modification (1, 7-9) or by chemical induction (10). Still, in these models, the low insulin production is not a consequence of the high insulin demand but is low already before the development of insulin resistance (Fig. 1D, grey). Thus, murine T2D etiology differs from the etiology in humans. There also exists efforts to implant human beta cells into mice (11), but these experiments are expensive, and still do not fully describe the common human T2D etiology. Nevertheless, in all murine models, it is possible to measure parameters at all three levels: whole-body parameters such as weight and body composition, organ-level parameters such as plasma glucose and insulin responses to a glucose tolerance test (GTT), and intracellular responses studied *in vitro* using tissue biopsies. To interpret these complex multi-level processes, all such data must be integrated into a cohesive framework. The arguably most powerful and rigorous way to do that interpretation is to use systems biology and mathematical modelling, also called *in silico* modelling.

In general, system biology mathematical modelling is based on two inputs: 1) experimental data, usually in the form of dose-response or time-course data, and 2) mathematical models that represent different mechanistic hypotheses serving as tentative *explanations* of the data (12, 13). The output of the analysis can come to one of two conclusions: I) the data cannot be explained by a specific hypothesis, which thereby should be rejected; or II) the hypothesis is a tentatively acceptable explanation to the currently existing data, and there are some associated new predictions, which can be used e.g., to guide new experiments, or to provide further insights regarding the way in which the mechanisms work. This type of mechanistic mathematical modeling has been used extensively to study different aspects of the insulin resistance and T2D. For instance, obesity- and weight-related mathematical models exist for rodents (14-16). These top-level and long-term models describe weight changes but do not propagate these effects to other metabolic parameters, such as insulin resistance. Instead, insulin resistance is commonly described in mathematical models at an exclusively organ and tissue level, dissecting interactions during a meal-response (measuring e.g., glucose and insulin levels) (17). Finally, for the cellular level, there are several in silico murine models describing e.g. insulin signaling in adipocytes (18), and regulation of insulin secretion in beta cells (19). However, these models are also single-level models, i.e., the models have not been connected to corresponding mathematical models for the other body levels.

In summary, while important advancements regarding the understanding of the progression of insulin resistance have been made, both experimentally and modelling wise, a more integrative view is still missing. To the best of our knowledge, there exists no multi-level and multi-timescale *in silico* model that integrates all three levels of insulin resistance: body composition, circulating glucose and insulin, and intracellular insulin signaling. Similarly, on the experimental side, no murine model fully captures the human diet-induced progression of T2D, where beta-cell failure is caused by increasing insulin resistance (Fig. 1D). In this work, we aim to overcome these two shortcomings, by integrating multi-level and multi-timescale murine data and modelling in a new framework: a multi-scale *in silico* mouse model.

Herein, we present a first multi-level, multi-timescale, and mechanistic mathematical model of insulin resistance development in mice. The model connects three different levels and models (Fig. 1B): whole-body weight and body-composition alterations (Guo *et al*. model (14)), meal-response glucose and insulin dynamics (Alskär *et al*. model (17)) and adipocyte intracellular insulin signaling (Bergqvist *et al*. model (18)). The three levels are connected by a new model for long- and short-term changes in insulin resistance. The *in silico* model was trained on our own previously published multi-level data (data from several body levels) from C57BL/6J mice (Fig. 1C, training data)(2, 20). Validation of the model predictions was done by collecting new experimental data generated for a longer feeding protocol (Fig. 1C, validation data). The potential of our validated *in silico* model is illustrated by creating a humanoid version, which can simulate the same processes on all three levels, and which – unlike any currently existing *in vivo* mouse - has the human-like progression of diet-induced T2D, caused by insulin resistance-induced failure of the pancreas (Fig. 1D, orange).

## Results

### We have assembled a mechanistic, multi-level, and multi-timescale model

Our multi-level model is comprised of three interconnected, previously published, dynamic models which are linked together using a new sub-model for insulin resistance (Fig. 2A). The whole-body model (Fig. 2B) describes body-composition changes (14), the tissue-level model (Fig. 2C) describes organ specific glucose uptake and insulin regulation (17), and the cellular level (Fig. 2D) describes intracellular insulin signaling. The body and organ levels are trained and validated on data from mice undergoing various HFD schemes, and the intracellular model is based on data from primary rodent adipocytes, taken from rodents undergoing HFD schemes (18). The new insulin resistance model is assumed to be driven by two known risk factors for insulin resistance: adiposity and diet (21). These two factors are quantified by the time spent on HFD (t_HFD_), and by the relative increase in fat mass (FM) above baseline (FM_0_), i.e. (FM/FM_0_). The variable t_HFD_ is a model variable which counts the number of days spent on an HFD, and FM comes from the whole-body level.

**Figure 2:**
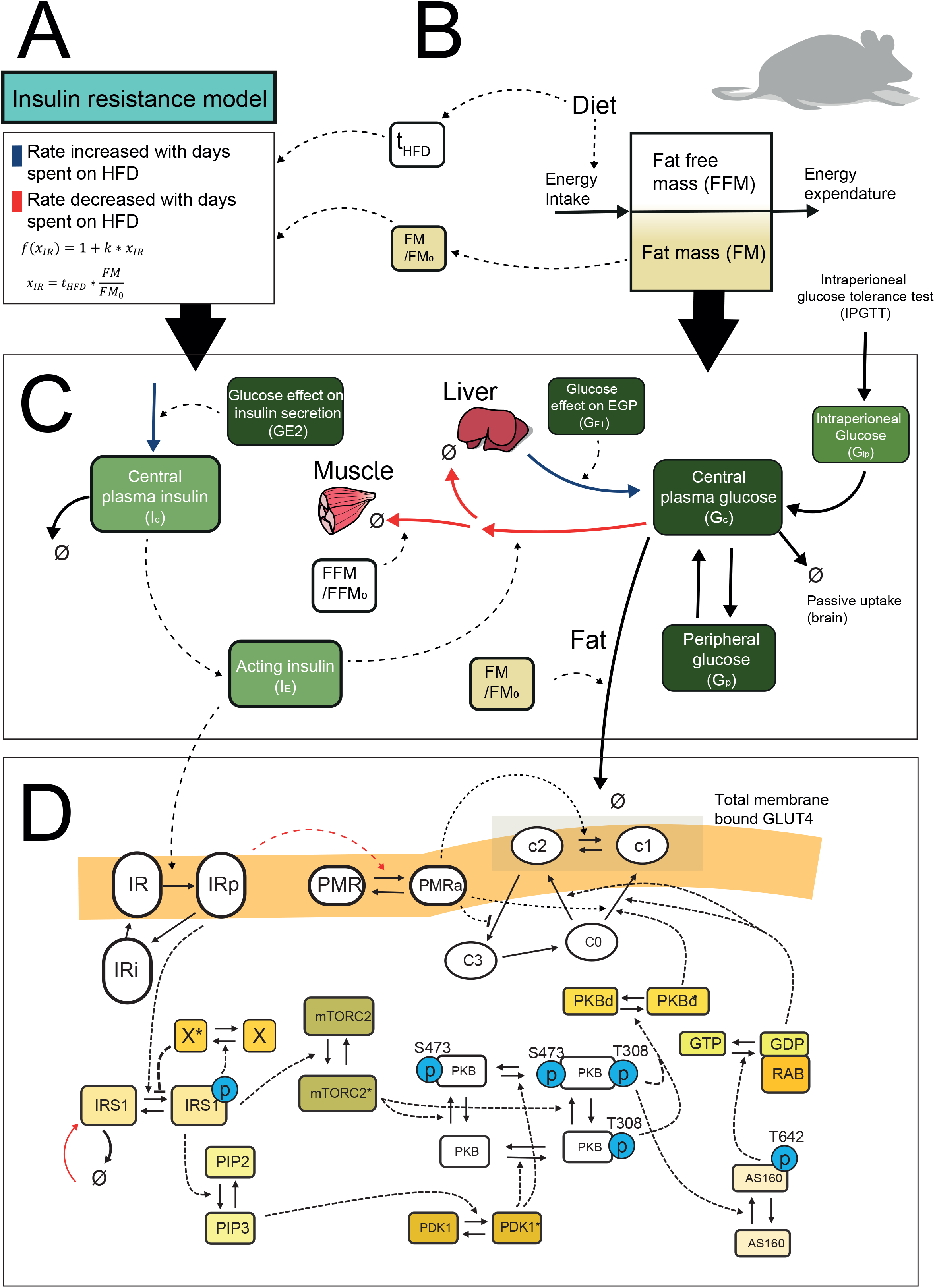
Multi-level and multi-timescale model structure featuring all model reactions and dependencies. **A)** Insulin resistance sub-model, and explanation of the color coding of the arrows. **B)** The structure of the body-level, body-weight composition model. **C)** The structure of the organ-level glucose and insulin model. **D)** The structure of the cell-level, adipocyte intracellular insulin signaling model.

The previously existing models on the three levels have been altered to different degrees (Fig. 2B-D). The body-level weight-regulation model provides input to the two lower levels, but is itself not altered by those levels. Note, however, that one of the parameters in the body-level model was re-optimized (physical activity, λ) to fit the experimental conditions in our studies. The tissue-level model was altered in a few ways. First, the glucose utilization was sub-divided into the contribution of the three insulin-responding organs: adipose, muscle, and liver (Supplementary Appendix, SA, Eq. 15,16,59). Second, when FM or fat-free mass (FFM) grows, the corresponding glucose uptakes (in fat and muscle, respectively) were scaled linearly with the increase in tissue (SA Eq. 59,15). Third, insulin resistance appears in three places in the tissue level model: at the glucose utilization, for endogenous glucose production (EGP), and for insulin secretion (SA Eq. 15,10,20). Fourth, the last alteration of the tissue-level model was that the glucose uptake in the adipose tissue (SA Eq. 59) was replaced by an intracellular adipose insulin signaling model (SA Eq. 24-58). Finally, the cellular level model was also altered in a few ways. First, we introduced a degradation and production of the insulin receptor substrate (IRS), to be able to describe long-term changes (SA Eq. 27). Second, the mechanistic target of rapamycin complex 2 (mTORC2) (SA Eq. 31) is now located downstream of IRS instead of directly downstream the insulin receptor (Fig. 2D). Third, insulin resistance impacts the signaling in two different ways: 1) the activity of the plasma membrane-located pathway (PMR) is reduced (SA Eq. 48), and 2) the production of IRS decreases with increasing insulin resistance (SA Eq. 27). The PMR pathway describes the insulin-induced effect of plasma membrane activity associated with changes in the rate of glucose transporters moving from monomer to cluster form (18). Also, at the tissue level, the parameters were refitted to our training data, with a few additional constraints to preserve realistic behaviors, and with the constraint of keeping the parameter values close to the original values. For the cellular level model, only the parameter governing the insulin receptor activation (*ik1*, SA Eq. 25) was re-optimized, to scale the input strength to our experiments. All model equations, and parameter values, with their biological explanations, are included in the appendix (SA).

### Our multi-level model was simultaneously fitted and tested to data from all three levels

We have merged new and previously generated data to a combined dataset obtained from our laboratory, which measures the processes occurring on all three levels. More specifically, we have induced different degrees of body weight and fat mass changes, and insulin resistance in mice, using various combinations of chow and HFD (Fig. 1C, Materials and methods). During these experiments, we measured body weight, fat mass, and performed an Intraperitoneal Glucose Tolerance Test (IPGTT) to measure insulin and time-varying glucose clearance. In parallel experiments, primary adipocytes were isolated to study insulin signaling. Details of these experiments are described in Materials and methods.

While the model was iteratively developed via numerous tests and modifications in its different parts, in the end, the model could describe all the data with one and the same simulation. In other words, while we below present simulations and data comparisons for each level one-by-one, the simulations come from the same run with the model, which provides an acceptable agreement with all the data. This is confirmed by a χ^2^ test (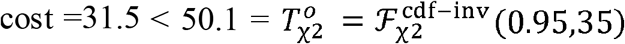, Materials and methods), which statistically confirms that the model should not be rejected. The parameter set producing the simulation with the best agreement to data is presented (Fig 3A-F, 4A-E, 5A, grey line). In practice, the model was estimated to some of the data. The remaining data was used in a subsequent validation step and these validation data comes from new experiments performed by us.

**Figure 3:**
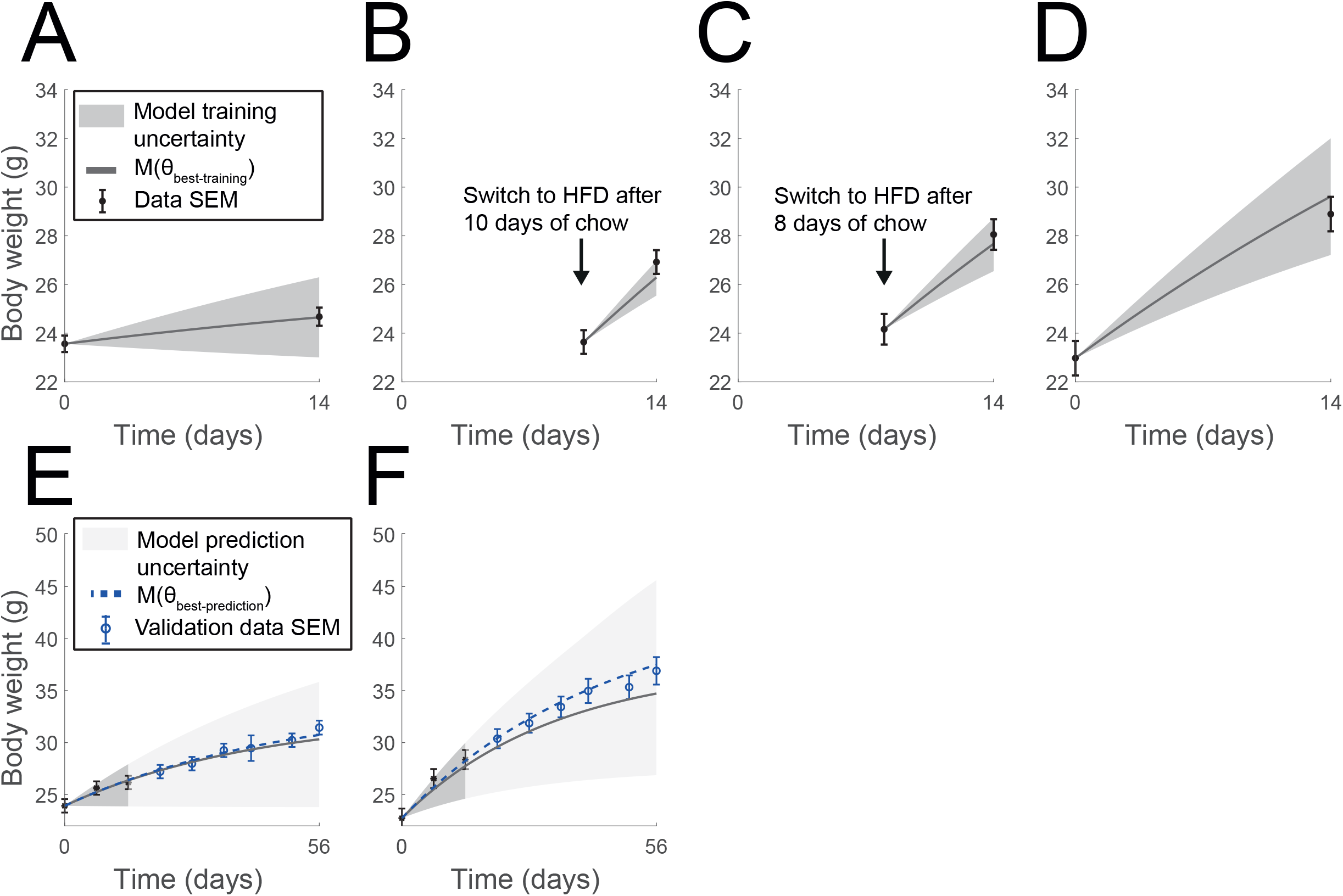
Result of model training and validation for body weight data in the whole-body level model. Model uncertainty is shown as the dark grey area and the dark grey line is the simulation yielding the best agreement to all training data (black error bars). **A)** two weeks of chow diet. **B)** mice following a switch diet with the first ten days being fed chow and then fed HFD for four days. **C)** mice following a switch diet with the first eight days being fed chow and then fed HFD for six days. **D)** two weeks of HFD. **E)** Eight weeks of chow diet where the data during the first two weeks (black error bars) was used for model training, and the rest (blue error bars) was used for model validation. The model prediction uncertainty is the light grey shaded area and the simulation which yielded the best agreement to the validation data can be seen as the dashed blue line. **F)** Eight weeks of HFD where the data during the first two weeks (black error bars) was used for model training, and the rest (blue error bars) was used for model validation.

**Figure 4:**
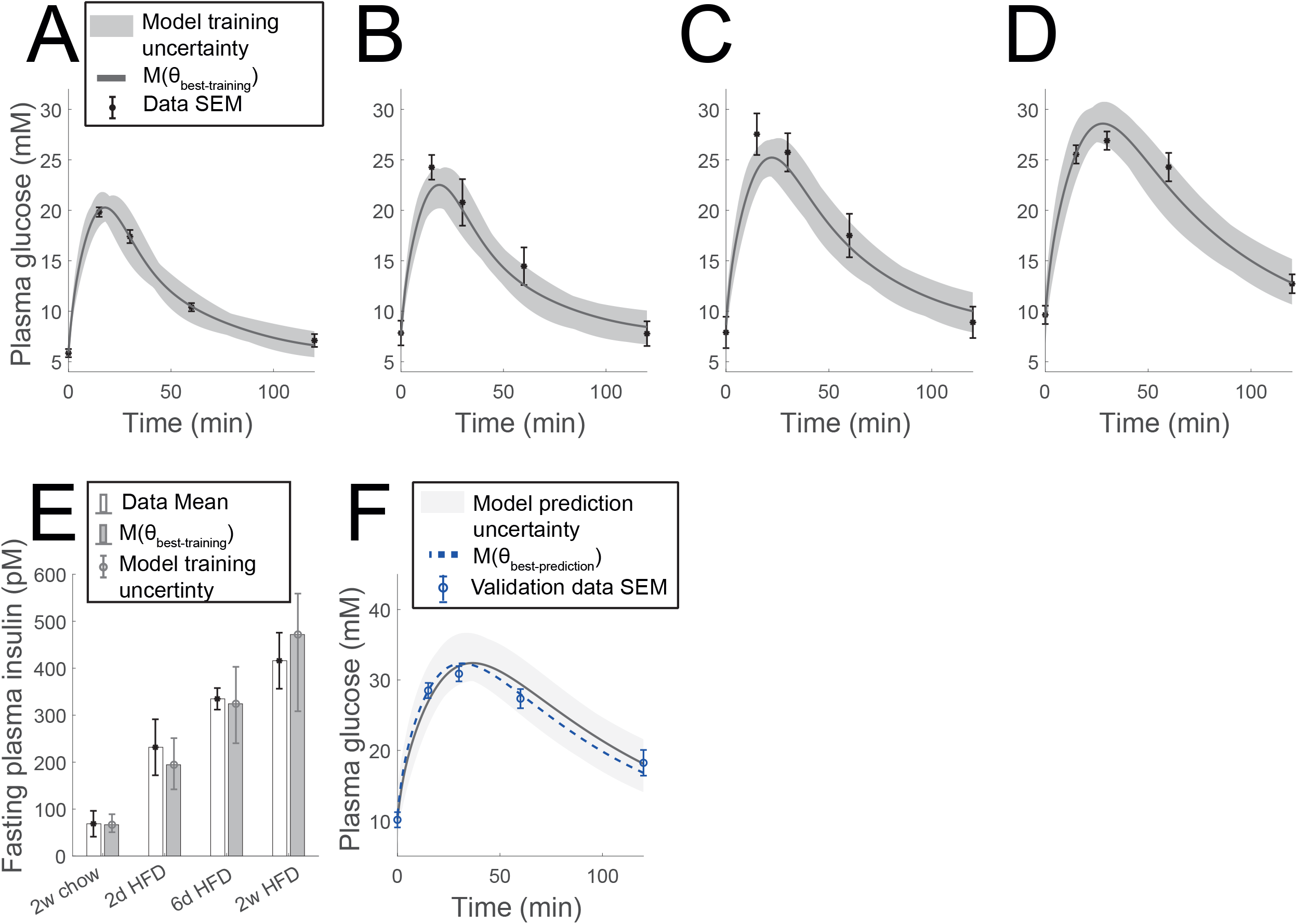
Result of model training and validation for IPGTT data and model training to fasting insulin in the tissue-level model. Model uncertainty is shown as the dark grey area and the dark grey line is the simulation yielding the best agreement to all training data (black error bars). **A)** IPGTT after two weeks of chow diet. **B)** IPGTT after 12 days chow diet and HFD for two days **C)** IPGTT after eight days chow diet and HFD for six days. **D)** IPGTT after two weeks of HFD. **E)** Model prediction of validation IPGTT data after 6 weeks of HFD (blue error bars). Light grey area shows the prediction uncertainty and the simulation which yielded the best agreement to the validation data can be seen as the dashed blue line.

**Figure 5:**
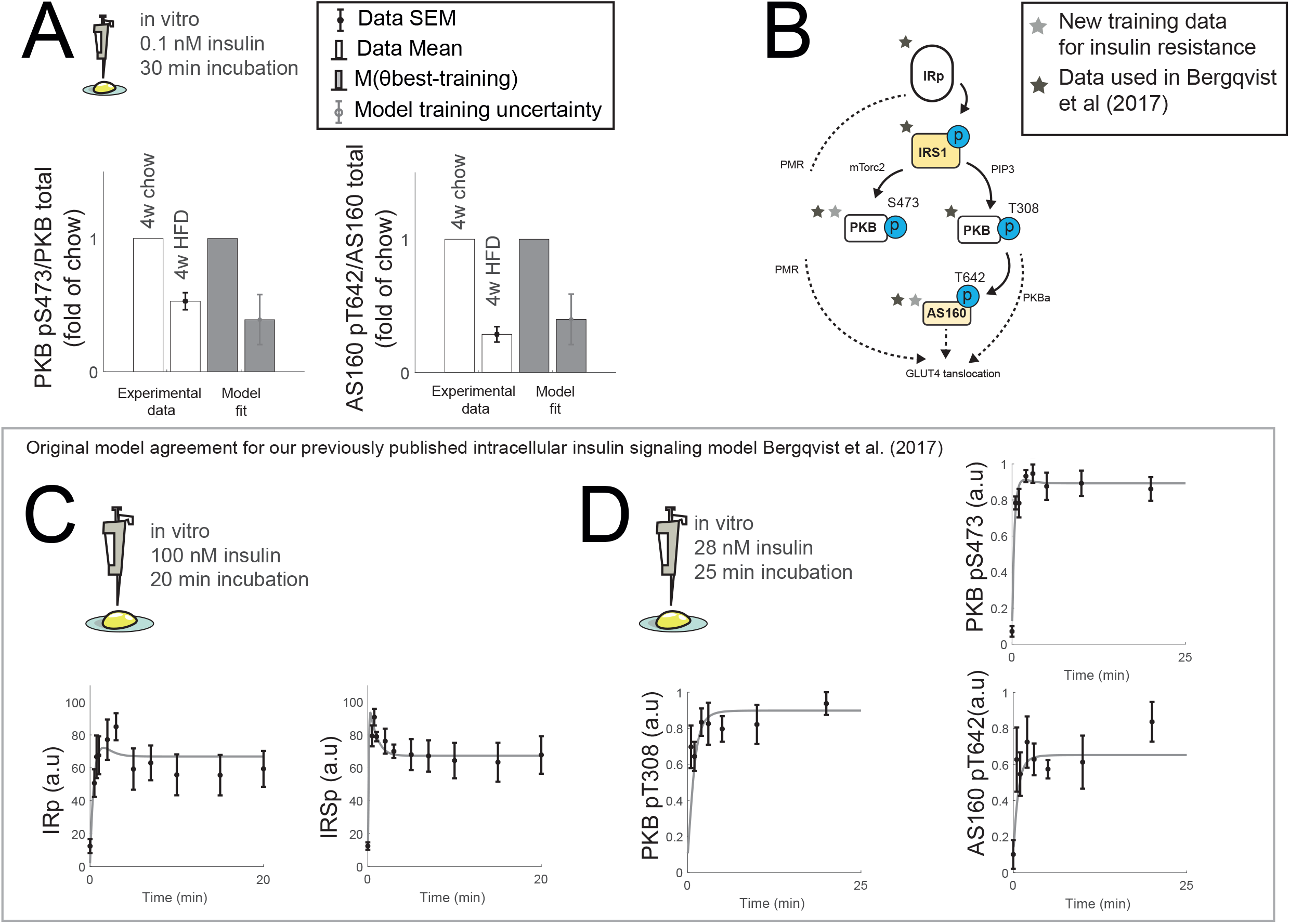
Result of model training in the intracellular insulin signaling model. **A)** Result of model training to intracellular data of PKB pS473 and AS160 pT6420 phosphorylation data after and 4-week duration of HFD or chow. The data (black error bar and white bar) describes the amount of phosphorylated protein fold over the total protein level, which was fold over the corresponding average for chow. The data comes from mice primary adipocytes stimulated with 0.1 nM insulin for 30 min. The grey bar represents the parameter-set that found the best agreement to the whole training dataset, with the corresponding model uncertainty shown as the grey error bars. **B)** Schematic structure over the insulin signaling pathway, highlighting the differences between the new data (light grey mark) and previously used training data (dark grey mark) in Bergqvist et al (2017)(18). **C)** Figure showing the model agreement to the data used in (18) for IR and IRS1 for experiments using primary rat adipocytes incubated with 100nM insulin for 20min. **D)** Figure showing the model agreement to the data used in (18) for PKB pT308, PKB pS473 and AS160 pT642 for experiments using primary rat adipocytes incubated with 28nM insulin for 25min.

### The whole-body part of our model describes body-weight alterations for different diets

We have previously performed dietary alteration experiments according to the setup shown in Fig. 1C, where male C57BL/6J mice were switched from a chow to an HFD at different times during a 2-week period (2). Almost all model parameters for the body-weight model have previously been experimentally validated (14) for male C57BL/6J mice. Similar to Gennemark *et al*., we let only the physical activity parameter (referred to as λ) be re-estimated (16). The model agrees well with training data (Fig. 3A-D, grey line). We also show the associated model uncertainty, which has been obtained by maximizing the model uncertainty at each data-point with the constraint that we must have agreement with all training data (Fig. 3A-D, grey area) (Materials and methods).

### The whole-body level model can predict new validation data not used during estimation

We further validated the trained model by predicting new unpublished data. Here, C57BL/6J mice were fed either chow diet or HFD for 8 weeks with IPGTT carried out at 6 weeks. The model was trained on the data for the first two-week duration to account for the varying experimental conditions from experiment to experiment. The model can predict the new data for the remaining duration of the 8 weeks period (Fig. 3E-F, blue error bars). As can be seen, the experimental data lies within the predicted bounds (light grey area). The blue dashed line shows the simulation which generated the lowest total cost when compared with these validation data 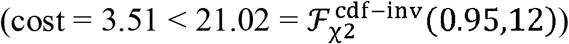. This successful prediction provides some faith in the quality of the whole-body part of the model.

### The tissue-level model can describe IPGTT data for different degrees of insulin resistance

In the same dataset as the initial body-weight training data (Hansson *et al*. 2018), IPGTT tests were preformed at the end of the diet period. The data indicates that exposure to HFD is correlated to higher basal glucose levels (first time-point in Fig. 4A-D), higher basal insulin levels (Fig. 4E), and lower glucose clearance (lower peak and faster decline in Fig. 4A-D), which all unanimously indicates that a longer time spent on HFD gives increasingly higher degrees of insulin resistance. As can be seen in Fig. 4, the model simulations (grey lines and areas) agree with all of these experimental observations (black error bars). In other words, our tissue-level model provides an acceptable explanation to the IPGTT data, collected at different stages of insulin resistance.

### The tissue-level model can predict new independent data not used for training

We used the same new experiment to validate the tissue-level part of the model, as we used to validate the whole-body level model. More specifically, we validated the trained model by predicting IPGTT responses in C57BL/6 mice after 6 weeks on HFD. As can be seen in Fig. 4F, the model predicts the new IPGTT data well (blue error bars), which again is statistically confirmed by a χ^2^ test 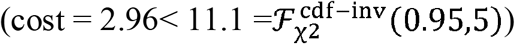.

### The cell-level model can describe phosphorylation data for key adipose insulin signaling intermediates

Finally, the model was simultaneously fitted also to insulin-signaling data, obtained from primary adipocytes extracted from the mice that had been on an HFD. This time, our already published data comes from an experiment where C57BL/6 mice were fed either chow or HFD for 4 weeks. The two experiments were compared, and the protein expression levels were normalized to the expressions in the chow data (Materials and methods, (20)). The model was trained to our previously published phosphorylation fold data of PKB-S473 and AS160-T642 from an *in vitro* system of primary mouse adipocytes incubated with 0.1 nM insulin for 30 min (20)(Fig. 5A). In Fig. 5B we present a summary of what signaling intermediates are included in our new data-set (light grey mark) compared to the corresponding data (dark grey mark) used for training in in our previous publication (18).

### The cell-level model has already been validated with respect to rodent data

We have previously developed and tested our intracellular insulin signaling model using our own data from another rodent system: rat adipocytes. These results were published in Bergqvist *et al*. (18), but we choose to include some of these previous results for completeness (Fig. 5C-D). Those *in vitro* data describes the intracellular response of different signaling intermediates to different doses of insulin. In (18), the model was estimated to a total of >140 data-points measuring the phosphorylation response in different proteins. Here, we include the comparison between two such estimation data and the model: IR-IRS1 phosphorylation during incubation with insulin (100 nM for 20 min, Fig. 5C) and phosphorylation of PKB and AS160 during incubation with insulin (28 nM for 25 min, Fig. 5CD). For the IR-IRS1 dynamics, the agreement between data and model was satisfactory, which is confirmed by a χ^2^ test 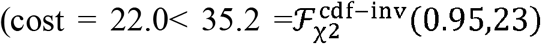, includes one data point for IR in plasma membrane). Also, for PKB and AS160 phosphorylation, the agreement between data and model was satisfactory, which is confirmed by a χ^2^ test 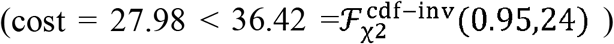. In Bergqvist *et al*. the model was also fitted and validated to GLUT4 translocation data, graphs not included here. Since these data from rat adipocytes are from another system than the rest of the data, the cell-level model has a scaling on the input signal to account for insulin doses and species-specific differences (Materials and methods).

### A humanoid version of the *in silico* mouse

We have now developed and validated all three levels in our multi-level and multi-timescale *in silico* mouse model and are ready to illustrate the potential with this multi-scale model. In principle, we could simulate various interventions, such as the response to a drug or another diet-scheme. Here, we choose to instead use the same dosing scheme and illustrate how we can improve the *in silico* mouse model, by making it more similar to human etiology. The reason why this is needed is that there are limitations in the existing *in vivo* mouse models for diet-induced T2D (Fig. 1D). Our new humanoid *in silico* mouse model has an additional phenomenological description of how beta-cell failure is induced in cases of long-term over-demand of insulin production (Fig. 6A). The exact behavior of this new beta-cell failure cannot be validated by experimental data – since there is no corresponding *in vivo* mouse model – but the main mechanisms and overall etiology mimic how human T2D develops, as can be seen in our simulations.

**Figure 6:**
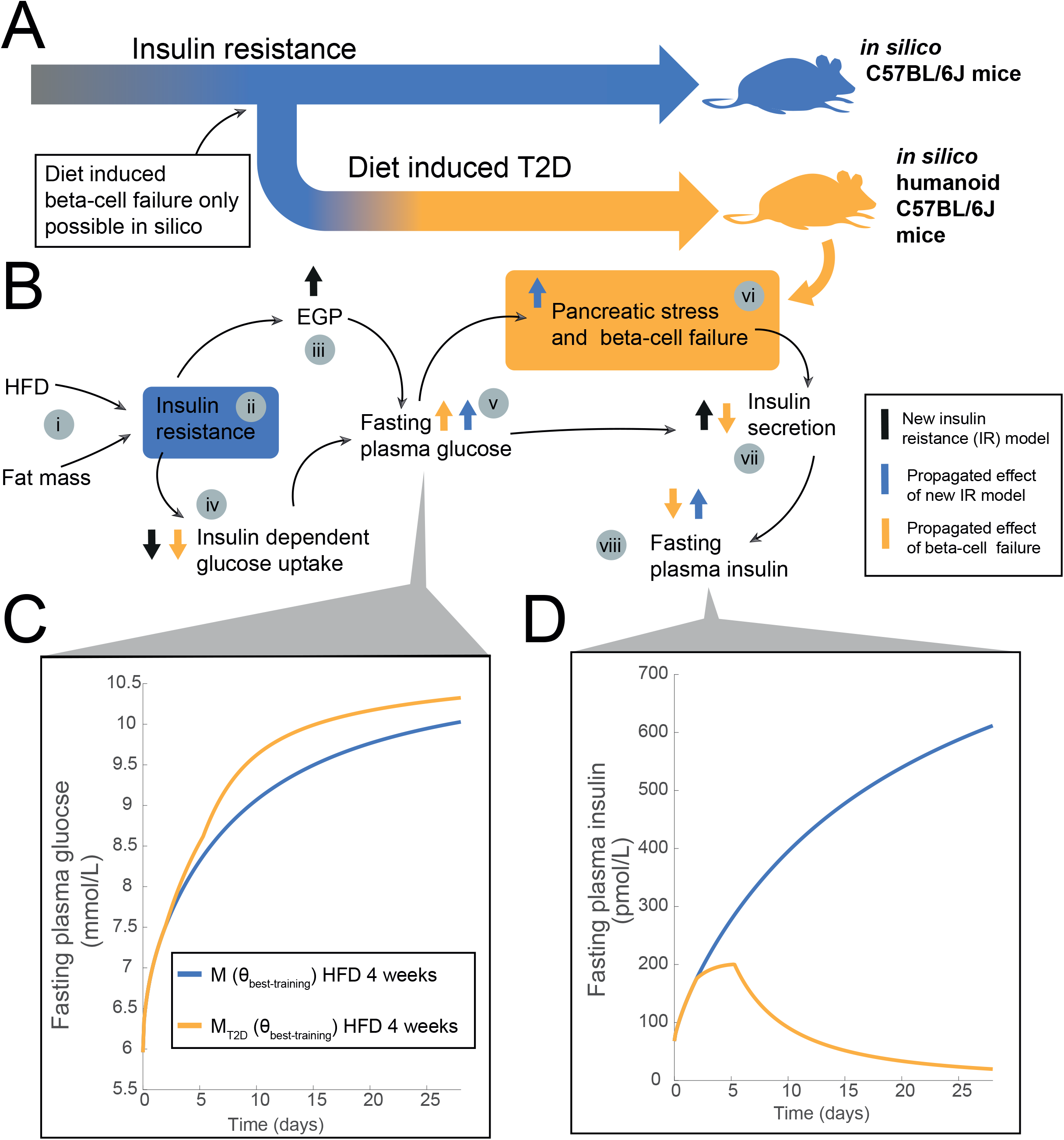
**A)** Visualization of the difference between the in silico mouse (blue) and the proposed in silico humanoid (orange) mouse. **B)** Schematic describing the disease progression in the two in silico mice. The broad arrows (black, blue, orange) describe the change in state values in response to disease progression for key components of the tissue-level model, and the narrow arrows describes how states interacts. Corresponding model simulations are found in Fig. SA2. The model insulin resistance (ii) is driven by time spent on HFD as well as the increase in fat mass (i), which is represented by the black arrows; increased EGP (iii), decreased insulin dependent glucose uptake(iv) and increased insulin secretion (vii). The blue arrows show the propagated effect from the changes brought on by the insulin resistance; increased fasting glucose (v), increased fasting insulin (viii). Exclusive to the humanoid mouse is the pancreatic stress and beta-cell failure (vi) which is driven by increased fasting glucose. The propagated effect of pancreatic stress and subsequent beta-cell failure is described by the broad orange arrows; decreased insulin secretion (vii), decreased fasting insulin (viii), increased fasting glucose (v) and decreased insulin dependent glucose uptake (iv). **C)** Fasting plasma glucose level during a model simulation of an in silico 4-week HFD intervention for the two mice. **D)** Fasting plasma insulin level during a model simulation of an in silico 4-week HFD intervention for the two mice. Here, the simulation for the humanoid mouse shows that fasting insulin goes below the initial baseline after some time of T2D.

The resulting simulated four-week responses for the two *in silico* mice are shown (Fig. 6C-D) for 1) the normal mice exposed to a HFD (light blue); and for 2) the new humanoid mice that can develop T2D (orange). The first part of the response (day 1-3) is identical in the two mice. During these first days, the HFD induces an increase of adipose tissue and both the diet and the increased adiposity led to an increased insulin resistance (Fig. 6Bii). In turn, this insulin resistance leads to increased fasting plasma glucose levels (Fig. 6C), due to an increased EGP (Fig. 6Biii) and a decreased insulin-dependent glucose uptake from all tissues (Fig. 6Biv). Finally, this increased glucose level leads to an increased insulin secretion and to a subsequent rise in insulin plasma levels (Fig. 6D). All these responses are identical for the two types of *in silico* mice during these initial days, and all these responses are based on the previous development and validation of the model (Fig. 2-7).

**Figure 7:**
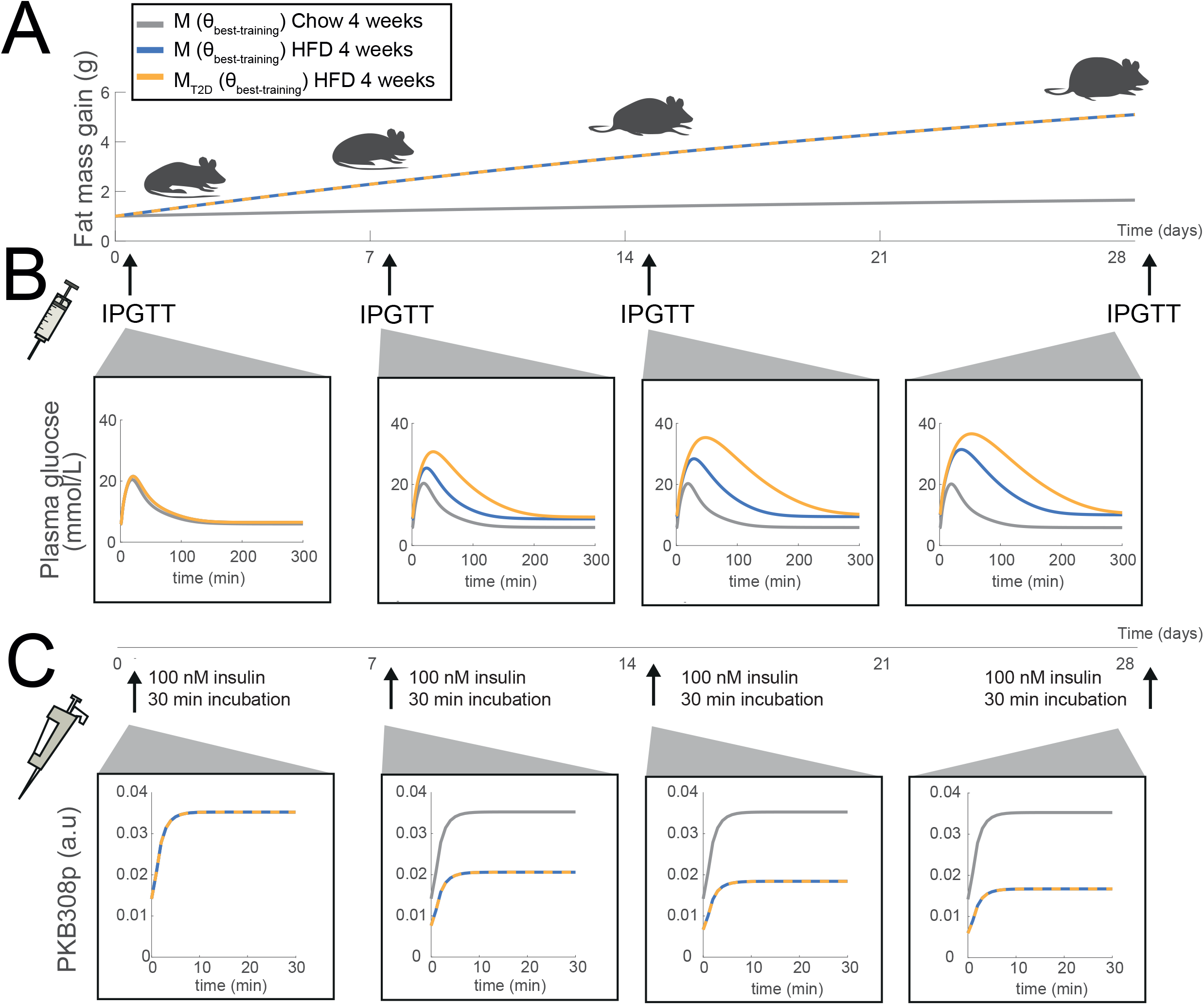
Model simulations for the same 4-week dietary intervention as in Fig 6, with the addition of a simulated chow control, as well as different intervention (IPGGT, insulin incubation) to study the short-term response during the disease progression. **A)** Model simulation for the predicted fat mass gain for the different mice; chow (grey), HFD regular in silico mouse (blue), and HFD with humanoid in silico mice (orange). The IPGTT (and insulin incubations) are marked with arrows and done at day; 1,7,14, 21. **B)** Plasma glucose during the IPGTTs. **C)** Simulations describing the in silico primary adipocyte experiments; with the cells being incubated with 100nM insulin for 30min. The response in phosphorylated PKB pT308 is shown.

The differences in the response in the two types of *in silico* mice can be seen after 3 days when the new mechanism kicks in: pancreatic stress. This pancreatic stress is modelled by an overdemand of insulin secretion (simulations of pancreatic stress and all other relevant variables are shown in SA Fig. SA2). More specifically, pancreatic stress starts to increase when the fasting plasma glucose level reaches a certain threshold value (7.5 mmol/L) and will inhibit insulin secretion to some degree (Fig 6vi). This leads to a plateau in the insulin secretion, representing a functional maximum secretion level (Fig 6vii). For the model to develop T2D, we created a second variable causing beta-cell failure. This variable starts to increase after some time of pancreatic stress (Fig 6vi). The beta-cell failure variable does not stop increasing and will thus create a point of no return resulting in a diminishing beta-cell mass; given sufficient time, the insulin secretion will be close to non-existent as seen by the severe reduction of fasting insulin levels in Fig 6D.

Using both types of *in silico* mice we can also simulate the short-term response during the proposed disease progression (Fig 6). In Fig 7, we show the same four-week *in silico* experiment as in Fig 6 C-D, but we here also chose to simulate a 4-week chow diet as control (grey line). To highlight the tissue-level response we preform four IPGTT’s at days 1, 7, 14 and 28 (Fig. 7B, black arrows). To illustrate the intracellular response, we performed *in silico* primary adipocyte experiments corresponding to the same type of experiments shown in Fig 5, where adipocytes were incubated with 100 nM insulin for 30 min. During the 4-week intervention, the HFD mice increased significantly more in fat-mass compared to the chow control (Fig. 7A). The tissue-level response is shown in the form of plasma glucose levels during each IPGTT (Fig 7B). As can be seen, the response dynamics is roughly the same during the first IPGTT, and changes over the 4-week duration. In the last IPGTT, the difference can clearly be seen between the different types of mice, with the humanoid mice having a much higher glucose peak and slower glucose clearance, which is due to the lack of insulin response. Similarly, during the 4-week duration, the HFD mice show an increasingly lower maximum amount of phosphorylated PKB (Fig. 7C). In this experiment the humanoid mice show no additional difference, as the insulin input is a set value (100 nM), and there is no modelled effect of T2D in the intracellular insulin signaling pathway. This *in silico* experiment is shown to highlight the difference between the chow control and mice developing insulin resistance.

Again, the diet-induced T2D progression shown in the humanoid mice model is not possible to validate, as no existing *in vivo* mice model can display this type of disease progression. Thus, these simulations highlight one of the key strengths with an *in silico* approach, namely to function as a complement to existing *in vivo* models, by enhancing and extending their capabilities to quantify and investigate complex disease progression.

## Discussion

Herein, we have presented the first multi-level, multi-timescale mechanistic model of insulin resistance progression: a new *in silico* mouse model. The new model can simultaneously find agreement to our multi-level dataset from C57BL/6J mice, describing insulin resistance development on three different biological levels: body composition (Fig. 3), plasma glucose and insulin (Fig. 4), and intracellular adipocyte insulin signaling (Fig. 5). The model can both describe estimation data and correctly predict independent validation data (Fig. 3-5). Lastly, we have modified this *in silico* mouse to create a humanoid *in silico* mouse model (Fig. 6-7), which can describe diet induced T2D, a feature not exhibited by any currently existing *in vivo* mouse model (Fig. 1D and Fig. 6).

### Our humanoid *in silico* mouse improves upon available *in vivo* mouse models

There exist several *in vivo* mouse models which can describe different aspects of T2D, but none of them captures the whole human T2D etiology. In humans, the progression starts with the development of peripheral insulin resistance (in e.g. muscle and adipose tissue) which leads to compensatory hyperinsulinemia and increased demand on the beta cells. This increased demand, possibly connected to inflammation and minimal ectopic fat storage in the beta cells, eventually leads to loss of beta-cell mass and beta-cell function (22). In mice, different aspects of this human etiology and pathology have been captured, using e.g., genetic manipulation, chemical injections, or external perturbations, such as dietary intervention or sleep deprivation (23, 24). Herein, we have studied C57BL/6J mice, which develop insulin resistance and compensatory insulin secretion following a HFD (3). However, this mice model keeps producing high levels of insulin, and thus do not develop T2D, but only pre-diabetes (Fig. 1D, black). This differs from mice having induced beta-cell failure, either by surgically removing the mice beta-cells (25), chemically inducing beta-cell failure (10), or by genetically perturbing the beta-cell function (26) (Fig. 1D, grey). Indeed, streptozotocin-induced diabetic mice displayed normalized glucose levels after receiving human beta-cell implants (25), pin-pointing the importance of maintained insulin secretion. In summary, there is thus no murine model that describes the initial increase in insulin production, due to insulin resistance, which then is followed by a resulting beta-cell failure and failed insulin production, as is produced by our *in silico* humanoid mice model (Fig. 1D and Fig. 6).

### The power of a mechanistic, multi-level, and multi-timescale model

The novel model presented herein opens the door to many new possibilities. One such possibility is that it can serve as a knowledge base. In other words, even if the model did not have any predictive capabilities, it is useful as a compact way of presenting a wide variety of data and knowledge. However, a model goes beyond mere visualization, it is an *explanation*. An explanation is a statistically acceptable hypothesis, which also is compatible with mechanistic knowledge about the system (12). Thus, our model presents a simplified view of the mechanisms behind the many different datasets presented herein and how they could have been generated. Because of the complex nature of the data, spanning different levels and timescales, such a testing of different tentative mechanistic explanations is almost impossible to do by mere reasoning around the data. Nevertheless, the arguably most important role of a mechanistic models is their ability to do simulations, i.e., predictions of what would happen in different scenarios. Our model could, e.g., be used to predict what would happen to these mice, if they would have been fed with a different diet protocol, or if they would have been exposed to a drug that modifies some of the processes described. In principle, the model can then describe how such modifications is propagated between the different levels, e.g. how an intracellular modification is propagated to the whole-body level. Such predictions can initially be used to test the current explanation for the data, by doing the same type of validation experiments that we have done here, comparing such predictions with new data (Fig. 4,6). Note that such predictions are useful also if the prediction fails, since that means that some critical mechanism is missing in the model (or that already included mechanisms needs to be modified). However, once such a multi-level multi-timescale model has been sufficiently validated, it will become increasingly more useful for e.g., *in silico* drug screening, or for identification of new drug targets. Finally, one additional potential with our *in silico* models is that they can be used to improve the realism of the original mouse model: e.g. by making them able to produce diet-induced T2D.

### Using the humanoid *in silico* mouse model to simulate diet-induced T2D: a new sub-step before translating to humans

The diet-induced T2D was introduced into the humanoid mice model by replacing the original beta-cell model with one that mimics the human etiology. This was done by the addition of two new model variables, pancreatic stress, *P*_*Stress*_, and beta-cell failure, *Beta*_*failure*_ (SA Eq. 69,70). The pancreatic stress in our model is caused by the increasing fasting glucose levels, representing the effect of glucotoxicity, and the stress starts when the fasting levels increase beyond a certain threshold level (Materials and methods). This is in line with the mathematical model for humans by Topp *et al*, where plasma glucose also is used as a driving force to control beta-cell mass (27). An even more detailed model for beta-cell failure is presented in Ha *et al*.(28). We choose not to use such detailed descriptions of beta-failure, since we wanted to keep the humanoid modifications as simple as possible, to illustrate proof of principle. Future *in silico* mouse models could use some of those more advanced beta-cell models.

Importantly, this new improved humanized mouse model, constitutes a new intermediary step, which lies in-between murine *in vivo* experiments and a full translation to human conditions. As described above, a mathematical model is useful to more correctly interpret experimental mouse data, by systematically testing and validating complex mechanistic explanations. Ultimately, one wants to translate these mathematical disease models to human conditions, as one already does routinely for e.g. pharmacokinetic properties, in drug development (29, 30). However, a corresponding translation also regarding the pharmacodynamic side, i.e., describing how a drug is interacting with the disease mechanisms, is still not available. The main reason for this is that there are many differences between humans and rodents, on many levels: intracellular signaling details differ, cell sizes differ, quantitative parameters differ, etc. (31, 32). These problems with scaling are exacerbated by the fact that we only know some of these interspecies differences. Before all those disease mechanisms are sufficiently elucidated to allow for a proper translation, we propose to use our type of *in silico* mouse models as a new intermediary level: a level where some human characteristics missing from rodent models have been introduced, but where the majority of the mechanisms still are consistent with the mouse mechanisms, which can be supported by numerous experimental data on all three levels: cell, organ, and whole-body.

Finally, there are some human *in silico* models with which our results should be compared. On the whole-body level, Hall *et al* have published some important models for weight regulation (e.g. (33)), but just as for the corresponding rodent models, these top-level models do not interconnect with more short-term models and models for the other two levels (Fig 1A). On the organ-level, there are numerous glucose-insulin meal response models (34, 35), and also some models that describe fatty acids (36), and other metabolites (37). There are also some such meal response models that connect with intracellular models for the pancreas and insulin signaling in the adipocytes (38-40), but these models do not interconnect with long-term disease progression whole-body models. Furthermore, the meal-response model that we use is a rescaled version of a human meal-response model (17, 35). Finally, there are some long-term disease-progression models, featuring multiple (41), but these do not describe short-term responses such as meals, and are thus not multi-timescale. In summary, our new model is the first data driven, mechanistic, multi-level (intracellular to whole-body) and multi-timescale model (minutes to months) for insulin resistance, not only for rodents, but for any species.

### Limitations and assumptions

Some limitations to the current results should be mentioned. To be able to find agreement to all data in our multi-level dataset, we had to re-estimate several of the model parameters (SA, Table 1-7). We believe this is reasonable since we describe experiments with different conditions compared to the original publications. Furthermore, our glucose load data were IPGTT-data which is different compared with data obtained from IVGTT and OGTT which the model-parameters originally were fitted to (17, 35). Another issue concerns the fact that we do not know the true uncertainty of the data points, since just looking at repeat variations, can sometimes give too small values, which ignores systematic errors. Therefore, within a dataset, all SEM values that were smaller than the mean SEM for the dataset, were replaced with mean SEM value. This was done to remove the high impact that those datapoints otherwise have, which we judged to be unwarranted. By keeping the original SEM values, we could have gotten an over-fitted model, which we wanted to avoid. Another limitation is that we did not have a lot of data available for intracellular responses in mice adipocytes, and that the original intracellular model was based on data obtained from rats. An important task for the future is thus to adopt and refine the intracellular model to the unique responses in mice. Another limitation is that we decided not to include insulin meal-response data in the model training. The model’s description of the insulin meal response is very simplified, and substantial model development would have been necessary to find agreement with data, especially data for the meal-response during different stage of insulin resistance. Finally, many simplifications have been done, and many central processes are not included in our model, for example intracellular signaling in muscle and liver, ectopic fat storage, other hormones such as glucagon and stress hormones.

In conclusion, while there are many assumptions and limitations done to produce this model, it is the first mechanistic, multi-level, and multi-timescale model for insulin resistance based on data from all three levels: intracellular, organ, and whole-body level. We hope that this model will become an important basis for future interpretation and integration of experimental data, which ultimately will help with the development of new treatments for insulin resistance and T2D.

## Materials and methods

### Animals and high fat diet intervention

Short-term overfeeding was carried out as previously described (2, 20, 42). Male C57BL/6J mice (Taconic, Ry, Denmark) were used at 8-9 weeks of age. Animals were on a 12 h light cycle with non-restricted food and water. Groups of animals (n=6-10 animals/group) were fed either chow or HFD (D12492 60 E% fat content; Research Diets, New Brunswick, NJ, USA) for 6-8 weeks. Body weight (BW) of individual animals were measured weekly.

### Animals and intraperitoneal glucose tolerance test

A subset of animals (n=10/group) were subjected to intraperitoneal glucose tolerance test (IPGTT). In short, mice fasted overnight (12 h) were injected intraperitoneally with glucose (50 mg/mouse) and serum samples were collected at indicated times. Blood glucose levels were measured (Accu-Chek, Lund, Sweden), and insulin levels were assayed using ELISA (Mercodia, Uppsala, Sweden). The supplier’s protocol was followed with an adjustment of the amount of sample volume (5µL of each sample) and the incubation time (30 min at room temperature).

All animal procedures were approved by the Malmö/Lund Committee for Animal Experiment Ethics, Lund, Sweden, and were carried out in accordance with the relevant guidelines and regulations.

### Mathematical modelling

The mathematical analysis, model simulation, numerical optimization of model parameters, was carried out using MATLAB 2018b and the SBtoolbox2 package (43).The model is based on *ordinary differential equations* (ODEs) with the general form:

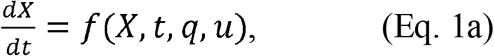

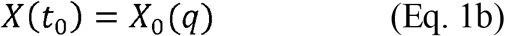

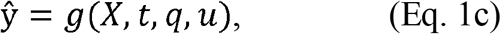

here, *X* represents a vector of state variables usually corresponding to concentrations of given system components; the functions *f* and *g* are non-linear smooth functions; *q* is a vector of model parameters (rate constants, scaling constants etc.); *u* is the input signal corresponding to the experimental data; *X* (*t*_0_), represents the initial condition value *X*_*0*_ (*q*), which are dependent on the model parameters *q*; and ŷ is the simulated model output. Parameter estimation was done by quantifying the model performance, using the model output ŷ to calculate the traditional weighed least squares cost function written as

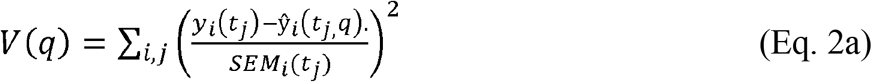

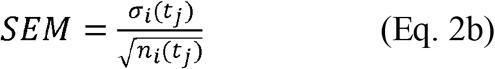

where, *q* is the model parameters; *y*_*i*_ (*t*_*j*_) is the measured data from an experimental setup *i*, at time point *j*; ŷ_*i*_ (*t*_*j*,_ *q*) is the simulation value for a given experiment setup *i* and time point *j*, and SEM is the standard error of the mean, which is the sample standard deviation, *σ*_*i*_ (*t*_*j*_)divided with the square root of the number of repeats, *n*_*i*_ (*t*_*j*_)at each time point. The value of the cost function, *V* (*q*), is then minimized by tuning the values of the parameters, typically referred to as parameter estimation. The parameter estimation was done using the extended scatter search (ESS) optimization algorithm from the MEIGO toolbox (44). The parameter estimation protocols were run at the Swedish national supercomputing center (NSC).

In order to evaluate the new model, we performed a *χ*^2^-test for the size of the residuals, with the null hypothesis that the experimental data have been generated by the model, and that the experimental noise is additive and normally distributed (12). In practice the cost function value was compared to a *χ*^2^ test statistic, 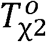 The test statistic value is given by the inverse cumulative density function,

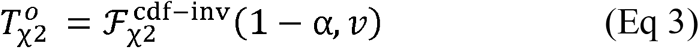

where 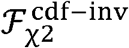 is the inverse density function; and α is the significance level (α = 0.05, was used) and *v* is the degrees of freedom, which was equal to the number of data points in the training dataset (36 in total, all timepoints over all experiments). In practice, the model is rejected if the model cost is larger than the *χ*^2^-threshold 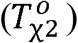.

The model uncertainty was estimated by reformulating the optimization problem, as proposed in Cedersund (2012) and implemented in Lövfors *et al*. (13, 45), to find the parameter sets that either maximized or minimized the model simulation at all given data points (36 in total), with the condition that the model should pass a *χ*^2^ test. The reformulated objective function can be written as

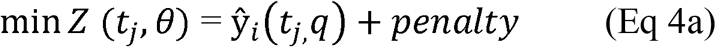

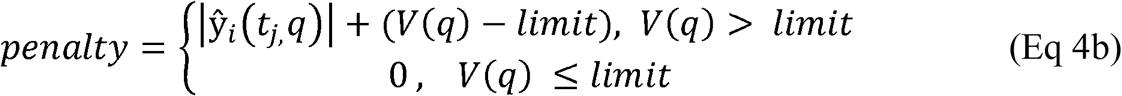

Here, Z (*t*_*j*_, *θ*) is the modified objective function; ŷ_*i*_ (*t*_*j*,_ *q*) is the simulation for a given timepoint *j* and experiment, *i*, the *penalty* is the condition that the cost function value should pass the *χ*^2^ test, and where *limit* is the *χ*^2^-threshold with α = 0.05 and 36 degrees of freedom (equal to the number of data points).

## Supporting information

Supplementary scripts

Appendix: Model description

## Author Contributions

C.S has done the model analysis, with support from WL, and with initial analysis by NB. Primary supervision and original conceptions has been done by GC, and additional supervision and inputs on parts of the modelling has been done by KS, EN, and PG. Experiments have been supervised and carried out in the lab of KS. The paper has been written primarily by CS and GC, with input from all authors. All authors have seen and approved of the final manuscript.

