## Supplementary scripts for "A multi-scale *in silico* mouse model for insulin resistance and humanoid type 2 diabetes": Plotscript_BQ2017.m

```
%% Notes %%%%%%%%%%%%%%%%%%%%%%%%%%%%%%%%%%%%%%%%%%%%%%%%%%%%%%%%%%%%%%%%
% This script simulates the model(s) with different insulin concentrations.
% See text document for details.
%%%%%%%%%%%%%%%%%
%% INITIAL STUFF
%%%%%%%%%%%%%%%%%
simOptions = [];
simOptions.method = 'stiff';
simOptions.maxnumsteps = 1e8;

modelName = 'GLUTcomplete';
optModel = SBmodel(strcat(modelName,'.txt'));
SBPDmakeMEXmodel(optModel,modelName);
modelNamekr = 'GLUTcompletekr';
optModelkr = SBmodel(strcat(modelNamekr,'.txt'));
SBPDmakeMEXmodel(optModelkr,modelNamekr);
modelNameBasal = 'GLUTPlosOne';
optModelbasal = SBmodel(strcat(modelNameBasal,'.txt'));
SBPDmakeMEXmodel(optModelbasal,modelNameBasal);
modelNameBasalkr = 'GLUTPlosOnekr';
optModelbasalkr = SBmodel(strcat(modelNameBasalkr,'.txt'));
SBPDmakeMEXmodel(optModelbasalkr,modelNameBasalkr);

[pNamesOpt, param] = SBparameters(optModel);
icOrig = SBinitialconditions(optModel);
load('EXPDATA.mat');
load('FractionData.mat');
% load('paramset1.mat');

GlutData=EXPDATA.GlutData;
ASData=EXPDATA.AS160;
PKBData=EXPDATA.PKB308;
PKB2Data=EXPDATA.PKB473;
mod1=EXPDATA.mod1;
C2DataB=EXPDATA.C2DataB;
C2DataB01=EXPDATA.C2DataB01;
C2DataB02=EXPDATA.C2DataB02;
C2DataB04=EXPDATA.C2DataB04;
C2Datakr=EXPDATA.C2Datakr;
C2Datakr01=EXPDATA.C2Datakr01;
C2Datakr02=EXPDATA.C2Datakr02;
C2Datakr04=EXPDATA.C2Datakr04;
fraction=FractionData.fraction;
PKBdf=FractionData.PKBdf;
RABGTPf=FractionData.RABGTPf;
PMRf=FractionData.PMAf;

%param=paramset1;
param(end)=0;
%%%%%%%%%%%%%%%%%%%%%%%%%%%%%%%%%%%%%%%%%%%%%%%%
%% MIFB1 INS=100 nM (MODULE 1)
%%%%%%%%%%%%%%%%%%%%%%%%%%%%%%%%%%%%%%%%%%%%%%%%
Time=mod1.Time;
IR_mean=mod1.IR_mean;
IR_std=mod1.IR_std;
IRS1_mean=mod1.IRS1_mean;
IRS1_std=mod1.IRS1_std;
InTime = [0:0.025:20];
try
  simDataSteadyState = SBPDsimulate(modelName,(0:1:200),icOrig,pNamesOpt,param,simOptions);
catch
    cost = inf;
    return 
end

Ic = simDataSteadyState.statevalues(end,:);

newparam=param;
newparam(end) = 100; %Insulin concentration during the experimental procedures
try
    simData = SBPDsimulate(modelName,InTime,Ic,pNamesOpt,newparam,simOptions);
catch
    disp('Now the simulation crashed! [1]');
    cost = inf;
    return 
end

maxIRmem   = max(simData.variablevalues(:,8));

simIRS1=1000*simData.variablevalues(:,6);
simIR=1000*simData.variablevalues(:,7);
simIRmem=100*simData.variablevalues(:,8)/maxIRmem;

%%%%%%%%%%%%%%%%%%%%%%%%%%%%%%%%%%%%%%%%%%%%%%%%%%%%%%%%%%%%%%%%%%%%
%% PKB-T308, PKB-S473, AS160. INS=28 nM (MODULE 2)
%%%%%%%%%%%%%%%%%%%%%%%%%%%%%%%%%%%%%%%%%%%%%%%%%%%%%%%%%%%%%%%%%%%%
try
  simDataSteadyState = SBPDsimulate(modelName,(0:1:200),icOrig,pNamesOpt,param,simOptions);
catch
    cost = inf;
    return 
end

Ic = simDataSteadyState.statevalues(end,:);

newparam = param;
newparam(end)=28; %Insulin concentration during the experimental procedures

try
    simData = SBPDsimulate(modelName,(200:0.001:230),Ic,pNamesOpt,newparam,simOptions);
catch
    disp('Now the simulation crashed! [2]');
    cost = inf;
    return 
end

PKB308=simData.variablevalues(:,end-3);
PKB473=simData.variablevalues(:,end-2);  
AS160p=simData.variablevalues(:,end-1);

%%%%%%%%%%%%%%%%%%%%%%%%%%%%%%%%%%%%%%%%%%%%%%%%%%%%
%% GLUT4 Transloacation (MODULE 3)
%%%%%%%%%%%%%%%%%%%%%%%%%%%%%%%%%%%%%%%%%%%%%%%%%%%%

try % % % Simulation of steadystate.% % % %
    simDatasteady = SBPDsimulate(modelName,(0:1:200),icOrig,pNamesOpt,param,simOptions);
    simDatasteadybasal = SBPDsimulate(modelNameBasal,(0:1:200),icOrig,pNamesOpt,param,simOptions);
catch
    cost = inf;
    disp('Now the simulation crashed! [3]');
    return
end

%Saving steadystatevalues for further simulations
Steadystatevaluesh=simDatasteady.statevalues(end,:);
Steadystatevaluesbasal=simDatasteadybasal.statevalues(end,:);

%% Simulation of basal plosOne values
%%P=1
init=param(end-1);
param(end-1)=1;
 try
  simDatab = SBPDsimulate(modelNameBasal,(200:0.01:205),Steadystatevaluesbasal,pNamesOpt,param,simOptions);
  simDatakr = SBPDsimulate(modelNameBasalkr,(200:0.01:205),Steadystatevaluesbasal,pNamesOpt,param,simOptions);
 catch 
 cost = inf; 
 disp('Now the simulation crashed! [4]');
 return 
 end 
SIMDATA1 = simDatab.variablevalues(:,end-4);
SIMDATAkr1 = simDatakr.variablevalues(:,end-4);

%%P=0.4
param(end-1)=0.4;
 try
  simDatab = SBPDsimulate(modelNameBasal,(200:0.01:205),Steadystatevaluesbasal,pNamesOpt,param,simOptions);
  simDatakr = SBPDsimulate(modelNameBasalkr,(200:0.01:205),Steadystatevaluesbasal,pNamesOpt,param,simOptions);
 catch 
 cost = inf; 
 disp('Now the simulation crashed! [5].');
 return 
 end 
SIMDATA04 = simDatab.variablevalues(:,end-4);
SIMDATAkr04 = simDatakr.variablevalues(:,end-4);

%%P=0.2
param(end-1)=0.2;
 try
  simDatab = SBPDsimulate(modelNameBasal,(200:0.01:205),Steadystatevaluesbasal,pNamesOpt,param,simOptions);
  simDatakr = SBPDsimulate(modelNameBasalkr,(200:0.01:205),Steadystatevaluesbasal,pNamesOpt,param,simOptions);
 catch 
 cost = inf; 
 disp('Now the simulation crashed! [6]');
 return 
 end 
SIMDATA02 = simDatab.variablevalues(:,end-4);
SIMDATAkr02 = simDatakr.variablevalues(:,end-4);

%%P=0.1
param(end-1)=0.1;
 try
  simDatab = SBPDsimulate(modelNameBasal,(200:0.01:205),Steadystatevaluesbasal,pNamesOpt,param,simOptions);
  simDatakr = SBPDsimulate(modelNameBasalkr,(200:0.01:205),Steadystatevaluesbasal,pNamesOpt,param,simOptions);
 catch 
 cost = inf; 
 disp('Now the simulation crashed! [7]');
 return 
 end 
SIMDATA01 = simDatab.variablevalues(:,end-4);
SIMDATAkr01 = simDatakr.variablevalues(:,end-4);
param(end-1)=init;

%% Simulation 30min for cell.met paper 
param(end)=70; %Insulin concentration during the experimental procedures,
%Cell.metabolism and plosOne.
 try
  simData = SBPDsimulate(modelName,(200:0.001:230),Steadystatevaluesh,pNamesOpt,param,simOptions); 
 catch 
 cost = inf; 
 disp('Now the simulation crashed! [8]');
 return 
 end 
SIMDATAy = simData.variablevalues(:,end); 

%% For further insulin simulations
Simulatedstatevalues=simData.statevalues(end,:);

%Sorting out important values
c0 = simData.statevalues(:,end-2);
c1 = simData.statevalues(:,end-1);
c2 = simData.statevalues(:,end);

v1 = simData.variablevalues(:,1);
v2 = simData.variablevalues(:,2);
v1exp = [0.0015 0.10 0.03]; %cell.met
v2exp = [0.03 0.05 0.016]; %cell.met
v1fold=max(v1)/v1(1);
v2fold=max(v2)/v2(1);


%% InsulinsteadystateSimulation with inputs to c2=0 and vexp active
%P=1
param(end)=70;
tmes=[1 84 168 251 334 418 501]; %Time in min times 100
param(end-1)=1;
try
  simDatainsb = SBPDsimulate(modelName,(230:0.01:235),Simulatedstatevalues,pNamesOpt,param,simOptions);
  simDatainskr = SBPDsimulate(modelNamekr,(230:0.01:235),Simulatedstatevalues,pNamesOpt,param,simOptions);
 catch 
 cost = inf;
 disp('Now the simulation crashed! [9]');
 return 
end 
C2data1b=simDatainsb.variablevalues(tmes,end-4);
C2data1kr=simDatainskr.variablevalues(tmes,end-4);

param(end-1)=0.4;
try
  simDatainsb = SBPDsimulate(modelName,(230:0.01:235),Simulatedstatevalues,pNamesOpt,param,simOptions);
  simDatainskr = SBPDsimulate(modelNamekr,(230:0.01:235),Simulatedstatevalues,pNamesOpt,param,simOptions);
 catch 
 cost = inf; 
 disp('Now the simulation crashed! [10]');
 return 
end 
C2data04b=simDatainsb.variablevalues(tmes,end-4);
C2data04kr=simDatainskr.variablevalues(tmes,end-4);

param(end-1)=0.2;
try
  simDatainsb = SBPDsimulate(modelName,(230:0.01:235),Simulatedstatevalues,pNamesOpt,param,simOptions);
  simDatainskr = SBPDsimulate(modelNamekr,(230:0.01:235),Simulatedstatevalues,pNamesOpt,param,simOptions);
 catch 
 cost = inf; 
 disp('Now the simulation crashed! [11]');
 return 
end 
C2data02b=simDatainsb.variablevalues(tmes,end-4);
C2data02kr=simDatainskr.variablevalues(tmes,end-4);

param(end-1)=0.1;
try
  simDatainsb = SBPDsimulate(modelName,(230:0.01:235),Simulatedstatevalues,pNamesOpt,param,simOptions);
  simDatainskr = SBPDsimulate(modelNamekr,(230:0.01:235),Simulatedstatevalues,pNamesOpt,param,simOptions);
 catch 
 cost = inf; 
 disp('Now the simulation crashed! [12]');
 return 
end 
C2data01b=simDatainsb.variablevalues(tmes,end-4);
C2data01kr=simDatainskr.variablevalues(tmes,end-4);

%% Transformation to dwell-time to match the units in the article
%%(plosOne)
for j=1:6
    dwellt1b(j)=C2data1b(j)-C2data1b(j+1);
end
for j=1:6
    dwellt1kr(j)=C2data1kr(j)-C2data1kr(j+1);
end
for j=1:6
    dwellt04b(j)=C2data04b(j)-C2data04b(j+1);
end
for j=1:6
    dwellt04kr(j)=C2data04kr(j)-C2data04kr(j+1);
end
for j=1:6
    dwellt02b(j)=C2data02b(j)-C2data02b(j+1);
end
for j=1:6
    dwellt02kr(j)=C2data02kr(j)-C2data02kr(j+1);
end
for j=1:6
    dwellt01b(j)=C2data01b(j)-C2data01b(j+1);
end
for j=1:6
    dwellt01kr(j)=C2data01kr(j)-C2data01kr(j+1);
end

%% Exponential fit to dwell time diagram and the computation of K:
dtime=[0.84 1.68 2.51 3.34 4.18 5.01];
f = fit(dtime',dwellt1b','exp1');
ab = coeffvalues(f);
K1b = -ab(2);
f = fit(dtime',dwellt1kr','exp1');
ab = coeffvalues(f);
K1kr = -ab(2);
f = fit(dtime',dwellt04b','exp1');
ab = coeffvalues(f);
K04b = -ab(2);
f = fit(dtime',dwellt04kr','exp1');
ab = coeffvalues(f);
K04kr = -ab(2);
f = fit(dtime',dwellt02b','exp1');
ab = coeffvalues(f);
K02b = -ab(2);
f = fit(dtime',dwellt02kr','exp1');
ab = coeffvalues(f);
K02kr = -ab(2);
f = fit(dtime',dwellt01b','exp1');
ab = coeffvalues(f);
K01b = -ab(2);
f = fit(dtime',dwellt01kr','exp1');
ab = coeffvalues(f);
K01kr = -ab(2);

%%%%%%%%%%%%%%%%%%%%%%%%%%%%%%%%%%%%%%%%%%%%%%%%%
%% Validation data simulation
%%%%%%%%%%%%%%%%%%%%%%%%%%%%%%%%%%%%%%%%%%%%%%%%%
%%Normal Simulation %%%%%%%%%%%%%%%%%%%%%%%%%%%%%%%%%%%%
param(end)=0;
simDatasteadynorm = SBPDsimulate(modelName,(0:1:200),icOrig,pNamesOpt,param);
Steadystatevaluesnorm=simDatasteadynorm.statevalues(end,:);
param(end)=7;
simDatanorm = SBPDsimulate(modelName,(200:0.001:230),Steadystatevaluesnorm,pNamesOpt,param,simOptions);
totPKBnorm=simDatanorm.statevalues(1,19)+simDatanorm.statevalues(1,20)+simDatanorm.statevalues(1,21)+simDatanorm.statevalues(1,22);
cmn7 = simDatanorm.variablevalues(end,end-5);
GLUT4norm=simDatanorm.variablevalues(:,end-5);
PKB308norm=simDatanorm.variablevalues(:,end-3); %Ploted
%Normal initial values but with reduced PKB-T308 activation (min)
h1param=param;
red=0.65;
h1param(17)=h1param(17)*red; %Parameter associated with PKB-T308 phosphorylation
h1param(21)=h1param(21)*red;
simDatanorm = SBPDsimulate(modelName,(200:0.001:230),Steadystatevaluesnorm,pNamesOpt,h1param,simOptions);
PKB308nmin=simDatanorm.variablevalues(:,end-3); %Ploted
%Normal initial values but with reduced PKB-T308 activation (max)
h2param=param;
red=0.45; %PKB-T308 inhibition factor
h2param(17)=h2param(17)*red; %Parameter associated with PKB-T308 phosphorylation
h2param(21)=h2param(21)*red;
simDatanorm = SBPDsimulate(modelName,(200:0.001:230),Steadystatevaluesnorm,pNamesOpt,h2param,simOptions);
PKB308nmax=simDatanorm.variablevalues(:,end-3); %Ploted

%%Altered simulation MIMIMUM %%%%%%%%%%%%%%%%%%%%%%%%%%
param(end)=0;
icOrigtemp = icOrig;
icOrigtemp(19) = icOrig(19)*0.51; %Initial reduction of total PKB
icOrigtemp(22) = icOrig(22)*0.51; %Initial reduction of total PKB
simDatasteadyh = SBPDsimulate(modelName,(0:1:200),icOrigtemp,pNamesOpt,param);
Steadystatevaluesh=simDatasteadyh.statevalues(end,:);
param(end)=7; %define ins conc
hparam=param;
red=0.65; %PKB-T308 inhibition factor
hparam(17)=hparam(17)*red; %Parameter associated with PKB-T308 phosphorylation
hparam(21)=hparam(21)*red; %Parameter associated with PKB-T308 phosphorylation
simDatah = SBPDsimulate(modelName,(200:0.001:230),Steadystatevaluesh,pNamesOpt,hparam,simOptions); 
totPKBh=simDatah.statevalues(1,19)+simDatah.statevalues(1,20)+simDatah.statevalues(1,21)+simDatah.statevalues(1,22);
cmh7 = simDatah.variablevalues(end,end-5);
totPKBdmin = totPKBh/totPKBnorm;
GLUT4mredmin = (cmn7-cmh7)/cmn7;
GLUT4min=simDatah.variablevalues(:,end-5);

%%Altered simulation MAXIMUM %%%%%%%%%%%%%%%%%%%%%%%%%%%
param(end)=0;
icOrigtemp = icOrig;
icOrigtemp(19) = icOrig(19)*0.37; %Initial reduction of total PKB
icOrigtemp(22) = icOrig(22)*0.37; %Initial reduction of total PKB
simDatasteadyh = SBPDsimulate(modelName,(0:1:200),icOrigtemp,pNamesOpt,param);
Steadystatevaluesh=simDatasteadyh.statevalues(end,:);
param(end)=7; %define ins conc
hparam=param;
red=0.45;
hparam(17)=hparam(17)*red; %Parameter associated with PKB-T308 phosphorylation
hparam(21)=hparam(21)*red; %Parameter associated with PKB-T308 phosphorylation
simDatah = SBPDsimulate(modelName,(200:0.001:230),Steadystatevaluesh,pNamesOpt,hparam,simOptions); 
totPKBh=simDatah.statevalues(1,19)+simDatah.statevalues(1,20)+simDatah.statevalues(1,21)+simDatah.statevalues(1,22);
cmh7 = simDatah.variablevalues(end,end-5);
totPKBdmax = totPKBh/totPKBnorm;
GLUT4mredmax = (cmn7-cmh7)/cmn7;
GLUT4max=simDatah.variablevalues(:,end-5);

%% Figures %%%%%%%%%%%%%%%%%%%%%%%%%%%%%%%%%%%%%%%%%%%%%%%%%%%%%%%%%%%%%%% 

err_color='k*';
tr_color = [ 0.5,0.5,0.5];

%% Figure2: Module 1
%%%%%%%%%%%%%%%%%%%%%%%%%%%%%%%%%%%%%%%%%%%%%%%%%%%%%%%%%%%%%%%%%%%%%%%%%%
%PLOT2: IR comparison with data
subplot(2,3,1);
h=gca;
hold on
plot(InTime,simIR,'color',tr_color,'linewidth',2)
errorbar(Time,IR_mean, IR_std,err_color,'linewidth',1.5) 
set(h,'box','off')
set(h,'fontsize',[13])
xlabel('time, min','fontsize',14,'fontweight','bold')
ylabel({'IR phosphorylation';'[a.e]'},'fontsize',14,'fontweight','bold')
axis([0 20 0 110])
box off
hold off
% a = get(gca,'XTickLabel');  
% set(gca,'XTickLabel',a,'fontsize',14,'FontWeight','bold')
% legend('1','2')

%PLOT1: IRS1 comparison with data
subplot(2,3,2);
h=gca;
hold on
plot(InTime,simIRS1,'color',tr_color,'linewidth',2)
errorbar(Time,IRS1_mean, IRS1_std,err_color,'linewidth',1.5) 
set(h,'box','off')
set(h,'fontsize',[13])
xlabel('time, min','fontsize',14,'fontweight','bold')
ylabel({'IRS1 phosphorylation';'[a.e]'},'fontsize',14,'fontweight','bold')
axis([0 20 0 110])
box off
hold off
% a = get(gca,'XTickLabel');  
% set(gca,'XTickLabel',a,'fontsize',14,'FontWeight','bold')


%% Figure 3: Module 2
%%%%%%%%%%%%%%%%%%%%%%%%%%%%%%%%%%%%%%%%%%%%%%%%%%%%%%%%%%%%%%%%%%%%%%%%%%
%PLOT4: AS160 phosphorylation in comparison with data
subplot(2,3,4);
plot((0:0.001:30),AS160p,'color',tr_color,'linewidth',2) 
hold on
errorbar(ASData.time,ASData.values,ASData.se,err_color,'linewidth',1.5)
hold on
xlabel('Time (min)','fontsize',14,'fontweight','bold')
ylabel({'AS160p';'[a.e]'},'fontsize',14,'fontweight','bold')
%title('Comparison simulation/experiment')
hold off
set(gca,'fontsize',13)
set(gca,'Xlim',[0 25])
set(gca,'Ylim',[0 1])
box off
% a = get(gca,'XTickLabel');  
% set(gca,'XTickLabel',a,'fontsize',14,'FontWeight','bold')
%PLOT5: PKB-T308 phosphorylation in comparison with data
subplot(2,3,5);
plot((0:0.001:30),PKB308,'color',tr_color,'linewidth',2) 
hold on
errorbar(PKBData.time,PKBData.values,PKBData.se,err_color,'linewidth',1.5)
xlabel('Time (min)','fontsize',14,'fontweight','bold')
ylabel({'PKB308p';'[a.e]'},'fontsize',14,'fontweight','bold')
%title('Comparison simulation/experiment')
hold off
set(gca,'fontsize',13)
set(gca,'Xlim',[0 25])
set(gca,'Ylim',[0 1])
box off
% a = get(gca,'XTickLabel');  
% set(gca,'XTickLabel',a,'fontsize',14,'FontWeight','bold')
%PLOT6: PKB-S473 phosphorylation in comparison with data
subplot(2,3,6);
plot((0:0.001:30),PKB473,'color',tr_color,'linewidth',2) 
hold on
errorbar(PKB2Data.time,PKB2Data.values,PKB2Data.se,err_color,'linewidth',1.5)
hold on
xlabel('Time (min)','fontsize',14,'fontweight','bold')
ylabel({'PKB473p';'[a.e]'},'fontsize',14,'fontweight','bold')
%title('Comparison simulation/experiment')
hold off
set(gca,'fontsize',13)
set(gca,'Xlim',[0 25])
set(gca,'Ylim',[0 1])
box off
% a = get(gca,'XTickLabel');  
% set(gca,'XTickLabel',a,'fontsize',14,'FontWeight','bold')

%%
% x=(0:0.001:30);
% lila =[0.8 0 0.6];
% %PLOT8: Fractions of GLUT4 in different states compared to data
% subplot(4,2,6);
% plot(x,c0,'b-',x,c1,'r-','linewidth',2)
% hold on
% plot(x,c2,'-','color',lila,'linewidth',2)
% xlabel('Time (min)')
% ylabel('Fraction of total GLUT4')
% title('Simulation of C0,C1,C2')
% legend('GSV-C0','Monomers-C1','Cluster-C2')
% hold on
% errorbar([0 25],[0.75 0.20],[0.05 0.05],'b*','linewidth',1)
% errorbar([0.4 24.6],[0.10 0.20],[0.05 0.05],'*','color',lila,'linewidth',1)
% errorbar([0 25],[0.10 0.40],[0.05 0.05],'r*','linewidth',1)
% set(gca,'fontsize',13)
% set(gca,'Xlim',[0 25])
% set(gca,'Ylim',[0 0.8])
% box off
% hold off
```
