## Supplementary scripts for "A multi-scale *in silico* mouse model for insulin resistance and humanoid type 2 diabetes": README.txt

IMPORTANT!: SBTOOLBOX2/SBPD/SBPOP package is required for model simulation.
http://www.sbtoolbox2.org/main.php
The SBtoolbox2 was recently replaced with IQM tools (http://www.intiquan.com/iqm-tools/)
It is possible to make the plotscript compatible with IQM tools by replacing the prefix SBPD with IQM throughout the script.
Furthermore, curve fitting toolbox is necessary. 


The folder consists of the following files:
- Plotscript
- EXPDATA (experimental data)
- FractionData (data from stretch analysis)
- paramset1 (the parameter set used when ploting the figures)
- paramsets (contains parameter sets from the stretch analysis)
- GLUTcomplete
- GLUTcompletekr
- GLUTPlosOne
- GLUTPlosOneKr
- simannealingSBAOClusteringL

The plotscript simulates the models (e.g. GLUT4complete.txt) multiple times for different insulin concentrations
and shows model output together with the corresponding experimental data.

There are 4 different model files, with the same model but events included to simulate the specific experimental details:
GLUT4complete.txt - all inputs to cluster removed at t=230min. 
GLUT4completeKr.txt - all inputs to cluster + endocytosis removed at t=230min.
GLUT4PlosOne.txt - all inputs to cluster removed at t=200min.
GLUT4PlosOneKr.txt - all inputs to cluster + endocytosis removed at t=200min.
(200 min steadystate simulation)

The plotscript is divided into parts corresponding to the different modules.

%%%%%%%%%%%%%%%%%%%%%%%%%%%%
First module : Ins = 100 nM 

Experimental data and original model:
Nyman, E., Fagerholm, S., Jullesson, D., Stralfors, P. & Cedersund, G. (2012) 
Mechanistic explanations for counter-intuitive phosphorylation dynamics of the insulin 
receptor and insulin receptor substrate-1 in response to insulin in murine adipocytes, 
The FEBS journal. 279, 987-99.

%%%%%%%%%%%%%%%%%%%%%%%%%%%%%
Second module : Ins = 28 nM. 

New experimental data by Karin G Stenkula, no model availible

%%%%%%%%%%%%%%%%%%%%%%%%%%%%%
Third module : Ins = 70 nM

Experimental data and original model:
Stenkula, K. G., Lizunov, V. A., Cushman, S. W. & Zimmerberg, J. (2010) 
Insulin controls the spatial distribution of GLUT4 on the cell surface through regulation of its postfusion dispersal, 
Cell metabolism. 12, 250-9. referred to as cell.met
More experimental data:
Lizunov, V. A., Stenkula, K., Troy, A., Cushman, S. W. & Zimmerberg, J. (2013) 
Insulin regulates Glut4 confinement in plasma membrane clusters in adipose cells, 
PloS one. referred to as plosOne

%%%%%%%%%%%%%%%%%%%%%%%%%%%%%
Validation data: Ins = 7 nM

Experimental data:
Bur�n J, Lai YC, Lundgren M, Eriksson JW, Jensen J. (2008)
Insulin action and signalling in fat and muscle from dexamethasone-treated rats. 
Arch Biochem Biophys 474: 91�101, 2008. doi:10.1016/j.abb.2008.02.034. pmid:18328801
