## Supplementary scripts for "A multi-scale *in silico* mouse model for insulin resistance and humanoid type 2 diabetes": simannealingSBAOClusteringL.m

```
function [X,FVAL,EXITFLAG] = simannealingSBAOClusteringL(FUN,X,varargin)
% simannealingSBAO: Minimization by simulated annealing. Algorithm partly
% based on section 10.4 in "Numerical Recipes in C", ISBN 0-521-43108-5.
% The implementation of this simmulated annealing method uses a nonlinear
% simplex search as basis. The algorithm has been modified in order to be
% able to take into account simple box constraints on parameter values.
%
% OBS! This algorithm have been modified due to demands to find even suboptimal
%      function minima. Hence, the algorithm have been modified from a singlestart,
%      i.e. one simplex is initiated at each temperature level in the best
%      point found so far,
%      to a multistart algorithm, i.e. multiple simplex may be initiated at each temperature level.
%      Suitable restart points is found with single-linkage clustering using an euclidian threshold.
%      Best Regards
%      Tobias Pettersson
%
%
% USAGE:
% ======
% [X,FVAL,EXITFLAG] = simannealingSBAO(FUN,X,OPTIONS)
%
% FUN: Function to optimize
% X: Starting Guess
% OPTIONS: structure containing options for the algorithm:
%        OPTIONS.tempstart: Starting temperature (should be around the
%           order of magnitude (or higher) than the cost function at the
%           initial guess)
%        OPTIONS.tempend: Ending temperature (When performed all iterations
%           for this temperature, the temperature is set to 0 and the
%           simannealingSBAO function converges to a normal simplex search)
%        OPTIONS.tempfactor: Reduction factor for temperature after running
%           through all iterations for current temperature
%        OPTIONS.maxitertemp: Number of iterations to carry put for each
%           non-zero temperature
%        OPTIONS.maxitertemp0: Number of iterations to carry out for 0
%           temperature
%        OPTIONS.maxtime: Maximum time (in minutes) for optimization
%        OPTIONS.tolx: Tolerance for max difference between the coordinates
%           of the vertices.
%        OPTIONS.tolfun: Tolerance for difference between best and worst
%           function evaluation in simplex
%        OPTIONS.highbounds: vector containing upper bounds for parameters
%           Instead of a vector highbounds can also be a scalar > 1. In the
%           latter case the highbounds are determined by multiplying this
%           scalar to the initial parameter guess X. This latter case works
%           only for positive X(i) ... if negative present then the user
%           needs to specify a whole lowbounds vector.
%        OPTIONS.lowbounds: vector containing lower bounds for parameters.
%           Instead of a vector lowbounds can also be a scalar < 1. In the
%           latter case the lowbounds are determined by multiplying this
%           scalar to the initial parameter guess X. This latter case works
%           only for positive X(i) ... if negative present then the user
%           needs to specify a whole highbounds vector.
%        OPTIONS.outputFunction: string with output function name. If
%           not given or if empty then no output function will be used.
%           This output function can be used to display data, or to
%           save optimization information to a file. The function needs
%           to be in the MATLAB path. Its calling syntax is:
%                   'outputFunction'(bestparameters,bestfunctionvalue,currentsimplex)
%        OPTIONS.silent: =0: output of info, =1: no output
%        OPTIONS.MaxRestartPoints: constant which specify the maximum number of points
%           to restart the algorithm in at each temperature level.
%
%
% DEFAULT VALUES:
% ===============
% OPTIONS.tempstart = 10*magnitude of function value at initial guess
% OPTIONS.tempend = 0.1
% OPTIONS.tempfactor = chosen such that 10 reductions of temperature
% OPTIONS.maxitertemp = 50*numberVariables
% OPTIONS.maxitertemp0 = 200*numberVariables
% OPTIONS.maxtime = 120
% OPTIONS.tolx = 1e-10
% OPTIONS.tolfun = 1e-10
% OPTIONS.lowbounds:    0.1  => lowbounds = 0.1*X
% OPTIONS.highbounds:    10  => highbounds = 10*X
% OPTIONS.outputFunction: no output function ('')
% OPTIONS.silent:         0 (no output of info)
% OPTIONS.MaxRestartPoints = 10;
%
% Output Arguments:
% =================
% X: Found solution
% FVAL: Value of the function FUN at X
% EXITFLAG: 1=success, 0=not found

% Information:
% ============
% SBaddon for the Systems Biology Toolbox
% Copyright 2006 by Fraunhofer-Chalmers Research Centre, Gothenburg, Sweden
% All rights reserved.
%
% The main author of SBaddon is Henning Schmidt.


% seed the random number generator
rand('state',sum(100*clock));

%%%%%%%%%%%%%%%%%%%%%%%%%%%%%%%%%%%%%%%%%%%%
% GLOBAL VARIABLES
%%%%%%%%%%%%%%%%%%%%%%%%%%%%%%%%%%%%%%%%%%%%
global ndim nfunk Xguess lowbounds highbounds Temp ybest pbest tempstart tempfactor MaxRestartPoints

%%%%%%%%%%%%%%%%%%%%%%%%%%%%%%%%%%%%%%%%%%%%
% Start
%%%%%%%%%%%%%%%%%%%%%%%%%%%%%%%%%%%%%%%%%%%%
if nargin == 2,
    OPTIONS = [];
elseif nargin == 3,
    OPTIONS = varargin{1};
else
    error('Incorrect number input of arguments.');
end

%%%%%%%%%%%%%%%%%%%%%%%%%%%%%%
% Set default values
%%%%%%%%%%%%%%%%%%%%%%%%%%%%%%
EXITFLAG = 1;
ndim = length(X);
tempstart = -1;     % default value calculation later
tempend = 0.1;
tempfactor = -1;    % default value calculation later
maxitertemp = 50*ndim;
maxitertemp0 = 200*ndim;
maxtime = 120;
tolx = 1e-10;
tolfun = 1e-10;
lowbounds = 0.1*X;
highbounds = 10*X;
outputFunction = '';
silent = 0;
sigma = 4;
MaxRestartPoints =10;
%%%%%%%%%%%%%%%%%%%%%%%%%%%%%%%%%%%%%%%%%%%%
% HANDLE OPTIONS
%%%%%%%%%%%%%%%%%%%%%%%%%%%%%%%%%%%%%%%%%%%%
% silent
if isfield(OPTIONS,'silent'),
    if ~isempty(OPTIONS.silent),
        silent = OPTIONS.silent;
    end
end
% tolfun
if isfield(OPTIONS,'tolfun'),
    if ~isempty(OPTIONS.tolfun),
        tolfun = OPTIONS.tolfun;
    end
end
% tolx
if isfield(OPTIONS,'tolx'),
    if ~isempty(OPTIONS.tolx),
        tolx = OPTIONS.tolx;
    end
end
% maxtime
if isfield(OPTIONS,'maxtime'),
    if ~isempty(OPTIONS.maxtime),
        maxtime = OPTIONS.maxtime;
    end
end
% maxitertemp0
if isfield(OPTIONS,'maxitertemp0'),
    if ~isempty(OPTIONS.maxitertemp0),
        maxitertemp0 = OPTIONS.maxitertemp0;
    end
end
% maxitertemp
if isfield(OPTIONS,'maxitertemp'),
    if ~isempty(OPTIONS.maxitertemp),
        maxitertemp = OPTIONS.maxitertemp;
    end
end
% tempfactor
if isfield(OPTIONS,'tempfactor'),
    if ~isempty(OPTIONS.tempfactor),
        tempfactor = OPTIONS.tempfactor;
    end
end
% tempend
if isfield(OPTIONS,'tempend'),
    if ~isempty(OPTIONS.tempend),
        tempend = OPTIONS.tempend;
    end
end
% tempstart
if isfield(OPTIONS,'tempstart'),
    if ~isempty(OPTIONS.tempstart),
        tempstart = OPTIONS.tempstart;
    end
end
% lowbounds
if isfield(OPTIONS,'lowbounds'),
    if ~isempty(OPTIONS.lowbounds),
        if length(OPTIONS.lowbounds) == ndim,
            lowbounds = OPTIONS.lowbounds;
        else
            if length(OPTIONS.lowbounds) == 1,
                if ~isempty(find(X<0)),
                    error('Please specify a vector for the lower bounds using OPTIONS.lowbounds.');
                end
                lowbounds = X*OPTIONS.lowbounds;
            else
                error('The OPTIONS.lowbounds setting is not correct.');
            end
        end
    end
else
    if ~isempty(find(X<0)),
        error('Please specify a vector for the lower bounds using OPTIONS.lowbounds.');
    end
end
% highbounds
if isfield(OPTIONS,'highbounds'),
    if ~isempty(OPTIONS.highbounds),
        if length(OPTIONS.highbounds) == ndim,
            highbounds = OPTIONS.highbounds;
        else
            if length(OPTIONS.highbounds) == 1,
                if ~isempty(find(X<0)),
                    error('Please specify a vector for the higher bounds using OPTIONS.highbounds.');
                end
                highbounds = X*OPTIONS.highbounds;
            else
                error('The OPTIONS.highbounds setting is not correct.');
            end
        end
    end
else
    if ~isempty(find(X<0)),
        error('Please specify a vector for the higher bounds using OPTIONS.highbounds.');
    end
end
lowbounds = lowbounds(:)';
highbounds = highbounds(:)';
% outputFunction
outputFunction = '';
if isfield(OPTIONS,'outputFunction'),
    if ~isempty(OPTIONS.outputFunction),
        outputFunction = OPTIONS.outputFunction;
    end
end

%%%%%%%%%%%%%%%%%%%%%%%%%%%%%%%%%%%%%%%%%%%%
% CHECK BOUND VIOLATION FOR INITIAL GUESS
%%%%%%%%%%%%%%%%%%%%%%%%%%%%%%%%%%%%%%%%%%%%
indexXhi = find(X > highbounds);
if ~isempty(indexXhi),
    error('Initial guess does violate high parameter bounds.');
end
indexXlo = find(X < lowbounds);
if ~isempty(indexXlo),
    error('Initial guess does violate low parameter bounds.');
end

%%%%%%%%%%%%%%%%%%%%%%%%%%%%%%%%%%%%%%%%%%%%
% INITIALIZE EMPTY SIMPLEX DATA
%%%%%%%%%%%%%%%%%%%%%%%%%%%%%%%%%%%%%%%%%%%%
p = zeros(ndim+1,ndim);     % vertice vectors in rows
y = zeros(ndim+1,1);


%%%%%%%%%%%%%%%%%%%%%%%%%%%%%%%%%%%%%%%%%%%%
% CALCULATE THE CLUSTERING VARIABLES
%%%%%%%%%%%%%%%%%%%%%%%%%%%%%%%%%%%%%%%%%%%%
Lebesgue = prod(highbounds-lowbounds);
NumberOfTempSteps = 0;

if isfield(OPTIONS,'sigma'),
    if ~isempty(OPTIONS.sigma),
        sigma = OPTIONS.sigma;
    end
end

if isfield(OPTIONS,'MaxRestartPoints'),
    if ~isempty(OPTIONS.MaxRestartPoints),
        MaxRestartPoints = OPTIONS.MaxRestartPoints;
    end
end


%%%%%%%%%%%%%%%%%%%%%%%%%%%%%%%%%%%%%%%%%%%%
% EVALUATE FUNCTION IN INITIAL GUESS
%%%%%%%%%%%%%%%%%%%%%%%%%%%%%%%%%%%%%%%%%%%%
pbest = ones(MaxRestartPoints+1,length(X));
pbest(1,:) = X(:)';
ybest = inf*ones(MaxRestartPoints+1,1);
ytemp = costFunction(FUN,pbest(1,:), 1);
ybest=ytemp*ones(MaxRestartPoints+1,1);
restartpoints = X(:)';
fpoints  = [];

%%%%%%%%%%%%%%%%%%%%%%%%%%%%%%%%%%%%%%%%%%%%
% DETERMINE START TEMP AND TEMPFACTOR (if not given by the user)
%%%%%%%%%%%%%%%%%%%%%%%%%%%%%%%%%%%%%%%%%%%%
if tempstart < 0,
    % tempstart not given by the user => determine here a default value
    tempstart = 10*abs(ybest(1));
end
if tempfactor < 0,
    % tempfactor not given by the user => determine here a default value
    tempfactor = (tempstart/tempend)^(-1/9);
end

%%%%%%%%%%%%%%%%%%%%%%%%%%%%%%%%%%%%%%%%%%%%
% TEMPERATURE LOOP - STARTING FROM BEST POINT
%%%%%%%%%%%%%%%%%%%%%%%%%%%%%%%%%%%%%%%%%%%%
Temp = tempstart;
tic % start timer
nfunk = 1;
nriterations = 0;
while(1),
    K = size(restartpoints,1);
    NumberOfTempSteps = NumberOfTempSteps +1;
    
    
    if Temp < tempend,
        Temp = 0;
        
        MAXITERTEMP = floor(maxitertemp0/K); %Ska det väl vara? Inte maxitertemp0 direkt, då blir ju aldrig klar (kommentar Gunnar ;)
    else
        MAXITERTEMP = floor(maxitertemp/K);
    end
test= find(ybest(:)==min(ybest));
    disp(sprintf(' Current best estimation: [%s]',sprintf('%g ', pbest(test(1),:))));
    
    %%%%%%%%%%%%%%%%%%%%%%%%%%%%%%%%%%%%%%%%%%%%
    % FIRST POINT OF INITIAL SIMPLEX
    %%%%%%%%%%%%%%%%%%%%%%%%%%%%%%%%%%%%%%%%%%%%
    evaluatedpoints = zeros(MAXITERTEMP, ndim);
    evaluatedfuncvalues = zeros(MAXITERTEMP,1);
    allpoints = [];
    allfvalues = [];
    indexlist = zeros(K,1);
    
    for ii = 1:K,
        
        disp(sprintf(' The algorithm restarts at: [%s]',sprintf('%g ', restartpoints(ii,:))));
        
        Xguess = restartpoints(ii,:);
        p(1,:) = Xguess;
        y(1) = costFunction(FUN,Xguess, ii);
        if ~silent,
            disp(' Nr Iter  Nr Fun Eval    Min function       Best function       Temp      Algorithm Step');
            if Temp == tempstart,
                disp(sprintf(' %5.0f   %5.0f       %12.6g       %12.6g   %12.6g       %s', nriterations, nfunk, y(1), ybest(ii), Temp, 'initial guess'));
            else
                disp(sprintf(' %5.0f   %5.0f       %12.6g       %12.6g   %12.6g       %s', nriterations, nfunk, y(1), ybest(ii), Temp, 'best point'));
            end
        end
        
        
        %%%%%%%%%%%%%%%%%%%%%%%%%%%%%%%%%%%%%%%%%%%%
        % REMAINING POINTS OF INITIAL SIMPLEX
        %%%%%%%%%%%%%%%%%%%%%%%%%%%%%%%%%%%%%%%%%%%%
        % construct vertices by modifying one element each
        % relative changes in case that elements are non-zero,
        % absolute changes in case that elements are zero
        relativeDelta = 0.25;
        absoluteDelta = 0.5;
        for k = 1:ndim
            Xmodify = Xguess;
            if Xmodify(k) == 0
                % absolute change
                if highbounds(k) > absoluteDelta,
                    Xmodify(k) = absoluteDelta;
                else
                    Xmodify(k) = -absoluteDelta;
                end
            else
                % relative change
                Xmodify(k) = (1 + relativeDelta)*Xmodify(k);
            end
            p(k+1,:) = Xmodify;
            y(k+1) = costFunction(FUN,Xmodify, ii);
        end
        algostep = 'initial simplex';
        nriterations = nriterations + 1;
        nfunk = nfunk + ndim + 1;
        
        % if output function given then run output function to plot
        % intermediate result
        if length(outputFunction) ~= 0,
            feval(outputFunction,p(1,:), y(1),p);
        end
        
        
        
        %%%%%%%%%%%%%%%%%%%%%%%%%%%%%%%%%%%%%%%%%%%%
        % MAIN ALGORITHM
        %%%%%%%%%%%%%%%%%%%%%%%%%%%%%%%%%%%%%%%%%%%%
        % reorder y and p so that the first row corresponds to the
        % lowest function value (after the temperature noise has been added, Gunnar's comment)
        for itertemp = 1:MAXITERTEMP,
            % add random thermal fluctuations
            yfluct = y + Temp*abs(log(rand(ndim+1,1)));
            % do sorting instead of determining the indices of the best, worst,
            % next worst
            help = sortrows([yfluct,y,p],1);
            yfluct = help(:,1);
            y = help(:,2);
            p = help(:,3:end);
            if ~silent,
                disp(sprintf(' %5.0f   %5.0f       %12.6g       %12.6g   %12.6g       %s', nriterations, nfunk, y(1), ybest(ii), Temp, algostep));
            end
            % if output function given then run output function to plot
            % intermediate result
            if length(outputFunction) ~= 0,
                feval(outputFunction,p(1,:), y(1),p);
            end
            % end the optimization if the difference between best and worst
            % function evaluation in simplex is smaller than tolfun and the
            % max difference between the coordinates of the verttices is less than
            % tolx
            if abs(max(y)-min(y)) < tolfun && max(max(abs(p(2:ndim+1)-p(1:ndim)))) < tolx,
                break;
            end
            % check number of iterations
            if toc/60 > maxtime,
                EXITFLAG = 0;
                disp('Exceeded maximum time.');
                break;
            end
            
            % Begin a new iteration. First extrapolate by a factor -1 through the face of the simplex
            % across from the high point, i.e., reflect the simplex from the high point.
            [yftry, ytry,ptry] = amotry(FUN, p, -1, ii);
            % check the result
            if yftry <= yfluct(1),
                % Gives a result better than the best point, so try an additional
                % extrapolation by a factor 2.
                [yftryexp, ytryexp,ptryexp] = amotry(FUN, p, -2, ii);
                if yftryexp < yftry,
                    p(end,:) = ptryexp;
                    y(end) = ytryexp;
                    algostep = 'extrapolation';
                else
                    p(end,:) = ptry;
                    y(end) = ytry;
                    algostep = 'reflection';
                end
            elseif yftry >= yfluct(ndim),
                % The reflected point is worse than the second-highest, so look
                % for an intermediate lower point, i.e., do a one-dimensional
                % contraction.
                [yftrycontr,ytrycontr,ptrycontr] = amotry(FUN, p, -0.5, ii);
                if yftrycontr < yfluct(end),
                    p(end,:) = ptrycontr;
                    y(end) = ytrycontr;
                    algostep = 'one dimensional contraction';
                else
                    % Can�t seem to get rid of that high point. Better contract
                    % around the lowest (best) point.
                    x = ones(ndim,ndim)*diag(p(1,:));
                    p(2:end,:) = 0.5*(p(2:end,:)+x);
                    for k=2:ndim,
                        y(k) = costFunction(FUN,p(k,:), ii);
                    end
                    algostep = 'contraction around best point';
                end
            else
                % if ytry better than second-highest point then use this point
                p(end,:) = ptry;
                y(end) = ytry;
                algostep = 'reflection';
            end
            nriterations = nriterations + 1;
            
            % Save the resulting value of the downhill simplex.
            evaluatedpoints(itertemp,:,ii) = p(end,:);
            evaluatedfuncvalues(itertemp,ii) = y(end);
            indexlist(ii) = indexlist(ii) +1;
        end
        
    end
    
    
    for jj = 1:K,
        allpoints = [allpoints; evaluatedpoints(:,:,jj)];
        allfvalues = [allfvalues; evaluatedfuncvalues(:,jj)];
    end
    
    % Break due to max number iterations, max time, minimum found
    % (tolerances)
    if itertemp < MAXITERTEMP,
        break;
    end
    Temp = Temp*tempfactor;
    % Break because 0 temperature has been run
    if Temp == 0,
        break;
    end
    
    % Calculate the euclidian distance used as critical distance for the clustering
    r = (pi)^(-0.5)*(gamma(1+0.5*ndim)*Lebesgue*(sigma*log(maxitertemp*NumberOfTempSteps))/...
        (NumberOfTempSteps*maxitertemp))^(1/ndim);
    
    % Find suitable restartpoints to use for next temperature level
    restartpoints = clusterpoints(allpoints, allfvalues, r);
    
end


% % Gather the best points from each existing simplex
% for kk = 1:K,
%     foundpoints(kk,:) = evaluatedpoints(indexlist(kk),:,kk);
%     foundfuncval(kk) = evaluatedfuncvalues(indexlist(kk),kk);
% end

% Sort the points with respect to quality
helpv = sortrows([ybest(:) pbest(:,:)],1);


X = helpv(:,2:end);
FVAL = helpv(:,1);

return


%%%%%%%%%%%%%%%%%%%%%%%%%%%%%%%%%%%%%%%%%%%%
% Use single-linkage clustering to find
% suitable restart points to use at next
% temperature level
%%%%%%%%%%%%%%%%%%%%%%%%%%%%%%%%%%%%%%%%%%%%
function [foundpoints] = clusterpoints(points, fvalues, r);

global MaxRestartPoints

% Initiations
foundpoints = [];

% Sort the points with respect to function value
help = sortrows([fvalues, points],1);

sfvalues = help(:,1);
spoints = help(:,2:end);

for ii = 1:size(spoints,1),
    % The best point is an obvious choice
    if ii == 1,
        foundpoints = [foundpoints; spoints(ii,:)];
        
        % include a point if there is not better point within the critical distance.
    elseif min(sqrt(sum((repmat(spoints(ii,:),ii-1,1)-spoints(1:ii-1,:)).^2,2))) > r,
        foundpoints = [foundpoints; spoints(ii,:)];
    end
    
    % Break if the number of restartpoints are greater than
    % the prescribed limit
    if size(foundpoints,1) > MaxRestartPoints,
        break;
    end
end
return


%%%%%%%%%%%%%%%%%%%%%%%%%%%%%%%%%%%%%%%%%%%%
% AMOTRY FUNCTION
%%%%%%%%%%%%%%%%%%%%%%%%%%%%%%%%%%%%%%%%%%%%
function [yftry,ytry,ptry] = amotry(FUN, p, fac,iter)
% Extrapolates by a factor fac through the face of the simplex across from
% the high point, tries it, and replaces the high point if the new point is
% better.
global ndim nfunk Temp lowbounds highbounds tempstart
psum = sum(p(1:ndim,:))/ndim;
ptry = psum*(1-fac) + p(end,:)*fac;

% Deal with low and high parameter bounds
indexXhi = find(ptry > highbounds);
indexXlo = find(ptry < lowbounds);
for k=1:length(indexXhi),
    ptry(indexXhi(k)) = highbounds(indexXhi(k))-rand(1)*(highbounds(indexXhi(k))-lowbounds(indexXhi(k)))*Temp/tempstart;
end
for k=1:length(indexXlo),
    ptry(indexXlo(k)) = lowbounds(indexXlo(k))+rand(1)*(highbounds(indexXlo(k))-lowbounds(indexXlo(k)))*Temp/tempstart;
end

% Evaluate the function at the trial point.
ytry = costFunction(FUN,ptry,iter);
yftry = ytry - Temp*abs(log(rand(1)));
nfunk = nfunk + 1;
return

%%%%%%%%%%%%%%%%%%%%%%%%%%%%%%%%%%%%%%%%%%%%
% COST FUNCTION EVALUATION
%%%%%%%%%%%%%%%%%%%%%%%%%%%%%%%%%%%%%%%%%%%%
function [ytry] = costFunction(FUN,ptry,iter)
global ybest pbest lowbounds highbounds
ytry = feval(FUN,ptry);
% save the best point ever (only if it is feasible, that is it fits the
% high and low bounds)
indexXhi = find(ptry > highbounds);
indexXlo = find(ptry < lowbounds);
if ytry < ybest(iter) && isempty(indexXhi) && isempty(indexXlo)
    ybest(iter) = ytry;
    pbest(iter,:) = ptry;
end
return
```
