## Supplementary scripts for "A multi-scale *in silico* mouse model for insulin resistance and humanoid type 2 diabetes": cost_function.m

```
function[cost]= cost_function( theta ,func_mex_model, D, utility)
ic0 = utility.ic0;
p = utility.p0;
objModel = utility.objModel; % variables used for indexing
% if nargin <9, e='hfd_6w_IPGTT'; end
% if nargin <10, x=120/(60*24)+56+0.5833; end

adhoc_cost =  0; % Should be reset every call to the objective function
pNames = utility.pNames ; 
id_FFM = ismember(SBstates(objModel),'FFM') ; id_FM = ismember(SBstates(objModel),'FM');
id_FFMname = ismember(pNames,'FFMinit') ; id_FMname = ismember(pNames,'FMinit');
% parameter and time vector setup
p(17:37)=[10.^theta(8:19) theta(20)  10.^theta(21:end)];  % GSS to ik1
if p(ismember(pNames,'ik1')) == 10^theta(end) % bounds check
else
    disp('check parameter vector')
end
try   
%% Steady state simulation
ic_ss = ic0; p_ss = p; p_ss( ismember(pNames, 'ss_x')) = 0 ;
ic_ss(id_FFM)=1 ; ic_ss(id_FM)= 1; p_ss(id_FFMname)=1; p_ss(id_FMname)=1; % no effect of Weight on tissue-level and a i s- models
p_ss(end) = 1000; %no IPGTT
ss=func_mex_model(0:100:300,ic_ss, p_ss); % steady state simulation
cost_ss_gcc = (( ss.variablevalues(end, end-1)  -  D.IPGTTgcc.Mean(1,1)  )^2 )./( D.IPGTTgcc.SEM(1,1) )^2 ;% basal glucose
cost_ss_icc = (( ss.variablevalues(end, end)  -  D.IPGTTicc.Mean(1,1)  )^2 )./( D.IPGTTicc.SEM(1,1) )^2   ;% basal insulin
ss_cost = cost_ss_gcc + cost_ss_icc;
ic0 = ss.statevalues(end,:); % overwrite ic0

%% Hansson et al (2018) 2w - Diet and IPGTT 
% Setting up parameters and time-vectors
t_diet=14; % diet time in days
t_fasting=[ t_diet  t_diet+0.5833 ]  ;  %fasting
t_IPGTT=(0:1:120)/(60*24)+t_fasting(end);
p(end) = t_fasting(end); % time of IPGTT (triggers after fasting)

% 2w chow
pchow=p;
pchow(1)=10.^theta(5);   % EI   theta(5) - Chow EI
pchow(2)=0.7;            % carb fraction 
pchow(16)= 10.^theta(1); % PA for chow
pchow(id_FFMname)=0.89*D.BW.mean(1,1); pchow(id_FMname)=0.11*D.BW.mean(1,1);
ic_chow = ic0; ic_chow(id_FFM)=0.89*D.BW.mean(1,1) ; ic_chow(id_FM)= 0.11*D.BW.mean(1,1);

% 12 d chow + 2d hfd
pchow12d=p; 
pchow12d(1)=18.1845; 	
pchow12d(2)=0.2;  
pchow12d(16)  = 10.^theta(2); % PA for hfd
% Here we dont have any data for starting weight.

% 10d chow + 4d hfd
pchow10d=p; 
pchow10d(1)=18.9311;    
pchow10d(2)=0.2;  
pchow10d(16)= 10.^theta(2); % PA for hfd
pchow10d(id_FFMname)=0.89*D.BW.mean(2,1); pchow10d(id_FMname)=0.11*D.BW.mean(2,1); 
ic_chow10d = ic0; ic_chow10d(id_FFM)=0.89*D.BW.mean(2,1) ; ic_chow10d(id_FM)= 0.11*D.BW.mean(2,1);

% 8d chow + 6d hfd
pchow8d=p;  
pchow8d(1)=18.1845;     
pchow8d(2)=0.2;   
pchow8d(16)= 10.^theta(2);  % PA for hfd
pchow8d(id_FFMname)=0.89*D.BW.mean(3,1);    pchow8d(id_FMname)=0.11*D.BW.mean(3,1);
ic_chow8d = ic0; ic_chow8d(id_FFM)=0.89*D.BW.mean(3,1) ; ic_chow8d(id_FM)= 0.11*D.BW.mean(3,1);

% 2w hfd
phfd=p;     
phfd(1)= 16.5217;       
phfd(2)=0.2;      
phfd(16)= 10.^theta(2);    % PA for hfd
phfd(id_FFMname)=0.89*D.BW.mean(4,1);    phfd(id_FMname)=0.11*D.BW.mean(4,1);
ic_hfd = ic0; ic_hfd(id_FFM)=0.89*D.BW.mean(4,1) ; ic_hfd(id_FM)= 0.11*D.BW.mean(4,1);

%%% Diet simulations %%%%
%2w chow
chow2w=func_mex_model( (0:1:t_diet),ic_chow, pchow);
% 12 d chow + 2d hfd
% Here we dont have any data for starting weight. Using values from 2 chow
% simulation
ic_chow12d = chow2w.statevalues(13,:) ; % 13 is index for day 12
ic_chow12d(id_FFM)=chow2w.statevalues(13,id_FFM) ; ic_chow12d(id_FM)= chow2w.statevalues(13,id_FM);
pchow12d(id_FFMname)=chow2w.statevalues(13,id_FFM); pchow12d(id_FMname)= chow2w.statevalues(13,id_FM);
chow12dHFD2d=func_mex_model((12:1:t_diet) ,ic_chow12d, pchow12d);
%10d chow + 4d HFD 
chow10dHFD4d=func_mex_model((10:1:t_diet) ,ic_chow10d, pchow10d);  
%8d chow + 6d HFD  
chow8dHFD6d=func_mex_model((8:1:t_diet) ,ic_chow8d, pchow8d); 
%2w hfd   
hfd2w=func_mex_model((0:1:t_diet),ic_hfd, phfd );   

%%% Fasting + IPGTT simulations 
ss_chow=func_mex_model( t_fasting, chow2w.statevalues(end,:), pchow);  % chow
IPGTT_chow=func_mex_model( t_IPGTT , ss_chow.statevalues(end,:), pchow ); 
ss_chow12dHFD2d=func_mex_model( t_fasting, chow12dHFD2d.statevalues(end,:), pchow12d);  % %12d chow + 2d HFD  
IPGTT_chow12dHFD2d=func_mex_model( t_IPGTT , ss_chow12dHFD2d.statevalues(end,:), pchow12d ); 
ss_chow8dHFD6d=func_mex_model( t_fasting, chow8dHFD6d.statevalues(end,:), pchow8d);  % %8d chow + 6d HFD  
IPGTT_chow8dHFD6d=func_mex_model( t_IPGTT , ss_chow8dHFD6d.statevalues(end,:), pchow8d ); 
ss_hfd14d=func_mex_model( t_fasting, hfd2w.statevalues(end,:), phfd);  % hfd
IPGTT_hfd14d=func_mex_model( t_IPGTT , ss_hfd14d.statevalues(end,:), phfd ); 

%%% Calculating cost
%%% Body weight cost
bw_cost1= nansum (  ( (chow2w.variablevalues([1,end],1)'   -  D.BW.mean(1,:) ).^2 )./( D.BW.SEM(1,end).^2 )); % chow
%bw_cost2= nansum (  ( (chow12dHFD2d.variablevalues([1,end],1)'   -  D.BW.mean(4,:) ).^2 )./( D.BW.SEM(4,:).^2 )); % % 2 d HFD 
bw_cost3= nansum (  ( (chow10dHFD4d.variablevalues(end,1)'   -  D.BW.mean(2,end) ).^2 )./( D.BW.SEM(2,end).^2 )); % 4 d HFD 
bw_cost4= nansum (  ( (chow8dHFD6d.variablevalues(end,1)'   -  D.BW.mean(3,end) ).^2 )./( D.BW.SEM(3,end).^2 )); % 6 d HFD 
bw_cost5= nansum (  ( (hfd2w.variablevalues([1,end],1)'   -  D.BW.mean(4,:) ).^2 )./( D.BW.SEM(4,end).^2 )); % 14 d HFD
cost_bw=bw_cost1 + bw_cost3  + bw_cost4 + bw_cost5 ;
%%% IPGTT cost ( Insulin, only basal, and Glucose data points)
id_gcc = [ 1 16 31 61 121 ];
id_icc = [ 1 16 121 ]; % only first data point.
yexp = D.IPGTTgcc.Mean(:,1) ; SEM=D.IPGTTgcc.SEM(:,1); ysim= IPGTT_chow.variablevalues(id_gcc, end-1);   %14d chow
IPGTT_glucose_cost1 =  nansum (  ( (yexp  -  ysim ).^2 )./( SEM.^2 ));  
yexp = D.IPGTTicc.Mean(:,1) ; SEM=D.IPGTTicc.SEM(:,1); ysim= IPGTT_chow.variablevalues(id_icc, end);   
IPGTT_insulin_cost1 =  nansum (  ( (yexp  -  ysim ).^2 )./( SEM.^2 ));

yexp = D.IPGTTgcc.Mean(:,2) ; SEM=D.IPGTTgcc.SEM(:,2); ysim= IPGTT_chow12dHFD2d.variablevalues(id_gcc, end-1);  % 12d chow + 2d HFD      
IPGTT_glucose_cost2 =  nansum (  ( (yexp  -  ysim ).^2 )./( SEM.^2 ));  
yexp = D.IPGTTicc.Mean(:,2) ; SEM=D.IPGTTicc.SEM(:,2); ysim= IPGTT_chow12dHFD2d.variablevalues(id_icc, end);   
IPGTT_insulin_cost2 =  nansum (  ( (yexp  -  ysim ).^2 )./( SEM.^2 ));

yexp = D.IPGTTgcc.Mean(:,3) ; SEM=D.IPGTTgcc.SEM(:,3); ysim= IPGTT_chow8dHFD6d.variablevalues(id_gcc, end-1);  % 8d chow + 6d HFD           
IPGTT_glucose_cost3 =  nansum (  ( (yexp  -  ysim ).^2 )./( SEM.^2 ));  
yexp = D.IPGTTicc.Mean(:,3) ; SEM=D.IPGTTicc.SEM(:,3); ysim= IPGTT_chow8dHFD6d.variablevalues(id_icc, end);   
IPGTT_insulin_cost3 =  nansum (  ( (yexp  -  ysim ).^2 )./( SEM.^2 ));

yexp = D.IPGTTgcc.Mean(:,4) ; SEM=D.IPGTTgcc.SEM(:,4); ysim= IPGTT_hfd14d.variablevalues(id_gcc, end-1);    %14d HFD
IPGTT_glucose_cost4 =  nansum (  ( (yexp  -  ysim ).^2 )./( SEM.^2 ));  
yexp = D.IPGTTicc.Mean(:,4) ; SEM=D.IPGTTicc.SEM(:,4); ysim= IPGTT_hfd14d.variablevalues(id_icc, end);   
IPGTT_insulin_cost4 =  nansum (  ( (yexp  -  ysim ).^2 )./( SEM.^2 ));

cost_IPGTT=  (IPGTT_glucose_cost1 + IPGTT_insulin_cost1) + (IPGTT_glucose_cost2 + IPGTT_insulin_cost2)...
           + (IPGTT_glucose_cost3 + IPGTT_insulin_cost3)  + (IPGTT_glucose_cost4 + IPGTT_insulin_cost4);
       
%% Unpublished diet data - 8w chow/hfd -  Stenkula et al 
t_diet_6w=56; % 2 weeks % diet time in days ( only training to the first two weeks)
% chow
pchow6w=p; 
pchow6w(1)= 10.^theta(6); % EI intake for 8w chow
pchow6w(2)=0.7; 
pchow6w(16)=  10.^theta(3); % PA for this 8w chow    
pchow6w(id_FFMname)=D.BW_unpublished_chow.Mean(1)*0.89;
pchow6w(id_FMname)= D.BW_unpublished_chow.Mean(1)*0.11;
ic_chow6w=ic0;
ic_chow6w( ismember(SBstates(objModel),'FFM')) = D.BW_unpublished_chow.Mean(1)*0.89;
ic_chow6w( ismember(SBstates(objModel),'FM')) = D.BW_unpublished_chow.Mean(1)*0.11;
% hfd
phfd6w=p;
phfd6w(1)=10.^theta(7); % EI for 8w HFD
phfd6w(2)=0.2;  
phfd6w(16)=  10.^theta(4); % setting up hfd specifc parameters
phfd6w(id_FFMname)=D.BW_unpublished_hfd.Mean(1)*0.89;
phfd6w(id_FMname)=D.BW_unpublished_hfd.Mean(1)*0.11;
ic_hfd6w=ic0;
ic_hfd6w( ismember(SBstates(objModel),'FFM')) = D.BW_unpublished_hfd.Mean(1)*0.89;
ic_hfd6w( ismember(SBstates(objModel),'FM')) = D.BW_unpublished_hfd.Mean(1)*0.11;
% simulations
chow_6w=func_mex_model( 0:1:t_diet_6w,ic_chow6w, pchow6w); % simulation 8 weeks chow diet
hfd_6w=func_mex_model((0:1:t_diet_6w),ic_hfd6w, phfd6w );     % simulation 8 weeks hfd diet
% cost - 2w BW
bw_cost1_up= nansum (  ( (chow_6w.variablevalues([1 8 15],1)'   -  D.BW_unpublished_chow.Mean(1:3) ).^2 )./( D.BW_unpublished_chow.SEM(1:3).^2 )); % chow
bw_cost2_up= nansum (  ( (hfd_6w.variablevalues([1 8 15],1)'   -  D.BW_unpublished_hfd.Mean(1:3) ).^2 )./( D.BW_unpublished_hfd.SEM(1:3).^2 )); % chow
bw_cost_up = bw_cost1_up + bw_cost2_up;

t_fasting_up=[ t_diet_6w  t_diet_6w+0.5833 ]  ; % fasting at 6 weeks
t_IPGTT_up=(0:1:120)/(60*24)+t_fasting_up(end);
% Prediction simulations for IPGTT
phfd6w(end) = t_fasting_up(end); % set time for IPGTT event after 6 weeks + fasting
hfd_6w_fast=func_mex_model( t_fasting_up, hfd_6w.statevalues(42,:), phfd6w);  % simulation  fasting before IPGTT at 6 weeks
hfd_6w_IPGTT=func_mex_model( t_IPGTT_up , hfd_6w_fast.statevalues(end,:), phfd6w ); % IPGTT 6 weeks

%% in vitro - primary adipcoytes - insulin signaling
t_diet=28; % 4 weeks
t_fasting=[ t_diet  t_diet+0.5833 ]  ;  % 14 hours fasting
t_in_vitro= [t_fasting(end) t_fasting(end) + (1/(24*60) )*30 ]; % 30 min (min expressed as days)
p(end) = t_fasting(end) + 1000; % no IPGTT

pchow_vitro=p; 
pchow_vitro(1)=10.^theta(5); 
pchow_vitro(2)=0.7; 
pchow_vitro(16)= 10.^theta(1); % using same BW parameters as Hanson et al 2018 simulations

phfd_vitro=p;  
phfd_vitro(1)= 16.5217; 
phfd_vitro(2)=0.2; 
phfd_vitro(16)= 10.^theta(2); % using same BW parameters as Hanson et al 2018 simulations
% Here we use Ic0 directley which use a start BW of 23g divded with 89/11 %
%fat/lean mass

%%% simulations 
chow4w=func_mex_model( (0:1:t_diet),ic0, pchow_vitro); %chow
hfd4w=func_mex_model((0:1:t_diet),ic0, phfd_vitro );  %HFD 14 days simulation
pchow_vitro(ismember(pNames,'Insact'))=0; % remvoing effect of I_E in vivo insulin
phfd_vitro(ismember(pNames,'Insact'))=0; 
ss_chow=func_mex_model( t_fasting, chow4w.statevalues(end,:), pchow_vitro);  
ss_hfd14d=func_mex_model( t_fasting, hfd4w.statevalues(end,:), phfd_vitro); 
pchow_vitro(ismember(pNames,'ins'))=0.1; % nM , experiment insulin dose
phfd_vitro(ismember(pNames,'ins'))=0.1; 
vitro_chow=func_mex_model( t_in_vitro , ss_chow.statevalues(end,:), pchow_vitro ); 
vitro_hfd14d=func_mex_model( t_in_vitro , ss_hfd14d.statevalues(end,:), phfd_vitro ); 

id_PKB_phos473 = ismember(SBvariables(objModel),'totalPKB473'); 
id_PKB = ismember(SBvariables(objModel),'totalPKB'); 
PKB_fold_chow = vitro_chow.variablevalues(end,id_PKB_phos473)/vitro_chow.variablevalues(end,id_PKB);
PKB_fold_hfd = vitro_hfd14d.variablevalues(end,id_PKB_phos473)/vitro_hfd14d.variablevalues(end,id_PKB);
fold_diet_PKB= PKB_fold_hfd/PKB_fold_chow;% fold between diets ( fold on fold )

id_AS = ismember(SBvariables(objModel),'totalAS160'); 
id_As_phos  = ismember(SBstates(objModel),'AS160p'); 
AS160_fold_chow = vitro_chow.statevalues(end,id_As_phos)/vitro_chow.variablevalues(end,id_AS);
AS160_fold_hfd = vitro_hfd14d.statevalues(end,id_As_phos)/vitro_hfd14d.variablevalues(end,id_AS);
fold_diet_AS160= AS160_fold_hfd/AS160_fold_chow;% fold between diets ( fold on fold )

cost_PKB=   ( ( D.Phos.MeanPKB(:,2) -  fold_diet_PKB ).^2 )./( D.Phos.SEMPKB(:,2) .^2 );    
cost_AS160=   ( ( D.Phos.MeanAS160(:,2) -  fold_diet_AS160 ).^2 )./( D.Phos.SEMAS160(:,2) .^2 );    
cost_in_vitro= cost_PKB + cost_AS160;

% limit = 20 + chi2inv(0.95,1); 
limit = chi2inv(0.95, (numel(D.BW.mean)-4 + numel(D.IPGTTgcc.Mean) + 4 + 2 + 6));
%% Adhoc 
%% Adhoc 
% U_A
IPGTT_hfd14d_UA= trapz(IPGTT_hfd14d.variablevalues(:, ismember(SBvariables(objModel),'U_id_A')) ); 
IPGTT_hfd14d_UM= trapz(IPGTT_hfd14d.variablevalues(:, ismember(SBvariables(objModel),'U_id_M')) ); 
adhoc_cost = adhoc_cost + Constraint(IPGTT_hfd14d_UA, IPGTT_hfd14d_UM*0.501, limit);
adhoc_cost = adhoc_cost + Constraint(IPGTT_hfd14d_UM*0.149, IPGTT_hfd14d_UA, limit);

IPGTT_chow_UA= trapz(IPGTT_chow.variablevalues(:, ismember(SBvariables(objModel),'U_id_A')) ); 
IPGTT_chow_UM= trapz(IPGTT_chow.variablevalues(:, ismember(SBvariables(objModel),'U_id_M')) ); 
adhoc_cost = adhoc_cost + Constraint(IPGTT_chow_UA, IPGTT_chow_UM*0.501, limit);
adhoc_cost = adhoc_cost + Constraint(IPGTT_chow_UM*0.15, IPGTT_chow_UA, limit);

hfd_6w_IPGTT_UA= trapz(hfd_6w_IPGTT.variablevalues(:, ismember(SBvariables(objModel),'U_id_A')) ); 
hfd_6w_IPGTT_UM= trapz(hfd_6w_IPGTT.variablevalues(:, ismember(SBvariables(objModel),'U_id_M')) ); 
adhoc_cost = adhoc_cost + Constraint(hfd_6w_IPGTT_UA, hfd_6w_IPGTT_UM*0.501, limit);
adhoc_cost = adhoc_cost + Constraint(hfd_6w_IPGTT_UM*0.149, hfd_6w_IPGTT_UA, limit);

% U_I  
IPGTT_hfd14d_U_id= trapz(IPGTT_hfd14d.variablevalues(:, ismember(SBvariables(objModel),'U_id')) ); 
IPGTT_hfd14d_U_ii= trapz(IPGTT_hfd14d.variablevalues(:, ismember(SBvariables(objModel),'U_ii')) ); 
adhoc_cost = adhoc_cost + Constraint(IPGTT_hfd14d_U_ii*0.99, IPGTT_hfd14d_U_id, limit);
adhoc_cost = adhoc_cost + Constraint(IPGTT_hfd14d_U_id*0.0499, IPGTT_hfd14d_U_ii, limit);

IPGTT_chow_U_id= trapz(IPGTT_chow.variablevalues(:, ismember(SBvariables(objModel),'U_id')) ); 
IPGTT_chow_U_ii= trapz(IPGTT_chow.variablevalues(:, ismember(SBvariables(objModel),'U_ii')) ); 
adhoc_cost = adhoc_cost + Constraint(IPGTT_chow_U_ii*0.99, IPGTT_chow_U_id, limit);
adhoc_cost = adhoc_cost + Constraint(IPGTT_chow_U_id*0.0499, IPGTT_chow_U_ii, limit);

hfd_6w_IPGTT_U_id = trapz(hfd_6w_IPGTT.variablevalues(:, ismember(SBvariables(objModel),'U_id')) ); 
hfd_6w_IPGTT_U_ii= trapz(hfd_6w_IPGTT.variablevalues(:, ismember(SBvariables(objModel),'U_ii')) ); 
adhoc_cost = adhoc_cost + Constraint(hfd_6w_IPGTT_U_ii*0.99, hfd_6w_IPGTT_U_id, limit);
adhoc_cost = adhoc_cost + Constraint(hfd_6w_IPGTT_U_id*0.0499, hfd_6w_IPGTT_U_ii, limit);

%% Total cost
cost= cost_bw +  cost_IPGTT + cost_in_vitro + bw_cost_up  + ss_cost  + adhoc_cost ;


catch 
   cost =1e99;
end


end


function [penalty] = Constraint(lower, upper, offset)
viol =  lower - upper  ; 
if viol > 0
    penalty = viol + offset; 
else
    penalty = 0;
end
end
```
