## Supplementary scripts for "A multi-scale *in silico* mouse model for insulin resistance and humanoid type 2 diabetes": Setup_data.m

```
function[D, Dpred] = Setup_data()
load('Stenkula_IPGTT_Mean.mat');
load('Stenkula_IPGTT_SEM.mat');
load('Stenkula_IPGTT_icc_SEM.mat');
load('Stenkula_IPGTT_icc_Mean.mat');
load('MeanSEM_SR2019_PKB_IRS1_AS160.mat');
load('Hanson_2018_BW.mat') 
load('Chow_BW_8W.mat')
load('HFD_BW_8W.mat')
load('Stenkula_IPGTT_6w.mat')

Stenkula_IPGTT_SEM( Stenkula_IPGTT_SEM(:,1) < mean(Stenkula_IPGTT_SEM(:,1)) , 1 ) = mean(Stenkula_IPGTT_SEM(:,1)); %14d chow
Stenkula_IPGTT_SEM( Stenkula_IPGTT_SEM(:,2) < mean(Stenkula_IPGTT_SEM(:,2)) , 2 ) = mean(Stenkula_IPGTT_SEM(:,2)); %2d HFD
Stenkula_IPGTT_SEM( Stenkula_IPGTT_SEM(:,3) < mean(Stenkula_IPGTT_SEM(:,3)) , 3 ) = mean(Stenkula_IPGTT_SEM(:,3));%6d HFD
Stenkula_IPGTT_SEM( Stenkula_IPGTT_SEM(:,4) < mean(Stenkula_IPGTT_SEM(:,4)) , 4 ) = mean(Stenkula_IPGTT_SEM(:,4)); %14d

Stenkula_IPGTT_6w.ChowSEM(1,  Stenkula_IPGTT_6w.ChowSEM(1,:) < mean(Stenkula_IPGTT_6w.ChowSEM(1,:)) )= mean(Stenkula_IPGTT_6w.ChowSEM(1,:)); % 6w Chow
Stenkula_IPGTT_6w.HFDSEM(1,  Stenkula_IPGTT_6w.HFDSEM(1,:) < mean(Stenkula_IPGTT_6w.HFDSEM(1,:)) ) = mean(Stenkula_IPGTT_6w.HFDSEM(1,:)); % 6w HFD

HFD_BW_8W.SEM( HFD_BW_8W.SEM < mean(HFD_BW_8W.SEM)  ) = mean(HFD_BW_8W.SEM) ;
Chow_BW_8W.SEM( Chow_BW_8W.SEM < mean(Chow_BW_8W.SEM)  ) = mean(Chow_BW_8W.SEM) ;

Dpred.IPGTTgcc.HFDMean = Stenkula_IPGTT_6w.HFDMean ;
Dpred.IPGTTgcc.HFDSEM = Stenkula_IPGTT_6w.HFDSEM ;
Dpred.IPGTTgcc.ChowMean = Stenkula_IPGTT_6w.ChowMean ; 
Dpred.IPGTTgcc.ChowSEM = Stenkula_IPGTT_6w.ChowSEM ;
Dpred.BW_chow = Chow_BW_8W;
Dpred.BW_HFD = HFD_BW_8W;

D.IPGTTgcc.Mean=Stenkula_IPGTT_Mean;
D.IPGTTgcc.SEM=Stenkula_IPGTT_SEM;
D.IPGTTicc.Mean=Stenkula_IPGTT_icc_Mean;
D.IPGTTicc.SEM=Stenkula_IPGTT_icc_SEM;
D.BW_unpublished_chow = Chow_BW_8W;
D.BW_unpublished_hfd = HFD_BW_8W;
D.BW = Hanson_2018_BW;
D.IPGTTicc.SEM(2:end,:)=inf;
D.Phos.MeanPKB= MeanSEM_SR2019_PKB_IRS1_AS160.Mean_pPKB;
D.Phos.SEMPKB= MeanSEM_SR2019_PKB_IRS1_AS160.SEM_pPKB;
D.Phos.MeanAS160= MeanSEM_SR2019_PKB_IRS1_AS160.Mean_pAS160;
D.Phos.SEMAS160= MeanSEM_SR2019_PKB_IRS1_AS160.SEM_pAS160;
clear("Stenkula*")
```
