## Supplementary scripts for "A multi-scale *in silico* mouse model for insulin resistance and humanoid type 2 diabetes": Figure_3_and_4.m

```
%% Setup
p=utility.p0;
id_FFM = ismember(SBstates(objModel),'FFM') ; id_FM = ismember(SBstates(objModel),'FM');
id_FFMname = ismember(pNames,'FFMinit') ; id_FMname = ismember(pNames,'FMinit');
t_diet_up=0:1:56;
t_fasting_up=[ 42  42+0.5833 ]  ;  %t_diet:0.0001:t_diet+0.5833;
t_IPGTT_up=(0:1:120)/(60*24)+t_fasting_up(end);

t_diet=14;
t_fasting=[ t_diet  t_diet+0.5833 ]  ;  %t_diet:0.0001:t_diet+0.5833;
t_IPGTT=(0:1:120)/(60*24)+t_fasting(end);

id_BW=ismember(SBvariables(objModel),'BW');
id_Glu = ismember(SBvariables(objModel),'Gconc');
id_Ins = ismember(SBvariables(objModel),'Iconc');

%% UNC
for i = 1:nfiles
load( strcat( filedir ,'/',files(i).name ) )
disp(strcat('Processing - ',files(i).name, int2str(i),'/',int2str(nfiles)))
theta=Results.xbest;


[ AllSimulations ] = Function_AllSimulations( theta ,func_mex_model, D, utility);

chow8w             = AllSimulations.chow8w;
hfd8w              = AllSimulations.hfd8w;
IPGTT_hfd6w        = AllSimulations.IPGTT_hfd6w;
chow2w              = AllSimulations.chow2w;
IPGTT_chow2w        = AllSimulations.IPGTT_chow2w;
chow12dHFD2d        = AllSimulations.chow12dHFD2d;
IPGTT_chow12dHFD2d  = AllSimulations.IPGTT_chow12dHFD2d;
chow10dHFD4d        = AllSimulations.chow10dHFD4d;
chow8dHFD6d         = AllSimulations.chow8dHFD6d;
IPGTT_chow8dHFD6d   = AllSimulations.IPGTT_chow8dHFD6d;
hfd2w               = AllSimulations.hfd2w;
IPGTT_hfd2w         = AllSimulations.IPGTT_hfd2w;

%% MIN and MAX value calculations

%% Predictions
if i == 1 % Setting up vectors
Max_chow8w_BW =  chow8w.variablevalues(:,id_BW);  Min_chow8w_BW =  chow8w.variablevalues(:,id_BW);     
Max_hfd8w_BW =  hfd8w.variablevalues(:,id_BW);  Min_hfd8w_BW =  hfd8w.variablevalues(:,id_BW);      
Max_hfd6w_Glu =  IPGTT_hfd6w.variablevalues(:,id_Glu);  Min_hfd6w_Glu =  IPGTT_hfd6w.variablevalues(:,id_Glu); 
end

Max_chow8w_BW( Max_chow8w_BW < chow8w.variablevalues(:,id_BW) ) =  chow8w.variablevalues( Max_chow8w_BW < chow8w.variablevalues(:,id_BW),id_BW);
Min_chow8w_BW( Min_chow8w_BW > chow8w.variablevalues(:,id_BW) ) =  chow8w.variablevalues( Min_chow8w_BW > chow8w.variablevalues(:,id_BW),id_BW);
Max_hfd8w_BW( Max_hfd8w_BW < hfd8w.variablevalues(:,id_BW) ) =  hfd8w.variablevalues( Max_hfd8w_BW < hfd8w.variablevalues(:,id_BW),id_BW);
Min_hfd8w_BW( Min_hfd8w_BW > hfd8w.variablevalues(:,id_BW) ) =  hfd8w.variablevalues( Min_hfd8w_BW > hfd8w.variablevalues(:,id_BW),id_BW);

Max_hfd6w_Glu( Max_hfd6w_Glu < IPGTT_hfd6w.variablevalues(:,id_Glu) ) =  IPGTT_hfd6w.variablevalues( Max_hfd6w_Glu < IPGTT_hfd6w.variablevalues(:,id_Glu),id_Glu);
Min_hfd6w_Glu( Min_hfd6w_Glu > IPGTT_hfd6w.variablevalues(:,id_Glu) ) =  IPGTT_hfd6w.variablevalues( Min_hfd6w_Glu > IPGTT_hfd6w.variablevalues(:,id_Glu),id_Glu);
% 
BW_id = [ 22 29 36 42 51 57];
bw_cost1_up_pred= nansum (  ( (chow8w.variablevalues(BW_id,1)'   -  D.BW_unpublished_chow.Mean(4:end) ).^2 )./( D.BW_unpublished_chow.SEM(4:end).^2 )); % chow
bw_cost2_up_pred= nansum (  ( (hfd8w.variablevalues(BW_id,1)'   -  D.BW_unpublished_hfd.Mean(4:end) ).^2 )./( D.BW_unpublished_hfd.SEM(4:end).^2 )); % chow
bw_cost_up_pred = bw_cost1_up_pred + bw_cost2_up_pred;

id_gcc = [ 1 16 31 61 121 ];
yexp = Dpred.IPGTTgcc.HFDMean ;
SEM= Dpred.IPGTTgcc.HFDSEM; 
ysim= IPGTT_hfd6w.variablevalues(id_gcc, end-1);           
IPGTT_glucose_cost_6w =  nansum (  ( (yexp  -  ysim' ).^2 )./( SEM.^2 )) ;

bw_cost1_up= nansum (  ( (chow8w.variablevalues([1 8 15],1)'   -  D.BW_unpublished_chow.Mean(1:3) ).^2 )./( D.BW_unpublished_chow.SEM(1:3).^2 )); % chow
bw_cost2_up= nansum (  ( (hfd8w.variablevalues([1 8 15],1)'   -  D.BW_unpublished_hfd.Mean(1:3) ).^2 )./( D.BW_unpublished_hfd.SEM(1:3).^2 )); % chow
bw_cost_up = bw_cost1_up + bw_cost2_up;

%% Hansson et al (2018)
if i == 1  % Setting up vectors
% 2w chow    
Max_chow2w_BW =  chow2w.variablevalues(:,id_BW);  Min_chow2w_BW =  chow2w.variablevalues(:,id_BW);     
Max_chow2w_Glu =  IPGTT_chow2w.variablevalues(:,id_Glu);  Min_chow2w_Glu =  IPGTT_chow2w.variablevalues(:,id_Glu); 
Max_chow2w_Ins = IPGTT_chow2w.variablevalues(1,id_Ins);  Min_chow2w_Ins =  IPGTT_chow2w.variablevalues(1,id_Ins); 
%10d chow + 4d HFD  
Max_chow10dHFD4d_BW =  chow10dHFD4d.variablevalues(:,id_BW);  Min_chow10dHFD4d_BW =  chow10dHFD4d.variablevalues(:,id_BW);     
%12d chow + 2d HFD  
Max_chow12dHFD2d_BW =  chow12dHFD2d.variablevalues(:,id_BW);  Min_chow12dHFD2d_BW =  chow12dHFD2d.variablevalues(:,id_BW);     
Max_chow12dHFD2d_Glu =  IPGTT_chow12dHFD2d.variablevalues(:,id_Glu);  Min_chow12dHFD2d_Glu =  IPGTT_chow12dHFD2d.variablevalues(:,id_Glu); 
Max_chow12dHFD2d_Ins = IPGTT_chow12dHFD2d.variablevalues(1,id_Ins);   Min_chow12dHFD2d_Ins =  IPGTT_chow12dHFD2d.variablevalues(1,id_Ins); 
%8d chow + 6d HFD  
Max_chow8dHFD6d_BW =  chow8dHFD6d.variablevalues(:,id_BW);  Min_chow8dHFD6d_BW =  chow8dHFD6d.variablevalues(:,id_BW);     
Max_chow8dHFD6d_Glu =  IPGTT_chow8dHFD6d.variablevalues(:,id_Glu);  Min_chow8dHFD6d_Glu =  IPGTT_chow8dHFD6d.variablevalues(:,id_Glu); 
Max_chow8dHFD6d_Ins = IPGTT_chow8dHFD6d.variablevalues(1,id_Ins);   Min_chow8dHFD6d_Ins =  IPGTT_chow8dHFD6d.variablevalues(1,id_Ins); 
% 2w hfd
Max_hfd2w_BW =  hfd2w.variablevalues(:,id_BW);  Min_hfd2w_BW =  hfd2w.variablevalues(:,id_BW);     
Max_hfd2w_Glu =  IPGTT_hfd2w.variablevalues(:,id_Glu);  Min_hfd2w_Glu =  IPGTT_hfd2w.variablevalues(:,id_Glu); 
Max_hfd2w_Ins = IPGTT_hfd2w.variablevalues(1,id_Ins);  Min_hfd2w_Ins =  IPGTT_hfd2w.variablevalues(1,id_Ins); 
end 

Max_chow2w_BW( Max_chow2w_BW < chow2w.variablevalues(:,id_BW) ) =  chow2w.variablevalues( Max_chow2w_BW < chow2w.variablevalues(:,id_BW),id_BW);
Min_chow2w_BW( Min_chow2w_BW > chow2w.variablevalues(:,id_BW) ) =  chow2w.variablevalues( Min_chow2w_BW > chow2w.variablevalues(:,id_BW),id_BW);
Max_chow2w_Glu( Max_chow2w_Glu < IPGTT_chow2w.variablevalues(:,id_Glu) ) =  IPGTT_chow2w.variablevalues( Max_chow2w_Glu < IPGTT_chow2w.variablevalues(:,id_Glu),id_Glu);
Min_chow2w_Glu( Min_chow2w_Glu > IPGTT_chow2w.variablevalues(:,id_Glu) ) =  IPGTT_chow2w.variablevalues( Min_chow2w_Glu > IPGTT_chow2w.variablevalues(:,id_Glu),id_Glu);
Max_chow2w_Ins( Max_chow2w_Ins < IPGTT_chow2w.variablevalues(1,id_Ins) ) =  IPGTT_chow2w.variablevalues( Max_chow2w_Ins < IPGTT_chow2w.variablevalues(1,id_Ins),id_Ins);
Min_chow2w_Ins( Min_chow2w_Ins > IPGTT_chow2w.variablevalues(1,id_Ins) ) =  IPGTT_chow2w.variablevalues( Min_chow2w_Ins > IPGTT_chow2w.variablevalues(1,id_Ins),id_Ins);

Max_chow12dHFD2d_BW( Max_chow12dHFD2d_BW < chow12dHFD2d.variablevalues(:,id_BW) ) =  chow12dHFD2d.variablevalues( Max_chow12dHFD2d_BW < chow12dHFD2d.variablevalues(:,id_BW),id_BW);
Min_chow12dHFD2d_BW( Min_chow12dHFD2d_BW > chow12dHFD2d.variablevalues(:,id_BW) ) =  chow12dHFD2d.variablevalues( Min_chow12dHFD2d_BW > chow12dHFD2d.variablevalues(:,id_BW),id_BW);
Max_chow12dHFD2d_Glu( Max_chow12dHFD2d_Glu < IPGTT_chow12dHFD2d.variablevalues(:,id_Glu) ) =  IPGTT_chow12dHFD2d.variablevalues( Max_chow12dHFD2d_Glu < IPGTT_chow12dHFD2d.variablevalues(:,id_Glu),id_Glu);
Min_chow12dHFD2d_Glu( Min_chow12dHFD2d_Glu > IPGTT_chow12dHFD2d.variablevalues(:,id_Glu) ) =  IPGTT_chow12dHFD2d.variablevalues( Min_chow12dHFD2d_Glu > IPGTT_chow12dHFD2d.variablevalues(:,id_Glu),id_Glu);
Max_chow12dHFD2d_Ins( Max_chow12dHFD2d_Ins < IPGTT_chow12dHFD2d.variablevalues(1,id_Ins) ) =  IPGTT_chow12dHFD2d.variablevalues( Max_chow12dHFD2d_Ins < IPGTT_chow12dHFD2d.variablevalues(1,id_Ins),id_Ins);
Min_chow12dHFD2d_Ins( Min_chow12dHFD2d_Ins > IPGTT_chow12dHFD2d.variablevalues(1,id_Ins) ) =  IPGTT_chow12dHFD2d.variablevalues( Min_chow12dHFD2d_Ins > IPGTT_chow12dHFD2d.variablevalues(1,id_Ins),id_Ins);

Max_chow10dHFD4d_BW( Max_chow10dHFD4d_BW < chow10dHFD4d.variablevalues(:,id_BW) ) =  chow10dHFD4d.variablevalues( Max_chow10dHFD4d_BW < chow10dHFD4d.variablevalues(:,id_BW),id_BW);
Min_chow10dHFD4d_BW( Min_chow10dHFD4d_BW > chow10dHFD4d.variablevalues(:,id_BW) ) =  chow10dHFD4d.variablevalues( Min_chow10dHFD4d_BW > chow10dHFD4d.variablevalues(:,id_BW),id_BW);

Max_chow8dHFD6d_BW( Max_chow8dHFD6d_BW < chow8dHFD6d.variablevalues(:,id_BW) ) =  chow8dHFD6d.variablevalues( Max_chow8dHFD6d_BW < chow8dHFD6d.variablevalues(:,id_BW),id_BW);
Min_chow8dHFD6d_BW( Min_chow8dHFD6d_BW > chow8dHFD6d.variablevalues(:,id_BW) ) =  chow8dHFD6d.variablevalues( Min_chow8dHFD6d_BW > chow8dHFD6d.variablevalues(:,id_BW),id_BW);
Max_chow8dHFD6d_Glu( Max_chow8dHFD6d_Glu < IPGTT_chow8dHFD6d.variablevalues(:,id_Glu) ) =  IPGTT_chow8dHFD6d.variablevalues( Max_chow8dHFD6d_Glu < IPGTT_chow8dHFD6d.variablevalues(:,id_Glu),id_Glu);
Min_chow8dHFD6d_Glu( Min_chow8dHFD6d_Glu > IPGTT_chow8dHFD6d.variablevalues(:,id_Glu) ) =  IPGTT_chow8dHFD6d.variablevalues( Min_chow8dHFD6d_Glu > IPGTT_chow8dHFD6d.variablevalues(:,id_Glu),id_Glu);
Max_chow8dHFD6d_Ins( Max_chow8dHFD6d_Ins < IPGTT_chow8dHFD6d.variablevalues(1,id_Ins) ) =  IPGTT_chow8dHFD6d.variablevalues( Max_chow8dHFD6d_Ins < IPGTT_chow8dHFD6d.variablevalues(1,id_Ins),id_Ins);
Min_chow8dHFD6d_Ins( Min_chow8dHFD6d_Ins > IPGTT_chow8dHFD6d.variablevalues(1,id_Ins) ) =  IPGTT_chow8dHFD6d.variablevalues( Min_chow8dHFD6d_Ins > IPGTT_chow8dHFD6d.variablevalues(1,id_Ins),id_Ins);

Max_hfd2w_BW( Max_hfd2w_BW < hfd2w.variablevalues(:,id_BW) ) =  hfd2w.variablevalues( Max_hfd2w_BW < hfd2w.variablevalues(:,id_BW),id_BW);
Min_hfd2w_BW( Min_hfd2w_BW > hfd2w.variablevalues(:,id_BW) ) =  hfd2w.variablevalues( Min_hfd2w_BW > hfd2w.variablevalues(:,id_BW),id_BW);
Max_hfd2w_Glu( Max_hfd2w_Glu < IPGTT_hfd2w.variablevalues(:,id_Glu) ) =  IPGTT_hfd2w.variablevalues( Max_hfd2w_Glu < IPGTT_hfd2w.variablevalues(:,id_Glu),id_Glu);
Min_hfd2w_Glu( Min_hfd2w_Glu > IPGTT_hfd2w.variablevalues(:,id_Glu) ) =  IPGTT_hfd2w.variablevalues( Min_hfd2w_Glu > IPGTT_hfd2w.variablevalues(:,id_Glu),id_Glu);
Max_hfd2w_Ins( Max_hfd2w_Ins < IPGTT_hfd2w.variablevalues(1,id_Ins) ) =  IPGTT_hfd2w.variablevalues( Max_hfd2w_Ins < IPGTT_hfd2w.variablevalues(1,id_Ins),id_Ins);
Min_hfd2w_Ins( Min_hfd2w_Ins > IPGTT_hfd2w.variablevalues(1,id_Ins) ) =  IPGTT_hfd2w.variablevalues( Min_hfd2w_Ins > IPGTT_hfd2w.variablevalues(1,id_Ins),id_Ins);

if i == 1
Max_P = p; Min_P =p;
end
Max_P( Max_P < p ) =  p( Max_P < p);
Min_P( Min_P > p ) =  p( Min_P > p);


%% Cost calculations

if i == 1 
    x_pred = bw_cost_up_pred + IPGTT_glucose_cost_6w;
    x_training = cost_function( theta ,func_mex_model, D, utility);
    best_param_train = theta;
    best_param_pred = theta;
end

cost_training = cost_function( theta ,func_mex_model, D, utility);
cost_pred = bw_cost_up_pred + IPGTT_glucose_cost_6w;

if x_training > cost_training
x_training = cost_training;
best_param_train = theta;
end

if x_pred > cost_pred
x_pred = cost_pred;
best_bw_cost_up_pred=bw_cost_up_pred;
best_IPGTT_glucose_cost_6w = IPGTT_glucose_cost_6w;
best_param_pred = theta;
end
end

%% Plotting

predColor =  [0.9,.9,0.9]; 
TrainColor = [ 0.5,0.5,0.5];
label_size = 25;
label_size_ax = 18;
marker_size = 5;
fig=0;

%% Simulations for best cost.
theta1 = best_param_pred;
[ AllSimulations_pred ] = Function_AllSimulations( theta1 ,func_mex_model, D, utility);

Estimation_folder ='Results/MLMM_final/Estimation';
S = dir(Estimation_folder);
load(strcat(Estimation_folder,'/',S(3).name));
theta2 = Results.xbest;
[ AllSimulations_train ] = Function_AllSimulations( theta2 ,func_mex_model, D, utility);

x_training_from_estimation = cost_function( theta2 ,func_mex_model, D, utility);
if x_training < x_training_from_estimation
    best_cost_training_data = x_training;
else
best_cost_training_data   = x_training_from_estimation;
disp(strcat('Best cost for training data = ', num2str(best_cost_training_data)));
end
best_cost_pred_data = x_pred ;
disp(strcat('Best cost for validation data = ', num2str(best_cost_pred_data)));


%% Figure 1 - Body weight - TRAINING AND PREDICTIONS
t_diet=0:1:14;
fill_time_2w = [ t_diet fliplr(t_diet) ];

fig=fig+1;
figure (fig)
set(figure(fig), 'outerposition',[0 0 2560 1440], 'PaperType','a4')

%%% Figure 1a  BW - Hansson 2018 et al
Experiments_to_plot = {'chow2w','chow10dHFD4d','chow8dHFD6d','hfd2w'};

for kk = 1:length(Experiments_to_plot)
subplot(2,4,kk) %% a)
values=[];
data.time=D.BW.time(kk,:);
data.mean = D.BW.mean(kk,:);
data.SEM = D.BW.SEM(kk,:);
eval(strcat('values(:,1)=Max_',Experiments_to_plot{kk},'_BW;')); 
eval(strcat('values(:,2)=Min_',Experiments_to_plot{kk},'_BW;')); 
if kk == 2
    time_plot = 10:1:t_diet(end);
    time_fill = [ time_plot fliplr(time_plot) ];
    
elseif kk == 3
    time_plot =8:1:t_diet(end);
    time_fill = [ time_plot fliplr(time_plot) ];
else
    time_plot = t_diet;
    time_fill = fill_time_2w;
end
f=fill(time_fill, [ values(:,2)' fliplr( values(:,1)' ) ], TrainColor, 'EdgeColor', 'none' );  
set(f,'facealpha',.5)
hold on
eval(strcat('plot( time_plot, AllSimulations_train.',Experiments_to_plot{kk}, ".variablevalues(:,id_BW),'color', TrainColor - 0.2 , 'linewidth', 2);"))

errorbar(data.time,data.mean, data.SEM,'k*' ,'linewidth',2) % 2wHFD
set(gca,'FontSize',label_size_ax, 'FontWeight','bold')
xlabel('Time (days)','FontSize', label_size )
ylabel('Body weight (g)','FontSize', label_size )
box off
xlim([0 14]); 
xticks([0 ,14])
xticklabels({'0','14'})
ylim([22 34]);
end
%%
Experiments_to_plot = {'chow8w','hfd8w'};
for kk = 1:length(Experiments_to_plot)
subplot(2,4,4+kk) %% a)
values=[];
if kk == 1
    data.time=Dpred.BW_chow.time; data.mean=Dpred.BW_chow.Mean; data.SEM=Dpred.BW_chow.SEM ;%Control    
elseif kk == 2
   data.time=Dpred.BW_HFD.time; data.mean=Dpred.BW_HFD.Mean; data.SEM=Dpred.BW_HFD.SEM;%Control
end
time_fill = fill_time_2w;
time_plot = 0:1:56; % plot both param set for full duration

eval(strcat('values(:,1)=Max_',Experiments_to_plot{kk},'_BW;')); 
eval(strcat('values(:,2)=Min_',Experiments_to_plot{kk},'_BW;')); 
f=fill(time_fill, [ values(1:15,2)' fliplr( values(1:15,1)' ) ], TrainColor, 'EdgeColor', 'none' );  
set(f,'facealpha',.5)
hold on
errorbar(data.time(1:3),data.mean(1:3), data.SEM(1:3),'k*' ,'linewidth',1.5) % 2wHFD

fill_time_8w = [ 14:1:56 fliplr(14:1:56) ];

f=fill(fill_time_8w, [ values(15:end,2)' fliplr( values(15:end,1)' ) ], predColor, 'EdgeColor', 'none' );  
set(f,'facealpha',.5)
hold on
errorbar(data.time(4:end),data.mean(4:end), data.SEM(4:end),'bO' ,'linewidth',1.5) % 2wHFD
% ploting best sim for training and pred
eval(strcat('plot( time_plot, AllSimulations_train.',Experiments_to_plot{kk}, ".variablevalues(:,id_BW),'color', TrainColor - 0.2 , 'linewidth', 2);"))
eval(strcat('plot( time_plot, AllSimulations_pred.',Experiments_to_plot{kk}, ".variablevalues(:,id_BW),'b--', 'linewidth', 2);"))


set(gca,'FontSize',label_size_ax, 'FontWeight','bold')
xlabel('Time (days)','FontSize', label_size )
ylabel('Body weight (g)','FontSize', label_size )
box off
xlim([0 56]); 
xticks([0 ,56])
xticklabels({'0','56'})
ylim([22 50]);

end


%% IPGTT
%% figure 2 - IPGTT - TRAINING AND PREDIKTION
fill_time_IPGTT = [ t_IPGTT-t_fasting(end) fliplr(t_IPGTT-t_fasting(end)) ]*24*60;
tp_IPGTT=[0 15 30 60 120];
time_best_cost = (t_IPGTT-t_fasting(end) )*24*60; 
fig=fig+1;
figure(fig)
set(figure(fig), 'outerposition',[0 0 2560 1440], 'PaperType','a4')


Experiments_to_plot = {'chow2w','chow12dHFD2d','chow8dHFD6d','hfd2w'};
IPGTT_sim_name = {'IPGTT_chow2w','IPGTT_chow12dHFD2d','IPGTT_chow8dHFD6d','IPGTT_hfd2w'};
for kk = 1:length(Experiments_to_plot)
subplot(2,4,kk) %% a)
values=[];
data.time=tp_IPGTT;
data.mean = D.IPGTTgcc.Mean(:,kk);
data.SEM = D.IPGTTgcc.SEM(:,kk);
eval(strcat('values(:,1)=Max_',Experiments_to_plot{kk},'_Glu;')); 
eval(strcat('values(:,2)=Min_',Experiments_to_plot{kk},'_Glu;')); 

time_fill = fill_time_IPGTT;
time_plot = (t_IPGTT-t_fasting(end))*24*60;

f=fill(time_fill, [ values(:,2)' fliplr( values(:,1)' ) ], TrainColor, 'EdgeColor', 'none' );  
set(f,'facealpha',.5)
hold on

errorbar(data.time,data.mean, data.SEM,'k*' ,'linewidth',1.5) % 2wHFD

eval(strcat('plot( time_plot, AllSimulations_train.',IPGTT_sim_name{kk}, ".variablevalues(:,id_Glu),'color', TrainColor - 0.2 , 'linewidth', 2);"))

box off
set(gca,'FontSize',label_size_ax, 'FontWeight','bold')
ylabel(' Plasma glucose (mM)','FontSize', label_size ,'FontWeight','bold')
xlabel('Time (min)','FontSize', label_size )
xlim([0 120]);
ylim([4 32]);

end
%%
subplot(2,4,5)
values=[];

ins_value_chow2w = AllSimulations_train.IPGTT_chow2w.variablevalues(1,id_Ins)  ;
ins_value_chow12dHFD2d = AllSimulations_train.IPGTT_chow12dHFD2d.variablevalues(1,id_Ins)  ;
ins_value_chow8dHFD6d = AllSimulations_train.IPGTT_chow8dHFD6d.variablevalues(1,id_Ins) ;
ins_value_hfd2w = AllSimulations_train.IPGTT_hfd2w.variablevalues(1,id_Ins)  ;

y = [ D.IPGTTicc.Mean(1,1) ins_value_chow2w       ;...
      D.IPGTTicc.Mean(1,2) ins_value_chow12dHFD2d ;...
      D.IPGTTicc.Mean(1,3) ins_value_chow8dHFD6d  ;...
      D.IPGTTicc.Mean(1,4) ins_value_hfd2w        ];
b=bar(y,'facecolor','flat','HandleVisibility','off');
b(1).CData(1,:) = [ 1 1 1];
b(1).CData(2,:) = [ 1 1 1];
b(1).CData(3,:) = [1 1 1];
b(1).CData(4,:) = [ 1 1 1];
b(2).CData(1,:) = [ 0.7 0.7 0.7];
b(2).CData(2,:) = [ 0.7 0.7 0.7];
b(2).CData(3,:) = [0.7 0.7 0.7];
b(2).CData(4,:) = [ 0.7 0.7 0.7];
hold on

errhigh = [Max_chow2w_Ins        - ins_value_chow2w,...  
           Max_chow12dHFD2d_Ins  - ins_value_chow12dHFD2d,...  
           Max_chow8dHFD6d_Ins   - ins_value_chow8dHFD6d,...  
           Max_hfd2w_Ins         - ins_value_hfd2w];

errlow =  [Min_chow2w_Ins        - ins_value_chow2w,...  
           Min_chow12dHFD2d_Ins  - ins_value_chow12dHFD2d,...  
           Min_chow8dHFD6d_Ins   - ins_value_chow8dHFD6d,...  
           Min_hfd2w_Ins         - ins_value_hfd2w];       
   

errorbar( 1.15:1:4.15, [ ins_value_chow2w  ins_value_chow12dHFD2d  ins_value_chow8dHFD6d ins_value_hfd2w],...
        errhigh,errlow,'O','color',[ 0.4 0.4 0.4],'linewidth',1.5)
errorbar( 0.85:1:3.85, [ D.IPGTTicc.Mean(1,1) D.IPGTTicc.Mean(1,2) D.IPGTTicc.Mean(1,3) D.IPGTTicc.Mean(1,4) ], [ D.IPGTTicc.SEM(1,1) D.IPGTTicc.SEM(1,2) D.IPGTTicc.SEM(1,3) D.IPGTTicc.SEM(1,4)],...
        'k*', 'linewidth',1.5) 
set(gca,'FontSize',label_size_ax, 'FontWeight','bold')
box off
ylabel('Plasma insulin (pM)','FontSize', label_size ,'FontWeight','bold')
xticks([1 2 3 4])
xticklabels({'2w chow','2d HFD',' 6d HFD',' 2w HFD'})
xtickangle(90)


%% 6w IPGTT prediction
values=[];
subplot(2,4,6)

hold on
data.time=tp_IPGTT; 

data.mean=Dpred.IPGTTgcc.HFDMean; data.SEM=Dpred.IPGTTgcc.HFDSEM;%Control
values(:,1)=Max_hfd6w_Glu; values(:,2)=Min_hfd6w_Glu; 

f=fill(fill_time_IPGTT, [ values(:,2)' fliplr( values(:,1)' ) ], predColor ,'EdgeColor', 'none' );
set(f,'facealpha',.5)
hold on

plot( time_plot, AllSimulations_train.IPGTT_hfd6w.variablevalues(:,id_Glu),'color', TrainColor - 0.2 , 'linewidth', 2);
plot( time_plot, AllSimulations_pred.IPGTT_hfd6w.variablevalues(:,id_Glu),'b--' , 'linewidth', 2);


errorbar(data.time,data.mean, data.SEM,'bO' ,'linewidth',1.5) % 2wHFD
set(gca,'FontSize',label_size_ax, 'FontWeight','bold')
ylabel(' Plasma glucose (mM)','FontSize', label_size ,'FontWeight','bold')
xlabel('Time (min)','FontSize', label_size )
xlim([0 120]);
ylim([5 45]);
```
