## Supplementary scripts for "A multi-scale *in silico* mouse model for insulin resistance and humanoid type 2 diabetes": Figure_5A.m

```
id_IRS = ismember(SBvariables(objModel),'totalIRS1'); 
id_PKB_phos473 = ismember(SBvariables(objModel),'totalPKB473'); 
id_PKB = ismember(SBvariables(objModel),'totalPKB'); 
id_AS = ismember(SBvariables(objModel),'totalAS160'); 
id_As_phos  = ismember(SBstates(objModel),'AS160p'); 
id_GLUT4m = ismember(SBvariables(objModel),'GLUT4m'); 
id_FFM = ismember(SBstates(objModel),'FFM') ; id_FM = ismember(SBstates(objModel),'FM');
id_FFMname = ismember(pNames,'FFMinit') ; id_FMname = ismember(pNames,'FMinit');

for i = 1:nfiles
load( strcat( filedir ,'/',files(i).name ) )
disp(strcat('Processing - ',files(i).name, int2str(i),'/',int2str(nfiles)))
theta=Results.xbest;

[ AllSimulations ] = Function_AllSimulations_4w_invitro( theta ,func_mex_model,utility);

chow4w       = AllSimulations.chow4w  ;
hfd4w        = AllSimulations.hfd4w   ;        
vitro_chow4w = AllSimulations.vitro_chow4w    ;   
vitro_hfd4w  = AllSimulations.vitro_hfd4w     ;


PKB_fold_chow = vitro_chow4w.variablevalues(end,id_PKB_phos473)/vitro_chow4w.variablevalues(end,id_PKB);
PKB_fold_hfd = vitro_hfd4w.variablevalues(end,id_PKB_phos473)/vitro_hfd4w.variablevalues(end,id_PKB);
fold_diet_PKB= PKB_fold_hfd/PKB_fold_chow;% fold between diets ( fold on fold )

AS160_fold_chow = vitro_chow4w.statevalues(end,id_As_phos)/vitro_chow4w.variablevalues(end,id_AS);
AS160_fold_hfd = vitro_hfd4w.statevalues(end,id_As_phos)/vitro_hfd4w.variablevalues(end,id_AS);
fold_diet_AS160= AS160_fold_hfd/AS160_fold_chow;% fold between diets ( fold on fold )


cost_PKB=   ( ( D.Phos.MeanPKB(:,2) -  fold_diet_PKB ).^2 )./( D.Phos.SEMPKB(:,2) .^2 );    
cost_AS160=   ( ( D.Phos.MeanAS160(:,2) -  fold_diet_AS160 ).^2 )./( D.Phos.SEMAS160(:,2) .^2 );    
cost_in_vitro= cost_PKB + cost_AS160;

if i == 1  % Setting up vectors    
Max_fold_diet_PKB = fold_diet_PKB;
Min_fold_diet_PKB= fold_diet_PKB;

Max_fold_diet_AS160= fold_diet_AS160;
Min_fold_diet_AS160= fold_diet_AS160;

x = cost_in_vitro;

end  

Max_fold_diet_PKB( Max_fold_diet_PKB < fold_diet_PKB) =  fold_diet_PKB( Max_fold_diet_PKB < fold_diet_PKB);
Min_fold_diet_PKB( Min_fold_diet_PKB > fold_diet_PKB) =  fold_diet_PKB( Min_fold_diet_PKB > fold_diet_PKB);

Max_fold_diet_AS160( Max_fold_diet_AS160 < fold_diet_AS160) =  fold_diet_AS160( Max_fold_diet_AS160 < fold_diet_AS160);
Min_fold_diet_AS160( Min_fold_diet_AS160 > fold_diet_AS160) =  fold_diet_AS160( Min_fold_diet_AS160 > fold_diet_AS160);

if cost_in_vitro < x
    x = cost_in_vitro    ;
end
end


Estimation_folder ='Results/MLMM_final/Estimation';
S = dir(Estimation_folder);
load(strcat(Estimation_folder,'/',S(3).name));
theta2 = Results.xbest;
[ AllSimulations2 ] = Function_AllSimulations_4w_invitro( theta2 ,func_mex_model,utility);

best_vitro_chow4w = AllSimulations2.vitro_chow4w    ;   
best_vitro_hfd4w  = AllSimulations2.vitro_hfd4w     ;

best_PKB_fold_chow = best_vitro_chow4w.variablevalues(end,id_PKB_phos473)/best_vitro_chow4w.variablevalues(end,id_PKB);
best_PKB_fold_hfd = best_vitro_hfd4w.variablevalues(end,id_PKB_phos473)/best_vitro_hfd4w.variablevalues(end,id_PKB);
best_fold_diet_PKB= best_PKB_fold_hfd/best_PKB_fold_chow;% fold between diets ( fold on fold )

best_AS160_fold_chow = best_vitro_chow4w.statevalues(end,id_As_phos)/best_vitro_chow4w.variablevalues(end,id_AS);
best_AS160_fold_hfd = best_vitro_hfd4w.statevalues(end,id_As_phos)/best_vitro_hfd4w.variablevalues(end,id_AS);
best_fold_diet_AS160= best_AS160_fold_hfd/best_AS160_fold_chow;% fold between diets ( fold on fold )


%% Plot
TrainColor = [ 0.5,0.5,0.5];
label_size = 25;
label_size_ax = 18;
marker_size = 5;
subplot(1,2,1)

y = [1 D.Phos.MeanPKB(:,2) 1 best_fold_diet_PKB ];
b = bar(y, 'grouped','FaceColor','flat' ,'HandleVisibility','off');
hold on
b(1).CData(1,:) = [1 1 1]; 
b(1).CData(2,:) = [1 1 1]; 
b(1).CData(3,:) = TrainColor; 
b(1).CData(4,:) = TrainColor; 
errhigh = [Max_fold_diet_PKB - best_fold_diet_PKB   ]  ;
errlow  = [Min_fold_diet_PKB - best_fold_diet_PKB   ]  ;
errorbar(4 , best_fold_diet_PKB ,errlow,errhigh,'O', 'color', [0.4 0.4 0.4], 'linewidth', 1.5 ,'markersize', 3 )
errorbar(2 ,D.Phos.MeanPKB(:,2) ,D.Phos.SEMPKB(:,2),'k*' ,'linewidth',1.5)

box off
ylim([0 1.5])
yticks([0 1])
xticks([1.5 3.5]);      
xticklabels({'Data','Model' })
a = get(gca,'XTickLabel');  
set(gca,'XTickLabel',a,'fontsize',label_size_ax,'FontWeight','bold')
ylabel({'PKB pS473/PKB total';'(fold of chow)'},'fontsize',label_size)

%%
subplot(1,2,2)
%figure(2) % PKB
y = [1 D.Phos.MeanAS160(:,2) 1 best_fold_diet_AS160 ];
b = bar(y, 'grouped','FaceColor','flat' ,'HandleVisibility','off');
hold on
b(1).CData(1,:) = [1 1 1]; 
b(1).CData(2,:) = [1 1 1]; 
b(1).CData(3,:) = TrainColor; 
b(1).CData(4,:) = TrainColor; 

errhigh = [Max_fold_diet_AS160 - best_fold_diet_AS160   ]  ;
errlow  = [Min_fold_diet_AS160 - best_fold_diet_AS160   ]  ;
errorbar(4 , best_fold_diet_AS160 ,errlow,errhigh,'O', 'color', [0.4 0.4 0.4], 'linewidth', 1.5 ,'markersize', 3 )
errorbar(2 ,D.Phos.MeanAS160(:,2) ,D.Phos.SEMAS160(:,2),'k*' ,'linewidth',1.5)

box off
ylim([0 1.5])
yticks([0 1])
xticks([1.5 3.5]);      
xticklabels({'Data','Model' })
a = get(gca,'XTickLabel');  
set(gca,'XTickLabel',a,'fontsize',label_size_ax,'FontWeight','bold')
ylabel({'AS160 pS473/AS160 total';'(fold of chow)'},'fontsize',label_size)
```
