## Supplementary scripts for "A multi-scale *in silico* mouse model for insulin resistance and humanoid type 2 diabetes": Figure_6_and_7AB.m

```
p=utility.p0; % for convenience 
id_IRS = ismember(SBvariables(objModel),'totalIRS1'); 
%id_PKB_phos308 = ismember(SBvariables(objModel),'totalPKB473'); 
id_PKB = ismember(SBvariables(objModel),'totalPKB'); 
id_AS = ismember(SBstates(objModel),'AS160'); 
id_As_phos  = ismember(SBstates(objModel),'AS160p'); 
id_RABGTP  =  ismember(SBstates(objModel),'RABGTP'); 
id_PMAa   =  ismember(SBstates(objModel),'PMAa'); 
id_GLUT4m = ismember(SBvariables(objModel),'GLUT4m'); 
id_CL_GIA = ismember(SBvariables(objModel),'CL_GIA'); 
id_PKBda   =  ismember(SBstates(objModel),'PKBda'); 
id_FFM = ismember(SBstates(objModel),'FFM') ; 
id_FM = ismember(SBstates(objModel),'FM');
id_FFMname = ismember(pNames,'FFMinit') ; id_FMname = ismember(pNames,'FMinit');
id_insulin_input = ismember(SBvariables(objModel),'insulin_input'); 
id_U_A = ismember(SBvariables(objModel),'U_id_A_plot'); 
id_U_M = ismember(SBvariables(objModel),'U_id_M'); 
id_U_H = ismember(SBvariables(objModel),'U_id_H'); 
id_U_id = ismember(SBvariables(objModel),'U_id'); 
id_U_ii = ismember(SBvariables(objModel),'U_ii'); 
id_ISEC = ismember(SBvariables(objModel),'ISEC'); 
id_Beta_failure = ismember(SBstates(objModel),'Beta_failure');
id_T2deffect = ismember(SBvariables(objModel),'T2deffect');
id_BW = ismember(SBvariables(objModel),'BW');
id_Iconc = ismember(SBvariables(objModel),'Iconc'); 
id_Gconc = ismember(SBvariables(objModel),'Gconc'); 

hfd_color = [ 0.2, 0.5, 0.98];
t2d_color = [1,0.7,0.2];

Estimation_folder ='Results/MLMM_final/Estimation';
S = dir(Estimation_folder);
load(strcat(Estimation_folder,'/',S(3).name));
theta2 = Results.xbest;
[ AllSimulations ] = Function_AllSimulations_t2d_prog( theta2 ,func_mex_model, D, utility);

chow4w = AllSimulations.chow4w;
hfd4w = AllSimulations.hfd4w ;
hfd4wt2d = AllSimulations.hfd4wt2d;

IPGTT_chow0 = AllSimulations.IPGTT_chow0 ;
IPGTT_hfd0 = AllSimulations.IPGTT_hfd0;
IPGTT_hfd0t2d = AllSimulations.IPGTT_hfd0t2d ;

IPGTT_chow1w = AllSimulations.IPGTT_chow1w ;
IPGTT_hfd1w = AllSimulations.IPGTT_hfd1w ;
IPGTT_hfd1wt2d = AllSimulations.IPGTT_hfd1wt2d;

IPGTT_chow2w = AllSimulations.IPGTT_chow2w;
IPGTT_hfd2w = AllSimulations.IPGTT_hfd2w   ;
IPGTT_hfd2wt2d = AllSimulations.IPGTT_hfd2wt2d ;

IPGTT_chow3w = AllSimulations.IPGTT_chow3w;
IPGTT_hfd3w = AllSimulations.IPGTT_hfd3w ;
IPGTT_hfd3wt2d = AllSimulations.IPGTT_hfd3wt2d ;

IPGTT_chow4w = AllSimulations.IPGTT_chow4w ;
IPGTT_hfd4w = AllSimulations.IPGTT_hfd4w ;
IPGTT_hfd4wt2d = AllSimulations.IPGTT_hfd4wt2d;


%% Plotting
TrainColor = [ 0.5,0.5,0.5];

xlabel_long ='time (days)';
%%
figure(1)
hold on
plot_line(1,2,1,hfd4w.variablevalues( : , id_Gconc ) ,hfd4w.time,'1',xlabel_long,{'Plasma gluocse';'(mmol/L)'},hfd_color)
plot_line(1,2,1,hfd4wt2d.variablevalues( : , id_Gconc ) ,hfd4wt2d.time,'1',xlabel_long,{'Plasma gluocse';'(mmol/L)'},t2d_color)
xlim([0 28])
xticks([ 0 7 14 21 28]);

hold on
plot_line(1,2,2,hfd4w.variablevalues( : , id_Iconc )  ,hfd4w.time,'1',xlabel_long,{'Plasma insulin';'(pmol/L)'},hfd_color)
plot_line(1,2,2,hfd4wt2d.variablevalues( : , id_Iconc )  ,hfd4wt2d.time,'1',xlabel_long,{'Plasma insulin';'(pmol/L)'},t2d_color)
xlim([0 28])
xticks([ 0 7 14 21 28]);

%%

figure(2)
set(figure(2), 'outerposition',[0 0 2560 1440], 'PaperType','a4')
plot_line(3,4,1:4,chow4w.statevalues( : , id_FM )./chow4w.statevalues( 1 , id_FM )  ,chow4w.time,'1','1','1',TrainColor)
hold on
plot_line(3,4,1:4,hfd4w.statevalues( : , id_FM )./hfd4w.statevalues( 1 , id_FM )  ,hfd4w.time,'1',xlabel_long,'Fat mass gain (g)',hfd_color)
plot_line(3,4,1:4,hfd4wt2d.statevalues( : , id_FM )./hfd4wt2d.statevalues( 1 , id_FM )  ,hfd4wt2d.time,'1',xlabel_long,'Fat mass(g)',t2d_color,'--')
xlim([0 28])
xticks([ 0 7 14 21 28]);

%% IPGTT 
%%% Glucose
% figure(2)
% set(figure(2), 'outerposition',[0 0 2560 1440], 'PaperType','a4')
xlabel_short ='time (min)';
time_short = (IPGTT_hfd4w.time - IPGTT_hfd4w.time(1))*24*60;

plot_line(3,4,5,IPGTT_chow0.variablevalues( : , id_Gconc ) ,time_short,'1',xlabel_short,{'Plasma gluocse';'(mmol/L)'},TrainColor)
plot_line(3,4,5,IPGTT_hfd0.variablevalues( : , id_Gconc ) ,time_short,'1',xlabel_short,{'Plasma gluocse';'(mmol/L)'},hfd_color)
plot_line(3,4,5,IPGTT_hfd0t2d.variablevalues( : , id_Gconc ) ,time_short,'1',xlabel_short,{'Plasma gluocse';'(mmol/L)'},t2d_color)
ylim([ 0 40])

plot_line(3,4,6,IPGTT_chow1w.variablevalues( : , id_Gconc ) ,time_short,'1',xlabel_short,{''},TrainColor)
plot_line(3,4,6,IPGTT_hfd1w.variablevalues( : , id_Gconc ) ,time_short,'1',xlabel_short,{''},hfd_color)
plot_line(3,4,6,IPGTT_hfd1wt2d.variablevalues( : , id_Gconc ) ,time_short,'1',xlabel_short,{''},t2d_color)
ylim([ 0 40])

plot_line(3,4,7,IPGTT_chow2w.variablevalues( : , id_Gconc ) ,time_short,'1',xlabel_short,{''},TrainColor)
plot_line(3,4,7,IPGTT_hfd2w.variablevalues( : , id_Gconc ) ,time_short,'1',xlabel_short,{''},hfd_color)
plot_line(3,4,7,IPGTT_hfd2wt2d.variablevalues( : , id_Gconc ) ,time_short,'1',xlabel_short,{''},t2d_color)
ylim([ 0 40])

plot_line(3,4,8,IPGTT_chow4w.variablevalues( : , id_Gconc ) ,time_short,'1',xlabel_short,{''},TrainColor)
plot_line(3,4,8,IPGTT_hfd4w.variablevalues( : , id_Gconc ) ,time_short,'1',xlabel_short,{''},hfd_color)
plot_line(3,4,8,IPGTT_hfd4wt2d.variablevalues( : , id_Gconc ) ,time_short,'1',xlabel_short,{''},t2d_color)
ylim([ 0 40])


%% insulin dependent uptake
plot_line(3,4,9,IPGTT_chow0.variablevalues( : , id_U_id ) ,time_short,'1',xlabel_short,{' Insulin dependent';'glucose uptake (mmol/h)'},TrainColor)
plot_line(3,4,9,IPGTT_hfd0.variablevalues( : , id_U_id ) ,time_short,'1',xlabel_short,{' Insulin dependent';'glucose uptake (mmol/h)'},hfd_color)
plot_line(3,4,9,IPGTT_hfd0t2d.variablevalues( : , id_U_id ) ,time_short,'1',xlabel_short,{' Insulin dependent';'glucose uptake (mmol/h)'},t2d_color)

plot_line(3,4,10,IPGTT_chow1w.variablevalues( : , id_U_id ) ,time_short,'1',xlabel_short,{''},TrainColor)
plot_line(3,4,10,IPGTT_hfd1w.variablevalues( : , id_U_id ) ,time_short,'1',xlabel_short,{''},hfd_color)
plot_line(3,4,10,IPGTT_hfd1wt2d.variablevalues( : , id_U_id ) ,time_short,'1',xlabel_short,{''},t2d_color)

plot_line(3,4,11,IPGTT_chow2w.variablevalues( : , id_U_id ) ,time_short,'1',xlabel_short,{''},TrainColor)
plot_line(3,4,11,IPGTT_hfd2w.variablevalues( : , id_U_id ) ,time_short,'1',xlabel_short,{''},hfd_color)
plot_line(3,4,11,IPGTT_hfd2wt2d.variablevalues( : , id_U_id ) ,time_short,'1',xlabel_short,{''},t2d_color)

plot_line(3,4,12,IPGTT_chow4w.variablevalues( : , id_U_id ) ,time_short,'1',xlabel_short,{''},TrainColor)
plot_line(3,4,12,IPGTT_hfd4w.variablevalues( : , id_U_id ) ,time_short,'1',xlabel_short,{''},hfd_color)
plot_line(3,4,12,IPGTT_hfd4wt2d.variablevalues( : , id_U_id ) ,time_short,'1',xlabel_short,{''},t2d_color)


%%

function [] = plot_line(x,y,z, values ,t ,titstring,xstring,ystring,x_color,style)


if nargin < 10
   style = '-'; 
end

label_size = 25;
label_size_ax = 18;

subplot(x,y,z)
hold on
plot(t,values,style,'color',x_color,'Linewidth',3)
hold on
%plot( 8:1:14,best_sim,'color', TrainColor + 0.1 , 'linewidth', 2)
xlabel(xstring,'FontSize', label_size )
ylabel(ystring,'FontSize', label_size )
box off
set(gca,'FontSize',label_size_ax, 'FontWeight','bold')
%t=title(titstring,'FontSize',16,'FontWeight','bold');
end
```
