## Supplementary scripts for "A multi-scale *in silico* mouse model for insulin resistance and humanoid type 2 diabetes": Figure_7C.m

```
p=utility.p0; % for convenience 
id_IRS = ismember(SBvariables(objModel),'totalIRS1'); 
%id_PKB_phos308 = ismember(SBvariables(objModel),'totalPKB473'); 
id_PKB = ismember(SBvariables(objModel),'totalPKB'); 
id_AS = ismember(SBstates(objModel),'AS160'); 
id_As_phos  = ismember(SBstates(objModel),'AS160p'); 
id_RABGTP  =  ismember(SBstates(objModel),'RABGTP'); 
id_PMAa   =  ismember(SBstates(objModel),'PMAa'); 
id_GLUT4m = ismember(SBvariables(objModel),'GLUT4m'); 
id_CL_GIA = ismember(SBvariables(objModel),'CL_GIA'); 
id_PKBda   =  ismember(SBstates(objModel),'PKBda'); 
id_FFM = ismember(SBstates(objModel),'FFM') ; 
id_FM = ismember(SBstates(objModel),'FM');
id_FFMname = ismember(pNames,'FFMinit') ; id_FMname = ismember(pNames,'FMinit');
id_insulin_input = ismember(SBvariables(objModel),'insulin_input'); 
id_U_A = ismember(SBvariables(objModel),'U_id_A_plot'); 
id_U_M = ismember(SBvariables(objModel),'U_id_M'); 
id_U_H = ismember(SBvariables(objModel),'U_id_H'); 
id_U_id = ismember(SBvariables(objModel),'U_id'); 
id_U_ii = ismember(SBvariables(objModel),'U_ii'); 
id_ISEC = ismember(SBvariables(objModel),'ISEC'); 
id_Beta_failure = ismember(SBstates(objModel),'Beta_failure');
id_T2deffect = ismember(SBvariables(objModel),'T2deffect');
id_BW = ismember(SBvariables(objModel),'BW');
id_Iconc = ismember(SBvariables(objModel),'Iconc'); 
id_Gconc = ismember(SBvariables(objModel),'Gconc'); 

hfd_color = [ 0.2, 0.5, 0.98];
t2d_color = [1,0.7,0.2];

Estimation_folder ='Results/MLMM_final/Estimation';
S = dir(Estimation_folder);
load(strcat(Estimation_folder,'/',S(3).name));
theta2 = Results.xbest;
[ AllSimulations ] = Function_AllSimulations_t2d_prog_invitro_exp( theta2 ,func_mex_model, D, utility);

chow4w = AllSimulations.chow4w;
hfd4w = AllSimulations.hfd4w ;
hfd4wt2d = AllSimulations.hfd4wt2d;

invitro_chow0 = AllSimulations.invitro_chow0 ;
invitro_hfd0 = AllSimulations.invitro_hfd0;
invitro_hfd0t2d = AllSimulations.invitro_hfd0t2d ;

invitro_chow1w = AllSimulations.invitro_chow1w ;
invitro_hfd1w = AllSimulations.invitro_hfd1w ;
invitro_hfd1wt2d = AllSimulations.invitro_hfd1wt2d;

invitro_chow2w = AllSimulations.invitro_chow2w;
invitro_hfd2w = AllSimulations.invitro_hfd2w   ;
invitro_hfd2wt2d = AllSimulations.invitro_hfd2wt2d ;

invitro_chow3w = AllSimulations.invitro_chow3w;
invitro_hfd3w = AllSimulations.invitro_hfd3w ;
invitro_hfd3wt2d = AllSimulations.invitro_hfd3wt2d ;

invitro_chow4w = AllSimulations.invitro_chow4w ;
invitro_hfd4w = AllSimulations.invitro_hfd4w ;
invitro_hfd4wt2d = AllSimulations.invitro_hfd4wt2d;

TrainColor = [ 0.5,0.5,0.5];


%% In vivo insulin simulation

figure(3)
time_short = (invitro_hfd4w.time - invitro_hfd4w.time(1))*24*60;
xlabel_short ='time (min)';
label_size = 26;

set(figure(3), 'outerposition',[0 0 2560 1440], 'PaperType','a4') 

id_PKB_phos308 = ismember(SBvariables(objModel),'totalPKB308'); 

plot_line_intracell(2,4,1,invitro_chow0.variablevalues( : , id_PKB_phos308 ) ,time_short,'1',xlabel_short,{'1'},TrainColor)
plot_line_intracell(2,4,1,invitro_hfd0.variablevalues( : , id_PKB_phos308 ) ,time_short,'1',xlabel_short,{'1'},hfd_color)
plot_line_intracell(2,4,1,invitro_hfd0t2d.variablevalues( : , id_PKB_phos308 ) ,time_short,'1',xlabel_short,{'1'},t2d_color,'--')
ylabel({'PKB308p';'(a.u)'},'FontSize',label_size, 'fontweight','bold')
ylim([0 0.04 ])

plot_line_intracell(2,4,2,invitro_chow1w.variablevalues( : , id_PKB_phos308 ) ,time_short,'1',xlabel_short,{'1'},TrainColor)
plot_line_intracell(2,4,2,invitro_hfd1w.variablevalues( : , id_PKB_phos308 ) ,time_short,'1',xlabel_short,{'1'},hfd_color)
plot_line_intracell(2,4,2,invitro_hfd1wt2d.variablevalues( : , id_PKB_phos308 ) ,time_short,'1',xlabel_short,{'1'},t2d_color,'--')
ylim([0 0.04 ])

plot_line_intracell(2,4,3,invitro_chow2w.variablevalues( : , id_PKB_phos308 ) ,time_short,'1',xlabel_short,{'1'},TrainColor)
plot_line_intracell(2,4,3,invitro_hfd2w.variablevalues( : , id_PKB_phos308 ) ,time_short,'1',xlabel_short,{'1'},hfd_color)
plot_line_intracell(2,4,3,invitro_hfd2wt2d.variablevalues( : , id_PKB_phos308 ) ,time_short,'1',xlabel_short,{'1'},t2d_color,'--')
ylim([0 0.04 ])

plot_line_intracell(2,4,4,invitro_chow4w.variablevalues( : , id_PKB_phos308 ) ,time_short,'1',xlabel_short,{'1'},TrainColor)
plot_line_intracell(2,4,4,invitro_hfd4w.variablevalues( : , id_PKB_phos308 ) ,time_short,'1',xlabel_short,{'1'},hfd_color)
plot_line_intracell(2,4,4,invitro_hfd4wt2d.variablevalues( : , id_PKB_phos308 ) ,time_short,'1',xlabel_short,{'1'},t2d_color,'--')
ylim([0 0.04 ])

id_As_phos  = ismember(SBstates(objModel),'AS160p'); 

plot_line_intracell(2,4,5,invitro_chow0.statevalues( : , id_As_phos ) ,time_short,'1',xlabel_short,{'1'},TrainColor)
plot_line_intracell(2,4,5,invitro_hfd0.statevalues( : , id_As_phos ) ,time_short,'1',xlabel_short,{'1'},hfd_color)
plot_line_intracell(2,4,5,invitro_hfd0t2d.statevalues( : , id_As_phos ) ,time_short,'1',xlabel_short,{'1'},t2d_color,'--')
ylabel({'AS160p';'(a.u)'},'FontSize',label_size, 'fontweight','bold')
ylim([0 0.16 ])

plot_line_intracell(2,4,6,invitro_chow1w.statevalues( : , id_As_phos ) ,time_short,'1',xlabel_short,{'1'},TrainColor)
plot_line_intracell(2,4,6,invitro_hfd1w.statevalues( : , id_As_phos ) ,time_short,'1',xlabel_short,{'1'},hfd_color)
plot_line_intracell(2,4,6,invitro_hfd1wt2d.statevalues( : , id_As_phos ) ,time_short,'1',xlabel_short,{'1'},t2d_color,'--')
ylim([0 0.16 ])

plot_line_intracell(2,4,7,invitro_chow2w.statevalues( : , id_As_phos ) ,time_short,'1',xlabel_short,{'1'},TrainColor)
plot_line_intracell(2,4,7,invitro_hfd2w.statevalues( : , id_As_phos ) ,time_short,'1',xlabel_short,{'1'},hfd_color)
plot_line_intracell(2,4,7,invitro_hfd2wt2d.statevalues( : , id_As_phos ) ,time_short,'1',xlabel_short,{'1'},t2d_color,'--')
ylim([0 0.16 ])

plot_line_intracell(2,4,8,invitro_chow4w.statevalues( : , id_As_phos ) ,time_short,'1',xlabel_short,{'1'},TrainColor)
plot_line_intracell(2,4,8,invitro_hfd4w.statevalues( : , id_As_phos ) ,time_short,'1',xlabel_short,{'1'},hfd_color)
plot_line_intracell(2,4,8,invitro_hfd4wt2d.statevalues( : , id_As_phos ) ,time_short,'1',xlabel_short,{'1'},t2d_color,'--')
ylim([0 0.16 ])


%%
function [] = plot_line_intracell(x,y,z, values ,t ,titstring,xstring,ystring,x_color,style)
label_size_ax = 18;

if nargin < 10
   style = '-'; 
end

subplot(x,y,z)
hold on
plot(t,values,style,'color',x_color,'Linewidth',3)
hold on
box off
set(gca,'FontSize',label_size_ax, 'FontWeight','bold')
xlabel(xstring,'FontSize',label_size_ax,'Fontweight','bold' )

%t=title(titstring,'FontSize',16,'FontWeight','bold');
end

function [] = plot_line(x,y,z, values ,t ,titstring,xstring,ystring,x_color)

label_size = 25;
label_size_ax = 18;

subplot(x,y,z)
hold on
plot(t,values,'color',x_color,'Linewidth',3)
hold on
%plot( 8:1:14,best_sim,'color', TrainColor + 0.1 , 'linewidth', 2)
%xlabel(xstring,'FontSize', label_size )
%ylabel(ystring,'FontSize', label_size )
box off
set(gca,'FontSize',label_size_ax, 'FontWeight','bold')
%t=title(titstring,'FontSize',16,'FontWeight','bold');
end
```
