## Supplementary scripts for "A multi-scale *in silico* mouse model for insulin resistance and humanoid type 2 diabetes": Figure_SA1.m

```
%% Setup
p=utility.p0;
id_FFM = ismember(SBstates(objModel),'FFM') ; id_FM = ismember(SBstates(objModel),'FM');
id_FFMname = ismember(pNames,'FFMinit') ; id_FMname = ismember(pNames,'FMinit');
t_diet_up=0:1:56;
t_fasting_up=[ 42  42+0.5833 ]  ;  %t_diet:0.0001:t_diet+0.5833;
t_IPGTT_up=(0:1:120)/(60*24)+t_fasting_up(end);

t_diet=14;
t_fasting=[ t_diet  t_diet+0.5833 ]  ;  %t_diet:0.0001:t_diet+0.5833;
t_IPGTT=(0:1:120)/(60*24)+t_fasting(end);

id_Uid = ismember(SBvariables(objModel),'U_id');
id_Uii = ismember(SBvariables(objModel),'U_ii');
id_UidA = ismember(SBvariables(objModel),'U_id_A');
id_UidM = ismember(SBvariables(objModel),'U_id_M');
id_UidH = ismember(SBvariables(objModel),'U_id_H');

%% UNC
for i = 1:nfiles
load( strcat( filedir ,'/',files(i).name ) )
disp(strcat('Processing - ',files(i).name, int2str(i),'/',int2str(nfiles)))
theta=Results.xbest;


[ AllSimulations ] = Function_AllSimulations( theta ,func_mex_model, D, utility);

chow8w             = AllSimulations.chow8w;
hfd8w              = AllSimulations.hfd8w;
IPGTT_hfd6w        = AllSimulations.IPGTT_hfd6w;
chow2w              = AllSimulations.chow2w;
IPGTT_chow2w        = AllSimulations.IPGTT_chow2w;
chow12dHFD2d        = AllSimulations.chow12dHFD2d;
IPGTT_chow12dHFD2d  = AllSimulations.IPGTT_chow12dHFD2d;
chow10dHFD4d        = AllSimulations.chow10dHFD4d;
chow8dHFD6d         = AllSimulations.chow8dHFD6d;
IPGTT_chow8dHFD6d   = AllSimulations.IPGTT_chow8dHFD6d;
hfd2w               = AllSimulations.hfd2w;
IPGTT_hfd2w         = AllSimulations.IPGTT_hfd2w;

%% MIN and MAX value calculations

%% Predictions
if i == 1 % Setting up vectors  
Max_hfd6w_Uid=  IPGTT_hfd6w.variablevalues(:,id_Uid);  Min_hfd6w_Uid =  IPGTT_hfd6w.variablevalues(:,id_Uid); 
Max_hfd6w_Uii =  IPGTT_hfd6w.variablevalues(:,id_Uii);  Min_hfd6w_Uii =  IPGTT_hfd6w.variablevalues(:,id_Uii); 

Max_hfd6w_UidA =  IPGTT_hfd6w.variablevalues(:,id_UidA);  Min_hfd6w_UidA  =  IPGTT_hfd6w.variablevalues(:,id_UidA); 
Max_hfd6w_UidM =  IPGTT_hfd6w.variablevalues(:,id_UidM);  Min_hfd6w_UidM  =  IPGTT_hfd6w.variablevalues(:,id_UidM); 
Max_hfd6w_UidH =  IPGTT_hfd6w.variablevalues(:,id_UidH);  Min_hfd6w_UidH  =  IPGTT_hfd6w.variablevalues(:,id_UidH); 
end
Max_hfd6w_Uid( Max_hfd6w_Uid < IPGTT_hfd6w.variablevalues(:,id_Uid) ) =  IPGTT_hfd6w.variablevalues( Max_hfd6w_Uid < IPGTT_hfd6w.variablevalues(:,id_Uid),id_Uid);
Min_hfd6w_Uid( Min_hfd6w_Uid > IPGTT_hfd6w.variablevalues(:,id_Uid) ) =  IPGTT_hfd6w.variablevalues( Min_hfd6w_Uid > IPGTT_hfd6w.variablevalues(:,id_Uid),id_Uid);

Max_hfd6w_Uii( Max_hfd6w_Uii < IPGTT_hfd6w.variablevalues(:,id_Uii) ) =  IPGTT_hfd6w.variablevalues( Max_hfd6w_Uii < IPGTT_hfd6w.variablevalues(:,id_Uii),id_Uii);
Min_hfd6w_Uii( Min_hfd6w_Uii > IPGTT_hfd6w.variablevalues(:,id_Uii) ) =  IPGTT_hfd6w.variablevalues( Min_hfd6w_Uii > IPGTT_hfd6w.variablevalues(:,id_Uii),id_Uii);

Max_hfd6w_UidA( Max_hfd6w_UidA < IPGTT_hfd6w.variablevalues(:,id_UidA) ) =  IPGTT_hfd6w.variablevalues( Max_hfd6w_UidA < IPGTT_hfd6w.variablevalues(:,id_UidA),id_UidA);
Min_hfd6w_UidA( Min_hfd6w_UidA > IPGTT_hfd6w.variablevalues(:,id_UidA) ) =  IPGTT_hfd6w.variablevalues( Min_hfd6w_UidA > IPGTT_hfd6w.variablevalues(:,id_UidA),id_UidA);

Max_hfd6w_UidM( Max_hfd6w_UidM < IPGTT_hfd6w.variablevalues(:,id_UidM) ) =  IPGTT_hfd6w.variablevalues( Max_hfd6w_UidM < IPGTT_hfd6w.variablevalues(:,id_UidM),id_UidM);
Min_hfd6w_UidM( Min_hfd6w_UidM > IPGTT_hfd6w.variablevalues(:,id_UidM) ) =  IPGTT_hfd6w.variablevalues( Min_hfd6w_UidM > IPGTT_hfd6w.variablevalues(:,id_UidM),id_UidM);

Max_hfd6w_UidH( Max_hfd6w_UidH < IPGTT_hfd6w.variablevalues(:,id_UidH) ) =  IPGTT_hfd6w.variablevalues( Max_hfd6w_UidH < IPGTT_hfd6w.variablevalues(:,id_UidH),id_UidH);
Min_hfd6w_UidH( Min_hfd6w_UidH > IPGTT_hfd6w.variablevalues(:,id_UidH) ) =  IPGTT_hfd6w.variablevalues( Min_hfd6w_UidH > IPGTT_hfd6w.variablevalues(:,id_UidH),id_UidH);


%% Hansson et al (2018)
if i == 1  % Setting up vectors
Max_chow2w_Uid=  IPGTT_chow2w.variablevalues(:,id_Uid);  Min_chow2w_Uid =  IPGTT_chow2w.variablevalues(:,id_Uid); 
Max_chow2w_Uii =  IPGTT_chow2w.variablevalues(:,id_Uii);  Min_chow2w_Uii =  IPGTT_chow2w.variablevalues(:,id_Uii); 
Max_chow2w_UidA =  IPGTT_chow2w.variablevalues(:,id_UidA);  Min_chow2w_UidA  =  IPGTT_chow2w.variablevalues(:,id_UidA); 
Max_chow2w_UidM =  IPGTT_chow2w.variablevalues(:,id_UidM);  Min_chow2w_UidM  =  IPGTT_chow2w.variablevalues(:,id_UidM); 
Max_chow2w_UidH =  IPGTT_chow2w.variablevalues(:,id_UidH);  Min_chow2w_UidH  =  IPGTT_chow2w.variablevalues(:,id_UidH); 

Max_hfd2w_Uid=  IPGTT_hfd2w.variablevalues(:,id_Uid);  Min_hfd2w_Uid =  IPGTT_hfd2w.variablevalues(:,id_Uid); 
Max_hfd2w_Uii =  IPGTT_hfd2w.variablevalues(:,id_Uii);  Min_hfd2w_Uii =  IPGTT_hfd2w.variablevalues(:,id_Uii); 
Max_hfd2w_UidA =  IPGTT_hfd2w.variablevalues(:,id_UidA);  Min_hfd2w_UidA  =  IPGTT_hfd2w.variablevalues(:,id_UidA); 
Max_hfd2w_UidM =  IPGTT_hfd2w.variablevalues(:,id_UidM);  Min_hfd2w_UidM  =  IPGTT_hfd2w.variablevalues(:,id_UidM); 
Max_hfd2w_UidH =  IPGTT_hfd2w.variablevalues(:,id_UidH);  Min_hfd2w_UidH  =  IPGTT_hfd2w.variablevalues(:,id_UidH); 
end 

Max_chow2w_Uid( Max_chow2w_Uid < IPGTT_chow2w.variablevalues(:,id_Uid) ) =  IPGTT_chow2w.variablevalues( Max_chow2w_Uid < IPGTT_chow2w.variablevalues(:,id_Uid),id_Uid);
Min_chow2w_Uid( Min_chow2w_Uid > IPGTT_chow2w.variablevalues(:,id_Uid) ) =  IPGTT_chow2w.variablevalues( Min_chow2w_Uid > IPGTT_chow2w.variablevalues(:,id_Uid),id_Uid);

Max_chow2w_Uii( Max_chow2w_Uii < IPGTT_chow2w.variablevalues(:,id_Uii) ) =  IPGTT_chow2w.variablevalues( Max_chow2w_Uii < IPGTT_chow2w.variablevalues(:,id_Uii),id_Uii);
Min_chow2w_Uii( Min_chow2w_Uii > IPGTT_chow2w.variablevalues(:,id_Uii) ) =  IPGTT_chow2w.variablevalues( Min_chow2w_Uii > IPGTT_chow2w.variablevalues(:,id_Uii),id_Uii);

Max_chow2w_UidA( Max_chow2w_UidA < IPGTT_chow2w.variablevalues(:,id_UidA) ) =  IPGTT_chow2w.variablevalues( Max_chow2w_UidA < IPGTT_chow2w.variablevalues(:,id_UidA),id_UidA);
Min_chow2w_UidA( Min_chow2w_UidA > IPGTT_chow2w.variablevalues(:,id_UidA) ) =  IPGTT_chow2w.variablevalues( Min_chow2w_UidA > IPGTT_chow2w.variablevalues(:,id_UidA),id_UidA);

Max_chow2w_UidM( Max_chow2w_UidM < IPGTT_chow2w.variablevalues(:,id_UidM) ) =  IPGTT_chow2w.variablevalues( Max_chow2w_UidM < IPGTT_chow2w.variablevalues(:,id_UidM),id_UidM);
Min_chow2w_UidM( Min_chow2w_UidM > IPGTT_chow2w.variablevalues(:,id_UidM) ) =  IPGTT_chow2w.variablevalues( Min_chow2w_UidM > IPGTT_chow2w.variablevalues(:,id_UidM),id_UidM);

Max_chow2w_UidH( Max_chow2w_UidH < IPGTT_chow2w.variablevalues(:,id_UidH) ) =  IPGTT_chow2w.variablevalues( Max_chow2w_UidH < IPGTT_chow2w.variablevalues(:,id_UidH),id_UidH);
Min_chow2w_UidH( Min_chow2w_UidH > IPGTT_chow2w.variablevalues(:,id_UidH) ) =  IPGTT_chow2w.variablevalues( Min_chow2w_UidH > IPGTT_chow2w.variablevalues(:,id_UidH),id_UidH);


Max_hfd2w_Uid( Max_hfd2w_Uid < IPGTT_hfd2w.variablevalues(:,id_Uid) ) =  IPGTT_hfd2w.variablevalues( Max_hfd2w_Uid < IPGTT_hfd2w.variablevalues(:,id_Uid),id_Uid);
Min_hfd2w_Uid( Min_hfd2w_Uid > IPGTT_hfd2w.variablevalues(:,id_Uid) ) =  IPGTT_hfd2w.variablevalues( Min_hfd2w_Uid > IPGTT_hfd2w.variablevalues(:,id_Uid),id_Uid);

Max_hfd2w_Uii( Max_hfd2w_Uii < IPGTT_hfd2w.variablevalues(:,id_Uii) ) =  IPGTT_hfd2w.variablevalues( Max_hfd2w_Uii < IPGTT_hfd2w.variablevalues(:,id_Uii),id_Uii);
Min_hfd2w_Uii( Min_hfd2w_Uii > IPGTT_hfd2w.variablevalues(:,id_Uii) ) =  IPGTT_hfd2w.variablevalues( Min_hfd2w_Uii > IPGTT_hfd2w.variablevalues(:,id_Uii),id_Uii);

Max_hfd2w_UidA( Max_hfd2w_UidA < IPGTT_hfd2w.variablevalues(:,id_UidA) ) =  IPGTT_hfd2w.variablevalues( Max_hfd2w_UidA < IPGTT_hfd2w.variablevalues(:,id_UidA),id_UidA);
Min_hfd2w_UidA( Min_hfd2w_UidA > IPGTT_hfd2w.variablevalues(:,id_UidA) ) =  IPGTT_hfd2w.variablevalues( Min_hfd2w_UidA > IPGTT_hfd2w.variablevalues(:,id_UidA),id_UidA);

Max_hfd2w_UidM( Max_hfd2w_UidM < IPGTT_hfd2w.variablevalues(:,id_UidM) ) =  IPGTT_hfd2w.variablevalues( Max_hfd2w_UidM < IPGTT_hfd2w.variablevalues(:,id_UidM),id_UidM);
Min_hfd2w_UidM( Min_hfd2w_UidM > IPGTT_hfd2w.variablevalues(:,id_UidM) ) =  IPGTT_hfd2w.variablevalues( Min_hfd2w_UidM > IPGTT_hfd2w.variablevalues(:,id_UidM),id_UidM);

Max_hfd2w_UidH( Max_hfd2w_UidH < IPGTT_hfd2w.variablevalues(:,id_UidH) ) =  IPGTT_hfd2w.variablevalues( Max_hfd2w_UidH < IPGTT_hfd2w.variablevalues(:,id_UidH),id_UidH);
Min_hfd2w_UidH( Min_hfd2w_UidH > IPGTT_hfd2w.variablevalues(:,id_UidH) ) =  IPGTT_hfd2w.variablevalues( Min_hfd2w_UidH > IPGTT_hfd2w.variablevalues(:,id_UidH),id_UidH);


end
%% Plotting

predColor =  [0.9,.9,0.9]; 
label_size = 25;
label_size_ax = 18;
marker_size = 5;
fig=0;

%% Simulations for best cost.

Estimation_folder ='Results/MLMM_final/Estimation';
S = dir(Estimation_folder);
load(strcat(Estimation_folder,'/',S(3).name));
theta2 = Results.xbest;
[ AllSimulations_train ] = Function_AllSimulations( theta2 ,func_mex_model, D, utility);


%% IPGTT
%% figure 1 - IPGTT - TRAINING AND PREDIKTION
fill_time_IPGTT = [ t_IPGTT-t_fasting(end) fliplr(t_IPGTT-t_fasting(end)) ]*24*60;
tp_IPGTT=[0 15 30 60 120];
time_best_cost = (t_IPGTT-t_fasting(end) )*24*60; 
fig=fig+1;
figure(fig)

col1 = [ 0.5,0.5,1];
col2 = [ 0.5,0.5,0.5];
col3 = [ 0.5,1,0.5];
col4 = [ 1,0.5,0.5];
col5 = [ 0.8,0.8,0.5];

Experiments_to_plot = {'chow2w','hfd2w','hfd6w'};
IPGTT_sim_name = {'IPGTT_chow2w','IPGTT_hfd2w','IPGTT_hfd6w'};
for kk = 1:length(Experiments_to_plot)
values=[];
data.time=tp_IPGTT;
data.mean = D.IPGTTgcc.Mean(:,kk);
data.SEM = D.IPGTTgcc.SEM(:,kk);
eval(strcat('values1(:,1)=Max_',Experiments_to_plot{kk},'_Uid;')); 
eval(strcat('values1(:,2)=Min_',Experiments_to_plot{kk},'_Uid;')); 

eval(strcat('values2(:,1)=Max_',Experiments_to_plot{kk},'_Uii;')); 
eval(strcat('values2(:,2)=Min_',Experiments_to_plot{kk},'_Uii;')); 

eval(strcat('values3(:,1)=Max_',Experiments_to_plot{kk},'_UidA;')); 
eval(strcat('values3(:,2)=Min_',Experiments_to_plot{kk},'_UidA;')); 

eval(strcat('values4(:,1)=Max_',Experiments_to_plot{kk},'_UidM;')); 
eval(strcat('values4(:,2)=Min_',Experiments_to_plot{kk},'_UidM;')); 

eval(strcat('values5(:,1)=Max_',Experiments_to_plot{kk},'_UidH;')); 
eval(strcat('values5(:,2)=Min_',Experiments_to_plot{kk},'_UidH;')); 


time_fill = fill_time_IPGTT;
time_plot = (t_IPGTT-t_fasting(end))*24*60;
subplot(2,3,kk) %% a)
f=fill(time_fill, [ values1(:,2)' fliplr( values1(:,1)' ) ], col1, 'EdgeColor', 'none' );  
set(f,'facealpha',.5)
hold on
f=fill(time_fill, [ values2(:,2)' fliplr( values2(:,1)' ) ], col2, 'EdgeColor', 'none' );  
set(f,'facealpha',.5)
hold on
eval(strcat('plot( time_plot, AllSimulations_train.',IPGTT_sim_name{kk}, ".variablevalues(:,id_Uid),'color', col1 - 0.2 , 'linewidth', 2);"))
eval(strcat('plot( time_plot, AllSimulations_train.',IPGTT_sim_name{kk}, ".variablevalues(:,id_Uii),'color', col2 - 0.2 , 'linewidth', 2);"))
xlim([0 120])
ylim([0 0.35])
title(Experiments_to_plot{kk})
if kk == 1
legend({'Uid','Uii'})
ylabel({'Insulin dependent';'glucose uptake (mmol/h)'},'FontSize',26, 'fontweight','bold')
end
box off
set(gca,'FontSize',label_size_ax, 'FontWeight','bold')
xlabel('Time (min)')


subplot(2,3,kk+3) %% a)
f=fill(time_fill, [ values3(:,2)' fliplr( values3(:,1)' ) ], col3, 'EdgeColor', 'none' );  
set(f,'facealpha',.5)
hold on
f=fill(time_fill, [ values4(:,2)' fliplr( values4(:,1)' ) ], col4, 'EdgeColor', 'none' );  
set(f,'facealpha',.5)
hold on
f=fill(time_fill, [ values5(:,2)' fliplr( values5(:,1)' ) ], col5, 'EdgeColor', 'none' );  
set(f,'facealpha',.5)
hold on

eval(strcat('plot( time_plot, AllSimulations_train.',IPGTT_sim_name{kk}, ".variablevalues(:,id_UidA),'color', col3 - 0.2 , 'linewidth', 2);"))
eval(strcat('plot( time_plot, AllSimulations_train.',IPGTT_sim_name{kk}, ".variablevalues(:,id_UidM),'color', col4 - 0.2 , 'linewidth', 2);"))
eval(strcat('plot( time_plot, AllSimulations_train.',IPGTT_sim_name{kk}, ".variablevalues(:,id_UidH),'color', col5 - 0.2 , 'linewidth', 2);"))
xlim([0 120])
ylim([0 0.18])
title(Experiments_to_plot{kk})

if kk == 1
legend({'UA','UM','UH'})
ylabel({'Insulin dependent';'glucose uptake (mmol/h)'},'FontSize',26, 'fontweight','bold')

end
box off
set(gca,'FontSize',label_size_ax, 'FontWeight','bold')
xlabel('Time (min)')


end
```
