## Supplementary scripts for "A multi-scale *in silico* mouse model for insulin resistance and humanoid type 2 diabetes": Figure_SA2.m

```
p=utility.p0; % for convenience 
id_IRS = ismember(SBvariables(objModel),'totalIRS1'); 
id_PKB_phos473 = ismember(SBvariables(objModel),'totalPKB473'); 
id_PKB = ismember(SBvariables(objModel),'totalPKB'); 
id_AS = ismember(SBstates(objModel),'AS160'); 
id_As_phos  = ismember(SBstates(objModel),'AS160p'); 
id_RABGTP  =  ismember(SBstates(objModel),'RABGTP'); 
id_PMAa   =  ismember(SBstates(objModel),'PMAa'); 
id_GLUT4m = ismember(SBvariables(objModel),'GLUT4m'); 
id_CL_GIA = ismember(SBvariables(objModel),'CL_GIA'); 
id_PKBda   =  ismember(SBstates(objModel),'PKBda'); 
id_FFM = ismember(SBstates(objModel),'FFM') ; 
id_FM = ismember(SBstates(objModel),'FM');
id_FFMname = ismember(pNames,'FFMinit') ; id_FMname = ismember(pNames,'FMinit');
id_insulin_input = ismember(SBvariables(objModel),'insulin_input'); 
id_U_A = ismember(SBvariables(objModel),'U_id_A_plot'); 
id_U_M = ismember(SBvariables(objModel),'U_id_M'); 
id_U_H = ismember(SBvariables(objModel),'U_id_H'); 
id_U_id = ismember(SBvariables(objModel),'U_id'); 
id_U_ii = ismember(SBvariables(objModel),'U_ii'); 
id_ISEC = ismember(SBvariables(objModel),'ISEC'); 
id_Beta_failure = ismember(SBstates(objModel),'Beta_failure');
id_T2deffect = ismember(SBvariables(objModel),'T2deffect');
id_BW = ismember(SBvariables(objModel),'BW');
id_Iconc = ismember(SBvariables(objModel),'Iconc'); 
id_Gconc = ismember(SBvariables(objModel),'Gconc'); 

hfd_color = [ 0.2, 0.5, 0.98];
t2d_color = [1,0.7,0.2];

Estimation_folder ='Results/MLMM_final/Estimation';
S = dir(Estimation_folder);
load(strcat(Estimation_folder,'/',S(3).name));
theta2 = Results.xbest;
[ AllSimulations ] = Function_AllSimulations_t2d( theta2 ,func_mex_model, D, utility);

chow4w         = AllSimulations.chow4w;
hfd4w          = AllSimulations.hfd4w;              
hfd4wt2d       = AllSimulations.hfd4wt2d;           
IPGTT_chow4w   = AllSimulations.IPGTT_chow4w; 
IPGTT_hfd4w    = AllSimulations.IPGTT_hfd4w;     
IPGTT_hfd4wt2d = AllSimulations.IPGTT_hfd4wt2d;
%% Plot
xlabel_long ='time (days)';
figure(1)
TrainColor = [ 0.5,0.5,0.5];

set(figure(1), 'outerposition',[0 0 2560 1440], 'PaperType','a4')

plot_line(2,5,1,chow4w.variablevalues( : , id_Gconc )  ,chow4w.time,'1','1','1',TrainColor)
plot_line(2,5,1,hfd4w.variablevalues( : , id_Gconc ) ,hfd4w.time,'1',xlabel_long,{'Fasting plasma gluocse';'(mmol/L)'},hfd_color)
plot_line(2,5,1,hfd4wt2d.variablevalues( : , id_Gconc )  ,hfd4wt2d.time,'1',xlabel_long,{'Fasting plasma gluocse';'(mmol/L)'},t2d_color)
xlim([0 28]) 

plot_line(2,5,2,chow4w.variablevalues( : , id_Iconc )  ,chow4w.time,'1','1','1',TrainColor)
plot_line(2,5,2,hfd4w.variablevalues( : , id_Iconc ) ,hfd4w.time,'1',xlabel_long,{'Fasting plasma insulin';'(pmol/L)'},hfd_color)
plot_line(2,5,2,hfd4wt2d.variablevalues( : , id_Iconc )  ,hfd4wt2d.time,'1',xlabel_long,{'Fasting plasma insulin';'(pmol/L)'},t2d_color)
xlim([0 28]) 

plot_line(2,5,3,chow4w.variablevalues( : , id_ISEC )  ,chow4w.time,'1','1','1',TrainColor)
plot_line(2,5,3,hfd4w.variablevalues( : , id_ISEC ) ,hfd4w.time,'1',xlabel_long,{'Fasting insulin secretion';'(pmol/L)'},hfd_color)
plot_line(2,5,3,hfd4wt2d.variablevalues( : , id_ISEC )  ,hfd4wt2d.time,'1',xlabel_long,{'Fasting insulin secretion';'(pmol/L)'},t2d_color)

plot_line(2,5,4,chow4w.variablevalues( : , id_T2deffect )  ,chow4w.time,'1','1','1',TrainColor)
plot_line(2,5,4,hfd4w.variablevalues( : , id_T2deffect ) ,hfd4w.time,'1',xlabel_long,{'T2D effect';'(a.e)'},hfd_color)
plot_line(2,5,4,hfd4wt2d.variablevalues( : , id_T2deffect )  ,hfd4wt2d.time,'1',xlabel_long,{'T2D effect';'(a.e)'},t2d_color)

id_EGP = ismember(SBreactions(objModel),'EGP'); 
plot_line(2,5,5,chow4w.reactionvalues( : , id_EGP )  ,chow4w.time,'1','1','1',TrainColor)
plot_line(2,5,5,hfd4w.reactionvalues( : , id_EGP )  ,hfd4w.time,'1',xlabel_long,{'EGP';'(mmol/h)'},hfd_color)
plot_line(2,5,5,hfd4wt2d.reactionvalues( : , id_EGP )  ,hfd4wt2d.time,'1',xlabel_long,{'EGP';'(mmol/h)'},t2d_color)
xlim([0 28]) 

%%
%figure(2)
%set(figure(2), 'outerposition',[0 0 2560 1440], 'PaperType','a4')
xlabel_short ='time (min)';
time_short = (IPGTT_hfd4w.time - IPGTT_hfd4w.time(1))*24*60;
plot_line(2,5,6,IPGTT_chow4w.variablevalues( : , id_Gconc ) ,time_short,'1',xlabel_short,{'Plasma gluocse';'(mmol/L)'},TrainColor)
plot_line(2,5,6,IPGTT_hfd4w.variablevalues( : , id_Gconc ) ,time_short,'1',xlabel_short,{'Plasma gluocse';'(mmol/L)'},hfd_color)
plot_line(2,5,6,IPGTT_hfd4wt2d.variablevalues( : , id_Gconc ) ,time_short,'1',xlabel_short,{'Plasma gluocse';'(mmol/L)'},t2d_color)

plot_line(2,5,7,IPGTT_chow4w.variablevalues( : , id_Iconc ) ,time_short,'1',xlabel_short,{'Plasma insulin';'(pmol/L)'},TrainColor)
plot_line(2,5,7,IPGTT_hfd4w.variablevalues( : , id_Iconc ) ,time_short,'1',xlabel_short,{'Plasma insulin';'(pmol/L)'},hfd_color)
plot_line(2,5,7,IPGTT_hfd4wt2d.variablevalues( : , id_Iconc ) ,time_short,'1',xlabel_short,{'Plasma insulin';'(pmol/L)'},t2d_color)

plot_line(2,5,8,IPGTT_chow4w.variablevalues( : , id_U_id ) ,time_short,'1',xlabel_short,{'Total insulin dependent';'glucose uptake (mmol/h)'},TrainColor)
plot_line(2,5,8,IPGTT_hfd4w.variablevalues( : , id_U_id ) ,time_short,'1',xlabel_short,{'Total insulin dependent';'glucose uptake (mmol/h)'},hfd_color)
plot_line(2,5,8,IPGTT_hfd4wt2d.variablevalues( : , id_U_id ) ,time_short,'1',xlabel_short,{'Total insulin dependent';'glucose uptake (mmol/h)'},t2d_color)

plot_line(2,5,9,IPGTT_chow4w.variablevalues( : , id_ISEC ) ,time_short,'1',xlabel_short,{'Insulin secreation';'(pmol/h)'},TrainColor)
plot_line(2,5,9,IPGTT_hfd4w.variablevalues( : , id_ISEC ) ,time_short,'1',xlabel_short,{'Insulin secreation';'(pmol/h)'},hfd_color)
plot_line(2,5,9,IPGTT_hfd4wt2d.variablevalues( : , id_ISEC ) ,time_short,'1',xlabel_short,{'Insulin secreation';'(pmol/h)'},t2d_color)

id_EGP = ismember(SBreactions(objModel),'EGP'); 

plot_line(2,5,10,IPGTT_chow4w.reactionvalues( : , id_EGP ) ,time_short,'1',xlabel_short,{'EGP';'(mmol/h)'},TrainColor)
plot_line(2,5,10,IPGTT_hfd4w.reactionvalues( : , id_EGP ) ,time_short,'1',xlabel_short,{'EGP';'(mmol/h)'},hfd_color)
plot_line(2,5,10,IPGTT_hfd4wt2d.reactionvalues( : , id_EGP ) ,time_short,'1',xlabel_short,{'EGP';'(mmol/h)'},t2d_color)


function [] = plot_line(x,y,z, values ,t ,titstring,xstring,ystring,x_color)

label_size = 25;
label_size_ax = 18;

subplot(x,y,z)
hold on
plot(t,values,'color',x_color,'Linewidth',3)
hold on
%plot( 8:1:14,best_sim,'color', TrainColor + 0.1 , 'linewidth', 2)
xlabel(xstring,'FontSize', label_size )
ylabel(ystring,'FontSize', label_size )
box off
set(gca,'FontSize',label_size_ax, 'FontWeight','bold')
%t=title(titstring,'FontSize',16,'FontWeight','bold');
end
```
