## Supplementary scripts for "A multi-scale *in silico* mouse model for insulin resistance and humanoid type 2 diabetes": Function_AllSimulations.m

```
function [AllSimulations] = Function_AllSimulations( theta ,func_mex_model, D, utility)

ic0 = utility.ic0;
p = utility.p0;
objModel = utility.objModel; % variables used for indexing

pNames = utility.pNames ; 
id_FFM = ismember(SBstates(objModel),'FFM') ; id_FM = ismember(SBstates(objModel),'FM');
id_FFMname = ismember(pNames,'FFMinit') ; id_FMname = ismember(pNames,'FMinit');
% parameter and time vector setup
p(17:37)=[10.^theta(8:19) theta(20)  10.^theta(21:end)];  % GSS to ik1
if p(ismember(pNames,'ik1')) == 10^theta(end) % bounds check
else
    disp('check parameter vector')
end
try   
%% Steady state simulation
ic_ss = ic0; p_ss = p; p_ss( ismember(pNames, 'ss_x')) = 0 ;
ic_ss(id_FFM)=1 ; ic_ss(id_FM)= 1; p_ss(id_FFMname)=1; p_ss(id_FMname)=1; % no effect of Weight on tissue-level and a i s- models
p_ss(end) = 1000; %no IPGTT
ss=func_mex_model(0:100:300,ic_ss, p_ss); % steady state simulation
cost_ss_gcc = (( ss.variablevalues(end, end-1)  -  D.IPGTTgcc.Mean(1,1)  )^2 )./( D.IPGTTgcc.SEM(1,1) )^2 ;% basal glucose
cost_ss_icc = (( ss.variablevalues(end, end)  -  D.IPGTTicc.Mean(1,1)  )^2 )./( D.IPGTTicc.SEM(1,1) )^2   ;% basal insulin
ss_cost = cost_ss_gcc + cost_ss_icc;
ic0 = ss.statevalues(end,:); % overwrite ic0

%% Hansson et al (2018) 2w - Diet and IPGTT 
% Setting up parameters and time-vectors
t_diet=14; % diet time in days
t_fasting=[ t_diet  t_diet+0.5833 ]  ;  %fasting
t_IPGTT=(0:1:120)/(60*24)+t_fasting(end);
p(end) = t_fasting(end); % time of IPGTT (triggers after fasting)

% 2w chow
pchow=p;
pchow(1)=10.^theta(5);   % EI   theta(5) - Chow EI
pchow(2)=0.7;            % carb fraction 
pchow(16)= 10.^theta(1); % PA for chow
pchow(id_FFMname)=0.89*D.BW.mean(1,1); pchow(id_FMname)=0.11*D.BW.mean(1,1);
ic_chow = ic0; ic_chow(id_FFM)=0.89*D.BW.mean(1,1) ; ic_chow(id_FM)= 0.11*D.BW.mean(1,1);

% 12 d chow + 2d hfd
pchow12d=p; 
pchow12d(1)=18.1845; 	
pchow12d(2)=0.2;  
pchow12d(16)  = 10.^theta(2); % PA for hfd
% Here we dont have any data for starting weight.

% 10d chow + 4d hfd
pchow10d=p; 
pchow10d(1)=18.9311;    
pchow10d(2)=0.2;  
pchow10d(16)= 10.^theta(2); % PA for hfd
pchow10d(id_FFMname)=0.89*D.BW.mean(2,1); pchow10d(id_FMname)=0.11*D.BW.mean(2,1); 
ic_chow10d = ic0; ic_chow10d(id_FFM)=0.89*D.BW.mean(2,1) ; ic_chow10d(id_FM)= 0.11*D.BW.mean(2,1);

% 8d chow + 6d hfd
pchow8d=p;  
pchow8d(1)=18.1845;     
pchow8d(2)=0.2;   
pchow8d(16)= 10.^theta(2);  % PA for hfd
pchow8d(id_FFMname)=0.89*D.BW.mean(3,1);    pchow8d(id_FMname)=0.11*D.BW.mean(3,1);
ic_chow8d = ic0; ic_chow8d(id_FFM)=0.89*D.BW.mean(3,1) ; ic_chow8d(id_FM)= 0.11*D.BW.mean(3,1);

% 2w hfd
phfd=p;     
phfd(1)= 16.5217;       
phfd(2)=0.2;      
phfd(16)= 10.^theta(2);    % PA for hfd
phfd(id_FFMname)=0.89*D.BW.mean(4,1);    phfd(id_FMname)=0.11*D.BW.mean(4,1);
ic_hfd = ic0; ic_hfd(id_FFM)=0.89*D.BW.mean(4,1) ; ic_hfd(id_FM)= 0.11*D.BW.mean(4,1);

%%% Diet simulations %%%%
%2w chow
chow2w=func_mex_model( (0:1:t_diet),ic_chow, pchow);
% 12 d chow + 2d hfd
% Here we dont have any data for starting weight. Using values from 2 chow
% simulation
ic_chow12d = chow2w.statevalues(13,:) ; % 13 is index for day 12
ic_chow12d(id_FFM)=chow2w.statevalues(13,id_FFM) ; ic_chow12d(id_FM)= chow2w.statevalues(13,id_FM);
pchow12d(id_FFMname)=chow2w.statevalues(13,id_FFM); pchow12d(id_FMname)= chow2w.statevalues(13,id_FM);
chow12dHFD2d=func_mex_model((12:1:t_diet) ,ic_chow12d, pchow12d);
%10d chow + 4d HFD 
chow10dHFD4d=func_mex_model((10:1:t_diet) ,ic_chow10d, pchow10d);  
%8d chow + 6d HFD  
chow8dHFD6d=func_mex_model((8:1:t_diet) ,ic_chow8d, pchow8d); 
%2w hfd   
hfd2w=func_mex_model((0:1:t_diet),ic_hfd, phfd );   
%%% Fasting + IPGTT simulations 
ss_chow2w=func_mex_model( t_fasting, chow2w.statevalues(end,:), pchow);  % chow
IPGTT_chow2w=func_mex_model( t_IPGTT , ss_chow2w.statevalues(end,:), pchow ); 
ss_chow12dHFD2d=func_mex_model( t_fasting, chow12dHFD2d.statevalues(end,:), pchow12d);  % %12d chow + 2d HFD  
IPGTT_chow12dHFD2d=func_mex_model( t_IPGTT , ss_chow12dHFD2d.statevalues(end,:), pchow12d ); 
ss_chow8dHFD6d=func_mex_model( t_fasting, chow8dHFD6d.statevalues(end,:), pchow8d);  % %8d chow + 6d HFD  
IPGTT_chow8dHFD6d=func_mex_model( t_IPGTT , ss_chow8dHFD6d.statevalues(end,:), pchow8d ); 
ss_hfd2w=func_mex_model( t_fasting, hfd2w.statevalues(end,:), phfd);  % hfd
IPGTT_hfd2w=func_mex_model( t_IPGTT , ss_hfd2w.statevalues(end,:), phfd );
%%

% Unpublished diet data - 8w chow/hfd -  Stenkula et al 
t_diet_6w=56; % 2 weeks % diet time in days ( only training to the first two weeks)
% chow
pchow8w=p; 
pchow8w(1)= 10.^theta(6); % EI intake for 8w chow
pchow8w(2)=0.7; 
pchow8w(16)=  10.^theta(3); % PA for this 8w chow    
pchow8w(id_FFMname)=D.BW_unpublished_chow.Mean(1)*0.89;
pchow8w(id_FMname)= D.BW_unpublished_chow.Mean(1)*0.11;
ic_chow8w=ic0;
ic_chow8w( ismember(SBstates(objModel),'FFM')) = D.BW_unpublished_chow.Mean(1)*0.89;
ic_chow8w( ismember(SBstates(objModel),'FM')) = D.BW_unpublished_chow.Mean(1)*0.11;
% hfd
phfd8w=p;
phfd8w(1)=10.^theta(7); % EI for 8w HFD
phfd8w(2)=0.2;  
phfd8w(16)=  10.^theta(4); % setting up hfd specifc parameters
phfd8w(id_FFMname)=D.BW_unpublished_hfd.Mean(1)*0.89;
phfd8w(id_FMname)=D.BW_unpublished_hfd.Mean(1)*0.11;
ic_hfd8w=ic0;
ic_hfd8w( ismember(SBstates(objModel),'FFM')) = D.BW_unpublished_hfd.Mean(1)*0.89;
ic_hfd8w( ismember(SBstates(objModel),'FM')) = D.BW_unpublished_hfd.Mean(1)*0.11;
% simulations
chow8w=func_mex_model( 0:1:t_diet_6w,ic_chow8w, pchow8w); % simulation 8 weeks chow diet
hfd8w=func_mex_model((0:1:t_diet_6w),ic_hfd8w, phfd8w );     % simulation 8 weeks hfd diet


t_fasting_up=[ t_diet_6w  t_diet_6w+0.5833 ]  ; % fasting at 6 weeks
t_IPGTT_up=(0:1:120)/(60*24)+t_fasting_up(end);

phfd8w(end) = t_fasting_up(end); % set time for IPGTT event after 6 weeks + fasting
hfd8w_fast=func_mex_model( t_fasting_up, hfd8w.statevalues(42,:), phfd8w);  % simulation  fasting before IPGTT at 6 weeks
IPGTT_hfd6w=func_mex_model( t_IPGTT_up , hfd8w_fast.statevalues(end,:), phfd8w ); % IPGTT 6 weeks
catch err 
    disp(err)
end

%%
AllSimulations.chow8w              = chow8w;
AllSimulations.hfd8w               = hfd8w;
AllSimulations.IPGTT_hfd6w         =IPGTT_hfd6w;
AllSimulations.chow2w               = chow2w;
AllSimulations.IPGTT_chow2w         = IPGTT_chow2w;
AllSimulations.chow12dHFD2d         = chow12dHFD2d;
AllSimulations.IPGTT_chow12dHFD2d   = IPGTT_chow12dHFD2d;
AllSimulations.chow10dHFD4d         = chow10dHFD4d;
AllSimulations.chow8dHFD6d          = chow8dHFD6d;
AllSimulations.IPGTT_chow8dHFD6d    = IPGTT_chow8dHFD6d;
AllSimulations.hfd2w                = hfd2w;
AllSimulations.IPGTT_hfd2w          = IPGTT_hfd2w;

end
```
