## Supplementary scripts for "A multi-scale *in silico* mouse model for insulin resistance and humanoid type 2 diabetes": Function_AllSimulations_t2d.m

```
function [AllSimulations] = Function_AllSimulations_t2d( theta ,func_mex_model, D, utility)
ic0 = utility.ic0;
p = utility.p0;
objModel = utility.objModel; % variables used for indexing

pNames = utility.pNames ; 
id_FFM = ismember(SBstates(objModel),'FFM') ; id_FM = ismember(SBstates(objModel),'FM');
id_FFMname = ismember(pNames,'FFMinit') ; id_FMname = ismember(pNames,'FMinit');
% parameter and time vector setup
p(17:37)=[10.^theta(8:19) theta(20)  10.^theta(21:end)];  % GSS to ik1
if p(ismember(pNames,'ik1')) == 10^theta(end) % bounds check
else
    disp('check parameter vector')
end
try   
    
    
%% Steady state simulation
ic_ss = ic0; p_ss = p; p_ss( ismember(pNames, 'ss_x')) = 0 ;
ic_ss(id_FFM)=1 ; ic_ss(id_FM)= 1; p_ss(id_FFMname)=1; p_ss(id_FMname)=1; % no effect of Weight on tissue-level and a i s- models
p_ss(end) = 1000; %no IPGTT
ss=func_mex_model(0:100:300,ic_ss, p_ss); % steady state simulation
cost_ss_gcc = (( ss.variablevalues(end, end-1)  -  D.IPGTTgcc.Mean(1,1)  )^2 )./( D.IPGTTgcc.SEM(1,1) )^2 ;% basal glucose
cost_ss_icc = (( ss.variablevalues(end, end)  -  D.IPGTTicc.Mean(1,1)  )^2 )./( D.IPGTTicc.SEM(1,1) )^2   ;% basal insulin
ss_cost = cost_ss_gcc + cost_ss_icc;
ic0 = ss.statevalues(end,:); % overwrite ic0
%% 4 weeks -chow -hfd - t2d hfd
t_diet=28; 
t_fasting=[ t_diet  t_diet+0.5833 ]  ;  %t_diet:0.0001:t_diet+0.5833;
t_IPGTT=(0:1:240)/(60*24)+t_fasting(end);
p(end) = t_fasting(end); % time of IPGTT (triggers after fasting)

%chow
pchow=p;
pchow(1)=10.^theta(5);   % EI   theta(5) - Chow EI
pchow(2)=0.7;            % carb fraction 
pchow(16)= 10.^theta(1); % PA for chow
pchow(id_FFMname)=0.89*D.BW.mean(1,1); pchow(id_FMname)=0.11*D.BW.mean(1,1);
ic_chow = ic0; ic_chow(id_FFM)=0.89*D.BW.mean(1,1) ; ic_chow(id_FM)= 0.11*D.BW.mean(1,1);
%hfd
phfd=p;     
phfd(1)= 16.5217;       
phfd(2)=0.2;      
phfd(16)      = 10.^theta(2);   
phfd(id_FFMname)=0.89*D.BW.mean(4,1);    
phfd(id_FMname)=0.11*D.BW.mean(4,1);
ic_hfd = ic0; ic_hfd(id_FFM)=0.89*D.BW.mean(4,1) ; ic_hfd(id_FM)= 0.11*D.BW.mean(4,1);
%Scenario 1 
psc1=p;     
psc1(1)= 16.5217;       
psc1(2)=0.2;      
psc1(16)      = 10.^theta(2);   
psc1(id_FFMname)=0.89*D.BW.mean(4,1);    psc1(id_FMname)=0.11*D.BW.mean(4,1);
psc1( ismember( pNames, 't2dswitch'))=1;
ic_sc1 = ic0; ic_sc1(id_FFM)=0.89*D.BW.mean(4,1) ; ic_sc1(id_FM)= 0.11*D.BW.mean(4,1);

chow4w=func_mex_model((0:0.1:t_diet),ic_chow, pchow );   
hfd4w=func_mex_model((0:0.1:t_diet),ic_hfd, phfd );   
hfd4wt2d=func_mex_model((0:0.1:t_diet),ic_sc1, psc1 ); 

ss_chow4w=func_mex_model( t_fasting, chow4w.statevalues(end,:), pchow);  % hf
IPGTT_chow4w=func_mex_model( t_IPGTT , ss_chow4w.statevalues(end,:), pchow ); 

ss_hfd4w=func_mex_model( t_fasting, hfd4w.statevalues(end,:), phfd);  % hf
IPGTT_hfd4w=func_mex_model( t_IPGTT , ss_hfd4w.statevalues(end,:), phfd ); 

ss_hfd4wt2d=func_mex_model( t_fasting, hfd4wt2d.statevalues(end,:), psc1);  % hf
IPGTT_hfd4wt2d=func_mex_model( t_IPGTT , ss_hfd4wt2d.statevalues(end,:), psc1 );      

catch err 
    disp(err)
end

%%
AllSimulations.chow4w             = chow4w;
AllSimulations.hfd4w              = hfd4w;
AllSimulations.hfd4wt2d           = hfd4wt2d;
AllSimulations.IPGTT_chow4w       = IPGTT_chow4w;
AllSimulations.IPGTT_hfd4w        = IPGTT_hfd4w;
AllSimulations.IPGTT_hfd4wt2d     = IPGTT_hfd4wt2d;


end
```
