## Supplementary scripts for "A multi-scale *in silico* mouse model for insulin resistance and humanoid type 2 diabetes": Plot_figures.m

```
%% Setup SBtoolbox
cd ('./SBPOPPACKAGE');
installSBPOPpackage
cd ..
%% Setup
addpath(genpath('./MLMM')); 
addpath('./MLMM/Models'); 
addpath('./MLMM/Data'); 
[D, Dpred] =   Setup_data; 

choice=input(strcat('What to plot ?',...
             '\n 1. Figure 3 and 4 ',...
             '\n 2. Figure 5 ',...
             '\n 3. Figure 6 and Figure 7',...
             '\n 4. Figure SA1',...
             '\n 5. Figure SA2',...
             '\n enter option (1-5):'));
          
if choice == 1 % Figure 3 and 4
modelName = 'MLMM_final'; 
cd('./MLMM/Models')
SBPDmakeMEXmodel(SBmodel(strcat(modelName,'.txt')))
cd ..
objModel = SBmodel(strcat(modelName,'.txt')); 
[pNames, p0] = SBparameters(objModel); ic0 = SBinitialconditions(objModel); func_mex_model = str2func(modelName);
utility.pNames = pNames;
utility.objModel = objModel;
utility.ic0=ic0;
utility.p0 = p0;

filedir = './Results/MLMM_final/Pred/minmax';
files=dir(fullfile(filedir, '*.mat'));
nfiles = length(files);
Figure_3_and_4
cd ..
elseif choice == 2 % Figure 5
modelName = 'MLMM_final'; 
cd('./MLMM/Models')
SBPDmakeMEXmodel(SBmodel(strcat(modelName,'.txt')))
cd ..
objModel = SBmodel(strcat(modelName,'.txt')); 
[pNames, p0] = SBparameters(objModel); ic0 = SBinitialconditions(objModel); func_mex_model = str2func(modelName);
utility.pNames = pNames;
utility.objModel = objModel;
utility.ic0=ic0;
utility.p0 = p0;

filedir = './Results/MLMM_final/Pred/minmax';
files=dir(fullfile(filedir, '*.mat'));
nfiles = length(files);
figure(1)        
Figure_5A    
cd ..
cd('./Bergqvist et al (2017)');
figure(2)
Plotscript_BQ2017
cd ..


elseif choice == 3 % Figure 6 and 7
modelName='MLMM_final_t2d';
cd('./MLMM/Models')
SBPDmakeMEXmodel(SBmodel(strcat(modelName,'.txt')))
cd ..

objModel = SBmodel(strcat(modelName,'.txt')); 
[pNames, p0] = SBparameters(objModel); ic0 = SBinitialconditions(objModel); func_mex_model = str2func(modelName);
utility.pNames = pNames;
utility.objModel = objModel;
utility.ic0=ic0;
utility.p0 = p0;

Figure_6_and_7AB
Figure_7C

cd ..

elseif choice == 4 
modelName = 'MLMM_final'; 
cd('./MLMM/Models')
SBPDmakeMEXmodel(SBmodel(strcat(modelName,'.txt')))
cd ..
objModel = SBmodel(strcat(modelName,'.txt')); 
[pNames, p0] = SBparameters(objModel); ic0 = SBinitialconditions(objModel); func_mex_model = str2func(modelName);
utility.pNames = pNames;
utility.objModel = objModel;
utility.ic0=ic0;
utility.p0 = p0;
filedir = './Results/MLMM_final/Pred/minmax';
files=dir(fullfile(filedir, '*.mat'));
nfiles = length(files);
Figure_SA1
cd ..
elseif choice == 5 % Figure 6
modelName='MLMM_final_t2d';
cd('./MLMM/Models')
SBPDmakeMEXmodel(SBmodel(strcat(modelName,'.txt')))
cd ..
objModel = SBmodel(strcat(modelName,'.txt')); 
[pNames, p0] = SBparameters(objModel); ic0 = SBinitialconditions(objModel); func_mex_model = str2func(modelName);
utility.pNames = pNames;
utility.objModel = objModel;
utility.ic0=ic0;
utility.p0 = p0;
Figure_SA2      
cd ..
end

clear all
disp('To plot another figure run the script again');
```
