## Supplementary scripts for "A multi-scale *in silico* mouse model for insulin resistance and humanoid type 2 diabetes": Readme.txt

To plot the figures please run the "Plot_figures" script. 

The script will first install the Systems Biology toolbox (located in the SBPOPPACKAGE folder) (http://www.sbtoolbox2.org/main.php). 

Secondly, a text prompt will appear in the command window asking what figure to be plotted.
Enter the number (1-5) for the selected figure. 

To plot another figure please run the "Plot_figures" script again.
