## Supplementary scripts for "A multi-scale *in silico* mouse model for insulin resistance and humanoid type 2 diabetes": installSBPOPpackage.m

```
function [] = installSBPOPpackage(varargin)
% installSBPOPpackage
% Installation function for the complete SBPOP package, including all
% additionally needed software. Note that this function needs to be run
% from the folder in which the function is located.
%
%       installSBPOPpackage
%
% This adds the complete SBPOP package and all subpackages and subdirectories 
% to the MATLAB path. If you are in a computer environment in which MATLAB
% is not able to store the path settings you can run this 
% installation function each time when starting MATLAB (ideally add the call
% to your startup.m script in the homefolder).

% Information:
% ============
% Copyright (c) 2012 Novartis Pharma AG
% 
% This program is Free Open Source Software: you can redistribute it and/or modify 
% it under the terms of the GNU General Public License as published by 
% the Free Software Foundation, either version 3 of the License, or 
% (at your option) any later version. 
% 
% This program is distributed in the hope that it will be useful, 
% but WITHOUT ANY WARRANTY; without even the implied warranty of 
% MERCHANTABILITY or FITNESS FOR A PARTICULAR PURPOSE. See the 
% GNU General Public License for more details. 
% 
% You should have received a copy of the GNU General Public License 
% along with this program. If not, see <http://www.gnu.org/licenses/>.

clc

%%%%%%%%%%%%%%%%%%%%%%%%%%%%%%%%%%%%%%%%%%%%%%%
% Check that installSBPOPpackage is started in the right folder
%%%%%%%%%%%%%%%%%%%%%%%%%%%%%%%%%%%%%%%%%%%%%%%
currentDir = pwd;
installSBPOPDir = fileparts(which('installSBPOPpackage.m'));
if ~strcmp(currentDir,installSBPOPDir),
    error('Run the ''installSBPOPpackage'' script from the folder where it is located.');
end

%%%%%%%%%%%%%%%%%%%%%%%%%%%%%%%%%%%%%%%%%%%%%%%
% Add folders to path
%%%%%%%%%%%%%%%%%%%%%%%%%%%%%%%%%%%%%%%%%%%%%%%
addpath(genpath(pwd));

%%%%%%%%%%%%%%%%%%%%%%%%%%%%%%%%%%%%%%%%%%%%%%%
% Add temp folder path
%%%%%%%%%%%%%%%%%%%%%%%%%%%%%%%%%%%%%%%%%%%%%%%
addpath(tempdirSB);

%%%%%%%%%%%%%%%%%%%%%%%%%%%%%%%%%%%%%%%%%%%%%%%
% Show all versions used
%%%%%%%%%%%%%%%%%%%%%%%%%%%%%%%%%%%%%%%%%%%%%%%
ver

%%%%%%%%%%%%%%%%%%%%%%%%%%%%%%%%%%%%%%%%%%%%%%%
% Final installation information, version, etc.
%%%%%%%%%%%%%%%%%%%%%%%%%%%%%%%%%%%%%%%%%%%%%%%
disp(' ');
disp(' ');
disp(sprintf('The SBPOP Package for MATLAB'));
disp(sprintf('Developed by Henning Schmidt,.'));
disp(sprintf('The SBPOP Package contains several third party packages, therefor'));
disp(sprintf('copyright statements are present in the individual functions and subpackages.'));
disp(sprintf(' '));
disp(sprintf('This program is Free Open Source Software: you can redistribute it and/or modify '));
disp(sprintf('it under the terms of the GNU General Public License as published by '));
disp(sprintf('the Free Software Foundation, either version 3 of the License, or '));
disp(sprintf('(at your option) any later version. '));
disp(sprintf(' '));
disp(sprintf('This program is distributed in the hope that it will be useful, '));
disp(sprintf('but WITHOUT ANY WARRANTY; without even the implied warranty of '));
disp(sprintf('MERCHANTABILITY or FITNESS FOR A PARTICULAR PURPOSE. See the '));
disp(sprintf('GNU General Public License for more details. '));
disp(sprintf(' '));
disp(sprintf('You should have received a copy of the GNU General Public License '));
disp(sprintf('along with this program. If not, see <http://www.gnu.org/licenses/>.'));

%%%%%%%%%%%%%%%%%%%%%%%%%%%%%%%%%%%%%%%%%%%%%%%
% Final installation information, version, etc.
%%%%%%%%%%%%%%%%%%%%%%%%%%%%%%%%%%%%%%%%%%%%%%%
disp(' ');
disp(' ');
disp(sprintf('The SBPOP PACKAGE was installed from path: ''%s''',pwd));

%%%%%%%%%%%%%%%%%%%%%%%%%%%%%%%%%%%%%%%%%%%%%%%
% Print out the TODO list
%%%%%%%%%%%%%%%%%%%%%%%%%%%%%%%%%%%%%%%%%%%%%%%
try
	help TODO
catch
end
```
