## Supplementary scripts for "A multi-scale *in silico* mouse model for insulin resistance and humanoid type 2 diabetes": installSBPOPpackageInitial.m

```
function [] = installSBPOPpackageInitial()
% installSBPOPpackageInitial
% Function to be run upon first installation of the SBPOP package.
% It will compile all necessary C, etc. files.
% In a server environment with a central installation of SBPOP package, this
% function will be run once by the systems administrator.
% Every user then has to run the installSBPOPpackage.m command to make 
% SBPOP available (see help text to installSBPOPpackage.m).
%
% Additionally, this adds the complete SBPOP package and all subpackages 
% and subdirectories to the MATLAB path. 
%
%       installSBPOPpackageInitial

% Information:
% ============
% Copyright � 2012 Novartis Pharma AG
% 
% This program is Free Open Source Software: you can redistribute it and/or modify 
% it under the terms of the GNU General Public License as published by 
% the Free Software Foundation, either version 3 of the License, or 
% (at your option) any later version. 
% 
% This program is distributed in the hope that it will be useful, 
% but WITHOUT ANY WARRANTY; without even the implied warranty of 
% MERCHANTABILITY or FITNESS FOR A PARTICULAR PURPOSE. See the 
% GNU General Public License for more details. 
% 
% You should have received a copy of the GNU General Public License 
% along with this program. If not, see <http://www.gnu.org/licenses/>.

clc

%%%%%%%%%%%%%%%%%%%%%%%%%%%%%%%%%%%%%%%%%%%%%%%
% Check that installSBPOPpackageInitial is started in the right folder
%%%%%%%%%%%%%%%%%%%%%%%%%%%%%%%%%%%%%%%%%%%%%%%
currentDir = pwd;
installSBPOPDir = fileparts(which('installSBPOPpackageInitial.m'));
if ~strcmp(currentDir,installSBPOPDir),
    error('Run the ''installSBPOPpackageInitial'' script from the folder where it is located.');
end

%%%%%%%%%%%%%%%%%%%%%%%%%%%%%%%%%%%%%%%%%%%%%%%
% Install sub packages
%%%%%%%%%%%%%%%%%%%%%%%%%%%%%%%%%%%%%%%%%%%%%%%
cd('SBTOOLBOX2');
installSB
cd('../SBPD');
installSBPD
cd('../SBPOP');
installSBPOP
cd('..');

%%%%%%%%%%%%%%%%%%%%%%%%%%%%%%%%%%%%%%%%%%%%%%%
% Add temp folder path
%%%%%%%%%%%%%%%%%%%%%%%%%%%%%%%%%%%%%%%%%%%%%%%
addpath(tempdirSB);

%%%%%%%%%%%%%%%%%%%%%%%%%%%%%%%%%%%%%%%%%%%%%%%
% Show all versions used
%%%%%%%%%%%%%%%%%%%%%%%%%%%%%%%%%%%%%%%%%%%%%%%
ver

%%%%%%%%%%%%%%%%%%%%%%%%%%%%%%%%%%%%%%%%%%%%%%%
% Final installation information, version, etc.
%%%%%%%%%%%%%%%%%%%%%%%%%%%%%%%%%%%%%%%%%%%%%%%
disp(' ');
disp(' ');
disp(sprintf('SBPOP PACKAGE installed from path: ''%s''',pwd));

%%%%%%%%%%%%%%%%%%%%%%%%%%%%%%%%%%%%%%%%%%%%%%%
% Print out the TODO list
%%%%%%%%%%%%%%%%%%%%%%%%%%%%%%%%%%%%%%%%%%%%%%%
try
	help TODO
catch
end
```
