## Supplementary scripts for "A multi-scale *in silico* mouse model for insulin resistance and humanoid type 2 diabetes": README.txt

Installation of the SBPOP Package:
==================================

- Start MATLAB
- The first time you install a new version of the SBPOP package, please run 
  the script: "installSBPOPpackageInitial". This will compile all needed 
  libraries and executables for your system.
- If you do not install it for the first time, then just execute the script
  "installSBPOPpackage". Alternatively you just need to add the SBPOP 
  Package folder and all subfolders to the MATLAB path.
- If your system is allowed to store the MATLAB path between MATLAB 
  sessions, you do not need to call these installation scripts again. If 
  your MATLAB path is not stored, then you will need to do that each time
  you start MATLAB and want to use the SBPOP Package.
