## Supplementary scripts for "A multi-scale *in silico* mouse model for insulin resistance and humanoid type 2 diabetes": convert2pdfSBPOP.m

```
function [] = convert2pdfSBPOP(filename)
% [DESCRIPTION]
% Old function but update to also work on windows by calling another
% function.
%
% [SYNTAX]
% [] = convert2pdfSBPOP(filename)
%
% [INPUT]
% filename:     filename to be used
% 
% [OUTPUT]
%
% [ASSUMPTIONS]
%
% [AUTHOR]
% Henning Schmidt,
%
% [DATE]
% 16th May 2010
%
% [PLATFORM]
% Windows XP Engine, MATLAB R2009a, MATLAB
%
% [KEYWORDS]
% MATLAB, SBPOP, median, averaging plot
% 
% [TOOLBOXES USED]
% Statistics Toolbox
%
% [VALIDATION HISTORY]
%
% [MODIFICATION HISTORY]

% Information:
% ============
% Copyright � 2012 Novartis Pharma AG
% 
% This program is Free Open Source Software: you can redistribute it and/or modify 
% it under the terms of the GNU General Public License as published by 
% the Free Software Foundation, either version 3 of the License, or 
% (at your option) any later version. 
% 
% This program is distributed in the hope that it will be useful, 
% but WITHOUT ANY WARRANTY; without even the implied warranty of 
% MERCHANTABILITY or FITNESS FOR A PARTICULAR PURPOSE. See the 
% GNU General Public License for more details. 
% 
% You should have received a copy of the GNU General Public License 
% along with this program. If not, see <http://www.gnu.org/licenses/>.

%%%%%%%%%%%%%%%%%%%%%%%%%%%%%%
% Convert to PDF and remove PS
%%%%%%%%%%%%%%%%%%%%%%%%%%%%%%
ps2pdf('psfile',[strrep(filename,'.ps','') '.ps'],'pdffile',[strrep(filename,'.pdf','') '.pdf'],'deletepsfile',1)

% %% Cold code
% if isunix && ~isempty(filename),
%     [path,file,ext] = fileparts(filename);    
%     oldPath = pwd;
%     if ~isempty(path), cd(path); end
%     system(sprintf('ps2pdfwr %s.ps %s.pdf',file,file));
%     warning off
%     delete([file '.ps']);
%     warning on
%     cd(oldPath);
% end
```
