## Supplementary scripts for "A multi-scale *in silico* mouse model for insulin resistance and humanoid type 2 diabetes": printFigureSBPOP.m

```
function [] = printFigureSBPOP(hfig, filename, varargin)
% [DESCRIPTION]
% Print figure "hfig" to file "filename". Supported formats: png, ps, jpg.
% PS is default. Several figures can be appended in the same file by using 
% format='ps'. Just run the function repeatedly with same filename.
% If format='png' or 'jpg', figures are not appended, but overwritten.
%
% Tips when using PS files:
%   1) If you want a PDF you need to generate it afterwards using the function: 
%             convert2pdfSBPOP (on Unix/Linux).
%   2) Dont use the PS format if you have transparency in your plots!
%   3) If you want to start a new file then remove the file first (by default 
%      plots are appended in PS mode). Files can be removed by: 
%             startNewPrintFigureSBPOP
%
% [SYNTAX]
% [] = printFigureSBPOP(hfig, filename)
% [] = printFigureSBPOP(hfig, filename, format)
%
% [INPUT]
% hfig:         handle of figure to be printed
% filename:     filename to be used 
% format:       'ps', 'png', 'jpg' (default: 'ps')
% 
% [OUTPUT]
%
% [ASSUMPTIONS]
%
% [AUTHOR]
% Henning Schmidt,
%
% [DATE]
% 16th May 2010
%
% [PLATFORM]
% Windows XP Engine, MATLAB R2009a, MATLAB
%
% [KEYWORDS]
% MATLAB, SBPOP, median, averaging plot
% 
% [TOOLBOXES USED]
% Statistics Toolbox
%
% [VALIDATION HISTORY]
%
% [MODIFICATION HISTORY]
 
% Information:
% ============
% Copyright � 2012 Novartis Pharma AG
% 
% This program is Free Open Source Software: you can redistribute it and/or modify 
% it under the terms of the GNU General Public License as published by 
% the Free Software Foundation, either version 3 of the License, or 
% (at your option) any later version. 
% 
% This program is distributed in the hope that it will be useful, 
% but WITHOUT ANY WARRANTY; without even the implied warranty of 
% MERCHANTABILITY or FITNESS FOR A PARTICULAR PURPOSE. See the 
% GNU General Public License for more details. 
% 
% You should have received a copy of the GNU General Public License 
% along with this program. If not, see <http://www.gnu.org/licenses/>.

imgformat   = 'psc2';
if nargin == 3,
    format = varargin{1};
    if strcmp(format,'ps'),
        imgformat   = 'psc2';
    elseif strcmp(format,'png'),
        imgformat   = 'png';        
    elseif strcmp(format,'jpg'),
        imgformat   = 'jpeg95';    
    elseif strcmp(format,'pdf'),
        imgformat   = 'pdf';    
    else
        error('Unknown image format');
    end
end
        
% Always PS-color
% Setting figure properties for printing
if strcmp(imgformat,'png'),
    set(hfig,'PaperOrientation','portrait');
else
    set(hfig,'PaperOrientation','landscape');
end    
set(hfig,'PaperType','A4');
set(hfig,'PaperPositionMode', 'manual');
set(hfig,'PaperUnits', 'centimeters');
set(hfig,'PaperPosition', [0 0 29.7 21]);
% Print
[path,file,ext] = fileparts(filename);
oldPath = pwd;
if ~isempty(path), cd(path); end
print(hfig,['-d',imgformat],file,'-append');
cd(oldPath);
```
