## Supplementary scripts for "A multi-scale *in silico* mouse model for insulin resistance and humanoid type 2 diabetes": ps2pdf.m

```
function ps2pdf(varargin)
%PS2PDF Function to convert a PostScript file to PDF using Ghostscript
%
%  Converts a postscript file into PDF. The resulting PDF file will contain
%  one page for each page defined in the postscript files, so a multi-page
%  postscript file, like those generated by using the '-append' option of 
%  MATLAB's print command, can be used to generate a multi-page PDF file. 
%
%   Ghostscript is a third-party application currently supplied with 
%   MATLAB. The caller may also specify a different version of Ghostscript
%   to use.
%
%   PS2PDF expects to be called with a set of parameter-value pairs. The
%   order of these is unimportant, but required parameters MUST be
%   specified
%
%   In the list below, required parameters are marked with an asterisk *
%        NOTE: ps2pdf can not use MATLAB's version of Ghostscript
%              in a deployed application; you MUST provide a 
%              the path to a separate instance of Ghostscript. This 
%              parameter is marked with a double asterisk **
%
%   Parameter:        Value:  
%   *  psfile         full or relative path to the postscript file to convert
%
%   *  pdffile        full or relative path to the pdf file to create
%
%   ** gscommand      path to Ghostscript executable to use; this will try
%                     to default to the version of Ghostscript shipped with 
%                     MATLAB, if any. If this value is specified you should 
%                     also specify the gsfontpath and gslibpath values.
%
%                     ** See note on deployed applications, above.
%
%      gsfontpath     full path to the Ghostscript font files
%                     If a gscommand is specified then this path should
%                     also be specified and reference the same Ghostscript
%                     version
%
%      gslibpath      full path to the Ghostscript library (.ps) files. 
%                     If a gscommand is specified then this path should
%                     also be specified and reference the same Ghostscript
%                     version
%
%                     If gscommand is NOT specified and we can determine
%                     the version of Ghostscript, if any, shipped with
%                     MATLAB, then this value will be overridden to use the
%                     path that references MATLAB's version of Ghostscript
%
%      gspapersize    paper size to use in the created .pdf file. If not 
%                     specified or the specified value is not recognized 
%                     it will use whatever default paper size is 
%                     built into the version of Ghostscript being run
%
%                         NOTE: no scaling of the input occurs - it's simply
%                         placed on a page with the specified paper size. 
%
%                         Valid values for gspapersize are: 
%                              'letter', 'ledger', 'legal', '11x17', 
%                              'archA', 'archB', 'archC', 'archD', 'archE', 
%                              'a0', 'a1', 'a2', 'a3','a4', 'a5',
%                              'a6', 'a7', 'a8', 'a9', 'a10'
%                         
%      deletepsfile   0 to keep the input ps file after creating pdf
%                     non-zero to delete the input ps file after creating pdf
%                     Default is 0: keep the input ps file (do NOT delete it)
%                        NOTE: if the pdf creation process fails, the input
%                        PS file will be kept regardless of this setting
%
%      verbose        0 to suppress display of status/progress info; 
%                     non-zero to allow display of status/progress info
%                     Default is 0 (no display)
%
% Example usage: 
%    use MATLAB's version of Ghostscript to generate an A4 pdf file
%      ps2pdf('psfile', 'input.ps', 'pdffile', 'output.pdf', 'gspapersize', 'a4')
%
%    use a local copy of Ghostcript to generate a file, and display some 
%    status/progress info while doing so.
%      ps2pdf('psfile', '../reports/input.ps', 'pdffile', 'c:\temp\output3.pdf', ...
%            'gspapersize', 'a4', 'verbose', 1, ...
%            'gscommand', 'C:\Program Files\GhostScript\bin\gswin32c.exe', ...
%            'gsfontpath', 'C:\Program Files\GhostScript\fonts', ...
%            'gslibpath', 'C:\Program Files\GhostScript\lib')
%
%    use MATLAB's version of Ghostscript to generate a pdf file and delete
%    the input.ps file when done 
%      ps2pdf('psfile', 'input.ps', 'pdffile', 'output.pdf', 'gspapersize', 'a4', 'deletepsfile', 1)

%   Update log: 
%      Jun 16, 2010: added check for deployed application
%      May 19, 2010: wrapped filenames sent to Ghostscript in quotes
%      May 06, 2010: updated how Ghostscript is found, don't rely on MATLAB version #
%      Aug 15, 2008: fixed bug where embedded space in location of 
%                    MATLAB's version of Ghostscript caused ps2pdf to fail.
%      Apr 16, 2008: added deletepsfile option

%   Copyright 2008-2010 The MathWorks, Inc.

   if nargin < 1 
      error('ps2pdf:parameters', 'No parameters specified. Type ''help ps2pdf'' for details on how to use this function.');
   end

   % parse input args
   gsData = LocalParseArgs(varargin{:});

   % setup the file that tells GS what we want it to do
   gsData = LocalCreateResponseFile(gsData); 

   gsDebug = 0;
   if gsData.verbose 
      fprintf('ps2pdf: input settings are:\n');
      if isfield(gsData, 'paperSizes')
          gsData = rmfield(gsData, 'paperSizes');
      end
      gsData  %#ok<NOPRT>
      fprintf('ps2pdf: response file for Ghostscript is:\n');
      type(gsData.responseFile);
      gsDebug = 1;
   end

   %to hold results/status from system call
   s = 0; %#ok<NASGU>
   r = ''; %#ok<NASGU>

   % run Ghostscript to convert the file
   if gsData.useBuiltin 
      [s, r] = gsData.cmd(['@' gsData.responseFile], gsData.psFile, gsDebug);
   else 
      [s, r] = system([gsData.cmd ' @"' gsData.responseFile '" "' gsData.psFile '"']);
   end

   if gsData.verbose
      disp( ['Ghostscript STDOUT: ' num2str(s) ] );
      disp( ['Ghostscript STDERR: ' r ] );
   else
      delete(gsData.responseFile)
   end

   if s && ~isempty(r)
      error('ps2pdf:ghostscript',  ['Problem converting PostScript. System returned error: ' num2str(s) '.' r]) 
   elseif s
      error('ps2pdf:ghostscript',  ['Problem calling GhostScript. System returned error: ' num2str(s)]) 
   end

   %if after all this we still couldn't create the file, report the error
   fid = fopen( gsData.pdfFile, 'r');
   if ( fid == -1 )
      error('ps2pdf:ghostscript', '%s', [ 'Ghostscript could not create ''' gsData.pfdFile '''.' ])
   else
      fclose( fid );
   end

   % if we get here, we successfully created pdf file; delete ps file if
   % requested to do so
   if gsData.deletePSFile 
       delete(gsData.psFile);
   end

end

%local function to parse arguments and fill in the gsData structure
%  .psFile - postscript file to convert
%  .pdfFile - pdf file to create
%  .cmd     - path/name of Ghostscript command to run or handle to gscript
%             builtin
%  .useBuiltin - 1 if using builtin gs command 
%  .fontPath - path to the Ghostscript fonts, if any
%  .libPath - path to the Ghostscript libs (Ghostscript .ps files), if any) 
%  .paperSize - paper size to set for resulting .pdf file 
%  .deletePSFile - 0 to keep (not delete) the input ps file if pdf created ok
%  .verbose - if non-zero, display some status/progress info to command window
function gsData = LocalParseArgs(varargin) 
    gsData.paperSizes = {'letter', 'ledger', 'legal', '11x17', 'archA', 'archB', ... 
                   'archC', 'archD', 'archE', 'a0', 'a1', 'a2', 'a3','a4', 'a5', ...
                   'a6', 'a7', 'a8', 'a9', 'a10'};

    %default values for some settings
    gsData.verbose      = 0; 
    gsData.useBuiltin   = 0;
    gsData.deletePSFile = 0; 
    
    for i = 1 : 2 : length(varargin)-1 
        param_arg = varargin{i};
        param_value = varargin{i+1};        
        switch(lower(param_arg)) 
            % path to ps file to conver
            case 'psfile'
               if ~exist(param_value, 'file')
                  error('print:ghostscript', ...
                      'Can not find postscript file <%s> to convert', ...
                      param_value)
               end
               gsData.psFile = param_value;

            % path to pdf file to create
            case 'pdffile'
                %verify we can create file at that location
                pdf_fid = fopen(param_value,'w');
                if pdf_fid < 0 
                    error('ps2pdf:invalidPDFFIle', ... 
                        'Can not open <%s> for writing', ...
                        param_value);
                end
                fclose(pdf_fid); 
                %delete temp file we created
                delete(param_value); 
                gsData.pdfFile = param_value;

            % full path to gs executable
            case 'gscommand' 
               if ~exist(param_value, 'file')
                  error('ps2pdf:ghostscriptCommand', ...
                      'Can not find Ghostscript executable (''gscommand'') <%s>',...
                      param_value)
               end
               if ispc && ~isempty(findstr(param_value, ' '))
                   param_value = ['"' param_value '"']; %#ok<AGROW>
               end
               gsData.cmd = param_value; 
                
            % full path to gs font dir
            case 'gsfontpath' 
               if ~exist(param_value, 'dir')
                   error('ps2pdf:ghostscriptFontPath', ...
                         'Can not find the directory <%s> for Ghostscript fonts (''gsfontpath'')', ...
                         param_value)
               end
               gsData.fontPath = param_value;

            % full path to gs lib dir
            case 'gslibpath' 
               if ~exist(param_value, 'dir')
                   error('ps2pdf:ghostscriptLibPath', ...
                         'Can not find the directory <%s> for Ghostscript library files (''gslibpath'')', ...
                         param_value)
               end
               gsData.libPath = param_value;
                
            % paper size 
            case 'gspapersize'
               idx = strcmpi(param_value, gsData.paperSizes);
               if ~any(idx)
                  warning('ps2pdf:papersize', ...
                        '''gspapersize'' value <%s> not found in the list of known sizes, ignoring it.', param_value);
               else
                  gsData.paperSize = gsData.paperSizes{idx};
               end

            % deletePSFile
            case 'deletepsfile'
               if isnumeric(param_value) 
                  gsData.deletePSFile = param_value; 
               else
                   warning('ps2pdf:deletepsfile', ...
                         '''deletepsfile'' value <%s> class <%s> should be numeric, defaulting to 0', ...
                         param_value, class(param_value));
               end
               
            % verbose
            case 'verbose'
               if isnumeric(param_value) 
                  gsData.verbose = param_value; 
               else
                   warning('ps2pdf:verbose', ...
                         '''verbose'' value <%s> class <%s> should be numeric, defaulting to 0', ...
                         param_value, class(param_value));
               end
               
            otherwise
               if isnumeric(param_value)
                   param_value = num2str(param_value);
               end
                  warning('ps2pdf:unknown', ...
                     'ignoring unknown parameter <%s> with value <%s>.', param_arg, param_value);
        end
    end    

    if ~isfield(gsData, 'psFile') 
        error('ps2pdf:noInputFile', ...
               'No input (psfile) file specified');
    end
    
    if ~isfield(gsData, 'pdfFile') 
        error('ps2pdf:noOutputFile', ...
               'No output (pdffile) file specified');
    end
    
    if ~isfield(gsData, 'cmd') 
        if isdeployed
            error('ps2pdf:deployedNeedsGhostscript', ...
                  'In order to use ''ps2pdf'' in a deployed application you must provide the path to a separate instance of Ghostscript.');
        end

        % updated code to find ghostscript - look for gs8x first, 
        % then try old location. Don't depend on MATLAB version #
        ghostDir = fullfile( matlabroot, 'sys', 'gs8x' );
        if ~exist(ghostDir, 'dir')
            [gsCmd, ghostDir] = Local_GetOldGhostscript();
            gsData.cmd = gsCmd;
        else
           gsData.cmd = Local_GetGscriptFcnHandle;
           if ~isempty(gsData.cmd)
              gsData.useBuiltin = 1; % use builtin Ghostscript
           end
        end
        if ~exist(ghostDir, 'dir')
           error('ps2pdf:ghostscriptCommand', ...
                 'Can not find Ghostscript installed with MATLAB in <%s>',...
                 ghostDir);
        end

        if ~isempty(gsData.cmd)
           % if using MATLAB's version of GhostScript, use same set of fonts and library files
           if isfield(gsData, 'fontPath') || isfield(gsData, 'libPath')
              warning('ps2pdf:ghostscriptPathOverride', ...
                    'Using MATLAB''s version of Ghostscript; overriding ''gsfontpath'' and ''gslibpath'' to use builtin MATLAB version');
           end
           gsData.fontPath = fullfile( ghostDir, 'fonts', '');
           gsData.libPath = fullfile( ghostDir, 'ps_files', '');
        else 
            error('ps2pdf:noGhostscriptCommand', ...
                  'Can not find Ghostscript program in MATLAB');
        end
    else
        % if gscommandpath was specified, 
        if ~isfield(gsData, 'fontPath') || ~isfield(gsData, 'libPath')
           warning('ps2pdf:ghostscriptCommandSuggestion', ...
                 ['When specifying a Ghostscript executable (''gscommand'') you should also '...
                 'specify both the ''gsfontpath'' and ''gslibpath'' locations']);
   
        end
    end
end

%local function to create the input file needed for Ghostscript
function gsData = LocalCreateResponseFile(gsData) 
   % open a response file to write out Ghostscript commands
   rsp_file = [tempname '.rsp'];
   rsp_fid = fopen (rsp_file, 'w');

   if (rsp_fid < 0)
      error('ps2pdf:responseFileCreate', 'Unable to create response file')
   end

   fprintf(rsp_fid, '-dBATCH -dNOPAUSE\n');
   if ~gsData.verbose
       fprintf(rsp_fid, '-q\n');
   end
   if isfield(gsData, 'libPath')
      fprintf(rsp_fid, '-I"%s"\n', gsData.libPath);
   end
   if isfield(gsData, 'fontPath')
      fprintf(rsp_fid, '-I"%s"\n', gsData.fontPath);
   end
   if isfield(gsData, 'paperSize') 
      fprintf( rsp_fid, '-sPAPERSIZE=%s\n', gsData.paperSize );
   end
   fprintf(rsp_fid, '-sOutputFile="%s"\n', gsData.pdfFile);
   fprintf(rsp_fid, '-sDEVICE=%s\n', 'pdfwrite');
   fclose(rsp_fid);
   gsData.responseFile = rsp_file;
end

%local function to get a handle to MATLAB's Ghostscript implementation
%NOTE: this may change or be removed in future releases
function gs = Local_GetGscriptFcnHandle()
  gs = '';
  p = which('-all', 'gscript');
  if ~isempty(p) 
      p = p{1};
      fpath = fileparts(p);
      olddir = cd(fpath);
      gs = @gscript;
      cd(olddir);
  end
end

% local function to try and get location of Ghostscript in older MATLAB
function [gsCmd, ghostDir] = Local_GetOldGhostscript
   ghostDir = fullfile( matlabroot, 'sys', 'ghostscript' );
   gsCmd = '';
   if ispc
      if exist(fullfile(ghostDir,'bin','win32','gs.exe'), 'file')
         gsCmd = fullfile(ghostDir,'bin','win32','gs.exe');
      end
   else 
      if exist(fullfile(ghostDir,'bin',lower(computer),'gs'), 'file')
         gsCmd = fullfile(ghostDir,'bin',lower(computer), 'gs');
      end
   end
   gsCmd = ['"' gsCmd '"'];
end
```
