## Supplementary scripts for "A multi-scale *in silico* mouse model for insulin resistance and humanoid type 2 diabetes": startNewPrintFigureSBPOP.m

```
function [] = startNewPrintFigureSBPOP(filename)
% [DESCRIPTION]
% Function starts a new file in which figures are to be printed.
% Basically this means, that it is checked if the file exists and if yes it is deleted.
% The function "printFigureSBPOP" can then be used to print a figure into this file.
% The function "convert2pdfSBPOP" can then be used to convert from PS to PDF (only unix).
% On Unix check is done for .PDF, on windows for .PS
%
% [SYNTAX]
% [] = startNewPrintFigureSBPOP(filename)
%
% [INPUT]
% filename:     filename to be used
% 
% [OUTPUT]
%
% [ASSUMPTIONS]
%
% [AUTHOR]
% Henning Schmidt,
%
% [DATE]
% 16th May 2010
%
% [PLATFORM]
% Windows XP Engine, MATLAB R2009a, MATLAB
%
% [KEYWORDS]
% MATLAB, SBPOP, median, averaging plot
% 
% [TOOLBOXES USED]
% Statistics Toolbox
%
% [VALIDATION HISTORY]
%
% [MODIFICATION HISTORY]
 
% Information:
% ============
% Copyright � 2012 Novartis Pharma AG
% 
% This program is Free Open Source Software: you can redistribute it and/or modify 
% it under the terms of the GNU General Public License as published by 
% the Free Software Foundation, either version 3 of the License, or 
% (at your option) any later version. 
% 
% This program is distributed in the hope that it will be useful, 
% but WITHOUT ANY WARRANTY; without even the implied warranty of 
% MERCHANTABILITY or FITNESS FOR A PARTICULAR PURPOSE. See the 
% GNU General Public License for more details. 
% 
% You should have received a copy of the GNU General Public License 
% along with this program. If not, see <http://www.gnu.org/licenses/>.

%%%%%%%%%%%%%%%%%%%%%%%%%%%%%%
% Create folder if it is not existing
%%%%%%%%%%%%%%%%%%%%%%%%%%%%%%
folder = fileparts(filename);
if ~isempty(folder),
    warning off
    mkdir(folder);
    warning on
end

%%%%%%%%%%%%%%%%%%%%%%%%%%%%%%
% If filename exists then delete it
%%%%%%%%%%%%%%%%%%%%%%%%%%%%%%
fclose all;
if ~isempty(filename),
    [path,file,ext] = fileparts(filename);
    filename_pdf = fullfile(path,[file '.pdf']);
    filename_ps  = fullfile(path,[file '.ps']);

    warning off;
    try
        delete(filename_pdf);
    catch
    end
    
    try
        delete(filename_ps);
    catch
    end
    warning on;
    
end
```
