## Supplementary scripts for "A multi-scale *in silico* mouse model for insulin resistance and humanoid type 2 diabetes": getcolorsSBPOP.m

```
function [colors,lines,dots,bwcolors] = getcolorsSBPOP()
% [DESCRIPTION]
% Help function to generate colors for plotting and for black and white
% plotting also line types.
%
% [SYNTAX]
% [colors,lines,dots,bwcolors] = getcolorsSBPOP()
%
% [INPUT]
% NONE
%
% [OUTPUT]
% colors: Nx3 matrix with colors to be used (see below in detail)
% lines:  cell-array with 48 unique line-styles
% dots:   cell-array with 12 unique dot-styles
%
% [ASSUMPTIONS]
%
% [AUTHOR]
% Henning Schmidt,
%
% [DATE]
% 15th April 2010
%
% [PLATFORM]
% Windows XP Engine, MATLAB R2009a, MATLAB
%
% [KEYWORDS]
% MATLAB, SBPOP, DV, PRED, IPRED, diagnostic, plot, DV vs PRED, DV vs IPRED
% 
% [TOOLBOXES USED]
% Statistics Toolbox
%
% [VALIDATION HISTORY]
%
% [MODIFICATION HISTORY]

% Information:
% ============
% Copyright � 2012 Novartis Pharma AG
% 
% This program is Free Open Source Software: you can redistribute it and/or modify 
% it under the terms of the GNU General Public License as published by 
% the Free Software Foundation, either version 3 of the License, or 
% (at your option) any later version. 
% 
% This program is distributed in the hope that it will be useful, 
% but WITHOUT ANY WARRANTY; without even the implied warranty of 
% MERCHANTABILITY or FITNESS FOR A PARTICULAR PURPOSE. See the 
% GNU General Public License for more details. 
% 
% You should have received a copy of the GNU General Public License 
% along with this program. If not, see <http://www.gnu.org/licenses/>.

colors = [0.87 0.49 0
           0.17 0.51 0.34
		   0.08 0.16 0.55
		   0.48 0.06 0.89
		   0.85 0.16 0
		   0.68 0.47 0
		   0.04 0.52 0.78
		   0.25 0.25 0.25 
		   0.6 0.2 0
		   0.75 0.75 0
		   0 0.75 0.75
		   0 0.5 0
		   0 0 1
		   0.75 0 0.75
		   1 0 0
		   0.04 0.14 0.42
		   0.31 0.31 0.31
		   0.5 0.5 0.5
		   0 0 0
		   1 0.69 0.39
		   0 1 0
		   0 1 1
		   1 0 1
		   0.7 0.78 1
		   1 1 0
		   0.68 0.92 1
		   0.85 0.7 1
		   1 0.6 0.78];
      
lines = {'o-','x-','+-','*-','s-','d-','v-','^-','<-','>-','p-','h-',       'o--','x--','+--','*--','s--','d--','v--','^--','<--','>--','p--','h--',     'o-.','x-.','+-.','*-.','s-.','d-.','v-.','^-.','<-.','>-.','p-.','h-.',   'o:','x:','+:','*:','s:','d:','v:','^:','<:','>:','p:','h:'   };
      
dots = {'o','x','+','*','s','d','v','^','<','>','p','h'};      

bwcolors = [0 0 0; 0.33 0.33 0.33; 0.66 0.66 0.66];
      
```
