## Supplementary scripts for "A multi-scale *in silico* mouse model for insulin resistance and humanoid type 2 diabetes": parseArgs.m

```
function ArgStruct=parseArgs(args,ArgStruct,varargin)
% Helper function for parsing varargin. 
%
%
% ArgStruct=parseArgs(varargin,ArgStruct[,FlagtypeParams[,Aliases]])
%
% * ArgStruct is the structure full of named arguments with default values.
% * Flagtype params is params that don't require a value. (the value will be set to 1 if it is present)
% * Aliases can be used to map one argument-name to several argstruct fields
%
%
% example usage: 
% --------------
% function parseargtest(varargin)
%
% %define the acceptable named arguments and assign default values
% Args=struct('Holdaxis',0, ...
%        'SpacingVertical',0.05,'SpacingHorizontal',0.05, ...
%        'PaddingLeft',0,'PaddingRight',0,'PaddingTop',0,'PaddingBottom',0, ...
%        'MarginLeft',.1,'MarginRight',.1,'MarginTop',.1,'MarginBottom',.1, ...
%        'rows',[],'cols',[]); 
%
% %The capital letters define abrreviations.  
% %  Eg. parseargtest('spacingvertical',0) is equivalent to  parseargtest('sv',0) 
%
% Args=parseArgs(varargin,Args, ... % fill the arg-struct with values entered by the user
%           {'Holdaxis'}, ... %this argument has no value (flag-type)
%           {'Spacing' {'sh','sv'}; 'Padding' {'pl','pr','pt','pb'}; 'Margin' {'ml','mr','mt','mb'}});
%
% disp(Args)
%
%
%
%
% % Aslak Grinsted 2003

Aliases={};
FlagTypeParams='';

if (length(varargin)>0) 
    FlagTypeParams=strvcat(varargin{1});
    if length(varargin)>1
        Aliases=varargin{2};
    end
end
 

%---------------Get "numeric" arguments
NumArgCount=1;
while (NumArgCount<=size(args,2))&(~ischar(args{NumArgCount}))
    NumArgCount=NumArgCount+1;
end
NumArgCount=NumArgCount-1;
if (NumArgCount>0)
    ArgStruct.NumericArguments={args{1:NumArgCount}};
else
    ArgStruct.NumericArguments={};
end 


%--------------Make an accepted fieldname matrix (case insensitive)
Fnames=fieldnames(ArgStruct);
for i=1:length(Fnames)
    name=lower(Fnames{i,1});
    Fnames{i,2}=name; %col2=lower
    AbbrevIdx=find(Fnames{i,1}~=name);
    Fnames{i,3}=[name(AbbrevIdx) ' ']; %col3=abreviation letters (those that are uppercase in the ArgStruct) e.g. SpacingHoriz->sh
    %the space prevents strvcat from removing empty lines
    Fnames{i,4}=isempty(strmatchSB(Fnames{i,2},FlagTypeParams)); %Does this parameter have a value? (e.g. not flagtype)
end
FnamesFull=strvcat(Fnames{:,2});
FnamesAbbr=strvcat(Fnames{:,3});

if length(Aliases)>0  
    for i=1:length(Aliases)
        name=lower(Aliases{i,1});
        FieldIdx=strmatchSB(name,FnamesAbbr,'exact'); %try abbreviations (must be exact)
        if isempty(FieldIdx) 
            FieldIdx=strmatchSB(name,FnamesFull); %&??????? exact or not? 
        end
        Aliases{i,2}=FieldIdx;
        AbbrevIdx=find(Aliases{i,1}~=name);
        Aliases{i,3}=[name(AbbrevIdx) ' ']; %the space prevents strvcat from removing empty lines
        Aliases{i,1}=name; %dont need the name in uppercase anymore for aliases
    end
    %Append aliases to the end of FnamesFull and FnamesAbbr
    FnamesFull=strvcat(FnamesFull,strvcat(Aliases{:,1})); 
    FnamesAbbr=strvcat(FnamesAbbr,strvcat(Aliases{:,3}));
end

%--------------get parameters--------------------
l=NumArgCount+1; 
while (l<=length(args))
    a=args{l};
    if ischar(a)
        paramHasValue=1; % assume that the parameter has is of type 'param',value
        a=lower(a);
        FieldIdx=strmatchSB(a,FnamesAbbr,'exact'); %try abbreviations (must be exact)
        if isempty(FieldIdx) 
            FieldIdx=strmatchSB(a,FnamesFull); 
        end
        if (length(FieldIdx)>1) %shortest fieldname should win 
            [mx,mxi]=max(sum(FnamesFull(FieldIdx,:)==' ',2));
            FieldIdx=FieldIdx(mxi);
        end
        if FieldIdx>length(Fnames) %then it's an alias type.
            FieldIdx=Aliases{FieldIdx-length(Fnames),2}; 
        end
        
        if isempty(FieldIdx) 
            error(['Unknown named parameter: ' a])
        end
        for curField=FieldIdx' %if it is an alias it could be more than one.
            if (Fnames{curField,4})
                val=args{l+1};
            else
                val=1; %parameter is of flag type and is set (1=true)....
            end
            ArgStruct.(Fnames{curField,1})=val;
        end
        l=l+1+Fnames{FieldIdx(1),4}; %if a wildcard matches more than one
    else
        error(['Expected a named parameter: ' num2str(a)])
    end
end
```
