## Supplementary scripts for "A multi-scale *in silico* mouse model for insulin resistance and humanoid type 2 diabetes": subaxis.m

```
function h=subaxis(varargin)
%SUBAXIS Create axes in tiled positions. (just like subplot)
%   Usage:
%      h=subaxis(rows,cols,cellno[,settings])
%      h=subaxis(rows,cols,cellx,celly[,settings])
%      h=subaxis(rows,cols,cellx,celly,spanx,spany[,settings])
%
% SETTINGS: Spacing,SpacingHoriz,SpacingVert
%           Padding,PaddingRight,PaddingLeft,PaddingTop,PaddingBottom
%           Margin,MarginRight,MarginLeft,MarginTop,MarginBottom
%           Holdaxis
%
%           all units are relative (e.g from 0 to 1)
%
%           Abbreviations of parameters can be used.. (Eg MR instead of MarginRight)
%           (holdaxis means that it wont delete any axes below.)
%
%
% Example:
%
%   >> subaxis(2,1,1,'SpacingVert',0,'MR',0); 
%   >> imagesc(magic(3))
%   >> subaxis(2,'p',.02);
%   >> imagesc(magic(4))
%
% 2001 / Aslak Grinsted  (Feel free to modify this code.)
f=gcf;


Args=[];
UserDataArgsOK=0;
Args=get(f,'UserData');
if isstruct(Args) 
    UserDataArgsOK=isfield(Args,'SpacingHorizontal')&isfield(Args,'Holdaxis')&isfield(Args,'rows')&isfield(Args,'cols');
end
OKToStoreArgs=isempty(Args)|UserDataArgsOK;

if isempty(Args)&(~UserDataArgsOK)
    Args=struct('Holdaxis',0, ...
        'SpacingVertical',0.05,'SpacingHorizontal',0.05, ...
        'PaddingLeft',0,'PaddingRight',0,'PaddingTop',0,'PaddingBottom',0, ...
        'MarginLeft',.1,'MarginRight',.1,'MarginTop',.1,'MarginBottom',.1, ...
        'rows',[],'cols',[]); 
end
Args=parseArgs(varargin,Args,{'Holdaxis'},{'Spacing' {'sh','sv'}; 'Padding' {'pl','pr','pt','pb'}; 'Margin' {'ml','mr','mt','mb'}});

if (length(Args.NumericArguments)>1)
    Args.rows=Args.NumericArguments{1};
    Args.cols=Args.NumericArguments{2};
%remove these 2 numerical arguments
    Args.NumericArguments={Args.NumericArguments{3:end}};
end

if OKToStoreArgs
    set(f,'UserData',Args);
end


    

switch length(Args.NumericArguments)
   case 0
       return % no arguments but rows/cols.... 
   case 1
      x1=mod((Args.NumericArguments{1}-1),Args.cols)+1; x2=x1;
      y1=floor((Args.NumericArguments{1}-1)/Args.cols)+1; y2=y1;
   case 2
      x1=Args.NumericArguments{1};x2=x1;
      y1=Args.NumericArguments{2};y2=y1;
   case 4
      x1=Args.NumericArguments{1};x2=x1+Args.NumericArguments{3}-1;
      y1=Args.NumericArguments{2};y2=y1+Args.NumericArguments{4}-1;
   otherwise
      error('subaxis argument error')
end
    

cellwidth=((1-Args.MarginLeft-Args.MarginRight)-(Args.cols-1)*Args.SpacingHorizontal)/Args.cols;
cellheight=((1-Args.MarginTop-Args.MarginBottom)-(Args.rows-1)*Args.SpacingVertical)/Args.rows;
xpos1=Args.MarginLeft+Args.PaddingLeft+cellwidth*(x1-1)+Args.SpacingHorizontal*(x1-1);
xpos2=Args.MarginLeft-Args.PaddingRight+cellwidth*x2+Args.SpacingHorizontal*(x2-1);
ypos1=Args.MarginTop+Args.PaddingTop+cellheight*(y1-1)+Args.SpacingVertical*(y1-1);
ypos2=Args.MarginTop-Args.PaddingBottom+cellheight*y2+Args.SpacingVertical*(y2-1);

if Args.Holdaxis
    h=axes('position',[xpos1 1-ypos2 xpos2-xpos1 ypos2-ypos1]);
else
    h=subplot('position',[xpos1 1-ypos2 xpos2-xpos1 ypos2-ypos1]);
end


set(h,'box','on');
%h=axes('position',[x1 1-y2 x2-x1 y2-y1]);
set(h,'units',get(gcf,'defaultaxesunits'));
set(h,'tag','subaxis');


if (nargout==0) clear h; end;
```
