## Supplementary scripts for "A multi-scale *in silico* mouse model for insulin resistance and humanoid type 2 diabetes": cellmaxlengthSBPOP.m

```
function [maxlength] = cellmaxlengthSBPOP(input)
% cellmaxlengthSBPOP: for a cell-array with only string entries the function
% determines the maxlength of these strings.

% Information:
% ============
% Copyright � 2012 Novartis Pharma AG
% 
% This program is Free Open Source Software: you can redistribute it and/or modify 
% it under the terms of the GNU General Public License as published by 
% the Free Software Foundation, either version 3 of the License, or 
% (at your option) any later version. 
% 
% This program is distributed in the hope that it will be useful, 
% but WITHOUT ANY WARRANTY; without even the implied warranty of 
% MERCHANTABILITY or FITNESS FOR A PARTICULAR PURPOSE. See the 
% GNU General Public License for more details. 
% 
% You should have received a copy of the GNU General Public License 
% along with this program. If not, see <http://www.gnu.org/licenses/>.

if ~iscell(input),
    error('Input is not a cell-array.');
end

maxlength = 0;
for k=1:length(input),
    if ~ischar(input{k}),
        error('Elements of cell-array need to be strings.');
    end
    if length(input{k}) > maxlength,
        maxlength = length(input{k});
    end
end
```
