## Supplementary scripts for "A multi-scale *in silico* mouse model for insulin resistance and humanoid type 2 diabetes": lookforSBPOP.m

```
function [] = lookforSBPOP(text)
% lookforSBPOP: searches all m-files in the SBPOP folder tree for the 
% text given as argument. It displays the filenames in which the text
% appears and additionally opens the files in the editor.

% Information:
% ============
% Copyright � 2012 Novartis Pharma AG
% 
% This program is Free Open Source Software: you can redistribute it and/or modify 
% it under the terms of the GNU General Public License as published by 
% the Free Software Foundation, either version 3 of the License, or 
% (at your option) any later version. 
% 
% This program is distributed in the hope that it will be useful, 
% but WITHOUT ANY WARRANTY; without even the implied warranty of 
% MERCHANTABILITY or FITNESS FOR A PARTICULAR PURPOSE. See the 
% GNU General Public License for more details. 
% 
% You should have received a copy of the GNU General Public License 
% along with this program. If not, see <http://www.gnu.org/licenses/>.

old = pwd;
cd([fileparts(which('installSBPOP')) '/..']);
recurseFolder('SBPOP',text);
cd(old);
return


function recurseFolder(folder,text)
% change folder
cd(folder);
% check all m files in folder for the text
mfiles = dir('*.m');
for k = 1:length(mfiles),
    % read file
    content = fileread(mfiles(k).name);
    if strfind(content,text),
        disp(mfiles(k).name);
        edit(mfiles(k).name);
    else
%        edit(mfiles(k).name);
    end
end
% recurse in all subfolders
allfiles = dir;
for k = 1:length(allfiles),
    if ~strcmp(allfiles(k).name,'.git') && ~strcmp(allfiles(k).name,'..') && ~strcmp(allfiles(k).name,'.') && allfiles(k).isdir == 1,
        recurseFolder(allfiles(k).name,text);
    end
end
% up we go
cd ..
return
```
