## Supplementary scripts for "A multi-scale *in silico* mouse model for insulin resistance and humanoid type 2 diabetes": setseedSBPOP.m

```
function [] = setseedSBPOP(defaultSeed)
% [DESCRIPTION]
% Set the default stream to defaultSeed. 
% In versions of MATLAB prior to 7.7, the control of the internal state of
% the random number stream used by rand is done by calling rand and randn
% directly with the 'seed'.
% For posterior version, the default stream is set by using RandStream
%
% [SYNTAX]
% [] = setseedSBPOP(defaultSeed)
%
% [INPUT]
% defaultSeed:   integer between 0 and 2^32.
%
% [OUTPUT]
%
% [ASSUMPTIONS]
% 
% [AUTHOR]
% SBPOP package 
% Copyright 2009 Novartis AG
% Author: Bruno Bieth
% Created: 2009-12-03
%
% [DATE]
%
% [PLATFORM]
% Windows XP Engine, MATLAB R2007a, MATLAB R2009a
%
% [KEYWORDS]
% seed, seed generation, default seed
% 
% [TOOLBOXES USED]
%
% [EXAMPLES]
%
% [VERIFIED BY]
% NOT YET

% Information:
% ============
% Copyright © 2012 Novartis Pharma AG
% 
% This program is Free Open Source Software: you can redistribute it and/or modify 
% it under the terms of the GNU General Public License as published by 
% the Free Software Foundation, either version 3 of the License, or 
% (at your option) any later version. 
% 
% This program is distributed in the hope that it will be useful, 
% but WITHOUT ANY WARRANTY; without even the implied warranty of 
% MERCHANTABILITY or FITNESS FOR A PARTICULAR PURPOSE. See the 
% GNU General Public License for more details. 
% 
% You should have received a copy of the GNU General Public License 
% along with this program. If not, see <http://www.gnu.org/licenses/>.

%%%%%%%%%%%%%%%%%%%%%%%%%%%%%%%%%%%%%%%%%%%%%%%%%%
% Check that seed is an integer between 0 and 2^32
%%%%%%%%%%%%%%%%%%%%%%%%%%%%%%%%%%%%%%%%%%%%%%%%%%
if (mod(defaultSeed,1)~=0) || defaultSeed<0 || defaultSeed>2^32,
    error('Seed must be an integer between 0 and 2^32');
end

%%%%%%%%%%%%%%%%%%%%%%%%%%%%%%%%%%%%%%%%%%%%%%%%%%
% Get the version release
%%%%%%%%%%%%%%%%%%%%%%%%%%%%%%%%%%%%%%%%%%%%%%%%%%
v=version('-release');

if str2num(v(1:3)) <7.7,
    randn('seed',seed);
    rand('seed',seed);
else
    s=RandStream.create('mrg32k3a','Seed',defaultSeed);
    RandStream.setGlobalStream(s);
end
```
