## Supplementary scripts for "A multi-scale *in silico* mouse model for insulin resistance and humanoid type 2 diabetes": SBPOPpublish.m

```
function [] = SBPOPpublish(scriptname,outputpath)

% scriptname  = 'SCRIPT_01_popPK';
% outputpath  = './';

outputname  = [scriptname '.html'];

%% Publish script using MATLAB function
options     = struct('format','html','outputDir',outputpath,'evalCode',false);
publish(scriptname,options);
try, movefile(fullfile(outputpath,[scriptname '.html']),fullfile(outputpath,outputname)); catch, end

%% Postprocess
content = fileread(fullfile(outputpath,outputname));

% Remove the Source code
ixstart = strfind(content,'##### SOURCE BEGIN #####');
ixend   = strfind(content,'##### SOURCE END #####');
content(ixstart-5:ixend+25) = [];
content = strrep(content,'<p class="footer"><br><a href="http://www.mathworks.com/products/matlab/">Published with MATLAB&reg; R2013a</a><br></p>','');

% Handle the Henning-Markup links
while ~isempty(strfind(content,'#_#'))
    ixLinks     = strfind(content,'#_#');
    ixstart     = ixLinks(1);
    ixend       = ixLinks(2)+2;
    link        = content(ixstart:ixend);
    link        = strrep(link,'#_#','');
    linkterms   = explodePCSB(link,'|');
    if length(linkterms) == 1,
        link = linkterms{1};
        name = linkterms{1};
    elseif length(linkterms) == 2,
        link = linkterms{2};
        name = linkterms{1};
    else
        error('Wrong Link');
    end
    linkText    = ['<a href="' link '" target="newwindow">' name '</a>'];
    content = [content(1:ixstart-1) linkText content(ixend+1:end)];
end

%% Handle Table of contents Henning markup

% Get the contents table elements
ixContentsStart         = strfind(content,'<h2>Contents</h2>')+22;
ixUL                    = strfind(content,'</ul>');
ixContentsEnd           = ixUL(find(ixUL-ixContentsStart>0)); ixContentsEnd = ixContentsEnd(1)+4;
contentsTextOriginal    = content(ixContentsStart:ixContentsEnd);
contentsTextReplace     = contentsTextOriginal;
% Remove <ul> tags
contentsTextOriginal    = strrep(contentsTextOriginal,'<ul>','');
contentsTextOriginal    = strrep(contentsTextOriginal,'</ul>','');
% Remove </li> tags
contentsTextOriginal    = strrep(contentsTextOriginal,'</li>','');
% Remove first <li> tag
contentsTextOriginal    = contentsTextOriginal(5:end);
% EXCHANGE <li> tags
contentsTextOriginal    = strrep(contentsTextOriginal,'<li>','$');
% Get contents items
terms = explodePCSB(contentsTextOriginal,'$');
% Piece them together again ... assume a === In the name means a main
% section and not === means a subsection
ix_main = [];
for k=1:length(terms),
    if ~isempty(strfind(terms{k},'===')),
        ix_main = [ix_main k];
    end
end
if isempty(ix_main),
    error('At least one main section needs to be available.');
end
if ix_main(1)~=1,
    error('The first section needs to be a main section.');
end
ix_sub = {};
for k=2:length(ix_main),
    ix_maincurrent  = ix_main(k);
    ix_mainprevious = ix_main(k-1);
    ix_subk = [ix_mainprevious+1:1:ix_maincurrent-1];
    ix_sub{k-1} = ix_subk;
end
ix_sub{end+1} = ix_main(end)+1:1:length(terms);
text = sprintf('<ul>\n');
for k=1:length(ix_main),
    text = sprintf('%s    <li>',text);
    text = sprintf('%s<b>%s</b>\n',text,strtrim(strrep(upper(terms{ix_main(k)}),'===','')));
    if ~isempty(ix_sub{k}),
        text = sprintf('%s        <ul>\n',text);
        for k2=1:length(ix_sub{k}),
            text = sprintf('%s            <li>',text);
            text = sprintf('%s%s',text,strtrim(strrep(terms{ix_sub{k}(k2)},'===','')));
            text = sprintf('%s</li>\n',text);
        end
        text = sprintf('%s        </ul>\n',text);
    end
    text = sprintf('%s    </li>\n',text);
end
text = sprintf('%s</ul>\n',text);
content = strrep(content,contentsTextReplace,text);
content = strrep(content,'<h2>===','<h2>');

% Save post-processed result
fid = fopen(fullfile(outputpath,outputname),'w');
fprintf(fid,'%s',content);
fclose(fid);

% %% Open result
% oldpath = pwd(); cd(outputpath); path = pwd; cd(oldpath)
% system(sprintf('"C:\\Program Files\\Internet Explorer\\iexplore" %s',fullfile(path,outputname)))
```
