## Supplementary scripts for "A multi-scale *in silico* mouse model for insulin resistance and humanoid type 2 diabetes": functionTemplateSBPOP.m

```
function [ output_args ] = functionTemplateSBPOP( input_args )
% [DESCRIPTION]
% In this section the purpose of the function should be described.
% Additionally, the different possible syntaxes. 
%
% [SYNTAX]
% Here you can show how the function can be called
%
% [INPUT]
% Define the input arguments or other inputs (e.g. required files, etc.)
% Please do NOT use hardcoded data in MATLAB functions.
%
% [OUTPUT]
% Define the output arguments or other outputs (e.g. produced log files,
% figures, etc.) 
%
% [ASSUMPTIONS]
% Here you can add any additional background information about assumptions.
%
% [AUTHOR]
% Original author, ideally with email address
%
% [DATE]
% Date of first creation
%
% [PLATFORM]
% Windows XP, MATLAB R2012a, ...
%
% [KEYWORDS]
% Any keyword you like ... comma separated
% 
% [TOOLBOXES USED]
% Any commercial and other MATLAB toolboxes that need to be available for
% this script to be run. PLEASE ADD ALSO THE VERSION/REVISION NUMBER OF THE
% USED TOOLBOXES, so you are always able to reproduce your results.
%
% [EXAMPLES]
% Here you can provide one or more examples that allow to run the
% function.
%
% [VALIDATION HISTORY]
% �
% [MODIFICATION HISTORY]
% including version numbering, changes, author and date of changes

% Here your function starts. The above header should be ended by an empty
% line.
```
