## Supplementary scripts for "A multi-scale *in silico* mouse model for insulin resistance and humanoid type 2 diabetes": newfunctionSBPOP.m

```
function [] = newfunctionSBPOP(varargin)
% newfunctionSBPOP: Creates a new SBPOP function that is compatible with the
% business guidelines in terms of header and additionally compatible with 
% the MODSPACE requirements for MATLAB file parsing.
%
% USAGE:
% ======
%   newfunctionsSBPOP
%   newfunctionsSBPOP('functionname')

% Information:
% ============
% Copyright � 2012 Novartis Pharma AG
% 
% This program is Free Open Source Software: you can redistribute it and/or modify 
% it under the terms of the GNU General Public License as published by 
% the Free Software Foundation, either version 3 of the License, or 
% (at your option) any later version. 
% 
% This program is distributed in the hope that it will be useful, 
% but WITHOUT ANY WARRANTY; without even the implied warranty of 
% MERCHANTABILITY or FITNESS FOR A PARTICULAR PURPOSE. See the 
% GNU General Public License for more details. 
% 
% You should have received a copy of the GNU General Public License 
% along with this program. If not, see <http://www.gnu.org/licenses/>.

file = which('functionTemplateSBPOP.m');
if nargin == 0,
    copyfile(file,[pwd '/newfunction.m']);
elseif nargin == 1,
    if ~ischar(varargin{1}),
        copyfile(file,[pwd '/newfunction.m']);
    else
        filename = varargin{1};
        filename = strrep(filename,'.m','');
        copyfile(file,[pwd '/' filename '.m']);
    end
else
    error('Incorrect number of input arguments.');
end
    
```
