## Supplementary scripts for "A multi-scale *in silico* mouse model for insulin resistance and humanoid type 2 diabetes": display.m

```
function [] = display(dos)
% display: Displays information about SBPOPdosing object. This function is 
% called by MATLAB whenever an object is the result of a statement that
% is not terminated by a semicolon. 

% Information:
% ============
% Copyright � 2012 Novartis Pharma AG
% 
% This program is Free Open Source Software: you can redistribute it and/or modify 
% it under the terms of the GNU General Public License as published by 
% the Free Software Foundation, either version 3 of the License, or 
% (at your option) any later version. 
% 
% This program is distributed in the hope that it will be useful, 
% but WITHOUT ANY WARRANTY; without even the implied warranty of 
% MERCHANTABILITY or FITNESS FOR A PARTICULAR PURPOSE. See the 
% GNU General Public License for more details. 
% 
% You should have received a copy of the GNU General Public License 
% along with this program. If not, see <http://www.gnu.org/licenses/>.

%%%%%%%%%%%%%%%%%%%%%%%%%%%%%%%%%%%%%%%%%%%%%%%
% COLLECT INFORMATION ABOUT THE EXPERIMENT
%%%%%%%%%%%%%%%%%%%%%%%%%%%%%%%%%%%%%%%%%%%%%%%
nrinputs = length(dos.inputs);
nameinputs = {dos.inputs.name};
typeinputs = {dos.inputs.type};

%%%%%%%%%%%%%%%%%%%%%%%%%%%%%%%%%%%%%%%%%%%%%%%
% DISPLAY INFORMATION
%%%%%%%%%%%%%%%%%%%%%%%%%%%%%%%%%%%%%%%%%%%%%%%
text = sprintf('\tSBPOPdosing\n\t===========\n');
text = sprintf('%s\tName:  %s\n',text,dos.name);
% text = sprintf('%s\tNotes: %s\n',text,dos.notes);
text = sprintf('%s\tNumber inputs:     %d',text,nrinputs);
disp(text);
for k=1:length(nameinputs),
    if length(dos.inputs(k).time) > 1,
        type2 = 'muliple';
    else 
        type2 = 'single';
    end
    if ~isempty(dos.inputs(k).Tlag),
        disp(sprintf('\t\t%s: %s %s\t\t(LAG)',dos.inputs(k).name,type2,dos.inputs(k).type));
    else
        disp(sprintf('\t\t%s: %s %s',dos.inputs(k).name,type2,dos.inputs(k).type));
    end
end
```
