## Supplementary scripts for "A multi-scale *in silico* mouse model for insulin resistance and humanoid type 2 diabetes": SBPOPdosing.m

```
function [dos] = SBPOPdosing(varargin)
% SBPOPdosing: creates an SBPOPdosing object defining a dosing schedule
%
% USAGE:
% ======
% [dos] = SBPOPdosing()             creates an empty SBPOPdosing object
% [dos] = SBPOPdosing(SBstructure)  creates an SBPOPdosing object from a MATLAB
%                                   structure in the internal SBPOPdosing format
% [dos] = SBPOPdosing(dosin)        construction from a given SBPOPdosing object (dosin)
% [dos] = SBPOPdosing('file.dos')   converting a SBPOPdosing text description 
%                                   to an SBPOPdosing object.
%
% Output Arguments:
% =================
% dos: SBPOPdosing object 

% Information:
% ============
% Copyright � 2012 Novartis Pharma AG
% 
% This program is Free Open Source Software: you can redistribute it and/or modify 
% it under the terms of the GNU General Public License as published by 
% the Free Software Foundation, either version 3 of the License, or 
% (at your option) any later version. 
% 
% This program is distributed in the hope that it will be useful, 
% but WITHOUT ANY WARRANTY; without even the implied warranty of 
% MERCHANTABILITY or FITNESS FOR A PARTICULAR PURPOSE. See the 
% GNU General Public License for more details. 
% 
% You should have received a copy of the GNU General Public License 
% along with this program. If not, see <http://www.gnu.org/licenses/>.

%%%%%%%%%%%%%%%%%%%%%%%%%%%%%%%%%%%%%%%%%%%%%%%
% CHECK THE TYPE OF THE INPUT
%%%%%%%%%%%%%%%%%%%%%%%%%%%%%%%%%%%%%%%%%%%%%%%
if nargin==0,
    inputType = 'empty';
elseif nargin == 1 || nargin == 2,
    if isSBPOPdosing(varargin{1}),
        inputType = 'SBPOPdosing';
        dosInput = varargin{1};
    elseif isstruct(varargin{1}),
        inputType = 'SBstructure';
        SBstructure = varargin{1};
    elseif ischar(varargin{1}),
        % check if '.dos' given as extension. If yes, then import text description
        filename = varargin{1};
        if ~isempty(strfind(filename,'.dos')),
            inputType = 'TextDosFile';
        elseif strcmp('DosingAsTextString', varargin{2}),
            inputType = varargin{2};
        else
            error('Input argument of unknown type');
        end
    else 
        error('Input argument of unknown type');
    end
else
    error('Wrong number of input arguments.');
end

%%%%%%%%%%%%%%%%%%%%%%%%%%%%%%%%%%%%%%%%%%%%%%%
% CONSTRUCT THE SBPOPdosing OBJECT 
%%%%%%%%%%%%%%%%%%%%%%%%%%%%%%%%%%%%%%%%%%%%%%%
if strcmp('empty',inputType),
    % Create empty SBstructure
    % parameters substructure
    parametersStruct = struct('name',{},'value',{},'notes',{});
    % inputs substructure
    inputsStruct = struct('name',{},'type',{},'time',{},'Tlag',{},'D',{},'parameters',parametersStruct,'TlagNotes',{},'notes',{});
    % Create SBstructure
    SBstructure = struct('type','SBPOPdosing','name','unnamed_dosing','notes','no notes','inputs',inputsStruct);
    % construct the dosing object
    dos = class(SBstructure,'SBPOPdosing');
elseif strcmp('SBPOPdosing',inputType),
    % copy the object
    dos = dosInput;
elseif strcmp('SBstructure',inputType),
    % check if the given structure is a SBstructure 
    if isfield(SBstructure,'type'),
        if ~strcmp(SBstructure.type,'SBPOPdosing'),
            error('Given structure is not a valid internal SBPOPdosing structure.');
        end
    else
        error('Given structure is not a valid internal SBPOPdosing structure.');
    end
    % construct the dosing object
    dos = class(SBstructure,'SBPOPdosing');
elseif strcmp('TextDosFile',inputType),
    % check if a file with given filename exists
    [path,filename,ext] = fileparts(filename);
    filename = fullfile(path, [filename '.dos']); 
    if ~exist(filename),
        error(sprintf('Dosing file, "%s", does not exist.', filename));
    end
    % If file exists then first load it
    dosText = fileread(filename);
    % then convert it to SBstructure
    [SBstructure, errorMsg] = convertTextToDosSBPOP(dosText);
    % Check if error occurred while importing the dosing description
    if ~isempty(errorMsg),
        error(errorMsg);
    end
    % construct the dosing object
    dos = class(SBstructure,'SBPOPdosing');
elseif strcmp('DosingAsTextString', inputType),
    dosText = varargin{1};
    % then convert text dosing to SBstructure
    [SBstructure, errorMsg] = convertTextToDosSBPOP(dosText);
    % Check if error occurred while importing the dosing definition
    if ~isempty(errorMsg),
        error(errorMsg);
    end
    % construct the dosing object
    dos = class(SBstructure,'SBPOPdosing');
else
    error('Wrong input arguments.');
end
return
```
