## Supplementary scripts for "A multi-scale *in silico* mouse model for insulin resistance and humanoid type 2 diabetes": dosing2doseeventSBPOP.m

```
function [doseeventstruct] = dosing2doseeventSBPOP(dosing)
% dosing2doseeventSBPOP: This function takes an SBPOP dosing object as
% input and returns a structure in which the dosing events, happening, are
% sorted according to the times at which they happen. Additionally, the
% output structure contains information about the dosing amount and the
% names of the parameters in a model to change in order to apply the
% dosing. Also lag and the other dosing type dependend informations are
% collected.
% 
% USAGE:
% ======
% [doseeventstruct] = dosing2doseeventSBPOP(dosing) 
%
% dosing: SBPOPdosing object
%
% Output Arguments:
% =================
% doseeventstruct: output argument with the following structure:
%   .time                   time of dosing application
%   .input 
%   .input.index            index of dosing input in dosing object to be applied
%   .input.timeparname      name of time parameter for this input
%   .input.parameternames   cell-array with other parameter names to change the values for
%   .input.parametervalues  vector with new parameter values (same order as
%                           parameternames cell-array)

% Information:
% ============
% Copyright � 2012 Novartis Pharma AG
% 
% This program is Free Open Source Software: you can redistribute it and/or modify 
% it under the terms of the GNU General Public License as published by 
% the Free Software Foundation, either version 3 of the License, or 
% (at your option) any later version. 
% 
% This program is distributed in the hope that it will be useful, 
% but WITHOUT ANY WARRANTY; without even the implied warranty of 
% MERCHANTABILITY or FITNESS FOR A PARTICULAR PURPOSE. See the 
% GNU General Public License for more details. 
% 
% You should have received a copy of the GNU General Public License 
% along with this program. If not, see <http://www.gnu.org/licenses/>.

%%%%%%%%%%%%%%%%%%%%%%%%%%%%%%%%%%%%%%%%%%%%%%%
% Check input
%%%%%%%%%%%%%%%%%%%%%%%%%%%%%%%%%%%%%%%%%%%%%%%
if ~isSBPOPdosing(dosing),
    error('Input argument is not an SBPOPdosing object.');
end

%%%%%%%%%%%%%%%%%%%%%%%%%%%%%%%%%%%%%%%%%%%%%%%
% Get dosing structure
%%%%%%%%%%%%%%%%%%%%%%%%%%%%%%%%%%%%%%%%%%%%%%%
ds = struct(dosing);

%%%%%%%%%%%%%%%%%%%%%%%%%%%%%%%%%%%%%%%%%%%%%%%
% Get a dosing matrix and dose/time parameter 
% names as added to a model
%%%%%%%%%%%%%%%%%%%%%%%%%%%%%%%%%%%%%%%%%%%%%%%
dosingmatrix = [];
inames = {}; 
for k=1:length(ds.inputs),
    dosingmatrix = [dosingmatrix; ds.inputs(k).time(:) ds.inputs(k).D(:) k*ones(length(ds.inputs(k).D),1)];
    inames{k} = ds.inputs(k).name;
    %%%%%%%%%%%%%%%%%%%%%%%%%%%%%%%%%%%%%%%%%%%%%%%
    % Check if dosing times unique
    %%%%%%%%%%%%%%%%%%%%%%%%%%%%%%%%%%%%%%%%%%%%%%%
    if length(ds.inputs(k).time) ~= length(unique(ds.inputs(k).time)),
        error('Dosing times in dosing object are not unique.');
    end
end
% sort the dosing matrix according to the time vector
dosingmatrix = sortrows(dosingmatrix,1);

%%%%%%%%%%%%%%%%%%%%%%%%%%%%%%%%%%%%%%%%%%%%%%%
% For each input in dosing scheme get info about 
% Tlag and the parameter
%%%%%%%%%%%%%%%%%%%%%%%%%%%%%%%%%%%%%%%%%%%%%%%
inputinfoadditional = [];
for k=1:length(ds.inputs),
    inputinfoadditional(k).input = ds.inputs(k).name; % unused but just to have it in there when looking at this structure
    inputinfoadditional(k).index = k; % unused but just to have it in there when looking at this structure
    try
        inputinfoadditional(k).paramname = ds.inputs(k).parameters.name;
        inputinfoadditional(k).param = ds.inputs(k).parameters.value;
    catch
        inputinfoadditional(k).paramname = {};
        inputinfoadditional(k).param = [];
    end
    inputinfoadditional(k).Tlag = ds.inputs(k).Tlag;
end

%%%%%%%%%%%%%%%%%%%%%%%%%%%%%%%%%%%%%%%%%%%%%%%
% convert dosing matrix to structure
% dosingevent.time
% dosingevent.input.index
% dosingevent.input.timeparname
% dosingevent.input.parameternames
% dosingevent.input.parametervalues
%% %%%%%%%%%%%%%%%%%%%%%%%%%%%%%%%%%%%%%%%%%%%%%
inputstruct = struct('index',{},'timeparname',{},'parameternames',{},'parametervalues',{});
dosingevent = struct('time',{},'input',inputstruct);
times = unique(dosingmatrix(:,1));
for k=1:length(times),
    % find dosings for current time
    di = find(dosingmatrix(:,1)==times(k));
    % add info to dosingevent structure
    dosingevent(k).time = times(k);
    for k2=1:length(di),
        index = dosingmatrix(di(k2),3);
        dosingevent(k).input(k2).index = index;
        dosingevent(k).input(k2).timeparname = ['Time_' lower(inames{index})]; % Name definition used in mergemoddosSBPOP
        % other parameters
        parameternames = {};
        parameternames{end+1} = ['Dose_' lower(inames{index})];
        if ~isempty(inputinfoadditional(index).Tlag),
            parameternames{end+1} = ['Tlag_' lower(inames{index})];
        end
        
        if ~isempty(inputinfoadditional(index).paramname),
            parameternames{end+1} = [inputinfoadditional(index).paramname '_' lower(inames{index})];
        end

        parametervalues = [];
        parametervalues(end+1) = dosingmatrix(di(k2),2);
        if ~isempty(inputinfoadditional(index).Tlag),
            parametervalues(end+1) = inputinfoadditional(index).Tlag;
        end

        if ~isempty(inputinfoadditional(index).param),
            timedoseinput = dosingevent(k).time;
            timeINPUT = ds.inputs(dosingevent(k).input(k2).index).time;
            paramvalueINPUT = ds.inputs(dosingevent(k).input(k2).index).parameters.value;
            pvalue = paramvalueINPUT(timeINPUT==timedoseinput);
            parametervalues(end+1) = pvalue;
        end
        
        dosingevent(k).input(k2).parameternames = parameternames;
        dosingevent(k).input(k2).parametervalues = parametervalues;
    end
end
%%
%%%%%%%%%%%%%%%%%%%%%%%%%%%%%%%%%%%%%%%%%%%%%%%
% Return the result
%%%%%%%%%%%%%%%%%%%%%%%%%%%%%%%%%%%%%%%%%%%%%%%
doseeventstruct = dosingevent;
return
```
