## Supplementary scripts for "A multi-scale *in silico* mouse model for insulin resistance and humanoid type 2 diabetes": getmoddosinputinfoSBPOP.m

```
function [inputinfo] = getmoddosinputinfoSBPOP(model,dosing)
% getmoddosinputinfoSBPOP: Checks the availability of dosing input
% definitions used in the model in the dosing object and returns a
% structure similar to the "input" field structure of an SBmodel, augmented
% with the dosing information, defined in the SBPOPdosing object.
%
% If the dosing object does not contain all inputs, required by the model,
% a warning is displayed and the undefined inputs are deleted from the
% "inputinfo" structure. This allows inputs in an SBmodel to be kept
% undefined.
%
% USAGE:
% ======
% [inputinfo] = getmoddosinputinfoSBPOP(model,dosing) 
%
% model: SBmodel
% dose: SBPOPdosing object
%
% Output Arguments:
% =================
% inputinfo: Same structure as defined in the SBmodel "input" field.
%   Augmented by the dosing input information, stroed in the dosing object.

% Information:
% ============
% Copyright � 2012 Novartis Pharma AG
% 
% This program is Free Open Source Software: you can redistribute it and/or modify 
% it under the terms of the GNU General Public License as published by 
% the Free Software Foundation, either version 3 of the License, or 
% (at your option) any later version. 
% 
% This program is distributed in the hope that it will be useful, 
% but WITHOUT ANY WARRANTY; without even the implied warranty of 
% MERCHANTABILITY or FITNESS FOR A PARTICULAR PURPOSE. See the 
% GNU General Public License for more details. 
% 
% You should have received a copy of the GNU General Public License 
% along with this program. If not, see <http://www.gnu.org/licenses/>.

%%%%%%%%%%%%%%%%%%%%%%%%%%%%%%%%%%%%%%%%%%%%%%%
% Check input
%%%%%%%%%%%%%%%%%%%%%%%%%%%%%%%%%%%%%%%%%%%%%%%
if ~isSBmodel(model),
    error('First input argument is not an SBmodel.');
end
if ~isSBPOPdosing(dosing),
    error('Second input argument is not an SBPOPdosing object.');
end

%%%%%%%%%%%%%%%%%%%%%%%%%%%%%%%%%%%%%%%%%%%%%%%
% Get model and dosing structures
%%%%%%%%%%%%%%%%%%%%%%%%%%%%%%%%%%%%%%%%%%%%%%%
ms = struct(model);
ds = struct(dosing);

%%%%%%%%%%%%%%%%%%%%%%%%%%%%%%%%%%%%%%%%%%%%%%%
% Check if model contains inputs.
% If not => error!
%%%%%%%%%%%%%%%%%%%%%%%%%%%%%%%%%%%%%%%%%%%%%%%    
if isempty(ms.inputs),
    error('The model does not contain any inputs. Merging with dosing objects does not make sense.');
end

%%%%%%%%%%%%%%%%%%%%%%%%%%%%%%%%%%%%%%%%%%%%%%%
% Initialize the inputinfo structure
%%%%%%%%%%%%%%%%%%%%%%%%%%%%%%%%%%%%%%%%%%%%%%%
inputinfo = [];

%%%%%%%%%%%%%%%%%%%%%%%%%%%%%%%%%%%%%%%%%%%%%%%
% Run through all inputs, defined in the model 
% and process the information in the dosing
% object accordingly
%%%%%%%%%%%%%%%%%%%%%%%%%%%%%%%%%%%%%%%%%%%%%%%
minputs = ms.inputs;
dinputs = ds.inputs;
for k=1:length(minputs),
    % First get the info stored in the model
    minfo = minputs(k);
    % Check if input exists in the dosing object
    index = strmatchSB(minfo.name,{dinputs.name},'exact');
    if ~isempty(index),
        % Input does exist in the dosing object => Now add the complete
        % input information to the inputinfo structure 
        % Get the info for this input stored in the dosing object
        dinfo = dinputs(index);
        % Add dosing schedule information to the minfo structure
        minfo.type = dinfo.type;
        minfo.time = dinfo.time;
        minfo.Tlag = dinfo.Tlag;
        minfo.D = dinfo.D;
        minfo.parameters = dinfo.parameters;
        minfo.notes = dinfo.notes;
        % Add minfo structure to the inputinfo output structure
        if isempty(inputinfo),
            inputinfo = minfo;
        else
            inputinfo(end+1) = minfo;
        end
    else
        % Input does not exist in the dosing object => warning to user 
        % and do not put it into the inputinfo structure
%         disp(sprintf('The input ''%s'', defined in the model, is not defined in the SBPOPdosing object.',minfo.name));
    end
end

%%%%%%%%%%%%%%%%%%%%%%%%%%%%%%%%%%%%%%%%%%%%%%%
% Check if any inputs of the models have been 
% found in the dosing object. If not => error!
%%%%%%%%%%%%%%%%%%%%%%%%%%%%%%%%%%%%%%%%%%%%%%%    
if isempty(inputinfo),
    error('No inputs defined in the model could be found in the dosing object. This should be wrong ...')        
end
    
```
