## Supplementary scripts for "A multi-scale *in silico* mouse model for insulin resistance and humanoid type 2 diabetes": convertDosToTextSBPOP.m

```
function [dosTextStructure] = convertDosToTextSBPOP(dos)
% convertDosToTextSBPOP: Converts an SBPOPdosing object to a structure
% containing the different parts of the text description of the dosing
% scheme.

% Information:
% ============
% Copyright � 2012 Novartis Pharma AG
% 
% This program is Free Open Source Software: you can redistribute it and/or modify 
% it under the terms of the GNU General Public License as published by 
% the Free Software Foundation, either version 3 of the License, or 
% (at your option) any later version. 
% 
% This program is distributed in the hope that it will be useful, 
% but WITHOUT ANY WARRANTY; without even the implied warranty of 
% MERCHANTABILITY or FITNESS FOR A PARTICULAR PURPOSE. See the 
% GNU General Public License for more details. 
% 
% You should have received a copy of the GNU General Public License 
% along with this program. If not, see <http://www.gnu.org/licenses/>.

% Initialize variables
dosTextStructure = [];
% Get SBstructure
ds = SBPOPstruct(dos);
% Parse structure into the dosTextStructure description

%%%%%%%%%%%%%%%%%%%%%%%%%%%%%%%%%%%%%%%%%%%%%%%
% Name
%%%%%%%%%%%%%%%%%%%%%%%%%%%%%%%%%%%%%%%%%%%%%%%
dosTextStructure.name = ds.name;

%%%%%%%%%%%%%%%%%%%%%%%%%%%%%%%%%%%%%%%%%%%%%%%
% Notes
%%%%%%%%%%%%%%%%%%%%%%%%%%%%%%%%%%%%%%%%%%%%%%%
dosTextStructure.notes = ds.notes;

%%%%%%%%%%%%%%%%%%%%%%%%%%%%%%%%%%%%%%%%%%%%%%%
% Inputs
%%%%%%%%%%%%%%%%%%%%%%%%%%%%%%%%%%%%%%%%%%%%%%%
dosTextStructure.inputs = {};
allInputText = {};
for k=1:length(ds.inputs),
    inputText = '';
    input = ds.inputs(k);
    % Write the limiter (name)
    inputText = sprintf('********** %s\n',input.name);
    % Type
    inputText = sprintf('%stype:         %s\n',inputText,input.type);
    % Time
    if length(input.time) == 1,
        inputText = sprintf('%stime:         %g\n',inputText,input.time);
    else
        timeText = sprintf('%g, ',input.time);
        inputText = sprintf('%stime:         [%s]\n',inputText,timeText(1:end-2));
    end
    % Tlag (write only if non-zero)
    if input.Tlag ~= 0,
        if isempty(input.TlagNotes),
            inputText = sprintf('%sTlag:         %g\n',inputText,input.Tlag);
        else
            inputText = sprintf('%sTlag:         %g    %% %s\n',inputText,input.Tlag,input.TlagNotes);
        end
    end
    % D
    if length(input.D) == 1,
        inputText = sprintf('%sD:            %g\n',inputText,input.D);
    else
        doseText = sprintf('%g, ',input.D);
        inputText = sprintf('%sD:            [%s]\n',inputText,doseText(1:end-2));
    end
    % parameters
    if ~isempty(input.parameters),
        for k2=1:length(input.parameters),
            parametersValueText = sprintf('%g, ',input.parameters(k2).value);
            if isempty(input.parameters(k2).notes),
                inputText = sprintf('%s%s:%s[%s]\n',inputText,input.parameters(k2).name,char(32*ones(1,14-length(input.parameters(k2).name)-1)),parametersValueText(1:end-2));
            else
                inputText = sprintf('%s%s:%s[%s]    %% %s\n',inputText,input.parameters(k2).name,char(32*ones(1,14-length(input.parameters(k2).name)-1)),parametersValueText(1:end-2),input.parameters(k2).notes);
            end
        end
    end
    % notes
    if ~isempty(input.notes),
        inputText = sprintf('%snotes:        %s\n',inputText,input.notes);
    end
    % save text
    dosTextStructure.inputs{k} = inputText;
end
```
