## Supplementary scripts for "A multi-scale *in silico* mouse model for insulin resistance and humanoid type 2 diabetes": setPartsToCompleteTextDosSBPOP.m

```
function [completeText] = setPartsToCompleteTextDosSBPOP(dosTextStructure)
% setPartsToCompleteTextDosSBPOP: Sets the different parts of a dosing 
% scheme description of an SBdosing object together to the complete text

% Information:
% ============
% Copyright � 2012 Novartis Pharma AG
% 
% This program is Free Open Source Software: you can redistribute it and/or modify 
% it under the terms of the GNU General Public License as published by 
% the Free Software Foundation, either version 3 of the License, or 
% (at your option) any later version. 
% 
% This program is distributed in the hope that it will be useful, 
% but WITHOUT ANY WARRANTY; without even the implied warranty of 
% MERCHANTABILITY or FITNESS FOR A PARTICULAR PURPOSE. See the 
% GNU General Public License for more details. 
% 
% You should have received a copy of the GNU General Public License 
% along with this program. If not, see <http://www.gnu.org/licenses/>.

completeText = sprintf('********** DOSING NAME\n%s\n\n',dosTextStructure.name);
completeText = sprintf('%s********** DOSING NOTES\n%s\n\n',completeText,dosTextStructure.notes);
if ~isempty(dosTextStructure.inputs),
    for k=1:length(dosTextStructure.inputs),
        completeText = sprintf('%s%s\n',completeText,dosTextStructure.inputs{k});
    end
else
    % If no inputs defined in the structure then print out a guideline on
    % how to represent the different type of input.
    completeText = sprintf('%s********** INPUT1\n',completeText);
    completeText = sprintf('%s%%%% Data required to define an Bolus (single, multiple)\n',completeText);
    completeText = sprintf('%s%%type: BOLUS\n',completeText);
    completeText = sprintf('%s%%notes:              %% (optional)\n',completeText);
    completeText = sprintf('%s%%time:               %% time for first application or all applications (scalar or vector)\n',completeText);
    completeText = sprintf('%s%%deltaT:             %% time inbetween applications (only if time scalar)\n',completeText);
    completeText = sprintf('%s%%nr_repetitions:  	%% number of applications (only if time scalar + optional if time scalar)\n',completeText);
    completeText = sprintf('%s%%D:                  %% dose (scalar or vector)\n',completeText);
    completeText = sprintf('%s%%Tlag:               %% lag time for input application (optional). Default: 0\n',completeText);
    completeText = sprintf('%s\n',completeText);
    completeText = sprintf('%s%%%% Data required to define an Infusion (with infusion rate as parameter) (single, multiple)\n',completeText);
    completeText = sprintf('%s%%type: INFUSION\n',completeText);
    completeText = sprintf('%s%%notes:              %% (optional)\n',completeText);
    completeText = sprintf('%s%%time:               %% time for first application or all applications (scalar or vector)\n',completeText);
    completeText = sprintf('%s%%deltaT:             %% time inbetween applications (only if time scalar)\n',completeText);
    completeText = sprintf('%s%%nr_repetitions: 	%% number of applications (only if time scalar + optional if time scalar)\n',completeText);
    completeText = sprintf('%s%%D:                  %% dose (scalar or vector)\n',completeText);
    completeText = sprintf('%s%%Rate:               %% Infusion rate (required)\n',completeText);
    completeText = sprintf('%s%%Tlag:               %% lag time for input application (optional). Default: 0\n',completeText);
    completeText = sprintf('%s\n',completeText);
    completeText = sprintf('%s%%%% ALTERNATIVE: Data required to define an Infusion (with infusion time as parameter) (single, multiple)\n',completeText);
    completeText = sprintf('%s%%type: INFUSION\n',completeText);
    completeText = sprintf('%s%%notes:              %% (optional)\n',completeText);
    completeText = sprintf('%s%%time:               %% time for first application or all applications (scalar or vector)\n',completeText);
    completeText = sprintf('%s%%deltaT:             %% time inbetween applications (only if time scalar)\n',completeText);
    completeText = sprintf('%s%%nr_repetitions: 	%% number of applications (only if time scalar + optional if time scalar)\n',completeText);
    completeText = sprintf('%s%%D:                  %% dose (scalar or vector)\n',completeText);
    completeText = sprintf('%s%%Tinf:               %% Infusion time (required)\n',completeText);
    completeText = sprintf('%s%%Tlag:               %% lag time for input application (optional). Default: 0\n',completeText);
    completeText = sprintf('%s\n',completeText);
    completeText = sprintf('%s%%%% Data required to define a 1st order absorption (single, multiple)\n',completeText);
    completeText = sprintf('%s%%type: ABSORPTION1\n',completeText);
    completeText = sprintf('%s%%notes:              %% (optional)\n',completeText);
    completeText = sprintf('%s%%time:               %% time for first application or all applications (scalar or vector)\n',completeText);
    completeText = sprintf('%s%%deltaT:             %% time inbetween applications (only if time scalar)\n',completeText);
    completeText = sprintf('%s%%nr_repetitions: 	%% number of applications (only if time scalar + optional if time scalar)\n',completeText);
    completeText = sprintf('%s%%D:                  %% dose (scalar or vector)\n',completeText);
    completeText = sprintf('%s%%ka:                 %% Absorption rate constant (required)\n',completeText);
    completeText = sprintf('%s%%Tlag:               %% lag time for input application (optional). Default: 0\n',completeText);
    completeText = sprintf('%s\n',completeText);
    completeText = sprintf('%s%%%% Data required to define a 0th order absorption (single, multiple)\n',completeText);
    completeText = sprintf('%s%%type: ABSORPTION0\n',completeText);
    completeText = sprintf('%s%%notes:              %% (optional)\n',completeText);
    completeText = sprintf('%s%%time:               %% time for first application or all applications (scalar or vector)\n',completeText);
    completeText = sprintf('%s%%deltaT:             %% time inbetween applications (only if time scalar)\n',completeText);
    completeText = sprintf('%s%%nr_repetitions: 	%% number of applications (only if time scalar + optional if time scalar)\n',completeText);
    completeText = sprintf('%s%%D:                  %% dose (scalar or vector)\n',completeText);
    completeText = sprintf('%s%%Tk0:                %% time for absorption (required)\n',completeText);
    completeText = sprintf('%s%%Tlag:               %% lag time for input application (optional). Default: 0\n',completeText);
    completeText = sprintf('%s\n',completeText);
end
return
```
