## Supplementary scripts for "A multi-scale *in silico* mouse model for insulin resistance and humanoid type 2 diabetes": convertTextToDosSBPOP.m

```
function [SBstructure,errorMsg] = convertTextToDosSBPOP(dosText)
% convertTextToDosSBPOP: Converts a text description of an SBPOPdosing
% object to the internal SBPOPdosing data structure representation.

% Information:
% ============
% Copyright � 2012 Novartis Pharma AG
% 
% This program is Free Open Source Software: you can redistribute it and/or modify 
% it under the terms of the GNU General Public License as published by 
% the Free Software Foundation, either version 3 of the License, or 
% (at your option) any later version. 
% 
% This program is distributed in the hope that it will be useful, 
% but WITHOUT ANY WARRANTY; without even the implied warranty of 
% MERCHANTABILITY or FITNESS FOR A PARTICULAR PURPOSE. See the 
% GNU General Public License for more details. 
% 
% You should have received a copy of the GNU General Public License 
% along with this program. If not, see <http://www.gnu.org/licenses/>.

% % initialize variables
errorMsg = '';
% errorConditions = '';
% errorParameterChanges = '';
% errorStateEvents = '';
SBstructure = [];

% cut text into pieces
dosTextStructure = getPartsFromCompleteTextDosSBPOP(dosText);

%%%%%%%%%%%%%%%%%%%%%%%%%%%%%%%%%%%%%%%%%%%%%%%
%%%%%%%%%%%%%%%%%% Initialize structure
%%%%%%%%%%%%%%%%%%%%%%%%%%%%%%%%%%%%%%%%%%%%%%%
SBstructure = struct(SBPOPdosing());

%%%%%%%%%%%%%%%%%%%%%%%%%%%%%%%%%%%%%%%%%%%%%%%
%%%%%%%%%%%%%%%%%% Name
%%%%%%%%%%%%%%%%%%%%%%%%%%%%%%%%%%%%%%%%%%%%%%%
SBstructure.name = strtrim(removeCharacters(dosTextStructure.name));

%%%%%%%%%%%%%%%%%%%%%%%%%%%%%%%%%%%%%%%%%%%%%%%
%%%%%%%%%%%%%%%%%% Notes
%%%%%%%%%%%%%%%%%%%%%%%%%%%%%%%%%%%%%%%%%%%%%%%
SBstructure.notes = strtrim(dosTextStructure.notes);

%%%%%%%%%%%%%%%%%%%%%%%%%%%%%%%%%%%%%%%%%%%%%%%
%%%%%%%%%%%%%%%%%% READ INPUTS
%%%%%%%%%%%%%%%%%%%%%%%%%%%%%%%%%%%%%%%%%%%%%%%
% 0) create empty input structure
parametersStruct = struct('name',{},'value',{},'notes',{});
inputStruct = struct('name',{},'type',{},'time',{},'Tlag',{},'D',{},'parameters',parametersStruct,'TlagNotes',{},'notes',{});
inputfieldnames = fieldnames(inputStruct);
timeDefStruct = struct('deltaT',{},'nr_repetitions',{});
for k=1:length(dosTextStructure.input),
    inputText = dosTextStructure.input{k};
    % all data for an input is organized in rows. each row holds a
    % statement. statements over several rows are NOT allowed.
    % ordering is arbitrary except that the name (INPUTx) will always be in
    % the first row. all other rows have an identifier followed by a colon
    % followed by a value.
    % 1) split text in rows
    rows = explodePCSB(inputText,char(10));
    % 2) INPUT name (check that "INPUT" is present)
    if isempty(regexp(rows{1},'INPUT', 'once' )),
        error('Please check input dose definitions. An "INPUTx" identifier seems to be corrupted.');
    end
    % 3) add name to struct
    inputStruct(k).name = strtrim(rows{1});
    % 3b) Initialize Tlag with default empty value
    inputStruct(k).Tlag = [];   % if undefined then leave empty!!!   
    % 4) add the rest (if row identifier equal to a input field ... add it
    % there, otherwise add it as a parameter).
    nr_repetitions = [];
    deltaT = [];
    for k2=2:length(rows),
        % check if row is empty dont consider it
        if ~isempty(strtrim(rows{k2})),
            % cut of everything from first percentage sign on (comment)
            % comment is saved since used for the additional parameters
            % (Rate, Tk0, ka)
            commentindex = strfind(rows{k2},'%');
            if ~isempty(commentindex),
                savenotes = strtrim(rows{k2}(commentindex(1)+1:end));
                rows{k2} = strtrim(rows{k2}(1:commentindex(1)-1));
            else
                savenotes = '';
            end
            % find first colon (if no colon then error
            colonindex = strfind(rows{k2},':');
            if isempty(colonindex),
                error('Syntax error in dose description. No colon in a row for input "%s".',inputStruct(k).name);
            end
            % get identifier and value
            identifier = strtrim(rows{k2}(1:colonindex-1));
            valuetry = strtrim(rows{k2}(colonindex+1:end));
            % evaluate the value (matlab notation allowed e.g. for vectors)
            % need to try and catch, since some values are strings :)
            try
                value = eval(valuetry);
            catch
                value = valuetry;
            end
            % Add value to field if identifier is fieldname
            if ~isempty(strmatchSB(identifier,inputfieldnames)),
                inputStruct(k) = setfield(inputStruct(k),identifier,value);
                if strcmp(identifier,'Tlag'),
                    inputStruct(k).TlagNotes = savenotes;
                end
            else
                % handle nr_repetitions and deltaT here
                if strcmp(identifier,'nr_repetitions'),
                    nr_repetitions = value;
                elseif strcmp(identifier,'deltaT'),
                    deltaT = value;
                else
                    % remaining non-field identifiers become parameters for the
                    % input to be handled later.
                    inputStruct(k).parameters(end+1).name = identifier;
                    inputStruct(k).parameters(end).value = value;
                    inputStruct(k).parameters(end).notes = savenotes;
                end
            end                
        end
    end
    % store deltaT and nr_repetitions information to be used in check
    timeDefStruct(k).deltaT = deltaT;
    timeDefStruct(k).nr_repetitions = nr_repetitions;
end
SBstructure.inputs = inputStruct;

%%%%%%%%%%%%%%%%%%%%%%%%%%%%%%%%%%%%%%%%%%%%%%%
%%%%%%%%%%%%%%%%%% CHECK INPUTS for consistency
%%%%%%%%%%%%%%%%%%%%%%%%%%%%%%%%%%%%%%%%%%%%%%%
for k=1:length(SBstructure.inputs),
    input = SBstructure.inputs(k);
    deltaT = timeDefStruct(k).deltaT;
    nr_repetitions = timeDefStruct(k).nr_repetitions;
   
    % Check general input consistency (time, deltaT, nr_repetitions, D,
    % notes)
    
    % 1) Dose D needs to be present and positive
    if isempty(input.D),
        error('No dose "D" defined for input "%s".',input.name);
    end
    if input.D < 0, 
        error('Dose "D" not allowed to be negative in input "%s".',input.name);
    end
    
    % 2) time needs to be defined
    if isempty(input.time),
        error('"time" needs to be defined in input "%s".',input.name);
    end
    
    % Check time against deltaT and nr_repetitions
    % 3a) If time is a vector deltaT and nr_rep need to be empty
    if length(input.time) > 1 && (~isempty(deltaT) || ~isempty(nr_repetitions)),
        error('When defining a time vector in "time", "nr_repetitions" and "deltaT" are not allowed to be defined in input "%s".',input.name);
    end
    % 3b) further checks
    if ~isempty(nr_repetitions),
        if nr_repetitions == 1,
            error('Do not use "nr_repetitions: 1" to specify single applications in input "%s".\nJust remove the "nr_repetitions" identifier.',input.name);
        end
        if isempty(deltaT),
            error('When defining "nr_repetitions", also "deltaT" needs to be defined in input "%s".',input.name);
        end
    else
        if ~isempty(deltaT),
            error('When defining "deltaT", also "nr_repetitions" needs to be defined in input "%s".',input.name);
        end
    end    
    
    % 4) Construct the time field (scalar for single or vector for multiple applications)
    if ~isempty(nr_repetitions),
        addTimeVector = [0:deltaT:deltaT*(nr_repetitions-1)];
        input.time = input.time+addTimeVector;
        % update in final SBPOPdosing structure
        SBstructure.inputs(k).time = input.time;
    end
    
    % 5) Check length of dose vector (either scalar or same length as time vector)
    if length(input.D) > 1,
        if length(input.D) ~= length(input.time),
            error('Length of "D" vector and "time" vector does not match in input "%s".',input.name);
        end
    elseif length(input.time) > 1,
        % If D scalar and time vector then expand D to a vector with the
        % same size as time ...
        D = input.D * ones(1,length(input.time));
        SBstructure.inputs(k).D = D;
    end
       
    % 5) Check that Tlag is positive
    if input.Tlag < 0,
        error('"Tlag" is not allowed to be negative in input "%s".',input.name);
    end

    % 6) Check parameter value of input
    if ~isempty(input.parameters),
        parametervalue = input.parameters.value;
        if length(parametervalue) == 1,
            % make vector out of it
            SBstructure.inputs(k).parameters.value = parametervalue*ones(1,length(SBstructure.inputs(k).time));
        else
            % Check length
            if length(parametervalue) ~= length(SBstructure.inputs(k).time),
                error('Length of "parameter value" vector and "time" vector does not match in input "%s".',input.name);
            end
        end
    end
    
    % Check input type and input type specific information
    if strcmp(input.type,'BOLUS'),
        % 1) Check Bolus inputs
        checkBolusInput(input);
    elseif strcmp(input.type,'INFUSION'),
        % 2) Check Infusion inputs
        checkInfusionInput(input)
    elseif strcmp(input.type,'ABSORPTION1'),
        % 3) Check 1st order absorbption inputs
        checkAbsorption1Input(input)
    elseif strcmp(input.type,'ABSORPTION0'),
        % 4) Check 0th order absorbption inputs
        checkAbsorption0Input(input)
    else
        % Error: unknown input type
        error('Unknown input type in dosing description in input "%s".',input.type);
    end
end

% Finally check if inputs with same names
a = {SBstructure.inputs.name};
if length(a) ~= length(unique(a)),
    error('At least two inputs are defined in the dosing scheme, which have the same name.');
end


%%%%%%%%%%%%%%%%%% END OF FUNCTION
return


%%%%%%%%%%%%%%%%%%%%%%%%%%%%%%%%%%%%%%%%%%%%%%%
% Check Bolus inputs
%%%%%%%%%%%%%%%%%%%%%%%%%%%%%%%%%%%%%%%%%%%%%%%
function [] = checkBolusInput(input)
    % no additional parameters are required => check that input.parameters
    % is empty
    if ~isempty(input.parameters),
        error('Bolus definition does not require additional parameters in input "%s".',input.name);
    end
    % Do check that Tlag is not to large. Tlag+(Tk0 or Dmax/Rate or deltaTBolus) < min delta T dosing is acceptable.
    % This limitation exists due to the implementation of the multiple
    % dosings for simulation using events to update the parameters that
    % perform the multiple inputs. (only if multiple inputs)
    if length(input.time) > 1,
        % Determine the minimum time between dosing instances
        minDeltaT = min(input.time(2:end)-input.time(1:end-1));
        if input.Tlag + 0.0001 >= minDeltaT,
            error('Tlag in ''%s'' is to large. Please choose it smaller than the min time between dosings.',input.name);
        end
    end
return

%%%%%%%%%%%%%%%%%%%%%%%%%%%%%%%%%%%%%%%%%%%%%%%
% Check Infusion inputs
%%%%%%%%%%%%%%%%%%%%%%%%%%%%%%%%%%%%%%%%%%%%%%%
% We need to check that multiple infusions do not lead to one infusion
% overlaying the previous one(s). This is done by checking that 
% max(D)/Rate < min delta time in application
function [] = checkInfusionInput(input)
    % Rate is required as additional input. 
    if length(input.parameters) ~= 1,
        error('Infusion definition does require a single additional parameter "Rate" or "Tinf" in input "%s".',input.name);
    end
    % Check name of parameter (required: Rate)
    if ~strcmp(input.parameters(1).name,'Rate') && ~strcmp(input.parameters(1).name,'Tinf'),
        error('Please use "Rate" or "Tinf" as infusion related parameter in input "%s".',input.name);
    end
    % Check non overlapping infusion inputs (only if multiple inputs)
    if length(input.time) > 1,
        if strcmp(input.parameters(1).name,'Rate'),
            % Determine minimum required time between dosing instances
            Rate = min(input.parameters(1).value);
            maxDose = max(input.D);
            minRequiredDeltaT = maxDose/Rate;
            % Determine the minimum time between dosing instances
            minDeltaT = min(input.time(2:end)-input.time(1:end-1));
            % Do the check
            if minRequiredDeltaT > minDeltaT,
                error('Infusion rate for input ''%s'' to low. Previous infusion would not be finished before the next one.',input.name);
            end
            % Do check that Tlag is not to large. Tlag+(Tk0 or Dmax/Rate or deltaTBolus) < min delta T dosing is acceptable.
            % This limitation exists due to the implementation of the multiple
            % dosings for simulation using events to update the parameters that
            % perform the multiple inputs. (only if multiple inputs)
            % Determine the minimum time between dosing instances
            if input.Tlag + minRequiredDeltaT >= minDeltaT,
                error('Tlag in ''%s'' is to large. Please choose it smaller than the min time between dosings minus max(Dose)/Rate.',input.name);
            end
        elseif strcmp(input.parameters(1).name,'Tinf'),
            % Determine minimum required time between dosing instances
            Tinf = max(input.parameters(1).value);
            minRequiredDeltaT = Tinf;
            % Determine the minimum time between dosing instances
            minDeltaT = min(input.time(2:end)-input.time(1:end-1));
            % Do the check
            if minRequiredDeltaT > minDeltaT,
                error('Infusion time for input ''%s'' to low. Previous infusion would not be finished before the next one.',input.name);
            end
            % Do check that Tlag is not to large. Tlag+(Tk0 or Dmax/Rate or deltaTBolus) < min delta T dosing is acceptable.
            % This limitation exists due to the implementation of the multiple
            % dosings for simulation using events to update the parameters that
            % perform the multiple inputs. (only if multiple inputs)
            % Determine the minimum time between dosing instances
            if input.Tlag + minRequiredDeltaT >= minDeltaT,
                error('Tlag in ''%s'' is to large. Please choose it smaller than the min time between dosings minus Tinf.',input.name);
            end
        end
    end
return

%%%%%%%%%%%%%%%%%%%%%%%%%%%%%%%%%%%%%%%%%%%%%%%
% Check 1st order Absorption inputs
%%%%%%%%%%%%%%%%%%%%%%%%%%%%%%%%%%%%%%%%%%%%%%%
function [] = checkAbsorption1Input(input)
    % ka is required as additional input.
    if length(input.parameters) ~= 1,
        error('1st order Absorption definition does require a single additional parameter "ka" in input "%s".',input.name);
    end
    % Check name of parameter (required: ka)
    if ~strcmp(input.parameters(1).name,'ka'),
        error('Please use "ka" as absorption rate parameter in input "%s".',input.name);
    end
    % Do check that Tlag is not to large. Tlag+(Tk0 or Dmax/Rate or deltaTBolus) < min delta T dosing is acceptable.
    % This limitation exists due to the implementation of the multiple
    % dosings for simulation using events to update the parameters that
    % perform the multiple inputs. (only if multiple inputs)
    if length(input.time) > 1,
        % Determine the minimum time between dosing instances
        minDeltaT = min(input.time(2:end)-input.time(1:end-1));
        if input.Tlag + 0.0001 >= minDeltaT,
            error('Tlag in ''%s'' is to large. Please choose it smaller than the min time between dosings.',input.name);
        end
    end    
return

%%%%%%%%%%%%%%%%%%%%%%%%%%%%%%%%%%%%%%%%%%%%%%%
% Check 0th order Absorption inputs
%%%%%%%%%%%%%%%%%%%%%%%%%%%%%%%%%%%%%%%%%%%%%%%
% We need to check that multiple 0 absorptions do not lead to one
% absorption overlaying the previous one(s). This is done by checking that 
% Tk0 < min delta time in application
function [] = checkAbsorption0Input(input)
    % Tk0 is required as additional input.
    if length(input.parameters) ~= 1,
        error('0th order Absorption definition does require a single additional parameter "Tk0" in input "%s".',input.name);
    end
    % Check name of parameter (required: Tk0)
    if ~strcmp(input.parameters(1).name,'Tk0'),
        error('Please use "Tk0" as absorption time parameter in input "%s".',input.name);
    end
    % Check non overlaying absorption0 inputs (only if multiple inputs)
    if length(input.time) > 1,
        % Determine minimum required time between dosing instances
        minRequiredDeltaT = max(input.parameters(1).value);  % corresponds to Tk0
        % Determine the minimum time between dosing instances
        minDeltaT = min(input.time(2:end)-input.time(1:end-1));
        % Do the check
        if minRequiredDeltaT > minDeltaT,
            error('Absorption time Tk0 for input ''%s'' to large. Previous dosing would not be finished before the next one.',input.name);
        end
        % Do check that Tlag is not to large. Tlag+(Tk0 or Dmax/Rate or deltaTBolus) < min delta T dosing is acceptable.
        % This limitation exists due to the implementation of the multiple
        % dosings for simulation using events to update the parameters that
        % perform the multiple inputs. (only if multiple inputs)
        % Determine the minimum time between dosing instances
        if input.Tlag + minRequiredDeltaT >= minDeltaT,
            error('Tlag in ''%s'' is to large. Please choose it smaller than the min time between dosings minus Tk0.',input.name);
        end        
    end
return

%%%%%%%%%%%%%%%%%%%%%%%%%%%%%%%%%%%%%%%%%%%%%%%
% Remove character function
%%%%%%%%%%%%%%%%%%%%%%%%%%%%%%%%%%%%%%%%%%%%%%%
function [output] = removeCharacters(input)
% delete all line breaks and tabs from the input string
temp = double(input);
temp(find(temp==13)) = 32;  % replace '\cr' by white space
temp(find(temp==10)) = 32;  % replace '\n' by white space
temp(find(temp==9)) = 32;   % replace '\t' by white space
output = char(temp);
% remove all spaces
%    output = strrep(output,' ','');
return
```
