## Supplementary scripts for "A multi-scale *in silico* mouse model for insulin resistance and humanoid type 2 diabetes": getPartsFromCompleteTextDosSBPOP.m

```
function [dosTextStructure] = getPartsFromCompleteTextDosSBPOP(dosText)
% getPartsFromCompleteTextDosSBPOP: Cuts a text description of an
% SBPOPdosing object into the different parts and returns them in a
% structure 

% Information:
% ============
% Copyright � 2012 Novartis Pharma AG
% 
% This program is Free Open Source Software: you can redistribute it and/or modify 
% it under the terms of the GNU General Public License as published by 
% the Free Software Foundation, either version 3 of the License, or 
% (at your option) any later version. 
% 
% This program is distributed in the hope that it will be useful, 
% but WITHOUT ANY WARRANTY; without even the implied warranty of 
% MERCHANTABILITY or FITNESS FOR A PARTICULAR PURPOSE. See the 
% GNU General Public License for more details. 
% 
% You should have received a copy of the GNU General Public License 
% along with this program. If not, see <http://www.gnu.org/licenses/>.

% take commented lines out of the dosing description
dosText = regexprep(dosText,'\n%[^\n]*','');

% Find the starts of the different sections. NAME and NOTES are required.
% Following these there can be an arbitrary number of input definitions. 
% The order of the inputs is defined through the naming of a section 
% ********** INPUT1  => corresponds to "INPUT1" in a model file.

% Required sections
nameStart = strfind(dosText,'********** DOSING NAME');
notesStart = strfind(dosText,'********** DOSING NOTES');

% Check if they are present
if isempty(nameStart),
    error('No "DOSING NAME" section in dosing description.');
end
if isempty(notesStart),
    error('No "DOSING NOTES" section in dosing description.');
end

% Parse the INPUTx sections. They are optional (No input is allowed).
% 1) Count all *****... etc.
nrAllSections = length(strfind(dosText,'**********'));
nrInputSections = length(strfind(dosText,'********** INPUT'));

% 2) Check the number of the sections (allows detecting syntax errors in
% input section definitions)
if nrInputSections+2 ~= nrAllSections,
    error('Please check the input section identifiers in the dose description file for errors.');
end

% 3) get startindices of input sections
startIndexInput = strfind(dosText,'********** INPUT');

% 4) Get all sections
dosTextStructure.name = strtrim(dosText(nameStart+length('********** DOSING NAME'):notesStart-1));
if isempty(startIndexInput),
    dosTextStructure.notes = strtrim(dosText(notesStart+length('********** DOSING NOTES'):end));
else
    dosTextStructure.notes = strtrim(dosText(notesStart+length('********** DOSING NOTES'):startIndexInput(1)-1));
end
dosTextStructure.input = {};
for k=1:length(startIndexInput)-1,
    dosTextStructure.input{k} = strtrim(dosText(startIndexInput(k)+length('********** '):startIndexInput(k+1)-1));
end
dosTextStructure.input{end+1} = strtrim(dosText(startIndexInput(end)+length('********** '):end));
return
```
