## Supplementary scripts for "A multi-scale *in silico* mouse model for insulin resistance and humanoid type 2 diabetes": SBPOPcreateDOSfile.m

```
function SBPOPcreateDOSfile(varargin)
% SBPOPcreateDOSfile: creates a *.dos file with the dosing text description
%
% USAGE:
% ======
% [] = SBPOPcreateDOSfile()         
% [] = SBPOPcreateDOSfile(filename)         
% [] = SBPOPcreateDOSfile(dos)         
% [] = SBPOPcreateDOSfile(dos,filename)
%
% dos: SBPOPdosing object to convert to a textfile description
% filename: filename for the created textfile 
%
% If dos is undefined, then an empty SBPOPdosing textfile will be created.
%
% DEFAULT VALUES:
% ===============
% dos: the SBPOPdosing object to be exported as DOS file. 
% filename: constructed from the SBPOPdosing object name

% Information:
% ============
% Copyright � 2012 Novartis Pharma AG
% 
% This program is Free Open Source Software: you can redistribute it and/or modify 
% it under the terms of the GNU General Public License as published by 
% the Free Software Foundation, either version 3 of the License, or 
% (at your option) any later version. 
% 
% This program is distributed in the hope that it will be useful, 
% but WITHOUT ANY WARRANTY; without even the implied warranty of 
% MERCHANTABILITY or FITNESS FOR A PARTICULAR PURPOSE. See the 
% GNU General Public License for more details. 
% 
% You should have received a copy of the GNU General Public License 
% along with this program. If not, see <http://www.gnu.org/licenses/>.

%%%%%%%%%%%%%%%%%%%%%%%%%%%%%%%%%%%%%%%%%%%%%%%
% PROCESS VARIABLE INPUT ARGUMENTS
%%%%%%%%%%%%%%%%%%%%%%%%%%%%%%%%%%%%%%%%%%%%%%%
if nargin == 0,
    % create empty SBPOPdosing file with the name "unnamed.dos"
    dos = SBPOPdosing();
    filename = 'unnamed.dos';
elseif nargin == 1,
    % check if first input argument dosing object or filename
    if isSBPOPdosing(varargin{1}),
        % dosing object given
        dos = varargin{1};
        % if no filename provided then use the name of the SBPOPdosing
        % object as filename but remove unwanted characters first
        ds = struct(dos);
        functionName = regexprep(ds.name,'\W','');
        filename = strcat(functionName,'.dos');
    else
        % filename given?
        if ~ischar(varargin{1}),
            error('Wrong input argument.');
        end
        filename = strcat(varargin{1},'.dos');
        dos = SBPOPdosing();
    end
elseif nargin == 2,
    dos = varargin{1};
    filename = strcat(varargin{2},'.dos');
else
    error('Incorrect number of input arguments.');
end

%%%%%%%%%%%%%%%%%%%%%%%%%%%%%%%%%%%%%%%%%%%%%%%
% CHECK INPUT
%%%%%%%%%%%%%%%%%%%%%%%%%%%%%%%%%%%%%%%%%%%%%%%
if ~isSBPOPdosing(dos),
    error('No SBPOPdosing object as first argument.');
end

%%%%%%%%%%%%%%%%%%%%%%%%%%%%%%%%%%%%%%%%%%%%%%%
% CONVERT DOSING TO TEXT
%%%%%%%%%%%%%%%%%%%%%%%%%%%%%%%%%%%%%%%%%%%%%%%
[dosTextStructure] = convertDosToTextSBPOP(dos);
[completeText] = setPartsToCompleteTextDosSBPOP(dosTextStructure);

%%%%%%%%%%%%%%%%%%%%%%%%%%%%%%%%%%%%%%%%%%%%%%%
% WRITE THE FILE
%%%%%%%%%%%%%%%%%%%%%%%%%%%%%%%%%%%%%%%%%%%%%%%
fid = fopen(filename,'w');
fwrite(fid,completeText);
fclose(fid);
return
```
