## Supplementary scripts for "A multi-scale *in silico* mouse model for insulin resistance and humanoid type 2 diabetes": SBPOPcreateDOSING.m

```
function [dosing] = SBPOPcreateDOSING(type,dose,time,parametervalue,varargin)
% SBPOPcreateDOSING: Creates a dosing scheme as desired based on inputs
%
% USAGE:
% ======
% [dosing] = SBPOPcreateDOSING(type,dose,time,parametervalue)         
% [dosing] = SBPOPcreateDOSING(type,dose,time,parametervalue,tlagvalue)         
%
% type:             cell-array with input types in the order as inputs will be defined
%                   possible types: "INFUSION", "ABSORPTION0", "ABSORPTION1", "BOLUS"
% dose:             cell-array with dose vectors for each defined input - if scalar then same dose will be used for each dosing time
% time:             cell-array with time vectors for each defined input
% parametervalue:   cell-array with TINF, ka, TK0 values - if scalar then same value assumed for each dosing time
% tlagvalue:        cell-array with lag time for each input definition
%
% dosing:           produced dosing scheme

% Information:
% ============
% Copyright � 2012 Novartis Pharma AG
% 
% This program is Free Open Source Software: you can redistribute it and/or modify 
% it under the terms of the GNU General Public License as published by 
% the Free Software Foundation, either version 3 of the License, or 
% (at your option) any later version. 
% 
% This program is distributed in the hope that it will be useful, 
% but WITHOUT ANY WARRANTY; without even the implied warranty of 
% MERCHANTABILITY or FITNESS FOR A PARTICULAR PURPOSE. See the 
% GNU General Public License for more details. 
% 
% You should have received a copy of the GNU General Public License 
% along with this program. If not, see <http://www.gnu.org/licenses/>.

%%%%%%%%%%%%%%%%%%%%%%%%%%%%%%%%%%%%%%%%%%%%%%%
% Create empty dosing scheme
%%%%%%%%%%%%%%%%%%%%%%%%%%%%%%%%%%%%%%%%%%%%%%%
ds = struct(SBPOPdosing);

%%%%%%%%%%%%%%%%%%%%%%%%%%%%%%%%%%%%%%%%%%%%%%%
% Variable input arguments
%%%%%%%%%%%%%%%%%%%%%%%%%%%%%%%%%%%%%%%%%%%%%%%
if nargin == 5,
    tlagvalue = varargin{1};
    if ~iscell(tlagvalue), tlagvalue = {tlagvalue}; end
else
    tlagvalue = [];
end

%%%%%%%%%%%%%%%%%%%%%%%%%%%%%%%%%%%%%%%%%%%%%%%
% Check inputs and convert to cells if needed
%%%%%%%%%%%%%%%%%%%%%%%%%%%%%%%%%%%%%%%%%%%%%%%
if ~iscell(type), type = {type}; end
if ~iscell(dose), dose = {dose}; end
if ~iscell(time), time = {time}; end
if ~iscell(parametervalue), parametervalue = {parametervalue}; end

%%%%%%%%%%%%%%%%%%%%%%%%%%%%%%%%%%%%%%%%%%%%%%%
% Update name fields
%%%%%%%%%%%%%%%%%%%%%%%%%%%%%%%%%%%%%%%%%%%%%%%
for k=1:length(type),
    ds.inputs(k).name = sprintf('INPUT%d',k);
end

%%%%%%%%%%%%%%%%%%%%%%%%%%%%%%%%%%%%%%%%%%%%%%%
% Update type fields
%%%%%%%%%%%%%%%%%%%%%%%%%%%%%%%%%%%%%%%%%%%%%%%
for k=1:length(type),
    ds.inputs(k).type = type{k};
end

%%%%%%%%%%%%%%%%%%%%%%%%%%%%%%%%%%%%%%%%%%%%%%%
% Update time fields
%%%%%%%%%%%%%%%%%%%%%%%%%%%%%%%%%%%%%%%%%%%%%%%
if length(time) ~= length(type),
    error('Number of entries in time cell-array different from number in type cell-array');
end
for k=1:length(time),
    ds.inputs(k).time = time{k};
end

%%%%%%%%%%%%%%%%%%%%%%%%%%%%%%%%%%%%%%%%%%%%%%%
% Update dose fields
%%%%%%%%%%%%%%%%%%%%%%%%%%%%%%%%%%%%%%%%%%%%%%%
if length(dose) ~= length(type),
    error('Number of entries in dose cell-array different from number in type cell-array');
end
for k=1:length(dose),
    dosek = dose{k};
    if length(dosek) == 1,
        ds.inputs(k).D = dosek*ones(1,length(ds.inputs(k).time));
    else
        if length(dosek) ~= length(ds.inputs(k).time),
            error('Different lengths of time and dose vectors for input %d',k);
        else
            ds.inputs(k).D = dosek;
        end
    end
end

%%%%%%%%%%%%%%%%%%%%%%%%%%%%%%%%%%%%%%%%%%%%%%%
% Update parameters fields
% Only handle if type not BOLUS
%%%%%%%%%%%%%%%%%%%%%%%%%%%%%%%%%%%%%%%%%%%%%%%
if length(parametervalue) ~= length(type),
    error('Number of entries in parametervalue cell-array different from number in type cell-array');
end
for k=1:length(type),
    if ~strcmp(type{k},'BOLUS'),
        pvk = parametervalue{k};
        if strcmp(type{k},'INFUSION'),
            ds.inputs(k).parameters.name = 'Tinf';
        elseif strcmp(type{k},'ABSORPTION1'),
            ds.inputs(k).parameters.name = 'ka';
        elseif strcmp(type{k},'ABSORPTION0'),
            ds.inputs(k).parameters.name = 'Tk0';
        end            
        ds.inputs(k).parameters.notes = '';
        
        if length(pvk) == 1,
            ds.inputs(k).parameters.value = pvk*ones(1,length(ds.inputs(k).time));
        else
            if length(pvk) ~= length(ds.inputs(k).time),
                error('Different lengths of parameter values and dose vectors for input %d',k);
            else
                ds.inputs(k).parameters.value = pvk;
            end
        end
    end
end

%%%%%%%%%%%%%%%%%%%%%%%%%%%%%%%%%%%%%%%%%%%%%%%
% Update Tlag fields
%%%%%%%%%%%%%%%%%%%%%%%%%%%%%%%%%%%%%%%%%%%%%%%
if ~isempty(tlagvalue),
    if length(tlagvalue) ~= length(type),
        error('Number of entries in tlagvalue cell-array different from number in type cell-array');
    end
    for k=1:length(tlagvalue),
        ds.inputs(k).Tlag = tlagvalue{k};
    end
end

%%%%%%%%%%%%%%%%%%%%%%%%%%%%%%%%%%%%%%%%%%%%%%%
% Create dosing scheme
%%%%%%%%%%%%%%%%%%%%%%%%%%%%%%%%%%%%%%%%%%%%%%%
dosing = SBPOPdosing(ds);
```
