## Supplementary scripts for "A multi-scale *in silico* mouse model for insulin resistance and humanoid type 2 diabetes": SBPOPdoseupdatevalue.m

```
function [dosingchanged] = SBPOPdoseupdatevalue(dosing,inputname,value)
% SBPOPdoseupdatevalue: Allows to update the dosing amount for a given
%   input, defined in an SBPOPdosing object.
%
% USAGE:
% ======
% [dosingchanged] = SBPOPdoseupdatevalue(dosing,inputname,value)         
%
% dosing: SBPOPdosing object 
% inputname: name of the input for which to change the dosing amount
% value: value to set the dosing amount to (only scalar value allowed, used
%   for all dosing instances in case of multiple doses)
%
% Output Arguments:
% =================
% dosingchanged: updated SBPOPdosing object with changed dosing amount

% Information:
% ============
% Copyright � 2012 Novartis Pharma AG
% 
% This program is Free Open Source Software: you can redistribute it and/or modify 
% it under the terms of the GNU General Public License as published by 
% the Free Software Foundation, either version 3 of the License, or 
% (at your option) any later version. 
% 
% This program is distributed in the hope that it will be useful, 
% but WITHOUT ANY WARRANTY; without even the implied warranty of 
% MERCHANTABILITY or FITNESS FOR A PARTICULAR PURPOSE. See the 
% GNU General Public License for more details. 
% 
% You should have received a copy of the GNU General Public License 
% along with this program. If not, see <http://www.gnu.org/licenses/>.

%%%%%%%%%%%%%%%%%%%%%%%%%%%%%%%%%%%%%%%%%%%%%%%
% Get dosing structure
%%%%%%%%%%%%%%%%%%%%%%%%%%%%%%%%%%%%%%%%%%%%%%%
ds = struct(dosing);

%%%%%%%%%%%%%%%%%%%%%%%%%%%%%%%%%%%%%%%%%%%%%%%
% Find input name
%%%%%%%%%%%%%%%%%%%%%%%%%%%%%%%%%%%%%%%%%%%%%%%
index = strmatchSB(inputname,{ds.inputs.name},'exact');

%%%%%%%%%%%%%%%%%%%%%%%%%%%%%%%%%%%%%%%%%%%%%%%
% Change Dosing amount if index found
%%%%%%%%%%%%%%%%%%%%%%%%%%%%%%%%%%%%%%%%%%%%%%%
if isempty(index),
    disp(['Input ' inputname ' does not exist in the dosing schedule.']);
else
    % update dosing amount with new value
    ds.inputs(index).D = value*ones(1,length(ds.inputs(index).time));
end

%%%%%%%%%%%%%%%%%%%%%%%%%%%%%%%%%%%%%%%%%%%%%%%
% DONE => Construct output
%%%%%%%%%%%%%%%%%%%%%%%%%%%%%%%%%%%%%%%%%%%%%%%
dosingchanged = SBPOPdosing(ds);
```
