## Supplementary scripts for "A multi-scale *in silico* mouse model for insulin resistance and humanoid type 2 diabetes": SBPOPmergemoddos.m

```
function [simmodel] = SBPOPmergemoddos(model,dosing)
% SBPOPmergemoddos: Based on a model and dosing object, a new SBmodel is
% generated, that implements the defined dosings. Multiple dosing schedules
% are realized by updating the parameters of subsequent dosings using
% events.
%
% This function is useful to simulate single dosing schedules. However, if
% you want to run trial simulations, which change the dosing amounts (or
% other things) between simulations, this function is not the best way to
% go, since you would need to apply these changes to the dosing schedule,
% run this function, recompile your model, etc.
%
% USAGE:
% ======
% [simmodel] = SBPOPmergemoddos(model,dosing) 
%
% model: SBmodel
% dosing: SBPOPdosing object
%
% Output Arguments:
% =================
% simmodel: Simulation model, implementing the dosing scheme, defined in
%   the model and the SBPOPdosing object.

% Information:
% ============
% Copyright � 2012 Novartis Pharma AG
% 
% This program is Free Open Source Software: you can redistribute it and/or modify 
% it under the terms of the GNU General Public License as published by 
% the Free Software Foundation, either version 3 of the License, or 
% (at your option) any later version. 
% 
% This program is distributed in the hope that it will be useful, 
% but WITHOUT ANY WARRANTY; without even the implied warranty of 
% MERCHANTABILITY or FITNESS FOR A PARTICULAR PURPOSE. See the 
% GNU General Public License for more details. 
% 
% You should have received a copy of the GNU General Public License 
% along with this program. If not, see <http://www.gnu.org/licenses/>.

%%%%%%%%%%%%%%%%%%%%%%%%%%%%%%%%%%%%%%%%%%%%%%%
% Check input
%%%%%%%%%%%%%%%%%%%%%%%%%%%%%%%%%%%%%%%%%%%%%%%
if ~isSBmodel(model),
    error('First input argument is not an SBmodel.');
end
if ~isSBPOPdosing(dosing),
    error('Second input argument is not an SBPOPdosing object.');
end

%%%%%%%%%%%%%%%%%%%%%%%%%%%%%%%%%%%%%%%%%%%%%%%
% Get model augmented with dosing related 
% components and the experiment description,
% implementing the dosing applications
%%%%%%%%%%%%%%%%%%%%%%%%%%%%%%%%%%%%%%%%%%%%%%%
[moddos,experiment] = mergemoddosSBPOP(model,dosing);

%%%%%%%%%%%%%%%%%%%%%%%%%%%%%%%%%%%%%%%%%%%%%%%
% Combine new model with experiment to get the
% simulation model
%%%%%%%%%%%%%%%%%%%%%%%%%%%%%%%%%%%%%%%%%%%%%%%
simmodel = SBmergemodexp(moddos,experiment);
```
