## Supplementary scripts for "A multi-scale *in silico* mouse model for insulin resistance and humanoid type 2 diabetes": SBPOPsimdosing.m

```
function [varargout] = SBPOPsimdosing(moddos,dosing,varargin)
% SBPOPsimdosing: Simulates the application of a dosing schedule to a model
% that has been prepared for it and either plots or returns the simulation
% results.
%
% USAGE:
% ======
% [output] = SBPOPsimdosing(moddos,dosing)         
% [output] = SBPOPsimdosing(moddos,dosing,time)         
% [output] = SBPOPsimdosing(moddos,dosing,time,ICs)         
% [output] = SBPOPsimdosing(moddos,dosing,time,ICs,paramnames,paramvalues)         
% [output] = SBPOPsimdosing(moddos,dosing,time,ICs,paramnames,paramvalues,OPTIONS)         
%
% moddos: SBmodel or MEXmodel which has been prepared using the
%   mergemoddosSBPOP function. Note that if you use here a model that does
%   not fit to the dosing input, the result will most certainly be wrong.
% dosing: SBPOPdosing object with which the moddos model has been prepared.
%   This input argument defines the dosing schedule to simulate
% time: timevector to simulate. If scalar than the value defines the end
%   simulation time and the time vector starts at 0. 
% ICs: vector with initial conditions
% paramnames: parameter name or cell-array with parameter-names 
%             Parameters which are passed to the function to be modified do have 
%             priority over parameters that are defined in the dosing scheme
%             (Tinf,ka, Tlag, etc.)
% paramvalues: vector with values for the parameters (in the same order as
%   paramnames)
% OPTIONS: CVODE integrator options
%
% DEFAULT VALUES:
% ===============
% time: If not defined or left empty ([]), then the timevector will be
%   chosen to cover the full dosing schedule +50%
%   If the only dosing event occurs at time = 0, then a default time of 20
%   is then used.
% ICs: [] (use initial conditions, stored in the model)
% paramnames: {}
% paramvalues: [] (use values stored in the model)
% OPTIONS: 
%   OPTIONs.method        = 'stiff'
%   OPTIONS.abstol        = 1e-6;
%   OPTIONS.reltol        = 1e-6;
%   OPTIONS.maxnumsteps   = 100000;
%
% Output Arguments:
% =================
% If an output argument is given, this variable will contain a standard
% simulation result structure containing the time and the state, variable,
% and reactions names and values. 
% If no output argument is given, the result is plotted using the SBplot
% function.

% Information:
% ============
% Copyright � 2012 Novartis Pharma AG
% 
% This program is Free Open Source Software: you can redistribute it and/or modify 
% it under the terms of the GNU General Public License as published by 
% the Free Software Foundation, either version 3 of the License, or 
% (at your option) any later version. 
% 
% This program is distributed in the hope that it will be useful, 
% but WITHOUT ANY WARRANTY; without even the implied warranty of 
% MERCHANTABILITY or FITNESS FOR A PARTICULAR PURPOSE. See the 
% GNU General Public License for more details. 
% 
% You should have received a copy of the GNU General Public License 
% along with this program. If not, see <http://www.gnu.org/licenses/>.

global doseeventstruct_sim

%%%%%%%%%%%%%%%%%%%%%%%%%%%%%%%%%%%%%%%%%%%%%%%
% SET DEFAULT VALUES
%%%%%%%%%%%%%%%%%%%%%%%%%%%%%%%%%%%%%%%%%%%%%%%
time = [];  % indicates to calculate timevector from dosing schedule
OPTIONS = [];
OPTIONS.method = 'stiff';
OPTIONS.abstol = 1e-6;
OPTIONS.reltol = 1e-6;
OPTIONS.maxnumstep = 100000;
paramnames = {};
paramvalues = [];
ICs = [];

%%%%%%%%%%%%%%%%%%%%%%%%%%%%%%%%%%%%%%%%%%%%%%%
% HANDLE VARIABLE INPUT ARGUMENTS
%%%%%%%%%%%%%%%%%%%%%%%%%%%%%%%%%%%%%%%%%%%%%%%
if nargin == 2,
elseif nargin == 3,
    time = varargin{1};
elseif nargin == 4,
    time = varargin{1};
    ICs = varargin{2};
elseif nargin == 6,
    time = varargin{1};
    ICs = varargin{2};
    paramnames = varargin{3};
    paramvalues = varargin{4};
elseif nargin == 7,
    time = varargin{1};
    ICs = varargin{2};
    paramnames = varargin{3};
    paramvalues = varargin{4};
    OPTIONS = varargin{5};
else
    error('Incorrect number of input arguments.');
end
% char => cell 
if ischar(paramnames),
    paramnames = {paramnames};
end

%%%%%%%%%%%%%%%%%%%%%%%%%%%%%%%%%%%%%%%%%%%%%%%
% CHECK IF SBmodel and create MEXmodel
%%%%%%%%%%%%%%%%%%%%%%%%%%%%%%%%%%%%%%%%%%%%%%%
if isSBmodel(moddos),
    [MEXmodel, MEXmodelfullpath] = makeTempMEXmodelSBPD(moddos);
else
    MEXmodel = moddos;
end

%%%%%%%%%%%%%%%%%%%%%%%%%%%%%%%%%%%%%%%%%%%%%%%
% Create doseeventstruct
%%%%%%%%%%%%%%%%%%%%%%%%%%%%%%%%%%%%%%%%%%%%%%%
doseeventstruct = dosing2doseeventSBPOP(dosing);

%%%%%%%%%%%%%%%%%%%%%%%%%%%%%%%%%%%%%%%%%%%%%%%
% Handle timevector definition
%%%%%%%%%%%%%%%%%%%%%%%%%%%%%%%%%%%%%%%%%%%%%%%
if isempty(time),
    time = round(1.5*doseeventstruct(end).time);
    % Could handle when the dosing event is at time 0
    % then set time to a default end time
    if time == 0
        time = 20;
    end
    timevector = [0:time/1000:time];    
elseif length(time) == 1,
    timevector = [0:time/1000:time];
else
    timevector = time;
end

%%%%%%%%%%%%%%%%%%%%%%%%%%%%%%%%%%%%%%%%%%%%%%%
% Handle the case when final simulation time is smaller than the last dosing
% times. Simply remove the corresponding entries from the doseeventstruct.
%%%%%%%%%%%%%%%%%%%%%%%%%%%%%%%%%%%%%%%%%%%%%%%
doseeventstruct_sim = doseeventstruct;
doseeventstruct_sim(find([doseeventstruct.time] > timevector(end))) = [];
if isempty(doseeventstruct_sim),
    error('Please use a final simulation time that is larger than the first dosing time.')
end 

%%%%%%%%%%%%%%%%%%%%%%%%%%%%%%%%%%%%%%%%%%%%%%%
% Prepare the simulation
%%%%%%%%%%%%%%%%%%%%%%%%%%%%%%%%%%%%%%%%%%%%%%%
% Split the time vector into pieces from 0 to start of dosing
% applications, during dosing applications and from last application to
% final time.
firstdosing = doseeventstruct_sim(1).time;
lastdosing = doseeventstruct_sim(end).time;
timevectorpre = timevector(find(timevector < firstdosing));
timevectorpost = timevector(find(timevector > lastdosing));
timevectorduring = setdiff(setdiff(timevector,timevectorpre),timevectorpost);
% Adjust time vectors to include dosing times. This leads to the first and
% last dosing times being included twice (need to handle this later).
if ~isempty(timevectorpre),
    % Add first dosing time only if pre time vector is not empty.
    timevectorpre = [timevectorpre(:); firstdosing];
end
if ~isempty(timevectorpost),
    % Add last dosing time only if post time vector is not empty.
    timevectorpost = [lastdosing; timevectorpost(:)];
end
timvectorduring = sort(unique([timevectorduring(:); [doseeventstruct_sim.time]']));

%%%%%%%%%%%%%%%%%%%%%%%%%%%%%%%%%%%%%%%%%%%%%%%
% Prepare the simulation output structure
%%%%%%%%%%%%%%%%%%%%%%%%%%%%%%%%%%%%%%%%%%%%%%%
simout = [];
simout.time = [];
simout.states = {};
simout.statevalues = [];
simout.variables = {};
simout.variablevalues = [];
simout.reactions = {};
simout.reactionvalues = [];

%%%%%%%%%%%%%%%%%%%%%%%%%%%%%%%%%%%%%%%%%%%%%%%
% Simulate the prepart (only if timevectorpre is not empty)
%%%%%%%%%%%%%%%%%%%%%%%%%%%%%%%%%%%%%%%%%%%%%%%
if ~isempty(timevectorpre),
    presim = SBPDsimulate(MEXmodel,timevectorpre,ICs,paramnames,paramvalues,OPTIONS);
    % add results to output structure (neglect last index, since it is also
    % available in the during timevector). But save last state as IC for
    % next stage simulation
    simout.time = [simout.time; presim.time(1:end-1)];
    simout.states = presim.states;
    simout.statevalues = [simout.statevalues; presim.statevalues(1:end-1,:)];
    simout.variables = presim.variables;
    simout.variablevalues = [simout.variablevalues; presim.variablevalues(1:end-1,:)];
    simout.reactions = presim.reactions;
    simout.reactionvalues = [simout.reactionvalues; presim.reactionvalues(1:end-1,:)];
    % Save info for next stage simulation
    ICnext_values = presim.statevalues(end,:);
else
    % No simulation of first part => define ICs as the ones given or stored
    % in the model
    ICnext_values = ICs;
end

%%%%%%%%%%%%%%%%%%%%%%%%%%%%%%%%%%%%%%%%%%%%%%%
% Simulate the part in which dosing applications occur. Here we need to
% simulate piecewise inbetween dosing times
%%%%%%%%%%%%%%%%%%%%%%%%%%%%%%%%%%%%%%%%%%%%%%%
for k=1:length([doseeventstruct_sim.time])-1,
    dosetime = doseeventstruct_sim(k).time;
    nextdosetime = doseeventstruct_sim(k+1).time;

    % Determine the changes to parameters to implement dosing and timing of doses
    paramchangevalues = [];
    paramchangenames = {};
    for k2 = 1:length(doseeventstruct_sim(k).input),
        paramchangevalues = [paramchangevalues(:)' doseeventstruct_sim(k).input(k2).parametervalues(:)' dosetime];
        paramchangenames = {paramchangenames{:} doseeventstruct_sim(k).input(k2).parameternames{:} doseeventstruct_sim(k).input(k2).timeparname};
    end
    % paramchangevalues and paramchangenames contains the values for dose, time, ka, Tinf, etc that are defined in the dosing scheme
    % we need to make sure that parameters that are defined in the model or in the parameters that are passed to the SBPOPsimdosing
    % function to have priority over the parameters defined in the dosing scheme.
    % This is done by checking if paramnames has same elements as paramchangenames and then remove these double definitions from the 
    % paramchangenames and paramchangevalues arrays.
    double_defined_parameters = intersect(paramchangenames,paramnames);
    for k2=1:length(double_defined_parameters),
        % Remove the double defined ones from the paramchangenames and values (parameters passed to the simulation function have priority over definition in 
        % dosing scheme).
        indexremove = strmatchSB(double_defined_parameters{k2},paramchangenames,'exact');
        paramchangevalues(indexremove) = [];
        paramchangenames(indexremove) = [];
        % Report what is done
%        fprintf('Parameter "%s" passed to SBPOPsimdosing function has priority over definition of same parameter in dosing scheme.',double_defined_parameters{k2});
    end

    % Combine parameter changes from dosing scheme and passed parameters to simulation function
    paramchangevalues = [paramchangevalues(:)' paramvalues(:)'];
    paramchangenames = {paramchangenames{:} paramnames{:}};
    % paramchangevalues = [paramvalues(:)' doseeventstruct_sim(k).input.parametervalues dosetime*ones(1,length(doseeventstruct_sim(k).input))];
    % paramchangenames = {paramnames{:} doseeventstruct_sim(k).input.parameternames{:} doseeventstruct_sim(k).input.timeparname};
    
    % Determine the timevector piece (from dosing time to next dosing time (wo application at the latter)
    timevectorpiece = timvectorduring;
    timevectorpiece(timevectorpiece<dosetime) = [];
    timevectorpiece(timevectorpiece>nextdosetime) = [];
    
    sim = SBPDsimulate(MEXmodel,timevectorpiece,ICnext_values,paramchangenames,paramchangevalues,OPTIONS);
    % add results to output structure (if timevectorpost not empty => 
    % neglect last index, since it is also available in the next timevector
    % piece. Do NOT neglect if timevectorpost is empty). In any case 
    % save last state as IC for next stage simulation
    if ~isempty(timevectorpost),
        simout.time = [simout.time; sim.time(1:end-1)];
        simout.statevalues = [simout.statevalues; sim.statevalues(1:end-1,:)];
        simout.variablevalues = [simout.variablevalues; sim.variablevalues(1:end-1,:)];
        simout.reactionvalues = [simout.reactionvalues; sim.reactionvalues(1:end-1,:)];
    else
        simout.time = [simout.time; sim.time(1:end)];
        simout.statevalues = [simout.statevalues; sim.statevalues(1:end,:)];
        simout.variablevalues = [simout.variablevalues; sim.variablevalues(1:end,:)];
        simout.reactionvalues = [simout.reactionvalues; sim.reactionvalues(1:end,:)];
    end
    % add them here also (since presim might not happen ifg dosing at 0).
    simout.states = sim.states;
    simout.variables = sim.variables;
    simout.reactions = sim.reactions;
    % Save info for next stage simulation
    ICnext_values = sim.statevalues(end,:);
end

%%%%%%%%%%%%%%%%%%%%%%%%%%%%%%%%%%%%%%%%%%%%%%%
% Simulate the postpart (only if timevectorpost is not empty)
% The postpart starts with the application of the LAST dosing amount
%%%%%%%%%%%%%%%%%%%%%%%%%%%%%%%%%%%%%%%%%%%%%%%
if ~isempty(timevectorpost),
    dosetime = doseeventstruct_sim(end).time;
    % Determine the changes to parameters to implement dosing and timing of doses
    
    paramchangevalues = [];
    paramchangenames = {};
    for k2 = 1:length(doseeventstruct_sim(end).input),
        paramchangevalues = [paramchangevalues(:)' doseeventstruct_sim(end).input(k2).parametervalues(:)' dosetime];
        paramchangenames = {paramchangenames{:} doseeventstruct_sim(end).input(k2).parameternames{:} doseeventstruct_sim(end).input(k2).timeparname};
    end
    % paramchangevalues and paramchangenames contains the values for dose, time, ka, Tinf, etc that are defined in the dosing scheme
    % we need to make sure that parameters that are defined in the model or in the parameters that are passed to the SBPOPsimdosing
    % function to have priority over the parameters defined in the dosing scheme.
    % This is done by checking if paramnames has same elements as paramchangenames and then remove these double definitions from the 
    % paramchangenames and paramchangevalues arrays.
    double_defined_parameters = intersect(paramchangenames,paramnames);
    for k2=1:length(double_defined_parameters),
        % Remove the double defined ones from the paramchangenames and values (parameters passed to the simulation function have priority over definition in 
        % dosing scheme).
        indexremove = strmatchSB(double_defined_parameters{k2},paramchangenames,'exact');
        paramchangevalues(indexremove) = [];
        paramchangenames(indexremove) = [];
        % Report what is done
%        fprintf('Parameter "%s" passed to SBPOPsimdosing function has priority over definition of same parameter in dosing scheme.',double_defined_parameters{k2});
    end
    
    paramchangevalues = [paramchangevalues(:)' paramvalues(:)'];
    paramchangenames = {paramchangenames{:} paramnames{:}};

    % Do simulation of last piece
    sim = SBPDsimulate(MEXmodel,timevectorpost,ICnext_values,paramchangenames,paramchangevalues,OPTIONS);
    % Collect simulation results
    simout.time = [simout.time; sim.time(1:end)];
    simout.statevalues = [simout.statevalues; sim.statevalues(1:end,:)];
    simout.variablevalues = [simout.variablevalues; sim.variablevalues(1:end,:)];
    simout.reactionvalues = [simout.reactionvalues; sim.reactionvalues(1:end,:)];
    % add them here also (since presim might not happen ifg dosing at 0).
    simout.states = sim.states;
    simout.variables = sim.variables;
    simout.reactions = sim.reactions;    
end

%%%%%%%%%%%%%%%%%%%%%%%%%%%%%%%%%%%%%%%%%%%%%%%
% Finally, do remove the timepoints that are dosing time points but not
% defined in the desired simulation time vector.
%%%%%%%%%%%%%%%%%%%%%%%%%%%%%%%%%%%%%%%%%%%%%%%
[values,removeindices] = setdiff(simout.time,timevector);
while ~isempty(removeindices),
    simout.time(removeindices) = [];
    simout.statevalues(removeindices,:) = [];
    if ~isempty(simout.variables),
        simout.variablevalues(removeindices,:) = [];
    end
    if ~isempty(simout.reactions),
        simout.reactionvalues(removeindices,:) = [];
    end
    [values,removeindices] = setdiff(simout.time,timevector);
end

%%%%%%%%%%%%%%%%%%%%%%%%%%%%%%%%%%%%%%%%%%%%%%%
% Really finally, do remove double definitions of
% time points, which can happen in the case that
% dosing and observation time points are the same
%%%%%%%%%%%%%%%%%%%%%%%%%%%%%%%%%%%%%%%%%%%%%%%
lengthOLD = Inf;
% seems that sometimes MATLABis not picking it up correctly, then 
% we need to redo the exercise :(
while length(simout.time) ~= lengthOLD,
    lengthOLD = length(simout.time);
    
    [B,I,J] = unique(simout.time);
    removeindices = setdiff(J,I);
    simout.time(removeindices) = [];
    simout.statevalues(removeindices,:) = [];
    if ~isempty(simout.variables),
        simout.variablevalues(removeindices,:) = [];
    end
    if ~isempty(simout.reactions),
        simout.reactionvalues(removeindices,:) = [];
    end
end

%%%%%%%%%%%%%%%%%%%%%%%%%%%%%%%%%%%%%%%%%%%%%%%
% Handle variable output arguments
%%%%%%%%%%%%%%%%%%%%%%%%%%%%%%%%%%%%%%%%%%%%%%%
output = simout;
if nargout == 1,
    varargout{1} = output;
elseif nargout == 0,
    % prepare data for plotting
    time = output.time;
    datanames = {};
    dataindex = 1;
    for k = 1:length(output.states),
        datanames{dataindex} = sprintf('%s (state)',output.states{k});
        dataindex = dataindex + 1;
    end
    for k = 1:length(output.variables),
        datanames{dataindex} = sprintf('%s (variable)',output.variables{k});
        dataindex = dataindex + 1;
    end
    for k = 1:length(output.reactions),
        datanames{dataindex} = sprintf('%s (reaction rate)',output.reactions{k});
        dataindex = dataindex + 1;
    end
    datavalues = [output.statevalues, output.variablevalues, output.reactionvalues];
    SBplot(createdatastructSBplotSB(time,datavalues,datanames));
else
    error('Incorrect number of input arguments.');
end

%%%%%%%%%%%%%%%%%%%%%%%%%%%%%%%%%%%%%%%%%%%%%%%
% Delete MEX file if created in this function
%%%%%%%%%%%%%%%%%%%%%%%%%%%%%%%%%%%%%%%%%%%%%%%
if isSBmodel(moddos),
    clear mex
    delete(MEXmodelfullpath);
end
```
