## Supplementary scripts for "A multi-scale *in silico* mouse model for insulin resistance and humanoid type 2 diabetes": Contents.m

```
% SBPOP 
% Version Rev1361 (R2013a) 03-Jul-2014
%
% Installation of the package
% ===========================
%   installSBPOP            - Installation script for the SBPOP package
%   lookforSBPOP            - Searches SBPOP functions for strings and
%                             opens the documents in which these strings
%                             appear 
%   newscriptSBPOP          - Function creating a template file for a new
%                             script 
%   newfunctionSBPOP        - Function creating a template file for a new
%                             function 
%   setseedSBPOP            - Set the default stream to defaultSeed
%
%
% Dosing schedule related functions
% =================================
%   SBPOPdosing             - Creates an SBPOPdosing object defining a
%                             dosing schedule
%   SBPOPstruct             - Returns the internal data structure of an
%                             SBPOPdosing object
%   isSBPOPdosing           - Checks if input argument is an SBPOPdosing
%                             object 
%   SBPOPcreateDOSfile      - Creates a *.dos file with the dosing text
%                             description 
%   SBPOPcreateDOSING       - Creating a desired SBPOPdosing scheme 
%   SBPOPdoseupdatevalue    - Allows to update the dosing amount for a
%                             given input, defined in an SBPOPdosing object
%   SBPOPsimulate           - Simulates a given SBmodel with a given
%                             SBPOPdosing scheme 
%   SBPOPmergemoddos        - Based on a model and dosing object, a new
%                             SBmodel is generated, that implements the
%                             defined dosings. Multiple dosing schedules
%                             are realized by updating the parameters of
%                             subsequent dosings using events. 
%   mergemoddosSBPOP        - This function takes an SBmodel and an
%                             SBPOPdosing scheme as inputs. It adds
%                             necessary elements to the model that are
%                             required to implement the different dosing
%                             inputs (bolus, infusion, absorption0,
%                             absorption1) for simulation purposes (e.g. by
%                             the SBPOPsimdosing function).
%   SBPOPsimdosing          - Simulates the application of a dosing
%                             schedule to a model that has been prepared
%                             for it (using the mergemoddosSBPOP function)
%                             and either plots or returns the simulation
%                             results.
%   getmoddosinputinfoSBPOP - Checks the availability of dosing input
%                             definitions used in the model in the dosing
%                             object and returns a structure similar to
%                             the "input" field structure of an SBmodel,
%                             augmented with the dosing information,
%                             defined in the SBPOPdosing object. 
%   dosing2doseeventSBPOP   - This function takes an SBPOP dosing object as
%                             input and returns a structure in which the
%                             dosing events, happening, are sorted
%                             according to the times at which they happen.
%                             Additionally, the output structure contains
%                             information about the dosing amount and the
%                             names of the parameters in a model to change
%                             in order to apply the dosing.
%   doseinputsSBPOP         - Extracts names and types of dosing inputs
%                             from a SBPOPdosing object
%
%%%%%%%%%%%%%%%%%%%%%%%%%%%%%%%%%%%%%%%%%%%%%%%%%%%%%%%%%%%%%%%%%%%%%%%%%%%
% TOOLS 
%%%%%%%%%%%%%%%%%%%%%%%%%%%%%%%%%%%%%%%%%%%%%%%%%%%%%%%%%%%%%%%%%%%%%%%%%%%
%
% General data file handling
% ==========================
% SBPOPloadCSVdataset 		- Loads a CSV dataset as a MATLAB dataset
% SBPOPloadNONCSVdataset 	- Loads a non-CSV dataset as a MATLAB dataset.
%                             Example: NONMEM and Monolix output tables 
% SBPOPexportCSVdataset 	- Exports a MATLAB dataset as a CSV file
% SBPOPdataset2wide         - Expands a dataset from a row based format to
%                             a column based format 
% SBPOPsas7bdat2csv         - Convert a sas7bdat file to a CSV file
%                             (Requires presence of SAS on your systems
%                             command line)
%
% Plotting Tools
% ==============
% SBPOPplotCovarianceCat    - This function plots the covariance
%                             relationship between a list of continuous
%                             variables and a list of categorical variabels 
% SBPOPplotfacetgrid        - This function allows to generate facet-grid
%                             plot with many different settings
% SBPOPplotfill             - Function allowing to plot shaded areas with
%                             transparency 
% SBPOPplotHistogram 		- This function plots histograms of the data
%                             provided, allows grouping and plotting
%                             several histograms on top of each other
% SBPOPplotpairwiseCorr     - Plots pairwise correlations of data
% SBPOPplotQQ 				- QQ plot for provided input data
% SBPOPplottrellis          - Nice function to do Trellis plots with many
%                             different settings
% SBPOPplotXY 	    		- This function plots Ydata vs. Xdata with many
%                             different settings
%
% Monolix related functions
% =========================
% SBPOPcreateMLXTRANfile 		- Create MLXTRAN structural model based on
%                                 SBmodel and SBPOPdosing scheme
% SBPOPcreateMONOLIXproject 	- Create Monolix project (includes structural
%                                 model creation) 
% SBPOPrunMONOLIXproject    	- Runs specified Monolix project
% SBPOPrunMONOLIXprojectFolder  - Runs all Monolix projects within a folder
% SBPOPsampleMONOLIXparam   	- Samples population and individual parameters
%                           	  from a Monolix fitting result. The resulting
%                           	  population and individual parameters can be
%                           	  used for model simulations
% SBPOPgetMonolixdataHeader     - Determines the data header for MONOLIX project generation
% isMONOLIXfitSBPOP         	- Checks if a NLME project is a Monolix one
% parseMONOLIXetasSBPOP     	- Parses the ETAs from a Monolix project
% parseMONOLIXpredictionsSBPOP  - Parses the predictions from a Monolix project
% parseMONOLIXindivparamSBPOP   - Parses the individual param from a Monolix project
%
% NONMEM related functions
% ========================
% SBPOPcreateNONMEMproject		- Creates a NONMEM project
% SBPOPrunNONMEMproject 		- Runs a NONMEM project
% SBPOPrunNONMEMprojectFolder   - Runs all NONMEM projects within a folder
% SBPOPreportNONMEMresults 		- Reports the results of a NONMEM run
% SBPOPplotConvergenceNONMEM 	- Plots the convergence trajectories
% SBPOPsampleNONMEMparam 		- Samples model parameters from a NONMEM project
% SBPOPgetNONMEMdataHeader 		- Determines the data header for NONMEM project generation
% isNONMEMfitSBPOP 				- Checks if a NLME project is a NONMEM one
% parseNONMEMetasSBPOP 			- Parses the ETAs from a NONMEM project
% parseNONMEMpredictionsSBPOP   - Parses the predictions from a NONMEM project
% parseNONMEMindivparamSBPOP    - Parses the individual param from a NONMEM project
%
% General NLME (MONOLIX and NONMEM) related functions
% ===================================================
% SBPOPsampleNLMEfitParam
%
%%%%%%%%%%%%%%%%%%%%%%%%%%%%%%%%%%%%%%%%%%%%%%%%%%%%%%%%%%%%%%%%%%%%%%%%%%%
% AUXILIARY FUNCTIONS
%%%%%%%%%%%%%%%%%%%%%%%%%%%%%%%%%%%%%%%%%%%%%%%%%%%%%%%%%%%%%%%%%%%%%%%%%%%
%
% Figure export tools
% ===================
% printFigureSBPOP          - Exports MATLAB figures as JPG, PNG, or PS
%                             files. For PS files it appends them in same
%                             file. 
% startNewPrintFigureSBPOP  - Removes PS file (windows) or PDF file (unix)
%                             if it exists (so new figures are not
%                             appended)  
% convert2pdfSBPOP          - Converts PS to PDF files on unix/linux
%
% Other
% =====
% getcolorsSBPOP 			- Function returns color settings and
%                             linestyles, etc. Useful for plots to be able
%                             to use better colors than the MATLAB default
% usernameSBPOP             - Get the name of the current user
%
%%%%%%%%%%%%%%%%%%%%%%%%%%%%%%%%%%%%%%%%%%%%%%%%%%%%%%%%%%%%%%%%%%%%%%%%%%%
% TASKS 
%%%%%%%%%%%%%%%%%%%%%%%%%%%%%%%%%%%%%%%%%%%%%%%%%%%%%%%%%%%%%%%%%%%%%%%%%%%
%
% Please note that the functions in the "tasks" folder within SBPOP assume
% a standard dataset format. Nothing really exciting but some standard is
% needed in order to simplify things.
%
%%%%%%%%%%%%%%%%%%%%%%%%%%%%%%%%%%%%%%%%%%%%%%%%%%%%%%%%%%%%%%%%%%%%%%%%%%%
% TASKS - popPKtoolbox
% Supporting both NONMEM and MONOLIX
%%%%%%%%%%%%%%%%%%%%%%%%%%%%%%%%%%%%%%%%%%%%%%%%%%%%%%%%%%%%%%%%%%%%%%%%%%%
% SBPOPcreatePopPKproject   - Set-up of a popPK workflow, automatically
%                             creating documented template scripts that can
%                             be used 
% SBPOPexplorePopPKdata     - Creates a wide range of data exploration
%                             plots specific for popPK analyses
% SBPOPcleanPopPKdata       - Supports simple data cleaning specific for
%                             popPK analyses
% SBPOPconvert2popPKdataset - Convert dataset to popPK dataset. Removing
%                             placebo patients, keeping only certain
%                             columns, keeping only dose and PK in the
%                             dataset
% SBPOPbuildPopPKModelSpace - Function allowing to generate a PK model
%                             subspace, running all the estimations,
%                             generating tables for comparison of models,
%                             and generating a wide range of fit analysis
%                             plots. For current limitations of possible
%                             models, please look at the help text to this
%                             function
% SBPOPcomparePopPKmodels   - Compares popPK model fits
% SBPOPcreatePopPKstratifiedVPC - Allows to generate a stratified VPC with 
% 								  even less input arguments, but limited to
% 							      popPK models built with the popPKPD toolbox 
% 								  in SBPOP
%
%%%%%%%%%%%%%%%%%%%%%%%%%%%%%%%%%%%%%%%%%%%%%%%%%%%%%%%%%%%%%%%%%%%%%%%%%%%
% TASKS - PDtoolbox
%%%%%%%%%%%%%%%%%%%%%%%%%%%%%%%%%%%%%%%%%%%%%%%%%%%%%%%%%%%%%%%%%%%%%%%%%%%
% SBPOPexplorePopPDdata     				- Creates a wide range of data exploration
%                             				  plots specific for popPD analyses
% SBPOPcleanPopPDdata       				- Supports simple data cleaning specific for
%                             				  popPD analyses
% SBPOPgraphicalExplorationContinuousPD		- Additional exploration of continuous PD data
% SBPOPgraphicalExplorationResponderRatePD  - Exploration of categorical PD data
% SBPOPconvert2popPDparametersDataset       - Convert dataset to popPD dataset, including individual 
%                                             PK parameters for simulation of the PK
%
%%%%%%%%%%%%%%%%%%%%%%%%%%%%%%%%%%%%%%%%%%%%%%%%%%%%%%%%%%%%%%%%%%%%%%%%%%%
% TASKS - General
%%%%%%%%%%%%%%%%%%%%%%%%%%%%%%%%%%%%%%%%%%%%%%%%%%%%%%%%%%%%%%%%%%%%%%%%%%%
% SBPOPcheckGeneralDataFormat 	 		- Checks if data in the general dataset format
% SBPOPconvertGeneralDataFormat         - Converts the general dataset format into an
% 										  augmented one 
% SBPOPcheckDataFormat      			- Checking the data format and reporting problems
% SBPOPexploreCovariateCorrelations     - Plots correlation information for covariates
% SBPOPexploreIndivData     			- This function allows to plot individual data
%                                     	  from a standardized PKPD dataset
% SBPOPexploreSummaryStats  			- This function produces summary statistics for
%                                     	  the provided dataset 
% SBPOPcleanImputeCovariates        	- Covariate imputation 
% SBPOPcleanRemoveFewObsSubjects    	- Removal of subjects with not more than Nobs observations
% SBPOPcleanRemovePlaceboSubjects   	- Removal of placebo subjects
% SBPOPcleanRemoveRecordsSUBJECTs   	- Removal of user defined subjects and records
% SBPOPhandleSameTimeObservationRecords - Auxiliary for VPC generation
% SBPOPfitanalysisETAvsCOV          	- Plots the individual variations over
%                                         covariates and categorical covariates
% SBPOPfitanalysisGOFplots          	- Produces several plots that can be
%                                         used for checking the goodness of fit
% SBPOPfitanalysisIndividualFits    	- Plots individual fits and population
%                                         prediction against observed data over
%                                         time 
% SBPOPfitanalysisOutlierDetection  	- Considers PWRES and searches for
%                                         outliers and displays info about them
% SBPOPfitanalysisRandomEffects     	- Plots information about the random
%                                         effects in different ways
% SBPOPfitanalysisProjectsFolderPlots 	- Creates fit analysis plots for all 
%                                         Monolix/NONMEM project folders in specified
% 										  folder. The underlying models do not 
% 										  need to contain the same parameters
% SBPOPfitanalysisProjectsFolderInfo  	- Creates model comparison tables for all 
%                                         Monolix/NONMEM project folders in specified
% 										  folder. The underlying models do not 
% 										  need to contain the same parameters
% SBPOPfitanalysisGeneralPlots  	 	- Plots fit analysis plots for a single specified 
% 								  	      Monolix/NONMEM project folder 
% SBPOPcovariateChangeAssessment    	- Assesses the changes that a
%                                     	  covariates introduces on the model
%                                         parameters, based on the contents of
%                                     	  the dataset (no uncertainty)
% SBPOPcovariateAssessmentUncertainty   - Assesses the changes that a
%                                     	  covariates introduces on the model
%                                     	  parameters, based on the contents of
%                                     	  the dataset (with uncertainty) 
% SBPOPcompareModels 					- This function allows to compare the same 
%							   	  		  structural model for different estimation 
%							   	  		  results from Monolix/NONMEM. 
% SBPOPassessInformationContent 		- This function allows to predict the 
% 							      	  	  information content in data of (a) future 
% 								  		  studies, given the planned dosing and 
% 								  		  observation schedule
% SBPOPcreateVPC 						- Generation and plotting of simple VPC
% SBPOPcreateStratifiedVPC      		- Allows to generate a stratified VPC
%
%%%%%%%%%%%%%%%%%%%%%%%%%%%%%%%%%%%%%%%%%%%%%%%%%%%%%%%%%%%%%%%%%%%%%%%%%%%
% TASKS - PDmedian
% Still undocumented :-)
%%%%%%%%%%%%%%%%%%%%%%%%%%%%%%%%%%%%%%%%%%%%%%%%%%%%%%%%%%%%%%%%%%%%%%%%%%%

% Information:
% ============
% Copyright � 2012 Novartis Pharma AG
% 
% This program is Free Open Source Software: you can redistribute it and/or modify 
% it under the terms of the GNU General Public License as published by 
% the Free Software Foundation, either version 3 of the License, or 
% (at your option) any later version. 
% 
% This program is distributed in the hope that it will be useful, 
% but WITHOUT ANY WARRANTY; without even the implied warranty of 
% MERCHANTABILITY or FITNESS FOR A PARTICULAR PURPOSE. See the 
% GNU General Public License for more details. 
% 
% You should have received a copy of the GNU General Public License 
% along with this program. If not, see <http://www.gnu.org/licenses/>.
```
