## Supplementary scripts for "A multi-scale *in silico* mouse model for insulin resistance and humanoid type 2 diabetes": installSBPOP.m

```
function [] = installSBPOP(varargin)
% installSBPOP
% Installation function for the complete SBPOP package. 
%
%       installSBPOP
%
% This adds the SBPOP package and all subdirectories to the
% MATLAB path. 

% Information:
% ============
% Copyright � 2012 Novartis Pharma AG
% 
% This program is Free Open Source Software: you can redistribute it and/or modify 
% it under the terms of the GNU General Public License as published by 
% the Free Software Foundation, either version 3 of the License, or 
% (at your option) any later version. 
% 
% This program is distributed in the hope that it will be useful, 
% but WITHOUT ANY WARRANTY; without even the implied warranty of 
% MERCHANTABILITY or FITNESS FOR A PARTICULAR PURPOSE. See the 
% GNU General Public License for more details. 
% 
% You should have received a copy of the GNU General Public License 
% along with this program. If not, see <http://www.gnu.org/licenses/>.

%%%%%%%%%%%%%%%%%%%%%%%%%%%%%%%%%%%%%%%%%%%%%%%
% Check that installSBPOP is started in the right folder
%%%%%%%%%%%%%%%%%%%%%%%%%%%%%%%%%%%%%%%%%%%%%%%
currentDir = pwd;
installSBPOPDir = fileparts(which('installSBPOP.m'));
if ~strcmp(currentDir,installSBPOPDir),
    error('Run the ''installSBPOP'' script from the folder where it is located.');
end

%%%%%%%%%%%%%%%%%%%%%%%%%%%%%%%%%%%%%%%%%%%%%%%
% Check that at least the SBTOOLBOX2 is installed 
%%%%%%%%%%%%%%%%%%%%%%%%%%%%%%%%%%%%%%%%%%%%%%%
SBT2present = ver('SBTOOLBOX2');
if isempty(SBT2present),
    error('Please install the SBTOOLBOX2 first.');
end

%%%%%%%%%%%%%%%%%%%%%%%%%%%%%%%%%%%%%%%%%%%%%%%
% Check that correct local path (network paths are not allowed)
%%%%%%%%%%%%%%%%%%%%%%%%%%%%%%%%%%%%%%%%%%%%%%%
if strcmp(currentDir(1:2),'\\'),
    error(sprintf('The installation can not be run from a network path (\\\\...).\nPlease run the installation from a local path.'));
end

%%%%%%%%%%%%%%%%%%%%%%%%%%%%%%%%%%%%%%%%%%%%%%%
% Check if SBPOP already installed
%%%%%%%%%%%%%%%%%%%%%%%%%%%%%%%%%%%%%%%%%%%%%%%
SBPOPver = ver('SBPOP');
if length(SBPOPver) >= 1,
    error('You seem to already have an installation of SBPOP. Please use "restoredefaultpath" before installing a different version.');
end

%%%%%%%%%%%%%%%%%%%%%%%%%%%%%%%%%%%%%%%%%%%%%%%
% Add SBPOP folder to path
%%%%%%%%%%%%%%%%%%%%%%%%%%%%%%%%%%%%%%%%%%%%%%%
addpath(genpath(pwd));

%%%%%%%%%%%%%%%%%%%%%%%%%%%%%%%%%%%%%%%%%%%%%%%
% Final installation information, version, etc.
%%%%%%%%%%%%%%%%%%%%%%%%%%%%%%%%%%%%%%%%%%%%%%%
disp(sprintf('SBPOP for SBTOOLBOX2 and SBPD'));
disp(sprintf(' '));
disp(sprintf('Copyright (c) 2013 Novartis Pharma AG'));
disp(sprintf(' '));
disp(sprintf('This program is Free Open Source Software: you can redistribute it and/or modify '));
disp(sprintf('it under the terms of the GNU General Public License as published by '));
disp(sprintf('the Free Software Foundation, either version 3 of the License, or '));
disp(sprintf('(at your option) any later version. '));
disp(sprintf(' '));
disp(sprintf('This program is distributed in the hope that it will be useful, '));
disp(sprintf('but WITHOUT ANY WARRANTY; without even the implied warranty of '));
disp(sprintf('MERCHANTABILITY or FITNESS FOR A PARTICULAR PURPOSE. See the '));
disp(sprintf('GNU General Public License for more details. '));
disp(sprintf(' '));
disp(sprintf('You should have received a copy of the GNU General Public License '));
disp(sprintf('along with this program. If not, see <http://www.gnu.org/licenses/>.'));
disp(sprintf(' '));
disp(sprintf(' '));
disp(sprintf(' '));
```
