## Supplementary scripts for "A multi-scale *in silico* mouse model for insulin resistance and humanoid type 2 diabetes": getdatatextstructSBPD.m

```
function text = getdatatextstructSBPD(root,roottext,text)
% getdatatextstructSBPD: converts a structure to a flat file

% Information:
% ============
% SBPD Package - Systems Biology Parameter Determination Package
% Copyright 2008 by Henning Schmidt,
academicWarningSBPD

% converting given root level of a structure into text
fn = fieldnames(root);
for k=1:length(fn),
    fv = getfield(root,fn{k});
    if isstruct(fv),
        text = getdatatextstructSBPD(fv,[roottext '.' fn{k}],text);
    else
        if isnumeric(fv),
            if length(fv) == 1,
                help = num2str(fv);
                text = sprintf('%s\n%s.%s = %s;',text,roottext,fn{k},help);
            else
                help = sprintf('%g, ',fv);
                text = sprintf('%s\n%s.%s = [%s];',text,roottext,fn{k},help(1:end-2));
            end
        end
        if ischar(fv),
            text = sprintf('%s\n%s.%s = ''%s'';',text,roottext,fn{k},fv);
        end
        if iscell(fv),
            help = '';
            for k2 = 1:length(fv),
                if ischar(fv{k2}),
                    help2 = ['''' fv{k2} ''''];
                elseif isnumeric(fv{k2}),
                    help2 = num2str(fv{k2});
                else
                    help2 = '';
                end
                help = sprintf('%s%s, ',help,help2);
            end
            text = sprintf('%s\n%s.%s = {%s};',text,roottext,fn{k},help(1:end-2));
        end
    end
end
return
```
