## Supplementary scripts for "A multi-scale *in silico* mouse model for insulin resistance and humanoid type 2 diabetes": helpparamictextSBPD.m

```
function [ictext,paramtext,paramlocaltext,maxlength] = helpparamictextSBPD(pgn,pglb,pgub,pln,pllb,plub,icn,iclb,icub)
% helpparamictextSBPD: help function to construct the text that is
% necessary to define parameters and initialconditions and associated high
% and low bounds

% Information:
% ============
% SBPD Package - Systems Biology Parameter Determination Package
% Copyright 2008 by Henning Schmidt,
academicWarningSBPD

% and lowbounds for the estimation
ictext = '';
paramtext = '';
paramlocaltext = '';
% get the formatting nice
maxlength = 0;
maxlength2 = 0;
for k=1:length(icn),
    l = length(icn{k});
    if l>maxlength,
        maxlength = l;
    end
    l = length(sprintf('%g',iclb(k)));
    if l>maxlength2,
        maxlength2 = l;
    end    
end
for k=1:length(pgn),
    l = length(pgn{k});
    if l>maxlength,
        maxlength = l;
    end
    l = length(sprintf('%g',pglb(k)));
    if l>maxlength2,
        maxlength2 = l;
    end    
end
for k=1:length(pln),
    l = length(pln{k});
    if l>maxlength,
        maxlength = l;
    end
    l = length(sprintf('%g',pllb(k)));
    if l>maxlength2,
        maxlength2 = l;
    end    
end
maxlength2 = max(maxlength2,length('Lower bounds'));
% states
ictext = '';
for k=1:length(icn),
    l = length(icn{k});
    add = maxlength - l;
    addText = char(32*ones(1,add));
    l = length(sprintf('%g',iclb(k)));
    add2 = maxlength2 - l;
    addText2 = char(32*ones(1,add2));
    text = sprintf('''%s''%s  %g%s  %g',icn{k},addText,iclb(k),addText2,icub(k));
    ictext = sprintf('%s%s\n',ictext,text);
end
% parameters global
paramtext = '';
for k=1:length(pgn),
    l = length(pgn{k});
    add = maxlength - l;
    addText = char(32*ones(1,add));
    l = length(sprintf('%g',pglb(k)));
    add2 = maxlength2 - l;
    addText2 = char(32*ones(1,add2));    
    text = sprintf('''%s''%s  %g%s  %g',pgn{k},addText,pglb(k),addText2,pgub(k));
    paramtext = sprintf('%s%s\n',paramtext,text);
end
% parameters local
paramlocaltext = '';
for k=1:length(pln),
    l = length(pln{k});
    add = maxlength - l;
    addText = char(32*ones(1,add));
    l = length(sprintf('%g',pllb(k)));
    add2 = maxlength2 - l;
    addText2 = char(32*ones(1,add2));    
    text = sprintf('''%s''%s  %g%s  %g',pln{k},addText,pllb(k),addText2,plub(k));
    paramlocaltext = sprintf('%s%s\n',paramlocaltext,text);
end
```
