## Supplementary scripts for "A multi-scale *in silico* mouse model for insulin resistance and humanoid type 2 diabetes": academicWarningSBPD.m

```
% Information:
% ============
% Copyright (C) 2005-2013 Henning Schmidt,
%
% This program is free software; you can redistribute it and/or modify
% it under the terms of the GNU General Public License as published by
% the Free Software Foundation; either version 2 of the License, or
% (at your option) any later version.
% 
% This program is distributed in the hope that it will be useful,
% but WITHOUT ANY WARRANTY; without even the implied warranty of
% MERCHANTABILITY or FITNESS FOR A PARTICULAR PURPOSE.  See the
% GNU General Public License for more details.
% 
% You should have received a copy of the GNU General Public License
% along with this program; if not, write to the Free Software
% Foundation, Inc., 59 Temple Place, Suite 330, Boston, MA  02111-1307  
% USA.

FLAGacademicWarning = 1;

if FLAGacademicWarning,
warning(sprintf('Function is located in the "_ACADEMIC" folder of the SBPD package.\n\nThis means:\n    - No unit tests have been developed for this function\n    - This function is not subject to validation requirements for use\n      in clinical drug development projects\n    - This function should NOT be used in clinical drug development projects\n      where validated software is required\n    - You can still use this function for academic or exploratory purposes\n\nIf you do not want to get this message everytime you run this function,\nplease set the flag in the "academicWarningSBPD.m" file from 1 to 0.\n\n'));
end
      
```
