## Supplementary scripts for "A multi-scale *in silico* mouse model for insulin resistance and humanoid type 2 diabetes": Contents.m

```
% SBPD
% Version Rev1361 (R2013a) 03-Jul-2014
%
% Installation of package
% =======================
%   installSBPD         - Installation script for the SBPD package. Just
%                         run it from within its folder
%   lookforSBPD         - Searches SBPD functions for strings and
%                         opens the documents in which these strings appear
%
% Project creation and handling
% =============================
%   SBPDproject             - Creating a new SBPDproject
%   SBPDgui                 - Graphical User Interface for parameter
%                             estimation on projects
%   SBPDstruct              - Returns the internal data structure of an SBPDproject
%   SBPDexportproject       - Export a project to a directory structure
%   SBPDsaveproject         - Save a project in a MAT file (extension: .sbp)
%   isSBPDproject           - Checks if the given argument is an SBPDproject
%   SBPDinfo                - Display information about the project
%   SBPDplotmeasurements    - Visualize all measurements in the project
%   SBPDgetmodel            - Get specified model from project
%   SBPDgetexperiment       - Get specified experiment from project
%   SBPDgetmeasurement      - Get specified measurement from project
%   SBPDcomparemeasurements - Compare measurements to model in project
%   SBPDupdatemodel         - Update or add a model in a project
%   SBPDupdateexperiment    - Update or add an experiment in a project
%   SBPDupdatemeasurement   - Update or add a measurement in a project
%   SBPDexportestimation    - Export estimation settings to a flat file
%
% MEX model creation and handling
% ===============================
%   SBPDmakeMEXmodel          - Creates an executable simulation MEX file 
%                                 for a specified SBmodel. The function can
%                                 also be used to create C model files (no
%                                 compilation)
%   makeTempMEXmodelSBPD      - Creates a temporary executable simulation MEX
%                                 file for a specified SBmodel
%   mexcompileSBPD         - Creates an executable simulation MEX file 
%                                 for an existing C model files. These files
%                                 might be written by hand or obtained by the 
%                                 SBPDmakeMEXmodel function
%   makeparamvecSBPD            - this function can be used to create a full
%                                 parameter vector for the given model,
%                                 where the given parameters are changed to
%                                 given values. All other parameters remain
%                                 unchanged
%   makeinicondvectorSBPD       - this function can be used to create a full
%                                 state initial condition vector for the
%                                 given model, where the given states are
%                                 changed to given values. All other states
%                                 remain unchanged
%   getparamindicesSBPD         - returns a vector containing the indices
%                                 of the given model parameters
%   getstateindicesSBPD         - returns a vector containing the indices
%                                 of the given model states
%
% Modelling
% =========
%   SBPDdirectoptss     - Tune a model to a desired steady-state by
%                         adjusting the velocity parameters of the reaction
%                         kinetics 
% 
% Additionally, the SBPD package features several inbuild kinetic rate
% laws, which are documented below. 
%
% Simulation & sensitivity analysis
% =================================
%   SBPDsimulate        - Wrapper function for simulation MEX files.
%                         Can also be used on SBmodels
%   SBPDsensitivity     - Function for computing sensitivities of states,
%                         variables, and reactions in the model with respect 
%                         to parameters 
%   SBPDinsilicoexp     - Performs an insilico experiment for given
%                         SBmodel and SBexperiment. Visualizes result
%                         or exports it to a SBmeasurement
%                         representation. Also Excel or CSV measurement
%                         files can directly be generated.
%   SBPDinsilicoexpproj - Performs an in-silico experiment directly on a
%                         model and experiment within a project. Per
%                         default a CSV measurement files is created.
%
% Parameter identifiability
% =========================
%   SBPDidentifiability       - Function for determining parameter
%                               identifiability based on correlation
%                               analysis of sensitivity trajectories.
%                               Directly applicable to SBPDprojects.   
%   SBPDparametercorrelation  - Function for determining parameter
%                               correlations based on parametric
%                               sensitivities. 
%
% Model reduction (requires the symbolic toolbox)
% ===============================================
%   SBPDreducerateexpression    - Reduce the reaction rate expression of a
%                                 model, defined in a project
%
% Parameter estimation
% ====================
% SBPDmanualtuning          - Graphical user interface allowing to manually
%                             tune the models present in a project
% SBPDmodeltuning           - Allows to compare and tune a model to one or
%                             more sets of data. No experiment description
%                             is required 
% SBPDparameterestimation   - Allows to estimate parameters for the models 
%                             in a project, using the experiments and
%                             measurements that are defined in the project
% getparamictextSBPD        - Auxiliary function to easily construct the
%                             text that is necessary to define the
%                             parameters and initial conditions to estimate
%                             + their upper and lower bounds
% createrunestimationscriptSBPD - Auxiliary function creating a template
%                             m-script for running parameter estimations on
%                             a given project 
% SBPDanalyzeresiduals      - Determines and analyzes the residuals for a 
%                             given project
% SBPDparameterfitanalysis  - Generates data to analyze the obtained
%                             parameter fit with respect to correlations, 
%                             local minima, etc.
% SBPDfahist                - Plots histograms for the parameter values that 
%                             have been estimated using the SBPDparameterfitanalysis 
%                             function 
% SBPDfaclustering          - Performs hierarchical clustering based on
%                             Euclidean distance of the parameter sets
%                             estimated using the SBPDparameterfitanalysis
%                             function
% SBPDfaboxplot             - Plots a box-and-whisker diagram for the
%                             estimation data, obtained using the
%                             SBPDparameterfitanalysis function 
% SBPDfacorr                - Determines the correlation matrix for the
%                             parameter sets determined with the
%                             SBPDparameterfitanalysis function 
% SBPDfasigncorr            - Determines a matrix of p-values for testing
%                             the hypothesis of no significant correlation
%                             based on the results generated by
%                             SBPDparameterfitanalysis 
% SBPDfadetcorr            - Plots detailed pairwise correlations between
%                             parameters.
% cutoffdataSBPD           - Used to select a cut-off threshold for the
%                             estimation data collected during the fit
%                             analysis
% 
% Symbolic Math Functions (require the symbolic toolbox)
% ======================================================
%   SBPDsymjacobian         - determines the Jacobian of an SBmodel
%                             symbolically
%
% Inbuild kinetic rate laws
% =========================
%   kin_allosteric_inihib_empirical_rev  - see function documentation
%   kin_allosteric_inihib_mwc_irr        - see function documentation 
%   kin_catalytic_activation_irr         - see function documentation
%   kin_catalytic_activation_rev         - see function documentation
%   kin_comp_inihib_irr                  - see function documentation
%   kin_comp_inihib_rev                  - see function documentation 
%   kin_constantflux                     - see function documentation 
%   kin_degradation                      - see function documentation 
%   kin_hill_1_modifier_rev              - see function documentation 
%   kin_hill_2_modifiers_rev             - see function documentation 
%   kin_hill_cooperativity_irr           - see function documentation 
%   kin_hill_rev                         - see function documentation 
%   kin_hyperbolic_modifier_irr          - see function documentation 
%   kin_hyperbolic_modifier_rev          - see function documentation 
%   kin_iso_uni_uni_rev                  - see function documentation 
%   kin_mass_action_irr                  - see function documentation 
%   kin_mass_action_rev                  - see function documentation 
%   kin_michaelis_menten_irr             - see function documentation 
%   kin_michaelis_menten_rev             - see function documentation 
%   kin_mixed_activation_irr             - see function documentation 
%   kin_mixed_activation_rev             - see function documentation 
%   kin_mixed_inihib_irr                 - see function documentation 
%   kin_mixed_inihib_rev                 - see function documentation 
%   kin_noncomp_inihib_irr               - see function documentation 
%   kin_noncomp_inihib_rev               - see function documentation 
%   kin_ordered_bi_bi_rev                - see function documentation 
%   kin_ordered_bi_uni_rev               - see function documentation 
%   kin_ordered_uni_bi_rev               - see function documentation 
%   kin_ping_pong_bi_bi_rev              - see function documentation 
%   kin_specific_activation_irr          - see function documentation 
%   kin_specific_activation_rev          - see function documentation 
%   kin_substrate_activation_irr         - see function documentation 
%   kin_substrate_inihib_irr             - see function documentation 
%   kin_substrate_inihib_rev             - see function documentation 
%   kin_uncomp_inihib_irr                - see function documentation 
%   kin_uncomp_inihib_rev                - see function documentation 
%   kin_uni_uni_rev                      - see function documentation 
%
% Parameter Estimation Benchmarks
% ===============================
% The SBPD package includes several SBPDprojects that can be used as
% benchmarks for the evaluation of parameter estimation methods.
% These projects are located in the SBPD/examples/benchmarkproblems
% folder and can be run using the following scripts:
%
%   run_ex1         - Isomerization of alpha-pinene
%   run_ex2         - Irreversible inhibition of HIV proteinase
%   run_ex3         - Three-step biochemical pathway
%
% To execute these scripts, please change into the
% SBPD/examples/benchmarkproblems folder, open the desired script and 
% run it step by step. (Here the Cell-mode of the MATLAB editor is very
% useful!).

% Information:
% ============
% Copyright (C) 2005-2013 Henning Schmidt,
%
% This program is free software; you can redistribute it and/or modify
% it under the terms of the GNU General Public License as published by
% the Free Software Foundation; either version 2 of the License, or
% (at your option) any later version.
% 
% This program is distributed in the hope that it will be useful,
% but WITHOUT ANY WARRANTY; without even the implied warranty of
% MERCHANTABILITY or FITNESS FOR A PARTICULAR PURPOSE.  See the
% GNU General Public License for more details.
% 
% You should have received a copy of the GNU General Public License
% along with this program; if not, write to the Free Software
% Foundation, Inc., 59 Temple Place, Suite 330, Boston, MA  02111-1307  
% USA.
```
