## Supplementary scripts for "A multi-scale *in silico* mouse model for insulin resistance and humanoid type 2 diabetes": installSBPD.m

```
function [] = installSBPD()
% installSBPD
% Installation function for the SBPD package. 
%
%       installSBPD
%
% This adds the SBPD and all subdirectories to the MATLAB path.
% Additionally, the command builds the necessary libraries.

% Information:
% ============
% Copyright (C) 2005-2013 Henning Schmidt,
%
% This program is free software; you can redistribute it and/or modify
% it under the terms of the GNU General Public License as published by
% the Free Software Foundation; either version 2 of the License, or
% (at your option) any later version.
% 
% This program is distributed in the hope that it will be useful,
% but WITHOUT ANY WARRANTY; without even the implied warranty of
% MERCHANTABILITY or FITNESS FOR A PARTICULAR PURPOSE.  See the
% GNU General Public License for more details.
% 
% You should have received a copy of the GNU General Public License
% along with this program; if not, write to the Free Software
% Foundation, Inc., 59 Temple Place, Suite 330, Boston, MA  02111-1307  
% USA.

%%%%%%%%%%%%%%%%%%%%%%%%%%%%%%%%%%%%%%%%%%%%%%%
% Check that installSBPD is started in the right folder
%%%%%%%%%%%%%%%%%%%%%%%%%%%%%%%%%%%%%%%%%%%%%%%
currentDir = pwd;
installSBPDDir = fileparts(which('installSBPD.m'));
if ~strcmp(currentDir,installSBPDDir),
    error('Run the ''installSBPD'' script from the folder where it is located.');
end

%%%%%%%%%%%%%%%%%%%%%%%%%%%%%%%%%%%%%%%%%%%%%%%
% Check that correct local path (network paths are not allowed)
%%%%%%%%%%%%%%%%%%%%%%%%%%%%%%%%%%%%%%%%%%%%%%%
if strcmp(currentDir(1:2),'\\'),
    error(sprintf('The installation can not be run from a network path (\\\\...).\nPlease run the installation from a local path.'));
end

%%%%%%%%%%%%%%%%%%%%%%%%%%%%%%%%%%%%%%%%%%%%%%%
% Check if SBPD already installed
%%%%%%%%%%%%%%%%%%%%%%%%%%%%%%%%%%%%%%%%%%%%%%%
SBPDver = ver('SBPD');
if length(SBPDver) >= 1,
    error('You seem to already have an installation of SBPD. Please use "restoredefaultpath" before installing a different version.');
end

%%%%%%%%%%%%%%%%%%%%%%%%%%%%%%%%%%%%%%%%%%%%%%%
% Add SBPD to the path 
%%%%%%%%%%%%%%%%%%%%%%%%%%%%%%%%%%%%%%%%%%%%%%%
addpath(genpath(pwd));

%%%%%%%%%%%%%%%%%%%%%%%%%%%%%%%%%%%%%%%%%%%%%%%
% Compile and install the needed packages 
% Compilation is done for Unix AND Windows systems
%%%%%%%%%%%%%%%%%%%%%%%%%%%%%%%%%%%%%%%%%%%%%%%
PATH_SBPD = pwd();
if ~ispc,
    cd(fileparts(which('buildCVODElibunix.m')));
    buildCVODElibunix
    cd(PATH_SBPD);
else
    cd(fileparts(which('buildCVODElib.m')));
    buildCVODElib
    cd(PATH_SBPD);
end

%%%%%%%%%%%%%%%%%%%%%%%%%%%%%%%%%%%%%%%%%%%%%%%
% Output license information
%%%%%%%%%%%%%%%%%%%%%%%%%%%%%%%%%%%%%%%%%%%%%%%
disp(sprintf('SBPD for SBTOOLBOX2'));
disp('Copyright (C) 2008-2013 Henning Schmidt,');
disp(' ');
disp(sprintf('This program is Free Open Source Software: you can redistribute it and/or modify '));
disp(sprintf('it under the terms of the GNU General Public License as published by '));
disp(sprintf('the Free Software Foundation, either version 3 of the License, or '));
disp(sprintf('(at your option) any later version. '));
disp(sprintf(' '));
disp(sprintf('This program is distributed in the hope that it will be useful, '));
disp(sprintf('but WITHOUT ANY WARRANTY; without even the implied warranty of '));
disp(sprintf('MERCHANTABILITY or FITNESS FOR A PARTICULAR PURPOSE. See the '));
disp(sprintf('GNU General Public License for more details. '));
disp(sprintf(' '));
disp(sprintf('You should have received a copy of the GNU General Public License '));
disp(sprintf('along with this program. If not, see <http://www.gnu.org/licenses/>.'));
disp(sprintf(' '));
disp(sprintf(' '));
disp(sprintf(' '));
```
