## Supplementary scripts for "A multi-scale *in silico* mouse model for insulin resistance and humanoid type 2 diabetes": allkineticlaws.m

```
allLawsNames = {    
    'kin_allosteric_inihib_empirical_rev'
    'kin_allosteric_inihib_mwc_irr'
    'kin_catalytic_activation_irr'
    'kin_catalytic_activation_rev'
    'kin_comp_inihib_irr'
    'kin_comp_inihib_rev'
    'kin_constantflux'
    'kin_degradation'
    'kin_hill_1_modifier_rev'
    'kin_hill_2_modifiers_rev'
    'kin_hill_cooperativity_irr'
    'kin_hill_rev'
    'kin_hyperbolic_modifier_irr'
    'kin_hyperbolic_modifier_rev'
    'kin_iso_uni_uni_rev'
    'kin_mass_action_irr'
    'kin_mass_action_rev'
    'kin_michaelis_menten_irr'
    'kin_michaelis_menten_rev'
    'kin_mixed_activation_irr'
    'kin_mixed_activation_rev'
    'kin_mixed_inihib_irr'
    'kin_mixed_inihib_rev'
    'kin_noncomp_inihib_irr'
    'kin_noncomp_inihib_rev'
    'kin_ordered_bi_bi_rev'
    'kin_ordered_bi_uni_rev'
    'kin_ordered_uni_bi_rev'
    'kin_ping_pong_bi_bi_rev'
    'kin_specific_activation_irr'
    'kin_specific_activation_rev'
    'kin_substrate_activation_irr'
    'kin_substrate_inihib_irr'
    'kin_substrate_inihib_rev'
    'kin_uncomp_inihib_irr'
    'kin_uncomp_inihib_rev'
    'kin_uni_uni_rev'
};

allLawsArguments = {
{'Vf','substrate','Kms','Vr','product','Kmp','inhibitor','Ki','n'}
{'V','substrate','Ks','n','L','inhibitor','Ki'}
{'V','substrate','activator','Kms','Ka'}
{'Vf','substrate','Kms','Vr','product','Kmp','activator','Ka'}
{'V','substrate','Km','inhibitor','Ki'}
{'Vf','substrate','Kms','Vr','product','Kmp','inhibitor','Ki'}
{'v'}
{'kdeg','substrate'}
{'Vf','substrate','Shalve','product','Keq','Phalve','h','modifier','Mhalve','alpha'}
{'Vf','substrate','Shalve','product','Keq','Phalve','h','modifierA','MAhalve','modifierB','MBhalve','alphaA','alphaB','alphaAB'}
{'V','substrate','h','Shalve'}
{'Vf','substrate','Shalve','product','Keq','Phalve','h'}
{'V','substrate','b','modifier','a','Kd','Km'}
{'Vf','substrate','Kms','Vr','product','Kmp','b','modifier','a','Kd'}
{'Vf','substrate','product','Keq','Kii','Kms','Kmp'}
{'k','substrate'}
{'k1','substrate','k2','product'}
{'V','substrate','Km'}
{'Vf','substrate','Kms','Vr','product','Kmp'}
{'V','substrate','activator','Kms','Kas','Kac'}
{'Vf','substrate','Kms','Vr','product','Kmp','activator','Kas','Kac'}
{'V','substrate','Km','inhibitor','Kis','Kic'}
{'Vf','substrate','Kms','Vr','product','Kmp','inhibitor','Kis','Kic'}
{'V','substrate','Km','inhibitor','Ki'}
{'Vf','substrate','Kms','Vr','product','Kmp','inhibitor','Ki'}
{'Vf','substratea','substrateb','productp','productq','Keq','Kip','Kma','Kmb','Kia','Vr','Kmq','Kmp','Kib'}
{'Vf','substratea','substrateb','product','Keq','Kma','Kmb','Vr','Kmp','Kia'}
{'Vf','substrate','productp','productq','Keq','Kms','Kip','Vr','Kmq','Kmp'}
{'Vf','substratea','substrateb','productp','productq','Keq','Kiq','Kma','Kmb','Vr','Kmq','Kia','Kmp'}
{'V','substrate','activator','Kms','Ka'}
{'Vf','substrate','Kms','Vr','product','Kmp','activator','Ka'}
{'V','substrate','Ksa','Ksc'}
{'V','substrate','Km','Ki'}
{'Vf','substrate','Kms','Vr','product','Kmp','Ki'}
{'V','substrate','Km','inhibitor','Ki'}
{'Vf','substrate','Kms','Vr','product','Kmp','inhibitor','Ki'}
{'Vf','substrate','product','Keq','Kms','Kmp'}
};

allLawsFormulas = {
'((Vf*substrate/Kms - Vr*product/Kmp) / (1 + substrate/Kms + product/Kmp + (inhibitor/Ki)^n))'
'(V*substrate*(Ks+substrate)^(n-1) / ( L*(Ks*(1+inhibitor/Ki))^n + (Ks+substrate)^n ))'
'(V*substrate*activator / ( (Kms + substrate)*(Ka+activator) )'
'(( Vf*substrate/Kms - Vr*product/Kmp )*activator / ( (1+substrate/Kms+product/Kmp)*(Ka+activator) ))'
'(V*substrate / ( Km*(1+inhibitor/Ki) + substrate ))'
'((Vf*substrate/Kms - Vr*product/Kmp) / ( 1+substrate/Kms+product/Kmp+inhibitor/Ki ))'
'(v)'
'(kdeg*substrate)'
'(Vf*substrate/Shalve*(1-product/(substrate*Keq))*(substrate/Shalve+product/Phalve)^(h-1) / ( (1+(modifier/Mhalve)^h)/(1+alpha*(modifier/Mhalve)^h)+(substrate/Shalve + product/Phalve)^h ))'
'(Vf*substrate/Shalve*(1-product/(substrate*Keq))*(substrate/Shalve+product/Phalve)^(h-1) / ( (1+(modifierA/MAhalve)^h + 1+(modifierB/MBhalve)^h) / ( 1+alphaA*(modifierA/MAhalve)^h+alphaB*(modifierB/MBhalve)^h+alphaA*alphaB*alphaAB*(modifierA/MAhalve)^h*(modifierB/MBhalve)^h ) + (substrate/Shalve + product/Phalve)^h ))'
'(V*substrate^h / ( Shalve^h + substrate^h ))'
'(Vf*substrate/Shalve*(1-product/(substrate*Keq))*(substrate/Shalve+product/Phalve)^(h-1) / ( 1+(substrate/Shalve + product/Phalve)^h ))'
'(V*substrate*(1+b*modifier/(a*Kd)) / ( Km*(1+modifier/Kd) + substrate*(1+modifier/(a*Kd)) ))'
'((Vf*substrate/Kms - Vr*product/Kmp)*(1+b*modifier/(a*Kd)) / ( 1+modifier/Kd+(substrate/Kms+product/Kmp)*(1+modifier/(a*Kd)) ))'
'(Vf*(substrate-product/Keq) / ( substrate*(1+product/Kii) + Kms*(1+product/Kmp) ))'
'(k*substrate)'
'(k1*substrate-k2*product)'
'(V*substrate / ( Km + substrate ))'
'((Vf*substrate/Kms+Vr*product/Kmp) / ( 1+substrate/Kms+product/Kmp ) )'
'(V*substrate*activator / ( Kms*(Kas+activator) + substrate*(Kac+activator) ))'
'((Vf*substrate/Kms - Vr*product/Kmp)*activator / ( Kas+activator+(substrate/Kms+product/Kmp)*(Kac+activator) ))'
'(V*substrate / ( (Km*(1+inhibitor/Kis) + substrate*(1+inhibitor/Kic) ))'
'((Vf*substrate/Kms - Vr*product/Kmp) / ( 1+inhibitor/Kis+(substrate/Kms+product/Kmp)*(1+inhibitor/Kic) ))'
'(V*substrate / ( (Km+substrate)*(1+inhibitor/Ki) ))'
'((Vf*substrate/Kms-Vr*product/Kmp) / ( (1+substrate/Kms+product/Kmp)*(1+inhibitor/Ki) ))'
'(Vf*(substratea*substrateb-productp*productq/Keq) / (substratea*substrateb*(1+productp/Kip) + Kma*substrateb + Kmb*(substratea+Kia)+Vf/(Vr*Keq)*(Kmq*productp*(1+substratea/Kia) + productq*(Kmp*(1+Kia*substrateb/(Kma*Kmb))+productp*(1+substrateb/Kib))) ))'
'(Vf*(substratea*substrateb-product/Keq) / ( substratea*substrateb+Kma*substrateb+Kmb*substratea+Vf/(Vr*Keq)*(Kmp+product*(1+substratea/Kia)) ))'
'(Vf*(substrate-productp*productq/Keq) / ( Kms+substrate*(1+productp/Kip)+Vf/(Vr*Keq)*(Kmq*productp+Kmp*productq+productp*productq) );)'
'(Vf*( substratea*substrateb-productp*productq/Keq ) / (substratea*substrateb*(1+productq/Kiq)+Kma*substrateb+Kmb*substratea+Vf/(Vr*Keq)*(Kmq*productp*(1+substratea/Kia)+productq*(Kmp+productp))))'
'(V*substrate*activator/( Kms*Ka+(Kms+substrate)*activator )'
'(( Vf*substrate/Kms-Vr*product/Kmp )*activator / ( Ka+(1+substrate/Kms+product/Kmp)*activator ))'
'(V*(substrate/Ksa)^2 / ( 1+substrate/Ksc+substrate/Ksa+(substrate/Ksa)^2 ))'
'(V*substrate / ( Km + substrate + Km*(substrate/Ki)^2 ))'
'((Vf*substrate/Kms-Vr*product/Kmp) / ( 1+substrate/Kms+product/Kmp+(substrate/Ki)^2 ))'
'(V*substrate / ( Km + substrate*(1+inhibitor/Ki) ))'
'(( Vf*substrate/Kms-Vr*product/Kmp ) / ( 1+(substrate/Kms+product/Kmp)*(1+inhibitor/Ki) ))'
'(Vf*( substrate-product/Keq ) / ( substrate+Kms*(1+product/Kmp) ))'
};
```
