## Supplementary scripts for "A multi-scale *in silico* mouse model for insulin resistance and humanoid type 2 diabetes": getkinformulaSBPD.m

```
function [R] = getkinformulaSBPD(R)
% getkinformulaSBPD: auxiliary function to convert rate law functions to
% formulas.

% Information:
% ============
% SBPD Package - Systems Biology Parameter Determination Package
% Copyright 2008 by Henning Schmidt,

Rorig = R;
R = ['#' R '#'];
% Load names of all kinetic laws
allkineticlaws

% Check if kinetic rate expression present
found = 0;
for k=1:length(allLawsNames),
    if ~isempty(regexp(R,['\W' allLawsNames{k} '\W'])),
        % it is present
        % get the arguments of the reaction
        ratelawname = allLawsNames{k};
        indexstart = strfind(R,ratelawname);
        arguments = R(indexstart+length(ratelawname):end);
        parentheses = 0;
        for k2=1:length(arguments),
            if arguments(k2) == '(',
                parentheses = parentheses+1;
            end
            if arguments(k2) == ')',
                parentheses = parentheses-1;
                if parentheses == 0,
                    break;
                end
            end
        end
        arguments = arguments(2:k2-1);
        arguments = explodePCSB(arguments);
        preLaw = R(1:indexstart-1);
        postLaw = R(k2+indexstart+length(ratelawname):end);
        R = [preLaw(2:end) allLawsFormulas{k} postLaw(1:end-1)];
        % finally exchange the argument names
        changearguments = allLawsArguments{k};
        R = regexprep(R,'(\W)',' $1 ');
        for k2=1:length(arguments),
            R = regexprep(R,['\W' changearguments{k2} '\W'], arguments{k2});
        end
        R = regexprep(R,' ','');
        found = 1;
    end
end
if found == 0,
    R = Rorig;
end
```
