## Supplementary scripts for "A multi-scale *in silico* mouse model for insulin resistance and humanoid type 2 diabetes": kin_constantflux.m

```
function [R] = kin_constantflux(v)
% kin_constantflux: Competitive inhibition (reversible) kinetics
%
% Application restrictions: 
% =========================
%   - Be careful that concentrations do not become negative!
%
% USAGE:
% ======
% R = kin_constantflux(v)
%
% Output Arguments:
% =================
% R = v

% Information:
% ============
% SBPD Package - Systems Biology Parameter Determination Package
% Copyright 2008 by Henning Schmidt,

R = v;
```
