## Supplementary scripts for "A multi-scale *in silico* mouse model for insulin resistance and humanoid type 2 diabetes": kin_hill_cooperativity_irr.m

```
function [R] = kin_hill_cooperativity_irr(V,substrate,h,Shalve)
% kin_hill_cooperativity_irr: Hill type (irreversible) kinetics
%
% Application restrictions: 
% =========================
%   - Only irreversible
%   - exactly 1 substrate
%
% USAGE:
% ======
% R = kin_hill_cooperativity_irr(V,substrate,h,Shalve)
%
% Output Arguments:
% =================
% R = V*substrate^h / ( Shalve^h + substrate^h )

% Information:
% ============
% SBPD Package - Systems Biology Parameter Determination Package
% Copyright 2008 by Henning Schmidt,

R = V*substrate^h / ( Shalve^h + substrate^h );
```
