## Supplementary scripts for "A multi-scale *in silico* mouse model for insulin resistance and humanoid type 2 diabetes": kin_hill_rev.m

```
function [R] = kin_hill_rev(Vf,substrate,Shalve,product,Keq,Phalve,h)
% kin_hill_rev: Hill type (reversible) kinetics
%
% Application restrictions: 
% =========================
%   - Only reversible
%   - exactly 1 substrate
%   - exactly 1 product
%
% USAGE:
% ======
% R = kin_hill_rev(Vf,substrate,Shalve,product,Keq,Phalve,h)
%
% Output Arguments:
% =================
% R = Vf*substrate/Shalve*(1-product/(substrate*Keq))*(substrate/Shalve+product/Phalve)^(h-1) / ( 1+(substrate/Shalve + product/Phalve)^h ) 

% Information:
% ============
% SBPD Package - Systems Biology Parameter Determination Package
% Copyright 2008 by Henning Schmidt, 

R = Vf*substrate/Shalve*(1-product/(substrate*Keq))*(substrate/Shalve+product/Phalve)^(h-1) / ( 1+(substrate/Shalve + product/Phalve)^h );
```
