## Supplementary scripts for "A multi-scale *in silico* mouse model for insulin resistance and humanoid type 2 diabetes": kin_hyperbolic_modifier_rev.m

```
function [R] = kin_hyperbolic_modifier_rev(Vf,substrate,Kms,Vr,product,Kmp,b,modifier,a,Kd)
% kin_hyperbolic_modifier_rev: Hyperbolic modifier (reversible) kinetics
%
% Application restrictions: 
% =========================
%   - Only reversible
%   - exactly 1 substrate
%   - exactly 1 product
%
% USAGE:
% ======
% R = kin_hyperbolic_modifier_rev(Vf,substrate,Kms,Vr,product,Kmp,b,modifier,a,Kd)
%
% Output Arguments:
% =================
% R = (Vf*substrate/Kms - Vr*product/Kmp)*(1+b*modifier/(a*Kd)) / ( 1+modifier/Kd+(substrate/Kms+product/Kmp)*(1+modifier/(a*Kd) )

% Information:
% ============
% SBPD Package - Systems Biology Parameter Determination Package
% Copyright 2008 by Henning Schmidt,

R = (Vf*substrate/Kms - Vr*product/Kmp)*(1+b*modifier/(a*Kd)) / ( 1+modifier/Kd+(substrate/Kms+product/Kmp)*(1+modifier/(a*Kd)) );
```
