## Supplementary scripts for "A multi-scale *in silico* mouse model for insulin resistance and humanoid type 2 diabetes": kin_iso_uni_uni_rev.m

```
function [R] = kin_iso_uni_uni_rev(Vf,substrate,product,Keq,Kii,Kms,Kmp)
% kin_iso_uni_uni_rev: enzyme isomerization product inhibition
%
% Application restrictions: 
% =========================
%   - Only reversible
%   - exactly 1 substrate
%   - exactly 1 product
%
% USAGE:
% ======
% R = kin_iso_uni_uni_rev(Vf,substrate,product,Keq,Kii,Kms,Kmp)
%
% Output Arguments:
% =================
% R = Vf*(substrate-product/Keq) / ( substrate*(1+product/Kii) + Kms*(1+product/Kmp) )

% Information:
% ============
% SBPD Package - Systems Biology Parameter Determination Package
% Copyright 2008 by Henning Schmidt,

R = Vf*(substrate-product/Keq) / ( substrate*(1+product/Kii) + Kms*(1+product/Kmp) );
```
