## Supplementary scripts for "A multi-scale *in silico* mouse model for insulin resistance and humanoid type 2 diabetes": kin_mixed_activation_irr.m

```
function [R] = kin_mixed_activation_irr(V,substrate,activator,Kms,Kas,Kac)
% kin_mixed_activation_irr: Mixed activation irreversible
%
% Application restrictions: 
% =========================
%   - Only irreversible
%   - exactly 1 substrate
%
% USAGE:
% ======
% R = kin_mixed_activation_irr(V,substrate,activator,Kms,Kas,Kac)
%
% Output Arguments:
% =================
% R = V*substrate*activator / ( Kms*(Kas+activator) + substrate*(Kac+activator) )

% Information:
% ============
% SBPD Package - Systems Biology Parameter Determination Package
% Copyright 2008 by Henning Schmidt,

R = V*substrate*activator / ( Kms*(Kas+activator) + substrate*(Kac+activator) );
```
