## Supplementary scripts for "A multi-scale *in silico* mouse model for insulin resistance and humanoid type 2 diabetes": kin_noncomp_inihib_rev.m

```
function [R] = kin_noncomp_inihib_rev(Vf,substrate,Kms,Vr,product,Kmp,inhibitor,Ki)
% kin_noncomp_inihib_rev: Noncompetitive inhibition (reversible) kinetics
%
% Application restrictions: 
% =========================
%   - Only reversible
%   - exactly 1 substrate
%   - exactly 1 product
%
% USAGE:
% ======
% R = kin_noncomp_inihib_rev(Vf,substrate,Kms,Vr,product,Kmp,inhibitor,Ki)
%
% Output Arguments:
% =================
% R = (Vf*substrate/Kms-Vr*product/Kmp) / ( (1+substrate/Kms+product/Kmp)*(1+inhibitor/Ki) )

% Information:
% ============
% SBPD Package - Systems Biology Parameter Determination Package
% Copyright 2008 by Henning Schmidt,

R = (Vf*substrate/Kms-Vr*product/Kmp) / ( (1+substrate/Kms+product/Kmp)*(1+inhibitor/Ki) );
```
