## Supplementary scripts for "A multi-scale *in silico* mouse model for insulin resistance and humanoid type 2 diabetes": kin_ordered_bi_bi_rev.m

```
function [R] = kin_ordered_bi_bi_rev(Vf,substratea,substrateb,productp,productq,Keq,Kip,Kma,Kmb,Kia,Vr,Kmq,Kmp,Kib)
% kin_ordered_bi_bi_rev: ordered bi-bi reversible
%
% Application restrictions: 
% =========================
%   - Only reversible
%   - exactly 2 substrates
%   - exactly 2 products
%
% USAGE:
% ======
% R = kin_ordered_bi_bi_rev(Vf,substratea,substrateb,productp,productq,Keq,Kip,Kma,Kmb,Kia,Vr,Kmq,Kmp,Kib)
%
% Output Arguments:
% =================
% R = Vf*(substratea*substrateb-productp*productq/Keq) / (substratea*substrateb*(1+productp/Kip) + Kma*substrateb + Kmb*(substratea+Kia)+Vf/(Vr*Keq)*(Kmq*productp*(1+substratea/Kia) + productq*(Kmp*(1+Kia*substrateb/(Kma*Kmb))+productp*(1+substrateb/Kib))) )

% Information:
% ============
% SBPD Package - Systems Biology Parameter Determination Package
% Copyright 2008 by Henning Schmidt,

R = Vf*(substratea*substrateb-productp*productq/Keq) / (substratea*substrateb*(1+productp/Kip) + Kma*substrateb + Kmb*(substratea+Kia)+Vf/(Vr*Keq)*(Kmq*productp*(1+substratea/Kia) + productq*(Kmp*(1+Kia*substrateb/(Kma*Kmb))+productp*(1+substrateb/Kib))) );
```
