## Supplementary scripts for "A multi-scale *in silico* mouse model for insulin resistance and humanoid type 2 diabetes": kin_ordered_bi_uni_rev.m

```
function [R] = kin_ordered_bi_uni_rev(Vf,substratea,substrateb,product,Keq,Kma,Kmb,Vr,Kmp,Kia)
% kin_ordered_bi_uni_rev: ordered bi-uni reversible
%
% Application restrictions: 
% =========================
%   - Only reversible
%   - exactly 2 substrates
%   - exactly 1 product
%
% USAGE:
% ======
% R = kin_ordered_bi_uni_rev(Vf,substratea,substrateb,product,Keq,Kma,Kmb,Vr,Kmp,Kia)
%
% Output Arguments:
% =================
% R = Vf*(substratea*substrateb-product/Keq) / ( substratea*substrateb+Kma*substrateb+Kmb*substratea+Vf/(Vr*Keq)*(Kmp+product*(1+substratea/Kia)) ) 

% Information:
% ============
% SBPD Package - Systems Biology Parameter Determination Package
% Copyright 2008 by Henning Schmidt,

R = Vf*(substratea*substrateb-product/Keq) / ( substratea*substrateb+Kma*substrateb+Kmb*substratea+Vf/(Vr*Keq)*(Kmp+product*(1+substratea/Kia)) );
```
