## Supplementary scripts for "A multi-scale *in silico* mouse model for insulin resistance and humanoid type 2 diabetes": kin_ordered_uni_bi_rev.m

```
function [R] = kin_ordered_uni_bi_rev(Vf,substrate,productp,productq,Keq,Kms,Kip,Vr,Kmq,Kmp)
% kin_ordered_uni_bi_rev: ordered uni-bi reversible
%
% Application restrictions: 
% =========================
%   - Only reversible
%   - exactly 1 substrate
%   - exactly 2 products
%
% USAGE:
% ======
% R = kin_ordered_uni_bi_rev(Vf,substrate,productp,productq,Keq,Kms,Kip,Vr,Kmq,Kmp)
%
% Output Arguments:
% =================
% R = Vf*(substrate-productp*productq/Keq) / ( Kms+substrate*(1+productp/Kip)+Vf/(Vr*Keq)*(Kmq*productp+Kmp*productq+productp*productq) ) 

% Information:
% ============
% SBPD Package - Systems Biology Parameter Determination Package
% Copyright 2008 by Henning Schmidt,

R = Vf*(substrate-productp*productq/Keq) / ( Kms+substrate*(1+productp/Kip)+Vf/(Vr*Keq)*(Kmq*productp+Kmp*productq+productp*productq) ); 
```
