## Supplementary scripts for "A multi-scale *in silico* mouse model for insulin resistance and humanoid type 2 diabetes": kin_ping_pong_bi_bi_rev.m

```
function [R] = kin_ping_pong_bi_bi_rev(Vf,substratea,substrateb,productp,productq,Keq,Kiq,Kma,Kmb,Vr,Kmq,Kia,Kmp)
% kin_ping_pong_bi_bi_rev: Ping pong bi bi kinetics (reversible) 
%
% Application restrictions: 
% =========================
%   - Only reversible
%   - exactly 2 substrates
%   - exactly 2 products
%
% USAGE:
% ======
% R = kin_ping_pong_bi_bi_rev(Vf,substratea,substrateb,productp,productq,Keq,Kiq,Kma,Kmb,Vr,Kmq,Kia,Kmp)
%
% Output Arguments:
% =================
% R = Vf*( substratea*substrateb-productp*productq/Keq ) / (substratea*substrateb*(1+productq/Kiq)+Kma*substrateb+Kmb*substratea+Vf/(Vr*Keq)*(Kmq*productp*(1+substratea/Kia)+productq*(Kmp+productp)))

% Information:
% ============
% SBPD Package - Systems Biology Parameter Determination Package
% Copyright 2008 by Henning Schmidt,

R = Vf*( substratea*substrateb-productp*productq/Keq ) / (substratea*substrateb*(1+productq/Kiq)+Kma*substrateb+Kmb*substratea+Vf/(Vr*Keq)*(Kmq*productp*(1+substratea/Kia)+productq*(Kmp+productp)));
```
