## Supplementary scripts for "A multi-scale *in silico* mouse model for insulin resistance and humanoid type 2 diabetes": kin_substrate_activation_irr.m

```
function [R] = kin_substrate_activation_irr(V,substrate,Ksa,Ksc)
% kin_substrate_activation_irr: Substrate activation (irreversible) kinetics
%
% Application restrictions: 
% =========================
%   - Only irreversible
%   - exactly 1 substrate
%
% USAGE:
% ======
% R = kin_substrate_activation_irr(V,substrate,Ksa,Ksc)
%
% Output Arguments:
% =================
% R = V*(substrate/Ksa)^2 / ( 1+substrate/Ksc+substrate/Ksa+(substrate/Ksa)^2 ) 

% Information:
% ============
% SBPD Package - Systems Biology Parameter Determination Package
% Copyright 2008 by Henning Schmidt,

R = V*(substrate/Ksa)^2 / ( 1+substrate/Ksc+substrate/Ksa+(substrate/Ksa)^2 );
```
