## Supplementary scripts for "A multi-scale *in silico* mouse model for insulin resistance and humanoid type 2 diabetes": kin_uncomp_inihib_irr.m

```
function [R] = kin_uncomp_inihib_irr(V,substrate,Km,inhibitor,Ki)
% kin_uncomp_inihib_irr: Uncompetitive inhibition (irreversible) kinetics
%
% Application restrictions: 
% =========================
%   - Only irreversible
%   - exactly 1 substrate
%
% USAGE:
% ======
% R = kin_uncomp_inihib_irr(V,substrate,Km,inhibitor,Ki)
%
% Output Arguments:
% =================
% R = V*substrate / ( Km + substrate*(1+inhibitor/Ki) )

% Information:
% ============
% SBPD Package - Systems Biology Parameter Determination Package
% Copyright 2008 by Henning Schmidt,

R = V*substrate / ( Km + substrate*(1+inhibitor/Ki) );
```
