## Supplementary scripts for "A multi-scale *in silico* mouse model for insulin resistance and humanoid type 2 diabetes": SBPDsensitivity.m

```
function [varargout] = SBPDsensitivity(model,timevector,varargin)
% SBPDsensitivity: The SBPDsensitivity function is able to determine the 
% sensitivities of states and variables with respect to initial conditions
% and parameters, using a final difference approach.
%
% USAGE:
% ======
% output = SBPDsensitivity(model, timevector)
% output = SBPDsensitivity(model, timevector, parameternames)
% output = SBPDsensitivity(model, timevector, parameternames, statenames)
% output = SBPDsensitivity(model, timevector, parameternames, statenames, options)
% output = SBPDsensitivity(model, timevector, parameternames, statenames, options, nomparamvalues, nomicvalues)
%
% model: SBmodel or MEX simulation function
% timevector: a vector of time steps at which to return the simulation data
% parameternames: cell-array containing the names of the parameters for which to 
%       calculate the sensitivities. (default: all parameters
%       in the model)
% statenames: cell-array containing the names of the states for whose initial
%       conditions to calculate the sensitivities. (default: no states)
% options: structure with options for the integrator and sensitivity
%          calculation
%               options.pertfactor: relative perturbation if parameter not
%                   equal to zero. Otherwise absolute peturbation
%               options.abstol: absolute tolerance
%               options.reltol: relative tolerance
%               options.minstep: minimal integrator step size
%               options.maxstep: maximal integrator step size
%               options.maxnumsteps: maximum number of steps to be
%                   taken by the solver in its attempt to reach the next
%                   output time 
% nomparamvalues: vector of nominal parameter values
% nomicvalues: vector of nominal initial condition values
%
% DEFAULT VALUES:
% ===============
% parameternames: (all parameters)
% statenames:     (no states)
% options.pertfactor: 1e-5 
% options.abstol: 1e-6
% options.reltol: 1e-6
% options.minstep: 0
% options.maxstep: inf
% options.maxnumsteps: 500
% nomparamvalues: parameter values stored in the model
% nomicvalues: initial condition values stored in the model
%
% Output Arguments:
% =================
% If no output arguments are given, the result of the simulation is plotted
% after finished simulation (only states, variables, reactions). The
% sensitivities are not plotted!
%
% The output of SBPDsensitivity is realized as a structure:
% output.time:              vector with time instants for the results
% output.states:            cell-array with state names
% output.statevalues:       matrix with state values. Each row corresponds to
%                           one time instant
%
% The following fields are only present in case options.simdata = 'all' is
% defined.
% output.variables:         cell-array with variable names
% output.variablevalues:    matrix with variable values. Each row corresponds to
%                           one time instant
% output.reactions:         cell-array with reaction names
% output.reactionvalues:    matrix with reaction values. Each row corresponds to
%                           one time instant
%
% In the case events are present in the model the following fields are
% present in the output structure additionally:
% output.events:            cell-arra with the names of the events
% output.eventtimes:        vector with time instants at which events
%                           happened
% output.eventflags:        matrix of dimension: 'number of events' x
%                           'number of event time instants'. Each column in
%                           this matrix correponds to one timeinstant in
%                           the output.eventtimes vector. The row numbers
%                           correspond to the numbers of the events. A "1"
%                           element in the matrix indicates that the
%                           corresponding event has happened at the
%                           corresponding time. A "0" element tells you
%                           that the event has not been fired at the
%                           corresponding instant.
%
% Sensitivity trajectoried are returned as follows:
% output.sensparameters:         cell-array containing the names of the
%                                parameters for which sensitivities are determined
% output.paramtrajectories.states:    cell-array containing one entry for each
%                                parameter in output.sensparameters (same order).
%                                Each entry consists of a matrix, where the
%                                columns correspond to state sensitivities wrt
%                                to the parameter and the rows correspond to
%                                the different integration time points, given in
%                                output.time.
% output.paramtrajectories.variables: same as above just for variables
% output.paramtrajectories.reactions: same as above just for reactions
% output.sensicstates:           cell-array containing the names of the
%                                states for which initial conditions
%                                sensitivities are determined
% output.ictrajectories.states:    cell-array containing one entry for each
%                                state in output.sensicstates (same order).
%                                Each entry consists of a matrix, where the
%                                columns correspond to state sensitivities wrt
%                                to the initial conditions and the rows correspond to
%                                the different integration time points, given in
%                                output.time.
% output.ictrajectories.variables: same as above just for variables
% output.ictrajectories.reactions: same as above just for reactions

% Information:
% ============
% Copyright (C) 2005-2013 Henning Schmidt,
%
% This program is free software; you can redistribute it and/or modify
% it under the terms of the GNU General Public License as published by
% the Free Software Foundation; either version 2 of the License, or
% (at your option) any later version.
% 
% This program is distributed in the hope that it will be useful,
% but WITHOUT ANY WARRANTY; without even the implied warranty of
% MERCHANTABILITY or FITNESS FOR A PARTICULAR PURPOSE.  See the
% GNU General Public License for more details.
% 
% You should have received a copy of the GNU General Public License
% along with this program; if not, write to the Free Software
% Foundation, Inc., 59 Temple Place, Suite 330, Boston, MA  02111-1307  
% USA.

pertfactor = 1e-5;
% HANDLE NON-NUMERIC INITIAL CONDITIONS
nomicvalues = SBcalcICvector(model);
[dummy,nomparamvalues] = SBparameters(model);

%%%%%%%%%%%%%%%%%%%%%%%%%%%%%%%%%%%%%%%%%%%%%%%
% CHECK TYPE OF MODEL
%%%%%%%%%%%%%%%%%%%%%%%%%%%%%%%%%%%%%%%%%%%%%%%
if ~isSBmodel(model) && ~ischar(model),
    error('Model is not an SBmodel or the name of a MEX simulation function.');
end
if length(timevector) <=1,
    error('''timevector'' input argument needs to be a vector.');
end

%%%%%%%%%%%%%%%%%%%%%%%%%%%%%%%%%%%%%%%%%%%%%%%
% HANDLE VARIABLE INPUT ARGUMENTS
%%%%%%%%%%%%%%%%%%%%%%%%%%%%%%%%%%%%%%%%%%%%%%%
parameternames = SBparameters(model);
statenames = [];
options = [];
if nargin >= 3,
    parameternames = varargin{1};
    if ischar(parameternames), 
        parameternames = {parameternames}; 
    end
end
if nargin >= 4,
    statenames = varargin{2};
    if ischar(statenames), 
        statenames = {statenames}; 
    end
end
if nargin >= 5,
    options = varargin{3};
    % reset the pertfactor if set in options
	if isfield(options,'pertfactor')
        pertfactor = options.pertfactor;
	end
end
if nargin >= 6,
    nomparamvalues = varargin{4};
end
if nargin == 7,
    nomicvalues = varargin{5};
end
if nargin > 7,
    error('Incorrect number of input arguments');
end

%%%%%%%%%%%%%%%%%%%%%%%%%%%%%%%%%%%%%%%%%%%%%%%
% CREATE THE MEX MODEL
%%%%%%%%%%%%%%%%%%%%%%%%%%%%%%%%%%%%%%%%%%%%%%%
if ischar(model),
    MEXmodel = model;
    MEXmodelfullpath = '';
else
    [MEXmodel, MEXmodelfullpath] = makeTempMEXmodelSBPD(model);
end

%%%%%%%%%%%%%%%%%%%%%%%%%%%%%%%%%%%%%%%%%%%%%%%
% GET NOMINAL VALUES AND INDICES
%%%%%%%%%%%%%%%%%%%%%%%%%%%%%%%%%%%%%%%%%%%%%%%
% nom values for all params and states
allparamvaluesnom = nomparamvalues;
allicvaluesnom = nomicvalues;
% nom values for params and states for which to compute the sensitivities
paramindices = getparamindicesSBPD(model,parameternames);
paramvaluesnom = allparamvaluesnom(paramindices);
icindices = getstateindicesSBPD(model,statenames);
icvaluesnom = allicvaluesnom(icindices);

%%%%%%%%%%%%%%%%%%%%%%%%%%%%%%%%%%%%%%%%%%%%%%%
% RUN A NOMINAL SIMULATION
%%%%%%%%%%%%%%%%%%%%%%%%%%%%%%%%%%%%%%%%%%%%%%%
nomsimdata = feval(MEXmodel,timevector,allicvaluesnom,allparamvaluesnom,options);

%%%%%%%%%%%%%%%%%%%%%%%%%%%%%%%%%%%%%%%%%%%%%%%
% RUN THE SENSITIVITY ANALYSIS (PARAMETERS)
%%%%%%%%%%%%%%%%%%%%%%%%%%%%%%%%%%%%%%%%%%%%%%%
parampertsimdata = {};
paramdeltapertvalue = [];
for k=1:length(parameternames),
    % perturb the parameter
    paramkindex = paramindices(k);
    paramk = paramvaluesnom(k);
    if paramk ~= 0,
        deltapert = paramk*pertfactor;
    else
        deltapert = pertfactor;
    end
    paramdeltapertvalue(end+1) = deltapert;
    pertparamk = paramk + deltapert;
    % create parameter vector
    pv = allparamvaluesnom;
    pv(paramkindex) = pertparamk;
    % simulate the perturbed model
    parampertsimdata{end+1} = feval(MEXmodel,timevector,allicvaluesnom,pv,options);
end

%%%%%%%%%%%%%%%%%%%%%%%%%%%%%%%%%%%%%%%%%%%%%%%
% RUN THE SENSITIVITY ANALYSIS (INITIAL CONDITIONS)
%%%%%%%%%%%%%%%%%%%%%%%%%%%%%%%%%%%%%%%%%%%%%%%
icpertsimdata = {};
icdeltapertvalue = [];
for k=1:length(statenames),
    % perturb the ic
    ickindex = icindices(k);
    ick = icvaluesnom(k);
    if ick ~= 0,
        deltapert = ick*pertfactor;
    else
        deltapert = pertfactor;
    end
    icdeltapertvalue(end+1) = deltapert;
    pertick = ick + deltapert;
    % create ic vector
    ic = allicvaluesnom;
    ic(ickindex) = pertick;
    % simulate the perturbed model
    icpertsimdata{end+1} = feval(MEXmodel,timevector,ic,allparamvaluesnom,options);
end

%%%%%%%%%%%%%%%%%%%%%%%%%%%%%%%%%%%%%%%%%%%%%%%
% DETERMINE SENSITIVITIES (STATES)
%%%%%%%%%%%%%%%%%%%%%%%%%%%%%%%%%%%%%%%%%%%%%%%
states_sensparamtrajectories = {};
nomstates = nomsimdata.statevalues;
for k=1:length(parameternames),
    parampertstates = parampertsimdata{k}.statevalues;
    states_sensparamtrajectories{end+1} = (parampertstates-nomstates)/paramdeltapertvalue(k);
end
states_sensictrajectories = {};
for k=1:length(statenames),
    icpertstates = icpertsimdata{k}.statevalues;
    states_sensictrajectories{end+1} = (icpertstates-nomstates)/icdeltapertvalue(k);
end

%%%%%%%%%%%%%%%%%%%%%%%%%%%%%%%%%%%%%%%%%%%%%%%
% DETERMINE SENSITIVITIES (VARIABLES)
%%%%%%%%%%%%%%%%%%%%%%%%%%%%%%%%%%%%%%%%%%%%%%%
variables_sensparamtrajectories = {};
nomvariables = nomsimdata.variablevalues;
for k=1:length(parameternames),
    parampertvariables = parampertsimdata{k}.variablevalues;
    variables_sensparamtrajectories{end+1} = (parampertvariables-nomvariables)/paramdeltapertvalue(k);
end
variables_sensictrajectories = {};
for k=1:length(statenames),
    icpertvariables = icpertsimdata{k}.variablevalues;
    variables_sensictrajectories{end+1} = (icpertvariables-nomvariables)/icdeltapertvalue(k);
end

%%%%%%%%%%%%%%%%%%%%%%%%%%%%%%%%%%%%%%%%%%%%%%%
% DETERMINE SENSITIVITIES (REACTIONS)
%%%%%%%%%%%%%%%%%%%%%%%%%%%%%%%%%%%%%%%%%%%%%%%
reactions_sensparamtrajectories = {};
nomreactions = nomsimdata.reactionvalues;
for k=1:length(parameternames),
    parampertreactions = parampertsimdata{k}.reactionvalues;
    reactions_sensparamtrajectories{end+1} = (parampertreactions-nomreactions)/paramdeltapertvalue(k);
end
reactions_sensictrajectories = {};
for k=1:length(statenames),
    icpertreactions = icpertsimdata{k}.reactionvalues;
    reactions_sensictrajectories{end+1} = (icpertreactions-nomreactions)/icdeltapertvalue(k);
end

%%%%%%%%%%%%%%%%%%%%%%%%%%%%%%%%%%%%%%%%%%%%%%%
% BUILD OUTPUT STRUCTURE
%%%%%%%%%%%%%%%%%%%%%%%%%%%%%%%%%%%%%%%%%%%%%%%
output = nomsimdata;
output.sensparameters = parameternames;
output.paramtrajectories.states = states_sensparamtrajectories;
output.paramtrajectories.variables = variables_sensparamtrajectories;
output.paramtrajectories.reactions = reactions_sensparamtrajectories;
output.sensicstates = statenames;
output.ictrajectories.states = states_sensictrajectories;
output.ictrajectories.variables = variables_sensictrajectories;
output.ictrajectories.reactions = reactions_sensictrajectories;

%%%%%%%%%%%%%%%%%%%%%%%%%%%%%%%%%%%%%%%%%%%%%%%
% HANDLE VARIABLE OUTPUT ARGUMENTS
%%%%%%%%%%%%%%%%%%%%%%%%%%%%%%%%%%%%%%%%%%%%%%%
if nargout == 0,
    % prepare data for plotting
    time = output.time;
    datanames = {};
    dataindex = 1;
    for k = 1:length(output.states),
        datanames{dataindex} = sprintf('%s (state)',output.states{k});
        dataindex = dataindex + 1;
    end
    if isfield(output,'variables'),
        for k = 1:length(output.variables),
            datanames{dataindex} = sprintf('%s (variable)',output.variables{k});
            dataindex = dataindex + 1;
        end
        for k = 1:length(output.reactions),
            datanames{dataindex} = sprintf('%s (reaction rate)',output.reactions{k});
            dataindex = dataindex + 1;
        end
        datavalues = [output.statevalues, output.variablevalues, output.reactionvalues];
    else
        datavalues = [output.statevalues];
    end
    SBplot(createdatastructSBplotSB(time(:),datavalues,datanames));
elseif nargout == 1,
    varargout{1} = output;
else 
    error('Incorrect number of output arguments!');
end

%%%%%%%%%%%%%%%%%%%%%%%%%%%%%%%%%%%%%%%%%%%%%%%
% DELETE TEMP MEX MODEL IF CREATED
%%%%%%%%%%%%%%%%%%%%%%%%%%%%%%%%%%%%%%%%%%%%%%%
if ~isempty(MEXmodelfullpath),
    clear mex;
    delete(MEXmodelfullpath);
end
```
