## Supplementary scripts for "A multi-scale *in silico* mouse model for insulin resistance and humanoid type 2 diabetes": SBPDsimulate.m

```
function [varargout] = SBPDsimulate(mmodel,varargin)
% SBPDsimulate: Function allowing to simulate SBmodels and MEX simulation
% functions. SBmodels are first converted to temporary MEX simulation
% functions, so that a conversion overhead is present.
%
% USAGE:
% ======
% output = SBPDsimulate(model)
% output = SBPDsimulate(model, time)
% output = SBPDsimulate(model, time, ICs)
% output = SBPDsimulate(model, time, ICs, paramNames, paramValues)
% output = SBPDsimulate(model, time, ICs, paramNames, paramValues, OPTIONS)
%
% model: Name of MEX simulation function OR SBmodel 
% time: Either simulation end time (scalar) or simulation time vector.
% ICs: Vector of initial conditions.
% paramNames: Cell-array with the names of the parameters for which the
%             values are to be changed to given values.
% paramValues: Vector of parameter values for the given parameters.
% OPTIONS: structure with integrator options.
%       OPTIONS.showIntegratorStats: =0 (off), =1 shows integrator
%               statistics in the MATLAB console window
%       OPTIONS.method:             'stiff' or 'nonstiff' (default: 'stiff')
%       OPTIONS.abstol:             abs tolerance (default: 1e-6)
%       OPTIONS.reltol:             rel tolerance (default: 1e-6)
%       OPTIONS.minstep:            min step-size integrator (default: 0)
%       OPTIONS.maxstep:            max step-size integrator (default: inf)
%       OPTIONS.maxnumsteps:        max number of steps between two output
%                                   points (default: 100000)
%       OPTIONS.maxerrtestfails:    maximum number of error test failures
%                                   permitted in attempting one step
%                                   (default: 50)
%       OPTIONS.maxorder:           maximum order of the linear multistep
%                                   method (default: 5 BDF, 12 ADAMS)
%       OPTIONS.maxconvfails:       maximum number of nonlinear solver convergence 
%                                   failures permitted during one step
%                                   (default: 10)
%       OPTIONS.initstep:           initial step size to be attempted
%                                   (default: 0)
%       OPTIONS.maxnonlineariter:   maximum number of nonlinear solver
%                                   iterations permitted per step 
%                                   (default: 3)
%
% DEFAULT VALUES:
% ===============
% time: 20 
% ICs: values stored in the model
% paramNames: {}  do not change any parameters
% paramValues: []   
%
% Output Arguments:
% =================
% If no output arguments are given, the result of the simulation is plotted
% after finished simulation.
%
% The output of SBPDsimulate is realized as a structure:
% output.time:              vector with time instants for the results
% output.states:            cell-array with state names
% output.statevalues:       matrix with state values. Each row corresponds to
%                           one time instant
%
% output.variables:         cell-array with variable names
% output.variablevalues:    matrix with variable values. Each row corresponds to
%                           one time instant
% output.reactions:         cell-array with reaction names
% output.reactionvalues:    matrix with reaction values. Each row corresponds to
%                           one time instant
%
% In the case events are present in the model the following fields are
% present in the output structure additionally:
% output.events:            cell-arra with the names of the events
% output.eventtimes:        vector with time instants at which events
%                           happened
% output.eventflags:        matrix of dimension: 'number of events' x
%                           'number of event time instants'. Each column in
%                           this matrix correponds to one timeinstant in
%                           the output.eventtimes vector. The row numbers
%                           correspond to the numbers of the events. A "1"
%                           element in the matrix indicates that the
%                           corresponding event has happened at the
%                           corresponding time. A "0" element tells you
%                           that the event has not been fired at the
%                           corresponding instant.

% Information:
% ============
% Copyright (C) 2005-2013 Henning Schmidt,
%
% This program is free software; you can redistribute it and/or modify
% it under the terms of the GNU General Public License as published by
% the Free Software Foundation; either version 2 of the License, or
% (at your option) any later version.
% 
% This program is distributed in the hope that it will be useful,
% but WITHOUT ANY WARRANTY; without even the implied warranty of
% MERCHANTABILITY or FITNESS FOR A PARTICULAR PURPOSE.  See the
% GNU General Public License for more details.
% 
% You should have received a copy of the GNU General Public License
% along with this program; if not, write to the Free Software
% Foundation, Inc., 59 Temple Place, Suite 330, Boston, MA  02111-1307  
% USA.

%%%%%%%%%%%%%%%%%%%%%%%%%%%%%%%%%%%%%%%%%%%%%%%
% FIRST CHECK THE GIVEN MODEL
%%%%%%%%%%%%%%%%%%%%%%%%%%%%%%%%%%%%%%%%%%%%%%%
if isSBmodel(mmodel)
    % create the MEX simulation function
    [MEXmodel, MEXmodelfullpath] = makeTempMEXmodelSBPD(mmodel);
elseif exist(mmodel) == 3
    MEXmodel = mmodel;
    MEXmodelfullpath = '';
else
    error('Model input argument is no SBmodel and no MEX simulation function.');
end

%%%%%%%%%%%%%%%%%%%%%%%%%%%%%%%%%%%%%%%%%%%%%%%
% HANDLE VARIABLE INPUT ARGUMENTS
%%%%%%%%%%%%%%%%%%%%%%%%%%%%%%%%%%%%%%%%%%%%%%%
timevector = [0:0.01:20];  % just the default timevector
time = [];
ICs = [];
paramvec = [];
paramNames = {};
OPTIONS = [];
if nargin<1 || nargin==4 || nargin > 6
    error('Incorrect number of input arguments');
elseif nargin == 2
    time = varargin{1};
elseif nargin == 3
    time = varargin{1};
    ICs = varargin{2};
elseif nargin == 5
    time = varargin{1};
    ICs = varargin{2};
    paramNames = varargin{3};
    paramValues = varargin{4};
elseif nargin == 6
    time = varargin{1};
    ICs = varargin{2};
    paramNames = varargin{3};
    paramValues = varargin{4};
    OPTIONS = varargin{5};
end

%%%%%%%%%%%%%%%%%%%%%%%%%%%%%%%%%%%%%%%%%%%%%%%
% PROCESS INPUTS
%%%%%%%%%%%%%%%%%%%%%%%%%%%%%%%%%%%%%%%%%%%%%%%
if ~isempty(time)
    if length(time) == 1
        timevector = [0:time/1000:time];
    elseif length(time) > 1
        timevector = time;
    end
end
if ~isempty(paramNames)
    paramvec = makeparamvecSBPD(MEXmodel,paramNames,paramValues);
end

%%%%%%%%%%%%%%%%%%%%%%%%%%%%%%%%%%%%%%%%%%%%%%%
% CHECK IF DELAY PRESENT ... THEN ERROR
%%%%%%%%%%%%%%%%%%%%%%%%%%%%%%%%%%%%%%%%%%%%%%%
if isSBmodel(mmodel)
    if usedelaySB(mmodel)
        error('The model contains delays. This is not supported for MEX file simulation.');
    end
end

%%%%%%%%%%%%%%%%%%%%%%%%%%%%%%%%%%%%%%%%%%%%%%%
% IF EVENTS ARE PRESENT THEN USE A DEFAULT MAXSTEP
% Only do that in case of an SBmodel
%%%%%%%%%%%%%%%%%%%%%%%%%%%%%%%%%%%%%%%%%%%%%%%
if isSBmodel(mmodel)
    if useeventSB(mmodel)
        if ~isfield(OPTIONS,'maxstep')
            % change only if not user provided 
            OPTIONS.maxstep = timevector(end)/1000;
        end
    end
end

%%%%%%%%%%%%%%%%%%%%%%%%%%%%%%%%%%%%%%%%%%%%%%%
% RUN THE MEX SIMULATION FUNCTION WITH GIVEN INPUT
%%%%%%%%%%%%%%%%%%%%%%%%%%%%%%%%%%%%%%%%%%%%%%%
simdata = feval(MEXmodel,timevector,ICs,paramvec,OPTIONS);

%%%%%%%%%%%%%%%%%%%%%%%%%%%%%%%%%%%%%%%%%%%%%%%
% HANDLE VARIABLE OUTPUT ARGUMENTS
%%%%%%%%%%%%%%%%%%%%%%%%%%%%%%%%%%%%%%%%%%%%%%%
if nargout == 0
    % NO OUTPUT => DO PLOT RESULTS
    time = simdata.time;
    datanames = {};
    dataindex = 1;
    for k = 1:length(simdata.states)
        datanames{dataindex} = sprintf('%s (state)',simdata.states{k});
        dataindex = dataindex + 1;
    end
    if isfield(simdata,'variables')
        for k = 1:length(simdata.variables)
            datanames{dataindex} = sprintf('%s (variable)',simdata.variables{k});
            dataindex = dataindex + 1;
        end
        for k = 1:length(simdata.reactions)
            datanames{dataindex} = sprintf('%s (reaction rate)',simdata.reactions{k});
            dataindex = dataindex + 1;
        end
        datavalues = [simdata.statevalues, simdata.variablevalues, simdata.reactionvalues];
    else
        datavalues = [simdata.statevalues];
    end
    SBplot(createdatastructSBplotSB(time(:),datavalues,datanames));
elseif nargout == 1
    varargout{1} = simdata;
else 
    error('Incorrect number of simdata arguments!');
end

%%%%%%%%%%%%%%%%%%%%%%%%%%%%%%%%%%%%%%%%%%%%%%%
% DELETE THE TEMPORARY MEX SIMULATION FUNCTION IF IT HAS BEEN CREATED
%%%%%%%%%%%%%%%%%%%%%%%%%%%%%%%%%%%%%%%%%%%%%%%
if ~isempty(MEXmodelfullpath)
    clear mex;
    delete(MEXmodelfullpath);
end
```
