## Supplementary scripts for "A multi-scale *in silico* mouse model for insulin resistance and humanoid type 2 diabetes": CVODEmex25.h

```
/*
 * CVODEmex25.h: MEX/CVODES Interface for Sundials CVODES version 2.5
 *
 * Information:
 * ============
 * SBPD Package - Systems Biology Parameter Determination Package
 * Copyright 2008 by Henning Schmidt,
 */

/* CVODE related flags */
#define DOFLAG_DDT 0
#define DOFLAG_VARREAC 1
#define DOFLAG_EVENTS 2
#define DOFLAG_EVENTASSIGN 3
#define DOFLAG_CALCICS 4

/* ParamData (contains pointer to parameter values passed to integrator) */
typedef struct {
    double *parametervector;
} ParamData;

/* Variables defined outside the library */
extern double defaultICs_num[], defaultParam[];
extern char  *defaultICs_nonnum[];
extern char  *stateNames[], *parameterNames[], *variableNames[], *reactionNames[], *eventNames[]; 
extern const int NRSTATES, NRPARAMETERS, NRVARIABLES, NRREACTIONS, NREVENTS;
extern const int hasOnlyNumericICs;
extern int   *interpcseSB_check; /* needed for spline function in mexsplineaddon.h */
extern int   *interpcseSlopeSB_check; /* needed for spline function in mexsplineaddon.h */

/* Functions containing the model equations */
extern void model(double time, double *stateVector, double *DDTvector, ParamData *paramdataPtr, int DOflag, double *variableVector, double *reactionVector, double *gout, int *eventVector);
```
