## Supplementary scripts for "A multi-scale *in silico* mouse model for insulin resistance and humanoid type 2 diabetes": kineticformulas.h

```
/*
 * kineticformulas.h: Include file for specification of useful kinetic formulas
 *
 * Information:
 * ============
 * SBPD Package - Systems Biology Parameter Determination Package
 * Copyright 2008 by Henning Schmidt,
 */

double kin_mass_action_irr(double k, double substrate)
{
    return k*substrate;    
}

double kin_mass_action_rev(double k1, double substrate, double k2, double product)
{
    return k1*substrate - k2*product;    
}

double kin_degradation(double kdeg, double substrate)
{
    return kdeg*substrate;    
}

double kin_michaelis_menten_irr(double V, double substrate, double Km)
{
    return V*substrate / ( Km + substrate );    
}

double kin_hill_cooperativity_irr(double V, double substrate, double h, double Shalve)
{
    return V*pow(substrate,h) / (pow(Shalve,h) + pow(substrate,h));    
}

double kin_mixed_inihib_irr(double V, double substrate, double Km, double inhibitor, double Kis, double Kic)
{
    return V*substrate / ( Km*(1+inhibitor/Kis) + substrate*(1+inhibitor/Kic) );    
}

double kin_noncomp_inihib_irr(double V, double substrate, double Km, double inhibitor, double Ki)
{
    return V*substrate / ( (Km+substrate)*(1+inhibitor/Ki) );
}

double kin_uncomp_inihib_irr(double V, double substrate, double Km, double inhibitor, double Ki)
{
    return V*substrate / ( Km + substrate*(1+inhibitor/Ki) );
}

double kin_comp_inihib_irr(double V, double substrate, double Km, double inhibitor, double Ki)
{
    return V*substrate / ( Km*(1+inhibitor/Ki) + substrate );
}

double kin_substrate_inihib_irr(double V, double substrate, double Km, double Ki)
{
    return V*substrate / ( Km + substrate + Km*pow((substrate/Ki),2.0) );
}

double kin_allosteric_inihib_mwc_irr(double V, double substrate, double Ks, double n, double L, double inhibitor, double Ki)
{
    return V*substrate*pow((Ks+substrate),(n-1.0)) / ( L*pow((Ks*(1+inhibitor/Ki)),n) + pow((Ks+substrate),n) );
}

double kin_allosteric_inihib_empirical_rev(double Vf, double substrate, double Kms, double Vr, double product, double Kmp, double inhibitor, double Ki, double n)
{
    return (Vf*substrate/Kms - Vr*product/Kmp) / (1 + substrate/Kms + product/Kmp + pow((inhibitor/Ki),n));
}

double kin_catalytic_activation_irr(double V, double substrate, double activator, double Kms, double Ka)
{
    return V*substrate*activator / ( (Kms + substrate)*(Ka+activator) );
}

double kin_catalytic_activation_rev(double Vf, double substrate, double Kms, double Vr, double product, double Kmp, double activator, double Ka)
{
    return ( Vf*substrate/Kms - Vr*product/Kmp )*activator / ( (1+substrate/Kms+product/Kmp)*(Ka+activator) );
}

double kin_comp_inihib_rev(double Vf, double substrate, double Kms, double Vr, double product, double Kmp, double inhibitor, double Ki)
{
    return (Vf*substrate/Kms - Vr*product/Kmp) / ( 1+substrate/Kms+product/Kmp+inhibitor/Ki );
}

double kin_constantflux(double v)
{
    return v;
}

double kin_hyperbolic_modifier_irr(double V,double substrate,double b,double modifier,double a,double Kd,double Km)
{
    return V*substrate*(1+b*modifier/(a*Kd)) / ( Km*(1+modifier/Kd) + substrate*(1+modifier/(a*Kd)) );
}

double kin_hyperbolic_modifier_rev(double Vf,double substrate,double Kms,double Vr,double product,double Kmp,double b,double modifier,double a,double Kd)
{
    return (Vf*substrate/Kms - Vr*product/Kmp)*(1+b*modifier/(a*Kd)) / ( 1+modifier/Kd+(substrate/Kms+product/Kmp)*(1+modifier/(a*Kd)) );
}

double kin_hill_rev(double Vf,double substrate,double Shalve,double product,double Keq,double Phalve,double h)
{
    return Vf*substrate/Shalve*(1-product/(substrate*Keq))*pow((substrate/Shalve+product/Phalve),(h-1.0)) / ( 1+pow((substrate/Shalve + product/Phalve),h) );
}

double kin_iso_uni_uni_rev(double Vf,double substrate,double product,double Keq,double Kii,double Kms,double Kmp)
{
    return Vf*(substrate-product/Keq) / ( substrate*(1+product/Kii) + Kms*(1+product/Kmp) );
}

double kin_mixed_activation_irr(double V,double substrate,double activator,double Kms,double Kas,double Kac)
{
    return V*substrate*activator / ( Kms*(Kas+activator) + substrate*(Kac+activator) );
}

double kin_mixed_activation_rev(double Vf,double substrate,double Kms,double Vr,double product,double Kmp,double activator,double Kas,double Kac)
{
    return (Vf*substrate/Kms - Vr*product/Kmp)*activator / ( Kas+activator+(substrate/Kms+product/Kmp)*(Kac+activator) );
}

double kin_mixed_inihib_rev(double Vf,double substrate,double Kms,double Vr,double product,double Kmp,double inhibitor,double Kis,double Kic)
{
    return (Vf*substrate/Kms - Vr*product/Kmp) / ( 1+inhibitor/Kis+(substrate/Kms+product/Kmp)*(1+inhibitor/Kic) );
}

double kin_noncomp_inihib_rev(double Vf,double substrate,double Kms,double Vr,double product,double Kmp,double inhibitor,double Ki)
{
    return (Vf*substrate/Kms-Vr*product/Kmp) / ( (1+substrate/Kms+product/Kmp)*(1+inhibitor/Ki) );
}

double kin_ordered_bi_bi_rev(double Vf,double substratea,double substrateb,double productp,double productq,double Keq,double Kip,double Kma,double Kmb,double Kia,double Vr,double Kmq,double Kmp,double Kib)
{
    return Vf*(substratea*substrateb-productp*productq/Keq) / (substratea*substrateb*(1+productp/Kip) + Kma*substrateb + Kmb*(substratea+Kia)+Vf/(Vr*Keq)*(Kmq*productp*(1+substratea/Kia) + productq*(Kmp*(1+Kia*substrateb/(Kma*Kmb))+productp*(1+substrateb/Kib))) );
}

double kin_ordered_bi_uni_rev(double Vf,double substratea,double substrateb,double product,double Keq,double Kma,double Kmb,double Vr,double Kmp,double Kia)
{
    return Vf*(substratea*substrateb-product/Keq) / ( substratea*substrateb+Kma*substrateb+Kmb*substratea+Vf/(Vr*Keq)*(Kmp+product*(1+substratea/Kia)) );
}

double kin_ordered_uni_bi_rev(double Vf,double substrate,double productp,double productq,double Keq,double Kms,double Kip,double Vr,double Kmq,double Kmp)
{
    return Vf*(substrate-productp*productq/Keq) / ( Kms+substrate*(1+productp/Kip)+Vf/(Vr*Keq)*(Kmq*productp+Kmp*productq+productp*productq) );
}

double kin_michaelis_menten_rev(double Vf,double substrate,double Kms,double Vr,double product,double Kmp)
{
    return (Vf*substrate/Kms+Vr*product/Kmp) / ( 1+substrate/Kms+product/Kmp );
}

double kin_substrate_inihib_rev(double Vf,double substrate,double Kms,double Vr,double product,double Kmp,double Ki)
{
    return (Vf*substrate/Kms-Vr*product/Kmp) / ( 1+substrate/Kms+product/Kmp+pow((substrate/Ki),2.0) );
}

double kin_uncomp_inihib_rev(double Vf,double substrate,double Kms,double Vr,double product,double Kmp,double inhibitor,double Ki)
{
    return ( Vf*substrate/Kms-Vr*product/Kmp ) / ( 1+(substrate/Kms+product/Kmp)*(1+inhibitor/Ki) );
}

double kin_uni_uni_rev(double Vf,double substrate,double product,double Keq,double Kms,double Kmp)
{
    return Vf*( substrate-product/Keq ) / ( substrate+Kms*(1+product/Kmp) );
}

double kin_specific_activation_irr(double V,double substrate,double activator,double Kms,double Ka)
{
    return V*substrate*activator/( Kms*Ka+(Kms+substrate)*activator );
}

double kin_specific_activation_rev(double Vf,double substrate,double Kms,double Vr,double product,double Kmp,double activator,double Ka)
{
    return ( Vf*substrate/Kms-Vr*product/Kmp )*activator / ( Ka+(1+substrate/Kms+product/Kmp)*activator );
}

double kin_substrate_activation_irr(double V,double substrate,double Ksa,double Ksc)
{
    return V*pow((substrate/Ksa),2.0) / ( 1+substrate/Ksc+substrate/Ksa+pow((substrate/Ksa),2.0) );
}

double kin_ping_pong_bi_bi_rev(double Vf,double substratea,double substrateb,double productp,double productq,double Keq,double Kiq,double Kma,double Kmb,double Vr,double Kmq,double Kia,double Kmp)
{
    return Vf*( substratea*substrateb-productp*productq/Keq ) / (substratea*substrateb*(1+productq/Kiq)+Kma*substrateb+Kmb*substratea+Vf/(Vr*Keq)*(Kmq*productp*(1+substratea/Kia)+productq*(Kmp+productp)));
}

double kin_hill_1_modifier_rev(double Vf,double substrate,double Shalve,double product,double Keq,double Phalve,double h,double modifier,double Mhalve,double alpha)
{
    return Vf*substrate/Shalve*(1-product/(substrate*Keq))*pow((substrate/Shalve+product/Phalve),(h-1.0)) / ( (1+pow((modifier/Mhalve),h))/(1+alpha*pow((modifier/Mhalve),h))+pow((substrate/Shalve + product/Phalve),h) );
}

double kin_hill_2_modifiers_rev(double Vf,double substrate,double Shalve,double product,double Keq,double Phalve,double h,double modifierA,double MAhalve,double modifierB,double MBhalve,double alphaA,double alphaB,double alphaAB)
{
    return Vf*substrate/Shalve*(1-product/(substrate*Keq))*pow((substrate/Shalve+product/Phalve),(h-1.0)) / ( (1+pow((modifierA/MAhalve),h) + 1+pow((modifierB/MBhalve),h)) / ( 1+alphaA*pow((modifierA/MAhalve),h)+alphaB*pow((modifierB/MBhalve),h)+alphaA*alphaB*alphaAB*pow((modifierA/MAhalve),h)*pow((modifierB/MBhalve),h) ) + pow((substrate/Shalve + product/Phalve),h) );
}
```
