## Supplementary scripts for "A multi-scale *in silico* mouse model for insulin resistance and humanoid type 2 diabetes": mexmathaddon.h

```
/*
 * mexmathaddon.h: Include file for automatically generated MEX model files.
 *
 * Information:
 * ============
 * SBPD Package - Systems Biology Parameter Determination Package
 * Copyright 2008 by Henning Schmidt,
 */

#define pi 3.141592653589793

#ifdef __LCC__
double max(double a, double b)
{
    if (a<b) return b;
    else return a;
}

double min(double a, double b)
{
    if (a>b) return b;
    else return a;
}
#endif

#ifdef __GNUC__
double max(double a, double b)
{
    return fmax(a,b);
}

double min(double a, double b)
{
    return fmin(a,b);
}
#endif


double minSB(int nargs, ...)
{
    int k;
    double minimum;
    double checkmin;
    va_list vararg;
    va_start(vararg, nargs);
    minimum = va_arg(vararg, double);
    for (k=1; k<nargs; k++ ) {
        checkmin = va_arg(vararg, double);
        if (checkmin < minimum) {
            minimum = checkmin;
        }
    }
    va_end (vararg);
    return minimum;
}

double maxSB(int nargs, ...)
{
    int k;
    double maximum;
    double checkmax;
    va_list vararg;
    va_start(vararg, nargs);
    maximum = va_arg(vararg, double);
    for (k=1; k<nargs; k++ ) {
        checkmax = va_arg(vararg, double);
        if (checkmax > maximum) {
            maximum = checkmax;
        }
    }
    va_end (vararg);
    return maximum;
}

double indexmaxSB(int nargs, ...)
{
    int k;
    double maximum;
    double indexmaximum;
    double checkmax;
    va_list vararg;
    va_start(vararg, nargs);
    maximum = va_arg(vararg, double);
    indexmaximum = 1;
    for (k=1; k<nargs; k++ ) {
        checkmax = va_arg(vararg, double);
        if (checkmax > maximum) {
            maximum = checkmax;
            indexmaximum = (double)k+1;
        }
    }
    va_end (vararg);
    return indexmaximum;
}

double sign(double a)
{
    if (a<0) return -1;
    else if (a==0) return 0;
    else if (a>0) return 1;
    else return a;
}

double absSB(double a)
{
    if (a < 0) return -a;
    else return a;
}

double mod(double a, double b)
{
    if (b == 0) return a;
    else if (a == b) return 0;
    return sign(b)*absSB(a-floor(a/b)*b);
}

double nthrootSB(double a, double n)
{
    return pow(a,1.0/n);
}

double andSB(int nargs, ...)
{
    int k;
    va_list vararg;
    va_start(vararg, nargs);
    for (k=0; k<nargs; k++ ) {
        if (va_arg(vararg, double) == 0) {
            va_end (vararg);
            return 0;
        }
    }
    va_end (vararg);
    return 1;
}

double orSB(int nargs, ...)
{
    int k;
    va_list vararg;
    va_start(vararg, nargs);
    for (k=0; k<nargs; k++ ) {
        if (va_arg(vararg, double) != 0) {
            va_end (vararg);
            return 1;
        }
    }
    va_end (vararg);
    return 0;
}

double piecewiseSB(int nargs, ...)
{
    int k, oddnumber;
    double *data;
    double returnvalue;
    va_list vararg;
    va_start(vararg, nargs);
    /* Read in all variable input arguments in double array */
    data = (double *)mxCalloc(nargs,sizeof(double));
    for (k=0; k<nargs; k++ ) {
        data[k] = va_arg(vararg, double);
    }
    /* 
       Determine result
       Check if odd or even number of input arguments
     */
    oddnumber = nargs % 2;
    for (k=0;k<nargs-oddnumber;k+=2) {
        if (data[k+1] != 0) {
            returnvalue = data[k];
            mxFree(data); /* free temporary array */
            return returnvalue;
        }
    }
    if (oddnumber != 0) {
            returnvalue = data[nargs-1];
            mxFree(data); /* free temporary array */
            return returnvalue;
    }
    else {
        mexErrMsgTxt("A piecewise statement is wrongly defined - missing (but needed) default value.");
        mxFree(data); /* free temporary array */
        return 0; /* statement never reached */
    }
}

double piecewiseT0SB(int nargs, ...) /* does the same as the piecewiseSB function but does not lead to added events in the simulation model */
{
    int k, oddnumber;
    double *data;
    double returnvalue;
    va_list vararg;
    va_start(vararg, nargs);
    /* Read in all variable input arguments in double array */
    data = (double *)mxCalloc(nargs,sizeof(double));
    for (k=0; k<nargs; k++ ) {
        data[k] = va_arg(vararg, double);
    }
    /* 
       Determine result
       Check if odd or even number of input arguments
     */
    oddnumber = nargs % 2;
    for (k=0;k<nargs-oddnumber;k+=2) {
        if (data[k+1] != 0) {
            returnvalue = data[k];
            mxFree(data); /* free temporary array */
            return returnvalue;
        }
    }
    if (oddnumber != 0) {
            returnvalue = data[nargs-1];
            mxFree(data); /* free temporary array */
            return returnvalue;
    }
    else {
        mexErrMsgTxt("A piecewise statement is wrongly defined - missing (but needed) default value.");
        mxFree(data); /* free temporary array */
        return 0; /* statement never reached */
    }
}


double piecewiseSmoothSB(double t,double y0,double y1,double alpha) 
{

return (y0+y1*exp(alpha*t))/(1+exp(alpha*t));

}
        

double gt(double a, double b)
{
    if (a > b) return 1;
    else return 0;
}

double ge(double a, double b)
{
    if (a >= b) return 1;
    else return 0;
}

double lt(double a, double b)
{
    if (a < b) return 1;
    else return 0;
}

double le(double a, double b)
{
    if (a <= b) return 1;
    else return 0;
}

double eq(double a, double b)
{
    if (a*b < 0) return 0;
    else if ( absSB(a-b) < 10.0*mxGetEps() ) return 1;
    else return 0;
}

/* THIS FUNCTION IS NOT NEEDED ANYMORE ... in SBPDmakeMEXmodel the relevant parts have been commented out 
 * and in SBPDsimulate a check has been included that gives an error is a model contains a delaySB function.
 *
double delaySB(double input, double tau, double time, char *queuename)
{
    mxArray *callArg[4];
    mxArray *outArg[1];
    callArg[0] = mxCreateDoubleScalar(input);
    callArg[1] = mxCreateDoubleScalar(tau);
    callArg[2] = mxCreateDoubleScalar(time);
    callArg[3] = mxCreateString(queuename);
    mexCallMATLAB(1, outArg, 4, callArg, "delaySB");
    return mxGetScalar(outArg[0]);
}
*/
```
