## Supplementary scripts for "A multi-scale *in silico* mouse model for insulin resistance and humanoid type 2 diabetes": mexsplineaddon.h

```
/*
 * mexsplineaddon.h: contains the function interpcseSB, interpcsexSB and '
 *      the function interpcsSB
 *      Cubic spline interpolation with (and without) endpoints for use 
 *      by the CVODE integrator in the SBPD package
 *
 * The interpcsSB is a function for cubic spline interpolation, but without the 
 * possibility of defining slopes at the endpoints.
 * 
 * The interpcseSB function allows to define slopes at the endpoints. The 
 * spline coefficients are only calculated once and then only evaluated (faster)
 * 
 * the interpcsexSB function determines the spline coefficients each time, allowing to 
 * implement time dependent spline functions (slower).
 *
 */

/* interpcseSB and interpcsexSB FUNCTIONS:
 * Original C-code for spline interpolation taken from:
 *   http://www.mech.uq.edu.au/staff/jacobs/nm_lib/cmathsrc/spline.c
 *
 * The syntax of this MEX function (to be used via CVODE interface) is as follows:
 *
 * yy = interpcseSB(nargs,n,x1,x2,...,xn,y1,y2,...,yn,xx)
 * yy = interpcseSB(nargs,n,e1,e2,x1,x2,...,xn,y1,y2,...,yn,xx)
 *
 * nargs:  number of arguments
 * n:      number of points in the x and y vectors
 * xi:     x-values 
 * yi:     y-values
 * xx:     x-value at which to evaluate the spline function (do NOT allow multiple)
 * e1,e2:  endpoint derivatives (if specified, both need to be given)
 * yy:     interpolated value
 */

/*
 *========================================================================
 * Initialization of the spline related functions
 *========================================================================
 */
void spline (int n, int e1, int e2, double s1, double s2, double x[], double y[], double b[], double c[], double d[], int *flag);
double seval (int n, double xx, double x[], double y[], double b[], double c[], double d[], int *last);
double deriv (int n, double xx, double x[], double b[], double c[], double d[], int *last);

/*
 *========================================================================
 * Declare variables for the interpcseSB function
 *========================================================================
 */
#define MAX_INTERPCSESB 10000    /* defined maximum number of interpcseSB functions in the model */
int*    interpcseSB_check = NULL; /* is initialized each new integration start to NULL ... required */
double* xvec[MAX_INTERPCSESB];
double* yvec[MAX_INTERPCSESB];
double* bvec[MAX_INTERPCSESB];
double* cvec[MAX_INTERPCSESB];
double* dvec[MAX_INTERPCSESB];

int*    interpcseSlopeSB_check = NULL; /* is initialized each new integration start to NULL ... required */
double* xvecslope[MAX_INTERPCSESB];
double* yvecslope[MAX_INTERPCSESB];
double* bvecslope[MAX_INTERPCSESB];
double* cvecslope[MAX_INTERPCSESB];
double* dvecslope[MAX_INTERPCSESB];

/*
 *========================================================================
 * INTERFACE FUNCTION for interpcseSB ... 
 *========================================================================
 */
double interpcseSB(int nargs, ...)
{
    /*************************************************************/
    /* Define the variables needed for the spline function calls */
    /*************************************************************/
    int    index_pw; /* index of the interpcseSB function */
    int    n;
    double *x;
    double *y;
    double xx;
    double s1,s2;
    int    e1,e2;
    double *b;
    double *c;
    double *d;
    int    flag;
    int    last;
    int    k;
    double yy;
    int    newCall;
    double b2,c2,d2,x0,x1,y0,y1;
    
    /***************************************/
    /* Handle the variable input arguments */
    /***************************************/
    /* Initialize the va_list */
    va_list vararg;
    va_start(vararg, nargs);
    /* Get the index of the current interpcseSB function */
    index_pw = (int) va_arg(vararg, double);
    /* Get the number of the x and y axis nodes for interpolation */
    n = (int) va_arg(vararg, double);
    /* Get the xx value at which to interpolate */
    xx = va_arg(vararg, double);
    
    /* Check if too many interpcseSB functions in the model (unlikely) */
    if (index_pw >= MAX_INTERPCSESB) {
        mexErrMsgTxt("interpcseSB_CVODE: To many calls to interpcseSB in the model.");
    }
    
    /* Check if memory already obtained for handling the different calls */
    if (interpcseSB_check == NULL) interpcseSB_check = (int *) mxCalloc(MAX_INTERPCSESB, sizeof(int));
    
    /* Check if new call or subsequent call */
    if (interpcseSB_check[index_pw] == 69) {
        newCall = 0;
    } else {
        newCall = 1;
        interpcseSB_check[index_pw] = 69;
    }

    if (newCall == 1 || n == 2) {
        /* First call to this function ... get the information */
        /* Check if the number of input argument is plausible and define slopes */
        if (nargs == 3+2*n) {
            /* Plausible, but no slopes defined */
            e1 = 0; s1 = 0; /* set s1 to 0 but it is not used */
            e2 = 0; s2 = 0; /* set s2 to 0 but it is not used */
        } else if (nargs == 3+2*n+2) {
            /* Plausible, slopes are defined */
            e1 = 1; s1 = va_arg(vararg, double);
            e2 = 1; s2 = va_arg(vararg, double);
        } else {
            mexErrMsgTxt("interpcseSB_CVODE: Incorrect number of input arguments (increase MAX_INTERPCSESB in mexsplineaddon.h).");
        }
        
        /***********/
        /* If n==2 */
        /***********/
        if (n == 2) {
            if (e1 == 0 || e2 == 0) mexErrMsgTxt("interpcseSB: n=2 requires the definition of slopes.");
            x0 = va_arg(vararg, double);
            x1 = va_arg(vararg, double);
            y0 = va_arg(vararg, double);
            y1 = va_arg(vararg, double);
            /* End the variable input handling */
            va_end (vararg);
            b2 = s1;
            c2 = -(-3.0*y1+3.0*y0+(s2+2.0*s1)*x1+(-s2-2.0*s1)*x0)/(pow(x1, 2.0)-2.0*x0*x1+pow(x0, 2.0));
            d2 = (-2.0*y1+2.0*y0+(s2+s1)*x1+(-s2-s1)*x0)/(pow(x1, 3.0)-3.0*x0*pow(x1, 2.0)+3.0*pow(x0, 2.0)*x1-pow(x0, 3.0));
            yy = y0 + b2*(xx-x0) + c2*pow(xx-x0, 2.0) + d2*pow(xx-x0, 3.0);   
        } else {
            /***********/
            /* If n!=2 */
            /***********/
            /* Allocate memory for x and y */
            x  = (double *) mxCalloc(n, sizeof(double));
            y  = (double *) mxCalloc(n, sizeof(double));
            /* Parse input arguments into x and y */
            for (k=0;k<n;k++) x[k] = va_arg(vararg, double);
            for (k=0;k<n;k++) y[k] = va_arg(vararg, double);
            /* End the variable input handling */
            va_end (vararg);
        
            /*****************************/
            /* Allocate memory for b,c,d */
            /*****************************/
            b  = (double *) mxCalloc(n, sizeof(double));
            c  = (double *) mxCalloc(n, sizeof(double));
            d  = (double *) mxCalloc(n, sizeof(double));
            
            /****************************/
            /* Call the spline function */
            /****************************/
            spline(n, e1, e2, s1, s2, x, y, b, c, d, &flag);
            
            /*********************************/
            /* Save b,c,d,x,y for next calls */
            /*********************************/
            xvec[index_pw] = x;
            yvec[index_pw] = y;
            bvec[index_pw] = b;
            cvec[index_pw] = c;
            dvec[index_pw] = d;
            
            /***************************************/
            /* Call the spline evaluation function */
            /***************************************/
            yy = seval (n, xx, x, y, b, c, d, &last);            
        }
    } else {
        /* get x,y,b,c,d pointers */
        x = xvec[index_pw];
        y = yvec[index_pw];
        b = bvec[index_pw];
        c = cvec[index_pw];
        d = dvec[index_pw];
        /***************************************/
        /* Call the spline evaluation function */
        /***************************************/
        yy = seval(n, xx, x, y, b, c, d, &last);
    }
    
    /*********************************/
    /* Return the interpolated value */
    /*********************************/
    return yy;
}


/*
 *========================================================================
 * INTERFACE FUNCTION for interpcseSB ... 
 *========================================================================
 */
double interpcseSlopeSB(int nargs, ...)
{
    /*************************************************************/
    /* Define the variables needed for the spline function calls */
    /*************************************************************/
    int    index_pw; /* index of the interpcseSlopeSB function */
    int    n;
    double *x;
    double *y;
    double xx;
    double s1,s2;
    int    e1,e2;
    double *b;
    double *c;
    double *d;
    int    flag;
    int    last;
    int    k;
    double yy;
    int    newCall;
    double b2,c2,d2,x0,x1,y0,y1;
    
    /***************************************/
    /* Handle the variable input arguments */
    /***************************************/
    /* Initialize the va_list */
    va_list vararg;
    va_start(vararg, nargs);
    /* Get the index of the current interpcseSlopeSB function */
    index_pw = (int) va_arg(vararg, double);
    /* Get the number of the x and y axis nodes for interpolation */
    n = (int) va_arg(vararg, double);
    /* Get the xx value at which to interpolate */
    xx = va_arg(vararg, double);
    
    /* Check if too many interpcseSlopeSB functions in the model (unlikely) */
    if (index_pw >= MAX_INTERPCSESB) {
        mexErrMsgTxt("interpcseSlopeSB_CVODE: To many calls to interpcseSB in the model.");
    }
    
    /* Check if memory already obtained for handling the different calls */
    if (interpcseSlopeSB_check == NULL) interpcseSlopeSB_check = (int *) mxCalloc(MAX_INTERPCSESB, sizeof(int));
    
    /* Check if new call or subsequent call */
    if (interpcseSlopeSB_check[index_pw] == 69) {
        newCall = 0;
    } else {
        newCall = 1;
        interpcseSlopeSB_check[index_pw] = 69;
    }

    if (newCall == 1 || n == 2) {
        /* First call to this function ... get the information */
        /* Check if the number of input argument is plausible and define slopes */
        if (nargs == 3+2*n) {
            /* Plausible, but no slopes defined */
            e1 = 0; s1 = 0; /* set s1 to 0 but it is not used */
            e2 = 0; s2 = 0; /* set s2 to 0 but it is not used */
        } else if (nargs == 3+2*n+2) {
            /* Plausible, slopes are defined */
            e1 = 1; s1 = va_arg(vararg, double);
            e2 = 1; s2 = va_arg(vararg, double);
        } else {
            mexErrMsgTxt("interpcseSlopeSB_CVODE: Incorrect number of input arguments (increase MAX_INTERPCSESB in mexsplineaddon.h).");
        }
        
        /***********/
        /* If n==2 */
        /***********/
        if (n == 2) {
            if (e1 == 0 || e2 == 0) mexErrMsgTxt("interpcseSlopeSB_CVODE: n=2 requires the definition of slopes.");
            x0 = va_arg(vararg, double);
            x1 = va_arg(vararg, double);
            y0 = va_arg(vararg, double);
            y1 = va_arg(vararg, double);
            /* End the variable input handling */
            va_end (vararg);
            b2 = s1;
            c2 = -(-3.0*y1+3.0*y0+(s2+2.0*s1)*x1+(-s2-2.0*s1)*x0)/(pow(x1, 2.0)-2.0*x0*x1+pow(x0, 2.0));
            d2 = (-2.0*y1+2.0*y0+(s2+s1)*x1+(-s2-s1)*x0)/(pow(x1, 3.0)-3.0*x0*pow(x1, 2.0)+3.0*pow(x0, 2.0)*x1-pow(x0, 3.0));
            yy = b2 + 2.0*c2*(xx-x0) + 3.0*d2*pow(xx-x0, 2.0);   
        } else {
            /***********/
            /* If n!=2 */
            /***********/
            /* Allocate memory for x and y */
            x  = (double *) mxCalloc(n, sizeof(double));
            y  = (double *) mxCalloc(n, sizeof(double));
            /* Parse input arguments into x and y */
            for (k=0;k<n;k++) x[k] = va_arg(vararg, double);
            for (k=0;k<n;k++) y[k] = va_arg(vararg, double);
            /* End the variable input handling */
            va_end (vararg);
        
            /*****************************/
            /* Allocate memory for b,c,d */
            /*****************************/
            b  = (double *) mxCalloc(n, sizeof(double));
            c  = (double *) mxCalloc(n, sizeof(double));
            d  = (double *) mxCalloc(n, sizeof(double));
            
            /****************************/
            /* Call the spline function */
            /****************************/
            spline(n, e1, e2, s1, s2, x, y, b, c, d, &flag);
            
            /*********************************/
            /* Save b,c,d,x,y for next calls */
            /*********************************/
            xvecslope[index_pw] = x;
            yvecslope[index_pw] = y;
            bvecslope[index_pw] = b;
            cvecslope[index_pw] = c;
            dvecslope[index_pw] = d;
            
            /***************************************/
            /* Call the spline evaluation function */
            /***************************************/
            yy = deriv(n, xx, x, b, c, d, &last);            
        }
    } else {
        /* get x,y,b,c,d pointers */
        x = xvecslope[index_pw];
        y = yvecslope[index_pw];
        b = bvecslope[index_pw];
        c = cvecslope[index_pw];
        d = dvecslope[index_pw];
        /***************************************/
        /* Call the spline evaluation function */
        /***************************************/
        yy = deriv(n, xx, x, b, c, d, &last);
    }
    
    /*********************************/
    /* Return the interpolated value */
    /*********************************/
    return yy;
}

/*
 *========================================================================
 * INTERFACE FUNCTION for interpcsexSB ... 
 *========================================================================
 */
double interpcsexSB(int nargs, ...)
{
    /*************************************************************/
    /* Define the variables needed for the spline function calls */
    /*************************************************************/
    int    n;
    double *x;
    double *y;
    double xx;
    double s1,s2;
    int    e1,e2;
    double *b;
    double *c;
    double *d;
    int    flag;
    int    last;
    int    k;
    double yy;
    double b2, c2, d2;
    double x0,x1,y0,y1;
    
    /***************************************/
    /* Handle the variable input arguments */
    /***************************************/
    /* Initialize the va_list */
    va_list vararg;
    va_start(vararg, nargs);
    /* Get the number of the x and y axis nodes for interpolation */
    n = (int) va_arg(vararg, double);
    /* Check if the number of input argument is plausible and define slopes */
    if (nargs == 1+2*n+1) {
        /* Plausible, but no slopes defined */
        e1 = 0; s1 = 0; /* set s1 to 0 but it is not used */
        e2 = 0; s2 = 0; /* set s2 to 0 but it is not used */
    } else if (nargs == 1+2*n+1+2) {
        /* Plausible, slopes are defined */
        e1 = 1; s1 = va_arg(vararg, double);
        e2 = 1; s2 = va_arg(vararg, double);
    } else {
        mexErrMsgTxt("interpcseSB_CVODE: Incorrect number of input arguments.");
    }
    
    /***********/
    /* If n==2 */
    /***********/
    if (n==2) {        
        if (e1 == 0 || e2 == 0) mexErrMsgTxt("interpcseSB: n=2 requires the definition of slopes.");
        x0 = va_arg(vararg, double);
        x1 = va_arg(vararg, double);
        y0 = va_arg(vararg, double);
        y1 = va_arg(vararg, double);

        /* Get the xx value at which to interpolate */
        xx = va_arg(vararg, double);
        /* End the variable input handling */
        va_end(vararg);

        b2 = s1;
        c2 = -(-3.0*y1+3.0*y0+(s2+2.0*s1)*x1+(-s2-2.0*s1)*x0)/(pow(x1,2.0)-2.0*x0*x1+pow(x0,2.0));
        d2 = (-2.0*y1+2.0*y0+(s2+s1)*x1+(-s2-s1)*x0)/(pow(x1,3.0)-3.0*x0*pow(x1,2.0)+3.0*pow(x0,2.0)*x1-pow(x0,3.0));
        yy = y0 + b2*(xx-x0) + c2*pow(xx-x0,2.0) + d2*pow(xx-x0,3.0);
    } else {
        /***********/
        /* If n!=2 */
        /***********/
        /* Allocate memory for x and y */
        x  = (double *) mxCalloc(n, sizeof(double));
        y  = (double *) mxCalloc(n, sizeof(double));
        /* Parse input arguments into x and y */
        for (k=0;k<n;k++) x[k] = va_arg(vararg, double);
        for (k=0;k<n;k++) y[k] = va_arg(vararg, double);
        
        /* Get the xx value at which to interpolate */
        xx = va_arg(vararg, double);
        /* End the variable input handling */
        va_end(vararg);
        
        /*****************************/
        /* Allocate memory for b,c,d */
        /*****************************/
        b  = (double *) mxCalloc(n, sizeof(double));
        c  = (double *) mxCalloc(n, sizeof(double));
        d  = (double *) mxCalloc(n, sizeof(double));
        
        /****************************/
        /* Call the spline function */
        /****************************/
        spline(n, e1, e2, s1, s2, x, y, b, c, d, &flag);
        
        /***************************************/
        /* Call the spline evaluation function */
        /***************************************/
        yy = seval(n, xx, x, y, b, c, d, &last);
        
        /***************/
        /* Free memory */
        /***************/
        mxFree(x);
        mxFree(y);
        mxFree(b);
        mxFree(c);
        mxFree(d);
    }
    
    /*********************************/
    /* Return the interpolated value */
    /*********************************/
    return yy;
}


/*
 *========================================================================
 * INTERFACE FUNCTION for interpcsexSlopeSB ... 
 *========================================================================
 */
double interpcsexSlopeSB(int nargs, ...)
{
    /*************************************************************/
    /* Define the variables needed for the spline function calls */
    /*************************************************************/
    int    n;
    double *x;
    double *y;
    double xx;
    double s1,s2;
    int    e1,e2;
    double *b;
    double *c;
    double *d;
    int    flag;
    int    last;
    int    k;
    double yy;
    double b2, c2, d2;
    double x0,x1,y0,y1;
    
    /***************************************/
    /* Handle the variable input arguments */
    /***************************************/
    /* Initialize the va_list */
    va_list vararg;
    va_start(vararg, nargs);
    /* Get the number of the x and y axis nodes for interpolation */
    n = (int) va_arg(vararg, double);
    /* Check if the number of input argument is plausible and define slopes */
    if (nargs == 1+2*n+1) {
        /* Plausible, but no slopes defined */
        e1 = 0; s1 = 0; /* set s1 to 0 but it is not used */
        e2 = 0; s2 = 0; /* set s2 to 0 but it is not used */
    } else if (nargs == 1+2*n+1+2) {
        /* Plausible, slopes are defined */
        e1 = 1; s1 = va_arg(vararg, double);
        e2 = 1; s2 = va_arg(vararg, double);
    } else {
        mexErrMsgTxt("interpcsexSlopeSB_CVODE: Incorrect number of input arguments.");
    }
    
    /***********/
    /* If n==2 */
    /***********/
    if (n==2) {        
        if (e1 == 0 || e2 == 0) mexErrMsgTxt("interpcsexSlopeSB: n=2 requires the definition of slopes.");
        x0 = va_arg(vararg, double);
        x1 = va_arg(vararg, double);
        y0 = va_arg(vararg, double);
        y1 = va_arg(vararg, double);

        /* Get the xx value at which to interpolate */
        xx = va_arg(vararg, double);
        /* End the variable input handling */
        va_end(vararg);

        b2 = s1;
        c2 = -(-3.0*y1+3.0*y0+(s2+2.0*s1)*x1+(-s2-2.0*s1)*x0)/(pow(x1,2.0)-2.0*x0*x1+pow(x0,2.0));
        d2 = (-2.0*y1+2.0*y0+(s2+s1)*x1+(-s2-s1)*x0)/(pow(x1,3.0)-3.0*x0*pow(x1,2.0)+3.0*pow(x0,2.0)*x1-pow(x0,3.0));
        yy = b2 + 2.0*c2*(xx-x0) + 3.0*d2*pow(xx-x0,2.0);
    } else {
        /***********/
        /* If n!=2 */
        /***********/
        /* Allocate memory for x and y */
        x  = (double *) mxCalloc(n, sizeof(double));
        y  = (double *) mxCalloc(n, sizeof(double));
        /* Parse input arguments into x and y */
        for (k=0;k<n;k++) x[k] = va_arg(vararg, double);
        for (k=0;k<n;k++) y[k] = va_arg(vararg, double);
        
        /* Get the xx value at which to interpolate */
        xx = va_arg(vararg, double);
        /* End the variable input handling */
        va_end(vararg);
        
        /*****************************/
        /* Allocate memory for b,c,d */
        /*****************************/
        b  = (double *) mxCalloc(n, sizeof(double));
        c  = (double *) mxCalloc(n, sizeof(double));
        d  = (double *) mxCalloc(n, sizeof(double));
        
        /****************************/
        /* Call the spline function */
        /****************************/
        spline(n, e1, e2, s1, s2, x, y, b, c, d, &flag);
        
        /***************************************/
        /* Call the spline evaluation function */
        /***************************************/
        yy = deriv(n, xx, x, b, c, d, &last);
        
        /***************/
        /* Free memory */
        /***************/
        mxFree(x);
        mxFree(y);
        mxFree(b);
        mxFree(c);
        mxFree(d);
    }
    
    /*********************************/
    /* Return the interpolated value */
    /*********************************/
    return yy;
}

/*=========================================================================
   Cubic spline coefficients
   -------------------------
   Evaluate the coefficients b[i], c[i], d[i], i = 0, 1, .. n-1 for
   a cubic interpolating spline

   S(xx) = Y[i] + b[i] * w + c[i] * w**2 + d[i] * w**3
   where w = xx - x[i]
   and   x[i] <= xx <= x[i+1]

   The n supplied data points are x[i], y[i], i = 0 ... n-1.

   Input :
   -------
   n       : The number of data points or knots (n >= 2)
   end1,
   end2    : = 1 to specify the slopes at the end points
             = 0 to obtain the default conditions
   slope1,
   slope2  : the slopes at the end points x[0] and x[n-1]
             respectively
   x[]     : the abscissas of the knots in strictly
             increasing order
   y[]     : the ordinates of the knots

   Output :
   --------
   b, c, d : arrays of spline coefficients as defined above
             (See note 2 for a definition.)
   iflag   : status flag
            = 0 normal return
            = 1 less than two data points; cannot interpolate
            = 2 x[] are not in ascending order

   This C code written by ...  Peter & Nigel,
   ----------------------      Design Software,
                               42 Gubberley St,
                               Kenmore, 4069,
                               Australia.

   Version ... 1.1, 30 September 1987
   -------     2.0, 6 April 1989    (start with zero subscript)
                                     remove ndim from parameter list
               2.1, 28 April 1989   (check on x[])
               2.2, 10 Oct   1989   change number order of matrix

   Notes ...
   -----
   (1) The accompanying function seval() may be used to evaluate the
       spline while deriv will provide the first derivative.
   (2) Using p to denote differentiation
       y[i] = S(X[i])
       b[i] = Sp(X[i])
       c[i] = Spp(X[i])/2
       d[i] = Sppp(X[i])/6  ( Derivative from the right )
   (3) Since the zero elements of the arrays ARE NOW used here,
       all arrays to be passed from the main program should be
       dimensioned at least [n].  These routines will use elements
       [0 .. n-1].
   (4) Adapted from the text
       Forsythe, G.E., Malcolm, M.A. and Moler, C.B. (1977)
       "Computer Methods for Mathematical Computations"
       Prentice Hall
   (5) Note that although there are only n-1 polynomial segments,
       n elements are requird in b, c, d.  The elements b[n-1],
       c[n-1] and d[n-1] are set to continue the last segment
       past x[n-1].
=========================================================================*/
void spline (int n, int end1, int end2,
            double slope1, double slope2,
            double x[], double y[],
            double b[], double c[], double d[],
            int *iflag)
{  /* begin procedure spline() */

int    nm1, ib, i;
double t;
int    ascend;

nm1    = n - 1;
*iflag = 0;

if (n < 2)
  {  /* no possible interpolation */
  *iflag = 1;
  return;
  }

ascend = 1;
for (i = 1; i < n; ++i) if (x[i] <= x[i-1]) ascend = 0;
if (!ascend)
   {
   *iflag = 2;
   return;
   }

if (n >= 3)
   {    /* ---- At least quadratic ---- */

   /* ---- Set up the symmetric tri-diagonal system
           b = diagonal
           d = offdiagonal
           c = right-hand-side  */
   d[0] = x[1] - x[0];
   c[1] = (y[1] - y[0]) / d[0];
   for (i = 1; i < nm1; ++i)
      {
      d[i]   = x[i+1] - x[i];
      b[i]   = 2.0 * (d[i-1] + d[i]);
      c[i+1] = (y[i+1] - y[i]) / d[i];
      c[i]   = c[i+1] - c[i];
      }

   /* ---- Default End conditions
           Third derivatives at x[0] and x[n-1] obtained
           from divided differences  */
   b[0]   = -d[0];
   b[nm1] = -d[n-2];
   c[0]   = 0.0;
   c[nm1] = 0.0;
   if (n != 3)
      {
      c[0]   = c[2] / (x[3] - x[1]) - c[1] / (x[2] - x[0]);
      c[nm1] = c[n-2] / (x[nm1] - x[n-3]) - c[n-3] / (x[n-2] - x[n-4]);
      c[0]   = c[0] * d[0] * d[0] / (x[3] - x[0]);
      c[nm1] = -c[nm1] * d[n-2] * d[n-2] / (x[nm1] - x[n-4]);
      }

   /* Alternative end conditions -- known slopes */
   if (end1 == 1)
      {
      b[0] = 2.0 * (x[1] - x[0]);
      c[0] = (y[1] - y[0]) / (x[1] - x[0]) - slope1;
      }
   if (end2 == 1)
      {
      b[nm1] = 2.0 * (x[nm1] - x[n-2]);
      c[nm1] = slope2 - (y[nm1] - y[n-2]) / (x[nm1] - x[n-2]);
      }

   /* Forward elimination */
   for (i = 1; i < n; ++i)
     {
     t    = d[i-1] / b[i-1];
     b[i] = b[i] - t * d[i-1];
     c[i] = c[i] - t * c[i-1];
     }

   /* Back substitution */
   c[nm1] = c[nm1] / b[nm1];
   for (ib = 0; ib < nm1; ++ib)
      {
      i    = n - ib - 2;
      c[i] = (c[i] - d[i] * c[i+1]) / b[i];
      }

   /* c[i] is now the sigma[i] of the text */

   /* Compute the polynomial coefficients */
   b[nm1] = (y[nm1] - y[n-2]) / d[n-2] + d[n-2] * (c[n-2] + 2.0 * c[nm1]);
   for (i = 0; i < nm1; ++i)
      {
      b[i] = (y[i+1] - y[i]) / d[i] - d[i] * (c[i+1] + 2.0 * c[i]);
      d[i] = (c[i+1] - c[i]) / d[i];
      c[i] = 3.0 * c[i];
      }
   c[nm1] = 3.0 * c[nm1];
   d[nm1] = d[n-2];

   }  /* at least quadratic */

else  /* if n >= 3 */
   {  /* linear segment only  */
   b[0] = (y[1] - y[0]) / (x[1] - x[0]);
   c[0] = 0.0;
   d[0] = 0.0;
   b[1] = b[0];
   c[1] = 0.0;
   d[1] = 0.0;
   }
}  /* end of spline() */


/*-------------------------------------------------------------------*/
/*-----------------------------------------------------------------*/
/*-----------------------------------------------------------------*/
double seval (int n, double u,
              double x[], double y[],
              double b[], double c[], double d[],
              int *last)
/*-----------------------------------------------------------------*/
/*Purpose ...
  -------
  Evaluate the cubic spline function

  S(xx) = y[i] + b[i] * w + c[i] * w**2 + d[i] * w**3
  where w = u - x[i]
  and   x[i] <= u <= x[i+1]
  Note that Horner's rule is used.
  If u < x[0]   then i = 0 is used.
  If u > x[n-1] then i = n-1 is used.

  Input :
  -------
  n       : The number of data points or knots (n >= 2)
  u       : the abscissa at which the spline is to be evaluated
  Last    : the segment that was last used to evaluate U
  x[]     : the abscissas of the knots in strictly increasing order
  y[]     : the ordinates of the knots
  b, c, d : arrays of spline coefficients computed by spline().

  Output :
  --------
  seval   : the value of the spline function at u
  Last    : the segment in which u lies

  Notes ...
  -----
  (1) If u is not in the same interval as the previous call then a
      binary search is performed to determine the proper interval.

*/
/*-------------------------------------------------------------------*/
{  /* begin function seval() */

int    i, j, k;
double w;

i = *last;
if (i >= n-1) i = 0;
if (i < 0)  i = 0;

if ((x[i] > u) || (x[i+1] < u))
  {  /* ---- perform a binary search ---- */
  i = 0;
  j = n;
  do
    {
    k = (i + j) / 2;         /* split the domain to search */
    if (u < x[k])  j = k;    /* move the upper bound */
    if (u >= x[k]) i = k;    /* move the lower bound */
    }                        /* there are no more segments to search */
  while (j > i+1);
  }
*last = i;

/* ---- Evaluate the spline ---- */
w = u - x[i];
w = y[i] + w * (b[i] + w * (c[i] + w * d[i]));
return (w);
}
/*-------------------------------------------------------------------*/

/*-------------------------------------------------------------------*/
/*-----------------------------------------------------------------*/
/*-----------------------------------------------------------------*/
double deriv (int n, double u,
              double x[],
              double b[], double c[], double d[],
              int *last)
/*-----------------------------------------------------------------*/
/* Purpose ...
   -------
   Evaluate the derivative of the cubic spline function

   S(x) = B[i] + 2.0 * C[i] * w + 3.0 * D[i] * w**2
   where w = u - X[i]
   and   X[i] <= u <= X[i+1]
   Note that Horner's rule is used.
   If U < X[0] then i = 0 is used.
   If U > X[n-1] then i = n-1 is used.

   Input :
   -------
   n       : The number of data points or knots (n >= 2)
   u       : the abscissa at which the derivative is to be evaluated
   last    : the segment that was last used
   x       : the abscissas of the knots in strictly increasing order
   b, c, d : arrays of spline coefficients computed by spline()

   Output :
   --------
   deriv : the value of the derivative of the spline
           function at u
   last  : the segment in which u lies

   Notes ...
   -----
   (1) If u is not in the same interval as the previous call then a
       binary search is performed to determine the proper interval.

*/
/*-------------------------------------------------------------------*/
{  /* begin function deriv() */

int    i, j, k;
double w;

i = *last;
if (i >= n-1) i = 0;
if (i < 0) i = 0;

if ((x[i] > u) || (x[i+1] < u))
  {  /* ---- perform a binary search ---- */
  i = 0;
  j = n;
  do
    {
    k = (i + j) / 2;          /* split the domain to search */
    if (u < x[k])  j = k;     /* move the upper bound */
    if (u >= x[k]) i = k;     /* move the lower bound */
    }                         /* there are no more segments to search */
  while (j > i+1);
  }
*last = i;

/* ---- Evaluate the derivative ---- */
w = u - x[i];
w = b[i] + w * (2.0 * c[i] + w * 3.0 * d[i]);
return (w);

} /* end of deriv() */
/*-------------------------------------------------------------------*/


/*************************************************************/
/*************************************************************/
/*************************************************************/
/*************************************************************/
/*************************************************************/
/* interpcsSB function ... MEX version for CVODE use in SBPD */
/*************************************************************/

void spline_pchip_set( int n, double x[], double f[], double d[] );
void spline_pchip_val( int n, double x[], double f[], double d[],  int ne, double xe, double *feptr );
int i4_max( int i1, int i2 );
int chfev( double x1, double x2, double f1, double f2, double d1, double d2, int ne, double xe, double *feptr, int next[] );
double r8_min( double x, double y );
double r8_max( double x, double y );
double pchst( double arg1, double arg2 );


/****************************************************************************/

double interpcsSB(int nargs, int n, double xe, ...)

/****************************************************************************

  Purpose:

    interpcsSB is the calling function to evaluate a piecewise cubic spline 
    function and calculte the Hermite interpolant.


  Modified:

    1 October 2007

  Author:

    Basti Bergdahl,
    Department of Applied Microbiology,
    Lund Insitute of Technology.

  Calling syntax:

    interpcsSB(NARGS, N, XE, X1,...,XN,F1,...,FN)

  Parameters:

    Input, int NARGS, the total number of input arguments (not counting NARGS itself).
    The number should be even, otherwise an error will occure.

    Input, int N, the number of data points (rows in the table).  N must be at least 2.

    Input, double XE, point at which the function is to be evaluated.

    Input, double X1,...,XN, the strictly increasing independent variable values.

    Input, double F1,...,FN, dependent variable values to be interpolated.  This
    routine is designed for monotonic data, but it will work for any F-array.
    It will force extrema at points where monotonicity switches direction.

    Output, double FE, the value of the cubic Hermite function at XE.
*/
{
    va_list vararg;
    int k, oddnumber;
    int ne = 1;
    double *data, *x, *f, *d, *feptr;
    double fe;

    /* Check if table data has been given */
    if (nargs == 2)
    {
        mexErrMsgTxt("interpcsSB error: Input incorrect. No data for the table has been supplied.");
        return 0;
    }

    /* Check if odd or even number of input arguments */
    oddnumber = nargs % 2;
    if (oddnumber != 0)
    {
        mexErrMsgTxt("interpcsSB error: Input incorrect. The number of inputs should be even.");
        return 0;
    }

    /* If number of input arguments in odd, check if
     * the X and Y vectors are of the same size */
    if ((nargs-2)/2 != n)
    {
        mexErrMsgTxt("interpcsSB error: Input incorrect. The X and Y vectors are either of different lengths or");
        mexErrMsgTxt("an incorrect number for the data points has been given.");
        return 0;
    }

    /* Read in all variable input arguments in double array */    
    va_start(vararg, xe);
    data = (double *)mxCalloc((n*2),sizeof(double));
    for (k=0; k<nargs-2; k++ )
    {
        data[k] = va_arg(vararg, double);
    }

    /* Create the X and Y arrays making up the table */
    x = (double *)mxCalloc(n,sizeof(double));
    f = (double *)mxCalloc(n,sizeof(double));
    for (k=0; k<n; k++)
    {
        x[k] = data[k];
        f[k] = data[k+n];
    }
    mxFree(data); /* free temporary array */

    /* handle off limit values */
    if (xe < x[0]) return(f[0]);
    if (xe > x[n-1]) return(f[n-1]);
    
    /* Calculate the derivatives for the piecewise cubic Hermite interpolant */
    d = (double *)mxCalloc(n,sizeof(double));
    spline_pchip_set(n, x, f, d);

    /* Evaluate the cubic Hermite function at x */    
    feptr = &fe;
    spline_pchip_val(n, x, f, d, ne, xe, feptr);

    return(fe);
}


/****************************************************************************/

void spline_pchip_set( int n, double x[], double f[], double d[] )

/****************************************************************************

  Purpose:

    SPLINE_PCHIP_SET sets derivatives for a piecewise cubic Hermite interpolant.

  Discussion:

    This routine computes what would normally be called a Hermite
    interpolant.  However, the user is only required to supply function
    values, not derivative values as well.  This routine computes
    "suitable" derivative values, so that the resulting Hermite interpolant
    has desirable shape and monotonicity properties.

    The interpolant will have an extremum at each point where
    monotonicity switches direction.

    The resulting piecewise cubic Hermite function may be evaluated
    by SPLINE_PCHIP_VAL..

    This routine was originally called "PCHIM".

  Modified:

    14 August 2005

  Author:

    Fred Fritsch,
    Mathematics and Statistics Division,
    Lawrence Livermore National Laboratory.

    C++ translation by John Burkardt.

  Reference:

    Fred Fritsch, Ralph Carlson,
    Monotone Piecewise Cubic Interpolation,
    SIAM Journal on Numerical Analysis,
    Volume 17, Number 2, April 1980, pages 238-246.

    Fred Fritsch, Judy Butland,
    A Method for Constructing Local Monotone Piecewise
    Cubic Interpolants,
    SIAM Journal on Scientific and Statistical Computing,
    Volume 5, Number 2, 1984, pages 300-304.

  Parameters:

    Input, int N, the number of data points.  N must be at least 2.

    Input, double X[N], the strictly increasing independent
    variable values.

    Input, double F[N], dependent variable values to be interpolated.  This
    routine is designed for monotonic data, but it will work for any F-array.
    It will force extrema at points where monotonicity switches direction.

    Output, double D[N], the derivative values at the
    data points.  If the data are monotonic, these values will determine
    a monotone cubic Hermite function.
*/
{
  double del1;
  double del2;
  double dmax;
  double dmin;
  double drat1;
  double drat2;
  double dsave;
  double h1;
  double h2;
  double hsum;
  double hsumt3;
  int i;
  int ierr;
  int nless1;
  double temp;
  double w1;
  double w2;

  /*  Check the arguments. */

  if ( n < 2 )
  {
    ierr = -1;
    mexErrMsgTxt("\n");
    mexErrMsgTxt("SPLINE_PCHIP_SET - Fatal error!\n");
    mexErrMsgTxt("  Number of data points less than 2.\n");
    exit ( ierr );
  }

  for ( i = 1; i < n; i++ )
  {
    if ( x[i] <= x[i-1] )
    {
      ierr = -3;
      mexErrMsgTxt("\n");
      mexErrMsgTxt("SPLINE_PCHIP_SET - Fatal error!\n");
      mexErrMsgTxt("  X array not strictly increasing.\n");
      exit ( ierr );
    }
  }

  ierr = 0;
  nless1 = n - 1;
  h1 = x[1] - x[0];
  del1 = ( f[1] - f[0] ) / h1;
  dsave = del1;

/*  Special case N=2, use linear interpolation. */

  if ( n == 2 )
  {
    d[0] = del1;
    d[n-1] = del1;
    return;
  }

/*  Normal case, 3 <= N. */

  h2 = x[2] - x[1];
  del2 = ( f[2] - f[1] ) / h2;

/*  Set D(1) via non-centered three point formula, adjusted to be
    shape preserving. */

  hsum = h1 + h2;
  w1 = ( h1 + hsum ) / hsum;
  w2 = -h1 / hsum;
  d[0] = w1 * del1 + w2 * del2;

  if ( pchst ( d[0], del1 ) <= 0.0 )
  {
    d[0] = 0.0;
  }

/*  Need do this check only if monotonicity switches. */

  else if ( pchst ( del1, del2 ) < 0.0 )
  {
     dmax = 3.0 * del1;

     if ( fabs ( dmax ) < fabs ( d[0] ) )
     {
       d[0] = dmax;
     }

  }

/*  Loop through interior points. */

  for ( i = 2; i <= nless1; i++ )
  {
    if ( 2 < i )
    {
      h1 = h2;
      h2 = x[i] - x[i-1];
      hsum = h1 + h2;
      del1 = del2;
      del2 = ( f[i] - f[i-1] ) / h2;
    }

/*  Set D(I)=0 unless data are strictly monotonic. */

    d[i-1] = 0.0;

    temp = pchst ( del1, del2 );

    if ( temp < 0.0 )
    {
      ierr = ierr + 1;
      dsave = del2;
    }

/*  Count number of changes in direction of monotonicity. */

    else if ( temp == 0.0 )
    {
      if ( del2 != 0.0 )
      {
        if ( pchst ( dsave, del2 ) < 0.0 )
        {
          ierr = ierr + 1;
        }
        dsave = del2;
      }
    }

/*  Use Brodlie modification of Butland formula. */

    else
    {
      hsumt3 = 3.0 * hsum;
      w1 = ( hsum + h1 ) / hsumt3;
      w2 = ( hsum + h2 ) / hsumt3;
      dmax = r8_max ( fabs ( del1 ), fabs ( del2 ) );
      dmin = r8_min ( fabs ( del1 ), fabs ( del2 ) );
      drat1 = del1 / dmax;
      drat2 = del2 / dmax;
      d[i-1] = dmin / ( w1 * drat1 + w2 * drat2 );
    }
  }

/*  Set D(N) via non-centered three point formula, adjusted to be
    shape preserving. */

  w1 = -h2 / hsum;
  w2 = ( h2 + hsum ) / hsum;
  d[n-1] = w1 * del1 + w2 * del2;

  if ( pchst ( d[n-1], del2 ) <= 0.0 )
  {
    d[n-1] = 0.0;
  }
  else if ( pchst ( del1, del2 ) < 0.0 )
  {

/*  Need do this check only if monotonicity switches. */

    dmax = 3.0 * del2;

    if ( fabs ( dmax ) < abs ( d[n-1] ) )
    {
      d[n-1] = dmax;
    }

  }
  return;
}

/****************************************************************************/

void spline_pchip_val( int n, double x[], double f[], double d[],
  int ne, double xe, double *feptr )

/****************************************************************************

  Purpose:

    SPLINE_PCHIP_VAL evaluates a piecewise cubic Hermite function.

  Description:

    This routine may be used by itself for Hermite interpolation, or as an
    evaluator for SPLINE_PCHIP_SET.

    This routine evaluates the cubic Hermite function at the points XE.

    Most of the coding between the call to CHFEV and the end of
    the IR loop could be eliminated if it were permissible to
    assume that XE is ordered relative to X.

    CHFEV does not assume that X1 is less than X2.  Thus, it would
    be possible to write a version of SPLINE_PCHIP_VAL that assumes a strictly
    decreasing X array by simply running the IR loop backwards
    and reversing the order of appropriate tests.

    The present code has a minor bug, which I have decided is not
    worth the effort that would be required to fix it.
    If XE contains points in [X(N-1),X(N)], followed by points less than
    X(N-1), followed by points greater than X(N), the extrapolation points
    will be counted (at least) twice in the total returned in IERR.

    The evaluation will be most efficient if the elements of XE are
    increasing relative to X; that is, for all J <= K,
      X(I) <= XE(J)
    implies
      X(I) <= XE(K).

    If any of the XE are outside the interval [X(1),X(N)],
    values are extrapolated from the nearest extreme cubic,
    and a warning error is returned.

    This routine was originally named "PCHFE".

  Modified:

    14 August 2005

  Author:

    Fred Fritsch,
    Mathematics and Statistics Division,
    Lawrence Livermore National Laboratory.

    C++ translation by John Burkardt.

  Reference:

    Fred Fritsch, Ralph Carlson,
    Monotone Piecewise Cubic Interpolation,
    SIAM Journal on Numerical Analysis,
    Volume 17, Number 2, April 1980, pages 238-246.

  Parameters:

    Input, int N, the number of data points.  N must be at least 2.

    Input, double X[N], the strictly increasing independent
    variable values.

    Input, double F[N], the function values.

    Input, double D[N], the derivative values.

    Input, int NE, the number of evaluation points.

    Input, double XE, point at which the function is to
    be evaluated.

    Output, double FE, the value of the cubic Hermite
    function at XE.
*/
{
  int i;
  int ierc;
  int ierr;
  int ir;
  int j;
  int j_first;
  int j_new;
  int j_save;
  int next[2];
  int nj;

/*  Check arguments. */

  if ( n < 2 )
  {
    ierr = -1;
    mexErrMsgTxt("\n");
    mexErrMsgTxt("SPLINE_PCHIP_VAL - Fatal error!\n");
    mexErrMsgTxt("Number of data points less than 2.\n");
    exit ( ierr );
  }

  for ( i = 1; i < n; i++ )
  {
    if ( x[i] <= x[i-1] )
    {
      ierr = -3;
      mexErrMsgTxt("\n");
      mexErrMsgTxt("SPLINE_PCHIP_VAL - Fatal error!\n");
      mexErrMsgTxt("X array not strictly increasing.\n");
      exit ( ierr );
    }
  }

  if ( ne < 1 )
  {
    ierr = -4;
    mexErrMsgTxt("\n");
    mexErrMsgTxt("SPLINE_PCHIP_VAL - Fatal error!\n");
    mexErrMsgTxt("Number of evaluation points less than 1.\n");
    return;
  }

  ierr = 0;

/*  Loop over intervals.
  The interval index is IL = IR-1.
  The interval is X(IL) <= X < X(IR).
*/
  j_first = 1;
  ir = 2;

  for ( ; ; )
  {
/*
  Skip out of the loop if have processed all evaluation points.
*/
    if ( ne < j_first )
    {
      break;
    }
/*
  Locate all points in the interval.
*/
    j_save = ne + 1;

    for ( j = j_first; j <= ne; j++ )
    {
      if ( x[ir-1] <= xe )
      {
        j_save = j;
        if ( ir == n )
        {
          j_save = ne + 1;
        }
        break;
      }
    }
/*
  Have located first point beyond interval.
*/
    j = j_save;

    nj = j - j_first;
/*
  Skip evaluation if no points in interval.
*/
    if ( nj != 0 )
    {
/*
  Evaluate cubic at XE(J_FIRST:J-1).
*/
      ierc = chfev ( x[ir-2], x[ir-1], f[ir-2], f[ir-1], d[ir-2], d[ir-1],
        nj, xe, feptr, next );

      if ( ierc < 0 )
      {
        ierr = -5;
        mexErrMsgTxt("\n");
        mexErrMsgTxt("SPLINE_PCHIP_VAL - Fatal error!\n");
        mexErrMsgTxt("Error return from CHFEV.\n");
        exit ( ierr );
      }
/*
  In the current set of XE points, there are NEXT(2) to the right of X(IR).
*/
      if ( next[1] != 0 )
      {
        if ( ir < n )
        {
          ierr = -5;
          mexErrMsgTxt("\n");
          mexErrMsgTxt("SPLINE_PCHIP_VAL - Fatal error!\n");
          mexErrMsgTxt("IR < N.\n");
          exit ( ierr );
        }
/*
  These are actually extrapolation points.
*/
        ierr = ierr + next[1];

      }
/*
  In the current set of XE points, there are NEXT(1) to the left of X(IR-1).
*/
      if ( next[0] != 0 )
      {
/*
  These are actually extrapolation points.
*/
        if ( ir <= 2 )
        {
          ierr = ierr + next[0];
        }
        else
        {
          j_new = -1;

          for ( i = j_first; i <= j-1; i++ )
          {
            if ( xe < x[ir-2] )
            {
              j_new = i;
              break;
            }
          }

          if ( j_new == -1 )
          {
            ierr = -5;
            mexErrMsgTxt("\n");
            mexErrMsgTxt("SPLINE_PCHIP_VAL - Fatal error!\n");
            mexErrMsgTxt("  Could not bracket the data point.\n");
            exit ( ierr );
          }
/*
  Reset J.  This will be the new J_FIRST.
*/
          j = j_new;
/*
  Now find out how far to back up in the X array.
*/
          for ( i = 1; i <= ir-1; i++ )
          {
            if ( xe < x[i-1] )
            {
              break;
            }
          }
/*
  At this point, either XE(J) < X(1) or X(i-1) <= XE(J) < X(I) .

  Reset IR, recognizing that it will be incremented before cycling.
*/
          ir = i4_max ( 1, i-1 );
        }
      }

      j_first = j;
    }

    ir = ir + 1;

    if ( n < ir )
    {
      break;
    }

  }

  return;
}

/****************************************************************************/

int chfev( double x1, double x2, double f1, double f2, double d1, double d2,
  int ne, double xe, double *feptr, int next[] )

/****************************************************************************

  Purpose:

    CHFEV evaluates a cubic polynomial given in Hermite form.

  Discussion:

    This routine evaluates a cubic polynomial given in Hermite form at an
    array of points.  While designed for use by SPLINE_PCHIP_VAL, it may
    be useful directly as an evaluator for a piecewise cubic
    Hermite function in applications, such as graphing, where
    the interval is known in advance.

    The cubic polynomial is determined by function values
    F1, F2 and derivatives D1, D2 on the interval [X1,X2].

  Modified:

    12 August 2005

  Author:

    Fred Fritsch,
    Mathematics and Statistics Division,
    Lawrence Livermore National Laboratory.

    C++ translation by John Burkardt.

  Reference:

    Fred Fritsch, Ralph Carlson,
    Monotone Piecewise Cubic Interpolation,
    SIAM Journal on Numerical Analysis,
    Volume 17, Number 2, April 1980, pages 238-246.

    David Kahaner, Cleve Moler, Steven Nash,
    Numerical Methods and Software,
    Prentice Hall, 1989,
    ISBN: 0-13-627258-4,
    LC: TA345.K34.

  Parameters:

    Input, double X1, X2, the endpoints of the interval of
    definition of the cubic.  X1 and X2 must be distinct.

    Input, double F1, F2, the values of the function at X1 and
    X2, respectively.

    Input, double D1, D2, the derivative values at X1 and
    X2, respectively.

    Input, int NE, the number of evaluation points.

    Input, double XE, the point at which the function is to
    be evaluated.  If the value of XE is outside the interval
    [X1,X2], a warning error is returned in NEXT.

    Output, double FE, the value of the cubic function
    at the point XE.

    Output, int NEXT[2], indicates the number of extrapolation points:
    NEXT[0] = number of evaluation points to the left of interval.
    NEXT[1] = number of evaluation points to the right of interval.

    Output, int CHFEV, error flag.
    0, no errors.
    -1, NE < 1.
    -2, X1 == X2.
*/
{
  double c2;
  double c3;
  double del1;
  double del2;
  double delta;
  double h;
  int ierr;
  double x;
  double fe;
  double xma;
  double xmi;

  if ( ne < 1 )
  {
    ierr = -1;
    mexErrMsgTxt("\n");
    mexErrMsgTxt("CHFEV - Fatal error!\n");
    mexErrMsgTxt("  Number of evaluation points is less than 1.\n");
    printf("  NE = %d\n", ne);
    return ierr;
  }

  h = x2 - x1;

  if ( h == 0.0 )
  {
    ierr = -2;
    mexErrMsgTxt("\n");
    mexErrMsgTxt("CHFEV - Fatal error!\n");
    mexErrMsgTxt("  The interval [X1,X2] is of zero length.\n");
    return ierr;
  }
/*
  Initialize.
*/
  ierr = 0;
  next[0] = 0;
  next[1] = 0;
  xmi = r8_min ( 0.0, h );
  xma = r8_max ( 0.0, h );
/*
  Compute cubic coefficients expanded about X1.
*/
  delta = ( f2 - f1 ) / h;
  del1 = ( d1 - delta ) / h;
  del2 = ( d2 - delta ) / h;
  c2 = -( del1 + del1 + del2 );
  c3 = ( del1 + del2 ) / h;
    x = xe - x1;
    fe = f1 + x * ( d1 + x * ( c2 + x * c3 ) );
    *feptr = fe;
/*
  Count the extrapolation points.
*/
    if ( x < xmi )
    {
      next[0] = next[0] + 1;
    }

    if ( xma < x )
    {
      next[1] = next[1] + 1;
    }


  return 0;
}

/****************************************************************************/

double pchst( double arg1, double arg2 )

/****************************************************************************

  Purpose:

    PCHST: PCHIP sign-testing routine.

  Discussion:

    This routine essentially computes the sign of ARG1 * ARG2.

    The object is to do this without multiplying ARG1 * ARG2, to avoid
    possible over/underflow problems.

  Modified:

    12 August 2005

  Author:

    Fred Fritsch,
    Mathematics and Statistics Division,
    Lawrence Livermore National Laboratory.

    C++ translation by John Burkardt.

  Reference:

    Fred Fritsch, Ralph Carlson,
    Monotone Piecewise Cubic Interpolation,
    SIAM Journal on Numerical Analysis,
    Volume 17, Number 2, April 1980, pages 238-246.

  Parameters:

    Input, double ARG1, ARG2, two values to check.

    Output, double PCHST,
    -1.0, if ARG1 and ARG2 are of opposite sign.
     0.0, if either argument is zero.
    +1.0, if ARG1 and ARG2 are of the same sign.
*/
{
  double value;

  if ( arg1 == 0.0 )
  {
    value = 0.0;
  }
  else if ( arg1 < 0.0 )
  {
    if ( arg2 < 0.0 )
    {
      value = 1.0;
    }
    else if ( arg2 == 0.0 )
    {
      value = 0.0;
    }
    else if ( 0.0 < arg2 )
    {
      value = -1.0;
    }
  }
  else if ( 0.0 < arg1 )
  {
    if ( arg2 < 0.0 )
    {
      value = -1.0;
    }
    else if ( arg2 == 0.0 )
    {
      value = 0.0;
    }
    else if ( 0.0 < arg2 )
    {
      value = 1.0;
    }
  }

  return(value);
}

/****************************************************************************/

double r8_max( double x, double y )

/****************************************************************************

  Purpose:

    R8_MAX returns the maximum of two R8's.

  Modified:

    10 January 2002

  Author:

    John Burkardt

  Parameters:

    Input, double X, Y, the quantities to compare.

    Output, double R8_MAX, the maximum of X and Y.
*/
{
  if ( y < x )
  {
    return(x);
  }
  else
  {
    return(y);
  }
}
/****************************************************************************/

double r8_min( double x, double y )

/****************************************************************************

  Purpose:

    R8_MIN returns the minimum of two R8's.

  Modified:

    09 May 2003

  Author:

    John Burkardt

  Parameters:

    Input, double X, Y, the quantities to compare.

    Output, double R8_MIN, the minimum of X and Y.
*/
{
  if ( y < x )
  {
    return(y);
  }
  else
  {
    return(x);
  }
}


/****************************************************************************/

int i4_max ( int i1, int i2 )

/****************************************************************************

  Purpose:

    I4_MAX returns the maximum of two I4's.

  Modified:

    13 October 1998

  Author:

    John Burkardt

  Parameters:

    Input, int I1, I2, are two integers to be compared.

    Output, int I4_MAX, the larger of I1 and I2.

*/
{
  if ( i2 < i1 )
  {
    return(i1);
  }
  else
  {
    return(i2);
  }

}
```
