## Supplementary scripts for "A multi-scale *in silico* mouse model for insulin resistance and humanoid type 2 diabetes": buildCVODElib.m

```
% This script generates a static library file for the CVODES integrator 
% package, including the MEX interface for MATLAB. Windows Version.
% Compiler used: LCC (MATLAB inbuild)
%
% Usually the user does not need to run this script, since for windows the 
% precompiled library is distributed along with the SBPD package.
% However, if the user wants to change the source code of the interface the
% new library can be build by running this script. The library file and 
% the header files are then automatically copied/installed at the correct 
% locations.

% Information:
% ============
% SBPD Package - Systems Biology Parameter Determination Package
% Copyright 2008 by Henning Schmidt,


% Copy files
disp('Copying files ...');
!copy CVODEmex25.c source_SUNDIALS_CVODES_25
!copy CVODEmex25.h include_SUNDIALS_CVODES_25
!copy mexmathaddon.h include_SUNDIALS_CVODES_25
!copy mexsplineaddon.h include_SUNDIALS_CVODES_25

% Change into source folder
cd source_SUNDIALS_CVODES_25

% Compile all files to object code using MinGW
disp('Compiling files ...');
mex -O -c -I../include_SUNDIALS_CVODES_25 CVODEmex25.c cvodea.c cvodea_io.c cvodes.c cvodes_band.c cvodes_bandpre.c cvodes_bbdpre.c cvodes_dense.c cvodes_diag.c cvodes_io.c cvodes_spbcgs.c cvodes_spbcgs.c cvodes_spils.c cvodes_sptfqmr.c nvector_serial.c sundials_band.c sundials_dense.c sundials_iterative.c sundials_math.c sundials_nvector.c sundials_smalldense.c sundials_spbcgs.c sundials_spgmr.c sundials_sptfqmr.c

% Delete the copied files again
disp('Deleting copied files ');
delete CVODEmex25.c
delete ../include_SUNDIALS_CVODES_25/CVODEmex25.h
delete ../include_SUNDIALS_CVODES_25/mexmathaddon.h
delete ../include_SUNDIALS_CVODES_25/mexsplineaddon.h

% Bind the object code files into a library.
disp('Generating library ');
% Use different ar's for 32 and 64 bit systems
if ~isempty(strfind(mexext,'32')),
    % 32 bit system
    system('"../ar32" rc CVODEmex25.lib *.obj');
elseif ~isempty(strfind(mexext,'64')),
    % 64 bit system
    system('"../ar64" rc CVODEmex25.lib CVODEmex25.obj cvodes_band.obj cvodes_diag.obj cvodes_sptfqmr.obj sundials_iterative.obj sundials_spbcgs.obj cvodea.obj cvodes_bandpre.obj cvodes_io.obj nvector_serial.obj sundials_math.obj sundials_spgmr.obj cvodea_io.obj cvodes_bbdpre.obj cvodes_spbcgs.obj sundials_band.obj sundials_nvector.obj sundials_sptfqmr.obj cvodes.obj cvodes_dense.obj cvodes_spils.obj sundials_dense.obj sundials_smalldense.obj');
else
    error('You got a strange system from the future ... or from the past?');
end

% Delete all object code files
disp('Deleting object files ');
delete *.obj

% Move generated lib file to parent folder
disp('Finishing ');
!move *.lib ../../lib
cd ..

% copy needed header files into the include folder of the package
!copy CVODEmex25.h "../include"
!copy mexmathaddon.h "../include"
!copy kineticformulas.h "../include"
!copy mexsplineaddon.h "../include"

% display success message
disp('The library has been build and all necessary files have been installed.');
disp(' ');
```
