## Supplementary scripts for "A multi-scale *in silico* mouse model for insulin resistance and humanoid type 2 diabetes": buildCVODElibunix.m

```
% This script generates a static library file for the CVODES integrator 
% package, including the MEX interface for MATLAB. Unix/Linux Version.
%
% Before using the SBaddon package to create MEX simulation functions the 
% user needs to build a static library that contains the SUNDIALS CVODES
% package and the MEX interface functions. This library can be build by
% running this script. The library file and the header files are then
% automatically copied/installed at the correct locations.

% Information:
% ============
% SBPD Package - Systems Biology Parameter Determination Package
% Copyright 2008 by Henning Schmidt,


% Copy files
disp('Copying files ...');
!cp CVODEmex25.c source_SUNDIALS_CVODES_25
!cp CVODEmex25.h include_SUNDIALS_CVODES_25
!cp mexmathaddon.h include_SUNDIALS_CVODES_25
!cp mexsplineaddon.h include_SUNDIALS_CVODES_25

% Change into source folder
cd source_SUNDIALS_CVODES_25

% Compile all files to object code
disp('Compiling files ...');
mex -c -O -I../include_SUNDIALS_CVODES_25 ...
    CVODEmex25.c ...   
    cvodea.c                cvodea_io.c ...
    cvodes.c                cvodes_band.c ...
    cvodes_bandpre.c        cvodes_bbdpre.c ...
    cvodes_dense.c          cvodes_diag.c ...
    cvodes_io.c             cvodes_spbcgs.c ...
    cvodes_spbcgs.c         cvodes_spils.c ...
    cvodes_sptfqmr.c        nvector_serial.c ...
    sundials_band.c         ...
    sundials_dense.c        sundials_iterative.c ...
    sundials_math.c         sundials_nvector.c ...
    sundials_smalldense.c   sundials_spbcgs.c ...
    sundials_spgmr.c        sundials_sptfqmr.c ...

% Delete the copied files again
disp('Deleting copied files ...');
delete CVODEmex25.c
delete ../include_SUNDIALS_CVODES_25/CVODEmex25.h
delete ../include_SUNDIALS_CVODES_25/mexmathaddon.h
delete ../include_SUNDIALS_CVODES_25/mexsplineaddon.h

% Bind the object code files into a library.
% To be able to do that the lib.exe needs to be 
% in the system path.
disp('Generating library ...');
!ar rc CVODEmex25.a *.o

% Delete all object code files
disp('Deleting object files ...');
delete *.o

% Move generated lib file to parent folder
disp('Finishing ...');
!mv *.a ../../lib
cd ..

% copy needed header files into the include folder of the package
!cp CVODEmex25.h ../include
!cp mexmathaddon.h ../include
!cp kineticformulas.h ../include
!cp mexsplineaddon.h "../include"

% display success message
disp('The library has been build and all necessary files have been installed.');
disp(' ');
```
