## Supplementary scripts for "A multi-scale *in silico* mouse model for insulin resistance and humanoid type 2 diabetes": CVODEmex25.c

```
/*
 * CVODEmex25.c: MEX/CVODES Interface for Sundials CVODES version 2.5
 *
 * Information:
 * ============
 * SBPD Package - Systems Biology Parameter Determination Package
 * Copyright 2008 by Henning Schmidt,
 */

#include <mex.h>
#include <stdio.h>
#include <stdlib.h>
#include <stddef.h>
#include <stdarg.h>
#include <string.h>
#include <matrix.h>
#include <math.h>

/* CVODES includes */
#include <cvodes/cvodes.h>
#include <nvector/nvector_serial.h>
#include <sundials/sundials_types.h>
#include <sundials/sundials_math.h>
#include "CVODEmex25.h"

/* Definitions for CVODES */
#define ABSTOL  1.0e-6  
#define RELTOL  1.0e-6  
#define MAXNUMSTEPS     100000  /* default in CVODE is 500 but higher is better for general purpose */
#define MAXERRTESTFAILS 50      /* default is 7 */
#define Ith(v,i) NV_Ith_S(v,i-1)

/* Initialize functions */
static void doInitilizationVariables();
static mxArray* handleInputArguments(int nlhs, mxArray *plhs[], int nrhs, const mxArray *prhs[]);
static mxArray* doXdotCalc();
static void allocSimMemory();
static void initCVODE();
static void integrate();
static void reportSTATS();
static mxArray* constructOutput();
static void addVec2Mat(double *matrix, double *rowvector, int row, int nrows, int ncols);
static int f(double time, N_Vector u, N_Vector udot, void *f_data);
static int g(double time, N_Vector y, double *gout, void *g_data);
static void errorMsg(char *text);
static void freeMem();

/* Some global variables for the interface */
int k,k2;   /* some loop variables */

int length2check;

char stringbuffer[256];

mxArray *timesimvectorMX = NULL;
double *timesimvector = NULL;

mxArray *parametervectorMX = NULL;
double *parametervector = NULL;
double *parametervectorCVODES = NULL; 

mxArray *initialconditionsMX = NULL;
double *initialconditions = NULL;

mxArray *optionsMX = NULL;

mxArray *showIntegratorStatsMX = NULL; 
int showIntegratorStats = 0;            /* Flag defining if integrator stats shown or not */


mxArray *minstepMX = NULL;
double minstep;
mxArray *maxstepMX = NULL;
double maxstep;
mxArray *maxnumstepsMX = NULL;
long int maxnumsteps;
mxArray *reltolMX = NULL;
double reltol;
mxArray *abstolMX = NULL;
double abstol;
mxArray *maxerrtestfailsMX = NULL;
int maxerrtestfails;
mxArray *maxorderMX = NULL;
int maxorder;
mxArray *maxconvfailsMX = NULL;
int maxconvfails;
mxArray *initstepMX = NULL;
double initstep;
mxArray *maxnonlineariterMX = NULL;
int maxnonlineariter;


mxArray *xdotcalcMX = NULL;
double xdotcalc;

int numbertimesteps;

mxArray *resultMX = NULL;
double *result = NULL;

mxArray *statesMX = NULL;

mxArray *statevaluesMX = NULL;
double *statevalues = NULL;

mxArray *variablesMX = NULL;

mxArray *variablevaluesMX = NULL;
double *variablevalues = NULL;

mxArray *reactionsMX = NULL;

mxArray *reactionvaluesMX = NULL;
double *reactionvalues = NULL;

mxArray *eventsMX = NULL;

mxArray *eventtimesMX = NULL;
double *eventtimes = NULL;

mxArray *eventflagsMX = NULL;
double *eventflags = NULL;

/* CVODES needed data */
ParamData modeldata;
N_Vector u = NULL;
void *cvode_mem;
int flag, flagroot;
double treturn;
double tendstep;

/* Interface to model RHS */
double *statevec = NULL;
double *variablevec = NULL;
double *reactionvec = NULL;
int *eventvec = NULL;
double *eventdataold = NULL;
int nreventshappend;
double *eventflagsdata = NULL;
double eventcorrecttest;

/* Error Flags */
int timevectorEmpty = 0;
int timevectorNotVector = 0;
int timevectorIsScalar = 0;

/* Method (stiff, nonstiff) Variables */
mxArray *methodMX = NULL;
int method;

/* Integrator Statistics */
long int STATS_nsteps;
long int STATS_nfevals;
long int STATS_netfails;
double STATS_hinused;
double STATS_tolsfac;
  
/*
 *========================================================================
 * MEX INTERFACE FOR MEX SIMULATION MODELS
 *========================================================================
 */
void CVODEmex25(int nlhs, mxArray *plhs[], int nrhs, const mxArray *prhs[])
{
    /* Initialize variables that are needed externally */
    interpcseSB_check = NULL; /* needed for spline function in mexsplineaddon.h */
    interpcseSlopeSB_check = NULL; /* needed for spline function in mexsplineaddon.h */

    /* Initialize all other variables that are initialized during their definition
     * Needs to be redone since not re-defined/initialized if MEX file recalled */
    doInitilizationVariables();
    
    /* Handle variable input arguments */
    resultMX = handleInputArguments(nlhs, plhs, nrhs, prhs);
    if (resultMX != NULL) {
        /* No integration requested */
        plhs[0] = resultMX;
        return;
    }
    
    /* Check if timevector defined */
    if (timevectorEmpty == 1) mexErrMsgTxt("Timevector is empty.");
        
    /* check if integration or xdot calculation */
    if (xdotcalc != 0) {
        /* return the RHS of the ODEs at given state and parameter values */
        resultMX = doXdotCalc();
        plhs[0] = resultMX;
        return;
    }
    
    /* Do error checks ... timevector needs to be defined if integration is to be done! */
    if (timevectorNotVector == 1 || timevectorIsScalar == 1) mexErrMsgTxt("'timevector' input argument needs to be a vector if you want to do a simulation.");
    /* Otherwise do the integration */
    /* Allocate memory for the simulation */
    allocSimMemory();
    /* Initialize CVODES */
    initCVODE();
    /* Integrate */
    integrate();
    /* Report Statistics of Integration */
    reportSTATS();
    /* Construct result */
    plhs[0] = constructOutput();
    /* Free allocated memory */
    freeMem();
}

/*
 *========================================================================
 * Integration statistics report function
 *========================================================================
 */
static void reportSTATS()
{
    /* Get the statistics */
    CVodeGetNumSteps(cvode_mem,&STATS_nsteps);
    CVodeGetNumRhsEvals(cvode_mem,&STATS_nfevals);
    CVodeGetNumErrTestFails(cvode_mem,&STATS_netfails);
    CVodeGetActualInitStep(cvode_mem,&STATS_hinused);
    CVodeGetTolScaleFactor(cvode_mem,&STATS_tolsfac);
    /* Report statistics */
    if (showIntegratorStats != 0) {
        mexPrintf("\nIntegrator Statistics\n");
        mexPrintf("=====================\n");
        mexPrintf("Cumulative number of internal steps:    %ld\n",STATS_nsteps);
        mexPrintf("No. of calls to r.h.s. function:        %ld\n",STATS_nfevals);
        mexPrintf("No. of local error test failures:       %ld\n",STATS_netfails);
        mexPrintf("Actual init step size used:             %g\n",STATS_hinused);
        mexPrintf("Suggested factor for tolerance scaling: %g\n\n",STATS_tolsfac);
    }
}

/*
 *========================================================================
 * Help function to add a row vector to a matrix
 *========================================================================
 */
static void addVec2Mat(double *matrix, double *rowvector, int row, int nrrows, int nrcols)
{
    int k;
    for (k=0;k<nrcols;k++) {
        matrix[row+k*nrrows] = rowvector[k];
    }
}

/*
 *========================================================================
 * RHS function f(t,u) 
 *========================================================================
 */
static int f(double time, N_Vector u, N_Vector udot, void *f_data)
{
    double *statevec, *DDTvector;
    ParamData *paramdataPtr;
    /* get pointer to modeldata */
    paramdataPtr = (ParamData*) f_data;
    /* connect input and result data */
    statevec = NV_DATA_S(u);
    DDTvector = NV_DATA_S(udot);
    /* run the model */
    model(time, statevec, DDTvector, paramdataPtr, DOFLAG_DDT, NULL, NULL, NULL, NULL);
    return(0);
}

/*
 *========================================================================
 * Event function 
 *========================================================================
 */
static int g(double time, N_Vector y, double *gout, void *g_data)
{
    double *statevec;
    ParamData *paramdataPtr;
    /* get pointer to model data */
    paramdataPtr = (ParamData*) g_data;
    /* connect input data */
    statevec = NV_DATA_S(y);
    /* run the event function */
    model(time, statevec, NULL, paramdataPtr, DOFLAG_EVENTS, NULL, NULL, gout, NULL);
    return(0);
}

/*
 *========================================================================
 * Error function 
 *========================================================================
 */
static void errorMsg(char *text)
{
    /* First free the allocated memory */
    freeMem();
    /* Then print error message and exit */
    mexErrMsgTxt(text);
}

/*
 *========================================================================
 * Free the memory 
 *========================================================================
 */
static void freeMem()
{
    /* Free all CVODE related memory */
    N_VDestroy_Serial(u);  /* Free the u vector */
    CVodeFree(&cvode_mem);  /* Free the integrator memory */
}

    /*
     * ==============================================
     * HANDLE THE VARIABLE INPUT ARGUMENTS
     * ==============================================
     */
static mxArray* handleInputArguments(int nlhs, mxArray *plhs[], int nrhs, const mxArray *prhs[])
{
    mxArray *rhs[1], *lhs[1]; /* for calling matlab function "convertNNCa2NANNCaSBPD" */

    /* need to set to NULL some things for checking */
    parametervectorMX = NULL;
    initialconditionsMX = NULL;
    optionsMX = NULL;
    
    if (nrhs == 0) {
        /***************************************************/
        /* no input arguments => return initial conditions */
        /***************************************************/
        
        /* check if only numeric initial conditions */
        if (hasOnlyNumericICs) {
            /* numeric */
            resultMX = mxCreateDoubleMatrix(NRSTATES, 1, mxREAL);
            result = mxGetPr(resultMX);
            for (k=0; k<NRSTATES; k++) result[k] = defaultICs_num[k];
            return resultMX;
        } else {
            /* non-numeric */
            resultMX = mxCreateCellMatrix(NRSTATES, 1);
            result = mxGetPr(resultMX);
            for (k=0; k<NRSTATES; k++) mxSetCell(resultMX,k,mxCreateString(defaultICs_nonnum[k]));
            /* call matlab function with resultMX as input to convert numeric strings to numbers ... */
            rhs[0] = resultMX;
            mexCallMATLAB(1, lhs, 1, rhs, "convertNNCa2NANNCaSBPD");
            return lhs[0];
        }
    } else if (nrhs == 1) {
        /***************************************************/
        /* handle single input argument                    */
        /***************************************************/
        
        if (mxIsEmpty(prhs[0])) {
            mexErrMsgTxt("Timevector is empty.");
        } 
        if (mxIsChar(prhs[0])) {
            length2check = (mxGetM(prhs[0]) * mxGetN(prhs[0]));
            if (length2check == 6) {       
                resultMX = mxCreateCellMatrix(NRSTATES, 1);
                for (k=0; k<NRSTATES; k++) mxSetCell(resultMX,k,mxCreateString(stateNames[k]));
                return resultMX;
            }
            else if (length2check == 10) { 
                resultMX = mxCreateCellMatrix(NRPARAMETERS, 1);
                for (k=0; k<NRPARAMETERS; k++) mxSetCell(resultMX,k,mxCreateString(parameterNames[k]));
                return resultMX;
            }
            else if (length2check == 15) { 
                resultMX = mxCreateDoubleMatrix(NRPARAMETERS, 1, mxREAL);
                result = mxGetPr(resultMX);
                for (k=0; k<NRPARAMETERS; k++) result[k] = defaultParam[k];
                return resultMX;
            } else mexErrMsgTxt("Incorrect input argument.");
        } else {
            if (!mxIsDouble(prhs[0])) mexErrMsgTxt("Check the 'timevector' (first) input argument.");
            /* if no error then the first argument is the timevector for simulation */
            timesimvectorMX = (mxArray *) prhs[0];
        }
    } else {
        /***************************************************/
        /* handle other numbers of input arguments         */
        /***************************************************/
        
        if (!mxIsDouble(prhs[0])) mexErrMsgTxt("Check 'timevector' (first) input argument.");
        timesimvectorMX = (mxArray *) prhs[0];
        if (nrhs >= 2) {
            if (!mxIsDouble(prhs[1]) && !mxIsEmpty(prhs[1])) mexErrMsgTxt("Check 'initialconditions' (second) input argument.");
            initialconditionsMX = (mxArray *) prhs[1];
        }
        if (nrhs >= 3) {
            if (!mxIsDouble(prhs[2]) && !mxIsEmpty(prhs[2])) mexErrMsgTxt("Check 'parametervector' (third) input argument.");
            parametervectorMX = (mxArray *) prhs[2];
        } 
        if (nrhs >= 4) {
            if (!mxIsStruct(prhs[3]) && !mxIsEmpty(prhs[3])) mexErrMsgTxt("Check 'options' (fourth) input argument.");
            optionsMX = (mxArray *) prhs[3];
        }
        if (nrhs >= 5) {
            mexErrMsgTxt("Incorrect number of input arguments.");
        }
    }
    /***************************************************/
    /* process input arguments                         */
    /***************************************************/

    /* process timevector only if not empty */
    if (!mxIsEmpty(timesimvectorMX)) {
        timesimvector = mxGetPr(timesimvectorMX);
        /* Get some flags to check usability of time vector */
        numbertimesteps = mxGetM(timesimvectorMX)*mxGetN(timesimvectorMX);
        if (numbertimesteps == 1) timevectorIsScalar = 1; /* needed if xdotcalc */
        if (mxGetN(timesimvectorMX) > 1 && mxGetM(timesimvectorMX) > 1) timevectorNotVector = 1; /* vector needed for simulation */
    } else {
        /* set error flag for simulation case */
        timevectorEmpty = 1;
    }
    /* check parametervector */
    /* needs to be done before initial conditions in order to be able to handle non-numeric ICs */
    /* we need to get new memory for the parametervector in order to create an independent copy of
     * the default parameter vector. Otherwise events on parameters do change the default parameter vector 
     * which definitely is not desired!!! */
    parametervector = (double *) mxCalloc(NRPARAMETERS, sizeof(double));
    /* Copy default parameters into the parametervector */
    for (k=0; k<NRPARAMETERS; k++) parametervector[k] = defaultParam[k];
    
    if (parametervectorMX != NULL) {
        if (!mxIsEmpty(parametervectorMX)) {
            if (mxGetN(parametervectorMX)*mxGetM(parametervectorMX) != NRPARAMETERS || (mxGetN(parametervectorMX) > 1 && mxGetM(parametervectorMX) > 1)) mexErrMsgTxt("'parametervector' needs to be a vector of 'number parameters' length.");
            parametervector = mxGetPr(parametervectorMX);
        }
    }
    /* assign parametervector to modeldata struct */
    modeldata.parametervector = parametervector;
    /* process initial conditions */
    initialconditions = (double *) mxCalloc(NRSTATES,sizeof(double));
    /* For the initial conditions we need to take care of eventual non-numeric initial conditions */
    /* These can depend on parameters (both nominal and user-provided ones */
    /* If the user provides initial conditions, these overwrite all other ones and the non-numeric equations */
    /* will not be used anymore. */
    if (hasOnlyNumericICs == 1) {
        for (k=0;k<NRSTATES;k++) initialconditions[k] = defaultICs_num[k];
    } else {
        calc_ic_model(initialconditions, &modeldata);
    }
    if (initialconditionsMX != NULL) {
        if (!mxIsEmpty(initialconditionsMX)) {
            if (mxGetN(initialconditionsMX)*mxGetM(initialconditionsMX) != NRSTATES || (mxGetN(initialconditionsMX) > 1 && mxGetM(initialconditionsMX) > 1)) mexErrMsgTxt("'initialconditions' needs to be a vector of 'number states' length.");
            initialconditions = mxGetPr(mxDuplicateArray(initialconditionsMX));
        }
    }    
    /* check options */
    showIntegratorStats = 0;
    minstep = 0.0;  /* default value in CVODE */
    maxstep = 0.0;  /* default value in CVODE => infinity */
    maxnumsteps = MAXNUMSTEPS;          /* we set a value different from CVODE default values as default */
    maxerrtestfails = MAXERRTESTFAILS;  /* we set a value different from CVODE default values as default */
    maxorder = 5; /* default value for default method (stiff) */
    maxconvfails = 10;      /* default value in CVODE */
    initstep = 0.0;         /* default value in CVODE */
    maxnonlineariter = 3;   /* default value in CVODE */
    method = 0;        /* 0 = stiff, 1 = nonstiff, default: 0 */
    reltol = RELTOL;   /* we set a value different from CVODE default values as default */
    abstol = ABSTOL;   /* we set a value different from CVODE default values as default */
    xdotcalc = 0;
    if (optionsMX != NULL) {
        if (!mxIsEmpty(optionsMX)) {
            showIntegratorStatsMX = mxGetField(optionsMX, 0, "showIntegratorStats");
            if (showIntegratorStatsMX != NULL) 
                if (mxIsDouble(showIntegratorStatsMX)) 
                    showIntegratorStats = (int)mxGetScalar(showIntegratorStatsMX); 
                else 
                    mexErrMsgTxt("'options.showIntegratorStats' wrongly defined.");

            minstepMX = mxGetField(optionsMX, 0, "minstep");
            if (minstepMX != NULL) 
                if (mxIsDouble(minstepMX)) 
                    minstep = mxGetScalar(minstepMX); 
                else 
                    mexErrMsgTxt("'options.minstep' wrongly defined.");
            maxstepMX = mxGetField(optionsMX, 0, "maxstep");
            if (maxstepMX != NULL) 
                if (mxIsDouble(maxstepMX)) 
                    maxstep = mxGetScalar(maxstepMX); 
                else 
                    mexErrMsgTxt("'options.maxstep' wrongly defined.");
            maxnumstepsMX = mxGetField(optionsMX, 0, "maxnumsteps");
            if (maxnumstepsMX != NULL) 
                if (mxIsDouble(maxnumstepsMX)) 
                    maxnumsteps = (long int)mxGetScalar(maxnumstepsMX); 
                else 
                    mexErrMsgTxt("'options.maxnumsteps' wrongly defined.");
            methodMX = mxGetField(optionsMX, 0, "method");
            if (methodMX != NULL) {
                if (mxIsChar(methodMX)) {
                    if (mxGetM(methodMX) * mxGetN(methodMX) == 8) {
                        method = 1; /* "nonstiff" method ... just check the length of the string */
                    } else if (mxGetM(methodMX) * mxGetN(methodMX) == 5) {
                        method = 0; /* "stiff method */
                    } else {
                        mexErrMsgTxt("'options.method' wrongly defined.");
                    }
                } else {
                    mexErrMsgTxt("'options.method' wrongly defined.");
                }
            }

            /* Set maxorder, depending on the method chosen to the default values */
			if (method==0) maxorder = 5;  /* stiff BDF */
			else maxorder = 12;           /* nonstiff Adams */
			
			reltolMX = mxGetField(optionsMX, 0, "reltol");
            if (reltolMX != NULL) 
                if (mxIsDouble(reltolMX)) 
                    reltol = mxGetScalar(reltolMX); 
                else 
                    mexErrMsgTxt("'options.reltol' wrongly defined.");
            abstolMX = mxGetField(optionsMX, 0, "abstol");
            if (abstolMX != NULL) 
                if (mxIsDouble(abstolMX)) 
                    abstol = mxGetScalar(abstolMX); 
                else 
                    mexErrMsgTxt("'options.abstol' wrongly defined.");
            xdotcalcMX = mxGetField(optionsMX, 0, "xdotcalc");
            if (xdotcalcMX != NULL) 
                if (mxIsDouble(xdotcalcMX)) 
                    xdotcalc = mxGetScalar(xdotcalcMX); 
                else 
                    mexErrMsgTxt("'options.xdotcalc' wrongly defined.");

            
            /* Additional Options */
            maxerrtestfailsMX = mxGetField(optionsMX, 0, "maxerrtestfails");
            if (maxerrtestfailsMX != NULL) 
                if (mxIsDouble(maxerrtestfailsMX)) 
                    maxerrtestfails = (int)mxGetScalar(maxerrtestfailsMX); 
                else 
                    mexErrMsgTxt("'options.maxerrtestfails' wrongly defined.");            

            maxorderMX = mxGetField(optionsMX, 0, "maxorder");
            if (maxorderMX != NULL) 
                if (mxIsDouble(maxorderMX)) 
                    maxorder = (int)mxGetScalar(maxorderMX); 
                else 
                    mexErrMsgTxt("'options.maxorder' wrongly defined.");            

            maxconvfailsMX = mxGetField(optionsMX, 0, "maxconvfails");
            if (maxconvfailsMX != NULL) 
                if (mxIsDouble(maxconvfailsMX)) 
                    maxconvfails = (int)mxGetScalar(maxconvfailsMX); 
                else 
                    mexErrMsgTxt("'options.maxconvfails' wrongly defined.");    
            
            initstepMX = mxGetField(optionsMX, 0, "initstep");
            if (initstepMX != NULL) 
                if (mxIsDouble(initstepMX)) 
                    initstep = (double)mxGetScalar(initstepMX); 
                else 
                    mexErrMsgTxt("'options.initstep' wrongly defined.");                

            maxnonlineariterMX = mxGetField(optionsMX, 0, "maxnonlineariter");
            if (maxnonlineariterMX != NULL) 
                if (mxIsDouble(maxnonlineariterMX)) 
                    maxnonlineariter = (double)mxGetScalar(maxnonlineariterMX); 
                else 
                    mexErrMsgTxt("'options.maxnonlineariter' wrongly defined.");                
        }
    }
    return NULL;
}

    /*
     * ==============================================
     * GET RHS OF ODEs (fir time = 0)
     * ==============================================
     */
static mxArray* doXdotCalc()
{
    /* Check if timevector is a scalar */
    if (timevectorIsScalar != 1) mexErrMsgTxt("'timevector' input argument needs to be a scalar if you want to calculate the RHS.");
    resultMX = mxCreateDoubleMatrix(NRSTATES, 1, mxREAL);
    result = mxGetPr(resultMX);
    model(timesimvector[0], initialconditions, result, &modeldata, DOFLAG_DDT, NULL, NULL, NULL, NULL);
    return resultMX;
}

    /*
     * ==============================================
     * ALLOCATE MEMORY 
     * ==============================================
     */
static void allocSimMemory()
{
    statevaluesMX = mxCreateDoubleMatrix(numbertimesteps, NRSTATES, mxREAL);
    statevalues = mxGetPr(statevaluesMX);
    if (NRVARIABLES > 0) {
        variablevec = (double *) mxCalloc(NRVARIABLES, sizeof(double));
        variablevaluesMX = mxCreateDoubleMatrix(numbertimesteps, NRVARIABLES, mxREAL);
        variablevalues = mxGetPr(variablevaluesMX);
    }
    if (NRREACTIONS > 0) {
        reactionvec = (double *) mxCalloc(NRREACTIONS, sizeof(double));
        reactionvaluesMX = mxCreateDoubleMatrix(numbertimesteps, NRREACTIONS, mxREAL);
        reactionvalues = mxGetPr(reactionvaluesMX);
    }
    if (NREVENTS > 0) {
        eventvec = (int *) mxCalloc(NREVENTS, sizeof(int));
        eventdataold = (double *) mxCalloc(NREVENTS, sizeof(double));
    }
}
    
    /*
     * ==============================================
     * INITIALIZE INTEGRATOR
     * ==============================================
     */
static void initCVODE()
{
    u = N_VMake_Serial(NRSTATES,initialconditions);
    /* Set integration method (stiff or non-stiff) */
    if (method == 0) cvode_mem = CVodeCreate(CV_BDF, CV_NEWTON);    /* default (stiff) */
    else cvode_mem = CVodeCreate(CV_ADAMS, CV_FUNCTIONAL);          /* nonstiff */
    CVodeMalloc(cvode_mem, f, timesimvector[0], u, CV_SS, reltol, &abstol);
    if (NREVENTS > 0) {
        CVodeRootInit(cvode_mem, NREVENTS, g, &modeldata);
    }
    CVodeSetFdata(cvode_mem, &modeldata);
    if (minstep > 0) CVodeSetMinStep(cvode_mem, minstep);
    if (maxstep > 0) CVodeSetMaxStep(cvode_mem, maxstep);
    if (maxnumsteps > 0) CVodeSetMaxNumSteps(cvode_mem, maxnumsteps);
    CVodeSetMaxErrTestFails(cvode_mem, maxerrtestfails);
    CVodeSetMaxOrd(cvode_mem, maxorder);
    CVodeSetMaxConvFails(cvode_mem, maxconvfails);
    CVodeSetInitStep(cvode_mem, initstep);
    CVodeSetMaxNonlinIters(cvode_mem, maxnonlineariter);
    CVDense(cvode_mem,NRSTATES);
}

    /*
     * ==============================================
     * DO THE INTEGRATION
     * ==============================================
     */
static void integrate()
{
    addVec2Mat(statevalues,initialconditions,0,numbertimesteps,NRSTATES);
    model(timesimvector[0], initialconditions, NULL, &modeldata, DOFLAG_VARREAC, variablevec, reactionvec,NULL,NULL);
    if (NRVARIABLES > 0) addVec2Mat(variablevalues,variablevec,0,numbertimesteps,NRVARIABLES);
    if (NRREACTIONS > 0) addVec2Mat(reactionvalues,reactionvec,0,numbertimesteps,NRREACTIONS);
    if (NREVENTS > 0) /* determine eventdataold to be able to detect directions of events */
        model(timesimvector[0], initialconditions, NULL, &modeldata, DOFLAG_EVENTS, NULL, NULL, eventdataold, NULL);
    k = 1;
    tendstep = timesimvector[k]; 
    nreventshappend = 0; 
    while(1) {
        flag = CVode(cvode_mem, tendstep, u, &treturn, CV_NORMAL);
        if (flag < 0) {
            if (flag == CV_TOO_MUCH_WORK) errorMsg("CVODE Error: CV_TOO_MUCH_WORK");
            else if (flag == CV_TOO_MUCH_ACC) errorMsg("CVODE Error: CV_TOO_MUCH_ACC");
            else if (flag == CV_ERR_FAILURE || flag == CV_CONV_FAILURE) errorMsg("CVODE Error: CV_ERR_FAILURE");
            else {
                sprintf(stringbuffer, "CVODE Error Flag: %d",flag);
                errorMsg(stringbuffer);
            }
        }
        statevec = NV_DATA_S(u);
        if (flag == CV_ROOT_RETURN) {
            /* Event happened */
            CVodeGetRootInfo(cvode_mem, eventvec);
            eventcorrecttest = 0;
            model(treturn, statevec, &eventcorrecttest, &modeldata, DOFLAG_EVENTASSIGN, NULL, NULL, eventdataold, eventvec);
            flag = CVodeReInit(cvode_mem, f, treturn, u, CV_SS, reltol, &abstol);
            if (eventcorrecttest > 0) {
                nreventshappend += 1;  
                if (nreventshappend == 1) {
                    eventtimes = (double *) mxCalloc(1, sizeof(double));
                    eventflagsdata = (double *) mxCalloc(NREVENTS, sizeof(double));
                } else {
                    eventtimes = (double *) mxRealloc((void *) eventtimes, nreventshappend*sizeof(double));
                    eventflagsdata = (double *) mxRealloc((void *) eventflagsdata, NREVENTS*nreventshappend*sizeof(double));
                }
                eventtimes[nreventshappend-1] = treturn;
                for (k2=0;k2<NREVENTS;k2++) eventflagsdata[NREVENTS*(nreventshappend-1)+k2] = (double)eventvec[k2];
            }
        }
        if (tendstep == treturn) { 
            addVec2Mat(statevalues,statevec,k,numbertimesteps,NRSTATES);
            model(tendstep, statevec, NULL, &modeldata, DOFLAG_VARREAC, variablevec, reactionvec, NULL, NULL);
            if (NRVARIABLES > 0) addVec2Mat(variablevalues,variablevec,k,numbertimesteps,NRVARIABLES);
            if (NRREACTIONS > 0) addVec2Mat(reactionvalues,reactionvec,k,numbertimesteps,NRREACTIONS);
            k = k+1;
            if (k < numbertimesteps) 
                tendstep = timesimvector[k];
            else 
                break; 
        }
        if (NREVENTS > 0) /* determine eventdataold to be able to detect directions of events */
            model(treturn, statevec, NULL, &modeldata, DOFLAG_EVENTS, NULL, NULL, eventdataold, NULL);
    }
}

    /*
     * ==============================================
     * CONSTRUCT OUTPUT VARIABLE
     * ==============================================
     */
static mxArray* constructOutput()
{
    resultMX = mxCreateStructMatrix(1,1,0,NULL);
    /* Create fields in result structure */
    mxAddField(resultMX,"time");
    mxAddField(resultMX,"states");
    mxAddField(resultMX,"statevalues");
    mxAddField(resultMX,"variables");
    mxAddField(resultMX,"variablevalues");
    mxAddField(resultMX,"reactions");
    mxAddField(resultMX,"reactionvalues");
    if (NREVENTS > 0) {
        mxAddField(resultMX,"events");
        mxAddField(resultMX,"eventtimes");
        mxAddField(resultMX,"eventflags");
    }
    mxSetField(resultMX, 0, "time", mxDuplicateArray(timesimvectorMX));
    mxSetField(resultMX, 0, "statevalues", statevaluesMX);
    if (NRVARIABLES > 0) mxSetField(resultMX, 0, "variablevalues", variablevaluesMX);
    if (NRREACTIONS > 0) mxSetField(resultMX, 0, "reactionvalues", reactionvaluesMX);
    statesMX = mxCreateCellMatrix(1, NRSTATES);
    for (k=0;k<NRSTATES;k++) mxSetCell(statesMX,k,mxCreateString(stateNames[k]));
    mxSetField(resultMX, 0, "states", statesMX);
    if (NRVARIABLES > 0) {
        variablesMX = mxCreateCellMatrix(1, NRVARIABLES);
        for (k=0;k<NRVARIABLES;k++) mxSetCell(variablesMX,k,mxCreateString(variableNames[k]));
        mxSetField(resultMX, 0, "variables", variablesMX);
    }
    if (NRREACTIONS > 0) {
        reactionsMX = mxCreateCellMatrix(1, NRREACTIONS);
        for (k=0;k<NRREACTIONS;k++) mxSetCell(reactionsMX,k,mxCreateString(reactionNames[k]));
        mxSetField(resultMX, 0, "reactions", reactionsMX);
    }
    if (NREVENTS > 0) {
        eventsMX = mxCreateCellMatrix(1, NREVENTS);
        for (k=0;k<NREVENTS;k++) mxSetCell(eventsMX,k,mxCreateString(eventNames[k]));
        mxSetField(resultMX, 0, "events", eventsMX);
        if (nreventshappend > 0) {
            eventtimesMX = mxCreateDoubleMatrix(1,nreventshappend,mxREAL);
            mxSetData(eventtimesMX, eventtimes);
            mxSetField(resultMX, 0, "eventtimes", eventtimesMX);
            eventflagsMX = mxCreateDoubleMatrix(NREVENTS,nreventshappend,mxREAL);
            mxSetData(eventflagsMX, eventflagsdata);
            mxSetField(resultMX, 0, "eventflags", eventflagsMX);
        }
    }
    return resultMX;
}

/*
 *========================================================================
 * Initialization of variables
 *========================================================================
 */
static void doInitilizationVariables()
 {
    timesimvectorMX = NULL;
    timesimvector = NULL;
    parametervectorMX = NULL;
    parametervector = NULL;
    parametervectorCVODES = NULL;
    initialconditionsMX = NULL;
    initialconditions = NULL;
    optionsMX = NULL;
    showIntegratorStatsMX = NULL;
    showIntegratorStats = 0;
    minstepMX = NULL;
    maxstepMX = NULL;
    maxnumstepsMX = NULL;
    reltolMX = NULL;
    abstolMX = NULL;
    maxerrtestfailsMX = NULL;
    maxorderMX = NULL;
    maxconvfailsMX = NULL;
    initstepMX = NULL;
    maxnonlineariterMX = NULL;
    xdotcalcMX = NULL;
    resultMX = NULL;
    result = NULL;
    statesMX = NULL;
    statevaluesMX = NULL;
    statevalues = NULL;
    variablesMX = NULL;
    variablevaluesMX = NULL;
    variablevalues = NULL;
    reactionsMX = NULL;
    reactionvaluesMX = NULL;
    reactionvalues = NULL;
    eventsMX = NULL;
    eventtimesMX = NULL;
    eventtimes = NULL;
    eventflagsMX = NULL;
    eventflags = NULL;
    u = NULL;
    statevec = NULL;
    variablevec = NULL;
    reactionvec = NULL;
    eventvec = NULL;
    eventdataold = NULL;
    eventflagsdata = NULL;
    timevectorEmpty = 0;
    timevectorNotVector = 0;
    timevectorIsScalar = 0;
    methodMX = NULL;
}
```
