## Supplementary scripts for "A multi-scale *in silico* mouse model for insulin resistance and humanoid type 2 diabetes": cvodes.h

```
/*
 * -----------------------------------------------------------------
 * $Revision: 749 $
 * $Date: 2011-06-22 14:49:10 +0200 (Wed, 22 Jun 2011) $
 * ----------------------------------------------------------------- 
 * Programmer(s): Radu Serban @ LLNL
 * -----------------------------------------------------------------
 * Copyright (c) 2005, The Regents of the University of California.
 * Produced at the Lawrence Livermore National Laboratory.
 * All rights reserved.
 * For details, see the LICENSE file.
 * -----------------------------------------------------------------
 * This is the interface file for the main CVODES integrator.
 * -----------------------------------------------------------------
 *
 * CVODES is used to solve numerically the ordinary initial value    
 * problem:                                                          
 *                                                                   
 *                 y' = f(t,y),                                      
 *                 y(t0) = y0,                                       
 *                                                                   
 * where t0, y0 in R^N, and f: R x R^N -> R^N are given.             
 *                                                                   
 * Optionally, CVODES can perform forward or adjoint sensitivity 
 * analysis to find sensitivities of the solution y with respect 
 * to parameters in the right hand side f and/or in the initial         
 * conditions y0.                                                    
 *
 * -----------------------------------------------------------------
 */

#ifndef _CVODES_H
#define _CVODES_H

#ifdef __cplusplus  /* wrapper to enable C++ usage */
extern "C" {
#endif

#include <stdio.h>

#include <sundials/sundials_nvector.h>

  /*
   * =================================================================
   *              C V O D E S     C O N S T A N T S
   * =================================================================
   */

  /*
   * -----------------------------------------------------------------
   * Enumerations for inputs to CVodeCreate, CVodeMalloc,
   * CVodeReInit, CVodeSensMalloc, CVodeSensReInit, CVodeQuadMalloc,
   * CVodeQuadReInit, CVodeSet*, CVode, and CVadjMalloc.
   * -----------------------------------------------------------------
   * Symbolic constants for the lmm, iter, and itol input parameters 
   * to CVodeMalloc and CVodeReInit, the input parameter itask to CVode, 
   * and the input parameter interp to CVadjMalloc, are given below.
   *
   * lmm:   The user of the CVODES package specifies whether to use
   *        the CV_ADAMS or CV_BDF (backward differentiation formula)
   *        linear multistep method. The BDF method is recommended
   *        for stiff problems, and the CV_ADAMS method is recommended
   *        for nonstiff problems.
   *
   * iter:  At each internal time step, a nonlinear equation must
   *        be solved. The user can specify either CV_FUNCTIONAL
   *        iteration, which does not require linear algebra, or a
   *        CV_NEWTON iteration, which requires the solution of linear
   *        systems. In the CV_NEWTON case, the user also specifies a
   *        CVODE linear solver. CV_NEWTON is recommended in case of
   *        stiff problems.
   *
   * itol:  This parameter specifies the relative and absolute
   *        tolerance types to be used. The CV_SS tolerance type means
   *        a scalar relative and absolute tolerance. The CV_SV
   *        tolerance type means a scalar relative tolerance and a
   *        vector absolute tolerance (a potentially different
   *        absolute tolerance for each vector component). The CV_WF
   *        tolerance type means that the user provides a function
   *        (of type CVEwtFn) to set the error weight vector.
   *
   * ism:   This parameter specifies the sensitivity corrector type
   *        to be used. In the CV_SIMULTANEOUS case, the nonlinear
   *        systems for states and all sensitivities are solved
   *        simultaneously. In the CV_STAGGERED case, the nonlinear
   *        system for states is solved first and then, the
   *        nonlinear systems for all sensitivities are solved
   *        at the same time. Finally, in the CV_STAGGERED1 approach
   *        all nonlinear systems are solved in a sequence.
   *
   * itask: The itask input parameter to CVode indicates the job
   *        of the solver for the next user step. The CV_NORMAL
   *        itask is to have the solver take internal steps until
   *        it has reached or just passed the user specified tout
   *        parameter. The solver then interpolates in order to
   *        return an approximate value of y(tout). The CV_ONE_STEP
   *        option tells the solver to just take one internal step
   *        and return the solution at the point reached by that
   *        step. The CV_NORMAL_TSTOP and CV_ONE_STEP_TSTOP modes are
   *        similar to CV_NORMAL and CV_ONE_STEP, respectively, except
   *        that the integration never proceeds past the value
   *        tstop (specified through the routine CVodeSetStopTime).
   *
   * interp: Specifies the interpolation type used to evaluate the
   *        forward solution during the backward integration phase.
   *        CV_HERMITE specifies cubic Hermite interpolation.
   *        CV_POYNOMIAL specifies the polynomial interpolation
   * -----------------------------------------------------------------
   */

  /* lmm */
#define CV_ADAMS          1
#define CV_BDF            2

  /* iter */
#define CV_FUNCTIONAL     1
#define CV_NEWTON         2

  /* itol */
#define CV_SS             1
#define CV_SV             2
#define CV_WF             3
#define CV_EE             4

  /* itask */
#define CV_NORMAL         1
#define CV_ONE_STEP       2
#define CV_NORMAL_TSTOP   3
#define CV_ONE_STEP_TSTOP 4

  /* ism */
#define CV_SIMULTANEOUS   1
#define CV_STAGGERED      2
#define CV_STAGGERED1     3

  /* DQtype */
#define CV_CENTERED       1
#define CV_FORWARD        2

  /* interp */
#define CV_HERMITE        1
#define CV_POLYNOMIAL     2

  /* 
   * ----------------------------------------
   * CVODES return flags
   * ----------------------------------------
   */

#define CV_SUCCESS               0
#define CV_TSTOP_RETURN          1
#define CV_ROOT_RETURN           2

#define CV_WARNING              99

#define CV_TOO_MUCH_WORK        -1
#define CV_TOO_MUCH_ACC         -2
#define CV_ERR_FAILURE          -3
#define CV_CONV_FAILURE         -4

#define CV_LINIT_FAIL           -5
#define CV_LSETUP_FAIL          -6
#define CV_LSOLVE_FAIL          -7
#define CV_RHSFUNC_FAIL         -8
#define CV_FIRST_RHSFUNC_ERR    -9
#define CV_REPTD_RHSFUNC_ERR    -10
#define CV_UNREC_RHSFUNC_ERR    -11
#define CV_RTFUNC_FAIL          -12

#define CV_MEM_FAIL             -20
#define CV_MEM_NULL             -21
#define CV_ILL_INPUT            -22
#define CV_NO_MALLOC            -23
#define CV_BAD_K                -24
#define CV_BAD_T                -25
#define CV_BAD_DKY              -26
#define CV_TOO_CLOSE            -27

#define CV_NO_QUAD              -30
#define CV_QRHSFUNC_FAIL        -31
#define CV_FIRST_QRHSFUNC_ERR   -32
#define CV_REPTD_QRHSFUNC_ERR   -33
#define CV_UNREC_QRHSFUNC_ERR   -34

#define CV_BAD_IS               -40
#define CV_NO_SENS              -41
#define CV_SRHSFUNC_FAIL        -42
#define CV_FIRST_SRHSFUNC_ERR   -43
#define CV_REPTD_SRHSFUNC_ERR   -44
#define CV_UNREC_SRHSFUNC_ERR   -45

  /* 
   * ----------------------------------------
   * CVODEA return flags
   * ----------------------------------------
   */

#define CV_ADJMEM_NULL         -101
#define CV_BAD_TB0             -103
#define CV_BCKMEM_NULL         -104
#define CV_REIFWD_FAIL         -105
#define CV_FWD_FAIL            -106
#define CV_BAD_ITASK           -107
#define CV_BAD_TBOUT           -108
#define CV_GETY_BADT           -109

  /*
   * =================================================================
   *              F U N C T I O N   T Y P E S
   * =================================================================
   */

  /*
   * -----------------------------------------------------------------
   * Type : CVRhsFn
   * -----------------------------------------------------------------
   * The f function which defines the right hand side of the ODE
   * system y' = f(t,y) must have type CVRhsFn.
   * f takes as input the independent variable value t, and the
   * dependent variable vector y.  It stores the result of f(t,y)
   * in the vector ydot.  The y and ydot arguments are of type
   * N_Vector.
   * (Allocation of memory for ydot is handled within CVODES)
   * The f_data parameter is the same as the f_data
   * parameter set by the user through the CVodeSetFdata routine.
   * This user-supplied pointer is passed to the user's f function
   * every time it is called.
   *
   * A CVRhsFn should return 0 if successful, a negative value if
   * an unrecoverable error occured, and a positive value if a 
   * recoverable error (e.g. invalid y values) occured. 
   * If an unrecoverable occured, the integration is halted. 
   * If a recoverable error occured, then (in most cases) CVODES
   * will try to correct and retry.
   * -----------------------------------------------------------------
   */

  typedef int (*CVRhsFn)(realtype t, N_Vector y,
                         N_Vector ydot, void *f_data);

  /*
   * -----------------------------------------------------------------
   * Type : CVRootFn
   * -----------------------------------------------------------------
   * A function g, which defines a set of functions g_i(t,y) whose
   * roots are sought during the integration, must have type CVRootFn.
   * The function g takes as input the independent variable value
   * t, and the dependent variable vector y.  It stores the nrtfn
   * values g_i(t,y) in the realtype array gout.
   * (Allocation of memory for gout is handled within CVODE.)
   * The g_data parameter is the same as that passed by the user
   * to the CVodeRootInit routine.  This user-supplied pointer is
   * passed to the user's g function every time it is called.
   *
   * A CVRootFn should return 0 if successful or a non-zero value
   * if an error occured (in which case the integration will be halted).
   * -----------------------------------------------------------------
   */

  typedef int (*CVRootFn)(realtype t, N_Vector y, realtype *gout,
                          void *g_data);

  /*
   * -----------------------------------------------------------------
   * Type : CVEwtFn
   * -----------------------------------------------------------------
   * A function e, which sets the error weight vector ewt, must have
   * type CVEwtFn.
   * The function e takes as input the current dependent variable y.
   * It must set the vector of error weights used in the WRMS norm:
   * 
   *   ||y||_WRMS = sqrt [ 1/N * sum ( ewt_i * y_i)^2 ]
   *
   * Typically, the vector ewt has components:
   * 
   *   ewt_i = 1 / (reltol * |y_i| + abstol_i)
   *
   * The e_data parameter is the same as that passed by the user
   * to the CVodeSetEdata routine.  This user-supplied pointer is
   * passed to the user's e function every time it is called.
   * A CVEwtFn e must return 0 if the error weight vector has been
   * successfuly set and a non-zero value otherwise.
   * -----------------------------------------------------------------
   */

  typedef int (*CVEwtFn)(N_Vector y, N_Vector ewt, void *e_data);


  /*
   * -----------------------------------------------------------------
   * Type : CVErrHandlerFn
   * -----------------------------------------------------------------
   * A function eh, which handles error messages, must have type
   * CVErrHandlerFn.
   * The function eh takes as input the error code, the name of the
   * module reporting the error, the error message, and a pointer to
   * user data, the same as that passed to CVodeSetErrHandlerFn.
   * 
   * All error codes are negative, except CV_WARNING which indicates 
   * a warning (the solver continues).
   *
   * A CVErrHandlerFn has no return value.
   * -----------------------------------------------------------------
   */
  
  typedef void (*CVErrHandlerFn)(int error_code, 
                                 const char *module, const char *function, 
                                 char *msg, void *eh_data); 

  /*
   * -----------------------------------------------------------------
   * Type : CVQuadRhsFn
   * -----------------------------------------------------------------
   * The fQ function which defines the right hand side of the
   * quadrature equations yQ' = fQ(t,y) must have type CVQuadRhsFn.
   * fQ takes as input the value of the independent variable t,
   * the vector of states y and must store the result of fQ in
   * yQdot. (Allocation of memory for yQdot is handled by CVODES).
   * The fQ_data parameter is the same as the fQ_data parameter
   * set by the user through the CVodeSetQuadFdata routine and is
   * passed to the fQ function every time it is called.
   *
   * A CVQuadRhsFn should return 0 if successful, a negative value if
   * an unrecoverable error occured, and a positive value if a 
   * recoverable error (e.g. invalid y values) occured. 
   * If an unrecoverable occured, the integration is halted. 
   * If a recoverable error occured, then (in most cases) CVODES
   * will try to correct and retry.
   * -----------------------------------------------------------------
   */

  typedef int (*CVQuadRhsFn)(realtype t, N_Vector y, N_Vector yQdot,
                             void *fQ_data);

  /*
   * -----------------------------------------------------------------
   * Type : CVSensRhsFn
   * -----------------------------------------------------------------
   * The fS function which defines the right hand side of the
   * sensitivity ODE systems s' = f_y * s + f_p must have type
   * CVSensRhsFn.
   * fS takes as input the number of sensitivities Ns, the
   * independent variable value t, the states y and the
   * corresponding value of f(t,y) in ydot, and the dependent
   * sensitivity vectors yS. It stores the result of fS in ySdot.
   * (Allocation of memory for ySdot is handled within CVODES)
   * The fS_data parameter is the same as the fS_data parameter
   * set by the user through the CVodeSetSensFdata routine and is
   * passed to the fS function every time it is called.
   *
   * A CVSensRhsFn should return 0 if successful, a negative value if
   * an unrecoverable error occured, and a positive value if a 
   * recoverable error (e.g. invalid y or yS values) occured. 
   * If an unrecoverable occured, the integration is halted. 
   * If a recoverable error occured, then (in most cases) CVODES
   * will try to correct and retry.
   * -----------------------------------------------------------------
   */

  typedef int (*CVSensRhsFn)(int Ns, realtype t,
                             N_Vector y, N_Vector ydot,
                             N_Vector *yS, N_Vector *ySdot,
                             void *fS_data,
                             N_Vector tmp1, N_Vector tmp2);

  /*
   * -----------------------------------------------------------------
   * Type : CVSensRhs1Fn
   * -----------------------------------------------------------------
   * The fS1 function which defines the right hand side of the i-th
   * sensitivity ODE system s_i' = f_y * s_i + f_p must have type
   * CVSensRhs1Fn.
   * fS1 takes as input the number of sensitivities Ns, the current
   * sensitivity iS, the independent variable value t, the states y
   * and the corresponding value of f(t,y) in ydot, and the
   * dependent sensitivity vector yS. It stores the result of fS in
   * ySdot.
   * (Allocation of memory for ySdot is handled within CVODES)
   * The fS_data parameter is the same as the fS_data parameter
   * set by the user through the CVodeSetSensFdata routine and is
   * passed to the fS1 function every time it is called.
   *
   * A CVSensRhs1Fn should return 0 if successful, a negative value if
   * an unrecoverable error occured, and a positive value if a 
   * recoverable error (e.g. invalid y or yS values) occured. 
   * If an unrecoverable occured, the integration is halted. 
   * If a recoverable error occured, then (in most cases) CVODES
   * will try to correct and retry.
   * -----------------------------------------------------------------
   */

  typedef int (*CVSensRhs1Fn)(int Ns, realtype t,
                              N_Vector y, N_Vector ydot,
                              int iS, N_Vector yS, N_Vector ySdot,
                              void *fS_data,
                              N_Vector tmp1, N_Vector tmp2);

  /*
   * -----------------------------------------------------------------
   * CVRhsFnB
   *    The fB function which defines the right hand side of the
   *    ODE systems to be integrated backwards must have type CVRhsFnB.
   * -----------------------------------------------------------------
   * CVQuadRhsFnB
   *    The fQB function which defines the quadratures to be integrated
   *    backwards must have type CVQuadRhsFnB.
   * -----------------------------------------------------------------
   */
  
  typedef int (*CVRhsFnB)(realtype t, N_Vector y,
                          N_Vector yB, N_Vector yBdot,
                          void *f_dataB);
  
  typedef int (*CVQuadRhsFnB)(realtype t, N_Vector y,
                              N_Vector yB, N_Vector qBdot,
                              void *fQ_dataB);

  /*
   * =================================================================
   *          U S E R - C A L L A B L E   R O U T I N E S
   * =================================================================
   */

  /*
   * -----------------------------------------------------------------
   * Function : CVodeCreate
   * -----------------------------------------------------------------
   * CVodeCreate creates an internal memory block for a problem to
   * be solved by CVODES.
   *
   * lmm  is the type of linear multistep method to be used.
   *      The legal values are CV_ADAMS and CV_BDF (see previous
   *      description).
   *
   * iter  is the type of iteration used to solve the nonlinear
   *       system that arises during each internal time step.
   *       The legal values are CV_FUNCTIONAL and CV_NEWTON.
   *
   * If successful, CVodeCreate returns a pointer to initialized
   * problem memory. This pointer should be passed to CVodeMalloc.
   * If an initialization error occurs, CVodeCreate prints an error
   * message to standard err and returns NULL.
   * -----------------------------------------------------------------
   */

  void *CVodeCreate(int lmm, int iter);

  /*
   * -----------------------------------------------------------------
   * Integrator optional input specification functions
   * -----------------------------------------------------------------
   * The following functions can be called to set optional inputs
   * to values other than the defaults given below:
   *
   * Function                |  Optional input / [ default value ]
   * -----------------------------------------------------------------
   *                         |
   * CVodeSetErrHandlerFn    | user-provided ErrHandler function.
   *                         | [internal]
   *                         |
   * CVodeSetErrFile         | the file pointer for an error file
   *                         | where all CVODE warning and error
   *                         | messages will be written if the default
   *                         | internal error handling function is used. 
   *                         | This parameter can be stdout (standard 
   *                         | output), stderr (standard error), or a 
   *                         | file pointer (corresponding to a user 
   *                         | error file opened for writing) returned 
   *                         | by fopen.
   *                         | If not called, then all messages will
   *                         | be written to the standard error stream.
   *                         | [stderr]
   *                         |
   * CVodeSetFdata           | a pointer to user data that will be
   *                         | passed to the user's f function every
   *                         | time f is called.
   *                         | [NULL]
   *                         |
   * CVodeSetEwtFn           | user-provide EwtSet function e and 
   *                         | a pointer to user data that will be
   *                         | passed to the user's e function every
   *                         | time e is called.
   *                         | [NULL]
   *                         | [NULL]
   *                         |
   * CVodeSetMaxOrd          | maximum lmm order to be used by the
   *                         | solver.
   *                         | [12 for Adams , 5 for BDF]
   *                         |
   * CVodeSetMaxNumSteps     | maximum number of internal steps to be
   *                         | taken by the solver in its attempt to
   *                         | reach tout.
   *                         | [500]
   *                         |
   * CVodeSetMaxHnilWarns    | maximum number of warning messages
   *                         | issued by the solver that t+h==t on the
   *                         | next internal step. A value of -1 means
   *                         | no such messages are issued.
   *                         | [10]
   *                         |
   * CVodeSetStabLimDet      | flag to turn on/off stability limit
   *                         | detection (TRUE = on, FALSE = off).
   *                         | When BDF is used and order is 3 or
   *                         | greater, CVsldet is called to detect
   *                         | stability limit.  If limit is detected,
   *                         | the order is reduced.
   *                         | [FALSE]
   *                         |
   * CVodeSetInitStep        | initial step size.
   *                         | [estimated by CVODES]
   *                         |
   * CVodeSetMinStep         | minimum absolute value of step size
   *                         | allowed.
   *                         | [0.0]
   *                         |
   * CVodeSetMaxStep         | maximum absolute value of step size
   *                         | allowed.
   *                         | [infinity]
   *                         |
   * CVodeSetStopTime        | the independent variable value past
   *                         | which the solution is not to proceed.
   *                         | [infinity]
   *                         |
   * CVodeSetMaxErrTestFails | Maximum number of error test failures
   *                         | in attempting one step.
   *                         | [7]
   *                         |
   * CVodeSetMaxNonlinIters  | Maximum number of nonlinear solver
   *                         | iterations at one solution.
   *                         | [3]
   *                         |
   * CVodeSetMaxConvFails    | Maximum number of allowable conv.
   *                         | failures in attempting one step.
   *                         | [10]
   *                         |
   * CVodeSetNonlinConvCoef  | Coeficient in the nonlinear conv.
   *                         | test.
   *                         | [0.1]
   *                         |
   * -----------------------------------------------------------------
   *                         |
   * CVodeSetIterType        | Changes the current nonlinear iteration
   *                         | type.
   *                         | [set by CVodecreate]
   *                         |
   * CVodeSetTolerances      | Changes the integration tolerances
   *                         | between calls to CVode().
   *                         | [set by CVodeMalloc/CVodeReInit]
   *                         |
   * -----------------------------------------------------------------
   * Return flag:
   *   CV_SUCCESS   if successful
   *   CV_MEM_NULL  if the cvode memory is NULL
   *   CV_ILL_INPUT if an argument has an illegal value
   * -----------------------------------------------------------------
   */

  int CVodeSetErrHandlerFn(void *cvode_mem, CVErrHandlerFn ehfun, void *eh_data);
  int CVodeSetErrFile(void *cvode_mem, FILE *errfp);
  int CVodeSetFdata(void *cvode_mem, void *f_data);
  int CVodeSetEwtFn(void *cvode_mem, CVEwtFn efun, void *e_data);
  int CVodeSetMaxOrd(void *cvode_mem, int maxord);
  int CVodeSetMaxNumSteps(void *cvode_mem, long int mxsteps);
  int CVodeSetMaxHnilWarns(void *cvode_mem, int mxhnil);
  int CVodeSetStabLimDet(void *cvode_mem, booleantype stldet);
  int CVodeSetInitStep(void *cvode_mem, realtype hin);
  int CVodeSetMinStep(void *cvode_mem, realtype hmin);
  int CVodeSetMaxStep(void *cvode_mem, realtype hmax);
  int CVodeSetStopTime(void *cvode_mem, realtype tstop);
  int CVodeSetMaxErrTestFails(void *cvode_mem, int maxnef);
  int CVodeSetMaxNonlinIters(void *cvode_mem, int maxcor);
  int CVodeSetMaxConvFails(void *cvode_mem, int maxncf);
  int CVodeSetNonlinConvCoef(void *cvode_mem, realtype nlscoef);

  int CVodeSetIterType(void *cvode_mem, int iter);
  int CVodeSetTolerances(void *cvode_mem,
                         int itol, realtype reltol, void *abstol);

  /*
   * -----------------------------------------------------------------
   * Function : CVodeMalloc
   * -----------------------------------------------------------------
   * CVodeMalloc allocates and initializes memory for a problem to
   * to be solved by CVODES.
   *
   * cvode_mem is pointer to CVODES memory returned by CVodeCreate.
   *
   * f       is the right hand side function in y' = f(t,y).
   *
   * t0      is the initial value of t.
   *
   * y0      is the initial condition vector y(t0).
   *
   * itol    is the type of tolerances to be used.
   *         The legal values are:
   *            CV_SS (scalar relative and absolute  tolerances),
   *            CV_SV (scalar relative tolerance and vector
   *                  absolute tolerance).
   *            CV_WF (indicates that the user will provide a
   *                function to evaluate the error weights.
   *                In this case, reltol and abstol are ignored.)
   *
   * reltol  is the relative tolerance scalar.
   *
   * abstol  is a pointer to the absolute tolerance scalar or
   *         an N_Vector of absolute tolerances.
   *
   * The parameters itol, reltol, and abstol define a vector of
   * error weights, ewt, with components
   *   ewt[i] = 1/(reltol*abs(y[i]) + abstol)   (if itol = CV_SS), or
   *   ewt[i] = 1/(reltol*abs(y[i]) + abstol[i])   (if itol = CV_SV).
   * This vector is used in all error and convergence tests, which
   * use a weighted RMS norm on all error-like vectors v:
   *    WRMSnorm(v) = sqrt( (1/N) sum(i=1..N) (v[i]*ewt[i])^2 ),
   * where N is the problem dimension.
   *
   * If successful, CVodeMalloc returns SUCCESS. If an argument has
   * an illegal value, CVodeMalloc prints an error message to the
   * file specified by errfp and returns one of the error flags
   * defined below.
   * -----------------------------------------------------------------
   */

  int CVodeMalloc(void *cvode_mem, CVRhsFn f,
                  realtype t0, N_Vector y0,
                  int itol, realtype reltol, void *abstol);

  /*
   * -----------------------------------------------------------------
   * Function : CVodeReInit
   * -----------------------------------------------------------------
   * CVodeReInit re-initializes CVode for the solution of a problem,
   * where a prior call to CVodeMalloc has been made with the same
   * problem size N. CVodeReInit performs the same input checking
   * and initializations that CVodeMalloc does.
   * But it does no memory allocation, assuming that the existing
   * internal memory is sufficient for the new problem.
   *
   * The use of CVodeReInit requires that the maximum method order,
   * maxord, is no larger for the new problem than for the problem
   * specified in the last call to CVodeMalloc.  This condition is
   * automatically fulfilled if the multistep method parameter lmm
   * is unchanged (or changed from CV_ADAMS to CV_BDF) and the default
   * value for maxord is specified.
   *
   * The first argument to CVodeReInit is:
   *
   * cvode_mem = pointer to CVODES memory returned by CVodeCreate.
   *
   * All the remaining arguments to CVodeReInit have names and
   * meanings identical to those of CVodeMalloc.
   *
   * The return value of CVodeReInit is equal to CV_SUCCESS = 0 if
   * there were no errors; otherwise it is a negative int equal to:
   *   CV_MEM_NULL  indicating cvode_mem was NULL (i.e.,
   *                CVodeCreate has not been called).
   *   CV_NO_MALLOC indicating that cvode_mem has not been
   *                allocated (i.e., CVodeMalloc has not been
   *                called).
   *   CV_ILL_INPUT indicating an input argument was illegal
   *                (including an attempt to increase maxord).
   * In case of an error return, an error message is also printed.
   * -----------------------------------------------------------------
   */

  int CVodeReInit(void *cvode_mem, CVRhsFn f,
                  realtype t0, N_Vector y0,
                  int itol, realtype reltol, void *abstol);

  /*
   * -----------------------------------------------------------------
   * Function : CVodeRootInit
   * -----------------------------------------------------------------
   * CVodeRootInit initializes a rootfinding problem to be solved
   * during the integration of the ODE system.  It must be called
   * after CVodeCreate, and before CVode.  The arguments are:
   *
   * cvode_mem = pointer to CVODE memory returned by CVodeCreate.
   *
   * nrtfn     = number of functions g_i, an int >= 0.
   *
   * g         = name of user-supplied function, of type CVRootFn,
   *             defining the functions g_i whose roots are sought.
   *
   * g_data    = a pointer to user data that will be passed to the 
   *             user's g function every time g is called.
   *
   * If a new problem is to be solved with a call to CVodeReInit,
   * where the new problem has no root functions but the prior one
   * did, then call CVodeRootInit with nrtfn = 0.
   *
   * The return value of CVodeRootInit is CV_SUCCESS = 0 if there were
   * no errors; otherwise it is a negative int equal to:
   *   CV_MEM_NULL    indicating cvode_mem was NULL, or
   *   CV_MEM_FAIL    indicating a memory allocation failed.
   *                  (including an attempt to increase maxord).
   *   CV_ILL_INPUT   indicating nrtfn > 0 but g = NULL.
   * In case of an error return, an error message is also printed.
   * -----------------------------------------------------------------
   */

  int CVodeRootInit(void *cvode_mem, int nrtfn, CVRootFn g, void *g_data);

  /*
   * -----------------------------------------------------------------
   * Quadrature optional input specification functions
   * -----------------------------------------------------------------
   * The following functions can be called to set optional inputs
   * to values other than the defaults given below:
   *
   * Function             |  Optional input / [ default value ]
   * --------------------------------------------------------------
   *                      |
   * CVodeSetQuadFdata    | a pointer to user data that will be
   *                      | passed to the user's fQ function every
   *                      | time fQ is called.
   *                      | [NULL]
   *                      |
   * CVodeSetQuadErrCon   | are quadrature variables considered in
   *                      | the error control?
   *                      | If yes, set tolerances for quadrature
   *                      | integration. 
   *                      | [errconQ = FALSE]
   *                      | [ not tolerances]
   *                      |
   * -----------------------------------------------------------------
   * If successful, these functions return CV_SUCCESS. If an argument
   * has an illegal value, they return one of the error flags
   * defined for the CVodeSet* routines.
   * -----------------------------------------------------------------
   */

  int CVodeSetQuadFdata(void *cvode_mem, void *fQ_data);
  int CVodeSetQuadErrCon(void *cvode_mem, booleantype errconQ, 
                         int itolQ, realtype reltolQ, void *abstolQ);

  /*
   * -----------------------------------------------------------------
   * Function : CVodeQuadMalloc
   * -----------------------------------------------------------------
   * CVodeQuadMalloc allocates and initializes memory related to
   * quadrature integration.
   *
   * cvode_mem is a pointer to CVODES memory returned by CVodeCreate
   *
   * fQ    is the user-provided integrand routine.
   *
   * yQ0   is an N_Vector with initial values for quadratures
   *       (typically yQ0 has all zero components).
   *
   * Return values:
   *  CV_SUCCESS if successful
   *  CV_MEM_NULL if the cvode memory was NULL
   *  CV_MEM_FAIL if a memory allocation failed
   * -----------------------------------------------------------------
   */

  int CVodeQuadMalloc(void *cvode_mem, CVQuadRhsFn fQ, N_Vector yQ0);

  /*
   * -----------------------------------------------------------------
   * Function : CVodeQuadReInit
   * -----------------------------------------------------------------
   * CVodeQuadReInit re-initializes CVODES's quadrature related
   * memory for a problem, assuming it has already been allocated
   * in prior calls to CVodeMalloc and CVodeQuadMalloc.
   *
   * All problem specification inputs are checked for errors.
   * The number of quadratures Nq is assumed to be unchanged
   * since the previous call to CVodeQuadMalloc.
   *
   * Return values:
   *  CV_SUCCESS  if successful
   *  CV_MEM_NULL if the cvode memory was NULL
   *  CV_NO_QUAD  if quadratures were not initialized
   * -----------------------------------------------------------------
   */

  int CVodeQuadReInit(void *cvode_mem, CVQuadRhsFn fQ, N_Vector yQ0);

  /*
   * -----------------------------------------------------------------
   * Forward sensitivity optional input specification functions
   * -----------------------------------------------------------------
   * The following functions can be called to set optional inputs
   * to other values than the defaults given below:
   *
   * Function                   |  Optional input / [ default value ]
   * -----------------------------------------------------------------
   *                            |
   * CVodeSetSensRhsFn          | sensitivity right hand side function
   *                            | and user data pointer.
   *                            | This function must compute right hand
   *                            | sides for all sensitivity equations.
   *                            | [CVODES difference quotient approx.]
   *                            | [internal]
   *                            |
   * CVodeSetSensRhs1Fn         | the sensitivity right hand side
   *                            | and user data pointer.
   *                            | This function must compute right hand
   *                            | sides for one sensitivity equation at a
   *                            | time.
   *                            | [CVODES difference quotient approx.]
   *                            | [internal]
   *                            |
   * CVodeSetSensDQMethod       | controls the selection of finite
   *                            | difference schemes used in evaluating
   *                            | the sensitivity right hand sides:
   *                            | (centered vs. forward and 
   *                            | simultaneous vs. separate)
   *                            | [DQtype=CV_CENTERED]
   *                            | [DQrhomax=0.0]
   *                            |
   * CVodeSetSensParams         | parameter information:
   *                            | p: pointer to problem parameters
   *                            | plist: list of parameters with respect
   *                            |        to which sensitivities are to be
   *                            |        computed.
   *                            | pbar: order of magnitude info. 
   *                            |       Typically, if p[plist[i]] is nonzero, 
   *                            |       pbar[i]=p[plist[i]].
   *                            | [p=NULL]
   *                            | [plist=NULL]
   *                            | [pbar=NULL]
   *                            |
   * CVodeSetSensErrCon         | are sensitivity variables considered in
   *                            | the error control?
   *                            | [FALSE]
   *                            |
   * CVodeSetSensTolerances     | type of sensi absolute tolerances.
   *                            |
   *                            | sensitivity relative tolerance scalar.
   *                            |
   *                            | pointer to the array of sensi
   *                            | abs tol scalars or a pointer
   *                            | to the array of N_Vector sensi
   *                            | absolute tolerances.
   *                            | [itolS = itol]
   *                            | [reltolS = reltol]
   *                            | [abstolS estimated by CVODES]
   *                            |
   * CVodeSetSensMaxNonlinIters | Maximum number of nonlinear solver
   *                            | iterations at one solution.
   *                            | [3]
   *                            |
   * -----------------------------------------------------------------
   * The return values are the same as for CVodeSet*
   * -----------------------------------------------------------------
   */

  int CVodeSetSensRhsFn(void *cvode_mem, CVSensRhsFn f, void *fS_dataS);
  int CVodeSetSensRhs1Fn(void *cvode_mem, CVSensRhs1Fn fS, void *fS_data);
  int CVodeSetSensDQMethod(void *cvode_mem, int DQtype, realtype DQrhomax);
  int CVodeSetSensErrCon(void *cvode_mem, booleantype errconS);
  int CVodeSetSensMaxNonlinIters(void *cvode_mem, int maxcorS);
  int CVodeSetSensParams(void *cvode_mem, realtype *p, realtype *pbar, int *plist);
  int CVodeSetSensTolerances(void *cvode_mem, int itolS,
                             realtype reltolS, void *abstolS);

  /*
   * -----------------------------------------------------------------
   * Function : CVodeSensMalloc
   * -----------------------------------------------------------------
   * CVodeSensMalloc allocates and initializes memory related to
   * sensitivity computations.
   *
   * cvode_mem is pointer to CVODES memory returned by CVodeCreate
   *
   * Ns        is the number of sensitivities to be computed.
   *
   * ism       is the type of corrector used in sensitivity
   *           analysis. The legal values are: CV_SIMULTANEOUS,
   *           CV_STAGGERED, and CV_STAGGERED1 (see previous description)
   *
   * yS0       is the array of initial condition vectors for
   *           sensitivity variables.
   *
   * Return values:
   *   CV_SUCCESS
   *   CV_MEM_NULL
   *   CV_ILL_INPUT 
   *   CV_MEM_FAIL
   * -----------------------------------------------------------------
   */

  int CVodeSensMalloc(void *cvode_mem, int Ns, int ism, N_Vector *yS0);
    
  /*
   * -----------------------------------------------------------------
   * Function : CVodeSensReInit
   * -----------------------------------------------------------------
   * CVodeSensReInit re-initializes CVODES's sensitivity related
   * memory for a problem, assuming it has already been allocated
   * in prior calls to CVodeMalloc and CvodeSensMalloc.
   *
   * All problem specification inputs are checked for errors.
   * The number of sensitivities Ns is assumed to be unchanged
   * since the previous call to CVodeSensMalloc.
   * If any error occurs during initialization, it is reported to
   * the file whose file pointer is errfp.
   *
   * CVodeSensReInit potentially does some minimal memory allocation
   * (for the sensitivity absolute tolerance and for arrays of
   * counters used by the CV_STAGGERED1 method).
   *
   * The return value is equal to CV_SUCCESS = 0 if there were no
   * errors; otherwise it is a negative int equal to:
   *   CV_MEM_NULL  indicating cvode_mem was NULL, or
   *   CV_NO_SENS   indicating there was not a prior call to
   *                CVodeSensMalloc.
   *   CV_ILL_INPUT indicating an input argument was illegal
   *                (including an attempt to increase maxord).
   *   CV_MEM_FAIL  indicating a memory request failed.
   * In case of an error return, an error message is also printed.
   * -----------------------------------------------------------------
   */

  int CVodeSensReInit(void *cvode_mem, int ism, N_Vector *yS0);

  /*
   * -----------------------------------------------------------------
   * Function : CVodeSensToggleOff
   * -----------------------------------------------------------------
   * CVodeSensToggleOff deactivates sensitivity calculations.
   * It does NOT deallocate sensitivity-related memory so that 
   * sensitivity computations can be later toggled ON (through
   * CVodeSensReInit).
   * 
   * The return value is equal to CV_SUCCESS = 0 if there were no
   * errors or CV_MEM_NULL if cvode_mem was NULL
   * -----------------------------------------------------------------
   */

  int CVodeSensToggleOff(void *cvode_mem);

  /*
   * -----------------------------------------------------------------
   * Function : CVode
   * -----------------------------------------------------------------
   * CVode integrates the ODE over an interval in t.
   * If itask is CV_NORMAL, then the solver integrates from its
   * current internal t value to a point at or beyond tout, then
   * interpolates to t = tout and returns y(tout) in the user-
   * allocated vector yout. If itask is CV_ONE_STEP, then the solver
   * takes one internal time step and returns in yout the value of
   * y at the new internal time. In this case, tout is used only
   * during the first call to CVode to determine the direction of
   * integration and the rough scale of the problem.  If itask is
   * CV_NORMAL_TSTOP or CV_ONE_STEP_TSTOP, then CVode returns the
   * solution at tstop if that comes sooner than tout or the end of
   * the next internal step, respectively.  In any case,
   * the time reached by the solver is placed in (*tret). The
   * user is responsible for allocating the memory for this value.
   *
   * cvode_mem is the pointer to CVODES memory returned by
   *           CVodeCreate.
   *
   * tout  is the next time at which a computed solution is desired.
   *
   * yout  is the computed solution vector. In CV_NORMAL mode with no
   *       errors and no roots found, yout=y(tout).
   *
   * tret  is a pointer to a real location. CVode sets (*tret) to
   *       the time reached by the solver and returns yout=y(*tret).
   *
   * itask is CV_NORMAL, CV_ONE_STEP, CV_NORMAL_TSTOP, or CV_ONE_STEP_TSTOP.
   *       These four modes are described above.
   *
   * Here is a brief description of each return value:
   *
   * CV_SUCCESS:     CVode succeeded and no roots were found.
   *
   * CV_ROOT_RETURN: CVode succeeded, and found one or more roots.
   *                 If nrtfn > 1, call CVodeGetRootInfo to see
   *                 which g_i were found to have a root at (*tret).
   *
   * CV_TSTOP_RETURN: CVode succeded and returned at tstop.
   *
   * CV_MEM_NULL:    The cvode_mem argument was NULL.
   *
   * CV_NO_MALLOC:   cvode_mem was not allocated.
   *
   * CV_ILL_INPUT:   One of the inputs to CVode is illegal. This
   *                 includes the situation when a component of the
   *                 error weight vectors becomes < 0 during
   *                 internal time-stepping. The ILL_INPUT flag
   *                 will also be returned if the linear solver
   *                 routine CV--- (called by the user after
   *                 calling CVodeCreate) failed to set one of the
   *                 linear solver-related fields in cvode_mem or
   *                 if the linear solver's init routine failed. In
   *                 any case, the user should see the printed
   *                 error message for more details.
   *
   * CV_TOO_MUCH_WORK: The solver took mxstep internal steps but
   *                 could not reach tout. The default value for
   *                 mxstep is MXSTEP_DEFAULT = 500.
   *
   * CV_TOO_MUCH_ACC: The solver could not satisfy the accuracy
   *                 demanded by the user for some internal step.
   *
   * CV_ERR_FAILURE: Error test failures occurred too many times
   *                 (= MXNEF = 7) during one internal time step or
   *                 occurred with |h| = hmin.
   *
   * CV_CONV_FAILURE: Convergence test failures occurred too many
   *                 times (= MXNCF = 10) during one internal time
   *                 step or occurred with |h| = hmin.
   *
   * CV_LINIT_FAIL:  The linear solver's initialization function 
   *                 failed.
   *
   * CV_LSETUP_FAIL: The linear solver's setup routine failed in an
   *                 unrecoverable manner.
   *
   * CV_LSOLVE_FAIL: The linear solver's solve routine failed in an
   *                 unrecoverable manner.
   * -----------------------------------------------------------------
   */

  int CVode(void *cvode_mem, realtype tout, N_Vector yout,
            realtype *tret, int itask);

  /*
   * -----------------------------------------------------------------
   * Function : CVodeGetDky
   * -----------------------------------------------------------------
   * CVodeGetDky computes the kth derivative of the y function at
   * time t, where tn-hu <= t <= tn, tn denotes the current
   * internal time reached, and hu is the last internal step size
   * successfully used by the solver. The user may request
   * k=0, 1, ..., qu, where qu is the current order. The
   * derivative vector is returned in dky. This vector must be
   * allocated by the caller. It is only legal to call this
   * function after a successful return from CVode.
   *
   * cvode_mem is the pointer to CVODES memory returned by
   *           CVodeCreate.
   *
   * t   is the time at which the kth derivative of y is evaluated.
   *     The legal range for t is [tn-hu,tn] as described above.
   *
   * k   is the order of the derivative of y to be computed. The
   *     legal range for k is [0,qu] as described above.
   *
   * dky is the output derivative vector [(D_k)y](t).
   *
   * The return values for CVodeGetDky are defined below.
   * Here is a brief description of each return value:
   *
   * CV_SUCCESS: CVodeGetDky succeeded.
   *
   * CV_BAD_K : k is not in the range 0, 1, ..., qu.
   *
   * CV_BAD_T : t is not in the interval [tn-hu,tn].
   *
   * CV_BAD_DKY : The dky argument was NULL.
   *
   * CV_MEM_NULL : The cvode_mem argument was NULL.
   * -----------------------------------------------------------------
   */

  int CVodeGetDky(void *cvode_mem, realtype t, int k, N_Vector dky);

  /*
   * -----------------------------------------------------------------
   * Integrator optional output extraction functions
   * -----------------------------------------------------------------
   * The following functions can be called to get optional outputs
   * and statistics related to the main integrator.
   * -----------------------------------------------------------------
   * CVodeGetWorkSpace returns the CVODES real and integer workspaces
   * CVodeGetNumSteps returns the cumulative number of internal
   *                  steps taken by the solver
   * CVodeGetNumRhsEvals returns the number of calls to the user's
   *                  f function
   * CVodeGetNumLinSolvSetups returns the number of calls made to
   *                  the linear solver's setup routine
   * CVodeGetNumErrTestFails returns the number of local error test
   *                  failures that have occured
   * CVodeGetLastOrder returns the order used during the last
   *                  internal step
   * CVodeGetCurrentOrder returns the order to be used on the next
   *                  internal step
   * CVodeGetNumStabLimOrderReds returns the number of order
   *                  reductions due to stability limit detection
   * CVodeGetActualInitStep returns the actual initial step size
   *                  used by CVODES
   * CVodeGetLastStep returns the step size for the last internal
   *                  step
   * CVodeGetCurrentStep returns the step size to be attempted on
   *                  the next internal step
   * CVodeGetCurrentTime returns the current internal time reached
   *                  by the solver
   * CVodeGetTolScaleFactor returns a suggested factor by which the
   *                  user's tolerances should be scaled when too
   *                  much accuracy has been requested for some
   *                  internal step
   * CVodeGetErrWeights returns the current error weight vector.
   *                    The user must allocate space for eweight.
   * CVodeGetEstLocalErrors returns the vector of estimated local
   *                  errors. The user must allocate space for ele.
   * CVodeGetNumGEvals returns the number of calls to the user's
   *                  g function (for rootfinding)
   * CVodeGetRootInfo returns the indices for which g_i was found to 
   *                  have a root. The user must allocate space for 
   *                  rootsfound. For i = 0 ... nrtfn-1, 
   *                  rootsfound[i] = 1 if g_i has a root, and = 0 if not.
   *
   * CVodeGet* return values:
   *   CV_SUCCESS   if succesful
   *   CV_MEM_NULL  if the cvode memory was NULL
   *   CV_NO_SLDET  if stability limit was not turned on
   * -----------------------------------------------------------------
   */

  int CVodeGetWorkSpace(void *cvode_mem, long int *lenrw, long int *leniw);
  int CVodeGetNumSteps(void *cvode_mem, long int *nsteps);
  int CVodeGetNumRhsEvals(void *cvode_mem, long int *nfevals);
  int CVodeGetNumLinSolvSetups(void *cvode_mem, long int *nlinsetups);
  int CVodeGetNumErrTestFails(void *cvode_mem, long int *netfails);
  int CVodeGetLastOrder(void *cvode_mem, int *qlast);
  int CVodeGetCurrentOrder(void *cvode_mem, int *qcur);
  int CVodeGetNumStabLimOrderReds(void *cvode_mem, long int *nslred);
  int CVodeGetActualInitStep(void *cvode_mem, realtype *hinused);
  int CVodeGetLastStep(void *cvode_mem, realtype *hlast);
  int CVodeGetCurrentStep(void *cvode_mem, realtype *hcur);
  int CVodeGetCurrentTime(void *cvode_mem, realtype *tcur);
  int CVodeGetTolScaleFactor(void *cvode_mem, realtype *tolsfac);
  int CVodeGetErrWeights(void *cvode_mem, N_Vector eweight);
  int CVodeGetEstLocalErrors(void *cvode_mem, N_Vector ele);
  int CVodeGetNumGEvals(void *cvode_mem, long int *ngevals);
  int CVodeGetRootInfo(void *cvode_mem, int *rootsfound);

  /*
   * -----------------------------------------------------------------
   * As a convenience, the following functions provides the
   * optional outputs in one group.
   * -----------------------------------------------------------------
   */

  int CVodeGetIntegratorStats(void *cvode_mem, long int *nsteps,
                              long int *nfevals, long int *nlinsetups,
                              long int *netfails, int *qlast,
                              int *qcur, realtype *hinused, realtype *hlast,
                              realtype *hcur, realtype *tcur);

  /*
   * -----------------------------------------------------------------
   * Nonlinear solver optional output extraction functions
   * -----------------------------------------------------------------
   * The following functions can be called to get optional outputs
   * and statistics related to the nonlinear solver.
   * -----------------------------------------------------------------
   * CVodeGetNumNonlinSolvIters returns the number of nonlinear
   *                            solver iterations performed.
   * CVodeGetNumNonlinSolvConvFails returns the number of nonlinear
   *                                convergence failures.
   * -----------------------------------------------------------------
   */

  int CVodeGetNumNonlinSolvIters(void *cvode_mem, long int *nniters);
  int CVodeGetNumNonlinSolvConvFails(void *cvode_mem, long int *nncfails);

  /*
   * -----------------------------------------------------------------
   * As a convenience, the following function provides the
   * nonlinear solver optional outputs in a group.
   * -----------------------------------------------------------------
   */

  int CVodeGetNonlinSolvStats(void *cvode_mem, long int *nniters,
                              long int *nncfails);


  /*
   * -----------------------------------------------------------------
   * The following function returns the name of the constant 
   * associated with a CVODES return flag
   * -----------------------------------------------------------------
   */

  char *CVodeGetReturnFlagName(int flag);

  /*
   * -----------------------------------------------------------------
   * Quadrature integration solution extraction routines
   * -----------------------------------------------------------------
   * The following functions can be called to obtain the quadrature
   * variables after a successful integration step.
   * If quadratures were not computed, they return CV_NO_QUAD.
   * -----------------------------------------------------------------
   */

  int CVodeGetQuad(void *cvode_mem, realtype t, N_Vector yQout);
  int CVodeGetQuadDky(void *cvode_mem, realtype t, int k, N_Vector dky);

  /*
   * -----------------------------------------------------------------
   * Quadrature integration optional output extraction routines
   * -----------------------------------------------------------------
   * The following functions can be called to get optional outputs
   * and statistics related to the integration of quadratures.
   * -----------------------------------------------------------------
   * CVodeGetQuadNumRhsEvals returns the number of calls to the
   *                         user function fQ defining the right hand
   *                         side of the quadrature variables.
   * CVodeGetQuadNumErrTestFails returns the number of local error
   *                             test failures for quadrature variables.
   * CVodeGetQuadErrWeights returns the vector of error weights for
   *                        the quadrature variables. The user must
   *                        allocate space for ewtQ.
   * -----------------------------------------------------------------
   */

  int CVodeGetQuadNumRhsEvals(void *cvode_mem, long int *nfQevals);
  int CVodeGetQuadNumErrTestFails(void *cvode_mem, long int *nQetfails);
  int CVodeGetQuadErrWeights(void *cvode_mem, N_Vector eQweight);

  /*
   * -----------------------------------------------------------------
   * As a convenience, the following function provides the
   * optional outputs in a group.
   * -----------------------------------------------------------------
   */

  int CVodeGetQuadStats(void *cvode_mem, long int *nfQevals,
                        long int *nQetfails);

  /*
   * -----------------------------------------------------------------
   * Forward sensitivity solution extraction routines
   * -----------------------------------------------------------------
   * CVodeGetSensDky1 computes the kth derivative of the is-th
   * sensitivity (is=1, 2, ..., Ns) of the y function at time t,
   * where tn-hu <= t <= tn, tn denotes the current internal time
   * reached, and hu is the last internal step size successfully
   * used by the solver. The user may request k=0, 1, ..., qu,
   * where qu is the current order.
   * The is-th sensitivity derivative vector is returned in dky.
   * This vector must be allocated by the caller. It is only legal
   * to call this function after a successful return from CVode
   * with sensitivty computations enabled.
   * Arguments have the same meaning as in CVodeDky.
   *
   * CVodeGetSensDky computes the k-th derivative of all
   * sensitivities of the y function at time t. It repeatedly calls
   * CVodeGetSensDky. The argument dkyA must be a pointer to
   * N_Vector and must be allocated by the user to hold at least Ns
   * vectors.
   *
   * CVodeGetSens1 returns the is-th sensitivity of the y function
   * at the time t. The argument ySout must be an N_Vector and must
   * be allocated by the user.
   *
   * CVodeGetSens returns sensitivities of the y function at
   * the time t. The argument ySout must be a pointer to N_Vector
   * and must be allocated by the user to hold at least Ns vectors.
   *
   * Return values are similar to those of CVodeDky. Additionally,
   * CVodeSensDky can return CV_NO_SENS if sensitivities were
   * not computed and CV_BAD_IS if is < 0 or is >= Ns.
   * -----------------------------------------------------------------
   */

  int CVodeGetSens(void *cvode_mem, realtype t, N_Vector *ySout);
  int CVodeGetSens1(void *cvode_mem, realtype t, int is, N_Vector ySout);
  int CVodeGetSensDky(void *cvode_mem, realtype t, int k,
                      N_Vector *dkyA);
  int CVodeGetSensDky1(void *cvode_mem, realtype t, int k,
                       int is, N_Vector dky);

  /*
   * -----------------------------------------------------------------
   * Forward sensitivity optional output extraction routines
   * -----------------------------------------------------------------
   * The following functions can be called to get optional outputs
   * and statistics related to the integration of sensitivities.
   * -----------------------------------------------------------------
   * CVodeGetNumSensRhsEvals returns the number of calls to the
   *                         sensitivity right hand side routine.
   * CVodeGetNumRhsEvalsSens returns the number of calls to the
   *                         user f routine due to finite difference
   *                         evaluations of the sensitivity equations.
   * CVodeGetNumSensErrTestFails returns the number of local error
   *                             test failures for sensitivity variables.
   * CVodeGetNumSensLinSolvSetups returns the number of calls made
   *                              to the linear solver's setup routine
   *                              due to sensitivity computations.
   * CVodeGetSensErrWeights returns the sensitivity error weight
   *                        vectors. The user need not allocate space
   *                        for ewtS.
   * -----------------------------------------------------------------
   */

  int CVodeGetNumSensRhsEvals(void *cvode_mem, long int *nfSevals);
  int CVodeGetNumRhsEvalsSens(void *cvode_mem, long int *nfevalsS);
  int CVodeGetNumSensErrTestFails(void *cvode_mem, long int *nSetfails);
  int CVodeGetNumSensLinSolvSetups(void *cvode_mem, long int *nlinsetupsS);
  int CVodeGetSensErrWeights(void *cvode_mem, N_Vector_S eSweight);

  /*
   * -----------------------------------------------------------------
   * As a convenience, the following function provides the
   * optional outputs in a group.
   * -----------------------------------------------------------------
   */

  int CVodeGetSensStats(void *cvode_mem, long int *nfSevals, long int *nfevalsS,
                        long int *nSetfails, long int *nlinsetupsS);

  /*
   * -----------------------------------------------------------------
   * Sensitivity nonlinear solver optional output extraction
   * -----------------------------------------------------------------
   * The following functions can be called to get optional outputs
   * and statistics related to the sensitivity nonlinear solver.
   * -----------------------------------------------------------------
   * CVodeGetNumSensNonlinSolvIters returns the total number of
   *                                nonlinear iterations for sensitivity
   *                                variables.
   * CVodeGetNumSensNonlinSolvConvFails returns the total number of
   *                                    nonlinear convergence failures
   *                                    for sensitivity variables
   * CVodeGetNumStgrSensNonlinSolvIters returns a vector of Ns
   *                                    nonlinear iteration counters
   *                                    for sensitivity variables
   *                                    in the CV_STAGGERED1 method.
   * CVodeGetNumStgrSensNonlinSolvConvFails returns a vector of Ns
   *                                        nonlinear solver convergence
   *                                        failure counters for
   *                                        sensitivity variables in
   *                                        the CV_STAGGERED1 method.
   * -----------------------------------------------------------------
   */

  int CVodeGetNumSensNonlinSolvIters(void *cvode_mem, long int *nSniters);
  int CVodeGetNumSensNonlinSolvConvFails(void *cvode_mem, long int *nSncfails);
  int CVodeGetNumStgrSensNonlinSolvIters(void *cvode_mem, long int *nSTGR1niters);
  int CVodeGetNumStgrSensNonlinSolvConvFails(void *cvode_mem, 
                                             long int *nSTGR1ncfails);

  /*
   * -----------------------------------------------------------------
   * As a convenience, the following function provides the      
   * optional outputs in groups.                                    
   * -----------------------------------------------------------------
   */

  int CVodeGetSensNonlinSolvStats(void *cvode_mem, long int *nSniters,
                                  long int *nSncfails);

  /*
   * -----------------------------------------------------------------
   * Function : CVodeFree
   * -----------------------------------------------------------------
   * CVodeFree frees the problem memory cvode_mem allocated by
   * CVodeMalloc.  Its only argument is the pointer cvode_mem
   * returned by CVodeCreate.
   * -----------------------------------------------------------------
   */

  void CVodeFree(void **cvode_mem);

  /*
   * -----------------------------------------------------------------
   * Function : CVodeQuadFree
   * -----------------------------------------------------------------
   * CVodeQuadFree frees the problem memory in cvode_mem allocated
   * for quadrature integration. Its only argument is the pointer
   * cvode_mem returned by CVodeCreate.
   * -----------------------------------------------------------------
   */

  void CVodeQuadFree(void *cvode_mem);

  /*
   * -----------------------------------------------------------------
   * Function : CVodeSensFree
   * -----------------------------------------------------------------
   * CVodeSensFree frees the problem memory in cvode_mem allocated
   * for sensitivity analysis. Its only argument is the pointer
   * cvode_mem returned by CVodeCreate.
   * -----------------------------------------------------------------
   */

  void CVodeSensFree(void *cvode_mem);

  /*
   * -----------------------------------------------------------------
   * CVadjMalloc
   * -----------------------------------------------------------------
   * CVadjMalloc specifies some parameters for the adjoint problem and
   * allocates space for the global CVODEA memory structure.
   * -----------------------------------------------------------------
   */
  
  void *CVadjMalloc(void *cvode_mem, long int steps, int interp);

  /*
   * -----------------------------------------------------------------
   * CVadjSetInterpType
   * -----------------------------------------------------------------
   * Changes the interpolation type. 
   * Must be called only after CVadjMalloc
   * -----------------------------------------------------------------
   */
  
  int CVadjSetInterpType(void *cvadj_mem, int interp);

  /*
   * -----------------------------------------------------------------
   * CVodeF
   * -----------------------------------------------------------------
   * CVodeF integrates towards tout and returns solution into yout.
   * In the same time, it stores check point data every 'steps'.
   *
   * CVodeF can be called repeatedly by the user.
   *
   * ncheckPtr points to the number of check points stored so far.
   *
   * Return values:
   *    CV_SUCCESS
   *    CVADJ_MEM_FAIL
   *    any CVode return value
   * -----------------------------------------------------------------
   */

  int CVodeF(void *cvadj_mem, realtype tout, N_Vector yout,
             realtype *tret, int itask, int *ncheckPtr);

  /*
   * -----------------------------------------------------------------
   * Interfaces to CVODES functions for setting-up the
   *  backward integration
   * -----------------------------------------------------------------
   * CVodeCreateB, CVodeMallocB, CVodeSet*B
   *    These functions are just wrappers around the corresponding
   *    functions in cvodes.h, with some particularizations for the
   *    backward integration.
   * -----------------------------------------------------------------
   * CVodeSetQuad*B, CVodeQuadMallocB, CVodeQuadReInitB
   * -----------------------------------------------------------------
   */

  int CVodeCreateB(void *cvadj_mem, int lmmB, int iterB);

  int CVodeMallocB(void *cvadj_mem, CVRhsFnB fB,
                   realtype tB0, N_Vector yB0,
                   int itolB, realtype reltolB, void *abstolB);
  
  int CVodeSetErrHandlerFnB(void *cvadj_mem, CVErrHandlerFn ehfunB, void *eh_dataB);
  int CVodeSetErrFileB(void *cvadj_mem, FILE *errfpB);
  int CVodeSetIterTypeB(void *cvadj_mem, int iterB);
  int CVodeSetFdataB(void *cvadj_mem, void *f_dataB);
  int CVodeSetMaxOrdB(void *cvadj_mem, int maxordB);
  int CVodeSetMaxNumStepsB(void *cvadj_mem, long int mxstepsB);
  int CVodeSetStabLimDetB(void *cvadj_mem, booleantype stldetB);
  int CVodeSetInitStepB(void *cvadj_mem, realtype hinB);
  int CVodeSetMinStepB(void *cvadj_mem, realtype hminB);
  int CVodeSetMaxStepB(void *cvadj_mem, realtype hmaxB);
  
  int CVodeReInitB(void *cvadj_mem, CVRhsFnB fB,
                   realtype tB0, N_Vector yB0,
                   int itolB, realtype reltolB, void *abstolB);
    
  int CVodeSetQuadFdataB(void *cvadj_mem, void *fQ_dataB);
  int CVodeSetQuadErrConB(void *cvadj_mem, booleantype errconQB,
                          int itolQB, realtype reltolQB, void *abstolQB);
  int CVodeQuadMallocB(void *cvadj_mem, CVQuadRhsFnB fQB, N_Vector yQB0);
  int CVodeQuadReInitB(void *cvadj_mem, CVQuadRhsFnB fQB, N_Vector yQB0);
    
  /*
   * -----------------------------------------------------------------
   * CVodeB
   * -----------------------------------------------------------------
   * CVodeB performs the backward integration from tfinal to
   * tinitial through a sequence of forward-backward runs in
   * between consecutive check points. It returns the values of
   * the adjoint variables and any existing quadrature variables
   * at tinitial.
   * -----------------------------------------------------------------
   */
  
  int CVodeB(void *cvadj_mem, realtype tBout, N_Vector yBout,
             realtype *tBret, int itaskB);
  
  /*
   * -----------------------------------------------------------------
   * CVodeGetQuadB
   * -----------------------------------------------------------------
   * CVodeGetQuadB extracts values for quadrature variables in
   * the N_Vector qB.
   * -----------------------------------------------------------------
   */
  
  int CVodeGetQuadB(void *cvadj_mem, N_Vector qB);
  
  /*
   * -----------------------------------------------------------------
   * CVadjFree
   * -----------------------------------------------------------------
   * CVadjFree frees the memory allocated by CVadjMalloc.
   * -----------------------------------------------------------------
   */
  
  void CVadjFree(void **cvadj_mem);
  
  /*
   * -----------------------------------------------------------------
   * CVadjGetCVodeBmem
   * -----------------------------------------------------------------
   * CVadjGetCVodeBmem returns a (void *) pointer to the CVODES
   * memory allocated for the backward problem. This pointer can
   * then be used to call any of the CVodeGet* CVODES routines to
   * extract optional output for the backward integration phase.
   * -----------------------------------------------------------------
   */
  
  void *CVadjGetCVodeBmem(void *cvadj_mem);

  /*
   * -----------------------------------------------------------------
   * The following function returns the name of the constant 
   * associated with a CVODEA-specific return flag
   * -----------------------------------------------------------------
   */
  
  char *CVadjGetReturnFlagName(int flag);


  /*
   * -----------------------------------------------------------------
   * CVadjGetY
   *    Returns the interpolated forward solution at time t. This
   *    function is a wrapper around the interpType-dependent internal
   *    function.
   *    The calling function must allocate space for y.
   * -----------------------------------------------------------------
   */

  int CVadjGetY(void *cvadj_mem, realtype t, N_Vector y);

  /*
   * -----------------------------------------------------------------
   * CVadjGetCheckPointsInfo
   *    Loads an array of nckpnts structures of type CVadjCheckPointRec.
   *    The user must allocate space for ckpnt (ncheck+1).
   * -----------------------------------------------------------------
   */

  typedef struct {
    void *my_addr;
    void *next_addr;
    realtype t0;
    realtype t1;
    long int nstep;
    int order;
    realtype step;
  } CVadjCheckPointRec;

  int CVadjGetCheckPointsInfo(void *cvadj_mem, CVadjCheckPointRec *ckpnt);

  /*
   * -----------------------------------------------------------------
   * CVadjGetDataPointHermite
   *    Returns the 2 vectors stored for cubic Hermite interpolation 
   *    at the data point 'which'. The user must allocate space for
   *    y and yd. Returns CVADJ_MEM_NULL if cvadj_mem is NULL.
   *    Returns CV_ILL_INPUT if interpType != CV_HERMITE.
   * CVadjGetDataPointPolynomial
   *    Returns the vector stored for polynomial interpolation 
   *    at the data point 'which'. The user must allocate space for
   *    y. Returns CVADJ_MEM_NULL if cvadj_mem is NULL.
   *    Returns CV_ILL_INPUT if interpType != CV_POLYNOMIAL.
   * -----------------------------------------------------------------
   */

  int CVadjGetDataPointHermite(void *cvadj_mem, long int which,
                               realtype *t, N_Vector y, N_Vector yd);
  
  int CVadjGetDataPointPolynomial(void *cvadj_mem, long int which,
                                  realtype *t, int *order, N_Vector y);

  /* 
   * ===============================================================
   * DEVELOPMENT USER-CALLABLE FUNCTIONS
   * ===============================================================
   */

  /*
   * -----------------------------------------------------------------
   * CVadjGetCurrentCheckPoint
   *    Returns the address of the 'active' check point.
   * -----------------------------------------------------------------
   */

  int CVadjGetCurrentCheckPoint(void *cvadj_mem, void **addr);

#ifdef __cplusplus
}
#endif

#endif
```
