## Supplementary scripts for "A multi-scale *in silico* mouse model for insulin resistance and humanoid type 2 diabetes": cvodes_band.h

```
/*
 * -----------------------------------------------------------------
 * $Revision: 749 $
 * $Date: 2011-06-22 14:49:10 +0200 (Wed, 22 Jun 2011) $
 * ----------------------------------------------------------------- 
 * Programmer(s): Radu Serban @ LLNL
 * -----------------------------------------------------------------
 * Copyright (c) 2005, The Regents of the University of California.
 * Produced at the Lawrence Livermore National Laboratory.
 * All rights reserved.
 * For details, see the LICENSE file.
 * -----------------------------------------------------------------
 * This is the header file for the CVODES band linear solver, CVBAND.
 *
 *
 * Part I contains type definitions and function prototypes for using
 * CVBAND on forward problems (IVP integration and/or FSA)
 *
 * Part II contains type definitions and function prototypes for using
 * CVBAND on adjoint (backward) problems
 * -----------------------------------------------------------------
 */

#ifndef _CVSBAND_H
#define _CVSBAND_H

#ifdef __cplusplus  /* wrapper to enable C++ usage */
extern "C" {
#endif

#include <sundials/sundials_band.h>
#include <sundials/sundials_nvector.h>

  /*
   * -----------------------------------------------------------------
   * CVBAND solver constants
   * -----------------------------------------------------------------
   * CVB_MSBJ : maximum number of steps between band Jacobian
   *            evaluations
   *
   * CVB_DGMAX : maximum change in gamma between band Jacobian
   *             evaluations
   * -----------------------------------------------------------------
   */

#define CVB_MSBJ  50
#define CVB_DGMAX RCONST(0.2)

  /*
   * -----------------------------------------------------------------
   * CVBAND return values
   * -----------------------------------------------------------------
   */

#define CVBAND_SUCCESS           0
#define CVBAND_MEM_NULL         -1
#define CVBAND_LMEM_NULL        -2
#define CVBAND_ILL_INPUT        -3
#define CVBAND_MEM_FAIL         -4

  /* Additional last_flag values */

#define CVBAND_JACFUNC_UNRECVR  -5
#define CVBAND_JACFUNC_RECVR    -6

  /* Return values for adjoint module */

#define CVBAND_ADJMEM_NULL      -101
#define CVBAND_LMEMB_NULL       -102

  /* 
   * -----------------------------------------------------------------
   * PART I - forward problems
   * -----------------------------------------------------------------
   */


  /*
   * -----------------------------------------------------------------
   * Type : CVBandJacFn
   * -----------------------------------------------------------------
   * A band Jacobian approximation function Jac must have the
   * prototype given below. Its parameters are:
   *
   * N is the length of all vector arguments.
   *
   * mupper is the upper half-bandwidth of the approximate banded
   * Jacobian. This parameter is the same as the mupper parameter
   * passed by the user to the CVBand function.
   *
   * mlower is the lower half-bandwidth of the approximate banded
   * Jacobian. This parameter is the same as the mlower parameter
   * passed by the user to the CVBand function.
   *
   * J is the band matrix (of type BandMat) that will be loaded
   * by a CVBandJacFn with an approximation to the Jacobian matrix
   * J = (df_i/dy_j) at the point (t,y).
   * J is preset to zero, so only the nonzero elements need to be
   * loaded. Three efficient ways to load J are:
   *
   * (1) (with macros - no explicit data structure references)
   *    for (j=0; j < n; j++) {
   *       col_j = BAND_COL(J,j);
   *       for (i=j-mupper; i <= j+mlower; i++) {
   *         generate J_ij = the (i,j)th Jacobian element
   *         BAND_COL_ELEM(col_j,i,j) = J_ij;
   *       }
   *     }
   *
   * (2) (with BAND_COL macro, but without BAND_COL_ELEM macro)
   *    for (j=0; j < n; j++) {
   *       col_j = BAND_COL(J,j);
   *       for (k=-mupper; k <= mlower; k++) {
   *         generate J_ij = the (i,j)th Jacobian element, i=j+k
   *         col_j[k] = J_ij;
   *       }
   *     }
   *
   * (3) (without macros - explicit data structure references)
   *     offset = J->smu;
   *     for (j=0; j < n; j++) {
   *       col_j = ((J->data)[j])+offset;
   *       for (k=-mupper; k <= mlower; k++) {
   *         generate J_ij = the (i,j)th Jacobian element, i=j+k
   *         col_j[k] = J_ij;
   *       }
   *     }
   * Caution: J->smu is generally NOT the same as mupper.
   *
   * The BAND_ELEM(A,i,j) macro is appropriate for use in small
   * problems in which efficiency of access is NOT a major concern.
   *
   * t is the current value of the independent variable.
   *
   * y is the current value of the dependent variable vector,
   *      namely the predicted value of y(t).
   *
   * fy is the vector f(t,y).
   *
   * jac_data is a pointer to user data - the same as the jac_data
   *          parameter passed to CVBand.
   *
   * NOTE: If the user's Jacobian routine needs other quantities,
   *     they are accessible as follows: hcur (the current stepsize)
   *     and ewt (the error weight vector) are accessible through
   *     CVodeGetCurrentStep and CVodeGetErrWeights, respectively
   *     (see cvode.h). The unit roundoff is available as
   *     UNIT_ROUNDOFF defined in sundials_types.h
   *
   * tmp1, tmp2, and tmp3 are pointers to memory allocated for
   * vectors of length N which can be used by a CVBandJacFn
   * as temporary storage or work space.
   *
   * A CVBandJacFn should return 0 if successful, a positive value if 
   * a recoverable error occurred, and a negative value if an 
   * unrecoverable error occurred.
   * -----------------------------------------------------------------
   */

  typedef int (*CVBandJacFn)(long int N, long int mupper, long int mlower,
                             BandMat J, realtype t,
                             N_Vector y, N_Vector fy, void *jac_data,
                             N_Vector tmp1, N_Vector tmp2, N_Vector tmp3);

  /*
   * -----------------------------------------------------------------
   * Function : CVBand
   * -----------------------------------------------------------------
   * A call to the CVBand function links the main CVODE integrator
   * with the CVBAND linear solver.
   *
   * cvode_mem is the pointer to the integrator memory returned by
   *           CVodeCreate.
   *
   * N is the size of the ODE system.
   *
   * mupper is the upper bandwidth of the band Jacobian
   *        approximation.
   *
   * mlower is the lower bandwidth of the band Jacobian
   *        approximation.
   *
   * The return value of CVBand is one of:
   *    CVBAND_SUCCESS   if successful
   *    CVBAND_MEM_NULL  if the cvode memory was NULL
   *    CVBAND_MEM_FAIL  if there was a memory allocation failure
   *    CVBAND_ILL_INPUT if a required vector operation is missing or
   *                     if a bandwidth has an illegal value.
   * -----------------------------------------------------------------
   */

  int CVBand(void *cvode_mem, long int N,
             long int mupper, long int mlower);

  /*
   * -----------------------------------------------------------------
   * Optional inputs to the CVBAND linear solver
   * -----------------------------------------------------------------
   *
   * CVBandSetJacFn specifies the band Jacobian approximation
   *                routine to be used. A user-supplied bjac routine
   *                must be of type CVBandJacFn. By default, a difference
   *                quotient routine CVBandDQJac, supplied with this
   *                solver is used.
   *                It also specifies a pointer to user data which is
   *                passed to the bjac routine every time it is called.
   *
   * The return value of CVBandSet* is one of:
   *    CVBAND_SUCCESS   if successful
   *    CVBAND_MEM_NULL  if the cvode memory was NULL
   *    CVBAND_LMEM_NULL if the cvband memory was NULL
   * -----------------------------------------------------------------
   */

  int CVBandSetJacFn(void *cvode_mem, CVBandJacFn bjac, void *jac_data);

  /*
   * -----------------------------------------------------------------
   * Optional outputs from the CVBAND linear solver
   * -----------------------------------------------------------------
   *
   * CVBandGetWorkSpace returns the real and integer workspace used
   *                    by CVBAND.
   * CVBandGetNumJacEvals returns the number of calls made to the
   *                      Jacobian evaluation routine bjac.
   * CVBandGetNumRhsEvals returns the number of calls to the user
   *                      f routine due to finite difference Jacobian
   *                      evaluation.
   * CVBandGetLastFlag returns the last error flag set by any of
   *                   the CVBAND interface functions.
   *
   * The return value of CVBandGet* is one of:
   *    CVBAND_SUCCESS   if successful
   *    CVBAND_MEM_NULL  if the cvode memory was NULL
   *    CVBAND_LMEM_NULL if the cvband memory was NULL
   * -----------------------------------------------------------------
   */

  int CVBandGetWorkSpace(void *cvode_mem, long int *lenrwLS, long int *leniwLS);
  int CVBandGetNumJacEvals(void *cvode_mem, long int *njevals);
  int CVBandGetNumRhsEvals(void *cvode_mem, long int *nfevalsLS);
  int CVBandGetLastFlag(void *cvode_mem, int *flag);

  /*
   * -----------------------------------------------------------------
   * The following function returns the name of the constant 
   * associated with a CVBAND return flag
   * -----------------------------------------------------------------
   */

  char *CVBandGetReturnFlagName(int flag);

  /* 
   * -----------------------------------------------------------------
   * PART II - backward problems
   * -----------------------------------------------------------------
   */

  /*
   * -----------------------------------------------------------------
   * Type : CVBandJacFnB
   * -----------------------------------------------------------------
   * A band Jacobian approximation function JacB for the adjoint 
   * (backward) problem must have the prototype given below. 
   * -----------------------------------------------------------------
   */

  typedef int (*CVBandJacFnB)(long int nB, long int mupperB,
                              long int mlowerB, BandMat JB,
                              realtype t, N_Vector y,
                              N_Vector yB, N_Vector fyB,
                              void *jac_dataB, N_Vector tmp1B,
                              N_Vector tmp2B, N_Vector tmp3B);
  
  /*
   * -----------------------------------------------------------------
   * Functions: CVBandB, CVBandSet*B
   * -----------------------------------------------------------------
   * CVBandB links the main CVODE integrator with the CVBAND
   * linear solver for the backward integration.
   * -----------------------------------------------------------------
   */

  int CVBandB(void *cvadj_mem, long int nB,
              long int mupperB, long int mlowerB);
  
  int CVBandSetJacFnB(void *cvadj_mem, CVBandJacFnB bjacB, void *jac_dataB);


#ifdef __cplusplus
}
#endif

#endif
```
