## Supplementary scripts for "A multi-scale *in silico* mouse model for insulin resistance and humanoid type 2 diabetes": cvodes_bandpre.h

```
/*
 * -----------------------------------------------------------------
 * $Revision: 749 $
 * $Date: 2011-06-22 14:49:10 +0200 (Wed, 22 Jun 2011) $
 * ----------------------------------------------------------------- 
 * Programmer(s): Radu Serban @ LLNL
 * -----------------------------------------------------------------
 * Copyright (c) 2005, The Regents of the University of California.
 * Produced at the Lawrence Livermore National Laboratory.
 * All rights reserved.
 * For details, see the LICENSE file.
 * -----------------------------------------------------------------
 * This is the header file for the CVBANDPRE module, which
 * provides a banded difference quotient Jacobian-based
 * preconditioner and solver routines for use with CVSPGMR,
 * CVSPBCG, or CVSPTFQMR.
 *
 *
 * Part I contains type definitions and function prototypes for using
 * CVBANDPRE on forward problems (IVP integration and/or FSA)
 *
 * Part II contains type definitions and function prototypes for using
 * CVBANDPRE on adjopint (backward) problems
 * -----------------------------------------------------------------
 */

#ifndef _CVSBANDPRE_H
#define _CVSBANDPRE_H

#ifdef __cplusplus  /* wrapper to enable C++ usage */
extern "C" {
#endif

  /* CVBANDPRE return values */

#define CVBANDPRE_SUCCESS           0
#define CVBANDPRE_PDATA_NULL      -11
#define CVBANDPRE_RHSFUNC_UNRECVR -12

#define CVBANDPRE_ADJMEM_NULL     -111
#define CVBANDPRE_MEM_FAIL        -112

#include <sundials/sundials_nvector.h>

  /* 
   * -----------------------------------------------------------------
   * PART I - forward problems
   * -----------------------------------------------------------------
   *
   * Summary:
   * These routines provide a band matrix preconditioner based on
   * difference quotients of the ODE right-hand side function f.
   * The user supplies parameters
   *   mu = upper half-bandwidth (number of super-diagonals)
   *   ml = lower half-bandwidth (number of sub-diagonals)
   * The routines generate a band matrix of bandwidth ml + mu + 1
   * and use this to form a preconditioner for use with the Krylov
   * linear solver in CVSP*. Although this matrix is intended to
   * approximate the Jacobian df/dy, it may be a very crude
   * approximation. The true Jacobian need not be banded, or its
   * true bandwith may be larger than ml + mu + 1, as long as the
   * banded approximation generated here is sufficiently accurate
   * to speed convergence as a preconditioner.
   *
   * Usage:
   *   The following is a summary of the usage of this module.
   *   Details of the calls to CVodeCreate, CVodeMalloc, CVSp*,
   *   and CVode are available in the User Guide.
   *   To use these routines, the sequence of calls in the user
   *   main program should be as follows:
   *
   *   #include <cvodes/cvodes_bandpre.h>
   *   #include <nvector_serial.h>
   *   ...
   *   void *bp_data;
   *   ...
   *   Set y0
   *   ...
   *   cvode_mem = CVodeCreate(...);
   *   ier = CVodeMalloc(...);
   *   ...
   *   bp_data = CVBandPrecAlloc(cvode_mem, N, mu, ml);
   *   ...
   *   flag = CVBPSptfqmr(cvode_mem, pretype, maxl, bp_data);
   *     -or-
   *   flag = CVBPSpgmr(cvode_mem, pretype, maxl, bp_data);
   *     -or-
   *   flag = CVBPSpbcg(cvode_mem, pretype, maxl, bp_data);
   *   ...
   *   flag = CVode(...);
   *   ...
   *   CVBandPrecFree(&bp_data);
   *   ...
   *   Free y0
   *   ...
   *   CVodeFree(cvode_mem);
   *
   * Notes:
   * (1) Include this file for the CVBandPrecData type definition.
   * (2) In the CVBandPrecAlloc call, the arguments N is the same
   *     as in the call to CVodeMalloc.
   * (3) In the CVBPSp* call, the user is free to specify
   *     the input pretype and the optional input maxl. The last
   *     argument must be the pointer returned by CVBandPrecAlloc.
   * -----------------------------------------------------------------
   */


  /*
   * -----------------------------------------------------------------
   * Function : CVBandPrecAlloc
   * -----------------------------------------------------------------
   * CVBandPrecAlloc allocates and initializes a CVBandPrecData
   * structure to be passed to CVSp* (and subsequently used
   * by CVBandPrecSetup and CVBandPrecSolve).
   *
   * The parameters of CVBandPrecAlloc are as follows:
   *
   * cvode_mem is the pointer to CVODE memory returned by CVodeCreate.
   *
   * N is the problem size.
   *
   * mu is the upper half bandwidth.
   *
   * ml is the lower half bandwidth.
   *
   * CVBandPrecAlloc returns the storage pointer of type
   * CVBandPrecData, or NULL if the request for storage cannot be
   * satisfied.
   *
   * NOTE: The band preconditioner assumes a serial implementation
   *       of the NVECTOR package. Therefore, CVBandPrecAlloc will
   *       first test for a compatible N_Vector internal
   *       representation by checking for required functions.
   * -----------------------------------------------------------------
   */

  void *CVBandPrecAlloc(void *cvode_mem, long int N,
                        long int mu, long int ml);

  /*
   * -----------------------------------------------------------------
   * Function : CVBPSptfqmr
   * -----------------------------------------------------------------
   * CVBPSptfqmr links the CVBANDPPRE preconditioner to the CVSPTFQMR
   * linear solver. It performs the following actions:
   *  1) Calls the CVSPTFQMR specification routine and attaches the
   *     CVSPTFQMR linear solver to the integrator memory;
   *  2) Sets the preconditioner data structure for CVSPTFQMR
   *  3) Sets the preconditioner setup routine for CVSPTFQMR
   *  4) Sets the preconditioner solve routine for CVSPTFQMR
   *
   * Its first 3 arguments are the same as for CVSptfqmr (see
   * cvsptfqmr.h). The last argument is the pointer to the CVBANDPPRE
   * memory block returned by CVBandPrecAlloc. Note that the user need
   * not call CVSptfqmr.
   *
   * Possible return values are:
   *    CVSPILS_SUCCESS      if successful
   *    CVSPILS_MEM_NULL     if the cvode memory was NULL
   *    CVSPILS_LMEM_NULL    if the cvsptfqmr memory was NULL
   *    CVSPILS_MEM_FAIL     if there was a memory allocation failure
   *    CVSPILS_ILL_INPUT    if a required vector operation is missing
   *    CVBANDPRE_PDATA_NULL if the bp_data was NULL
   * -----------------------------------------------------------------
   */

  int CVBPSptfqmr(void *cvode_mem, int pretype, int maxl, void *p_data);

  /*
   * -----------------------------------------------------------------
   * Function : CVBPSpbcg
   * -----------------------------------------------------------------
   * CVBPSpbcg links the CVBANDPPRE preconditioner to the CVSPBCG
   * linear solver. It performs the following actions:
   *  1) Calls the CVSPBCG specification routine and attaches the
   *     CVSPBCG linear solver to the integrator memory;
   *  2) Sets the preconditioner data structure for CVSPBCG
   *  3) Sets the preconditioner setup routine for CVSPBCG
   *  4) Sets the preconditioner solve routine for CVSPBCG
   *
   * Its first 3 arguments are the same as for CVSpbcg (see
   * cvspbcgs.h). The last argument is the pointer to the CVBANDPPRE
   * memory block returned by CVBandPrecAlloc. Note that the user need
   * not call CVSpbcg.
   *
   * Possible return values are:
   *    CVSPILS_SUCCESS       if successful
   *    CVSPILS_MEM_NULL      if the cvode memory was NULL
   *    CVSPILS_LMEM_NULL     if the cvspbcg memory was NULL
   *    CVSPILS_MEM_FAIL      if there was a memory allocation failure
   *    CVSPILS_ILL_INPUT     if a required vector operation is missing
   *    CVBANDPRE_PDATA_NULL  if the bp_data was NULL
   * -----------------------------------------------------------------
   */

  int CVBPSpbcg(void *cvode_mem, int pretype, int maxl, void *p_data);

  /*
   * -----------------------------------------------------------------
   * Function : CVBPSpgmr
   * -----------------------------------------------------------------
   * CVBPSpgmr links the CVBANDPPRE preconditioner to the CVSPGMR
   * linear solver. It performs the following actions:
   *  1) Calls the CVSPGMR specification routine and attaches the
   *     CVSPGMR linear solver to the integrator memory;
   *  2) Sets the preconditioner data structure for CVSPGMR
   *  3) Sets the preconditioner setup routine for CVSPGMR
   *  4) Sets the preconditioner solve routine for CVSPGMR
   *
   * Its first 3 arguments are the same as for CVSpgmr (see
   * cvspgmr.h). The last argument is the pointer to the CVBANDPPRE
   * memory block returned by CVBandPrecAlloc. Note that the user need
   * not call CVSpgmr.
   *
   * Possible return values are:
   *    CVSPILS_SUCCESS       if successful
   *    CVSPILS_MEM_NULL      if the cvode memory was NULL
   *    CVSPILS_LMEM_NULL     if the cvspgmr memory was NULL
   *    CVSPILS_MEM_FAIL      if there was a memory allocation failure
   *    CVSPILS_ILL_INPUT     if a required vector operation is missing
   *    CVBANDPRE_PDATA_NULL  if the bp_data was NULL
   * -----------------------------------------------------------------
   */

  int CVBPSpgmr(void *cvode_mem, int pretype, int maxl, void *p_data);

  /*
   * -----------------------------------------------------------------
   * Function : CVBandPrecFree
   * -----------------------------------------------------------------
   * CVBandPrecFree frees the memory allocated by CVBandPrecAlloc
   * in the argument bp_data.
   * -----------------------------------------------------------------
   */

  void CVBandPrecFree(void **bp_data);

  /*
   * -----------------------------------------------------------------
   * Optional output functions : CVBandPrecGet*
   * -----------------------------------------------------------------
   * CVBandPrecGetWorkSpace returns the real and integer work space used
   *                        by CVBANDPRE.
   * CVBandPrecGetNumRhsEvals returns the number of calls made from
   *                          CVBANDPRE to the user's right-hand side
   *                          routine f.
   *
   * The return value of CVBandPrecGet* is one of:
   *    CVBANDPRE_SUCCESS    if successful
   *    CVBANDPRE_PDATA_NULL if the bp_data memory was NULL
   * -----------------------------------------------------------------
   */

  int CVBandPrecGetWorkSpace(void *bp_data, long int *lenrwLS, long int *leniwLS);
  int CVBandPrecGetNumRhsEvals(void *bp_data, long int *nfevalsBP);

  /*
   * -----------------------------------------------------------------
   * The following function returns the name of the constant 
   * associated with a CVBANDPRE return flag
   * -----------------------------------------------------------------
   */
  
  char *CVBandPrecGetReturnFlagName(int flag);

  /* 
   * -----------------------------------------------------------------
   * PART II - backward problems
   * -----------------------------------------------------------------
   */

  /*
   * -----------------------------------------------------------------
   * Functions: CVBandPrecAllocB, CVBPSp*B
   * -----------------------------------------------------------------
   * Interface functions for the CVBANDPRE preconditioner to be used
   * on the backward phase.
   *
   * CVBandPrecAllocB interfaces to the CVBANDPRE preconditioner
   * for the backward integration. The pointer to the structure
   * returned by this routine should then be used in the call to
   * CVBPSp*B which interfaces to CVBPSPGMR/CVBPSPBCG/CVSPTFQMR.
   * -----------------------------------------------------------------
   */

  int CVBandPrecAllocB(void *cvadj_mem, long int nB,
                       long int muB, long int mlB);

  int CVBPSptfqmrB(void *cvadj_mem, int pretypeB, int maxlB);
  int CVBPSpbcgB(void *cvadj_mem, int pretypeB, int maxlB);
  int CVBPSpgmrB(void *cvadj_mem, int pretypeB, int maxlB);

  void CVBandPrecFreeB(void *cvadj_mem);

#ifdef __cplusplus
}
#endif

#endif
```
