## Supplementary scripts for "A multi-scale *in silico* mouse model for insulin resistance and humanoid type 2 diabetes": cvodes_bbdpre.h

```
/*
 * -----------------------------------------------------------------
 * $Revision: 749 $
 * $Date: 2011-06-22 14:49:10 +0200 (Wed, 22 Jun 2011) $
 * ----------------------------------------------------------------- 
 * Programmer(s): Radu Serban @ LLNL
 * -----------------------------------------------------------------
 * Copyright (c) 2005, The Regents of the University of California.
 * Produced at the Lawrence Livermore National Laboratory.
 * All rights reserved.
 * For details, see the LICENSE file.
 * -----------------------------------------------------------------
 * This is the header file for the CVBBDPRE module, for a
 * band-block-diagonal preconditioner, i.e. a block-diagonal
 * matrix with banded blocks, for use with CVSPGMR/CVSPBCG/CVSPTFQMR, 
 * and the parallel implementation of the NVECTOR module.
 *
 *
 * Part I contains type definitions and function prototypes for using
 * CVBBDPRE on forward problems (IVP integration and/or FSA)
 *
 * Part II contains type definitions and function prototypes for using
 * CVBBDPRE on adjopint (backward) problems
 * -----------------------------------------------------------------
 */

#ifndef _CVSBBDPRE_H
#define _CVSBBDPRE_H

#ifdef __cplusplus  /* wrapper to enable C++ usage */
extern "C" {
#endif

#include <sundials/sundials_nvector.h>

  /* CVBBDPRE return values */

#define CVBBDPRE_SUCCESS            0
#define CVBBDPRE_PDATA_NULL       -11
#define CVBBDPRE_FUNC_UNRECVR     -12

#define CVBBDPRE_ADJMEM_NULL      -111
#define CVBBDPRE_PDATAB_NULL      -112
#define CVBBDPRE_MEM_FAIL         -113


  /* 
   * -----------------------------------------------------------------
   * PART I - forward problems
   * -----------------------------------------------------------------
   *
   * Summary:
   *
   * These routines provide a preconditioner matrix that is
   * block-diagonal with banded blocks. The blocking corresponds
   * to the distribution of the dependent variable vector y among
   * the processors. Each preconditioner block is generated from
   * the Jacobian of the local part (on the current processor) of a
   * given function g(t,y) approximating f(t,y). The blocks are
   * generated by a difference quotient scheme on each processor
   * independently. This scheme utilizes an assumed banded
   * structure with given half-bandwidths, mudq and mldq.
   * However, the banded Jacobian block kept by the scheme has
   * half-bandwiths mukeep and mlkeep, which may be smaller.
   *
   * The user's calling program should have the following form:
   *
   *   #include <cvodes/cvodes_bbdpre.h>
   *   #include <nvector_parallel.h>
   *   ...
   *   void *cvode_mem;
   *   void *bbd_data;
   *   ...
   *   Set y0
   *   ...
   *   cvode_mem = CVodeCreate(...);
   *   ier = CVodeMalloc(...);
   *   ...
   *   bbd_data = CVBBDPrecAlloc(cvode_mem, Nlocal, mudq ,mldq,
   *                             mukeep, mlkeep, dqrely, gloc, cfn);
   *   flag = CVBBDSpgmr(cvode_mem, pretype, maxl, bbd_data);
   *      -or-
   *   flag = CVBBDSpbcg(cvode_mem, pretype, maxl, bbd_data);
   *      -or-
   *   flag = CVBBDSptfqmr(cvode_mem, pretype, maxl, bbd_data);
   *   ...
   *   ier = CVode(...);
   *   ...
   *   CVBBDPrecFree(&bbd_data);
   *   ...                                                           
   *   CVodeFree(...);
   * 
   *   Free y0
   *
   * The user-supplied routines required are:
   *
   *   f    = function defining the ODE right-hand side f(t,y).
   *
   *   gloc = function defining the approximation g(t,y).
   *
   *   cfn  = function to perform communication need for gloc.
   *
   * Notes:
   *
   * 1) This header file is included by the user for the definition
   *    of the CVBBDData type and for needed function prototypes.
   *
   * 2) The CVBBDPrecAlloc call includes half-bandwiths mudq and mldq
   *    to be used in the difference quotient calculation of the
   *    approximate Jacobian. They need not be the true
   *    half-bandwidths of the Jacobian of the local block of g,
   *    when smaller values may provide a greater efficiency.
   *    Also, the half-bandwidths mukeep and mlkeep of the retained
   *    banded approximate Jacobian block may be even smaller,
   *    to reduce storage and computation costs further.
   *    For all four half-bandwidths, the values need not be the
   *    same on every processor.
   *
   * 3) The actual name of the user's f function is passed to
   *    CVodeMalloc, and the names of the user's gloc and cfn
   *    functions are passed to CVBBDPrecAlloc.
   *
   * 4) The pointer to the user-defined data block f_data, which is
   *    set through CVodeSetFdata is also available to the user in
   *    gloc and cfn.
   *
   * 5) For the CVSpgmr solver, the Gram-Schmidt type gstype,
   *    is left to the user to specify through CVSpgmrSetGStype.
   *
   * 6) Optional outputs specific to this module are available by
   *    way of routines listed below. These include work space sizes
   *    and the cumulative number of gloc calls. The costs
   *    associated with this module also include nsetups banded LU
   *    factorizations, nlinsetups cfn calls, and npsolves banded
   *    backsolve calls, where nlinsetups and npsolves are
   *    integrator/CVSPGMR/CVSPBCG/CVSPTFQMR optional outputs.
   * -----------------------------------------------------------------
   */

  /*
   * -----------------------------------------------------------------
   * Type : CVLocalFn
   * -----------------------------------------------------------------
   * The user must supply a function g(t,y) which approximates the
   * right-hand side function f for the system y'=f(t,y), and which
   * is computed locally (without interprocess communication).
   * (The case where g is mathematically identical to f is allowed.)
   * The implementation of this function must have type CVLocalFn.
   *
   * This function takes as input the local vector size Nlocal, the
   * independent variable value t, the local real dependent
   * variable vector y, and a pointer to the user-defined data
   * block f_data. It is to compute the local part of g(t,y) and
   * store this in the vector g.
   * (Allocation of memory for y and g is handled within the
   * preconditioner module.)
   * The f_data parameter is the same as that specified by the user
   * through the CVodeSetFdata routine.
   *
   * A CVLocalFn should return 0 if successful, a positive value if 
   * a recoverable error occurred, and a negative value if an 
   * unrecoverable error occurred.
   * -----------------------------------------------------------------
   */

  typedef int (*CVLocalFn)(long int Nlocal, realtype t, N_Vector y,
                           N_Vector g, void *f_data);

  /*
   * -----------------------------------------------------------------
   * Type : CVCommFn
   * -----------------------------------------------------------------
   * The user may supply a function of type CVCommFn which performs
   * all interprocess communication necessary to evaluate the
   * approximate right-hand side function described above.
   *
   * This function takes as input the local vector size Nlocal,
   * the independent variable value t, the dependent variable
   * vector y, and a pointer to the user-defined data block f_data.
   * The f_data parameter is the same as that specified by the user
   * through the CVodeSetFdata routine. The CVCommFn cfn is
   * expected to save communicated data in space defined within the
   * structure f_data. Note: A CVCommFn cfn does not have a return value.
   *
   * Each call to the CVCommFn cfn is preceded by a call to the
   * CVRhsFn f with the same (t,y) arguments. Thus cfn can omit any
   * communications done by f if relevant to the evaluation of g.
   * If all necessary communication was done by f, the user can
   * pass NULL for cfn in CVBBDPrecAlloc (see below).
   *
   * A CVCommFn should return 0 if successful, a positive value if 
   * a recoverable error occurred, and a negative value if an 
   * unrecoverable error occurred.
   * -----------------------------------------------------------------
   */

  typedef int (*CVCommFn)(long int Nlocal, realtype t, N_Vector y,
                          void *f_data);

  /*
   * -----------------------------------------------------------------
   * Function : CVBBDPrecAlloc
   * -----------------------------------------------------------------
   * CVBBDPrecAlloc allocates and initializes a CVBBDData structure
   * to be passed to CVSp* (and used by CVBBDPrecSetup and
   * CVBBDPrecSolve).
   *
   * The parameters of CVBBDPrecAlloc are as follows:
   *
   * cvode_mem is the pointer to the integrator memory.
   *
   * Nlocal is the length of the local block of the vectors y etc.
   *        on the current processor.
   *
   * mudq, mldq are the upper and lower half-bandwidths to be used
   *            in the difference quotient computation of the local
   *            Jacobian block.
   *
   * mukeep, mlkeep are the upper and lower half-bandwidths of the
   *                retained banded approximation to the local Jacobian
   *                block.
   *
   * dqrely is an optional input. It is the relative increment
   *        in components of y used in the difference quotient
   *        approximations. To specify the default, pass 0.
   *        The default is dqrely = sqrt(unit roundoff).
   *
   * gloc is the name of the user-supplied function g(t,y) that
   *      approximates f and whose local Jacobian blocks are
   *      to form the preconditioner.
   *
   * cfn is the name of the user-defined function that performs
   *     necessary interprocess communication for the
   *     execution of gloc.
   *
   * CVBBDPrecAlloc returns the storage allocated (type *void),
   * or NULL if the request for storage cannot be satisfied.
   * -----------------------------------------------------------------
   */

  void *CVBBDPrecAlloc(void *cvode_mem, long int Nlocal, 
                       long int mudq, long int mldq, 
                       long int mukeep, long int mlkeep, 
                       realtype dqrely,
                       CVLocalFn gloc, CVCommFn cfn);

  /*
   * -----------------------------------------------------------------
   * Function : CVBBDSptfqmr
   * -----------------------------------------------------------------
   * CVBBDSptfqmr links the CVBBDPRE preconditioner to the CVSPTFQMR
   * linear solver. It performs the following actions:
   *  1) Calls the CVSPTFQMR specification routine and attaches the
   *     CVSPTFQMR linear solver to the integrator memory;
   *  2) Sets the preconditioner data structure for CVSPTFQMR
   *  3) Sets the preconditioner setup routine for CVSPTFQMR
   *  4) Sets the preconditioner solve routine for CVSPTFQMR
   *
   * Its first 3 arguments are the same as for CVSptfqmr (see
   * cvsptfqmr.h). The last argument is the pointer to the CVBBDPRE
   * memory block returned by CVBBDPrecAlloc. Note that the user need
   * not call CVSptfqmr.
   *
   * Possible return values are:
   *    CVSPILS_SUCCESS     if successful
   *    CVSPILS_MEM_NULL    if the cvode memory was NULL
   *    CVSPILS_LMEM_NULL   if the cvspbcg memory was NULL
   *    CVSPILS_MEM_FAIL    if there was a memory allocation failure
   *    CVSPILS_ILL_INPUT   if a required vector operation is missing
   *    CVBBDPRE_PDATA_NULL if the bbd_data was NULL
   * -----------------------------------------------------------------
   */

  int CVBBDSptfqmr(void *cvode_mem, int pretype, int maxl, void *bbd_data);

  /*
   * -----------------------------------------------------------------
   * Function : CVBBDSpbcg
   * -----------------------------------------------------------------
   * CVBBDSpbcg links the CVBBDPRE preconditioner to the CVSPBCG
   * linear solver. It performs the following actions:
   *  1) Calls the CVSPBCG specification routine and attaches the
   *     CVSPBCG linear solver to the integrator memory;
   *  2) Sets the preconditioner data structure for CVSPBCG
   *  3) Sets the preconditioner setup routine for CVSPBCG
   *  4) Sets the preconditioner solve routine for CVSPBCG
   *
   * Its first 3 arguments are the same as for CVSpbcg (see
   * cvspbcgs.h). The last argument is the pointer to the CVBBDPRE
   * memory block returned by CVBBDPrecAlloc. Note that the user need
   * not call CVSpbcg.
   *
   * Possible return values are:
   *    CVSPILS_SUCCESS      if successful
   *    CVSPILS_MEM_NULL     if the cvode memory was NULL
   *    CVSPILS_LMEM_NULL    if the cvspbcg memory was NULL
   *    CVSPILS_MEM_FAIL     if there was a memory allocation failure
   *    CVSPILS_ILL_INPUT    if a required vector operation is missing
   *    CVBBDPRE_PDATA_NULL  if the bbd_data was NULL
   * -----------------------------------------------------------------
   */

  int CVBBDSpbcg(void *cvode_mem, int pretype, int maxl, void *bbd_data);

  /*
   * -----------------------------------------------------------------
   * Function : CVBBDSpgmr
   * -----------------------------------------------------------------
   * CVBBDSpgmr links the CVBBDPRE preconditioner to the CVSPGMR
   * linear solver. It performs the following actions:
   *  1) Calls the CVSPGMR specification routine and attaches the
   *     CVSPGMR linear solver to the integrator memory;
   *  2) Sets the preconditioner data structure for CVSPGMR
   *  3) Sets the preconditioner setup routine for CVSPGMR
   *  4) Sets the preconditioner solve routine for CVSPGMR
   *
   * Its first 3 arguments are the same as for CVSpgmr (see
   * cvspgmr.h). The last argument is the pointer to the CVBBDPRE
   * memory block returned by CVBBDPrecAlloc. Note that the user need
   * not call CVSpgmr.
   *
   * Possible return values are:
   *    CVSPILS_SUCCESS      if successful
   *    CVSPILS_MEM_NULL     if the cvode memory was NULL
   *    CVSPILS_LMEM_NULL    if the cvspgmr memory was NULL
   *    CVSPILS_MEM_FAIL     if there was a memory allocation failure
   *    CVSPILS_ILL_INPUT    if a required vector operation is missing
   *    CVBBDPRE_PDATA_NULL  if the bbd_data was NULL
   * -----------------------------------------------------------------
   */

  int CVBBDSpgmr(void *cvode_mem, int pretype, int maxl, void *bbd_data);

  /*
   * -----------------------------------------------------------------
   * Function : CVBBDPrecReInit
   * -----------------------------------------------------------------
   * CVBBDPrecReInit re-initializes the BBDPRE module when solving a
   * sequence of problems of the same size with CVSPGMR/CVBBDPRE,
   * CVSPBCG/CVBBDPRE, or CVSPTFQMR/CVBBDPRE provided there is no change 
   * in Nlocal, mukeep, or mlkeep. After solving one problem, and after 
   * calling CVodeReInit to re-initialize the integrator for a subsequent 
   * problem, call CVBBDPrecReInit. Then call CVSpgmrSet*, CVSpbcgSet*,
   * or CVSptfqmrSet*  functions if necessary for any changes to CVSPGMR,
   * CVSPBCG, or CVSPTFQMR parameters, before calling CVode.
   *
   * The first argument to CVBBDPrecReInit must be the pointer pdata
   * that was returned by CVBBDPrecAlloc. All other arguments have
   * the same names and meanings as those of CVBBDPrecAlloc.
   *
   * The return value of CVBBDPrecReInit is CVBBDPRE_SUCCESS, indicating
   * success, or CVBBDPRE_PDATA_NULL if bbd_data was NULL.
   * -----------------------------------------------------------------
   */

  int CVBBDPrecReInit(void *bbd_data, long int mudq, long int mldq,
                      realtype dqrely, CVLocalFn gloc, CVCommFn cfn);

  /*
   * -----------------------------------------------------------------
   * Function : CVBBDPrecFree
   * -----------------------------------------------------------------
   * CVBBDPrecFree frees the memory block bbd_data allocated by the
   * call to CVBBDAlloc.
   * -----------------------------------------------------------------
   */

  void CVBBDPrecFree(void **bbd_data);

  /*
   * -----------------------------------------------------------------
   * BBDPRE optional output extraction routines
   * -----------------------------------------------------------------
   * CVBBDPrecGetWorkSpace returns the BBDPRE real and integer work space
   *                       sizes.
   * CVBBDPrecGetNumGfnEvals returns the number of calls to gfn.
   *
   * The return value of CVBBDPrecGet* is one of:
   *    CVBBDPRE_SUCCESS    if successful
   *    CVBBDPRE_PDATA_NULL if the bbd_data memory was NULL
   * -----------------------------------------------------------------
   */

  int CVBBDPrecGetWorkSpace(void *bbd_data, long int *lenrwBBDP, 
                            long int *leniwBBDP);
  int CVBBDPrecGetNumGfnEvals(void *bbd_data, long int *ngevalsBBDP);

  /*
   * -----------------------------------------------------------------
   * The following function returns the name of the constant 
   * associated with a CVBBDPRE return flag
   * -----------------------------------------------------------------
   */
  
  char *CVBBDPrecGetReturnFlagName(int flag);

  /* 
   * -----------------------------------------------------------------
   * PART II - backward problems
   * -----------------------------------------------------------------
   */

  /*
   * -----------------------------------------------------------------
   * Types: CVLocalFnB and CVCommFnB
   * -----------------------------------------------------------------
   * Local approximation function and inter-process communication
   * function for the BBD preconditioner on the backward phase.
   * -----------------------------------------------------------------
   */

  typedef int (*CVLocalFnB)(long int NlocalB, realtype t,
                            N_Vector y, N_Vector yB, N_Vector gB,
                            void *f_dataB);
  
  typedef int (*CVCommFnB)(long int NlocalB, realtype t,
                           N_Vector y, N_Vector yB,
                           void *f_dataB);

  /*
   * -----------------------------------------------------------------
   * Functions: CVBBDPrecAllocB, CVBBDSp*B, CVBBDPrecReInit
   * -----------------------------------------------------------------
   * Interface functions for the CVBBDPRE preconditioner to be used on
   * the backward phase.
   * -----------------------------------------------------------------
   */

  int CVBBDPrecAllocB(void *cvadj_mem, long int NlocalB,
                      long int mudqB, long int mldqB,
                      long int mukeepB, long int mlkeepB,
                      realtype dqrelyB,
                      CVLocalFnB glocB, CVCommFnB cfnB);

  int CVBBDSptfqmrB(void *cvadj_mem, int pretypeB, int maxlB);
  int CVBBDSpbcgB(void *cvadj_mem, int pretypeB, int maxlB);
  int CVBBDSpgmrB(void *cvadj_mem, int pretypeB, int maxlB);
  
  int CVBBDPrecReInitB(void *cvadj_mem, long int mudqB, long int mldqB,
                       realtype dqrelyB, CVLocalFnB glocB, CVCommFnB cfnB);

  void CVBBDPrecFreeB(void *cvadj_mem);

#ifdef __cplusplus
}
#endif

#endif
```
