## Supplementary scripts for "A multi-scale *in silico* mouse model for insulin resistance and humanoid type 2 diabetes": cvodes_dense.h

```
/*
 * -----------------------------------------------------------------
 * $Revision: 749 $
 * $Date: 2011-06-22 14:49:10 +0200 (Wed, 22 Jun 2011) $
 * ----------------------------------------------------------------- 
 * Programmer(s): Radu Serban @ LLNL
 * -----------------------------------------------------------------
 * Copyright (c) 2005, The Regents of the University of California.
 * Produced at the Lawrence Livermore National Laboratory.
 * All rights reserved.
 * For details, see the LICENSE file.
 * -----------------------------------------------------------------
 * This is the header file for the CVODES dense linear solver, CVDENSE.
 *
 *
 * Part I contains type definitions and function prototypes for using
 * CVDENSE on forward problems (IVP integration and/or FSA)
 *
 * Part II contains type definitions and function prototypes for using
 * CVDENSE on adjoint (backward) problems
 * -----------------------------------------------------------------
 */

#ifndef _CVSDENSE_H
#define _CVSDENSE_H

#ifdef __cplusplus  /* wrapper to enable C++ usage */
extern "C" {
#endif

#include <sundials/sundials_dense.h>
#include <sundials/sundials_nvector.h>

  /*
   * -----------------------------------------------------------------
   * CVDENSE solver constants
   * -----------------------------------------------------------------
   * CVD_MSBJ : maximum number of steps between dense Jacobian
   *            evaluations
   *
   * CVD_DGMAX : maximum change in gamma between dense Jacobian
   *             evaluations
   * -----------------------------------------------------------------
   */

#define CVD_MSBJ  50
#define CVD_DGMAX RCONST(0.2)

  /*
   * -----------------------------------------------------------------
   * CVDENSE return values
   * -----------------------------------------------------------------
   */

#define CVDENSE_SUCCESS           0
#define CVDENSE_MEM_NULL         -1
#define CVDENSE_LMEM_NULL        -2
#define CVDENSE_ILL_INPUT        -3
#define CVDENSE_MEM_FAIL         -4

  /* Additional last_flag values */

#define CVDENSE_JACFUNC_UNRECVR  -5
#define CVDENSE_JACFUNC_RECVR    -6

  /* Return values for the adjoint module */

#define CVDENSE_ADJMEM_NULL      -101
#define CVDENSE_LMEMB_NULL       -102

  /* 
   * -----------------------------------------------------------------
   * PART I - forward problems
   * -----------------------------------------------------------------
   */

  /*
   * -----------------------------------------------------------------
   * Type : CVDenseJacFn
   * -----------------------------------------------------------------
   * A dense Jacobian approximation function Jac must have the
   * prototype given below. Its parameters are:
   *
   * N is the problem size.
   *
   * J is the dense matrix (of type DenseMat) that will be loaded
   * by a CVDenseJacFn with an approximation to the Jacobian matrix
   * J = (df_i/dy_j) at the point (t,y).
   * J is preset to zero, so only the nonzero elements need to be
   * loaded. Two efficient ways to load J are:
   *
   * (1) (with macros - no explicit data structure references)
   *     for (j=0; j < n; j++) {
   *       col_j = DENSE_COL(J,j);
   *       for (i=0; i < n; i++) {
   *         generate J_ij = the (i,j)th Jacobian element
   *         col_j[i] = J_ij;
   *       }
   *     }
   *
   * (2) (without macros - explicit data structure references)
   *     for (j=0; j < n; j++) {
   *       col_j = (J->data)[j];
   *       for (i=0; i < n; i++) {
   *         generate J_ij = the (i,j)th Jacobian element
   *         col_j[i] = J_ij;
   *       }
   *     }
   *
   * The DENSE_ELEM(A,i,j) macro is appropriate for use in small
   * problems in which efficiency of access is NOT a major concern.
   *
   * t is the current value of the independent variable.
   *
   * y is the current value of the dependent variable vector,
   *   namely the predicted value of y(t).
   *
   * fy is the vector f(t,y).
   *
   * jac_data is a pointer to user data - the same as the jac_data
   *          parameter passed to CVDense.
   *
   * NOTE: If the user's Jacobian routine needs other quantities,
   *       they are accessible as follows: hcur (the current stepsize)
   *       and ewt (the error weight vector) are accessible through
   *       CVodeGetCurrentStep and CVodeGetErrWeights, respectively
   *       (see cvode.h). The unit roundoff is available as
   *       UNIT_ROUNDOFF defined in sundials_types.h.
   *
   * tmp1, tmp2, and tmp3 are pointers to memory allocated for
   * vectors of length N which can be used by a CVDenseJacFn
   * as temporary storage or work space.
   *
   * A CVDenseJacFn should return 0 if successful, a positive value if 
   * a recoverable error occurred, and a negative value if an 
   * unrecoverable error occurred.
   * -----------------------------------------------------------------
   */

  typedef int (*CVDenseJacFn)(long int N, DenseMat J, realtype t,
                              N_Vector y, N_Vector fy, void *jac_data,
                              N_Vector tmp1, N_Vector tmp2, N_Vector tmp3);

  /*
   * -----------------------------------------------------------------
   * Function : CVDense
   * -----------------------------------------------------------------
   * A call to the CVDense function links the main integrator with
   * the CVDENSE linear solver.
   *
   * cvode_mem is the pointer to the integrator memory returned by
   *           CVodeCreate.
   *
   * N is the size of the ODE system.
   *
   * The return value of CVDense is one of:
   *    CVDENSE_SUCCESS   if successful
   *    CVDENSE_MEM_NULL  if the cvode memory was NULL
   *    CVDENSE_MEM_FAIL  if there was a memory allocation failure
   *    CVDENSE_ILL_INPUT if a required vector operation is missing
   * -----------------------------------------------------------------
   */

  int CVDense(void *cvode_mem, long int N);

  /*
   * -----------------------------------------------------------------
   * Optional inputs to the CVDENSE linear solver
   * -----------------------------------------------------------------
   *
   * CVDenseSetJacFn specifies the dense Jacobian approximation
   *                 routine to be used. A user-supplied djac routine
   *                 must be of type CVDenseJacFn. By default, a
   *                 difference quotient routine CVDenseDQJac, supplied
   *                 with this solver is used.                     
   *                 It also specifies a pointer to user data which is
   *                 passed to the djac routine every time it is called.
   *
   * The return value of CVDenseSet* is one of:
   *    CVDENSE_SUCCESS   if successful
   *    CVDENSE_MEM_NULL  if the cvode memory was NULL
   *    CVDENSE_LMEM_NULL if the cvdense memory was NULL
   * -----------------------------------------------------------------
   */

  int CVDenseSetJacFn(void *cvode_mem, CVDenseJacFn djac, void *jac_data);

  /*
   * -----------------------------------------------------------------
   * Optional outputs from the CVDENSE linear solver
   * -----------------------------------------------------------------
   *
   * CVDenseGetWorkSpace returns the real and integer workspace used
   *                     by CVDENSE.
   * CVDenseGetNumJacEvals returns the number of calls made to the
   *                       Jacobian evaluation routine djac.
   * CVDenseGetNumRhsEvals returns the number of calls to the user
   *                       f routine due to finite difference Jacobian
   *                       evaluation.
   * CVDenseGetLastFlag returns the last error flag set by any of
   *                    the CVDENSE interface functions.
   *
   * The return value of CVDenseGet* is one of:
   *    CVDENSE_SUCCESS   if successful
   *    CVDENSE_MEM_NULL  if the cvode memory was NULL
   *    CVDENSE_LMEM_NULL if the cvdense memory was NULL
   * -----------------------------------------------------------------
   */

  int CVDenseGetWorkSpace(void *cvode_mem, long int *lenrwLS, long int *leniwLS);
  int CVDenseGetNumJacEvals(void *cvode_mem, long int *njevals);
  int CVDenseGetNumRhsEvals(void *cvode_mem, long int *nfevalsLS);
  int CVDenseGetLastFlag(void *cvode_mem, int *flag);

  /*
   * -----------------------------------------------------------------
   * The following function returns the name of the constant 
   * associated with a CVDENSE return flag
   * -----------------------------------------------------------------
   */

  char *CVDenseGetReturnFlagName(int flag);

  /* 
   * -----------------------------------------------------------------
   * PART II - backward problems
   * -----------------------------------------------------------------
   */

  /*
   * -----------------------------------------------------------------
   * Type: CVDenseJacFnB
   * -----------------------------------------------------------------
   * A dense Jacobian approximation function JacB for the adjoint
   * (backward) problem must have the prototype given below. 
   * -----------------------------------------------------------------
   */

  typedef int (*CVDenseJacFnB)(long int nB, DenseMat JB, realtype t,
                               N_Vector y, N_Vector yB, N_Vector fyB,
                               void *jac_dataB, N_Vector tmp1B,
                               N_Vector tmp2B, N_Vector tmp3B);

  /*
   * -----------------------------------------------------------------
   * Functions: CVDenseB, CVDenseSetJacFnB
   * -----------------------------------------------------------------
   * CVDenseB links the main CVODE integrator with the CVDENSE
   * linear solver for the backward integration.
   * -----------------------------------------------------------------
   */

  int CVDenseB(void *cvadj_mem, long int nB);
  
  int CVDenseSetJacFnB(void *cvadj_mem, CVDenseJacFnB djacB, void *jac_dataB);

#ifdef __cplusplus
}
#endif

#endif
```
