## Supplementary scripts for "A multi-scale *in silico* mouse model for insulin resistance and humanoid type 2 diabetes": cvodes_diag.h

```
/*
 * -----------------------------------------------------------------
 * $Revision: 749 $
 * $Date: 2011-06-22 14:49:10 +0200 (Wed, 22 Jun 2011) $
 * ----------------------------------------------------------------- 
 * Programmer(s): Radu Serban @ LLNL
 * -----------------------------------------------------------------
 * Copyright (c) 2005, The Regents of the University of California.
 * Produced at the Lawrence Livermore National Laboratory.
 * All rights reserved.
 * For details, see the LICENSE file.
 * -----------------------------------------------------------------
 * This is the header file for the CVODES diagonal linear solver, 
 * CVDIAG.
 *
 *
 * Part I contains type definitions and function prototypes for using
 * CVDIAG on forward problems (IVP integration and/or FSA)
 *
 * Part II contains type definitions and function prototypes for using
 * CVDIAG on adjoint (backward) problems
 * -----------------------------------------------------------------
 */

#ifndef _CVSDIAG_H
#define _CVSDIAG_H

#ifdef __cplusplus  /* wrapper to enable C++ usage */
extern "C" {
#endif

#include <sundials/sundials_nvector.h>

  /*
   * -----------------------------------------------------------------
   * CVDIAG return values
   * -----------------------------------------------------------------
   */

#define CVDIAG_SUCCESS          0
#define CVDIAG_MEM_NULL        -1
#define CVDIAG_LMEM_NULL       -2
#define CVDIAG_ILL_INPUT       -3
#define CVDIAG_MEM_FAIL        -4

  /* Additional last_flag values */

#define CVDIAG_INV_FAIL        -5
#define CVDIAG_RHSFUNC_UNRECVR -6
#define CVDIAG_RHSFUNC_RECVR   -7

  /* Return values for adjoint module */

#define CVDIAG_ADJMEM_NULL     -101

  /* 
   * -----------------------------------------------------------------
   * PART I - forward problems
   * -----------------------------------------------------------------
   */

  /*
   * -----------------------------------------------------------------
   * Function : CVDiag
   * -----------------------------------------------------------------
   * A call to the CVDiag function links the main integrator with
   * the CVDIAG linear solver.
   *
   * cvode_mem is the pointer to the integrator memory returned by
   *           CVodeCreate.
   *
   * The return value of CVDiag is one of:
   *    CVDIAG_SUCCESS   if successful
   *    CVDIAG_MEM_NULL  if the cvode memory was NULL
   *    CVDIAG_MEM_FAIL  if there was a memory allocation failure
   *    CVDIAG_ILL_INPUT if a required vector operation is missing
   * -----------------------------------------------------------------
   */

  int CVDiag(void *cvode_mem);
  
  /*
   * -----------------------------------------------------------------
   * Optional outputs from the CVDIAG linear solver
   * -----------------------------------------------------------------
   *
   * CVDiagGetWorkSpace returns the real and integer workspace used
   *                    by CVDIAG.
   * CVDiagGetNumRhsEvals returns the number of calls to the user
   *                      f routine due to finite difference Jacobian
   *                      evaluation.
   *                      Note: The number of diagonal approximate
   *                      Jacobians formed is equal to the number of
   *                      CVDiagSetup calls. This number is available
   *                      through CVodeGetNumLinSolvSetups.
   * CVDiagGetLastFlag returns the last error flag set by any of
   *                   the CVDIAG interface functions.
   *
   * The return value of CVDiagGet* is one of:
   *    CVDIAG_SUCCESS   if successful
   *    CVDIAG_MEM_NULL  if the cvode memory was NULL
   *    CVDIAG_LMEM_NULL if the cvdiag memory was NULL
   * -----------------------------------------------------------------
   */

  int CVDiagGetWorkSpace(void *cvode_mem, long int *lenrwLS, long int *leniwLS);
  int CVDiagGetNumRhsEvals(void *cvode_mem, long int *nfevalsLS);
  int CVDiagGetLastFlag(void *cvode_mem, int *flag);

  /*
   * -----------------------------------------------------------------
   * The following function returns the name of the constant 
   * associated with a CVDIAG return flag
   * -----------------------------------------------------------------
   */
  
  char *CVDiagGetReturnFlagName(int flag);

  /* 
   * -----------------------------------------------------------------
   * PART II - backward problems
   * -----------------------------------------------------------------
   */

  /*
   * -----------------------------------------------------------------
   * Function: CVDiagB
   * -----------------------------------------------------------------
   * CVDiagB links the main CVODE integrator with the CVDIAG
   * linear solver for the backward integration.
   * -----------------------------------------------------------------
   */
  
  int CVDiagB(void *cvadj_mem);
  

#ifdef __cplusplus
}
#endif

#endif
```
