## Supplementary scripts for "A multi-scale *in silico* mouse model for insulin resistance and humanoid type 2 diabetes": cvodes_spils.h

```
/*
 * -----------------------------------------------------------------
 * $Revision: 749 $
 * $Date: 2011-06-22 14:49:10 +0200 (Wed, 22 Jun 2011) $
 * ----------------------------------------------------------------- 
 * Programmer(s): Radu Serban @ LLNL
 * -----------------------------------------------------------------
 * Copyright (c) 2005, The Regents of the University of California.
 * Produced at the Lawrence Livermore National Laboratory.
 * All rights reserved.
 * For details, see the LICENSE file.
 * -----------------------------------------------------------------
 * This is the common header file for the Scaled, Preconditioned
 * Iterative Linear Solvers in CVODE/CVODES.
 *
 * Part I contains type definitions and functions for using the 
 * iterative linear solvers on forward problems 
 * (IVP integration and/or FSA)
 *
 * Part II contains type definitions and functions for using the 
 * iterative linear solvers on adjoint (backward) problems
 * -----------------------------------------------------------------
 */

#ifndef _CVSSPILS_H
#define _CVSSPILS_H

#ifdef __cplusplus  /* wrapper to enable C++ usage */
extern "C" {
#endif

#include <sundials/sundials_iterative.h>
#include <sundials/sundials_nvector.h>

  /*
   * -----------------------------------------------------------------
   * CVSPILS return values
   * -----------------------------------------------------------------
   */

#define CVSPILS_SUCCESS          0
#define CVSPILS_MEM_NULL        -1
#define CVSPILS_LMEM_NULL       -2
#define CVSPILS_ILL_INPUT       -3
#define CVSPILS_MEM_FAIL        -4

  /* Return values for the adjoint module */

#define CVSPILS_ADJMEM_NULL     -101
#define CVSPILS_LMEMB_NULL      -102

  /*
   * -----------------------------------------------------------------
   * CVSPILS solver constants
   * -----------------------------------------------------------------
   * CVSPILS_MAXL   : default value for the maximum Krylov
   *                  dimension
   *
   * CVSPILS_MSBPRE : maximum number of steps between
   *                  preconditioner evaluations
   *
   * CVSPILS_DGMAX  : maximum change in gamma between
   *                  preconditioner evaluations
   *
   * CVSPILS_DELT   : default value for factor by which the
   *                  tolerance on the nonlinear iteration is
   *                  multiplied to get a tolerance on the linear
   *                  iteration
   * -----------------------------------------------------------------
   */

#define CVSPILS_MAXL   5
#define CVSPILS_MSBPRE 50
#define CVSPILS_DGMAX  RCONST(0.2)
#define CVSPILS_DELT   RCONST(0.05)

  /* 
   * -----------------------------------------------------------------
   * PART I - forward problems
   * -----------------------------------------------------------------
   */

  /*
   * -----------------------------------------------------------------
   * Type : CVSpilsPrecSetupFn
   * -----------------------------------------------------------------
   * The user-supplied preconditioner setup function PrecSetup and
   * the user-supplied preconditioner solve function PrecSolve
   * together must define left and right preconditoner matrices
   * P1 and P2 (either of which may be trivial), such that the
   * product P1*P2 is an approximation to the Newton matrix
   * M = I - gamma*J.  Here J is the system Jacobian J = df/dy,
   * and gamma is a scalar proportional to the integration step
   * size h.  The solution of systems P z = r, with P = P1 or P2,
   * is to be carried out by the PrecSolve function, and PrecSetup
   * is to do any necessary setup operations.
   *
   * The user-supplied preconditioner setup function PrecSetup
   * is to evaluate and preprocess any Jacobian-related data
   * needed by the preconditioner solve function PrecSolve.
   * This might include forming a crude approximate Jacobian,
   * and performing an LU factorization on the resulting
   * approximation to M.  This function will not be called in
   * advance of every call to PrecSolve, but instead will be called
   * only as often as necessary to achieve convergence within the
   * Newton iteration.  If the PrecSolve function needs no
   * preparation, the PrecSetup function can be NULL.
   *
   * For greater efficiency, the PrecSetup function may save
   * Jacobian-related data and reuse it, rather than generating it
   * from scratch.  In this case, it should use the input flag jok
   * to decide whether to recompute the data, and set the output
   * flag *jcurPtr accordingly.
   *
   * Each call to the PrecSetup function is preceded by a call to
   * the RhsFn f with the same (t,y) arguments.  Thus the PrecSetup
   * function can use any auxiliary data that is computed and
   * saved by the f function and made accessible to PrecSetup.
   *
   * A function PrecSetup must have the prototype given below.
   * Its parameters are as follows:
   *
   * t       is the current value of the independent variable.
   *
   * y       is the current value of the dependent variable vector,
   *          namely the predicted value of y(t).
   *
   * fy      is the vector f(t,y).
   *
   * jok     is an input flag indicating whether Jacobian-related
   *         data needs to be recomputed, as follows:
   *           jok == FALSE means recompute Jacobian-related data
   *                  from scratch.
   *           jok == TRUE  means that Jacobian data, if saved from
   *                  the previous PrecSetup call, can be reused
   *                  (with the current value of gamma).
   *         A Precset call with jok == TRUE can only occur after
   *         a call with jok == FALSE.
   *
   * jcurPtr is a pointer to an output integer flag which is
   *         to be set by PrecSetup as follows:
   *         Set *jcurPtr = TRUE if Jacobian data was recomputed.
   *         Set *jcurPtr = FALSE if Jacobian data was not recomputed,
   *                        but saved data was reused.
   *
   * gamma   is the scalar appearing in the Newton matrix.
   *
   * P_data  is a pointer to user data - the same as the P_data
   *         parameter passed to the CV*SetPreconditioner function.
   *
   * tmp1, tmp2, and tmp3 are pointers to memory allocated
   *                      for N_Vectors which can be used by
   *                      CVSpilsPrecSetupFn as temporary storage or
   *                      work space.
   *
   * NOTE: If the user's preconditioner needs other quantities,
   *       they are accessible as follows: hcur (the current stepsize)
   *       and ewt (the error weight vector) are accessible through
   *       CVodeGetCurrentStep and CVodeGetErrWeights, respectively).
   *       The unit roundoff is available as UNIT_ROUNDOFF defined in
   *       sundials_types.h.
   *
   * Returned value:
   * The value to be returned by the PrecSetup function is a flag
   * indicating whether it was successful.  This value should be
   *   0   if successful,
   *   > 0 for a recoverable error (step will be retried),
   *   < 0 for an unrecoverable error (integration is halted).
   * -----------------------------------------------------------------
   */

  typedef int (*CVSpilsPrecSetupFn)(realtype t, N_Vector y, N_Vector fy,
                                    booleantype jok, booleantype *jcurPtr,
                                    realtype gamma, void *P_data,
                                    N_Vector tmp1, N_Vector tmp2,
                                    N_Vector tmp3);

  /*
   * -----------------------------------------------------------------
   * Type : CVSpilsPrecSolveFn
   * -----------------------------------------------------------------
   * The user-supplied preconditioner solve function PrecSolve
   * is to solve a linear system P z = r in which the matrix P is
   * one of the preconditioner matrices P1 or P2, depending on the
   * type of preconditioning chosen.
   *
   * A function PrecSolve must have the prototype given below.
   * Its parameters are as follows:
   *
   * t      is the current value of the independent variable.
   *
   * y      is the current value of the dependent variable vector.
   *
   * fy     is the vector f(t,y).
   *
   * r      is the right-hand side vector of the linear system.
   *
   * z      is the output vector computed by PrecSolve.
   *
   * gamma  is the scalar appearing in the Newton matrix.
   *
   * delta  is an input tolerance for use by PSolve if it uses
   *        an iterative method in its solution.  In that case,
   *        the residual vector Res = r - P z of the system
   *        should be made less than delta in weighted L2 norm,
   *        i.e., sqrt [ Sum (Res[i]*ewt[i])^2 ] < delta.
   *        Note: the error weight vector ewt can be obtained
   *        through a call to the routine CVodeGetErrWeights.
   *
   * lr     is an input flag indicating whether PrecSolve is to use
   *        the left preconditioner P1 or right preconditioner
   *        P2: lr = 1 means use P1, and lr = 2 means use P2.
   *
   * P_data is a pointer to user data - the same as the P_data
   *        parameter passed to the CV*SetPreconditioner function.
   *
   * tmp    is a pointer to memory allocated for an N_Vector
   *        which can be used by PSolve for work space.
   *
   * Returned value:
   * The value to be returned by the PrecSolve function is a flag
   * indicating whether it was successful.  This value should be
   *   0 if successful,
   *   positive for a recoverable error (step will be retried),
   *   negative for an unrecoverable error (integration is halted).
   * -----------------------------------------------------------------
   */

  typedef int (*CVSpilsPrecSolveFn)(realtype t, N_Vector y, N_Vector fy,
                                    N_Vector r, N_Vector z,
                                    realtype gamma, realtype delta,
                                    int lr, void *P_data, N_Vector tmp);

  /*
   * -----------------------------------------------------------------
   * Type : CVSpilsJacTimesVecFn
   * -----------------------------------------------------------------
   * The user-supplied function jtimes is to generate the product
   * J*v for given v, where J is the Jacobian df/dy, or an
   * approximation to it, and v is a given vector. It should return
   * 0 if successful a positive value for a recoverable error or 
   * a negative value for an unrecoverable failure.
   *
   * A function jtimes must have the prototype given below. Its
   * parameters are as follows:
   *
   *   v        is the N_Vector to be multiplied by J.
   *
   *   Jv       is the output N_Vector containing J*v.
   *
   *   t        is the current value of the independent variable.
   *
   *   y        is the current value of the dependent variable
   *            vector.
   *
   *   fy       is the vector f(t,y).
   *
   *   jac_data is a pointer to user Jacobian data, the same as the
   *            jac_data parameter passed to the CV*SetJacTimesVecFn 
   *            function.
   *
   *   tmp      is a pointer to memory allocated for an N_Vector
   *            which can be used by Jtimes for work space.
   * -----------------------------------------------------------------
   */

  typedef int (*CVSpilsJacTimesVecFn)(N_Vector v, N_Vector Jv, realtype t,
                                      N_Vector y, N_Vector fy,
                                      void *jac_data, N_Vector tmp);


  /*
   * -----------------------------------------------------------------
   * Optional inputs to the CVSPILS linear solver
   * -----------------------------------------------------------------
   *
   * CVSpilsSetPrecType resets the type of preconditioner, pretype,
   *                from the value previously set.
   *                This must be one of PREC_NONE, PREC_LEFT, 
   *                PREC_RIGHT, or PREC_BOTH.
   *
   * CVSpilsSetGSType specifies the type of Gram-Schmidt
   *                orthogonalization to be used. This must be one of
   *                the two enumeration constants MODIFIED_GS or
   *                CLASSICAL_GS defined in iterative.h. These correspond
   *                to using modified Gram-Schmidt and classical
   *                Gram-Schmidt, respectively.
   *                Default value is MODIFIED_GS.
   *
   * CVSpilsSetMaxl resets the maximum Krylov subspace size, maxl,
   *                from the value previously set.
   *                An input value <= 0, gives the default value.
   *
   * CVSpilsSetDelt specifies the factor by which the tolerance on
   *                the nonlinear iteration is multiplied to get a
   *                tolerance on the linear iteration.
   *                Default value is 0.05.
   *
   * CVSpilsSetPreconditioner specifies the PrecSetup and PrecSolve functions.
   *                as well as a pointer to user preconditioner data.
   *                This pointer is passed to PrecSetup and PrecSolve
   *                every time these routines are called.
   *                Default is NULL for al three arguments.
   *
   * CVSpilsSetJacTimesVecFn specifies the jtimes function and a pointer to
   *                user Jacobian data. This pointer is passed to jtimes every 
   *                time the jtimes routine is called.
   *                Default is to use an internal finite difference
   *                approximation routine.
   *
   * The return value of CVSpilsSet* is one of:
   *    CVSPILS_SUCCESS   if successful
   *    CVSPILS_MEM_NULL  if the cvode memory was NULL
   *    CVSPILS_LMEM_NULL if the cvspgmr memory was NULL
   *    CVSPILS_ILL_INPUT if an input has an illegal value
   * -----------------------------------------------------------------
   */

  int CVSpilsSetPrecType(void *cvode_mem, int pretype);
  int CVSpilsSetGSType(void *cvode_mem, int gstype);
  int CVSpilsSetMaxl(void *cvode_mem, int maxl);
  int CVSpilsSetDelt(void *cvode_mem, realtype delt);
  int CVSpilsSetPreconditioner(void *cvode_mem, CVSpilsPrecSetupFn pset, 
                               CVSpilsPrecSolveFn psolve, void *P_data);
  int CVSpilsSetJacTimesVecFn(void *cvode_mem, 
                              CVSpilsJacTimesVecFn jtimes, void *jac_data);

  /*
   * -----------------------------------------------------------------
   * Optional outputs from the CVSPILS linear solver
   * -----------------------------------------------------------------
   * CVSpilsGetWorkSpace returns the real and integer workspace used
   *                by the SPILS module.
   *
   * CVSpilsGetNumPrecEvals returns the number of preconditioner
   *                 evaluations, i.e. the number of calls made
   *                 to PrecSetup with jok==FALSE.
   *
   * CVSpilsGetNumPrecSolves returns the number of calls made to
   *                 PrecSolve.
   *
   * CVSpilsGetNumLinIters returns the number of linear iterations.
   *
   * CVSpilsGetNumConvFails returns the number of linear
   *                 convergence failures.
   *
   * CVSpilsGetNumJtimesEvals returns the number of calls to jtimes.
   *
   * CVSpilsGetNumRhsEvals returns the number of calls to the user
   *                 f routine due to finite difference Jacobian
   *                 times vector evaluation.
   *
   * CVSpilsGetLastFlag returns the last error flag set by any of
   *                 the CVSPILS interface functions.
   *
   * The return value of CVSpilsGet* is one of:
   *    CVSPILS_SUCCESS   if successful
   *    CVSPILS_MEM_NULL  if the cvode memory was NULL
   *    CVSPILS_LMEM_NULL if the cvspgmr memory was NULL
   * -----------------------------------------------------------------
   */

  int CVSpilsGetWorkSpace(void *cvode_mem, long int *lenrwLS, long int *leniwLS);
  int CVSpilsGetNumPrecEvals(void *cvode_mem, long int *npevals);
  int CVSpilsGetNumPrecSolves(void *cvode_mem, long int *npsolves);
  int CVSpilsGetNumLinIters(void *cvode_mem, long int *nliters);
  int CVSpilsGetNumConvFails(void *cvode_mem, long int *nlcfails);
  int CVSpilsGetNumJtimesEvals(void *cvode_mem, long int *njvevals);
  int CVSpilsGetNumRhsEvals(void *cvode_mem, long int *nfevalsLS); 
  int CVSpilsGetLastFlag(void *cvode_mem, int *flag);

  /*
   * -----------------------------------------------------------------
   * The following function returns the name of the constant 
   * associated with a CVSPILS return flag
   * -----------------------------------------------------------------
   */
  
  char *CVSpilsGetReturnFlagName(int flag);


  /* 
   * -----------------------------------------------------------------
   * PART II - backward problems
   * -----------------------------------------------------------------
   */

  /*
   * -----------------------------------------------------------------
   * Type : CVSpilsPrecSetupFnB
   * -----------------------------------------------------------------
   * A function PrecSetupB for the adjoint (backward) problem must have 
   * the prototype given below.
   * -----------------------------------------------------------------
   */

  typedef int (*CVSpilsPrecSetupFnB)(realtype t, N_Vector y,
                                     N_Vector yB, N_Vector fyB,
                                     booleantype jokB,
                                     booleantype *jcurPtrB, realtype gammaB,
                                     void *P_dataB,
                                     N_Vector tmp1B, N_Vector tmp2B,
                                     N_Vector tmp3B);


  /*
   * -----------------------------------------------------------------
   * Type : CVSpilsPrecSolveFnB
   * -----------------------------------------------------------------
   * A function PrecSolveB for the adjoint (backward) problem  must 
   * have the prototype given below.
   * -----------------------------------------------------------------
   */

  typedef int (*CVSpilsPrecSolveFnB)(realtype t, N_Vector y,
                                     N_Vector yB, N_Vector fyB,
                                     N_Vector rB, N_Vector zB,
                                     realtype gammaB, realtype deltaB,
                                     int lrB, void *P_dataB, N_Vector tmpB);

  /*
   * -----------------------------------------------------------------
   * Type : CVSpilsJacTimesVecFnB
   * -----------------------------------------------------------------
   * A function jtimesB for the adjoint (backward) problem must have 
   * the prototype given below.
   * -----------------------------------------------------------------
   */

  typedef int (*CVSpilsJacTimesVecFnB)(N_Vector vB, N_Vector JvB, realtype t,
                                       N_Vector y, N_Vector yB, N_Vector fyB,
                                       void *jac_dataB, N_Vector tmpB);

  /*
   * -----------------------------------------------------------------
   * Functions
   * -----------------------------------------------------------------
   */

  int CVSpilsSetPrecTypeB(void *cvadj_mem, int pretypeB);
  int CVSpilsSetGSTypeB(void *cvadj_mem, int gstypeB);
  int CVSpilsSetDeltB(void *cvadj_mem, realtype deltB);
  int CVSpilsSetMaxlB(void *cvadj_mem, int maxlB);
  int CVSpilsSetPreconditionerB(void *cvadj_mem, CVSpilsPrecSetupFnB psetB,
                                CVSpilsPrecSolveFnB psolveB, void *P_dataB);
  int CVSpilsSetJacTimesVecFnB(void *cvadj_mem, CVSpilsJacTimesVecFnB jtimesB,
                               void *jac_dataB);


#ifdef __cplusplus
}
#endif

#endif
```
