## Supplementary scripts for "A multi-scale *in silico* mouse model for insulin resistance and humanoid type 2 diabetes": cvodes_sptfqmr.h

```
/*
 * -----------------------------------------------------------------
 * $Revision: 749 $
 * $Date: 2011-06-22 14:49:10 +0200 (Wed, 22 Jun 2011) $
 * ----------------------------------------------------------------- 
 * Programmer(s): Aaron Collier and Radu Serban @ LLNL
 * -----------------------------------------------------------------
 * Copyright (c) 2005, The Regents of the University of California.
 * Produced at the Lawrence Livermore National Laboratory.
 * All rights reserved.
 * For details, see the LICENSE file.
 * -----------------------------------------------------------------
 * This is the header file for the CVODES scaled preconditioned TFQMR 
 * linear solver, CVSPTFQMR.
 *
 * Part I contains function prototypes for using CVSPTFQMR on forward 
 * problems (IVP integration and/or FSA)
 *
 * Part II contains function prototypes for using CVSPTFQMR on adjoint 
 * (backward) problems
 * -----------------------------------------------------------------
 */

#ifndef _CVSSPTFQMR_H
#define _CVSSPTFQMR_H

#ifdef __cplusplus  /* wrapper to enable C++ usage */
extern "C" {
#endif

#include <cvodes/cvodes_spils.h>
#include <sundials/sundials_sptfqmr.h>

  /* 
   * -----------------------------------------------------------------
   * PART I - forward problems
   * -----------------------------------------------------------------
   */

  /*
   * -----------------------------------------------------------------
   * Function : CVSptfqmr
   * -----------------------------------------------------------------
   * A call to the CVSptfqmr function links the main CVODE integrator
   * with the CVSPTFQMR linear solver.
   *
   * cvode_mem is the pointer to the integrator memory returned by
   *           CVodeCreate.
   *
   * pretype   is the type of user preconditioning to be done.
   *           This must be one of the four enumeration constants
   *           PREC_NONE, PREC_LEFT, PREC_RIGHT, or PREC_BOTH defined
   *           in iterative.h. These correspond to no preconditioning,
   *           left preconditioning only, right preconditioning
   *           only, and both left and right preconditioning,
   *           respectively.
   *
   * maxl      is the maximum Krylov dimension. This is an
   *           optional input to the CVSPTFQMR solver. Pass 0 to
   *           use the default value CVSPILS_MAXL=5.
   *
   * The return value of CVSptfqmr is one of:
   *    CVSPILS_SUCCESS   if successful
   *    CVSPILS_MEM_NULL  if the cvode memory was NULL
   *    CVSPILS_MEM_FAIL  if there was a memory allocation failure
   *    CVSPILS_ILL_INPUT if a required vector operation is missing
   * The above constants are defined in cvodes_spils.h
   * -----------------------------------------------------------------
   */

  int CVSptfqmr(void *cvode_mem, int pretype, int maxl);


  /* 
   * -----------------------------------------------------------------
   * PART II - backward problems
   * -----------------------------------------------------------------
   */

  int CVSptfqmrB(void *cvadj_mem, int pretypeB, int maxlB);


#ifdef __cplusplus
}
#endif

#endif
```
