## Supplementary scripts for "A multi-scale *in silico* mouse model for insulin resistance and humanoid type 2 diabetes": nvector_serial.h

```
/*
 * -----------------------------------------------------------------
 * $Revision: 749 $
 * $Date: 2011-06-22 14:49:10 +0200 (Wed, 22 Jun 2011) $
 * ----------------------------------------------------------------- 
 * Programmer(s): Scott D. Cohen, Alan C. Hindmarsh, Radu Serban,
 *                and Aaron Collier @ LLNL
 * -----------------------------------------------------------------
 * Copyright (c) 2002, The Regents of the University of California.
 * Produced at the Lawrence Livermore National Laboratory.
 * All rights reserved.
 * For details, see the LICENSE file.
 * -----------------------------------------------------------------
 * This is the header file for the serial implementation of the
 * NVECTOR module.
 *
 * Part I contains declarations specific to the serial
 * implementation of the supplied NVECTOR module.
 *
 * Part II defines accessor macros that allow the user to
 * efficiently use the type N_Vector without making explicit
 * references to the underlying data structure.
 *
 * Part III contains the prototype for the constructor N_VNew_Serial
 * as well as implementation-specific prototypes for various useful
 * vector operations.
 *
 * Notes:
 *
 *   - The definition of the generic N_Vector structure can be found
 *     in the header file sundials_nvector.h.
 *
 *   - The definition of the type 'realtype' can be found in the
 *     header file sundials_types.h, and it may be changed (at the 
 *     configuration stage) according to the user's needs. 
 *     The sundials_types.h file also contains the definition
 *     for the type 'booleantype'.
 *
 *   - N_Vector arguments to arithmetic vector operations need not
 *     be distinct. For example, the following call:
 *
 *       N_VLinearSum_Serial(a,x,b,y,y);
 *
 *     (which stores the result of the operation a*x+b*y in y)
 *     is legal.
 * -----------------------------------------------------------------
 */

#ifndef _NVECTOR_SERIAL_H
#define _NVECTOR_SERIAL_H

#ifdef __cplusplus  /* wrapper to enable C++ usage */
extern "C" {
#endif

#include <sundials/sundials_nvector.h>

/*
 * -----------------------------------------------------------------
 * PART I: SERIAL implementation of N_Vector
 * -----------------------------------------------------------------
 */

/* serial implementation of the N_Vector 'content' structure
   contains the length of the vector, a pointer to an array
   of 'realtype' components, and a flag indicating ownership of
   the data */

struct _N_VectorContent_Serial {
  long int length;
  booleantype own_data;
  realtype *data;
};

typedef struct _N_VectorContent_Serial *N_VectorContent_Serial;

/*
 * -----------------------------------------------------------------
 * PART II: macros NV_CONTENT_S, NV_DATA_S, NV_OWN_DATA_S,
 *          NV_LENGTH_S, and NV_Ith_S
 * -----------------------------------------------------------------
 * In the descriptions below, the following user declarations
 * are assumed:
 *
 * N_Vector v;
 * long int i;
 *
 * (1) NV_CONTENT_S
 *
 *     This routines gives access to the contents of the serial
 *     vector N_Vector.
 *
 *     The assignment v_cont = NV_CONTENT_S(v) sets v_cont to be
 *     a pointer to the serial N_Vector content structure.
 *
 * (2) NV_DATA_S NV_OWN_DATA_S and NV_LENGTH_S
 *
 *     These routines give access to the individual parts of
 *     the content structure of a serial N_Vector.
 *
 *     The assignment v_data = NV_DATA_S(v) sets v_data to be
 *     a pointer to the first component of v. The assignment
 *     NV_DATA_S(v) = data_V sets the component array of v to
 *     be data_v by storing the pointer data_v.
 *
 *     The assignment v_len = NV_LENGTH_S(v) sets v_len to be
 *     the length of v. The call NV_LENGTH_S(v) = len_v sets
 *     the length of v to be len_v.
 *
 * (3) NV_Ith_S
 *
 *     In the following description, the components of an
 *     N_Vector are numbered 0..n-1, where n is the length of v.
 *
 *     The assignment r = NV_Ith_S(v,i) sets r to be the value of
 *     the ith component of v. The assignment NV_Ith_S(v,i) = r
 *     sets the value of the ith component of v to be r.
 *
 * Note: When looping over the components of an N_Vector v, it is
 * more efficient to first obtain the component array via
 * v_data = NV_DATA_S(v) and then access v_data[i] within the
 * loop than it is to use NV_Ith_S(v,i) within the loop.
 * -----------------------------------------------------------------
 */

#define NV_CONTENT_S(v)  ( (N_VectorContent_Serial)(v->content) )

#define NV_LENGTH_S(v)   ( NV_CONTENT_S(v)->length )

#define NV_OWN_DATA_S(v) ( NV_CONTENT_S(v)->own_data )

#define NV_DATA_S(v)     ( NV_CONTENT_S(v)->data )

#define NV_Ith_S(v,i)    ( NV_DATA_S(v)[i] )

/*
 * -----------------------------------------------------------------
 * PART III: functions exported by nvector_serial
 * 
 * CONSTRUCTORS:
 *    N_VNew_Serial
 *    N_VNewEmpty_Serial
 *    N_VMake_Serial
 *    N_VCloneVectorArray_Serial
 *    N_VCloneVectorArrayEmpty_Serial
 * DESTRUCTORS:
 *    N_VDestroy_Serial
 *    N_VDestroyVectorArray_Serial
 * OTHER:
 *    N_VPrint_Serial
 * -----------------------------------------------------------------
 */

/*
 * -----------------------------------------------------------------
 * Function : N_VNew_Serial
 * -----------------------------------------------------------------
 * This function creates and allocates memory for a serial vector.
 * -----------------------------------------------------------------
 */

N_Vector N_VNew_Serial(long int vec_length);

/*
 * -----------------------------------------------------------------
 * Function : N_VNewEmpty_Serial
 * -----------------------------------------------------------------
 * This function creates a new serial N_Vector with an empty (NULL)
 * data array.
 * -----------------------------------------------------------------
 */

N_Vector N_VNewEmpty_Serial(long int vec_length);

/*
 * -----------------------------------------------------------------
 * Function : N_VMake_Serial
 * -----------------------------------------------------------------
 * This function creates and allocates memory for a serial vector
 * with a user-supplied data array.
 * -----------------------------------------------------------------
 */

N_Vector N_VMake_Serial(long int vec_length, realtype *v_data);

/*
 * -----------------------------------------------------------------
 * Function : N_VCloneVectorArray_Serial
 * -----------------------------------------------------------------
 * This function creates an array of 'count' SERIAL vectors by
 * cloning a given vector w.
 * -----------------------------------------------------------------
 */

N_Vector *N_VCloneVectorArray_Serial(int count, N_Vector w);

/*
 * -----------------------------------------------------------------
 * Function : N_VCloneVectorArrayEmpty_Serial
 * -----------------------------------------------------------------
 * This function creates an array of 'count' SERIAL vectors each
 * with an empty (NULL) data array by cloning w.
 * -----------------------------------------------------------------
 */

N_Vector *N_VCloneVectorArrayEmpty_Serial(int count, N_Vector w);

/*
 * -----------------------------------------------------------------
 * Function : N_VDestroyVectorArray_Serial
 * -----------------------------------------------------------------
 * This function frees an array of SERIAL vectors created with 
 * N_VCloneVectorArray_Serial or N_VCloneVectorArrayEmpty_Serial.
 * -----------------------------------------------------------------
 */

void N_VDestroyVectorArray_Serial(N_Vector *vs, int count);

/*
 * -----------------------------------------------------------------
 * Function : N_VPrint_Serial
 * -----------------------------------------------------------------
 * This function prints the content of a serial vector to stdout.
 * -----------------------------------------------------------------
 */

void N_VPrint_Serial(N_Vector v);

/*
 * -----------------------------------------------------------------
 * serial implementations of various useful vector operations
 * -----------------------------------------------------------------
 */

N_Vector N_VCloneEmpty_Serial(N_Vector w);
N_Vector N_VClone_Serial(N_Vector w);
void N_VDestroy_Serial(N_Vector v);
void N_VSpace_Serial(N_Vector v, long int *lrw, long int *liw);
realtype *N_VGetArrayPointer_Serial(N_Vector v);
void N_VSetArrayPointer_Serial(realtype *v_data, N_Vector v);
void N_VLinearSum_Serial(realtype a, N_Vector x, realtype b, N_Vector y, N_Vector z);
void N_VConst_Serial(realtype c, N_Vector z);
void N_VProd_Serial(N_Vector x, N_Vector y, N_Vector z);
void N_VDiv_Serial(N_Vector x, N_Vector y, N_Vector z);
void N_VScale_Serial(realtype c, N_Vector x, N_Vector z);
void N_VAbs_Serial(N_Vector x, N_Vector z);
void N_VInv_Serial(N_Vector x, N_Vector z);
void N_VAddConst_Serial(N_Vector x, realtype b, N_Vector z);
realtype N_VDotProd_Serial(N_Vector x, N_Vector y);
realtype N_VMaxNorm_Serial(N_Vector x);
realtype N_VWrmsNorm_Serial(N_Vector x, N_Vector w);
realtype N_VWrmsNormMask_Serial(N_Vector x, N_Vector w, N_Vector id);
realtype N_VMin_Serial(N_Vector x);
realtype N_VWL2Norm_Serial(N_Vector x, N_Vector w);
realtype N_VL1Norm_Serial(N_Vector x);
void N_VCompare_Serial(realtype c, N_Vector x, N_Vector z);
booleantype N_VInvTest_Serial(N_Vector x, N_Vector z);
booleantype N_VConstrMask_Serial(N_Vector c, N_Vector x, N_Vector m);
realtype N_VMinQuotient_Serial(N_Vector num, N_Vector denom);

#ifdef __cplusplus
}
#endif

#endif
```
