## Supplementary scripts for "A multi-scale *in silico* mouse model for insulin resistance and humanoid type 2 diabetes": sundials_band.h

```
/*
 * -----------------------------------------------------------------
 * $Revision: 749 $
 * $Date: 2011-06-22 14:49:10 +0200 (Wed, 22 Jun 2011) $
 * -----------------------------------------------------------------
 * Programmer(s): Alan C. Hindmarsh and Radu Serban @ LLNL
 * -----------------------------------------------------------------
 * Copyright (c) 2002, The Regents of the University of California.
 * Produced at the Lawrence Livermore National Laboratory.
 * All rights reserved.
 * For details, see the LICENSE file.
 * -----------------------------------------------------------------
 * This is the header file for a generic BAND linear solver
 * package. There are two sets of band solver routines listed in
 * this file: one set uses type BandMat defined below and the
 * other set uses the type realtype ** for band matrix arguments.
 * The two sets of band solver routines make it easy to work
 * with two types of band matrices:
 *
 * (1) The BandMat type is intended for use with large
 *     band matrices whose elements/columns may be stored in
 *     non-contiguous memory locations or even distributed across
 *     different process memory spaces. This type may be modified
 *     to include such distribution information. If this is done,
 *     then all the routines that use BandMat must be modified to
 *     reflect the new data structure.
 *
 * (2) The set of routines that use realtype ** (and NOT the
 *     BandMat type) is intended for use with small matrices
 *     which can easily be allocated within a contiguous block of
 *     memory for a single process.
 *
 * Routines that work with the type BandMat begin with "Band".
 * The BandAllocMat function allocates a band matrix for use in
 * the other matrix routines listed in this file. Matrix storage
 * details are given in the documentation for the type BandMat.
 * The BandAllocPiv function allocates memory for pivot
 * information. The storage allocated by BandAllocMat and
 * BandAllocPiv is deallocated by the routines BandFreeMat and
 * BandFreePiv, respectively. The BandGBTRF and BandGBTRS
 * routines perform the actual solution of a band linear system.
 *
 * Routines that work with realtype ** begin with "band" (except
 * for the factor and solve routines which are called bandGBTRF and
 * bandGBTRS, respectively). The underlying matrix storage is
 * described in the documentation for bandalloc.
 * -----------------------------------------------------------------
 */

#ifndef _BAND_H
#define _BAND_H

#ifdef __cplusplus  /* wrapper to enable C++ usage */
extern "C" {
#endif

#include <sundials/sundials_types.h>

/*
 * -----------------------------------------------------------------
 * Type: BandMat
 * -----------------------------------------------------------------
 * The type BandMat is the type of a large (possibly distributed)
 * band matrix. It is defined to be a pointer to a structure
 * with the following fields:
 *
 * size is the number of columns (== number of rows)
 *
 * mu   is the upper bandwidth, 0 <= mu <= size-1
 *
 * ml   is the lower bandwidth, 0 <= ml <= size-1
 *
 * smu  is the storage upper bandwidth, mu <= smu <= size-1.
 *      The BandGBTRF routine writes the LU factors
 *      into the storage for A. The upper triangular factor U,
 *      however, may have an upper bandwidth as big as
 *      MIN(size-1,mu+ml) because of partial pivoting. The smu
 *      field holds the upper bandwidth allocated for A.
 *
 * data is a two dimensional array used for component storage.
 *      The elements of a band matrix of type BandMat are
 *      stored columnwise (i.e. columns are stored one on top
 *      of the other in memory). Only elements within the
 *      specified bandwidths are stored.
 *
 * If we number rows and columns in the band matrix starting
 * from 0, then
 *
 * data[0] is a pointer to (smu+ml+1)*size contiguous locations
 *         which hold the elements within the band of A
 *
 * data[j] is a pointer to the uppermost element within the band
 *         in the jth column. This pointer may be treated as
 *         an array indexed from smu-mu (to access the
 *         uppermost element within the band in the jth
 *         column) to smu+ml (to access the lowest element
 *         within the band in the jth column). (Indices from 0
 *         to smu-mu-1 give access to extra storage elements
 *         required by BandGBTRF.)
 *
 * data[j][i-j+smu] is the (i,j)th element, j-mu <= i <= j+ml.
 *
 * The macros below allow a user to access individual matrix
 * elements without writing out explicit data structure
 * references and without knowing too much about the underlying
 * element storage. The only storage assumption needed is that
 * elements are stored columnwise and that a pointer into the jth
 * column of elements can be obtained via the BAND_COL macro. The
 * BAND_COL_ELEM macro selects an element from a column which has
 * already been isolated via BAND_COL. BAND_COL_ELEM allows the
 * user to avoid the translation from the matrix location (i,j)
 * to the index in the array returned by BAND_COL at which the
 * (i,j)th element is stored. See the documentation for BAND_COL
 * and BAND_COL_ELEM for usage details. Users should use these
 * macros whenever possible.
 * -----------------------------------------------------------------
 */

typedef struct _BandMat {
  long int size;
  long int mu, ml, smu;
  realtype **data;
} *BandMat;

/* BandMat accessor macros */

/*
 * -----------------------------------------------------------------
 * Macro : BAND_ELEM
 * -----------------------------------------------------------------
 * Usage : BAND_ELEM(A,i,j) = a_ij;  OR
 *         a_ij = BAND_ELEM(A,i,j);
 * -----------------------------------------------------------------
 * BAND_ELEM(A,i,j) references the (i,j)th element of the
 * N by N band matrix A, where 0 <= i,j <= N-1. The location
 * (i,j) should further satisfy j-(A->mu) <= i <= j+(A->ml).
 * -----------------------------------------------------------------
 */

#define BAND_ELEM(A,i,j) ((A->data)[j][(i)-(j)+(A->smu)])

/*
 * -----------------------------------------------------------------
 * Macro : BAND_COL
 * -----------------------------------------------------------------
 * Usage : col_j = BAND_COL(A,j);
 * -----------------------------------------------------------------
 * BAND_COL(A,j) references the diagonal element of the jth
 * column of the N by N band matrix A, 0 <= j <= N-1. The type of
 * the expression BAND_COL(A,j) is realtype *. The pointer
 * returned by the call BAND_COL(A,j) can be treated as an array
 * which is indexed from -(A->mu) to (A->ml).
 * -----------------------------------------------------------------
 */

#define BAND_COL(A,j) (((A->data)[j])+(A->smu))

/*
 * -----------------------------------------------------------------
 * Macro : BAND_COL_ELEM
 * -----------------------------------------------------------------
 * Usage : col_j = BAND_COL(A,j);
 *         BAND_COL_ELEM(col_j,i,j) = a_ij;  OR
 *         a_ij = BAND_COL_ELEM(col_j,i,j);
 * -----------------------------------------------------------------
 * This macro references the (i,j)th entry of the band matrix A
 * when used in conjunction with BAND_COL as shown above. The
 * index (i,j) should satisfy j-(A->mu) <= i <= j+(A->ml).
 * -----------------------------------------------------------------
 */

#define BAND_COL_ELEM(col_j,i,j) (col_j[(i)-(j)])

/* Functions that use the BandMat representation for a band matrix */

/*
 * -----------------------------------------------------------------
 * Function : BandAllocMat
 * -----------------------------------------------------------------
 * Usage : A = BandAllocMat(N, mu, ml, smu);
 *         if (A == NULL) ... memory request failed
 * -----------------------------------------------------------------
 * BandAllocMat allocates memory for an N by N band matrix with
 * upper bandwidth mu, lower bandwidth ml, and storage upper
 * bandwidth smu. Pass smu as follows depending on whether A will
 * be factored by BandGBTRF:
 *
 * (1) Pass smu = mu if A will not be factored.
 *
 * (2) Pass smu = MIN(N-1,mu+ml) if A will be factored.
 *
 * BandAllocMat returns the storage allocated (type BandMat) or
 * NULL if the request for matrix storage cannot be satisfied.
 * See the documentation for the type BandMat for matrix storage
 * details.
 * -----------------------------------------------------------------
 */

BandMat BandAllocMat(long int N, long int mu, long int ml, 
                     long int smu);

/*
 * -----------------------------------------------------------------
 * Function : BandAllocPiv
 * -----------------------------------------------------------------
 * Usage : p = BandAllocPiv(N);
 *         if (p == NULL) ... memory request failed
 * -----------------------------------------------------------------
 * BandAllocPiv allocates memory for pivot information to be
 * filled in by the BandGBTRF routine during the factorization
 * of an N by N band matrix. The underlying type for pivot
 * information is an array of N integers and this routine returns
 * the pointer to the memory it allocates. If the request for
 * pivot storage cannot be satisfied, BandAllocPiv returns NULL.
 * -----------------------------------------------------------------
 */

long int *BandAllocPiv(long int N);

/*
 * -----------------------------------------------------------------
 * Function : BandGBTRF
 * -----------------------------------------------------------------
 * Usage : ier = BandGBTRF(A, p);
 *         if (ier != 0) ... A is singular
 * -----------------------------------------------------------------
 * BandGBTRF performs the LU factorization of the N by N band
 * matrix A. This is done using standard Gaussian elimination
 * with partial pivoting.
 *
 * A successful LU factorization leaves the "matrix" A and the
 * pivot array p with the following information:
 *
 * (1) p[k] contains the row number of the pivot element chosen
 *     at the beginning of elimination step k, k=0, 1, ..., N-1.
 *
 * (2) If the unique LU factorization of A is given by PA = LU,
 *     where P is a permutation matrix, L is a lower triangular
 *     matrix with all 1's on the diagonal, and U is an upper
 *     triangular matrix, then the upper triangular part of A
 *     (including its diagonal) contains U and the strictly lower
 *     triangular part of A contains the multipliers, I-L.
 *
 * BandGBTRF returns 0 if successful. Otherwise it encountered
 * a zero diagonal element during the factorization. In this case
 * it returns the column index (numbered from one) at which
 * it encountered the zero.
 *
 * Important Note: A must be allocated to accommodate the increase
 * in upper bandwidth that occurs during factorization. If
 * mathematically, A is a band matrix with upper bandwidth mu and
 * lower bandwidth ml, then the upper triangular factor U can
 * have upper bandwidth as big as smu = MIN(n-1,mu+ml). The lower
 * triangular factor L has lower bandwidth ml. Allocate A with
 * call A = BandAllocMat(N,mu,ml,smu), where mu, ml, and smu are
 * as defined above. The user does not have to zero the "extra"
 * storage allocated for the purpose of factorization. This will
 * handled by the BandGBTRF routine.
 * -----------------------------------------------------------------
 */

long int BandGBTRF(BandMat A, long int *p);

/*
 * -----------------------------------------------------------------
 * Function : BandGBTRS
 * -----------------------------------------------------------------
 * Usage : BandGBTRS(A, p, b);
 * -----------------------------------------------------------------
 * BandGBTRS solves the N-dimensional system A x = b using
 * the LU factorization in A and the pivot information in p
 * computed in BandGBTRF. The solution x is returned in b. This
 * routine cannot fail if the corresponding call to BandGBTRF
 * did not fail.
 * -----------------------------------------------------------------
 */

void BandGBTRS(BandMat A, long int *p, realtype *b);

/*
 * -----------------------------------------------------------------
 * Function : BandZero
 * -----------------------------------------------------------------
 * Usage : BandZero(A);
 * -----------------------------------------------------------------
 * A(i,j) <- 0.0,    j-(A->mu) <= i <= j+(A->ml).
 * -----------------------------------------------------------------
 */

void BandZero(BandMat A);

/*
 * -----------------------------------------------------------------
 * Function : BandCopy
 * -----------------------------------------------------------------
 * Usage : BandCopy(A, B, copymu, copyml);
 * -----------------------------------------------------------------
 * BandCopy copies the submatrix with upper and lower bandwidths
 * copymu, copyml of the N by N band matrix A into the N by N
 * band matrix B.
 * -----------------------------------------------------------------
 */

void BandCopy(BandMat A, BandMat B, long int copymu, long int copyml);

/*
 * -----------------------------------------------------------------
 * Function: BandScale
 * -----------------------------------------------------------------
 * Usage : BandScale(c, A);
 * -----------------------------------------------------------------
 * A(i,j) <- c*A(i,j),   j-(A->mu) <= i <= j+(A->ml).
 * -----------------------------------------------------------------
 */

void BandScale(realtype c, BandMat A);

/*
 * -----------------------------------------------------------------
 * Function : BandAddI
 * -----------------------------------------------------------------
 * Usage : BandAddI(A);
 * -----------------------------------------------------------------
 * A(j,j) <- A(j,j)+1.0,   0 <= j <= (A->size)-1.
 * -----------------------------------------------------------------
 */

void BandAddI(BandMat A);

/*
 * -----------------------------------------------------------------
 * Function : BandFreeMat
 * -----------------------------------------------------------------
 * Usage : BandFreeMat(A);
 * -----------------------------------------------------------------
 * BandFreeMat frees the memory allocated by BandAllocMat for
 * the band matrix A.
 * -----------------------------------------------------------------
 */

void BandFreeMat(BandMat A);

/*
 * -----------------------------------------------------------------
 * Function : BandFreePiv
 * -----------------------------------------------------------------
 * Usage : BandFreePiv(p);
 * -----------------------------------------------------------------
 * BandFreePiv frees the memory allocated by BandAllocPiv for
 * the pivot information array p.
 * -----------------------------------------------------------------
 */

void BandFreePiv(long int *p);

/*
 * -----------------------------------------------------------------
 * Function : BandPrint
 * -----------------------------------------------------------------
 * Usage : BandPrint(A);
 * -----------------------------------------------------------------
 * This routine prints the N by N band matrix A (upper and lower
 * bandwidths A->mu and A->ml, respectively) to standard output
 * as it would normally appear on paper. It is intended as a
 * debugging tool with small values of N. The elements are
 * printed using the %g/lg/Lg option. A blank line is printed
 * before and after the matrix.
 * -----------------------------------------------------------------
 */

void BandPrint(BandMat A);

/* Functions that use the realtype ** representation for a band matrix */

/*
 * -----------------------------------------------------------------
 * Function : bandalloc
 * -----------------------------------------------------------------
 * Usage : realtype **a;
 *         a = bandalloc(n, smu, ml);
 *         if (a == NULL) ... memory request failed
 * -----------------------------------------------------------------
 * bandalloc(n, smu, ml) allocates storage for an n by n band
 * matrix A with storage upper bandwidth smu and lower bandwidth
 * ml. It returns a pointer to the newly allocated storage if
 * successful. If the memory request cannot be satisfied, then
 * bandalloc returns NULL. If, mathematically, A has upper and
 * lower bandwidths mu and ml, respectively, then the value
 * passed to bandalloc for smu may need to be greater than mu.
 * The bandGBTRF routine writes the LU factors into the storage
 * (named "a" in the above usage documentation) for A (thus destroying
 * the original elements of A). The upper triangular factor U,
 * however, may have a larger upper bandwidth than the upper
 * bandwidth mu of A. Thus some "extra" storage for A must be
 * allocated if A is to be factored by bandGBTRF. Pass smu as follows:
 *
 * (1) Pass smu = mu if A will not be factored.
 *
 * (2) Pass smu = MIN(n-1,mu+ml) if A will be factored.
 *
 * The underlying type of the band matrix returned is realtype**.
 * If we allocate a band matrix A in realtype **a by
 * a = bandalloc(n,smu,ml), then a[0] is a pointer to
 * n * (smu + ml + 1) contiguous storage locations and a[j] is a
 * pointer to the uppermost element in the storage for the jth
 * column. The expression a[j][i-j+smu] references the (i,j)th
 * element of A, where 0 <= i,j <= n-1 and j-mu <= i <= j+ml.
 * (The elements a[j][0], a[j][1], ..., a[j][smu-mu-1] are used
 * by bandGBTRF and bandGBTRS.)
 * -----------------------------------------------------------------
 */

realtype **bandalloc(long int n, long int smu, long int ml);

/*
 * -----------------------------------------------------------------
 * Function : bandallocpiv
 * -----------------------------------------------------------------
 * Usage : long int *pivot;
 *         pivot = bandallocpiv(n);
 *         if (pivot == NULL) ... memory request failed
 * -----------------------------------------------------------------
 * bandallocpiv(n) allocates an array of n integers. It returns a
 * pointer to the first element in the array if successful. It
 * returns NULL if the memory request could not be satisfied.
 * -----------------------------------------------------------------
 */

long int *bandallocpiv(long int n);

/*
 * -----------------------------------------------------------------
 * Function : bandGBTRF
 * -----------------------------------------------------------------
 * Usage : long int ier;
 *         ier = bandGBTRF(a,n,mu,ml,smu,p);
 *         if (ier > 0) ... zero element encountered during
 *                          the factorization
 * -----------------------------------------------------------------
 * bandGBTRF(a,n,mu,ml,smu,p) factors the n by n band matrix A 
 * (upper and lower bandwidths mu and ml, storage upper bandwidth smu)
 * stored in "a". It overwrites the elements of A with the LU
 * factors and it keeps track of the pivot rows chosen in the
 * pivot array p.
 *
 * A successful LU factorization leaves a and pivot array p with
 * the following information:
 *
 * (1) p[k] contains the row number of the pivot element chosen
 *     at the beginning of elimination step k, k=0, 1, ..., n-1.
 *
 * (2) If the unique LU factorization of A is given by PA = LU,
 *     where P is a permutation matrix, L is a lower triangular
 *     matrix with all 1's on the diagonal, and U is an upper
 *     triangular matrix, then the upper triangular part of A
 *     (including its diagonal) contains U and the strictly lower
 *     triangular part of A contains the multipliers, I-L.
 *
 * bandGBTRF returns 0 if successful. Otherwise it encountered a zero
 * diagonal element during the factorization. In this case it
 * returns the column index (numbered from one) at which it
 * encountered the zero.
 *
 * IMPORTANT NOTE: Suppose A is a band matrix with upper
 * bandwidth mu and lower bandwidth ml, then the upper triangular
 * factor U can have upper bandwidth as big as MIN(n-1,mu+ml)
 * because of partial pivoting. The lower triangular factor L has
 * lower bandwidth ml. Thus, if A is to be factored and
 * backsolved using bandGBTRF and bandGBTRS, then it should be allocated
 * as a = bandalloc(n,smu,ml), where smu = MIN(n-1,mu+ml). The
 * call to bandGBTRF is ier = bandGBTRF(a,n,mu,ml,smu,p). The
 * corresponding call to bandGBTRS is bandGBTRS(a,n,smu,ml,p,b). The user 
 * does not need to zero the "extra" storage allocated for the 
 * purpose of factorization. This is handled by the bandGBTRF routine. 
 * If A is not going to be factored and backsolved, then it can be
 * allocated as a = bandalloc(n,smu,ml). In either case, all
 * routines in this section use the parameter name smu for a
 * parameter which must be the "storage upper bandwidth" which
 * was passed to bandalloc.
 * -----------------------------------------------------------------
 */

long int bandGBTRF(realtype **a, long int n, long int mu, long int ml, 
                   long int smu, long int *p);

/*
 * -----------------------------------------------------------------
 * Function : bandGBTRS
 * -----------------------------------------------------------------
 * Usage : realtype *b;
 *         ier = bandGBTRF(a,n,mu,ml,smu,p);
 *         if (ier == 0) bandGBTRS(a,n,smu,ml,p,b);
 * -----------------------------------------------------------------
 * bandGBTRS(a,n,smu,ml,p,b) solves the n by n linear system
 * Ax = b, where A is band matrix stored in "a" with storage
 * upper bandwidth smu and lower bandwidth ml. It assumes that A
 * has been LU factored and the pivot array p has been set by a
 * successful call bandGBTRF(a,n,mu,ml,smu,p). The solution x is
 * written into the b array.
 * -----------------------------------------------------------------
 */

void bandGBTRS(realtype **a, long int n, long int smu, 
               long int ml, long int *p, realtype *b);

/*
 * -----------------------------------------------------------------
 * Function : bandzero
 * -----------------------------------------------------------------
 * Usage : bandzero(a,n,mu,ml,smu);
 * -----------------------------------------------------------------
 * a(i,j) <- 0.0,   0 <= i,j <= n-1, j-mu <= i <= j+ml.
 * -----------------------------------------------------------------
 */

void bandzero(realtype **a, long int n, long int mu, 
              long int ml, long int smu);

/*
 * -----------------------------------------------------------------
 * Function : bandcopy
 * -----------------------------------------------------------------
 * Usage : bandcopy(a,b,n,a_smu,b_smu,copymu,copyml);
 * -----------------------------------------------------------------
 * b(i,j) <- a(i,j), 0 <= i,j <= n-1, j-copymu <= i <= j+copyml.
 * -----------------------------------------------------------------
 */

void bandcopy(realtype **a, realtype **b, long int n, 
              long int a_smu, long int b_smu,
              long int copymu, long int copyml);

/*
 * -----------------------------------------------------------------
 * Function : bandscale
 * -----------------------------------------------------------------
 * Usage : bandscale(c,a,n,mu,ml);
 * -----------------------------------------------------------------
 * a(i,j) <- c*a(i,j),   0 <= i,j <= n-1, j-mu <= i <= j+ml.
 * -----------------------------------------------------------------
 */

void bandscale(realtype c, realtype **a, long int n, 
               long int mu, long int ml, long int smu);

/*
 * -----------------------------------------------------------------
 * Function : bandaddI
 * -----------------------------------------------------------------
 * Usage : bandaddI(a,n,smu);
 * -----------------------------------------------------------------
 * a(j,j) <- a(j,j)+1.0,   0 <= j <= n-1.
 * -----------------------------------------------------------------
 */

void bandaddI(realtype **a, long int n, long int smu);

/*
 * -----------------------------------------------------------------
 * Function : bandfreepiv
 * -----------------------------------------------------------------
 * Usage : bandfreepiv(p);
 * -----------------------------------------------------------------
 * bandfreepiv(p) frees the pivot array p allocated by
 * bandallocpiv.
 * -----------------------------------------------------------------
 */

void bandfreepiv(long int *p);

/*
 * -----------------------------------------------------------------
 * Function : bandfree
 * -----------------------------------------------------------------
 * Usage : bandfree(a);
 * -----------------------------------------------------------------
 * bandfree(a) frees the band matrix a allocated by bandalloc.
 * -----------------------------------------------------------------
 */

void bandfree(realtype **a);

/*
 * -----------------------------------------------------------------
 * Function : bandprint
 * -----------------------------------------------------------------
 * Usage : bandprint(a,n,mu,ml,smu);
 * -----------------------------------------------------------------
 * bandprint(a,n,mu,ml,smu) prints the n by n band matrix stored
 * in a (with upper bandwidth mu and lower bandwidth ml) to
 * standard output as it would normally appear on paper. It is
 * intended as a debugging tool with small values of n. The
 * elements are printed using the %g/lg/Lg option. A blank line
 * is printed before and after the matrix.
 * -----------------------------------------------------------------
 */

void bandprint(realtype **a, long int n, long int mu, long int ml, 
               long int smu);

#ifdef __cplusplus
}
#endif

#endif
```
