## Supplementary scripts for "A multi-scale *in silico* mouse model for insulin resistance and humanoid type 2 diabetes": sundials_dense.h

```
/*
 * -----------------------------------------------------------------
 * $Revision: 749 $
 * $Date: 2011-06-22 14:49:10 +0200 (Wed, 22 Jun 2011) $
 * -----------------------------------------------------------------
 * Programmer: Radu Serban @ LLNL
 * -----------------------------------------------------------------
 * Copyright (c) 2006, The Regents of the University of California.
 * Produced at the Lawrence Livermore National Laboratory.
 * All rights reserved.
 * For details, see the LICENSE file.
 * -----------------------------------------------------------------
 * This is the header file for a generic package of DENSE matrix
 * operations.  The routines listed in this file all use type
 * DenseMat, defined below, for M by N matrices.
 * These routines in turn call routines in the smalldense module,
 * which use the type realtype** for matrices. This separation allows
 * for possible modifications in which matrices of type DenseMat
 * may not be stored contiguously, while small matrices can still
 * be treated with the routines in smalldense.
 *
 * Routines that work with the type DenseMat begin with "Dense".
 * The DenseAllocMat function allocates a dense matrix for use in
 * the other DenseMat routines listed in this file. Matrix
 * storage details are given in the documentation for the type
 * DenseMat. The DenseAllocPiv function allocates memory for
 * pivot information. The storage allocated by DenseAllocMat and
 * DenseAllocPiv is deallocated by the routines DenseFreeMat and
 * DenseFreePiv, respectively. The DenseGETRF and DenseGETRS
 * routines perform the actual solution of a dense linear system.
 *
 * Routines that work with realtype** begin with "den". 
 * The underlying matrix storage is described in the documentation 
 * for denalloc in smalldense.h
 * -----------------------------------------------------------------
 */

#ifndef _DENSE_H
#define _DENSE_H

#ifdef __cplusplus  /* wrapper to enable C++ usage */
extern "C" {
#endif

#include <sundials/sundials_types.h>
#include <sundials/sundials_smalldense.h>

  /*
   * ==================================================================
   * Type definitions
   * ==================================================================
   */

  /*
   * -----------------------------------------------------------------
   * Type : DenseMat
   * -----------------------------------------------------------------
   * The type DenseMat is defined to be a pointer to a structure
   * with sizes (M and N) and a data field. The M and N fields 
   * indicates the number of rows and columns, respectively of a dense 
   * matrix, while the data field is a two dimensional array used for 
   * component storage. The elements of a dense matrix are stored 
   * columnwise (i.e columns are stored one on top of the other in 
   * memory). If A is of type DenseMat, then the (i,j)th element 
   * of A (with 0 <= i < M and 0 <= j < N) is given by the expression 
   * (A->data)[j][i] or by the expression (A->data)[0][j*n+i]. 
   * The macros below allow a user to access efficiently individual
   * matrix elements without writing out explicit data structure
   * references and without knowing too much about the underlying
   * element storage. The only storage assumption needed is that
   * elements are stored columnwise and that a pointer to the jth
   * column of elements can be obtained via the DENSE_COL macro.
   * Users should use these macros whenever possible.
   * -----------------------------------------------------------------
   */

  typedef struct _DenseMat {
    long int M;
    long int N;
    realtype **data;
  } *DenseMat;

  /*
   * ==================================================================
   * Data accessor macros
   * ==================================================================
   */

  /*
   * -----------------------------------------------------------------
   * Macro : DENSE_ELEM
   * -----------------------------------------------------------------
   * Usage : DENSE_ELEM(A,i,j) = a_ij;  OR
   *         a_ij = DENSE_ELEM(A,i,j);
   * -----------------------------------------------------------------
   * DENSE_ELEM(A,i,j) references the (i,j)th element of the M by N
   * DenseMat A, 0 <= i < M ; 0 <= j < N.
   * -----------------------------------------------------------------
   */

#define DENSE_ELEM(A,i,j) ((A->data)[j][i])

  /*
   * -----------------------------------------------------------------
   * Macro : DENSE_COL
   * -----------------------------------------------------------------
   * Usage : col_j = DENSE_COL(A,j);
   * -----------------------------------------------------------------
   * DENSE_COL(A,j) references the jth column of the M by N
   * DenseMat A, 0 <= j < N. The type of the expression DENSE_COL(A,j) 
   * is (realtype *). After the assignment in the usage above, col_j 
   * may be treated as an array indexed from 0 to M-1. 
   * The (i,j)-th element of A is thus referenced by col_j[i].
   * -----------------------------------------------------------------
   */

#define DENSE_COL(A,j) ((A->data)[j])

  /*
   * ==================================================================
   * Function prototypes
   * ==================================================================
   */

  /*
   * -----------------------------------------------------------------
   * Function : DenseAllocMat
   * -----------------------------------------------------------------
   * Usage : A = DenseAllocMat(M, N);
   *         if (A == NULL) ... memory request failed
   * -----------------------------------------------------------------
   * DenseAllocMat allocates memory for an M by N dense matrix and
   * returns the storage allocated (type DenseMat). DenseAllocMat
   * returns NULL if the request for matrix storage cannot be
   * satisfied. See the above documentation for the type DenseMat
   * for matrix storage details.
   * -----------------------------------------------------------------
   */

  DenseMat DenseAllocMat(long int M, long int N);

  /*
   * -----------------------------------------------------------------
   * Function : DenseAllocPiv
   * -----------------------------------------------------------------
   * Usage : p = DenseAllocPiv(N);
   *         if (p == NULL) ... memory request failed
   * -----------------------------------------------------------------
   * DenseAllocPiv allocates memory for pivot information to be
   * filled in by the DenseGETRF routine during the factorization
   * of an N by N dense matrix. The underlying type for pivot
   * information is an array of N integers and this routine returns
   * the pointer to the memory it allocates. If the request for
   * pivot storage cannot be satisfied, DenseAllocPiv returns NULL.
   * -----------------------------------------------------------------
   */

  long int *DenseAllocPiv(long int N);

  /*
   * -----------------------------------------------------------------
   * Function : DenseGETRF
   * -----------------------------------------------------------------
   * Usage : ier = DenseGETRF(A, p);
   *         if (ier != 0) ... A is singular
   * -----------------------------------------------------------------
   * DenseGETRF performs the LU factorization of the M by N dense
   * matrix A. This is done using standard Gaussian elimination
   * with partial (row) pivoting. Note that this applies only
   * to matrices with M >= N and full column rank.
   *
   * A successful LU factorization leaves the matrix A and the
   * pivot array p with the following information:
   *
   * (1) p[k] contains the row number of the pivot element chosen
   *     at the beginning of elimination step k, k=0, 1, ..., N-1.
   *
   * (2) If the unique LU factorization of A is given by PA = LU,
   *     where P is a permutation matrix, L is a lower trapezoidal
   *     matrix with all 1's on the diagonal, and U is an upper
   *     triangular matrix, then the upper triangular part of A
   *     (including its diagonal) contains U and the strictly lower
   *     trapezoidal part of A contains the multipliers, I-L.
   *
   * For square matrices (M=N), L is unit lower triangular.
   *
   * DenseGETRF returns 0 if successful. Otherwise it encountered
   * a zero diagonal element during the factorization. In this case
   * it returns the column index (numbered from one) at which
   * it encountered the zero.
   * -----------------------------------------------------------------
   */

  long int DenseGETRF(DenseMat A, long int *p);

  /*
   * -----------------------------------------------------------------
   * Function : DenseGETRS
   * -----------------------------------------------------------------
   * Usage : DenseGETRS(A, p, b);
   * -----------------------------------------------------------------
   * DenseGETRS solves the N-dimensional system A x = b using
   * the LU factorization in A and the pivot information in p
   * computed in DenseGETRF. The solution x is returned in b. This
   * routine cannot fail if the corresponding call to DenseGETRF
   * did not fail.
   * DenseGETRS does NOT check for a square matrix!
   * -----------------------------------------------------------------
   */

  void DenseGETRS(DenseMat A, long int *p, realtype *b);

  /*
   * -----------------------------------------------------------------
   * Function : DenseZero
   * -----------------------------------------------------------------
   * Usage : DenseZero(A);
   * -----------------------------------------------------------------
   * DenseZero sets all the elements of the M by N matrix A to 0.0.
   * -----------------------------------------------------------------
   */

  void DenseZero(DenseMat A);

  /*
   * -----------------------------------------------------------------
   * Function : DenseCopy
   * -----------------------------------------------------------------
   * Usage : DenseCopy(A, B);
   * -----------------------------------------------------------------
   * DenseCopy copies the contents of the M by N matrix A into the
   * M by N matrix B.
   * -----------------------------------------------------------------
   */

  void DenseCopy(DenseMat A, DenseMat B);

  /*
   * -----------------------------------------------------------------
   * Function: DenseScale
   * -----------------------------------------------------------------
   * Usage : DenseScale(c, A);
   * -----------------------------------------------------------------
   * DenseScale scales the elements of the M by N matrix A by the
   * constant c and stores the result back in A.
   * -----------------------------------------------------------------
   */

  void DenseScale(realtype c, DenseMat A);

  /*
   * -----------------------------------------------------------------
   * Function : DenseAddI
   * -----------------------------------------------------------------
   * Usage : DenseAddI(A);
   * -----------------------------------------------------------------
   * DenseAddI adds 1.0 to the main diagonal (A_ii, i=1,2,...,N-1) of
   * the M by N matrix A (M >= N) and stores the result back in A.
   * DenseAddI is typically used with square matrices.
   * DenseAddI does not check for M >= N and therefore a segmentation
   * fault will occur if M < N!
   * -----------------------------------------------------------------
   */

  void DenseAddI(DenseMat A);

  /*
   * -----------------------------------------------------------------
   * Function : DenseFreeMat
   * -----------------------------------------------------------------
   * Usage : DenseFreeMat(A);
   * -----------------------------------------------------------------
   * DenseFreeMat frees the memory allocated by DenseAllocMat for
   * the M by N matrix A.
   * -----------------------------------------------------------------
   */

  void DenseFreeMat(DenseMat A);

  /*
   * -----------------------------------------------------------------
   * Function : DenseFreePiv
   * -----------------------------------------------------------------
   * Usage : DenseFreePiv(p);
   * -----------------------------------------------------------------
   * DenseFreePiv frees the memory allocated by DenseAllocPiv for
   * the pivot information array p.
   * -----------------------------------------------------------------
   */

  void DenseFreePiv(long int *p);

  /*
   * -----------------------------------------------------------------
   * Function : DensePrint
   * -----------------------------------------------------------------
   * Usage : DensePrint(A);
   * -----------------------------------------------------------------
   * This routine prints the M by N dense matrix A to standard output
   * as it would normally appear on paper. It is intended as a 
   * debugging tool with small values of M and N. The elements are
   * printed using the %g/%lg/%Lg option. A blank line is printed
   * before and after the matrix.
   * -----------------------------------------------------------------
   */

  void DensePrint(DenseMat A);

#ifdef __cplusplus
}
#endif

#endif
```
