## Supplementary scripts for "A multi-scale *in silico* mouse model for insulin resistance and humanoid type 2 diabetes": sundials_fnvector.h

```
/*
 * -----------------------------------------------------------------
 * $Revision: 749 $
 * $Date: 2011-06-22 14:49:10 +0200 (Wed, 22 Jun 2011) $
 * ----------------------------------------------------------------- 
 * Programmer(s): Radu Serban and Aaron Collier @ LLNL
 * -----------------------------------------------------------------
 * Copyright (c) 2002, The Regents of the University of California.
 * Produced at the Lawrence Livermore National Laboratory.
 * All rights reserved.
 * For details, see the LICENSE file.
 * -----------------------------------------------------------------
 * This file (companion of nvector.h) contains definitions 
 * needed for the initialization of vector operations in Fortran.
 * -----------------------------------------------------------------
 */


#ifndef _FNVECTOR_H
#define _FNVECTOR_H

#ifdef __cplusplus  /* wrapper to enable C++ usage */
extern "C" {
#endif

#ifndef _SUNDIALS_CONFIG_H
#define _SUNDIALS_CONFIG_H
#include <sundials/sundials_config.h>
#endif

/* SUNDIALS solver IDs */

#define FCMIX_CVODE   1
#define FCMIX_IDA     2
#define FCMIX_KINSOL  3

#ifdef __cplusplus
}
#endif

#endif
```
