## Supplementary scripts for "A multi-scale *in silico* mouse model for insulin resistance and humanoid type 2 diabetes": sundials_iterative.h

```
/*
 * -----------------------------------------------------------------
 * $Revision: 749 $
 * $Date: 2011-06-22 14:49:10 +0200 (Wed, 22 Jun 2011) $
 * ----------------------------------------------------------------- 
 * Programmer(s): Scott D. Cohen and Alan C. Hindmarsh @ LLNL
 * -----------------------------------------------------------------
 * Copyright (c) 2002, The Regents of the University of California.
 * Produced at the Lawrence Livermore National Laboratory.
 * All rights reserved.
 * For details, see the LICENSE file.
 * -----------------------------------------------------------------
 * This header file contains declarations intended for use by
 * generic iterative solvers of Ax = b. The enumeration gives
 * symbolic names for the type  of preconditioning to be used.
 * The function type declarations give the prototypes for the
 * functions to be called within an iterative linear solver, that
 * are responsible for
 *    multiplying A by a given vector v (ATimesFn), and
 *    solving the preconditioner equation Pz = r (PSolveFn).
 * -----------------------------------------------------------------
 */

#ifndef _ITERATIVE_H
#define _ITERATIVE_H

#ifdef __cplusplus  /* wrapper to enable C++ usage */
extern "C" {
#endif

#include <sundials/sundials_nvector.h>


/*
 * -----------------------------------------------------------------
 * enum : types of preconditioning                                
 * -----------------------------------------------------------------
 * PREC_NONE  : The iterative linear solver should not use             
 *              preconditioning.                                       
 *                                                                
 * PREC_LEFT  : The iterative linear solver uses preconditioning on    
 *              the left only.                                         
 *                                                                
 * PREC_RIGHT : The iterative linear solver uses preconditioning on    
 *              the right only.                                        
 *                                                                
 * PREC_BOTH  : The iterative linear solver uses preconditioning on    
 *              both the left and the right.                           
 * -----------------------------------------------------------------
 */

enum { PREC_NONE, PREC_LEFT, PREC_RIGHT, PREC_BOTH };

/*
 * -----------------------------------------------------------------
 * enum : types of Gram-Schmidt routines                          
 * -----------------------------------------------------------------
 * MODIFIED_GS  : The iterative solver uses the modified          
 *                Gram-Schmidt routine ModifiedGS listed in this  
 *                file.                                           
 *                                                                
 * CLASSICAL_GS : The iterative solver uses the classical         
 *                Gram-Schmidt routine ClassicalGS listed in this 
 *                file.                                           
 * -----------------------------------------------------------------
 */

enum { MODIFIED_GS = 1, CLASSICAL_GS = 2 };

/*
 * -----------------------------------------------------------------
 * Type: ATimesFn                                                 
 * -----------------------------------------------------------------
 * An ATimesFn multiplies Av and stores the result in z. The      
 * caller is responsible for allocating memory for the z vector.  
 * The parameter A_data is a pointer to any information about A   
 * which the function needs in order to do its job. The vector v  
 * is unchanged. An ATimesFn returns 0 if successful and a        
 * non-zero value if unsuccessful.                                
 * -----------------------------------------------------------------
 */

typedef int (*ATimesFn)(void *A_data, N_Vector v, N_Vector z);

/*
 * -----------------------------------------------------------------
 * Type: PSolveFn                                                 
 * -----------------------------------------------------------------
 * A PSolveFn solves the preconditioner equation Pz = r for the   
 * vector z. The caller is responsible for allocating memory for  
 * the z vector. The parameter P_data is a pointer to any         
 * information about P which the function needs in order to do    
 * its job. The parameter lr is input, and indicates whether P    
 * is to be taken as the left preconditioner or the right         
 * preconditioner: lr = 1 for left and lr = 2 for right.          
 * If preconditioning is on one side only, lr can be ignored.     
 * The vector r is unchanged.                                     
 * A PSolveFn returns 0 if successful and a non-zero value if     
 * unsuccessful.  On a failure, a negative return value indicates 
 * an unrecoverable condition, while a positive value indicates   
 * a recoverable one, in which the calling routine may reattempt  
 * the solution after updating preconditioner data.               
 * -----------------------------------------------------------------
 */

typedef int (*PSolveFn)(void *P_data, N_Vector r, N_Vector z, int lr);

/*
 * -----------------------------------------------------------------
 * Function: ModifiedGS                                           
 * -----------------------------------------------------------------
 * ModifiedGS performs a modified Gram-Schmidt orthogonalization  
 * of the N_Vector v[k] against the p unit N_Vectors at           
 * v[k-1], v[k-2], ..., v[k-p].                                   
 *                                                                
 * v is an array of (k+1) N_Vectors v[i], i=0, 1, ..., k.         
 * v[k-1], v[k-2], ..., v[k-p] are assumed to have L2-norm        
 * equal to 1.                                                    
 *                                                                
 * h is the output k by k Hessenberg matrix of inner products.    
 * This matrix must be allocated row-wise so that the (i,j)th     
 * entry is h[i][j]. The inner products (v[i],v[k]),              
 * i=i0, i0+1, ..., k-1, are stored at h[i][k-1]. Here            
 * i0=MAX(0,k-p).                                                 
 *                                                                
 * k is the index of the vector in the v array that needs to be   
 * orthogonalized against previous vectors in the v array.        
 *                                                                
 * p is the number of previous vectors in the v array against     
 * which v[k] is to be orthogonalized.                            
 *                                                                
 * new_vk_norm is a pointer to memory allocated by the caller to  
 * hold the Euclidean norm of the orthogonalized vector v[k].     
 *                                                                
 * If (k-p) < 0, then ModifiedGS uses p=k. The orthogonalized     
 * v[k] is NOT normalized and is stored over the old v[k]. Once   
 * the orthogonalization has been performed, the Euclidean norm   
 * of v[k] is stored in (*new_vk_norm).                           
 *                                                                
 * ModifiedGS returns 0 to indicate success. It cannot fail.      
 * -----------------------------------------------------------------
 */                                                                

int ModifiedGS(N_Vector *v, realtype **h, int k, int p, 
               realtype *new_vk_norm);

/*
 * -----------------------------------------------------------------
 * Function: ClassicalGS                                          
 * -----------------------------------------------------------------
 * ClassicalGS performs a classical Gram-Schmidt                  
 * orthogonalization of the N_Vector v[k] against the p unit      
 * N_Vectors at v[k-1], v[k-2], ..., v[k-p]. The parameters v, h, 
 * k, p, and new_vk_norm are as described in the documentation    
 * for ModifiedGS.                                                
 *                                                                
 * temp is an N_Vector which can be used as workspace by the      
 * ClassicalGS routine.                                           
 *                                                                
 * s is a length k array of realtype which can be used as         
 * workspace by the ClassicalGS routine.                          
 *
 * ClassicalGS returns 0 to indicate success. It cannot fail.     
 * -----------------------------------------------------------------
 */

int ClassicalGS(N_Vector *v, realtype **h, int k, int p, 
                realtype *new_vk_norm, N_Vector temp, realtype *s);

/*
 * -----------------------------------------------------------------
 * Function: QRfact                                               
 * -----------------------------------------------------------------
 * QRfact performs a QR factorization of the Hessenberg matrix H. 
 *                                                                
 * n is the problem size; the matrix H is (n+1) by n.             
 *                                                                
 * h is the (n+1) by n Hessenberg matrix H to be factored. It is  
 * stored row-wise.                                               
 *                                                                
 * q is an array of length 2*n containing the Givens rotations    
 * computed by this function. A Givens rotation has the form:     
 * | c  -s |                                                      
 * | s   c |.                                                     
 * The components of the Givens rotations are stored in q as      
 * (c, s, c, s, ..., c, s).                                       
 *                                                                
 * job is a control flag. If job==0, then a new QR factorization  
 * is performed. If job!=0, then it is assumed that the first     
 * n-1 columns of h have already been factored and only the last  
 * column needs to be updated.                                    
 *                                                                
 * QRfact returns 0 if successful. If a zero is encountered on    
 * the diagonal of the triangular factor R, then QRfact returns   
 * the equation number of the zero entry, where the equations are 
 * numbered from 1, not 0. If QRsol is subsequently called in     
 * this situation, it will return an error because it could not   
 * divide by the zero diagonal entry.                             
 * -----------------------------------------------------------------
 */                                                                

int QRfact(int n, realtype **h, realtype *q, int job);

/*                                                                
 * -----------------------------------------------------------------
 * Function: QRsol                                                
 * -----------------------------------------------------------------
 * QRsol solves the linear least squares problem                  
 *                                                                
 * min (b - H*x, b - H*x), x in R^n,                              
 *                                                                
 * where H is a Hessenberg matrix, and b is in R^(n+1).           
 * It uses the QR factors of H computed by QRfact.                
 *                                                                
 * n is the problem size; the matrix H is (n+1) by n.             
 *                                                                
 * h is a matrix (computed by QRfact) containing the upper        
 * triangular factor R of the original Hessenberg matrix H.       
 *                                                                
 * q is an array of length 2*n (computed by QRfact) containing    
 * the Givens rotations used to factor H.                         
 *                                                                
 * b is the (n+1)-vector appearing in the least squares problem   
 * above.                                                         
 *                                                                
 * On return, b contains the solution x of the least squares      
 * problem, if QRsol was successful.                              
 *                                                                
 * QRsol returns a 0 if successful.  Otherwise, a zero was        
 * encountered on the diagonal of the triangular factor R.        
 * In this case, QRsol returns the equation number (numbered      
 * from 1, not 0) of the zero entry.                              
 * -----------------------------------------------------------------
 */                                                                

int QRsol(int n, realtype **h, realtype *q, realtype *b);

#ifdef __cplusplus
}
#endif

#endif
```
