## Supplementary scripts for "A multi-scale *in silico* mouse model for insulin resistance and humanoid type 2 diabetes": sundials_math.h

```
/*
 * -----------------------------------------------------------------
 * $Revision: 749 $
 * $Date: 2011-06-22 14:49:10 +0200 (Wed, 22 Jun 2011) $
 * -----------------------------------------------------------------
 * Programmer(s): Scott D. Cohen, Alan C. Hindmarsh and
 *                Aaron Collier @ LLNL
 * -----------------------------------------------------------------
 * Copyright (c) 2002, The Regents of the University of California.
 * Produced at the Lawrence Livermore National Laboratory.
 * All rights reserved.
 * For details, see the LICENSE file.
 * -----------------------------------------------------------------
 * This is the header file for a simple C-language math library. The
 * routines listed here work with the type realtype as defined in
 * the header file sundials_types.h.
 * -----------------------------------------------------------------
 */

#ifndef _SUNDIALSMATH_H
#define _SUNDIALSMATH_H

#ifdef __cplusplus  /* wrapper to enable C++ usage */
extern "C" {
#endif

#include <sundials/sundials_types.h>

/*
 * -----------------------------------------------------------------
 * Macros : MIN and MAX
 * -----------------------------------------------------------------
 * MIN(A,B) returns the minimum of A and B
 *
 * MAX(A,B) returns the maximum of A and B
 *
 * SQR(A) returns A^2
 * -----------------------------------------------------------------
 */

#ifndef MIN
#define MIN(A, B) ((A) < (B) ? (A) : (B))
#endif

#ifndef MAX
#define MAX(A, B) ((A) > (B) ? (A) : (B))
#endif

#ifndef SQR
#define SQR(A) ((A)*(A))
#endif

#ifndef ABS
#define ABS RAbs
#endif

#ifndef SQRT
#define SQRT RSqrt
#endif

#ifndef EXP
#define EXP RExp
#endif

/*
 * -----------------------------------------------------------------
 * Function : RPowerI
 * -----------------------------------------------------------------
 * Usage : int exponent;
 *         realtype base, ans;
 *         ans = RPowerI(base,exponent);
 * -----------------------------------------------------------------
 * RPowerI returns the value of base^exponent, where base is of type
 * realtype and exponent is of type int.
 * -----------------------------------------------------------------
 */

realtype RPowerI(realtype base, int exponent);

/*
 * -----------------------------------------------------------------
 * Function : RPowerR
 * -----------------------------------------------------------------
 * Usage : realtype base, exponent, ans;
 *         ans = RPowerR(base,exponent);
 * -----------------------------------------------------------------
 * RPowerR returns the value of base^exponent, where both base and
 * exponent are of type realtype. If base < ZERO, then RPowerR
 * returns ZERO.
 * -----------------------------------------------------------------
 */

realtype RPowerR(realtype base, realtype exponent);

/*
 * -----------------------------------------------------------------
 * Function : RSqrt
 * -----------------------------------------------------------------
 * Usage : realtype sqrt_x;
 *         sqrt_x = RSqrt(x);
 * -----------------------------------------------------------------
 * RSqrt(x) returns the square root of x. If x < ZERO, then RSqrt
 * returns ZERO.
 * -----------------------------------------------------------------
 */

realtype RSqrt(realtype x);

/*
 * -----------------------------------------------------------------
 * Function : RAbs (a.k.a. ABS)
 * -----------------------------------------------------------------
 * Usage : realtype abs_x;
 *         abs_x = RAbs(x);
 * -----------------------------------------------------------------
 * RAbs(x) returns the absolute value of x.
 * -----------------------------------------------------------------
 */

realtype RAbs(realtype x);

/*
 * -----------------------------------------------------------------
 * Function : RExp (a.k.a. EXP)
 * -----------------------------------------------------------------
 * Usage : realtype exp_x;
 *         exp_x = RExp(x);
 * -----------------------------------------------------------------
 * RExp(x) returns e^x (base-e exponential function).
 * -----------------------------------------------------------------
 */

realtype RExp(realtype x);

#ifdef __cplusplus
}
#endif

#endif
```
