## Supplementary scripts for "A multi-scale *in silico* mouse model for insulin resistance and humanoid type 2 diabetes": sundials_nvector.h

```
/*
 * -----------------------------------------------------------------
 * $Revision: 749 $
 * $Date: 2011-06-22 14:49:10 +0200 (Wed, 22 Jun 2011) $
 * ----------------------------------------------------------------- 
 * Programmer(s): Radu Serban and Aaron Collier @ LLNL
 * -----------------------------------------------------------------
 * Copyright (c) 2002, The Regents of the University of California.
 * Produced at the Lawrence Livermore National Laboratory.
 * All rights reserved.
 * For details, see the LICENSE file.
 * -----------------------------------------------------------------
 * This is the header file for a generic NVECTOR package.
 * It defines the N_Vector structure (_generic_N_Vector) which
 * contains the following fields:
 *   - an implementation-dependent 'content' field which contains
 *     the description and actual data of the vector
 *   - an 'ops' filed which contains a structure listing operations
 *     acting on such vectors
 *
 * Part I of this file contains type declarations for the
 * _generic_N_Vector and _generic_N_Vector_Ops structures, as well
 * as references to pointers to such structures (N_Vector).
 *
 * Part II of this file contains the prototypes for the vector
 * functions which operate on N_Vector.
 *
 * At a minimum, a particular implementation of an NVECTOR must
 * do the following:
 *  - specify the 'content' field of N_Vector,
 *  - implement the operations on those N_Vectors,
 *  - provide a constructor routine for new vectors
 *
 * Additionally, an NVECTOR implementation may provide the following:
 *  - macros to access the underlying N_Vector data
 *  - a constructor for an array of N_Vectors
 *  - a constructor for an empty N_Vector (i.e., a new N_Vector with
 *    a NULL data pointer).
 *  - a routine to print the content of an N_Vector
 * -----------------------------------------------------------------
 */

#ifndef _NVECTOR_H
#define _NVECTOR_H

#ifdef __cplusplus  /* wrapper to enable C++ usage */
extern "C" {
#endif

#include <sundials/sundials_types.h>

/*
 * -----------------------------------------------------------------
 * Generic definition of N_Vector
 * -----------------------------------------------------------------
 */

/* Forward reference for pointer to N_Vector_Ops object */
typedef struct _generic_N_Vector_Ops *N_Vector_Ops;

/* Forward reference for pointer to N_Vector object */
typedef struct _generic_N_Vector *N_Vector;

/* Define array of N_Vectors */
typedef N_Vector *N_Vector_S;

/* Structure containing function pointers to vector operations  */  
struct _generic_N_Vector_Ops {
  N_Vector    (*nvclone)(N_Vector);
  N_Vector    (*nvcloneempty)(N_Vector);
  void        (*nvdestroy)(N_Vector);
  void        (*nvspace)(N_Vector, long int *, long int *);
  realtype*   (*nvgetarraypointer)(N_Vector);
  void        (*nvsetarraypointer)(realtype *, N_Vector);
  void        (*nvlinearsum)(realtype, N_Vector, realtype, N_Vector, N_Vector); 
  void        (*nvconst)(realtype, N_Vector);
  void        (*nvprod)(N_Vector, N_Vector, N_Vector);
  void        (*nvdiv)(N_Vector, N_Vector, N_Vector);
  void        (*nvscale)(realtype, N_Vector, N_Vector);
  void        (*nvabs)(N_Vector, N_Vector);
  void        (*nvinv)(N_Vector, N_Vector);
  void        (*nvaddconst)(N_Vector, realtype, N_Vector);
  realtype    (*nvdotprod)(N_Vector, N_Vector);
  realtype    (*nvmaxnorm)(N_Vector);
  realtype    (*nvwrmsnorm)(N_Vector, N_Vector);
  realtype    (*nvwrmsnormmask)(N_Vector, N_Vector, N_Vector);
  realtype    (*nvmin)(N_Vector);
  realtype    (*nvwl2norm)(N_Vector, N_Vector);
  realtype    (*nvl1norm)(N_Vector);
  void        (*nvcompare)(realtype, N_Vector, N_Vector);
  booleantype (*nvinvtest)(N_Vector, N_Vector);
  booleantype (*nvconstrmask)(N_Vector, N_Vector, N_Vector);
  realtype    (*nvminquotient)(N_Vector, N_Vector);
};

/*
 * -----------------------------------------------------------------
 * A vector is a structure with an implementation-dependent
 * 'content' field, and a pointer to a structure of vector
 * operations corresponding to that implementation.
 * -----------------------------------------------------------------
 */

struct _generic_N_Vector {
  void *content;
  struct _generic_N_Vector_Ops *ops;
};
  
/*
 * -----------------------------------------------------------------
 * Functions exported by NVECTOR module
 * -----------------------------------------------------------------
 */

/*
 * -----------------------------------------------------------------
 * N_VClone
 *   Creates a new vector of the same type as an existing vector.
 *   It does not copy the vector, but rather allocates storage for
 *   the new vector.
 *
 * N_VCloneEmpty
 *   Creates a new vector of the same type as an existing vector,
 *   but does not allocate storage.
 *
 * N_VDestroy
 *   Destroys a vector created with N_VClone.
 *
 * N_VSpace
 *   Returns space requirements for one N_Vector (type 'realtype' in
 *   lrw and type 'long int' in liw).
 *
 * N_VGetArrayPointer
 *   Returns a pointer to the data component of the given N_Vector.
 *   NOTE: This function assumes that the internal data is stored
 *   as a contiguous 'realtype' array. This routine is only used in
 *   the solver-specific interfaces to the dense and banded linear
 *   solvers, as well as the interfaces to  the banded preconditioners
 *   distributed with SUNDIALS.
 *   
 * N_VSetArrayPointer
 *   Overwrites the data field in the given N_Vector with a user-supplied
 *   array of type 'realtype'.
 *   NOTE: This function assumes that the internal data is stored
 *   as a contiguous 'realtype' array. This routine is only used in
 *   the interfaces to the dense linear solver.
 *
 * N_VLinearSum
 *   Performs the operation z = a*x + b*y
 *
 * N_VConst
 *   Performs the operation z[i] = c for i = 0, 1, ..., N-1
 *
 * N_VProd
 *   Performs the operation z[i] = x[i]*y[i] for i = 0, 1, ..., N-1
 *
 * N_VDiv
 *   Performs the operation z[i] = x[i]/y[i] for i = 0, 1, ..., N-1
 *
 * N_VScale
 *   Performs the operation z = c*x
 *
 * N_VAbs
 *   Performs the operation z[i] = |x[i]| for i = 0, 1, ..., N-1
 *
 * N_VInv
 *   Performs the operation z[i] = 1/x[i] for i = 0, 1, ..., N-1
 *   This routine does not check for division by 0. It should be
 *   called only with an N_Vector x which is guaranteed to have
 *   all non-zero components.
 *
 * N_VAddConst
 *   Performs the operation z[i] = x[i] + b   for i = 0, 1, ..., N-1
 *
 * N_VDotProd
 *   Returns the dot product of two vectors:
 *         sum (i = 0 to N-1) {x[i]*y[i]}
 *
 * N_VMaxNorm
 *   Returns the maximum norm of x:
 *         max (i = 0 to N-1) ABS(x[i])
 *
 * N_VWrmsNorm
 *   Returns the weighted root mean square norm of x with weight 
 *   vector w:
 *         sqrt [(sum (i = 0 to N-1) {(x[i]*w[i])^2})/N]
 *
 * N_VWrmsNormMask
 *   Returns the weighted root mean square norm of x with weight
 *   vector w, masked by the elements of id:
 *         sqrt [(sum (i = 0 to N-1) {(x[i]*w[i]*msk[i])^2})/N]
 *   where msk[i] = 1.0 if id[i] > 0 and
 *         msk[i] = 0.0 if id[i] < 0
 *
 * N_VMin
 *   Returns the smallest element of x:
 *         min (i = 0 to N-1) x[i]
 *
 * N_VWL2Norm
 *   Returns the weighted Euclidean L2 norm of x with weight 
 *   vector w:
 *         sqrt [(sum (i = 0 to N-1) {(x[i]*w[i])^2})]
 *
 * N_VL1Norm
 *   Returns the L1 norm of x:
 *         sum (i = 0 to N-1) {ABS(x[i])}
 *
 * N_VCompare
 *   Performs the operation
 *          z[i] = 1.0 if ABS(x[i]) >= c   i = 0, 1, ..., N-1
 *                 0.0 otherwise
 *
 * N_VInvTest
 *   Performs the operation z[i] = 1/x[i] with a test for 
 *   x[i] == 0.0 before inverting x[i].
 *   This routine returns TRUE if all components of x are non-zero 
 *   (successful inversion) and returns FALSE otherwise.
 *
 * N_VConstrMask
 *   Performs the operation : 
 *       m[i] = 1.0 if constraint test fails for x[i]
 *       m[i] = 0.0 if constraint test passes for x[i]
 *   where the constraint tests are as follows:
 *      If c[i] = +2.0, then x[i] must be >  0.0.
 *      If c[i] = +1.0, then x[i] must be >= 0.0.
 *      If c[i] = -1.0, then x[i] must be <= 0.0.
 *      If c[i] = -2.0, then x[i] must be <  0.0.
 *   This routine returns a boolean FALSE if any element failed
 *   the constraint test, TRUE if all passed. It also sets a
 *   mask vector m, with elements equal to 1.0 where the
 *   corresponding constraint test failed, and equal to 0.0
 *   where the constraint test passed.
 *   This routine is specialized in that it is used only for
 *   constraint checking.
 *
 * N_VMinQuotient
 *   Performs the operation : 
 *       minq  = min ( num[i]/denom[i]) over all i such that   
 *       denom[i] != 0.
 *   This routine returns the minimum of the quotients obtained
 *   by term-wise dividing num[i] by denom[i]. A zero element
 *   in denom will be skipped. If no such quotients are found,
 *   then the large value BIG_REAL is returned.
 *
 * -----------------------------------------------------------------
 *
 * The following table lists the vector functions used by
 * different modules in SUNDIALS. The symbols in the table
 * have the following meaning:
 * S    -  called by the solver;
 * D    -  called by the dense linear solver module
 * B    -  called by the band linear solver module
 * Di   -  called by the diagonal linear solver module
 * I    -  called by the iterative linear solver module
 * BP   -  called by the band preconditioner module
 * BBDP -  called by the band-block diagonal preconditioner module
 * F    -  called by the Fortran-to-C interface
 *
 *                  ------------------------------------------------
 *                                         MODULES                  
 * NVECTOR          ------------------------------------------------
 * FUNCTIONS          CVODE/CVODES          IDA             KINSOL    
 * -----------------------------------------------------------------
 * N_VClone           S Di I                S I BBDP        S I BBDP
 * -----------------------------------------------------------------
 * N_VCloneEmpty      F                     F               F
 * -----------------------------------------------------------------
 * N_VDestroy         S Di I                S I BBDP        S I BBDP
 * -----------------------------------------------------------------
 * N_VSpace           S                     S               S         
 * -----------------------------------------------------------------
 * N_VGetArrayPointer D B BP BBDP F         D B BBDP        BBDP F     
 * -----------------------------------------------------------------
 * N_VSetArrayPointer D F                   D               F
 * -----------------------------------------------------------------
 * N_VLinearSum       S D Di I              S D I           S I       
 * -----------------------------------------------------------------
 * N_VConst           S I                   S I             I       
 * -----------------------------------------------------------------
 * N_VProd            S Di I                S I             S I       
 * -----------------------------------------------------------------
 * N_VDiv             S Di I                S I             S I
 * -----------------------------------------------------------------
 * N_VScale           S D B Di I BP BBDP    S D B I BBDP    S I BBDP  
 * -----------------------------------------------------------------
 * N_VAbs             S                     S               S         
 * -----------------------------------------------------------------
 * N_VInv             S Di                  S               S         
 * -----------------------------------------------------------------
 * N_VAddConst        S Di                  S                        
 * -----------------------------------------------------------------
 * N_VDotProd         I                     I               I         
 * -----------------------------------------------------------------
 * N_VMaxNorm         S                     S               S         
 * -----------------------------------------------------------------
 * N_VWrmsNorm        S D B I BP BBDP       S                         
 * -----------------------------------------------------------------
 * N_VWrmsNormMask                          S                         
 * -----------------------------------------------------------------
 * N_VMin             S                     S               S         
 * -----------------------------------------------------------------
 * N_VWL2Norm                                               S I       
 * -----------------------------------------------------------------
 * N_VL1Norm                                                I
 * -----------------------------------------------------------------
 * N_VCompare         Di                    S                         
 * -----------------------------------------------------------------
 * N_VInvTest         Di                                              
 * -----------------------------------------------------------------
 * N_VConstrMask                            S               S         
 * -----------------------------------------------------------------
 * N_VMinQuotient                           S               S         
 * -----------------------------------------------------------------
 */
  
N_Vector N_VClone(N_Vector w);
N_Vector N_VCloneEmpty(N_Vector w);
void N_VDestroy(N_Vector v);
void N_VSpace(N_Vector v, long int *lrw, long int *liw);
realtype *N_VGetArrayPointer(N_Vector v);
void N_VSetArrayPointer(realtype *v_data, N_Vector v);
void N_VLinearSum(realtype a, N_Vector x, realtype b, N_Vector y, N_Vector z);
void N_VConst(realtype c, N_Vector z);
void N_VProd(N_Vector x, N_Vector y, N_Vector z);
void N_VDiv(N_Vector x, N_Vector y, N_Vector z);
void N_VScale(realtype c, N_Vector x, N_Vector z);
void N_VAbs(N_Vector x, N_Vector z);
void N_VInv(N_Vector x, N_Vector z);
void N_VAddConst(N_Vector x, realtype b, N_Vector z);
realtype N_VDotProd(N_Vector x, N_Vector y);
realtype N_VMaxNorm(N_Vector x);
realtype N_VWrmsNorm(N_Vector x, N_Vector w);
realtype N_VWrmsNormMask(N_Vector x, N_Vector w, N_Vector id);
realtype N_VMin(N_Vector x);
realtype N_VWL2Norm(N_Vector x, N_Vector w);
realtype N_VL1Norm(N_Vector x);
void N_VCompare(realtype c, N_Vector x, N_Vector z);
booleantype N_VInvTest(N_Vector x, N_Vector z);
booleantype N_VConstrMask(N_Vector c, N_Vector x, N_Vector m);
realtype N_VMinQuotient(N_Vector num, N_Vector denom);

/*
 * -----------------------------------------------------------------
 * Additional functions exported by NVECTOR module
 * -----------------------------------------------------------------
 */

/*
 * -----------------------------------------------------------------
 * N_VCloneEmptyVectorArray
 *   Creates (by cloning 'w') an array of 'count' empty N_Vectors 
 *
 * N_VCloneVectorArray
 *   Creates (by cloning 'w') an array of 'count' N_Vectors 
 *
 * N_VDestroyVectorArray
 *   Frees memory for an array of 'count' N_Vectors that was
 *   created by a call to N_VCloneVectorArray
 *
 * These functions are used by the SPGMR iterative linear solver 
 * module and by the CVODES and IDAS solvers.
 * -----------------------------------------------------------------
 */

N_Vector *N_VCloneEmptyVectorArray(int count, N_Vector w);
N_Vector *N_VCloneVectorArray(int count, N_Vector w);
void N_VDestroyVectorArray(N_Vector *vs, int count);

#ifdef __cplusplus
}
#endif

#endif
```
