## Supplementary scripts for "A multi-scale *in silico* mouse model for insulin resistance and humanoid type 2 diabetes": sundials_smalldense.h

```
/*
 * -----------------------------------------------------------------
 * $Revision: 749 $
 * $Date: 2011-06-22 14:49:10 +0200 (Wed, 22 Jun 2011) $
 * -----------------------------------------------------------------
 * Programmer: Radu Serban @ LLNL
 * -----------------------------------------------------------------
 * Copyright (c) 2006, The Regents of the University of California.
 * Produced at the Lawrence Livermore National Laboratory.
 * All rights reserved.
 * For details, see the LICENSE file.
 * -----------------------------------------------------------------
 * This is the header file for a generic DENSE linear solver
 * package, intended for small m by n dense matrices, with m<=n.
 * These routines use the type realtype** for dense matrix arguments.
 * -----------------------------------------------------------------
 */

#ifndef _SMALLDENSE_H
#define _SMALLDENSE_H

#ifdef __cplusplus  /* wrapper to enable C++ usage */
extern "C" {
#endif

#include <sundials/sundials_types.h>

  /*
   * -----------------------------------------------------------------
   * Function : denalloc
   * -----------------------------------------------------------------
   * Usage : realtype **a;
   *         a = denalloc(m, n);
   *         if (a == NULL) ... memory request failed
   * -----------------------------------------------------------------
   * denalloc(m, n) allocates storage for an m by n dense matrix.
   * It returns a pointer to the newly allocated storage if successful. 
   * If the memory request cannot be satisfied, then denalloc returns 
   * NULL. The underlying type of the dense matrix returned is a double
   * pointer to realtype. If we allocate a dense matrix a by 
   * a = denalloc(m, n), then a[j][i] references the (i,j)-th element 
   * of the matrix a, 0 <= i < m, 0 <= j < n,  and a[j] is a pointer 
   * to the first element in the jth column of a. The location a[0] 
   * contains a pointer to m*n contiguous locations which contain the 
   * elements of a.
   * -----------------------------------------------------------------
   */

  realtype **denalloc(long int m, long int n);

  /*
   * -----------------------------------------------------------------
   * Function : denallocpiv
   * -----------------------------------------------------------------
   * Usage : long int *pivot;
   *         pivot = denallocpiv(n);
   *         if (pivot == NULL) ... memory request failed
   * -----------------------------------------------------------------
   * denallocpiv(n) allocates an array of n long int. It returns
   * a pointer to the first element in the array if successful.
   * It returns NULL if the memory request could not be satisfied.
   * -----------------------------------------------------------------
   */

  long int *denallocpiv(long int n);

  /*
   * -----------------------------------------------------------------
   * Function : denGETRF
   * -----------------------------------------------------------------
   * Usage : long int ier;
   *         ier = denGETRF(a,m,n,p);
   *         if (ier > 0) ... zero element encountered during
   *                          the factorization
   * -----------------------------------------------------------------
   * denGETRF(a,m,n,p) factors the m by n dense matrix a, m>=n.
   * It overwrites the elements of a with its LU factors and keeps 
   * track of the pivot rows chosen in the pivot array p.
   *
   * A successful LU factorization leaves the matrix a and the
   * pivot array p with the following information:
   *
   * (1) p[k] contains the row number of the pivot element chosen
   *     at the beginning of elimination step k, k=0, 1, ..., n-1.
   *
   * (2) If the unique LU factorization of a is given by Pa = LU,
   *     where P is a permutation matrix, L is a lower trapezoidal
   *     matrix with all 1.0 on the diagonal, and U is an upper
   *     triangular matrix, then the upper triangular part of a
   *     (including its diagonal) contains U and the strictly lower
   *     trapezoidal part of a contains the multipliers, I-L.
   *
   * Note that for square matrices (m=n), L is unit lower triangular.
   *
   * denGETRF returns 0 if successful. Otherwise it encountered a
   * zero diagonal element during the factorization. In this case
   * it returns the column index (numbered from one) at which it
   * encountered the zero.
   * -----------------------------------------------------------------
   */

  long int denGETRF(realtype **a, long int m, long int n, long int *p);

  /*
   * -----------------------------------------------------------------
   * Function : denGETRS
   * -----------------------------------------------------------------
   * Usage : realtype *b;
   *         ier = denGETRF(a,n,n,p);
   *         if (ier == 0) denGETRS(a,n,p,b);
   * -----------------------------------------------------------------
   * denGETRS(a,n,p,b) solves the n by n linear system a*x = b.
   * It assumes that a has been LU factored and the pivot array p has
   * been set by a successful call to denGETRF(a,n,n,p). 
   * denGETRS does not check whether a is square!
   * The solution x is written into the b array.
   * -----------------------------------------------------------------
   */

  void denGETRS(realtype **a, long int n, long int *p, realtype *b);

  /*
   * -----------------------------------------------------------------
   * Function : denzero
   * -----------------------------------------------------------------
   * Usage : denzero(a,m,n);
   * -----------------------------------------------------------------
   * denzero(a,m,n) sets all the elements of the m by n dense matrix
   * a to be 0.0.
   * -----------------------------------------------------------------
   */

  void denzero(realtype **a, long int m, long int n);

  /*
   * -----------------------------------------------------------------
   * Function : dencopy
   * -----------------------------------------------------------------
   * Usage : dencopy(a,b,m,n);
   * -----------------------------------------------------------------
   * dencopy(a,b,n) copies the m by n dense matrix a into the
   * m by n dense matrix b.
   * -----------------------------------------------------------------
   */

  void dencopy(realtype **a, realtype **b, long int m, long int n);

  /*
   * -----------------------------------------------------------------
   * Function : denscale
   * -----------------------------------------------------------------
   * Usage : denscale(c,a,m,n);
   * -----------------------------------------------------------------
   * denscale(c,a,m,n) scales every element in the m by n dense
   * matrix a by c.
   * -----------------------------------------------------------------
   */

  void denscale(realtype c, realtype **a, long int m, long int n);

  /*
   * -----------------------------------------------------------------
   * Function : denaddI
   * -----------------------------------------------------------------
   * Usage : denaddI(a,n);
   * -----------------------------------------------------------------
   * denaddI(a,n) increments the diagonal elements of the dense 
   * m by n matrix a by 1.0. (a_ii <= a_ii + 1, i=1,2,..n-1.)
   * denaddI is typically used with square matrices.
   * denaddI does NOT check for m >= n! Therefore, a segmentation 
   * fault will occur if m<n!
   * -----------------------------------------------------------------
   */

  void denaddI(realtype **a, long int n);

  /*
   * -----------------------------------------------------------------
   * Function : denfreepiv
   * -----------------------------------------------------------------
   * Usage : denfreepiv(p);
   * -----------------------------------------------------------------
   * denfreepiv(p) frees the pivot array p allocated by
   * denallocpiv.
   * -----------------------------------------------------------------
   */

  void denfreepiv(long int *p);

  /*
   * -----------------------------------------------------------------
   * Function : denfree
   * -----------------------------------------------------------------
   * Usage : denfree(a);
   * -----------------------------------------------------------------
   * denfree(a) frees the dense matrix a allocated by denalloc.
   * -----------------------------------------------------------------
   */

  void denfree(realtype **a);

  /*
   * -----------------------------------------------------------------
   * Function : denprint
   * -----------------------------------------------------------------
   * Usage : denprint(a,m,n);
   * -----------------------------------------------------------------
   * denprint(a,m,n) prints the m by n dense matrix a to standard
   * output as it would normally appear on paper. It is intended as
   * a debugging tool with small values of m and n. The elements are
   * printed using the %g/%lg/%Lg option. A blank line is printed
   * before and after the matrix.
   * -----------------------------------------------------------------
   */

  void denprint(realtype **a, long int m, long int n);

#ifdef __cplusplus
}
#endif

#endif
```
