## Supplementary scripts for "A multi-scale *in silico* mouse model for insulin resistance and humanoid type 2 diabetes": sundials_spbcgs.h

```
/*
 * -----------------------------------------------------------------
 * $Revision: 749 $
 * $Date: 2011-06-22 14:49:10 +0200 (Wed, 22 Jun 2011) $
 * -----------------------------------------------------------------
 * Programmer(s): Peter Brown and Aaron Collier @ LLNL
 * -----------------------------------------------------------------
 * Copyright (c) 2004, The Regents of the University of California.
 * Produced at the Lawrence Livermore National Laboratory.
 * All rights reserved.
 * For details, see the LICENSE file.
 * -----------------------------------------------------------------
 * This is the header file for the implementation of the scaled,
 * preconditioned Bi-CGSTAB (SPBCG) iterative linear solver.
 * -----------------------------------------------------------------
 */

#ifndef _SPBCG_H
#define _SPBCG_H

#ifdef __cplusplus  /* wrapper to enable C++ usage */
extern "C" {
#endif

#include <sundials/sundials_iterative.h>

/*
 * -----------------------------------------------------------------
 * Types: struct SpbcgMemRec and struct *SpbcgMem
 * -----------------------------------------------------------------
 * A variable declaration of type struct *SpbcgMem denotes a pointer
 * to a data structure of type struct SpbcgMemRec. The SpbcgMemRec
 * structure contains numerous fields that must be accessed by the
 * SPBCG linear solver module.
 *
 *  l_max  maximum Krylov subspace dimension that SpbcgSolve will
 *         be permitted to use
 *
 *  r  vector (type N_Vector) which holds the scaled, preconditioned
 *     linear system residual
 *
 *  r_star  vector (type N_Vector) which holds the initial scaled,
 *          preconditioned linear system residual
 *
 *  p, q, u and Ap  vectors (type N_Vector) used for workspace by
 *                  the SPBCG algorithm
 *
 *  vtemp  scratch vector (type N_Vector) used as temporary vector
 *         storage
 * -----------------------------------------------------------------
 */

typedef struct {

  int l_max;

  N_Vector r_star;
  N_Vector r;
  N_Vector p;
  N_Vector q;
  N_Vector u;
  N_Vector Ap;
  N_Vector vtemp;

} SpbcgMemRec, *SpbcgMem;

/*
 * -----------------------------------------------------------------
 * Function : SpbcgMalloc
 * -----------------------------------------------------------------
 * SpbcgMalloc allocates additional memory needed by the SPBCG
 * linear solver module.
 *
 *  l_max  maximum Krylov subspace dimension that SpbcgSolve will
 *         be permitted to use
 *
 *  vec_tmpl  implementation-specific template vector (type N_Vector)
 *            (created using either N_VNew_Serial or N_VNew_Parallel)
 *
 * If successful, SpbcgMalloc returns a non-NULL memory pointer. If
 * an error occurs, then a NULL pointer is returned.
 * -----------------------------------------------------------------
 */

SpbcgMem SpbcgMalloc(int l_max, N_Vector vec_tmpl);

/*
 * -----------------------------------------------------------------
 * Function : SpbcgSolve
 * -----------------------------------------------------------------
 * SpbcgSolve solves the linear system Ax = b by means of a scaled
 * preconditioned Bi-CGSTAB (SPBCG) iterative method.
 *
 *  mem  pointer to an internal memory block allocated during a
 *       prior call to SpbcgMalloc
 *
 *  A_data  pointer to a data structure containing information
 *          about the coefficient matrix A (passed to user-supplied
 *          function referenced by atimes (function pointer))
 *
 *  x  vector (type N_Vector) containing initial guess x_0 upon
 *     entry, but which upon return contains an approximate solution
 *     of the linear system Ax = b (solution only valid if return
 *     value is either SPBCG_SUCCESS or SPBCG_RES_REDUCED)
 *
 *  b  vector (type N_Vector) set to the right-hand side vector b
 *     of the linear system (undisturbed by function)
 *
 *  pretype  variable (type int) indicating the type of
 *           preconditioning to be used (see sundials_iterative.h)
 *
 *  delta  tolerance on the L2 norm of the scaled, preconditioned
 *         residual (if return value == SPBCG_SUCCESS, then
 *         ||sb*P1_inv*(b-Ax)||_L2 <= delta)
 *
 *  P_data  pointer to a data structure containing preconditioner
 *          information (passed to user-supplied function referenced
 *          by psolve (function pointer))
 *
 *  sx  vector (type N_Vector) containing positive scaling factors
 *      for x (pass sx == NULL if scaling NOT required)
 *
 *  sb  vector (type N_Vector) containing positive scaling factors
 *      for b (pass sb == NULL if scaling NOT required)
 *
 *  atimes  user-supplied routine responsible for computing the
 *          matrix-vector product Ax (see sundials_iterative.h)
 *
 *  psolve  user-supplied routine responsible for solving the
 *          preconditioned linear system Pz = r (ignored if
 *          pretype == PREC_NONE) (see sundials_iterative.h)
 *
 *  res_norm  pointer (type realtype*) to the L2 norm of the
 *            scaled, preconditioned residual (if return value
 *            is either SPBCG_SUCCESS or SPBCG_RES_REDUCED, then
 *            *res_norm = ||sb*P1_inv*(b-Ax)||_L2, where x is
 *            the computed approximate solution, sb is the diagonal
 *            scaling matrix for the right-hand side b, and P1_inv
 *            is the inverse of the left-preconditioner matrix)
 *
 *  nli  pointer (type int*) to the total number of linear
 *       iterations performed
 *
 *  nps  pointer (type int*) to the total number of calls made
 *       to the psolve routine
 * -----------------------------------------------------------------
 */

int SpbcgSolve(SpbcgMem mem, void *A_data, N_Vector x, N_Vector b,
               int pretype, realtype delta, void *P_data, N_Vector sx,
               N_Vector sb, ATimesFn atimes, PSolveFn psolve,
               realtype *res_norm, int *nli, int *nps);

/* Return values for SpbcgSolve */

#define SPBCG_SUCCESS            0  /* SPBCG algorithm converged          */
#define SPBCG_RES_REDUCED        1  /* SPBCG did NOT converge, but the
				       residual was reduced               */
#define SPBCG_CONV_FAIL          2  /* SPBCG algorithm failed to converge */
#define SPBCG_PSOLVE_FAIL_REC    3  /* psolve failed recoverably          */
#define SPBCG_ATIMES_FAIL_REC    4  /* atimes failed recoverably          */
#define SPBCG_PSET_FAIL_REC      5  /* pset faild recoverably             */

#define SPBCG_MEM_NULL          -1  /* mem argument is NULL               */
#define SPBCG_ATIMES_FAIL_UNREC -2  /* atimes returned failure flag       */
#define SPBCG_PSOLVE_FAIL_UNREC -3  /* psolve failed unrecoverably        */
#define SPBCG_PSET_FAIL_UNREC   -4  /* pset failed unrecoverably          */

/*
 * -----------------------------------------------------------------
 * Function : SpbcgFree
 * -----------------------------------------------------------------
 * SpbcgFree frees the memory allocated by a call to SpbcgMalloc.
 * It is illegal to use the pointer mem after a call to SpbcgFree.
 * -----------------------------------------------------------------
 */

void SpbcgFree(SpbcgMem mem);

/*
 * -----------------------------------------------------------------
 * Macro : SPBCG_VTEMP
 * -----------------------------------------------------------------
 * This macro provides access to the vector r in the
 * memory block of the SPBCG module. The argument mem is the
 * memory pointer returned by SpbcgMalloc, of type SpbcgMem,
 * and the macro value is of type N_Vector.
 *
 * Note: Only used by IDA (r contains P_inverse F if nli_inc == 0).
 * -----------------------------------------------------------------
 */

#define SPBCG_VTEMP(mem) (mem->r)

#ifdef __cplusplus
}
#endif

#endif
```
