## Supplementary scripts for "A multi-scale *in silico* mouse model for insulin resistance and humanoid type 2 diabetes": sundials_spgmr.h

```
/*
 * -----------------------------------------------------------------
 * $Revision: 749 $
 * $Date: 2011-06-22 14:49:10 +0200 (Wed, 22 Jun 2011) $
 * ----------------------------------------------------------------- 
 * Programmer(s): Scott D. Cohen, Alan C. Hindmarsh and
 *                Radu Serban @ LLNL
 * -----------------------------------------------------------------
 * Copyright (c) 2002, The Regents of the University of California.
 * Produced at the Lawrence Livermore National Laboratory.
 * All rights reserved.
 * For details, see the LICENSE file.
 * -----------------------------------------------------------------
 * This is the header file for the implementation of SPGMR Krylov
 * iterative linear solver.  The SPGMR algorithm is based on the
 * Scaled Preconditioned GMRES (Generalized Minimal Residual)
 * method.
 *
 * The SPGMR algorithm solves a linear system A x = b.
 * Preconditioning is allowed on the left, right, or both.
 * Scaling is allowed on both sides, and restarts are also allowed.
 * We denote the preconditioner and scaling matrices as follows:
 *   P1 = left preconditioner
 *   P2 = right preconditioner
 *   S1 = diagonal matrix of scale factors for P1-inverse b
 *   S2 = diagonal matrix of scale factors for P2 x
 * The matrices A, P1, and P2 are not required explicitly; only
 * routines that provide A, P1-inverse, and P2-inverse as
 * operators are required.
 *
 * In this notation, SPGMR applies the underlying GMRES method to
 * the equivalent transformed system
 *   Abar xbar = bbar , where
 *   Abar = S1 (P1-inverse) A (P2-inverse) (S2-inverse) ,
 *   bbar = S1 (P1-inverse) b , and   xbar = S2 P2 x .
 *
 * The scaling matrices must be chosen so that vectors S1
 * P1-inverse b and S2 P2 x have dimensionless components.
 * If preconditioning is done on the left only (P2 = I), by a
 * matrix P, then S2 must be a scaling for x, while S1 is a
 * scaling for P-inverse b, and so may also be taken as a scaling
 * for x.  Similarly, if preconditioning is done on the right only
 * (P1 = I, P2 = P), then S1 must be a scaling for b, while S2 is
 * a scaling for P x, and may also be taken as a scaling for b.
 *
 * The stopping test for the SPGMR iterations is on the L2 norm of
 * the scaled preconditioned residual:
 *      || bbar - Abar xbar ||_2  <  delta
 * with an input test constant delta.
 *
 * The usage of this SPGMR solver involves supplying two routines
 * and making three calls.  The user-supplied routines are
 *    atimes (A_data, x, y) to compute y = A x, given x,
 * and
 *    psolve (P_data, x, y, lr)
 *                to solve P1 x = y or P2 x = y for x, given y.
 * The three user calls are:
 *    mem  = SpgmrMalloc(lmax, vec_tmpl);
 *           to initialize memory,
 *    flag = SpgmrSolve(mem,A_data,x,b,...,
 *                      P_data,s1,s2,atimes,psolve,...);
 *           to solve the system, and
 *    SpgmrFree(mem);
 *           to free the memory created by SpgmrMalloc.
 * Complete details for specifying atimes and psolve and for the
 * usage calls are given in the paragraphs below and in iterative.h.
 * -----------------------------------------------------------------
 */

#ifndef _SPGMR_H
#define _SPGMR_H

#ifdef __cplusplus  /* wrapper to enable C++ usage */
extern "C" {
#endif

#include <sundials/sundials_iterative.h>

/*
 * -----------------------------------------------------------------
 * Types: SpgmrMemRec, SpgmrMem
 * -----------------------------------------------------------------
 * SpgmrMem is a pointer to an SpgmrMemRec which contains
 * the memory needed by SpgmrSolve. The SpgmrMalloc routine
 * returns a pointer of type SpgmrMem which should then be passed
 * in subsequent calls to SpgmrSolve. The SpgmrFree routine frees
 * the memory allocated by SpgmrMalloc.
 *
 * l_max is the maximum Krylov dimension that SpgmrSolve will be
 * permitted to use.
 *
 * V is the array of Krylov basis vectors v_1, ..., v_(l_max+1),
 * stored in V[0], ..., V[l_max], where l_max is the second
 * parameter to SpgmrMalloc. Each v_i is a vector of type
 * N_Vector.
 *
 * Hes is the (l_max+1) x l_max Hessenberg matrix. It is stored
 * row-wise so that the (i,j)th element is given by Hes[i][j].
 *
 * givens is a length 2*l_max array which represents the
 * Givens rotation matrices that arise in the algorithm. The
 * Givens rotation matrices F_0, F_1, ..., F_j, where F_i is
 *
 *             1
 *               1
 *                 c_i  -s_i      <--- row i
 *                 s_i   c_i
 *                           1
 *                             1
 *
 * are represented in the givens vector as
 * givens[0]=c_0, givens[1]=s_0, givens[2]=c_1, givens[3]=s_1,
 * ..., givens[2j]=c_j, givens[2j+1]=s_j.
 *
 * xcor is a vector (type N_Vector) which holds the scaled,
 * preconditioned correction to the initial guess.
 *
 * yg is a length (l_max+1) array of realtype used to hold "short"
 * vectors (e.g. y and g).
 *
 * vtemp is a vector (type N_Vector) used as temporary vector
 * storage during calculations.
 * -----------------------------------------------------------------
 */
  
typedef struct _SpgmrMemRec {

  int l_max;

  N_Vector *V;
  realtype **Hes;
  realtype *givens;
  N_Vector xcor;
  realtype *yg;
  N_Vector vtemp;

} SpgmrMemRec, *SpgmrMem;

/*
 * -----------------------------------------------------------------
 * Function : SpgmrMalloc
 * -----------------------------------------------------------------
 * SpgmrMalloc allocates the memory used by SpgmrSolve. It
 * returns a pointer of type SpgmrMem which the user of the
 * SPGMR package should pass to SpgmrSolve. The parameter l_max
 * is the maximum Krylov dimension that SpgmrSolve will be
 * permitted to use. The parameter vec_tmpl is a pointer to an
 * N_Vector used as a template to create new vectors by duplication.
 * This routine returns NULL if there is a memory request failure.
 * -----------------------------------------------------------------
 */

SpgmrMem SpgmrMalloc(int l_max, N_Vector vec_tmpl);

/*
 * -----------------------------------------------------------------
 * Function : SpgmrSolve
 * -----------------------------------------------------------------
 * SpgmrSolve solves the linear system Ax = b using the SPGMR
 * method. The return values are given by the symbolic constants
 * below. The first SpgmrSolve parameter is a pointer to memory
 * allocated by a prior call to SpgmrMalloc.
 *
 * mem is the pointer returned by SpgmrMalloc to the structure
 * containing the memory needed by SpgmrSolve.
 *
 * A_data is a pointer to information about the coefficient
 * matrix A. This pointer is passed to the user-supplied function
 * atimes.
 *
 * x is the initial guess x_0 upon entry and the solution
 * N_Vector upon exit with return value SPGMR_SUCCESS or
 * SPGMR_RES_REDUCED. For all other return values, the output x
 * is undefined.
 *
 * b is the right hand side N_Vector. It is undisturbed by this
 * function.
 *
 * pretype is the type of preconditioning to be used. Its
 * legal possible values are enumerated in iterativ.h. These
 * values are PREC_NONE=0, PREC_LEFT=1, PREC_RIGHT=2, and
 * PREC_BOTH=3.
 *
 * gstype is the type of Gram-Schmidt orthogonalization to be
 * used. Its legal values are enumerated in iterativ.h. These
 * values are MODIFIED_GS=0 and CLASSICAL_GS=1.
 *
 * delta is the tolerance on the L2 norm of the scaled,
 * preconditioned residual. On return with value SPGMR_SUCCESS,
 * this residual satisfies || s1 P1_inv (b - Ax) ||_2 <= delta.
 *
 * max_restarts is the maximum number of times the algorithm is
 * allowed to restart.
 *
 * P_data is a pointer to preconditioner information. This
 * pointer is passed to the user-supplied function psolve.
 *
 * s1 is an N_Vector of positive scale factors for P1-inv b, where
 * P1 is the left preconditioner. (Not tested for positivity.)
 * Pass NULL if no scaling on P1-inv b is required.
 *
 * s2 is an N_Vector of positive scale factors for P2 x, where
 * P2 is the right preconditioner. (Not tested for positivity.)
 * Pass NULL if no scaling on P2 x is required.
 *
 * atimes is the user-supplied function which performs the
 * operation of multiplying A by a given vector. Its description
 * is given in iterative.h.
 *
 * psolve is the user-supplied function which solves a
 * preconditioner system Pz = r, where P is P1 or P2. Its full
 * description is  given in iterativ.h. The psolve function will
 * not be called if pretype is NONE; in that case, the user
 * should pass NULL for psolve.
 *
 * res_norm is a pointer to the L2 norm of the scaled,
 * preconditioned residual. On return with value SPGMR_SUCCESS or
 * SPGMR_RES_REDUCED, (*res_norm) contains the value
 * || s1 P1_inv (b - Ax) ||_2 for the computed solution x.
 * For all other return values, (*res_norm) is undefined. The
 * caller is responsible for allocating the memory (*res_norm)
 * to be filled in by SpgmrSolve.
 *
 * nli is a pointer to the number of linear iterations done in
 * the execution of SpgmrSolve. The caller is responsible for
 * allocating the memory (*nli) to be filled in by SpgmrSolve.
 *
 * nps is a pointer to the number of calls made to psolve during
 * the execution of SpgmrSolve. The caller is responsible for
 * allocating the memory (*nps) to be filled in by SpgmrSolve.
 *
 * Note: Repeated calls can be made to SpgmrSolve with varying
 * input arguments. If, however, the problem size N or the
 * maximum Krylov dimension l_max changes, then a call to
 * SpgmrMalloc must be made to obtain new memory for SpgmrSolve
 * to use.
 * -----------------------------------------------------------------
 */                                                                
     
int SpgmrSolve(SpgmrMem mem, void *A_data, N_Vector x, N_Vector b,
               int pretype, int gstype, realtype delta, 
               int max_restarts, void *P_data, N_Vector s1, 
               N_Vector s2, ATimesFn atimes, PSolveFn psolve, 
               realtype *res_norm, int *nli, int *nps);


/* Return values for SpgmrSolve */

#define SPGMR_SUCCESS            0  /* Converged                     */
#define SPGMR_RES_REDUCED        1  /* Did not converge, but reduced
                                       norm of residual              */
#define SPGMR_CONV_FAIL          2  /* Failed to converge            */
#define SPGMR_QRFACT_FAIL        3  /* QRfact found singular matrix  */
#define SPGMR_PSOLVE_FAIL_REC    4  /* psolve failed recoverably     */
#define SPGMR_ATIMES_FAIL_REC    5  /* atimes failed recoverably     */
#define SPGMR_PSET_FAIL_REC      6  /* pset faild recoverably        */

#define SPGMR_MEM_NULL          -1  /* mem argument is NULL          */
#define SPGMR_ATIMES_FAIL_UNREC -2  /* atimes returned failure flag  */
#define SPGMR_PSOLVE_FAIL_UNREC -3  /* psolve failed unrecoverably   */
#define SPGMR_GS_FAIL           -4  /* Gram-Schmidt routine faiuled  */        
#define SPGMR_QRSOL_FAIL        -5  /* QRsol found singular R        */
#define SPGMR_PSET_FAIL_UNREC   -6  /* pset failed unrecoverably     */

/*
 * -----------------------------------------------------------------
 * Function : SpgmrFree
 * -----------------------------------------------------------------
 * SpgmrMalloc frees the memory allocated by SpgmrMalloc. It is
 * illegal to use the pointer mem after a call to SpgmrFree.
 * -----------------------------------------------------------------
 */                                                                

void SpgmrFree(SpgmrMem mem);

/*
 * -----------------------------------------------------------------
 * Macro: SPGMR_VTEMP
 * -----------------------------------------------------------------
 * This macro provides access to the work vector vtemp in the
 * memory block of the SPGMR module.  The argument mem is the
 * memory pointer returned by SpgmrMalloc, of type SpgmrMem,
 * and the macro value is of type N_Vector.
 * On a return from SpgmrSolve with *nli = 0, this vector
 * contains the scaled preconditioned initial residual,
 * s1 * P1_inverse * (b - A x_0).
 * -----------------------------------------------------------------
 */

#define SPGMR_VTEMP(mem) (mem->vtemp)

#ifdef __cplusplus
}
#endif

#endif
```
