## Supplementary scripts for "A multi-scale *in silico* mouse model for insulin resistance and humanoid type 2 diabetes": sundials_sptfqmr.h

```
/*
 * -----------------------------------------------------------------
 * $Revision: 749 $
 * $Date: 2011-06-22 14:49:10 +0200 (Wed, 22 Jun 2011) $
 * -----------------------------------------------------------------
 * Programmer(s): Aaron Collier @ LLNL
 * -----------------------------------------------------------------
 * Copyright (c) 2005, The Regents of the University of California.
 * Produced at the Lawrence Livermore National Laboratory.
 * All rights reserved.
 * For details, see the LICENSE file.
 * -----------------------------------------------------------------
 * This is the header file for the implementation of the scaled
 * preconditioned Transpose-Free Quasi-Minimal Residual (SPTFQMR)
 * linear solver.
 *
 * The SPTFQMR algorithm solves a linear system of the form Ax = b.
 * Preconditioning is allowed on the left (PREC_LEFT), right
 * (PREC_RIGHT), or both (PREC_BOTH).  Scaling is allowed on both
 * sides.  We denote the preconditioner and scaling matrices as
 * follows:
 *   P1 = left preconditioner
 *   P2 = right preconditioner
 *   S1 = diagonal matrix of scale factors for P1-inverse b
 *   S2 = diagonal matrix of scale factors for P2 x
 * The matrices A, P1, and P2 are not required explicitly; only
 * routines that provide A, P1-inverse, and P2-inverse as operators
 * are required.
 *
 * In this notation, SPTFQMR applies the underlying TFQMR method to
 * the equivalent transformed system:
 *   Abar xbar = bbar, where
 *   Abar = S1 (P1-inverse) A (P2-inverse) (S2-inverse),
 *   bbar = S1 (P1-inverse) b, and
 *   xbar = S2 P2 x.
 *
 * The scaling matrices must be chosen so that vectors
 * S1 P1-inverse b and S2 P2 x have dimensionless components.  If
 * preconditioning is done on the left only (P2 = I), by a matrix P,
 * then S2 must be a scaling for x, while S1 is a scaling for
 * P-inverse b, and so may also be taken as a scaling for x.
 * Similarly, if preconditioning is done on the right only (P1 = I,
 * P2 = P), then S1 must be a scaling for b, while S2 is a scaling
 * for P x, and may also be taken as a scaling for b.
 *
 * The stopping test for the SPTFQMR iterations is on the L2-norm of
 * the scaled preconditioned residual:
 *   || bbar - Abar xbar ||_2 < delta
 * with an input test constant delta.
 *
 * The usage of this SPTFQMR solver involves supplying two routines
 * and making three calls.  The user-supplied routines are:
 *   atimes(A_data, x, y) to compute y = A x, given x,
 * and
 *   psolve(P_data, x, y, lr) to solve P1 x = y or P2 x = y for x,
 *                            given y.
 * The three user calls are:
 *   mem  = SptfqmrMalloc(lmax, vec_tmpl);
 *          to initialize memory
 *   flag = SptfqmrSolve(mem, A_data, x, b, pretype, delta, P_data,
 *                       sx, sb, atimes, psolve, res_norm, nli, nps);
 *          to solve the system, and
 *   SptfqmrFree(mem);
 *          to free the memory allocated by SptfqmrMalloc().
 * Complete details for specifying atimes() and psolve() and for the
 * usage calls are given in the paragraphs below and in the header
 * file sundials_iterative.h.
 * -----------------------------------------------------------------
 */

#ifndef _SPTFQMR_H
#define _SPTFQMR_H

#ifdef __cplusplus  /* wrapper to enable C++ usage */
extern "C" {
#endif

#include <sundials/sundials_iterative.h>

/*
 * -----------------------------------------------------------------
 * Types: struct SptfqmrMemRec and struct *SptfqmrMem
 * -----------------------------------------------------------------
 * A variable declaration of type struct *SptfqmrMem denotes a pointer
 * to a data structure of type struct SptfqmrMemRec. The SptfqmrMemRec
 * structure contains numerous fields that must be accessed by the
 * SPTFQMR linear solver module.
 *
 *  l_max  maximum Krylov subspace dimension that SptfqmrSolve will
 *         be permitted to use
 *
 *  r_star  vector (type N_Vector) which holds the initial scaled,
 *          preconditioned linear system residual
 *
 *  q/d/v/p/u/r  vectors (type N_Vector) used for workspace by
 *               the SPTFQMR algorithm
 *
 *  vtemp1/vtemp2/vtemp3  scratch vectors (type N_Vector) used as
 *                        temporary storage
 * -----------------------------------------------------------------
 */

typedef struct {

  int l_max;

  N_Vector r_star;
  N_Vector q;
  N_Vector d;
  N_Vector v;
  N_Vector p;
  N_Vector *r;
  N_Vector u;
  N_Vector vtemp1;
  N_Vector vtemp2;
  N_Vector vtemp3;

} SptfqmrMemRec, *SptfqmrMem;

/*
 * -----------------------------------------------------------------
 * Function : SptfqmrMalloc
 * -----------------------------------------------------------------
 * SptfqmrMalloc allocates additional memory needed by the SPTFQMR
 * linear solver module.
 *
 *  l_max  maximum Krylov subspace dimension that SptfqmrSolve will
 *         be permitted to use
 *
 *  vec_tmpl  implementation-specific template vector (type N_Vector)
 *            (created using either N_VNew_Serial or N_VNew_Parallel)
 *
 * If successful, SptfqmrMalloc returns a non-NULL memory pointer. If
 * an error occurs, then a NULL pointer is returned.
 * -----------------------------------------------------------------
 */

SptfqmrMem SptfqmrMalloc(int l_max, N_Vector vec_tmpl);

/*
 * -----------------------------------------------------------------
 * Function : SptfqmrSolve
 * -----------------------------------------------------------------
 * SptfqmrSolve solves the linear system Ax = b by means of a scaled
 * preconditioned Transpose-Free Quasi-Minimal Residual (SPTFQMR)
 * method.
 *
 *  mem  pointer to an internal memory block allocated during a
 *       prior call to SptfqmrMalloc
 *
 *  A_data  pointer to a data structure containing information
 *          about the coefficient matrix A (passed to user-supplied
 *          function referenced by atimes (function pointer))
 *
 *  x  vector (type N_Vector) containing initial guess x_0 upon
 *     entry, but which upon return contains an approximate solution
 *     of the linear system Ax = b (solution only valid if return
 *     value is either SPTFQMR_SUCCESS or SPTFQMR_RES_REDUCED)
 *
 *  b  vector (type N_Vector) set to the right-hand side vector b
 *     of the linear system (undisturbed by function)
 *
 *  pretype  variable (type int) indicating the type of
 *           preconditioning to be used (see sundials_iterative.h)
 *
 *  delta  tolerance on the L2 norm of the scaled, preconditioned
 *         residual (if return value == SPTFQMR_SUCCESS, then
 *         ||sb*P1_inv*(b-Ax)||_L2 <= delta)
 *
 *  P_data  pointer to a data structure containing preconditioner
 *          information (passed to user-supplied function referenced
 *          by psolve (function pointer))
 *
 *  sx  vector (type N_Vector) containing positive scaling factors
 *      for x (pass sx == NULL if scaling NOT required)
 *
 *  sb  vector (type N_Vector) containing positive scaling factors
 *      for b (pass sb == NULL if scaling NOT required)
 *
 *  atimes  user-supplied routine responsible for computing the
 *          matrix-vector product Ax (see sundials_iterative.h)
 *
 *  psolve  user-supplied routine responsible for solving the
 *          preconditioned linear system Pz = r (ignored if
 *          pretype == PREC_NONE) (see sundials_iterative.h)
 *
 *  res_norm  pointer (type realtype*) to the L2 norm of the
 *            scaled, preconditioned residual (if return value
 *            is either SPTFQMR_SUCCESS or SPTFQMR_RES_REDUCED, then
 *            *res_norm = ||sb*P1_inv*(b-Ax)||_L2, where x is
 *            the computed approximate solution, sb is the diagonal
 *            scaling matrix for the right-hand side b, and P1_inv
 *            is the inverse of the left-preconditioner matrix)
 *
 *  nli  pointer (type int*) to the total number of linear
 *       iterations performed
 *
 *  nps  pointer (type int*) to the total number of calls made
 *       to the psolve routine
 * -----------------------------------------------------------------
 */

int SptfqmrSolve(SptfqmrMem mem, void *A_data, N_Vector x, N_Vector b,
		 int pretype, realtype delta, void *P_data, N_Vector sx,
		 N_Vector sb, ATimesFn atimes, PSolveFn psolve,
		 realtype *res_norm, int *nli, int *nps);

/* Return values for SptfqmrSolve */

#define SPTFQMR_SUCCESS            0  /* SPTFQMR algorithm converged          */
#define SPTFQMR_RES_REDUCED        1  /* SPTFQMR did NOT converge, but the
				         residual was reduced                 */
#define SPTFQMR_CONV_FAIL          2  /* SPTFQMR algorithm failed to converge */
#define SPTFQMR_PSOLVE_FAIL_REC    3  /* psolve failed recoverably            */
#define SPTFQMR_ATIMES_FAIL_REC    4  /* atimes failed recoverably            */
#define SPTFQMR_PSET_FAIL_REC      5  /* pset faild recoverably               */

#define SPTFQMR_MEM_NULL          -1  /* mem argument is NULL                 */
#define SPTFQMR_ATIMES_FAIL_UNREC -2  /* atimes returned failure flag         */
#define SPTFQMR_PSOLVE_FAIL_UNREC -3  /* psolve failed unrecoverably          */
#define SPTFQMR_PSET_FAIL_UNREC   -4  /* pset failed unrecoverably            */

/*
 * -----------------------------------------------------------------
 * Function : SptfqmrFree
 * -----------------------------------------------------------------
 * SptfqmrFree frees the memory allocated by a call to SptfqmrMalloc.
 * It is illegal to use the pointer mem after a call to SptfqmrFree.
 * -----------------------------------------------------------------
 */

void SptfqmrFree(SptfqmrMem mem);

/*
 * -----------------------------------------------------------------
 * Macro : SPTFQMR_VTEMP
 * -----------------------------------------------------------------
 * This macro provides access to the work vector vtemp1 in the
 * memory block of the SPTFQMR module. The argument mem is the
 * memory pointer returned by SptfqmrMalloc, of type SptfqmrMem,
 * and the macro value is of type N_Vector.
 *
 * Note: Only used by IDA (vtemp1 contains P_inverse F if
 *       nli_inc == 0).
 * -----------------------------------------------------------------
 */

#define SPTFQMR_VTEMP(mem) (mem->vtemp1)

#ifdef __cplusplus
}
#endif

#endif
```
