## Supplementary scripts for "A multi-scale *in silico* mouse model for insulin resistance and humanoid type 2 diabetes": sundials_types.h

```
/*
 * -----------------------------------------------------------------
 * $Revision: 749 $
 * $Date: 2011-06-22 14:49:10 +0200 (Wed, 22 Jun 2011) $
 * ----------------------------------------------------------------- 
 * Programmer(s): Scott Cohen, Alan Hindmarsh, Radu Serban, and
 *                Aaron Collier @ LLNL
 * -----------------------------------------------------------------
 * Copyright (c) 2002, The Regents of the University of California.
 * Produced at the Lawrence Livermore National Laboratory.
 * All rights reserved.
 * For details, see the LICENSE file.
 *------------------------------------------------------------------
 * This header file exports two types: realtype and booleantype,
 * as well as the constants TRUE and FALSE.
 *
 * Users should include the header file sundials_types.h in every
 * program file and use the exported name realtype instead of
 * float, double or long double.
 *
 * The constants SUNDIALS_SINGLE_PRECISION, SUNDIALS_DOUBLE_PRECISION
 * and SUNDIALS_LONG_DOUBLE_PRECISION indicate the underlying data
 * type of realtype. It is set at the configuration stage.
 *
 * The legal types for realtype are float, double and long double.
 *
 * The macro RCONST gives the user a convenient way to define
 * real-valued constants. To use the constant 1.0, for example,
 * the user should write the following:
 *
 *   #define ONE RCONST(1.0)
 *
 * If realtype is defined as a double, then RCONST(1.0) expands
 * to 1.0. If realtype is defined as a float, then RCONST(1.0)
 * expands to 1.0F. If realtype is defined as a long double,
 * then RCONST(1.0) expands to 1.0L. There is never a need to
 * explicitly cast 1.0 to (realtype).
 *------------------------------------------------------------------
 */
  
#ifndef _SUNDIALSTYPES_H
#define _SUNDIALSTYPES_H

#ifdef __cplusplus  /* wrapper to enable C++ usage */
extern "C" {
#endif

#ifndef _SUNDIALS_CONFIG_H
#define _SUNDIALS_CONFIG_H
#include <sundials/sundials_config.h>
#endif

#include <float.h>
  
#if defined(SUNDIALS_SINGLE_PRECISION)

typedef float realtype;
#define RCONST(x) x##F
#define BIG_REAL FLT_MAX
#define SMALL_REAL FLT_MIN
#define UNIT_ROUNDOFF FLT_EPSILON

#elif defined(SUNDIALS_DOUBLE_PRECISION)

typedef double realtype;
#define RCONST(x) x
#define BIG_REAL DBL_MAX
#define SMALL_REAL DBL_MIN
#define UNIT_ROUNDOFF DBL_EPSILON

#elif defined(SUNDIALS_EXTENDED_PRECISION)

typedef long double realtype;
#define RCONST(x) x##L
#define BIG_REAL LDBL_MAX
#define SMALL_REAL LDBL_MIN
#define UNIT_ROUNDOFF LDBL_EPSILON

#endif

/*
 *------------------------------------------------------------------
 * Type : booleantype
 *------------------------------------------------------------------
 * Constants : FALSE and TRUE
 *------------------------------------------------------------------
 * ANSI C does not have a built-in boolean data type. Below is the
 * definition for a new type called booleantype. The advantage of
 * using the name booleantype (instead of int) is an increase in
 * code readability. It also allows the programmer to make a
 * distinction between int and boolean data. Variables of type
 * booleantype are intended to have only the two values FALSE and
 * TRUE which are defined below to be equal to 0 and 1,
 * respectively.
 *------------------------------------------------------------------
 */

#ifndef booleantype
#define booleantype int
#endif

#ifndef FALSE
#define FALSE 0
#endif

#ifndef TRUE
#define TRUE 1
#endif

#ifdef __cplusplus
}
#endif

#endif
```
