## Supplementary scripts for "A multi-scale *in silico* mouse model for insulin resistance and humanoid type 2 diabetes": cvodea.c

```
/*
 * -----------------------------------------------------------------
 * $Revision: 749 $
 * $Date: 2011-06-22 14:49:10 +0200 (Wed, 22 Jun 2011) $
 * ----------------------------------------------------------------- 
 * Programmer(s): Radu Serban @ LLNL
 * -----------------------------------------------------------------
 * Copyright (c) 2006, The Regents of the University of California.
 * Produced at the Lawrence Livermore National Laboratory.
 * All rights reserved.
 * For details, see the LICENSE file.
 * -----------------------------------------------------------------
 * This is the implementation file for the CVODEA adjoint integrator.
 * -----------------------------------------------------------------
 */

/* 
 * =================================================================
 * IMPORTED HEADER FILES
 * =================================================================
 */

#include <stdio.h>
#include <stdlib.h>

#include "cvodes_impl.h"

#include <sundials/sundials_math.h>
#include <sundials/sundials_types.h>

/* 
 * =================================================================
 * MACRO DEFINITIONS
 * =================================================================
 */

#define loop for(;;)

/* 
 * =================================================================
 * CVODEA PRIVATE CONSTANTS
 * =================================================================
 */

#define ZERO        RCONST(0.0)        /* real 0.0   */
#define ONE         RCONST(1.0)        /* real 1.0   */
#define TWO         RCONST(2.0)        /* real 2.0   */
#define HUNDRED     RCONST(100.0)      /* real 100.0 */
#define FUZZ_FACTOR RCONST(1000000.0)  /* fuzz factor for CVadjGetY */

/* 
 * =================================================================
 * PRIVATE FUNCTION PROTOTYPES
 * =================================================================
 */

static booleantype CVAallocVectors(CVadjMem ca_mem);
static void CVAfreeVectors(CVadjMem ca_mem);

static CkpntMem CVAckpntInit(CVodeMem cv_mem);
static CkpntMem CVAckpntNew(CVodeMem cv_mem);
static void CVAckpntDelete(CkpntMem *ck_memPtr);

static int  CVAdataStore(CVadjMem ca_mem, CkpntMem ck_mem);
static int  CVAckpntGet(CVodeMem cv_mem, CkpntMem ck_mem); 

static int CVAfindIndex(CVadjMem ca_mem, realtype t, 
                        long int *indx, booleantype *newpoint);

static booleantype CVAhermiteMalloc(CVadjMem ca_mem, long int steps);
static void CVAhermiteFree(DtpntMem *dt_mem, long int steps);
static int CVAhermiteGetY(CVadjMem ca_mem, realtype t, N_Vector y);
static int CVAhermiteStorePnt(CVodeMem cv_mem, DtpntMem d);

static booleantype CVApolynomialMalloc(CVadjMem ca_mem, long int steps);
static void CVApolynomialFree(DtpntMem *dt_mem, long int steps);
static int CVApolynomialGetY(CVadjMem ca_mem, realtype t, N_Vector y);
static int CVApolynomialStorePnt(CVodeMem cv_mem, DtpntMem d);

/* Wrappers */

static int CVArhs(realtype t, N_Vector yB, 
                  N_Vector yBdot, void *cvadj_mem);

static int CVArhsQ(realtype t, N_Vector yB, 
                   N_Vector qBdot, void *cvadj_mem);

/* 
 * =================================================================
 * EXPORTED FUNCTIONS IMPLEMENTATION
 * =================================================================
 */

/* 
 * -----------------------------------------------------------------
 * Readibility Constants
 * -----------------------------------------------------------------
 */

#define uround      (ca_mem->ca_uround)
#define tinitial    (ca_mem->ca_tinitial)
#define tfinal      (ca_mem->ca_tfinal)
#define nckpnts     (ca_mem->ca_nckpnts)
#define nsteps      (ca_mem->ca_nsteps)
#define ckpntData   (ca_mem->ca_ckpntData)
#define newData     (ca_mem->ca_newData)
#define np          (ca_mem->ca_np)
#define ytmp        (ca_mem->ca_ytmp)
#define f_B         (ca_mem->ca_fB)
#define f_data_B    (ca_mem->ca_f_dataB)
#define fQ_B        (ca_mem->ca_fQB)
#define fQ_data_B   (ca_mem->ca_fQ_dataB)
#define t_for_quad  (ca_mem->ca_t_for_quad)
#define Y0          (ca_mem->ca_Y0)
#define Y1          (ca_mem->ca_Y1)
#define Y           (ca_mem->ca_Y)
#define T           (ca_mem->ca_T)
#define interpType  (ca_mem->ca_interpType)
#define getY        (ca_mem->ca_getY)
#define storePnt    (ca_mem->ca_storePnt)
#define lfreeB      (ca_mem->ca_lfreeB)

#define zn         (cv_mem->cv_zn)
#define nst        (cv_mem->cv_nst)
#define q          (cv_mem->cv_q)
#define qu         (cv_mem->cv_qu)
#define qprime     (cv_mem->cv_qprime)
#define qwait      (cv_mem->cv_qwait)
#define L          (cv_mem->cv_L)
#define gammap     (cv_mem->cv_gammap)
#define h          (cv_mem->cv_h)
#define hprime     (cv_mem->cv_hprime)
#define hscale     (cv_mem->cv_hscale)
#define eta        (cv_mem->cv_eta)
#define etamax     (cv_mem->cv_etamax)
#define tn         (cv_mem->cv_tn)
#define tretlast   (cv_mem->cv_tretlast)
#define tau        (cv_mem->cv_tau)
#define tq         (cv_mem->cv_tq)
#define l          (cv_mem->cv_l)
#define saved_tq5  (cv_mem->cv_saved_tq5)
#define forceSetup (cv_mem->cv_forceSetup)
#define f          (cv_mem->cv_f)
#define lmm        (cv_mem->cv_lmm)
#define iter       (cv_mem->cv_iter)
#define itol       (cv_mem->cv_itol)
#define reltol     (cv_mem->cv_reltol)
#define Sabstol    (cv_mem->cv_Sabstol)
#define Vabstol    (cv_mem->cv_Vabstol)
#define efun       (cv_mem->cv_efun)
#define f_data     (cv_mem->cv_f_data)
#define errfp      (cv_mem->cv_errfp)
#define h0u        (cv_mem->cv_h0u)
#define quadr      (cv_mem->cv_quadr)
#define errconQ    (cv_mem->cv_errconQ)
#define znQ        (cv_mem->cv_znQ)
#define itolQ      (cv_mem->cv_itolQ)
#define reltolQ    (cv_mem->cv_reltolQ)
#define SabstolQ   (cv_mem->cv_SabstolQ)
#define VabstolQ   (cv_mem->cv_VabstolQ)
#define fQ         (cv_mem->cv_fQ)
#define tempv      (cv_mem->cv_tempv)
#define tempvQ     (cv_mem->cv_tempvQ)

#define t0_        (ck_mem->ck_t0)
#define t1_        (ck_mem->ck_t1)
#define zn_        (ck_mem->ck_zn)
#define znQ_       (ck_mem->ck_znQ)
#define quadr_     (ck_mem->ck_quadr)
#define zqm_       (ck_mem->ck_zqm)
#define nst_       (ck_mem->ck_nst)
#define tretlast_  (ck_mem->ck_tretlast)
#define q_         (ck_mem->ck_q)
#define qprime_    (ck_mem->ck_qprime)
#define qwait_     (ck_mem->ck_qwait)
#define L_         (ck_mem->ck_L)
#define gammap_    (ck_mem->ck_gammap)
#define h_         (ck_mem->ck_h)
#define hprime_    (ck_mem->ck_hprime)
#define hscale_    (ck_mem->ck_hscale)
#define eta_       (ck_mem->ck_eta)
#define etamax_    (ck_mem->ck_etamax)
#define tau_       (ck_mem->ck_tau)
#define tq_        (ck_mem->ck_tq)
#define l_         (ck_mem->ck_l)
#define saved_tq5_ (ck_mem->ck_saved_tq5)
#define next_      (ck_mem->ck_next)

/*
 * CVadjMalloc
 *
 * This routine allocates space for the global CVODEA memory
 * structure.
 */

void *CVadjMalloc(void *cvode_mem, long int steps, int interp)
{
  CVadjMem ca_mem;
  CVodeMem cv_mem;
  booleantype allocOK;
  long int i, ii;

  /* Check arguments */
  if (cvode_mem == NULL) {
    CVProcessError(NULL, 0, "CVODEA", "CVadjMalloc", MSGAM_NULL_CVMEM);
    return(NULL);
  }
  cv_mem = (CVodeMem)cvode_mem;

  if (steps <= 0) {
    CVProcessError(NULL, 0, "CVODEA", "CVadjMalloc", MSGAM_BAD_STEPS);
    return(NULL);
  }

  if ( (interp != CV_HERMITE) && (interp != CV_POLYNOMIAL) ) {
    CVProcessError(NULL, 0, "CVODEA", "CVadjMalloc", MSGAM_BAD_INTERP);
    return(NULL);
  } 

  /* Allocate memory block */
  ca_mem = NULL;
  ca_mem = (CVadjMem) malloc(sizeof(struct CVadjMemRec));
  if (ca_mem == NULL) {
    CVProcessError(NULL, 0, "CVODEA", "CVadjMalloc", MSGAM_MEM_FAIL);
    return(NULL);
  }

  /* Attach CVODES memory for forward runs */
  ca_mem->cv_mem = cv_mem;

  /* Set interpType */
  interpType = interp;

  /* Allocate memory for workspace vectors */
  allocOK = CVAallocVectors(ca_mem);
  if (!allocOK) {
    free(ca_mem); ca_mem = NULL;
    CVProcessError(NULL, 0, "CVODEA", "CVadjMalloc", MSGAM_MEM_FAIL);
    return(NULL);
  }

  /* Initialize Check Points linked list */
  ca_mem->ck_mem = NULL;
  ca_mem->ck_mem = CVAckpntInit(cv_mem);
  if (ca_mem->ck_mem == NULL) {
    CVAfreeVectors(ca_mem);
    free(ca_mem); ca_mem = NULL;
    CVProcessError(NULL, 0, "CVODEA", "CVadjMalloc", MSGAM_MEM_FAIL);
    return(NULL);
  }

  /* Allocate space for the array of Data Point structures */
  
  ca_mem->dt_mem = NULL;
  ca_mem->dt_mem = (DtpntMem *) malloc((steps+1)*sizeof(struct DtpntMemRec *));
  if (ca_mem->dt_mem == NULL) {
    while (ca_mem->ck_mem != NULL) CVAckpntDelete(&(ca_mem->ck_mem));
    CVAfreeVectors(ca_mem);
    free(ca_mem); ca_mem = NULL;
    CVProcessError(NULL, 0, "CVODEA", "CVadjMalloc", MSGAM_MEM_FAIL);
    return(NULL);
  }

  for (i=0; i<=steps; i++) { 
    ca_mem->dt_mem[i] = NULL;
    ca_mem->dt_mem[i] = (DtpntMem) malloc(sizeof(struct DtpntMemRec));
    if (ca_mem->dt_mem[i] == NULL) {
      for(ii=0; ii<i; ii++) {free(ca_mem->dt_mem[ii]); ca_mem->dt_mem[ii] = NULL;}
      free(ca_mem->dt_mem); ca_mem->dt_mem = NULL;
      while (ca_mem->ck_mem != NULL) CVAckpntDelete(&(ca_mem->ck_mem));
      CVAfreeVectors(ca_mem);
      free(ca_mem); ca_mem = NULL;
      CVProcessError(NULL, 0, "CVODEA", "CVadjMalloc", MSGAM_MEM_FAIL);
      return(NULL);
    }
  }

  switch (interpType) {

  case CV_HERMITE:
    /* Allocate Data Points memory */
    allocOK = CVAhermiteMalloc(ca_mem, steps);
    if (!allocOK) {
      for(i=0; i<=steps; i++) {free(ca_mem->dt_mem[i]); ca_mem->dt_mem[i] = NULL;}
      free(ca_mem->dt_mem); ca_mem->dt_mem = NULL;
      while (ca_mem->ck_mem != NULL) CVAckpntDelete(&(ca_mem->ck_mem));
      CVAfreeVectors(ca_mem);
      free(ca_mem); ca_mem = NULL;
      CVProcessError(NULL, 0, "CVODEA", "CVadjMalloc", MSGAM_MEM_FAIL);
      return(NULL);
    }
    /* Attach interpolation functions getY and storePnt */
    getY = CVAhermiteGetY;
    storePnt = CVAhermiteStorePnt;
    /* Rename zn[0] and zn[1] for use in interpolation */
    Y0 = zn[0];
    Y1 = zn[1];
    break;
  case CV_POLYNOMIAL:
    /* Allocate Data Points memory */
    allocOK = CVApolynomialMalloc(ca_mem, steps);
    if (!allocOK) {
      for(i=0; i<=steps; i++) {free(ca_mem->dt_mem[i]); ca_mem->dt_mem[i] = NULL;}
      free(ca_mem->dt_mem); ca_mem->dt_mem = NULL;
      while (ca_mem->ck_mem != NULL) CVAckpntDelete(&(ca_mem->ck_mem));
      CVAfreeVectors(ca_mem);
      free(ca_mem); ca_mem = NULL;
      CVProcessError(NULL, 0, "CVODEA", "CVadjMalloc", MSGAM_MEM_FAIL);
      return(NULL);
    }
    /* Attach interpolation functions getY and storePnt */
    getY = CVApolynomialGetY;
    storePnt = CVApolynomialStorePnt;
    /* Rename zn for use in interpolation */
    for (i=0;i<L_MAX;i++) Y[i] = zn[i];
    break;
  }

  /* Other entries in ca_mem */
  uround   = cv_mem->cv_uround;
  nsteps   = steps;
  tinitial = tn; 

  /* Initialize nckpnts to ZERO */
  nckpnts = 0;

  /* Initialize backward cvode memory to NULL */
  ca_mem->cvb_mem = NULL;

  ca_mem->ca_lmemB = NULL;
  ca_mem->ca_lfreeB = NULL;
  ca_mem->ca_pmemB = NULL;
  
  ca_mem->ca_f_dataB = NULL;
  ca_mem->ca_fQ_dataB = NULL;

  return((void *)ca_mem);
} 

/* 
 * CVadjSetInterpType
 *
 * Changes the interpolation type.
 * Possible return values: CV_SUCCESS, CV_ADJMEM_NULL, CV_ILL_INPUT, CV_MEM_FAIL
 */

int CVadjSetInterpType(void *cvadj_mem, int interp)
{
  CVadjMem ca_mem;
  CVodeMem cv_mem;
  booleantype allocOK;
  long int i;
  
  if (cvadj_mem == NULL) {
    CVProcessError(NULL, CV_ADJMEM_NULL, "CVODEA", "CVadjSetInterpType", MSGAM_NULL_CAMEM);
    return(CV_ADJMEM_NULL);
  }
  ca_mem = (CVadjMem) cvadj_mem;

  cv_mem = ca_mem->cv_mem;

  if ( (interp != CV_HERMITE) && (interp != CV_POLYNOMIAL) ) {
    CVProcessError(NULL, CV_ILL_INPUT, "CVODEA", "CVadjSetInterpType", MSGAM_BAD_INTERP);
    return(CV_ILL_INPUT);
  } 

  if (interp == interpType) return(CV_SUCCESS);

  interpType = interp;


  switch (interpType) {

  case CV_HERMITE:
    /* Delete Data Points memory */
    CVApolynomialFree(ca_mem->dt_mem, nsteps);
    /* Allocate Data Points memory */
    allocOK = CVAhermiteMalloc(ca_mem, nsteps);
    if (!allocOK) {
      CVProcessError(NULL, CV_MEM_FAIL, "CVODEA", "CVadjSetInterpType", MSGAM_MEM_FAIL);
      return(CV_MEM_FAIL);
    }
    /* Attach interpolation functions getY and storePnt */
    getY = CVAhermiteGetY;
    storePnt = CVAhermiteStorePnt;
    /* Rename zn[0] and zn[1] for use in interpolation */
    Y0 = zn[0];
    Y1 = zn[1];
    break;
  case CV_POLYNOMIAL:
    /* Delete Data Points memory */
    CVAhermiteFree(ca_mem->dt_mem, nsteps);
    /* Allocate Data Points memory */
    allocOK = CVApolynomialMalloc(ca_mem, nsteps);
    if (!allocOK) {
      CVProcessError(NULL, CV_MEM_FAIL, "CVODEA", "CVadjSetInterpType", MSGAM_MEM_FAIL);
      return(CV_MEM_FAIL);
    }
    /* Attach interpolation functions getY and storePnt */
    getY = CVApolynomialGetY;
    storePnt = CVApolynomialStorePnt;
    /* Rename zn for use in interpolation */
    for (i=0;i<L_MAX;i++) Y[i] = zn[i];
    break;
  }

  return(CV_SUCCESS);

}

/*
 * CVadjFree
 *
 * This routine frees the memory allocated by CVadjMalloc.
 */

void CVadjFree(void **cvadj_mem)
{
  void *cvode_bmem;
  CVadjMem ca_mem;
  long int i;

  if (*cvadj_mem == NULL) return;

  ca_mem = (CVadjMem) (*cvadj_mem);

  /* Delete check points one by one */
  while (ca_mem->ck_mem != NULL) CVAckpntDelete(&(ca_mem->ck_mem));

  /* Free vectors at each data point */
  switch (interpType) {
  case CV_HERMITE:
    CVAhermiteFree(ca_mem->dt_mem, nsteps);
    break;
  case CV_POLYNOMIAL:
    CVApolynomialFree(ca_mem->dt_mem, nsteps);
    break;
  }
  for(i=0; i<=nsteps; i++) {free(ca_mem->dt_mem[i]); ca_mem->dt_mem[i] = NULL;}
  free(ca_mem->dt_mem); ca_mem->dt_mem = NULL;

  /* Free workspace vectors in ca_mem */
  CVAfreeVectors(ca_mem);

  /* Free memory allocated by the linear solver */
  if (lfreeB != NULL) lfreeB(ca_mem);

  /* Free CVODES memory for backward run */
  cvode_bmem = (void *)ca_mem->cvb_mem;
  CVodeFree(&cvode_bmem);

  /* Free CVODEA memory */
  free(*cvadj_mem);
  *cvadj_mem = NULL;
}

/*
 * CVodeF
 *
 * This routine integrates to tout and returns solution into yout.
 * In the same time, it stores check point data every 'steps' steps. 
 * 
 * CVodeF can be called repeatedly by the user.
 *
 * ncheckPtr points to the number of check points stored so far.
 */

int CVodeF(void *cvadj_mem, realtype tout, N_Vector yout, 
           realtype *tret, int itask, int *ncheckPtr)
{
  CVadjMem ca_mem;
  CVodeMem cv_mem;
  CkpntMem tmp;
  DtpntMem *dt_mem;
  int cv_itask, flag;
  booleantype iret;

  if (cvadj_mem == NULL) {
    CVProcessError(NULL, CV_ADJMEM_NULL, "CVODEA", "CVodeF", MSGAM_NULL_CAMEM);
    return(CV_ADJMEM_NULL);
  }
  ca_mem = (CVadjMem) cvadj_mem;

  cv_mem = ca_mem->cv_mem;
  dt_mem = ca_mem->dt_mem;

  iret = TRUE;
  cv_itask = CV_ONE_STEP;

  /* Interpret itask */
  switch (itask) {
  case CV_NORMAL:
    iret = FALSE;
    cv_itask = CV_ONE_STEP;
    break;
  case CV_ONE_STEP:
    iret = TRUE;
    cv_itask = CV_ONE_STEP;
    break;
  case CV_NORMAL_TSTOP:
    iret = FALSE;
    cv_itask = CV_ONE_STEP_TSTOP;
    break;
  case CV_ONE_STEP_TSTOP:
    iret = TRUE;
    cv_itask = CV_ONE_STEP_TSTOP;
    break;
  }

  /* On the first step, load dt_mem[0].
     On subsequent steps, test if taking a new step is necessary. */
  if ( nst == 0) {

    dt_mem[0]->t = ca_mem->ck_mem->ck_t0;
    storePnt(cv_mem, dt_mem[0]);

  } else if ( (tn - tout)*h >= ZERO ) {

    /* If tout was passed, return interpolated solution. 
       No changes to ck_mem or dt_mem are needed. */
    *tret = tout;
    flag = CVodeGetDky(cv_mem, tout, 0, yout);
    *ncheckPtr = nckpnts;
    newData = TRUE;
    ckpntData = ca_mem->ck_mem;
    np = nst % nsteps + 1;
    return(flag);

  }

  /* Integrate to tout (in CV_ONE_STEP mode) while loading check points */
  loop {

    /* Perform one step of the integration */

    flag = CVode(cv_mem, tout, yout, tret, cv_itask);
    if (flag < 0) break;

    /* Test if a new check point is needed */

    if ( nst % nsteps == 0 ) {

      ca_mem->ck_mem->ck_t1 = *tret;

      /* Create a new check point, load it, and append it to the list */
      tmp = CVAckpntNew(cv_mem);
      if (tmp == NULL) {
        CVProcessError(cv_mem, CV_MEM_FAIL, "CVODEA", "CVodeF", MSGAM_MEM_FAIL);
        flag = CV_MEM_FAIL;
        break;
      }
      tmp->ck_next = ca_mem->ck_mem;
      ca_mem->ck_mem = tmp;
      nckpnts++;
      forceSetup = TRUE;
      
      /* Reset i=0 and load dt_mem[0] */
      dt_mem[0]->t = ca_mem->ck_mem->ck_t0;
      storePnt(cv_mem, dt_mem[0]);

    } else {
      
      /* Load next point in dt_mem */
      dt_mem[nst%nsteps]->t = *tret;
      storePnt(cv_mem, dt_mem[nst%nsteps]);

    }

    /* Set t1 field of the current ckeck point structure
       for the case in which there will be no future
       check points */
    ca_mem->ck_mem->ck_t1 = *tret;

    /* tfinal is now set to *tret */
    tfinal = *tret;

    /* Return if in CV_ONE_STEP mode */
    if (iret) break;

    /* Return if tout reached */
    if ( (*tret - tout)*h >= ZERO ) {
      *tret = tout;
      CVodeGetDky(cv_mem, tout, 0, yout);
      break;
    }

  } /* end of loop() */

  /* Get ncheck from ca_mem */ 
  *ncheckPtr = nckpnts;

  /* Data is available for the last interval */
  newData = TRUE;
  ckpntData = ca_mem->ck_mem;
  np = nst % nsteps + 1;

  return(flag);
}

/*
 * CVodeCreateB, CVodeMallocB, and CVodeReInitB
 *
 * Wrappers for the backward phase around the corresponding 
 * CVODES functions
 */

int CVodeCreateB(void *cvadj_mem, int lmmB, int iterB)
{
  CVadjMem ca_mem;
  void *cvode_mem;

  if (cvadj_mem == NULL) {
    CVProcessError(NULL, CV_ADJMEM_NULL, "CVODEA", "CVodeCreateB", MSGAM_NULL_CAMEM);
    return(CV_ADJMEM_NULL);
  }

  ca_mem = (CVadjMem) cvadj_mem;

  cvode_mem = CVodeCreate(lmmB, iterB);

  if (cvode_mem == NULL)
    return(CV_MEM_FAIL);

  ca_mem->cvb_mem = (CVodeMem) cvode_mem;

  return(CV_SUCCESS);

}

int CVodeMallocB(void *cvadj_mem, CVRhsFnB fB, 
                 realtype tB0, N_Vector yB0,
                 int itolB, realtype reltolB, void *abstolB)
{
  CVadjMem ca_mem;
  CVodeMem cv_mem;
  int sign, flag;

  if (cvadj_mem == NULL) {
    CVProcessError(NULL, CV_ADJMEM_NULL, "CVODEA", "CVodeMallocB", MSGAM_NULL_CAMEM);
    return(CV_ADJMEM_NULL);
  }
  ca_mem = (CVadjMem) cvadj_mem;

  cv_mem = ca_mem->cvb_mem;

  flag = CVodeMalloc(cv_mem, CVArhs, tB0, yB0,
                     itolB, reltolB, abstolB);
  if (flag != CV_SUCCESS) return(flag);

  sign = (tfinal - tinitial > ZERO) ? 1 : -1;
  if ( (sign*(tB0-tinitial) < ZERO) || (sign*(tfinal-tB0) < ZERO) ) {
    CVProcessError(cv_mem, CV_BAD_TB0, "CVODEA", "CVodeMallocB", MSGAM_BAD_TB0);
    return(CV_BAD_TB0);
  }

  f_B = fB;

  CVodeSetMaxHnilWarns(cv_mem, -1);
  CVodeSetFdata(cv_mem, cvadj_mem);

  return(CV_SUCCESS);

}

int CVodeReInitB(void *cvadj_mem, CVRhsFnB fB, 
                 realtype tB0, N_Vector yB0, 
                 int itolB, realtype reltolB, void *abstolB)
{
  CVadjMem ca_mem;
  CVodeMem cv_mem;
  int sign, flag;

  if (cvadj_mem == NULL) {
    CVProcessError(NULL, CV_ADJMEM_NULL, "CVODEA", "CVodeReInitB", MSGAM_NULL_CAMEM);
    return(CV_ADJMEM_NULL);
  }
  ca_mem = (CVadjMem) cvadj_mem;

  cv_mem = ca_mem->cvb_mem;

  flag = CVodeReInit(cv_mem, CVArhs, tB0, yB0,
                     itolB, reltolB, abstolB);

  if (flag != CV_SUCCESS) return(flag);

  sign = (tfinal - tinitial > ZERO) ? 1 : -1;
  if ( (sign*(tB0-tinitial) < ZERO) || (sign*(tfinal-tB0) < ZERO) ) {
    CVProcessError(cv_mem, CV_BAD_TB0, "CVODEA", "CVodeReInitB", MSGAM_BAD_TB0);
    return(CV_BAD_TB0);
  }
  
  f_B  = fB;

  CVodeSetMaxHnilWarns(cv_mem, -1);
  CVodeSetFdata(cv_mem, cvadj_mem);

  return(CV_SUCCESS);

}

/*
 * CVodeQuadMallocB, and CVodeQuadReInitB
 *
 * Wrappers for the backward phase around the corresponding 
 * CVODES functions
 */

int CVodeQuadMallocB(void *cvadj_mem, CVQuadRhsFnB fQB, N_Vector yQB0)
{
  CVadjMem ca_mem;
  void *cvode_mem;
  int flag;

  if (cvadj_mem == NULL) {
    CVProcessError(NULL, CV_ADJMEM_NULL, "CVODEA", "CVodeQuadMallocB", MSGAM_NULL_CAMEM);
    return(CV_ADJMEM_NULL);
  }
  ca_mem = (CVadjMem) cvadj_mem;

  fQ_B = fQB;

  cvode_mem = (void *) ca_mem->cvb_mem;

  flag = CVodeQuadMalloc(cvode_mem, CVArhsQ, yQB0);
  if (flag != CV_SUCCESS) return(flag);

  flag = CVodeSetQuadFdata(cvode_mem, cvadj_mem);

  return(flag);

}

int CVodeQuadReInitB(void *cvadj_mem, CVQuadRhsFnB fQB, N_Vector yQB0)
{
  CVadjMem ca_mem;
  void *cvode_mem;
  int flag;

  if (cvadj_mem == NULL) {
    CVProcessError(NULL, CV_ADJMEM_NULL, "CVODEA", "CVodeQuadReInitB", MSGAM_NULL_CAMEM);
    return(CV_ADJMEM_NULL);
  }
  ca_mem = (CVadjMem) cvadj_mem;

  fQ_B = fQB;

  cvode_mem = (void *) ca_mem->cvb_mem;

  flag = CVodeQuadReInit(cvode_mem, CVArhsQ, yQB0);

  return(flag);

}

/*
 * CVodeB
 *
 * This routine performs the backward integration towards tBout. 
 * When necessary, it performs a forward integration between two 
 * consecutive check points to update interpolation data.
 * itask can be CV_NORMAL or CV_ONE_STEP only.
 *
 * On a successful return, CVodeB returns either CV_SUCCESS
 * (in ONE_STEP mode or if tBout was reached in NORMAL mode)
 * unless the return time happens to be a checkpoint, in which
 * case it returns CV_TSTOP_RETURN)
 */

int CVodeB(void *cvadj_mem, realtype tBout, N_Vector yBout, 
           realtype *tBret, int itaskB)
{
  CVadjMem ca_mem;
  CkpntMem ck_mem;
  CVodeMem cvb_mem;
  int sign, flag, cv_itask;
  realtype tBn, hB, troundoff;
  
  if (cvadj_mem == NULL) {
    CVProcessError(NULL, CV_ADJMEM_NULL, "CVODEA", "CVodeB", MSGAM_NULL_CAMEM);
    return(CV_ADJMEM_NULL);
  }
  ca_mem  = (CVadjMem) cvadj_mem;

  cvb_mem = ca_mem->cvb_mem;
  if (cvb_mem == NULL) {
    CVProcessError(NULL, CV_BCKMEM_NULL, "CVODEA", "CVodeB", MSGAM_NULL_CVMEM);
    return(CV_BCKMEM_NULL);
  }

  if (itaskB == CV_NORMAL)
    cv_itask = CV_NORMAL_TSTOP;
  else if (itaskB == CV_ONE_STEP)
    cv_itask = CV_ONE_STEP_TSTOP;
  else {
    CVProcessError(cvb_mem, CV_BAD_ITASK, "CVODEA", "CVodeB", MSGAM_BAD_ITASKB);
    return(CV_BAD_ITASK);
  }

  ck_mem = ca_mem->ck_mem;

  sign = (tfinal - tinitial > ZERO) ? 1 : -1;

  if ( (sign*(tBout-tinitial) < ZERO) || (sign*(tfinal-tBout) < ZERO) ) {
    CVProcessError(cvb_mem, CV_BAD_TBOUT, "CVODEA", "CVodeB", MSGAM_BAD_TBOUT);
    return(CV_BAD_TBOUT);
  }

  /* Move ck_mem pointer to the checkpoint appropriate for
     tBn (the current time reached by CVODES) */
  tBn = cvb_mem->cv_tn;
  troundoff = HUNDRED*uround*(ABS(tBn) + ABS(hB));
  while ( sign*(tBn - t0_) <= troundoff ) {
    if (next_ == NULL) break;
    ck_mem = next_;
  }

  loop {

    /* Store interpolation data if not available 
       This is the 2nd forward integration pass */
    if (ck_mem != ckpntData) {
      flag = CVAdataStore(ca_mem, ck_mem);
      if (flag != CV_SUCCESS) break;
    }

    /* Backward integration */
    CVodeSetStopTime((void *)cvb_mem, t0_);
    flag = CVode(cvb_mem, tBout, yBout, tBret, cv_itask);

    /* If an error occured, return now */
    if (flag < 0) break;

    /* Set the time at which CVodeGetQuadB will evaluate any quadratures */
    t_for_quad = *tBret;

    /* If in CV_ONE_STEP mode, return now (flag=CV_SUCCESS or flag=CV_TSTOP_RETURN) */
    if (itaskB == CV_ONE_STEP) break;

    /* If succesfully reached tBout, return now */
    if ( sign*(*tBret - tBout) <= ZERO) break;

    /* Move check point in linked list to next one */
    ck_mem = next_;

  } 

  return(flag);
}

/* 
 * =================================================================
 * PRIVATE FUNCTIONS
 * =================================================================
 */

/*
 * CVAallocVectors
 *
 * Allocate memory for adjoint workspace N_Vectors
 */

static booleantype CVAallocVectors(CVadjMem ca_mem)
{
  CVodeMem cv_mem;

  cv_mem = ca_mem->cv_mem;

  ytmp = NULL;
  ytmp = N_VClone(tempv);
  if (ytmp == NULL) {
    return(FALSE);
  }

  return(TRUE);

}

/*
 * CVAfreeVectors
 * 
 * Free memory allocated by CVAallocVectors
 */

static void CVAfreeVectors(CVadjMem ca_mem)
{
  N_VDestroy(ytmp);
}


/*
 * CVAckpntInit
 *
 * This routine initializes the check point linked list with 
 * information from the initial time.
 */

static CkpntMem CVAckpntInit(CVodeMem cv_mem)
{
  CkpntMem ck_mem;

  /* Allocate space for ckdata */
  ck_mem = NULL;
  ck_mem = (CkpntMem) malloc(sizeof(struct CkpntMemRec));
  if (ck_mem == NULL) return(NULL);

  zn_[0] = NULL;
  zn_[0] = N_VClone(tempv);
  if (zn_[0] == NULL) {
    free(ck_mem); ck_mem = NULL;
    return(NULL);
  }
  
  zn_[1] = NULL;
  zn_[1] = N_VClone(tempv);
  if (zn_[1] == NULL) {
    N_VDestroy(zn_[0]);
    free(ck_mem); ck_mem = NULL;
    return(NULL);
  }

  /* zn_[qmax] was not allocated */
  zqm_ = 0;

  /* Load ckdata from cv_mem */
  N_VScale(ONE, zn[0], zn_[0]);
  t0_    = tn;
  nst_   = 0;
  q_     = 1;
  h_     = 0.0;
  
  /* Do we need to carry quadratures */
  quadr_ = quadr && errconQ;

  if (quadr_) {
    znQ_[0] = NULL;
    znQ_[0] = N_VClone(tempvQ);
    if (znQ_[0] == NULL) {
      N_VDestroy(zn_[0]);
      N_VDestroy(zn_[1]);
      free(ck_mem); ck_mem = NULL;
      return(NULL);
    }
    N_VScale(ONE, znQ[0], znQ_[0]);
  }

  /* Next in list */
  next_  = NULL;

  return(ck_mem);
}

/*
 * CVAckpntNew
 *
 * This routine allocates space for a new check point and sets 
 * its data from current values in cv_mem.
 */

static CkpntMem CVAckpntNew(CVodeMem cv_mem)
{
  CkpntMem ck_mem;
  int j, jj;
  int qmax; 

  /* Allocate space for ckdata */
  ck_mem = NULL;
  ck_mem = (CkpntMem) malloc(sizeof(struct CkpntMemRec));
  if (ck_mem == NULL) return(NULL);

  /* Test if we need to allocate space for the last zn.
     NOTE: zn(qmax) may be needed for a hot restart, if an order
     increase is deemed necessary at the first step after a check 
     point */
  qmax = cv_mem->cv_qmax;
  zqm_ = (q < qmax) ? qmax : 0;

  for (j=0; j<=q; j++) {
    zn_[j] = NULL;
    zn_[j] = N_VClone(tempv);
    if (zn_[j] == NULL) {
      for (jj=0; jj<j; jj++) N_VDestroy(zn_[jj]);
      free(ck_mem); ck_mem = NULL;
      return(NULL);
    }
  }

  if (q < qmax) {
    zn_[qmax] = NULL;
    zn_[qmax] = N_VClone(tempv);
    if (zn_[qmax] == NULL) {
      for (j=0; j<=q; j++) N_VDestroy(zn_[j]);
      free(ck_mem); ck_mem = NULL;
      return(NULL);
    }
  }

  /* Test if we need to carry quadratures */
  quadr_ = quadr && errconQ;

  if (quadr_) {

    for (j=0; j<=q; j++) {
      znQ_[j] = NULL;
      znQ_[j] = N_VClone(tempvQ);
      if(znQ_[j] == NULL) {
        for (jj=0; jj<j; jj++) N_VDestroy(znQ_[jj]);
        if (q < qmax) N_VDestroy(zn_[qmax]);
        for (j=0; j<=q; j++) N_VDestroy(zn_[j]);
        free(ck_mem); ck_mem = NULL;
        return(NULL);
      }
    }

    if ( q < qmax) {
      znQ_[qmax] = NULL;
      znQ_[qmax] = N_VClone(tempvQ);
      if (znQ_[qmax] == NULL) {
        for (j=0; j<=q; j++) N_VDestroy(znQ_[j]);
        N_VDestroy(zn_[qmax]);
        for (j=0; j<=q; j++) N_VDestroy(zn_[j]);
        free(ck_mem); ck_mem = NULL;
        return(NULL);
      }
    }
  }

  /* Load check point data from cv_mem */

  for (j=0; j<=q; j++) N_VScale(ONE, zn[j], zn_[j]);
  if ( q < qmax ) N_VScale(ONE, zn[qmax], zn_[qmax]);

  if(quadr_) {
    for (j=0; j<=q; j++) N_VScale(ONE, znQ[j], znQ_[j]);
    if ( q < qmax ) N_VScale(ONE, znQ[qmax], znQ_[qmax]);
  }

  for (j=0; j<=L_MAX; j++)     tau_[j] = tau[j];
  for (j=0; j<=NUM_TESTS; j++) tq_[j] = tq[j];
  for (j=0; j<=q; j++)         l_[j] = l[j];
  nst_       = nst;
  tretlast_  = tretlast;
  q_         = q;
  qprime_    = qprime;
  qwait_     = qwait;
  L_         = L;
  gammap_    = gammap;
  h_         = h;
  hprime_    = hprime;
  hscale_    = hscale;
  eta_       = eta;
  etamax_    = etamax;
  t0_        = tn;
  saved_tq5_ = saved_tq5;

  return(ck_mem);
}

/*
 * CVAckpntDelete
 *
 * This routine deletes the first check point in list.
 */

static void CVAckpntDelete(CkpntMem *ck_memPtr)
{
  CkpntMem tmp;
  int j;

  if (*ck_memPtr != NULL) {

    /* store head of list */
    tmp = *ck_memPtr;

    /* move head of list */
    *ck_memPtr = (*ck_memPtr)->ck_next;

    /* free N_Vectors in tmp */
    for (j=0;j<=tmp->ck_q;j++) N_VDestroy(tmp->ck_zn[j]);
    if (tmp->ck_zqm != 0) N_VDestroy(tmp->ck_zn[tmp->ck_zqm]);

    /* free N_Vectors for quadratures in tmp 
       Note that at the check point at t_initial, only znQ_[0] 
       was allocated*/
    if(tmp->ck_quadr) {
      if(tmp->ck_next != NULL) {
        for (j=0;j<=tmp->ck_q;j++) N_VDestroy(tmp->ck_znQ[j]);
        if (tmp->ck_zqm != 0) N_VDestroy(tmp->ck_znQ[tmp->ck_zqm]);
      } else {
        N_VDestroy(tmp->ck_znQ[0]);
      }
    }

    free(tmp); tmp = NULL;

  }
}

/*
 * CVAdataStore
 *
 * This routine integrates the forward model starting at the check
 * point ck_mem and stores y and yprime at all intermediate steps.
 *
 * Return values:
 * CV_SUCCESS
 * CV_REIFWD_FAIL
 * CV_FWD_FAIL
 */

static int CVAdataStore(CVadjMem ca_mem, CkpntMem ck_mem)
{
  CVodeMem cv_mem;
  DtpntMem *dt_mem;
  realtype t;
  long int i;
  int flag;

  cv_mem = ca_mem->cv_mem;
  dt_mem = ca_mem->dt_mem;

  /* Initialize cv_mem with data from ck_mem */
  flag = CVAckpntGet(cv_mem, ck_mem);
  if (flag != CV_SUCCESS)
    return(CV_REIFWD_FAIL);

  /* Set first structure in dt_mem[0] */
  dt_mem[0]->t = t0_;
  storePnt(cv_mem, dt_mem[0]);

  /* Run CVode to set following structures in dt_mem[i] */

  i = 1;
  do {
    flag = CVode(cv_mem, t1_, ytmp, &t, CV_ONE_STEP);
    if (flag < 0) 
      return(CV_FWD_FAIL);
    dt_mem[i]->t = t;
    storePnt(cv_mem, dt_mem[i]);
    i++;
  } while (t<t1_);


  newData = TRUE;       /* New data is now available    */
  ckpntData = ck_mem;   /* starting at this check point */
  np = i;               /* and we have this many points */

  return(CV_SUCCESS);
}

/*
 * CVAckpntGet
 *
 * This routine prepares CVODES for a hot restart from
 * the check point ck_mem
 */

static int CVAckpntGet(CVodeMem cv_mem, CkpntMem ck_mem) 
{
  int j;
  int flag;
  int qmax;
  void *abstol;

  abstol = NULL;

  if (next_ == NULL) {

    /* In this case, we just call the reinitialization routine,
       but make sure we use the same initial stepsize as on 
       the first run. */

    CVodeSetInitStep(cv_mem, h0u);

    switch (itol) {
    case CV_SS:
      abstol = (void *) &Sabstol;
      break;
    case CV_SV:
      abstol = (void *)Vabstol;
      break;
    case CV_EE:
      abstol = (void *)efun;
      break;
    }
    flag = CVodeReInit(cv_mem, f, t0_, zn_[0], itol, reltol, abstol);
    if (flag != CV_SUCCESS) return(flag);

    if(quadr_) {
      flag = CVodeQuadReInit(cv_mem, fQ, znQ_[0]);
      if (flag != CV_SUCCESS) return(flag);
    }

  } else {
    
    qmax = cv_mem->cv_qmax;

    /* Copy parameters from check point data structure */
    nst       = nst_;
    tretlast  = tretlast_;
    q         = q_;
    qprime    = qprime_;
    qwait     = qwait_;
    L         = L_;
    gammap    = gammap_;
    h         = h_;
    hprime    = hprime_;
    hscale    = hscale_;
    eta       = eta_;
    etamax    = etamax_;
    tn        = t0_;
    saved_tq5 = saved_tq5_;
    
    /* Copy the arrays from check point data structure */
    for (j=0; j<=q; j++) N_VScale(ONE, zn_[j], zn[j]);
    if ( q < qmax ) N_VScale(ONE, zn_[qmax], zn[qmax]);
    if(quadr_) {
      for (j=0; j<=q; j++) N_VScale(ONE, znQ_[j], znQ[j]);
      if ( q < qmax ) N_VScale(ONE, znQ_[qmax], znQ[qmax]);
    }
    for (j=0; j<=L_MAX; j++)     tau[j] = tau_[j];
    for (j=0; j<=NUM_TESTS; j++) tq[j] = tq_[j];
    for (j=0; j<=q; j++)         l[j] = l_[j];
    
    /* Force a call to setup */
    forceSetup = TRUE;

  }

  return(CV_SUCCESS);
}

/* 
 * -----------------------------------------------------------------
 * Functions for interpolation
 * -----------------------------------------------------------------
 */

/*
 * CVAfindIndex
 *
 * Finds the index in the array of data point strctures such that
 *     dt_mem[indx-1].t <= t < dt_mem[indx].t
 * If indx is changed from the previous invocation, then newpoint = TRUE
 *
 * If t is beyond the leftmost limit, but close enough, indx=0.
 *
 * Returns CV_SUCCESS if successful and CV_GETY_BADT if unable to
 * find indx (t is too far beyond limits).
 */

static int CVAfindIndex(CVadjMem ca_mem, realtype t, 
                        long int *indx, booleantype *newpoint)
{
  static long int ilast;
  DtpntMem *dt_mem;
  int sign;
  booleantype to_left, to_right;

  dt_mem = ca_mem->dt_mem;

  *newpoint = FALSE;

  /* Find the direction of integration */
  sign = (tfinal - tinitial > ZERO) ? 1 : -1;

  /* If this is the first time we use new data */
  if (newData) {
    ilast     = np-1;
    *newpoint = TRUE;
    newData   = FALSE;
  }

  /* Search for indx starting from ilast */
  to_left  = ( sign*(t - dt_mem[ilast-1]->t) < ZERO);
  to_right = ( sign*(t - dt_mem[ilast]->t)   > ZERO);

  if ( to_left ) {
    /* look for a new indx to the left */

    *newpoint = TRUE;
    
    *indx = ilast;
    loop {
      if ( *indx == 0 ) break;
      if ( sign*(t - dt_mem[*indx-1]->t) <= ZERO ) (*indx)--;
      else                                         break;
    }

    if ( *indx == 0 )
      ilast = 1;
    else
      ilast = *indx;

    if ( *indx == 0 ) {
      /* t is beyond leftmost limit. Is it too far? */  
      if ( ABS(t - dt_mem[0]->t) > FUZZ_FACTOR * uround ) return(CV_GETY_BADT);
    }

  } else if ( to_right ) {
    /* look for a new indx to the right */

    *newpoint = TRUE;

    *indx = ilast;
    loop {
      if ( sign*(t - dt_mem[*indx]->t) > ZERO) (*indx)++;
      else                                     break;
    }

    ilast = *indx;


  } else {
    /* ilast is still OK */

    *indx = ilast;

  }

  return(CV_SUCCESS);


}

/*
 * CVadjGetY
 *
 * This routine returns the interpolated forward solution at time t.
 * The user must allocate space for y.
 */

int CVadjGetY(void *cvadj_mem, realtype t, N_Vector y)
{
  int flag;
  CVadjMem ca_mem;

  if (cvadj_mem == NULL) {
    CVProcessError(NULL, CV_ADJMEM_NULL, "CVODEA", "CVadjGetY", MSGAM_NULL_CAMEM);
    return(CV_ADJMEM_NULL);
  }
  ca_mem  = (CVadjMem) cvadj_mem;
  
  flag = getY(ca_mem, t, y);

  return(flag);
}

/* 
 * -----------------------------------------------------------------
 * Functions specific to cubic Hermite interpolation
 * -----------------------------------------------------------------
 */

/*
 * CVAhermiteMalloc
 *
 * This routine allocates memory for storing information at all
 * intermediate points between two consecutive check points. 
 * This data is then used to interpolate the forward solution 
 * at any other time.
 */

static booleantype CVAhermiteMalloc(CVadjMem ca_mem, long int steps)
{
  CVodeMem cv_mem;
  DtpntMem *dt_mem;
  HermiteDataMem content;
  long int i, ii=0;
  booleantype allocOK;

  allocOK = TRUE;

  dt_mem = ca_mem->dt_mem;
  cv_mem = ca_mem->cv_mem;

  for (i=0; i<=steps; i++) {

    content = NULL;
    content = (HermiteDataMem) malloc(sizeof(struct HermiteDataMemRec));
    if (content == NULL) {
      ii = i;
      allocOK = FALSE;
      break;
    }

    content->y = NULL;
    content->y = N_VClone(tempv);
    if (content->y == NULL) {
      free(content); content = NULL;
      ii = i;
      allocOK = FALSE;
      break;
    }

    content->yd = NULL;
    content->yd = N_VClone(tempv);
    if (content->yd == NULL) {
      N_VDestroy(content->y);
      free(content); content = NULL;
      ii = i;
      allocOK = FALSE;
      break;
    }
    
    dt_mem[i]->content = content;

  } 

  if (!allocOK) {
    for (i=0; i<ii; i++) {
      content = (HermiteDataMem) (dt_mem[i]->content);
      N_VDestroy(content->y);
      N_VDestroy(content->yd);
      free(dt_mem[i]->content); dt_mem[i]->content = NULL;
    }
  }

  return(allocOK);
}

/*
 * CVAhermiteFree
 *
 * This routine frees the memory allocated for data storage.
 */

static void CVAhermiteFree(DtpntMem *dt_mem, long int steps)
{
  long int i;
  HermiteDataMem content;

  for (i=0; i<=steps; i++) {
    content = (HermiteDataMem) (dt_mem[i]->content);
    N_VDestroy(content->y);
    N_VDestroy(content->yd);
    free(dt_mem[i]->content); dt_mem[i]->content = NULL;
  }
}

/*
 * CVAhermiteStorePnt ( -> storePnt )
 *
 * This routine stores a new point (y,yd) in the structure d for use
 * in the cubic Hermite interpolation.
 * Note that the time is already stored.
 */

static int CVAhermiteStorePnt(CVodeMem cv_mem, DtpntMem d)
{
  int retval;
  HermiteDataMem content;

  content = (HermiteDataMem) d->content;

  N_VScale(ONE, zn[0], content->y);
  
  if (nst == 0)
    retval = f(tn, content->y, content->yd, f_data);
  else
    N_VScale(ONE/h, zn[1], content->yd);

  return(0);
}

/*
 * CVAhermiteGetY ( -> getY )
 *
 * This routine uses cubic piece-wise Hermite interpolation for 
 * the forward solution vector. 
 * It is typically called by the wrapper routines before calling
 * user provided routines (fB, djacB, bjacB, jtimesB, psolB) but
 * can be directly called by the user through CVadjGetY
 */

static int CVAhermiteGetY(CVadjMem ca_mem, realtype t, N_Vector y)
{
  DtpntMem *dt_mem;
  HermiteDataMem content0, content1;
  realtype t0, t1, delta, factor;
  N_Vector y0, yd0, y1, yd1;
  int flag;
  long int indx;
  booleantype newpoint;

  dt_mem = ca_mem->dt_mem;
  
  /* Get the index in dt_mem */

  flag = CVAfindIndex(ca_mem, t, &indx, &newpoint);
  if (flag != CV_SUCCESS) return(flag);

  /* If we are beyond the left limit but close enough,
     then return y at the left limit. */

  if (indx == 0) {
    content0 = (HermiteDataMem) (dt_mem[0]->content);
    N_VScale(ONE, content0->y, y);
    return(CV_SUCCESS);
  }

  /* Extract stuff from the appropriate data points */

  t0 = dt_mem[indx-1]->t;
  t1 = dt_mem[indx]->t;
  delta = t1 - t0;

  content0 = (HermiteDataMem) (dt_mem[indx-1]->content);
  y0  = content0->y;
  yd0 = content0->yd;

  if (newpoint) {
    
    /* Recompute Y0 and Y1 */

    content1 = (HermiteDataMem) (dt_mem[indx]->content);
    y1  = content1->y;
    yd1 = content1->yd;

    N_VLinearSum(ONE, y1, -ONE, y0, Y0);
    N_VLinearSum(ONE, yd1,  ONE, yd0, Y1);
    N_VLinearSum(delta, Y1, -TWO, Y0, Y1);
    N_VLinearSum(ONE, Y0, -delta, yd0, Y0);

  }

  /* Perform the actual interpolation. */

  factor = t - t0;
  N_VLinearSum(ONE, y0, factor, yd0, y);

  factor = factor/delta;
  factor = factor*factor;
  N_VLinearSum(ONE, y, factor, Y0, y);

  factor = factor*(t-t1)/delta;
  N_VLinearSum(ONE, y, factor, Y1, y);

  return(CV_SUCCESS);
}

/* 
 * -----------------------------------------------------------------
 * Functions specific to Polynomial interpolation
 * -----------------------------------------------------------------
 */

/*
 * CVApolynomialMalloc
 *
 * This routine allocates memory for storing information at all
 * intermediate points between two consecutive check points. 
 * This data is then used to interpolate the forward solution 
 * at any other time.
 */

static booleantype CVApolynomialMalloc(CVadjMem ca_mem, long int steps)
{
  CVodeMem cv_mem;
  DtpntMem *dt_mem;
  PolynomialDataMem content;
  long int i, ii=0;
  booleantype allocOK;

  allocOK = TRUE;

  dt_mem = ca_mem->dt_mem;
  cv_mem = ca_mem->cv_mem;

  for (i=0; i<=steps; i++) {
    content = NULL;
    content = (PolynomialDataMem) malloc(sizeof(struct PolynomialDataMemRec));
    if (content == NULL) {
      ii = i;
      allocOK = FALSE;
      break;
    }

    content->y = NULL;
    content->y = N_VClone(tempv);
    if (content->y == NULL) {
      free(content); content = NULL;
      ii = i;
      allocOK = FALSE;
      break;
    }

    dt_mem[i]->content = content;
  } 

  if (!allocOK) {
    for (i=0; i<ii; i++) {
      content = (PolynomialDataMem) (dt_mem[i]->content);
      N_VDestroy(content->y);
      free(dt_mem[i]->content); dt_mem[i]->content = NULL;
    }
  }

  return(allocOK);

}

/*
 * CVApolynomialFree
 *
 * This routine frees the memeory allocated for data storage.
 */

static void CVApolynomialFree(DtpntMem *dt_mem, long int steps)
{
  long int i;
  PolynomialDataMem content;

  for (i=0; i<=steps; i++) {
    content = (PolynomialDataMem) (dt_mem[i]->content);
    N_VDestroy(content->y);
    free(dt_mem[i]->content); dt_mem[i]->content = NULL;
  }
}

/*
 * CVApolynomialStorePnt ( -> storePnt )
 *
 * This routine stores a new point y in the structure d for use
 * in the Polynomial interpolation.
 * Note that the time is already stored.
 */

static int CVApolynomialStorePnt(CVodeMem cv_mem, DtpntMem d)
{
  PolynomialDataMem content;

  content = (PolynomialDataMem) d->content;

  N_VScale(ONE, zn[0], content->y);
  content->order = qu;

  return(0);
}

/*
 * CVApolynomialGetY ( -> getY )
 *
 * This routine uses polynomial interpolation for the forward solution vector. 
 * It is typically called by the wrapper routines before calling
 * user provided routines (fB, djacB, bjacB, jtimesB, psolB)) but
 * can be directly called by the user through CVadjGetY.
 */

static int CVApolynomialGetY(CVadjMem ca_mem, realtype t, N_Vector y)
{
  DtpntMem *dt_mem;
  PolynomialDataMem content;
  int flag, dir, order, i, j;
  long int indx, base;
  booleantype newpoint;
  realtype dt, factor;

  dt_mem = ca_mem->dt_mem;
  
  /* Get the index in dt_mem */

  flag = CVAfindIndex(ca_mem, t, &indx, &newpoint);
  if (flag != CV_SUCCESS) return(flag);

  /* If we are beyond the left limit but close enough,
     then return y at the left limit. */

  if (indx == 0) {
    content = (PolynomialDataMem) (dt_mem[0]->content);
    N_VScale(ONE, content->y, y);
    return(CV_SUCCESS);
  }

  /* Scaling factor */

  dt = ABS(dt_mem[indx]->t - dt_mem[indx-1]->t);

  /* Find the direction of the forward integration */

  dir = (tfinal - tinitial > ZERO) ? 1 : -1;

  /* Establish the base point depending on the integration direction.
     Modify the base if there are not enough points for the current order */

  if (dir == 1) {
    base = indx;
    content = (PolynomialDataMem) (dt_mem[base]->content);
    order = content->order;
    if(indx < order) base += order-indx;
  } else {
    base = indx-1;
    content = (PolynomialDataMem) (dt_mem[base]->content);
    order = content->order;
    if (np-indx > order) base -= indx+order-np;
  }

  /* Recompute Y (divided differences for Newton polynomial) if needed */

  if (newpoint) {

    /* Store 0-th order DD */
    if (dir == 1) {
      for(j=0;j<=order;j++) {
        T[j] = dt_mem[base-j]->t;
        content = (PolynomialDataMem) (dt_mem[base-j]->content);
        N_VScale(ONE, content->y, Y[j]);
      }
    } else {
      for(j=0;j<=order;j++) {
        T[j] = dt_mem[base-1+j]->t;
        content = (PolynomialDataMem) (dt_mem[base-1+j]->content);
        N_VScale(ONE, content->y, Y[j]);
      }
    }

    /* Compute higher-order DD */
    for(i=1;i<=order;i++) {
      for(j=order;j>=i;j--) {
        factor = dt/(T[j]-T[j-i]);
        N_VLinearSum(factor, Y[j], -factor, Y[j-1], Y[j]);
      }
    }
  }

  /* Perform the actual interpolation using nested multiplications */

  N_VScale(ONE, Y[order], y);
  for (i=order-1; i>=0; i--) {
    factor = (t-T[i])/dt;
    N_VLinearSum(factor, y, ONE, Y[i], y);
  }

  return(CV_SUCCESS);

}

/* 
 * =================================================================
 * WRAPPERS FOR ADJOINT SYSTEM
 * =================================================================
 */
/*
 * CVArhs
 *
 * This routine interfaces to the CVRhsFnB routine provided by
 * the user.
 * NOTE: f_data actually contains cvadj_mem
 */

static int CVArhs(realtype t, N_Vector yB, 
                  N_Vector yBdot, void *cvadj_mem)
{
  CVadjMem ca_mem;
  CVodeMem cv_mem;
  int flag, retval;

  ca_mem = (CVadjMem) cvadj_mem;
  cv_mem = ca_mem->cvb_mem;

  /* Forward solution from interpolation */
  flag = getY(ca_mem, t, ytmp);
  if (flag != CV_SUCCESS) {
    CVProcessError(cv_mem, -1, "CVODEA", "CVArhs", MSGAM_BAD_T);
    return(-1);
  }

  /* Call user's adjoint RHS routine */
  retval = f_B(t, ytmp, yB, yBdot, f_data_B);

  return(retval);
}

/*
 * CVArhsQ
 *
 * This routine interfaces to the CVQuadRhsFnB routine provided by
 * the user.
 * NOTE: fQ_data actually contains cvadj_mem
 */

static int CVArhsQ(realtype t, N_Vector yB, 
                   N_Vector qBdot, void *cvadj_mem)
{
  CVadjMem ca_mem;
  CVodeMem cv_mem;
  int flag, retval;

  ca_mem = (CVadjMem) cvadj_mem;
  cv_mem = ca_mem->cvb_mem;

  /* Forward solution from interpolation */
  flag = getY(ca_mem, t, ytmp);
  if (flag != CV_SUCCESS) {
    CVProcessError(cv_mem, -1, "CVODEA", "CVArhsQ", MSGAM_BAD_T);
    return(-1);
  }

  /* Call user's adjoint RHS routine */
  retval = fQ_B(t, ytmp, yB, qBdot, fQ_data_B);

  return(retval);
}
```
