## Supplementary scripts for "A multi-scale *in silico* mouse model for insulin resistance and humanoid type 2 diabetes": cvodea_io.c

```
/*
 * -----------------------------------------------------------------
 * $Revision: 749 $
 * $Date: 2011-06-22 14:49:10 +0200 (Wed, 22 Jun 2011) $
 * -----------------------------------------------------------------
 * Programmer: Radu Serban @ LLNL
 * -----------------------------------------------------------------
 * Copyright (c) 2006, The Regents of the University of California.
 * Produced at the Lawrence Livermore National Laboratory.
 * All rights reserved.
 * For details, see the LICENSE file.
 * -----------------------------------------------------------------
 * This is the implementation file for the optional input and output
 * functions for the adjoint module in the CVODES solver.
 * -----------------------------------------------------------------
 */

/* 
 * =================================================================
 * IMPORTED HEADER FILES
 * =================================================================
 */

#include <stdio.h>
#include <stdlib.h>

#include "cvodes_impl.h"
#include <sundials/sundials_types.h>

/* 
 * =================================================================
 * CVODEA PRIVATE CONSTANTS
 * =================================================================
 */

#define ONE         RCONST(1.0) 

/* 
 * =================================================================
 * EXPORTED FUNCTIONS IMPLEMENTATION
 * =================================================================
 */

/* 
 * -----------------------------------------------------------------
 * Readibility Constants
 * -----------------------------------------------------------------
 */

#define interpType  (ca_mem->ca_interpType)
#define f_data_B    (ca_mem->ca_f_dataB)
#define fQ_data_B   (ca_mem->ca_fQ_dataB)
#define t_for_quad  (ca_mem->ca_t_for_quad)
#define ckpntData   (ca_mem->ca_ckpntData)

#define t0_         (ck_mem->ck_t0)
#define t1_         (ck_mem->ck_t1)
#define nst_        (ck_mem->ck_nst)
#define q_          (ck_mem->ck_q)
#define h_          (ck_mem->ck_h)
#define next_       (ck_mem->ck_next)

/* 
 * -----------------------------------------------------------------
 * Optional input functions for backward integration
 * -----------------------------------------------------------------
 */

/*
 * CVodeSet***B
 *
 * Wrappers for the backward phase around the corresponding 
 * CVODES optional input functions
 */

int CVodeSetErrHandlerFnB(void *cvadj_mem, CVErrHandlerFn ehfunB, void *eh_dataB)
{
  CVadjMem ca_mem;
  void *cvode_mem;
  int flag;

  if (cvadj_mem == NULL) {
    CVProcessError(NULL, CV_ADJMEM_NULL, "CVODEA", "CVodeSetErrHandlerB", MSGAM_NULL_CAMEM);
    return(CV_ADJMEM_NULL);
  }
  ca_mem = (CVadjMem) cvadj_mem;
  cvode_mem = (void *)ca_mem->cvb_mem;

  flag = CVodeSetErrHandlerFn(cvode_mem, ehfunB, eh_dataB);

  return(flag);
}

int CVodeSetErrFileB(void *cvadj_mem, FILE *errfpB)
{
  CVadjMem ca_mem;
  void *cvode_mem;
  int flag;

  if (cvadj_mem == NULL) {
    CVProcessError(NULL, CV_ADJMEM_NULL, "CVODEA", "CVodeSetErrFileB", MSGAM_NULL_CAMEM);
    return(CV_ADJMEM_NULL);
  }
  ca_mem = (CVadjMem) cvadj_mem;
  cvode_mem = (void *)ca_mem->cvb_mem;

  flag = CVodeSetErrFile(cvode_mem, errfpB);

  return(flag);
}

int CVodeSetIterTypeB(void *cvadj_mem, int iterB)
{
  CVadjMem ca_mem;
  void *cvode_mem;
  int flag;

  if (cvadj_mem == NULL) {
    CVProcessError(NULL, CV_ADJMEM_NULL, "CVODEA", "CVodeSetIterTypeB", MSGAM_NULL_CAMEM);
    return(CV_ADJMEM_NULL);
  }
  ca_mem = (CVadjMem) cvadj_mem;
  cvode_mem = (void *)ca_mem->cvb_mem;

  flag = CVodeSetIterType(cvode_mem, iterB);
  
  return(flag);
}

int CVodeSetFdataB(void *cvadj_mem, void *f_dataB)
{
  CVadjMem ca_mem;

  if (cvadj_mem == NULL) {
    CVProcessError(NULL, CV_ADJMEM_NULL, "CVODEA", "CVodeSetFdataB", MSGAM_NULL_CAMEM);
    return(CV_ADJMEM_NULL);
  }
  ca_mem = (CVadjMem) cvadj_mem;

  f_data_B = f_dataB;

  return(CV_SUCCESS);
}

int CVodeSetMaxOrdB(void *cvadj_mem, int maxordB)
{
  CVadjMem ca_mem;
  void *cvode_mem;
  int flag;

  if (cvadj_mem == NULL) {
    CVProcessError(NULL, CV_ADJMEM_NULL, "CVODEA", "CVodeSetMaxOrdB", MSGAM_NULL_CAMEM);
    return(CV_ADJMEM_NULL);
  }
  ca_mem = (CVadjMem) cvadj_mem;
  cvode_mem = (void *)ca_mem->cvb_mem;

  flag = CVodeSetMaxOrd(cvode_mem, maxordB);

  return(flag);
}


int CVodeSetMaxNumStepsB(void *cvadj_mem, long int mxstepsB)
{
  CVadjMem ca_mem;
  void *cvode_mem;
  int flag;

  if (cvadj_mem == NULL) {
    CVProcessError(NULL, CV_ADJMEM_NULL, "CVODEA", "CVodeSetMaxNumStepsB", MSGAM_NULL_CAMEM);
    return(CV_ADJMEM_NULL);
  }
  ca_mem = (CVadjMem) cvadj_mem;
  cvode_mem = (void *)ca_mem->cvb_mem;

  flag = CVodeSetMaxNumSteps(cvode_mem, mxstepsB);

  return(flag);
}

int CVodeSetStabLimDetB(void *cvadj_mem, booleantype stldetB)
{
  CVadjMem ca_mem;
  void *cvode_mem;
  int flag;

  if (cvadj_mem == NULL) {
    CVProcessError(NULL, CV_ADJMEM_NULL, "CVODEA", "CVodeSetStabLimDetB", MSGAM_NULL_CAMEM);
    return(CV_ADJMEM_NULL);
  }
  ca_mem = (CVadjMem) cvadj_mem;
  cvode_mem = (void *)ca_mem->cvb_mem;

  flag = CVodeSetStabLimDet(cvode_mem, stldetB);

  return(flag);
}

int CVodeSetInitStepB(void *cvadj_mem, realtype hinB)
{
  CVadjMem ca_mem;
  void *cvode_mem;
  int flag;

  if (cvadj_mem == NULL) {
    CVProcessError(NULL, CV_ADJMEM_NULL, "CVODEA", "CVodeSetInitStepB", MSGAM_NULL_CAMEM);
    return(CV_ADJMEM_NULL);
  }
  ca_mem = (CVadjMem) cvadj_mem;
  cvode_mem = (void *)ca_mem->cvb_mem;

  flag = CVodeSetInitStep(cvode_mem, hinB);

  return(flag);
}

int CVodeSetMinStepB(void *cvadj_mem, realtype hminB)
{
  CVadjMem ca_mem;
  void *cvode_mem;
  int flag;

  if (cvadj_mem == NULL) {
    CVProcessError(NULL, CV_ADJMEM_NULL, "CVODEA", "CVodeSetMinStepB", MSGAM_NULL_CAMEM);
    return(CV_ADJMEM_NULL);
  }
  ca_mem = (CVadjMem) cvadj_mem;
  cvode_mem = (void *)ca_mem->cvb_mem;

  flag = CVodeSetMinStep(cvode_mem, hminB);

  return(flag);
}

int CVodeSetMaxStepB(void *cvadj_mem, realtype hmaxB)
{
  CVadjMem ca_mem;
  void *cvode_mem;
  int flag;

  if (cvadj_mem == NULL) {
    CVProcessError(NULL, CV_ADJMEM_NULL, "CVODEA", "CVodeSetMaxStepB", MSGAM_NULL_CAMEM);
    return(CV_ADJMEM_NULL);
  }
  ca_mem = (CVadjMem) cvadj_mem;
  cvode_mem = (void *)ca_mem->cvb_mem;

  flag = CVodeSetMaxStep(cvode_mem, hmaxB);

  return(flag);
}

/*
 * CVodeSetQuad*B
 *
 * Wrappers for the backward phase around the corresponding 
 * CVODES quadrature optional input functions
 */

int CVodeSetQuadFdataB(void *cvadj_mem, void *fQ_dataB)
{
  CVadjMem ca_mem;

  if (cvadj_mem == NULL) {
    CVProcessError(NULL, CV_ADJMEM_NULL, "CVODEA", "CVodeSetQuadFdataB", MSGAM_NULL_CAMEM);
    return(CV_ADJMEM_NULL);
  }
  ca_mem = (CVadjMem) cvadj_mem;

  fQ_data_B = fQ_dataB;

  return(CV_SUCCESS);
}

int CVodeSetQuadErrConB(void *cvadj_mem, booleantype errconQB,
                        int itolQB, realtype reltolQB, void *abstolQB)
{
  CVadjMem ca_mem;
  void *cvode_mem;
  int flag;

  if (cvadj_mem == NULL) {
    CVProcessError(NULL, CV_ADJMEM_NULL, "CVODEA", "CVodeSetQuadErrConB", MSGAM_NULL_CAMEM);
    return(CV_ADJMEM_NULL);
  }
  ca_mem = (CVadjMem) cvadj_mem;

  cvode_mem = (void *)ca_mem->cvb_mem;

  flag = CVodeSetQuadErrCon(cvode_mem, errconQB, itolQB, reltolQB, abstolQB);

  return(flag);
}

/* 
 * -----------------------------------------------------------------
 * Optional output functions for backward integration
 * -----------------------------------------------------------------
 */

/*
 * CVodeGetQuadB
 */

int CVodeGetQuadB(void *cvadj_mem, N_Vector qB)
{
  CVadjMem ca_mem;
  void *cvode_mem;
  int flag;
  
  if (cvadj_mem == NULL) {
    CVProcessError(NULL, CV_ADJMEM_NULL, "CVODEA", "CVodeGetQuadB", MSGAM_NULL_CAMEM);
    return(CV_ADJMEM_NULL);
  }
  ca_mem  = (CVadjMem) cvadj_mem;
  cvode_mem = (void *) ca_mem->cvb_mem;
  
  flag = CVodeGetQuad(cvode_mem, t_for_quad, qB);

  return(flag);
}

/*
 * CVadjGetCVodeBmem
 *
 * CVadjGetCVodeBmem returns a (void *) pointer to the CVODES     
 * memory allocated for the backward problem. This pointer can    
 * then be used to call any of the CVodeGet* CVODES routines to  
 * extract optional output for the backward integration phase.
 */

void *CVadjGetCVodeBmem(void *cvadj_mem)
{
  CVadjMem ca_mem;
  void *cvode_mem;
  
  if (cvadj_mem == NULL) {
    CVProcessError(NULL, 0, "CVODEA", "CVadjGetCVodeBmem", MSGAM_NULL_CAMEM);
    return(NULL);
  }
  ca_mem  = (CVadjMem) cvadj_mem;
  cvode_mem = (void *) ca_mem->cvb_mem;

  return(cvode_mem);
}

/*
 * CVadjGetReturnFlagName
 *
 * The following function returns the name of the constant 
 * associated with a CVODEA-specific return flag
 */

char *CVadjGetReturnFlagName(int flag)
{
  char *name;

  name = (char *)malloc(30*sizeof(char));

  switch(flag) {
  case CV_SUCCESS:
    sprintf(name,"CV_SUCCESS");
    break;
  case CV_ADJMEM_NULL:
    sprintf(name,"CV_ADJMEM_NULL");
    break;
  case CV_BAD_TB0:
    sprintf(name,"CV_BAD_TB0");
    break;
  case CV_BCKMEM_NULL:
    sprintf(name,"CV_BCKMEM_NULL");
    break;
  case CV_REIFWD_FAIL:
    sprintf(name,"CV_REIFWD_FAIL");
    break;
  case CV_FWD_FAIL:
    sprintf(name,"CV_FWD_FAIL");
    break;
  case CV_BAD_ITASK:
    sprintf(name,"CV_BAD_ITASK");
    break;
  case CV_BAD_TBOUT:
    sprintf(name,"CV_BAD_TBOUT");
    break;
  case CV_GETY_BADT:
    sprintf(name,"CV_GETY_BADT");
    break;
  default:
    sprintf(name,"NONE");
  }

  return(name);
}

/*
 * CVadjGetCheckPointsInfo
 *
 * This routine loads an array of nckpnts structures of type CVadjCheckPointRec.
 * The user must allocate space for ckpnt.
 */

int CVadjGetCheckPointsInfo(void *cvadj_mem, CVadjCheckPointRec *ckpnt)
{
  CVadjMem ca_mem;
  CkpntMem ck_mem;
  int i;

  if (cvadj_mem == NULL) {
    CVProcessError(NULL, CV_ADJMEM_NULL, "CVODEA", "CVadjGetCheckPointsInfo", MSGAM_NULL_CAMEM);
    return(CV_ADJMEM_NULL);
  }
  ca_mem = (CVadjMem) cvadj_mem;
  ck_mem = ca_mem->ck_mem;
  i = 0;

  while (ck_mem != NULL) {

    ckpnt[i].my_addr = (void *) ck_mem;
    ckpnt[i].next_addr = (void *) next_;
    ckpnt[i].t0 = t0_;
    ckpnt[i].t1 = t1_;
    ckpnt[i].nstep = nst_;
    ckpnt[i].order = q_;
    ckpnt[i].step = h_;

    ck_mem = next_;
    i++;

  }

  return(CV_SUCCESS);

}

/*
 * CVadjGetDataPointHermite
 *
 * This routine returns the solution stored in the data structure
 * at the 'which' data point. Cubic Hermite interpolation.
 */

int CVadjGetDataPointHermite(void *cvadj_mem, long int which, 
                             realtype *t, N_Vector y, N_Vector yd)
{
  CVadjMem ca_mem;
  DtpntMem *dt_mem;
  HermiteDataMem content;

  if (cvadj_mem == NULL) {
    CVProcessError(NULL, CV_ADJMEM_NULL, "CVODEA", "CVadjGetDataPointHermite", MSGAM_NULL_CAMEM);
    return(CV_ADJMEM_NULL);
  }
  ca_mem = (CVadjMem) cvadj_mem;
  dt_mem = ca_mem->dt_mem;

  if (interpType != CV_HERMITE) {
    CVProcessError(NULL, CV_ILL_INPUT, "CVODEA", "CVadjGetDataPointHermite", MSGAM_WRONG_INTERP);
    return(CV_ILL_INPUT);
  }

  *t = dt_mem[which]->t;

  content = (HermiteDataMem) (dt_mem[which]->content);

  if (y != NULL)
    N_VScale(ONE, content->y, y);

  if (yd != NULL)
    N_VScale(ONE, content->yd, yd);

  return(CV_SUCCESS);
}

/*
 * CVadjGetDataPointPolynomial
 *
 * This routine returns the solution stored in the data structure
 * at the 'which' data point. Polynomial interpolation.
 */

int CVadjGetDataPointPolynomial(void *cvadj_mem, long int which, 
                                realtype *t, int *order, N_Vector y)
{
  CVadjMem ca_mem;
  DtpntMem *dt_mem;
  PolynomialDataMem content;

  if (cvadj_mem == NULL) {
    CVProcessError(NULL, CV_ADJMEM_NULL, "CVODEA", "CVadjGetDataPointPolynomial", MSGAM_NULL_CAMEM);
    return(CV_ADJMEM_NULL);
  }
  ca_mem = (CVadjMem) cvadj_mem;
  dt_mem = ca_mem->dt_mem;

  if (interpType != CV_POLYNOMIAL) {
    CVProcessError(NULL, CV_ILL_INPUT, "CVODEA", "CVadjGetDataPointPolynomial", MSGAM_WRONG_INTERP);
    return(CV_ILL_INPUT);
  }

  *t = dt_mem[which]->t;

  content = (PolynomialDataMem) (dt_mem[which]->content);

  if (y != NULL)
    N_VScale(ONE, content->y, y);

  *order = content->order;

  return(CV_SUCCESS);
}

/* 
 * -----------------------------------------------------------------
 * UNDOCUMENTED development user-callable functions
 * -----------------------------------------------------------------
 */

/*
 * CVadjGetCurrentCheckPoint
 *
 * Returns the address of the 'active' check point.
 */

int CVadjGetCurrentCheckPoint(void *cvadj_mem, void **addr)
{
  CVadjMem ca_mem;

  if (cvadj_mem == NULL) {
    CVProcessError(NULL, CV_ADJMEM_NULL, "CVODEA", "CVadjGetCurrentCheckPoint", MSGAM_NULL_CAMEM);
    return(CV_ADJMEM_NULL);
  }
  ca_mem = (CVadjMem) cvadj_mem;

  *addr = (void *) ckpntData;

  return(CV_SUCCESS);
}
```
