## Supplementary scripts for "A multi-scale *in silico* mouse model for insulin resistance and humanoid type 2 diabetes": cvodes.c

```
/*
 * -----------------------------------------------------------------
 * $Revision: 749 $
 * $Date: 2011-06-22 14:49:10 +0200 (Wed, 22 Jun 2011) $
 * ----------------------------------------------------------------- 
 * Programmer(s): Alan C. Hindmarsh and Radu Serban @ LLNL
 * -----------------------------------------------------------------
 * Copyright (c) 2006, The Regents of the University of California.
 * Produced at the Lawrence Livermore National Laboratory.
 * All rights reserved.
 * For details, see the LICENSE file.
 * -----------------------------------------------------------------
 * This is the implementation file for the main CVODES integrator
 * with sensitivity analysis capabilities.
 * -----------------------------------------------------------------
 * 
 * EXPORTED FUNCTIONS
 * ------------------
 *
 *   Creation, allocation and re-initialization functions
 *      CVodeCreate
 *      CVodeMalloc 
 *      CVodeReInit
 *      CVodeRootInit      
 *      CVodeQuadMalloc
 *      CVodeQuadReInit   
 *      CVodeSensMalloc
 *      CVodeSensReInit             
 *      CVodeSensToggleOff
 *
 *   Main solver function
 *      CVode
 *
 *   Interpolated output and extraction functions
 *      CVodeGetDky
 *      CVodeGetQuad             
 *      CVodeGetQuadDky
 *      CVodeGetSens             
 *      CVodeGetSens1
 *      CVodeGetSensDky         
 *      CVodeGetSensDky1
 *
 *   Deallocation functions
 *      CVodeFree              
 *      CVodeQuadFree
 *      CVodeSensFree  
 *
 * PRIVATE FUNCTIONS
 * -----------------
 *
 *      CVCheckNvector
 *
 *   Memory allocation/deallocation
 *      CVAllocVectors         
 *      CVFreeVectors
 *      CVQuadAllocVectors      
 *      CVQuadFreeVectors
 *      CVSensAllocVectors     
 *      CVSensFreeVectors
 *
 *   Initial stepsize calculation
 *      CVHin                 
 *      CVUpperBoundH0
 *      CVYddNorm     
 *
 *   Initial setup
 *      CVInitialSetup
 *      CVEwtSet           
 *      CVEwtSetSS
 *      CVEwtSetSV         
 *      CVQuadEwtSet
 *      CVQuadEwtSetSS     
 *      CVQuadEwtSetSV
 *      CVSensEwtSet      
 *      CVSensEwtSetEE
 *      CVSensEwtSetSS    
 *      CVSensEwtSetSV
 *
 *   Main CVStep function
 *      CVStep
 *
 *   Functions called at beginning of step
 *      CVAdjustParams
 *      CVAdjustOrder    
 *      CVAdjustAdams
 *      CVAdjustBDF       
 *      CVIncreaseBDF
 *      CVDecreaseBDF     
 *      CVRescale
 *      CVPredict         
 *      CVSet
 *      CVSetAdams        
 *      CVAdamsStart
 *      CVAdamsFinish      
 *      CVAltSum
 *      CVSetBDF           
 *      CVSetTqBDF
 *
 *   Nonlinear solver functions
 *      CVNls              
 *      CVNlsFunctional
 *      CVNlsNewton        
 *      CVNewtonIteration
 *      CVQuadNls           
 *      CVStgrNls
 *      CVStgrNlsFunctional     
 *      CVStgrNlsNewton
 *      CVStgrNewtonIteration    
 *      CVStgr1Nls
 *      CVStgr1NlsFunctional    
 *      CVStgr1NlsNewton
 *      CVStgr1NewtonIteration   
 *      CVHandleNFlag
 *      CVRestore             
 *
 *   Error Test
 *      CVDoErrorTest
 *
 *   Functions called after a successful step
 *      CVCompleteStep      
 *      CVPrepareNextStep
 *      CVSetEta          
 *      CVComputeEtaqm1
 *      CVComputeEtaqp1   
 *      CVChooseEta
 *
 *   Function to handle failures
 *      CVHandleFailure   
 *
 *   Functions for BDF Stability Limit Detection  
 *      CVBDFStab
 *      CVsldet   
 *
 *   Functions for rootfinding
 *      CVRcheck1
 *      CVRcheck2         
 *      CVRcheck3
 *      CVRootfind  
 *
 *   Functions for combined norms
 *      CVQuadUpdateNorm
 *      CVQuadUpdateDsm     
 *      CVSensNorm
 *      CVSensUpdateNorm    
 *      CVStgrUpdateDsm
 *
 *   Wrappers for sensitivity RHS
 *      CVSensRhs           
 *      CVSensRhs1
 *
 *   Internal DQ approximations for sensitivity RHS
 *      CVSensRhsDQ         
 *      CVSensRhs1DQ
 *
 *   Error message handling functions
 *      CVProcessError      
 *      CVErrHandler
 *
 * -----------------------------------------------------------------
 */

/* 
 * =================================================================
 * IMPORTED HEADER FILES
 * =================================================================
 */

#include <stdio.h>
#include <stdlib.h>
#include <stdarg.h>

#include "cvodes_impl.h"
#include <sundials/sundials_math.h>
#include <sundials/sundials_types.h>

/* 
 * =================================================================
 * MACRO DEFINITIONS
 * =================================================================
 */

/* Macro: loop */
#define loop for(;;)

/* 
 * =================================================================
 * CVODES PRIVATE CONSTANTS
 * =================================================================
 */

#define ZERO   RCONST(0.0)
#define TINY   RCONST(1.0e-10)
#define TENTH  RCONST(0.1)
#define POINT2 RCONST(0.2)
#define FOURTH RCONST(0.25)
#define HALF   RCONST(0.5)
#define ONE    RCONST(1.0)
#define TWO    RCONST(2.0)
#define THREE  RCONST(3.0)
#define FOUR   RCONST(4.0)
#define FIVE   RCONST(5.0)
#define TWELVE RCONST(12.0)
#define HUN    RCONST(100.0)

/* 
 * =================================================================
 * CVODES ROUTINE-SPECIFIC CONSTANTS
 * =================================================================
 */

/* 
 * Control constants for lower-level functions used by CVStep 
 * ----------------------------------------------------------
 *
 * CVHin return values:
 *    CV_SUCCESS,
 *    CV_RHSFUNC_FAIL,  CV_RPTD_RHSFUNC_ERR,
 *    CV_QRHSFUNC_FAIL, CV_RPTD_QRHSFUNC_ERR,
 *    CV_SRHSFUNC_FAIL, CV_RPTD_SRHSFUNC_ERR,
 *    CV_TOO_CLOSE
 *
 * CVStep control constants:
 *    DO_ERROR_TEST
 *    PREDICT_AGAIN
 *
 * CVStep return values: 
 *    CV_SUCCESS,
 *    CV_CONV_FAILURE,      CV_ERR_FAILURE,
 *    CV_LSETUP_FAIL,       CV_LSOLVE_FAIL, 
 *    CV_RTFUNC_FAIL,
 *    CV_RHSFUNC_FAIL,      CV_QRHSFUNC_FAIL,      CV_SRHSFUNC_FAIL,
 *    CV_FIRST_RHSFUNC_ERR, CV_FIRST_QRHSFUNC_ERR, CV_FIRST_SRHSFUNC_ERR
 *    CV_UNREC_RHSFUNC_ERR, CV_UNREC_QRHSFUNC_ERR, CV_UNREC_SRHSFUNC_ERR,
 *    CV_REPTD_RHSFUNC_ERR, CV_REPTD_QRHSFUNC_ERR, CV_REPTD_SRHSFUNC_ERR 
 *
 * CVNls input nflag values:
 *    FIRST_CALL
 *    PREV_CONV_FAIL
 *    PREV_ERR_FAIL
 *    
 * CVNls return values: 
 *    CV_SUCCESS,
 *    CV_LSETUP_FAIL,  CV_LSOLVE_FAIL, 
 *    CV_RHSFUNC_FAIL, CV_SRHSFUNC_FAIL,
 *    CONV_FAIL, 
 *    RHSFUNC_RECVR,   SRHSFUNC_RECVR
 * 
 * CVNewtonIteration return values:
 *    CV_SUCCESS, 
 *    CV_LSOLVE_FAIL, 
 *    CV_RHSFUNC_FAIL, CV_SRHSFUNC_FAIL,
 *    CONV_FAIL,       TRY_AGAIN
 *    RHSFUNC_RECVR,   SRHSFUNC_RECVR
 *
 */

#define DO_ERROR_TEST    +2
#define PREDICT_AGAIN    +3

#define CONV_FAIL        +4 
#define TRY_AGAIN        +5

#define FIRST_CALL       +6
#define PREV_CONV_FAIL   +7
#define PREV_ERR_FAIL    +8

#define RHSFUNC_RECVR    +9

#define QRHSFUNC_RECVR   +11
#define SRHSFUNC_RECVR   +12

/*
 * Control constants for lower-level rootfinding functions
 * -------------------------------------------------------
 *
 * CVRcheck1 return values:
 *    CV_SUCCESS,
 *    CV_RTFUNC_FAIL,
 *    INITROOT
 * CVRcheck2 return values:
 *    CV_SUCCESS,
 *    CV_RTFUNC_FAIL,
 *    CLOSERT,
 *    RTFOUND
 * CVRcheck3 return values:
 *    CV_SUCCESS,
 *    CV_RTFUNC_FAIL,
 *    RTFOUND
 * CVRootFind return values:
 *    CV_SUCCESS,
 *    CV_RTFUNC_FAIL,
 *    RTFOUND
 */

#define RTFOUND          +1
#define INITROOT         +2
#define CLOSERT          +3

/*
 * Control constants for sensitivity DQ
 * ------------------------------------
 */

#define CENTERED1        +1
#define CENTERED2        +2
#define FORWARD1         +3
#define FORWARD2         +4


/*
 * Algorithmic constants
 * ---------------------
 *
 * CVodeGetDky and CVStep
 *
 *    FUZZ_FACTOR fuzz factor used to estimate infinitesimal time intervals
 *
 * CVHin
 *
 *    HLB_FACTOR  factor for upper bound on initial step size
 *    HUB_FACTOR  factor for lower bound on initial step size
 *    H_BIAS      bias factor in selection of intial step size
 *    MAX_ITERS   maximum attempts to compute the initial step size
 *
 * CVodeCreate 
 *
 *   CORTES       constant in nonlinear iteration convergence test
 *
 * CVStep
 *
 *    THRESH      if eta < THRESH reject a change in step size or order
 *    ETAMX1      -+
 *    ETAMX2       |
 *    ETAMX3       |-> bounds for eta (step size change)
 *    ETAMXF       |
 *    ETAMIN       |
 *    ETACF       -+
 *    ADDON       safety factor in computing eta
 *    BIAS1       -+
 *    BIAS2        |-> bias factors in eta selection
 *    BIAS3       -+
 *    ONEPSM      (1+epsilon) used in testing if the step size is below its bound
 *
 *    SMALL_NST   nst > SMALL_NST => use ETAMX3 
 *    MXNCF       max no. of convergence failures during one step try
 *    MXNEF       max no. of error test failures during one step try
 *    MXNEF1      max no. of error test failures before forcing a reduction of order
 *    SMALL_NEF   if an error failure occurs and SMALL_NEF <= nef <= MXNEF1, then
 *                reset eta =  MIN(eta, ETAMXF)
 *    LONG_WAIT   number of steps to wait before considering an order change when
 *                q==1 and MXNEF1 error test failures have occurred
 *
 * CVNls
 *    
 *    NLS_MAXCOR  maximum no. of corrector iterations for the nonlinear solver
 *    CRDOWN      constant used in the estimation of the convergence rate (crate)
 *                of the iterates for the nonlinear equation
 *    DGMAX       iter == CV_NEWTON, |gamma/gammap-1| > DGMAX => call lsetup
 *    RDIV        declare divergence if ratio del/delp > RDIV
 *    MSBP        max no. of steps between lsetup calls
 *    
 */


#define FUZZ_FACTOR RCONST(100.0)

#define HLB_FACTOR RCONST(100.0)
#define HUB_FACTOR RCONST(0.1)
#define H_BIAS     HALF
#define MAX_ITERS  4

#define CORTES RCONST(0.1)

#define THRESH RCONST(1.5)
#define ETAMX1 RCONST(10000.0) 
#define ETAMX2 RCONST(10.0)
#define ETAMX3 RCONST(10.0)
#define ETAMXF RCONST(0.2)
#define ETAMIN RCONST(0.1)
#define ETACF  RCONST(0.25)
#define ADDON  RCONST(0.000001)
#define BIAS1  RCONST(6.0)
#define BIAS2  RCONST(6.0)
#define BIAS3  RCONST(10.0)
#define ONEPSM RCONST(1.000001)

#define SMALL_NST    10
#define MXNCF        10
#define MXNEF         7
#define MXNEF1        3
#define SMALL_NEF     2
#define LONG_WAIT    10

#define NLS_MAXCOR 3
#define CRDOWN RCONST(0.3)
#define DGMAX  RCONST(0.3)

#define RDIV      TWO
#define MSBP       20

/* 
 * =================================================================
 * PRIVATE FUNCTION PROTOTYPES
 * =================================================================
 */

static booleantype CVCheckNvector(N_Vector tmpl);

/* Memory allocation/deallocation */

static booleantype CVAllocVectors(CVodeMem cv_mem, N_Vector tmpl, int tol);
static void CVFreeVectors(CVodeMem cv_mem);

static booleantype CVQuadAllocVectors(CVodeMem cv_mem, N_Vector tmpl);
static void CVQuadFreeVectors(CVodeMem cv_mem);

static booleantype CVSensAllocVectors(CVodeMem cv_mem, N_Vector tmpl);
static void CVSensFreeVectors(CVodeMem cv_mem);

/* Initial stepsize calculation */

static int CVHin(CVodeMem cv_mem, realtype tout);
static realtype CVUpperBoundH0(CVodeMem cv_mem, realtype tdist);
static int CVYddNorm(CVodeMem cv_mem, realtype hg, realtype *yddnrm);

/* Initial setup */

static int CVInitialSetup(CVodeMem cv_mem);

static int CVEwtSetSS(CVodeMem cv_mem, N_Vector ycur, N_Vector weight);
static int CVEwtSetSV(CVodeMem cv_mem, N_Vector ycur, N_Vector weight);

static int CVQuadEwtSet(CVodeMem cv_mem, N_Vector qcur, N_Vector weightQ);
static int CVQuadEwtSetSS(CVodeMem cv_mem, N_Vector qcur, N_Vector weightQ);
static int CVQuadEwtSetSV(CVodeMem cv_mem, N_Vector qcur, N_Vector weightQ);

static int CVSensEwtSet(CVodeMem cv_mem, N_Vector *yScur, N_Vector *weightS);
static int CVSensEwtSetEE(CVodeMem cv_mem, N_Vector *yScur, N_Vector *weightS);
static int CVSensEwtSetSS(CVodeMem cv_mem, N_Vector *yScur, N_Vector *weightS);
static int CVSensEwtSetSV(CVodeMem cv_mem, N_Vector *yScur, N_Vector *weightS);

/* Main CVStep function */

static int CVStep(CVodeMem cv_mem);

/* Function called at beginning of step */

static void CVAdjustParams(CVodeMem cv_mem);
static void CVAdjustOrder(CVodeMem cv_mem, int deltaq);
static void CVAdjustAdams(CVodeMem cv_mem, int deltaq);
static void CVAdjustBDF(CVodeMem cv_mem, int deltaq);
static void CVIncreaseBDF(CVodeMem cv_mem);
static void CVDecreaseBDF(CVodeMem cv_mem);
static void CVRescale(CVodeMem cv_mem);
static void CVPredict(CVodeMem cv_mem);
static void CVSet(CVodeMem cv_mem);
static void CVSetAdams(CVodeMem cv_mem);
static realtype CVAdamsStart(CVodeMem cv_mem, realtype m[]);
static void CVAdamsFinish(CVodeMem cv_mem, realtype m[], realtype M[], realtype hsum);
static realtype CVAltSum(int iend, realtype a[], int k);
static void CVSetBDF(CVodeMem cv_mem);
static void CVSetTqBDF(CVodeMem cv_mem, realtype hsum, realtype alpha0,
                       realtype alpha0_hat, realtype xi_inv, realtype xistar_inv);

/* Nonlinear solver functions */

static int CVNls(CVodeMem cv_mem, int nflag);
static int CVNlsFunctional(CVodeMem cv_mem);
static int CVNlsNewton(CVodeMem cv_mem, int nflag);
static int CVNewtonIteration(CVodeMem cv_mem);

static int CVQuadNls(CVodeMem cv_mem);

static int CVStgrNls(CVodeMem cv_mem);
static int CVStgrNlsFunctional(CVodeMem cv_mem);
static int CVStgrNlsNewton(CVodeMem cv_mem);
static int CVStgrNewtonIteration(CVodeMem cv_mem);

static int CVStgr1Nls(CVodeMem cv_mem, int is);
static int CVStgr1NlsFunctional(CVodeMem cv_mem, int is);
static int CVStgr1NlsNewton(CVodeMem cv_mem, int is);
static int CVStgr1NewtonIteration(CVodeMem cv_mem, int is);

static int CVHandleNFlag(CVodeMem cv_mem, int *nflagPtr, realtype saved_t,
                         int *ncfPtr, long int *ncfnPtr);

static void CVRestore(CVodeMem cv_mem, realtype saved_t);

/* Error Test */

static int CVDoErrorTest(CVodeMem cv_mem, int *nflagPtr, realtype saved_t, 
                         realtype acor_nrm,
                         int *nefPtr, long int *netfPtr, realtype *dsmPtr);

/* Function called after a successful step */

static void CVCompleteStep(CVodeMem cv_mem);
static void CVPrepareNextStep(CVodeMem cv_mem, realtype dsm);
static void CVSetEta(CVodeMem cv_mem);
static realtype CVComputeEtaqm1(CVodeMem cv_mem);
static realtype CVComputeEtaqp1(CVodeMem cv_mem);
static void CVChooseEta(CVodeMem cv_mem);

/* Function to handle failures */

static int CVHandleFailure(CVodeMem cv_mem,int flag);

/* Functions for BDF Stability Limit Detection */

static void CVBDFStab(CVodeMem cv_mem);
static int CVsldet(CVodeMem cv_mem);

/* Functions for rootfinding */

static int CVRcheck1(CVodeMem cv_mem);
static int CVRcheck2(CVodeMem cv_mem);
static int CVRcheck3(CVodeMem cv_mem);
static int CVRootfind(CVodeMem cv_mem);

/* Function for combined norms */

static realtype CVQuadUpdateNorm(CVodeMem cv_mem, realtype old_nrm,
                                 N_Vector xQ, N_Vector wQ);
static realtype CVQuadUpdateDsm(CVodeMem cv_mem, realtype old_dsm, 
                                realtype dsmQ);
static realtype CVSensNorm(CVodeMem cv_mem, N_Vector *xS, N_Vector *wS);
static realtype CVSensUpdateNorm(CVodeMem cv_mem, realtype old_nrm,
                                 N_Vector *xS, N_Vector *wS);
static realtype CVStgrUpdateDsm(CVodeMem cv_mem, realtype old_dsm, 
                                realtype dsmS);

/* Wrappers for sensitivity RHS */

static int CVSensRhs(CVodeMem cv_mem, realtype time, 
                     N_Vector ycur, N_Vector fcur, 
                     N_Vector *yScur, N_Vector *fScur,
                     N_Vector temp1, N_Vector temp2);
static int CVSensRhs1(CVodeMem cv_mem, realtype time, 
                      N_Vector ycur, N_Vector fcur, 
                      int is, N_Vector yScur, N_Vector fScur,
                      N_Vector temp1, N_Vector temp2);

/* 
 * =================================================================
 * EXPORTED FUNCTIONS IMPLEMENTATION
 * =================================================================
 */

/* 
 * -----------------------------------------------------------------
 * Creation, allocation and re-initialization functions
 * -----------------------------------------------------------------
 */

/* 
 * CVodeCreate
 *
 * CVodeCreate creates an internal memory block for a problem to 
 * be solved by CVODES.
 * If successful, CVodeCreate returns a pointer to the problem memory. 
 * This pointer should be passed to CVodeMalloc.  
 * If an initialization error occurs, CVodeCreate prints an error 
 * message to standard err and returns NULL. 
 */

void *CVodeCreate(int lmm, int iter)
{
  int maxord;
  CVodeMem cv_mem;

  /* Test inputs */

  if ((lmm != CV_ADAMS) && (lmm != CV_BDF)) {
    CVProcessError(NULL, 0, "CVODES", "CVodeCreate", MSGCV_BAD_LMM);
    return(NULL);
  }
  
  if ((iter != CV_FUNCTIONAL) && (iter != CV_NEWTON)) {
    CVProcessError(NULL, 0, "CVODES", "CVodeCreate", MSGCV_BAD_ITER);
    return(NULL);
  }

  cv_mem = NULL;
  cv_mem = (CVodeMem) malloc(sizeof(struct CVodeMemRec));
  if (cv_mem == NULL) {
    CVProcessError(NULL, 0, "CVODES", "CVodeCreate", MSGCV_CVMEM_FAIL);
    return(NULL);
  }

  maxord = (lmm == CV_ADAMS) ? ADAMS_Q_MAX : BDF_Q_MAX;

  /* copy input parameters into cv_mem */
  cv_mem->cv_lmm  = lmm;
  cv_mem->cv_iter = iter;

  /* Set uround */
  cv_mem->cv_uround = UNIT_ROUNDOFF;

  /* Set default values for integrator optional inputs */
  cv_mem->cv_f        = NULL;
  cv_mem->cv_f_data   = NULL;
  cv_mem->cv_efun     = NULL;
  cv_mem->cv_e_data   = NULL;
  cv_mem->cv_ehfun    = CVErrHandler;
  cv_mem->cv_eh_data  = (void *) cv_mem;
  cv_mem->cv_errfp    = stderr;
  cv_mem->cv_qmax     = maxord;
  cv_mem->cv_mxstep   = MXSTEP_DEFAULT;
  cv_mem->cv_mxhnil   = MXHNIL_DEFAULT;
  cv_mem->cv_sldeton  = FALSE;
  cv_mem->cv_hin      = ZERO;
  cv_mem->cv_hmin     = HMIN_DEFAULT;
  cv_mem->cv_hmax_inv = HMAX_INV_DEFAULT;
  cv_mem->cv_tstopset = FALSE;
  cv_mem->cv_maxcor   = NLS_MAXCOR;
  cv_mem->cv_maxnef   = MXNEF;
  cv_mem->cv_maxncf   = MXNCF;
  cv_mem->cv_nlscoef  = CORTES;

  /* Initialize root finding variables */

  cv_mem->cv_glo    = NULL;
  cv_mem->cv_ghi    = NULL;
  cv_mem->cv_grout  = NULL;
  cv_mem->cv_iroots = NULL;
  cv_mem->cv_gfun   = NULL;
  cv_mem->cv_g_data = NULL;
  cv_mem->cv_nrtfn  = 0;  

  /* Set default values for quad. optional inputs */
  cv_mem->cv_quadr    = FALSE;
  cv_mem->cv_fQ       = NULL;
  cv_mem->cv_fQ_data  = NULL;
  cv_mem->cv_errconQ  = FALSE;

  /* Set default values for sensi. optional inputs */
  cv_mem->cv_sensi        = FALSE;
  cv_mem->cv_fS_data      = (void *)cv_mem;
  cv_mem->cv_fS           = CVSensRhsDQ;
  cv_mem->cv_fS1          = CVSensRhs1DQ;
  cv_mem->cv_fSDQ         = TRUE;
  cv_mem->cv_ifS          = CV_ONESENS;
  cv_mem->cv_DQtype       = CV_CENTERED;
  cv_mem->cv_DQrhomax     = ZERO;
  cv_mem->cv_p            = NULL;
  cv_mem->cv_pbar         = NULL;
  cv_mem->cv_plist        = NULL;
  cv_mem->cv_errconS      = FALSE;
  cv_mem->cv_maxcorS      = NLS_MAXCOR;
  cv_mem->cv_ncfS1        = NULL;
  cv_mem->cv_ncfnS1       = NULL;
  cv_mem->cv_nniS1        = NULL;
  cv_mem->cv_itolS        = CV_EE;

  /* Set the saved values for qmax_alloc */

  cv_mem->cv_qmax_alloc  = maxord;
  cv_mem->cv_qmax_allocQ = maxord;
  cv_mem->cv_qmax_allocS = maxord;

  /* Initialize lrw and liw */

  cv_mem->cv_lrw = 65 + 2*L_MAX + NUM_TESTS;
  cv_mem->cv_liw = 52;

  /* No mallocs have been done yet */

  cv_mem->cv_VabstolMallocDone  = FALSE;
  cv_mem->cv_MallocDone         = FALSE;

  cv_mem->cv_VabstolQMallocDone = FALSE;
  cv_mem->cv_quadMallocDone     = FALSE;

  cv_mem->cv_VabstolSMallocDone = FALSE;
  cv_mem->cv_SabstolSMallocDone = FALSE;
  cv_mem->cv_sensMallocDone     = FALSE;

  /* Return pointer to CVODES memory block */

  return((void *)cv_mem);
}

/*-----------------------------------------------------------------*/

#define iter (cv_mem->cv_iter)  
#define lmm  (cv_mem->cv_lmm) 
#define lrw  (cv_mem->cv_lrw)
#define liw  (cv_mem->cv_liw)

/*-----------------------------------------------------------------*/

/*
 * CVodeMalloc
 * 
 * CVodeMalloc allocates and initializes memory for a problem. All 
 * problem inputs are checked for errors. If any error occurs during 
 * initialization, it is reported to the file whose file pointer is 
 * errfp and an error flag is returned. Otherwise, it returns CV_SUCCESS
 */

int CVodeMalloc(void *cvode_mem, CVRhsFn f, realtype t0, N_Vector y0, 
                int itol, realtype reltol, void *abstol)
{
  CVodeMem cv_mem;
  booleantype nvectorOK, allocOK, neg_abstol;
  long int lrw1, liw1;
  int i,k;

  /* Check cvode_mem */

  if (cvode_mem==NULL) {
    CVProcessError(NULL, CV_MEM_NULL, "CVODES", "CVodeMalloc", MSGCV_NO_MEM);
    return(CV_MEM_NULL);
  }
  cv_mem = (CVodeMem) cvode_mem;

  /* Check for legal input parameters */

  if (y0==NULL) {
    CVProcessError(cv_mem, CV_ILL_INPUT, "CVODES", "CVodeMalloc", MSGCV_NULL_Y0);
    return(CV_ILL_INPUT);
  }

  if ((itol != CV_SS) && (itol != CV_SV) && (itol != CV_WF)) {
    CVProcessError(cv_mem, CV_ILL_INPUT, "CVODES", "CVodeMalloc", MSGCV_BAD_ITOL);
    return(CV_ILL_INPUT);
  }

  if (f == NULL) {
    CVProcessError(cv_mem, CV_ILL_INPUT, "CVODES", "CVodeMalloc", MSGCV_NULL_F);
    return(CV_ILL_INPUT);
  }

  /* Test if all required vector operations are implemented */

  nvectorOK = CVCheckNvector(y0);
  if(!nvectorOK) {
    CVProcessError(cv_mem, CV_ILL_INPUT, "CVODES", "CVodeMalloc", MSGCV_BAD_NVECTOR);
    return(CV_ILL_INPUT);
  }

  /* Test tolerances */

  if (itol != CV_WF) {

    if (abstol == NULL) {
      CVProcessError(cv_mem, CV_ILL_INPUT, "CVODES", "CVodeMalloc", MSGCV_NULL_ABSTOL);
      return(CV_ILL_INPUT);
    }

    if (reltol < ZERO) {
      CVProcessError(cv_mem, CV_ILL_INPUT, "CVODES", "CVodeMalloc", MSGCV_BAD_RELTOL);
      return(CV_ILL_INPUT);
    }

    if (itol == CV_SS)
      neg_abstol = (*((realtype *)abstol) < ZERO);
    else
      neg_abstol = (N_VMin((N_Vector)abstol) < ZERO);

    if (neg_abstol) {
      CVProcessError(cv_mem, CV_ILL_INPUT, "CVODES", "CVodeMalloc", MSGCV_BAD_ABSTOL);
      return(CV_ILL_INPUT);
    }

  }

  /* Set space requirements for one N_Vector */

  if (y0->ops->nvspace != NULL) {
    N_VSpace(y0, &lrw1, &liw1);
  } else {
    lrw1 = 0;
    liw1 = 0;
  }
  cv_mem->cv_lrw1 = lrw1;
  cv_mem->cv_liw1 = liw1;

  /* Allocate the vectors (using y0 as a template) */

  allocOK = CVAllocVectors(cv_mem, y0, itol);
  if (!allocOK) {
    CVProcessError(cv_mem, CV_MEM_FAIL, "CVODES", "CVodeMalloc", MSGCV_MEM_FAIL);
    return(CV_MEM_FAIL);
  }

  /* Copy tolerances into memory */

  cv_mem->cv_itol   = itol;
  cv_mem->cv_reltol = reltol;      

  if (itol == CV_SS)
    cv_mem->cv_Sabstol = *((realtype *)abstol);
  else if (itol == CV_SV)
    N_VScale(ONE, (N_Vector)abstol, cv_mem->cv_Vabstol);

  /* All error checking is complete at this point */

  /* Copy the input parameters into CVODES state */

  cv_mem->cv_f  = f;
  cv_mem->cv_tn = t0;

  /* Set step parameters */

  cv_mem->cv_q      = 1;
  cv_mem->cv_L      = 2;
  cv_mem->cv_qwait  = cv_mem->cv_L;
  cv_mem->cv_etamax = ETAMX1;

  cv_mem->cv_qu     = 0;
  cv_mem->cv_hu     = ZERO;
  cv_mem->cv_tolsf  = ONE;

  /* Set the linear solver addresses to NULL.
     (We check != NULL later, in CVode, if using CV_NEWTON.) */

  cv_mem->cv_linit  = NULL;
  cv_mem->cv_lsetup = NULL;
  cv_mem->cv_lsolve = NULL;
  cv_mem->cv_lfree  = NULL;
  cv_mem->cv_lmem   = NULL;

  /* Set forceSetup to FALSE */

  cv_mem->cv_forceSetup = FALSE;

  /* Initialize zn[0] in the history array */

  N_VScale(ONE, y0, cv_mem->cv_zn[0]);

  /* Initialize all the counters */

  cv_mem->cv_nst     = 0;
  cv_mem->cv_nfe     = 0;
  cv_mem->cv_ncfn    = 0;
  cv_mem->cv_netf    = 0;
  cv_mem->cv_nni     = 0;
  cv_mem->cv_nsetups = 0;
  cv_mem->cv_nhnil   = 0;
  cv_mem->cv_nstlp   = 0;
  cv_mem->cv_nscon   = 0;
  cv_mem->cv_nge     = 0;

  /* Initialize other integrator optional outputs */

  cv_mem->cv_h0u      = ZERO;
  cv_mem->cv_next_h   = ZERO;
  cv_mem->cv_next_q   = 0;

  /* Initialize Stablilty Limit Detection data */
  /* NOTE: We do this even if stab lim det was not
     turned on yet. This way, the user can turn it
     on at any time */

  cv_mem->cv_nor = 0;
  for (i = 1; i <= 5; i++)
    for (k = 1; k <= 3; k++) 
      cv_mem->cv_ssdat[i-1][k-1] = ZERO;

  /* Problem has been successfully initialized */

  cv_mem->cv_MallocDone = TRUE;

  return(CV_SUCCESS);
}

/*-----------------------------------------------------------------*/

#define lrw1 (cv_mem->cv_lrw1)
#define liw1 (cv_mem->cv_liw1)

/*-----------------------------------------------------------------*/

/*
 * CVodeReInit
 *
 * CVodeReInit re-initializes CVODES's memory for a problem, assuming
 * it has already been allocated in a prior CVodeMalloc call.
 * All problem specification inputs are checked for errors.
 * If any error occurs during initialization, it is reported to the
 * file whose file pointer is errfp.
 * The return value is CV_SUCCESS = 0 if no errors occurred, or
 * a negative value otherwise.
 */

int CVodeReInit(void *cvode_mem, CVRhsFn f, realtype t0, N_Vector y0, 
                int itol, realtype reltol, void *abstol)
{
  CVodeMem cv_mem;
  booleantype neg_abstol;
  int i,k;
 
  /* Check cvode_mem */

  if (cvode_mem==NULL) {
    CVProcessError(NULL, CV_MEM_NULL, "CVODES", "CVodeReInit", MSGCV_NO_MEM);
    return(CV_MEM_NULL);
  }
  cv_mem = (CVodeMem) cvode_mem;

  /* Check if cvode_mem was allocated */

  if (cv_mem->cv_MallocDone == FALSE) {
    CVProcessError(cv_mem, CV_NO_MALLOC, "CVODES", "CVodeReInit", MSGCV_NO_MALLOC);
    return(CV_NO_MALLOC);
  }

  /* Check for legal input parameters */

  if (y0 == NULL) {
    CVProcessError(cv_mem, CV_ILL_INPUT, "CVODES", "CVodeReInit", MSGCV_NULL_Y0);
    return(CV_ILL_INPUT);
  }
  
  if ((itol != CV_SS) && (itol != CV_SV) && (itol != CV_WF)) {
    CVProcessError(cv_mem, CV_ILL_INPUT, "CVODES", "CVodeReInit", MSGCV_BAD_ITOL);
    return(CV_ILL_INPUT);
  }

  if (f == NULL) {
    CVProcessError(cv_mem, CV_ILL_INPUT, "CVODES", "CVodeReInit", MSGCV_NULL_F);
    return(CV_ILL_INPUT);
  }

  /* Test tolerances */
   
  if (itol != CV_WF) {
    
    if (abstol == NULL) {
      CVProcessError(cv_mem, CV_ILL_INPUT, "CVODES", "CVodeReInit", MSGCV_NULL_ABSTOL);
      return(CV_ILL_INPUT);
    }

    if (reltol < ZERO) {
      CVProcessError(cv_mem, CV_ILL_INPUT, "CVODES", "CVodeReInit", MSGCV_BAD_RELTOL);
      return(CV_ILL_INPUT);
    }

    if (itol == CV_SS)
      neg_abstol = (*((realtype *)abstol) < ZERO);
    else
      neg_abstol = (N_VMin((N_Vector)abstol) < ZERO);

    if (neg_abstol) {
      CVProcessError(cv_mem, CV_ILL_INPUT, "CVODES", "CVodeReInit", MSGCV_BAD_ABSTOL);
      return(CV_ILL_INPUT);
    }

  }

  /* Copy tolerances into memory */

  if ( (itol != CV_SV) && (cv_mem->cv_VabstolMallocDone) ) {
    N_VDestroy(cv_mem->cv_Vabstol);
    lrw -= lrw1;
    liw -= liw1;
    cv_mem->cv_VabstolMallocDone = FALSE;
  }

  if ( (itol == CV_SV) && !(cv_mem->cv_VabstolMallocDone) ) {
    cv_mem->cv_Vabstol = NULL;
    cv_mem->cv_Vabstol = N_VClone(y0);
    lrw += lrw1;
    liw += liw1;
    cv_mem->cv_VabstolMallocDone = TRUE;
  }

  cv_mem->cv_itol   = itol;
  cv_mem->cv_reltol = reltol;

  if (itol == CV_SS)
    cv_mem->cv_Sabstol = *((realtype *)abstol);
  else if (itol == CV_SV)
    N_VScale(ONE, (N_Vector)abstol, cv_mem->cv_Vabstol);

  /* All error checking is complete at this point */
  
  /* Copy the input parameters into CVODES state */

  cv_mem->cv_f = f;
  cv_mem->cv_tn = t0;

  /* Set step parameters */

  cv_mem->cv_q      = 1;
  cv_mem->cv_L      = 2;
  cv_mem->cv_qwait  = cv_mem->cv_L;
  cv_mem->cv_etamax = ETAMX1;

  cv_mem->cv_qu     = 0;
  cv_mem->cv_hu     = ZERO;
  cv_mem->cv_tolsf  = ONE;

  /* Set forceSetup to FALSE */

  cv_mem->cv_forceSetup = FALSE;

  /* Initialize zn[0] in the history array */

  N_VScale(ONE, y0, cv_mem->cv_zn[0]);
 
  /* Initialize all the counters */

  cv_mem->cv_nst     = 0;
  cv_mem->cv_nfe     = 0;
  cv_mem->cv_ncfn    = 0;
  cv_mem->cv_netf    = 0;
  cv_mem->cv_nni     = 0;
  cv_mem->cv_nsetups = 0;
  cv_mem->cv_nhnil   = 0;
  cv_mem->cv_nstlp   = 0;
  cv_mem->cv_nscon   = 0;
  cv_mem->cv_nge     = 0;

  /* Initialize other integrator optional outputs */

  cv_mem->cv_h0u      = ZERO;
  cv_mem->cv_next_h   = ZERO;
  cv_mem->cv_next_q   = 0;

  /* Initialize Stablilty Limit Detection data */

  cv_mem->cv_nor = 0;
  for (i = 1; i <= 5; i++)
    for (k = 1; k <= 3; k++) 
      cv_mem->cv_ssdat[i-1][k-1] = ZERO;
  
  /* Problem has been successfully re-initialized */

  return(CV_SUCCESS);
}

/*-----------------------------------------------------------------*/

#define gfun   (cv_mem->cv_gfun)
#define g_data (cv_mem->cv_g_data) 
#define glo    (cv_mem->cv_glo)
#define ghi    (cv_mem->cv_ghi)
#define grout  (cv_mem->cv_grout)
#define iroots (cv_mem->cv_iroots)

/*-----------------------------------------------------------------*/

/*
 * CVodeRootInit
 *
 * CVodeRootInit initializes a rootfinding problem to be solved
 * during the integration of the ODE system.  It loads the root
 * function pointer and the number of root functions, and allocates
 * workspace memory.  The return value is CV_SUCCESS = 0 if no errors
 * occurred, or a negative value otherwise.
 */

int CVodeRootInit(void *cvode_mem, int nrtfn, CVRootFn g, void *gdata)
{
  CVodeMem cv_mem;
  int nrt;

  /* Check cvode_mem */
  if (cvode_mem==NULL) {
    CVProcessError(NULL, CV_MEM_NULL, "CVODES", "CVodeRootInit", MSGCV_NO_MEM);
    return(CV_MEM_NULL);
  }
  cv_mem = (CVodeMem) cvode_mem;

  nrt = (nrtfn < 0) ? 0 : nrtfn;

  /* If rerunning CVodeRootInit() with a different number of root
     functions (changing number of gfun components), then free
     currently held memory resources */
  if ((nrt != cv_mem->cv_nrtfn) && (cv_mem->cv_nrtfn > 0)) {
    free(glo); glo = NULL;
    free(ghi); ghi = NULL;
    free(grout); grout = NULL;
    free(iroots); iroots = NULL;

    lrw -= 3* (cv_mem->cv_nrtfn);
    liw -= cv_mem->cv_nrtfn;

  }

  /* If CVodeRootInit() was called with nrtfn == 0, then set cv_nrtfn to
     zero and cv_gfun to NULL before returning */
  if (nrt == 0) {
    cv_mem->cv_nrtfn = nrt;
    gfun = NULL;
    g_data = NULL;
    return(CV_SUCCESS);
  }

  /* Store user's data pointer */
  g_data = gdata;

  /* If rerunning CVodeRootInit() with the same number of root functions
     (not changing number of gfun components), then check if the root
     function argument has changed */
  /* If g != NULL then return as currently reserved memory resources
     will suffice */
  if (nrt == cv_mem->cv_nrtfn) {
    if (g != gfun) {
      if (g == NULL) {
	free(glo); glo = NULL;
	free(ghi); ghi = NULL;
	free(grout); grout = NULL;
	free(iroots); iroots = NULL;
        
        lrw -= 3*nrt;
        liw -= nrt;

        CVProcessError(cv_mem, CV_ILL_INPUT, "CVODES", "CVodeRootInit", MSGCV_NULL_G);
	return(CV_ILL_INPUT);
      }
      else {
	gfun = g;
	return(CV_SUCCESS);
      }
    }
    else return(CV_SUCCESS);
  }

  /* Set variable values in CVode memory block */
  cv_mem->cv_nrtfn = nrt;
  if (g == NULL) {
    CVProcessError(cv_mem, CV_ILL_INPUT, "CVODES", "CVodeRootInit", MSGCV_NULL_G);
    return(CV_ILL_INPUT);
  }
  else gfun = g;

  /* Allocate necessary memory and return */
  glo = NULL;
  glo = (realtype *) malloc(nrt*sizeof(realtype));
  if (glo == NULL) {
    CVProcessError(cv_mem, CV_MEM_FAIL, "CVODES", "CVodeRootInit", MSGCV_MEM_FAIL);
    return(CV_MEM_FAIL);
  }
    
  ghi = NULL;
  ghi = (realtype *) malloc(nrt*sizeof(realtype));
  if (ghi == NULL) {
    free(glo); glo = NULL;
    CVProcessError(cv_mem, CV_MEM_FAIL, "CVODES", "CVodeRootInit", MSGCV_MEM_FAIL);
    return(CV_MEM_FAIL);
  }
    
  grout = NULL;
  grout = (realtype *) malloc(nrt*sizeof(realtype));
  if (grout == NULL) {
    free(glo); glo = NULL;
    free(ghi); ghi = NULL;
    CVProcessError(cv_mem, CV_MEM_FAIL, "CVODES", "CVodeRootInit", MSGCV_MEM_FAIL);
    return(CV_MEM_FAIL);
  }

  iroots = NULL;
  iroots = (int *) malloc(nrt*sizeof(int));
  if (iroots == NULL) {
    free(glo); glo = NULL;
    free(ghi); ghi = NULL;
    free(grout); grout = NULL;
    CVProcessError(cv_mem, CV_MEM_FAIL, "CVODES", "CVodeRootInit", MSGCV_MEM_FAIL);
    return(CV_MEM_FAIL);
  }

  lrw += 3*nrt;
  liw += nrt;

  return(CV_SUCCESS);
}

/*
 * CVodeQuadMalloc
 *
 * CVodeQuadMalloc allocates and initializes quadrature related 
 * memory for a problem. All problem specification inputs are 
 * checked for errors. If any error occurs during initialization, 
 * it is reported to the file whose file pointer is errfp. 
 * The return value is CV_SUCCESS = 0 if no errors occurred, or
 * a negative value otherwise.
 */

int CVodeQuadMalloc(void *cvode_mem, CVQuadRhsFn fQ, N_Vector yQ0)
{
  CVodeMem cv_mem;
  booleantype allocOK;
  long int lrw1Q, liw1Q;

  /* Check cvode_mem */
  if (cvode_mem==NULL) {
    CVProcessError(NULL, CV_MEM_NULL, "CVODES", "CVodeQuadMalloc", MSGCV_NO_MEM);
    return(CV_MEM_NULL);
  }
  cv_mem = (CVodeMem) cvode_mem;

  /* Set space requirements for one N_Vector */
  N_VSpace(yQ0, &lrw1Q, &liw1Q);
  cv_mem->cv_lrw1Q = lrw1Q;
  cv_mem->cv_liw1Q = liw1Q;

  /* Allocate the vectors (using yQ0 as a template) */
  allocOK = CVQuadAllocVectors(cv_mem, yQ0);
  if (!allocOK) {
    CVProcessError(cv_mem, CV_MEM_FAIL, "CVODES", "CVodeQuadMalloc", MSGCV_MEM_FAIL);
    return(CV_MEM_FAIL);
  }

  /* Initialize znQ[0] in the history array */
  N_VScale(ONE, yQ0, cv_mem->cv_znQ[0]);

  /* Copy the input parameters into CVODES state */
  cv_mem->cv_fQ = fQ;

  /* Initialize counters */
  cv_mem->cv_nfQe  = 0;
  cv_mem->cv_netfQ = 0;

  /* Quadrature integration turned ON */
  cv_mem->cv_quadr = TRUE;
  cv_mem->cv_quadMallocDone = TRUE;

  /* Quadrature initialization was successfull */
  return(CV_SUCCESS);
}

/*-----------------------------------------------------------------*/

#define lrw1Q (cv_mem->cv_lrw1Q)
#define liw1Q (cv_mem->cv_liw1Q)

/*-----------------------------------------------------------------*/

/*
 * CVodeQuadReInit
 *
 * CVodeQuadReInit re-initializes CVODES's quadrature related memory 
 * for a problem, assuming it has already been allocated in prior 
 * calls to CVodeMalloc and CVodeQuadMalloc. 
 * All problem specification inputs are checked for errors.
 * If any error occurs during initialization, it is reported to the
 * file whose file pointer is errfp.
 * The return value is CV_SUCCESS = 0 if no errors occurred, or
 * a negative value otherwise.
 */

int CVodeQuadReInit(void *cvode_mem, CVQuadRhsFn fQ, N_Vector yQ0)
{
  CVodeMem cv_mem;

  /* Check cvode_mem */
  if (cvode_mem==NULL) {
    CVProcessError(NULL, CV_MEM_NULL, "CVODES", "CVodeQuadReInit", MSGCV_NO_MEM);
    return(CV_MEM_NULL);
  }
  cv_mem = (CVodeMem) cvode_mem;

  /* Ckeck if quadrature was initialized? */
  if (cv_mem->cv_quadMallocDone == FALSE) {
    CVProcessError(cv_mem, CV_NO_QUAD, "CVODES", "CVodeQuadReInit", MSGCV_NO_QUAD);
    return(CV_NO_QUAD);
  }

  /* Initialize znQ[0] in the history array */
  N_VScale(ONE, yQ0, cv_mem->cv_znQ[0]);

  /* Copy the input parameters into CVODES state */
  cv_mem->cv_fQ = fQ;

  /* Initialize counters */
  cv_mem->cv_nfQe  = 0;
  cv_mem->cv_netfQ = 0;

  /* Quadrature integration turned ON */
  cv_mem->cv_quadr = TRUE;

  /* Quadrature re-initialization was successfull */
  return(CV_SUCCESS);
}

/*-----------------------------------------------------------------*/

#define ifS          (cv_mem->cv_ifS)
#define fSDQ         (cv_mem->cv_fSDQ)
#define stgr1alloc   (cv_mem->cv_stgr1alloc)
#define nniS1        (cv_mem->cv_nniS1)
#define ncfnS1       (cv_mem->cv_ncfnS1)
#define ncfS1        (cv_mem->cv_ncfS1)

/*-----------------------------------------------------------------*/

/*
 * CVodeSenMalloc
 *
 * CVodeSensMalloc allocates and initializes sensitivity related 
 * memory for a problem. All problem specification inputs are 
 * checked for errors. If any error occurs during initialization, 
 * it is reported to the file whose file pointer is errfp. 
 * The return value is CV_SUCCESS = 0 if no errors occurred, or
 * a negative value otherwise.
 */

int CVodeSensMalloc(void *cvode_mem, int Ns, int ism, N_Vector *yS0)
{
  CVodeMem    cv_mem;
  booleantype allocOK;
  int is;
  
  /* Check cvode_mem */
  if (cvode_mem==NULL) {
    CVProcessError(NULL, CV_MEM_NULL, "CVODES", "CVodeSensMalloc", MSGCV_NO_MEM);
    return(CV_MEM_NULL);
  }
  cv_mem = (CVodeMem) cvode_mem;

  /* Check if Ns is legal */
  if (Ns<=0) {
    CVProcessError(cv_mem, CV_ILL_INPUT, "CVODES", "CVodeSensMalloc", MSGCV_BAD_NS);
    return(CV_ILL_INPUT);
  }
  cv_mem->cv_Ns = Ns;

  /* Check if ism is legal */
  if ((ism!=CV_SIMULTANEOUS) && (ism!=CV_STAGGERED) && (ism!=CV_STAGGERED1)) {
    CVProcessError(cv_mem, CV_ILL_INPUT, "CVODES", "CVodeSensMalloc", MSGCV_BAD_ISM);
    return(CV_ILL_INPUT);
  }
  cv_mem->cv_ism = ism;
   
  /* Check if yS0 is non-null */
  if (yS0 == NULL) {
    CVProcessError(cv_mem, CV_ILL_INPUT, "CVODES", "CVodeSensMalloc", MSGCV_NULL_YS0);
    return(CV_ILL_INPUT);
  }

  /* Allocate ncfS1, ncfnS1, and nniS1 if needed */
  if (ism == CV_STAGGERED1) {
    stgr1alloc = TRUE;
    ncfS1 = NULL;
    ncfS1 = (int*)malloc(Ns*sizeof(int));
    ncfnS1 = NULL;
    ncfnS1 = (long int*)malloc(Ns*sizeof(long int));
    nniS1 = NULL;
    nniS1 = (long int*)malloc(Ns*sizeof(long int));
    if ( (ncfS1 == NULL) || (ncfnS1 == NULL) || (nniS1 == NULL) ) {
      CVProcessError(cv_mem, CV_MEM_FAIL, "CVODES", "CVodeSensMalloc", MSGCV_MEM_FAIL);
      return(CV_MEM_FAIL);
    }
  } else {
    stgr1alloc = FALSE;
  }

  /* Allocate the vectors (using yS0[0] as a template) */
  allocOK = CVSensAllocVectors(cv_mem, yS0[0]);
  if (!allocOK) {
    if (stgr1alloc) {
      free(ncfS1); ncfS1 = NULL;
      free(ncfnS1); ncfnS1 = NULL;
      free(nniS1); nniS1 = NULL;
    }
    CVProcessError(cv_mem, CV_MEM_FAIL, "CVODES", "CVodeSensMalloc", MSGCV_MEM_FAIL);
    return(CV_MEM_FAIL);
  }
  
  /*---------------------------------------------- 
    All error checking is complete at this point 
    -----------------------------------------------*/

  /* Initialize znS[0] in the history array */
  for (is=0; is<Ns; is++) 
    N_VScale(ONE, yS0[is], cv_mem->cv_znS[0][is]);

  /* Initialize all sensitivity related counters */
  cv_mem->cv_nfSe     = 0;
  cv_mem->cv_nfeS     = 0;
  cv_mem->cv_ncfnS    = 0;
  cv_mem->cv_netfS    = 0;
  cv_mem->cv_nniS     = 0;
  cv_mem->cv_nsetupsS = 0;
  if (ism==CV_STAGGERED1)
    for (is=0; is<Ns; is++) {
      ncfnS1[is] = 0;
      nniS1[is] = 0;
    }

  /* Set default values for plist and pbar */
  for (is=0; is<Ns; is++) {
    cv_mem->cv_plist[is] = is;
    cv_mem->cv_pbar[is] = ONE;
  }

  /* Sensitivities will be computed */
  cv_mem->cv_sensi = TRUE;
  cv_mem->cv_sensMallocDone = TRUE;

  /* Sensitivity initialization was successfull */
  return(CV_SUCCESS);
}

/*-----------------------------------------------------------------*/

#define Ns (cv_mem->cv_Ns)

/*-----------------------------------------------------------------*/

/*
 * CVodeSensReInit
 *
 * CVodeSensReInit re-initializes CVODES's sensitivity related memory 
 * for a problem, assuming it has already been allocated in prior 
 * calls to CVodeMalloc and CVodeSensMalloc. 
 * All problem specification inputs are checked for errors.
 * The number of sensitivities Ns is assumed to be unchanged since
 * the previous call to CVodeSensMalloc.
 * If any error occurs during initialization, it is reported to the
 * file whose file pointer is errfp.
 * The return value is CV_SUCCESS = 0 if no errors occurred, or
 * a negative value otherwise.
 */ 

int CVodeSensReInit(void *cvode_mem, int ism, N_Vector *yS0)
{
  CVodeMem    cv_mem;
  int is;  

  /* Check cvode_mem */
  if (cvode_mem==NULL) {
    CVProcessError(NULL, CV_MEM_NULL, "CVODES", "CVodeSensReInit", MSGCV_NO_MEM);
    return(CV_MEM_NULL);
  }
  cv_mem = (CVodeMem) cvode_mem;

  /* Was sensitivity initialized? */
  if (cv_mem->cv_sensMallocDone == FALSE) {
    CVProcessError(cv_mem, CV_NO_SENS, "CVODES", "CVodeSensReInit", MSGCV_NO_SENSI);
    return(CV_NO_SENS);
  } 

  /* Check if ism is legal */
  if ((ism!=CV_SIMULTANEOUS) && (ism!=CV_STAGGERED) && (ism!=CV_STAGGERED1)) {
    CVProcessError(cv_mem, CV_ILL_INPUT, "CVODES", "CVodeSensReInit", MSGCV_BAD_ISM);
    return(CV_ILL_INPUT);
  }
  cv_mem->cv_ism = ism;
  
  /* Check if yS0 is non-null */
  if (yS0 == NULL) {
    CVProcessError(cv_mem, CV_ILL_INPUT, "CVODES", "CVodeSensReInit", MSGCV_NULL_YS0);
    return(CV_ILL_INPUT);
  }  
  
  /* Allocate ncfS1, ncfnS1, and nniS1 if needed */
  if ( (ism==CV_STAGGERED1) && (stgr1alloc==FALSE) ) {
    stgr1alloc = TRUE;
    ncfS1 = NULL;
    ncfS1 = (int*)malloc(Ns*sizeof(int));
    ncfnS1 = NULL;
    ncfnS1 = (long int*)malloc(Ns*sizeof(long int));
    nniS1 = NULL;
    nniS1 = (long int*)malloc(Ns*sizeof(long int));
    if ( (ncfS1==NULL) || (ncfnS1==NULL) || (nniS1==NULL) ) {
      CVProcessError(cv_mem, CV_MEM_FAIL, "CVODES", "CVodeSensReInit", MSGCV_MEM_FAIL);
      return(CV_MEM_FAIL);
    }
  }

  /*---------------------------------------------- 
    All error checking is complete at this point 
    -----------------------------------------------*/

  /* Initialize znS[0] in the history array */
  for (is=0; is<Ns; is++) 
    N_VScale(ONE, yS0[is], cv_mem->cv_znS[0][is]);

  /* Initialize all sensitivity related counters */
  cv_mem->cv_nfSe     = 0;
  cv_mem->cv_nfeS     = 0;
  cv_mem->cv_ncfnS    = 0;
  cv_mem->cv_netfS    = 0;
  cv_mem->cv_nniS     = 0;
  cv_mem->cv_nsetupsS = 0;
  if (ism==CV_STAGGERED1)
    for (is=0; is<Ns; is++) {
      ncfnS1[is] = 0;
      nniS1[is] = 0;
    }

  /* Problem has been successfully re-initialized */
  cv_mem->cv_sensi = TRUE;
  return(CV_SUCCESS);
}

/*
 * CVodeSensToggleOff
 *
 * CVodeSensToggleOff deactivates sensitivity calculations.
 * It does NOT deallocate sensitivity-related memory.
 */

int CVodeSensToggleOff(void *cvode_mem)
{
  CVodeMem cv_mem;

  /* Check cvode_mem */
  if (cvode_mem==NULL) {
    CVProcessError(NULL, CV_MEM_NULL, "CVODES", "CVodeSensToggleOff", MSGCV_NO_MEM);
    return(CV_MEM_NULL);
  }
  cv_mem = (CVodeMem) cvode_mem;

  /* Disable sensitivities */
  cv_mem->cv_sensi = FALSE;

  return(CV_SUCCESS);
}

/* 
 * -----------------------------------------------------------------
 * Readibility Constants
 * -----------------------------------------------------------------
 */

#define f              (cv_mem->cv_f)      
#define f_data         (cv_mem->cv_f_data) 
#define efun           (cv_mem->cv_efun)
#define e_data         (cv_mem->cv_e_data) 
#define qmax           (cv_mem->cv_qmax)
#define mxstep         (cv_mem->cv_mxstep)
#define mxhnil         (cv_mem->cv_mxhnil)
#define sldeton        (cv_mem->cv_sldeton)
#define hin            (cv_mem->cv_hin)
#define hmin           (cv_mem->cv_hmin)
#define hmax_inv       (cv_mem->cv_hmax_inv)
#define istop          (cv_mem->cv_istop)
#define tstop          (cv_mem->cv_tstop)
#define tstopset       (cv_mem->cv_tstopset)
#define maxnef         (cv_mem->cv_maxnef)
#define maxncf         (cv_mem->cv_maxncf)
#define maxcor         (cv_mem->cv_maxcor)
#define nlscoef        (cv_mem->cv_nlscoef)
#define itol           (cv_mem->cv_itol)         
#define reltol         (cv_mem->cv_reltol)       
#define Sabstol        (cv_mem->cv_Sabstol) 
#define Vabstol        (cv_mem->cv_Vabstol) 

#define fQ             (cv_mem->cv_fQ)
#define fQ_data        (cv_mem->cv_fQ_data)
#define errconQ        (cv_mem->cv_errconQ)
#define itolQ          (cv_mem->cv_itolQ)
#define reltolQ        (cv_mem->cv_reltolQ)
#define SabstolQ       (cv_mem->cv_SabstolQ)
#define VabstolQ       (cv_mem->cv_VabstolQ)

#define fS             (cv_mem->cv_fS)
#define fS1            (cv_mem->cv_fS1)
#define fS_data        (cv_mem->cv_fS_data)
#define DQtype         (cv_mem->cv_DQtype)
#define DQrhomax       (cv_mem->cv_DQrhomax)
#define pbar           (cv_mem->cv_pbar)
#define errconS        (cv_mem->cv_errconS)
#define maxcorS        (cv_mem->cv_maxcorS)
#define itolS          (cv_mem->cv_itolS)
#define reltolS        (cv_mem->cv_reltolS)
#define SabstolS       (cv_mem->cv_SabstolS)
#define VabstolS       (cv_mem->cv_VabstolS)
#define ism            (cv_mem->cv_ism)
#define p              (cv_mem->cv_p)
#define plist          (cv_mem->cv_plist)

#define uround         (cv_mem->cv_uround)  
#define zn             (cv_mem->cv_zn) 
#define ewt            (cv_mem->cv_ewt)  
#define y              (cv_mem->cv_y)
#define acor           (cv_mem->cv_acor)
#define tempv          (cv_mem->cv_tempv)
#define ftemp          (cv_mem->cv_ftemp) 
#define q              (cv_mem->cv_q)
#define qprime         (cv_mem->cv_qprime)
#define next_q         (cv_mem->cv_next_q)
#define qwait          (cv_mem->cv_qwait)
#define L              (cv_mem->cv_L)
#define h              (cv_mem->cv_h)
#define hprime         (cv_mem->cv_hprime)
#define next_h         (cv_mem->cv_next_h)
#define eta            (cv_mem->cv_eta) 
#define etaqm1         (cv_mem->cv_etaqm1) 
#define etaq           (cv_mem->cv_etaq) 
#define etaqp1         (cv_mem->cv_etaqp1) 
#define nscon          (cv_mem->cv_nscon)
#define hscale         (cv_mem->cv_hscale)
#define tn             (cv_mem->cv_tn)
#define tau            (cv_mem->cv_tau)
#define tq             (cv_mem->cv_tq)
#define l              (cv_mem->cv_l)
#define rl1            (cv_mem->cv_rl1)
#define gamma          (cv_mem->cv_gamma) 
#define gammap         (cv_mem->cv_gammap) 
#define gamrat         (cv_mem->cv_gamrat)
#define crate          (cv_mem->cv_crate)
#define acnrm          (cv_mem->cv_acnrm)
#define mnewt          (cv_mem->cv_mnewt)
#define etamax         (cv_mem->cv_etamax)
#define nst            (cv_mem->cv_nst)
#define nfe            (cv_mem->cv_nfe)
#define ncfn           (cv_mem->cv_ncfn)
#define netf           (cv_mem->cv_netf)
#define nni            (cv_mem->cv_nni)
#define nsetups        (cv_mem->cv_nsetups)
#define nhnil          (cv_mem->cv_nhnil)
#define linit          (cv_mem->cv_linit)
#define lsetup         (cv_mem->cv_lsetup)
#define lsolve         (cv_mem->cv_lsolve) 
#define lfree          (cv_mem->cv_lfree) 
#define lmem           (cv_mem->cv_lmem) 
#define qu             (cv_mem->cv_qu)          
#define nstlp          (cv_mem->cv_nstlp)  
#define h0u            (cv_mem->cv_h0u)
#define hu             (cv_mem->cv_hu)         
#define saved_tq5      (cv_mem->cv_saved_tq5)  
#define indx_acor      (cv_mem->cv_indx_acor)
#define jcur           (cv_mem->cv_jcur)         
#define tolsf          (cv_mem->cv_tolsf)      
#define setupNonNull   (cv_mem->cv_setupNonNull) 
#define forceSetup     (cv_mem->cv_forceSetup)
#define nor            (cv_mem->cv_nor)
#define ssdat          (cv_mem->cv_ssdat)

#define nrtfn          (cv_mem->cv_nrtfn)
#define tlo            (cv_mem->cv_tlo)
#define thi            (cv_mem->cv_thi)
#define tretlast       (cv_mem->cv_tretlast)
#define toutc          (cv_mem->cv_toutc)
#define trout          (cv_mem->cv_trout)
#define ttol           (cv_mem->cv_ttol)
#define taskc          (cv_mem->cv_taskc)
#define irfnd          (cv_mem->cv_irfnd)
#define nge            (cv_mem->cv_nge)

#define sensi          (cv_mem->cv_sensi)
#define znS            (cv_mem->cv_znS)
#define ewtS           (cv_mem->cv_ewtS)
#define acorS          (cv_mem->cv_acorS)
#define yS             (cv_mem->cv_yS)
#define tempvS         (cv_mem->cv_tempvS)
#define ftempS         (cv_mem->cv_ftempS)
#define crateS         (cv_mem->cv_crateS)
#define acnrmS         (cv_mem->cv_acnrmS)
#define nfSe           (cv_mem->cv_nfSe)
#define nfeS           (cv_mem->cv_nfeS)
#define nniS           (cv_mem->cv_nniS)
#define ncfnS          (cv_mem->cv_ncfnS)
#define netfS          (cv_mem->cv_netfS)
#define nsetupsS       (cv_mem->cv_nsetupsS)
#define stgr1alloc     (cv_mem->cv_stgr1alloc)
#define sensMallocDone (cv_mem->cv_sensMallocDone)

#define quadr          (cv_mem->cv_quadr)
#define znQ            (cv_mem->cv_znQ)
#define ewtQ           (cv_mem->cv_ewtQ)
#define acorQ          (cv_mem->cv_acorQ)
#define yQ             (cv_mem->cv_yQ)
#define tempvQ         (cv_mem->cv_tempvQ)
#define acnrmQ         (cv_mem->cv_acnrmQ)
#define nfQe           (cv_mem->cv_nfQe)
#define netfQ          (cv_mem->cv_netfQ)
#define quadMallocDone (cv_mem->cv_quadMallocDone)

/* 
 * -----------------------------------------------------------------
 * Main solver function
 * -----------------------------------------------------------------
 */

/*
 * CVode
 *
 * This routine is the main driver of the CVODES package. 
 *
 * It integrates over a time interval defined by the user, by calling
 * CVStep to do internal time steps.
 *
 * The first time that CVode is called for a successfully initialized
 * problem, it computes a tentative initial step size h.
 *
 * CVode supports four modes, specified by itask: CV_NORMAL, CV_ONE_STEP,
 * CV_NORMAL_TSTOP, and CV_ONE_STEP_TSTOP.
 * In the CV_NORMAL mode, the solver steps until it reaches or passes tout
 * and then interpolates to obtain y(tout).
 * In the CV_ONE_STEP mode, it takes one internal step and returns.
 * CV_NORMAL_TSTOP and CV_ONE_STEP_TSTOP are similar to CV_NORMAL and CV_ONE_STEP,
 * respectively, but the integration never proceeds past tstop (which
 * must have been defined through a call to CVodeSetStopTime).
 */

int CVode(void *cvode_mem, realtype tout, N_Vector yout, 
          realtype *tret, int itask)
{
  CVodeMem cv_mem;
  long int nstloc; 
  int retval, hflag, kflag, istate, is, ier, task, irfndp;
  realtype troundoff, rh, nrm;

  /*
   * -------------------------------------
   * 1. Check and process inputs
   * -------------------------------------
   */

  /* Check if cvode_mem exists */
  if (cvode_mem == NULL) {
    CVProcessError(NULL, CV_MEM_NULL, "CVODES", "CVode", MSGCV_NO_MEM);
    return(CV_MEM_NULL);
  }
  cv_mem = (CVodeMem) cvode_mem;

  /* Check if cvode_mem was allocated */
  if (cv_mem->cv_MallocDone == FALSE) {
    CVProcessError(cv_mem, CV_NO_MALLOC, "CVODES", "CVode", MSGCV_NO_MALLOC);
    return(CV_NO_MALLOC);
  }
  
  /* Check for yout != NULL */
  if ((y = yout) == NULL) {
    CVProcessError(cv_mem, CV_ILL_INPUT, "CVODES", "CVode", MSGCV_YOUT_NULL);
    return(CV_ILL_INPUT);
  }
  
  /* Check for tret != NULL */
  if (tret == NULL) {
    CVProcessError(cv_mem, CV_ILL_INPUT, "CVODES", "CVode", MSGCV_TRET_NULL);
    return(CV_ILL_INPUT);
  }

  /* Check for valid itask */
  if ((itask != CV_NORMAL)       && 
      (itask != CV_ONE_STEP)     &&
      (itask != CV_NORMAL_TSTOP) &&
      (itask != CV_ONE_STEP_TSTOP) ) {
    CVProcessError(cv_mem, CV_ILL_INPUT, "CVODES", "CVode", MSGCV_BAD_ITASK);
    return(CV_ILL_INPUT);
  }

  /* Split itask into task and istop */
  if ((itask == CV_NORMAL_TSTOP) || (itask == CV_ONE_STEP_TSTOP)) {
    if ( tstopset == FALSE ) {
      CVProcessError(cv_mem, CV_ILL_INPUT, "CVODES", "CVode", MSGCV_NO_TSTOP);
      return(CV_ILL_INPUT);
    }
    istop = TRUE;
  } else {
    istop = FALSE;
  }
  if ((itask == CV_NORMAL) || (itask == CV_NORMAL_TSTOP)) {
    task = CV_NORMAL; toutc = tout;
  } else {
    task = CV_ONE_STEP;
  }
  taskc = task;


  /* Sensitivity-specific tests */

  if (sensi && fSDQ) {

    /* If using internal DQ function for sensitivity RHS */

    /* Set internal DQ functions */
    fS  = CVSensRhsDQ;
    fS1 = CVSensRhs1DQ;

    /* Make sure we have the right 'user data' */
    fS_data = cvode_mem;
    
    /* Test if we have the problem parameters */
    if(p == NULL) {
      CVProcessError(cv_mem, CV_ILL_INPUT, "CVODES", "CVode", MSGCV_NULL_P);
      return(CV_ILL_INPUT);
    }

    /* Set ifS to CV_ONESENS so that any method can be used */
    ifS = CV_ONESENS;
    
  }

  /*
   * ----------------------------------------
   * 2. Initializations performed only at
   *    the first step (nst=0):
   *    - initial setup
   *    - initialize Nordsieck history array
   *    - compute initial step size
   *    - check for approach to tstop
   *    - check for approach to a root
   * ----------------------------------------
   */

  if (nst == 0) {

    /* Check inputs for corectness */

    ier = CVInitialSetup(cv_mem);
    if (ier!= CV_SUCCESS) return(ier);

    /* 
       Call f at (t0,y0), set zn[1] = y'(t0). 
       If computing any quadratures, call fQ at (t0,y0), set znQ[1] = yQ'(t0)
       If computing sensitivities, call fS at (t0,y0,yS0), set znS[1][is] = yS'(t0), is=1,...,Ns.
    */

    retval = f(tn, zn[0], zn[1], f_data); 
    nfe++;
    if (retval < 0) {
      CVProcessError(cv_mem, CV_RHSFUNC_FAIL, "CVODES", "CVode", MSGCV_RHSFUNC_FAILED, tn);
      return(CV_RHSFUNC_FAIL);
    }
    if (retval > 0) {
      CVProcessError(cv_mem, CV_FIRST_RHSFUNC_ERR, "CVODES", "CVode", MSGCV_RHSFUNC_FIRST);
      return(CV_FIRST_RHSFUNC_ERR);
    }

    if (quadr) {
      retval = fQ(tn, zn[0], znQ[1], fQ_data);
      nfQe++;
      if (retval < 0) {
        CVProcessError(cv_mem, CV_QRHSFUNC_FAIL, "CVODES", "CVode", MSGCV_QRHSFUNC_FAILED, tn);
        return(CV_QRHSFUNC_FAIL);
      }
      if (retval > 0) {
        CVProcessError(cv_mem, CV_FIRST_QRHSFUNC_ERR, "CVODES", "CVode", MSGCV_QRHSFUNC_FIRST);
        return(CV_FIRST_QRHSFUNC_ERR);
      }
    }

    if (sensi) {
      retval = CVSensRhs(cv_mem, tn, zn[0], zn[1], znS[0], znS[1], tempv, ftemp);
      if (retval < 0) {
        CVProcessError(cv_mem, CV_SRHSFUNC_FAIL, "CVODES", "CVSensRhs", MSGCV_SRHSFUNC_FAILED, tn);
        return(CV_SRHSFUNC_FAIL);
      } 
      if (retval > 0) {
        CVProcessError(cv_mem, CV_FIRST_SRHSFUNC_ERR, "CVODES", "CVSensRhs", MSGCV_SRHSFUNC_FIRST);
        return(CV_FIRST_SRHSFUNC_ERR);
      }
    }

    /* Set initial h (from H0 or CVHin). */
    
    h = hin;
    if ( (h != ZERO) && ((tout-tn)*h < ZERO) ) {
      CVProcessError(cv_mem, CV_ILL_INPUT, "CVODES", "CVode", MSGCV_BAD_H0);
      return(CV_ILL_INPUT);
    }
    if (h == ZERO) {
      hflag = CVHin(cv_mem, tout);
      if (hflag != CV_SUCCESS) {
        istate = CVHandleFailure(cv_mem, hflag);
        return(istate);
      }
    }
    rh = ABS(h)*hmax_inv;
    if (rh > ONE) h /= rh;
    if (ABS(h) < hmin) h *= hmin/ABS(h);

    /* Check for approach to tstop */

    if (istop) {
      if ( (tstop - tn)*h < ZERO ) {
        CVProcessError(cv_mem, CV_ILL_INPUT, "CVODES", "CVode", MSGCV_BAD_TSTOP, tn);
        return(CV_ILL_INPUT);
      }
      if ( (tn + h - tstop)*h > ZERO ) 
        h = (tstop - tn)*(ONE-FOUR*uround);
    }

    /* 
       Scale zn[1] by h.
       If computing any quadratures, scale znQ[1] by h.
       If computing sensitivities,  scale znS[1][is] by h. 
    */

    hscale = h;
    h0u    = h;
    hprime = h;

    N_VScale(h, zn[1], zn[1]);
    
    if (sensi)
      for (is=0; is<Ns; is++) 
        N_VScale(h, znS[1][is], znS[1][is]);
    
    if (quadr)
      N_VScale(h, znQ[1], znQ[1]);

    /* Check for zeros of root function g at and near t0. */

    if (nrtfn > 0) {

      retval = CVRcheck1(cv_mem);

      if (retval == INITROOT) {
        CVProcessError(cv_mem, CV_ILL_INPUT, "CVODES", "CVRcheck1", MSGCV_BAD_INIT_ROOT);
        return(CV_ILL_INPUT);
      } else if (retval == CV_RTFUNC_FAIL) {
        CVProcessError(cv_mem, CV_RTFUNC_FAIL, "CVODES", "CVRcheck1", MSGCV_RTFUNC_FAILED, tn);
        return(CV_RTFUNC_FAIL);
      }

    }

  } /* end first call block */

  /*
   * ------------------------------------------------------
   * 3. At following steps, perform stop tests:
   *    - check for root in last step
   *    - check if we passed tstop
   *    - check if we passed tout (NORMAL mode)
   *    - check if current tn was returned (ONE_STEP mode)
   *    - check if we are close to tstop
   *      (adjust step size if needed)
   * -------------------------------------------------------
   */

  if (nst > 0) {

    /* Estimate an infinitesimal time interval to be used as
       a roundoff for time quantities (based on current time 
       and step size) */
    troundoff = FUZZ_FACTOR*uround*(ABS(tn) + ABS(h));

    /* First check for a root in the last step taken, other than the
       last root found, if any.  If task = CV_ONE_STEP and y(tn) was not
       returned because of an intervening root, return y(tn) now.     */
    if (nrtfn > 0) {
      
      irfndp = irfnd;
      
      retval = CVRcheck2(cv_mem);

      if (retval == CLOSERT) {
        CVProcessError(cv_mem, CV_ILL_INPUT, "CVODES", "CVRcheck2", MSGCV_CLOSE_ROOTS, tlo);
        return(CV_ILL_INPUT);
      } else if (retval == CV_RTFUNC_FAIL) {
        CVProcessError(cv_mem, CV_RTFUNC_FAIL, "CVODES", "CVRcheck2", MSGCV_RTFUNC_FAILED, tlo);
        return(CV_RTFUNC_FAIL);
      } else if (retval == RTFOUND) {
        tretlast = *tret = tlo;
        return(CV_ROOT_RETURN);
      }
      
      /* If tn is distinct from tretlast (within roundoff),
         check remaining interval for roots */
      if ( ABS(tn - tretlast) > troundoff ) {

        retval = CVRcheck3(cv_mem);

        if (retval == CV_SUCCESS) {     /* no root found */
          irfnd = 0;
          if ((irfndp == 1) && (task == CV_ONE_STEP)) {
            tretlast = *tret = tn;
            N_VScale(ONE, zn[0], yout);
            return(CV_SUCCESS);
          }
        } else if (retval == RTFOUND) {  /* a new root was found */
          irfnd = 1;
          tretlast = *tret = tlo;
          return(CV_ROOT_RETURN);
        } else if (retval == CV_RTFUNC_FAIL) {  /* g failed */
          CVProcessError(cv_mem, CV_RTFUNC_FAIL, "CVODES", "CVRcheck3", MSGCV_RTFUNC_FAILED, tlo);
          return(CV_RTFUNC_FAIL);
        }

      }
      
    } /* end of root stop check */
    
    /* Test for tn past tstop */
    if ( istop && ((tstop - tn)*h < ZERO) ) {
      CVProcessError(cv_mem, CV_ILL_INPUT, "CVODES", "CVode", MSGCV_BAD_TSTOP, tn);
      return(CV_ILL_INPUT);
    }
    
    /* In CV_NORMAL mode, test if tout was reached */
    if ( (task == CV_NORMAL) && ((tn-tout)*h >= ZERO) ) {
      tretlast = *tret = tout;
      ier =  CVodeGetDky(cv_mem, tout, 0, yout);
      if (ier != CV_SUCCESS) {
        CVProcessError(cv_mem, CV_ILL_INPUT, "CVODES", "CVode", MSGCV_BAD_TOUT, tout);
        return(CV_ILL_INPUT);
      }
      return(CV_SUCCESS);
    }
    
    /* In CV_ONE_STEP mode, test if tn was returned */
    if ( task == CV_ONE_STEP && ABS(tn - tretlast) > troundoff ) {
      tretlast = *tret = tn;
      N_VScale(ONE, zn[0], yout);
      return(CV_SUCCESS);
    }
    
    /* Test for tn at tstop or near tstop */
    if ( istop ) {
      
      if ( ABS(tn - tstop) <= troundoff ) {
        ier =  CVodeGetDky(cv_mem, tstop, 0, yout);
        if (ier != CV_SUCCESS) {
          CVProcessError(cv_mem, CV_ILL_INPUT, "CVODES", "CVode", MSGCV_BAD_TSTOP, tn);
          return(CV_ILL_INPUT);
        }
        tretlast = *tret = tstop;
        return(CV_TSTOP_RETURN);
      }
      
      /* If next step would overtake tstop, adjust stepsize */
      if ( (tn + hprime - tstop)*h > ZERO ) {
        hprime = (tstop - tn)*(ONE-FOUR*uround);
        eta = hprime/h;
      }
      
    } /* end of istop tests block */
    
  } /* end stopping tests block at nst>0 */  
  
  /*
   * --------------------------------------------------
   * 4. Looping point for internal steps
   *
   *    4.1. check for errors (too many steps, too much
   *         accuracy requested, step size too small)
   *    4.2. take a new step (call CVStep)
   *    4.3. stop on error 
   *    4.4. perform stop tests:
   *         - check for root in last step
   *         - check if tout was passed
   *         - check if close to tstop
   *         - check if in ONE_STEP mode (must return)
   * --------------------------------------------------
   */  
  
  nstloc = 0;
  loop {
   
    next_h = h;
    next_q = q;
    
    /* Reset and check ewt, ewtQ, ewtS */   
    if (nst > 0) {

      ier = efun(zn[0], ewt, e_data);
      if(ier != 0) {
        if (itol == CV_WF) CVProcessError(cv_mem, CV_ILL_INPUT, "CVODES", "CVode", MSGCV_EWT_NOW_FAIL, tn);
        else               CVProcessError(cv_mem, CV_ILL_INPUT, "CVODES", "CVode", MSGCV_EWT_NOW_BAD, tn);
        istate = CV_ILL_INPUT;
        tretlast = *tret = tn;
        N_VScale(ONE, zn[0], yout);
        break;
      }

      if (quadr && errconQ) {
        ier = CVQuadEwtSet(cv_mem, znQ[0], ewtQ);
        if(ier != 0) {
          CVProcessError(cv_mem, CV_ILL_INPUT, "CVODES", "CVode", MSGCV_EWTQ_NOW_BAD, tn);
          istate = CV_ILL_INPUT;
          tretlast = *tret = tn;
          N_VScale(ONE, zn[0], yout);
          break;
        }
      }

      if (sensi) {
        ier = CVSensEwtSet(cv_mem, znS[0], ewtS);
        if (ier != 0) {
          CVProcessError(cv_mem, CV_ILL_INPUT, "CVODES", "CVode", MSGCV_EWTS_NOW_BAD, tn);
          istate = CV_ILL_INPUT;
          tretlast = *tret = tn;
          N_VScale(ONE, zn[0], yout);
          break;
        }
      }

    }

    /* Check for too many steps */
    if (nstloc >= mxstep) {
      CVProcessError(cv_mem, CV_TOO_MUCH_WORK, "CVODES", "CVode", MSGCV_MAX_STEPS, tn);
      istate = CV_TOO_MUCH_WORK;
      tretlast = *tret = tn;
      N_VScale(ONE, zn[0], yout);
      break;
    }

    /* Check for too much accuracy requested */
    nrm = N_VWrmsNorm(zn[0], ewt);
    if (quadr && errconQ) {
      nrm = CVQuadUpdateNorm(cv_mem, nrm, znQ[0], ewtQ); 
    }
    if (sensi && errconS) {
      nrm = CVSensUpdateNorm(cv_mem, nrm, znS[0], ewtS);
    }
    tolsf = uround * nrm;
    if (tolsf > ONE) {
      CVProcessError(cv_mem, CV_TOO_MUCH_ACC, "CVODES", "CVode", MSGCV_TOO_MUCH_ACC, tn);
      istate = CV_TOO_MUCH_ACC;
      tretlast = *tret = tn;
      N_VScale(ONE, zn[0], yout);
      tolsf *= TWO;
      break;
    } else {
      tolsf = ONE;
    }
    
    /* Check for h below roundoff level in tn */
    if (tn + h == tn) {
      nhnil++;
      if (nhnil <= mxhnil) 
        CVProcessError(cv_mem, CV_WARNING, "CVODES", "CVode", MSGCV_HNIL, tn, h);
      if (nhnil == mxhnil) 
        CVProcessError(cv_mem, CV_WARNING, "CVODES", "CVode", MSGCV_HNIL_DONE);
    }

    /* Call CVStep to take a step */
    kflag = CVStep(cv_mem);

    /* Process failed step cases, and exit loop */
    if (kflag != CV_SUCCESS) {
      istate = CVHandleFailure(cv_mem, kflag);
      tretlast = *tret = tn;
      N_VScale(ONE, zn[0], yout);
      break;
    }
    
    nstloc++;

    /* Check for root in last step taken. */    
    if (nrtfn > 0) {
      
      retval = CVRcheck3(cv_mem);
      
      if (retval == RTFOUND) {  /* A new root was found */
        irfnd = 1;
        istate = CV_ROOT_RETURN;
        tretlast = *tret = tlo;
        break;
      } else if (retval == CV_RTFUNC_FAIL) { /* g failed */
        CVProcessError(cv_mem, CV_RTFUNC_FAIL, "CVODES", "CVRcheck3", MSGCV_RTFUNC_FAILED, tlo);
        istate = CV_RTFUNC_FAIL;
        break;
      }

    }

    /* In NORMAL mode, check if tout reached */
    if ( (task == CV_NORMAL) &&  (tn-tout)*h >= ZERO ) {
      istate = CV_SUCCESS;
      tretlast = *tret = tout;
      (void) CVodeGetDky(cv_mem, tout, 0, yout);
      next_q = qprime;
      next_h = hprime;
      break;
    }

    /* Check if tn is at tstop or near tstop */
    if ( istop ) {

      troundoff = FUZZ_FACTOR*uround*(ABS(tn) + ABS(h));
      if ( ABS(tn - tstop) <= troundoff) {
        (void) CVodeGetDky(cv_mem, tstop, 0, yout);
        tretlast = *tret = tstop;
        istate = CV_TSTOP_RETURN;
        break;
      }

      if ( (tn + hprime - tstop)*h > ZERO ) {
        hprime = (tstop - tn)*(ONE-FOUR*uround);
        eta = hprime/h;
      }

    }

    /* In ONE_STEP mode, copy y and exit loop */
    if (task == CV_ONE_STEP) {
      istate = CV_SUCCESS;
      tretlast = *tret = tn;
      N_VScale(ONE, zn[0], yout);
      next_q = qprime;
      next_h = hprime;
      break;
    }

  } /* end looping for internal steps */
  
  /* Load optional output */
  if (sensi && (ism==CV_STAGGERED1)) { 
    nniS  = 0;
    ncfnS = 0;
    for (is=0; is<Ns; is++) {
      nniS  += nniS1[is];
      ncfnS += ncfnS1[is];
    }
  }
  
  return(istate);

}

/* 
 * -----------------------------------------------------------------
 * Interpolated output and extraction functions
 * -----------------------------------------------------------------
 */

/*
 * CVodeGetDky
 *
 * This routine computes the k-th derivative of the interpolating
 * polynomial at the time t and stores the result in the vector dky.
 * The formula is:
 *         q 
 *  dky = SUM c(j,k) * (t - tn)^(j-k) * h^(-j) * zn[j] , 
 *        j=k 
 * where c(j,k) = j*(j-1)*...*(j-k+1), q is the current order, and
 * zn[j] is the j-th column of the Nordsieck history array.
 *
 * This function is called by CVode with k = 0 and t = tout, but
 * may also be called directly by the user.
 */

int CVodeGetDky(void *cvode_mem, realtype t, int k, N_Vector dky)
{
  realtype s, c, r;
  realtype tfuzz, tp, tn1;
  int i, j;
  CVodeMem cv_mem;
  
  /* Check all inputs for legality */
 
  if (cvode_mem == NULL) {
    CVProcessError(NULL, CV_MEM_NULL, "CVODES", "CVodeGetDky", MSGCV_NO_MEM);
    return(CV_MEM_NULL);
  }
  cv_mem = (CVodeMem) cvode_mem;

  if (dky == NULL) {
    CVProcessError(cv_mem, CV_BAD_DKY, "CVODES", "CVodeGetDky", MSGCV_NULL_DKY);
    return(CV_BAD_DKY);
  }

  if ((k < 0) || (k > q)) {
    CVProcessError(cv_mem, CV_BAD_K, "CVODES", "CVodeGetDky", MSGCV_BAD_K);
    return(CV_BAD_K);
  }
  
  /* Allow for some slack */
  tfuzz = FUZZ_FACTOR * uround * (ABS(tn) + ABS(hu));
  if (hu < ZERO) tfuzz = -tfuzz;
  tp = tn - hu - tfuzz;
  tn1 = tn + tfuzz;
  if ((t-tp)*(t-tn1) > ZERO) {
    CVProcessError(cv_mem, CV_BAD_T, "CVODES", "CVodeGetDky", MSGCV_BAD_T, t, tn-hu, tn);
    return(CV_BAD_T);
  }

  /* Sum the differentiated interpolating polynomial */

  s = (t - tn) / h;
  for (j=q; j >= k; j--) {
    c = ONE;
    for (i=j; i >= j-k+1; i--) c *= i;
    if (j == q) {
      N_VScale(c, zn[q], dky);
    } else {
      N_VLinearSum(c, zn[j], s, dky, dky);
    }
  }
  if (k == 0) return(CV_SUCCESS);
  r = RPowerI(h,-k);
  N_VScale(r, dky, dky);
  return(CV_SUCCESS);
}

/* 
 * CVodeGetQuad
 *
 * This routine extracts quadrature solution into yQout.
 * This is just a wrapper that calls CVodeGetQuadDky with k=0                    
 */
 
int CVodeGetQuad(void *cvode_mem, realtype t, N_Vector yQout)
{
  return(CVodeGetQuadDky(cvode_mem,t,0,yQout));
}

/*
 * CVodeGetQuadDky
 *
 * CVodeQuadDky computes the kth derivative of the yQ function at
 * time t, where tn-hu <= t <= tn, tn denotes the current         
 * internal time reached, and hu is the last internal step size   
 * successfully used by the solver. The user may request 
 * k=0, 1, ..., qu, where qu is the current order. 
 * The derivative vector is returned in dky. This vector 
 * must be allocated by the caller. It is only legal to call this         
 * function after a successful return from CVode with quadrature
 * computation enabled.
 */

int CVodeGetQuadDky(void *cvode_mem, realtype t, int k, N_Vector dkyQ)
{ 
  realtype s, c, r;
  realtype tfuzz, tp, tn1;
  int i, j;
  CVodeMem cv_mem;
  
  /* Check all inputs for legality */
  
  if (cvode_mem == NULL) {
    CVProcessError(NULL, CV_MEM_NULL, "CVODES", "CVodeGetQuadDky", MSGCV_NO_MEM);
    return(CV_MEM_NULL);
  }
  cv_mem = (CVodeMem) cvode_mem;  

  if(quadr != TRUE) {
    CVProcessError(cv_mem, CV_NO_QUAD, "CVODES", "CVodeGetQuadDky", MSGCV_NO_QUAD);
    return(CV_NO_QUAD);
  }

  if (dkyQ == NULL) {
    CVProcessError(cv_mem, CV_BAD_DKY, "CVODES", "CVodeGetQuadDky", MSGCV_NULL_DKY);
    return(CV_BAD_DKY);
  }
  
  if ((k < 0) || (k > q)) {
    CVProcessError(cv_mem, CV_BAD_K, "CVODES", "CVodeGetQuadDky", MSGCV_BAD_K);
    return(CV_BAD_K);
  }
  
  /* Allow for some slack */
  tfuzz = FUZZ_FACTOR * uround * (ABS(tn) + ABS(hu));
  if (hu < ZERO) tfuzz = -tfuzz;
  tp = tn - hu - tfuzz;
  tn1 = tn + tfuzz;
  if ((t-tp)*(t-tn1) > ZERO) {
    CVProcessError(cv_mem, CV_BAD_T, "CVODES", "CVodeGetQuadDky", MSGCV_BAD_T);
    return(CV_BAD_T);
  }
  
  /* Sum the differentiated interpolating polynomial */
  
  s = (t - tn) / h;
  for (j=q; j >= k; j--) {
    c = ONE;
    for (i=j; i >= j-k+1; i--) c *= i;
    if (j == q) {
      N_VScale(c, znQ[q], dkyQ);
    } else {
      N_VLinearSum(c, znQ[j], s, dkyQ, dkyQ);
    }
  }
  if (k == 0) return(CV_SUCCESS);
  r = RPowerI(h,-k);
  N_VScale(r, dkyQ, dkyQ);
  return(CV_SUCCESS);
  
}

/* 
 * CVodeGetSens
 *
 * This routine extracts sensitivity solution into ySout.
 * This is just a wrapper that calls CVodeSensDky with k=0                    
 */
 
int CVodeGetSens(void *cvode_mem, realtype t, N_Vector *ySout)
{
  return(CVodeGetSensDky(cvode_mem,t,0,ySout));
}
    
/* 
 * CVodeGetSens1
 *
 * This routine extracts the is-th sensitivity solution into ySout.
 * This is just a wrapper that calls CVodeSensDky1 with k=0                    
 */
 
int CVodeGetSens1(void *cvode_mem, realtype t, int is, N_Vector ySout)
{
  return(CVodeGetSensDky1(cvode_mem,t,0,is,ySout));
}
    
/*
 * CVodeGetSensDky
 *
 * If the user calls directly CVodeSensDky then s must be allocated
 * prior to this call. When CVodeSensDky is called by 
 * CVodeGetSens, only ier=CV_SUCCESS, ier=CV_NO_SENS, or 
 * ier=CV_BAD_T are possible.
 */

int CVodeGetSensDky(void *cvode_mem, realtype t, int k, N_Vector *dkyS)
{
  int ier=CV_SUCCESS;
  int is;
  CVodeMem cv_mem;
  
  if (cvode_mem == NULL) {
    CVProcessError(NULL, CV_MEM_NULL, "CVODES", "CVodeGetSensDky", MSGCV_NO_MEM);
    return(CV_MEM_NULL);
  }
  cv_mem = (CVodeMem) cvode_mem;  
  
  if (dkyS == NULL) {
    CVProcessError(cv_mem, CV_BAD_DKY, "CVODES", "CVodeGetSensDky", MSGCV_NULL_DKYA);
    return(CV_BAD_DKY);
  }
  
  for (is=0; is<Ns; is++) {
    ier = CVodeGetSensDky1(cvode_mem,t,k,is,dkyS[is]);
    if (ier!=CV_SUCCESS) break;
  }
  
  return(ier);
}
    
/*
 * CVodeGetSensDky1
 *
 * CVodeSensDky1 computes the kth derivative of the yS[is] function at
 * time t, where tn-hu <= t <= tn, tn denotes the current         
 * internal time reached, and hu is the last internal step size   
 * successfully used by the solver. The user may request 
 * is=0, 1, ..., Ns-1 and k=0, 1, ..., qu, where qu is the current
 * order. The derivative vector is returned in dky. This vector 
 * must be allocated by the caller. It is only legal to call this         
 * function after a successful return from CVode with sensitivity 
 * computation enabled.
 */

int CVodeGetSensDky1(void *cvode_mem, realtype t, int k, int is, N_Vector dkyS)
{ 
  realtype s, c, r;
  realtype tfuzz, tp, tn1;
  int i, j;
  CVodeMem cv_mem;
  
  /* Check all inputs for legality */
  
  if (cvode_mem == NULL) {
    CVProcessError(NULL, CV_MEM_NULL, "CVODES", "CVodeGetSensDky1", MSGCV_NO_MEM);
    return(CV_MEM_NULL);
  }
  cv_mem = (CVodeMem) cvode_mem;  
  
  if(sensi != TRUE) {
    CVProcessError(cv_mem, CV_NO_SENS, "CVODES", "CVodeGetSensDky1", MSGCV_NO_SENSI);
    return(CV_NO_SENS);
  }

  if (dkyS == NULL) {
    CVProcessError(cv_mem, CV_BAD_DKY, "CVODES", "CVodeGetSensDky1", MSGCV_NULL_DKY);
    return(CV_BAD_DKY);
  }
  
  if ((k < 0) || (k > q)) {
    CVProcessError(cv_mem, CV_BAD_K, "CVODES", "CVodeGetSensDky1", MSGCV_BAD_K);
    return(CV_BAD_K);
  }
  
  if ((is < 0) || (is > Ns-1)) {
    CVProcessError(cv_mem, CV_BAD_IS, "CVODES", "CVodeGetSensDky1", MSGCV_BAD_IS);
    return(CV_BAD_IS);
  }
  
  /* Allow for some slack */
  tfuzz = FUZZ_FACTOR * uround * (ABS(tn) + ABS(hu));
  if (hu < ZERO) tfuzz = -tfuzz;
  tp = tn - hu - tfuzz;
  tn1 = tn + tfuzz;
  if ((t-tp)*(t-tn1) > ZERO) {
    CVProcessError(cv_mem, CV_BAD_T, "CVODES", "CVodeGetSensDky1", MSGCV_BAD_T);
    return(CV_BAD_T);
  }
  
  /* Sum the differentiated interpolating polynomial */
  
  s = (t - tn) / h;
  for (j=q; j >= k; j--) {
    c = ONE;
    for (i=j; i >= j-k+1; i--) c *= i;
    if (j == q) {
      N_VScale(c, znS[q][is], dkyS);
    } else {
      N_VLinearSum(c, znS[j][is], s, dkyS, dkyS);
    }
  }
  if (k == 0) return(CV_SUCCESS);
  r = RPowerI(h,-k);
  N_VScale(r, dkyS, dkyS);
  return(CV_SUCCESS);
  
}

/* 
 * -----------------------------------------------------------------
 * Deallocation functions
 * -----------------------------------------------------------------
 */

/*
 * CVodeFree
 *
 * This routine frees the problem memory allocated by CVodeMalloc.
 * Such memory includes all the vectors allocated by CVAllocVectors,
 * and the memory lmem for the linear solver (deallocated by a call
 * to lfree), as well as (if Ns!=0) all memory allocated for 
 * sensitivity computations by CVodeSensMalloc.
 */

void CVodeFree(void **cvode_mem)
{
  CVodeMem cv_mem;

  if (*cvode_mem == NULL) return;

  cv_mem = (CVodeMem) (*cvode_mem);
  
  CVFreeVectors(cv_mem);

  CVodeQuadFree(cv_mem);

  CVodeSensFree(cv_mem);

  if (iter == CV_NEWTON && lfree != NULL) lfree(cv_mem);

  if (nrtfn > 0) {
    free(glo); glo = NULL; 
    free(ghi);  ghi = NULL;
    free(grout);  grout = NULL;
    free(iroots); iroots = NULL;
  }

  free(*cvode_mem);
  *cvode_mem = NULL;
}

/*
 * CVodeQuadFree
 *
 * CVodeQuadFree frees the problem memory in cvode_mem allocated
 * for quadrature integration. Its only argument is the pointer
 * cvode_mem returned by CVodeCreate. 
 */

void CVodeQuadFree(void *cvode_mem)
{
  CVodeMem cv_mem;
  
  if (cvode_mem == NULL) return;
  cv_mem = (CVodeMem) cvode_mem;

  if(quadMallocDone) {
    CVQuadFreeVectors(cv_mem);
    quadMallocDone = FALSE;
    quadr = FALSE;
  }
}

/*
 * CVodeSensFree
 *
 * CVodeSensFree frees the problem memory in cvode_mem allocated
 * for sensitivity analysis. Its only argument is the pointer
 * cvode_mem returned by CVodeCreate. 
 */

void CVodeSensFree(void *cvode_mem)
{
  CVodeMem cv_mem;
  
  if (cvode_mem == NULL) return;
  cv_mem = (CVodeMem) cvode_mem;

  if(sensMallocDone) {
    if (stgr1alloc) {
      free(ncfS1); ncfS1 = NULL;
      free(ncfnS1); ncfnS1 = NULL;
      free(nniS1); nniS1 = NULL;
      stgr1alloc = FALSE;
    }
    CVSensFreeVectors(cv_mem);
    sensMallocDone = FALSE;
    sensi = FALSE;
  }
}

/* 
 * =================================================================
 * PRIVATE FUNCTIONS
 * =================================================================
 */

/*
 * CVCheckNvector
 * This routine checks if all required vector operations are present.
 * If any of them is missing it returns FALSE.
 */

static booleantype CVCheckNvector(N_Vector tmpl)
{
  if((tmpl->ops->nvclone     == NULL) ||
     (tmpl->ops->nvdestroy   == NULL) ||
     (tmpl->ops->nvlinearsum == NULL) ||
     (tmpl->ops->nvconst     == NULL) ||
     (tmpl->ops->nvprod      == NULL) ||
     (tmpl->ops->nvdiv       == NULL) ||
     (tmpl->ops->nvscale     == NULL) ||
     (tmpl->ops->nvabs       == NULL) ||
     (tmpl->ops->nvinv       == NULL) ||
     (tmpl->ops->nvaddconst  == NULL) ||
     (tmpl->ops->nvmaxnorm   == NULL) ||
     (tmpl->ops->nvwrmsnorm  == NULL) ||
     (tmpl->ops->nvmin       == NULL))
    return(FALSE);
  else
    return(TRUE);
}

/* 
 * -----------------------------------------------------------------
 * Memory allocation/deallocation
 * -----------------------------------------------------------------
 */

/*
 * CVAllocVectors
 *
 * This routine allocates the CVODES vectors ewt, acor, tempv, ftemp, and
 * zn[0], ..., zn[maxord]. If itol=CV_SV, it also allocates space for Vabstol.
 * If all memory allocations are successful, CVAllocVectors returns TRUE. 
 * Otherwise all allocated memory is freed and CVAllocVectors returns FALSE.
 * This routine also sets the optional outputs lrw and liw, which are
 * (respectively) the lengths of the real and integer work spaces
 * allocated here.
 */

static booleantype CVAllocVectors(CVodeMem cv_mem, N_Vector tmpl, int tol)
{
  int i, j;

  /* Allocate ewt, acor, tempv, ftemp */
  
  ewt = NULL;
  ewt = N_VClone(tmpl);
  if (ewt == NULL) return(FALSE);

  acor = NULL;
  acor = N_VClone(tmpl);
  if (acor == NULL) {
    N_VDestroy(ewt);
    return(FALSE);
  }

  tempv = NULL;
  tempv = N_VClone(tmpl);
  if (tempv == NULL) {
    N_VDestroy(ewt);
    N_VDestroy(acor);
    return(FALSE);
  }

  ftemp = NULL;
  ftemp = N_VClone(tmpl);
  if (ftemp == NULL) {
    N_VDestroy(tempv);
    N_VDestroy(ewt);
    N_VDestroy(acor);
    return(FALSE);
  }

  /* Allocate zn[0] ... zn[qmax] */

  for (j=0; j <= qmax; j++) {
    zn[j] = NULL;
    zn[j] = N_VClone(tmpl);
    if (zn[j] == NULL) {
      N_VDestroy(ewt);
      N_VDestroy(acor);
      N_VDestroy(tempv);
      N_VDestroy(ftemp);
      for (i=0; i < j; i++) N_VDestroy(zn[i]);
      return(FALSE);
    }
  }

  /* Update solver workspace lengths  */
  lrw += (qmax + 5)*lrw1;
  liw += (qmax + 5)*liw1;

  if (tol == CV_SV) {
    Vabstol = NULL;
    Vabstol = N_VClone(tmpl);
    if (Vabstol == NULL) {
      N_VDestroy(ewt);
      N_VDestroy(acor);
      N_VDestroy(tempv);
      N_VDestroy(ftemp);
      for (i=0; i <= qmax; i++) N_VDestroy(zn[i]);
      return(FALSE);
    }
    lrw += lrw1;
    liw += liw1;
    cv_mem->cv_VabstolMallocDone = TRUE;
  }  

  /* Store the value of qmax used here */
  cv_mem->cv_qmax_alloc = qmax;

  return(TRUE);
}

/*  
 * CVFreeVectors
 *
 * This routine frees the CVODES vectors allocated in CVAllocVectors.
 */

static void CVFreeVectors(CVodeMem cv_mem)
{
  int j, maxord;
  
  maxord = cv_mem->cv_qmax_alloc;

  N_VDestroy(ewt);
  N_VDestroy(acor);
  N_VDestroy(tempv);
  N_VDestroy(ftemp);
  for (j=0; j <= maxord; j++) 
    N_VDestroy(zn[j]);

  lrw -= (maxord + 5)*lrw1;
  liw -= (maxord + 5)*liw1;

  if (cv_mem->cv_VabstolMallocDone) {
    N_VDestroy(Vabstol);
    lrw -= lrw1;
    liw -= liw1;
  }
}

/*
 * CVodeQuadAllocVectors
 *
 * NOTE: Space for ewtQ is allocated even when errconQ=FALSE, 
 * although in this case, ewtQ is never used. The reason for this
 * decision is to allow the user to re-initialize the quadrature
 * computation with errconQ=TRUE, after an initialization with
 * errconQ=FALSE, without new memory allocation within 
 * CVodeQuadReInit.
 */

static booleantype CVQuadAllocVectors(CVodeMem cv_mem, N_Vector tmpl) 
{
  int i, j;

  /* Allocate ewtQ */
  ewtQ = NULL;
  ewtQ = N_VClone(tmpl);
  if (ewtQ == NULL) {
    return(FALSE);
  }
  
  /* Allocate acorQ */
  acorQ = NULL;
  acorQ = N_VClone(tmpl);
  if (acorQ == NULL) {
    N_VDestroy(ewtQ);
    return(FALSE);
  }

  /* Allocate yQ */
  yQ = NULL;
  yQ = N_VClone(tmpl);
  if (yQ == NULL) {
    N_VDestroy(ewtQ);
    N_VDestroy(acorQ);
    return(FALSE);
  }

  /* Allocate tempvQ */
  tempvQ = NULL;
  tempvQ = N_VClone(tmpl);
  if (tempvQ == NULL) {
    N_VDestroy(ewtQ);
    N_VDestroy(acorQ);
    N_VDestroy(yQ);
    return(FALSE);
  }

  /* Allocate zQn[0] ... zQn[maxord] */

  for (j=0; j <= qmax; j++) {
    znQ[j] = NULL;
    znQ[j] = N_VClone(tmpl);
    if (znQ[j] == NULL) {
      N_VDestroy(ewtQ);
      N_VDestroy(acorQ);
      N_VDestroy(yQ);
      N_VDestroy(tempvQ);
      for (i=0; i < j; i++) N_VDestroy(znQ[i]);
      return(FALSE);
    }
  }

  /* Store the value of qmax used here */
  cv_mem->cv_qmax_allocQ = qmax;

  /* Update solver workspace lengths */
  lrw += (qmax + 5)*lrw1Q;
  liw += (qmax + 5)*liw1Q;

  return(TRUE);
}

/*
 * CVQuadFreeVectors
 *
 * This routine frees the CVODES vectors allocated in CVQuadAllocVectors.
 */

static void CVQuadFreeVectors(CVodeMem cv_mem)
{
  int j, maxord;
  
  maxord = cv_mem->cv_qmax_allocQ;

  N_VDestroy(ewtQ);
  N_VDestroy(acorQ);
  N_VDestroy(yQ);
  N_VDestroy(tempvQ);
  
  for (j=0; j<=maxord; j++) N_VDestroy(znQ[j]);

  lrw -= (maxord + 5)*lrw1Q;
  liw -= (maxord + 5)*liw1Q;

  if (cv_mem->cv_VabstolQMallocDone) {
    N_VDestroy(VabstolQ);
    lrw -= lrw1Q;
    liw -= liw1Q;
  }

  cv_mem->cv_VabstolQMallocDone = FALSE;
}

/*
 * CVSensAllocVectors
 *
 * Create (through duplication) N_Vectors used for sensitivity analysis, 
 * using the N_Vector 'tmpl' as a template.
 */

static booleantype CVSensAllocVectors(CVodeMem cv_mem, N_Vector tmpl) 
{
  int i, j;
  
  /* Allocate yS */
  yS = NULL;
  yS = N_VCloneVectorArray(Ns, tmpl);
  if (yS == NULL) {
    return(FALSE);
  }

  /* Allocate ewtS */
  ewtS = NULL;
  ewtS = N_VCloneVectorArray(Ns, tmpl);
  if (ewtS == NULL) {
    N_VDestroyVectorArray(yS, Ns);
    return(FALSE);
  }
  
  /* Allocate acorS */
  acorS = NULL;
  acorS = N_VCloneVectorArray(Ns, tmpl);
  if (acorS == NULL) {
    N_VDestroyVectorArray(yS, Ns);
    N_VDestroyVectorArray(ewtS, Ns);
    return(FALSE);
  }
  
  /* Allocate tempvS */
  tempvS = NULL;
  tempvS = N_VCloneVectorArray(Ns, tmpl);
  if (tempvS == NULL) {
    N_VDestroyVectorArray(yS, Ns);
    N_VDestroyVectorArray(ewtS, Ns);
    N_VDestroyVectorArray(acorS, Ns);
    return(FALSE);
  }
    
  /* Allocate ftempS */
  ftempS = NULL;
  ftempS = N_VCloneVectorArray(Ns, tmpl);
  if (ftempS == NULL) {
    N_VDestroyVectorArray(yS, Ns);
    N_VDestroyVectorArray(ewtS, Ns);
    N_VDestroyVectorArray(acorS, Ns);
    N_VDestroyVectorArray(tempvS, Ns);
    return(FALSE);
  }
  
  /* Allocate znS */
  for (j=0; j<=qmax; j++) {
    znS[j] = NULL;
    znS[j] = N_VCloneVectorArray(Ns, tmpl);
    if (znS[j] == NULL) {
      N_VDestroyVectorArray(yS, Ns);
      N_VDestroyVectorArray(ewtS, Ns);
      N_VDestroyVectorArray(acorS, Ns);
      N_VDestroyVectorArray(tempvS, Ns);
      N_VDestroyVectorArray(ftempS, Ns);
      for (i=0; i<j; i++) N_VDestroyVectorArray(znS[i], Ns);
      return(FALSE);
    }
  }
  
  /* Allocate space for pbar and plist */
  pbar = NULL;
  pbar = (realtype *)malloc(Ns*sizeof(realtype));
  if (pbar == NULL) {
    N_VDestroyVectorArray(yS, Ns);
    N_VDestroyVectorArray(ewtS, Ns);
    N_VDestroyVectorArray(acorS, Ns);
    N_VDestroyVectorArray(tempvS, Ns);
    N_VDestroyVectorArray(ftempS, Ns);
    for (i=0; i<=qmax; i++) N_VDestroyVectorArray(znS[i], Ns);
    return(FALSE);
  }

  plist = NULL;
  plist = (int *)malloc(Ns*sizeof(int));
  if (plist == NULL) {
    N_VDestroyVectorArray(yS, Ns);
    N_VDestroyVectorArray(ewtS, Ns);
    N_VDestroyVectorArray(acorS, Ns);
    N_VDestroyVectorArray(tempvS, Ns);
    N_VDestroyVectorArray(ftempS, Ns);
    for (i=0; i<=qmax; i++) N_VDestroyVectorArray(znS[i], Ns);
    free(pbar); pbar = NULL;
    return(FALSE);
  }

  /* Update solver workspace lengths */
  lrw += (qmax + 6)*Ns*lrw1 + Ns;
  liw += (qmax + 6)*Ns*liw1 + Ns;
  
  /* Store the value of qmax used here */
  cv_mem->cv_qmax_allocS = qmax;

  return(TRUE);
}

/*
 * CVSensFreeVectors
 *
 * This routine frees the CVODES vectors allocated in CVSensAllocVectors.
 */

static void CVSensFreeVectors(CVodeMem cv_mem) 
{
  int j, maxord;
  
  maxord = cv_mem->cv_qmax_allocS;

  N_VDestroyVectorArray(yS, Ns);
  N_VDestroyVectorArray(ewtS, Ns);
  N_VDestroyVectorArray(acorS, Ns);
  N_VDestroyVectorArray(tempvS, Ns);
  N_VDestroyVectorArray(ftempS, Ns);
  
  for (j=0; j<=maxord; j++) N_VDestroyVectorArray(znS[j], Ns);  

  free(pbar); pbar = NULL;
  free(plist); plist = NULL;

  lrw -= (maxord + 6)*Ns*lrw1 + Ns;
  liw -= (maxord + 6)*Ns*liw1 + Ns;

  if (cv_mem->cv_VabstolSMallocDone) {
    N_VDestroyVectorArray(VabstolS, Ns);
    lrw -= Ns*lrw1;
    liw -= Ns*liw1;
  }
  if (cv_mem->cv_SabstolSMallocDone) {
    free(SabstolS); SabstolS = NULL;
    lrw -= Ns;
  }
  cv_mem->cv_VabstolSMallocDone = FALSE;
  cv_mem->cv_SabstolSMallocDone = FALSE;
}

/* 
 * -----------------------------------------------------------------
 * Initial stepsize calculation
 * -----------------------------------------------------------------
 */

/*
 * CVHin
 *
 * This routine computes a tentative initial step size h0. 
 * If tout is too close to tn (= t0), then CVHin returns CV_TOO_CLOSE
 * and h remains uninitialized. 
 * If any RHS function fails unrecoverably, CVHin returns CV_*RHSFUNC_FAIL.
 * If any RHS function fails recoverably too many times and recovery is
 * not possible, CVHin returns CV_REPTD_*RHSFUNC_ERR.
 * Otherwise, CVHin sets h to the chosen value h0 and returns CV_SUCCESS.
 *
 * The algorithm used seeks to find h0 as a solution of
 *       (WRMS norm of (h0^2 ydd / 2)) = 1, 
 * where ydd = estimated second derivative of y. Here, y includes
 * all variables considered in the error test.
 *
 * We start with an initial estimate equal to the geometric mean of the
 * lower and upper bounds on the step size.
 *
 * Loop up to MAX_ITERS times to find h0.
 * Stop if new and previous values differ by a factor < 2.
 * Stop if hnew/hg > 2 after one iteration, as this probably means
 * that the ydd value is bad because of cancellation error.        
 *  
 * For each new proposed hg, we allow MAX_ITERS attempts to
 * resolve a possible recoverable failure from f() by reducing
 * the proposed stepsize by a factor of 0.2. If a legal stepsize
 * still cannot be found, fall back on a previous value if possible,
 * or else return CV_REPTD_RHSFUNC_ERR.
 *
 * Finally, we apply a bias (0.5) and verify that h0 is within bounds.
 */

static int CVHin(CVodeMem cv_mem, realtype tout)
{
  int retval, sign, count1, count2;
  realtype tdiff, tdist, tround, hlb, hub;
  realtype hg, hgs, hs, hnew, hrat, h0, yddnrm;
  booleantype hgOK, hnewOK;

  /* If tout is too close to tn, give up */
  
  if ((tdiff = tout-tn) == ZERO) return(CV_TOO_CLOSE);
  
  sign = (tdiff > ZERO) ? 1 : -1;
  tdist = ABS(tdiff);
  tround = uround * MAX(ABS(tn), ABS(tout));

  if (tdist < TWO*tround) return(CV_TOO_CLOSE);
  
  /* 
     Set lower and upper bounds on h0, and take geometric mean 
     as first trial value.
     Exit with this value if the bounds cross each other.
  */

  hlb = HLB_FACTOR * tround;
  hub = CVUpperBoundH0(cv_mem, tdist);

  hg  = RSqrt(hlb*hub);

  if (hub < hlb) {
    if (sign == -1) h = -hg;
    else            h =  hg;
    return(CV_SUCCESS);
  }
  
  /* Outer loop */

  hnewOK = FALSE;
  hs = hg;         /* safeguard against 'uninitialized variable' warning */

  for(count1 = 1; count1 <= MAX_ITERS; count1++) {

    /* Attempts to estimate ydd */

    hgOK = FALSE;

    for (count2 = 1; count2 <= MAX_ITERS; count2++) {
      hgs = hg*sign;
      retval = CVYddNorm(cv_mem, hgs, &yddnrm);
      /* If a RHS function failed unrecoverably, give up */
      if (retval < 0) return(retval);
      /* If successful, we can use ydd */
      if (retval == CV_SUCCESS) {hgOK = TRUE; break;}
      /* A RHS function failed recoverably; cut step size and test it again */
      hg *= POINT2;
    }

    /* If a RHS function failed recoverably MAX_ITERS times */

    if (!hgOK) {
      /* Exit if this is the first or second pass. No recovery possible */
      if (count1 <= 2) 
        if (retval == RHSFUNC_RECVR)  return(CV_REPTD_RHSFUNC_ERR);
        if (retval == QRHSFUNC_RECVR) return(CV_REPTD_QRHSFUNC_ERR);
        if (retval == SRHSFUNC_RECVR) return(CV_REPTD_SRHSFUNC_ERR);
      /* We have a fall-back option. The value hs is a previous hnew which
         passed through f(). Use it and break */
      hnew = hs;
      break;
    }

    /* The proposed step size is feasible. Save it. */
    hs = hg;

    /* If the stopping criteria was met, or if this is the last pass, stop */
    if ( (hnewOK) || (count1 == MAX_ITERS))  {hnew = hg; break;}

    /* Propose new step size */
    hnew = (yddnrm*hub*hub > TWO) ? RSqrt(TWO/yddnrm) : RSqrt(hg*hub);
    hrat = hnew/hg;
    
    /* Accept hnew if it does not differ from hg by more than a factor of 2 */
    if ((hrat > HALF) && (hrat < TWO)) {
      hnewOK = TRUE;
    }

    /* After one pass, if ydd seems to be bad, use fall-back value. */
    if ((count1 > 1) && (hrat > TWO)) {
      hnew = hg;
      hnewOK = TRUE;
    }

    /* Send this value back through f() */
    hg = hnew;

  }

  /* Apply bounds, bias factor, and attach sign */

  h0 = H_BIAS*hnew;
  if (h0 < hlb) h0 = hlb;
  if (h0 > hub) h0 = hub;
  if (sign == -1) h0 = -h0;
  h = h0;

  return(CV_SUCCESS);
}

/*
 * CVUpperBoundH0
 *
 * This routine sets an upper bound on abs(h0) based on
 * tdist = tn - t0 and the values of y[i]/y'[i].
 */

static realtype CVUpperBoundH0(CVodeMem cv_mem, realtype tdist)
{
  realtype hub_inv, hubQ_inv, hubS_inv, hub;
  N_Vector temp1, temp2;
  N_Vector tempQ1, tempQ2;
  N_Vector *tempS1;
  int is;

  /* 
   * Bound based on |y|/|y'| -- allow at most an increase of
   * HUB_FACTOR in y0 (based on a forward Euler step). The weight 
   * factor is used as a safeguard against zero components in y0. 
   */

  temp1 = tempv;
  temp2 = acor;

  N_VAbs(zn[0], temp2);
  efun(zn[0], temp1, e_data);
  N_VInv(temp1, temp1);
  N_VLinearSum(HUB_FACTOR, temp2, ONE, temp1, temp1);

  N_VAbs(zn[1], temp2);

  N_VDiv(temp2, temp1, temp1);
  hub_inv = N_VMaxNorm(temp1);

  /* Bound based on |yQ|/|yQ'| */
  
  if (quadr && errconQ) {

    tempQ1 = tempvQ;
    tempQ2 = acorQ;

    N_VAbs(znQ[0], tempQ2);
    CVQuadEwtSet(cv_mem, znQ[0], tempQ1);
    N_VInv(tempQ1, tempQ1);
    N_VLinearSum(HUB_FACTOR, tempQ2, ONE, tempQ1, tempQ1);
    
    N_VAbs(znQ[1], tempQ2);
    
    N_VDiv(tempQ2, tempQ1, tempQ1);
    hubQ_inv = N_VMaxNorm(tempQ1);

    if (hubQ_inv > hub_inv) hub_inv = hubQ_inv;

  }

  /* Bound based on |yS|/|yS'| */

  if (sensi && errconS) {

    tempS1 = acorS;
    CVSensEwtSet(cv_mem, znS[0], tempS1);

    for (is=0; is<Ns; is++) {

      N_VAbs(znS[0][is], temp2);
      N_VInv(tempS1[is], temp1);
      N_VLinearSum(HUB_FACTOR, temp2, ONE, temp1, temp1);
      
      N_VAbs(znS[1][is], temp2);
      
      N_VDiv(temp2, temp1, temp1);
      hubS_inv = N_VMaxNorm(temp1);

      if (hubS_inv > hub_inv) hub_inv = hubS_inv;

    }

  }

  /*
   * bound based on tdist -- allow at most a step of magnitude
   * HUB_FACTOR * tdist
   */
  
  hub = HUB_FACTOR*tdist;

  /* Use the smaler of the two */

  if (hub*hub_inv > ONE) hub = ONE/hub_inv;

  return(hub);
}

/*
 * CVYddNorm
 *
 * This routine computes an estimate of the second derivative of Y
 * using a difference quotient, and returns its WRMS norm.
 *
 * Y contains all variables included in the error test. 
 */

static int CVYddNorm(CVodeMem cv_mem, realtype hg, realtype *yddnrm)
{
  int retval, is;
  N_Vector wrk1, wrk2;
  
  /* y     <- h*y'(t) + y(t) */
  
  N_VLinearSum(hg, zn[1], ONE, zn[0], y);
  
  if (sensi && errconS) 
    for (is=0; is<Ns; is++)
      N_VLinearSum(hg, znS[1][is], ONE, znS[0][is], yS[is]);
  
  /* tempv <- f(t+h, h*y'(t)+y(t)) */

  retval = f(tn+hg, y, tempv, f_data);
  nfe++;
  if (retval < 0) return(CV_RHSFUNC_FAIL);
  if (retval > 0) return(RHSFUNC_RECVR);

  if (quadr && errconQ) {
    retval = fQ(tn+hg, y, tempvQ, fQ_data);
    nfQe++;
    if (retval < 0) return(CV_QRHSFUNC_FAIL);
    if (retval > 0) return(QRHSFUNC_RECVR);
  }

  if (sensi && errconS) {
    wrk1 = ftemp;
    wrk2 = acor;
    retval = CVSensRhs(cv_mem, tn+hg, y, tempv, yS, tempvS, wrk1, wrk2);
    if (retval < 0) return(CV_SRHSFUNC_FAIL);
    if (retval > 0) return(SRHSFUNC_RECVR);
  }  

  /* tempv <- f(t+h, h*y'(t)+y(t)) - y'(t) */
  /* tempv <- ydd */
  
  N_VLinearSum(ONE, tempv, -ONE, zn[1], tempv);
  N_VScale(ONE/hg, tempv, tempv);
  
  if (quadr && errconQ) {
    N_VLinearSum(ONE, tempvQ, -ONE, znQ[1], tempvQ);
    N_VScale(ONE/hg, tempvQ, tempvQ);
  }

  if (sensi && errconS)
    for (is=0; is<Ns; is++) {
      N_VLinearSum(ONE, tempvS[is], -ONE, znS[1][is], tempvS[is]);
      N_VScale(ONE/hg, tempvS[is], tempvS[is]);
    }

  /* Estimate ||y''|| */
  
  *yddnrm = N_VWrmsNorm(tempv, ewt);
  if (quadr && errconQ) {
    *yddnrm = CVQuadUpdateNorm(cv_mem, *yddnrm, tempvQ, ewtQ);
  }
  if (sensi && errconS) {
    *yddnrm = CVSensUpdateNorm(cv_mem, *yddnrm, tempvS, ewtS);
  }

  return(CV_SUCCESS);
}

/* 
 * -----------------------------------------------------------------
 * Initial setup
 * -----------------------------------------------------------------
 */

/*  
 * CVInitialSetup
 *
 * This routine performs input consistency checks at the first step.
 * If needed, it also checks the linear solver module and calls the
 * linear solver initialization routine.
 */

static int CVInitialSetup(CVodeMem cv_mem)
{
  int ier;

  /* Did the user provide efun? */

  if (itol != CV_WF) {
    efun = CVEwtSet;
    e_data = (void *)cv_mem;
  } else {
    if (efun == NULL) {
      CVProcessError(cv_mem, CV_ILL_INPUT, "CVODES", "CVInitialSetup", MSGCV_NO_EFUN);
      return(CV_ILL_INPUT);
    }
  }

  ier = efun(zn[0], ewt, e_data);
  if (ier != 0) {

    if (itol == CV_WF) 
      CVProcessError(cv_mem, CV_ILL_INPUT, "CVODES", "CVInitialSetup", MSGCV_EWT_FAIL);
    else
      CVProcessError(cv_mem, CV_ILL_INPUT, "CVODES", "CVInitialSetup", MSGCV_BAD_EWT);

    return(CV_ILL_INPUT);
  }
  
  /* Quadrature initial setup */

  if (quadr && errconQ) {

    /* Load ewtQ */
    ier = CVQuadEwtSet(cv_mem, znQ[0], ewtQ);
    if (ier != 0) {
      CVProcessError(cv_mem, CV_ILL_INPUT, "CVODES", "CVInitialSetup", MSGCV_BAD_EWTQ);
      return(CV_ILL_INPUT);
    }

  }

  if (!quadr) errconQ = FALSE;

  /* Forward sensitivity initial setup */

  if (sensi) {

    /* Check if ism and ifS agree */
    if ((ism==CV_STAGGERED1) && (ifS==CV_ALLSENS)) {
      CVProcessError(cv_mem, CV_ILL_INPUT, "CVODES", "CVInitialSetup", MSGCV_BAD_ISM_IFS);
      return(CV_ILL_INPUT);
    }    

    /* Load ewtS */
    ier = CVSensEwtSet(cv_mem, znS[0], ewtS);
    if (ier != 0) {
      CVProcessError(cv_mem, CV_ILL_INPUT, "CVODES", "CVInitialSetup", MSGCV_BAD_EWTS);
      return(CV_ILL_INPUT);
    }

  }

  /* Check if lsolve function exists (if needed)
     and call linit function (if it exists) */

  if (iter == CV_NEWTON) {
    if (lsolve == NULL) {
      CVProcessError(cv_mem, CV_ILL_INPUT, "CVODES", "CVInitialSetup", MSGCV_LSOLVE_NULL);
      return(CV_ILL_INPUT);
    }
    if (linit != NULL) {
      ier = linit(cv_mem);
      if (ier != 0) {
        CVProcessError(cv_mem, CV_LINIT_FAIL, "CVODES", "CVInitialSetup", MSGCV_LINIT_FAIL);
        return(CV_LINIT_FAIL);
      }
    }
  }
    
  return(CV_SUCCESS);
}

/*
 * CVEwtSet
 *
 * This routine is responsible for setting the error weight vector ewt,
 * according to tol_type, as follows:
 *
 * (1) ewt[i] = 1 / (reltol * ABS(ycur[i]) + *abstol), i=0,...,neq-1
 *     if tol_type = CV_SS
 * (2) ewt[i] = 1 / (reltol * ABS(ycur[i]) + abstol[i]), i=0,...,neq-1
 *     if tol_type = CV_SV
 *
 * CVEwtSet returns 0 if ewt is successfully set as above to a
 * positive vector and -1 otherwise. In the latter case, ewt is
 * considered undefined.
 *
 * All the real work is done in the routines CVEwtSetSS, CVEwtSetSV.
 */

int CVEwtSet(N_Vector ycur, N_Vector weight, void *data)
{
  CVodeMem cv_mem;
  int flag = 0;

  /* data points to cv_mem here */

  cv_mem = (CVodeMem) data;

  switch(itol) {
  case CV_SS: 
    flag = CVEwtSetSS(cv_mem, ycur, weight);
    break;
  case CV_SV: 
    flag = CVEwtSetSV(cv_mem, ycur, weight);
    break;
  }

  return(flag);
}

/*
 * CVEwtSetSS
 *
 * This routine sets ewt as decribed above in the case tol_type = CV_SS.
 * It tests for non-positive components before inverting. CVEwtSetSS
 * returns 0 if ewt is successfully set to a positive vector
 * and -1 otherwise. In the latter case, ewt is considered undefined.
 */

static int CVEwtSetSS(CVodeMem cv_mem, N_Vector ycur, N_Vector weight)
{
  N_VAbs(ycur, tempv);
  N_VScale(reltol, tempv, tempv);
  N_VAddConst(tempv, Sabstol, tempv);
  if (N_VMin(tempv) <= ZERO) return(-1);
  N_VInv(tempv, weight);

  return(0);
}

/*
 * CVEwtSetSV
 *
 * This routine sets ewt as decribed above in the case tol_type = CV_SV.
 * It tests for non-positive components before inverting. CVEwtSetSV
 * returns 0 if ewt is successfully set to a positive vector
 * and -1 otherwise. In the latter case, ewt is considered undefined.
 */

static int CVEwtSetSV(CVodeMem cv_mem, N_Vector ycur, N_Vector weight)
{
  N_VAbs(ycur, tempv);
  N_VLinearSum(reltol, tempv, ONE, Vabstol, tempv);
  if (N_VMin(tempv) <= ZERO) return(-1);
  N_VInv(tempv, weight);
  return(0);
}

/*
 * CVQuadEwtSet
 *
 */

static int CVQuadEwtSet(CVodeMem cv_mem, N_Vector qcur, N_Vector weightQ)
{
  int flag=0;

  switch (itolQ) {
  case CV_SS: 
    flag = CVQuadEwtSetSS(cv_mem, qcur, weightQ);
    break;
  case CV_SV: 
    flag = CVQuadEwtSetSV(cv_mem, qcur, weightQ);
    break;
  }

  return(flag);

}

/*
 * CVQuadEwtSetSS
 *
 */

static int CVQuadEwtSetSS(CVodeMem cv_mem, N_Vector qcur, N_Vector weightQ)
{
  N_VAbs(qcur, tempvQ);
  N_VScale(reltolQ, tempvQ, tempvQ);
  N_VAddConst(tempvQ, SabstolQ, tempvQ);
  if (N_VMin(tempvQ) <= ZERO) return(-1);
  N_VInv(tempvQ, weightQ);

  return(0);
}

/*
 * CVQuadEwtSetSV
 *
 */

static int CVQuadEwtSetSV(CVodeMem cv_mem, N_Vector qcur, N_Vector weightQ)
{
  N_VAbs(qcur, tempvQ);
  N_VLinearSum(reltolQ, tempvQ, ONE, VabstolQ, tempvQ);
  if (N_VMin(tempvQ) <= ZERO) return(-1);
  N_VInv(tempvQ, weightQ);

  return(0);
}

/*
 * CVSensEwtSet
 *
 */

static int CVSensEwtSet(CVodeMem cv_mem, N_Vector *yScur, N_Vector *weightS)
{
  int flag=0;

  switch (itolS) {
  case CV_EE:
    flag = CVSensEwtSetEE(cv_mem, yScur, weightS);
    break;
  case CV_SS: 
    flag = CVSensEwtSetSS(cv_mem, yScur, weightS);
    break;
  case CV_SV: 
    flag = CVSensEwtSetSV(cv_mem, yScur, weightS);
    break;
  }

  return(flag);

}

/*
 * CVSensEwtSetEE
 *
 * In this case, the error weight vector for the i-th sensitivity is set to
 *
 * ewtS_i = pbar_i * efun(pbar_i*yS_i)
 *
 * In other words, the scaled sensitivity pbar_i * yS_i has the same error
 * weight vector calculation as the solution vector.
 *
 */

static int CVSensEwtSetEE(CVodeMem cv_mem, N_Vector *yScur, N_Vector *weightS)
{
  int is;
  N_Vector pyS;
  int flag;

  /* Use tempvS[0] as temporary storage for the scaled sensitivity */
  pyS = tempvS[0];

  for (is=0; is<Ns; is++) {
    N_VScale(pbar[is], yScur[is], pyS);
    flag = efun(pyS, weightS[is], e_data);
    if (flag != 0) return(-1);
    N_VScale(pbar[is], weightS[is], weightS[is]);
  }

  return(0);
}

/*
 * CVSensEwtSetSS
 *
 */

static int CVSensEwtSetSS(CVodeMem cv_mem, N_Vector *yScur, N_Vector *weightS)
{
  int is;
  
  for (is=0; is<Ns; is++) {
    N_VAbs(yScur[is], tempv);
    N_VScale(reltolS, tempv, tempv);
    N_VAddConst(tempv, SabstolS[is], tempv);
    if (N_VMin(tempv) <= ZERO) return(-1);
    N_VInv(tempv, weightS[is]);
  }
  return(0);
}

/*
 * CVSensEwtSetSV
 *
 */

static int CVSensEwtSetSV(CVodeMem cv_mem, N_Vector *yScur, N_Vector *weightS)
{
  int is;
  
  for (is=0; is<Ns; is++) {
    N_VAbs(yScur[is], tempv);
    N_VLinearSum(reltolS, tempv, ONE, VabstolS[is], tempv);
    if (N_VMin(tempv) <= ZERO) return(-1);
    N_VInv(tempv, weightS[is]);
  }

  return(0);
}

/* 
 * -----------------------------------------------------------------
 * Main CVStep function
 * -----------------------------------------------------------------
 */

/* 
 * CVStep
 *
 * This routine performs one internal cvode step, from tn to tn + h.
 * It calls other routines to do all the work.
 *
 * The main operations done here are as follows:
 * - preliminary adjustments if a new step size was chosen;
 * - prediction of the Nordsieck history array zn at tn + h;
 * - setting of multistep method coefficients and test quantities;
 * - solution of the nonlinear system;
 * - testing the local error;
 * - updating zn and other state data if successful;
 * - resetting stepsize and order for the next step.
 * - if SLDET is on, check for stability, reduce order if necessary.
 * On a failure in the nonlinear system solution or error test, the
 * step may be reattempted, depending on the nature of the failure.
 */

static int CVStep(CVodeMem cv_mem)
{
  realtype saved_t, dsm, dsmS, dsmQ;
  booleantype do_sensi_stg, do_sensi_stg1;
  int ncf, nef, ncfS, nefS, nefQ;
  int nflag, kflag, eflag;
  int retval, is;

  /* Are we computing sensitivities with a staggered approach? */

  do_sensi_stg  = (sensi && (ism==CV_STAGGERED));
  do_sensi_stg1 = (sensi && (ism==CV_STAGGERED1));

  /* Initialize local counters for convergence and error test failures */

  ncf  = nef  = 0;
  nefQ = 0;
  ncfS = nefS = 0;
  if (do_sensi_stg1)
    for (is=0; is<Ns; is++) ncfS1[is] = 0;

  /* If needed, adjust method parameters */

  if ((nst > 0) && (hprime != h)) CVAdjustParams(cv_mem);

  /* Looping point for attempts to take a step */

  saved_t = tn;
  nflag = FIRST_CALL;

  loop {  

    CVPredict(cv_mem);  
    CVSet(cv_mem);

    /* ------ Correct state variables ------ */
    
    nflag = CVNls(cv_mem, nflag);
    kflag = CVHandleNFlag(cv_mem, &nflag, saved_t, &ncf, &ncfn);

    /* Go back in loop if we need to predict again (nflag=PREV_CONV_FAIL) */
    if (kflag == PREDICT_AGAIN) continue;

    /* Return if nonlinear solve failed and recovery not possible. */
    if (kflag != DO_ERROR_TEST) return(kflag);

    /* Perform error test (nflag=CV_SUCCESS) */
    eflag = CVDoErrorTest(cv_mem, &nflag, saved_t, acnrm, &nef, &netf, &dsm);

    /* Go back in loop if we need to predict again (nflag=PREV_ERR_FAIL) */
    if (eflag == TRY_AGAIN) continue;

    /* Return if error test failed and recovery not possible. */
    if (eflag != CV_SUCCESS) return(eflag);

    /* Error test passed (eflag=CV_SUCCESS, nflag=CV_SUCCESS), go on */

    /* ------ Correct the quadrature variables ------ */

    if (quadr) {

      ncf = nef = 0; /* reset counters for states */

      nflag = CVQuadNls(cv_mem);
      kflag = CVHandleNFlag(cv_mem, &nflag, saved_t, &ncf, &ncfn);

      if (kflag == PREDICT_AGAIN) continue;
      if (kflag != DO_ERROR_TEST) return(kflag);

      /* Error test on quadratures */
      if (errconQ) {
        acnrmQ = N_VWrmsNorm(acorQ, ewtQ);
        eflag = CVDoErrorTest(cv_mem, &nflag, saved_t, acnrmQ, &nefQ, &netfQ, &dsmQ);

        if (eflag == TRY_AGAIN)  continue;
        if (eflag != CV_SUCCESS) return(eflag);

        /* update 'dsm' with 'dsmQ' (to be used in CVPrepareNextStep) */
        dsm = CVQuadUpdateDsm(cv_mem, dsm, dsmQ);
      }

    }

    /* ------ Correct the sensitivity variables (STAGGERED or STAGGERED1) ------- */

    if (do_sensi_stg || do_sensi_stg1) {

      ncf = nef = 0;        /* reset counters for states     */
      if (quadr) nefQ = 0;  /* reset counter for quadratures */

      /* Evaluate f at converged y, needed for future evaluations of sens. RHS 
         If f() fails recoverably, treat it as a convergence failure and 
         attempt the step again */

      retval = f(tn, y, ftemp, f_data);
      nfe++;
      if (retval < 0) return(CV_RHSFUNC_FAIL);
      if (retval > 0) {
        nflag = PREV_CONV_FAIL;
        continue;
      }

      if (do_sensi_stg) {
        /* Nonlinear solve for sensitivities (all-at-once) */
        nflag = CVStgrNls(cv_mem);
        kflag = CVHandleNFlag(cv_mem, &nflag, saved_t, &ncfS, &ncfnS);
      } else {
        /* Nonlinear solve for sensitivities (one-by-one) */
        for (is=0; is<Ns; is++) { 
          nflag = CVStgr1Nls(cv_mem, is); 
          kflag = CVHandleNFlag(cv_mem, &nflag, saved_t, &ncfS1[is], &ncfnS1[is]);
          if (kflag != DO_ERROR_TEST) break; 
        }
      }

      if (kflag == PREDICT_AGAIN) continue;
      if (kflag != DO_ERROR_TEST) return(kflag);

      /* Error test on sensitivities */
      if (errconS) {

        if (do_sensi_stg1) acnrmS = CVSensNorm(cv_mem, acorS, ewtS);

        eflag = CVDoErrorTest(cv_mem, &nflag, saved_t, acnrmS, &nefS, &netfS, &dsmS);

        if (eflag == TRY_AGAIN)  continue;
        if (eflag != CV_SUCCESS) return(eflag);

        /* update 'dsm' with 'dsmS' (to be used in CVPrepareNextStep) */
        dsm = CVStgrUpdateDsm(cv_mem, dsm, dsmS);
      }

    }

    /* Everything went fine; exit loop */ 
    break;

  }

  /* Nonlinear system solve and error test were both successful.
     Update data, and consider change of step and/or order.       */

  CVCompleteStep(cv_mem); 

  CVPrepareNextStep(cv_mem, dsm); 

  /* If Stablilty Limit Detection is turned on, call stability limit
     detection routine for possible order reduction. */

  if (sldeton) CVBDFStab(cv_mem);

  etamax = (nst <= SMALL_NST) ? ETAMX2 : ETAMX3;

  /*  Finally, we rescale the acor array to be the 
      estimated local error vector. */

  N_VScale(ONE/tq[2], acor, acor);

  if (quadr)
    N_VScale(ONE/tq[2], acorQ, acorQ);

  if (sensi)
    for (is=0; is<Ns; is++)
      N_VScale(ONE/tq[2], acorS[is], acorS[is]);

  return(CV_SUCCESS);
      
}

/* 
 * -----------------------------------------------------------------
 * Function called at beginning of step
 * -----------------------------------------------------------------
 */

/*
 * CVAdjustParams
 *
 * This routine is called when a change in step size was decided upon,
 * and it handles the required adjustments to the history array zn.
 * If there is to be a change in order, we call CVAdjustOrder and reset
 * q, L = q+1, and qwait.  Then in any case, we call CVRescale, which
 * resets h and rescales the Nordsieck array.
 */

static void CVAdjustParams(CVodeMem cv_mem)
{
  if (qprime != q) {
    CVAdjustOrder(cv_mem, qprime-q);
    q = qprime;
    L = q+1;
    qwait = L;
  }
  CVRescale(cv_mem);
}

/*
 * CVAdjustOrder
 *
 * This routine is a high level routine which handles an order
 * change by an amount deltaq (= +1 or -1). If a decrease in order
 * is requested and q==2, then the routine returns immediately.
 * Otherwise CVAdjustAdams or CVAdjustBDF is called to handle the
 * order change (depending on the value of lmm).
 */

static void CVAdjustOrder(CVodeMem cv_mem, int deltaq)
{
  if ((q==2) && (deltaq != 1)) return;
  
  switch(lmm){
  case CV_ADAMS: 
    CVAdjustAdams(cv_mem, deltaq);
    break;
  case CV_BDF:   
    CVAdjustBDF(cv_mem, deltaq);
    break;
  }
}

/*
 * CVAdjustAdams
 *
 * This routine adjusts the history array on a change of order q by
 * deltaq, in the case that lmm == CV_ADAMS.
 */

static void CVAdjustAdams(CVodeMem cv_mem, int deltaq)
{
  int i, j;
  int is;
  realtype xi, hsum;

  /* On an order increase, set new column of zn to zero and return */
  
  if (deltaq==1) {
    N_VConst(ZERO, zn[L]);
    if (quadr)
      N_VConst(ZERO, znQ[L]);
    if (sensi)
      for (is=0; is<Ns; is++)
        N_VConst(ZERO, znS[L][is]);
    return;
  }

  /*
   * On an order decrease, each zn[j] is adjusted by a multiple of zn[q].
   * The coeffs. in the adjustment are the coeffs. of the polynomial:
   *        x
   * q * INT { u * ( u + xi_1 ) * ... * ( u + xi_{q-2} ) } du 
   *        0
   * where xi_j = [t_n - t_(n-j)]/h => xi_0 = 0
   */

  for (i=0; i <= qmax; i++) l[i] = ZERO;
  l[1] = ONE;
  hsum = ZERO;
  for (j=1; j <= q-2; j++) {
    hsum += tau[j];
    xi = hsum / hscale;
    for (i=j+1; i >= 1; i--) l[i] = l[i]*xi + l[i-1];
  }
  
  for (j=1; j <= q-2; j++) l[j+1] = q * (l[j] / (j+1));
  
  for (j=2; j < q; j++)
    N_VLinearSum(-l[j], zn[q], ONE, zn[j], zn[j]);

  if (quadr)
    for (j=2; j < q; j++)
      N_VLinearSum(-l[j], znQ[q], ONE, znQ[j], znQ[j]);

  if (sensi)
    for (is=0; is<Ns; is++)
      for (j=2; j < q; j++)
        N_VLinearSum(-l[j], znS[q][is], ONE, znS[j][is], znS[j][is]);

}

/*
 * CVAdjustBDF
 *
 * This is a high level routine which handles adjustments to the
 * history array on a change of order by deltaq in the case that 
 * lmm == CV_BDF.  CVAdjustBDF calls CVIncreaseBDF if deltaq = +1 and 
 * CVDecreaseBDF if deltaq = -1 to do the actual work.
 */

static void CVAdjustBDF(CVodeMem cv_mem, int deltaq)
{
  switch(deltaq) {
  case 1: 
    CVIncreaseBDF(cv_mem);
    return;
  case -1: 
    CVDecreaseBDF(cv_mem);
    return;
  }
}

/*
 * CVIncreaseBDF
 *
 * This routine adjusts the history array on an increase in the 
 * order q in the case that lmm == CV_BDF.  
 * A new column zn[q+1] is set equal to a multiple of the saved 
 * vector (= acor) in zn[indx_acor].  Then each zn[j] is adjusted by
 * a multiple of zn[q+1].  The coefficients in the adjustment are the 
 * coefficients of the polynomial x*x*(x+xi_1)*...*(x+xi_j),
 * where xi_j = [t_n - t_(n-j)]/h.
 */

static void CVIncreaseBDF(CVodeMem cv_mem)
{
  realtype alpha0, alpha1, prod, xi, xiold, hsum, A1;
  int i, j;
  int is;

  for (i=0; i <= qmax; i++) l[i] = ZERO;
  l[2] = alpha1 = prod = xiold = ONE;
  alpha0 = -ONE;
  hsum = hscale;
  if (q > 1) {
    for (j=1; j < q; j++) {
      hsum += tau[j+1];
      xi = hsum / hscale;
      prod *= xi;
      alpha0 -= ONE / (j+1);
      alpha1 += ONE / xi;
      for (i=j+2; i >= 2; i--) l[i] = l[i]*xiold + l[i-1];
      xiold = xi;
    }
  }
  A1 = (-alpha0 - alpha1) / prod;

  /* 
     zn[indx_acor] contains the value Delta_n = y_n - y_n(0) 
     This value was stored there at the previous successful
     step (in CVCompleteStep) 
     
     A1 contains dbar = (1/xi* - 1/xi_q)/prod(xi_j)
  */
  
  N_VScale(A1, zn[indx_acor], zn[L]);
  for (j=2; j <= q; j++)
    N_VLinearSum(l[j], zn[L], ONE, zn[j], zn[j]);

  if (quadr) {
    N_VScale(A1, znQ[indx_acor], znQ[L]);
    for (j=2; j <= q; j++)
      N_VLinearSum(l[j], znQ[L], ONE, znQ[j], znQ[j]);
  }

  if (sensi) {
    for (is=0; is<Ns; is++) {
      N_VScale(A1, znS[indx_acor][is], znS[L][is]);
      for (j=2; j <= q; j++)
        N_VLinearSum(l[j], znS[L][is], ONE, znS[j][is], znS[j][is]);
    }
  }

}

/*
 * CVDecreaseBDF
 *
 * This routine adjusts the history array on a decrease in the 
 * order q in the case that lmm == CV_BDF.  
 * Each zn[j] is adjusted by a multiple of zn[q].  The coefficients
 * in the adjustment are the coefficients of the polynomial
 *   x*x*(x+xi_1)*...*(x+xi_j), where xi_j = [t_n - t_(n-j)]/h.
 */

static void CVDecreaseBDF(CVodeMem cv_mem)
{
  realtype hsum, xi;
  int i, j;
  int is;
  
  for (i=0; i <= qmax; i++) l[i] = ZERO;
  l[2] = ONE;
  hsum = ZERO;
  for (j=1; j <= q-2; j++) {
    hsum += tau[j];
    xi = hsum /hscale;
    for (i=j+2; i >= 2; i--) l[i] = l[i]*xi + l[i-1];
  }
  
  for (j=2; j < q; j++)
    N_VLinearSum(-l[j], zn[q], ONE, zn[j], zn[j]);

  if (quadr) {
    for (j=2; j < q; j++)
      N_VLinearSum(-l[j], znQ[q], ONE, znQ[j], znQ[j]);
  }

  if (sensi) {
    for (is=0; is<Ns; is++) 
      for (j=2; j < q; j++)
        N_VLinearSum(-l[j], znS[q][is], ONE, znS[j][is], znS[j][is]);
  }
}

/*
 * CVRescale
 *
 * This routine rescales the Nordsieck array by multiplying the
 * jth column zn[j] by eta^j, j = 1, ..., q.  Then the value of
 * h is rescaled by eta, and hscale is reset to h.
 */

static void CVRescale(CVodeMem cv_mem)
{
  int j;
  int is;
  realtype factor;

  factor = eta;
  for (j=1; j <= q; j++) {

    N_VScale(factor, zn[j], zn[j]);

    if (quadr)
      N_VScale(factor, znQ[j], znQ[j]);

    if (sensi)
      for (is=0; is<Ns; is++)
        N_VScale(factor, znS[j][is], znS[j][is]);

    factor *= eta;

  }
  h = hscale * eta;
  next_h = h;
  hscale = h;
  nscon = 0;
}

/*
 * CVPredict
 *
 * This routine advances tn by the tentative step size h, and computes
 * the predicted array z_n(0), which is overwritten on zn.  The
 * prediction of zn is done by repeated additions.
 * In TSTOP mode, it is possible for tn + h to be past tstop by roundoff,
 * and in that case, we reset tn (after incrementing by h) to tstop.
 */

static void CVPredict(CVodeMem cv_mem)
{
  int j, k;
  int is;

  tn += h;
  if (istop) {
    if ((tn - tstop)*h > ZERO) tn = tstop;
  }

  for (k = 1; k <= q; k++)
    for (j = q; j >= k; j--) 
      N_VLinearSum(ONE, zn[j-1], ONE, zn[j], zn[j-1]); 

  if (quadr) {
    for (k = 1; k <= q; k++)
      for (j = q; j >= k; j--) 
        N_VLinearSum(ONE, znQ[j-1], ONE, znQ[j], znQ[j-1]);
  }

  if (sensi) {
    for (is=0; is<Ns; is++) {
      for (k = 1; k <= q; k++)
        for (j = q; j >= k; j--) 
          N_VLinearSum(ONE, znS[j-1][is], ONE, znS[j][is], znS[j-1][is]);
    }
  }
}

/*
 * CVSet
 *
 * This routine is a high level routine which calls CVSetAdams or
 * CVSetBDF to set the polynomial l, the test quantity array tq, 
 * and the related variables  rl1, gamma, and gamrat.
 */

static void CVSet(CVodeMem cv_mem)
{
  switch(lmm) {
  case CV_ADAMS: 
    CVSetAdams(cv_mem);
    break;
  case CV_BDF: 
    CVSetBDF(cv_mem);
    break;
  }
  rl1 = ONE / l[1];
  gamma = h * rl1;
  if (nst == 0) gammap = gamma;
  gamrat = (nst > 0) ? gamma / gammap : ONE;  /* protect x / x != 1.0 */
}

/*
 * CVSetAdams
 *
 * This routine handles the computation of l and tq for the
 * case lmm == CV_ADAMS.
 *
 * The components of the array l are the coefficients of a
 * polynomial Lambda(x) = l_0 + l_1 x + ... + l_q x^q, given by
 *                          q-1
 * (d/dx) Lambda(x) = c * PRODUCT (1 + x / xi_i) , where
 *                          i=1
 *  Lambda(-1) = 0, Lambda(0) = 1, and c is a normalization factor.
 * Here xi_i = [t_n - t_(n-i)] / h.
 *
 * The array tq is set to test quantities used in the convergence
 * test, the error test, and the selection of h at a new order.
 */

static void CVSetAdams(CVodeMem cv_mem)
{
  realtype m[L_MAX], M[3], hsum;
  
  if (q == 1) {
    l[0] = l[1] = tq[1] = tq[5] = ONE;
    tq[2] = TWO;
    tq[3] = TWELVE;
    tq[4] = nlscoef * tq[2];       /* = 0.1 * tq[2] */
    return;
  }
  
  hsum = CVAdamsStart(cv_mem, m);
  
  M[0] = CVAltSum(q-1, m, 1);
  M[1] = CVAltSum(q-1, m, 2);
  
  CVAdamsFinish(cv_mem, m, M, hsum);
}

/*
 * CVAdamsStart
 *
 * This routine generates in m[] the coefficients of the product
 * polynomial needed for the Adams l and tq coefficients for q > 1.
 */

static realtype CVAdamsStart(CVodeMem cv_mem, realtype m[])
{
  realtype hsum, xi_inv, sum;
  int i, j;
  
  hsum = h;
  m[0] = ONE;
  for (i=1; i <= q; i++) m[i] = ZERO;
  for (j=1; j < q; j++) {
    if ((j==q-1) && (qwait == 1)) {
      sum = CVAltSum(q-2, m, 2);
      tq[1] = m[q-2] / (q * sum);
    }
    xi_inv = h / hsum;
    for (i=j; i >= 1; i--) m[i] += m[i-1] * xi_inv;
    hsum += tau[j];
    /* The m[i] are coefficients of product(1 to j) (1 + x/xi_i) */
  }
  return(hsum);
}

/*
 * CVAdamsFinish
 *
 * This routine completes the calculation of the Adams l and tq.
 */

static void CVAdamsFinish(CVodeMem cv_mem, realtype m[], realtype M[], realtype hsum)
{
  int i;
  realtype M0_inv, xi, xi_inv;
  
  M0_inv = ONE / M[0];
  
  l[0] = ONE;
  for (i=1; i <= q; i++) l[i] = M0_inv * (m[i-1] / i);
  xi = hsum / h;
  xi_inv = ONE / xi;
  
  tq[2] = xi * M[0] / M[1];
  tq[5] = xi / l[q];

  if (qwait == 1) {
    for (i=q; i >= 1; i--) m[i] += m[i-1] * xi_inv;
    M[2] = CVAltSum(q, m, 2);
    tq[3] = L * M[0] / M[2];
  }

  tq[4] = nlscoef * tq[2];
}

/*  
 * CVAltSum
 *
 * CVAltSum returns the value of the alternating sum
 *   sum (i= 0 ... iend) [ (-1)^i * (a[i] / (i + k)) ].
 * If iend < 0 then CVAltSum returns 0.
 * This operation is needed to compute the integral, from -1 to 0,
 * of a polynomial x^(k-1) M(x) given the coefficients of M(x).
 */

static realtype CVAltSum(int iend, realtype a[], int k)
{
  int i, sign;
  realtype sum;
  
  if (iend < 0) return(ZERO);
  
  sum = ZERO;
  sign = 1;
  for (i=0; i <= iend; i++) {
    sum += sign * (a[i] / (i+k));
    sign = -sign;
  }
  return(sum);
}

/*
 * CVSetBDF
 *
 * This routine computes the coefficients l and tq in the case
 * lmm == CV_BDF.  CVSetBDF calls CVSetTqBDF to set the test
 * quantity array tq. 
 * 
 * The components of the array l are the coefficients of a
 * polynomial Lambda(x) = l_0 + l_1 x + ... + l_q x^q, given by
 *                                 q-1
 * Lambda(x) = (1 + x / xi*_q) * PRODUCT (1 + x / xi_i) , where
 *                                 i=1
 *  xi_i = [t_n - t_(n-i)] / h.
 *
 * The array tq is set to test quantities used in the convergence
 * test, the error test, and the selection of h at a new order.
 */

static void CVSetBDF(CVodeMem cv_mem)
{
  realtype alpha0, alpha0_hat, xi_inv, xistar_inv, hsum;
  int i,j;
  
  l[0] = l[1] = xi_inv = xistar_inv = ONE;
  for (i=2; i <= q; i++) l[i] = ZERO;
  alpha0 = alpha0_hat = -ONE;
  hsum = h;
  if (q > 1) {
    for (j=2; j < q; j++) {
      hsum += tau[j-1];
      xi_inv = h / hsum;
      alpha0 -= ONE / j;
      for (i=j; i >= 1; i--) l[i] += l[i-1]*xi_inv;
      /* The l[i] are coefficients of product(1 to j) (1 + x/xi_i) */
    }
    
    /* j = q */
    alpha0 -= ONE / q;
    xistar_inv = -l[1] - alpha0;
    hsum += tau[q-1];
    xi_inv = h / hsum;
    alpha0_hat = -l[1] - xi_inv;
    for (i=q; i >= 1; i--) l[i] += l[i-1]*xistar_inv;
  }

  CVSetTqBDF(cv_mem, hsum, alpha0, alpha0_hat, xi_inv, xistar_inv);
}

/*
 * CVSetTqBDF
 *
 * This routine sets the test quantity array tq in the case
 * lmm == CV_BDF.
 */

static void CVSetTqBDF(CVodeMem cv_mem, realtype hsum, realtype alpha0,
                       realtype alpha0_hat, realtype xi_inv, realtype xistar_inv)
{
  realtype A1, A2, A3, A4, A5, A6;
  realtype C, CPrime, CPrimePrime;
  
  A1 = ONE - alpha0_hat + alpha0;
  A2 = ONE + q * A1;
  tq[2] = ABS(alpha0 * (A2 / A1));
  tq[5] = ABS((A2) / (l[q] * xi_inv/xistar_inv));
  if (qwait == 1) {
    C = xistar_inv / l[q];
    A3 = alpha0 + ONE / q;
    A4 = alpha0_hat + xi_inv;
    CPrime = A3 / (ONE - A4 + A3);
    tq[1] = ABS(CPrime / C);
    hsum += tau[q];
    xi_inv = h / hsum;
    A5 = alpha0 - (ONE / (q+1));
    A6 = alpha0_hat - xi_inv;
    CPrimePrime = A2 / (ONE - A6 + A5);
    tq[3] = ABS(CPrimePrime * xi_inv * (q+2) * A5);
  }
  tq[4] = nlscoef * tq[2];
}

/* 
 * -----------------------------------------------------------------
 * Nonlinear solver functions
 * -----------------------------------------------------------------
 */

/*
 * CVNls
 *
 * This routine attempts to solve the nonlinear system associated
 * with a single implicit step of the linear multistep method.
 * Depending on iter, it calls CVNlsFunctional or CVNlsNewton
 * to do the work.
 */

static int CVNls(CVodeMem cv_mem, int nflag)
{
  int flag=CV_SUCCESS;

  switch(iter) {
  case CV_FUNCTIONAL: 
    flag = CVNlsFunctional(cv_mem);
    break;
  case CV_NEWTON: 
    flag = CVNlsNewton(cv_mem, nflag);
    break;
  }
  
  return(flag);

}

/*
 * CVNlsFunctional
 *
 * This routine attempts to solve the nonlinear system using 
 * functional iteration (no matrices involved).
 * 
 * This routine also handles the functional iteration of the
 * combined system (states + sensitivities) when sensitivities are 
 * computed using the CV_SIMULTANEOUS approach.
 *
 * Possible return values are:
 *
 *   CV_SUCCESS       ---> continue with error test
 *
 *   CV_RHSFUNC_FAIL  -+   
 *   CV_SRHSFUNC_FAIL -+-> halt the integration 
 *
 *   CONV_FAIL        -+
 *   RHSFUNC_RECVR     |-> predict again or stop if too many
 *   SRHSFUNC_RECVR   -+
 *
 */

static int CVNlsFunctional(CVodeMem cv_mem)
{
  int m;
  realtype del, delS, Del, Delp, dcon;
  int retval, is;
  booleantype do_sensi_sim;
  N_Vector wrk1, wrk2;

  /* Are we computing sensitivities with the CV_SIMULTANEOUS approach? */
  do_sensi_sim = (sensi && (ism==CV_SIMULTANEOUS));

  /* Initialize counter and evaluate f at predicted y */
  crate = ONE;
  m = 0;

  /* Initialize delS and Delp to avoid compiler warning message */
  delS = Delp = ZERO;

  retval = f(tn, zn[0], tempv, f_data);
  nfe++;
  if (retval < 0) return(CV_RHSFUNC_FAIL);
  if (retval > 0) return(RHSFUNC_RECVR);

  if (do_sensi_sim) {
    wrk1 = ftemp;
    wrk2 = ftempS[0];
    retval = CVSensRhs(cv_mem, tn, zn[0], tempv, znS[0], tempvS, wrk1, wrk2);
    if (retval < 0) return(CV_SRHSFUNC_FAIL);
    if (retval > 0) return(SRHSFUNC_RECVR);
  }

  /* Initialize correction to zero */

  N_VConst(ZERO, acor);
  if (do_sensi_sim) {
    for (is=0; is<Ns; is++)
      N_VConst(ZERO,acorS[is]);
  }

  /* Loop until convergence; accumulate corrections in acor */

  loop {

    nni++;

    /* Correct y directly from the last f value */

    N_VLinearSum(h, tempv, -ONE, zn[1], tempv);
    N_VScale(rl1, tempv, tempv);
    N_VLinearSum(ONE, zn[0], ONE, tempv, y);

    if (do_sensi_sim)
      for (is=0; is<Ns; is++) {
        N_VLinearSum(h, tempvS[is], -ONE, znS[1][is], tempvS[is]);
        N_VScale(rl1, tempvS[is], tempvS[is]);
        N_VLinearSum(ONE, znS[0][is], ONE, tempvS[is], yS[is]);
      }
    
    /* Get WRMS norm of current correction to use in convergence test */

    N_VLinearSum(ONE, tempv, -ONE, acor, acor);
    if (do_sensi_sim)
      for (is=0; is<Ns; is++)
        N_VLinearSum(ONE, tempvS[is], -ONE, acorS[is], acorS[is]);

    del = N_VWrmsNorm(acor, ewt);
    if (do_sensi_sim)
      delS = CVSensUpdateNorm(cv_mem, del, acorS, ewtS);

    N_VScale(ONE, tempv, acor);
    if (do_sensi_sim) 
      for (is=0; is<Ns; is++)
        N_VScale(ONE, tempvS[is], acorS[is]);
    
    /* Test for convergence.  If m > 0, an estimate of the convergence
       rate constant is stored in crate, and used in the test. 

       Recall that, even when errconS=FALSE, all variables are used in the
       convergence test. Hence, we use Del (and not del). However, acnrm
       is used in the error test and thus it has different forms
       depending on errconS (and this explains why we have to carry around
       del and delS)
    */
    
    Del = (do_sensi_sim) ? delS : del;
    if (m > 0) crate = MAX(CRDOWN * crate, Del / Delp);
    dcon = Del * MIN(ONE, crate) / tq[4];

    if (dcon <= ONE) {
      if (m == 0)
        if (do_sensi_sim && errconS) acnrm = delS;
        else                         acnrm = del;
      else {
        acnrm = N_VWrmsNorm(acor, ewt);
        if (do_sensi_sim && errconS)
          acnrm = CVSensUpdateNorm(cv_mem, acnrm, acorS, ewtS);
      }
      return(CV_SUCCESS);  /* Convergence achieved */
    }

    /* Stop at maxcor iterations or if iter. seems to be diverging */

    m++;
    if ((m==maxcor) || ((m >= 2) && (Del > RDIV * Delp))) return(CONV_FAIL);

    /* Save norm of correction, evaluate f, and loop again */

    Delp = Del;

    retval = f(tn, y, tempv, f_data);
    nfe++;
    if (retval < 0) return(CV_RHSFUNC_FAIL);
    if (retval > 0) return(RHSFUNC_RECVR);

    if (do_sensi_sim) {
      wrk1 = ftemp;
      wrk2 = ftempS[0];
      retval = CVSensRhs(cv_mem, tn, y, tempv, yS, tempvS, wrk1, wrk2);
      if (retval < 0) return(CV_SRHSFUNC_FAIL);
      if (retval > 0) return(SRHSFUNC_RECVR);
    }      

  } /* end loop */

}

/*
 * CVNlsNewton
 *
 * This routine handles the Newton iteration. It calls lsetup if 
 * indicated, calls CVNewtonIteration to perform the iteration, and 
 * retries a failed attempt at Newton iteration if that is indicated.
 * See return values at top of this file.
 *
 * This routine also handles the Newton iteration of the combined 
 * system when sensitivities are computed using the CV_SIMULTANEOUS
 * approach. Since in that case we use a quasi-Newton on the 
 * combined system (by approximating the Jacobian matrix by its
 * block diagonal) and thus only solve linear systems with 
 * multiple right hand sides (all sharing the same coefficient
 * matrix - whatever iteration matrix we decide on) we set-up
 * the linear solver to handle N equations at a time.
 *
 * Possible return values:
 *
 *   CV_SUCCESS       ---> continue with error test
 *
 *   CV_RHSFUNC_FAIL  -+  
 *   CV_LSETUP_FAIL    |
 *   CV_LSOLVE_FAIL    |-> halt the integration 
 *   CV_SRHSFUNC_FAIL -+
 *
 *   CONV_FAIL        -+
 *   RHSFUNC_RECVR     |-> predict again or stop if too many
 *   SRHSFUNC_RECVR   -+
 *
 */

static int CVNlsNewton(CVodeMem cv_mem, int nflag)
{
  N_Vector vtemp1, vtemp2, vtemp3, wrk1, wrk2;
  int convfail, ier;
  booleantype callSetup, do_sensi_sim;
  int retval, is;
  
  /* Are we computing sensitivities with the CV_SIMULTANEOUS approach? */
  do_sensi_sim = (sensi && (ism==CV_SIMULTANEOUS));

  vtemp1 = acor;  /* rename acor as vtemp1 for readability  */
  vtemp2 = y;     /* rename y as vtemp2 for readability     */
  vtemp3 = tempv; /* rename tempv as vtemp3 for readability */
  
  /* Set flag convfail, input to lsetup for its evaluation decision */
  convfail = ((nflag == FIRST_CALL) || (nflag == PREV_ERR_FAIL)) ?
    CV_NO_FAILURES : CV_FAIL_OTHER;

  /* Decide whether or not to call setup routine (if one exists) */
  if (setupNonNull) {      
    callSetup = (nflag == PREV_CONV_FAIL) || (nflag == PREV_ERR_FAIL) ||
      (nst == 0) || (nst >= nstlp + MSBP) || (ABS(gamrat-ONE) > DGMAX);

    /* Decide whether to force a call to setup */
    if (forceSetup) {
      callSetup = TRUE;
      convfail = CV_FAIL_OTHER;
    }

  } else {  
    crate = ONE;
    crateS = ONE;  /* if NO lsetup all conv. rates are set to ONE */
    callSetup = FALSE;
  }
  
  /* Looping point for the solution of the nonlinear system.
     Evaluate f at the predicted y, call lsetup if indicated, and
     call CVNewtonIteration for the Newton iteration itself.      */

  loop {

    retval = f(tn, zn[0], ftemp, f_data);
    nfe++; 
    if (retval < 0) return(CV_RHSFUNC_FAIL);
    if (retval > 0) return(RHSFUNC_RECVR);

    if (do_sensi_sim) {
      wrk1 = tempv;
      wrk2 = tempvS[0];
      retval = CVSensRhs(cv_mem, tn, zn[0], ftemp, znS[0], ftempS, wrk1, wrk2);
      if (retval < 0) return(CV_SRHSFUNC_FAIL);
      if (retval > 0) return(SRHSFUNC_RECVR);
    }

    if (callSetup) {
      ier = lsetup(cv_mem, convfail, zn[0], ftemp, &jcur, 
                   vtemp1, vtemp2, vtemp3);
      nsetups++;
      callSetup = FALSE;
      forceSetup = FALSE;
      gamrat = ONE; 
      gammap = gamma;
      crate = ONE;
      crateS = ONE; /* after lsetup all conv. rates are reset to ONE */
      nstlp = nst;
      /* Return if lsetup failed */
      if (ier < 0) return(CV_LSETUP_FAIL);
      if (ier > 0) return(CONV_FAIL);
    }

    /* Set acor to zero and load prediction into y vector */
    N_VConst(ZERO, acor);
    N_VScale(ONE, zn[0], y);

    if (do_sensi_sim)
      for (is=0; is<Ns; is++) {
        N_VConst(ZERO, acorS[is]);
        N_VScale(ONE, znS[0][is], yS[is]);
      }

    /* Do the Newton iteration */
    ier = CVNewtonIteration(cv_mem);

    /* If there is a convergence failure and the Jacobian-related 
       data appears not to be current, loop again with a call to lsetup
       in which convfail=CV_FAIL_BAD_J.  Otherwise return.                 */
    if (ier != TRY_AGAIN) return(ier);
    
    callSetup = TRUE;
    convfail = CV_FAIL_BAD_J;
  }
}

/*
 * CVNewtonIteration
 *
 * This routine performs the Newton iteration. If the iteration succeeds,
 * it returns the value CV_SUCCESS. If not, it may signal the CVNlsNewton 
 * routine to call lsetup again and reattempt the iteration, by
 * returning the value TRY_AGAIN. (In this case, CVNlsNewton must set 
 * convfail to CV_FAIL_BAD_J before calling setup again). 
 * Otherwise, this routine returns one of the appropriate values 
 * CV_LSOLVE_FAIL, CV_RHSFUNC_FAIL, CV_SRHSFUNC_FAIL, CONV_FAIL,
 * RHSFUNC_RECVR, or SRHSFUNC_RECVR back to CVNlsNewton.
 *
 * If sensitivities are computed using the CV_SIMULTANEOUS approach, this
 * routine performs a quasi-Newton on the combined nonlinear system.
 * The iteration matrix of the combined system is block diagonal with
 * each block being the iteration matrix of the original system. Thus
 * we solve linear systems with the same matrix but different right
 * hand sides.
 */

static int CVNewtonIteration(CVodeMem cv_mem)
{
  int m;
  realtype del, delS, Del, Delp, dcon;
  N_Vector b, *bS=NULL, wrk1, wrk2;
  booleantype do_sensi_sim;
  int retval, is;
  
  /* Are we computing sensitivities with the CV_SIMULTANEOUS approach? */
  do_sensi_sim = (sensi && (ism==CV_SIMULTANEOUS));

  mnewt = m = 0;

  /* Initialize delS and Delp to avoid compiler warning message */
  delS = Delp = ZERO;

  /* At this point, ftemp  <- f(t_n, y_n(0))
     ftempS <- fS(t_n, y_n(0), s_n(0))
     acor   <- 0
     acorS  <- 0
     y      <- y_n(0)
     yS     <- yS_n(0)                 */

  /* Looping point for Newton iteration */
  loop {
    
    /* Evaluate the residual of the nonlinear system */
    N_VLinearSum(rl1, zn[1], ONE, acor, tempv);
    N_VLinearSum(gamma, ftemp, -ONE, tempv, tempv);

    /* Call the lsolve function */
    b = tempv;

    retval = lsolve(cv_mem, b, ewt, y, ftemp); 
    nni++;

    if (retval < 0) return(CV_LSOLVE_FAIL);
    
    /* If lsolve had a recoverable failure and Jacobian data is
       not current, signal to try the solution again            */
    if (retval > 0) { 
      if ((!jcur) && (setupNonNull)) return(TRY_AGAIN);
      else                           return(CONV_FAIL);
    }

    /* Solve the sensitivity linear systems and do the same 
       tests on the return value of lsolve. */
 
    if (do_sensi_sim) {

      for (is=0; is<Ns; is++) {
        N_VLinearSum(rl1, znS[1][is], ONE, acorS[is], tempvS[is]);
        N_VLinearSum(gamma, ftempS[is], -ONE, tempvS[is], tempvS[is]);
      }
      bS = tempvS;
      for (is=0; is<Ns; is++) {
        retval = lsolve(cv_mem, bS[is], ewtS[is], y, ftemp);
        if (retval < 0) return(CV_LSOLVE_FAIL);
        if (retval > 0) { 
          if ((!jcur) && (setupNonNull)) return(TRY_AGAIN);
          else                           return(CONV_FAIL);
        }
      }
    }
    
    /* Get WRMS norm of correction; add correction to acor and y */

    del = N_VWrmsNorm(b, ewt);
    N_VLinearSum(ONE, acor, ONE, b, acor);
    N_VLinearSum(ONE, zn[0], ONE, acor, y);

    if (do_sensi_sim) {
      delS = CVSensUpdateNorm(cv_mem, del, bS, ewtS);
      for (is=0; is<Ns; is++) {
        N_VLinearSum(ONE, acorS[is], ONE, bS[is], acorS[is]);
        N_VLinearSum(ONE, znS[0][is], ONE, acorS[is], yS[is]);
      }
    }

    /* Test for convergence.  If m > 0, an estimate of the convergence
       rate constant is stored in crate, and used in the test.        */

    Del = (do_sensi_sim) ? delS : del;
    if (m > 0) crate = MAX(CRDOWN * crate, Del/Delp);
    dcon = Del * MIN(ONE, crate) / tq[4];
    
    if (dcon <= ONE) {
      if (m == 0)
        if (do_sensi_sim && errconS) acnrm = delS;
        else                         acnrm = del;
      else {
        acnrm = N_VWrmsNorm(acor, ewt);
        if (do_sensi_sim && errconS)
          acnrm = CVSensUpdateNorm(cv_mem, acnrm, acorS, ewtS);
      }
      jcur = FALSE;
      return(CV_SUCCESS);  /* Convergence achieved */
    }

    mnewt = ++m;
    
    /* Stop at maxcor iterations or if iter. seems to be diverging.
       If still not converged and Jacobian data is not current, 
       signal to try the solution again                            */
    if ((m == maxcor) || ((m >= 2) && (Del > RDIV * Delp))) {
      if ((!jcur) && (setupNonNull)) return(TRY_AGAIN);
      else                           return(CONV_FAIL);
    }
    
    /* Save norm of correction, evaluate f, and loop again */
    Delp = Del;
    retval = f(tn, y, ftemp, f_data);
    nfe++;
    if (retval < 0) return(CV_RHSFUNC_FAIL);
    if (retval > 0) {
      if ((!jcur) && (setupNonNull)) return(TRY_AGAIN);
      else                           return(RHSFUNC_RECVR);
    }

    if (do_sensi_sim) {
      wrk1 = tempv;
      wrk2 = tempvS[0];
      retval = CVSensRhs(cv_mem, tn, y, ftemp, yS, ftempS, wrk1, wrk2);
      if (retval < 0) return(CV_SRHSFUNC_FAIL);
      if (retval > 0) {
        if ((!jcur) && (setupNonNull)) return(TRY_AGAIN);
        else                           return(SRHSFUNC_RECVR);
      }
    }

  } /* end loop */

}

/*
 * CVQuadNls
 * 
 * This routine solves for the quadrature variables at the new step.
 * It does not solve a nonlinear system, but rather updates the
 * quadrature variables. The name for this function is just for 
 * uniformity purposes.
 *
 * Possible return values (interpreted by CVHandleNFlag)
 *
 *   CV_SUCCESS       -> continue with error test
 *   CV_QRHSFUNC_FAIL -> halt the integration 
 *   QRHSFUNC_RECVR   -> predict again or stop if too many
 *   
 */

static int CVQuadNls(CVodeMem cv_mem)
{
  int retval;

  /* Save quadrature correction in acorQ */
  retval = fQ(tn, y, acorQ, fQ_data);
  nfQe++;
  if (retval < 0) return(CV_QRHSFUNC_FAIL);
  if (retval > 0) return(QRHSFUNC_RECVR);

  N_VLinearSum(h, acorQ, -ONE, znQ[1], acorQ);
  N_VScale(rl1, acorQ, acorQ);

  /* Apply correction to quadrature variables */
  N_VLinearSum(ONE, znQ[0], ONE, acorQ, yQ);

  return(CV_SUCCESS);
}

/*
 * CVStgrNls
 *
 * This is a high-level routine that attempts to solve the 
 * sensitivity linear systems using nonlinear iterations (CV_FUNCTIONAL
 * or CV_NEWTON - depending on the value of iter) once the states y_n
 * were obtained and passed the error test.
 */

static int CVStgrNls(CVodeMem cv_mem)
{
  int flag=CV_SUCCESS;

  switch(iter) {
  case CV_FUNCTIONAL: 
    flag = CVStgrNlsFunctional(cv_mem);
    break;
  case CV_NEWTON:     
    flag = CVStgrNlsNewton(cv_mem);
    break;
  }

  return(flag);

}

/*
 * CVStgrNlsfunctional
 *
 * This routine attempts to solve the sensitivity linear systems 
 * using functional iteration (no matrices involved).
 *
 * Possible return values:
 *  CV_SUCCESS,
 *  CV_SRHSFUNC_FAIL,
 *  CONV_FAIL, SRHSFUNC_RECVR
 */

static int CVStgrNlsFunctional(CVodeMem cv_mem)
{
  int m;
  int retval, is;
  realtype Del, Delp, dcon;
  N_Vector wrk1, wrk2;

  /* Initialize estimated conv. rate and counter */
  crateS = ONE;
  m = 0;

  /* Initialize Delp to avoid compiler warning message */
  Delp = ZERO;

  /* Evaluate fS at predicted yS but with converged y (and corresponding f) */
  wrk1 = tempv;
  wrk2 = ftempS[0];
  retval = CVSensRhs(cv_mem, tn, y, ftemp, znS[0], tempvS, wrk1, wrk2);
  if (retval < 0) return(CV_SRHSFUNC_FAIL);
  if (retval > 0) return(SRHSFUNC_RECVR);

  /* Initialize correction to zero */
  for (is=0; is<Ns; is++)
    N_VConst(ZERO,acorS[is]);

  /* Loop until convergence; accumulate corrections in acorS */

  loop {
    
    nniS++;
    
    /* Correct yS from last fS value */
    for (is=0; is<Ns; is++) {
      N_VLinearSum(h, tempvS[is], -ONE, znS[1][is], tempvS[is]);
      N_VScale(rl1, tempvS[is], tempvS[is]);
      N_VLinearSum(ONE, znS[0][is], ONE, tempvS[is], yS[is]);
    }
    /* Get norm of current correction to use in convergence test */
    for (is=0; is<Ns; is++)
      N_VLinearSum(ONE, tempvS[is], -ONE, acorS[is], acorS[is]);
    Del = CVSensNorm(cv_mem, acorS, ewtS);
    for (is=0; is<Ns; is++)
      N_VScale(ONE, tempvS[is], acorS[is]);

    /* Test for convergence.  If m > 0, an estimate of the convergence
       rate constant is stored in crateS, and used in the test. 
       acnrmS contains the norm of the corrections (yS_n-yS_n(0)) and
       will be used in the error test (if errconS==TRUE)              */
    if (m > 0) crateS = MAX(CRDOWN * crateS, Del / Delp);
    dcon = Del * MIN(ONE, crateS) / tq[4];
    
    if (dcon <= ONE) {
      if (errconS)
        acnrmS = (m==0)? Del : CVSensNorm(cv_mem, acorS, ewtS);
      return(CV_SUCCESS);  /* Convergence achieved */
    }

    /* Stop at maxcor iterations or if iter. seems to be diverging */
    m++;
    if ((m==maxcorS) || ((m >= 2) && (Del > RDIV * Delp))) return(CONV_FAIL);

    /* Save norm of correction, evaluate f, and loop again */
    Delp = Del;

    wrk1 = tempv;
    wrk2 = ftempS[0];
    retval = CVSensRhs(cv_mem, tn, y, ftemp, yS, tempvS, wrk1, wrk2);
    if (retval < 0) return(CV_SRHSFUNC_FAIL);
    if (retval > 0) return(SRHSFUNC_RECVR);

  } /* end loop */
  
}

/*
 * CVStgrNlsNewton
 *
 * This routine attempts to solve the sensitivity linear systems using 
 * Newton iteration. It calls CVStgrNlsNewton to perform the actual 
 * iteration. If the Newton iteration fails with out-of-date Jacobian 
 * data (ier=TRY_AGAIN), it calls lsetup and retries the Newton iteration. 
 * This second try is unlikely to happen when using a Krylov linear solver.
 *
 * Possible return values:
 *
 *   CV_SUCCESS
 *
 *   CV_LSOLVE_FAIL   -+
 *   CV_LSETUP_FAIL    |
 *   CV_SRHSFUNC_FAIL -+
 *
 *   CONV_FAIL        -+
 *   SRHSFUNC_RECVR   -+
 */

static int CVStgrNlsNewton(CVodeMem cv_mem)
{
  int retval, is;
  int convfail, ier;
  N_Vector vtemp1, vtemp2, vtemp3, wrk1, wrk2;

  loop {

    /* Set acorS to zero and load prediction into yS vector */
    for (is=0; is<Ns; is++) {
      N_VConst(ZERO, acorS[is]);
      N_VScale(ONE, znS[0][is], yS[is]);
    }
 
    /* Evaluate fS at predicted yS but with converged y (and corresponding f) */
    wrk1 = tempv;
    wrk2 = tempvS[0];
    retval = CVSensRhs(cv_mem, tn, y, ftemp, yS, ftempS, wrk1, wrk2);
    if (retval < 0) return(CV_SRHSFUNC_FAIL);
    if (retval > 0) return(SRHSFUNC_RECVR);

    /* Do the Newton iteration */
    ier = CVStgrNewtonIteration(cv_mem);

    /* If the solve was successful (ier=CV_SUCCESS) or if an error 
       that cannot be fixed by a call to lsetup occured
       (ier = CV_LSOLVE_FAIL or CONV_FAIL) return */
    if (ier != TRY_AGAIN) return(ier);

    /* There was a convergence failure and the Jacobian-related data
       appears not to be current. Call lsetup with convfail=CV_FAIL_BAD_J
       and then loop again */
    convfail = CV_FAIL_BAD_J;

    /* Rename some vectors for readibility */
    vtemp1 = tempv;
    vtemp2 = yS[0];
    vtemp3 = ftempS[0];

    /* Call linear solver setup at converged y */
    ier = lsetup(cv_mem, convfail, y, ftemp, &jcur, 
                 vtemp1, vtemp2, vtemp3);
    nsetups++;
    nsetupsS++;
    gamrat = ONE;
    gammap = gamma;
    crate = ONE;
    crateS = ONE; /* after lsetup all conv. rates are reset to ONE */
    nstlp = nst;

    /* Return if lsetup failed */
    if (ier < 0) return(CV_LSETUP_FAIL);
    if (ier > 0) return(CONV_FAIL);

  } /* end loop */

}

/*
 * CVstgrNewtonIteration
 *
 * This routine performs the Newton iteration for all sensitivities. 
 * If the iteration succeeds, it returns the value CV_SUCCESS. 
 * If not, it may signal the CVStgrNlsNewton routine to call lsetup and 
 * reattempt the iteration, by returning the value TRY_AGAIN. (In this case, 
 * CVStgrNlsNewton must set convfail to CV_FAIL_BAD_J before calling setup again). 
 * Otherwise, this routine returns one of the appropriate values 
 * CV_LSOLVE_FAIL or CONV_FAIL back to CVStgrNlsNewton.
 */

static int CVStgrNewtonIteration(CVodeMem cv_mem)
{
  int m, retval;
  realtype Del, Delp, dcon;
  N_Vector *bS, wrk1, wrk2;
  int is;

  m = 0;

  /* Initialize Delp to avoid compiler warning message */
  Delp = ZERO;

  /* ftemp  <- f(t_n, y_n)
     y      <- y_n
     ftempS <- fS(t_n, y_n(0), s_n(0))
     acorS  <- 0
     yS     <- yS_n(0)                   */

  loop {

    /* Evaluate the residual of the nonlinear systems */
    for (is=0; is<Ns; is++) {
      N_VLinearSum(rl1, znS[1][is], ONE, acorS[is], tempvS[is]);
      N_VLinearSum(gamma, ftempS[is], -ONE, tempvS[is], tempvS[is]);
    }

    /* Call the lsolve function */
    bS = tempvS;
    nniS++;
    for (is=0; is<Ns; is++) {

      retval = lsolve(cv_mem, bS[is], ewtS[is], y, ftemp);

      /* Unrecoverable error in lsolve */
      if (retval < 0) return(CV_LSOLVE_FAIL);

      /* Recoverable error in lsolve and Jacobian data not current */
      if (retval > 0) { 
        if ((!jcur) && (setupNonNull)) return(TRY_AGAIN);
        else                           return(CONV_FAIL);
      }

    }
 
    /* Get norm of correction; add correction to acorS and yS */
    Del = CVSensNorm(cv_mem, bS, ewtS);
    for (is=0; is<Ns; is++) {
      N_VLinearSum(ONE, acorS[is], ONE, bS[is], acorS[is]);
      N_VLinearSum(ONE, znS[0][is], ONE, acorS[is], yS[is]);
    }

    /* Test for convergence.  If m > 0, an estimate of the convergence
       rate constant is stored in crateS, and used in the test.        */
    if (m > 0) crateS = MAX(CRDOWN * crateS, Del/Delp);
    dcon = Del * MIN(ONE, crateS) / tq[4];
    if (dcon <= ONE) {
      if (errconS)
        acnrmS = (m==0) ? Del : CVSensNorm(cv_mem, acorS, ewtS);
      jcur = FALSE;
      return(CV_SUCCESS);  /* Convergence achieved */
    }

    m++;

    /* Stop at maxcor iterations or if iter. seems to be diverging.
       If still not converged and Jacobian data is not current, 
       signal to try the solution again                            */
    if ((m == maxcorS) || ((m >= 2) && (Del > RDIV * Delp))) {
      if ((!jcur) && (setupNonNull)) return(TRY_AGAIN);
      else                           return(CONV_FAIL);
    }
    
    /* Save norm of correction, evaluate fS, and loop again */
    Delp = Del;
    
    wrk1 = tempv;
    wrk2 = tempvS[0];
    retval = CVSensRhs(cv_mem, tn, y, ftemp, yS, ftempS, wrk1, wrk2);
    if (retval < 0) return(CV_SRHSFUNC_FAIL);
    if (retval > 0) {
      if ((!jcur) && (setupNonNull)) return(TRY_AGAIN);
      else                           return(SRHSFUNC_RECVR);
    }

  } /* end loop */

}

/*
 * CVStgr1Nls
 *
 * This is a high-level routine that attempts to solve the i-th 
 * sensitivity linear system using nonlinear iterations (CV_FUNCTIONAL
 * or CV_NEWTON - depending on the value of iter) once the states y_n
 * were obtained and passed the error test.
 */

static int CVStgr1Nls(CVodeMem cv_mem, int is)
{
  int flag=CV_SUCCESS;

  switch(iter) {
  case CV_FUNCTIONAL: 
    flag = CVStgr1NlsFunctional(cv_mem,is);
    break;
  case CV_NEWTON:     
    flag = CVStgr1NlsNewton(cv_mem,is);
    break;
  }

  return(flag);

}

/*
 * CVStgr1NlsFunctional
 *
 * This routine attempts to solve the i-th sensitivity linear system
 * using functional iteration (no matrices involved).
 *
 * Possible return values:
 *   CV_SUCCESS,
 *   CV_SRHSFUNC_FAIL,
 *   CONV_FAIL, SRHSFUNC_RECVR
 */

static int CVStgr1NlsFunctional(CVodeMem cv_mem, int is)
{
  int retval, m;
  realtype Del, Delp, dcon;
  N_Vector wrk1, wrk2;

  /* Initialize estimated conv. rate and counter */
  crateS = ONE;
  m = 0;

  /* Initialize Delp to avoid compiler warning message */
  Delp = ZERO;

  /* Evaluate fS at predicted yS but with converged y (and corresponding f) */
  wrk1 = tempv;
  wrk2 = ftempS[0];
  retval = CVSensRhs1(cv_mem, tn, y, ftemp, is, znS[0][is], tempvS[is], wrk1, wrk2);
  if (retval < 0) return(CV_SRHSFUNC_FAIL);
  if (retval > 0) return(SRHSFUNC_RECVR);

  /* Initialize correction to zero */
  N_VConst(ZERO,acorS[is]);

  /* Loop until convergence; accumulate corrections in acorS */

  loop {

    nniS1[is]++;

    /* Correct yS from last fS value */
    N_VLinearSum(h, tempvS[is], -ONE, znS[1][is], tempvS[is]);
    N_VScale(rl1, tempvS[is], tempvS[is]);
    N_VLinearSum(ONE, znS[0][is], ONE, tempvS[is], yS[is]);

    /* Get WRMS norm of current correction to use in convergence test */
    N_VLinearSum(ONE, tempvS[is], -ONE, acorS[is], acorS[is]);
    Del = N_VWrmsNorm(acorS[is], ewtS[is]);
    N_VScale(ONE, tempvS[is], acorS[is]);

    /* Test for convergence.  If m > 0, an estimate of the convergence
       rate constant is stored in crateS, and used in the test. */

    if (m > 0) crateS = MAX(CRDOWN * crateS, Del / Delp);
    dcon = Del * MIN(ONE, crateS) / tq[4];

    if (dcon <= ONE) {
      return(CV_SUCCESS);  /* Convergence achieved */
    }

    /* Stop at maxcor iterations or if iter. seems to be diverging */
    m++;
    if ((m==maxcorS) || ((m >= 2) && (Del > RDIV * Delp))) return(CONV_FAIL);

    /* Save norm of correction, evaluate f, and loop again */
    Delp = Del;

    wrk1 = tempv;
    wrk2 = ftempS[0];

    retval = CVSensRhs1(cv_mem, tn, y, ftemp, is, yS[is], tempvS[is], wrk1, wrk2);
    if (retval < 0) return(CV_SRHSFUNC_FAIL);
    if (retval > 0) return(SRHSFUNC_RECVR);

  } /* end loop */
  
}

/*
 * CVStgr1NlsNewton
 *
 * This routine attempts to solve the i-th sensitivity linear system 
 * using Newton iteration. It calls CVStgr1NlsNewton to perform the 
 * actual iteration. If the Newton iteration fails with out-of-date 
 * Jacobian data (ier=TRY_AGAIN), it calls lsetup and retries the 
 * Newton iteration. This second try is unlikely to happen when 
 * using a Krylov linear solver.
 *
 * Possible return values:
 *
 *   CV_SUCCESS
 *
 *   CV_LSOLVE_FAIL
 *   CV_LSETUP_FAIL
 *   CV_SRHSFUNC_FAIL
 *
 *   CONV_FAIL
 *   SRHSFUNC_RECVR
 */

static int CVStgr1NlsNewton(CVodeMem cv_mem, int is)
{
  int convfail, retval, ier;
  N_Vector vtemp1, vtemp2, vtemp3, wrk1, wrk2;

  loop {

    /* Set acorS to zero and load prediction into yS vector */
    N_VConst(ZERO, acorS[is]);
    N_VScale(ONE, znS[0][is], yS[is]);
 
    /* Evaluate fS at predicted yS but with converged y (and corresponding f) */
    wrk1 = tempv;
    wrk2 = tempvS[0];
    retval = CVSensRhs1(cv_mem, tn, y, ftemp, is, yS[is], ftempS[is], wrk1, wrk2);
    if (retval < 0) return(CV_SRHSFUNC_FAIL);
    if (retval > 0) return(SRHSFUNC_RECVR);
  
    /* Do the Newton iteration */
    ier = CVStgr1NewtonIteration(cv_mem, is);

    /* If the solve was successful (ier=CV_SUCCESS) or if an error 
       that cannot be fixed by a call to lsetup occured
       (ier = CV_LSOLVE_FAIL or CONV_FAIL) return */
    if (ier != TRY_AGAIN) return(ier);

    /* There was a convergence failure and the Jacobian-related data
       appears not to be current. Call lsetup with convfail=CV_FAIL_BAD_J
       and then loop again */
    convfail = CV_FAIL_BAD_J;

    /* Rename some vectors for readibility */
    vtemp1 = tempv;
    vtemp2 = yS[0];
    vtemp3 = ftempS[0];

    /* Call linear solver setup at converged y */
    ier = lsetup(cv_mem, convfail, y, ftemp, &jcur, 
                 vtemp1, vtemp2, vtemp3);
    nsetups++;
    nsetupsS++;
    gamrat = ONE;
    crate = ONE;
    crateS = ONE; /* after lsetup all conv. rates are reset to ONE */
    gammap = gamma;
    nstlp = nst;

    /* Return if lsetup failed */
    if (ier < 0) return(CV_LSETUP_FAIL);
    if (ier > 0) return(CONV_FAIL);

  } /* end loop */

}

/*
 * CVStgr1NewtonIteration
 *
 * This routine performs the Newton iteration for the i-th sensitivity. 
 * If the iteration succeeds, it returns the value CV_SUCCESS. 
 * If not, it may signal the CVStgr1NlsNewton routine to call lsetup 
 * and reattempt the iteration, by returning the value TRY_AGAIN. 
 * (In this case, CVStgr1NlsNewton must set convfail to CV_FAIL_BAD_J 
 * before calling setup again). Otherwise, this routine returns one 
 * of the appropriate values CV_LSOLVE_FAIL or CONV_FAIL back to 
 * CVStgr1NlsNewton.
 */

static int CVStgr1NewtonIteration(CVodeMem cv_mem, int is)
{
  int m, retval;
  realtype Del, Delp, dcon;
  N_Vector *bS, wrk1, wrk2;

  m = 0;

  /* Initialize Delp to avoid compiler warning message */
  Delp = ZERO;

  /* ftemp      <- f(t_n, y_n)
     y          <- y_n
     ftempS[is] <- fS(is, t_n, y_n(0), s_n(0))
     acorS[is]  <- 0
     yS[is]     <- yS_n(0)[is]                 */

  loop {

    /* Evaluate the residual of the nonlinear systems */
    N_VLinearSum(rl1, znS[1][is], ONE, acorS[is], tempvS[is]);
    N_VLinearSum(gamma, ftempS[is], -ONE, tempvS[is], tempvS[is]);

    /* Call the lsolve function */
    bS = tempvS;

    nniS1[is]++;

    retval = lsolve(cv_mem, bS[is], ewtS[is], y, ftemp);

    /* Unrecoverable error in lsolve */
    if (retval < 0) return(CV_LSOLVE_FAIL);

    /* Recoverable error in lsolve and Jacobian data not current */
    if (retval > 0) { 
      if ((!jcur) && (setupNonNull)) return(TRY_AGAIN);
      else                           return(CONV_FAIL);
    }

    /* Get norm of correction; add correction to acorS and yS */
    Del = N_VWrmsNorm(bS[is], ewtS[is]);
    N_VLinearSum(ONE, acorS[is], ONE, bS[is], acorS[is]);
    N_VLinearSum(ONE, znS[0][is], ONE, acorS[is], yS[is]);

    /* Test for convergence.  If m > 0, an estimate of the convergence
       rate constant is stored in crateS, and used in the test.        */
    if (m > 0) crateS = MAX(CRDOWN * crateS, Del/Delp);
    dcon = Del * MIN(ONE, crateS) / tq[4];
    if (dcon <= ONE) {
      jcur = FALSE;
      return(CV_SUCCESS);  /* Convergence achieved */
    }

    m++;

    /* Stop at maxcor iterations or if iter. seems to be diverging.
       If still not converged and Jacobian data is not current, 
       signal to try the solution again                            */
    if ((m == maxcorS) || ((m >= 2) && (Del > RDIV * Delp))) {
      if ((!jcur) && (setupNonNull)) return(TRY_AGAIN);
      else                           return(CONV_FAIL);
    }

    /* Save norm of correction, evaluate fS, and loop again */
    Delp = Del;

    wrk1 = tempv;
    wrk2 = tempvS[0];
    retval = CVSensRhs1(cv_mem, tn, y, ftemp, is, yS[is], ftempS[is], wrk1, wrk2);
    if (retval < 0) return(CV_SRHSFUNC_FAIL);
    if (retval > 0) {
      if ((!jcur) && (setupNonNull)) return(TRY_AGAIN);
      else                           return(SRHSFUNC_RECVR);
    }

  } /* end loop */

}

/*
 * CVHandleNFlag
 *
 * This routine takes action on the return value nflag = *nflagPtr
 * returned by CVNls, as follows:
 *
 * If CVNls succeeded in solving the nonlinear system, then
 * CVHandleNFlag returns the constant DO_ERROR_TEST, which tells CVStep
 * to perform the error test.
 *
 * If the nonlinear system was not solved successfully, then ncfn and
 * ncf = *ncfPtr are incremented and Nordsieck array zn is restored.
 *
 * If the solution of the nonlinear system failed due to an
 * unrecoverable failure by setup, we return the value CV_LSETUP_FAIL.
 * 
 * If it failed due to an unrecoverable failure in solve, then we return
 * the value CV_LSOLVE_FAIL.
 *
 * If it failed due to an unrecoverable failure in rhs, then we return
 * the value CV_RHSFUNC_FAIL.
 *
 * If it failed due to an unrecoverable failure in quad rhs, then we return
 * the value CV_QRHSFUNC_FAIL.
 *
 * If it failed due to an unrecoverable failure in sensi rhs, then we return
 * the value CV_SRHSFUNC_FAIL.
 *
 * Otherwise, a recoverable failure occurred when solving the 
 * nonlinear system (CVNls returned nflag = CONV_FAIL, RHSFUNC_RECVR, or
 * SRHSFUNC_RECVR). 
 * In this case, if ncf is now equal to maxncf or |h| = hmin, 
 * we return the value CV_CONV_FAILURE (if nflag=CONV_FAIL), or
 * CV_REPTD_RHSFUNC_ERR (if nflag=RHSFUNC_RECVR), or CV_REPTD_SRHSFUNC_ERR
 * (if nflag=SRHSFUNC_RECVR).
 * If not, we set *nflagPtr = PREV_CONV_FAIL and return the value
 * PREDICT_AGAIN, telling CVStep to reattempt the step.
 *
 */

static int CVHandleNFlag(CVodeMem cv_mem, int *nflagPtr, realtype saved_t,
                         int *ncfPtr, long int *ncfnPtr)
{
  int nflag;
  
  nflag = *nflagPtr;
  
  if (nflag == CV_SUCCESS) return(DO_ERROR_TEST);

  /* The nonlinear soln. failed; increment ncfn and restore zn */
  (*ncfnPtr)++;
  CVRestore(cv_mem, saved_t);
  
  /* Return if lsetup, lsolve, or some rhs failed unrecoverably */
  if (nflag == CV_LSETUP_FAIL)   return(CV_LSETUP_FAIL);
  if (nflag == CV_LSOLVE_FAIL)   return(CV_LSOLVE_FAIL);
  if (nflag == CV_RHSFUNC_FAIL)  return(CV_RHSFUNC_FAIL);
  if (nflag == CV_QRHSFUNC_FAIL) return(CV_QRHSFUNC_FAIL);
  if (nflag == CV_SRHSFUNC_FAIL) return(CV_SRHSFUNC_FAIL);

  /* At this point, nflag = CONV_FAIL, RHSFUNC_RECVR, or SRHSFUNC_RECVR; 
     increment ncf */
  
  (*ncfPtr)++;
  etamax = ONE;

  /* If we had maxncf failures or |h| = hmin, 
     return CV_CONV_FAILURE, CV_REPTD_RHSFUNC_ERR, 
     CV_REPTD_QRHSFUNC_ERR, or CV_REPTD_SRHSFUNC_ERR */

  if ((ABS(h) <= hmin*ONEPSM) || (*ncfPtr == maxncf)) {
    if (nflag == CONV_FAIL)      return(CV_CONV_FAILURE);
    if (nflag == RHSFUNC_RECVR)  return(CV_REPTD_RHSFUNC_ERR);    
    if (nflag == QRHSFUNC_RECVR) return(CV_REPTD_QRHSFUNC_ERR);    
    if (nflag == SRHSFUNC_RECVR) return(CV_REPTD_SRHSFUNC_ERR);    
  }

  /* Reduce step size; return to reattempt the step */

  eta = MAX(ETACF, hmin / ABS(h));
  *nflagPtr = PREV_CONV_FAIL;
  CVRescale(cv_mem);

  return(PREDICT_AGAIN);
}

/*
 * CVRestore
 *
 * This routine restores the value of tn to saved_t and undoes the
 * prediction.  After execution of CVRestore, the Nordsieck array zn has
 * the same values as before the call to CVPredict.
 */

static void CVRestore(CVodeMem cv_mem, realtype saved_t)
{
  int j, k;
  int is;

  tn = saved_t;
  for (k = 1; k <= q; k++)
    for (j = q; j >= k; j--)
      N_VLinearSum(ONE, zn[j-1], -ONE, zn[j], zn[j-1]);

  if (quadr) {
    for (k = 1; k <= q; k++)
      for (j = q; j >= k; j--)
        N_VLinearSum(ONE, znQ[j-1], -ONE, znQ[j], znQ[j-1]);
  }

  if (sensi) {
    for (is=0; is<Ns; is++) {
      for (k = 1; k <= q; k++)
        for (j = q; j >= k; j--)
          N_VLinearSum(ONE, znS[j-1][is], -ONE, znS[j][is], znS[j-1][is]);
    }
  }
}

/* 
 * -----------------------------------------------------------------
 * Error Test
 * -----------------------------------------------------------------
 */

/*
 * CVDoErrorTest
 *
 * This routine performs the local error test, for the state, quadrature, 
 * or sensitivity variables. Its last three arguments change depending
 * on which variables the error test is to be performed on.
 * 
 * The weighted local error norm dsm is loaded into *dsmPtr, and 
 * the test dsm ?<= 1 is made.
 *
 * If the test passes, CVDoErrorTest returns CV_SUCCESS. 
 *
 * If the test fails, we undo the step just taken (call CVRestore) and 
 *
 *   - if maxnef error test failures have occurred or if ABS(h) = hmin,
 *     we return CV_ERR_FAILURE.
 *
 *   - if more than MXNEF1 error test failures have occurred, an order
 *     reduction is forced. If already at order 1, restart by reloading 
 *     zn from scratch (also znQ and znS if appropriate).
 *     If f() fails, we return CV_RHSFUNC_FAIL or CV_UNREC_RHSFUNC_ERR;
 *     if fQ() fails, we return CV_QRHSFUNC_FAIL or CV_UNREC_QRHSFUNC_ERR;
 *     if CVSensRhs() fails, we return CV_SRHSFUNC_FAIL or CV_UNREC_SRHSFUNC_ERR;
 *     (no recovery is possible at this stage).
 *
 *   - otherwise, set *nflagPtr to PREV_ERR_FAIL, and return TRY_AGAIN. 
 *
 */

static int CVDoErrorTest(CVodeMem cv_mem, int *nflagPtr, realtype saved_t, 
                         realtype acor_nrm,
                         int *nefPtr, long int *netfPtr, realtype *dsmPtr)
{
  realtype dsm;
  int retval, is;
  N_Vector wrk1, wrk2;

  dsm = acor_nrm / tq[2];

  /* If est. local error norm dsm passes test, return CV_SUCCESS */  
  *dsmPtr = dsm; 
  if (dsm <= ONE) return(CV_SUCCESS);
  
  /* Test failed; increment counters, set nflag, and restore zn array */
  (*nefPtr)++;
  (*netfPtr)++;
  *nflagPtr = PREV_ERR_FAIL;
  CVRestore(cv_mem, saved_t);

  /* At maxnef failures or |h| = hmin, return CV_ERR_FAILURE */
  if ((ABS(h) <= hmin*ONEPSM) || (*nefPtr == maxnef)) return(CV_ERR_FAILURE);

  /* Set etamax = 1 to prevent step size increase at end of this step */
  etamax = ONE;

  /* Set h ratio eta from dsm, rescale, and return for retry of step */
  if (*nefPtr <= MXNEF1) {
    eta = ONE / (RPowerR(BIAS2*dsm,ONE/L) + ADDON);
    eta = MAX(ETAMIN, MAX(eta, hmin / ABS(h)));
    if (*nefPtr >= SMALL_NEF) eta = MIN(eta, ETAMXF);
    CVRescale(cv_mem);
    return(TRY_AGAIN);
  }
  
  /* After MXNEF1 failures, force an order reduction and retry step */
  if (q > 1) {
    eta = MAX(ETAMIN, hmin / ABS(h));
    CVAdjustOrder(cv_mem,-1);
    L = q;
    q--;
    qwait = L;
    CVRescale(cv_mem);
    return(TRY_AGAIN);
  }

  /* If already at order 1, restart: reload zn, znQ, znS from scratch */
  eta = MAX(ETAMIN, hmin / ABS(h));
  h *= eta;
  next_h = h;
  hscale = h;
  qwait = LONG_WAIT;
  nscon = 0;

  retval = f(tn, zn[0], tempv, f_data);
  nfe++;
  if (retval < 0) return(CV_RHSFUNC_FAIL);
  if (retval > 0) return(CV_UNREC_RHSFUNC_ERR);

  N_VScale(h, tempv, zn[1]);

  if (quadr) {

    retval = fQ(tn, zn[0], tempvQ, fQ_data);
    nfQe++;
    if (retval < 0) return(CV_QRHSFUNC_FAIL);
    if (retval > 0) return(CV_UNREC_QRHSFUNC_ERR);

    N_VScale(h, tempvQ, znQ[1]);

  }

  if (sensi) {

    wrk1 = ftemp;
    wrk2 = ftempS[0];

    retval = CVSensRhs(cv_mem, tn, zn[0], tempv, znS[0], tempvS, wrk1, wrk2);
    if (retval < 0) return(CV_SRHSFUNC_FAIL);
    if (retval > 0) return(CV_UNREC_SRHSFUNC_ERR);

    for (is=0; is<Ns; is++) 
      N_VScale(h, tempvS[is], znS[1][is]);

  }
  
  return(TRY_AGAIN);
}

/* 
 * -----------------------------------------------------------------
 * Functions called after a successful step
 * -----------------------------------------------------------------
 */

/*
 * CVCompleteStep
 *
 * This routine performs various update operations when the solution
 * to the nonlinear system has passed the local error test. 
 * We increment the step counter nst, record the values hu and qu,
 * update the tau array, and apply the corrections to the zn array.
 * The tau[i] are the last q values of h, with tau[1] the most recent.
 * The counter qwait is decremented, and if qwait == 1 (and q < qmax)
 * we save acor and tq[5] for a possible order increase.
 */

static void CVCompleteStep(CVodeMem cv_mem)
{
  int i, j;
  int is;
  
  nst++;
  nscon++;
  hu = h;
  qu = q;

  for (i=q; i >= 2; i--)  tau[i] = tau[i-1];
  if ((q==1) && (nst > 1)) tau[2] = tau[1];
  tau[1] = h;

  /* Apply correction to column j of zn: l_j * Delta_n */

  for (j=0; j <= q; j++) 
    N_VLinearSum(l[j], acor, ONE, zn[j], zn[j]);

  if (quadr) {
    for (j=0; j <= q; j++) 
      N_VLinearSum(l[j], acorQ, ONE, znQ[j], znQ[j]);
  }

  if (sensi) {
    for (is=0; is<Ns; is++)
      for (j=0; j <= q; j++) 
        N_VLinearSum(l[j], acorS[is], ONE, znS[j][is], znS[j][is]);
  }

  /* If necessary, store Delta_n in zn[qmax] to be used in order increase
  
  This actually will be Delta_{n-1} in the ELTE at q+1 since it happens at
  the next to last step of order q before a possible one at order q+1
  */

  qwait--;
  if ((qwait == 1) && (q != qmax)) {
    
    N_VScale(ONE, acor, zn[qmax]);
    
    if (quadr)
      N_VScale(ONE, acorQ, znQ[qmax]);

    if (sensi)
      for (is=0; is<Ns; is++)
        N_VScale(ONE, acorS[is], znS[qmax][is]);
    
    saved_tq5 = tq[5];
    indx_acor = qmax;
  }

}

/*
 * CVprepareNextStep
 *
 * This routine handles the setting of stepsize and order for the
 * next step -- hprime and qprime.  Along with hprime, it sets the
 * ratio eta = hprime/h.  It also updates other state variables 
 * related to a change of step size or order. 
 */

static void CVPrepareNextStep(CVodeMem cv_mem, realtype dsm)
{
  /* If etamax = 1, defer step size or order changes */
  if (etamax == ONE) {
    qwait = MAX(qwait, 2);
    qprime = q;
    hprime = h;
    eta = ONE;
    return;
  }

  /* etaq is the ratio of new to old h at the current order */  
  etaq = ONE /(RPowerR(BIAS2*dsm,ONE/L) + ADDON);
  
  /* If no order change, adjust eta and acor in CVSetEta and return */
  if (qwait != 0) {
    eta = etaq;
    qprime = q;
    CVSetEta(cv_mem);
    return;
  }
  
  /* If qwait = 0, consider an order change.   etaqm1 and etaqp1 are 
     the ratios of new to old h at orders q-1 and q+1, respectively.
     CVChooseEta selects the largest; CVSetEta adjusts eta and acor */
  qwait = 2;
  etaqm1 = CVComputeEtaqm1(cv_mem);
  etaqp1 = CVComputeEtaqp1(cv_mem);  
  CVChooseEta(cv_mem); 
  CVSetEta(cv_mem);
}

/*
 * CVsetEta
 *
 * This routine adjusts the value of eta according to the various
 * heuristic limits and the optional input hmax.  It also resets
 * etamax to be the estimated local error vector.
 */

static void CVSetEta(CVodeMem cv_mem)
{

  /* If eta below the threshhold THRESH, reject a change of step size */
  if (eta < THRESH) {
    eta = ONE;
    hprime = h;
  } else {
    /* Limit eta by etamax and hmax, then set hprime */
    eta = MIN(eta, etamax);
    eta /= MAX(ONE, ABS(h)*hmax_inv*eta);
    hprime = h * eta;
    if (qprime < q) nscon = 0;
  }

  /* Reset etamx for the next step size change, and scale acor */
}

/*
 * CVComputeEtaqm1
 *
 * This routine computes and returns the value of etaqm1 for a
 * possible decrease in order by 1.
 */

static realtype CVComputeEtaqm1(CVodeMem cv_mem)
{
  realtype ddn;
  
  etaqm1 = ZERO;

  if (q > 1) {

    ddn = N_VWrmsNorm(zn[q], ewt);

    if ( quadr && errconQ) {
      ddn = CVQuadUpdateNorm(cv_mem, ddn, znQ[q], ewtQ);
    }

    if ( sensi && errconS ) {
      ddn = CVSensUpdateNorm(cv_mem, ddn, znS[q], ewtS);
    }

    ddn = ddn/tq[1];

    etaqm1 = ONE/(RPowerR(BIAS1*ddn, ONE/q) + ADDON);

  }

  return(etaqm1);
}

/*
 * CVComputeEtaqp1
 *
 * This routine computes and returns the value of etaqp1 for a
 * possible increase in order by 1.
 */

static realtype CVComputeEtaqp1(CVodeMem cv_mem)
{
  realtype dup, cquot;
  int is;
  
  etaqp1 = ZERO;

  if (q != qmax) {

    cquot = (tq[5] / saved_tq5) * RPowerI(h/tau[2], L);

    N_VLinearSum(-cquot, zn[qmax], ONE, acor, tempv);

    dup = N_VWrmsNorm(tempv, ewt);

    if ( quadr && errconQ ) {
      N_VLinearSum(-cquot, znQ[qmax], ONE, acorQ, tempvQ);
      dup = CVQuadUpdateNorm(cv_mem, dup, tempvQ, ewtQ);
    }

    if ( sensi && errconS ) {
      for (is=0; is<Ns; is++) 
        N_VLinearSum(-cquot, znS[qmax][is], ONE, acorS[is], tempvS[is]);
      dup = CVSensUpdateNorm(cv_mem, dup, tempvS, ewtS);
    }

    dup = dup / tq[3];

    etaqp1 = ONE / (RPowerR(BIAS3*dup, ONE/(L+1)) + ADDON);

  }

  return(etaqp1);
}

/*
 * CVChooseEta
 * Given etaqm1, etaq, etaqp1 (the values of eta for qprime =
 * q - 1, q, or q + 1, respectively), this routine chooses the 
 * maximum eta value, sets eta to that value, and sets qprime to the
 * corresponding value of q.  If there is a tie, the preference
 * order is to (1) keep the same order, then (2) decrease the order,
 * and finally (3) increase the order.  If the maximum eta value
 * is below the threshhold THRESH, the order is kept unchanged and
 * eta is set to 1.
 */

static void CVChooseEta(CVodeMem cv_mem)
{
  realtype etam;
  int is;
  
  etam = MAX(etaqm1, MAX(etaq, etaqp1));
  
  if (etam < THRESH) {
    eta = ONE;
    qprime = q;
    return;
  }

  if (etam == etaq) {

    eta = etaq;
    qprime = q;

  } else if (etam == etaqm1) {

    eta = etaqm1;
    qprime = q - 1;

  } else {

    eta = etaqp1;
    qprime = q + 1;

    if (lmm == CV_BDF) {

      /* 
       * Store Delta_n in zn[qmax] to be used in order increase 
       *
       * This happens at the last step of order q before an increase
       * to order q+1, so it represents Delta_n in the ELTE at q+1
       */
      
      N_VScale(ONE, acor, zn[qmax]);
      
      if (quadr && errconQ)
        N_VScale(ONE, acorQ, znQ[qmax]);
      
      if (sensi && errconS)
        for (is=0; is<Ns; is++)
          N_VScale(ONE, acorS[is], znS[qmax][is]);
    }

  }

}

/* 
 * -----------------------------------------------------------------
 * Function to handle failures
 * -----------------------------------------------------------------
 */

/*
 * CVHandleFailure
 *
 * This routine prints error messages for all cases of failure by
 * CVHin or CVStep. 
 * It returns to CVode the value that CVode is to return to the user.
 */

static int CVHandleFailure(CVodeMem cv_mem, int flag)
{

  /* Set vector of  absolute weighted local errors */
  /*
  N_VProd(acor, ewt, tempv);
  N_VAbs(tempv, tempv);
  */  

  /* Depending on flag, print error message and return error flag */
  switch (flag) {
  case CV_ERR_FAILURE:  
    CVProcessError(cv_mem, CV_ERR_FAILURE, "CVODES", "CVode", MSGCV_ERR_FAILS, tn, h);
    break;
  case CV_CONV_FAILURE: 
    CVProcessError(cv_mem, CV_CONV_FAILURE, "CVODES", "CVode", MSGCV_CONV_FAILS, tn, h);
    break;
  case CV_LSETUP_FAIL:  
    CVProcessError(cv_mem, CV_LSETUP_FAIL, "CVODES", "CVode", MSGCV_SETUP_FAILED, tn);
    break;
  case CV_LSOLVE_FAIL:  
    CVProcessError(cv_mem, CV_LSOLVE_FAIL, "CVODES", "CVode", MSGCV_SOLVE_FAILED, tn);
    break;
  case CV_RHSFUNC_FAIL:
    CVProcessError(cv_mem, CV_RHSFUNC_FAIL, "CVODES", "CVode", MSGCV_RHSFUNC_FAILED, tn);
    break;
  case CV_UNREC_RHSFUNC_ERR:
    CVProcessError(cv_mem, CV_UNREC_RHSFUNC_ERR, "CVODES", "CVode", MSGCV_RHSFUNC_UNREC, tn);
    break;
  case CV_REPTD_RHSFUNC_ERR:
    CVProcessError(cv_mem, CV_REPTD_RHSFUNC_ERR, "CVODES", "CVode", MSGCV_RHSFUNC_REPTD, tn);
    break;
  case CV_RTFUNC_FAIL:    
    CVProcessError(cv_mem, CV_RTFUNC_FAIL, "CVODES", "CVode", MSGCV_RTFUNC_FAILED, tn);
    break;
  case CV_QRHSFUNC_FAIL:
    CVProcessError(cv_mem, CV_QRHSFUNC_FAIL, "CVODES", "CVode", MSGCV_QRHSFUNC_FAILED, tn);
    break;
  case CV_UNREC_QRHSFUNC_ERR:
    CVProcessError(cv_mem, CV_UNREC_QRHSFUNC_ERR, "CVODES", "CVode", MSGCV_QRHSFUNC_UNREC, tn);
    break;
  case CV_REPTD_QRHSFUNC_ERR:
    CVProcessError(cv_mem, CV_REPTD_QRHSFUNC_ERR, "CVODES", "CVode", MSGCV_QRHSFUNC_REPTD, tn);
    break;
  case CV_SRHSFUNC_FAIL:
    CVProcessError(cv_mem, CV_SRHSFUNC_FAIL, "CVODES", "CVode", MSGCV_SRHSFUNC_FAILED, tn);
    break;
  case CV_UNREC_SRHSFUNC_ERR:
    CVProcessError(cv_mem, CV_UNREC_SRHSFUNC_ERR, "CVODES", "CVode", MSGCV_SRHSFUNC_UNREC, tn);
    break;
  case CV_REPTD_SRHSFUNC_ERR:
    CVProcessError(cv_mem, CV_REPTD_SRHSFUNC_ERR, "CVODES", "CVode", MSGCV_SRHSFUNC_REPTD, tn);
    break;
  case CV_TOO_CLOSE:
    CVProcessError(cv_mem, CV_TOO_CLOSE, "CVODES", "CVode", MSGCV_TOO_CLOSE);
  default:
    return(CV_SUCCESS);
  }

  return(flag);

}

/* 
 * -----------------------------------------------------------------
 * Functions for BDF Stability Limit Detection   
 * -----------------------------------------------------------------
 */

/*
 * CVBDFStab
 *
 * This routine handles the BDF Stability Limit Detection Algorithm
 * STALD.  It is called if lmm = CV_BDF and the SLDET option is on.
 * If the order is 3 or more, the required norm data is saved.
 * If a decision to reduce order has not already been made, and
 * enough data has been saved, CVsldet is called.  If it signals
 * a stability limit violation, the order is reduced, and the step
 * size is reset accordingly.
 */

static void CVBDFStab(CVodeMem cv_mem)
{
  int i,k, ldflag, factorial;
  realtype sq, sqm1, sqm2;
      
  /* If order is 3 or greater, then save scaled derivative data,
     push old data down in i, then add current values to top.    */

  if (q >= 3) {
    for (k = 1; k <= 3; k++)
      for (i = 5; i >= 2; i--) 
        ssdat[i][k] = ssdat[i-1][k]; 
    factorial = 1;
    for (i = 1; i <= q-1; i++) factorial *= i;
    sq = factorial*q*(q+1)*acnrm/tq[5];
    sqm1 = factorial*q*N_VWrmsNorm(zn[q], ewt);
    sqm2 = factorial*N_VWrmsNorm(zn[q-1], ewt);
    ssdat[1][1] = sqm2*sqm2;
    ssdat[1][2] = sqm1*sqm1;
    ssdat[1][3] = sq*sq;
  }  

  if (qprime >= q) {

    /* If order is 3 or greater, and enough ssdat has been saved,
       nscon >= q+5, then call stability limit detection routine.  */

    if ( (q >= 3) && (nscon >= q+5) ) {
      ldflag = CVsldet(cv_mem);
      if (ldflag > 3) {
        /* A stability limit violation is indicated by
           a return flag of 4, 5, or 6.
           Reduce new order.                     */
        qprime = q-1;
        eta = etaqm1; 
        eta = MIN(eta,etamax);
        eta = eta/MAX(ONE,ABS(h)*hmax_inv*eta);
        hprime = h*eta;
        nor = nor + 1;
      }
    }
  }
  else {
    /* Otherwise, let order increase happen, and 
       reset stability limit counter, nscon.     */
    nscon = 0;
  }
}

/*
 * CVsldet
 *
 * This routine detects stability limitation using stored scaled 
 * derivatives data. CVsldet returns the magnitude of the
 * dominate characteristic root, rr. The presents of a stability
 * limit is indicated by rr > "something a little less then 1.0",  
 * and a positive kflag. This routine should only be called if
 * order is greater than or equal to 3, and data has been collected
 * for 5 time steps. 
 * 
 * Returned values:
 *    kflag = 1 -> Found stable characteristic root, normal matrix case
 *    kflag = 2 -> Found stable characteristic root, quartic solution
 *    kflag = 3 -> Found stable characteristic root, quartic solution,
 *                 with Newton correction
 *    kflag = 4 -> Found stability violation, normal matrix case
 *    kflag = 5 -> Found stability violation, quartic solution
 *    kflag = 6 -> Found stability violation, quartic solution,
 *                 with Newton correction
 *
 *    kflag < 0 -> No stability limitation, 
 *                 or could not compute limitation.
 *
 *    kflag = -1 -> Min/max ratio of ssdat too small.
 *    kflag = -2 -> For normal matrix case, vmax > vrrt2*vrrt2
 *    kflag = -3 -> For normal matrix case, The three ratios
 *                  are inconsistent.
 *    kflag = -4 -> Small coefficient prevents elimination of quartics.  
 *    kflag = -5 -> R value from quartics not consistent.
 *    kflag = -6 -> No corrected root passes test on qk values
 *    kflag = -7 -> Trouble solving for sigsq.
 *    kflag = -8 -> Trouble solving for B, or R via B.
 *    kflag = -9 -> R via sigsq[k] disagrees with R from data.
 */

static int CVsldet(CVodeMem cv_mem)
{
  int i, k, j, it, kmin=0, kflag=0;
  realtype rat[5][4], rav[4], qkr[4], sigsq[4], smax[4], ssmax[4];
  realtype drr[4], rrc[4],sqmx[4], qjk[4][4], vrat[5], qc[6][4], qco[6][4];
  realtype rr, rrcut, vrrtol, vrrt2, sqtol, rrtol;
  realtype smink, smaxk, sumrat, sumrsq, vmin, vmax, drrmax, adrr;
  realtype tem, sqmax, saqk, qp, s, sqmaxk, saqj, sqmin;
  realtype rsa, rsb, rsc, rsd, rd1a, rd1b, rd1c;
  realtype rd2a, rd2b, rd3a, cest1, corr1; 
  realtype ratp, ratm, qfac1, qfac2, bb, rrb;

  /* The following are cutoffs and tolerances used by this routine */

  rrcut  = RCONST(0.98);
  vrrtol = RCONST(1.0e-4);
  vrrt2  = RCONST(5.0e-4);
  sqtol  = RCONST(1.0e-3);
  rrtol  = RCONST(1.0e-2);

  rr = ZERO;

  /*  Index k corresponds to the degree of the interpolating polynomial. */
  /*      k = 1 -> q-1          */
  /*      k = 2 -> q            */
  /*      k = 3 -> q+1          */

  /*  Index i is a backward-in-time index, i = 1 -> current time, */
  /*      i = 2 -> previous step, etc    */

  /* get maxima, minima, and variances, and form quartic coefficients  */

  for (k=1; k<=3; k++) {
    smink = ssdat[1][k];
    smaxk = ZERO;
    
    for (i=1; i<=5; i++) {
      smink = MIN(smink,ssdat[i][k]);
      smaxk = MAX(smaxk,ssdat[i][k]);
    }
    
    if (smink < TINY*smaxk) {
      kflag = -1;  
      return(kflag);
    }
    smax[k] = smaxk;
    ssmax[k] = smaxk*smaxk;
    
    sumrat = ZERO;
    sumrsq = ZERO;
    for (i=1; i<=4; i++) {
      rat[i][k] = ssdat[i][k]/ssdat[i+1][k];
      sumrat = sumrat + rat[i][k];
      sumrsq = sumrsq + rat[i][k]*rat[i][k];
    } 
    rav[k] = FOURTH*sumrat;
    vrat[k] = ABS(FOURTH*sumrsq - rav[k]*rav[k]);
      
    qc[5][k] = ssdat[1][k]*ssdat[3][k] - ssdat[2][k]*ssdat[2][k];
    qc[4][k] = ssdat[2][k]*ssdat[3][k] - ssdat[1][k]*ssdat[4][k];
    qc[3][k] = ZERO;
    qc[2][k] = ssdat[2][k]*ssdat[5][k] - ssdat[3][k]*ssdat[4][k];
    qc[1][k] = ssdat[4][k]*ssdat[4][k] - ssdat[3][k]*ssdat[5][k];
    
    for (i=1; i<=5; i++) {
      qco[i][k] = qc[i][k];
    }
  }                            /* End of k loop */
  
  /* Isolate normal or nearly-normal matrix case. Three quartic will
     have common or nearly-common roots in this case. 
     Return a kflag = 1 if this procedure works. If three root 
     differ more than vrrt2, return error kflag = -3.    */
  
  vmin = MIN(vrat[1],MIN(vrat[2],vrat[3]));
  vmax = MAX(vrat[1],MAX(vrat[2],vrat[3]));
  
  if (vmin < vrrtol*vrrtol) {

    if (vmax > vrrt2*vrrt2) {
      kflag = -2;  
      return(kflag);
    } else {
      rr = (rav[1] + rav[2] + rav[3])/THREE;
      drrmax = ZERO;
      for (k = 1;k<=3;k++) {
        adrr = ABS(rav[k] - rr);
        drrmax = MAX(drrmax, adrr);
      }
      if (drrmax > vrrt2)
        kflag = -3;    
      kflag = 1;
      /*  can compute charactistic root, drop to next section   */
    }

  } else {
      
    /* use the quartics to get rr. */
      
    if (ABS(qco[1][1]) < TINY*ssmax[1]) {
      kflag = -4;    
      return(kflag);
    }
      
    tem = qco[1][2]/qco[1][1];
    for (i=2; i<=5; i++) {
      qco[i][2] = qco[i][2] - tem*qco[i][1];
    }
      
    qco[1][2] = ZERO;
    tem = qco[1][3]/qco[1][1];
    for (i=2; i<=5; i++) {
      qco[i][3] = qco[i][3] - tem*qco[i][1];
    }
    qco[1][3] = ZERO;
      
    if (ABS(qco[2][2]) < TINY*ssmax[2]) {
      kflag = -4;    
      return(kflag);
    }
      
    tem = qco[2][3]/qco[2][2];
    for (i=3; i<=5; i++) {
      qco[i][3] = qco[i][3] - tem*qco[i][2];
    }
      
    if (ABS(qco[4][3]) < TINY*ssmax[3]) {
      kflag = -4;    
      return(kflag);
    }
      
    rr = -qco[5][3]/qco[4][3];
      
    if (rr < TINY || rr > HUN) {
      kflag = -5;   
      return(kflag);
    }
      
    for (k=1; k<=3; k++) {
      qkr[k] = qc[5][k] + rr*(qc[4][k] + rr*rr*(qc[2][k] + rr*qc[1][k]));
    }  
      
    sqmax = ZERO;
    for (k=1; k<=3; k++) {
      saqk = ABS(qkr[k])/ssmax[k];
      if (saqk > sqmax) sqmax = saqk;
    } 
      
    if (sqmax < sqtol) {
      kflag = 2;
        
      /*  can compute charactistic root, drop to "given rr,etc"   */
        
    } else {
          
      /* do Newton corrections to improve rr.  */
        
      for (it=1; it<=3; it++) {
        for (k=1; k<=3; k++) {
          qp = qc[4][k] + rr*rr*(THREE*qc[2][k] + rr*FOUR*qc[1][k]);
          drr[k] = ZERO;
          if (ABS(qp) > TINY*ssmax[k]) drr[k] = -qkr[k]/qp;
          rrc[k] = rr + drr[k];
        } 
          
        for (k=1; k<=3; k++) {
          s = rrc[k];
          sqmaxk = ZERO;
          for (j=1; j<=3; j++) {
            qjk[j][k] = qc[5][j] + s*(qc[4][j] + 
                                      s*s*(qc[2][j] + s*qc[1][j]));
            saqj = ABS(qjk[j][k])/ssmax[j];
            if (saqj > sqmaxk) sqmaxk = saqj;
          } 
          sqmx[k] = sqmaxk;
        } 
              
        sqmin = sqmx[1] + ONE;
        for (k=1; k<=3; k++) {
          if (sqmx[k] < sqmin) {
            kmin = k;
            sqmin = sqmx[k];
          }
        } 
        rr = rrc[kmin];
              
        if (sqmin < sqtol) {
          kflag = 3;
          /*  can compute charactistic root   */
          /*  break out of Newton correction loop and drop to "given rr,etc" */ 
          break;
        } else {
          for (j=1; j<=3; j++) {
            qkr[j] = qjk[j][kmin];
          }
        }     
      } /*  end of Newton correction loop  */ 
          
      if (sqmin > sqtol) {
        kflag = -6;
        return(kflag);
      }
    } /*  end of if (sqmax < sqtol) else   */
  } /*  end of if (vmin < vrrtol*vrrtol) else, quartics to get rr. */
  
  /* given rr, find sigsq[k] and verify rr.  */
  /* All positive kflag drop to this section  */
  
  for (k=1; k<=3; k++) {
    rsa = ssdat[1][k];
    rsb = ssdat[2][k]*rr;
    rsc = ssdat[3][k]*rr*rr;
    rsd = ssdat[4][k]*rr*rr*rr;
    rd1a = rsa - rsb;
    rd1b = rsb - rsc;
    rd1c = rsc - rsd;
    rd2a = rd1a - rd1b;
    rd2b = rd1b - rd1c;
    rd3a = rd2a - rd2b;
    
    if (ABS(rd1b) < TINY*smax[k]) {
      kflag = -7;
      return(kflag);
    }
    
    cest1 = -rd3a/rd1b;
    if (cest1 < TINY || cest1 > FOUR) {
      kflag = -7;
      return(kflag);
    }
    corr1 = (rd2b/cest1)/(rr*rr);
    sigsq[k] = ssdat[3][k] + corr1;
  }
  
  if (sigsq[2] < TINY) {
    kflag = -8;
    return(kflag);
  }
  
  ratp = sigsq[3]/sigsq[2];
  ratm = sigsq[1]/sigsq[2];
  qfac1 = FOURTH*(q*q - ONE);
  qfac2 = TWO/(q - ONE);
  bb = ratp*ratm - ONE - qfac1*ratp;
  tem = ONE - qfac2*bb;
  
  if (ABS(tem) < TINY) {
    kflag = -8;
    return(kflag);
  }
  
  rrb = ONE/tem;
  
  if (ABS(rrb - rr) > rrtol) {
    kflag = -9;
    return(kflag);
  }
  
  /* Check to see if rr is above cutoff rrcut  */
  if (rr > rrcut) {
    if (kflag == 1) kflag = 4;
    if (kflag == 2) kflag = 5;
    if (kflag == 3) kflag = 6;
  }
  
  /* All positive kflag returned at this point  */
  
  return(kflag);
  
}

/* 
 * -----------------------------------------------------------------
 * Functions for rootfinding
 * -----------------------------------------------------------------
 */

/* 
 * CVRcheck1
 *
 * This routine completes the initialization of rootfinding memory
 * information, and checks whether g has a zero both at and very near
 * the initial point of the IVP.
 *
 * This routine returns an int equal to:
 *  INITROOT = -1 if a close pair of zeros was found, and
 *  CV_SUCCESS     =  0 otherwise.
 */

static int CVRcheck1(CVodeMem cv_mem)
{
  int i, retval;
  realtype smallh, hratio;
  booleantype zroot;

  for (i = 0; i < nrtfn; i++) iroots[i] = 0;
  tlo = tn;
  ttol = (ABS(tn) + ABS(h))*uround*HUN;

  /* Evaluate g at initial t and check for zero values. */
  retval = gfun(tlo, zn[0], glo, g_data);
  nge = 1;
  if (retval != 0) return(CV_RTFUNC_FAIL);
  
  zroot = FALSE;
  for (i = 0; i < nrtfn; i++) {
    if (ABS(glo[i]) == ZERO) zroot = TRUE;
  }
  if (!zroot) return(CV_SUCCESS);

  /* Some g_i is zero at t0; look at g at t0+(small increment). */
  hratio = MAX(ttol/ABS(h), TENTH);
  smallh = hratio*h;
  tlo += smallh;
  N_VLinearSum(ONE, zn[0], hratio, zn[1], y);
  retval = gfun(tlo, y, glo, g_data);
  nge++;
  if (retval != 0) return(CV_RTFUNC_FAIL);

  zroot = FALSE;
  for (i = 0; i < nrtfn; i++) {
    if (ABS(glo[i]) == ZERO) {
      zroot = TRUE;
      iroots[i] = 1;
    }
  }
  if (zroot) return(INITROOT);
  return(CV_SUCCESS);

}

/*
 * CVRcheck2
 *
 * This routine checks for exact zeros of g at the last root found,
 * if the last return was a root.  It then checks for a close
 * pair of zeros (an error condition), and for a new root at a
 * nearby point.  The left endpoint (tlo) of the search interval
 * is adjusted if necessary to assure that all g_i are nonzero
 * there, before returning to do a root search in the interval.
 *
 * On entry, tlo = tretlast is the last value of tret returned by
 * CVode.  This may be the previous tn, the previous tout value, or
 * the last root location.
 *
 * This routine returns an int equal to:
 *      CLOSERT = -2 if a close pair of zeros was found,
 *      RTFOUND =  1 if a new zero of g was found near tlo, or
 *      CV_SUCCESS    =  0 otherwise.
 */

static int CVRcheck2(CVodeMem cv_mem)
{
  int i, retval;
  realtype smallh, hratio;
  booleantype zroot;

  if (irfnd == 0) return(CV_SUCCESS);

  (void) CVodeGetDky(cv_mem, tlo, 0, y);
  retval = gfun(tlo, y, glo, g_data);
  nge++;
  if (retval != 0) return(CV_RTFUNC_FAIL);

  zroot = FALSE;
  for (i = 0; i < nrtfn; i++) iroots[i] = 0;
  for (i = 0; i < nrtfn; i++) {
    if (ABS(glo[i]) == ZERO) {
      zroot = TRUE;
      iroots[i] = 1;
    }
  }
  if (!zroot) return(CV_SUCCESS);

  /* One or more g_i has a zero at tlo.  Check g at tlo+smallh. */
  ttol = (ABS(tn) + ABS(h))*uround*HUN;
  smallh = (h > ZERO) ? ttol : -ttol;
  tlo += smallh;
  if ( (tlo - tn)*h >= ZERO) {
    hratio = smallh/h;
    N_VLinearSum(ONE, y, hratio, zn[1], y);
  } else {
    (void) CVodeGetDky(cv_mem, tlo, 0, y);
  }
  retval = gfun(tlo, y, glo, g_data);
  nge++;
  if (retval != 0) return(CV_RTFUNC_FAIL);

  zroot = FALSE;
  for (i = 0; i < nrtfn; i++) {
    if (ABS(glo[i]) == ZERO) {
      if (iroots[i] == 1) return(CLOSERT);
      zroot = TRUE;
      iroots[i] = 1;
    }
  }
  if (zroot) return(RTFOUND);
  return(CV_SUCCESS);

}

/*
 * CVRcheck3
 *
 * This routine interfaces to CVRootfind to look for a root of g
 * between tlo and either tn or tout, whichever comes first.
 * Only roots beyond tlo in the direction of integration are sought.
 *
 * This routine returns an int equal to:
 *      RTFOUND =  1 if a root of g was found, or
 *      CV_SUCCESS    =  0 otherwise.
 */

static int CVRcheck3(CVodeMem cv_mem)
{
  int i, retval, ier;

  /* Set thi = tn or tout, whichever comes first; set y = y(thi). */
  if (taskc == CV_ONE_STEP) {
    thi = tn;
    N_VScale(ONE, zn[0], y);
  }
  if (taskc == CV_NORMAL) {
    if ( (toutc - tn)*h >= ZERO) {
      thi = tn; 
      N_VScale(ONE, zn[0], y);
    } else {
      thi = toutc;
      (void) CVodeGetDky(cv_mem, thi, 0, y);
    }
  }

  /* Set ghi = g(thi) and call CVRootfind to search (tlo,thi) for roots. */
  retval = gfun(thi, y, ghi, g_data);
  nge++;
  if (retval != 0) return(CV_RTFUNC_FAIL);

  ttol = (ABS(tn) + ABS(h))*uround*HUN;
  ier = CVRootfind(cv_mem);
  tlo = trout;
  for (i = 0; i < nrtfn; i++) glo[i] = grout[i];

  /* If no root found, return CV_SUCCESS. */  
  if (ier == CV_SUCCESS) return(CV_SUCCESS);

  /* If a root was found, interpolate to get y(trout) and return.  */
  (void) CVodeGetDky(cv_mem, trout, 0, y);
  return(RTFOUND);

}

/*
 * CVRootFind
 *
 * This routine solves for a root of g(t) between tlo and thi, if
 * one exists.  Only roots of odd multiplicity (i.e. with a change
 * of sign in one of the g_i), or exact zeros, are found.
 * Here the sign of tlo - thi is arbitrary, but if multiple roots
 * are found, the one closest to tlo is returned.
 *
 * The method used is the Illinois algorithm, a modified secant method.
 * Reference: Kathie L. Hiebert and Lawrence F. Shampine, Implicitly
 * Defined Output Points for Solutions of ODEs, Sandia National
 * Laboratory Report SAND80-0180, February 1980.
 *
 * This routine uses the following parameters for communication:
 *
 * nrtfn    = number of functions g_i, or number of components of
 *            the vector-valued function g(t).  Input only.
 *
 * gfun     = user-defined function for g(t).  Its form is
 *            (void) gfun(t, y, gt, g_data)
 *
 * nge      = cumulative counter for gfun calls.
 *
 * ttol     = a convergence tolerance for trout.  Input only.
 *            When a root at trout is found, it is located only to
 *            within a tolerance of ttol.  Typically, ttol should
 *            be set to a value on the order of
 *               100 * UROUND * max (ABS(tlo), ABS(thi))
 *            where UROUND is the unit roundoff of the machine.
 *
 * tlo, thi = endpoints of the interval in which roots are sought.
 *            On input, and must be distinct, but tlo - thi may
 *            be of either sign.  The direction of integration is
 *            assumed to be from tlo to thi.  On return, tlo and thi
 *            are the endpoints of the final relevant interval.
 *
 * glo, ghi = arrays of length nrtfn containing the vectors g(tlo)
 *            and g(thi) respectively.  Input and output.  On input,
 *            none of the glo[i] should be zero.
 *
 * trout    = root location, if a root was found, or thi if not.
 *            Output only.  If a root was found other than an exact
 *            zero of g, trout is the endpoint thi of the final
 *            interval bracketing the root, with size at most ttol.
 *
 * grout    = array of length nrtfn containing g(trout) on return.
 *
 * iroots   = int array of length nrtfn with root information.
 *            Output only.  If a root was found, iroots indicates
 *            which components g_i have a root at trout.  For
 *            i = 0, ..., nrtfn-1, iroots[i] = 1 if g_i has a root
 *            and iroots[i] = 0 otherwise.
 *
 * This routine returns an int equal to:
 *      RTFOUND =  1 if a root of g was found, or
 *      CV_SUCCESS    =  0 otherwise.
 */

static int CVRootfind(CVodeMem cv_mem)
{
  realtype alpha, tmid, gfrac, maxfrac, fracint, fracsub;
  int i, retval, imax, side, sideprev;
  booleantype zroot, sgnchg;

  imax = 0;

  /* First check for change in sign in ghi or for a zero in ghi. */
  maxfrac = ZERO;
  zroot = FALSE;
  sgnchg = FALSE;
  for (i = 0;  i < nrtfn; i++) {
    if (ABS(ghi[i]) == ZERO) {
      zroot = TRUE;
    } else {
      if (glo[i]*ghi[i] < ZERO) {
        gfrac = ABS(ghi[i]/(ghi[i] - glo[i]));
        if (gfrac > maxfrac) {
          sgnchg = TRUE;
          maxfrac = gfrac;
          imax = i;
        }
      }
    }
  }

  /* If no sign change was found, reset trout and grout.  Then return
     CV_SUCCESS if no zero was found, or set iroots and return RTFOUND.  */ 
  if (!sgnchg) {
    trout = thi;
    for (i = 0; i < nrtfn; i++) grout[i] = ghi[i];
    if (!zroot) return(CV_SUCCESS);
    for (i = 0; i < nrtfn; i++) {
      iroots[i] = 0;
      if (ABS(ghi[i]) == ZERO) iroots[i] = 1;
    }
    return(RTFOUND);
  }

  /* Initialize alpha to avoid compiler warning */

  alpha = ONE;

  /* A sign change was found.  Loop to locate nearest root. */

  side = 0;  sideprev = -1;
  loop {                                    /* Looping point */

    /* Set weight alpha.
       On the first two passes, set alpha = 1.  Thereafter, reset alpha
       according to the side (low vs high) of the subinterval in which
       the sign change was found in the previous two passes.
       If the sides were opposite, set alpha = 1.
       If the sides were the same, then double alpha (if high side),
       or halve alpha (if low side).
       The next guess tmid is the secant method value if alpha = 1, but
       is closer to tlo if alpha < 1, and closer to thi if alpha > 1.    */

    if (sideprev == side) {
      alpha = (side == 2) ? alpha*TWO : alpha*HALF;
    } else {
      alpha = ONE;
    }

    /* Set next root approximation tmid and get g(tmid).
       If tmid is too close to tlo or thi, adjust it inward,
       by a fractional distance that is between 0.1 and 0.5.  */
    tmid = thi - (thi - tlo)*ghi[imax]/(ghi[imax] - alpha*glo[imax]);
    if (ABS(tmid - tlo) < HALF*ttol) {
      fracint = ABS(thi - tlo)/ttol;
      fracsub = (fracint > FIVE) ? TENTH : HALF/fracint;
      tmid = tlo + fracsub*(thi - tlo);
    }
    if (ABS(thi - tmid) < HALF*ttol) {
      fracint = ABS(thi - tlo)/ttol;
      fracsub = (fracint > FIVE) ? TENTH : HALF/fracint;
      tmid = thi - fracsub*(thi - tlo);
    }

    (void) CVodeGetDky(cv_mem, tmid, 0, y);
    retval = gfun(tmid, y, grout, g_data);
    nge++;
    if (retval != 0) return(CV_RTFUNC_FAIL);

    /* Check to see in which subinterval g changes sign, and reset imax.
       Set side = 1 if sign change is on low side, or 2 if on high side.  */  
    maxfrac = ZERO;
    zroot = FALSE;
    sgnchg = FALSE;
    sideprev = side;
    for (i = 0;  i < nrtfn; i++) {
      if (ABS(grout[i]) == ZERO) {
        zroot = TRUE;
      } else {
        if (glo[i]*grout[i] < ZERO) {
          gfrac = ABS(grout[i]/(grout[i] - glo[i]));
          if (gfrac > maxfrac) {
            sgnchg = TRUE;
            maxfrac = gfrac;
            imax = i;
          }
        }
      }
    }
    if (sgnchg) {
      /* Sign change found in (tlo,tmid); replace thi with tmid. */
      thi = tmid;
      for (i = 0; i < nrtfn; i++) ghi[i] = grout[i];
      side = 1;
      /* Stop at root thi if converged; otherwise loop. */
      if (ABS(thi - tlo) <= ttol) break;
      continue;  /* Return to looping point. */
    }

    if (zroot) {
      /* No sign change in (tlo,tmid), but g = 0 at tmid; return root tmid. */
      thi = tmid;
      for (i = 0; i < nrtfn; i++) ghi[i] = grout[i];
      break;
    }

    /* No sign change in (tlo,tmid), and no zero at tmid.
       Sign change must be in (tmid,thi).  Replace tlo with tmid. */
    tlo = tmid;
    for (i = 0; i < nrtfn; i++) glo[i] = grout[i];
    side = 2;
    /* Stop at root thi if converged; otherwise loop back. */
    if (ABS(thi - tlo) <= ttol) break;

  } /* End of root-search loop */

  /* Reset trout and grout, set iroots, and return RTFOUND. */
  trout = thi;
  for (i = 0; i < nrtfn; i++) {
    grout[i] = ghi[i];
    iroots[i] = 0;
    if (ABS(ghi[i]) == ZERO) iroots[i] = 1;
    if (glo[i]*ghi[i] < ZERO) iroots[i] = 1;
  }
  return(RTFOUND);
}

/* 
 * -----------------------------------------------------------------
 * Functions for combined norms
 * -----------------------------------------------------------------
 */

/*
 * CVQuadUpdateNorm
 *
 * Updates the norm old_nrm to account for all quadratures.
 */

static realtype CVQuadUpdateNorm(CVodeMem cv_mem, realtype old_nrm,
                                 N_Vector xQ, N_Vector wQ)
{
  realtype qnrm;

  qnrm = N_VWrmsNorm(xQ, wQ);
  if (old_nrm > qnrm) return(old_nrm);
  else                return(qnrm);
}

/*
 * CVQuadUpdateDsm
 *
 *  Usage    : dms = CVQuadUpdateDsm(cv_mem, dsm, dsmQ);
 *
 * This routine updates the local error norm dsm with quadrature
 * related information. Used only if quadratures are computed
 * with FULL error control.
 *
 * Returns the maximum over the wheighted local error norms.
 */

static realtype CVQuadUpdateDsm(CVodeMem cv_mem, realtype old_dsm, 
                                realtype dsmQ)
{
  if ( old_dsm > dsmQ ) return(old_dsm);
  else                  return(dsmQ);
}

/*
 * CVSensNorm
 *
 * This routine returns the maximum over the weighted root mean 
 * square norm of xS with weight vectors wS:
 *
 *  max { wrms(xS[0],wS[0]) ... wrms(xS[Ns-1],wS[Ns-1]) }    
 *
 * Called by CVSensUpdateNorm or directly in the CV_STAGGERED approach 
 * during the NLS solution and before the error test.
 */

static realtype CVSensNorm(CVodeMem cv_mem, N_Vector *xS, N_Vector *wS)
{
  int is;
  realtype nrm, snrm;

  nrm = N_VWrmsNorm(xS[0],wS[0]);
  for (is=1; is<Ns; is++) {
    snrm = N_VWrmsNorm(xS[is],wS[is]);
    if ( snrm > nrm ) nrm = snrm;
  }

  return(nrm);

}

/*
 * CVSensUpdateNorm
 *
 * Updates the norm old_nrm to account for all sensitivities.
 */

static realtype CVSensUpdateNorm(CVodeMem cv_mem, realtype old_nrm,
                                 N_Vector *xS, N_Vector *wS)
{
  realtype snrm;
  
  snrm = CVSensNorm(cv_mem, xS, wS);
  if (old_nrm > snrm) return(old_nrm);
  else                return(snrm);
}

/*
 * CVStgrUpdateNorm
 *
 *  Usage    : dms = CVStgrUpdateDsm(cv_mem, old_dsm, dsmS);
 *
 * This routine updates the local error norm old_dsm with sensitivity 
 * related information. Used only in the CV_STAGGERED or CV_STAGGERED1 
 * approach with FULL error control.This value is consistent with 
 * the one computed in CVDoErrorTest when ism=CV_SIMULTANEOUS and 
 * errconS=TRUE.
 *
 * Returns the maximum over the wheighted local error norms.
 */

static realtype CVStgrUpdateDsm(CVodeMem cv_mem, realtype old_dsm, 
                                realtype dsmS)
{
  if ( old_dsm > dsmS ) return(old_dsm);
  else                  return(dsmS);
}

/* 
 * -----------------------------------------------------------------
 * Wrappers for sensitivity RHS
 * -----------------------------------------------------------------
 */

/*
 * CVSensRhs
 *
 * CVSensRhs is a high level routine that returns right hand side 
 * of sensitivity equations. Depending on the 'ifS' flag, it either 
 * calls directly the fS routine (ifS=CV_ALLSENS) or (if ifS=CV_ONESENS) 
 * calls the fS1 routine in a loop over all sensitivities.
 *
 * CVSensRhs is called:
 *  (*) by CVode at the first step
 *  (*) by CVYddNorm if errcon=TRUE
 *  (*) by CVNlsFunctional, CVNlsNewton, and CVNewtonIteration
 *      if ism=CV_SIMULTANEOUS
 *  (*) by CVDoErrorTest when restarting from scratch
 *  (*) in the corrector loop if ism=CV_STAGGERED
 *  (*) by CVStgrDoErrorTest when restarting from scratch 
 *
 * The return value is that of the sensitivity RHS function fS,
 *
 */

static int CVSensRhs(CVodeMem cv_mem, realtype time, 
                     N_Vector ycur, N_Vector fcur, 
                     N_Vector *yScur, N_Vector *fScur,
                     N_Vector temp1, N_Vector temp2)
{
  int retval=0, is;

  if (ifS==CV_ALLSENS) {
    retval = fS(Ns, time, ycur, fcur, yScur, fScur, 
                fS_data, temp1, temp2);
    nfSe++;
  } else {
    for (is=0; is<Ns; is++) {
      retval = fS1(Ns, time, ycur, fcur, is, yScur[is], fScur[is], 
                   fS_data, temp1, temp2);
      nfSe++;
      if (retval != 0) break;
    }
  }

  return(retval);
}

/*
 * CVSensRhs1
 *
 * CVSensRhs1 is a high level routine that returns right hand side 
 * of the is-th sensitivity equation. 
 *
 * CVSensRhs1 is called only during the CV_STAGGERED1 corrector loop
 * (ifS must be CV_ONESENS, otherwise CVodeSensMalloc would have 
 * issued an error message).
 *
 * The return value is that of the sensitivity RHS function fS1,
 */

static int CVSensRhs1(CVodeMem cv_mem, realtype time, 
                      N_Vector ycur, N_Vector fcur, 
                      int is, N_Vector yScur, N_Vector fScur,
                      N_Vector temp1, N_Vector temp2)
{
  int retval;

  retval = fS1(Ns, time, ycur, fcur, is, yScur, fScur, 
               fS_data, temp1, temp2);
  nfSe++;

  return(retval);
}

/* 
 * -----------------------------------------------------------------
 * Internal DQ approximations for sensitivity RHS
 * -----------------------------------------------------------------
 */

/* Undefine Readibility Constants */

#undef Ns
#undef y
#undef yS
#undef fS_data
#undef ftemp

/*
 * CVSensRhsDQ   - internal CVSensRhsFn
 *
 * CVSensRhsDQ computes right hand side of all sensitivity equations
 * by finite differences
 */

int CVSensRhsDQ(int Ns, realtype t, 
                N_Vector y, N_Vector ydot, 
                N_Vector *yS, N_Vector *ySdot, 
                void *fS_data,  
                N_Vector ytemp, N_Vector ftemp)
{
  int is, retval;
  
  for (is=0; is<Ns; is++) {
    retval = CVSensRhs1DQ(Ns, t, y, ydot, is, yS[is], ySdot[is],
                          fS_data, ytemp, ftemp);
    if (retval!=0) return(retval);
  }

  return(0);
}

/*
 * CVSensRhs1DQ   - internal CVSensRhs1Fn
 *
 * CVSensRhs1DQ computes the right hand side of the is-th sensitivity 
 * equation by finite differences
 *
 * CVSensRhs1DQ returns 0 if successful. Otherwise it returns the 
 * non-zero return value from f().
 */

int CVSensRhs1DQ(int Ns, realtype t, 
                 N_Vector y, N_Vector ydot, 
                 int is, N_Vector yS, N_Vector ySdot, 
                 void *fS_data,
                 N_Vector ytemp, N_Vector ftemp)
{
  CVodeMem cv_mem;
  int retval, method;
  int nfel = 0, which;
  realtype psave, pbari;
  realtype delta , rdelta;
  realtype Deltap, rDeltap, r2Deltap;
  realtype Deltay, rDeltay, r2Deltay;
  realtype Delta , rDelta , r2Delta ;
  realtype norms, ratio;
  
  /* fS_data points to cv_mem */
  cv_mem = (CVodeMem) fS_data;

  delta = RSqrt(MAX(reltol, uround));
  rdelta = ONE/delta;
  
  pbari = pbar[is];

  which = plist[is];

  psave = p[which];
  
  Deltap  = pbari * delta;
  rDeltap = ONE/Deltap;
  norms   = N_VWrmsNorm(yS, ewt) * pbari;
  rDeltay = MAX(norms, rdelta) / pbari;
  Deltay  = ONE/rDeltay;
  
  if (DQrhomax == ZERO) {
    /* No switching */
    method = (DQtype==CV_CENTERED) ? CENTERED1 : FORWARD1;
  } else {
    /* switch between simultaneous/separate DQ */
    ratio = Deltay * rDeltap;
    if ( MAX(ONE/ratio, ratio) <= DQrhomax ) 
      method = (DQtype==CV_CENTERED) ? CENTERED1 : FORWARD1;
    else
      method = (DQtype==CV_CENTERED) ? CENTERED2 : FORWARD2;
  }

  switch(method) {
    
  case CENTERED1:
    
    Delta = MIN(Deltay, Deltap);
    r2Delta = HALF/Delta;
    
    N_VLinearSum(ONE,y,Delta,yS,ytemp);
    p[which] = psave + Delta;

    retval = f(t, ytemp, ySdot, f_data);
    nfel++;
    if (retval != 0) return(retval);
    
    N_VLinearSum(ONE,y,-Delta,yS,ytemp);
    p[which] = psave - Delta;

    retval = f(t, ytemp, ftemp, f_data);
    nfel++;
    if (retval != 0) return(retval);

    N_VLinearSum(r2Delta,ySdot,-r2Delta,ftemp,ySdot);
    
    break;
    
  case CENTERED2:
    
    r2Deltap = HALF/Deltap;
    r2Deltay = HALF/Deltay;
    
    N_VLinearSum(ONE,y,Deltay,yS,ytemp);

    retval = f(t, ytemp, ySdot, f_data);
    nfel++;
    if (retval != 0) return(retval);

    N_VLinearSum(ONE,y,-Deltay,yS,ytemp);

    retval = f(t, ytemp, ftemp, f_data);
    nfel++;
    if (retval != 0) return(retval);

    N_VLinearSum(r2Deltay, ySdot, -r2Deltay, ftemp, ySdot);
    
    p[which] = psave + Deltap;
    retval = f(t, y, ytemp, f_data);
    nfel++;
    if (retval != 0) return(retval);

    p[which] = psave - Deltap;
    retval = f(t, y, ftemp, f_data);
    nfel++;
    if (retval != 0) return(retval);

    N_VLinearSum(r2Deltap,ytemp,-r2Deltap,ftemp,ftemp);
    
    N_VLinearSum(ONE,ySdot,ONE,ftemp,ySdot);
    
    break;
    
  case FORWARD1:
    
    Delta = MIN(Deltay, Deltap);
    rDelta = ONE/Delta;
    
    N_VLinearSum(ONE,y,Delta,yS,ytemp);
    p[which] = psave + Delta;

    retval = f(t, ytemp, ySdot, f_data);
    nfel++;
    if (retval != 0) return(retval);
    
    N_VLinearSum(rDelta,ySdot,-rDelta,ydot,ySdot);
    
    break;
    
  case FORWARD2:
    
    N_VLinearSum(ONE,y,Deltay,yS,ytemp);

    retval = f(t, ytemp, ySdot, f_data);
    nfel++;
    if (retval != 0) return(retval);

    N_VLinearSum(rDeltay, ySdot, -rDeltay, ydot, ySdot);
    
    p[which] = psave + Deltap;
    retval = f(t, y, ytemp, f_data);
    nfel++;
    if (retval != 0) return(retval);

    N_VLinearSum(rDeltap,ytemp,-rDeltap,ydot,ftemp);
      
    N_VLinearSum(ONE,ySdot,ONE,ftemp,ySdot);
    
    break;
    
  }
  
  p[which] = psave;
  
  /* Increment counter nfeS */
  nfeS += nfel;
  
  return(0);
}

/* 
 * -----------------------------------------------------------------
 * Error message handling functions
 * -----------------------------------------------------------------
 */

/* 
 * CVProcessError is a high level error handling function
 * - if cv_mem==NULL it prints the error message to stderr
 * - otherwise, it sets-up and calls the error hadling function 
 *   pointed to by cv_ehfun
 */

#define ehfun    (cv_mem->cv_ehfun)
#define eh_data  (cv_mem->cv_eh_data)

void CVProcessError(CVodeMem cv_mem, 
                    int error_code, const char *module, const char *fname, 
                    const char *msgfmt, ...)
{
  va_list ap;
  char msg[256];

  /* Initialize the argument pointer variable 
     (msgfmt is the last required argument to CVProcessError) */

  va_start(ap, msgfmt);

  if (cv_mem == NULL) {    /* We write to stderr */

#ifndef NO_FPRINTF_OUTPUT
    fprintf(stderr, "\n[%s ERROR]  %s\n  ", module, fname);
    fprintf(stderr, msgfmt);
    fprintf(stderr, "\n\n");
#endif

  } else {                 /* We can call ehfun */

    /* Compose the message */

    vsprintf(msg, msgfmt, ap);

    /* Call ehfun */

    ehfun(error_code, module, fname, msg, eh_data);

  }

  /* Finalize argument processing */
  
  va_end(ap);

  return;

}

/* 
 * CVErrHandler is the default error handling function.
 * It sends the error message to the stream pointed to by cv_errfp 
 */

#define errfp    (cv_mem->cv_errfp)

void CVErrHandler(int error_code, const char *module,
                  const char *function, char *msg, void *data)
{
  CVodeMem cv_mem;
  char err_type[10];

  /* data points to cv_mem here */

  cv_mem = (CVodeMem) data;

  if (error_code == CV_WARNING)
    sprintf(err_type,"WARNING");
  else
    sprintf(err_type,"ERROR");

#ifndef NO_FPRINTF_OUTPUT
  if (errfp!=NULL) {
    fprintf(errfp,"\n[%s %s]  %s\n",module,err_type,function);
    fprintf(errfp,"  %s\n\n",msg);
  }
#endif

  return;
}
```
