## Supplementary scripts for "A multi-scale *in silico* mouse model for insulin resistance and humanoid type 2 diabetes": cvodes_band.c

```
/*
 * -----------------------------------------------------------------
 * $Revision: 749 $
 * $Date: 2011-06-22 14:49:10 +0200 (Wed, 22 Jun 2011) $
 * ----------------------------------------------------------------- 
 * Programmer(s): Radu Serban @ LLNL
 * -----------------------------------------------------------------
 * Copyright (c) 2005, The Regents of the University of California.
 * Produced at the Lawrence Livermore National Laboratory.
 * All rights reserved.
 * For details, see the LICENSE file.
 * -----------------------------------------------------------------
 * This is the implementation file for the CVBAND linear solver.
 * -----------------------------------------------------------------
 */

#include <stdio.h>
#include <stdlib.h>

#include "cvodes_band_impl.h"
#include "cvodes_impl.h"

#include <sundials/sundials_math.h>

/* Other Constants */

#define MIN_INC_MULT RCONST(1000.0)
#define ZERO         RCONST(0.0)
#define ONE          RCONST(1.0)
#define TWO          RCONST(2.0)

/* CVBAND linit, lsetup, lsolve, and lfree routines */

static int CVBandInit(CVodeMem cv_mem);

static int CVBandSetup(CVodeMem cv_mem, int convfail, N_Vector ypred,
                       N_Vector fpred, booleantype *jcurPtr, N_Vector vtemp1,
                       N_Vector vtemp2, N_Vector vtemp3);

static int CVBandSolve(CVodeMem cv_mem, N_Vector b, N_Vector weight,
                       N_Vector ycur, N_Vector fcur);

static void CVBandFree(CVodeMem cv_mem);

/* CVBAND lfreeB function */

static void CVBandFreeB(CVadjMem ca_mem);

/* Wrapper function for adjoint code */

static int CVAbandJac(long int nB, long int mupperB, 
                      long int mlowerB, BandMat JB, realtype t, 
                      N_Vector yB, N_Vector fyB, void *cvadj_mem, 
                      N_Vector tmp1B, N_Vector tmp2B, N_Vector tmp3B);

/* CVBAND DQJac routine */

static int CVBandDQJac(long int n, long int mupper, long int mlower,
                       BandMat J, realtype t,
                       N_Vector y, N_Vector fy, void *jac_data,
                       N_Vector tmp1, N_Vector tmp2, N_Vector tmp3);


/* 
 * ================================================================
 *
 *                   PART I - forward problems
 *
 * ================================================================
 */


/* Readability Replacements */

#define lmm       (cv_mem->cv_lmm)
#define f         (cv_mem->cv_f)
#define f_data    (cv_mem->cv_f_data)
#define uround    (cv_mem->cv_uround)
#define nst       (cv_mem->cv_nst)
#define tn        (cv_mem->cv_tn)
#define h         (cv_mem->cv_h)
#define gamma     (cv_mem->cv_gamma)
#define gammap    (cv_mem->cv_gammap)
#define gamrat    (cv_mem->cv_gamrat)
#define ewt       (cv_mem->cv_ewt)
#define nfe       (cv_mem->cv_nfe)
#define linit     (cv_mem->cv_linit)
#define lsetup    (cv_mem->cv_lsetup)
#define lsolve    (cv_mem->cv_lsolve)
#define lfree     (cv_mem->cv_lfree)
#define lmem      (cv_mem->cv_lmem)
#define vec_tmpl      (cv_mem->cv_tempv)
#define setupNonNull  (cv_mem->cv_setupNonNull)

#define n          (cvband_mem->b_n)
#define jac        (cvband_mem->b_jac)
#define M          (cvband_mem->b_M)
#define mu         (cvband_mem->b_mu)
#define ml         (cvband_mem->b_ml)
#define storage_mu (cvband_mem->b_storage_mu)
#define pivots     (cvband_mem->b_pivots)
#define savedJ     (cvband_mem->b_savedJ)
#define nstlj      (cvband_mem->b_nstlj)
#define nje        (cvband_mem->b_nje)
#define nfeB       (cvband_mem->b_nfeB)
#define J_data     (cvband_mem->b_J_data)
#define last_flag  (cvband_mem->b_last_flag)


/*
 * -----------------------------------------------------------------
 * CVBand
 * -----------------------------------------------------------------
 * This routine initializes the memory record and sets various function
 * fields specific to the band linear solver module.  CVBand first calls
 * the existing lfree routine if this is not NULL.  It then sets the
 * cv_linit, cv_lsetup, cv_lsolve, and cv_lfree fields in (*cvode_mem)
 * to be CVBandInit, CVBandSetup, CVBandSolve, and CVBandFree,
 * respectively.  It allocates memory for a structure of type
 * CVBandMemRec and sets the cv_lmem field in (*cvode_mem) to the
 * address of this structure.  It sets setupNonNull in (*cvode_mem) to be
 * TRUE, b_mu to be mupper, b_ml to be mlower, and the b_jac field to be 
 * CVBandDQJac.
 * Finally, it allocates memory for M, savedJ, and pivot.  The CVBand
 * return value is SUCCESS = 0, LMEM_FAIL = -1, or LIN_ILL_INPUT = -2.
 *
 * NOTE: The band linear solver assumes a serial implementation
 *       of the NVECTOR package. Therefore, CVBand will first 
 *       test for compatible a compatible N_Vector internal
 *       representation by checking that the function 
 *       N_VGetArrayPointer exists.
 * -----------------------------------------------------------------
 */
                  
int CVBand(void *cvode_mem, long int N,
           long int mupper, long int mlower)
{
  CVodeMem cv_mem;
  CVBandMem cvband_mem;

  /* Return immediately if cvode_mem is NULL */
  if (cvode_mem == NULL) {
    CVProcessError(NULL, CVBAND_MEM_NULL, "CVBAND", "CVBand", MSGB_CVMEM_NULL);
    return(CVBAND_MEM_NULL);
  }
  cv_mem = (CVodeMem) cvode_mem;

  /* Test if the NVECTOR package is compatible with the BAND solver */
  if (vec_tmpl->ops->nvgetarraypointer == NULL) {
    CVProcessError(cv_mem, CVBAND_ILL_INPUT, "CVBAND", "CVBand", MSGB_BAD_NVECTOR);
    return(CVBAND_ILL_INPUT);
  }

  if (lfree != NULL) lfree(cv_mem);

  /* Set four main function fields in cv_mem */  
  linit  = CVBandInit;
  lsetup = CVBandSetup;
  lsolve = CVBandSolve;
  lfree  = CVBandFree;
  
  /* Get memory for CVBandMemRec */
  cvband_mem = NULL;
  cvband_mem = (CVBandMem) malloc(sizeof(CVBandMemRec));
  if (cvband_mem == NULL) {
    CVProcessError(cv_mem, CVBAND_MEM_FAIL, "CVBAND", "CVBand", MSGB_MEM_FAIL);
    return(CVBAND_MEM_FAIL);
  }
  
  /* Set default Jacobian routine and Jacobian data */
  jac = CVBandDQJac;
  J_data = cvode_mem;
  last_flag = CVBAND_SUCCESS;

  setupNonNull = TRUE;
  
  /* Load problem dimension */
  n = N;

  /* Load half-bandwiths in cvband_mem */
  ml = mlower;
  mu = mupper;

  /* Test ml and mu for legality */
  if ((ml < 0) || (mu < 0) || (ml >= N) || (mu >= N)) {
    CVProcessError(cv_mem, CVBAND_ILL_INPUT, "CVBAND", "CVBand", MSGB_BAD_SIZES);
    return(CVBAND_ILL_INPUT);
  }

  /* Set extended upper half-bandwith for M (required for pivoting) */
  storage_mu = MIN(N-1, mu + ml);

  /* Allocate memory for M, savedJ, and pivot arrays */
  M = NULL;
  M = BandAllocMat(N, mu, ml, storage_mu);
  if (M == NULL) {
    CVProcessError(cv_mem, CVBAND_MEM_FAIL, "CVBAND", "CVBand", MSGB_MEM_FAIL);
    free(cvband_mem); cvband_mem = NULL;
    return(CVBAND_MEM_FAIL);
  }
  savedJ = NULL;
  savedJ = BandAllocMat(N, mu, ml, mu);
  if (savedJ == NULL) {
    CVProcessError(cv_mem, CVBAND_MEM_FAIL, "CVBAND", "CVBand", MSGB_MEM_FAIL);
    BandFreeMat(M);
    free(cvband_mem); cvband_mem = NULL;
    return(CVBAND_MEM_FAIL);
  }
  pivots = NULL;
  pivots = BandAllocPiv(N);
  if (pivots == NULL) {
    CVProcessError(cv_mem, CVBAND_MEM_FAIL, "CVBAND", "CVBand", MSGB_MEM_FAIL);
    BandFreeMat(M);
    BandFreeMat(savedJ);
    free(cvband_mem); cvband_mem = NULL;
    return(CVBAND_MEM_FAIL);
  }

  /* Attach linear solver memory to integrator memory */
  lmem = cvband_mem;

  return(CVBAND_SUCCESS);
}

/*
 * -----------------------------------------------------------------
 * CVBandSetJacFn
 * -----------------------------------------------------------------
 */

int CVBandSetJacFn(void *cvode_mem, CVBandJacFn bjac, void *jac_data)
{
  CVodeMem cv_mem;
  CVBandMem cvband_mem;

  /* Return immediately if cvode_mem is NULL */
  if (cvode_mem == NULL) {
    CVProcessError(NULL, CVBAND_MEM_NULL, "CVBAND", "CVBandSetJacFn", MSGB_CVMEM_NULL);
    return(CVBAND_MEM_NULL);
  }
  cv_mem = (CVodeMem) cvode_mem;

  if (lmem == NULL) {
    CVProcessError(cv_mem, CVBAND_LMEM_NULL, "CVBAND", "CVBandSetJacFn", MSGB_LMEM_NULL);
    return(CVBAND_LMEM_NULL);
  }
  cvband_mem = (CVBandMem) lmem;

  jac = bjac;
  if (bjac != NULL) J_data = jac_data;

  return(CVBAND_SUCCESS);
}

/*
 * -----------------------------------------------------------------
 * CVBandGetWorkSpace
 * -----------------------------------------------------------------
 */

int CVBandGetWorkSpace(void *cvode_mem, long int *lenrwLS, long int *leniwLS)
{
  CVodeMem cv_mem;
  CVBandMem cvband_mem;

  /* Return immediately if cvode_mem is NULL */
  if (cvode_mem == NULL) {
    CVProcessError(NULL, CVBAND_MEM_NULL, "CVBAND", "CVBandGetWorkSpace", MSGB_CVMEM_NULL);
    return(CVBAND_MEM_NULL);
  }
  cv_mem = (CVodeMem) cvode_mem;

  if (lmem == NULL) {
    CVProcessError(cv_mem, CVBAND_LMEM_NULL, "CVBAND", "CVBandGetWorkSpace", MSGB_LMEM_NULL);
    return(CVBAND_LMEM_NULL);
  }
  cvband_mem = (CVBandMem) lmem;

  *lenrwLS = n*(storage_mu + mu + 2*ml + 2);
  *leniwLS = n;

  return(CVBAND_SUCCESS);
}

/*
 * -----------------------------------------------------------------
 * CVBandGetNumJacEvals
 * -----------------------------------------------------------------
 */

int CVBandGetNumJacEvals(void *cvode_mem, long int *njevals)
{
  CVodeMem cv_mem;
  CVBandMem cvband_mem;

  /* Return immediately if cvode_mem is NULL */
  if (cvode_mem == NULL) {
    CVProcessError(NULL, CVBAND_MEM_NULL, "CVBAND", "CVBandGetNumJacEvals", MSGB_CVMEM_NULL);
    return(CVBAND_MEM_NULL);
  }
  cv_mem = (CVodeMem) cvode_mem;

  if (lmem == NULL) {
    CVProcessError(cv_mem, CVBAND_LMEM_NULL, "CVBAND", "CVBandGetNumJacEvals", MSGB_LMEM_NULL);
    return(CVBAND_LMEM_NULL);
  }
  cvband_mem = (CVBandMem) lmem;

  *njevals = nje;

  return(CVBAND_SUCCESS);
}

/*
 * -----------------------------------------------------------------
 * CVBandGetNumRhsEvals
 * -----------------------------------------------------------------
 */

int CVBandGetNumRhsEvals(void *cvode_mem, long int *nfevalsLS)
{
  CVodeMem cv_mem;
  CVBandMem cvband_mem;

  /* Return immediately if cvode_mem is NULL */
  if (cvode_mem == NULL) {
    CVProcessError(NULL, CVBAND_MEM_NULL, "CVBAND", "CVBandGetNumRhsEvals", MSGB_CVMEM_NULL);
    return(CVBAND_MEM_NULL);
  }
  cv_mem = (CVodeMem) cvode_mem;

  if (lmem == NULL) {
    CVProcessError(cv_mem, CVBAND_LMEM_NULL, "CVBAND", "CVBandGetNumRhsEvals", MSGB_LMEM_NULL);
    return(CVBAND_LMEM_NULL);
  }
  cvband_mem = (CVBandMem) lmem;

  *nfevalsLS = nfeB;

  return(CVBAND_SUCCESS);
}

/*
 * -----------------------------------------------------------------
 * CVBandGetLastFlag
 * -----------------------------------------------------------------
 */

int CVBandGetLastFlag(void *cvode_mem, int *flag)
{
  CVodeMem cv_mem;
  CVBandMem cvband_mem;

  /* Return immediately if cvode_mem is NULL */
  if (cvode_mem == NULL) {
    CVProcessError(NULL, CVBAND_MEM_NULL, "CVBAND", "CVBandGetLastFlag", MSGB_CVMEM_NULL);
    return(CVBAND_MEM_NULL);
  }
  cv_mem = (CVodeMem) cvode_mem;

  if (lmem == NULL) {
    CVProcessError(cv_mem, CVBAND_LMEM_NULL, "CVBAND", "CVBandGetLastFlag", MSGB_LMEM_NULL);
    return(CVBAND_LMEM_NULL);
  }
  cvband_mem = (CVBandMem) lmem;

  *flag = last_flag;

  return(CVBAND_SUCCESS);
}

/*
 * -----------------------------------------------------------------
 * CVBandGetReturnFlagName
 * -----------------------------------------------------------------
 */

char *CVBandGetReturnFlagName(int flag)
{
  char *name;

  name = (char *)malloc(30*sizeof(char));

  switch(flag) {
  case CVBAND_SUCCESS:
    sprintf(name,"CVBAND_SUCCESS");
    break;  
  case CVBAND_MEM_NULL:
    sprintf(name,"CVBAND_MEM_NULL");
    break;
  case CVBAND_LMEM_NULL:
    sprintf(name,"CVBAND_LMEM_NULL");
    break;
  case CVBAND_ILL_INPUT:
    sprintf(name,"CVBAND_ILL_INPUT");
    break;
  case CVBAND_MEM_FAIL:
    sprintf(name,"CVBAND_MEM_FAIL");
    break;
  case CVBAND_JACFUNC_UNRECVR:
    sprintf(name,"CVBAND_JACFUNC_UNRECVR");
    break;
  case CVBAND_JACFUNC_RECVR:
    sprintf(name,"CVBAND_JACFUNC_RECVR");
    break;
  case CVBAND_ADJMEM_NULL:
    sprintf(name,"CVBAND_ADJMEM_NULL");
    break;
  case CVBAND_LMEMB_NULL:
    sprintf(name,"CVBAND_LMEMB_NULL");
    break;
  default:
    sprintf(name,"NONE");
  }

  return(name);
}

/*
 * -----------------------------------------------------------------
 * CVBandInit
 * -----------------------------------------------------------------
 * This routine does remaining initializations specific to the band
 * linear solver.
 * -----------------------------------------------------------------
 */

static int CVBandInit(CVodeMem cv_mem)
{
  CVBandMem cvband_mem;

  cvband_mem = (CVBandMem) lmem;

  nje   = 0;
  nfeB  = 0;
  nstlj = 0;

  if (jac == NULL) {
    jac = CVBandDQJac;
    J_data = cv_mem;
  }

  last_flag = CVBAND_SUCCESS;
  return(0);
}

/*
 * -----------------------------------------------------------------
 * CVBandSetup
 * -----------------------------------------------------------------
 * This routine does the setup operations for the band linear solver.
 * It makes a decision whether or not to call the Jacobian evaluation
 * routine based on various state variables, and if not it uses the 
 * saved copy.  In any case, it constructs the Newton matrix 
 * M = I - gamma*J, updates counters, and calls the band LU 
 * factorization routine.
 * -----------------------------------------------------------------
 */

static int CVBandSetup(CVodeMem cv_mem, int convfail, N_Vector ypred,
                       N_Vector fpred, booleantype *jcurPtr, N_Vector vtemp1,
                       N_Vector vtemp2, N_Vector vtemp3)
{
  booleantype jbad, jok;
  realtype dgamma;
  long int ier;
  CVBandMem cvband_mem;
  int retval;

  cvband_mem = (CVBandMem) lmem;

  /* Use nst, gamma/gammap, and convfail to set J eval. flag jok */

  dgamma = ABS((gamma/gammap) - ONE);
  jbad = (nst == 0) || (nst > nstlj + CVB_MSBJ) ||
         ((convfail == CV_FAIL_BAD_J) && (dgamma < CVB_DGMAX)) ||
         (convfail == CV_FAIL_OTHER);
  jok = !jbad;
  
  if (jok) {

    /* If jok = TRUE, use saved copy of J */
    *jcurPtr = FALSE;
    BandCopy(savedJ, M, mu, ml);

  } else {

    /* If jok = FALSE, call jac routine for new J value */
    nje++;
    nstlj = nst;
    *jcurPtr = TRUE;
    BandZero(M); 

    retval = jac(n, mu, ml, M, tn, ypred, fpred, J_data, vtemp1, vtemp2, vtemp3);
    if (retval < 0) {
      CVProcessError(cv_mem, CVBAND_JACFUNC_UNRECVR, "CVBAND", "CVBandSetup", MSGB_JACFUNC_FAILED);
      last_flag = CVBAND_JACFUNC_UNRECVR;
      return(-1);
    }
    if (retval > 0) {
      last_flag = CVBAND_JACFUNC_RECVR;
      return(1);
    }

    BandCopy(M, savedJ, mu, ml);

  }
  
  /* Scale and add I to get M = I - gamma*J */
  BandScale(-gamma, M);
  BandAddI(M);

  /* Do LU factorization of M */
  ier = BandGBTRF(M, pivots);

  /* Return 0 if the LU was complete; otherwise return 1 */
  if (ier > 0) {
    last_flag = ier;
    return(1);
  }
  last_flag = CVBAND_SUCCESS;
  return(0);
}

/*
 * -----------------------------------------------------------------
 * CVBandSolve
 * -----------------------------------------------------------------
 * This routine handles the solve operation for the band linear solver
 * by calling the band backsolve routine.  The return value is 0.
 * -----------------------------------------------------------------
 */

static int CVBandSolve(CVodeMem cv_mem, N_Vector b, N_Vector weight,
                       N_Vector ycur, N_Vector fcur)
{
  CVBandMem cvband_mem;
  realtype *bd;

  cvband_mem = (CVBandMem) lmem;

  bd = N_VGetArrayPointer(b);

  BandGBTRS(M, pivots, bd);

  /* If CV_BDF, scale the correction to account for change in gamma */
  if ((lmm == CV_BDF) && (gamrat != ONE)) {
    N_VScale(TWO/(ONE + gamrat), b, b);
  }

  last_flag = CVBAND_SUCCESS;
  return(0);
}

/*
 * -----------------------------------------------------------------
 * CVBandFree
 * -----------------------------------------------------------------
 * This routine frees memory specific to the band linear solver.
 * -----------------------------------------------------------------
 */

static void CVBandFree(CVodeMem cv_mem)
{
  CVBandMem cvband_mem;

  cvband_mem = (CVBandMem) lmem;

  BandFreeMat(M);
  BandFreeMat(savedJ);
  BandFreePiv(pivots);
  free(cvband_mem); cvband_mem = NULL;
}

/*
 * -----------------------------------------------------------------
 * CVBandDQJac
 * -----------------------------------------------------------------
 * This routine generates a banded difference quotient approximation to
 * the Jacobian of f(t,y).  It assumes that a band matrix of type
 * BandMat is stored column-wise, and that elements within each column
 * are contiguous. This makes it possible to get the address of a column
 * of J via the macro BAND_COL and to write a simple for loop to set
 * each of the elements of a column in succession.
 * -----------------------------------------------------------------
 */

static int CVBandDQJac(long int N, long int mupper, long int mlower,
                       BandMat J, realtype t,
                       N_Vector y, N_Vector fy, void *jac_data,
                       N_Vector tmp1, N_Vector tmp2, N_Vector tmp3)
{
  N_Vector ftemp, ytemp;
  realtype fnorm, minInc, inc, inc_inv, srur;
  realtype *col_j, *ewt_data, *fy_data, *ftemp_data, *y_data, *ytemp_data;
  long int group, i, j, width, ngroups, i1, i2;
  int retval = 0;

  CVodeMem cv_mem;
  CVBandMem cvband_mem;

  /* jac_dat points to cvode_mem */
  cv_mem = (CVodeMem) jac_data;
  cvband_mem = (CVBandMem) lmem;

  /* Rename work vectors for use as temporary values of y and f */
  ftemp = tmp1;
  ytemp = tmp2;

  /* Obtain pointers to the data for ewt, fy, ftemp, y, ytemp */
  ewt_data   = N_VGetArrayPointer(ewt);
  fy_data    = N_VGetArrayPointer(fy);
  ftemp_data = N_VGetArrayPointer(ftemp);
  y_data     = N_VGetArrayPointer(y);
  ytemp_data = N_VGetArrayPointer(ytemp);

  /* Load ytemp with y = predicted y vector */
  N_VScale(ONE, y, ytemp);

  /* Set minimum increment based on uround and norm of f */
  srur = RSqrt(uround);
  fnorm = N_VWrmsNorm(fy, ewt);
  minInc = (fnorm != ZERO) ?
           (MIN_INC_MULT * ABS(h) * uround * N * fnorm) : ONE;

  /* Set bandwidth and number of column groups for band differencing */
  width = mlower + mupper + 1;
  ngroups = MIN(width, N);
  
  for (group=1; group <= ngroups; group++) {
    
    /* Increment all y_j in group */
    for(j=group-1; j < N; j+=width) {
      inc = MAX(srur*ABS(y_data[j]), minInc/ewt_data[j]);
      ytemp_data[j] += inc;
    }

    /* Evaluate f with incremented y */

    retval = f(tn, ytemp, ftemp, f_data);
    nfeB++;
    if (retval != 0) break;

    /* Restore ytemp, then form and load difference quotients */
    for (j=group-1; j < N; j+=width) {
      ytemp_data[j] = y_data[j];
      col_j = BAND_COL(J,j);
      inc = MAX(srur*ABS(y_data[j]), minInc/ewt_data[j]);
      inc_inv = ONE/inc;
      i1 = MAX(0, j-mupper);
      i2 = MIN(j+mlower, N-1);
      for (i=i1; i <= i2; i++)
        BAND_COL_ELEM(col_j,i,j) =
          inc_inv * (ftemp_data[i] - fy_data[i]);
    }
  }
  
  return(retval);
}


/* 
 * ================================================================
 *
 *                   PART II - backward problems
 *
 * ================================================================
 */


/* Additional readability replacements */

#define ytmp        (ca_mem->ca_ytmp)
#define getY        (ca_mem->ca_getY)
#define lmemB       (ca_mem->ca_lmemB)
#define lfreeB      (ca_mem->ca_lfreeB)

#define bjac_B      (cvbandB_mem->b_bjacB)
#define jac_data_B  (cvbandB_mem->b_jac_dataB)

/*
 * CVBandB and CVBandSet*B
 *
 * Wrappers for the backward phase around the corresponding 
 * CVODES functions
 */

int CVBandB(void *cvadj_mem, long int nB, 
            long int mupperB, long int mlowerB)
{
  CVadjMem ca_mem;
  CVBandMemB cvbandB_mem;
  CVodeMem cvB_mem;
  int flag;

  if (cvadj_mem == NULL) {
    CVProcessError(NULL, CVBAND_ADJMEM_NULL, "CVBAND", "CVBandB", MSGB_CAMEM_NULL);
    return(CVBAND_ADJMEM_NULL);
  }
  ca_mem = (CVadjMem) cvadj_mem;

  cvB_mem = ca_mem->cvb_mem;

  /* Get memory for CVBandMemRecB */
  cvbandB_mem = (CVBandMemB) malloc(sizeof(CVBandMemRecB));
  if (cvbandB_mem == NULL) {
    CVProcessError(cvB_mem, CVBAND_MEM_FAIL, "CVBAND", "CVBandB", MSGB_MEM_FAIL);
    return(CVBAND_MEM_FAIL);
  }

  bjac_B = NULL;
  jac_data_B = NULL;

  /* attach lmemB and lfreeB */
  lmemB = cvbandB_mem;
  lfreeB = CVBandFreeB;

  flag = CVBand(cvB_mem, nB, mupperB, mlowerB);

  if (flag != CVBAND_SUCCESS) {
    free(cvbandB_mem);
    cvbandB_mem = NULL;
  }

  return(flag);
}

int CVBandSetJacFnB(void *cvadj_mem, CVBandJacFnB bjacB, void *jac_dataB)
{
  CVadjMem ca_mem;
  CVBandMemB cvbandB_mem;
  CVodeMem cvB_mem;
  int flag;

  if (cvadj_mem == NULL) {
    CVProcessError(NULL, CVBAND_ADJMEM_NULL, "CVBAND", "CVBandSetJacFnB", MSGB_CAMEM_NULL);
    return(CVBAND_ADJMEM_NULL);
  }
  ca_mem = (CVadjMem) cvadj_mem;

  cvB_mem = ca_mem->cvb_mem;

  if (lmemB == NULL) {
    CVProcessError(cvB_mem, CVBAND_LMEMB_NULL, "CVBAND", "CVBandSetJacFnB", MSGB_LMEMB_NULL);
    return(CVBAND_LMEMB_NULL);
  }
  cvbandB_mem = (CVBandMemB) lmemB;

  bjac_B     = bjacB;
  jac_data_B = jac_dataB;

  flag = CVBandSetJacFn(cvB_mem, CVAbandJac, cvadj_mem);

  return(flag);
}

/*
 * CVBandFreeB 
 */

static void CVBandFreeB(CVadjMem ca_mem)
{
  CVBandMemB cvbandB_mem;

  cvbandB_mem = (CVBandMemB) lmemB;

  free(cvbandB_mem);
}

/*
 * CVAbandJac
 *
 * This routine interfaces to the CVBandJacFnB routine provided 
 * by the user.
 * NOTE: jac_data actually contains cvadj_mem
 */

static int CVAbandJac(long int nB, long int mupperB, 
                      long int mlowerB, BandMat JB, realtype t, 
                      N_Vector yB, N_Vector fyB, void *cvadj_mem, 
                      N_Vector tmp1B, N_Vector tmp2B, N_Vector tmp3B)
{
  CVadjMem ca_mem;
  CVodeMem cvB_mem;
  CVBandMemB cvbandB_mem;
  int retval, flag;

  ca_mem = (CVadjMem) cvadj_mem;
  cvB_mem = ca_mem->cvb_mem;
  cvbandB_mem = (CVBandMemB) lmemB;

  /* Forward solution from interpolation */
  flag = getY(ca_mem, t, ytmp);
  if (flag != CV_SUCCESS) {
    CVProcessError(cvB_mem, -1, "CVBAND", "CVAbandJac", MSGB_BAD_T);
    return(-1);
  }

  /* Call user's adjoint band bjacB routine */
  retval = bjac_B(nB, mupperB, mlowerB, JB, t, ytmp, yB, fyB, jac_data_B,
                  tmp1B, tmp2B, tmp3B);

  return(retval);
}
```
