## Supplementary scripts for "A multi-scale *in silico* mouse model for insulin resistance and humanoid type 2 diabetes": cvodes_band_impl.h

```
/*
 * -----------------------------------------------------------------
 * $Revision: 749 $
 * $Date: 2011-06-22 14:49:10 +0200 (Wed, 22 Jun 2011) $
 * ----------------------------------------------------------------- 
 * Programmer(s): Radu Serban @ LLNL
 * -----------------------------------------------------------------
 * Copyright (c) 2005, The Regents of the University of California.
 * Produced at the Lawrence Livermore National Laboratory.
 * All rights reserved.
 * For details, see the LICENSE file.
 * -----------------------------------------------------------------
 * Implementation header file for the band linear solver, CVBAND.
 * -----------------------------------------------------------------
 */

#ifndef _CVSBAND_IMPL_H
#define _CVSBAND_IMPL_H

#ifdef __cplusplus  /* wrapper to enable C++ usage */
extern "C" {
#endif

#include <cvodes/cvodes_band.h>

  /*
   * -----------------------------------------------------------------
   * Types: CVBandMemRec, CVBandMem                                
   * -----------------------------------------------------------------
   * The type CVBandMem is pointer to a CVBandMemRec.
   * This structure contains CVBand solver-specific data.                
   *
   * CVBand attaches such a structure to the lmem field of CVodeMem
   * -----------------------------------------------------------------
   */                                                                

  typedef struct {

    long int b_n;           /* N = problem dimension                    */

    CVBandJacFn b_jac;      /* jac = Jacobian routine to be called      */

    long int b_ml;          /* b_ml = lower bandwidth of savedJ         */
  
    long int b_mu;          /* b_mu = upper bandwidth of savedJ         */ 
  
    long int b_storage_mu;  /* upper bandwith of M = MIN(N-1,b_mu+b_ml) */
  
    BandMat b_M;            /* M = I - gamma J, gamma = h / l1          */
  
    long int *b_pivots;     /* pivots = pivot array for PM = LU         */
  
    BandMat b_savedJ;       /* savedJ = old Jacobian                    */
  
    long int b_nstlj;       /* nstlj = nst at last Jacobian eval.       */
  
    long int b_nje;         /* nje = no. of calls to jac                */
  
    long int b_nfeB;        /* nfeB = no. of calls to f due to difference
                               quotient band Jacobian approximation     */

    void *b_J_data;         /* J_data is passed to jac                  */

    int b_last_flag;        /* last error return flag                   */
  
  } CVBandMemRec, *CVBandMem;


  /*
   * -----------------------------------------------------------------
   * Types : CVBandMemRecB, CVBandMemB       
   * -----------------------------------------------------------------
   * CVBandB attaches such a structure to the lmemB filed of CVadjMem
   * -----------------------------------------------------------------
   */

  typedef struct {

    CVBandJacFnB b_bjacB;
    void *b_jac_dataB;

  } CVBandMemRecB, *CVBandMemB;


  /*
   * -----------------------------------------------------------------
   * Error Messages 
   * -----------------------------------------------------------------
   */

#define MSGB_CVMEM_NULL "Integrator memory is NULL."
#define MSGB_MEM_FAIL "A memory request failed."
#define MSGB_BAD_SIZES "Illegal bandwidth parameter(s). Must have 0 <=  ml, mu <= N-1."
#define MSGB_BAD_NVECTOR "A required vector operation is not implemented."
#define MSGB_LMEM_NULL "CVBAND memory is NULL."
#define MSGB_JACFUNC_FAILED "The Jacobian routine failed in an unrecoverable manner."

#define MSGB_CAMEM_NULL "cvadj_mem = NULL illegal."
#define MSGB_LMEMB_NULL "CVBAND memory is NULL for the backward integration."
#define MSGB_BAD_T "Bad t for interpolation."

#ifdef __cplusplus
}
#endif

#endif
```
