## Supplementary scripts for "A multi-scale *in silico* mouse model for insulin resistance and humanoid type 2 diabetes": cvodes_bandpre.c

```
/*
 * -----------------------------------------------------------------
 * $Revision: 749 $
 * $Date: 2011-06-22 14:49:10 +0200 (Wed, 22 Jun 2011) $
 * ----------------------------------------------------------------- 
 * Programmer(s): Radu Serban and Aaron Collier @ LLNL
 * -----------------------------------------------------------------
 * Copyright (c) 2005, The Regents of the University of California.
 * Produced at the Lawrence Livermore National Laboratory.
 * All rights reserved.
 * For details, see the LICENSE file.
 * -----------------------------------------------------------------
 * This file contains implementations of the banded difference
 * quotient Jacobian-based preconditioner and solver routines for
 * use with the CVSPILS linear solvers.
 * -----------------------------------------------------------------
 */

#include <stdio.h>
#include <stdlib.h>

#include "cvodes_impl.h"
#include "cvodes_bandpre_impl.h"

#include <cvodes/cvodes_sptfqmr.h>
#include <cvodes/cvodes_spbcgs.h>
#include <cvodes/cvodes_spgmr.h>

#include <sundials/sundials_math.h>

#define MIN_INC_MULT RCONST(1000.0)
#define ZERO         RCONST(0.0)
#define ONE          RCONST(1.0)

/* Prototypes of CVBandPrecSetup and CVBandPrecSolve */
  
static int CVBandPrecSetup(realtype t, N_Vector y, N_Vector fy, 
                           booleantype jok, booleantype *jcurPtr, 
                           realtype gamma, void *bp_data,
                           N_Vector tmp1, N_Vector tmp2, N_Vector tmp3);

static int CVBandPrecSolve(realtype t, N_Vector y, N_Vector fy, 
                           N_Vector r, N_Vector z, 
                           realtype gamma, realtype delta,
                           int lr, void *bp_data, N_Vector tmp);

/* Prototype for difference quotient Jacobian calculation routine */

static int CVBandPDQJac(CVBandPrecData pdata, 
                        realtype t, N_Vector y, N_Vector fy, 
                        N_Vector ftemp, N_Vector ytemp);

/* 
 * ================================================================
 *
 *                   PART I - forward problems
 *
 * ================================================================
 */

/* Redability replacements */

#define vec_tmpl (cv_mem->cv_tempv)

/*
 * -----------------------------------------------------------------
 * Malloc, Free, and Get Functions
 * NOTE: The band linear solver assumes a serial implementation
 *       of the NVECTOR package. Therefore, CVBandPrecAlloc will
 *       first test for a compatible N_Vector internal representation
 *       by checking that the function N_VGetArrayPointer exists.
 * -----------------------------------------------------------------
 */

void *CVBandPrecAlloc(void *cvode_mem, long int N, 
                      long int mu, long int ml)
{
  CVodeMem cv_mem;
  CVBandPrecData pdata;
  long int mup, mlp, storagemu;

  if (cvode_mem == NULL) {
    CVProcessError(NULL, 0, "CVBANDPRE", "CVBandPrecAlloc", MSGBP_CVMEM_NULL);
    return(NULL);
  }
  cv_mem = (CVodeMem) cvode_mem;

  /* Test if the NVECTOR package is compatible with the BAND preconditioner */
  if(vec_tmpl->ops->nvgetarraypointer == NULL) {
    CVProcessError(cv_mem, 0, "CVBANDPRE", "CVBandPrecAlloc", MSGBP_BAD_NVECTOR);
    return(NULL);
  }

  pdata = NULL;
  pdata = (CVBandPrecData) malloc(sizeof *pdata);  /* Allocate data memory */
  if (pdata == NULL) {
    CVProcessError(cv_mem, 0, "CVBANDPRE", "CVBandPrecAlloc", MSGBP_MEM_FAIL);
    return(NULL);
  }

  /* Load pointers and bandwidths into pdata block. */
  pdata->cvode_mem = cvode_mem;
  pdata->N = N;
  pdata->mu = mup = MIN(N-1, MAX(0,mu));
  pdata->ml = mlp = MIN(N-1, MAX(0,ml));

  /* Initialize nfeBP counter */
  pdata->nfeBP = 0;

  /* Allocate memory for saved banded Jacobian approximation. */
  pdata->savedJ = NULL;
  pdata->savedJ = BandAllocMat(N, mup, mlp, mup);
  if (pdata->savedJ == NULL) {
    free(pdata); pdata = NULL;
    CVProcessError(cv_mem, 0, "CVBANDPRE", "CVBandPrecAlloc", MSGBP_MEM_FAIL);
    return(NULL);
  }

  /* Allocate memory for banded preconditioner. */
  storagemu = MIN(N-1, mup+mlp);
  pdata->savedP = NULL;
  pdata->savedP = BandAllocMat(N, mup, mlp, storagemu);
  if (pdata->savedP == NULL) {
    BandFreeMat(pdata->savedJ);
    free(pdata); pdata = NULL;
    CVProcessError(cv_mem, 0, "CVBANDPRE", "CVBandPrecAlloc", MSGBP_MEM_FAIL);
    return(NULL);
  }

  /* Allocate memory for pivot array. */
  pdata->pivots = NULL;
  pdata->pivots = BandAllocPiv(N);
  if (pdata->savedJ == NULL) {
    BandFreeMat(pdata->savedP);
    BandFreeMat(pdata->savedJ);
    free(pdata); pdata = NULL;
    CVProcessError(cv_mem, 0, "CVBANDPRE", "CVBandPrecAlloc", MSGBP_MEM_FAIL);
    return(NULL);
  }

  return((void *) pdata);
}

int CVBPSptfqmr(void *cvode_mem, int pretype, int maxl, void *p_data)
{
  CVodeMem cv_mem;
  int flag;

  flag = CVSptfqmr(cvode_mem, pretype, maxl);
  if(flag != CVSPILS_SUCCESS) return(flag);
  
  cv_mem = (CVodeMem) cvode_mem;

  if ( p_data == NULL ) {
    CVProcessError(cv_mem, CVBANDPRE_PDATA_NULL, "CVBANDPRE", "CVBPSptfqmr", MSGBP_PDATA_NULL);
    return(CVBANDPRE_PDATA_NULL);
  } 

  flag = CVSpilsSetPreconditioner(cvode_mem, CVBandPrecSetup, CVBandPrecSolve, p_data);
  if(flag != CVSPILS_SUCCESS) return(flag);

  return(CVSPILS_SUCCESS);
}

int CVBPSpbcg(void *cvode_mem, int pretype, int maxl, void *p_data)
{
  CVodeMem cv_mem;
  int flag;

  flag = CVSpbcg(cvode_mem, pretype, maxl);
  if(flag != CVSPILS_SUCCESS) return(flag);

  cv_mem = (CVodeMem) cvode_mem;

  if ( p_data == NULL ) {
    CVProcessError(cv_mem, CVBANDPRE_PDATA_NULL, "CVBANDPRE", "CVBPSpbcg", MSGBP_PDATA_NULL);
    return(CVBANDPRE_PDATA_NULL);
  } 

  flag = CVSpilsSetPreconditioner(cvode_mem, CVBandPrecSetup, CVBandPrecSolve, p_data);
  if(flag != CVSPILS_SUCCESS) return(flag);

  return(CVSPILS_SUCCESS);
}

int CVBPSpgmr(void *cvode_mem, int pretype, int maxl, void *p_data)
{
  CVodeMem cv_mem;
  int flag;

  flag = CVSpgmr(cvode_mem, pretype, maxl);
  if(flag != CVSPILS_SUCCESS) return(flag);

  cv_mem = (CVodeMem) cvode_mem;

  if ( p_data == NULL ) {
    CVProcessError(cv_mem, CVBANDPRE_PDATA_NULL, "CVBANDPRE", "CVBPSpgmr", MSGBP_PDATA_NULL);
    return(CVBANDPRE_PDATA_NULL);
  }

  flag = CVSpilsSetPreconditioner(cvode_mem, CVBandPrecSetup, CVBandPrecSolve, p_data);
  if(flag != CVSPILS_SUCCESS) return(flag);

  return(CVSPILS_SUCCESS);
}

void CVBandPrecFree(void **bp_data)
{
  CVBandPrecData pdata;

  if (*bp_data == NULL) return;

  pdata = (CVBandPrecData) (*bp_data);
  BandFreeMat(pdata->savedJ);
  BandFreeMat(pdata->savedP);
  BandFreePiv(pdata->pivots);

  free(*bp_data);
  *bp_data = NULL;

}

int CVBandPrecGetWorkSpace(void *bp_data, long int *lenrwBP, long int *leniwBP)
{
  CVBandPrecData pdata;
  long int N, ml, mu, smu;

  if ( bp_data == NULL ) {
    CVProcessError(NULL, CVBANDPRE_PDATA_NULL, "CVBANDPRE", "CVBandPrecGetWorkSpace", MSGBP_PDATA_NULL);
    return(CVBANDPRE_PDATA_NULL);
  } 

  pdata = (CVBandPrecData) bp_data;

  N   = pdata->N;
  mu  = pdata->mu;
  ml  = pdata->ml;
  smu = MIN( N-1, mu + ml);

  *leniwBP = pdata->N;
  *lenrwBP = N * ( 2*ml + smu + mu + 2 );

  return(CVBANDPRE_SUCCESS);
}

int CVBandPrecGetNumRhsEvals(void *bp_data, long int *nfevalsBP)
{
  CVBandPrecData pdata;

  if (bp_data == NULL) {
    CVProcessError(NULL, CVBANDPRE_PDATA_NULL, "CVBANDPRE", "CVBandPrecGetNumRhsEvals", MSGBP_PDATA_NULL);
    return(CVBANDPRE_PDATA_NULL);
  } 

  pdata = (CVBandPrecData) bp_data;

  *nfevalsBP = pdata->nfeBP;

  return(CVBANDPRE_SUCCESS);
}

/*
 * -----------------------------------------------------------------
 * CVBandPrecGetReturnFlagName
 * -----------------------------------------------------------------
 */

char *CVBandPrecGetReturnFlagName(int flag)
{
  char *name;

  name = (char *)malloc(30*sizeof(char));

  switch(flag) {
  case CVBANDPRE_SUCCESS:
    sprintf(name,"CVBANDPRE_SUCCESS");
    break;    
  case CVBANDPRE_PDATA_NULL:
    sprintf(name,"CVBANDPRE_PDATA_NULL");
    break;
  case CVBANDPRE_RHSFUNC_UNRECVR:
    sprintf(name,"CVBANDPRE_RHSFUNC_UNRECVR");
    break;
  case CVBANDPRE_ADJMEM_NULL:
    sprintf(name,"CVBANDPRE_ADJMEM_NULL");
    break;
  case CVBANDPRE_MEM_FAIL:
    sprintf(name,"CVBANDPRE_MEM_FAIL");
    break;
  default:
    sprintf(name,"NONE");
  }

  return(name);
}

/* Readability Replacements */

#define N      (pdata->N)
#define mu     (pdata->mu)
#define ml     (pdata->ml)
#define pivots (pdata->pivots)
#define savedJ (pdata->savedJ)
#define savedP (pdata->savedP)
#define nfeBP  (pdata->nfeBP)

/*
 * -----------------------------------------------------------------
 * CVBandPrecSetup
 * -----------------------------------------------------------------
 * Together CVBandPrecSetup and CVBandPrecSolve use a banded
 * difference quotient Jacobian to create a preconditioner.
 * CVBandPrecSetup calculates a new J, if necessary, then
 * calculates P = I - gamma*J, and does an LU factorization of P.
 *
 * The parameters of CVBandPrecSetup are as follows:
 *
 * t       is the current value of the independent variable.
 *
 * y       is the current value of the dependent variable vector,
 *         namely the predicted value of y(t).
 *
 * fy      is the vector f(t,y).
 *
 * jok     is an input flag indicating whether Jacobian-related
 *         data needs to be recomputed, as follows:
 *           jok == FALSE means recompute Jacobian-related data
 *                  from scratch.
 *           jok == TRUE means that Jacobian data from the
 *                  previous PrecSetup call will be reused
 *                  (with the current value of gamma).
 *         A CVBandPrecSetup call with jok == TRUE should only
 *         occur after a call with jok == FALSE.
 *
 * *jcurPtr is a pointer to an output integer flag which is
 *          set by CVBandPrecond as follows:
 *            *jcurPtr = TRUE if Jacobian data was recomputed.
 *            *jcurPtr = FALSE if Jacobian data was not recomputed,
 *                       but saved data was reused.
 *
 * gamma   is the scalar appearing in the Newton matrix.
 *
 * bp_data is a pointer to preconditoner data - the same as the
 *         bp_data parameter passed to CVSp*.
 *
 * tmp1, tmp2, and tmp3 are pointers to memory allocated
 *           for vectors of length N for work space. This
 *           routine uses only tmp1 and tmp2.
 *
 * The value to be returned by the CVBandPrecSetup function is
 *   0  if successful, or
 *   1  if the band factorization failed.
 * -----------------------------------------------------------------
 */

static int CVBandPrecSetup(realtype t, N_Vector y, N_Vector fy, 
                           booleantype jok, booleantype *jcurPtr, 
                           realtype gamma, void *bp_data,
                           N_Vector tmp1, N_Vector tmp2, N_Vector tmp3)
{
  long int ier;
  CVBandPrecData pdata;
  CVodeMem cv_mem;
  int retval;

  /* Assume matrix and pivots have already been allocated. */
  pdata = (CVBandPrecData) bp_data;

  cv_mem = (CVodeMem) pdata->cvode_mem;

  if (jok) {

    /* If jok = TRUE, use saved copy of J. */
    *jcurPtr = FALSE;
    BandCopy(savedJ, savedP, mu, ml);

  } else {

    /* If jok = FALSE, call CVBandPDQJac for new J value. */
    *jcurPtr = TRUE;
    BandZero(savedJ);

    retval = CVBandPDQJac(pdata, t, y, fy, tmp1, tmp2);
    if (retval < 0) {
      CVProcessError(cv_mem, CVBANDPRE_RHSFUNC_UNRECVR, "CVBANDPRE", "CVBandPrecSetup", MSGBP_RHSFUNC_FAILED);
      return(-1);
    }
    if (retval > 0) {
      return(1);
    }

    BandCopy(savedJ, savedP, mu, ml);

  }
  
  /* Scale and add I to get savedP = I - gamma*J. */
  BandScale(-gamma, savedP);
  BandAddI(savedP);
 
  /* Do LU factorization of matrix. */
  ier = BandGBTRF(savedP, pivots);
 
  /* Return 0 if the LU was complete; otherwise return 1. */
  if (ier > 0) return(1);
  return(0);
}

/*
 * -----------------------------------------------------------------
 * CVBandPrecSolve
 * -----------------------------------------------------------------
 * CVBandPrecSolve solves a linear system P z = r, where P is the
 * matrix computed by CVBandPrecond.
 *
 * The parameters of CVBandPrecSolve used here are as follows:
 *
 * r       is the right-hand side vector of the linear system.
 *
 * bp_data is a pointer to preconditioner data - the same as the
 *         bp_data parameter passed to CVSp*.
 *
 * z       is the output vector computed by CVBandPrecSolve.
 *
 * The value returned by the CVBandPrecSolve function is always 0,
 * indicating success.
 * -----------------------------------------------------------------
 */ 

static int CVBandPrecSolve(realtype t, N_Vector y, N_Vector fy, 
                           N_Vector r, N_Vector z, 
                           realtype gamma, realtype delta,
                           int lr, void *bp_data, N_Vector tmp)
{
  CVBandPrecData pdata;
  realtype *zd;

  /* Assume matrix and pivots have already been allocated. */
  pdata = (CVBandPrecData) bp_data;

  /* Copy r to z. */
  N_VScale(ONE, r, z);

  /* Do band backsolve on the vector z. */
  zd = N_VGetArrayPointer(z);

  BandGBTRS(savedP, pivots, zd);

  return(0);
}

#define ewt    (cv_mem->cv_ewt)
#define uround (cv_mem->cv_uround)
#define h      (cv_mem->cv_h)
#define f      (cv_mem->cv_f)
#define f_data (cv_mem->cv_f_data)

/*
 * -----------------------------------------------------------------
 * CVBandPDQJac
 * -----------------------------------------------------------------
 * This routine generates a banded difference quotient approximation to
 * the Jacobian of f(t,y). It assumes that a band matrix of type
 * BandMat is stored column-wise, and that elements within each column
 * are contiguous. This makes it possible to get the address of a column
 * of J via the macro BAND_COL and to write a simple for loop to set
 * each of the elements of a column in succession.
 * -----------------------------------------------------------------
 */

static int CVBandPDQJac(CVBandPrecData pdata, 
                        realtype t, N_Vector y, N_Vector fy, 
                        N_Vector ftemp, N_Vector ytemp)
{
  CVodeMem cv_mem;
  realtype fnorm, minInc, inc, inc_inv, srur;
  long int group, i, j, width, ngroups, i1, i2;
  realtype *col_j, *ewt_data, *fy_data, *ftemp_data, *y_data, *ytemp_data;
  int retval;

  cv_mem = (CVodeMem) pdata->cvode_mem;

  /* Obtain pointers to the data for ewt, fy, ftemp, y, ytemp. */
  ewt_data   = N_VGetArrayPointer(ewt);
  fy_data    = N_VGetArrayPointer(fy);
  ftemp_data = N_VGetArrayPointer(ftemp);
  y_data     = N_VGetArrayPointer(y);
  ytemp_data = N_VGetArrayPointer(ytemp);

  /* Load ytemp with y = predicted y vector. */
  N_VScale(ONE, y, ytemp);

  /* Set minimum increment based on uround and norm of f. */
  srur = RSqrt(uround);
  fnorm = N_VWrmsNorm(fy, ewt);
  minInc = (fnorm != ZERO) ?
           (MIN_INC_MULT * ABS(h) * uround * N * fnorm) : ONE;

  /* Set bandwidth and number of column groups for band differencing. */
  width = ml + mu + 1;
  ngroups = MIN(width, N);
  
  for (group = 1; group <= ngroups; group++) {
    
    /* Increment all y_j in group. */
    for(j = group-1; j < N; j += width) {
      inc = MAX(srur*ABS(y_data[j]), minInc/ewt_data[j]);
      ytemp_data[j] += inc;
    }

    /* Evaluate f with incremented y. */

    retval = f(t, ytemp, ftemp, f_data);
    nfeBP++;
    if (retval != 0) return(retval);

    /* Restore ytemp, then form and load difference quotients. */
    for (j = group-1; j < N; j += width) {
      ytemp_data[j] = y_data[j];
      col_j = BAND_COL(savedJ,j);
      inc = MAX(srur*ABS(y_data[j]), minInc/ewt_data[j]);
      inc_inv = ONE/inc;
      i1 = MAX(0, j-mu);
      i2 = MIN(j+ml, N-1);
      for (i=i1; i <= i2; i++)
        BAND_COL_ELEM(col_j,i,j) =
          inc_inv * (ftemp_data[i] - fy_data[i]);
    }
  }

  return(0);
}

/* 
 * ================================================================
 *
 *                   PART II - backward problems
 *
 * ================================================================
 */

/* Additional readability replacements */

#define bp_data_B   (ca_mem->ca_pmemB)

/*
 * CVBandPrecAllocB, CVBPSp*B
 *
 * Wrappers for the backward phase around the corresponding 
 * CVODES functions
 */

int CVBandPrecAllocB(void *cvadj_mem, long int nB, 
                     long int muB, long int mlB)
{
  CVadjMem ca_mem;
  CVodeMem cvB_mem;
  void *bp_dataB;

  if (cvadj_mem == NULL) {
    CVProcessError(NULL, CVBANDPRE_ADJMEM_NULL, "CVBANDPRE", "CVBandPrecAlloc", MSGBP_CAMEM_NULL);
    return(CVBANDPRE_ADJMEM_NULL);
  }
  ca_mem = (CVadjMem) cvadj_mem;

  cvB_mem = ca_mem->cvb_mem;

  bp_dataB = CVBandPrecAlloc(cvB_mem, nB, muB, mlB);
  if (bp_dataB == NULL) {
    CVProcessError(cvB_mem, CVBANDPRE_MEM_FAIL, "CVBANDPRE", "CVBandPrecAlloc", MSGBP_MEM_FAIL);
    return(CVBANDPRE_MEM_FAIL);
  }

  bp_data_B = bp_dataB;

  return(CVBANDPRE_SUCCESS);

}

int CVBPSptfqmrB(void *cvadj_mem, int pretypeB, int maxlB)
{
  CVadjMem ca_mem;
  CVodeMem cvB_mem;
  int flag;

  if (cvadj_mem == NULL) {
    CVProcessError(NULL, CVBANDPRE_ADJMEM_NULL, "CVBANDPRE", "CVBPSptfqmrAlloc", MSGBP_CAMEM_NULL);
    return(CVBANDPRE_ADJMEM_NULL);
  }
  ca_mem = (CVadjMem) cvadj_mem;

  cvB_mem = ca_mem->cvb_mem;
  
  flag = CVBPSptfqmr(cvB_mem, pretypeB, maxlB, bp_data_B);

  return(flag);
}

int CVBPSpbcgB(void *cvadj_mem, int pretypeB, int maxlB)
{
  CVadjMem ca_mem;
  CVodeMem cvB_mem;
  int flag;

  if (cvadj_mem == NULL) {
    CVProcessError(NULL, CVBANDPRE_ADJMEM_NULL, "CVBANDPRE", "CVBPSptbcgAlloc", MSGBP_CAMEM_NULL);
    return(CVBANDPRE_ADJMEM_NULL);
  }
  ca_mem = (CVadjMem) cvadj_mem;

  cvB_mem = ca_mem->cvb_mem;
  
  flag = CVBPSpbcg(cvB_mem, pretypeB, maxlB, bp_data_B);

  return(flag);
}

int CVBPSpgmrB(void *cvadj_mem, int pretypeB, int maxlB)
{
  CVadjMem ca_mem;
  CVodeMem cvB_mem;
  int flag;

  if (cvadj_mem == NULL) {
    CVProcessError(NULL, CVBANDPRE_ADJMEM_NULL, "CVBANDPRE", "CVBPSpgmrAlloc", MSGBP_CAMEM_NULL);
    return(CVBANDPRE_ADJMEM_NULL);
  }
  ca_mem = (CVadjMem) cvadj_mem;

  cvB_mem = ca_mem->cvb_mem;
  
  flag = CVBPSpgmr(cvB_mem, pretypeB, maxlB, bp_data_B);

  return(flag);
}


void CVBandPrecFreeB(void *cvadj_mem)
{
  CVadjMem ca_mem;

  if (cvadj_mem == NULL) return;
  ca_mem = (CVadjMem) cvadj_mem;

  CVBandPrecFree(&bp_data_B);
}
```
