## Supplementary scripts for "A multi-scale *in silico* mouse model for insulin resistance and humanoid type 2 diabetes": cvodes_bandpre_impl.h

```
/*
 * -----------------------------------------------------------------
 * $Revision: 749 $
 * $Date: 2011-06-22 14:49:10 +0200 (Wed, 22 Jun 2011) $
 * ----------------------------------------------------------------- 
 * Programmer(s): Radu Serban @ LLNL
 * -----------------------------------------------------------------
 * Copyright (c) 2005, The Regents of the University of California.
 * Produced at the Lawrence Livermore National Laboratory.
 * All rights reserved.
 * For details, see the LICENSE file.
 * -----------------------------------------------------------------
 * Implementation header file for the CVBANDPRE module.
 * -----------------------------------------------------------------
 */

#ifndef _CVSBANDPRE_IMPL_H
#define _CVSBANDPRE_IMPL_H

#ifdef __cplusplus  /* wrapper to enable C++ usage */
extern "C" {
#endif

#include <cvodes/cvodes_bandpre.h>
#include <sundials/sundials_band.h>

  /*
   * -----------------------------------------------------------------
   * Type: CVBandPrecData
   * -----------------------------------------------------------------
   */

  typedef struct {

    /* Data set by user in CVBandPrecAlloc */

    long int N;
    long int ml, mu;

    /* Data set by CVBandPrecSetup */

    BandMat savedJ;
    BandMat savedP;
    long int *pivots;

    /* Rhs calls */

    long int nfeBP;

    /* Pointer to cvode_mem */

    void *cvode_mem;

  } *CVBandPrecData;

  /*
   * -----------------------------------------------------------------
   * CVBANDPRE error messages
   * -----------------------------------------------------------------
   */

#define MSGBP_CVMEM_NULL "Integrator memory is NULL."
#define MSGBP_MEM_FAIL "A memory request failed."
#define MSGBP_BAD_NVECTOR "A required vector operation is not implemented."
#define MSGBP_PDATA_NULL "CVBANDPRE memory is NULL."
#define MSGBP_RHSFUNC_FAILED "The right-hand side routine failed in an unrecoverable manner."

#define MSGBP_CAMEM_NULL "cvadj_mem = NULL illegal."

#ifdef __cplusplus
}
#endif

#endif
```
