## Supplementary scripts for "A multi-scale *in silico* mouse model for insulin resistance and humanoid type 2 diabetes": cvodes_bbdpre.c

```
/*
 * -----------------------------------------------------------------
 * $Revision: 749 $
 * $Date: 2011-06-22 14:49:10 +0200 (Wed, 22 Jun 2011) $
 * ----------------------------------------------------------------- 
 * Programmer(s): Radu Serban and Aaron Collier @ LLNL
 * -----------------------------------------------------------------
 * Copyright (c) 2005, The Regents of the University of California.
 * Produced at the Lawrence Livermore National Laboratory.
 * All rights reserved.
 * For details, see the LICENSE file.
 * -----------------------------------------------------------------
 * This file contains implementations of routines for a
 * band-block-diagonal preconditioner, i.e. a block-diagonal
 * matrix with banded blocks, for use with CVODE, a CVSPILS linear
 * solver, and the parallel implementation of NVECTOR.
 * -----------------------------------------------------------------
 */

#include <stdio.h>
#include <stdlib.h>

#include "cvodes_impl.h"
#include "cvodes_bbdpre_impl.h"

#include <cvodes/cvodes_sptfqmr.h>
#include <cvodes/cvodes_spbcgs.h>
#include <cvodes/cvodes_spgmr.h>

#include <sundials/sundials_math.h>

#define MIN_INC_MULT RCONST(1000.0)
#define ZERO         RCONST(0.0)
#define ONE          RCONST(1.0)

/* Prototypes of functions CVBBDPrecSetup and CVBBDPrecSolve */

static int CVBBDPrecSetup(realtype t, N_Vector y, N_Vector fy, 
                          booleantype jok, booleantype *jcurPtr, 
                          realtype gamma, void *bbd_data, 
                          N_Vector tmp1, N_Vector tmp2, N_Vector tmp3);

static int CVBBDPrecSolve(realtype t, N_Vector y, N_Vector fy, 
                          N_Vector r, N_Vector z, 
                          realtype gamma, realtype delta,
                          int lr, void *bbd_data, N_Vector tmp);

/* Wrapper functions for adjoint code */

static int CVAgloc(long int NlocalB, realtype t, N_Vector yB, N_Vector gB, 
                   void *cvadj_mem);

static int CVAcfn(long int NlocalB, realtype t, N_Vector yB,
                  void *cvadj_mem);

/* Prototype for difference quotient Jacobian calculation routine */

static int CVBBDDQJac(CVBBDPrecData pdata, realtype t, 
                      N_Vector y, N_Vector gy, 
                      N_Vector ytemp, N_Vector gtemp);


/* 
 * ================================================================
 *
 *                   PART I - forward problems
 *
 * ================================================================
 */

/* Redability replacements */

#define uround   (cv_mem->cv_uround)
#define vec_tmpl (cv_mem->cv_tempv)

/*
 * -----------------------------------------------------------------
 * User-Callable Functions: malloc, reinit and free
 * -----------------------------------------------------------------
 */

void *CVBBDPrecAlloc(void *cvode_mem, long int Nlocal, 
                     long int mudq, long int mldq,
                     long int mukeep, long int mlkeep, 
                     realtype dqrely, 
                     CVLocalFn gloc, CVCommFn cfn)
{
  CVodeMem cv_mem;
  CVBBDPrecData pdata;
  long int muk, mlk, storage_mu;

  if (cvode_mem == NULL) {
    CVProcessError(NULL, 0, "CVBBDPRE", "CVBBDPrecAlloc", MSGBBDP_CVMEM_NULL);
    return(NULL);
  }
  cv_mem = (CVodeMem) cvode_mem;

  /* Test if the NVECTOR package is compatible with the BLOCK BAND preconditioner */
  if(vec_tmpl->ops->nvgetarraypointer == NULL) {
    CVProcessError(cv_mem, 0, "CVBBDPRE", "CVBBDPrecAlloc", MSGBBDP_BAD_NVECTOR);
    return(NULL);
  }

  /* Allocate data memory */
  pdata = NULL;
  pdata = (CVBBDPrecData) malloc(sizeof *pdata);  
  if (pdata == NULL) {
    CVProcessError(cv_mem, 0, "CVBBDPRE", "CVBBDPrecAlloc", MSGBBDP_MEM_FAIL);
    return(NULL);
  }

  /* Set pointers to gloc and cfn; load half-bandwidths */
  pdata->cvode_mem = cvode_mem;
  pdata->gloc = gloc;
  pdata->cfn = cfn;
  pdata->mudq = MIN(Nlocal-1, MAX(0,mudq));
  pdata->mldq = MIN(Nlocal-1, MAX(0,mldq));
  muk = MIN(Nlocal-1, MAX(0,mukeep));
  mlk = MIN(Nlocal-1, MAX(0,mlkeep));
  pdata->mukeep = muk;
  pdata->mlkeep = mlk;

  /* Allocate memory for saved Jacobian */
  pdata->savedJ = BandAllocMat(Nlocal, muk, mlk, muk);
  if (pdata->savedJ == NULL) { 
    free(pdata); pdata = NULL; 
    CVProcessError(cv_mem, 0, "CVBBDPRE", "CVBBDPrecAlloc", MSGBBDP_MEM_FAIL);
    return(NULL); 
  }

  /* Allocate memory for preconditioner matrix */
  storage_mu = MIN(Nlocal-1, muk + mlk);
  pdata->savedP = NULL;
  pdata->savedP = BandAllocMat(Nlocal, muk, mlk, storage_mu);
  if (pdata->savedP == NULL) {
    BandFreeMat(pdata->savedJ);
    free(pdata); pdata = NULL;
    CVProcessError(cv_mem, 0, "CVBBDPRE", "CVBBDPrecAlloc", MSGBBDP_MEM_FAIL);
    return(NULL);
  }
  /* Allocate memory for pivots */
  pdata->pivots = NULL;
  pdata->pivots = BandAllocPiv(Nlocal);
  if (pdata->savedJ == NULL) {
    BandFreeMat(pdata->savedP);
    BandFreeMat(pdata->savedJ);
    free(pdata); pdata = NULL;
    CVProcessError(cv_mem, 0, "CVBBDPRE", "CVBBDPrecAlloc", MSGBBDP_MEM_FAIL);
    return(NULL);
  }

  /* Set pdata->dqrely based on input dqrely (0 implies default). */
  pdata->dqrely = (dqrely > ZERO) ? dqrely : RSqrt(uround);

  /* Store Nlocal to be used in CVBBDPrecSetup */
  pdata->n_local = Nlocal;

  /* Set work space sizes and initialize nge */
  pdata->rpwsize = Nlocal*(muk + 2*mlk + storage_mu + 2);
  pdata->ipwsize = Nlocal;
  pdata->nge = 0;

  return((void *)pdata);
}

int CVBBDSptfqmr(void *cvode_mem, int pretype, int maxl, void *bbd_data)
{
  CVodeMem cv_mem;
  int flag;

  flag = CVSptfqmr(cvode_mem, pretype, maxl);
  if(flag != CVSPILS_SUCCESS) return(flag);

  cv_mem = (CVodeMem) cvode_mem;

  if (bbd_data == NULL) {
    CVProcessError(cv_mem, CVBBDPRE_PDATA_NULL, "CVBBDPRE", "CVBBDSptfqmr", MSGBBDP_PDATA_NULL);
    return(CVBBDPRE_PDATA_NULL);
  } 

  flag = CVSpilsSetPreconditioner(cvode_mem, CVBBDPrecSetup, CVBBDPrecSolve, bbd_data);
  if(flag != CVSPILS_SUCCESS) return(flag);

  return(CVSPILS_SUCCESS);
}

int CVBBDSpbcg(void *cvode_mem, int pretype, int maxl, void *bbd_data)
{
  CVodeMem cv_mem;
  int flag;

  flag = CVSpbcg(cvode_mem, pretype, maxl);
  if(flag != CVSPILS_SUCCESS) return(flag);

  cv_mem = (CVodeMem) cvode_mem;

  if (bbd_data == NULL) {
    CVProcessError(cv_mem, CVBBDPRE_PDATA_NULL, "CVBBDPRE", "CVBBDSpbcg", MSGBBDP_PDATA_NULL);
    return(CVBBDPRE_PDATA_NULL);
  } 

  flag = CVSpilsSetPreconditioner(cvode_mem, CVBBDPrecSetup, CVBBDPrecSolve, bbd_data);
  if(flag != CVSPILS_SUCCESS) return(flag);

  return(CVSPILS_SUCCESS);
}

int CVBBDSpgmr(void *cvode_mem, int pretype, int maxl, void *bbd_data)
{
  CVodeMem cv_mem;
  int flag;

  flag = CVSpgmr(cvode_mem, pretype, maxl);
  if(flag != CVSPILS_SUCCESS) return(flag);

  cv_mem = (CVodeMem) cvode_mem;

  if (bbd_data == NULL) {
    CVProcessError(cv_mem, CVBBDPRE_PDATA_NULL, "CVBBDPRE", "CVBBDSpgmr", MSGBBDP_PDATA_NULL);
    return(CVBBDPRE_PDATA_NULL);
  } 

  flag = CVSpilsSetPreconditioner(cvode_mem, CVBBDPrecSetup, CVBBDPrecSolve, bbd_data);
  if(flag != CVSPILS_SUCCESS) return(flag);

  return(CVSPILS_SUCCESS);
}

int CVBBDPrecReInit(void *bbd_data, 
                    long int mudq, long int mldq, 
                    realtype dqrely, 
                    CVLocalFn gloc, CVCommFn cfn)
{
  CVBBDPrecData pdata;
  CVodeMem cv_mem;
  long int Nlocal;

  if (bbd_data == NULL) {
    CVProcessError(NULL, CVBBDPRE_PDATA_NULL, "CVBBDPRE", "CVBBDPrecReInit", MSGBBDP_PDATA_NULL);
    return(CVBBDPRE_PDATA_NULL);
  } 

  pdata  = (CVBBDPrecData) bbd_data;
  cv_mem = (CVodeMem) pdata->cvode_mem;

  /* Set pointers to gloc and cfn; load half-bandwidths */
  pdata->gloc = gloc;
  pdata->cfn = cfn;
  Nlocal = pdata->n_local;
  pdata->mudq = MIN(Nlocal-1, MAX(0,mudq));
  pdata->mldq = MIN(Nlocal-1, MAX(0,mldq));

  /* Set pdata->dqrely based on input dqrely (0 implies default). */
  pdata->dqrely = (dqrely > ZERO) ? dqrely : RSqrt(uround);

  /* Re-initialize nge */
  pdata->nge = 0;

  return(CVBBDPRE_SUCCESS);
}

void CVBBDPrecFree(void **bbd_data)
{
  CVBBDPrecData pdata;
  
  if (*bbd_data == NULL) return;

  pdata = (CVBBDPrecData) (*bbd_data);
  BandFreeMat(pdata->savedJ);
  BandFreeMat(pdata->savedP);
  BandFreePiv(pdata->pivots);

  free(*bbd_data);
  *bbd_data = NULL;

}

int CVBBDPrecGetWorkSpace(void *bbd_data, long int *lenrwBBDP, long int *leniwBBDP)
{
  CVBBDPrecData pdata;

  if (bbd_data == NULL) {
    CVProcessError(NULL, CVBBDPRE_PDATA_NULL, "CVBBDPRE", "CVBBDPrecGetWorkSpace", MSGBBDP_PDATA_NULL);    
    return(CVBBDPRE_PDATA_NULL);
  } 

  pdata = (CVBBDPrecData) bbd_data;

  *lenrwBBDP = pdata->rpwsize;
  *leniwBBDP = pdata->ipwsize;

  return(CVBBDPRE_SUCCESS);
}

int CVBBDPrecGetNumGfnEvals(void *bbd_data, long int *ngevalsBBDP)
{
  CVBBDPrecData pdata;

  if (bbd_data == NULL) {
    CVProcessError(NULL, CVBBDPRE_PDATA_NULL, "CVBBDPRE", "CVBBDPrecGetNumGfnEvals", MSGBBDP_PDATA_NULL);
    return(CVBBDPRE_PDATA_NULL);
  } 

  pdata = (CVBBDPrecData) bbd_data;

  *ngevalsBBDP = pdata->nge;

  return(CVBBDPRE_SUCCESS);
}

/*
 * -----------------------------------------------------------------
 * CVBBDPrecGetReturnFlagName
 * -----------------------------------------------------------------
 */

char *CVBBDPrecGetReturnFlagName(int flag)
{
  char *name;

  name = (char *)malloc(30*sizeof(char));

  switch(flag) {
  case CVBBDPRE_SUCCESS:
    sprintf(name,"CVBBDPRE_SUCCESS");
    break; 
  case CVBBDPRE_PDATA_NULL:
    sprintf(name,"CVBBDPRE_PDATA_NULL");
    break;
  case CVBBDPRE_FUNC_UNRECVR:
    sprintf(name,"CVBBDPRE_FUNC_UNRECVR");
    break;
  case CVBBDPRE_ADJMEM_NULL:
    sprintf(name,"CVBBDPRE_ADJMEM_NULL");
    break;
  case CVBBDPRE_PDATAB_NULL:
    sprintf(name,"CVBBDPRE_PDATAB_NULL");
    break;
  case CVBBDPRE_MEM_FAIL:
    sprintf(name,"CVBBDPRE_MEM_FAIL");
    break;
  default:
    sprintf(name,"NONE");
  }

  return(name);
}

/* Readability Replacements */

#define Nlocal (pdata->n_local)
#define mudq   (pdata->mudq)
#define mldq   (pdata->mldq)
#define mukeep (pdata->mukeep)
#define mlkeep (pdata->mlkeep)
#define dqrely (pdata->dqrely)
#define gloc   (pdata->gloc)
#define cfn    (pdata->cfn)
#define savedJ (pdata->savedJ)
#define savedP (pdata->savedP)
#define pivots (pdata->pivots)
#define nge    (pdata->nge)

/*
 * -----------------------------------------------------------------
 * Function : CVBBDPrecSetup                                      
 * -----------------------------------------------------------------
 * CVBBDPrecSetup generates and factors a banded block of the
 * preconditioner matrix on each processor, via calls to the
 * user-supplied gloc and cfn functions. It uses difference
 * quotient approximations to the Jacobian elements.
 *
 * CVBBDPrecSetup calculates a new J,if necessary, then calculates
 * P = I - gamma*J, and does an LU factorization of P.
 *
 * The parameters of CVBBDPrecSetup used here are as follows:
 *
 * t       is the current value of the independent variable.
 *
 * y       is the current value of the dependent variable vector,
 *         namely the predicted value of y(t).
 *
 * fy      is the vector f(t,y).
 *
 * jok     is an input flag indicating whether Jacobian-related
 *         data needs to be recomputed, as follows:
 *           jok == FALSE means recompute Jacobian-related data
 *                  from scratch.
 *           jok == TRUE  means that Jacobian data from the
 *                  previous CVBBDPrecon call can be reused
 *                  (with the current value of gamma).
 *         A CVBBDPrecon call with jok == TRUE should only occur
 *         after a call with jok == FALSE.
 *
 * jcurPtr is a pointer to an output integer flag which is
 *         set by CVBBDPrecon as follows:
 *           *jcurPtr = TRUE if Jacobian data was recomputed.
 *           *jcurPtr = FALSE if Jacobian data was not recomputed,
 *                      but saved data was reused.
 *
 * gamma   is the scalar appearing in the Newton matrix.
 *
 * bbd_data  is a pointer to user data - the same as the P_data
 *           parameter passed to CVSp*. For CVBBDPrecon,
 *           this should be of type CVBBDData.
 *
 * tmp1, tmp2, and tmp3 are pointers to memory allocated
 *           for NVectors which are be used by CVBBDPrecSetup
 *           as temporary storage or work space.
 *
 * Return value:
 * The value returned by this CVBBDPrecSetup function is the int
 *   0  if successful,
 *   1  for a recoverable error (step will be retried).
 * -----------------------------------------------------------------
 */

static int CVBBDPrecSetup(realtype t, N_Vector y, N_Vector fy, 
                          booleantype jok, booleantype *jcurPtr, 
                          realtype gamma, void *bbd_data, 
                          N_Vector tmp1, N_Vector tmp2, N_Vector tmp3)
{
  long int ier;
  CVBBDPrecData pdata;
  CVodeMem cv_mem;
  int retval;

  pdata = (CVBBDPrecData) bbd_data;

  cv_mem = (CVodeMem) pdata->cvode_mem;

  if (jok) {

    /* If jok = TRUE, use saved copy of J */
    *jcurPtr = FALSE;
    BandCopy(savedJ, savedP, mukeep, mlkeep);

  } else {

    /* Otherwise call CVBBDDQJac for new J value */
    *jcurPtr = TRUE;
    BandZero(savedJ);

    retval = CVBBDDQJac(pdata, t, y, tmp1, tmp2, tmp3);
    if (retval < 0) {
      CVProcessError(cv_mem, CVBBDPRE_FUNC_UNRECVR, "CVBBDPRE", "CVBBDPrecSetup", MSGBBDP_FUNC_FAILED);
      return(-1);
    }
    if (retval > 0) {
      return(1);
    }

    BandCopy(savedJ, savedP, mukeep, mlkeep);

  }
  
  /* Scale and add I to get P = I - gamma*J */
  BandScale(-gamma, savedP);
  BandAddI(savedP);
 
  /* Do LU factorization of P in place */
  ier = BandGBTRF(savedP, pivots);
 
  /* Return 0 if the LU was complete; otherwise return 1 */
  if (ier > 0) return(1);
  return(0);
}

/*
 * -----------------------------------------------------------------
 * Function : CVBBDPrecSolve
 * -----------------------------------------------------------------
 * CVBBDPrecSolve solves a linear system P z = r, with the
 * band-block-diagonal preconditioner matrix P generated and
 * factored by CVBBDPrecSetup.
 *
 * The parameters of CVBBDPrecSolve used here are as follows:
 *
 * r      is the right-hand side vector of the linear system.
 *
 * bbd_data is a pointer to the preconditioner data returned by
 *          CVBBDPrecAlloc.
 *
 * z      is the output vector computed by CVBBDPrecSolve.
 *
 * The value returned by the CVBBDPrecSolve function is always 0,
 * indicating success.
 * -----------------------------------------------------------------
 */

static int CVBBDPrecSolve(realtype t, N_Vector y, N_Vector fy, 
                          N_Vector r, N_Vector z, 
                          realtype gamma, realtype delta,
                          int lr, void *bbd_data, N_Vector tmp)
{
  CVBBDPrecData pdata;
  realtype *zd;

  pdata = (CVBBDPrecData) bbd_data;

  /* Copy r to z, then do backsolve and return */
  N_VScale(ONE, r, z);
  
  zd = N_VGetArrayPointer(z);

  BandGBTRS(savedP, pivots, zd);

  return(0);
}

#define ewt    (cv_mem->cv_ewt)
#define h      (cv_mem->cv_h)
#define f_data (cv_mem->cv_f_data)

/*
 * -----------------------------------------------------------------
 * Function : CVBBDDQJac
 * -----------------------------------------------------------------
 * This routine generates a banded difference quotient approximation
 * to the local block of the Jacobian of g(t,y). It assumes that a
 * band matrix of type BandMat is stored columnwise, and that elements
 * within each column are contiguous. All matrix elements are generated
 * as difference quotients, by way of calls to the user routine gloc.
 * By virtue of the band structure, the number of these calls is
 * bandwidth + 1, where bandwidth = mldq + mudq + 1.
 * But the band matrix kept has bandwidth = mlkeep + mukeep + 1.
 * This routine also assumes that the local elements of a vector are
 * stored contiguously.
 * -----------------------------------------------------------------
 */

static int CVBBDDQJac(CVBBDPrecData pdata, realtype t, 
                      N_Vector y, N_Vector gy, 
                      N_Vector ytemp, N_Vector gtemp)
{
  CVodeMem cv_mem;
  realtype gnorm, minInc, inc, inc_inv;
  long int group, i, j, width, ngroups, i1, i2;
  realtype *y_data, *ewt_data, *gy_data, *gtemp_data, *ytemp_data, *col_j;
  int retval;

  cv_mem = (CVodeMem) pdata->cvode_mem;

  /* Load ytemp with y = predicted solution vector */
  N_VScale(ONE, y, ytemp);

  /* Call cfn and gloc to get base value of g(t,y) */
  if (cfn != NULL) {
    retval = cfn(Nlocal, t, y, f_data);
    if (retval != 0) return(retval);
  }

  retval = gloc(Nlocal, t, ytemp, gy, f_data);
  nge++;
  if (retval != 0) return(retval);

  /* Obtain pointers to the data for various vectors */
  y_data     =  N_VGetArrayPointer(y);
  gy_data    =  N_VGetArrayPointer(gy);
  ewt_data   =  N_VGetArrayPointer(ewt);
  ytemp_data =  N_VGetArrayPointer(ytemp);
  gtemp_data =  N_VGetArrayPointer(gtemp);

  /* Set minimum increment based on uround and norm of g */
  gnorm = N_VWrmsNorm(gy, ewt);
  minInc = (gnorm != ZERO) ?
           (MIN_INC_MULT * ABS(h) * uround * Nlocal * gnorm) : ONE;

  /* Set bandwidth and number of column groups for band differencing */
  width = mldq + mudq + 1;
  ngroups = MIN(width, Nlocal);

  /* Loop over groups */  
  for (group=1; group <= ngroups; group++) {
    
    /* Increment all y_j in group */
    for(j=group-1; j < Nlocal; j+=width) {
      inc = MAX(dqrely*ABS(y_data[j]), minInc/ewt_data[j]);
      ytemp_data[j] += inc;
    }

    /* Evaluate g with incremented y */
    retval = gloc(Nlocal, t, ytemp, gtemp, f_data);
    nge++;
    if (retval != 0) return(retval);

    /* Restore ytemp, then form and load difference quotients */
    for (j=group-1; j < Nlocal; j+=width) {
      ytemp_data[j] = y_data[j];
      col_j = BAND_COL(savedJ,j);
      inc = MAX(dqrely*ABS(y_data[j]), minInc/ewt_data[j]);
      inc_inv = ONE/inc;
      i1 = MAX(0, j-mukeep);
      i2 = MIN(j+mlkeep, Nlocal-1);
      for (i=i1; i <= i2; i++)
        BAND_COL_ELEM(col_j,i,j) =
          inc_inv * (gtemp_data[i] - gy_data[i]);
    }
  }

  return(0);
}


/* 
 * ================================================================
 *
 *                   PART II - backward problems
 *
 * ================================================================
 */


/* Additional readability replacements */

#define ytmp        (ca_mem->ca_ytmp)
#define f_data_B    (ca_mem->ca_f_dataB)
#define getY        (ca_mem->ca_getY)
#define pmemB       (ca_mem->ca_pmemB)

#define bbd_data_B  (cvbbdB_mem->bbd_dataB)
#define gloc_B      (cvbbdB_mem->glocB)
#define cfn_B       (cvbbdB_mem->cfnB)


/*
 * CVBBDPrecAllocB, CVBPSp*B
 *
 * Wrappers for the backward phase around the corresponding 
 * CVODES functions
 */

int CVBBDPrecAllocB(void *cvadj_mem, long int NlocalB, 
                    long int mudqB, long int mldqB, 
                    long int mukeepB, long int mlkeepB, 
                    realtype dqrelyB,
                    CVLocalFnB glocB, CVCommFnB cfnB)
{
  CVadjMem ca_mem;
  CVBBDPrecDataB cvbbdB_mem;
  CVodeMem cvB_mem;
  void *bbd_dataB;

  if (cvadj_mem == NULL) {
    CVProcessError(NULL, CVBBDPRE_ADJMEM_NULL, "CVBBDPRE", "CVBBDPrecAllocB", MSGBBDP_CAMEM_NULL);
    return(CVBBDPRE_ADJMEM_NULL);
  }
  ca_mem = (CVadjMem) cvadj_mem;

  cvB_mem = ca_mem->cvb_mem;

  /* Get memory for CVBBDPrecDataB */
  cvbbdB_mem = NULL;
  cvbbdB_mem = (CVBBDPrecDataB) malloc(sizeof(* cvbbdB_mem));
  if (cvbbdB_mem == NULL) {
    CVProcessError(cvB_mem, CVBBDPRE_MEM_FAIL, "CVBBDPRE", "CVBBDPrecAllocB", MSGBBDP_MEM_FAIL);
    return(CVBBDPRE_MEM_FAIL);
  }

  gloc_B = glocB;
  cfn_B  = cfnB;

  bbd_dataB = CVBBDPrecAlloc(cvB_mem, NlocalB, 
                             mudqB, mldqB,
                             mukeepB, mlkeepB, 
                             dqrelyB, 
                             CVAgloc, CVAcfn);
  if (bbd_dataB == NULL) {
    free(cvbbdB_mem); cvbbdB_mem = NULL;

    return(CVBBDPRE_MEM_FAIL);
  }

  bbd_data_B = bbd_dataB;

  /* attach pmemB */
  pmemB = cvbbdB_mem;

  return(CVBBDPRE_SUCCESS);

}

int CVBBDSptfqmrB(void *cvadj_mem, int pretypeB, int maxlB)
{

  CVadjMem ca_mem;
  CVBBDPrecDataB cvbbdB_mem;
  CVodeMem cvB_mem;
  int flag;
  
  if (cvadj_mem == NULL) {
    CVProcessError(NULL, CVBBDPRE_ADJMEM_NULL, "CVBBDPRE", "CVBBDSptfqmrB", MSGBBDP_CAMEM_NULL);
    return(CVBBDPRE_ADJMEM_NULL);
  }
  ca_mem = (CVadjMem) cvadj_mem;
  
  cvB_mem = ca_mem->cvb_mem;
  
  if (pmemB == NULL) {
    CVProcessError(cvB_mem, CVBBDPRE_PDATAB_NULL, "CVBBDPRE", "CVBBDSptfqmrB", MSGBBDP_PDATAB_NULL);
    return(CVBBDPRE_PDATAB_NULL);
  }
  cvbbdB_mem = (CVBBDPrecDataB) pmemB;

  flag = CVBBDSptfqmr(cvB_mem, pretypeB, maxlB, bbd_data_B);

  return(flag);

}

int CVBBDSpbcgB(void *cvadj_mem, int pretypeB, int maxlB)
{

  CVadjMem ca_mem;
  CVBBDPrecDataB cvbbdB_mem;
  CVodeMem cvB_mem;
  int flag;
  
  if (cvadj_mem == NULL) {
    CVProcessError(NULL, CVBBDPRE_ADJMEM_NULL, "CVBBDPRE", "CVBBDSpbcgB", MSGBBDP_CAMEM_NULL);
    return(CVBBDPRE_ADJMEM_NULL);
  }
  ca_mem = (CVadjMem) cvadj_mem;
  
  cvB_mem = ca_mem->cvb_mem;
  
  if (pmemB == NULL) {
    CVProcessError(cvB_mem, CVBBDPRE_PDATAB_NULL, "CVBBDPRE", "CVBBDSpbcgB", MSGBBDP_PDATAB_NULL);
    return(CVBBDPRE_PDATAB_NULL);
  }
  cvbbdB_mem = (CVBBDPrecDataB) pmemB;

  flag = CVBBDSpbcg(cvB_mem, pretypeB, maxlB, bbd_data_B);

  return(flag);

}

int CVBBDSpgmrB(void *cvadj_mem, int pretypeB, int maxlB)
{

  CVadjMem ca_mem;
  CVBBDPrecDataB cvbbdB_mem;
  CVodeMem cvB_mem;
  int flag;
  
  if (cvadj_mem == NULL) {
    CVProcessError(NULL, CVBBDPRE_ADJMEM_NULL, "CVBBDPRE", "CVBBDSpgmrB", MSGBBDP_CAMEM_NULL);
    return(CV_ADJMEM_NULL);
  }
  ca_mem = (CVadjMem) cvadj_mem;
  
  cvB_mem = ca_mem->cvb_mem;
  
  if (pmemB == NULL) {
    CVProcessError(cvB_mem, CVBBDPRE_PDATAB_NULL, "CVBBDPRE", "CVBBDSpgmrB", MSGBBDP_PDATAB_NULL);
    return(CVBBDPRE_PDATAB_NULL);
  }
  cvbbdB_mem = (CVBBDPrecDataB) pmemB;

  flag = CVBBDSpgmr(cvB_mem, pretypeB, maxlB, bbd_data_B);

  return(flag);

}

int CVBBDPrecReInitB(void *cvadj_mem, long int mudqB, long int mldqB,
                     realtype dqrelyB, CVLocalFnB glocB, CVCommFnB cfnB)
{
  CVadjMem ca_mem;
  CVodeMem cvB_mem;
  CVBBDPrecDataB cvbbdB_mem;
  int flag;

  if (cvadj_mem == NULL) {
    CVProcessError(NULL, CVBBDPRE_ADJMEM_NULL, "CVBBDPRE", "CVBBDPrecReInitB", MSGBBDP_CAMEM_NULL);
    return(CVBBDPRE_ADJMEM_NULL);
  }
  ca_mem = (CVadjMem) cvadj_mem;
  
  cvB_mem = ca_mem->cvb_mem;

  if (pmemB == NULL) {
    CVProcessError(cvB_mem, CVBBDPRE_PDATAB_NULL, "CVBBDPRE", "CVBBDPrecReInitB", MSGBBDP_PDATAB_NULL);
    return(CVBBDPRE_PDATAB_NULL);
  }
  cvbbdB_mem = (CVBBDPrecDataB) pmemB;

  gloc_B = glocB;
  cfn_B  = cfnB;

  flag = CVBBDPrecReInit(bbd_data_B, mudqB, mldqB,
                         dqrelyB, CVAgloc, CVAcfn);

  return(flag);
}


void CVBBDPrecFreeB(void *cvadj_mem)
{
  CVadjMem ca_mem;
  CVBBDPrecDataB cvbbdB_mem;

  if (cvadj_mem == NULL) return;
  ca_mem = (CVadjMem) cvadj_mem;
  
  if (pmemB == NULL) return;
  cvbbdB_mem = (CVBBDPrecDataB) pmemB;

  CVBBDPrecFree(&bbd_data_B);
  
  free(pmemB); pmemB = NULL;
}


/*
 * CVAgloc
 *
 * This routine interfaces to the CVLocalFnB routine 
 * provided by the user.
 * NOTE: f_data actually contains cvadj_mem
 */

static int CVAgloc(long int NlocalB, realtype t, N_Vector yB, N_Vector gB, 
                   void *cvadj_mem)
{
  CVadjMem ca_mem;
  CVodeMem cvB_mem;
  CVBBDPrecDataB cvbbdB_mem;
  int retval, flag;

  ca_mem = (CVadjMem) cvadj_mem;
  cvB_mem = ca_mem->cvb_mem;
  cvbbdB_mem = (CVBBDPrecDataB) pmemB;

  /* Forward solution from interpolation */
  flag = getY(ca_mem, t, ytmp);
  if (flag != CV_SUCCESS) {
    CVProcessError(cvB_mem, -1, "CVBBDPRE", "CVAgloc", MSGBBDP_BAD_T);
    return(-1);
  } 

  /* Call user's adjoint glocB routine */
  retval = gloc_B(NlocalB, t, ytmp, yB, gB, f_data_B);

  return(retval);
}

/*
 * CVAcfn
 *
 * This routine interfaces to the CVCommFnB routine 
 * provided by the user.
 * NOTE: f_data actually contains cvadj_mem
 */

static int CVAcfn(long int NlocalB, realtype t, N_Vector yB,
                  void *cvadj_mem)
{
  CVadjMem ca_mem;
  CVodeMem cvB_mem;
  CVBBDPrecDataB cvbbdB_mem;
  int retval, flag;

  ca_mem = (CVadjMem) cvadj_mem;
  cvB_mem = ca_mem->cvb_mem;
  cvbbdB_mem = (CVBBDPrecDataB) pmemB;

  if (cfn_B == NULL) return(0);

  /* Forward solution from interpolation */
  flag = getY(ca_mem, t, ytmp);
  if (flag != CV_SUCCESS) {
    CVProcessError(cvB_mem, -1, "CVBBDPRE", "CVAcfn", MSGBBDP_BAD_T);
    return(-1);
  } 

  /* Call user's adjoint cfnB routine */
  retval = cfn_B(NlocalB, t, ytmp, yB, f_data_B);

  return(retval);
}
```
