## Supplementary scripts for "A multi-scale *in silico* mouse model for insulin resistance and humanoid type 2 diabetes": cvodes_bbdpre_impl.h

```
/*
 * -----------------------------------------------------------------
 * $Revision: 749 $
 * $Date: 2011-06-22 14:49:10 +0200 (Wed, 22 Jun 2011) $
 * ----------------------------------------------------------------- 
 * Programmer(s): Radu Serban @ LLNL
 * -----------------------------------------------------------------
 * Copyright (c) 2005, The Regents of the University of California.
 * Produced at the Lawrence Livermore National Laboratory.
 * All rights reserved.
 * For details, see the LICENSE file.
 * -----------------------------------------------------------------
 * Implementation header file for the CVBBDPRE module.
 * -----------------------------------------------------------------
 */

#ifndef _CVSBBDPRE_IMPL_H
#define _CVSBBDPRE_IMPL_H

#ifdef __cplusplus  /* wrapper to enable C++ usage */
extern "C" {
#endif

#include <cvodes/cvodes_bbdpre.h>
#include <sundials/sundials_band.h>

  /*
   * -----------------------------------------------------------------
   * Type: CVBBDPrecData
   * -----------------------------------------------------------------
   */

  typedef struct {

    /* passed by user to CVBBDPrecAlloc and used by PrecSetup/PrecSolve */

    long int mudq, mldq, mukeep, mlkeep;
    realtype dqrely;
    CVLocalFn gloc;
    CVCommFn cfn;

    /* set by CVBBDPrecSetup and used by CVBBDPrecSolve */

    BandMat savedJ;
    BandMat savedP;
    long int *pivots;

    /* set by CVBBDPrecAlloc and used by CVBBDPrecSetup */

    long int n_local;

    /* available for optional output */

    long int rpwsize;
    long int ipwsize;
    long int nge;

    /* pointer to cvode_mem */

    void *cvode_mem;

  } *CVBBDPrecData;


  /*
   * -----------------------------------------------------------------
   * Type: CVBBDPrecDataB
   * -----------------------------------------------------------------
   */

  typedef struct {

    /* BBD user functions (glocB and cfnB) for backward run */
    CVLocalFnB glocB;
    CVCommFnB  cfnB;
    
    /* BBD prec data */
    void *bbd_dataB;

  } *CVBBDPrecDataB;

  /*
   * -----------------------------------------------------------------
   * CVBBDPRE error messages
   * -----------------------------------------------------------------
   */

#define MSGBBDP_CVMEM_NULL  "Integrator memory is NULL."
#define MSGBBDP_MEM_FAIL    "A memory request failed."
#define MSGBBDP_BAD_NVECTOR "A required vector operation is not implemented."
#define MSGBBDP_PDATA_NULL  "CVBBDPRE memory is NULL."
#define MSGBBDP_FUNC_FAILED "The gloc or cfn routine failed in an unrecoverable manner."

#define MSGBBDP_CAMEM_NULL  "cvadj_mem = NULL illegal."
#define MSGBBDP_PDATAB_NULL "CVBBDPRE memory is NULL for the backward integration."
#define MSGBBDP_BAD_T       "Bad t for interpolation."


#ifdef __cplusplus
}
#endif

#endif
```
