## Supplementary scripts for "A multi-scale *in silico* mouse model for insulin resistance and humanoid type 2 diabetes": cvodes_dense.c

```
/*
 * -----------------------------------------------------------------
 * $Revision: 749 $
 * $Date: 2011-06-22 14:49:10 +0200 (Wed, 22 Jun 2011) $
 * ----------------------------------------------------------------- 
 * Programmer(s): Radu Serban @ LLNL
 * -----------------------------------------------------------------
 * Copyright (c) 2005, The Regents of the University of California.
 * Produced at the Lawrence Livermore National Laboratory.
 * All rights reserved.
 * For details, see the LICENSE file.
 * -----------------------------------------------------------------
 * This is the implementation file for the CVDENSE linear solver.
 * -----------------------------------------------------------------
 */

#include <stdio.h>
#include <stdlib.h>

#include "cvodes_dense_impl.h"
#include "cvodes_impl.h"

#include <sundials/sundials_math.h>

/* Other Constants */

#define MIN_INC_MULT RCONST(1000.0)
#define ZERO         RCONST(0.0)
#define ONE          RCONST(1.0)
#define TWO          RCONST(2.0)

/* CVDENSE linit, lsetup, lsolve, and lfree routines */
 
static int CVDenseInit(CVodeMem cv_mem);

static int CVDenseSetup(CVodeMem cv_mem, int convfail, N_Vector ypred,
                        N_Vector fpred, booleantype *jcurPtr, 
                        N_Vector vtemp1, N_Vector vtemp2, N_Vector vtemp3);

static int CVDenseSolve(CVodeMem cv_mem, N_Vector b, N_Vector weight,
                        N_Vector ycur, N_Vector fcur);

static void CVDenseFree(CVodeMem cv_mem);

/* CVDENSE lfreeB function */

static void CVDenseFreeB(CVadjMem ca_mem);

/* Wrapper function for adjoint code */

static int CVAdenseJac(long int nB, DenseMat JB, realtype t, 
                       N_Vector yB, N_Vector fyB, void *cvadj_mem,
                       N_Vector tmp1B, N_Vector tmp2B, N_Vector tmp3B);


/* CVDENSE DQJac routine */

static int CVDenseDQJac(long int n, DenseMat J, realtype t, 
                        N_Vector y, N_Vector fy, void *jac_data,
                        N_Vector tmp1, N_Vector tmp2, N_Vector tmp3);


/* 
 * ================================================================
 *
 *                   PART I - forward problems
 *
 * ================================================================
 */


/* Readability Replacements */

#define lmm       (cv_mem->cv_lmm)
#define f         (cv_mem->cv_f)
#define f_data    (cv_mem->cv_f_data)
#define uround    (cv_mem->cv_uround)
#define nst       (cv_mem->cv_nst)
#define tn        (cv_mem->cv_tn)
#define h         (cv_mem->cv_h)
#define gamma     (cv_mem->cv_gamma)
#define gammap    (cv_mem->cv_gammap)
#define gamrat    (cv_mem->cv_gamrat)
#define ewt       (cv_mem->cv_ewt)
#define linit     (cv_mem->cv_linit)
#define lsetup    (cv_mem->cv_lsetup)
#define lsolve    (cv_mem->cv_lsolve)
#define lfree     (cv_mem->cv_lfree)
#define lmem      (cv_mem->cv_lmem)
#define vec_tmpl     (cv_mem->cv_tempv)
#define setupNonNull (cv_mem->cv_setupNonNull)

#define n         (cvdense_mem->d_n)
#define jac       (cvdense_mem->d_jac)
#define M         (cvdense_mem->d_M)
#define pivots    (cvdense_mem->d_pivots)
#define savedJ    (cvdense_mem->d_savedJ)
#define nstlj     (cvdense_mem->d_nstlj)
#define nje       (cvdense_mem->d_nje)
#define nfeD      (cvdense_mem->d_nfeD)
#define J_data    (cvdense_mem->d_J_data)
#define last_flag (cvdense_mem->d_last_flag)

/*
 * -----------------------------------------------------------------
 * CVDense
 * -----------------------------------------------------------------
 * This routine initializes the memory record and sets various function
 * fields specific to the dense linear solver module.  CVDense first
 * calls the existing lfree routine if this is not NULL.  Then it sets
 * the cv_linit, cv_lsetup, cv_lsolve, cv_lfree fields in (*cvode_mem)
 * to be CVDenseInit, CVDenseSetup, CVDenseSolve, and CVDenseFree,
 * respectively.  It allocates memory for a structure of type
 * CVDenseMemRec and sets the cv_lmem field in (*cvode_mem) to the
 * address of this structure.  It sets setupNonNull in (*cvode_mem) to
 * TRUE, and the d_jac field to the default CVDenseDQJac.
 * Finally, it allocates memory for M, savedJ, and pivots.
 * The return value is SUCCESS = 0, or LMEM_FAIL = -1.
 *
 * NOTE: The dense linear solver assumes a serial implementation
 *       of the NVECTOR package. Therefore, CVDense will first 
 *       test for compatible a compatible N_Vector internal
 *       representation by checking that N_VGetArrayPointer and
 *       N_VSetArrayPointer exist.
 * -----------------------------------------------------------------
 */

int CVDense(void *cvode_mem, long int N)
{
  CVodeMem cv_mem;
  CVDenseMem cvdense_mem;

  /* Return immediately if cvode_mem is NULL */
  if (cvode_mem == NULL) {
    CVProcessError(NULL, CVDENSE_MEM_NULL, "CVDENSE", "CVDense", MSGDS_CVMEM_NULL);
    return(CVDENSE_MEM_NULL);
  }
  cv_mem = (CVodeMem) cvode_mem;

  /* Test if the NVECTOR package is compatible with the DENSE solver */
  if (vec_tmpl->ops->nvgetarraypointer == NULL ||
      vec_tmpl->ops->nvsetarraypointer == NULL) {
    CVProcessError(cv_mem, CVDENSE_ILL_INPUT, "CVDENSE", "CVDense", MSGDS_BAD_NVECTOR);
    return(CVDENSE_ILL_INPUT);
  }

  if (lfree !=NULL) lfree(cv_mem);

  /* Set four main function fields in cv_mem */
  linit  = CVDenseInit;
  lsetup = CVDenseSetup;
  lsolve = CVDenseSolve;
  lfree  = CVDenseFree;

  /* Get memory for CVDenseMemRec */
  cvdense_mem = NULL;
  cvdense_mem = (CVDenseMem) malloc(sizeof(CVDenseMemRec));
  if (cvdense_mem == NULL) {
    CVProcessError(cv_mem, CVDENSE_MEM_FAIL, "CVDENSE", "CVDense", MSGDS_MEM_FAIL);
    return(CVDENSE_MEM_FAIL);
  }

  /* Set default Jacobian routine and Jacobian data */
  jac = CVDenseDQJac;
  J_data = cvode_mem;
  last_flag = CVDENSE_SUCCESS;

  setupNonNull = TRUE;

  /* Set problem dimension */
  n = N;

  /* Allocate memory for M, savedJ, and pivot array */

  M = NULL;
  M = DenseAllocMat(N, N);
  if (M == NULL) {
    CVProcessError(cv_mem, CVDENSE_MEM_FAIL, "CVDENSE", "CVDense", MSGDS_MEM_FAIL);
    free(cvdense_mem); cvdense_mem = NULL;
    return(CVDENSE_MEM_FAIL);
  }
  savedJ = NULL;
  savedJ = DenseAllocMat(N, N);
  if (savedJ == NULL) {
    CVProcessError(cv_mem, CVDENSE_MEM_FAIL, "CVDENSE", "CVDense", MSGDS_MEM_FAIL);
    DenseFreeMat(M);
    free(cvdense_mem); cvdense_mem = NULL;
    return(CVDENSE_MEM_FAIL);
  }
  pivots = NULL;
  pivots = DenseAllocPiv(N);
  if (pivots == NULL) {
    CVProcessError(cv_mem, CVDENSE_MEM_FAIL, "CVDENSE", "CVDense", MSGDS_MEM_FAIL);
    DenseFreeMat(M);
    DenseFreeMat(savedJ);
    free(cvdense_mem); cvdense_mem = NULL;
    return(CVDENSE_MEM_FAIL);
  }

  /* Attach linear solver memory to integrator memory */
  lmem = cvdense_mem;

  return(CVDENSE_SUCCESS);
}

/*
 * -----------------------------------------------------------------
 * CVDenseSetJacFn
 * -----------------------------------------------------------------
 */

int CVDenseSetJacFn(void *cvode_mem, CVDenseJacFn djac, void *jac_data)
{
  CVodeMem cv_mem;
  CVDenseMem cvdense_mem;

  /* Return immediately if cvode_mem is NULL */
  if (cvode_mem == NULL) {
    CVProcessError(NULL, CVDENSE_MEM_NULL, "CVDENSE", "CVDenseSetJacFn", MSGDS_CVMEM_NULL);
    return(CVDENSE_MEM_NULL);
  }
  cv_mem = (CVodeMem) cvode_mem;

  if (lmem == NULL) {
    CVProcessError(cv_mem, CVDENSE_LMEM_NULL, "CVDENSE", "CVDenseSetJacFn", MSGDS_LMEM_NULL);
    return(CVDENSE_LMEM_NULL);
  }
  cvdense_mem = (CVDenseMem) lmem;

  jac = djac;
  if (djac != NULL) J_data = jac_data;

  return(CVDENSE_SUCCESS);
}

/*
 * -----------------------------------------------------------------
 * CVDenseGetWorkSpace
 * -----------------------------------------------------------------
 */

int CVDenseGetWorkSpace(void *cvode_mem, long int *lenrwLS, long int *leniwLS)
{
  CVodeMem cv_mem;
  CVDenseMem cvdense_mem;

  /* Return immediately if cvode_mem is NULL */
  if (cvode_mem == NULL) {
    CVProcessError(NULL, CVDENSE_MEM_NULL, "CVDENSE", "CVDenseGetWorkSpace", MSGDS_CVMEM_NULL);
    return(CVDENSE_MEM_NULL);
  }
  cv_mem = (CVodeMem) cvode_mem;

  if (lmem == NULL) {
    CVProcessError(cv_mem, CVDENSE_LMEM_NULL, "CVDENSE", "CVDenseGetWorkSpace", MSGDS_LMEM_NULL);
    return(CVDENSE_LMEM_NULL);
  }
  cvdense_mem = (CVDenseMem) lmem;

  *lenrwLS = 2*n*n;
  *leniwLS = n;

  return(CVDENSE_SUCCESS);
}

/*
 * -----------------------------------------------------------------
 * CVDenseGetNumJacEvals
 * -----------------------------------------------------------------
 */

int CVDenseGetNumJacEvals(void *cvode_mem, long int *njevals)
{
  CVodeMem cv_mem;
  CVDenseMem cvdense_mem;

  /* Return immediately if cvode_mem is NULL */
  if (cvode_mem == NULL) {
    CVProcessError(NULL, CVDENSE_MEM_NULL, "CVDENSE", "CVDenseGetNumJacEvals", MSGDS_CVMEM_NULL);
    return(CVDENSE_MEM_NULL);
  }
  cv_mem = (CVodeMem) cvode_mem;

  if (lmem == NULL) {
    CVProcessError(cv_mem, CVDENSE_LMEM_NULL, "CVDENSE", "CVDenseGetNumJacEvals", MSGDS_LMEM_NULL);
    return(CVDENSE_LMEM_NULL);
  }
  cvdense_mem = (CVDenseMem) lmem;

  *njevals = nje;

  return(CVDENSE_SUCCESS);
}

/*
 * -----------------------------------------------------------------
 * CVDenseGetNumRhsEvals
 * -----------------------------------------------------------------
 */

int CVDenseGetNumRhsEvals(void *cvode_mem, long int *nfevalsLS)
{
  CVodeMem cv_mem;
  CVDenseMem cvdense_mem;

  /* Return immediately if cvode_mem is NULL */
  if (cvode_mem == NULL) {
    CVProcessError(NULL, CVDENSE_MEM_NULL, "CVDENSE", "CVDenseGetNumRhsEvals", MSGDS_CVMEM_NULL);
    return(CVDENSE_MEM_NULL);
  }
  cv_mem = (CVodeMem) cvode_mem;

  if (lmem == NULL) {
    CVProcessError(cv_mem, CVDENSE_LMEM_NULL, "CVDENSE", "CVDenseGetNumRhsEvals", MSGDS_LMEM_NULL);
    return(CVDENSE_LMEM_NULL);
  }
  cvdense_mem = (CVDenseMem) lmem;

  *nfevalsLS = nfeD;

  return(CVDENSE_SUCCESS);
}

/*
 * -----------------------------------------------------------------
 * CVDenseGetLastFlag
 * -----------------------------------------------------------------
 */

int CVDenseGetLastFlag(void *cvode_mem, int *flag)
{
  CVodeMem cv_mem;
  CVDenseMem cvdense_mem;

  /* Return immediately if cvode_mem is NULL */
  if (cvode_mem == NULL) {
    CVProcessError(NULL, CVDENSE_MEM_NULL, "CVDENSE", "CVDenseGetLastFlag", MSGDS_CVMEM_NULL);
    return(CVDENSE_MEM_NULL);
  }
  cv_mem = (CVodeMem) cvode_mem;

  if (lmem == NULL) {
    CVProcessError(cv_mem, CVDENSE_LMEM_NULL, "CVDENSE", "CVDenseGetLastFlag", MSGDS_LMEM_NULL);
    return(CVDENSE_LMEM_NULL);
  }
  cvdense_mem = (CVDenseMem) lmem;

  *flag = last_flag;

  return(CVDENSE_SUCCESS);
}

/*
 * -----------------------------------------------------------------
 * CVDenseGetReturnFlagName
 * -----------------------------------------------------------------
 */

char *CVDenseGetReturnFlagName(int flag)
{
  char *name;

  name = (char *)malloc(30*sizeof(char));

  switch(flag) {
  case CVDENSE_SUCCESS:
    sprintf(name,"CVDENSE_SUCCESS");
    break; 
  case CVDENSE_MEM_NULL:
    sprintf(name,"CVDENSE_MEM_NULL");
    break;
  case CVDENSE_LMEM_NULL:
    sprintf(name,"CVDENSE_LMEM_NULL");
    break;
  case CVDENSE_ILL_INPUT:
    sprintf(name,"CVDENSE_ILL_INPUT");
    break;
  case CVDENSE_MEM_FAIL:
    sprintf(name,"CVDENSE_MEM_FAIL");
    break;
  case CVDENSE_JACFUNC_UNRECVR:
    sprintf(name,"CVDENSE_JACFUNC_UNRECVR");
    break;
  case CVDENSE_JACFUNC_RECVR:
    sprintf(name,"CVDENSE_JACFUNC_RECVR");
    break;
  case CVDENSE_ADJMEM_NULL:
    sprintf(name,"CVDENSE_ADJMEM_NULL");
    break;
  case CVDENSE_LMEMB_NULL:
    sprintf(name,"CVDENSE_LMEMB_NULL");
    break;
  default:
    sprintf(name,"NONE");
  }

  return(name);
}

/*
 * -----------------------------------------------------------------
 * CVDenseInit
 * -----------------------------------------------------------------
 * This routine does remaining initializations specific to the dense
 * linear solver.
 * -----------------------------------------------------------------
 */

static int CVDenseInit(CVodeMem cv_mem)
{
  CVDenseMem cvdense_mem;

  cvdense_mem = (CVDenseMem) lmem;
  
  nje   = 0;
  nfeD  = 0;
  nstlj = 0;
  
  if (jac == NULL) {
    jac = CVDenseDQJac;
    J_data = cv_mem;
  }

  last_flag = CVDENSE_SUCCESS;
  return(0);
}

/*
 * -----------------------------------------------------------------
 * CVDenseSetup
 * -----------------------------------------------------------------
 * This routine does the setup operations for the dense linear solver.
 * It makes a decision whether or not to call the Jacobian evaluation
 * routine based on various state variables, and if not it uses the 
 * saved copy.  In any case, it constructs the Newton matrix 
 * M = I - gamma*J, updates counters, and calls the dense LU 
 * factorization routine.
 * -----------------------------------------------------------------
 */

static int CVDenseSetup(CVodeMem cv_mem, int convfail, N_Vector ypred,
                        N_Vector fpred, booleantype *jcurPtr, 
                        N_Vector vtemp1, N_Vector vtemp2, N_Vector vtemp3)
{
  booleantype jbad, jok;
  realtype dgamma;
  long int ier;
  CVDenseMem cvdense_mem;
  int retval;

  cvdense_mem = (CVDenseMem) lmem;
 
  /* Use nst, gamma/gammap, and convfail to set J eval. flag jok */
 
  dgamma = ABS((gamma/gammap) - ONE);
  jbad = (nst == 0) || (nst > nstlj + CVD_MSBJ) ||
         ((convfail == CV_FAIL_BAD_J) && (dgamma < CVD_DGMAX)) ||
         (convfail == CV_FAIL_OTHER);
  jok = !jbad;
 
  if (jok) {

    /* If jok = TRUE, use saved copy of J */
    *jcurPtr = FALSE;
    DenseCopy(savedJ, M);

  } else {

    /* If jok = FALSE, call jac routine for new J value */
    nje++;
    nstlj = nst;
    *jcurPtr = TRUE;
    DenseZero(M);

    retval = jac(n, M, tn, ypred, fpred, J_data, vtemp1, vtemp2, vtemp3);
    if (retval < 0) {
      CVProcessError(cv_mem, CVDENSE_JACFUNC_UNRECVR, "CVDENSE", "CVDenseSetup", MSGDS_JACFUNC_FAILED);
      last_flag = CVDENSE_JACFUNC_UNRECVR;
      return(-1);
    }
    if (retval > 0) {
      last_flag = CVDENSE_JACFUNC_RECVR;
      return(1);
    }

    DenseCopy(M, savedJ);

  }
  
  /* Scale and add I to get M = I - gamma*J */
  DenseScale(-gamma, M);
  DenseAddI(M);

  /* Do LU factorization of M */
  ier = DenseGETRF(M, pivots); 

  /* Return 0 if the LU was complete; otherwise return 1 */
  last_flag = ier;
  if (ier > 0) return(1);
  return(0);
}

/*
 * -----------------------------------------------------------------
 * CVDenseSolve
 * -----------------------------------------------------------------
 * This routine handles the solve operation for the dense linear solver
 * by calling the dense backsolve routine.  The returned value is 0.
 * -----------------------------------------------------------------
 */

static int CVDenseSolve(CVodeMem cv_mem, N_Vector b, N_Vector weight,
                        N_Vector ycur, N_Vector fcur)
{
  CVDenseMem cvdense_mem;
  realtype *bd;

  cvdense_mem = (CVDenseMem) lmem;
  
  bd = N_VGetArrayPointer(b);

  DenseGETRS(M, pivots, bd);

  /* If CV_BDF, scale the correction to account for change in gamma */
  if ((lmm == CV_BDF) && (gamrat != ONE)) {
    N_VScale(TWO/(ONE + gamrat), b, b);
  }
  
  last_flag = CVDENSE_SUCCESS;
  return(0);
}

/*
 * -----------------------------------------------------------------
 * CVDenseFree
 * -----------------------------------------------------------------
 * This routine frees memory specific to the dense linear solver.
 * -----------------------------------------------------------------
 */

static void CVDenseFree(CVodeMem cv_mem)
{
  CVDenseMem  cvdense_mem;

  cvdense_mem = (CVDenseMem) lmem;
  
  DenseFreeMat(M);
  DenseFreeMat(savedJ);
  DenseFreePiv(pivots);
  free(cvdense_mem); cvdense_mem = NULL;
}

/*
 * -----------------------------------------------------------------
 * CVDenseDQJac 
 * -----------------------------------------------------------------
 * This routine generates a dense difference quotient approximation to
 * the Jacobian of f(t,y). It assumes that a dense matrix of type
 * DenseMat is stored column-wise, and that elements within each column
 * are contiguous. The address of the jth column of J is obtained via
 * the macro DENSE_COL and this pointer is associated with an N_Vector
 * using the N_VGetArrayPointer/N_VSetArrayPointer functions. 
 * Finally, the actual computation of the jth column of the Jacobian is 
 * done with a call to N_VLinearSum.
 * -----------------------------------------------------------------
 */
 
static int CVDenseDQJac(long int N, DenseMat J, realtype t, 
                        N_Vector y, N_Vector fy, void *jac_data,
                        N_Vector tmp1, N_Vector tmp2, N_Vector tmp3)
{
  realtype fnorm, minInc, inc, inc_inv, yjsaved, srur;
  realtype *tmp2_data, *y_data, *ewt_data;
  N_Vector ftemp, jthCol;
  long int j;
  int retval = 0;

  CVodeMem cv_mem;
  CVDenseMem  cvdense_mem;

  /* jac_data points to cvode_mem */
  cv_mem = (CVodeMem) jac_data;
  cvdense_mem = (CVDenseMem) lmem;

  /* Save pointer to the array in tmp2 */
  tmp2_data = N_VGetArrayPointer(tmp2);

  /* Rename work vectors for readibility */
  ftemp = tmp1; 
  jthCol = tmp2;

  /* Obtain pointers to the data for ewt, y */
  ewt_data = N_VGetArrayPointer(ewt);
  y_data   = N_VGetArrayPointer(y);

  /* Set minimum increment based on uround and norm of f */
  srur = RSqrt(uround);
  fnorm = N_VWrmsNorm(fy, ewt);
  minInc = (fnorm != ZERO) ?
           (MIN_INC_MULT * ABS(h) * uround * N * fnorm) : ONE;

  /* This is the only for loop for 0..N-1 in CVODE */

  for (j = 0; j < N; j++) {

    /* Generate the jth col of J(tn,y) */

    N_VSetArrayPointer(DENSE_COL(J,j), jthCol);

    yjsaved = y_data[j];
    inc = MAX(srur*ABS(yjsaved), minInc/ewt_data[j]);
    y_data[j] += inc;

    retval = f(tn, y, ftemp, f_data);
    nfeD++;
    if (retval != 0) break;

    y_data[j] = yjsaved;

    inc_inv = ONE/inc;
    N_VLinearSum(inc_inv, ftemp, -inc_inv, fy, jthCol);

    DENSE_COL(J,j) = N_VGetArrayPointer(jthCol);
  }

  /* Restore original array pointer in tmp2 */
  N_VSetArrayPointer(tmp2_data, tmp2);

  return(retval);
}


/* 
 * ================================================================
 *
 *                   PART II - backward problems
 *
 * ================================================================
 */


/* Additional readability replacements */

#define ytmp        (ca_mem->ca_ytmp)
#define getY        (ca_mem->ca_getY)
#define lmemB       (ca_mem->ca_lmemB)
#define lfreeB      (ca_mem->ca_lfreeB)

#define djac_B      (cvdenseB_mem->d_djacB)
#define jac_data_B  (cvdenseB_mem->d_jac_dataB)


/*
 * CVDenseB and CVdenseSet*B
 *
 * Wrappers for the backward phase around the corresponding 
 * CVODES functions
 */

int CVDenseB(void *cvadj_mem, long int nB)
{
  CVadjMem ca_mem;
  CVDenseMemB cvdenseB_mem;
  CVodeMem cvB_mem;
  int flag;

  if (cvadj_mem == NULL) {
    CVProcessError(NULL, CVDENSE_ADJMEM_NULL, "CVDENSE", "CVDenseB", MSGDS_CAMEM_NULL);
    return(CVDENSE_ADJMEM_NULL);
  }
  ca_mem = (CVadjMem) cvadj_mem;

  cvB_mem = ca_mem->cvb_mem;

  /* Get memory for CVDenseMemRecB */
  cvdenseB_mem = (CVDenseMemB) malloc(sizeof(CVDenseMemRecB));
  if (cvdenseB_mem == NULL) {
    CVProcessError(cvB_mem, CVDENSE_MEM_FAIL, "CVDENSE", "CVDenseB", MSGDS_MEM_FAIL);
    return(CVDENSE_MEM_FAIL);
  }

  djac_B = NULL;
  jac_data_B = NULL;

  /* attach lmemB and lfreeB */
  lmemB = cvdenseB_mem;
  lfreeB = CVDenseFreeB;

  flag = CVDense(cvB_mem, nB);

  if (flag != CVDENSE_SUCCESS) {
    free(cvdenseB_mem);
    cvdenseB_mem = NULL;
  }

  return(flag);
}

int CVDenseSetJacFnB(void *cvadj_mem, CVDenseJacFnB djacB, void *jac_dataB)
{
  CVadjMem ca_mem;
  CVDenseMemB cvdenseB_mem;
  CVodeMem cvB_mem;
  int flag;

  if (cvadj_mem == NULL) {
    CVProcessError(NULL, CVDENSE_ADJMEM_NULL, "CVDENSE", "CVDenseSetJacFnB", MSGDS_CAMEM_NULL);
    return(CVDENSE_ADJMEM_NULL);
  }
  ca_mem = (CVadjMem) cvadj_mem;

  cvB_mem = ca_mem->cvb_mem;

  if (lmemB == NULL) {
    CVProcessError(cvB_mem, CVDENSE_LMEMB_NULL, "CVDENSE", "CVDenseSetJacFnB", MSGDS_LMEMB_NULL);
    return(CVDENSE_LMEMB_NULL);
  }
  cvdenseB_mem = (CVDenseMemB) lmemB;

  djac_B     = djacB;
  jac_data_B = jac_dataB;

  flag = CVDenseSetJacFn(cvB_mem, CVAdenseJac, cvadj_mem);

  return(flag);
}

/*
 * CVDenseFreeB 
 */


static void CVDenseFreeB(CVadjMem ca_mem)
{
  CVDenseMemB cvdenseB_mem;

  cvdenseB_mem = (CVDenseMemB) lmemB;

  free(cvdenseB_mem);
}


/*
 * CVAdenseJac
 *
 * This routine interfaces to the CVDenseJacFnB routine provided 
 * by the user.
 * NOTE: jac_data actually contains cvadj_mem
 */

static int CVAdenseJac(long int nB, DenseMat JB, realtype t, 
                       N_Vector yB, N_Vector fyB, void *cvadj_mem,
                       N_Vector tmp1B, N_Vector tmp2B, N_Vector tmp3B)
{
  CVadjMem ca_mem;
  CVodeMem cvB_mem;
  CVDenseMemB cvdenseB_mem;
  int retval, flag;

  ca_mem = (CVadjMem) cvadj_mem;
  cvB_mem = ca_mem->cvb_mem;
  cvdenseB_mem = (CVDenseMemB) lmemB;

  /* Forward solution from interpolation */
  flag = getY(ca_mem, t, ytmp);
  if (flag != CV_SUCCESS) {
    CVProcessError(cvB_mem, -1, "CVDENSE", "CVAdenseJac", MSGDS_BAD_T);
    return(-1);
  }

  /* Call user's adjoint dense djacB routine */
  retval = djac_B(nB, JB, t, ytmp, yB, fyB, jac_data_B, 
                  tmp1B, tmp2B, tmp3B);

  return(retval);

}
```
