## Supplementary scripts for "A multi-scale *in silico* mouse model for insulin resistance and humanoid type 2 diabetes": cvodes_dense_impl.h

```
/*
 * -----------------------------------------------------------------
 * $Revision: 749 $
 * $Date: 2011-06-22 14:49:10 +0200 (Wed, 22 Jun 2011) $
 * ----------------------------------------------------------------- 
 * Programmer(s): Radu Serban @ LLNL
 * -----------------------------------------------------------------
 * Copyright (c) 2005, The Regents of the University of California.
 * Produced at the Lawrence Livermore National Laboratory.
 * All rights reserved.
 * For details, see the LICENSE file.
 * -----------------------------------------------------------------
 * Implementation header file for the dense linear solver, CVDENSE.
 * -----------------------------------------------------------------
 */

#ifndef _CVSDENSE_IMPL_H
#define _CVSDENSE_IMPL_H

#ifdef __cplusplus  /* wrapper to enable C++ usage */
extern "C" {
#endif

#include <cvodes/cvodes_dense.h>

  /*
   * -----------------------------------------------------------------
   * Types : CVDenseMemRec, CVDenseMem                             
   * -----------------------------------------------------------------
   * The type CVDenseMem is pointer to a CVDenseMemRec.
   * This structure contains CVDense solver-specific data.
   *
   * CVDense attaches such a structure to the lmem field of CVodeMem
   * -----------------------------------------------------------------
   */

  typedef struct {

    long int d_n;        /* problem dimension                       */

    CVDenseJacFn d_jac;  /* jac = Jacobian routine to be called     */

    DenseMat d_M;        /* M = I - gamma J, gamma = h / l1         */
  
    long int *d_pivots;  /* pivots = pivot array for PM = LU        */
  
    DenseMat d_savedJ;   /* savedJ = old Jacobian                   */
  
    long int  d_nstlj;   /* nstlj = nst at last Jacobian eval.      */
  
    long int d_nje;      /* nje = no. of calls to jac               */

    long int d_nfeD;     /* nfeD = no. of calls to f due to
                            difference quotient approximation of J  */
  
    void *d_J_data;      /* J_data is passed to jac                 */

    int d_last_flag;     /* last error return flag                  */
  
  } CVDenseMemRec, *CVDenseMem;


  /*
   * -----------------------------------------------------------------
   * Types : CVDenseMemRecB, CVDenseMemB       
   * -----------------------------------------------------------------
   * CVDenseB attaches such a structure to the lmemB filed of CVadjMem
   * -----------------------------------------------------------------
   */

  typedef struct {

    CVDenseJacFnB d_djacB;
    void *d_jac_dataB;

  } CVDenseMemRecB, *CVDenseMemB;

  /*
   * -----------------------------------------------------------------
   * Error Messages 
   * -----------------------------------------------------------------
   */

#define MSGDS_CVMEM_NULL "Integrator memory is NULL."
#define MSGDS_MEM_FAIL "A memory request failed."
#define MSGDS_BAD_NVECTOR "A required vector operation is not implemented."
#define MSGDS_LMEM_NULL "CVDENSE memory is NULL."
#define MSGDS_JACFUNC_FAILED "The Jacobian routine failed in an unrecoverable manner."

#define MSGDS_CAMEM_NULL "cvadj_mem = NULL illegal."
#define MSGDS_LMEMB_NULL "CVDENSE memory is NULL for the backward integration."
#define MSGDS_BAD_T "Bad t for interpolation."


#ifdef __cplusplus
}
#endif

#endif
```
