## Supplementary scripts for "A multi-scale *in silico* mouse model for insulin resistance and humanoid type 2 diabetes": cvodes_diag.c

```
/*
 * -----------------------------------------------------------------
 * $Revision: 749 $
 * $Date: 2011-06-22 14:49:10 +0200 (Wed, 22 Jun 2011) $
 * ----------------------------------------------------------------- 
 * Programmer(s): Radu Serban @ LLNL
 * -----------------------------------------------------------------
 * Copyright (c) 2005, The Regents of the University of California.
 * Produced at the Lawrence Livermore National Laboratory.
 * All rights reserved.
 * For details, see the LICENSE file.
 * -----------------------------------------------------------------
 * This is the implementation file for the CVDIAG linear solver.
 * -----------------------------------------------------------------
 */

#include <stdio.h>
#include <stdlib.h>

#include "cvodes_diag_impl.h"
#include "cvodes_impl.h"

/* Other Constants */
  
#define FRACT RCONST(0.1)
#define ONE   RCONST(1.0)

/* CVDIAG linit, lsetup, lsolve, and lfree routines */

static int CVDiagInit(CVodeMem cv_mem);

static int CVDiagSetup(CVodeMem cv_mem, int convfail, N_Vector ypred,
                       N_Vector fpred, booleantype *jcurPtr, N_Vector vtemp1,
                       N_Vector vtemp2, N_Vector vtemp3);

static int CVDiagSolve(CVodeMem cv_mem, N_Vector b, N_Vector weight,
                       N_Vector ycur, N_Vector fcur);

static void CVDiagFree(CVodeMem cv_mem);


/* 
 * ================================================================
 *
 *                   PART I - forward problems
 *
 * ================================================================
 */


/* Readability Replacements */

#define lrw1      (cv_mem->cv_lrw1)
#define liw1      (cv_mem->cv_liw1)
#define f         (cv_mem->cv_f)
#define f_data    (cv_mem->cv_f_data)
#define uround    (cv_mem->cv_uround)
#define tn        (cv_mem->cv_tn)
#define h         (cv_mem->cv_h)
#define rl1       (cv_mem->cv_rl1)
#define gamma     (cv_mem->cv_gamma)
#define ewt       (cv_mem->cv_ewt)
#define nfe       (cv_mem->cv_nfe)
#define zn        (cv_mem->cv_zn)
#define linit     (cv_mem->cv_linit)
#define lsetup    (cv_mem->cv_lsetup)
#define lsolve    (cv_mem->cv_lsolve)
#define lfree     (cv_mem->cv_lfree)
#define lmem      (cv_mem->cv_lmem)
#define vec_tmpl  (cv_mem->cv_tempv)
#define setupNonNull   (cv_mem->cv_setupNonNull)

#define gammasv   (cvdiag_mem->di_gammasv)
#define M         (cvdiag_mem->di_M)
#define bit       (cvdiag_mem->di_bit)
#define bitcomp   (cvdiag_mem->di_bitcomp)
#define nfeDI     (cvdiag_mem->di_nfeDI)
#define last_flag (cvdiag_mem->di_last_flag)


/*
 * -----------------------------------------------------------------
 * CVDiag 
 * -----------------------------------------------------------------
 * This routine initializes the memory record and sets various function
 * fields specific to the diagonal linear solver module.  CVDense first
 * calls the existing lfree routine if this is not NULL.  Then it sets
 * the cv_linit, cv_lsetup, cv_lsolve, cv_lfree fields in (*cvode_mem)
 * to be CVDiagInit, CVDiagSetup, CVDiagSolve, and CVDiagFree,
 * respectively.  It allocates memory for a structure of type
 * CVDiagMemRec and sets the cv_lmem field in (*cvode_mem) to the
 * address of this structure.  It sets setupNonNull in (*cvode_mem) to
 * TRUE.  Finally, it allocates memory for M, bit, and bitcomp.
 * The CVDiag return value is SUCCESS = 0, LMEM_FAIL = -1, or 
 * LIN_ILL_INPUT=-2.
 * -----------------------------------------------------------------
 */
  
int CVDiag(void *cvode_mem)
{
  CVodeMem cv_mem;
  CVDiagMem cvdiag_mem;

  /* Return immediately if cvode_mem is NULL */
  if (cvode_mem == NULL) {
    CVProcessError(NULL, CVDIAG_MEM_NULL, "CVDIAG", "CVDiag", MSGDG_CVMEM_NULL);
    return(CVDIAG_MEM_NULL);
  }
  cv_mem = (CVodeMem) cvode_mem;

  /* Check if N_VCompare and N_VInvTest are present */
  if(vec_tmpl->ops->nvcompare == NULL ||
     vec_tmpl->ops->nvinvtest == NULL) {
    CVProcessError(cv_mem, CVDIAG_ILL_INPUT, "CVDIAG", "CVDiag", MSGDG_BAD_NVECTOR);
    return(CVDIAG_ILL_INPUT);
  }

  if (lfree != NULL) lfree(cv_mem);
  
  /* Set four main function fields in cv_mem */
  linit  = CVDiagInit;
  lsetup = CVDiagSetup;
  lsolve = CVDiagSolve;
  lfree  = CVDiagFree;

  /* Get memory for CVDiagMemRec */
  cvdiag_mem = NULL;
  cvdiag_mem = (CVDiagMem) malloc(sizeof(CVDiagMemRec));
  if (cvdiag_mem == NULL) {
    CVProcessError(cv_mem, CVDIAG_MEM_FAIL, "CVDIAG", "CVDiag", MSGDG_MEM_FAIL);
    return(CVDIAG_MEM_FAIL);
  }

  last_flag = CVDIAG_SUCCESS;

  /* Set flag setupNonNull = TRUE */
  setupNonNull = TRUE;

  /* Allocate memory for M, bit, and bitcomp */
    
  M = NULL;
  M = N_VClone(vec_tmpl);
  if (M == NULL) {
    CVProcessError(cv_mem, CVDIAG_MEM_FAIL, "CVDIAG", "CVDiag", MSGDG_MEM_FAIL);
    free(cvdiag_mem); cvdiag_mem = NULL;
    return(CVDIAG_MEM_FAIL);
  }
  bit = NULL;
  bit = N_VClone(vec_tmpl);
  if (bit == NULL) {
    CVProcessError(cv_mem, CVDIAG_MEM_FAIL, "CVDIAG", "CVDiag", MSGDG_MEM_FAIL);
    N_VDestroy(M);
    free(cvdiag_mem); cvdiag_mem = NULL;
    return(CVDIAG_MEM_FAIL);
  }
  bitcomp = NULL;
  bitcomp = N_VClone(vec_tmpl);
  if (bitcomp == NULL) {
    CVProcessError(cv_mem, CVDIAG_MEM_FAIL, "CVDIAG", "CVDiag", MSGDG_MEM_FAIL);
    N_VDestroy(M);
    N_VDestroy(bit);
    free(cvdiag_mem); cvdiag_mem = NULL;
    return(CVDIAG_MEM_FAIL);
  }

  /* Attach linear solver memory to integrator memory */
  lmem = cvdiag_mem;

  return(CVDIAG_SUCCESS);
}

/*
 * -----------------------------------------------------------------
 * CVDiagGetWorkSpace
 * -----------------------------------------------------------------
 */

int CVDiagGetWorkSpace(void *cvode_mem, long int *lenrwLS, long int *leniwLS)
{
  CVodeMem cv_mem;

  /* Return immediately if cvode_mem is NULL */
  if (cvode_mem == NULL) {
    CVProcessError(NULL, CVDIAG_MEM_NULL, "CVDIAG", "CVDiagGetWorkSpace", MSGDG_CVMEM_NULL);
    return(CVDIAG_MEM_NULL);
  }
  cv_mem = (CVodeMem) cvode_mem;

  *lenrwLS = 3*lrw1;
  *leniwLS = 3*liw1;

  return(CVDIAG_SUCCESS);
}

/*
 * -----------------------------------------------------------------
 * CVDiagGetNumRhsEvals
 * -----------------------------------------------------------------
 */

int CVDiagGetNumRhsEvals(void *cvode_mem, long int *nfevalsLS)
{
  CVodeMem cv_mem;
  CVDiagMem cvdiag_mem;

  /* Return immediately if cvode_mem is NULL */
  if (cvode_mem == NULL) {
    CVProcessError(NULL, CVDIAG_MEM_NULL, "CVDIAG", "CVDiagGetNumRhsEvals", MSGDG_CVMEM_NULL);
    return(CVDIAG_MEM_NULL);
  }
  cv_mem = (CVodeMem) cvode_mem;

  if (lmem == NULL) {
    CVProcessError(cv_mem, CVDIAG_LMEM_NULL, "CVDIAG", "CVDiagGetNumRhsEvals", MSGDG_LMEM_NULL);
    return(CVDIAG_LMEM_NULL);
  }
  cvdiag_mem = (CVDiagMem) lmem;

  *nfevalsLS = nfeDI;

  return(CVDIAG_SUCCESS);
}

/*
 * -----------------------------------------------------------------
 * CVDiagGetLastFlag
 * -----------------------------------------------------------------
 */

int CVDiagGetLastFlag(void *cvode_mem, int *flag)
{
  CVodeMem cv_mem;
  CVDiagMem cvdiag_mem;

  /* Return immediately if cvode_mem is NULL */
  if (cvode_mem == NULL) {
    CVProcessError(NULL, CVDIAG_MEM_NULL, "CVDIAG", "CVDiagGetLastFlag", MSGDG_CVMEM_NULL);
    return(CVDIAG_MEM_NULL);
  }
  cv_mem = (CVodeMem) cvode_mem;

  if (lmem == NULL) {
    CVProcessError(cv_mem, CVDIAG_LMEM_NULL, "CVDIAG", "CVDiagGetLastFlag", MSGDG_LMEM_NULL);
    return(CVDIAG_LMEM_NULL);
  }
  cvdiag_mem = (CVDiagMem) lmem;

  *flag = last_flag;

  return(CVDIAG_SUCCESS);
}

/*
 * -----------------------------------------------------------------
 * CVDiagGetReturnFlagName
 * -----------------------------------------------------------------
 */

char *CVDiagGetReturnFlagName(int flag)
{
  char *name;

  name = (char *)malloc(30*sizeof(char));

  switch(flag) {
  case CVDIAG_SUCCESS:
    sprintf(name,"CVDIAG_SUCCESS");
    break;  
  case CVDIAG_MEM_NULL:
    sprintf(name,"CVDIAG_MEM_NULL");
    break;
  case CVDIAG_LMEM_NULL:
    sprintf(name,"CVDIAG_LMEM_NULL");
    break;
  case CVDIAG_ILL_INPUT:
    sprintf(name,"CVDIAG_ILL_INPUT");
    break;
  case CVDIAG_MEM_FAIL:
    sprintf(name,"CVDIAG_MEM_FAIL");
    break;
  case CVDIAG_INV_FAIL:
    sprintf(name,"CVDIAG_INV_FAIL");
    break;
  case CVDIAG_RHSFUNC_UNRECVR:
    sprintf(name,"CVDIAG_RHSFUNC_UNRECVR");
    break;
  case CVDIAG_RHSFUNC_RECVR:
    sprintf(name,"CVDIAG_RHSFUNC_RECVR");
    break;
  case CVDIAG_ADJMEM_NULL:
    sprintf(name,"CVDIAG_ADJMEM_NULL");
    break;
  default:
    sprintf(name,"NONE");
  }

  return(name);
}

/*
 * -----------------------------------------------------------------
 * CVDiagInit
 * -----------------------------------------------------------------
 * This routine does remaining initializations specific to the diagonal
 * linear solver.
 * -----------------------------------------------------------------
 */

static int CVDiagInit(CVodeMem cv_mem)
{
  CVDiagMem cvdiag_mem;

  cvdiag_mem = (CVDiagMem) lmem;

  nfeDI = 0;

  last_flag = CVDIAG_SUCCESS;
  return(0);
}

/*
 * -----------------------------------------------------------------
 * CVDiagSetup
 * -----------------------------------------------------------------
 * This routine does the setup operations for the diagonal linear 
 * solver.  It constructs a diagonal approximation to the Newton matrix 
 * M = I - gamma*J, updates counters, and inverts M.
 * -----------------------------------------------------------------
 */

static int CVDiagSetup(CVodeMem cv_mem, int convfail, N_Vector ypred,
                       N_Vector fpred, booleantype *jcurPtr, N_Vector vtemp1,
                       N_Vector vtemp2, N_Vector vtemp3)
{
  realtype r;
  N_Vector ftemp, y;
  booleantype invOK;
  CVDiagMem cvdiag_mem;
  int retval;

  cvdiag_mem = (CVDiagMem) lmem;

  /* Rename work vectors for use as temporary values of y and f */
  ftemp = vtemp1;
  y     = vtemp2;

  /* Form y with perturbation = FRACT*(func. iter. correction) */
  r = FRACT * rl1;
  N_VLinearSum(h, fpred, -ONE, zn[1], ftemp);
  N_VLinearSum(r, ftemp, ONE, ypred, y);

  /* Evaluate f at perturbed y */
  retval = f(tn, y, M, f_data);
  nfeDI++;
  if (retval < 0) {
    CVProcessError(cv_mem, CVDIAG_RHSFUNC_UNRECVR, "CVDIAG", "CVDiagSetup", MSGDG_RHSFUNC_FAILED);
    last_flag = CVDIAG_RHSFUNC_UNRECVR;
    return(-1);
  }
  if (retval > 0) {
    last_flag = CVDIAG_RHSFUNC_RECVR;
    return(1);
  }

  /* Construct M = I - gamma*J with J = diag(deltaf_i/deltay_i) */
  N_VLinearSum(ONE, M, -ONE, fpred, M);
  N_VLinearSum(FRACT, ftemp, -h, M, M);
  N_VProd(ftemp, ewt, y);
  /* Protect against deltay_i being at roundoff level */
  N_VCompare(uround, y, bit);
  N_VAddConst(bit, -ONE, bitcomp);
  N_VProd(ftemp, bit, y);
  N_VLinearSum(FRACT, y, -ONE, bitcomp, y);
  N_VDiv(M, y, M);
  N_VProd(M, bit, M);
  N_VLinearSum(ONE, M, -ONE, bitcomp, M);

  /* Invert M with test for zero components */
  invOK = N_VInvTest(M, M);
  if (!invOK) {
    last_flag = CVDIAG_INV_FAIL;
    return(1);
  }

  /* Set jcur = TRUE, save gamma in gammasv, and return */
  *jcurPtr = TRUE;
  gammasv = gamma;
  last_flag = CVDIAG_SUCCESS;
  return(0);
}

/*
 * -----------------------------------------------------------------
 * CVDiagSolve
 * -----------------------------------------------------------------
 * This routine performs the solve operation for the diagonal linear
 * solver.  If necessary it first updates gamma in M = I - gamma*J.
 * -----------------------------------------------------------------
 */

static int CVDiagSolve(CVodeMem cv_mem, N_Vector b, N_Vector weight,
                       N_Vector ycur, N_Vector fcur)
{
  booleantype invOK;
  realtype r;
  CVDiagMem cvdiag_mem;

  cvdiag_mem = (CVDiagMem) lmem;
  
  /* If gamma has changed, update factor in M, and save gamma value */

  if (gammasv != gamma) {
    r = gamma / gammasv;
    N_VInv(M, M);
    N_VAddConst(M, -ONE, M);
    N_VScale(r, M, M);
    N_VAddConst(M, ONE, M);
    invOK = N_VInvTest(M, M);
    if (!invOK) {
      last_flag = CVDIAG_INV_FAIL;
      return (1);
    }
    gammasv = gamma;
  }

  /* Apply M-inverse to b */
  N_VProd(b, M, b);

  last_flag = CVDIAG_SUCCESS;
  return(0);
}

/*
 * -----------------------------------------------------------------
 * CVDiagFree
 * -----------------------------------------------------------------
 * This routine frees memory specific to the diagonal linear solver.
 * -----------------------------------------------------------------
 */

static void CVDiagFree(CVodeMem cv_mem)
{
  CVDiagMem cvdiag_mem;
  
  cvdiag_mem = (CVDiagMem) lmem;

  N_VDestroy(M);
  N_VDestroy(bit);
  N_VDestroy(bitcomp);
  free(cvdiag_mem); cvdiag_mem = NULL;
}


/* 
 * ================================================================
 *
 *                   PART II - backward problems
 *
 * ================================================================
 */


/*
 * CVDiagB
 *
 * Wrappers for the backward phase around the corresponding 
 * CVODES functions
 */

int CVDiagB(void *cvadj_mem)
{
  CVadjMem ca_mem;
  CVodeMem cvB_mem;
  int flag;

  if (cvadj_mem == NULL) {
    CVProcessError(NULL, CVDIAG_ADJMEM_NULL, "CVDIAG", "CVDiagB", MSGDG_CAMEM_NULL);
    return(CVDIAG_ADJMEM_NULL);
  }
  ca_mem = (CVadjMem) cvadj_mem;

  cvB_mem = ca_mem->cvb_mem;
  
  flag = CVDiag(cvB_mem);

  return(flag);
}
```
