## Supplementary scripts for "A multi-scale *in silico* mouse model for insulin resistance and humanoid type 2 diabetes": cvodes_diag_impl.h

```
/*
 * -----------------------------------------------------------------
 * $Revision: 749 $
 * $Date: 2011-06-22 14:49:10 +0200 (Wed, 22 Jun 2011) $
 * ----------------------------------------------------------------- 
 * Programmer(s): Radu Serban @ LLNL
 * -----------------------------------------------------------------
 * Copyright (c) 2005, The Regents of the University of California.
 * Produced at the Lawrence Livermore National Laboratory.
 * All rights reserved.
 * For details, see the LICENSE file.
 * -----------------------------------------------------------------
 * Implementation header file for the diagonal linear solver, CVDIAG.
 * -----------------------------------------------------------------
 */

#ifndef _CVSDIAG_IMPL_H
#define _CVSDIAG_IMPL_H

#ifdef __cplusplus  /* wrapper to enable C++ usage */
extern "C" {
#endif

#include <cvodes/cvodes_diag.h>

  /*
   * -----------------------------------------------------------------
   * Types: CVDiagMemRec, CVDiagMem
   * -----------------------------------------------------------------
   * The type CVDiagMem is pointer to a CVDiagMemRec.
   * This structure contains CVDiag solver-specific data.
   * -----------------------------------------------------------------
   */

  typedef struct {

    realtype di_gammasv; /* gammasv = gamma at the last call to setup */
    /* or solve                                  */

    N_Vector di_M;       /* M = (I - gamma J)^{-1} , gamma = h / l1   */

    N_Vector di_bit;     /* temporary storage vector                  */

    N_Vector di_bitcomp; /* temporary storage vector                  */

    long int di_nfeDI;   /* no. of calls to f due to difference 
                            quotient diagonal Jacobian approximation  */

    int di_last_flag;    /* last error return flag                    */

  } CVDiagMemRec, *CVDiagMem;

  /* Error Messages */

#define MSGDG_CVMEM_NULL "Integrator memory is NULL."
#define MSGDG_MEM_FAIL "A memory request failed."
#define MSGDG_BAD_NVECTOR "A required vector operation is not implemented."
#define MSGDG_LMEM_NULL "CVDIAG memory is NULL."
#define MSGDG_RHSFUNC_FAILED "The right-hand side routine failed in an unrecoverable manner."

#define MSGDG_CAMEM_NULL "cvadj_mem = NULL illegal."

#ifdef __cplusplus
}
#endif

#endif
```
