## Supplementary scripts for "A multi-scale *in silico* mouse model for insulin resistance and humanoid type 2 diabetes": cvodes_impl.h

```
/*
 * -----------------------------------------------------------------
 * $Revision: 749 $
 * $Date: 2011-06-22 14:49:10 +0200 (Wed, 22 Jun 2011) $
 * ----------------------------------------------------------------- 
 * Programmer(s): Radu Serban @ LLNL
 * -----------------------------------------------------------------
 * Copyright (c) 2005, The Regents of the University of California.
 * Produced at the Lawrence Livermore National Laboratory.
 * All rights reserved.
 * For details, see the LICENSE file.
 * -----------------------------------------------------------------
 * Implementation header file for the main CVODES integrator.
 * -----------------------------------------------------------------
 */

#ifndef _CVODES_IMPL_H
#define _CVODES_IMPL_H

#ifdef __cplusplus  /* wrapper to enable C++ usage */
extern "C" {
#endif

#include <stdarg.h>

#include <cvodes/cvodes.h>
#include <sundials/sundials_nvector.h>
#include <sundials/sundials_types.h>

  /* 
   * =================================================================
   *   M A I N    I N T E G R A T O R    M E M O R Y    B L O C K
   * =================================================================
   */

  /* Basic CVODES constants */

#define ADAMS_Q_MAX 12      /* max value of q for lmm == ADAMS    */
#define BDF_Q_MAX    5      /* max value of q for lmm == BDF      */
#define Q_MAX  ADAMS_Q_MAX  /* max value of q for either lmm      */
#define L_MAX  (Q_MAX+1)    /* max value of L for either lmm      */
#define NUM_TESTS    5      /* number of error test quantities    */

#define HMIN_DEFAULT     RCONST(0.0)    /* hmin default value     */
#define HMAX_INV_DEFAULT RCONST(0.0)    /* hmax_inv default value */
#define MXHNIL_DEFAULT   10             /* mxhnil default value   */
#define MXSTEP_DEFAULT   500            /* mxstep default value   */

  /*                                                                
   * ifS is the type of the function returning the sensitivity
   * right-hand side. ifS can be either CV_ALLSENS if the function    
   * (of type CVSensRhsFn) returns right hand sides for all    
   * sensitivity systems at once, or CV_ONESENS if the function 
   * (of type SensRhs1Fn) returns the right hand side of one 
   *  sensitivity system at a time.                           
   */

#define CV_ONESENS 1
#define CV_ALLSENS 2

  /*
   * -----------------------------------------------------------------
   * Types: struct CVodeMemRec, CVodeMem
   * -----------------------------------------------------------------
   * The type CVodeMem is type pointer to struct CVodeMemRec.
   * This structure contains fields to keep track of problem state.
   * -----------------------------------------------------------------
   */
  
  typedef struct CVodeMemRec {
    
    realtype cv_uround;      /* machine unit roundoff                        */   

    /*-------------------------- 
      Problem Specification Data 
      --------------------------*/

    CVRhsFn cv_f;            /* y' = f(t,y(t))                               */
    void *cv_f_data;         /* user pointer passed to f                     */
    int cv_lmm;              /* lmm = ADAMS or BDF                           */
    int cv_iter;             /* iter = FUNCTIONAL or NEWTON                  */
    int cv_itol;             /* itol = SS or SV                              */

    realtype cv_reltol;      /* relative tolerance                           */
    realtype cv_Sabstol;     /* scalar absolute tolerance                    */
    N_Vector cv_Vabstol;     /* vector absolute tolerance                    */
    CVEwtFn cv_efun;         /* function to set ewt                          */
    void *cv_e_data;         /* user pointer passed to efun                  */

    /*-----------------------
      Quadrature Related Data 
      -----------------------*/

    booleantype cv_quadr;    /* TRUE if integrating quadratures              */

    CVQuadRhsFn cv_fQ;
    void *cv_fQ_data;        /* user pointer passed to fQ                    */
    int cv_itolQ;
    booleantype cv_errconQ;

    realtype cv_reltolQ;     /* relative tolerance for quadratures           */
    realtype cv_SabstolQ;    /* scalar absolute tolerance for quadratures    */
    N_Vector cv_VabstolQ;    /* vector absolute tolerance for quadratures    */

    /*------------------------
      Sensitivity Related Data 
      ------------------------*/

    booleantype cv_sensi;    /* TRUE if computing sensitivities              */

    int cv_Ns;               /* Number of sensitivities                      */

    int cv_ism;              /* ism = SIMULTANEOUS or STAGGERED              */

    CVSensRhsFn cv_fS;       /* fS = (df/dy)*yS + (df/dp)                    */
    CVSensRhs1Fn cv_fS1;     /* fS1 = (df/dy)*yS_i + (df/dp)                 */
    void *cv_user_fS_data;   /* user data pointer for fS                     */
    void *cv_fS_data;        /* actual data pointer passed to fS             */
    booleantype cv_fSDQ;
    int cv_ifS;              /* ifS = ALLSENS or ONESENS                     */

    realtype *cv_p;          /* parameters in f(t,y,p)                       */
    realtype *cv_pbar;       /* scale factors for parameters                 */
    int *cv_plist;           /* list of sensitivities                        */
    int cv_DQtype;           /* central/forward finite differences           */
    realtype cv_DQrhomax;    /* cut-off value for separate/simultaneous FD   */

    booleantype cv_errconS;  /* TRUE if sensitivities are in err. control    */

    int cv_itolS;
    realtype cv_reltolS;     /* relative tolerance for sensitivities         */
    realtype *cv_SabstolS;   /* scalar absolute tolerances for sensi.        */
    N_Vector *cv_VabstolS;   /* vector absolute tolerances for sensi.        */

    /*-----------------------
      Nordsieck History Array 
      -----------------------*/

    N_Vector cv_zn[L_MAX];   /* Nordsieck array, of size N x (q+1).
                                zn[j] is a vector of length N (j=0,...,q)
                                zn[j] = [1/factorial(j)] * h^j * 
                                (jth derivative of the interpolating 
                                polynomial                                   */

    /*-------------------
      Vectors of length N 
      -------------------*/

    N_Vector cv_ewt;         /* error weight vector                          */
    N_Vector cv_y;           /* y is used as temporary storage by the solver.
                                The memory is provided by the user to CVode 
                                where the vector is named yout.              */
    N_Vector cv_acor;        /* In the context of the solution of the
                                nonlinear equation, acor = y_n(m) - y_n(0).
                                On return, this vector is scaled to give
                                the estimated local error in y.              */
    N_Vector cv_tempv;       /* temporary storage vector                     */
    N_Vector cv_ftemp;       /* temporary storage vector                     */

    /*--------------------------
      Quadrature Related Vectors 
      --------------------------*/

    N_Vector cv_znQ[L_MAX];  /* Nordsieck arrays for quadratures             */
    N_Vector cv_ewtQ;        /* error weight vector for quadratures          */
    N_Vector cv_yQ;          /* Unlike y, yQ is not allocated by the user    */
    N_Vector cv_acorQ;       /* acorQ = yQ_n(m) - yQ_n(0)                    */
    N_Vector cv_tempvQ;      /* temporary storage vector (~ tempv)           */

    /*---------------------------
      Sensitivity Related Vectors 
      ---------------------------*/

    N_Vector *cv_znS[L_MAX]; /* Nordsieck arrays for sensitivities           */
    N_Vector *cv_ewtS;       /* error weight vectors for sensitivities       */
    N_Vector *cv_yS;         /* yS=yS0 (allocated by the user)               */
    N_Vector *cv_acorS;      /* acorS = yS_n(m) - yS_n(0)                    */
    N_Vector *cv_tempvS;     /* temporary storage vector (~ tempv)           */
    N_Vector *cv_ftempS;     /* temporary storage vector (~ ftemp)           */

    /*-----------------------------------------------
      Does CVodeSensMalloc allocate additional space?
      -----------------------------------------------*/  

    booleantype cv_stgr1alloc; /* Did we allocate ncfS1, ncfnS1, and nniS1?  */

    /*-----------------
      Tstop information
      -----------------*/
    booleantype cv_istop;
    booleantype cv_tstopset;
    realtype cv_tstop;

    /*---------
      Step Data 
      ---------*/

    int cv_q;                    /* current order                            */
    int cv_qprime;               /* order to be used on the next step        */ 
    /* = q-1, q, or q+1                         */
    int cv_next_q;               /* order to be used on the next step        */
    int cv_qwait;                /* number of internal steps to wait before  */
    /* considering a change in q                */
    int cv_L;                    /* L = q + 1                                */

    realtype cv_hin;
    realtype cv_h;               /* current step size                        */
    realtype cv_hprime;          /* step size to be used on the next step    */ 
    realtype cv_next_h;          /* step size to be used on the next step    */ 
    realtype cv_eta;             /* eta = hprime / h                         */
    realtype cv_hscale;          /* value of h used in zn                    */
    realtype cv_tn;              /* current internal value of t              */
    realtype cv_tretlast;        /* last value of t returned                 */

    realtype cv_tau[L_MAX+1];    /* array of previous q+1 successful step
                                    sizes indexed from 1 to q+1              */
    realtype cv_tq[NUM_TESTS+1]; /* array of test quantities indexed from
                                    1 to NUM_TESTS(=5)                       */
    realtype cv_l[L_MAX];        /* coefficients of l(x) (degree q poly)     */

    realtype cv_rl1;             /* the scalar 1/l[1]                        */
    realtype cv_gamma;           /* gamma = h * rl1                          */
    realtype cv_gammap;          /* gamma at the last setup call             */
    realtype cv_gamrat;          /* gamma / gammap                           */

    realtype cv_crate;           /* est. corrector conv. rate in Nls         */
    realtype cv_crateS;          /* est. corrector conv. rate in NlsStgr     */
    realtype cv_acnrm;           /* | acor |                                 */
    realtype cv_acnrmS;          /* | acorS |                                */
    realtype cv_acnrmQ;          /* | acorQ |                                */
    realtype cv_nlscoef;         /* coeficient in nonlinear convergence test */
    int  cv_mnewt;               /* Newton iteration counter                 */
    int  *cv_ncfS1;              /* Array of Ns local counters for conv.  
                                    failures (used in CVStep for STAGGERED1) */

    /*------
      Limits 
      ------*/

    int cv_qmax;             /* q <= qmax                                    */
    long int cv_mxstep;      /* maximum number of internal steps for one 
                                user call                                    */
    int cv_maxcor;           /* maximum number of corrector iterations for 
                                the solution of the nonlinear equation       */
    int cv_maxcorS;
    int cv_mxhnil;           /* maximum number of warning messages issued to 
                                the user that t + h == t for the next 
                                internal step                                */
    int cv_maxnef;           /* maximum number of error test failures        */
    int cv_maxncf;           /* maximum number of nonlinear conv. failures   */

    realtype cv_hmin;        /* |h| >= hmin                                  */
    realtype cv_hmax_inv;    /* |h| <= 1/hmax_inv                            */
    realtype cv_etamax;      /* eta <= etamax                                */

    /*----------
      Counters 
      ----------*/

    long int cv_nst;         /* number of internal steps taken               */
    long int cv_nfe;         /* number of f calls                            */
    long int cv_nfSe;        /* number of fS calls                           */
    long int cv_nfQe;        /* number of fQ calls                           */
    long int cv_nfeS;        /* number of f calls from sensi DQ              */

    long int cv_ncfn;        /* number of corrector convergence failures     */
    long int cv_ncfnS;       /* number of total sensi. corr. conv. failures  */
    long int *cv_ncfnS1;     /* number of sensi. corrector conv. failures    */

    long int cv_nni;         /* number of nonlinear iterations performed     */
    long int cv_nniS;        /* number of total sensi. nonlinear iterations  */
    long int *cv_nniS1;      /* number of sensi. nonlinear iterations        */

    long int cv_netf;        /* number of error test failures                */
    long int cv_netfS;       /* number of sensi. error test failures         */
    long int cv_netfQ;       /* number of quadr. error test failures         */

    long int cv_nsetups;     /* number of setup calls                        */
    long int cv_nsetupsS;    /* number of setup calls due to sensitivities   */

    int cv_nhnil;            /* number of messages issued to the user that
                                t + h == t for the next iternal step    */

    /*-----------------------------
      Space requirements for CVODES 
      -----------------------------*/

    long int cv_lrw1;        /* no. of realtype words in 1 N_Vector y        */ 
    long int cv_liw1;        /* no. of integer words in 1 N_Vector y         */ 
    long int cv_lrw1Q;       /* no. of realtype words in 1 N_Vector yQ       */ 
    long int cv_liw1Q;       /* no. of integer words in 1 N_Vector yQ        */ 
    long int cv_lrw;         /* no. of realtype words in CVODES work vectors */
    long int cv_liw;         /* no. of integer words in CVODES work vectors  */

    /*----------------
      Step size ratios
      ----------------*/

    realtype cv_etaqm1;      /* ratio of new to old h for order q-1          */
    realtype cv_etaq;        /* ratio of new to old h for order q            */
    realtype cv_etaqp1;      /* ratio of new to old h for order q+1          */

    /*------------------
      Linear Solver Data 
      ------------------*/

    /* Linear Solver functions to be called */

    int (*cv_linit)(struct CVodeMemRec *cv_mem);

    int (*cv_lsetup)(struct CVodeMemRec *cv_mem, int convfail, 
                     N_Vector ypred, N_Vector fpred, booleantype *jcurPtr, 
                     N_Vector vtemp1, N_Vector vtemp2, N_Vector vtemp3); 

    int (*cv_lsolve)(struct CVodeMemRec *cv_mem, N_Vector b, N_Vector weight,
                     N_Vector ycur, N_Vector fcur);

    void (*cv_lfree)(struct CVodeMemRec *cv_mem);

    /* Linear Solver specific memory */

    void *cv_lmem;           

    /* Flag to request a call to the setup routine */

    booleantype cv_forceSetup;

    /*------------
      Saved Values
      ------------*/

    int cv_qu;             /* last successful q value used                        */
    long int cv_nstlp;     /* step number of last setup call                      */
    realtype cv_h0u;       /* actual initial stepsize                             */
    realtype cv_hu;        /* last successful h value used                        */
    realtype cv_saved_tq5; /* saved value of tq[5]                                */
    booleantype cv_jcur;   /* Is the Jacobian info used by linear solver current? */
    realtype cv_tolsf;     /* tolerance scale factor                              */
    int cv_qmax_alloc;     /* value of qmax used when allocating memory           */
    int cv_qmax_allocQ;    /* value of qmax used when allocating quad. memory     */
    int cv_qmax_allocS;    /* value of qmax used when allocating sensi. memory    */
    int cv_indx_acor;      /* index of the zn vector in which acor is saved       */
    booleantype cv_setupNonNull; /* Does setup do something?                      */

    /*--------------------------------------------------------------------
      Flags turned ON by CVodeMalloc, CVodeSensMalloc, and CVodeQuadMalloc 
      and read by CVodeReInit, CVodeSensReInit, and CVodeQuadReInit
      --------------------------------------------------------------------*/

    booleantype cv_VabstolMallocDone;
    booleantype cv_MallocDone;

    booleantype cv_VabstolQMallocDone;
    booleantype cv_quadMallocDone;

    booleantype cv_VabstolSMallocDone;
    booleantype cv_SabstolSMallocDone;
    booleantype cv_sensMallocDone;

    /*-------------------------------------------
      Error handler function and error ouput file 
      -------------------------------------------*/

    CVErrHandlerFn cv_ehfun;    /* Error messages are handled by ehfun       */
    void *cv_eh_data;           /* user pointer passed to ehfun              */
    FILE *cv_errfp;             /* CVODES error messages are sent to errfp   */    

    /*-------------------------
      Stability Limit Detection
      -------------------------*/

    booleantype cv_sldeton;     /* Is Stability Limit Detection on?          */
    realtype cv_ssdat[6][4];    /* scaled data array for STALD               */
    int cv_nscon;               /* counter for STALD method                  */
    long int cv_nor;            /* counter for number of order reductions    */

    /*----------------
      Rootfinding Data
      ----------------*/

    CVRootFn cv_gfun;     /* Function g for roots sought                     */
    int cv_nrtfn;         /* number of components of g                       */
    void *cv_g_data;      /* pointer to user data for g                      */
    int *cv_iroots;       /* int array for root information                  */
    realtype cv_tlo;      /* nearest endpoint of interval in root search     */
    realtype cv_thi;      /* farthest endpoint of interval in root search    */
    realtype cv_trout;    /* t value returned by rootfinding routine         */
    realtype *cv_glo;     /* saved array of g values at t = tlo              */
    realtype *cv_ghi;     /* saved array of g values at t = thi              */
    realtype *cv_grout;   /* array of g values at t = trout                  */
    realtype cv_toutc;    /* copy of tout (if NORMAL mode)                   */
    realtype cv_ttol;     /* tolerance on root location trout                */
    int cv_taskc;         /* copy of parameter task                          */
    int cv_irfnd;         /* flag showing whether last step had a root       */
    long int cv_nge;      /* counter for g evaluations                       */

  } *CVodeMem;


  /* 
   * =================================================================
   *   A D J O I N T   M O D U L E    M E M O R Y    B L O C K
   * =================================================================
   */

  /*
   * -----------------------------------------------------------------
   * Forward references for pointers to various structures 
   * -----------------------------------------------------------------
   */

  typedef struct CVadjMemRec *CVadjMem;
  typedef struct CkpntMemRec *CkpntMem;
  typedef struct DtpntMemRec *DtpntMem;

  /*
   * -----------------------------------------------------------------
   * Types for functions provided by an interpolation module
   * -----------------------------------------------------------------
   * CVAGetYFn:   Function type for a function that returns the 
   *              interpolated forward solution.
   * CVAStorePnt: Function type for a function that stores a new
   *              point in the structure d
   * -----------------------------------------------------------------
   */

  typedef int (*CVAGetYFn)(CVadjMem ca_mem, realtype t, N_Vector y);
  typedef int (*CVAStorePntFn)(CVodeMem cv_mem, DtpntMem d);

  /*
   * -----------------------------------------------------------------
   * Types : struct CkpntMemRec, CkpntMem
   * -----------------------------------------------------------------
   * The type CkpntMem is type pointer to struct CkpntMemRec.
   * This structure contains fields to store all information at a
   * check point that is needed to 'hot' start cvodes.
   * -----------------------------------------------------------------
   */

  struct CkpntMemRec {

    /* Integration limits */
    realtype ck_t0;
    realtype ck_t1;
    
    /* Nordsieck History Array */
    N_Vector ck_zn[L_MAX];
    
    /* Nordsieck History Array for quadratures */
    N_Vector ck_znQ[L_MAX];
    
    /* Do we need to carry quadratures? */
    booleantype ck_quadr;
    
    /* Was ck_zn[qmax] allocated?
       ck_zqm = 0    - no
       ck_zqm = qmax - yes      */
    int ck_zqm;
    
    /* Step data */
    long int ck_nst;
    realtype ck_tretlast;
    int      ck_q;
    int      ck_qprime;
    int      ck_qwait;
    int      ck_L;
    realtype ck_gammap;
    realtype ck_h;
    realtype ck_hprime;
    realtype ck_hscale;
    realtype ck_eta;
    realtype ck_etamax;
    realtype ck_tau[L_MAX+1];
    realtype ck_tq[NUM_TESTS+1];
    realtype ck_l[L_MAX];
    
    /* Saved values */
    realtype ck_saved_tq5;
    
    /* Pointer to next structure in list */
    struct CkpntMemRec *ck_next;
    
  };
  
  /*
   * -----------------------------------------------------------------
   * Type : struct DtpntMemRec
   * -----------------------------------------------------------------
   * This structure contains fields to store all information at a
   * data point that is needed to interpolate solution of forward
   * simulations. Its content field is interpType-dependent.
   * -----------------------------------------------------------------
   */
  
  struct DtpntMemRec {
    realtype t;    /* time */
    void *content; /* interpType-dependent content */
  };

  /* Data for cubic Hermite interpolation */
  typedef struct HermiteDataMemRec {
    N_Vector y;
    N_Vector yd;
  } *HermiteDataMem;

  /* Data for polynomial interpolation */
  typedef struct PolynomialDataMemRec {
    N_Vector y;
    int order;
  } *PolynomialDataMem;

  /*
   * -----------------------------------------------------------------
   * Type : struct CVadjMemRec
   * -----------------------------------------------------------------
   * The type CVadjMem is type pointer to struct CVadjMemRec.
   * This structure contins fields to store all information
   * necessary for adjoint sensitivity analysis.
   * -----------------------------------------------------------------
   */

  struct CVadjMemRec {
    
    /* CVODE memory for forward runs */
    struct CVodeMemRec *cv_mem;
    
    /* CVODE memory for backward run */
    struct CVodeMemRec *cvb_mem;
    
    /* Storage for check point information */
    struct CkpntMemRec *ck_mem;

    /* Interpolation type */
    int ca_interpType;

    /* Storage for data from forward runs */
    struct DtpntMemRec **dt_mem;
    
    /* Functions set by the interpolation module */
    CVAStorePntFn ca_storePnt; /* store a new interpolation point */
    CVAGetYFn     ca_getY;     /* interpolate forward solution    */
    
    /* Right hand side function (fB) for backward run */
    CVRhsFnB ca_fB;
    
    /* Right hand side quadrature function (fQB) for backward run */
    CVQuadRhsFnB ca_fQB;

    /* User f_dataB */
    void *ca_f_dataB;
    
    /* User fQ_dataB */
    void *ca_fQ_dataB;
    
    /* Memory block for a linear solver's interface to CVODEA */
    void *ca_lmemB;

    /* Function to free any memory allocated by the linear solver */
    void (*ca_lfreeB)(CVadjMem ca_mem);

    /* Memory block for a preconditioner's module interface to CVODEA */ 
    void *ca_pmemB;
    
    /* Unit roundoff */
    realtype ca_uround;
    
    /* Integration interval */
    realtype ca_tinitial, ca_tfinal;
    
    /* Time at which to extract quadratures */
    realtype ca_t_for_quad;
    
    /* Number of check points */
    int ca_nckpnts;
    
    /* Number of steps between 2 check points */
    long int ca_nsteps;
    
    /* Flag to indicate that data in dt_mem is new */
    booleantype ca_newData;
    
    /* address of the check point structure for which data is available */
    struct CkpntMemRec *ca_ckpntData;
    
    /* Actual number of data points saved in current dt_mem */
    /* Commonly, np = nsteps+1                              */
    long int ca_np;
    
    /* Workspace used by the Hermite interpolation */
    N_Vector ca_Y0, ca_Y1;    /* pointers to zn[0] and zn[1] */

    /* Workspace for polynomial interpolation */
    N_Vector ca_Y[L_MAX];     /* pointers to zn[i] */
    realtype ca_T[L_MAX];

    /* Workspace for wrapper functions */
    N_Vector ca_ytmp;
    
  };
  

  /*
   * =================================================================
   *     I N T E R F A C E   T O    L I N E A R   S O L V E R S
   * =================================================================
   */

  /*
   * -----------------------------------------------------------------
   * Communication between CVODE and a CVODE Linear Solver
   * -----------------------------------------------------------------
   * convfail (input to cv_lsetup)
   *
   * CV_NO_FAILURES : Either this is the first cv_setup call for this
   *                  step, or the local error test failed on the
   *                  previous attempt at this step (but the Newton
   *                  iteration converged).
   *
   * CV_FAIL_BAD_J  : This value is passed to cv_lsetup if
   *
   *                  (a) The previous Newton corrector iteration
   *                      did not converge and the linear solver's
   *                      setup routine indicated that its Jacobian-
   *                      related data is not current
   *                                   or
   *                  (b) During the previous Newton corrector
   *                      iteration, the linear solver's solve routine
   *                      failed in a recoverable manner and the
   *                      linear solver's setup routine indicated that
   *                      its Jacobian-related data is not current.
   *
   * CV_FAIL_OTHER  : During the current internal step try, the
   *                  previous Newton iteration failed to converge
   *                  even though the linear solver was using current
   *                  Jacobian-related data.
   * -----------------------------------------------------------------
   */

  /* Constants for convfail (input to cv_lsetup) */

#define CV_NO_FAILURES 0
#define CV_FAIL_BAD_J  1
#define CV_FAIL_OTHER  2

  /*
   * -----------------------------------------------------------------
   * int (*cv_linit)(CVodeMem cv_mem);
   * -----------------------------------------------------------------
   * The purpose of cv_linit is to complete initializations for a
   * specific linear solver, such as counters and statistics.
   * An LInitFn should return 0 if it has successfully initialized the
   * CVODE linear solver and a negative value otherwise.
   * If an error does occur, an appropriate message should be sent to
   * the error handler function.
   * -----------------------------------------------------------------
   */

  /*
   * -----------------------------------------------------------------
   * int (*cv_lsetup)(CVodeMem cv_mem, int convfail, N_Vector ypred,
   *                 N_Vector fpred, booleantype *jcurPtr,
   *                 N_Vector vtemp1, N_Vector vtemp2,
   *                 N_Vector vtemp3);
   * -----------------------------------------------------------------
   * The job of cv_lsetup is to prepare the linear solver for
   * subsequent calls to cv_lsolve. It may recompute Jacobian-
   * related data is it deems necessary. Its parameters are as
   * follows:
   *
   * cv_mem - problem memory pointer of type CVodeMem. See the
   *          typedef earlier in this file.
   *
   * convfail - a flag to indicate any problem that occurred during
   *            the solution of the nonlinear equation on the
   *            current time step for which the linear solver is
   *            being used. This flag can be used to help decide
   *            whether the Jacobian data kept by a CVODE linear
   *            solver needs to be updated or not.
   *            Its possible values have been documented above.
   *
   * ypred - the predicted y vector for the current CVODE internal
   *         step.
   *
   * fpred - f(tn, ypred).
   *
   * jcurPtr - a pointer to a boolean to be filled in by cv_lsetup.
   *           The function should set *jcurPtr=TRUE if its Jacobian
   *           data is current after the call and should set
   *           *jcurPtr=FALSE if its Jacobian data is not current.
   *           Note: If cv_lsetup calls for re-evaluation of
   *           Jacobian data (based on convfail and CVODE state
   *           data), it should return *jcurPtr=TRUE always;
   *           otherwise an infinite loop can result.
   *
   * vtemp1 - temporary N_Vector provided for use by cv_lsetup.
   *
   * vtemp3 - temporary N_Vector provided for use by cv_lsetup.
   *
   * vtemp3 - temporary N_Vector provided for use by cv_lsetup.
   *
   * The cv_lsetup routine should return 0 if successful, a positive
   * value for a recoverable error, and a negative value for an
   * unrecoverable error.
   * -----------------------------------------------------------------
   */

  /*
   * -----------------------------------------------------------------
   * int (*cv_lsolve)(CVodeMem cv_mem, N_Vector b, N_Vector weight,
   *                  N_Vector ycur, N_Vector fcur);
   * -----------------------------------------------------------------
   * cv_lsolve must solve the linear equation P x = b, where
   * P is some approximation to (I - gamma J), J = (df/dy)(tn,ycur)
   * and the RHS vector b is input. The N-vector ycur contains
   * the solver's current approximation to y(tn) and the vector
   * fcur contains the N_Vector f(tn,ycur). The solution is to be
   * returned in the vector b. cv_lsolve returns a positive value
   * for a recoverable error and a negative value for an
   * unrecoverable error. Success is indicated by a 0 return value.
   * -----------------------------------------------------------------
   */

  /*
   * -----------------------------------------------------------------
   * void (*cv_lfree)(CVodeMem cv_mem);
   * -----------------------------------------------------------------
   * cv_lfree should free up any memory allocated by the linear
   * solver. This routine is called once a problem has been
   * completed and the linear solver is no longer needed.
   * -----------------------------------------------------------------
   */

  /*
   * =================================================================
   *   C V O D E S    I N T E R N A L   F U N C T I O N S
   * =================================================================
   */
  
  /* Prototype of internal ewtSet function */

  int CVEwtSet(N_Vector ycur, N_Vector weight, void *e_data);

  /* High level error handler */

  void CVProcessError(CVodeMem cv_mem, 
                      int error_code, const char *module, const char *fname, 
                      const char *msgfmt, ...);

  /* Prototype of internal errHandler function */

  void CVErrHandler(int error_code, const char *module, const char *function, 
                    char *msg, void *eh_data);

  /* Prototypes for internal sensitivity rhs DQ functions */

  int CVSensRhsDQ(int Ns, realtype t, 
                  N_Vector y, N_Vector ydot, 
                  N_Vector *yS, N_Vector *ySdot, 
                  void *fS_data,  
                  N_Vector tempv, N_Vector ftemp);

  int CVSensRhs1DQ(int Ns, realtype t, 
                   N_Vector y, N_Vector ydot, 
                   int is, N_Vector yS, N_Vector ySdot, 
                   void *fS_data,
                   N_Vector tempv, N_Vector ftemp);


  /* 
   * =================================================================
   *   C V O D E S    E R R O R    M E S S A G E S
   * =================================================================
   */

#if defined(SUNDIALS_EXTENDED_PRECISION)

#define MSG_TIME      "t = %Lg"
#define MSG_TIME_H    "t = %Lg and h = %Lg"
#define MSG_TIME_INT  "t = %Lg is not between tcur - hu = %Lg and tcur = %Lg."
#define MSG_TIME_TOUT "tout = %Lg"

#elif defined(SUNDIALS_DOUBLE_PRECISION)

#define MSG_TIME      "t = %lg"
#define MSG_TIME_H    "t = %lg and h = %lg"
#define MSG_TIME_INT  "t = %lg is not between tcur - hu = %lg and tcur = %lg."
#define MSG_TIME_TOUT "tout = %lg"

#else

#define MSG_TIME      "t = %g"
#define MSG_TIME_H    "t = %g and h = %g"
#define MSG_TIME_INT  "t = %g is not between tcur - hu = %g and tcur = %g."
#define MSG_TIME_TOUT "tout = %g"

#endif


/* Initialization and I/O error messages */

#define MSGCV_NO_MEM "cvode_mem = NULL illegal."
#define MSGCV_CVMEM_FAIL "Allocation of cvode_mem failed."
#define MSGCV_MEM_FAIL "A memory request failed."
#define MSGCV_BAD_LMM  "Illegal value for lmm. The legal values are CV_ADAMS and CV_BDF."
#define MSGCV_BAD_ITER  "Illegal value for iter. The legal values are CV_FUNCTIONAL and CV_NEWTON."
#define MSGCV_BAD_ITOL "Illegal value for itol. The legal values are CV_SS, CV_SV, and CV_WF."
#define MSGCV_NO_MALLOC "Attempt to call before CVodeMalloc."
#define MSGCV_NEG_MAXORD "maxord <= 0 illegal."
#define MSGCV_BAD_MAXORD  "Illegal attempt to increase maximum method order."
#define MSGCV_NEG_MXSTEPS "mxsteps < 0 illegal."
#define MSGCV_SET_SLDET  "Attempt to use stability limit detection with the CV_ADAMS method illegal."
#define MSGCV_NEG_HMIN "hmin < 0 illegal."
#define MSGCV_NEG_HMAX "hmax < 0 illegal."
#define MSGCV_BAD_HMIN_HMAX "Inconsistent step size limits: hmin > hmax."
#define MSGCV_BAD_RELTOL "reltol < 0 illegal."
#define MSGCV_BAD_ABSTOL "abstol has negative component(s) (illegal)."
#define MSGCV_NULL_ABSTOL "abstol = NULL illegal."
#define MSGCV_NULL_Y0 "y0 = NULL illegal."
#define MSGCV_NULL_F "f = NULL illegal."
#define MSGCV_NULL_G "g = NULL illegal."
#define MSGCV_BAD_NVECTOR "A required vector operation is not implemented."
#define MSGCV_BAD_K "Illegal value for k."
#define MSGCV_NULL_DKY "dky = NULL illegal."
#define MSGCV_BAD_T "Illegal value for t." MSG_TIME_INT

#define MSGCV_NO_QUAD  "Illegal attempt to call before calling CVodeQuadMalloc."
#define MSGCV_BAD_ITOLQ "Illegal value for itolQ. The legal values are CV_SS and CV_SV."
#define MSGCV_NULL_ABSTOLQ "abstolQ = NULL illegal."
#define MSGCV_BAD_RELTOLQ "reltolQ < 0 illegal."
#define MSGCV_BAD_ABSTOLQ "abstolQ has negative component(s) (illegal)."  

#define MSGCV_NO_SENSI  "Illegal attempt to call before calling CVodeSensMalloc."
#define MSGCV_BAD_ITOLS "Illegal value for itolS. The legal values are CV_SS, CV_SV, and CV_EE."
#define MSGCV_NULL_ABSTOLS "abstolS = NULL illegal."
#define MSGCV_BAD_RELTOLS "reltolS < 0 illegal."
#define MSGCV_BAD_ABSTOLS "abstolS has negative component(s) (illegal)."  
#define MSGCV_BAD_PBAR "pbar has zero component(s) (illegal)."
#define MSGCV_BAD_PLIST "plist has negative component(s) (illegal)."
#define MSGCV_BAD_NS "NS <= 0 illegal."
#define MSGCV_NULL_YS0 "yS0 = NULL illegal."
#define MSGCV_BAD_ISM "Illegal value for ism. Legal values are: CV_SIMULTANEOUS, CV_STAGGERED and CV_STAGGERED1."
#define MSGCV_BAD_IS "Illegal value for is."
#define MSGCV_NULL_DKYA "dkyA = NULL illegal."
#define MSGCV_BAD_DQTYPE "Illegal value for DQtype. Legal values are: CV_CENTERED and CV_FORWARD."
#define MSGCV_BAD_DQRHO "DQrhomax < 0 illegal."

/* CVode Error Messages */

#define MSGCV_LSOLVE_NULL "The linear solver's solve routine is NULL."
#define MSGCV_YOUT_NULL "yout = NULL illegal."
#define MSGCV_TRET_NULL "tret = NULL illegal."
#define MSGCV_BAD_EWT "Initial ewt has component(s) equal to zero (illegal)."
#define MSGCV_EWT_NOW_BAD "At " MSG_TIME ", a component of ewt has become <= 0."
#define MSGCV_BAD_ITASK "Illegal value for itask."
#define MSGCV_BAD_H0 "h0 and tout - t0 inconsistent."
#define MSGCV_BAD_INIT_ROOT "Root found at and very near initial t."
#define MSGCV_BAD_TOUT "Trouble interpolating at " MSG_TIME_TOUT ". tout too far back in direction of integration"
#define MSGCV_NO_EFUN "itol = CV_WF but no EwtSet function was provided."
#define MSGCV_NO_TSTOP "itask = CV_NORMAL_TSTOP or itask = CV_ONE_STEP_TSTOP but tstop was not set."
#define MSGCV_EWT_FAIL "The user-provide EwtSet function failed."
#define MSGCV_EWT_NOW_FAIL "At " MSG_TIME ", the user-provide EwtSet function failed."
#define MSGCV_LINIT_FAIL "The linear solver's init routine failed."
#define MSGCV_HNIL_DONE "The above warning has been issued mxhnil times and will not be issued again for this problem."
#define MSGCV_TOO_CLOSE "tout too close to t0 to start integration."
#define MSGCV_MAX_STEPS "At " MSG_TIME ", mxstep steps taken before reaching tout."
#define MSGCV_TOO_MUCH_ACC "At " MSG_TIME ", too much accuracy requested."
#define MSGCV_HNIL "Internal " MSG_TIME_H " are such that t + h = t on the next step. The solver will continue anyway."
#define MSGCV_ERR_FAILS "At " MSG_TIME_H ", the error test failed repeatedly or with |h| = hmin."
#define MSGCV_CONV_FAILS "At " MSG_TIME_H ", the corrector convergence test failed repeatedly or with |h| = hmin."
#define MSGCV_SETUP_FAILED "At " MSG_TIME ", the setup routine failed in an unrecoverable manner."
#define MSGCV_SOLVE_FAILED "At " MSG_TIME ", the solve routine failed in an unrecoverable manner."
#define MSGCV_RHSFUNC_FAILED "At " MSG_TIME ", the right-hand side routine failed in an unrecoverable manner."
#define MSGCV_RHSFUNC_UNREC "At " MSG_TIME ", the right-hand side failed in a recoverable manner, but no recovery is possible."
#define MSGCV_RHSFUNC_REPTD "At " MSG_TIME "repeated recoverable right-hand side function errors."
#define MSGCV_RHSFUNC_FIRST "The right-hand side routine failed at the first call."
#define MSGCV_RTFUNC_FAILED "At " MSG_TIME ", the rootfinding routine failed in an unrecoverable manner."
#define MSGCV_CLOSE_ROOTS "Root found at and very near " MSG_TIME "."
#define MSGCV_BAD_TSTOP "tstop is behind current " MSG_TIME "in the direction of integration."

#define MSGCV_BAD_EWTQ "Initial ewtQ has component(s) equal to zero (illegal)."
#define MSGCV_EWTQ_NOW_BAD "At " MSG_TIME ", a component of ewtQ has become <= 0."
#define MSGCV_QRHSFUNC_FAILED "At " MSG_TIME ", the quadrature right-hand side routine failed in an unrecoverable manner."
#define MSGCV_QRHSFUNC_UNREC "At " MSG_TIME ", the quadrature right-hand side failed in a recoverable manner, but no recovery is possible."
#define MSGCV_QRHSFUNC_REPTD "At " MSG_TIME "repeated recoverable quadrature right-hand side function errors."
#define MSGCV_QRHSFUNC_FIRST "The quadrature right-hand side routine failed at the first call."

#define MSGCV_BAD_ISM_IFS "Illegal sens. rhs for ism = CV_STAGGERED1."
#define MSGCV_NULL_P "p = NULL when using internal DQ for sensitivity RHS illegal."
#define MSGCV_BAD_EWTS "Initial ewtS has component(s) equal to zero (illegal)."
#define MSGCV_EWTS_NOW_BAD "At " MSG_TIME ", a component of ewtS has become <= 0."
#define MSGCV_SRHSFUNC_FAILED "At " MSG_TIME ", the sensitivity right-hand side routine failed in an unrecoverable manner."
#define MSGCV_SRHSFUNC_UNREC "At " MSG_TIME ", the sensitivity right-hand side failed in a recoverable manner, but no recovery is possible."
#define MSGCV_SRHSFUNC_REPTD "At " MSG_TIME "repeated recoverable sensitivity right-hand side function errors."
#define MSGCV_SRHSFUNC_FIRST "The sensitivity right-hand side routine failed at the first call."

  /* 
   * =================================================================
   *   C V O D E A    E R R O R    M E S S A G E S
   * =================================================================
   */

#define MSGAM_NULL_CVMEM   "cvode_mem = NULL illegal."
#define MSGAM_NULL_CAMEM   "cvadj_mem = NULL illegal."
#define MSGAM_BAD_STEPS    "Steps nonpositive illegal."
#define MSGAM_MEM_FAIL     "A memory request failed."
#define MSGAM_BAD_INTERP   "Illegal value for interp."
#define MSGAM_BAD_ITASKB   "Illegal value for itaskB. Legal values are CV_NORMAL and CV_ONE_STEP."
#define MSGAM_BAD_TB0      "The initial time tB0 is outside the interval over which the forward problem was solved."
#define MSGAM_BAD_TBOUT    "The final time tBout is outside the interval over which the forward problem was solved."
#define MSGAM_BAD_T        "Bad t for interpolation."
#define MSGAM_WRONG_INTERP "This function cannot be called for the specified interp type."


#ifdef __cplusplus
}
#endif

#endif
```
