## Supplementary scripts for "A multi-scale *in silico* mouse model for insulin resistance and humanoid type 2 diabetes": cvodes_io.c

```
/*
 * -----------------------------------------------------------------
 * $Revision: 749 $
 * $Date: 2011-06-22 14:49:10 +0200 (Wed, 22 Jun 2011) $
 * -----------------------------------------------------------------
 * Programmer(s): Alan C. Hindmarsh and Radu Serban @ LLNL
 * -----------------------------------------------------------------
 * Copyright (c) 2005, The Regents of the University of California.
 * Produced at the Lawrence Livermore National Laboratory.
 * All rights reserved.
 * For details, see the LICENSE file.
 * -----------------------------------------------------------------
 * This is the implementation file for the optional input and output
 * functions for the CVODES solver.
 * -----------------------------------------------------------------
 */

#include <stdio.h>
#include <stdlib.h>

#include "cvodes_impl.h"

#include <sundials/sundials_math.h>
#include <sundials/sundials_types.h>

#define ZERO RCONST(0.0)
#define ONE  RCONST(1.0)

/* 
 * =================================================================
 * CVODES optional input functions
 * =================================================================
 */

/* 
 * Readability constants
 */

#define lrw   (cv_mem->cv_lrw)
#define liw   (cv_mem->cv_liw)
#define lrw1  (cv_mem->cv_lrw1)
#define liw1  (cv_mem->cv_liw1)

/* 
 * CVodeSetErrHandlerFn
 *
 * Specifies the error handler function
 */

int CVodeSetErrHandlerFn(void *cvode_mem, CVErrHandlerFn ehfun, void *eh_data)
{
  CVodeMem cv_mem;

  if (cvode_mem==NULL) {
    CVProcessError(NULL, CV_MEM_NULL, "CVODES", "CVodeSetErrHandlerFn", MSGCV_NO_MEM);
    return(CV_MEM_NULL);
  }

  cv_mem = (CVodeMem) cvode_mem;

  cv_mem->cv_ehfun = ehfun;
  cv_mem->cv_eh_data = eh_data;

  return(CV_SUCCESS);
}

/* 
 * CVodeSetErrFile
 *
 * Specifies the FILE pointer for output (NULL means no messages)
 */

int CVodeSetErrFile(void *cvode_mem, FILE *errfp)
{
  CVodeMem cv_mem;

  if (cvode_mem==NULL) {
    CVProcessError(NULL, CV_MEM_NULL, "CVODES", "CVodeSetErrFile", MSGCV_NO_MEM);
    return(CV_MEM_NULL);
  }

  cv_mem = (CVodeMem) cvode_mem;

  cv_mem->cv_errfp = errfp;

  return(CV_SUCCESS);
}

/* 
 * CVodeSetIterType
 *
 * Specifies the iteration type (CV_FUNCTIONAL or CV_NEWTON)
 */

int CVodeSetIterType(void *cvode_mem, int iter)
{
  CVodeMem cv_mem;

  if (cvode_mem==NULL) {
    CVProcessError(NULL, CV_MEM_NULL, "CVODES", "CVodeSetIterType", MSGCV_NO_MEM);
    return(CV_MEM_NULL);
  }

  cv_mem = (CVodeMem) cvode_mem;

  if ((iter != CV_FUNCTIONAL) && (iter != CV_NEWTON)) {
    CVProcessError(cv_mem, CV_ILL_INPUT, "CVODES", "CVodeSetIterType", MSGCV_BAD_ITER);
    return (CV_ILL_INPUT);
  }

  cv_mem->cv_iter = iter;

  return(CV_SUCCESS);
}

/* 
 * CVodeSetFdata
 *
 * Specifies the user data pointer for f
 */

int CVodeSetFdata(void *cvode_mem, void *f_data)
{
  CVodeMem cv_mem;

  if (cvode_mem==NULL) {
    CVProcessError(NULL, CV_MEM_NULL, "CVODES", "CVodeSetFdata", MSGCV_NO_MEM);
    return(CV_MEM_NULL);
  }

  cv_mem = (CVodeMem) cvode_mem;

  cv_mem->cv_f_data = f_data;

  return(CV_SUCCESS);
}

/* 
 * CVodeSetMaxOrd
 *
 * Specifies the maximum method order
 */

int CVodeSetMaxOrd(void *cvode_mem, int maxord)
{
  CVodeMem cv_mem;
  int qmax_alloc;

  if (cvode_mem==NULL) {
    CVProcessError(NULL, CV_MEM_NULL, "CVODES", "CVodeSetMaxOrd", MSGCV_NO_MEM);
    return(CV_MEM_NULL);
  }

  cv_mem = (CVodeMem) cvode_mem;

  if (maxord <= 0) {
    CVProcessError(cv_mem, CV_ILL_INPUT, "CVODES", "CVodeSetMaxOrd", MSGCV_NEG_MAXORD);
    return(CV_ILL_INPUT);
  }
  
  /* Cannot increase maximum order beyond the value that
     was used when allocating memory */
  qmax_alloc = cv_mem->cv_qmax_alloc;
  qmax_alloc = MIN(qmax_alloc, cv_mem->cv_qmax_allocQ);
  qmax_alloc = MIN(qmax_alloc, cv_mem->cv_qmax_allocS);  

  if (maxord > qmax_alloc) {
    CVProcessError(cv_mem, CV_ILL_INPUT, "CVODES", "CVodeSetMaxOrd", MSGCV_BAD_MAXORD);
    return(CV_ILL_INPUT);
  }

  cv_mem->cv_qmax = maxord;

  return(CV_SUCCESS);
}

/* 
 * CVodeSetMaxNumSteps
 *
 * Specifies the maximum number of integration steps
 */

int CVodeSetMaxNumSteps(void *cvode_mem, long int mxsteps)
{
  CVodeMem cv_mem;

  if (cvode_mem==NULL) {
    CVProcessError(NULL, CV_MEM_NULL, "CVODES", "CVodeSetMaxNumSteps", MSGCV_NO_MEM);
    return(CV_MEM_NULL);
  }

  cv_mem = (CVodeMem) cvode_mem;

  if (mxsteps < 0) {
    CVProcessError(cv_mem, CV_ILL_INPUT, "CVODES", "CVodeSetMaxNumSteps", MSGCV_NEG_MXSTEPS);
    return(CV_ILL_INPUT);
  }

  /* Passing 0 sets the default */
  if (mxsteps == 0)
    cv_mem->cv_mxstep = MXSTEP_DEFAULT;
  else
    cv_mem->cv_mxstep = mxsteps;

  return(CV_SUCCESS);
}

/* 
 * CVodeSetMaxHnilWarns
 *
 * Specifies the maximum number of warnings for small h
 */

int CVodeSetMaxHnilWarns(void *cvode_mem, int mxhnil)
{
  CVodeMem cv_mem;

  if (cvode_mem==NULL) {
    CVProcessError(NULL, CV_MEM_NULL, "CVODES", "CVodeSetMaxHnilWarns", MSGCV_NO_MEM);
    return(CV_MEM_NULL);
  }

  cv_mem = (CVodeMem) cvode_mem;

  cv_mem->cv_mxhnil = mxhnil;

  return(CV_SUCCESS);
}

/* 
 *CVodeSetStabLimDet
 *
 * Turns on/off the stability limit detection algorithm
 */

int CVodeSetStabLimDet(void *cvode_mem, booleantype sldet)
{
  CVodeMem cv_mem;

  if (cvode_mem==NULL) {
    CVProcessError(NULL, CV_MEM_NULL, "CVODES", "CVodeSetStabLimDet", MSGCV_NO_MEM);
    return(CV_MEM_NULL);
  }

  cv_mem = (CVodeMem) cvode_mem;

  if( sldet && (cv_mem->cv_lmm != CV_BDF) ) {
    CVProcessError(cv_mem, CV_ILL_INPUT, "CVODES", "CVodeSetStabLimDet", MSGCV_SET_SLDET);
    return(CV_ILL_INPUT);
  }

  cv_mem->cv_sldeton = sldet;

  return(CV_SUCCESS);
}

/* 
 * CVodeSetInitStep
 *
 * Specifies the initial step size
 */

int CVodeSetInitStep(void *cvode_mem, realtype hin)
{
  CVodeMem cv_mem;

  if (cvode_mem==NULL) {
    CVProcessError(NULL, CV_MEM_NULL, "CVODES", "CVodeSetInitStep", MSGCV_NO_MEM);
    return(CV_MEM_NULL);
  }

  cv_mem = (CVodeMem) cvode_mem;

  cv_mem->cv_hin = hin;

  return(CV_SUCCESS);
}

/* 
 * CVodeSetMinStep
 *
 * Specifies the minimum step size
 */

int CVodeSetMinStep(void *cvode_mem, realtype hmin)
{
  CVodeMem cv_mem;

  if (cvode_mem==NULL) {
    CVProcessError(NULL, CV_MEM_NULL, "CVODES", "CVodeSetMinStep", MSGCV_NO_MEM);
    return(CV_MEM_NULL);
  }

  cv_mem = (CVodeMem) cvode_mem;

  if (hmin<0) {
    CVProcessError(cv_mem, CV_ILL_INPUT, "CVODES", "CVodeSetMinStep", MSGCV_NEG_HMIN);
    return(CV_ILL_INPUT);
  }

  /* Passing 0 sets hmin = zero */
  if (hmin == ZERO) {
    cv_mem->cv_hmin = HMIN_DEFAULT;
    return(CV_SUCCESS);
  }

  if (hmin * cv_mem->cv_hmax_inv > ONE) {
    CVProcessError(cv_mem, CV_ILL_INPUT, "CVODES", "CVodeSetMinStep", MSGCV_BAD_HMIN_HMAX);
    return(CV_ILL_INPUT);
  }

  cv_mem->cv_hmin = hmin;

  return(CV_SUCCESS);
}

/* 
 * CVodeSetMaxStep
 *
 * Specifies the maximum step size
 */

int CVodeSetMaxStep(void *cvode_mem, realtype hmax)
{
  realtype hmax_inv;
  CVodeMem cv_mem;

  if (cvode_mem==NULL) {
    CVProcessError(NULL, CV_MEM_NULL, "CVODES", "CVodeSetMaxStep", MSGCV_NO_MEM);
    return (CV_MEM_NULL);
  }

  cv_mem = (CVodeMem) cvode_mem;

  if (hmax < 0) {
    CVProcessError(cv_mem, CV_ILL_INPUT, "CVODES", "CVodeSetMaxStep", MSGCV_NEG_HMAX);
    return(CV_ILL_INPUT);
  }

  /* Passing 0 sets hmax = infinity */
  if (hmax == ZERO) {
    cv_mem->cv_hmax_inv = HMAX_INV_DEFAULT;
    return(CV_SUCCESS);
  }

  hmax_inv = ONE/hmax;
  if (hmax_inv * cv_mem->cv_hmin > ONE) {
    CVProcessError(cv_mem, CV_ILL_INPUT, "CVODES", "CVodeSetMaxStep", MSGCV_BAD_HMIN_HMAX);
    return(CV_ILL_INPUT);
  }

  cv_mem->cv_hmax_inv = hmax_inv;

  return(CV_SUCCESS);
}

/* 
 * CVodeSetStopTime
 *
 * Specifies the time beyond which the integration is not to
 * proceed
 */

int CVodeSetStopTime(void *cvode_mem, realtype tstop)
{
  CVodeMem cv_mem;

  if (cvode_mem==NULL) {
    CVProcessError(NULL, CV_MEM_NULL, "CVODES", "CVodeSetStopTime", MSGCV_NO_MEM);
    return (CV_MEM_NULL);
  }

  cv_mem = (CVodeMem) cvode_mem;

  cv_mem->cv_tstop = tstop;
  cv_mem->cv_tstopset = TRUE;

  return(CV_SUCCESS);
}

/* 
 * CVodeSetMaxErrTestFails
 *
 * Specifies the maximum number of error test failures during one
 * step try.
 */

int CVodeSetMaxErrTestFails(void *cvode_mem, int maxnef)
{
  CVodeMem cv_mem;

  if (cvode_mem==NULL) {
    CVProcessError(NULL, CV_MEM_NULL, "CVODES", "CVodeSetMaxErrTestFails", MSGCV_NO_MEM);
    return (CV_MEM_NULL);
  }

  cv_mem = (CVodeMem) cvode_mem;

  cv_mem->cv_maxnef = maxnef;

  return(CV_SUCCESS);
}

/* 
 * CVodeSetMaxConvFails
 *
 * Specifies the maximum number of nonlinear convergence failures 
 * during one step try.
 */

int CVodeSetMaxConvFails(void *cvode_mem, int maxncf)
{
  CVodeMem cv_mem;

  if (cvode_mem==NULL) {
    CVProcessError(NULL, CV_MEM_NULL, "CVODES", "CVodeSetMaxConvFails", MSGCV_NO_MEM);
    return (CV_MEM_NULL);
  }

  cv_mem = (CVodeMem) cvode_mem;

  cv_mem->cv_maxncf = maxncf;

  return(CV_SUCCESS);
}

/* 
 * CVodeSetMaxNonlinIters
 *
 * Specifies the maximum number of nonlinear iterations during
 * one solve.
 */

int CVodeSetMaxNonlinIters(void *cvode_mem, int maxcor)
{
  CVodeMem cv_mem;

  if (cvode_mem==NULL) {
    CVProcessError(NULL, CV_MEM_NULL, "CVODES", "CVodeSetMaxNonlinIters", MSGCV_NO_MEM);
    return (CV_MEM_NULL);
  }

  cv_mem = (CVodeMem) cvode_mem;

  cv_mem->cv_maxcor = maxcor;

  return(CV_SUCCESS);
}

/* 
 * CVodeSetNonlinConvCoef
 *
 * Specifies the coeficient in the nonlinear solver convergence
 * test
 */

int CVodeSetNonlinConvCoef(void *cvode_mem, realtype nlscoef)
{
  CVodeMem cv_mem;

  if (cvode_mem==NULL) {
    CVProcessError(NULL, CV_MEM_NULL, "CVODES", "CVodeSetNonlinConvCoef", MSGCV_NO_MEM);
    return(CV_MEM_NULL);
  }

  cv_mem = (CVodeMem) cvode_mem;

  cv_mem->cv_nlscoef = nlscoef;

  return(CV_SUCCESS);
}

/*
 * CVodeSetTolerances
 *
 * Changes the integration tolerances between calls to CVode()
 */

int CVodeSetTolerances(void *cvode_mem, 
                       int itol, realtype reltol, void *abstol)
{
  CVodeMem cv_mem;
  booleantype neg_abstol;

  if (cvode_mem==NULL) {
    CVProcessError(NULL, CV_MEM_NULL, "CVODES", "CVodeSetTolerances", MSGCV_NO_MEM);
    return(CV_MEM_NULL);
  }

  cv_mem = (CVodeMem) cvode_mem;

  /* Check if cvode_mem was allocated */

  if (cv_mem->cv_MallocDone == FALSE) {
    CVProcessError(cv_mem, CV_NO_MALLOC, "CVODES", "CVodeSetTolerances", MSGCV_NO_MALLOC);
    return(CV_NO_MALLOC);
  }

  /* Check inputs */

  if ( (itol != CV_SS) && (itol != CV_SV) ) {
    CVProcessError(cv_mem, CV_ILL_INPUT, "CVODES", "CVodeSetTolerances", MSGCV_BAD_ITOL);
    return(CV_ILL_INPUT);
  }

  if (abstol == NULL) {
    CVProcessError(cv_mem, CV_ILL_INPUT, "CVODES", "CVodeSetTolerances", MSGCV_NULL_ABSTOL);
    return(CV_ILL_INPUT);
  }

  if (reltol < ZERO) {
    CVProcessError(cv_mem, CV_ILL_INPUT, "CVODES", "CVodeSetTolerances", MSGCV_BAD_RELTOL);
    return(CV_ILL_INPUT);
  }

  if (itol == CV_SS)
    neg_abstol = (*((realtype *)abstol) < ZERO);
  else
    neg_abstol = (N_VMin((N_Vector)abstol) < ZERO);
    
  if (neg_abstol) {
    CVProcessError(cv_mem, CV_ILL_INPUT, "CVODES", "CVodeSetTolerances", MSGCV_BAD_ABSTOL);
    return(CV_ILL_INPUT);
  }

  /* Copy tolerances into memory */

  if ( (itol != CV_SV) && (cv_mem->cv_VabstolMallocDone) ) {
    N_VDestroy(cv_mem->cv_Vabstol);
    lrw -= lrw1;
    liw -= liw1;
    cv_mem->cv_VabstolMallocDone = FALSE;
  }

  if ( (itol == CV_SV) && !(cv_mem->cv_VabstolMallocDone) ) {
    cv_mem->cv_Vabstol = NULL;
    cv_mem->cv_Vabstol = N_VClone(cv_mem->cv_ewt);
    lrw += lrw1;
    liw += liw1;
    cv_mem->cv_VabstolMallocDone = TRUE;
  }

  cv_mem->cv_itol   = itol;
  cv_mem->cv_reltol = reltol;
  if (itol == CV_SS)
    cv_mem->cv_Sabstol = *((realtype *)abstol);
  else
    N_VScale(ONE, (N_Vector)abstol, cv_mem->cv_Vabstol);

  cv_mem->cv_efun = CVEwtSet;
  cv_mem->cv_e_data = cvode_mem;


  return(CV_SUCCESS);
}

/* 
 * CVodeSetEwtFn
 *
 * Specifies the user-provide EwtSet function and data pointer for e
 */

int CVodeSetEwtFn(void *cvode_mem, CVEwtFn efun, void *e_data)
{
  CVodeMem cv_mem;

  if (cvode_mem==NULL) {
    CVProcessError(NULL, CV_MEM_NULL, "CVODES", "CVodeSetEwtFn", MSGCV_NO_MEM);
    return(CV_MEM_NULL);
  }

  cv_mem = (CVodeMem) cvode_mem;

  if ( cv_mem->cv_VabstolMallocDone ) {
    N_VDestroy(cv_mem->cv_Vabstol);
    lrw -= lrw1;
    liw -= liw1;
    cv_mem->cv_VabstolMallocDone = FALSE;
  }

  cv_mem->cv_itol = CV_WF;
  cv_mem->cv_efun = efun;
  cv_mem->cv_e_data = e_data;

  return(CV_SUCCESS);
}


/* 
 * =================================================================
 * Quadrature optional input functions
 * =================================================================
 */

/* 
 * Readability constants
 */

#define lrw1Q (cv_mem->cv_lrw1Q)
#define liw1Q (cv_mem->cv_liw1Q)

/*-----------------------------------------------------------------*/

int CVodeSetQuadFdata(void *cvode_mem, void *fQ_data)
{
  CVodeMem cv_mem;

  if (cvode_mem==NULL) {
    CVProcessError(NULL, CV_MEM_NULL, "CVODES", "CVodeSetQuadFdata", MSGCV_NO_MEM);    
    return(CV_MEM_NULL);
  }

  cv_mem = (CVodeMem) cvode_mem;

  cv_mem->cv_fQ_data = fQ_data;

  return(CV_SUCCESS);
}

/*-----------------------------------------------------------------*/

int CVodeSetQuadErrCon(void *cvode_mem, booleantype errconQ, 
                       int itolQ, realtype reltolQ, void *abstolQ)
{
  CVodeMem cv_mem;
  booleantype neg_abstol;

  if (cvode_mem==NULL) {
    CVProcessError(NULL, CV_MEM_NULL, "CVODES", "CVodeSetQuadErrCon", MSGCV_NO_MEM);    
    return(CV_MEM_NULL);
  }
  
  cv_mem = (CVodeMem) cvode_mem;

  cv_mem->cv_errconQ = errconQ;

  /* Ckeck if quadrature was initialized? */

  if (cv_mem->cv_quadMallocDone == FALSE) {
    CVProcessError(cv_mem, CV_NO_QUAD, "CVODES", "CVodeSetQuadErrCon", MSGCV_NO_QUAD); 
    return(CV_NO_QUAD);
  }

  /* Check inputs */

  if(errconQ == FALSE) {
    if (cv_mem->cv_VabstolQMallocDone) {
      N_VDestroy(cv_mem->cv_VabstolQ);
      lrw -= lrw1Q;
      liw -= liw1Q;
      cv_mem->cv_VabstolQMallocDone = FALSE;
    }
    return(CV_SUCCESS);
  }
  
  if ((itolQ != CV_SS) && (itolQ != CV_SV)) {
    CVProcessError(cv_mem, CV_ILL_INPUT, "CVODES", "CVodeSetQuadErrCon", MSGCV_BAD_ITOLQ);
    return(CV_ILL_INPUT);
  }

  if (abstolQ == NULL) {
    CVProcessError(cv_mem, CV_ILL_INPUT, "CVODES", "CVodeSetQuadErrCon", MSGCV_NULL_ABSTOLQ);
    return(CV_ILL_INPUT);
  }

  if (reltolQ < ZERO) {
    CVProcessError(cv_mem, CV_ILL_INPUT, "CVODES", "CVodeSetQuadErrCon", MSGCV_BAD_RELTOLQ);
    return(CV_ILL_INPUT);
  }

  if (itolQ == CV_SS)
    neg_abstol = (*((realtype *)abstolQ) < ZERO);
  else
    neg_abstol = (N_VMin((N_Vector)abstolQ) < ZERO);

  if (neg_abstol) {
    CVProcessError(cv_mem, CV_ILL_INPUT, "CVODES", "CVodeSetQuadErrCon", MSGCV_BAD_ABSTOLQ);
    return(CV_ILL_INPUT);
  }

  /* See if we need to free or allocate memory */

  if ( (itolQ != CV_SV) && (cv_mem->cv_VabstolQMallocDone) ) {
    N_VDestroy(cv_mem->cv_VabstolQ);
    lrw -= lrw1Q;
    liw -= liw1Q;
    cv_mem->cv_VabstolQMallocDone = FALSE;
  }

  if ( (itolQ == CV_SV) && !(cv_mem->cv_VabstolQMallocDone) ) {
    cv_mem->cv_VabstolQ = NULL;
    cv_mem->cv_VabstolQ = N_VClone(cv_mem->cv_tempvQ);
    lrw += lrw1Q;
    liw += liw1Q;
    cv_mem->cv_VabstolQMallocDone = TRUE;
  }

  /* Copy tolerances into memory */

  cv_mem->cv_itolQ    = itolQ;
  cv_mem->cv_reltolQ  = reltolQ;

  if (itolQ == CV_SS)
    cv_mem->cv_SabstolQ = *((realtype *)abstolQ);
  else
    N_VScale(ONE, (N_Vector)abstolQ, cv_mem->cv_VabstolQ);
  
  return(CV_SUCCESS);
}

/* 
 * =================================================================
 * FSA optional input functions
 * =================================================================
 */


int CVodeSetSensRhsFn(void *cvode_mem, CVSensRhsFn fS, void *fS_data)
{
  CVodeMem cv_mem;

  if (cvode_mem==NULL) {
    CVProcessError(NULL, CV_MEM_NULL, "CVODES", "CVodeSetSensRhsFn", MSGCV_NO_MEM);    
    return(CV_MEM_NULL);
  }

  cv_mem = (CVodeMem) cvode_mem;

  if (fS != NULL) {
    cv_mem->cv_ifS  = CV_ALLSENS;
    cv_mem->cv_fS      = fS;
    cv_mem->cv_fS_data = fS_data;
    cv_mem->cv_fSDQ    = FALSE;
  } else {
    cv_mem->cv_fSDQ    = TRUE;
  }

  return(CV_SUCCESS);
}

/*-----------------------------------------------------------------*/

int CVodeSetSensRhs1Fn(void *cvode_mem, CVSensRhs1Fn fS1, void *fS_data)
{
  CVodeMem cv_mem;

  if (cvode_mem==NULL) {
    CVProcessError(NULL, CV_MEM_NULL, "CVODES", "CVodeSetSensRhs1Fn", MSGCV_NO_MEM);    
    return(CV_MEM_NULL);
  }

  cv_mem = (CVodeMem) cvode_mem;
  
  if(fS1 != NULL) {
    cv_mem->cv_ifS  = CV_ONESENS;
    cv_mem->cv_fS1     = fS1;
    cv_mem->cv_fS_data = fS_data;
    cv_mem->cv_fSDQ    = FALSE;
  } else {
    cv_mem->cv_fSDQ    = TRUE;
  }

  return(CV_SUCCESS);
}

/*-----------------------------------------------------------------*/

int CVodeSetSensDQMethod(void *cvode_mem, int DQtype, realtype DQrhomax)
{
  CVodeMem cv_mem;

  if (cvode_mem==NULL) {
    CVProcessError(NULL, CV_MEM_NULL, "CVODES", "CVodeSetSensDQMethod", MSGCV_NO_MEM);    
    return(CV_MEM_NULL);
  }

  cv_mem = (CVodeMem) cvode_mem;

  if ( (DQtype != CV_CENTERED) && (DQtype != CV_FORWARD) ) {
    CVProcessError(cv_mem, CV_ILL_INPUT, "CVODES", "CVodeSetSensDQMethod", MSGCV_BAD_DQTYPE);    
    return(CV_ILL_INPUT);
  }

  if (DQrhomax < ZERO ) {
    CVProcessError(cv_mem, CV_ILL_INPUT, "CVODES", "CVodeSetSensDQMethod", MSGCV_BAD_DQRHO);    
    return(CV_ILL_INPUT);
  }

  cv_mem->cv_DQtype = DQtype;
  cv_mem->cv_DQrhomax = DQrhomax;

  return(CV_SUCCESS);
}

/*-----------------------------------------------------------------*/

int CVodeSetSensErrCon(void *cvode_mem, booleantype errconS)
{
  CVodeMem cv_mem;

  if (cvode_mem==NULL) {
    CVProcessError(NULL, CV_MEM_NULL, "CVODES", "CVodeSetSensErrCon", MSGCV_NO_MEM);    
    return(CV_MEM_NULL);
  }

  cv_mem = (CVodeMem) cvode_mem;

  cv_mem->cv_errconS = errconS;

  return(CV_SUCCESS);
}

/*-----------------------------------------------------------------*/

int CVodeSetSensMaxNonlinIters(void *cvode_mem, int maxcorS)
{
  CVodeMem cv_mem;

  if (cvode_mem==NULL) {
    CVProcessError(NULL, CV_MEM_NULL, "CVODES", "CVodeSetSensMaxNonlinIters", MSGCV_NO_MEM);    
    return (CV_MEM_NULL);
  }

  cv_mem = (CVodeMem) cvode_mem;

  cv_mem->cv_maxcorS = maxcorS;

  return(CV_SUCCESS);
}

/*-----------------------------------------------------------------*/

int CVodeSetSensParams(void *cvode_mem, realtype *p, realtype *pbar, int *plist)
{
  CVodeMem cv_mem;
  int is, Ns;

  if (cvode_mem==NULL) {
    CVProcessError(NULL, CV_MEM_NULL, "CVODES", "CVodeSetSensParams", MSGCV_NO_MEM);    
    return(CV_MEM_NULL);
  }

  cv_mem = (CVodeMem) cvode_mem;

  /* Was sensitivity initialized? */

  if (cv_mem->cv_sensMallocDone == FALSE) {
    CVProcessError(cv_mem, CV_NO_SENS, "CVODES", "CVodeSetSensParams", MSGCV_NO_SENSI);
    return(CV_NO_SENS);
  }

  Ns = cv_mem->cv_Ns;

  /* Parameters */

  cv_mem->cv_p = p;

  /* pbar */

  if (pbar != NULL)
    for (is=0; is<Ns; is++) {
      if (pbar[is] == ZERO) {
        CVProcessError(cv_mem, CV_ILL_INPUT, "CVODES", "CVodeSetSensParams", MSGCV_BAD_PBAR);
        return(CV_ILL_INPUT);
      }
      cv_mem->cv_pbar[is] = ABS(pbar[is]);
    }
  else
    for (is=0; is<Ns; is++)
      cv_mem->cv_pbar[is] = ONE;

  /* plist */

  if (plist != NULL)
    for (is=0; is<Ns; is++) {
      if ( plist[is] < 0 ) {
        CVProcessError(cv_mem, CV_ILL_INPUT, "CVODES", "CVodeSetSensParams", MSGCV_BAD_PLIST);
        return(CV_ILL_INPUT);
      }
      cv_mem->cv_plist[is] = plist[is];
    }
  else
    for (is=0; is<Ns; is++)
      cv_mem->cv_plist[is] = is;

  return(CV_SUCCESS);
}

/*-----------------------------------------------------------------*/

int CVodeSetSensTolerances(void *cvode_mem, int itolS,
                           realtype reltolS, void *abstolS)
{
  CVodeMem cv_mem;
  booleantype neg_abstol;
  realtype *atolSS;
  N_Vector *atolSV;
  int is, Ns;

  atolSS = NULL;
  atolSV = NULL;

  if (cvode_mem==NULL) {
    CVProcessError(NULL, CV_MEM_NULL, "CVODES", "CVodeSetSensTolerances", MSGCV_NO_MEM);    
    return(CV_MEM_NULL);
  }

  cv_mem = (CVodeMem) cvode_mem;

  /* Was sensitivity initialized? */

  if (cv_mem->cv_sensMallocDone == FALSE) {
    CVProcessError(cv_mem, CV_NO_SENS, "CVODES", "CVodeSetSensTolerances", MSGCV_NO_SENSI);
    return(CV_NO_SENS);
  } 

  /* Check inputs */

  Ns = cv_mem->cv_Ns;

  if ((itolS != CV_SS) && (itolS != CV_SV) && (itolS != CV_EE)) {
    CVProcessError(cv_mem, CV_ILL_INPUT, "CVODES", "CVodeSetSensTolerances", MSGCV_BAD_ITOLS);
    return(CV_ILL_INPUT);
  }

  if (itolS != CV_EE) {

    /* Test user-supplied tolerances */
    
    if (reltolS < ZERO) {
      CVProcessError(cv_mem, CV_ILL_INPUT, "CVODES", "CVodeSetSensTolerances", MSGCV_BAD_RELTOLS);
      return(CV_ILL_INPUT);
    }

    if (abstolS == NULL) {
      CVProcessError(cv_mem, CV_ILL_INPUT, "CVODES", "CVodeSetSensTolerances", MSGCV_NULL_ABSTOLS);
      return(CV_ILL_INPUT);
    }

    neg_abstol = FALSE;

    if (itolS == CV_SS) {
      atolSS = (realtype *) abstolS;
      for (is=0; is<Ns; is++)
        if (atolSS[is] < ZERO) {neg_abstol = TRUE; break;}
    } else {
      atolSV = (N_Vector *) abstolS;
      for (is=0; is<Ns; is++) 
        if (N_VMin(atolSV[is]) < ZERO) {neg_abstol = TRUE; break;}
    }

    if (neg_abstol) {
      CVProcessError(cv_mem, CV_ILL_INPUT, "CVODES", "CVodeSetSensTolerances", MSGCV_BAD_ABSTOLS);
      return(CV_ILL_INPUT);
    }
    
  }

  /* See if we should release some memory */

  if ( (itolS != CV_SV) && (cv_mem->cv_VabstolSMallocDone) ) {
    N_VDestroyVectorArray(cv_mem->cv_VabstolS, Ns);
    lrw -= Ns*lrw1;
    liw -= Ns*liw1;
    cv_mem->cv_VabstolSMallocDone = FALSE;
  }

  if ( (itolS != CV_SS) && (cv_mem->cv_SabstolSMallocDone) ) {
    free(cv_mem->cv_SabstolS); cv_mem->cv_SabstolS = NULL;
    lrw -= Ns;
    cv_mem->cv_SabstolSMallocDone = FALSE;
  }

  /* If tolerances will be estimated, return now */

  if (itolS == CV_EE) return(CV_SUCCESS);

  /* See if we need to allocate some memory */

  if ( (itolS == CV_SV) && !(cv_mem->cv_VabstolSMallocDone) ) {
    cv_mem->cv_VabstolS = NULL;
    cv_mem->cv_VabstolS = N_VCloneVectorArray(Ns, cv_mem->cv_tempv);
    lrw += Ns*lrw1;
    liw += Ns*liw1;
    cv_mem->cv_VabstolSMallocDone = TRUE;
  }

  if ( (itolS == CV_SS) && !(cv_mem->cv_SabstolSMallocDone) ) {
    cv_mem->cv_SabstolS = NULL;
    cv_mem->cv_SabstolS = (realtype *)malloc(Ns*sizeof(realtype));
    lrw += Ns;
    cv_mem->cv_SabstolSMallocDone = TRUE;
  }

  /* Copy tolerances into memory */

  cv_mem->cv_itolS   = itolS;
  cv_mem->cv_reltolS = reltolS;

  if (itolS == CV_SS)
    for (is=0; is<Ns; is++)
      cv_mem->cv_SabstolS[is] = atolSS[is];
  else
    for (is=0; is<Ns; is++)    
      N_VScale(ONE, atolSV[is], cv_mem->cv_VabstolS[is]);

  return(CV_SUCCESS);
}

/* 
 * =================================================================
 * CVODES optional output functions
 * =================================================================
 */

/* 
 * Readability constants
 */

#define nst            (cv_mem->cv_nst)
#define nfe            (cv_mem->cv_nfe)
#define ncfn           (cv_mem->cv_ncfn)
#define netf           (cv_mem->cv_netf)
#define nni            (cv_mem->cv_nni)
#define nsetups        (cv_mem->cv_nsetups)
#define qu             (cv_mem->cv_qu)
#define next_q         (cv_mem->cv_next_q)
#define ewt            (cv_mem->cv_ewt)  
#define hu             (cv_mem->cv_hu)
#define next_h         (cv_mem->cv_next_h)
#define h0u            (cv_mem->cv_h0u)
#define tolsf          (cv_mem->cv_tolsf)  
#define acor           (cv_mem->cv_acor)
#define lrw            (cv_mem->cv_lrw)
#define liw            (cv_mem->cv_liw)
#define nge            (cv_mem->cv_nge)
#define iroots         (cv_mem->cv_iroots)
#define nor            (cv_mem->cv_nor)
#define sldeton        (cv_mem->cv_sldeton)
#define tn             (cv_mem->cv_tn)
#define efun           (cv_mem->cv_efun)
#define e_data         (cv_mem->cv_e_data) 

/*
 * CVodeGetNumSteps
 *
 * Returns the current number of integration steps
 */

int CVodeGetNumSteps(void *cvode_mem, long int *nsteps)
{
  CVodeMem cv_mem;

  if (cvode_mem==NULL) {
    CVProcessError(NULL, CV_MEM_NULL, "CVODES", "CVodeGetNumSteps", MSGCV_NO_MEM);
    return(CV_MEM_NULL);
  }

  cv_mem = (CVodeMem) cvode_mem;

  *nsteps = nst;

  return(CV_SUCCESS);
}

/* 
 * CVodeGetNumRhsEvals
 *
 * Returns the current number of calls to f
 */

int CVodeGetNumRhsEvals(void *cvode_mem, long int *nfevals)
{
  CVodeMem cv_mem;

  if (cvode_mem==NULL) {
    CVProcessError(NULL, CV_MEM_NULL, "CVODES", "CVodeGetNumRhsEvals", MSGCV_NO_MEM);
    return(CV_MEM_NULL);
  }

  cv_mem = (CVodeMem) cvode_mem;

  *nfevals = nfe;

  return(CV_SUCCESS);
}

/* 
 * CVodeGetNumLinSolvSetups
 *
 * Returns the current number of calls to the linear solver setup routine
 */

int CVodeGetNumLinSolvSetups(void *cvode_mem, long int *nlinsetups)
{
  CVodeMem cv_mem;

  if (cvode_mem==NULL) {
    CVProcessError(NULL, CV_MEM_NULL, "CVODES", "CVodeGetNumLinSolvSetups", MSGCV_NO_MEM);
    return(CV_MEM_NULL);
  }

  cv_mem = (CVodeMem) cvode_mem;

  *nlinsetups = nsetups;

  return(CV_SUCCESS);
}

/*
 * CVodeGetNumErrTestFails
 *
 * Returns the current number of error test failures
 */

int CVodeGetNumErrTestFails(void *cvode_mem, long int *netfails)
{
  CVodeMem cv_mem;

  if (cvode_mem==NULL) {
    CVProcessError(NULL, CV_MEM_NULL, "CVODES", "CVodeGetNumErrTestFails", MSGCV_NO_MEM);
    return(CV_MEM_NULL);
  }

  cv_mem = (CVodeMem) cvode_mem;

  *netfails = netf;

  return(CV_SUCCESS);
}

/* 
 * CVodeGetLastOrder
 *
 * Returns the order on the last succesful step
 */

int CVodeGetLastOrder(void *cvode_mem, int *qlast)
{
  CVodeMem cv_mem;

  if (cvode_mem==NULL) {
    CVProcessError(NULL, CV_MEM_NULL, "CVODES", "CVodeGetLastOrder", MSGCV_NO_MEM);
    return(CV_MEM_NULL);
  }

  cv_mem = (CVodeMem) cvode_mem;

  *qlast = qu;

  return(CV_SUCCESS);
}

/* 
 * CVodeGetCurrentOrder
 *
 * Returns the order to be attempted on the next step
 */

int CVodeGetCurrentOrder(void *cvode_mem, int *qcur)
{
  CVodeMem cv_mem;

  if (cvode_mem==NULL) {
    CVProcessError(NULL, CV_MEM_NULL, "CVODES", "CVodeGetCurrentOrder", MSGCV_NO_MEM);
    return(CV_MEM_NULL);
  }

  cv_mem = (CVodeMem) cvode_mem;

  *qcur = next_q;

  return(CV_SUCCESS);
}

/* 
 * CVodeGetNumStabLimOrderReds
 *
 * Returns the number of order reductions triggered by the stability
 * limit detection algorithm
 */

int CVodeGetNumStabLimOrderReds(void *cvode_mem, long int *nslred)
{
  CVodeMem cv_mem;

  if (cvode_mem==NULL) {
    CVProcessError(NULL, CV_MEM_NULL, "CVODES", "CVodeGetNumStabLimOrderReds", MSGCV_NO_MEM);
    return(CV_MEM_NULL);
  }

  cv_mem = (CVodeMem) cvode_mem;

  if (sldeton==FALSE)
    *nslred = 0;
  else
    *nslred = nor;

  return(CV_SUCCESS);
}

/* 
 * CVodeGetActualInitStep
 *
 * Returns the step size used on the first step
 */

int CVodeGetActualInitStep(void *cvode_mem, realtype *hinused)
{
  CVodeMem cv_mem;

  if (cvode_mem==NULL) {
    CVProcessError(NULL, CV_MEM_NULL, "CVODES", "CVodeGetActualInitStep", MSGCV_NO_MEM);
    return(CV_MEM_NULL);
  }

  cv_mem = (CVodeMem) cvode_mem;

  *hinused = h0u;

  return(CV_SUCCESS);
}

/*
 * CVodeGetLastStep
 *
 * Returns the step size used on the last successful step
 */

int CVodeGetLastStep(void *cvode_mem, realtype *hlast)
{
  CVodeMem cv_mem;

  if (cvode_mem==NULL) {
    CVProcessError(NULL, CV_MEM_NULL, "CVODES", "CVodeGetLastStep", MSGCV_NO_MEM);
    return(CV_MEM_NULL);
  }

  cv_mem = (CVodeMem) cvode_mem;

  *hlast = hu;

  return(CV_SUCCESS);
}

/* 
 * CVodeGetCurrentStep
 *
 * Returns the step size to be attempted on the next step
 */

int CVodeGetCurrentStep(void *cvode_mem, realtype *hcur)
{
  CVodeMem cv_mem;

  if (cvode_mem==NULL) {
    CVProcessError(NULL, CV_MEM_NULL, "CVODES", "CVodeGetCurrentStep", MSGCV_NO_MEM);
    return(CV_MEM_NULL);
  }

  cv_mem = (CVodeMem) cvode_mem;
  
  *hcur = next_h;

  return(CV_SUCCESS);
}

/* 
 * CVodeGetCurrentTime
 *
 * Returns the current value of the independent variable
 */

int CVodeGetCurrentTime(void *cvode_mem, realtype *tcur)
{
  CVodeMem cv_mem;

  if (cvode_mem==NULL) {
    CVProcessError(NULL, CV_MEM_NULL, "CVODES", "CVodeGetCurrentTime", MSGCV_NO_MEM);
    return(CV_MEM_NULL);
  }

  cv_mem = (CVodeMem) cvode_mem;

  *tcur = tn;

  return(CV_SUCCESS);
}

/* 
 * CVodeGetTolScaleFactor
 *
 * Returns a suggested factor for scaling tolerances
 */

int CVodeGetTolScaleFactor(void *cvode_mem, realtype *tolsfact)
{
  CVodeMem cv_mem;

  if (cvode_mem==NULL) {
    CVProcessError(NULL, CV_MEM_NULL, "CVODES", "CVodeGetTolScaleFactor", MSGCV_NO_MEM);
    return(CV_MEM_NULL);
  }

  cv_mem = (CVodeMem) cvode_mem;

  *tolsfact = tolsf;

  return(CV_SUCCESS);
}

/* 
 * CVodeGetErrWeights
 *
 * This routine returns the current weight vector.
 */

int CVodeGetErrWeights(void *cvode_mem, N_Vector eweight)
{
  CVodeMem cv_mem;

  if (cvode_mem==NULL) {
    CVProcessError(NULL, CV_MEM_NULL, "CVODES", "CVodeGetErrWeights", MSGCV_NO_MEM);
    return(CV_MEM_NULL);
  }

  cv_mem = (CVodeMem) cvode_mem;

  N_VScale(ONE, ewt, eweight);

  return(CV_SUCCESS);
}

/*
 * CVodeGetEstLocalErrors
 *
 * Returns an estimate of the local error
 */

int CVodeGetEstLocalErrors(void *cvode_mem, N_Vector ele)
{
  CVodeMem cv_mem;

  if (cvode_mem==NULL) {
    CVProcessError(NULL, CV_MEM_NULL, "CVODES", "CVodeGetEstLocalErrors", MSGCV_NO_MEM);
    return(CV_MEM_NULL);
  }

  cv_mem = (CVodeMem) cvode_mem;

  N_VScale(ONE, acor, ele);

  return(CV_SUCCESS);
}

/* 
 * CVodeGetWorkSpace
 *
 * Returns integrator work space requirements
 */

int CVodeGetWorkSpace(void *cvode_mem, long int *lenrw, long int *leniw)
{
  CVodeMem cv_mem;

  if (cvode_mem==NULL) {
    CVProcessError(NULL, CV_MEM_NULL, "CVODES", "CVodeGetWorkSpace", MSGCV_NO_MEM);
    return(CV_MEM_NULL);
  }

  cv_mem = (CVodeMem) cvode_mem;

  *leniw = liw;
  *lenrw = lrw;

  return(CV_SUCCESS);
}

/* 
 * CVodeGetIntegratorStats
 *
 * Returns integrator statistics
 */

int CVodeGetIntegratorStats(void *cvode_mem, long int *nsteps, long int *nfevals, 
                            long int *nlinsetups, long int *netfails, int *qlast, 
                            int *qcur, realtype *hinused, realtype *hlast, 
                            realtype *hcur, realtype *tcur)
{
  CVodeMem cv_mem;

  if (cvode_mem==NULL) {
    CVProcessError(NULL, CV_MEM_NULL, "CVODES", "CVodeGetIntegratorStats", MSGCV_NO_MEM);
    return(CV_MEM_NULL);
  }

  cv_mem = (CVodeMem) cvode_mem;

  *nsteps = nst;
  *nfevals = nfe;
  *nlinsetups = nsetups;
  *netfails = netf;
  *qlast = qu;
  *qcur = next_q;
  *hinused = h0u;
  *hlast = hu;
  *hcur = next_h;
  *tcur = tn;

  return(CV_SUCCESS);
}

/* 
 * CVodeGetNumGEvals
 *
 * Returns the current number of calls to g (for rootfinding)
 */

int CVodeGetNumGEvals(void *cvode_mem, long int *ngevals)
{
  CVodeMem cv_mem;

  if (cvode_mem==NULL) {
    CVProcessError(NULL, CV_MEM_NULL, "CVODES", "CVodeGetNumGEvals", MSGCV_NO_MEM);
    return(CV_MEM_NULL);
  }

  cv_mem = (CVodeMem) cvode_mem;

  *ngevals = nge;

  return(CV_SUCCESS);
}

/* 
 * CVodeGetRootInfo
 *
 * Returns pointer to array rootsfound showing roots found
 */

int CVodeGetRootInfo(void *cvode_mem, int *rootsfound)
{
  CVodeMem cv_mem;
  int i, nrt;

  if (cvode_mem==NULL) {
    CVProcessError(NULL, CV_MEM_NULL, "CVODES", "CVodeGetRootInfo", MSGCV_NO_MEM);
    return(CV_MEM_NULL);
  }

  cv_mem = (CVodeMem) cvode_mem;

  nrt = cv_mem->cv_nrtfn;

  for (i=0; i<nrt; i++) rootsfound[i] = iroots[i];

  return(CV_SUCCESS);
}


/* 
 * CVodeGetNumNonlinSolvIters
 *
 * Returns the current number of iterations in the nonlinear solver
 */

int CVodeGetNumNonlinSolvIters(void *cvode_mem, long int *nniters)
{
  CVodeMem cv_mem;

  if (cvode_mem==NULL) {
    CVProcessError(NULL, CV_MEM_NULL, "CVODES", "CVodeGetNumNonlinSolvIters", MSGCV_NO_MEM);
    return(CV_MEM_NULL);
  }

  cv_mem = (CVodeMem) cvode_mem;

  *nniters = nni;

  return(CV_SUCCESS);
}

/* 
 * CVodeGetNumNonlinSolvConvFails
 *
 * Returns the current number of convergence failures in the
 * nonlinear solver
 */

int CVodeGetNumNonlinSolvConvFails(void *cvode_mem, long int *nncfails)
{
  CVodeMem cv_mem;

  if (cvode_mem==NULL) {
    CVProcessError(NULL, CV_MEM_NULL, "CVODES", "CVodeGetNumNonlinSolvConvFails", MSGCV_NO_MEM);
    return(CV_MEM_NULL);
  }

  cv_mem = (CVodeMem) cvode_mem;

  *nncfails = ncfn;

  return(CV_SUCCESS);
}

/* 
 * CVodeGetNonlinSolvStats
 *
 * Returns nonlinear solver statistics
 */

int CVodeGetNonlinSolvStats(void *cvode_mem, long int *nniters, 
                            long int *nncfails)
{
  CVodeMem cv_mem;

  if (cvode_mem==NULL) {
    CVProcessError(NULL, CV_MEM_NULL, "CVODES", "CVodeGetNonlinSolvStats", MSGCV_NO_MEM);
    return(CV_MEM_NULL);
  }

  cv_mem = (CVodeMem) cvode_mem;

  *nniters = nni;
  *nncfails = ncfn;

  return(CV_SUCCESS);
}


/* 
 * =================================================================
 * Quadrature optional output functions
 * =================================================================
 */

/* 
 * Readability constants
 */

#define quadr          (cv_mem->cv_quadr)
#define nfQe           (cv_mem->cv_nfQe)
#define netfQ          (cv_mem->cv_netfQ)
#define ewtQ           (cv_mem->cv_ewtQ)
#define errconQ        (cv_mem->cv_errconQ)

/*-----------------------------------------------------------------*/

int CVodeGetQuadNumRhsEvals(void *cvode_mem, long int *nfQevals)
{
  CVodeMem cv_mem;

  if (cvode_mem==NULL) {
    CVProcessError(NULL, CV_MEM_NULL, "CVODES", "CVodeGetQuadNumRhsEvals", MSGCV_NO_MEM);    
    return(CV_MEM_NULL);
  }

  cv_mem = (CVodeMem) cvode_mem;

  if (quadr==FALSE) {
    CVProcessError(cv_mem, CV_NO_QUAD, "CVODES", "CVodeGetQuadNumRhsEvals", MSGCV_NO_QUAD); 
    return(CV_NO_QUAD);
  }

  *nfQevals = nfQe;

  return(CV_SUCCESS);
}

/*-----------------------------------------------------------------*/

int CVodeGetQuadNumErrTestFails(void *cvode_mem, long int *nQetfails)
{
  CVodeMem cv_mem;

  if (cvode_mem==NULL) {
    CVProcessError(NULL, CV_MEM_NULL, "CVODES", "CVodeGetQuadNumErrTestFails", MSGCV_NO_MEM);    
    return(CV_MEM_NULL);
  }

  cv_mem = (CVodeMem) cvode_mem;

  if (quadr==FALSE) {
    CVProcessError(cv_mem, CV_NO_QUAD, "CVODES", "CVodeGetQuadNumErrTestFails", MSGCV_NO_QUAD); 
    return(CV_NO_QUAD);
  }

  *nQetfails = netfQ;

  return(CV_SUCCESS);
}

/*-----------------------------------------------------------------*/

int CVodeGetQuadErrWeights(void *cvode_mem, N_Vector eQweight)
{
  CVodeMem cv_mem;

  if (cvode_mem==NULL) {
    CVProcessError(NULL, CV_MEM_NULL, "CVODES", "CVodeGetQuadErrWeights", MSGCV_NO_MEM); 
    return(CV_MEM_NULL);
  }

  cv_mem = (CVodeMem) cvode_mem;

  if (quadr==FALSE) {
    CVProcessError(cv_mem, CV_NO_QUAD, "CVODES", "CVodeGetQuadErrWeights", MSGCV_NO_QUAD); 
    return(CV_NO_QUAD);
  }

  if(errconQ) N_VScale(ONE, ewtQ, eQweight);

  return(CV_SUCCESS);
}

/*-----------------------------------------------------------------*/

int CVodeGetQuadStats(void *cvode_mem, long int *nfQevals, long int *nQetfails)
{
  CVodeMem cv_mem;

  if (cvode_mem==NULL) {
    CVProcessError(NULL, CV_MEM_NULL, "CVODES", "CVodeGetQuadStats", MSGCV_NO_MEM);    
    return(CV_MEM_NULL);
  }

  cv_mem = (CVodeMem) cvode_mem;

  if (quadr==FALSE) {
    CVProcessError(cv_mem, CV_NO_QUAD, "CVODES", "CVodeGetQuadStats", MSGCV_NO_QUAD); 
    return(CV_NO_QUAD);
  }

  *nfQevals = nfQe;
  *nQetfails = netfQ;

  return(CV_SUCCESS);
}

/* 
 * =================================================================
 * FSA optional output functions
 * =================================================================
 */

/* 
 * Readability constants
 */

#define sensi          (cv_mem->cv_sensi)
#define ism            (cv_mem->cv_ism)
#define ewtS           (cv_mem->cv_ewtS)
#define nfSe           (cv_mem->cv_nfSe)
#define nfeS           (cv_mem->cv_nfeS)
#define nniS           (cv_mem->cv_nniS)
#define ncfnS          (cv_mem->cv_ncfnS)
#define netfS          (cv_mem->cv_netfS)
#define nsetupsS       (cv_mem->cv_nsetupsS)
#define nniS1          (cv_mem->cv_nniS1)
#define ncfnS1         (cv_mem->cv_ncfnS1)
#define ncfS1          (cv_mem->cv_ncfS1)

/*-----------------------------------------------------------------*/

int CVodeGetNumSensRhsEvals(void *cvode_mem, long int *nfSevals)
{
  CVodeMem cv_mem;

  if (cvode_mem==NULL) {
    CVProcessError(NULL, CV_MEM_NULL, "CVODES", "CVodeGetNumSensRhsEvals", MSGCV_NO_MEM);    
    return(CV_MEM_NULL);
  }

  cv_mem = (CVodeMem) cvode_mem;

  if (sensi==FALSE) {
    CVProcessError(cv_mem, CV_NO_SENS, "CVODES", "CVodeGetNumSensRhsEvals", MSGCV_NO_SENSI);
    return(CV_NO_SENS);
  }

  *nfSevals = nfSe;

  return(CV_SUCCESS);
}

/*-----------------------------------------------------------------*/

int CVodeGetNumRhsEvalsSens(void *cvode_mem, long int *nfevalsS)
{
  CVodeMem cv_mem;

  if (cvode_mem==NULL) {
    CVProcessError(NULL, CV_MEM_NULL, "CVODES", "CVodeGetNumRhsEvalsSens", MSGCV_NO_MEM);    
    return(CV_MEM_NULL);
  }

  cv_mem = (CVodeMem) cvode_mem;

  if (sensi==FALSE) {
    CVProcessError(cv_mem, CV_NO_SENS, "CVODES", "CVodeGetNumRhsEvalsSens", MSGCV_NO_SENSI);
    return(CV_NO_SENS);
  }

  *nfevalsS = nfeS;

  return(CV_SUCCESS);
}

/*-----------------------------------------------------------------*/

int CVodeGetNumSensErrTestFails(void *cvode_mem, long int *nSetfails)
{
  CVodeMem cv_mem;

  if (cvode_mem==NULL) {
    CVProcessError(NULL, CV_MEM_NULL, "CVODES", "CVodeGetNumSensErrTestFails", MSGCV_NO_MEM);    
    return(CV_MEM_NULL);
  }

  cv_mem = (CVodeMem) cvode_mem;

  if (sensi==FALSE) {
    CVProcessError(cv_mem, CV_NO_SENS, "CVODES", "CVodeGetNumSensErrTestFails", MSGCV_NO_SENSI);
    return(CV_NO_SENS);
  }

  *nSetfails = netfS;

  return(CV_SUCCESS);
}

/*-----------------------------------------------------------------*/

int CVodeGetNumSensLinSolvSetups(void *cvode_mem, long int *nlinsetupsS)
{
  CVodeMem cv_mem;

  if (cvode_mem==NULL) {
    CVProcessError(NULL, CV_MEM_NULL, "CVODES", "CVodeGetNumSensLinSolvSetups", MSGCV_NO_MEM);    
    return(CV_MEM_NULL);
  }

  cv_mem = (CVodeMem) cvode_mem;

  if (sensi==FALSE) {
    CVProcessError(cv_mem, CV_NO_SENS, "CVODES", "CVodeGetNumSensLinSolvSetups", MSGCV_NO_SENSI);
    return(CV_NO_SENS);
  }

  *nlinsetupsS = nsetupsS;

  return(CV_SUCCESS);
}

/*-----------------------------------------------------------------*/

int CVodeGetSensErrWeights(void *cvode_mem, N_Vector_S eSweight)
{
  CVodeMem cv_mem;
  int is, Ns;

  if (cvode_mem==NULL) {
    CVProcessError(NULL, CV_MEM_NULL, "CVODES", "CVodeGetSensErrWeights", MSGCV_NO_MEM);    
    return(CV_MEM_NULL);
  }

  cv_mem = (CVodeMem) cvode_mem;

  if (sensi==FALSE) {
    CVProcessError(cv_mem, CV_NO_SENS, "CVODES", "CVodeGetSensErrWeights", MSGCV_NO_SENSI);
    return(CV_NO_SENS);
  }

  Ns = cv_mem->cv_Ns;

  for (is=0; is<Ns; is++)
    N_VScale(ONE, ewtS[is], eSweight[is]);

  return(CV_SUCCESS);
}

/*-----------------------------------------------------------------*/

int CVodeGetSensStats(void *cvode_mem, long int *nfSevals, long int *nfevalsS, 
                      long int *nSetfails, long int *nlinsetupsS)
{
  CVodeMem cv_mem;

  if (cvode_mem==NULL) {
    CVProcessError(NULL, CV_MEM_NULL, "CVODES", "CVodeGetSensStats", MSGCV_NO_MEM);
    return(CV_MEM_NULL);
  }

  cv_mem = (CVodeMem) cvode_mem;

  if (sensi==FALSE) {
    CVProcessError(cv_mem, CV_NO_SENS, "CVODES", "CVodeGetSensStats", MSGCV_NO_SENSI);
    return(CV_NO_SENS);
  }

  *nfSevals = nfSe;
  *nfevalsS = nfeS;
  *nSetfails = netfS;
  *nlinsetupsS = nsetupsS;

  return(CV_SUCCESS);
}

/*-----------------------------------------------------------------*/

int CVodeGetNumSensNonlinSolvIters(void *cvode_mem, long int *nSniters)
{
  CVodeMem cv_mem;

  if (cvode_mem==NULL) {
    CVProcessError(NULL, CV_MEM_NULL, "CVODES", "CVodeGetNumSensNonlinSolvIters", MSGCV_NO_MEM);
    return(CV_MEM_NULL);
  }

  cv_mem = (CVodeMem) cvode_mem;

  if (sensi==FALSE) {
    CVProcessError(cv_mem, CV_NO_SENS, "CVODES", "CVodeGetNumSensNonlinSolvIters", MSGCV_NO_SENSI);
    return(CV_NO_SENS);
  }

  *nSniters = nniS;

  return(CV_SUCCESS);
}

/*-----------------------------------------------------------------*/

int CVodeGetNumSensNonlinSolvConvFails(void *cvode_mem, long int *nSncfails)
{
  CVodeMem cv_mem;

  if (cvode_mem==NULL) {
    CVProcessError(NULL, CV_MEM_NULL, "CVODES", "CVodeGetNumSensNonlinSolvConvFails", MSGCV_NO_MEM);
    return(CV_MEM_NULL);
  }

  cv_mem = (CVodeMem) cvode_mem;

  if (sensi==FALSE) {
    CVProcessError(cv_mem, CV_NO_SENS, "CVODES", "CVodeGetNumSensNonlinSolvConvFails", MSGCV_NO_SENSI);
    return(CV_NO_SENS);
  }

  *nSncfails = ncfnS;

  return(CV_SUCCESS);
}

/*-----------------------------------------------------------------*/

int CVodeGetNumStgrSensNonlinSolvIters(void *cvode_mem, long int *nSTGR1niters)
{
  CVodeMem cv_mem;
  int is, Ns;

  if (cvode_mem==NULL) {
    CVProcessError(NULL, CV_MEM_NULL, "CVODES", "CVodeGetNumStgrSensNonlinSolvIters", MSGCV_NO_MEM);
    return(CV_MEM_NULL);
  }

  cv_mem = (CVodeMem) cvode_mem;

  Ns = cv_mem->cv_Ns;

  if (sensi==FALSE) {
    CVProcessError(cv_mem, CV_NO_SENS, "CVODES", "CVodeGetNumStgrSensNonlinSolvIters", MSGCV_NO_SENSI);
    return(CV_NO_SENS);
  }

  if(ism==CV_STAGGERED1) 
    for(is=0; is<Ns; is++) nSTGR1niters[is] = nniS1[is];

  return(CV_SUCCESS);
}

/*-----------------------------------------------------------------*/

int CVodeGetNumStgrSensNonlinSolvConvFails(void *cvode_mem, long int *nSTGR1ncfails)
{
  CVodeMem cv_mem;
  int is, Ns;

  if (cvode_mem==NULL) {
    CVProcessError(NULL, CV_MEM_NULL, "CVODES", "CVodeGetNumStgrSensNonlinSolvConvFails", MSGCV_NO_MEM);
    return(CV_MEM_NULL);
  }

  cv_mem = (CVodeMem) cvode_mem;

  Ns = cv_mem->cv_Ns;

  if (sensi==FALSE) {
    CVProcessError(cv_mem, CV_NO_SENS, "CVODES", "CVodeGetNumStgrSensNonlinSolvConvFails", MSGCV_NO_SENSI);
    return(CV_NO_SENS);
  }

  if(ism==CV_STAGGERED1) 
    for(is=0; is<Ns; is++) nSTGR1ncfails[is] = ncfnS1[is];

  return(CV_SUCCESS);
}

/*-----------------------------------------------------------------*/

int CVodeGetSensNonlinSolvStats(void *cvode_mem, long int *nSniters, 
                                long int *nSncfails)
{
  CVodeMem cv_mem;

  if (cvode_mem==NULL) {
    CVProcessError(NULL, CV_MEM_NULL, "CVODES", "CVodeGetSensNonlinSolvstats", MSGCV_NO_MEM);
    return(CV_MEM_NULL);
  }

  cv_mem = (CVodeMem) cvode_mem;

  if (sensi==FALSE) {
    CVProcessError(cv_mem, CV_NO_SENS, "CVODES", "CVodeGetSensNonlinSolvStats", MSGCV_NO_SENSI);
    return(CV_NO_SENS);
  }

  *nSniters = nniS;
  *nSncfails = ncfnS;

  return(CV_SUCCESS);
}

/*-----------------------------------------------------------------*/

char *CVodeGetReturnFlagName(int flag)
{
  char *name;

  name = (char *)malloc(24*sizeof(char));

  switch(flag) {
  case CV_SUCCESS:
    sprintf(name,"CV_SUCCESS");
    break;
  case CV_TSTOP_RETURN:
    sprintf(name,"CV_TSTOP_RETURN");
    break;
  case CV_ROOT_RETURN:
    sprintf(name,"CV_ROOT_RETURN");
    break;
  case CV_TOO_MUCH_WORK:
    sprintf(name,"CV_TOO_MUCH_WORK");
    break;
  case CV_TOO_MUCH_ACC:
    sprintf(name,"CV_TOO_MUCH_ACC");
    break;
  case CV_ERR_FAILURE:
    sprintf(name,"CV_ERR_FAILURE");
    break;
  case CV_CONV_FAILURE:
    sprintf(name,"CV_CONV_FAILURE");
    break;
  case CV_LINIT_FAIL:
    sprintf(name,"CV_LINIT_FAIL");
    break;
  case CV_LSETUP_FAIL:
    sprintf(name,"CV_LSETUP_FAIL");
    break;
  case CV_LSOLVE_FAIL:
    sprintf(name,"CV_LSOLVE_FAIL");
    break;
  case CV_RHSFUNC_FAIL:
    sprintf(name,"CV_RHSFUNC_FAIL");
    break;
  case CV_FIRST_RHSFUNC_ERR:
    sprintf(name,"CV_FIRST_RHSFUNC_ERR");
    break;
  case CV_REPTD_RHSFUNC_ERR:
    sprintf(name,"CV_REPTD_RHSFUNC_ERR");
    break;
  case CV_UNREC_RHSFUNC_ERR:
    sprintf(name,"CV_UNREC_RHSFUNC_ERR");
    break;
  case CV_RTFUNC_FAIL:
    sprintf(name,"CV_RTFUNC_FAIL");
    break;
  case CV_MEM_FAIL:
    sprintf(name,"CV_MEM_FAIL");
    break;
  case CV_MEM_NULL:
    sprintf(name,"CV_MEM_NULL");
    break;
  case CV_ILL_INPUT:
    sprintf(name,"CV_ILL_INPUT");
    break;
  case CV_NO_MALLOC:
    sprintf(name,"CV_NO_MALLOC");
    break;
  case CV_BAD_K:
    sprintf(name,"CV_BAD_K");
    break;
  case CV_BAD_T:
    sprintf(name,"CV_BAD_T");
    break;
  case CV_BAD_DKY:
    sprintf(name,"CV_BAD_DKY");
    break;
  case CV_NO_QUAD:
    sprintf(name,"CV_NO_QUAD");
    break;
  case CV_QRHSFUNC_FAIL:
    sprintf(name,"CV_QRHSFUNC_FAIL");
    break;
  case CV_FIRST_QRHSFUNC_ERR:
    sprintf(name,"CV_FIRST_QRHSFUNC_ERR");
    break;
  case CV_REPTD_QRHSFUNC_ERR:
    sprintf(name,"CV_REPTD_QRHSFUNC_ERR");
    break;
  case CV_UNREC_QRHSFUNC_ERR:
    sprintf(name,"CV_UNREC_QRHSFUNC_ERR");
    break;
  case CV_BAD_IS:
    sprintf(name,"CV_BAD_IS");
    break;
  case CV_NO_SENS:
    sprintf(name,"CV_NO_SENS");
    break;
  case CV_SRHSFUNC_FAIL:
    sprintf(name,"CV_SRHSFUNC_FAIL");
    break;
  case CV_FIRST_SRHSFUNC_ERR:
    sprintf(name,"CV_FIRST_SRHSFUNC_ERR");
    break;
  case CV_REPTD_SRHSFUNC_ERR:
    sprintf(name,"CV_REPTD_SRHSFUNC_ERR");
    break;
  case CV_UNREC_SRHSFUNC_ERR:
    sprintf(name,"CV_UNREC_SRHSFUNC_ERR");
    break;
  case CV_TOO_CLOSE:
    sprintf(name,"CV_TOO_CLOSE");
    break;    
  default:
    sprintf(name,"NONE");
  }

  return(name);
}
```
