## Supplementary scripts for "A multi-scale *in silico* mouse model for insulin resistance and humanoid type 2 diabetes": cvodes_spgmr.c

```
/*
 * -----------------------------------------------------------------
 * $Revision: 749 $
 * $Date: 2011-06-22 14:49:10 +0200 (Wed, 22 Jun 2011) $
 * ----------------------------------------------------------------- 
 * Programmer(s): Radu Serban @ LLNL
 * -----------------------------------------------------------------
 * Copyright (c) 2005, The Regents of the University of California.
 * Produced at the Lawrence Livermore National Laboratory.
 * All rights reserved.
 * For details, see the LICENSE file.
 * -----------------------------------------------------------------
 * This is the implementation file for the CVSPGMR linear solver.
 * -----------------------------------------------------------------
 */

#include <stdio.h>
#include <stdlib.h>

#include <cvodes/cvodes_spgmr.h>
#include "cvodes_spils_impl.h"
#include "cvodes_impl.h"

#include <sundials/sundials_spgmr.h>
#include <sundials/sundials_math.h>

/* Constants */

#define ZERO RCONST(0.0)
#define ONE  RCONST(1.0)

/* CVSPGMR linit, lsetup, lsolve, and lfree routines */

static int CVSpgmrInit(CVodeMem cv_mem);

static int CVSpgmrSetup(CVodeMem cv_mem, int convfail, N_Vector ypred,
                        N_Vector fpred, booleantype *jcurPtr, N_Vector vtemp1,
                        N_Vector vtemp2, N_Vector vtemp3);

static int CVSpgmrSolve(CVodeMem cv_mem, N_Vector b, N_Vector weight,
                        N_Vector ynow, N_Vector fnow);

static void CVSpgmrFree(CVodeMem cv_mem);

/* CVSPGMR lfreeB function */

static void CVSpgmrFreeB(CVadjMem ca_mem);

/* 
 * ================================================================
 *
 *                   PART I - forward problems
 *
 * ================================================================
 */


/* Readability Replacements */

#define uround  (cv_mem->cv_uround)
#define tq      (cv_mem->cv_tq)
#define nst     (cv_mem->cv_nst)
#define tn      (cv_mem->cv_tn)
#define h       (cv_mem->cv_h)
#define gamma   (cv_mem->cv_gamma)
#define gammap  (cv_mem->cv_gammap)   
#define f       (cv_mem->cv_f)
#define f_data  (cv_mem->cv_f_data)
#define ewt     (cv_mem->cv_ewt)
#define mnewt   (cv_mem->cv_mnewt)
#define ropt    (cv_mem->cv_ropt)
#define linit   (cv_mem->cv_linit)
#define lsetup  (cv_mem->cv_lsetup)
#define lsolve  (cv_mem->cv_lsolve)
#define lfree   (cv_mem->cv_lfree)
#define lmem    (cv_mem->cv_lmem)
#define vec_tmpl     (cv_mem->cv_tempv)
#define setupNonNull (cv_mem->cv_setupNonNull)

#define sqrtN   (cvspils_mem->s_sqrtN)   
#define ytemp   (cvspils_mem->s_ytemp)
#define x       (cvspils_mem->s_x)
#define ycur    (cvspils_mem->s_ycur)
#define fcur    (cvspils_mem->s_fcur)
#define delta   (cvspils_mem->s_delta)
#define deltar  (cvspils_mem->s_deltar)
#define npe     (cvspils_mem->s_npe)
#define nli     (cvspils_mem->s_nli)
#define nps     (cvspils_mem->s_nps)
#define ncfl    (cvspils_mem->s_ncfl)
#define nstlpre (cvspils_mem->s_nstlpre)
#define njtimes (cvspils_mem->s_njtimes)
#define nfes   (cvspils_mem->s_nfes)
#define spils_mem (cvspils_mem->s_spils_mem)
#define last_flag (cvspils_mem->s_last_flag)

/*
 * -----------------------------------------------------------------
 * CVSpgmr
 * -----------------------------------------------------------------
 * This routine initializes the memory record and sets various function
 * fields specific to the Spgmr linear solver module. CVSpgmr first
 * calls the existing lfree routine if this is not NULL.  It then sets
 * the cv_linit, cv_lsetup, cv_lsolve, cv_lfree fields in (*cvode_mem)
 * to be CVSpgmrInit, CVSpgmrSetup, CVSpgmrSolve, and CVSpgmrFree,
 * respectively.  It allocates memory for a structure of type
 * CVSpilsMemRec and sets the cv_lmem field in (*cvode_mem) to the
 * address of this structure.  It sets setupNonNull in (*cvode_mem),
 * and sets the following fields in the CVSpilsMemRec structure:
 *   s_pretype = pretype                                       
 *   s_gstype  = gstype                                       
 *   s_maxl    = MIN(N,CVSPILS_MAXL  if maxl <= 0             
 *             = maxl                 if maxl > 0              
 *   s_delt    = CVSPILS_DELT if delt == 0.0                     
 *             = delt         if delt != 0.0                     
 *   s_P_data  = P_data                                        
 *   s_pset    = pset                                       
 *   s_psolve  = psolve                                        
 *   s_jtimes  = input parameter jtimes  if jtimes != NULL
 *             = CVSpilsDQJtimes         otherwise
 *   s_j_data  = input parameter jac_data
 * Finally, CVSpgmr allocates memory for ytemp and x, and calls
 * SpgmrMalloc to allocate memory for the Spgmr solver.
 * -----------------------------------------------------------------
 */

int CVSpgmr(void *cvode_mem, int pretype, int maxl)
{
  CVodeMem cv_mem;
  CVSpilsMem cvspils_mem;
  SpgmrMem spgmr_mem;
  int mxl;

  /* Return immediately if cvode_mem is NULL */
  if (cvode_mem == NULL) {
    CVProcessError(NULL, CVSPILS_MEM_NULL, "CVSPGMR", "CVSpgmr", MSGS_CVMEM_NULL);
    return(CVSPILS_MEM_NULL);
  }
  cv_mem = (CVodeMem) cvode_mem;

  /* Check if N_VDotProd is present */
  if(vec_tmpl->ops->nvdotprod == NULL) {
    CVProcessError(cv_mem, CVSPILS_ILL_INPUT, "CVSPGMR", "CVSpgmr", MSGS_BAD_NVECTOR);
    return(CVSPILS_ILL_INPUT);
  }

  if (lfree != NULL) lfree(cv_mem);

  /* Set four main function fields in cv_mem */
  linit  = CVSpgmrInit;
  lsetup = CVSpgmrSetup;
  lsolve = CVSpgmrSolve;
  lfree  = CVSpgmrFree;

  /* Get memory for CVSpilsMemRec */
  cvspils_mem = NULL;
  cvspils_mem = (CVSpilsMem) malloc(sizeof(CVSpilsMemRec));
  if (cvspils_mem == NULL) {
    CVProcessError(cv_mem, CVSPILS_MEM_FAIL, "CVSPGMR", "CVSpgmr", MSGS_MEM_FAIL);
    return(CVSPILS_MEM_FAIL);
  }

  /* Set ILS type */
  cvspils_mem->s_type = SPILS_SPGMR;

  /* Set Spgmr parameters that have been passed in call sequence */
  cvspils_mem->s_pretype    = pretype;
  mxl = cvspils_mem->s_maxl = (maxl <= 0) ? CVSPILS_MAXL : maxl;

  /* Set default values for the rest of the Spgmr parameters */
  cvspils_mem->s_gstype     = MODIFIED_GS;
  cvspils_mem->s_delt       = CVSPILS_DELT;
  cvspils_mem->s_P_data     = NULL;
  cvspils_mem->s_pset       = NULL;
  cvspils_mem->s_psolve     = NULL;
  cvspils_mem->s_jtimes     = CVSpilsDQJtimes;
  cvspils_mem->s_j_data     = cvode_mem;
  cvspils_mem->s_last_flag  = CVSPILS_SUCCESS;


  setupNonNull = FALSE;

  /* Check for legal pretype */ 
  if ((pretype != PREC_NONE) && (pretype != PREC_LEFT) &&
      (pretype != PREC_RIGHT) && (pretype != PREC_BOTH)) {
    CVProcessError(cv_mem, CVSPILS_ILL_INPUT, "CVSPGMR", "CVSpgmr", MSGS_BAD_PRETYPE);
    return(CVSPILS_ILL_INPUT);
  }

  /* Allocate memory for ytemp and x */
  ytemp = NULL;
  ytemp = N_VClone(vec_tmpl);
  if (ytemp == NULL) {
    CVProcessError(cv_mem, CVSPILS_MEM_FAIL, "CVSPGMR", "CVSpgmr", MSGS_MEM_FAIL);
    free(cvspils_mem); cvspils_mem = NULL;
    return(CVSPILS_MEM_FAIL);
  }
  x = NULL;
  x = N_VClone(vec_tmpl);
  if (x == NULL) {
    CVProcessError(cv_mem, CVSPILS_MEM_FAIL, "CVSPGMR", "CVSpgmr", MSGS_MEM_FAIL);
    N_VDestroy(ytemp);
    free(cvspils_mem); cvspils_mem = NULL;
    return(CVSPILS_MEM_FAIL);
  }

  /* Compute sqrtN from a dot product */
  N_VConst(ONE, ytemp);
  sqrtN = RSqrt( N_VDotProd(ytemp, ytemp) );

  /* Call SpgmrMalloc to allocate workspace for Spgmr */
  spgmr_mem = NULL;
  spgmr_mem = SpgmrMalloc(mxl, vec_tmpl);
  if (spgmr_mem == NULL) {
    CVProcessError(cv_mem, CVSPILS_MEM_FAIL, "CVSPGMR", "CVSpgmr", MSGS_MEM_FAIL);
    N_VDestroy(ytemp);
    N_VDestroy(x);
    free(cvspils_mem); cvspils_mem = NULL;
    return(CVSPILS_MEM_FAIL);
  }
  
  /* Attach SPGMR memory to spils memory structure */
  spils_mem = (void *) spgmr_mem;

  /* Attach linear solver memory to integrator memory */
  lmem = cvspils_mem;

  return(CVSPILS_SUCCESS);
}


/* Additional readability Replacements */

#define pretype (cvspils_mem->s_pretype)
#define gstype  (cvspils_mem->s_gstype)
#define delt    (cvspils_mem->s_delt)
#define maxl    (cvspils_mem->s_maxl)
#define psolve  (cvspils_mem->s_psolve)
#define pset    (cvspils_mem->s_pset)
#define P_data  (cvspils_mem->s_P_data)
#define jtimes  (cvspils_mem->s_jtimes)
#define j_data  (cvspils_mem->s_j_data)

/*
 * -----------------------------------------------------------------
 * CVSpgmrInit
 * -----------------------------------------------------------------
 * This routine does remaining initializations specific to the Spgmr 
 * linear solver.
 * -----------------------------------------------------------------
 */

static int CVSpgmrInit(CVodeMem cv_mem)
{
  CVSpilsMem cvspils_mem;
  cvspils_mem = (CVSpilsMem) lmem;

  /* Initialize counters */
  npe = nli = nps = ncfl = nstlpre = 0;
  njtimes = nfes = 0;

  /* Check for legal combination pretype - psolve */
  if ((pretype != PREC_NONE) && (psolve == NULL)) {
    CVProcessError(cv_mem, -1, "CVSPGMR", "CVSpgmrInit", MSGS_PSOLVE_REQ);
    last_flag = CVSPILS_ILL_INPUT;
    return(-1);
  }

  /* Set setupNonNull = TRUE iff there is preconditioning (pretype != PREC_NONE)
     and there is a preconditioning setup phase (pset != NULL)             */
  setupNonNull = (pretype != PREC_NONE) && (pset != NULL);

  /* If jtimes is NULL at this time, set it to DQ */
  if (jtimes == NULL) {
    jtimes = CVSpilsDQJtimes;
    j_data = cv_mem;
  }

  last_flag = CVSPILS_SUCCESS;
  return(0);
}

/*
 * -----------------------------------------------------------------
 * CVSpgmrSetup
 * -----------------------------------------------------------------
 * This routine does the setup operations for the Spgmr linear solver.
 * It makes a decision as to whether or not to signal for re-evaluation
 * of Jacobian data in the pset routine, based on various state
 * variables, then it calls pset.  If we signal for re-evaluation,
 * then we reset jcur = *jcurPtr to TRUE, regardless of the pset output.
 * In any case, if jcur == TRUE, we increment npe and save nst in nstlpre.
 * -----------------------------------------------------------------
 */

static int CVSpgmrSetup(CVodeMem cv_mem, int convfail, N_Vector ypred,
                        N_Vector fpred, booleantype *jcurPtr, N_Vector vtemp1,
                        N_Vector vtemp2, N_Vector vtemp3)
{
  booleantype jbad, jok;
  realtype dgamma;
  int  retval;
  CVSpilsMem cvspils_mem;

  cvspils_mem = (CVSpilsMem) lmem;

  /* Use nst, gamma/gammap, and convfail to set J eval. flag jok */
  dgamma = ABS((gamma/gammap) - ONE);
  jbad = (nst == 0) || (nst > nstlpre + CVSPILS_MSBPRE) ||
    ((convfail == CV_FAIL_BAD_J) && (dgamma < CVSPILS_DGMAX)) ||
    (convfail == CV_FAIL_OTHER);
  *jcurPtr = jbad;
  jok = !jbad;

  /* Call pset routine and possibly reset jcur */
  retval = pset(tn, ypred, fpred, jok, jcurPtr, gamma, P_data, 
                vtemp1, vtemp2, vtemp3);
  if (retval < 0) {
    CVProcessError(cv_mem, SPGMR_PSET_FAIL_UNREC, "CVSPGMR", "CVSpgmrSetup", MSGS_PSET_FAILED);
    last_flag = SPGMR_PSET_FAIL_UNREC;
  }
  if (retval > 0) {
    last_flag = SPGMR_PSET_FAIL_REC;
  }

  if (jbad) *jcurPtr = TRUE;

  /* If jcur = TRUE, increment npe and save nst value */
  if (*jcurPtr) {
    npe++;
    nstlpre = nst;
  }

  last_flag = SPGMR_SUCCESS;

  /* Return the same value that pset returned */
  return(retval);
}

/*
 * -----------------------------------------------------------------
 * CVSpgmrSolve
 * -----------------------------------------------------------------
 * This routine handles the call to the generic solver SpgmrSolve
 * for the solution of the linear system Ax = b with the SPGMR method,
 * without restarts.  The solution x is returned in the vector b.
 *
 * If the WRMS norm of b is small, we return x = b (if this is the first
 * Newton iteration) or x = 0 (if a later Newton iteration).
 *
 * Otherwise, we set the tolerance parameter and initial guess (x = 0),
 * call SpgmrSolve, and copy the solution x into b.  The x-scaling and
 * b-scaling arrays are both equal to weight, and no restarts are allowed.
 *
 * The counters nli, nps, and ncfl are incremented, and the return value
 * is set according to the success of SpgmrSolve.  The success flag is
 * returned if SpgmrSolve converged, or if this is the first Newton
 * iteration and the residual norm was reduced below its initial value.
 * -----------------------------------------------------------------
 */

static int CVSpgmrSolve(CVodeMem cv_mem, N_Vector b, N_Vector weight,
                        N_Vector ynow, N_Vector fnow)
{
  realtype bnorm, res_norm;
  CVSpilsMem cvspils_mem;
  SpgmrMem spgmr_mem;
  int nli_inc, nps_inc, retval;
  
  cvspils_mem = (CVSpilsMem) lmem;

  spgmr_mem = (SpgmrMem) spils_mem;

  /* Test norm(b); if small, return x = 0 or x = b */
  deltar = delt*tq[4]; 

  bnorm = N_VWrmsNorm(b, weight);
  if (bnorm <= deltar) {
    if (mnewt > 0) N_VConst(ZERO, b); 
    return(0);
  }

  /* Set vectors ycur and fcur for use by the Atimes and Psolve routines */
  ycur = ynow;
  fcur = fnow;

  /* Set inputs delta and initial guess x = 0 to SpgmrSolve */  
  delta = deltar * sqrtN;
  N_VConst(ZERO, x);
  
  /* Call SpgmrSolve and copy x to b */
  retval = SpgmrSolve(spgmr_mem, cv_mem, x, b, pretype, gstype, delta, 0,
                   cv_mem, weight, weight, CVSpilsAtimes, CVSpilsPSolve,
                   &res_norm, &nli_inc, &nps_inc);

  N_VScale(ONE, x, b);
  
  /* Increment counters nli, nps, and ncfl */
  nli += nli_inc;
  nps += nps_inc;
  if (retval != SPGMR_SUCCESS) ncfl++;

  /* Interpret return value from SpgmrSolve */

  last_flag = retval;

  switch(retval) {

  case SPGMR_SUCCESS:
    return(0);
    break;
  case SPGMR_RES_REDUCED:
    if (mnewt == 0) return(0);
    else            return(1);
    break;
  case SPGMR_CONV_FAIL:
    return(1);
    break;
  case SPGMR_QRFACT_FAIL:
    return(1);
    break;
  case SPGMR_PSOLVE_FAIL_REC:
    return(1);
    break;
  case SPGMR_ATIMES_FAIL_REC:
    return(1);
    break;
  case SPGMR_MEM_NULL:
    return(-1);
    break;
  case SPGMR_ATIMES_FAIL_UNREC:
    CVProcessError(cv_mem, SPGMR_ATIMES_FAIL_UNREC, "CVSPGMR", "CVSpgmrSolve", MSGS_JTIMES_FAILED);    
    return(-1);
    break;
  case SPGMR_PSOLVE_FAIL_UNREC:
    CVProcessError(cv_mem, SPGMR_PSOLVE_FAIL_UNREC, "CVSPGMR", "CVSpgmrSolve", MSGS_PSOLVE_FAILED);
    return(-1);
    break;
  case SPGMR_GS_FAIL:
    return(-1);
    break;
  case SPGMR_QRSOL_FAIL:
    return(-1);
    break;
  }

  return(0);  

}

/*
 * -----------------------------------------------------------------
 * CVSpgmrFree
 * -----------------------------------------------------------------
 * This routine frees memory specific to the Spgmr linear solver.
 * -----------------------------------------------------------------
 */

static void CVSpgmrFree(CVodeMem cv_mem)
{
  CVSpilsMem cvspils_mem;
  SpgmrMem spgmr_mem;

  cvspils_mem = (CVSpilsMem) lmem;
  
  spgmr_mem = (SpgmrMem) spils_mem;

  N_VDestroy(ytemp);
  N_VDestroy(x);
  SpgmrFree(spgmr_mem);
  free(cvspils_mem); cvspils_mem = NULL;
}


/* 
 * ================================================================
 *
 *                   PART II - backward problems
 *
 * ================================================================
 */


/* Additional readability replacements */

#define lmemB       (ca_mem->ca_lmemB)
#define lfreeB      (ca_mem->ca_lfreeB)

#define pset_B      (cvspilsB_mem->s_psetB)
#define psolve_B    (cvspilsB_mem->s_psolveB)
#define jtimes_B    (cvspilsB_mem->s_jtimesB)
#define P_data_B    (cvspilsB_mem->s_P_dataB)
#define jac_data_B  (cvspilsB_mem->s_jac_dataB)

/*
 * CVSpgmrB
 *
 * Wrapper for the backward phase
 *
 */

int CVSpgmrB(void *cvadj_mem, int pretypeB, int maxlB)
{
  CVadjMem ca_mem;
  CVSpilsMemB cvspilsB_mem;
  CVodeMem cvB_mem;
  int flag;

  if (cvadj_mem == NULL) {
    CVProcessError(NULL, CVSPILS_ADJMEM_NULL, "CVSPGMR", "CVSpgmrB", MSGS_CAMEM_NULL);
    return(CVSPILS_ADJMEM_NULL);
  }
  ca_mem = (CVadjMem) cvadj_mem;

  cvB_mem = ca_mem->cvb_mem;

  /* Get memory for CVSpilsMemRecB */
  cvspilsB_mem = NULL;
  cvspilsB_mem = (CVSpilsMemB) malloc(sizeof(CVSpilsMemRecB));
  if (cvspilsB_mem == NULL) {
    CVProcessError(cvB_mem, CVSPILS_MEM_FAIL, "CVSPGMR", "CVSpgmrB", MSGS_MEM_FAIL);
    return(CVSPILS_MEM_FAIL);
  }

  pset_B = NULL;
  psolve_B = NULL;
  jtimes_B = NULL;
  P_data_B = NULL;
  jac_data_B = NULL;

  /* attach lmemB and lfreeB */
  lmemB = cvspilsB_mem;
  lfreeB = CVSpgmrFreeB;
  
  flag = CVSpgmr(cvB_mem, pretypeB, maxlB);

  if (flag != CVSPILS_SUCCESS) {
    free(cvspilsB_mem);
    cvspilsB_mem = NULL;
  }

  return(flag);
}


/*
 * CVSpgmrFreeB 
 */


static void CVSpgmrFreeB(CVadjMem ca_mem)
{
  CVSpilsMemB cvspilsB_mem;

  cvspilsB_mem = (CVSpilsMemB) lmemB;

  free(cvspilsB_mem);
}
```
