## Supplementary scripts for "A multi-scale *in silico* mouse model for insulin resistance and humanoid type 2 diabetes": cvodes_spils.c

```
/*
 * -----------------------------------------------------------------
 * $Revision: 749 $
 * $Date: 2011-06-22 14:49:10 +0200 (Wed, 22 Jun 2011) $
 * ----------------------------------------------------------------- 
 * Programmer(s):Radu Serban @ LLNL
 * -----------------------------------------------------------------
 * Copyright (c) 2005, The Regents of the University of California.
 * Produced at the Lawrence Livermore National Laboratory.
 * All rights reserved.
 * For details, see the LICENSE file.
 * -----------------------------------------------------------------
 * This is the implementation file for the CVSPILS linear solvers.
 *
 * Part II contains wrappers for using the CVODES iterative linear 
 * solvers on adjoint (backward) problems.
 * -----------------------------------------------------------------
 */

#include <stdio.h>
#include <stdlib.h>

#include "cvodes_impl.h"
#include "cvodes_spils_impl.h"

/* Private constants */

#define ZERO   RCONST(0.0)
#define PT25   RCONST(0.25)
#define ONE    RCONST(1.0)

/* Algorithmic constants */

#define MAX_ITERS  3  /* max. number of attempts to recover in DQ J*v */

/* 
 * ================================================================
 *
 *                   PART I - forward problems
 *
 * ================================================================
 */

/* Readability Replacements */

#define lrw1    (cv_mem->cv_lrw1)
#define liw1    (cv_mem->cv_liw1)
#define tq      (cv_mem->cv_tq)
#define tn      (cv_mem->cv_tn)
#define h       (cv_mem->cv_h)
#define gamma   (cv_mem->cv_gamma)
#define nfe     (cv_mem->cv_nfe)
#define f       (cv_mem->cv_f)
#define f_data  (cv_mem->cv_f_data)
#define ewt     (cv_mem->cv_ewt)
#define lmem    (cv_mem->cv_lmem)

#define ils_type (cvspils_mem->s_type)
#define sqrtN   (cvspils_mem->s_sqrtN)   
#define ytemp   (cvspils_mem->s_ytemp)
#define x       (cvspils_mem->s_x)
#define ycur    (cvspils_mem->s_ycur)
#define fcur    (cvspils_mem->s_fcur)
#define delta   (cvspils_mem->s_delta)
#define npe     (cvspils_mem->s_npe)
#define nli     (cvspils_mem->s_nli)
#define nps     (cvspils_mem->s_nps)
#define ncfl    (cvspils_mem->s_ncfl)
#define njtimes (cvspils_mem->s_njtimes)
#define nfes    (cvspils_mem->s_nfes)

#define last_flag (cvspils_mem->s_last_flag)


/*
 * -----------------------------------------------------------------
 * OPTIONAL INPUT and OUTPUT FUNCTIONS
 * -----------------------------------------------------------------
 */


/*
 * -----------------------------------------------------------------
 * CVSpilsSetPrecType
 * -----------------------------------------------------------------
 */

int CVSpilsSetPrecType(void *cvode_mem, int pretype)
{
  CVodeMem cv_mem;
  CVSpilsMem cvspils_mem;

  /* Return immediately if cvode_mem is NULL */
  if (cvode_mem == NULL) {
    CVProcessError(NULL, CVSPILS_MEM_NULL, "CVSPILS", "CVSpilsSetPrecType", MSGS_CVMEM_NULL);
    return(CVSPILS_MEM_NULL);
  }
  cv_mem = (CVodeMem) cvode_mem;

  if (lmem == NULL) {
    CVProcessError(cv_mem, CVSPILS_LMEM_NULL, "CVSPILS", "CVSpilsSetPrecType", MSGS_LMEM_NULL);
    return(CVSPILS_LMEM_NULL);
  }
  cvspils_mem = (CVSpilsMem) lmem;

  /* Check for legal pretype */ 
  if ((pretype != PREC_NONE) && (pretype != PREC_LEFT) &&
      (pretype != PREC_RIGHT) && (pretype != PREC_BOTH)) {
    CVProcessError(cv_mem, CVSPILS_ILL_INPUT, "CVSPILS", "CVSpilsSetPrecType", MSGS_BAD_PRETYPE);
    return(CVSPILS_ILL_INPUT);
  }

  cvspils_mem->s_pretype = pretype;

  return(CVSPILS_SUCCESS);
}

/*
 * -----------------------------------------------------------------
 * CVSpilsSetGSType
 * -----------------------------------------------------------------
 */

int CVSpilsSetGSType(void *cvode_mem, int gstype)
{
  CVodeMem cv_mem;
  CVSpilsMem cvspils_mem;

  /* Return immediately if cvode_mem is NULL */
  if (cvode_mem == NULL) {
    CVProcessError(NULL, CVSPILS_MEM_NULL, "CVSPILS", "CVSpilsSetGSType", MSGS_CVMEM_NULL);
    return(CVSPILS_MEM_NULL);
  }
  cv_mem = (CVodeMem) cvode_mem;

  if (lmem == NULL) {
    CVProcessError(cv_mem, CVSPILS_LMEM_NULL, "CVSPILS", "CVSpilsSetGSType", MSGS_LMEM_NULL);
    return(CVSPILS_LMEM_NULL);
  }
  cvspils_mem = (CVSpilsMem) lmem;

  if (ils_type != SPILS_SPGMR) {
    CVProcessError(cv_mem, CVSPILS_ILL_INPUT, "CVSPILS", "CVSpilsSetGSType", MSGS_BAD_LSTYPE);
    return(CVSPILS_ILL_INPUT);
  }

  /* Check for legal gstype */
  if ((gstype != MODIFIED_GS) && (gstype != CLASSICAL_GS)) {
    CVProcessError(cv_mem, CVSPILS_ILL_INPUT, "CVSPILS", "CVSpilsSetGSType", MSGS_BAD_GSTYPE);
    return(CVSPILS_ILL_INPUT);
  }

  cvspils_mem->s_gstype = gstype;

  return(CVSPILS_SUCCESS);
}

/*
 * -----------------------------------------------------------------
 * Function : CVSpilsSetMaxl
 * -----------------------------------------------------------------
 */

int CVSpilsSetMaxl(void *cvode_mem, int maxl)
{
  CVodeMem cv_mem;
  CVSpilsMem cvspils_mem;
  int mxl;

  /* Return immediately if cvode_mem is NULL */
  if (cvode_mem == NULL) {
    CVProcessError(NULL, CVSPILS_MEM_NULL, "CVSPILS", "CVSpilsSetMaxl", MSGS_CVMEM_NULL);
    return(CVSPILS_MEM_NULL);
  }
  cv_mem = (CVodeMem) cvode_mem;

  if (lmem == NULL) {
    CVProcessError(NULL, CVSPILS_LMEM_NULL, "CVSPILS", "CVSpilsSetMaxl", MSGS_LMEM_NULL);
    return(CVSPILS_LMEM_NULL);
  }
  cvspils_mem = (CVSpilsMem) lmem;

  if (ils_type == SPILS_SPGMR) {
    CVProcessError(cv_mem, CVSPILS_ILL_INPUT, "CVSPILS", "CVSpilsSetMaxl", MSGS_BAD_LSTYPE);
    return(CVSPILS_ILL_INPUT);
  }

  mxl = (maxl <= 0) ? CVSPILS_MAXL : maxl;
  cvspils_mem->s_maxl = mxl;

  /*  spbcg_mem->l_max  = mxl; */

  return(CVSPILS_SUCCESS);
}


/*
 * -----------------------------------------------------------------
 * CVSpilsSetDelt
 * -----------------------------------------------------------------
 */

int CVSpilsSetDelt(void *cvode_mem, realtype delt)
{
  CVodeMem cv_mem;
  CVSpilsMem cvspils_mem;

  /* Return immediately if cvode_mem is NULL */
  if (cvode_mem == NULL) {
    CVProcessError(NULL, CVSPILS_MEM_NULL, "CVSPILS", "CVSpilsSetDelt", MSGS_CVMEM_NULL);
    return(CVSPILS_MEM_NULL);
  }
  cv_mem = (CVodeMem) cvode_mem;

  if (lmem == NULL) {
    CVProcessError(cv_mem, CVSPILS_LMEM_NULL, "CVSPILS", "CVSpilsSetDelt", MSGS_LMEM_NULL);
    return(CVSPILS_LMEM_NULL);
  }
  cvspils_mem = (CVSpilsMem) lmem;

  /* Check for legal delt */
  if(delt < ZERO) {
    CVProcessError(cv_mem, CVSPILS_ILL_INPUT, "CVSPILS", "CVSpilsSetDelt", MSGS_BAD_DELT);
    return(CVSPILS_ILL_INPUT);
  }

  cvspils_mem->s_delt = (delt == ZERO) ? CVSPILS_DELT : delt;

  return(CVSPILS_SUCCESS);
}

/*
 * -----------------------------------------------------------------
 * CVSpilsSetPrecSetupFn
 * -----------------------------------------------------------------
 */

int CVSpilsSetPreconditioner(void *cvode_mem, CVSpilsPrecSetupFn pset, 
                             CVSpilsPrecSolveFn psolve, void *P_data)
{
  CVodeMem cv_mem;
  CVSpilsMem cvspils_mem;

  /* Return immediately if cvode_mem is NULL */
  if (cvode_mem == NULL) {
    CVProcessError(NULL, CVSPILS_MEM_NULL, "CVSPILS", "CVSpilsSetPreconditioner", MSGS_CVMEM_NULL);
    return(CVSPILS_MEM_NULL);
  }
  cv_mem = (CVodeMem) cvode_mem;

  if (lmem == NULL) {
    CVProcessError(cv_mem, CVSPILS_LMEM_NULL, "CVSPILS", "CVSpilsSetPreconditioner", MSGS_LMEM_NULL);
    return(CVSPILS_LMEM_NULL);
  }
  cvspils_mem = (CVSpilsMem) lmem;

  cvspils_mem->s_pset = pset;
  cvspils_mem->s_psolve = psolve;
  if (psolve != NULL) cvspils_mem->s_P_data = P_data;

  return(CVSPILS_SUCCESS);
}

/*
 * -----------------------------------------------------------------
 * CVSpilsSetJacTimesVecFn
 * -----------------------------------------------------------------
 */

int CVSpilsSetJacTimesVecFn(void *cvode_mem, CVSpilsJacTimesVecFn jtimes, void *jac_data)
{
  CVodeMem cv_mem;
  CVSpilsMem cvspils_mem;

  /* Return immediately if cvode_mem is NULL */
  if (cvode_mem == NULL) {
    CVProcessError(NULL, CVSPILS_MEM_NULL, "CVSPILS", "CVSpilsSetJacTimesVecFn", MSGS_CVMEM_NULL);
    return(CVSPILS_MEM_NULL);
  }
  cv_mem = (CVodeMem) cvode_mem;

  if (lmem == NULL) {
    CVProcessError(cv_mem, CVSPILS_LMEM_NULL, "CVSPILS", "CVSpilsSetJacTimesVecFn", MSGS_LMEM_NULL);
    return(CVSPILS_LMEM_NULL);
  }
  cvspils_mem = (CVSpilsMem) lmem;

  cvspils_mem->s_jtimes = jtimes;
  if (jtimes != NULL) cvspils_mem->s_j_data = jac_data;

  return(CVSPILS_SUCCESS);
}

/*
 * -----------------------------------------------------------------
 * CVSpilsGetWorkSpace
 * -----------------------------------------------------------------
 */

int CVSpilsGetWorkSpace(void *cvode_mem, long int *lenrwLS, long int *leniwLS)
{
  CVodeMem cv_mem;
  CVSpilsMem cvspils_mem;
  int maxl;

  /* Return immediately if cvode_mem is NULL */
  if (cvode_mem == NULL) {
    CVProcessError(NULL, CVSPILS_MEM_NULL, "CVSPILS", "CVSpilsGetWorkSpace", MSGS_CVMEM_NULL);
    return(CVSPILS_MEM_NULL);
  }
  cv_mem = (CVodeMem) cvode_mem;

  if (lmem == NULL) {
    CVProcessError(cv_mem, CVSPILS_LMEM_NULL, "CVSPILS", "CVSpilsGetWorkSpace", MSGS_LMEM_NULL);
    return(CVSPILS_LMEM_NULL);
  }
  cvspils_mem = (CVSpilsMem) lmem;

  
  switch(ils_type) {
  case SPILS_SPGMR:
    maxl = cvspils_mem->s_maxl;
    *lenrwLS = lrw1*(maxl + 5) + maxl*(maxl + 4) + 1;
    *leniwLS = liw1*(maxl + 5);
    break;
  case SPILS_SPBCG:
    *lenrwLS = lrw1 * 9;
    *leniwLS = liw1 * 9;
    break;
  case SPILS_SPTFQMR:
    *lenrwLS = lrw1*11;
    *leniwLS = liw1*11;
    break;
  }


  return(CVSPILS_SUCCESS);
}

/*
 * -----------------------------------------------------------------
 * CVSpilsGetNumPrecEvals
 * -----------------------------------------------------------------
 */

int CVSpilsGetNumPrecEvals(void *cvode_mem, long int *npevals)
{
  CVodeMem cv_mem;
  CVSpilsMem cvspils_mem;

  /* Return immediately if cvode_mem is NULL */
  if (cvode_mem == NULL) {
    CVProcessError(NULL, CVSPILS_MEM_NULL, "CVSPILS", "CVSpilsGetNumPrecEvals", MSGS_CVMEM_NULL);
    return(CVSPILS_MEM_NULL);
  }
  cv_mem = (CVodeMem) cvode_mem;

  if (lmem == NULL) {
    CVProcessError(cv_mem, CVSPILS_LMEM_NULL, "CVSPILS", "CVSpilsGetNumPrecEvals", MSGS_LMEM_NULL);
    return(CVSPILS_LMEM_NULL);
  }
  cvspils_mem = (CVSpilsMem) lmem;

  *npevals = npe;

  return(CVSPILS_SUCCESS);
}

/*
 * -----------------------------------------------------------------
 * CVSpilsGetNumPrecSolves
 * -----------------------------------------------------------------
 */

int CVSpilsGetNumPrecSolves(void *cvode_mem, long int *npsolves)
{
  CVodeMem cv_mem;
  CVSpilsMem cvspils_mem;

  /* Return immediately if cvode_mem is NULL */
  if (cvode_mem == NULL) {
    CVProcessError(NULL, CVSPILS_MEM_NULL, "CVSPILS", "CVSpilsGetNumPrecSolves", MSGS_CVMEM_NULL);
    return(CVSPILS_MEM_NULL);
  }
  cv_mem = (CVodeMem) cvode_mem;

  if (lmem == NULL) {
    CVProcessError(cv_mem, CVSPILS_LMEM_NULL, "CVSPILS", "CVSpilsGetNumPrecSolves", MSGS_LMEM_NULL);
    return(CVSPILS_LMEM_NULL);
  }
  cvspils_mem = (CVSpilsMem) lmem;

  *npsolves = nps;

  return(CVSPILS_SUCCESS);
}

/*
 * -----------------------------------------------------------------
 * CVSpilsGetNumLinIters
 * -----------------------------------------------------------------
 */

int CVSpilsGetNumLinIters(void *cvode_mem, long int *nliters)
{
  CVodeMem cv_mem;
  CVSpilsMem cvspils_mem;

  /* Return immediately if cvode_mem is NULL */
  if (cvode_mem == NULL) {
    CVProcessError(NULL, CVSPILS_MEM_NULL, "CVSPILS", "CVSpilsGetNumLinIters", MSGS_CVMEM_NULL);
    return(CVSPILS_MEM_NULL);
  }
  cv_mem = (CVodeMem) cvode_mem;

  if (lmem == NULL) {
    CVProcessError(cv_mem, CVSPILS_LMEM_NULL, "CVSPILS", "CVSpilsGetNumLinIters", MSGS_LMEM_NULL);
    return(CVSPILS_LMEM_NULL);
  }
  cvspils_mem = (CVSpilsMem) lmem;

  *nliters = nli;

  return(CVSPILS_SUCCESS);
}

/*
 * -----------------------------------------------------------------
 * CVSpilsGetNumConvFails
 * -----------------------------------------------------------------
 */

int CVSpilsGetNumConvFails(void *cvode_mem, long int *nlcfails)
{
  CVodeMem cv_mem;
  CVSpilsMem cvspils_mem;

  /* Return immediately if cvode_mem is NULL */
  if (cvode_mem == NULL) {
    CVProcessError(NULL, CVSPILS_MEM_NULL, "CVSPILS", "CVSpilsGetNumConvFails", MSGS_CVMEM_NULL);
    return(CVSPILS_MEM_NULL);
  }
  cv_mem = (CVodeMem) cvode_mem;

  if (lmem == NULL) {
    CVProcessError(cv_mem, CVSPILS_LMEM_NULL, "CVSPILS", "CVSpilsGetNumConvFails", MSGS_LMEM_NULL);
    return(CVSPILS_LMEM_NULL);
  }
  cvspils_mem = (CVSpilsMem) lmem;

  *nlcfails = ncfl;

  return(CVSPILS_SUCCESS);
}

/*
 * -----------------------------------------------------------------
 * CVSpilsGetNumJtimesEvals
 * -----------------------------------------------------------------
 */

int CVSpilsGetNumJtimesEvals(void *cvode_mem, long int *njvevals)
{
  CVodeMem cv_mem;
  CVSpilsMem cvspils_mem;

  /* Return immediately if cvode_mem is NULL */
  if (cvode_mem == NULL) {
    CVProcessError(NULL, CVSPILS_MEM_NULL, "CVSPILS", "CVSpilsGetNumJtimesEvals", MSGS_CVMEM_NULL);
    return(CVSPILS_MEM_NULL);
  }
  cv_mem = (CVodeMem) cvode_mem;

  if (lmem == NULL) {
    CVProcessError(cv_mem, CVSPILS_LMEM_NULL, "CVSPILS", "CVSpilsGetNumJtimesEvals", MSGS_LMEM_NULL);
    return(CVSPILS_LMEM_NULL);
  }
  cvspils_mem = (CVSpilsMem) lmem;

  *njvevals = njtimes;

  return(CVSPILS_SUCCESS);
}

/*
 * -----------------------------------------------------------------
 * CVSpilsGetNumRhsEvals
 * -----------------------------------------------------------------
 */

int CVSpilsGetNumRhsEvals(void *cvode_mem, long int *nfevalsLS)
{
  CVodeMem cv_mem;
  CVSpilsMem cvspils_mem;

  /* Return immediately if cvode_mem is NULL */
  if (cvode_mem == NULL) {
    CVProcessError(NULL, CVSPILS_MEM_NULL, "CVSPILS", "CVSpilsGetNumRhsEvals", MSGS_CVMEM_NULL);
    return(CVSPILS_MEM_NULL);
  }
  cv_mem = (CVodeMem) cvode_mem;

  if (lmem == NULL) {
    CVProcessError(cv_mem, CVSPILS_LMEM_NULL, "CVSPILS", "CVSpilsGetNumRhsEvals", MSGS_LMEM_NULL);
    return(CVSPILS_LMEM_NULL);
  }
  cvspils_mem = (CVSpilsMem) lmem;

  *nfevalsLS = nfes;

  return(CVSPILS_SUCCESS);
}

/*
 * -----------------------------------------------------------------
 * CVSpilsGetLastFlag
 * -----------------------------------------------------------------
 */

int CVSpilsGetLastFlag(void *cvode_mem, int *flag)
{
  CVodeMem cv_mem;
  CVSpilsMem cvspils_mem;

  /* Return immediately if cvode_mem is NULL */
  if (cvode_mem == NULL) {
    CVProcessError(NULL, CVSPILS_MEM_NULL, "CVSPILS", "CVSpilsGetLastFlag", MSGS_CVMEM_NULL);
    return(CVSPILS_MEM_NULL);
  }
  cv_mem = (CVodeMem) cvode_mem;

  if (lmem == NULL) {
    CVProcessError(cv_mem, CVSPILS_LMEM_NULL, "CVSPILS", "CVSpilsGetLastFlag", MSGS_LMEM_NULL);
    return(CVSPILS_LMEM_NULL);
  }
  cvspils_mem = (CVSpilsMem) lmem;

  *flag = last_flag;

  return(CVSPILS_SUCCESS);
}

/*
 * -----------------------------------------------------------------
 * CVSpilsGetReturnFlagName
 * -----------------------------------------------------------------
 */

char *CVSpilsGetReturnFlagName(int flag)
{
  char *name;

  name = (char *)malloc(30*sizeof(char));

  switch(flag) {
  case CVSPILS_SUCCESS:
    sprintf(name,"CVSPILS_SUCCESS");
    break;    
  case CVSPILS_MEM_NULL:
    sprintf(name,"CVSPILS_MEM_NULL");
    break;
  case CVSPILS_LMEM_NULL:
    sprintf(name,"CVSPILS_LMEM_NULL");
    break;
  case CVSPILS_ILL_INPUT:
    sprintf(name,"CVSPILS_ILL_INPUT");
    break;
  case CVSPILS_MEM_FAIL:
    sprintf(name,"CVSPILS_MEM_FAIL");
    break;
  case CVSPILS_ADJMEM_NULL:
    sprintf(name,"CVSPILS_ADJMEM_NULL");
    break;
  case CVSPILS_LMEMB_NULL:
    sprintf(name,"CVSPILS_LMEMB_NULL");
    break;
  default:
    sprintf(name,"NONE");
  }

  return(name);
}

/*
 * -----------------------------------------------------------------
 * CVSPILS private functions
 * -----------------------------------------------------------------
 */


/* Additional readability Replacements */

#define pretype (cvspils_mem->s_pretype)
#define delt    (cvspils_mem->s_delt)
#define maxl    (cvspils_mem->s_maxl)
#define psolve  (cvspils_mem->s_psolve)
#define P_data  (cvspils_mem->s_P_data)
#define jtimes  (cvspils_mem->s_jtimes)
#define j_data  (cvspils_mem->s_j_data)

/*
 * -----------------------------------------------------------------
 * CVSpilsAtimes
 * -----------------------------------------------------------------
 * This routine generates the matrix-vector product z = Mv, where
 * M = I - gamma*J. The product J*v is obtained by calling the jtimes 
 * routine. It is then scaled by -gamma and added to v to obtain M*v.
 * The return value is the same as the value returned by jtimes --
 * 0 if successful, nonzero otherwise.
 * -----------------------------------------------------------------
 */

int CVSpilsAtimes(void *cvode_mem, N_Vector v, N_Vector z)
{
  CVodeMem   cv_mem;
  CVSpilsMem cvspils_mem;
  int retval;

  cv_mem = (CVodeMem) cvode_mem;
  cvspils_mem = (CVSpilsMem) lmem;

  retval = jtimes(v, z, tn, ycur, fcur, j_data, ytemp);
  njtimes++;
  if (retval != 0) return(retval);

  N_VLinearSum(ONE, v, -gamma, z, z);

  return(0);
}

/*
 * -----------------------------------------------------------------
 * CVSpilsPSolve
 * -----------------------------------------------------------------
 * This routine interfaces between the generic SpgmrSolve routine and
 * the user's psolve routine.  It passes to psolve all required state 
 * information from cvode_mem.  Its return value is the same as that
 * returned by psolve. Note that the generic SPGMR solver guarantees
 * that CVSpilsPSolve will not be called in the case in which
 * preconditioning is not done. This is the only case in which the
 * user's psolve routine is allowed to be NULL.
 * -----------------------------------------------------------------
 */

int CVSpilsPSolve(void *cvode_mem, N_Vector r, N_Vector z, int lr)
{
  CVodeMem   cv_mem;
  CVSpilsMem cvspils_mem;
  int retval;

  cv_mem = (CVodeMem) cvode_mem;
  cvspils_mem = (CVSpilsMem)lmem;

  /* This call is counted in nps within the CVSp***Solve routine */
  retval = psolve(tn, ycur, fcur, r, z, gamma, delta, lr, P_data, ytemp);

  return(retval);     
}

/*
 * -----------------------------------------------------------------
 * CVSpilsDQJtimes
 * -----------------------------------------------------------------
 * This routine generates a difference quotient approximation to
 * the Jacobian times vector f_y(t,y) * v. The approximation is 
 * Jv = vnrm[f(y + v/vnrm) - f(y)], where vnrm = (WRMS norm of v) is
 * input, i.e. the WRMS norm of v/vnrm is 1.
 * -----------------------------------------------------------------
 */

int CVSpilsDQJtimes(N_Vector v, N_Vector Jv, realtype t, 
                    N_Vector y, N_Vector fy,
                    void *jac_data, N_Vector work)
{
  CVodeMem cv_mem;
  CVSpilsMem cvspils_mem;
  realtype sig, siginv;
  int iter, retval;

  /* jac_data is cvode_mem */
  cv_mem = (CVodeMem) jac_data;
  cvspils_mem = (CVSpilsMem) lmem;

  /* Initialize perturbation to 1/||v|| */
  sig = ONE/N_VWrmsNorm(v, ewt);

  for (iter=0; iter<MAX_ITERS; iter++) {

    /* Set work = y + sig*v */
    N_VLinearSum(sig, v, ONE, y, work);

    /* Set Jv = f(tn, y+sig*v) */
    retval = f(t, work, Jv, f_data); 
    nfes++;
    if (retval == 0) break;
    if (retval < 0)  return(-1);

    sig *= PT25;
  }

  if (retval > 0) return(+1);

  /* Replace Jv by (Jv - fy)/sig */
  siginv = ONE/sig;
  N_VLinearSum(siginv, Jv, -siginv, fy, Jv);

  return(0);
}

/* 
 * ================================================================
 *
 *                   PART II - backward problems
 *
 * ================================================================
 */

/* Readability replacements */

#define ytmp        (ca_mem->ca_ytmp)
#define getY        (ca_mem->ca_getY)
#define lmemB       (ca_mem->ca_lmemB)

#define pset_B      (cvspilsB_mem->s_psetB)
#define psolve_B    (cvspilsB_mem->s_psolveB)
#define jtimes_B    (cvspilsB_mem->s_jtimesB)
#define P_data_B    (cvspilsB_mem->s_P_dataB)
#define jac_data_B  (cvspilsB_mem->s_jac_dataB)

/*
 * -----------------------------------------------------------------
 * OPTIONAL INPUT and OUTPUT FUNCTIONS
 * -----------------------------------------------------------------
 */

int CVSpilsSetPrecTypeB(void *cvadj_mem, int pretypeB)
{
  CVadjMem ca_mem;
  CVodeMem cvB_mem;
  int flag;

  if (cvadj_mem == NULL) {
    CVProcessError(NULL, CVSPILS_ADJMEM_NULL, "CVSPILS", "CVSpilsSetPrecTypeB", MSGS_CAMEM_NULL);
    return(CVSPILS_ADJMEM_NULL);
  }
  ca_mem = (CVadjMem) cvadj_mem;

  cvB_mem = ca_mem->cvb_mem;

  flag = CVSpilsSetPrecType(cvB_mem, pretypeB);

  return(flag);
}

int CVSpilsSetGSTypeB(void *cvadj_mem, int gstypeB)
{
  CVadjMem ca_mem;
  CVodeMem cvB_mem;
  int flag;

  if (cvadj_mem == NULL) {
    CVProcessError(NULL, CVSPILS_ADJMEM_NULL, "CVSPILS", "CVSpilsSetGSTypeB", MSGS_CAMEM_NULL);
    return(CVSPILS_ADJMEM_NULL);
  }
  ca_mem = (CVadjMem) cvadj_mem;

  cvB_mem = ca_mem->cvb_mem;

  flag = CVSpilsSetGSType(cvB_mem,gstypeB);

  return(flag);
}

int CVSpilsSetDeltB(void *cvadj_mem, realtype deltB)
{
  CVadjMem ca_mem;
  CVodeMem cvB_mem;
  int flag;

  if (cvadj_mem == NULL) {
    CVProcessError(NULL, CVSPILS_ADJMEM_NULL, "CVSPILS", "CVSpilsSetDeltB", MSGS_CAMEM_NULL);
    return(CVSPILS_ADJMEM_NULL);
  }
  ca_mem = (CVadjMem) cvadj_mem;

  cvB_mem = ca_mem->cvb_mem;

  flag = CVSpilsSetDelt(cvB_mem,deltB);

  return(flag);
}

int CVSpilsSetMaxlB(void *cvadj_mem, int maxlB)
{
  CVadjMem ca_mem;
  CVodeMem cvB_mem;
  int flag;

  if (cvadj_mem == NULL) {
    CVProcessError(NULL, CVSPILS_ADJMEM_NULL, "CVSPILS", "CVSpilsSetMaxlB", MSGS_CAMEM_NULL);
    return(CVSPILS_ADJMEM_NULL);
  }
  ca_mem = (CVadjMem) cvadj_mem;

  cvB_mem = ca_mem->cvb_mem;

  flag = CVSpilsSetMaxl(cvB_mem,maxlB);

  return(flag);
}

int CVSpilsSetPreconditionerB(void *cvadj_mem, CVSpilsPrecSetupFnB psetB,
                              CVSpilsPrecSolveFnB psolveB, void *P_dataB)
{
  CVadjMem ca_mem;
  CVSpilsMemB cvspilsB_mem; 
  CVodeMem cvB_mem;
  int flag;

  if (cvadj_mem == NULL) {
    CVProcessError(NULL, CVSPILS_ADJMEM_NULL, "CVSPILS", "CVSpilsSetPreconditionerB", MSGS_CAMEM_NULL);
    return(CVSPILS_ADJMEM_NULL);
  }
  ca_mem = (CVadjMem) cvadj_mem;

  cvB_mem = ca_mem->cvb_mem;

  if (lmemB == NULL) {
    CVProcessError(cvB_mem, CVSPILS_LMEMB_NULL, "CVSPILS", "CVSpilsSetPreconditonerB", MSGS_LMEMB_NULL);
    return(CVSPILS_LMEMB_NULL);
  }
  cvspilsB_mem = (CVSpilsMemB) lmemB;

  pset_B   = psetB;
  psolve_B = psolveB;
  P_data_B = P_dataB;

  flag = CVSpilsSetPreconditioner(cvB_mem, CVAspilsPrecSetup, CVAspilsPrecSolve, cvadj_mem);

  return(flag);
}

int CVSpilsSetJacTimesVecFnB(void *cvadj_mem, CVSpilsJacTimesVecFnB jtimesB,
                             void *jac_dataB)
{
  CVadjMem ca_mem;
  CVSpilsMemB cvspilsB_mem; 
  CVodeMem cvB_mem;
  int flag;

  if (cvadj_mem == NULL) {
    CVProcessError(NULL, CVSPILS_ADJMEM_NULL, "CVSPILS", "CVSpilsSetJacTimesVecFnB", MSGS_CAMEM_NULL);
    return(CVSPILS_ADJMEM_NULL);
  }
  ca_mem = (CVadjMem) cvadj_mem;

  cvB_mem = ca_mem->cvb_mem;

  if (lmemB == NULL) {
    CVProcessError(cvB_mem, CVSPILS_LMEMB_NULL, "CVSPILS", "CVSpilsSetJacTimesVecFnB", MSGS_LMEMB_NULL);
    return(CVSPILS_LMEMB_NULL);
  }
  cvspilsB_mem = (CVSpilsMemB) lmemB;

  jtimes_B   = jtimesB;
  jac_data_B = jac_dataB;

  flag = CVSpilsSetJacTimesVecFn(cvB_mem, CVAspilsJacTimesVec, cvadj_mem);

  return(flag);
}


/*
 * -----------------------------------------------------------------
 * CVSPILS private functions
 * -----------------------------------------------------------------
 */

/*
 * CVAspilsPrecSetup
 *
 * This routine interfaces to the CVSpilsPrecSetupFnB routine 
 * provided by the user.
 * NOTE: p_data actually contains cvadj_mem
 */

int CVAspilsPrecSetup(realtype t, N_Vector yB, 
                      N_Vector fyB, booleantype jokB, 
                      booleantype *jcurPtrB, realtype gammaB,
                      void *cvadj_mem,
                      N_Vector tmp1B, N_Vector tmp2B, N_Vector tmp3B)
{
  CVadjMem ca_mem;
  CVodeMem cvB_mem;
  CVSpilsMemB cvspilsB_mem;
  int retval, flag;

  ca_mem = (CVadjMem) cvadj_mem;
  cvB_mem = ca_mem->cvb_mem;
  cvspilsB_mem = (CVSpilsMemB) lmemB;

  /* Forward solution from interpolation */
  flag = getY(ca_mem, t, ytmp);
  if (flag != CV_SUCCESS) {
    CVProcessError(cvB_mem, -1, "CVSPILS", "CVAspilsPrecSetup", MSGS_BAD_T);
    return(-1);
  } 

  /* Call user's adjoint precondB routine */
  retval = pset_B(t, ytmp, yB, fyB, jokB, jcurPtrB, gammaB,
                  P_data_B, tmp1B, tmp2B, tmp3B);

  return(retval);
}


/*
 * CVAspilsPrecSolve
 *
 * This routine interfaces to the CVSpilsPrecSolveFnB routine 
 * provided by the user.
 * NOTE: p_data actually contains cvadj_mem
 */

int CVAspilsPrecSolve(realtype t, N_Vector yB, N_Vector fyB,
                      N_Vector rB, N_Vector zB,
                      realtype gammaB, realtype deltaB,
                      int lrB, void *cvadj_mem, N_Vector tmpB)
{
  CVadjMem ca_mem;
  CVodeMem cvB_mem;
  CVSpilsMemB cvspilsB_mem;
  int retval, flag;

  ca_mem = (CVadjMem) cvadj_mem;
  cvB_mem = ca_mem->cvb_mem;
  cvspilsB_mem = (CVSpilsMemB) lmemB;

  /* Forward solution from interpolation */
  flag = getY(ca_mem, t, ytmp);
  if (flag != CV_SUCCESS) {
    CVProcessError(cvB_mem, -1, "CVSPILS", "CVAspilsPrecSolve", MSGS_BAD_T);
    return(-1);
  }

  /* Call user's adjoint psolveB routine */
  retval = psolve_B(t, ytmp, yB, fyB, rB, zB, gammaB, deltaB, 
                    lrB, P_data_B, tmpB);

  return(retval);
}


/*
 * CVAspilsJacTimesVec
 *
 * This routine interfaces to the CVSpilsJacTimesVecFnB routine 
 * provided by the user.
 * NOTE: jac_data actually contains cvadj_mem
 */

int CVAspilsJacTimesVec(N_Vector vB, N_Vector JvB, realtype t, 
                        N_Vector yB, N_Vector fyB, 
                        void *cvadj_mem, N_Vector tmpB)
{
  CVadjMem ca_mem;
  CVodeMem cvB_mem;
  CVSpilsMemB cvspilsB_mem;
  int retval, flag;

  ca_mem = (CVadjMem) cvadj_mem;
  cvB_mem = ca_mem->cvb_mem;
  cvspilsB_mem = (CVSpilsMemB) lmemB;

  /* Forward solution from interpolation */
  flag = getY(ca_mem, t, ytmp);
  if (flag != CV_SUCCESS) {
    CVProcessError(cvB_mem, -1, "CVSPILS", "CVAspilsJacTimesVec", MSGS_BAD_T);
    return(-1);
  } 

  /* Call user's adjoint jtimesB routine */
  retval = jtimes_B(vB, JvB, t, ytmp, yB, fyB, jac_data_B, tmpB);

  return(retval);
}
```
