## Supplementary scripts for "A multi-scale *in silico* mouse model for insulin resistance and humanoid type 2 diabetes": cvodes_spils_impl.h

```
/*
 * -----------------------------------------------------------------
 * $Revision: 749 $
 * $Date: 2011-06-22 14:49:10 +0200 (Wed, 22 Jun 2011) $
 * ----------------------------------------------------------------- 
 * Programmer(s): Radu Serban @ LLNL
 * -----------------------------------------------------------------
 * Copyright (c) 2005, The Regents of the University of California.
 * Produced at the Lawrence Livermore National Laboratory.
 * All rights reserved.
 * For details, see the LICENSE file.
 * -----------------------------------------------------------------
 * Common implementation header file for the scaled, preconditioned
 * iterative linear solvers
 * -----------------------------------------------------------------
 */

#ifndef _CVSSPILS_IMPL_H
#define _CVSSPILS_IMPL_H

#ifdef __cplusplus  /* wrapper to enable C++ usage */
extern "C" {
#endif

#include <cvodes/cvodes_spils.h>

  /* Types of iterative linear solvers */

#define SPILS_SPGMR   1
#define SPILS_SPBCG   2
#define SPILS_SPTFQMR 3


  /* 
   * -----------------------------------------------------------------
   * PART I - forward problems
   * -----------------------------------------------------------------
   */

  /*
   * -----------------------------------------------------------------
   * Types : CVSpilsMemRec, CVSpilsMem
   * -----------------------------------------------------------------
   * The type CVSpilsMem is pointer to a CVSpilsMemRec.
   * -----------------------------------------------------------------
   */

  typedef struct {

    int s_type;           /* type of scaled preconditioned iterative LS   */

    int  s_pretype;       /* type of preconditioning                      */
    int  s_gstype;        /* type of Gram-Schmidt orthogonalization       */
    realtype s_sqrtN;     /* sqrt(N)                                      */
    realtype s_delt;      /* delt = user specified or DELT_DEFAULT        */
    realtype s_deltar;    /* deltar = delt * tq4                          */
    realtype s_delta;     /* delta = deltar * sqrtN                       */
    int  s_maxl;          /* maxl = maximum dimension of the Krylov space */

    long int s_nstlpre;   /* value of nst at the last pset call           */
    long int s_npe;       /* npe = total number of pset calls             */
    long int s_nli;       /* nli = total number of linear iterations      */
    long int s_nps;       /* nps = total number of psolve calls           */
    long int s_ncfl;      /* ncfl = total number of convergence failures  */
    long int s_njtimes;   /* njtimes = total number of calls to jtimes    */
    long int s_nfes;      /* nfeSG = total number of calls to f for     
                             difference quotient Jacobian-vector products */

    N_Vector s_ytemp;     /* temp vector passed to jtimes and psolve      */
    N_Vector s_x;         /* temp vector used by CVSpilsSolve             */
    N_Vector s_ycur;      /* CVODE current y vector in Newton Iteration   */
    N_Vector s_fcur;      /* fcur = f(tn, ycur)                           */

    CVSpilsPrecSetupFn s_pset; 
    /* pset = user-supplied routine to compute      */
    /* a preconditioner                             */

    CVSpilsPrecSolveFn s_psolve;   
    /* psolve = user-supplied routine to solve      */
    /* preconditioner linear system                 */

    void *s_P_data;       /* P_data passed to psolve and pset             */

    void* s_spils_mem;    /* memory used by the generic solver            */

    CVSpilsJacTimesVecFn s_jtimes;  
    /* jtimes = Jacobian * vector routine           */
    void *s_j_data;       /* j_data is passed to jtimes                   */

    int s_last_flag;      /* last error flag returned by any function     */

  } CVSpilsMemRec, *CVSpilsMem;

  /*
   * -----------------------------------------------------------------
   * Prototypes of internal functions
   * -----------------------------------------------------------------
   */

  /* Atimes and PSolve routines called by generic solver */

  int CVSpilsAtimes(void *cv_mem, N_Vector v, N_Vector z);

  int CVSpilsPSolve(void *cv_mem, N_Vector r, N_Vector z, int lr);

  /* Difference quotient approximation for Jac times vector */

  int CVSpilsDQJtimes(N_Vector v, N_Vector Jv, realtype t,
                      N_Vector y, N_Vector fy, void *jac_data,
                      N_Vector work);


  /*
   * -----------------------------------------------------------------
   * Error Messages
   * -----------------------------------------------------------------
   */

#define MSGS_CVMEM_NULL  "Integrator memory is NULL."
#define MSGS_MEM_FAIL    "A memory request failed."
#define MSGS_BAD_NVECTOR "A required vector operation is not implemented."
#define MSGS_BAD_LSTYPE  "Incompatible linear solver type."
#define MSGS_BAD_PRETYPE "Illegal value for pretype. Legal values are PREC_NONE, PREC_LEFT, PREC_RIGHT, and PREC_BOTH."
#define MSGS_PSOLVE_REQ  "pretype != PREC_NONE, but PSOLVE = NULL is illegal."
#define MSGS_LMEM_NULL   "Linear solver memory is NULL."
#define MSGS_BAD_GSTYPE  "Illegal value for gstype. Legal values are MODIFIED_GS and CLASSICAL_GS."
#define MSGS_BAD_DELT    "delt < 0 illegal."
  
#define MSGS_PSET_FAILED "The preconditioner setup routine failed in an unrecoverable manner."
#define MSGS_PSOLVE_FAILED "The preconditioner solve routine failed in an unrecoverable manner."
#define MSGS_JTIMES_FAILED "The Jacobian x vector routine failed in an unrecoverable manner."


  /* 
   * -----------------------------------------------------------------
   * PART II - backward problems
   * -----------------------------------------------------------------
   */

  /*
   * -----------------------------------------------------------------
   * Types : CVSpilsMemRecB, CVSpilsMemB       
   * -----------------------------------------------------------------
   * CVSpgmrB, CVSpbcgB, and CVSptfqmr attach such a structure to the 
   * lmemB filed of CVadjMem
   * -----------------------------------------------------------------
   */

  typedef struct {

    CVSpilsJacTimesVecFnB s_jtimesB;
    CVSpilsPrecSetupFnB s_psetB;
    CVSpilsPrecSolveFnB s_psolveB;
    void *s_P_dataB;
    void *s_jac_dataB;

  } CVSpilsMemRecB, *CVSpilsMemB;


  /*
   * ------------------------------------------------
   * Wrapper functions for using the iterative linear 
   * solvers on adjoint (backward) problems
   * ------------------------------------------------
   */

  /* 
   * CVAspilsPrecSetup has type CVSpilsPrecSetupFn
   * It wraps around the user-provided function of type CVSpilsPrecSetupFnB
   */

  int CVAspilsPrecSetup(realtype t, N_Vector yB, 
                        N_Vector fyB, booleantype jokB, 
                        booleantype *jcurPtrB, realtype gammaB,
                        void *cvadj_mem,
                        N_Vector tmp1B, N_Vector tmp2B, N_Vector tmp3B);

  /* 
   * CVAspilsPrecSolve has type CVSpilsPrecSolveFn 
   * It wraps around the user-provided function of type CVSpilsPrecSolveFnB
   */

  int CVAspilsPrecSolve(realtype t, N_Vector yB, N_Vector fyB,
                        N_Vector rB, N_Vector zB,
                        realtype gammaB, realtype deltaB,
                        int lrB, void *cvadj_mem, N_Vector tmpB);
  
  /* 
   * CVAspilsJacTimesVec has type CVSpilsJacTimesVecFn 
   * It wraps around the user-provided function of type CVSpilsJacTimesVecFnB
   */

  int CVAspilsJacTimesVec(N_Vector vB, N_Vector JvB, realtype t, 
                          N_Vector yB, N_Vector fyB, 
                          void *cvadj_mem, N_Vector tmpB);

  /*
   * -----------------------------------------------------------------
   * Error Messages 
   * -----------------------------------------------------------------
   */

#define MSGS_CAMEM_NULL "cvadj_mem = NULL illegal."
#define MSGS_LMEMB_NULL "Linear solver memory is NULL for the backward integration."
#define MSGS_BAD_T      "Bad t for interpolation."


#ifdef __cplusplus
}
#endif

#endif
```
