## Supplementary scripts for "A multi-scale *in silico* mouse model for insulin resistance and humanoid type 2 diabetes": fnvector_parallel.h

```
/*
 * -----------------------------------------------------------------
 * $Revision: 749 $
 * $Date: 2011-06-22 14:49:10 +0200 (Wed, 22 Jun 2011) $
 * ----------------------------------------------------------------- 
 * Programmer(s): Radu Serban and Aaron Collier @ LLNL
 * -----------------------------------------------------------------
 * Copyright (c) 2002, The Regents of the University of California.
 * Produced at the Lawrence Livermore National Laboratory.
 * All rights reserved.
 * For details, see the LICENSE file.
 * -----------------------------------------------------------------
 * This file (companion of nvector_parallel.c) contains the
 * definitions needed for the initialization of parallel
 * vector operations in Fortran.
 * -----------------------------------------------------------------
 */

#ifndef _FNVECTOR_PARALLEL_H
#define _FNVECTOR_PARALLEL_H

#ifdef __cplusplus  /* wrapper to enable C++ usage */
extern "C" {
#endif

#include <nvector/nvector_parallel.h>  
#include <sundials/sundials_fnvector.h>

#if defined(F77_FUNC)

#define FNV_INITP    F77_FUNC(fnvinitp, FNVINITP)
#define FNV_INITP_Q  F77_FUNC_(fnvinitp_q, FNVINITP_Q)
#define FNV_INITP_S  F77_FUNC_(fnvinitp_s, FNVINITP_S)
#define FNV_INITP_B  F77_FUNC_(fnvinitp_b, FNVINITP_B)
#define FNV_INITP_QB F77_FUNC_(fnvinitp_qb, FNVINITP_QB)

#elif defined(SUNDIALS_UNDERSCORE_NONE) && defined(SUNDIALS_CASE_LOWER)

#define FNV_INITP    fnvinitp
#define FNV_INITP_Q  fnvinitp_q
#define FNV_INITP_S  fnvinitp_s
#define FNV_INITP_B  fnvinitp_b
#define FNV_INITP_QB fnvinitp_qb

#elif defined(SUNDIALS_UNDERSCORE_NONE) && defined(SUNDIALS_CASE_UPPER)

#define FNV_INITP    FNVINITP
#define FNV_INITP_Q  FNVINITP_Q
#define FNV_INITP_S  FNVINITP_S
#define FNV_INITP_B  FNVINITP_B
#define FNV_INITP_QB FNVINITP_QB

#elif defined(SUNDIALS_UNDERSCORE_ONE) && defined(SUNDIALS_CASE_LOWER)

#define FNV_INITP    fnvinitp_
#define FNV_INITP_Q  fnvinitp_q_
#define FNV_INITP_S  fnvinitp_s_
#define FNV_INITP_B  fnvinitp_b_
#define FNV_INITP_QB fnvinitp_qb_

#elif defined(SUNDIALS_UNDERSCORE_ONE) && defined(SUNDIALS_CASE_UPPER)

#define FNV_INITP    FNVINITP_
#define FNV_INITP_Q  FNVINITP_Q_
#define FNV_INITP_S  FNVINITP_S_
#define FNV_INITP_B  FNVINITP_B_
#define FNV_INITP_QB FNVINITP_QB_

#elif defined(SUNDIALS_UNDERSCORE_TWO) && defined(SUNDIALS_CASE_LOWER)

#define FNV_INITP    fnvinitp__
#define FNV_INITP_Q  fnvinitp_q__
#define FNV_INITP_S  fnvinitp_s__
#define FNV_INITP_B  fnvinitp_b__
#define FNV_INITP_QB fnvinitp_qb__

#elif defined(SUNDIALS_UNDERSCORE_TWO) && defined(SUNDIALS_CASE_UPPER)

#define FNV_INITP    FNVINITP__
#define FNV_INITP_Q  FNVINITP_Q__
#define FNV_INITP_S  FNVINITP_S__
#define FNV_INITP_B  FNVINITP_B__
#define FNV_INITP_QB FNVINITP_QB__

#endif

  /* Declarations of global variables */

  extern N_Vector F2C_CVODE_vec;
  extern N_Vector F2C_CVODE_vecQ;
  extern N_Vector *F2C_CVODE_vecS;
  extern N_Vector F2C_CVODE_vecB;
  extern N_Vector F2C_CVODE_vecQB;

  extern N_Vector F2C_IDA_vec;
  extern N_Vector F2C_IDA_vecQ;
  extern N_Vector *F2C_IDA_vecS;
  extern N_Vector F2C_IDA_vecB;
  extern N_Vector F2C_IDA_vecQB;

  extern N_Vector F2C_KINSOL_vec;

  /* 
   * Prototypes of exported functions 
   *
   * FNV_INITP    - initializes parallel vector operations for main problem
   * FNV_INITP_Q  - initializes parallel vector operations for quadratures
   * FNV_INITP_S  - initializes parallel vector operations for sensitivities
   * FNV_INITP_B  - initializes parallel vector operations for adjoint problem
   * FNV_INITP_QB - initializes parallel vector operations for adjoint quadratures
   *
   */

#ifndef SUNDIALS_MPI_COMM_F2C
#define MPI_Fint int
#endif

  void FNV_INITP(MPI_Fint *comm, int *code, long int *L, long int *N, int *ier);
  void FNV_INITP_Q(MPI_Fint *comm, int *code, long int *Lq, long int *Nq, int *ier);
  void FNV_INITP_B(MPI_Fint *comm, int *code, long int *LB, long int *NB, int *ier);
  void FNV_INITP_QB(MPI_Fint *comm, int *code, long int *LqB, long int *NqB, int *ier);
  void FNV_INITP_S(int *code, int *Ns, int *ier);

#ifdef __cplusplus
}
#endif

#endif
```
