## Supplementary scripts for "A multi-scale *in silico* mouse model for insulin resistance and humanoid type 2 diabetes": fnvector_serial.c

```
/*
 * -----------------------------------------------------------------
 * $Revision: 749 $
 * $Date: 2011-06-22 14:49:10 +0200 (Wed, 22 Jun 2011) $
 * ----------------------------------------------------------------- 
 * Programmer(s): Radu Serban @ LLNL
 * -----------------------------------------------------------------
 * Copyright (c) 2002, The Regents of the University of California.
 * Produced at the Lawrence Livermore National Laboratory.
 * All rights reserved.
 * For details, see the LICENSE file.
 * -----------------------------------------------------------------
 * This file (companion of nvector_serial.h) contains the
 * implementation needed for the Fortran initialization of serial
 * vector operations.
 * -----------------------------------------------------------------
 */

#include <stdio.h>
#include <stdlib.h>

#include "fnvector_serial.h"

/* Define global vector variables */

N_Vector F2C_CVODE_vec;
N_Vector F2C_CVODE_vecQ;
N_Vector *F2C_CVODE_vecS;
N_Vector F2C_CVODE_vecB;
N_Vector F2C_CVODE_vecQB;

N_Vector F2C_IDA_vec;
N_Vector F2C_IDA_vecQ;
N_Vector *F2C_IDA_vecS;
N_Vector F2C_IDA_vecB;
N_Vector F2C_IDA_vecQB;

N_Vector F2C_KINSOL_vec;

/* Fortran callable interfaces */

void FNV_INITS(int *code, long int *N, int *ier)
{
  *ier = 0;

  switch(*code) {
  case FCMIX_CVODE:
    F2C_CVODE_vec = NULL;
    F2C_CVODE_vec = N_VNewEmpty_Serial(*N);
    if (F2C_CVODE_vec == NULL) *ier = -1;
    break;
  case FCMIX_IDA:
    F2C_IDA_vec = NULL;
    F2C_IDA_vec = N_VNewEmpty_Serial(*N);
    if (F2C_IDA_vec == NULL) *ier = -1;
    break;
  case FCMIX_KINSOL:
    F2C_KINSOL_vec = NULL;
    F2C_KINSOL_vec = N_VNewEmpty_Serial(*N);
    if (F2C_KINSOL_vec == NULL) *ier = -1;
    break;
  default:
    *ier = -1;
  }
}

void FNV_INITS_Q(int *code, long int *Nq, int *ier)
{
  *ier = 0;

  switch(*code) {
  case FCMIX_CVODE:
    F2C_CVODE_vecQ = NULL;
    F2C_CVODE_vecQ = N_VNewEmpty_Serial(*Nq);
    if (F2C_CVODE_vecQ == NULL) *ier = -1;
    break;
  case FCMIX_IDA:
    F2C_IDA_vecQ = NULL;
    F2C_IDA_vecQ = N_VNewEmpty_Serial(*Nq);
    if (F2C_IDA_vecQ == NULL) *ier = -1;
    break;
  default:
    *ier = -1;
  }
}

void FNV_INITS_B(int *code, long int *NB, int *ier)
{
  *ier = 0;

  switch(*code) {
  case FCMIX_CVODE:
    F2C_CVODE_vecB = NULL;
    F2C_CVODE_vecB = N_VNewEmpty_Serial(*NB);
    if (F2C_CVODE_vecB == NULL) *ier = -1;
    break;
  case FCMIX_IDA:
    F2C_IDA_vecB = NULL;
    F2C_IDA_vecB = N_VNewEmpty_Serial(*NB);
    if (F2C_IDA_vecB == NULL) *ier = -1;
    break;
  default:
    *ier = -1;
  }
}

void FNV_INITS_QB(int *code, long int *NqB, int *ier)
{
  *ier = 0;

  switch(*code) {
  case FCMIX_CVODE:
    F2C_CVODE_vecQB = NULL;
    F2C_CVODE_vecQB = N_VNewEmpty_Serial(*NqB);
    if (F2C_CVODE_vecQB == NULL) *ier = -1;
    break;
  case FCMIX_IDA:
    F2C_IDA_vecQB = NULL;
    F2C_IDA_vecQB = N_VNewEmpty_Serial(*NqB);
    if (F2C_IDA_vecQB == NULL) *ier = -1;
    break;
  default:
    *ier = -1;
  }
}

void FNV_INITS_S(int *code, int *Ns, int *ier)
{
  *ier = 0;

  switch(*code) {
  case FCMIX_CVODE:
    F2C_CVODE_vecS = NULL;
    F2C_CVODE_vecS = (N_Vector *) N_VCloneVectorArrayEmpty_Serial(*Ns, F2C_CVODE_vec);
    if (F2C_CVODE_vecS == NULL) *ier = -1;
    break;
  case FCMIX_IDA:
    F2C_IDA_vecS = NULL;
    F2C_IDA_vecS = (N_Vector *) N_VCloneVectorArrayEmpty_Serial(*Ns, F2C_IDA_vec);
    if (F2C_IDA_vecS == NULL) *ier = -1;
    break;
  default:
    *ier = -1;
  }
}
```
