## Supplementary scripts for "A multi-scale *in silico* mouse model for insulin resistance and humanoid type 2 diabetes": fnvector_serial.h

```
/*
 * -----------------------------------------------------------------
 * $Revision: 749 $
 * $Date: 2011-06-22 14:49:10 +0200 (Wed, 22 Jun 2011) $
 * ----------------------------------------------------------------- 
 * Programmer(s): Radu Serban and Aaron Collier @ LLNL
 * -----------------------------------------------------------------
 * Copyright (c) 2002, The Regents of the University of California.
 * Produced at the Lawrence Livermore National Laboratory.
 * All rights reserved.
 * For details, see the LICENSE file.
 * -----------------------------------------------------------------
 * This file (companion of nvector_serial.h) contains the
 * definitions needed for the initialization of serial
 * vector operations in Fortran.
 * -----------------------------------------------------------------
 */

#ifndef _FNVECTOR_SERIAL_H
#define _FNVECTOR_SERIAL_H

#ifdef __cplusplus  /* wrapper to enable C++ usage */
extern "C" {
#endif

#include <nvector/nvector_serial.h>  
#include <sundials/sundials_fnvector.h>

#if defined(F77_FUNC)

#define FNV_INITS    F77_FUNC(fnvinits, FNVINITS)
#define FNV_INITS_Q  F77_FUNC_(fnvinits_q, FNVINITS_Q)
#define FNV_INITS_S  F77_FUNC_(fnvinits_s, FNVINITS_S)
#define FNV_INITS_B  F77_FUNC_(fnvinits_b, FNVINITS_B)
#define FNV_INITS_QB F77_FUNC_(fnvinits_qb, FNVINITS_QB)

#elif defined(SUNDIALS_UNDERSCORE_NONE) && defined(SUNDIALS_CASE_LOWER)

#define FNV_INITS    fnvinits
#define FNV_INITS_Q  fnvinits_q
#define FNV_INITS_S  fnvinits_s
#define FNV_INITS_B  fnvinits_b
#define FNV_INITS_QB fnvinits_qb

#elif defined(SUNDIALS_UNDERSCORE_NONE) && defined(SUNDIALS_CASE_UPPER)

#define FNV_INITS    FNVINITS
#define FNV_INITS_Q  FNVINITS_Q
#define FNV_INITS_S  FNVINITS_S
#define FNV_INITS_B  FNVINITS_B
#define FNV_INITS_QB FNVINITS_QB

#elif defined(SUNDIALS_UNDERSCORE_ONE) && defined(SUNDIALS_CASE_LOWER)

#define FNV_INITS    fnvinits_
#define FNV_INITS_Q  fnvinits_q_
#define FNV_INITS_S  fnvinits_s_
#define FNV_INITS_B  fnvinits_b_
#define FNV_INITS_QB fnvinits_qb_

#elif defined(SUNDIALS_UNDERSCORE_ONE) && defined(SUNDIALS_CASE_UPPER)

#define FNV_INITS    FNVINITS_
#define FNV_INITS_Q  FNVINITS_Q_
#define FNV_INITS_S  FNVINITS_S_
#define FNV_INITS_B  FNVINITS_B_
#define FNV_INITS_QB FNVINITS_QB_

#elif defined(SUNDIALS_UNDERSCORE_TWO) && defined(SUNDIALS_CASE_LOWER)

#define FNV_INITS    fnvinits__
#define FNV_INITS_Q  fnvinits_q__
#define FNV_INITS_S  fnvinits_s__
#define FNV_INITS_B  fnvinits_b__
#define FNV_INITS_QB fnvinits_qb__

#elif defined(SUNDIALS_UNDERSCORE_TWO) && defined(SUNDIALS_CASE_UPPER)

#define FNV_INITS    FNVINITS__
#define FNV_INITS_Q  FNVINITS_Q__
#define FNV_INITS_S  FNVINITS_S__
#define FNV_INITS_B  FNVINITS_B__
#define FNV_INITS_QB FNVINITS_QB__

#endif

  /* Declarations of global variables */

  extern N_Vector F2C_CVODE_vec;
  extern N_Vector F2C_CVODE_vecQ;
  extern N_Vector *F2C_CVODE_vecS;
  extern N_Vector F2C_CVODE_vecB;
  extern N_Vector F2C_CVODE_vecQB;

  extern N_Vector F2C_IDA_vec;
  extern N_Vector F2C_IDA_vecQ;
  extern N_Vector *F2C_IDA_vecS;
  extern N_Vector F2C_IDA_vecB;
  extern N_Vector F2C_IDA_vecQB;

  extern N_Vector F2C_KINSOL_vec;

  /* 
   * Prototypes of exported functions 
   *
   * FNV_INITS    - initializes serial vector operations for main problem
   * FNV_INITS_Q  - initializes serial vector operations for quadratures
   * FNV_INITS_S  - initializes serial vector operations for sensitivities
   * FNV_INITS_B  - initializes serial vector operations for adjoint problem
   * FNV_INITS_QB - initializes serial vector operations for adjoint quadratures
   *
   */

  void FNV_INITS(int *code, long int *neq, int *ier);
  void FNV_INITS_Q(int *code, long int *Nq, int *ier);
  void FNV_INITS_S(int *code, int *Ns, int *ier);
  void FNV_INITS_B(int *code, long int *NB, int *ier);
  void FNV_INITS_QB(int *code, long int *NqB, int *ier);

#ifdef __cplusplus
}
#endif

#endif
```
