## Supplementary scripts for "A multi-scale *in silico* mouse model for insulin resistance and humanoid type 2 diabetes": nvector_serial.c

```
/*
 * -----------------------------------------------------------------
 * $Revision: 749 $
 * $Date: 2011-06-22 14:49:10 +0200 (Wed, 22 Jun 2011) $
 * ----------------------------------------------------------------- 
 * Programmer(s): Scott D. Cohen, Alan C. Hindmarsh, Radu Serban,
 *                and Aaron Collier @ LLNL
 * -----------------------------------------------------------------
 * Copyright (c) 2002, The Regents of the University of California.
 * Produced at the Lawrence Livermore National Laboratory.
 * All rights reserved.
 * For details, see the LICENSE file.
 * -----------------------------------------------------------------
 * This is the implementation file for a serial implementation
 * of the NVECTOR package.
 * -----------------------------------------------------------------
 */

#include <stdio.h>
#include <stdlib.h>

#include <nvector/nvector_serial.h>
#include <sundials/sundials_math.h>

#define ZERO   RCONST(0.0)
#define HALF   RCONST(0.5)
#define ONE    RCONST(1.0)
#define ONEPT5 RCONST(1.5)

/* Private function prototypes */
/* z=x */
static void VCopy_Serial(N_Vector x, N_Vector z);
/* z=x+y */
static void VSum_Serial(N_Vector x, N_Vector y, N_Vector z);
/* z=x-y */
static void VDiff_Serial(N_Vector x, N_Vector y, N_Vector z);
/* z=-x */
static void VNeg_Serial(N_Vector x, N_Vector z);
/* z=c(x+y) */
static void VScaleSum_Serial(realtype c, N_Vector x, N_Vector y, N_Vector z);
/* z=c(x-y) */
static void VScaleDiff_Serial(realtype c, N_Vector x, N_Vector y, N_Vector z); 
/* z=ax+y */
static void VLin1_Serial(realtype a, N_Vector x, N_Vector y, N_Vector z);
/* z=ax-y */
static void VLin2_Serial(realtype a, N_Vector x, N_Vector y, N_Vector z);
/* y <- ax+y */
static void Vaxpy_Serial(realtype a, N_Vector x, N_Vector y);
/* x <- ax */
static void VScaleBy_Serial(realtype a, N_Vector x);

/*
 * -----------------------------------------------------------------
 * exported functions
 * -----------------------------------------------------------------
 */

/* ----------------------------------------------------------------------------
 * Function to create a new empty serial vector 
 */

N_Vector N_VNewEmpty_Serial(long int length)
{
  N_Vector v;
  N_Vector_Ops ops;
  N_VectorContent_Serial content;

  /* Create vector */
  v = NULL;
  v = (N_Vector) malloc(sizeof *v);
  if (v == NULL) return(NULL);
  
  /* Create vector operation structure */
  ops = NULL;
  ops = (N_Vector_Ops) malloc(sizeof(struct _generic_N_Vector_Ops));
  if (ops == NULL) { free(v); return(NULL); }

  ops->nvclone           = N_VClone_Serial;
  ops->nvcloneempty      = N_VCloneEmpty_Serial;
  ops->nvdestroy         = N_VDestroy_Serial;
  ops->nvspace           = N_VSpace_Serial;
  ops->nvgetarraypointer = N_VGetArrayPointer_Serial;
  ops->nvsetarraypointer = N_VSetArrayPointer_Serial;
  ops->nvlinearsum       = N_VLinearSum_Serial;
  ops->nvconst           = N_VConst_Serial;
  ops->nvprod            = N_VProd_Serial;
  ops->nvdiv             = N_VDiv_Serial;
  ops->nvscale           = N_VScale_Serial;
  ops->nvabs             = N_VAbs_Serial;
  ops->nvinv             = N_VInv_Serial;
  ops->nvaddconst        = N_VAddConst_Serial;
  ops->nvdotprod         = N_VDotProd_Serial;
  ops->nvmaxnorm         = N_VMaxNorm_Serial;
  ops->nvwrmsnormmask    = N_VWrmsNormMask_Serial;
  ops->nvwrmsnorm        = N_VWrmsNorm_Serial;
  ops->nvmin             = N_VMin_Serial;
  ops->nvwl2norm         = N_VWL2Norm_Serial;
  ops->nvl1norm          = N_VL1Norm_Serial;
  ops->nvcompare         = N_VCompare_Serial;
  ops->nvinvtest         = N_VInvTest_Serial;
  ops->nvconstrmask      = N_VConstrMask_Serial;
  ops->nvminquotient     = N_VMinQuotient_Serial;

  /* Create content */
  content = NULL;
  content = (N_VectorContent_Serial) malloc(sizeof(struct _N_VectorContent_Serial));
  if (content == NULL) { free(ops); free(v); return(NULL); }

  content->length   = length;
  content->own_data = FALSE;
  content->data     = NULL;

  /* Attach content and ops */
  v->content = content;
  v->ops     = ops;

  return(v);
}

/* ----------------------------------------------------------------------------
 * Function to create a new serial vector 
 */

N_Vector N_VNew_Serial(long int length)
{
  N_Vector v;
  realtype *data;

  v = NULL;
  v = N_VNewEmpty_Serial(length);
  if (v == NULL) return(NULL);

  /* Create data */
  if (length > 0) {

    /* Allocate memory */
    data = NULL;
    data = (realtype *) malloc(length * sizeof(realtype));
    if(data == NULL) { N_VDestroy_Serial(v); return(NULL); }

    /* Attach data */
    NV_OWN_DATA_S(v) = TRUE;
    NV_DATA_S(v)     = data;

  }

  return(v);
}

/* ----------------------------------------------------------------------------
 * Function to create a serial N_Vector with user data component 
 */

N_Vector N_VMake_Serial(long int length, realtype *v_data)
{
  N_Vector v;

  v = NULL;
  v = N_VNewEmpty_Serial(length);
  if (v == NULL) return(NULL);

  if (length > 0) {
    /* Attach data */
    NV_OWN_DATA_S(v) = FALSE;
    NV_DATA_S(v)     = v_data;
  }

  return(v);
}

/* ----------------------------------------------------------------------------
 * Function to create an array of new serial vectors. 
 */

N_Vector *N_VCloneVectorArray_Serial(int count, N_Vector w)
{
  N_Vector *vs;
  int j;

  if (count <= 0) return(NULL);

  vs = NULL;
  vs = (N_Vector *) malloc(count * sizeof(N_Vector));
  if(vs == NULL) return(NULL);

  for (j = 0; j < count; j++) {
    vs[j] = NULL;
    vs[j] = N_VClone_Serial(w);
    if (vs[j] == NULL) {
      N_VDestroyVectorArray_Serial(vs, j-1);
      return(NULL);
    }
  }

  return(vs);
}

/* ----------------------------------------------------------------------------
 * Function to create an array of new serial vectors with NULL data array. 
 */

N_Vector *N_VCloneVectorArrayEmpty_Serial(int count, N_Vector w)
{
  N_Vector *vs;
  int j;

  if (count <= 0) return(NULL);

  vs = NULL;
  vs = (N_Vector *) malloc(count * sizeof(N_Vector));
  if(vs == NULL) return(NULL);

  for (j = 0; j < count; j++) {
    vs[j] = NULL;
    vs[j] = N_VCloneEmpty_Serial(w);
    if (vs[j] == NULL) {
      N_VDestroyVectorArray_Serial(vs, j-1);
      return(NULL);
    }
  }

  return(vs);
}

/* ----------------------------------------------------------------------------
 * Function to free an array created with N_VCloneVectorArray_Serial
 */

void N_VDestroyVectorArray_Serial(N_Vector *vs, int count)
{
  int j;

  for (j = 0; j < count; j++) N_VDestroy_Serial(vs[j]);

  free(vs); vs = NULL;

  return;
}

/* ----------------------------------------------------------------------------
 * Function to print the a serial vector 
 */
 
void N_VPrint_Serial(N_Vector x)
{
  long int i, N;
  realtype *xd;

  xd = NULL;

  N  = NV_LENGTH_S(x);
  xd = NV_DATA_S(x);

  for (i = 0; i < N; i++) {
#if defined(SUNDIALS_EXTENDED_PRECISION)
    printf("%11.8Lg\n", xd[i]);
#elif defined(SUNDIALS_DOUBLE_PRECISION)
    printf("%11.8lg\n", xd[i]);
#else
    printf("%11.8g\n", xd[i]);
#endif
  }
  printf("\n");

  return;
}

/*
 * -----------------------------------------------------------------
 * implementation of vector operations
 * -----------------------------------------------------------------
 */

N_Vector N_VCloneEmpty_Serial(N_Vector w)
{
  N_Vector v;
  N_Vector_Ops ops;
  N_VectorContent_Serial content;

  if (w == NULL) return(NULL);

  /* Create vector */
  v = NULL;
  v = (N_Vector) malloc(sizeof *v);
  if (v == NULL) return(NULL);

  /* Create vector operation structure */
  ops = NULL;
  ops = (N_Vector_Ops) malloc(sizeof(struct _generic_N_Vector_Ops));
  if (ops == NULL) { free(v); return(NULL); }
  
  ops->nvclone           = w->ops->nvclone;
  ops->nvcloneempty      = w->ops->nvcloneempty;
  ops->nvdestroy         = w->ops->nvdestroy;
  ops->nvspace           = w->ops->nvspace;
  ops->nvgetarraypointer = w->ops->nvgetarraypointer;
  ops->nvsetarraypointer = w->ops->nvsetarraypointer;
  ops->nvlinearsum       = w->ops->nvlinearsum;
  ops->nvconst           = w->ops->nvconst;  
  ops->nvprod            = w->ops->nvprod;   
  ops->nvdiv             = w->ops->nvdiv;
  ops->nvscale           = w->ops->nvscale; 
  ops->nvabs             = w->ops->nvabs;
  ops->nvinv             = w->ops->nvinv;
  ops->nvaddconst        = w->ops->nvaddconst;
  ops->nvdotprod         = w->ops->nvdotprod;
  ops->nvmaxnorm         = w->ops->nvmaxnorm;
  ops->nvwrmsnormmask    = w->ops->nvwrmsnormmask;
  ops->nvwrmsnorm        = w->ops->nvwrmsnorm;
  ops->nvmin             = w->ops->nvmin;
  ops->nvwl2norm         = w->ops->nvwl2norm;
  ops->nvl1norm          = w->ops->nvl1norm;
  ops->nvcompare         = w->ops->nvcompare;    
  ops->nvinvtest         = w->ops->nvinvtest;
  ops->nvconstrmask      = w->ops->nvconstrmask;
  ops->nvminquotient     = w->ops->nvminquotient;

  /* Create content */
  content = NULL;
  content = (N_VectorContent_Serial) malloc(sizeof(struct _N_VectorContent_Serial));
  if (content == NULL) { free(ops); free(v); return(NULL); }

  content->length   = NV_LENGTH_S(w);
  content->own_data = FALSE;
  content->data     = NULL;

  /* Attach content and ops */
  v->content = content;
  v->ops     = ops;

  return(v);
}

N_Vector N_VClone_Serial(N_Vector w)
{
  N_Vector v;
  realtype *data;
  long int length;

  v = NULL;
  v = N_VCloneEmpty_Serial(w);
  if (v == NULL) return(NULL);

  length = NV_LENGTH_S(w);

  /* Create data */
  if (length > 0) {

    /* Allocate memory */
    data = NULL;
    data = (realtype *) malloc(length * sizeof(realtype));
    if(data == NULL) { N_VDestroy_Serial(v); return(NULL); }

    /* Attach data */
    NV_OWN_DATA_S(v) = TRUE;
    NV_DATA_S(v)     = data;

  }

  return(v);
}

void N_VDestroy_Serial(N_Vector v)
{
  if (NV_OWN_DATA_S(v) == TRUE) {
    free(NV_DATA_S(v));
    NV_DATA_S(v) = NULL;
  }
  free(v->content); v->content = NULL;
  free(v->ops); v->ops = NULL;
  free(v); v = NULL;

  return;
}

void N_VSpace_Serial(N_Vector v, long int *lrw, long int *liw)
{
  *lrw = NV_LENGTH_S(v);
  *liw = 1;

  return;
}

realtype *N_VGetArrayPointer_Serial(N_Vector v)
{
  return((realtype *) NV_DATA_S(v));
}

void N_VSetArrayPointer_Serial(realtype *v_data, N_Vector v)
{
  if (NV_LENGTH_S(v) > 0) NV_DATA_S(v) = v_data;

  return;
}

void N_VLinearSum_Serial(realtype a, N_Vector x, realtype b, N_Vector y, N_Vector z)
{
  long int i, N;
  realtype c, *xd, *yd, *zd;
  N_Vector v1, v2;
  booleantype test;

  xd = yd = zd = NULL;

  if ((b == ONE) && (z == y)) {    /* BLAS usage: axpy y <- ax+y */
    Vaxpy_Serial(a,x,y);
    return;
  }

  if ((a == ONE) && (z == x)) {    /* BLAS usage: axpy x <- by+x */
    Vaxpy_Serial(b,y,x);
    return;
  }

  /* Case: a == b == 1.0 */

  if ((a == ONE) && (b == ONE)) {
    VSum_Serial(x, y, z);
    return;
  }

  /* Cases: (1) a == 1.0, b = -1.0, (2) a == -1.0, b == 1.0 */

  if ((test = ((a == ONE) && (b == -ONE))) || ((a == -ONE) && (b == ONE))) {
    v1 = test ? y : x;
    v2 = test ? x : y;
    VDiff_Serial(v2, v1, z);
    return;
  }

  /* Cases: (1) a == 1.0, b == other or 0.0, (2) a == other or 0.0, b == 1.0 */
  /* if a or b is 0.0, then user should have called N_VScale */

  if ((test = (a == ONE)) || (b == ONE)) {
    c  = test ? b : a;
    v1 = test ? y : x;
    v2 = test ? x : y;
    VLin1_Serial(c, v1, v2, z);
    return;
  }

  /* Cases: (1) a == -1.0, b != 1.0, (2) a != 1.0, b == -1.0 */

  if ((test = (a == -ONE)) || (b == -ONE)) {
    c = test ? b : a;
    v1 = test ? y : x;
    v2 = test ? x : y;
    VLin2_Serial(c, v1, v2, z);
    return;
  }

  /* Case: a == b */
  /* catches case both a and b are 0.0 - user should have called N_VConst */

  if (a == b) {
    VScaleSum_Serial(a, x, y, z);
    return;
  }

  /* Case: a == -b */

  if (a == -b) {
    VScaleDiff_Serial(a, x, y, z);
    return;
  }

  /* Do all cases not handled above:
     (1) a == other, b == 0.0 - user should have called N_VScale
     (2) a == 0.0, b == other - user should have called N_VScale
     (3) a,b == other, a !=b, a != -b */
  
  N  = NV_LENGTH_S(x);
  xd = NV_DATA_S(x);
  yd = NV_DATA_S(y);
  zd = NV_DATA_S(z);

  for (i = 0; i < N; i++)
    zd[i] = (a*xd[i])+(b*yd[i]);

  return;
}

void N_VConst_Serial(realtype c, N_Vector z)
{
  long int i, N;
  realtype *zd;

  zd = NULL;

  N  = NV_LENGTH_S(z);
  zd = NV_DATA_S(z);

  for (i = 0; i < N; i++) zd[i] = c;

  return;
}

void N_VProd_Serial(N_Vector x, N_Vector y, N_Vector z)
{
  long int i, N;
  realtype *xd, *yd, *zd;

  xd = yd = zd = NULL;

  N  = NV_LENGTH_S(x);
  xd = NV_DATA_S(x);
  yd = NV_DATA_S(y);
  zd = NV_DATA_S(z);

  for (i = 0; i < N; i++)
    zd[i] = xd[i]*yd[i];

  return;
}

void N_VDiv_Serial(N_Vector x, N_Vector y, N_Vector z)
{
  long int i, N;
  realtype *xd, *yd, *zd;

  xd = yd = zd = NULL;

  N  = NV_LENGTH_S(x);
  xd = NV_DATA_S(x);
  yd = NV_DATA_S(y);
  zd = NV_DATA_S(z);

  for (i = 0; i < N; i++)
    zd[i] = xd[i]/yd[i];

  return;
}

void N_VScale_Serial(realtype c, N_Vector x, N_Vector z)
{
  long int i, N;
  realtype *xd, *zd;

  xd = zd = NULL;

  if (z == x) {  /* BLAS usage: scale x <- cx */
    VScaleBy_Serial(c, x);
    return;
  }

  if (c == ONE) {
    VCopy_Serial(x, z);
  } else if (c == -ONE) {
    VNeg_Serial(x, z);
  } else {
    N  = NV_LENGTH_S(x);
    xd = NV_DATA_S(x);
    zd = NV_DATA_S(z);
    for (i = 0; i < N; i++) 
      zd[i] = c*xd[i];
  }

  return;
}

void N_VAbs_Serial(N_Vector x, N_Vector z)
{
  long int i, N;
  realtype *xd, *zd;

  xd = zd = NULL;

  N  = NV_LENGTH_S(x);
  xd = NV_DATA_S(x);
  zd = NV_DATA_S(z);

  for (i = 0; i < N; i++)
    zd[i] = ABS(xd[i]);

  return;
}

void N_VInv_Serial(N_Vector x, N_Vector z)
{
  long int i, N;
  realtype *xd, *zd;

  xd = zd = NULL;

  N  = NV_LENGTH_S(x);
  xd = NV_DATA_S(x);
  zd = NV_DATA_S(z);

  for (i = 0; i < N; i++)
    zd[i] = ONE/xd[i];

  return;
}

void N_VAddConst_Serial(N_Vector x, realtype b, N_Vector z)
{
  long int i, N;
  realtype *xd, *zd;

  xd = zd = NULL;

  N  = NV_LENGTH_S(x);
  xd = NV_DATA_S(x);
  zd = NV_DATA_S(z);

  for (i = 0; i < N; i++) 
    zd[i] = xd[i]+b;

  return;
}

realtype N_VDotProd_Serial(N_Vector x, N_Vector y)
{
  long int i, N;
  realtype sum, *xd, *yd;

  sum = ZERO;
  xd = yd = NULL;

  N  = NV_LENGTH_S(x);
  xd = NV_DATA_S(x);
  yd = NV_DATA_S(y);

  for (i = 0; i < N; i++)
    sum += xd[i]*yd[i];
  
  return(sum);
}

realtype N_VMaxNorm_Serial(N_Vector x)
{
  long int i, N;
  realtype max, *xd;

  max = ZERO;
  xd = NULL;

  N  = NV_LENGTH_S(x);
  xd = NV_DATA_S(x);

  for (i = 0; i < N; i++) {
    if (ABS(xd[i]) > max) max = ABS(xd[i]);
  }

  return(max);
}

realtype N_VWrmsNorm_Serial(N_Vector x, N_Vector w)
{
  long int i, N;
  realtype sum, prodi, *xd, *wd;

  sum = ZERO;
  xd = wd = NULL;

  N  = NV_LENGTH_S(x);
  xd = NV_DATA_S(x);
  wd = NV_DATA_S(w);

  for (i = 0; i < N; i++) {
    prodi = xd[i]*wd[i];
    sum += SQR(prodi);
  }

  return(RSqrt(sum/N));
}

realtype N_VWrmsNormMask_Serial(N_Vector x, N_Vector w, N_Vector id)
{
  long int i, N;
  realtype sum, prodi, *xd, *wd, *idd;

  sum = ZERO;
  xd = wd = idd = NULL;

  N  = NV_LENGTH_S(x);
  xd  = NV_DATA_S(x);
  wd  = NV_DATA_S(w);
  idd = NV_DATA_S(id);

  for (i = 0; i < N; i++) {
    if (idd[i] > ZERO) {
      prodi = xd[i]*wd[i];
      sum += SQR(prodi);
    }
  }

  return(RSqrt(sum / N));
}

realtype N_VMin_Serial(N_Vector x)
{
  long int i, N;
  realtype min, *xd;

  xd = NULL;

  N  = NV_LENGTH_S(x);
  xd = NV_DATA_S(x);

  min = xd[0];

  for (i = 1; i < N; i++) {
    if (xd[i] < min) min = xd[i];
  }

  return(min);
}

realtype N_VWL2Norm_Serial(N_Vector x, N_Vector w)
{
  long int i, N;
  realtype sum, prodi, *xd, *wd;

  sum = ZERO;
  xd = wd = NULL;

  N  = NV_LENGTH_S(x);
  xd = NV_DATA_S(x);
  wd = NV_DATA_S(w);

  for (i = 0; i < N; i++) {
    prodi = xd[i]*wd[i];
    sum += SQR(prodi);
  }

  return(RSqrt(sum));
}

realtype N_VL1Norm_Serial(N_Vector x)
{
  long int i, N;
  realtype sum, *xd;

  sum = ZERO;
  xd = NULL;

  N  = NV_LENGTH_S(x);
  xd = NV_DATA_S(x);
  
  for (i = 0; i<N; i++)  
    sum += ABS(xd[i]);

  return(sum);
}

void N_VCompare_Serial(realtype c, N_Vector x, N_Vector z)
{
  long int i, N;
  realtype *xd, *zd;

  xd = zd = NULL;

  N  = NV_LENGTH_S(x);
  xd = NV_DATA_S(x);
  zd = NV_DATA_S(z);

  for (i = 0; i < N; i++) {
    zd[i] = (ABS(xd[i]) >= c) ? ONE : ZERO;
  }

  return;
}

booleantype N_VInvTest_Serial(N_Vector x, N_Vector z)
{
  long int i, N;
  realtype *xd, *zd;

  xd = zd = NULL;

  N  = NV_LENGTH_S(x);
  xd = NV_DATA_S(x);
  zd = NV_DATA_S(z);

  for (i = 0; i < N; i++) {
    if (xd[i] == ZERO) return(FALSE);
    zd[i] = ONE/xd[i];
  }

  return(TRUE);
}

booleantype N_VConstrMask_Serial(N_Vector c, N_Vector x, N_Vector m)
{
  long int i, N;
  booleantype test;
  realtype *cd, *xd, *md;

  cd = xd = md = NULL;

  N  = NV_LENGTH_S(x);
  xd = NV_DATA_S(x);
  cd = NV_DATA_S(c);
  md = NV_DATA_S(m);

  test = TRUE;

  for (i = 0; i < N; i++) {
    md[i] = ZERO;
    if (cd[i] == ZERO) continue;
    if (cd[i] > ONEPT5 || cd[i] < -ONEPT5) {
      if ( xd[i]*cd[i] <= ZERO) { test = FALSE; md[i] = ONE; }
      continue;
    }
    if ( cd[i] > HALF || cd[i] < -HALF) {
      if (xd[i]*cd[i] < ZERO ) { test = FALSE; md[i] = ONE; }
    }
  }

  return(test);
}

realtype N_VMinQuotient_Serial(N_Vector num, N_Vector denom)
{
  booleantype notEvenOnce;
  long int i, N;
  realtype *nd, *dd, min;

  nd = dd = NULL;

  N  = NV_LENGTH_S(num);
  nd = NV_DATA_S(num);
  dd = NV_DATA_S(denom);

  notEvenOnce = TRUE;
  min = BIG_REAL;

  for (i = 0; i < N; i++) {
    if (dd[i] == ZERO) continue;
    else {
      if (!notEvenOnce) min = MIN(min, nd[i]/dd[i]);
      else {
	min = nd[i]/dd[i];
        notEvenOnce = FALSE;
      }
    }
  }

  return(min);
}

/*
 * -----------------------------------------------------------------
 * private functions
 * -----------------------------------------------------------------
 */

static void VCopy_Serial(N_Vector x, N_Vector z)
{
  long int i, N;
  realtype *xd, *zd;

  xd = zd = NULL;

  N  = NV_LENGTH_S(x);
  xd = NV_DATA_S(x);
  zd = NV_DATA_S(z);

  for (i = 0; i < N; i++)
    zd[i] = xd[i]; 

  return;
}

static void VSum_Serial(N_Vector x, N_Vector y, N_Vector z)
{
  long int i, N;
  realtype *xd, *yd, *zd;

  xd = yd = zd = NULL;

  N  = NV_LENGTH_S(x);
  xd = NV_DATA_S(x);
  yd = NV_DATA_S(y);
  zd = NV_DATA_S(z);

  for (i = 0; i < N; i++)
    zd[i] = xd[i]+yd[i];

  return;
}

static void VDiff_Serial(N_Vector x, N_Vector y, N_Vector z)
{
  long int i, N;
  realtype *xd, *yd, *zd;

  xd = yd = zd = NULL;

  N  = NV_LENGTH_S(x);
  xd = NV_DATA_S(x);
  yd = NV_DATA_S(y);
  zd = NV_DATA_S(z);

  for (i = 0; i < N; i++)
    zd[i] = xd[i]-yd[i];

  return;
}

static void VNeg_Serial(N_Vector x, N_Vector z)
{
  long int i, N;
  realtype *xd, *zd;

  xd = zd = NULL;

  N  = NV_LENGTH_S(x);
  xd = NV_DATA_S(x);
  zd = NV_DATA_S(z);
  
  for (i = 0; i < N; i++)
    zd[i] = -xd[i];

  return;
}

static void VScaleSum_Serial(realtype c, N_Vector x, N_Vector y, N_Vector z)
{
  long int i, N;
  realtype *xd, *yd, *zd;

  xd = yd = zd = NULL;

  N  = NV_LENGTH_S(x);
  xd = NV_DATA_S(x);
  yd = NV_DATA_S(y);
  zd = NV_DATA_S(z);

  for (i = 0; i < N; i++)
    zd[i] = c*(xd[i]+yd[i]);

  return;
}

static void VScaleDiff_Serial(realtype c, N_Vector x, N_Vector y, N_Vector z)
{
  long int i, N;
  realtype *xd, *yd, *zd;

  xd = yd = zd = NULL;

  N  = NV_LENGTH_S(x);
  xd = NV_DATA_S(x);
  yd = NV_DATA_S(y);
  zd = NV_DATA_S(z);

  for (i = 0; i < N; i++)
    zd[i] = c*(xd[i]-yd[i]);

  return;
}

static void VLin1_Serial(realtype a, N_Vector x, N_Vector y, N_Vector z)
{
  long int i, N;
  realtype *xd, *yd, *zd;

  xd = yd = zd = NULL;

  N  = NV_LENGTH_S(x);
  xd = NV_DATA_S(x);
  yd = NV_DATA_S(y);
  zd = NV_DATA_S(z);

  for (i = 0; i < N; i++)
    zd[i] = (a*xd[i])+yd[i];

  return;
}

static void VLin2_Serial(realtype a, N_Vector x, N_Vector y, N_Vector z)
{
  long int i, N;
  realtype *xd, *yd, *zd;

  xd = yd = zd = NULL;

  N  = NV_LENGTH_S(x);
  xd = NV_DATA_S(x);
  yd = NV_DATA_S(y);
  zd = NV_DATA_S(z);

  for (i = 0; i < N; i++)
    zd[i] = (a*xd[i])-yd[i];

  return;
}

static void Vaxpy_Serial(realtype a, N_Vector x, N_Vector y)
{
  long int i, N;
  realtype *xd, *yd;

  xd = yd = NULL;

  N  = NV_LENGTH_S(x);
  xd = NV_DATA_S(x);
  yd = NV_DATA_S(y);

  if (a == ONE) {
    for (i = 0; i < N; i++)
      yd[i] += xd[i];
    return;
  }

  if (a == -ONE) {
    for (i = 0; i < N; i++)
      yd[i] -= xd[i];
    return;
  }    

  for (i = 0; i < N; i++)
    yd[i] += a*xd[i];

  return;
}

static void VScaleBy_Serial(realtype a, N_Vector x)
{
  long int i, N;
  realtype *xd;

  xd = NULL;

  N  = NV_LENGTH_S(x);
  xd = NV_DATA_S(x);

  for (i = 0; i < N; i++)
    xd[i] *= a;

  return;
}
```
