## Supplementary scripts for "A multi-scale *in silico* mouse model for insulin resistance and humanoid type 2 diabetes": sundials_band.c

```
/*
 * -----------------------------------------------------------------
 * $Revision: 749 $
 * $Date: 2011-06-22 14:49:10 +0200 (Wed, 22 Jun 2011) $
 * -----------------------------------------------------------------
 * Programmer(s): Alan C. Hindmarsh and Radu Serban @ LLNL
 * -----------------------------------------------------------------
 * Copyright (c) 2002, The Regents of the University of California.
 * Produced at the Lawrence Livermore National Laboratory.
 * All rights reserved.
 * For details, see the LICENSE file.
 * -----------------------------------------------------------------
 * This is the implementation file for a generic BAND linear
 * solver package.
 * -----------------------------------------------------------------
 */

#include <stdio.h>
#include <stdlib.h>

#include <sundials/sundials_band.h>
#include <sundials/sundials_math.h>

#define ZERO RCONST(0.0)
#define ONE  RCONST(1.0)

#define ROW(i,j,smu) (i-j+smu)

/* Implementation */

BandMat BandAllocMat(long int N, long int mu, long int ml, long int smu)
{
  BandMat A;

  if (N <= 0) return(NULL);
  
  A = NULL;
  A = (BandMat) malloc(sizeof *A);
  if (A == NULL) return (NULL);

  A->data = NULL;
  A->data = bandalloc(N, smu, ml);
  if (A->data == NULL) {
    free(A); A = NULL;
    return(NULL);
  }
  
  A->size = N;
  A->mu = mu;
  A->ml = ml;
  A->smu = smu;

  return(A);
}

long int *BandAllocPiv(long int N)
{
  return(bandallocpiv(N));
}

long int BandGBTRF(BandMat A, long int *p)
{
  return(bandGBTRF(A->data, A->size, A->mu, A->ml, A->smu, p));
}

void BandGBTRS(BandMat A, long int *p, realtype *b)
{
  bandGBTRS(A->data, A->size, A->smu, A->ml, p, b);
}

void BandZero(BandMat A)
{
  bandzero(A->data, A->size, A->mu, A->ml, A->smu);
}

void BandCopy(BandMat A, BandMat B, long int copymu, 
              long int copyml)
{
  bandcopy(A->data, B->data, A->size, A->smu, B->smu, copymu, copyml);
}

void BandScale(realtype c, BandMat A)
{
  bandscale(c, A->data, A->size, A->mu, A->ml, A->smu);
}

void BandAddI(BandMat A)
{
  bandaddI(A->data, A->size, A->smu);
}

void BandFreeMat(BandMat A)
{
  bandfree(A->data);
  free(A); A = NULL;
}

void BandFreePiv(long int *p)
{ 
  bandfreepiv(p);
}

void BandPrint(BandMat A)
{
  bandprint(A->data, A->size, A->mu, A->ml, A->smu);
}

realtype **bandalloc(long int n, long int smu, long int ml)
{
  realtype **a;
  long int j, colSize;

  if (n <= 0) return(NULL);

  a = NULL;
  a = (realtype **) malloc(n * sizeof(realtype *));
  if (a == NULL) return(NULL);

  colSize = smu + ml + 1;
  a[0] = NULL;
  a[0] = (realtype *) malloc(n * colSize * sizeof(realtype));
  if (a[0] == NULL) {
    free(a); a = NULL;
    return(NULL);
  }

  for (j=1; j < n; j++) a[j] = a[0] + j * colSize;

  return(a);
}

long int *bandallocpiv(long int n)
{
  long int *piv;

  if (n <= 0) return(NULL);

  piv = NULL;
  piv = (long int *) malloc(n * sizeof(long int));
  if (piv == NULL) return(NULL);

  return(piv);
}

long int bandGBTRF(realtype **a, long int n, long int mu, long int ml, 
                   long int smu, long int *p)
{
  long int c, r, num_rows;
  long int i, j, k, l, storage_l, storage_k, last_col_k, last_row_k;
  realtype *a_c, *col_k, *diag_k, *sub_diag_k, *col_j, *kptr, *jptr;
  realtype max, temp, mult, a_kj;
  booleantype swap;

  /* zero out the first smu - mu rows of the rectangular array a */

  num_rows = smu - mu;
  if (num_rows > 0) {
    for (c=0; c < n; c++) {
      a_c = a[c];
      for (r=0; r < num_rows; r++) {
	a_c[r] = ZERO;
      }
    }
  }

  /* k = elimination step number */

  for (k=0; k < n-1; k++, p++) {
    
    col_k     = a[k];
    diag_k    = col_k + smu;
    sub_diag_k = diag_k + 1;
    last_row_k = MIN(n-1,k+ml);

    /* find l = pivot row number */

    l=k;
    max = ABS(*diag_k);
    for (i=k+1, kptr=sub_diag_k; i <= last_row_k; i++, kptr++) { 
      if (ABS(*kptr) > max) {
	l=i;
	max = ABS(*kptr);
      }
    }
    storage_l = ROW(l, k, smu);
    *p = l;
    
    /* check for zero pivot element */

    if (col_k[storage_l] == ZERO) return(k+1);
    
    /* swap a(l,k) and a(k,k) if necessary */
    
    if ( (swap = (l != k) )) {
      temp = col_k[storage_l];
      col_k[storage_l] = *diag_k;
      *diag_k = temp;
    }

    /* Scale the elements below the diagonal in         */
    /* column k by -1.0 / a(k,k). After the above swap, */
    /* a(k,k) holds the pivot element. This scaling     */
    /* stores the pivot row multipliers -a(i,k)/a(k,k)  */
    /* in a(i,k), i=k+1, ..., MIN(n-1,k+ml).            */
    
    mult = -ONE / (*diag_k);
    for (i=k+1, kptr = sub_diag_k; i <= last_row_k; i++, kptr++)
      (*kptr) *= mult;

    /* row_i = row_i - [a(i,k)/a(k,k)] row_k, i=k+1, ..., MIN(n-1,k+ml) */
    /* row k is the pivot row after swapping with row l.                */
    /* The computation is done one column at a time,                    */
    /* column j=k+1, ..., MIN(k+smu,n-1).                               */
    
    last_col_k = MIN(k+smu,n-1);
    for (j=k+1; j <= last_col_k; j++) {
      
      col_j = a[j];
      storage_l = ROW(l,j,smu); 
      storage_k = ROW(k,j,smu); 
      a_kj = col_j[storage_l];

      /* Swap the elements a(k,j) and a(k,l) if l!=k. */
      
      if (swap) {
	col_j[storage_l] = col_j[storage_k];
	col_j[storage_k] = a_kj;
      }

      /* a(i,j) = a(i,j) - [a(i,k)/a(k,k)]*a(k,j) */
      /* a_kj = a(k,j), *kptr = - a(i,k)/a(k,k), *jptr = a(i,j) */

      if (a_kj != ZERO) {
	for (i=k+1, kptr=sub_diag_k, jptr=col_j+ROW(k+1,j,smu);
	     i <= last_row_k;
	     i++, kptr++, jptr++)
	  (*jptr) += a_kj * (*kptr);
      }
    }    
  }
  
  /* set the last pivot row to be n-1 and check for a zero pivot */

  *p = n-1; 
  if (a[n-1][smu] == ZERO) return(n);

  /* return 0 to indicate success */

  return(0);
}

void bandGBTRS(realtype **a, long int n, long int smu, long int ml, 
               long int *p, realtype *b)
{
  long int k, l, i, first_row_k, last_row_k;
  realtype mult, *diag_k;
  
  /* Solve Ly = Pb, store solution y in b */
  
  for (k=0; k < n-1; k++) {
    l = p[k];
    mult = b[l];
    if (l != k) {
      b[l] = b[k];
      b[k] = mult;
    }
    diag_k = a[k]+smu;
    last_row_k = MIN(n-1,k+ml);
    for (i=k+1; i <= last_row_k; i++)
      b[i] += mult * diag_k[i-k];
  }
  
  /* Solve Ux = y, store solution x in b */
  
  for (k=n-1; k >= 0; k--) {
    diag_k = a[k]+smu;
    first_row_k = MAX(0,k-smu);
    b[k] /= (*diag_k);
    mult = -b[k];
    for (i=first_row_k; i <= k-1; i++)
      b[i] += mult*diag_k[i-k];
  }
}

void bandzero(realtype **a, long int n, long int mu, long int ml, 
              long int smu)
{
  long int i, j, colSize;
  realtype *col_j;

  colSize = mu + ml + 1;
  for (j=0; j < n; j++) {
    col_j = a[j]+smu-mu;
    for (i=0; i < colSize; i++)
      col_j[i] = ZERO;
  }
}

void bandcopy(realtype **a, realtype **b, long int n, long int a_smu, 
              long int b_smu, long int copymu, long int copyml)
{
  long int i, j, copySize;
  realtype *a_col_j, *b_col_j;

  copySize = copymu + copyml + 1;
 
  for (j=0; j < n; j++) {
    a_col_j = a[j]+a_smu-copymu;
    b_col_j = b[j]+b_smu-copymu;
    for (i=0; i < copySize; i++)
      b_col_j[i] = a_col_j[i];
  }
}

void bandscale(realtype c, realtype **a, long int n, long int mu, 
               long int ml, long int smu)
{
  long int i, j, colSize;
  realtype *col_j;

  colSize = mu + ml + 1;

  for(j=0; j < n; j++) {
    col_j = a[j]+smu-mu;
    for (i=0; i < colSize; i++)
      col_j[i] *= c;
  }
}

void bandaddI(realtype **a, long int n, long int smu)
{
  long int j;
 
  for(j=0; j < n; j++)
    a[j][smu] += ONE;
}

void bandfreepiv(long int *p)
{
  free(p); p = NULL;
}

void bandfree(realtype **a)
{
  free(a[0]); a[0] = NULL;
  free(a); a = NULL;
}

void bandprint(realtype **a, long int n, long int mu, long int ml, 
               long int smu)
{
  long int i, j, start, finish;
 
  printf("\n");
  for (i=0; i < n; i++) {
    start = MAX(0,i-ml);
    finish = MIN(n-1,i+mu);
    for (j=0; j < start; j++) printf("%12s  ","");
    for (j=start; j <= finish; j++) {
#if defined(SUNDIALS_EXTENDED_PRECISION)
      printf("%12Lg  ", a[j][i-j+smu]);
#elif defined(SUNDIALS_DOUBLE_PRECISION)
      printf("%12lg  ", a[j][i-j+smu]);
#else
      printf("%12g  ", a[j][i-j+smu]);
#endif
    }
    printf("\n");
  }
  printf("\n");
}
```
