## Supplementary scripts for "A multi-scale *in silico* mouse model for insulin resistance and humanoid type 2 diabetes": sundials_dense.c

```
/*
 * -----------------------------------------------------------------
 * $Revision: 749 $
 * $Date: 2011-06-22 14:49:10 +0200 (Wed, 22 Jun 2011) $
 * -----------------------------------------------------------------
 * Programmer(s): Scott D. Cohen, Alan C. Hindmarsh and
 *                Radu Serban @ LLNL
 * -----------------------------------------------------------------
 * Copyright (c) 2002, The Regents of the University of California.
 * Produced at the Lawrence Livermore National Laboratory.
 * All rights reserved.
 * For details, see the LICENSE file.
 * -----------------------------------------------------------------
 * This is the implementation file for a generic package of dense
 * matrix operations.
 * -----------------------------------------------------------------
 */ 

#include <stdio.h>
#include <stdlib.h>

#include <sundials/sundials_dense.h>
#include <sundials/sundials_math.h>

#define ZERO RCONST(0.0)
#define ONE  RCONST(1.0)

/* Implementation */

DenseMat DenseAllocMat(long int M, long int N)
{
  DenseMat A;

  /* Note that M and N are tested in denalloc */

  A = NULL;
  A = (DenseMat) malloc(sizeof *A);
  if (A==NULL) return (NULL);
  
  A->data = NULL;
  A->data = denalloc(M, N);
  if (A->data == NULL) {
    free(A); A = NULL;
    return(NULL);
  }

  A->M = M;
  A->N = N;

  return(A);
}

long int *DenseAllocPiv(long int N)
{
  return(denallocpiv(N));
}

long int DenseGETRF(DenseMat A, long int *p)
{
  return(denGETRF(A->data, A->M, A->N, p));
}

void DenseGETRS(DenseMat A, long int *p, realtype *b)
{
  denGETRS(A->data, A->N, p, b);
}

void DenseZero(DenseMat A)
{
  denzero(A->data, A->M, A->N);
}

void DenseCopy(DenseMat A, DenseMat B)
{
  dencopy(A->data, B->data, A->M, A->N);
}

void DenseScale(realtype c, DenseMat A)
{
  denscale(c, A->data, A->M, A->N);
}

void DenseAddI(DenseMat A)
{
  denaddI(A->data, A->N);
}

void DenseFreeMat(DenseMat A)
{
  denfree(A->data);
  free(A); A = NULL;
}

void DenseFreePiv(long int *p)
{  
  denfreepiv(p);
}

void DensePrint(DenseMat A)
{
  denprint(A->data, A->M, A->N);
}
```
