## Supplementary scripts for "A multi-scale *in silico* mouse model for insulin resistance and humanoid type 2 diabetes": sundials_iterative.c

```
/*
 * -----------------------------------------------------------------
 * $Revision: 749 $
 * $Date: 2011-06-22 14:49:10 +0200 (Wed, 22 Jun 2011) $
 * ----------------------------------------------------------------- 
 * Programmer(s): Scott D. Cohen, Alan C. Hindmarsh and
 *                Radu Serban @ LLNL
 * -----------------------------------------------------------------
 * Copyright (c) 2002, The Regents of the University of California.
 * Produced at the Lawrence Livermore National Laboratory.
 * All rights reserved.
 * For details, see the LICENSE file.
 * -----------------------------------------------------------------
 * This is the implementation file for the iterative.h header
 * file. It contains the implementation of functions that may be
 * useful for many different iterative solvers of A x = b.
 * -----------------------------------------------------------------
 */

#include <stdio.h>

#include <sundials/sundials_iterative.h>
#include <sundials/sundials_math.h>

#define FACTOR RCONST(1000.0)
#define ZERO   RCONST(0.0)
#define ONE    RCONST(1.0)

/*
 * -----------------------------------------------------------------
 * Function : ModifiedGS
 * -----------------------------------------------------------------
 * This implementation of ModifiedGS is a slight modification of a
 * previous modified Gram-Schmidt routine (called mgs) written by
 * Milo Dorr.
 * -----------------------------------------------------------------
 */
 
int ModifiedGS(N_Vector *v, realtype **h, int k, int p, 
               realtype *new_vk_norm)
{
  int  i, k_minus_1, i0;
  realtype new_norm_2, new_product, vk_norm, temp;
  
  vk_norm = RSqrt(N_VDotProd(v[k],v[k]));
  k_minus_1 = k - 1;
  i0 = MAX(k-p, 0);
  
  /* Perform modified Gram-Schmidt */
  
  for (i=i0; i < k; i++) {
    h[i][k_minus_1] = N_VDotProd(v[i], v[k]);
    N_VLinearSum(ONE, v[k], -h[i][k_minus_1], v[i], v[k]);
  }

  /* Compute the norm of the new vector at v[k] */

  *new_vk_norm = RSqrt(N_VDotProd(v[k], v[k]));

  /* If the norm of the new vector at v[k] is less than
     FACTOR (== 1000) times unit roundoff times the norm of the
     input vector v[k], then the vector will be reorthogonalized
     in order to ensure that nonorthogonality is not being masked
     by a very small vector length. */

  temp = FACTOR * vk_norm;
  if ((temp + (*new_vk_norm)) != temp) return(0);
  
  new_norm_2 = ZERO;

  for (i=i0; i < k; i++) {
    new_product = N_VDotProd(v[i], v[k]);
    temp = FACTOR * h[i][k_minus_1];
    if ((temp + new_product) == temp) continue;
    h[i][k_minus_1] += new_product;
    N_VLinearSum(ONE, v[k],-new_product, v[i], v[k]);
    new_norm_2 += SQR(new_product);
  }

  if (new_norm_2 != ZERO) {
    new_product = SQR(*new_vk_norm) - new_norm_2;
    *new_vk_norm = (new_product > ZERO) ? RSqrt(new_product) : ZERO;
  }

  return(0);
}

/*
 * -----------------------------------------------------------------
 * Function : ClassicalGS
 * -----------------------------------------------------------------
 * This implementation of ClassicalGS was contributed by Homer Walker
 * and Peter Brown.
 * -----------------------------------------------------------------
 */

int ClassicalGS(N_Vector *v, realtype **h, int k, int p, 
                realtype *new_vk_norm, N_Vector temp, realtype *s)
{
  int  i, k_minus_1, i0;
  realtype vk_norm;

  k_minus_1 = k - 1;
  
  /* Perform Classical Gram-Schmidt */

  vk_norm = RSqrt(N_VDotProd(v[k], v[k]));

  i0 = MAX(k-p, 0);
  for (i=i0; i < k; i++) {
    h[i][k_minus_1] = N_VDotProd(v[i], v[k]);
  }

  for (i=i0; i < k; i++) {
    N_VLinearSum(ONE, v[k], -h[i][k_minus_1], v[i], v[k]);
  }

  /* Compute the norm of the new vector at v[k] */

  *new_vk_norm = RSqrt(N_VDotProd(v[k], v[k]));

  /* Reorthogonalize if necessary */

  if ((FACTOR * (*new_vk_norm)) < vk_norm) {

    for (i=i0; i < k; i++) {
      s[i] = N_VDotProd(v[i], v[k]);
    }

    if (i0 < k) {
      N_VScale(s[i0], v[i0], temp);
      h[i0][k_minus_1] += s[i0];
    }
    for (i=i0+1; i < k; i++) {
      N_VLinearSum(s[i], v[i], ONE, temp, temp);
      h[i][k_minus_1] += s[i];
    }
    N_VLinearSum(ONE, v[k], -ONE, temp, v[k]);

    *new_vk_norm = RSqrt(N_VDotProd(v[k],v[k]));
  }

  return(0);
}

/*
 * -----------------------------------------------------------------
 * Function : QRfact
 * -----------------------------------------------------------------
 * This implementation of QRfact is a slight modification of a
 * previous routine (called qrfact) written by Milo Dorr.
 * -----------------------------------------------------------------
 */

int QRfact(int n, realtype **h, realtype *q, int job)
{
  realtype c, s, temp1, temp2, temp3;
  int i, j, k, q_ptr, n_minus_1, code=0;

  switch (job) {
  case 0:

    /* Compute a new factorization of H */

    code = 0;
    for (k=0; k < n; k++) {
      
      /* Multiply column k by the previous k-1 Givens rotations */

      for (j=0; j < k-1; j++) {
	i = 2*j;
	temp1 = h[j][k];
	temp2 = h[j+1][k];
	c = q[i];
	s = q[i+1];
	h[j][k] = c*temp1 - s*temp2;
	h[j+1][k] = s*temp1 + c*temp2;
      }
      
      /* Compute the Givens rotation components c and s */

      q_ptr = 2*k;
      temp1 = h[k][k];
      temp2 = h[k+1][k];
      if( temp2 == ZERO) {
	c = ONE;
	s = ZERO;
      } else if (ABS(temp2) >= ABS(temp1)) {
	temp3 = temp1/temp2;
	s = -ONE/RSqrt(ONE+SQR(temp3));
	c = -s*temp3;
      } else {
	temp3 = temp2/temp1;
	c = ONE/RSqrt(ONE+SQR(temp3));
	s = -c*temp3;
      }
      q[q_ptr] = c;
      q[q_ptr+1] = s;
      if( (h[k][k] = c*temp1 - s*temp2) == ZERO) code = k+1;
    }
    break;

  default:

    /* Update the factored H to which a new column has been added */

    n_minus_1 = n - 1;
    code = 0;
    
    /* Multiply the new column by the previous n-1 Givens rotations */

    for (k=0; k < n_minus_1; k++) {
      i = 2*k;
      temp1 = h[k][n_minus_1];
      temp2 = h[k+1][n_minus_1];
      c = q[i];
      s = q[i+1];
      h[k][n_minus_1] = c*temp1 - s*temp2;
      h[k+1][n_minus_1] = s*temp1 + c*temp2;
    }
    
    /* Compute new Givens rotation and multiply it times the last two
       entries in the new column of H.  Note that the second entry of 
       this product will be 0, so it is not necessary to compute it. */

    temp1 = h[n_minus_1][n_minus_1];
    temp2 = h[n][n_minus_1];
    if (temp2 == ZERO) {
      c = ONE;
      s = ZERO;
    } else if (ABS(temp2) >= ABS(temp1)) {
      temp3 = temp1/temp2;
      s = -ONE/RSqrt(ONE+SQR(temp3));
      c = -s*temp3;
    } else {
      temp3 = temp2/temp1;
      c = ONE/RSqrt(ONE+SQR(temp3));
      s = -c*temp3;
    }
    q_ptr = 2*n_minus_1;
    q[q_ptr] = c;
    q[q_ptr+1] = s;
    if ((h[n_minus_1][n_minus_1] = c*temp1 - s*temp2) == ZERO)
      code = n;
  }
  
  return (code);
}

/*
 * -----------------------------------------------------------------
 * Function : QRsol
 * -----------------------------------------------------------------
 * This implementation of QRsol is a slight modification of a
 * previous routine (called qrsol) written by Milo Dorr.
 * -----------------------------------------------------------------
 */

int QRsol(int n, realtype **h, realtype *q, realtype *b)
{
  realtype c, s, temp1, temp2;
  int i, k, q_ptr, code=0;

  /* Compute Q*b */
  
  for (k=0; k < n; k++) {
    q_ptr = 2*k;
    c = q[q_ptr];
    s = q[q_ptr+1];
    temp1 = b[k];
    temp2 = b[k+1];
    b[k] = c*temp1 - s*temp2;
    b[k+1] = s*temp1 + c*temp2;
  }

  /* Solve  R*x = Q*b */

  for (k=n-1; k >= 0; k--) {
    if (h[k][k] == ZERO) {
      code = k + 1;
      break;
    }
    b[k] /= h[k][k];
    for (i=0; i < k; i++) b[i] -= b[k]*h[i][k];
  }
  
  return (code);
}
```
