## Supplementary scripts for "A multi-scale *in silico* mouse model for insulin resistance and humanoid type 2 diabetes": sundials_math.c

```
/*
 * -----------------------------------------------------------------
 * $Revision: 749 $
 * $Date: 2011-06-22 14:49:10 +0200 (Wed, 22 Jun 2011) $
 * -----------------------------------------------------------------
 * Programmer(s): Scott D. Cohen, Alan C. Hindmarsh and
 *                Aaron Collier @ LLNL
 * -----------------------------------------------------------------
 * Copyright (c) 2002, The Regents of the University of California.
 * Produced at the Lawrence Livermore National Laboratory.
 * All rights reserved.
 * For details, see the LICENSE file.
 * -----------------------------------------------------------------
 * This is the implementation file for a simple C-language math
 * library.
 * -----------------------------------------------------------------
 */

#include <stdio.h>
#include <stdlib.h>
#include <math.h>

#include <sundials/sundials_math.h>

#define ZERO RCONST(0.0)
#define ONE  RCONST(1.0)

realtype RPowerI(realtype base, int exponent)
{
  int i, expt;
  realtype prod;

  prod = ONE;
  expt = abs(exponent);
  for(i = 1; i <= expt; i++) prod *= base;
  if (exponent < 0) prod = ONE/prod;
  return(prod);
}

realtype RPowerR(realtype base, realtype exponent)
{
  if (base <= ZERO) return(ZERO);

#if defined(SUNDIALS_USE_GENERIC_MATH)
  return((realtype) pow((double) base, (double) exponent));
#elif defined(SUNDIALS_DOUBLE_PRECISION)
  return(pow(base, exponent));
#elif defined(SUNDIALS_SINGLE_PRECISION)
  return(powf(base, exponent));
#elif defined(SUNDIALS_EXTENDED_PRECISION)
  return(powl(base, exponent));
#endif
}

realtype RSqrt(realtype x)
{
  if (x <= ZERO) return(ZERO);

#if defined(SUNDIALS_USE_GENERIC_MATH)
  return((realtype) sqrt((double) x));
#elif defined(SUNDIALS_DOUBLE_PRECISION)
  return(sqrt(x));
#elif defined(SUNDIALS_SINGLE_PRECISION)
  return(sqrtf(x));
#elif defined(SUNDIALS_EXTENDED_PRECISION)
  return(sqrtl(x));
#endif
}

realtype RAbs(realtype x)
{
#if defined(SUNDIALS_USE_GENERIC_MATH)
  return((realtype) fabs((double) x));
#elif defined(SUNDIALS_DOUBLE_PRECISION)
  return(fabs(x));
#elif defined(SUNDIALS_SINGLE_PRECISION)
  return(fabsf(x));
#elif defined(SUNDIALS_EXTENDED_PRECISION)
  return(fabsl(x));
#endif
}

realtype RExp(realtype x)
{
#if defined(SUNDIALS_USE_GENERIC_MATH)
  return((realtype) exp((double) x));
#elif defined(SUNDIALS_DOUBLE_PRECISION)
  return(exp(x));
#elif defined(SUNDIALS_SINGLE_PRECISION)
  return(expf(x));
#elif defined(SUNDIALS_EXTENDED_PRECISION)
  return(expl(x));
#endif
}
```
