## Supplementary scripts for "A multi-scale *in silico* mouse model for insulin resistance and humanoid type 2 diabetes": sundials_nvector.c

```
/*
 * -----------------------------------------------------------------
 * $Revision: 749 $
 * $Date: 2011-06-22 14:49:10 +0200 (Wed, 22 Jun 2011) $
 * ----------------------------------------------------------------- 
 * Programmer(s): Radu Serban and Aaron Collier @ LLNL                               
 * -----------------------------------------------------------------
 * Copyright (c) 2002, The Regents of the University of California.
 * Produced at the Lawrence Livermore National Laboratory.
 * All rights reserved.
 * For details, see the LICENSE file.
 * -----------------------------------------------------------------
 * This is the implementation file for a generic NVECTOR package.
 * It contains the implementation of the N_Vector operations listed
 * in nvector.h.
 * -----------------------------------------------------------------
 */

#include <stdlib.h>

#include <sundials/sundials_nvector.h>

/*
 * -----------------------------------------------------------------
 * Functions in the 'ops' structure
 * -----------------------------------------------------------------
 */

N_Vector N_VClone(N_Vector w)
{
  return(w->ops->nvclone(w));
}

N_Vector N_VCloneEmpty(N_Vector w)
{
  return(w->ops->nvcloneempty(w));
}

void N_VDestroy(N_Vector v)
{
  v->ops->nvdestroy(v);
  return;
}

void N_VSpace(N_Vector v, long int *lrw, long int *liw)
{
  v->ops->nvspace(v, lrw, liw);
  return;
}

realtype *N_VGetArrayPointer(N_Vector v)
{
  return((realtype *) v->ops->nvgetarraypointer(v));
}

void N_VSetArrayPointer(realtype *v_data, N_Vector v)
{
  v->ops->nvsetarraypointer(v_data, v);
  return;
}

void N_VLinearSum(realtype a, N_Vector x, realtype b, N_Vector y, N_Vector z)
{
  z->ops->nvlinearsum(a, x, b, y, z);
  return;
}

void N_VConst(realtype c, N_Vector z)
{
  z->ops->nvconst(c, z);
  return;
}

void N_VProd(N_Vector x, N_Vector y, N_Vector z)
{
  z->ops->nvprod(x, y, z);
  return;
}

void N_VDiv(N_Vector x, N_Vector y, N_Vector z)
{
  z->ops->nvdiv(x, y, z);
  return;
}

void N_VScale(realtype c, N_Vector x, N_Vector z) 
{
  z->ops->nvscale(c, x, z);
  return;
}

void N_VAbs(N_Vector x, N_Vector z)
{
  z->ops->nvabs(x, z);
  return;
}

void N_VInv(N_Vector x, N_Vector z)
{
  z->ops->nvinv(x, z);
  return;
}

void N_VAddConst(N_Vector x, realtype b, N_Vector z)
{
  z->ops->nvaddconst(x, b, z);
  return;
}

realtype N_VDotProd(N_Vector x, N_Vector y)
{
  return((realtype) y->ops->nvdotprod(x, y));
}

realtype N_VMaxNorm(N_Vector x)
{
  return((realtype) x->ops->nvmaxnorm(x));
}

realtype N_VWrmsNorm(N_Vector x, N_Vector w)
{
  return((realtype) x->ops->nvwrmsnorm(x, w));
}

realtype N_VWrmsNormMask(N_Vector x, N_Vector w, N_Vector id)
{
  return((realtype) x->ops->nvwrmsnormmask(x, w, id));
}

realtype N_VMin(N_Vector x)
{
  return((realtype) x->ops->nvmin(x));
}

realtype N_VWL2Norm(N_Vector x, N_Vector w)
{
  return((realtype) x->ops->nvwl2norm(x, w));
}

realtype N_VL1Norm(N_Vector x)
{
  return((realtype) x->ops->nvl1norm(x));
}

void N_VCompare(realtype c, N_Vector x, N_Vector z)
{
  z->ops->nvcompare(c, x, z);
  return;
}

booleantype N_VInvTest(N_Vector x, N_Vector z)
{
  return((booleantype) z->ops->nvinvtest(x, z));
}

booleantype N_VConstrMask(N_Vector c, N_Vector x, N_Vector m)
{
  return((booleantype) x->ops->nvconstrmask(c, x, m));
}

realtype N_VMinQuotient(N_Vector num, N_Vector denom)
{
  return((realtype) num->ops->nvminquotient(num, denom));
}

/*
 * -----------------------------------------------------------------
 * Additional functions exported by the generic NVECTOR:
 *   N_VCloneEmptyVectorArray
 *   N_VCloneVectorArray
 *   N_VDestroyVectorArray
 * -----------------------------------------------------------------
 */

N_Vector *N_VCloneEmptyVectorArray(int count, N_Vector w)
{
  N_Vector *vs;
  int j;

  if (count <= 0) return(NULL);

  vs = NULL;
  vs = (N_Vector *) malloc(count * sizeof(N_Vector));
  if(vs == NULL) return(NULL);

  for (j = 0; j < count; j++) {
    vs[j] = NULL;
    vs[j] = N_VCloneEmpty(w);
    if (vs[j] == NULL) {
      N_VDestroyVectorArray(vs, j-1);
      return(NULL);
    }
  }

  return(vs);
}

N_Vector *N_VCloneVectorArray(int count, N_Vector w)
{
  N_Vector *vs;
  int j;

  if (count <= 0) return(NULL);

  vs = NULL;
  vs = (N_Vector *) malloc(count * sizeof(N_Vector));
  if(vs == NULL) return(NULL);

  for (j = 0; j < count; j++) {
    vs[j] = NULL;
    vs[j] = N_VClone(w);
    if (vs[j] == NULL) {
      N_VDestroyVectorArray(vs, j-1);
      return(NULL);
    }
  }

  return(vs);
}

void N_VDestroyVectorArray(N_Vector *vs, int count)
{
  int j;

  for (j = 0; j < count; j++) N_VDestroy(vs[j]);

  free(vs); vs = NULL;

  return;
}
```
