## Supplementary scripts for "A multi-scale *in silico* mouse model for insulin resistance and humanoid type 2 diabetes": sundials_smalldense.c

```
/*
 * -----------------------------------------------------------------
 * $Revision: 749 $
 * $Date: 2011-06-22 14:49:10 +0200 (Wed, 22 Jun 2011) $
 * -----------------------------------------------------------------
 * Programmer: Radu Serban @ LLNL
 * -----------------------------------------------------------------
 * Copyright (c) 2006, The Regents of the University of California.
 * Produced at the Lawrence Livermore National Laboratory.
 * All rights reserved.
 * For details, see the LICENSE file.
 * -----------------------------------------------------------------
 * This is the implementation file for a generic DENSE linear
 * solver package, intended for small dense matrices.
 * -----------------------------------------------------------------
 */

#include <stdio.h>
#include <stdlib.h>

#include <sundials/sundials_smalldense.h>
#include <sundials/sundials_math.h>

#define ZERO RCONST(0.0)
#define ONE  RCONST(1.0)

/* Implementation */


realtype **denalloc(long int m, long int n)
{
  long int j;
  realtype **a;

  if ( (n <= 0) || (m <= 0) ) return(NULL);

  a = NULL;
  a = (realtype **) malloc(n * sizeof(realtype *));
  if (a == NULL) return(NULL);

  a[0] = NULL;
  a[0] = (realtype *) malloc(m * n * sizeof(realtype));
  if (a[0] == NULL) {
    free(a); a = NULL;
    return(NULL);
  }

  for (j=1; j < n; j++) a[j] = a[0] + j * m;

  return(a);
}

long int *denallocpiv(long int n)
{
  long int *piv;

  if (n <= 0) return(NULL);

  piv = NULL;
  piv = (long int *) malloc(n * sizeof(long int));

  return(piv);
}

long int denGETRF(realtype **a, long int m, long int n, long int *p)
{
  long int i, j, k, l;
  realtype *col_j, *col_k;
  realtype temp, mult, a_kj;

  /* k-th elimination step number */
  for (k=0; k < n; k++) {

    col_k  = a[k];

    /* find l = pivot row number */
    l=k;
    for (i=k+1; i < m; i++)
      if (ABS(col_k[i]) > ABS(col_k[l])) l=i;
    p[k] = l;

    /* check for zero pivot element */
    if (col_k[l] == ZERO) return(k+1);
    
    /* swap a(k,1:n) and a(l,1:n) if necessary */    
    if ( l!= k ) {
      for (i=0; i<n; i++) {
        temp = a[i][l];
        a[i][l] = a[i][k];
        a[i][k] = temp;
      }
    }

    /* Scale the elements below the diagonal in
     * column k by 1.0/a(k,k). After the above swap
     * a(k,k) holds the pivot element. This scaling
     * stores the pivot row multipliers a(i,k)/a(k,k)
     * in a(i,k), i=k+1, ..., m-1.                      
     */
    mult = ONE/col_k[k];
    for(i=k+1; i < m; i++) col_k[i] *= mult;

    /* row_i = row_i - [a(i,k)/a(k,k)] row_k, i=k+1, ..., m-1 */
    /* row k is the pivot row after swapping with row l.      */
    /* The computation is done one column at a time,          */
    /* column j=k+1, ..., n-1.                                */

    for (j=k+1; j < n; j++) {

      col_j = a[j];
      a_kj = col_j[k];

      /* a(i,j) = a(i,j) - [a(i,k)/a(k,k)]*a(k,j)  */
      /* a_kj = a(k,j), col_k[i] = - a(i,k)/a(k,k) */

      if (a_kj != ZERO) {
	for (i=k+1; i < m; i++)
	  col_j[i] -= a_kj * col_k[i];
      }
    }
  }

  /* return 0 to indicate success */

  return(0);
}

void denGETRS(realtype **a, long int n, long int *p, realtype *b)
{
  long int i, k, pk;
  realtype *col_k, tmp;

  /* Permute b, based on pivot information in p */
  for (k=0; k<n; k++) {
    pk = p[k];
    if(pk != k) {
      tmp = b[k];
      b[k] = b[pk];
      b[pk] = tmp;
    }
  }

  /* Solve Ly = b, store solution y in b */
  for (k=0; k<n-1; k++) {
    col_k = a[k];
    for (i=k+1; i<n; i++) b[i] -= col_k[i]*b[k];
  }

  /* Solve Ux = y, store solution x in b */
  for (k = n-1; k > 0; k--) {
    col_k = a[k];
    b[k] /= col_k[k];
    for (i=0; i<k; i++) b[i] -= col_k[i]*b[k];
  }
  b[0] /= a[0][0];

}

void denzero(realtype **a, long int m, long int n)
{
  long int i, j;
  realtype *col_j;

  for (j=0; j < n; j++) {
    col_j = a[j];
    for (i=0; i < m; i++)
      col_j[i] =  ZERO;
  }
}

void dencopy(realtype **a, realtype **b, long int m, long int n)
{
  long int i, j;
  realtype *a_col_j, *b_col_j;

  for (j=0; j < n; j++) {
    a_col_j = a[j];
    b_col_j = b[j];
    for (i=0; i < m; i++)
      b_col_j[i] = a_col_j[i];
  }

}

void denscale(realtype c, realtype **a, long int m, long int n)
{
  long int i, j;
  realtype *col_j;

  for (j=0; j < n; j++) {
    col_j = a[j];
    for (i=0; i < m; i++)
      col_j[i] *= c;
  }
}

void denaddI(realtype **a, long int n)
{
  long int i;
  
  for (i=0; i < n; i++) a[i][i] += ONE;
}

void denfreepiv(long int *p)
{
  free(p); p = NULL;
}

void denfree(realtype **a)
{
  free(a[0]); a[0] = NULL;
  free(a); a = NULL;
}

void denprint(realtype **a, long int m, long int n)
{
  long int i, j;

  printf("\n");
  for (i=0; i < m; i++) {
    for (j=0; j < n; j++) {
#if defined(SUNDIALS_EXTENDED_PRECISION)
      printf("%12Lg  ", a[j][i]);
#elif defined(SUNDIALS_DOUBLE_PRECISION)
      printf("%12lg  ", a[j][i]);
#else
      printf("%12g  ", a[j][i]);
#endif
    }
    printf("\n");
  }
  printf("\n");
}
```
