## Supplementary scripts for "A multi-scale *in silico* mouse model for insulin resistance and humanoid type 2 diabetes": sundials_spbcgs.c

```
/*
 * -----------------------------------------------------------------
 * $Revision: 749 $
 * $Date: 2011-06-22 14:49:10 +0200 (Wed, 22 Jun 2011) $
 * -----------------------------------------------------------------
 * Programmer(s): Peter Brown and Aaron Collier @ LLNL
 * -----------------------------------------------------------------
 * Copyright (c) 2004, The Regents of the University of California.
 * Produced at the Lawrence Livermore National Laboratory.
 * All rights reserved.
 * For details, see the LICENSE file.
 * -----------------------------------------------------------------
 * This is the implementation file for the scaled, preconditioned
 * Bi-CGSTAB (SPBCG) iterative linear solver.
 * -----------------------------------------------------------------
 */

#include <stdio.h>
#include <stdlib.h>

#include <sundials/sundials_spbcgs.h>
#include <sundials/sundials_math.h>

/*
 * -----------------------------------------------------------------
 * private constants
 * -----------------------------------------------------------------
 */

#define ZERO RCONST(0.0)
#define ONE  RCONST(1.0)

/*
 * -----------------------------------------------------------------
 * Function : SpbcgMalloc
 * -----------------------------------------------------------------
 */

SpbcgMem SpbcgMalloc(int l_max, N_Vector vec_tmpl)
{
  SpbcgMem mem;
  N_Vector r_star, r, p, q, u, Ap, vtemp;

  /* Check the input parameters */

  if (l_max <= 0) return(NULL);

  /* Get arrays to hold temporary vectors */

  r_star = NULL;
  r_star = N_VClone(vec_tmpl);
  if (r_star == NULL) {
    return(NULL);
  }

  r = NULL;
  r = N_VClone(vec_tmpl);
  if (r == NULL) {
    N_VDestroy(r_star);
    return(NULL);
  }

  p = NULL;
  p = N_VClone(vec_tmpl);
  if (p == NULL) {
    N_VDestroy(r_star);
    N_VDestroy(r);
    return(NULL);
  }

  q = NULL;
  q = N_VClone(vec_tmpl);
  if (q == NULL) {
    N_VDestroy(r_star);
    N_VDestroy(r);
    N_VDestroy(p);
    return(NULL);
  }

  u = NULL;
  u = N_VClone(vec_tmpl);
  if (u == NULL) {
    N_VDestroy(r_star);
    N_VDestroy(r);
    N_VDestroy(p);
    N_VDestroy(q);
    return(NULL);
  }

  Ap = NULL;
  Ap = N_VClone(vec_tmpl);
  if (Ap == NULL) {
    N_VDestroy(r_star);
    N_VDestroy(r);
    N_VDestroy(p);
    N_VDestroy(q);
    N_VDestroy(u);
    return(NULL);
  }

  vtemp = NULL;
  vtemp = N_VClone(vec_tmpl);
  if (vtemp == NULL) {
    N_VDestroy(r_star);
    N_VDestroy(r);
    N_VDestroy(p);
    N_VDestroy(q);
    N_VDestroy(u);
    N_VDestroy(Ap);
    return(NULL);
  }

  /* Get memory for an SpbcgMemRec containing SPBCG matrices and vectors */

  mem = NULL;
  mem = (SpbcgMem) malloc(sizeof(SpbcgMemRec));
  if (mem == NULL) {
    N_VDestroy(r_star);
    N_VDestroy(r);
    N_VDestroy(p);
    N_VDestroy(q);
    N_VDestroy(u);
    N_VDestroy(Ap);
    N_VDestroy(vtemp);
    return(NULL);
  }

  /* Set the fields of mem */

  mem->l_max  = l_max;
  mem->r_star = r_star;
  mem->r      = r;
  mem->p      = p;
  mem->q      = q;
  mem->u      = u;
  mem->Ap     = Ap;
  mem->vtemp  = vtemp;

  /* Return the pointer to SPBCG memory */

  return(mem);
}

/*
 * -----------------------------------------------------------------
 * Function : SpbcgSolve
 * -----------------------------------------------------------------
 */

int SpbcgSolve(SpbcgMem mem, void *A_data, N_Vector x, N_Vector b,
               int pretype, realtype delta, void *P_data, N_Vector sx,
               N_Vector sb, ATimesFn atimes, PSolveFn psolve,
               realtype *res_norm, int *nli, int *nps)
{
  realtype alpha, beta, omega, omega_denom, beta_num, beta_denom, r_norm, rho;
  N_Vector r_star, r, p, q, u, Ap, vtemp;
  booleantype preOnLeft, preOnRight, scale_x, scale_b, converged;
  int l, l_max, ier;

  if (mem == NULL) return(SPBCG_MEM_NULL);

  /* Make local copies of mem variables */

  l_max  = mem->l_max;
  r_star = mem->r_star;
  r      = mem->r;
  p      = mem->p;
  q      = mem->q;
  u      = mem->u;
  Ap     = mem->Ap;
  vtemp  = mem->vtemp;

  *nli = *nps = 0;    /* Initialize counters */
  converged = FALSE;  /* Initialize converged flag */

  if ((pretype != PREC_LEFT) && (pretype != PREC_RIGHT) && (pretype != PREC_BOTH)) pretype = PREC_NONE;

  preOnLeft  = ((pretype == PREC_BOTH) || (pretype == PREC_LEFT));
  preOnRight = ((pretype == PREC_BOTH) || (pretype == PREC_RIGHT));

  scale_x = (sx != NULL);
  scale_b = (sb != NULL);

  /* Set r_star to initial (unscaled) residual r_0 = b - A*x_0 */

  if (N_VDotProd(x, x) == ZERO) N_VScale(ONE, b, r_star);
  else {
    ier = atimes(A_data, x, r_star);
    if (ier != 0)
      return((ier < 0) ? SPBCG_ATIMES_FAIL_UNREC : SPBCG_ATIMES_FAIL_REC);
    N_VLinearSum(ONE, b, -ONE, r_star, r_star);
  }

  /* Apply left preconditioner and b-scaling to r_star = r_0 */

  if (preOnLeft) {
    ier = psolve(P_data, r_star, r, PREC_LEFT);
    (*nps)++;
    if (ier != 0) return((ier < 0) ? SPBCG_PSOLVE_FAIL_UNREC : SPBCG_PSOLVE_FAIL_REC);
  }
  else N_VScale(ONE, r_star, r);

  if (scale_b) N_VProd(sb, r, r_star);
  else N_VScale(ONE, r, r_star);

  /* Initialize beta_denom to the dot product of r0 with r0 */

  beta_denom = N_VDotProd(r_star, r_star);

  /* Set r_norm to L2 norm of r_star = sb P1_inv r_0, and
     return if small */

  *res_norm = r_norm = rho = RSqrt(beta_denom);
  if (r_norm <= delta) return(SPBCG_SUCCESS);

  /* Copy r_star to r and p */

  N_VScale(ONE, r_star, r);
  N_VScale(ONE, r_star, p);

  /* Begin main iteration loop */

  for(l = 0; l < l_max; l++) {

    (*nli)++;

    /* Generate Ap = A-tilde p, where A-tilde = sb P1_inv A P2_inv sx_inv */

    /*   Apply x-scaling: vtemp = sx_inv p */

    if (scale_x) N_VDiv(p, sx, vtemp);
    else N_VScale(ONE, p, vtemp);

    /*   Apply right preconditioner: vtemp = P2_inv sx_inv p */

    if (preOnRight) {
      N_VScale(ONE, vtemp, Ap);
      ier = psolve(P_data, Ap, vtemp, PREC_RIGHT);
      (*nps)++;
      if (ier != 0) return((ier < 0) ? SPBCG_PSOLVE_FAIL_UNREC : SPBCG_PSOLVE_FAIL_REC);
    }

    /*   Apply A: Ap = A P2_inv sx_inv p */

    ier = atimes(A_data, vtemp, Ap );
    if (ier != 0)
      return((ier < 0) ? SPBCG_ATIMES_FAIL_UNREC : SPBCG_ATIMES_FAIL_REC);

    /*   Apply left preconditioner: vtemp = P1_inv A P2_inv sx_inv p */

    if (preOnLeft) {
      ier = psolve(P_data, Ap, vtemp, PREC_LEFT);
      (*nps)++;
      if (ier != 0) return((ier < 0) ? SPBCG_PSOLVE_FAIL_UNREC : SPBCG_PSOLVE_FAIL_REC);
    }
    else N_VScale(ONE, Ap, vtemp);

    /*   Apply b-scaling: Ap = sb P1_inv A P2_inv sx_inv p */

    if (scale_b) N_VProd(sb, vtemp, Ap);
    else N_VScale(ONE, vtemp, Ap);


    /* Calculate alpha = <r,r_star>/<Ap,r_star> */

    alpha = ((N_VDotProd(r, r_star) / N_VDotProd(Ap, r_star)));

    /* Update q = r - alpha*Ap = r - alpha*(sb P1_inv A P2_inv sx_inv p) */

    N_VLinearSum(ONE, r, -alpha, Ap, q);

    /* Generate u = A-tilde q */

    /*   Apply x-scaling: vtemp = sx_inv q */

    if (scale_x) N_VDiv(q, sx, vtemp);
    else N_VScale(ONE, q, vtemp);

    /*   Apply right preconditioner: vtemp = P2_inv sx_inv q */

    if (preOnRight) {
      N_VScale(ONE, vtemp, u);
      ier = psolve(P_data, u, vtemp, PREC_RIGHT);
      (*nps)++;
      if (ier != 0) return((ier < 0) ? SPBCG_PSOLVE_FAIL_UNREC : SPBCG_PSOLVE_FAIL_REC);
    }

    /*   Apply A: u = A P2_inv sx_inv u */

    ier = atimes(A_data, vtemp, u );
    if (ier != 0)
      return((ier < 0) ? SPBCG_ATIMES_FAIL_UNREC : SPBCG_ATIMES_FAIL_REC);

    /*   Apply left preconditioner: vtemp = P1_inv A P2_inv sx_inv p */

    if (preOnLeft) {
      ier = psolve(P_data, u, vtemp, PREC_LEFT);
      (*nps)++;
      if (ier != 0) return((ier < 0) ? SPBCG_PSOLVE_FAIL_UNREC : SPBCG_PSOLVE_FAIL_REC);
    }
    else N_VScale(ONE, u, vtemp);

    /*   Apply b-scaling: u = sb P1_inv A P2_inv sx_inv u */

    if (scale_b) N_VProd(sb, vtemp, u);
    else N_VScale(ONE, vtemp, u);


    /* Calculate omega = <u,q>/<u,u> */

    omega_denom = N_VDotProd(u, u);
    if (omega_denom == ZERO) omega_denom = ONE;
    omega = (N_VDotProd(u, q) / omega_denom);

    /* Update x = x + alpha*p + omega*q */

    N_VLinearSum(alpha, p, omega, q, vtemp);
    N_VLinearSum(ONE, x, ONE, vtemp, x);

    /* Update the residual r = q - omega*u */

    N_VLinearSum(ONE, q, -omega, u, r);

    /* Set rho = norm(r) and check convergence */

    *res_norm = rho = RSqrt(N_VDotProd(r, r));
    if (rho <= delta) {
      converged = TRUE;
      break;
    }

    /* Not yet converged, continue iteration */
    /* Update beta = <rnew,r_star> / <rold,r_start> * alpha / omega */

    beta_num = N_VDotProd(r, r_star);
    beta = ((beta_num / beta_denom) * (alpha / omega));
    beta_denom = beta_num;

    /* Update p = r + beta*(p - omega*Ap) */

    N_VLinearSum(ONE, p, -omega, Ap, vtemp);
    N_VLinearSum(ONE, r, beta, vtemp, p);

  }

  /* Main loop finished */

  if ((converged == TRUE) || (rho < r_norm)) {

    /* Apply the x-scaling and right preconditioner: x = P2_inv sx_inv x */

    if (scale_x) N_VDiv(x, sx, x);
    if (preOnRight) {
      ier = psolve(P_data, x, vtemp, PREC_RIGHT);
      (*nps)++;
      if (ier != 0) return((ier < 0) ? SPBCG_PSOLVE_FAIL_UNREC : SPBCG_PSOLVE_FAIL_REC);
      N_VScale(ONE, vtemp, x);
    }

    if (converged == TRUE) return(SPBCG_SUCCESS);
    else return(SPBCG_RES_REDUCED);
  }
  else return(SPBCG_CONV_FAIL);
}

/*
 * -----------------------------------------------------------------
 * Function : SpbcgFree
 * -----------------------------------------------------------------
 */

void SpbcgFree(SpbcgMem mem)
{

  if (mem == NULL) return;

  N_VDestroy(mem->r_star);
  N_VDestroy(mem->r);
  N_VDestroy(mem->p);
  N_VDestroy(mem->q);
  N_VDestroy(mem->u);
  N_VDestroy(mem->Ap);
  N_VDestroy(mem->vtemp);

  free(mem); mem = NULL;
}
```
