## Supplementary scripts for "A multi-scale *in silico* mouse model for insulin resistance and humanoid type 2 diabetes": sundials_spgmr.c

```
/*
 * -----------------------------------------------------------------
 * $Revision: 749 $
 * $Date: 2011-06-22 14:49:10 +0200 (Wed, 22 Jun 2011) $
 * -----------------------------------------------------------------
 * Programmer(s): Scott D. Cohen, Alan C. Hindmarsh and
 *                Radu Serban @ LLNL
 * -----------------------------------------------------------------
 * Copyright (c) 2002, The Regents of the University of California.
 * Produced at the Lawrence Livermore National Laboratory.
 * All rights reserved.
 * For details, see the LICENSE file.
 * -----------------------------------------------------------------
 * This is the implementation file for the scaled preconditioned
 * GMRES (SPGMR) iterative linear solver.
 * -----------------------------------------------------------------
 */

#include <stdio.h>
#include <stdlib.h>

#include <sundials/sundials_spgmr.h>
#include <sundials/sundials_math.h>

/*
 * -----------------------------------------------------------------
 * private constants
 * -----------------------------------------------------------------
 */

#define ZERO RCONST(0.0)
#define ONE  RCONST(1.0)

/*
 * -----------------------------------------------------------------
 * Function : SpgmrMalloc
 * -----------------------------------------------------------------
 */

SpgmrMem SpgmrMalloc(int l_max, N_Vector vec_tmpl)
{
  SpgmrMem mem;
  N_Vector *V, xcor, vtemp;
  realtype **Hes, *givens, *yg;
  int k, i;
 
  /* Check the input parameters. */

  if (l_max <= 0) return(NULL);

  /* Get memory for the Krylov basis vectors V[0], ..., V[l_max]. */

  V = NULL;
  V = N_VCloneVectorArray(l_max+1, vec_tmpl);
  if (V == NULL) return(NULL);

  /* Get memory for the Hessenberg matrix Hes. */

  Hes = NULL;
  Hes = (realtype **) malloc((l_max+1)*sizeof(realtype *)); 
  if (Hes == NULL) {
    N_VDestroyVectorArray(V, l_max+1);
    return(NULL);
  }

  for (k = 0; k <= l_max; k++) {
    Hes[k] = NULL;
    Hes[k] = (realtype *) malloc(l_max*sizeof(realtype));
    if (Hes[k] == NULL) {
      for (i = 0; i < k; i++) {free(Hes[i]); Hes[i] = NULL;}
      free(Hes); Hes = NULL;
      N_VDestroyVectorArray(V, l_max+1);
      return(NULL);
    }
  }
  
  /* Get memory for Givens rotation components. */
  
  givens = NULL;
  givens = (realtype *) malloc(2*l_max*sizeof(realtype));
  if (givens == NULL) {
    for (i = 0; i <= l_max; i++) {free(Hes[i]); Hes[i] = NULL;}
    free(Hes); Hes = NULL;
    N_VDestroyVectorArray(V, l_max+1);
    return(NULL);
  }

  /* Get memory to hold the correction to z_tilde. */

  xcor = NULL;
  xcor = N_VClone(vec_tmpl);
  if (xcor == NULL) {
    free(givens); givens = NULL;
    for (i = 0; i <= l_max; i++) {free(Hes[i]); Hes[i] = NULL;}
    free(Hes); Hes = NULL;
    N_VDestroyVectorArray(V, l_max+1);
    return(NULL);
  }

  /* Get memory to hold SPGMR y and g vectors. */

  yg = NULL;
  yg = (realtype *) malloc((l_max+1)*sizeof(realtype));
  if (yg == NULL) {
    N_VDestroy(xcor);
    free(givens); givens = NULL;
    for (i = 0; i <= l_max; i++) {free(Hes[i]); Hes[i] = NULL;}
    free(Hes); Hes = NULL;
    N_VDestroyVectorArray(V, l_max+1);
    return(NULL);
  }

  /* Get an array to hold a temporary vector. */

  vtemp = NULL;
  vtemp = N_VClone(vec_tmpl);
  if (vtemp == NULL) {
    free(yg); yg = NULL;
    N_VDestroy(xcor);
    free(givens); givens = NULL;
    for (i = 0; i <= l_max; i++) {free(Hes[i]); Hes[i] = NULL;}
    free(Hes); Hes = NULL;
    N_VDestroyVectorArray(V, l_max+1);
    return(NULL);
  }

  /* Get memory for an SpgmrMemRec containing SPGMR matrices and vectors. */

  mem = NULL;
  mem = (SpgmrMem) malloc(sizeof(SpgmrMemRec));
  if (mem == NULL) {
    N_VDestroy(vtemp);
    free(yg); yg = NULL;
    N_VDestroy(xcor);
    free(givens); givens = NULL;
    for (i = 0; i <= l_max; i++) {free(Hes[i]); Hes[i] = NULL;}
    free(Hes); Hes = NULL;
    N_VDestroyVectorArray(V, l_max+1);
    return(NULL); 
  }

  /* Set the fields of mem. */

  mem->l_max = l_max;
  mem->V = V;
  mem->Hes = Hes;
  mem->givens = givens;
  mem->xcor = xcor;
  mem->yg = yg;
  mem->vtemp = vtemp;

  /* Return the pointer to SPGMR memory. */

  return(mem);
}

/*
 * -----------------------------------------------------------------
 * Function : SpgmrSolve
 * -----------------------------------------------------------------
 */

int SpgmrSolve(SpgmrMem mem, void *A_data, N_Vector x, N_Vector b,
               int pretype, int gstype, realtype delta, int max_restarts,
               void *P_data, N_Vector s1, N_Vector s2, ATimesFn atimes,
               PSolveFn psolve, realtype *res_norm, int *nli, int *nps)
{
  N_Vector *V, xcor, vtemp;
  realtype **Hes, *givens, *yg;
  realtype beta, rotation_product, r_norm, s_product, rho;
  booleantype preOnLeft, preOnRight, scale2, scale1, converged;
  int i, j, k, l, l_plus_1, l_max, krydim, ier, ntries;

  if (mem == NULL) return(SPGMR_MEM_NULL);

  /* Initialize some variables */

  l_plus_1 = 0;
  krydim = 0;

  /* Make local copies of mem variables. */

  l_max  = mem->l_max;
  V      = mem->V;
  Hes    = mem->Hes;
  givens = mem->givens;
  xcor   = mem->xcor;
  yg     = mem->yg;
  vtemp  = mem->vtemp;

  *nli = *nps = 0;    /* Initialize counters */
  converged = FALSE;  /* Initialize converged flag */

  if (max_restarts < 0) max_restarts = 0;

  if ((pretype != PREC_LEFT) && (pretype != PREC_RIGHT) && (pretype != PREC_BOTH))
    pretype = PREC_NONE;
  
  preOnLeft  = ((pretype == PREC_LEFT) || (pretype == PREC_BOTH));
  preOnRight = ((pretype == PREC_RIGHT) || (pretype == PREC_BOTH));
  scale1 = (s1 != NULL);
  scale2 = (s2 != NULL);

  /* Set vtemp and V[0] to initial (unscaled) residual r_0 = b - A*x_0. */

  if (N_VDotProd(x, x) == ZERO) {
    N_VScale(ONE, b, vtemp);
  } else {
    ier = atimes(A_data, x, vtemp);
    if (ier != 0)
      return((ier < 0) ? SPGMR_ATIMES_FAIL_UNREC : SPGMR_ATIMES_FAIL_REC);
    N_VLinearSum(ONE, b, -ONE, vtemp, vtemp);
  }
  N_VScale(ONE, vtemp, V[0]);

  /* Apply left preconditioner and left scaling to V[0] = r_0. */
  
  if (preOnLeft) {
    ier = psolve(P_data, V[0], vtemp, PREC_LEFT);
    (*nps)++;
    if (ier != 0)
      return((ier < 0) ? SPGMR_PSOLVE_FAIL_UNREC : SPGMR_PSOLVE_FAIL_REC);
  } else {
    N_VScale(ONE, V[0], vtemp);
  }
  
  if (scale1) {
    N_VProd(s1, vtemp, V[0]);   
  } else {
    N_VScale(ONE, vtemp, V[0]);
  }

  /* Set r_norm = beta to L2 norm of V[0] = s1 P1_inv r_0, and
     return if small.  */

  *res_norm = r_norm = beta = RSqrt(N_VDotProd(V[0], V[0])); 
  if (r_norm <= delta)
    return(SPGMR_SUCCESS);

  /* Initialize rho to avoid compiler warning message */

  rho = beta;

  /* Set xcor = 0. */

  N_VConst(ZERO, xcor);


  /* Begin outer iterations: up to (max_restarts + 1) attempts. */
  
  for (ntries = 0; ntries <= max_restarts; ntries++) {
    
    /* Initialize the Hessenberg matrix Hes and Givens rotation
       product.  Normalize the initial vector V[0].             */
    
    for (i = 0; i <= l_max; i++)
      for (j = 0; j < l_max; j++)
        Hes[i][j] = ZERO;
    
    rotation_product = ONE;
    
    N_VScale(ONE/r_norm, V[0], V[0]);
    
    /* Inner loop: generate Krylov sequence and Arnoldi basis. */
    
    for (l = 0; l < l_max; l++) {
      
      (*nli)++;
      
      krydim = l_plus_1 = l + 1;
      
      /* Generate A-tilde V[l], where A-tilde = s1 P1_inv A P2_inv s2_inv. */
      
      /* Apply right scaling: vtemp = s2_inv V[l]. */

      if (scale2) N_VDiv(V[l], s2, vtemp);
      else N_VScale(ONE, V[l], vtemp);
      
      /* Apply right preconditioner: vtemp = P2_inv s2_inv V[l]. */ 

      if (preOnRight) {
        N_VScale(ONE, vtemp, V[l_plus_1]);
        ier = psolve(P_data, V[l_plus_1], vtemp, PREC_RIGHT);
        (*nps)++;
        if (ier != 0)
          return((ier < 0) ? SPGMR_PSOLVE_FAIL_UNREC : SPGMR_PSOLVE_FAIL_REC);
      }
      
      /* Apply A: V[l+1] = A P2_inv s2_inv V[l]. */

      ier = atimes(A_data, vtemp, V[l_plus_1] );
      if (ier != 0)
        return((ier < 0) ? SPGMR_ATIMES_FAIL_UNREC : SPGMR_ATIMES_FAIL_REC);
      
      /* Apply left preconditioning: vtemp = P1_inv A P2_inv s2_inv V[l]. */

      if (preOnLeft) {
        ier = psolve(P_data, V[l_plus_1], vtemp, PREC_LEFT);
        (*nps)++;
        if (ier != 0)
          return((ier < 0) ? SPGMR_PSOLVE_FAIL_UNREC : SPGMR_PSOLVE_FAIL_REC);
      } else {
        N_VScale(ONE, V[l_plus_1], vtemp);
      }
      
      /* Apply left scaling: V[l+1] = s1 P1_inv A P2_inv s2_inv V[l]. */

      if (scale1) {
        N_VProd(s1, vtemp, V[l_plus_1]);
      } else {
        N_VScale(ONE, vtemp, V[l_plus_1]);
      }
      
      /*  Orthogonalize V[l+1] against previous V[i]: V[l+1] = w_tilde. */
      
      if (gstype == CLASSICAL_GS) {
        if (ClassicalGS(V, Hes, l_plus_1, l_max, &(Hes[l_plus_1][l]),
                        vtemp, yg) != 0)
          return(SPGMR_GS_FAIL);
      } else {
        if (ModifiedGS(V, Hes, l_plus_1, l_max, &(Hes[l_plus_1][l])) != 0) 
          return(SPGMR_GS_FAIL);
      }
      
      /*  Update the QR factorization of Hes. */
      
      if(QRfact(krydim, Hes, givens, l) != 0 )
        return(SPGMR_QRFACT_FAIL);
      
      /*  Update residual norm estimate; break if convergence test passes. */
      
      rotation_product *= givens[2*l+1];
      *res_norm = rho = ABS(rotation_product*r_norm);
      
      if (rho <= delta) { converged = TRUE; break; }
      
      /* Normalize V[l+1] with norm value from the Gram-Schmidt routine. */

      N_VScale(ONE/Hes[l_plus_1][l], V[l_plus_1], V[l_plus_1]);
    }
    
    /* Inner loop is done.  Compute the new correction vector xcor. */
    
    /* Construct g, then solve for y. */

    yg[0] = r_norm;
    for (i = 1; i <= krydim; i++) yg[i]=ZERO;
    if (QRsol(krydim, Hes, givens, yg) != 0)
      return(SPGMR_QRSOL_FAIL);
    
    /* Add correction vector V_l y to xcor. */

    for (k = 0; k < krydim; k++)
      N_VLinearSum(yg[k], V[k], ONE, xcor, xcor);
    
    /* If converged, construct the final solution vector x and return. */

    if (converged) {
      
      /* Apply right scaling and right precond.: vtemp = P2_inv s2_inv xcor. */
      
      if (scale2) N_VDiv(xcor, s2, xcor);
      if (preOnRight) {
        ier = psolve(P_data, xcor, vtemp, PREC_RIGHT);
        (*nps)++;
        if (ier != 0)
          return((ier < 0) ? SPGMR_PSOLVE_FAIL_UNREC : SPGMR_PSOLVE_FAIL_REC);
      } else {
        N_VScale(ONE, xcor, vtemp);
      }
      
      /* Add vtemp to initial x to get final solution x, and return */

      N_VLinearSum(ONE, x, ONE, vtemp, x);
      
      return(SPGMR_SUCCESS);
    }
    
    /* Not yet converged; if allowed, prepare for restart. */
    
    if (ntries == max_restarts) break;
    
    /* Construct last column of Q in yg. */

    s_product = ONE;
    for (i = krydim; i > 0; i--) {
      yg[i] = s_product*givens[2*i-2];
      s_product *= givens[2*i-1];
    }
    yg[0] = s_product;
    
    /* Scale r_norm and yg. */
    r_norm *= s_product;
    for (i = 0; i <= krydim; i++)
      yg[i] *= r_norm;
    r_norm = ABS(r_norm);
    
    /* Multiply yg by V_(krydim+1) to get last residual vector; restart. */
    N_VScale(yg[0], V[0], V[0]);
    for (k = 1; k <= krydim; k++)
      N_VLinearSum(yg[k], V[k], ONE, V[0], V[0]);
    
  }
  
  /* Failed to converge, even after allowed restarts.
     If the residual norm was reduced below its initial value, compute
     and return x anyway.  Otherwise return failure flag. */

  if (rho < beta) {
    
    /* Apply right scaling and right precond.: vtemp = P2_inv s2_inv xcor. */
    
    if (scale2) N_VDiv(xcor, s2, xcor);
    if (preOnRight) {
      ier = psolve(P_data, xcor, vtemp, PREC_RIGHT);
      (*nps)++;
      if (ier != 0)
        return((ier < 0) ? SPGMR_PSOLVE_FAIL_UNREC : SPGMR_PSOLVE_FAIL_REC);
      } else {
      N_VScale(ONE, xcor, vtemp);
    }

    /* Add vtemp to initial x to get final solution x, and return. */

    N_VLinearSum(ONE, x, ONE, vtemp, x);
    
    return(SPGMR_RES_REDUCED);
  }

  return(SPGMR_CONV_FAIL); 
}

/*
 * -----------------------------------------------------------------
 * Function : SpgmrFree
 * -----------------------------------------------------------------
 */

void SpgmrFree(SpgmrMem mem)
{
  int i, l_max;
  realtype **Hes, *givens, *yg;
  
  if (mem == NULL) return;

  l_max  = mem->l_max;
  Hes    = mem->Hes;
  givens = mem->givens;
  yg     = mem->yg;

  for (i = 0; i <= l_max; i++) {free(Hes[i]); Hes[i] = NULL;}
  free(Hes); Hes = NULL;
  free(mem->givens); givens = NULL; 
  free(mem->yg); yg = NULL;

  N_VDestroyVectorArray(mem->V, l_max+1);
  N_VDestroy(mem->xcor);
  N_VDestroy(mem->vtemp);

  free(mem); mem = NULL;
}
```
