## Supplementary scripts for "A multi-scale *in silico* mouse model for insulin resistance and humanoid type 2 diabetes": sundials_sptfqmr.c

```
/*
 * -----------------------------------------------------------------
 * $Revision: 749 $
 * $Date: 2011-06-22 14:49:10 +0200 (Wed, 22 Jun 2011) $
 * -----------------------------------------------------------------
 * Programmer(s): Aaron Collier @ LLNL
 * -----------------------------------------------------------------
 * Copyright (c) 2005, The Regents of the University of California.
 * Produced at the Lawrence Livermore National Laboratory.
 * All rights reserved.
 * For details, see the LICENSE file.
 * -----------------------------------------------------------------
 * This is the implementation file for the scaled preconditioned
 * Transpose-Free Quasi-Minimal Residual (SPTFQMR) linear solver.
 * -----------------------------------------------------------------
 */

#include <stdio.h>
#include <stdlib.h>

#include <sundials/sundials_sptfqmr.h>
#include <sundials/sundials_math.h>

/*
 * -----------------------------------------------------------------
 * private constants
 * -----------------------------------------------------------------
 */

#define ZERO RCONST(0.0)
#define ONE  RCONST(1.0)

/*
 * -----------------------------------------------------------------
 * Function : SptfqmrMalloc
 * -----------------------------------------------------------------
 */

SptfqmrMem SptfqmrMalloc(int l_max, N_Vector vec_tmpl)
{
  SptfqmrMem mem;
  N_Vector *r;
  N_Vector q, d, v, p, u;
  N_Vector r_star, vtemp1, vtemp2, vtemp3;

  /* Check the input parameters */
  if ((l_max <= 0) || (vec_tmpl == NULL)) return(NULL);

  /* Allocate space for vectors */
  r_star = NULL;
  r_star = N_VClone(vec_tmpl);
  if (r_star == NULL) return(NULL);

  q = NULL;
  q = N_VClone(vec_tmpl);
  if (q == NULL) {
    N_VDestroy(r_star);
    return(NULL);
  }

  d = NULL;
  d = N_VClone(vec_tmpl);
  if (d == NULL) {
    N_VDestroy(r_star);
    N_VDestroy(q);
    return(NULL);
  }

  v = NULL;
  v = N_VClone(vec_tmpl);
  if (v == NULL) {
    N_VDestroy(r_star);
    N_VDestroy(q);
    N_VDestroy(d);
    return(NULL);
  }

  p = NULL;
  p = N_VClone(vec_tmpl);
  if (p == NULL) {
    N_VDestroy(r_star);
    N_VDestroy(q);
    N_VDestroy(d);
    N_VDestroy(v);
    return(NULL);
  }

  r = NULL;
  r = N_VCloneVectorArray(2, vec_tmpl);
  if (r == NULL) {
    N_VDestroy(r_star);
    N_VDestroy(q);
    N_VDestroy(d);
    N_VDestroy(v);
    N_VDestroy(p);
    return(NULL);
  }

  u = NULL;
  u = N_VClone(vec_tmpl);
  if (u == NULL) {
    N_VDestroy(r_star);
    N_VDestroy(q);
    N_VDestroy(d);
    N_VDestroy(v);
    N_VDestroy(p);
    N_VDestroyVectorArray(r, 2);
    return(NULL);
  }

  vtemp1 = NULL;
  vtemp1 = N_VClone(vec_tmpl);
  if (vtemp1 == NULL) {
    N_VDestroy(r_star);
    N_VDestroy(q);
    N_VDestroy(d);
    N_VDestroy(v);
    N_VDestroy(p);
    N_VDestroyVectorArray(r, 2);
    N_VDestroy(u);
    return(NULL);
  }

  vtemp2 = NULL;
  vtemp2 = N_VClone(vec_tmpl);
  if (vtemp2 == NULL) {
    N_VDestroy(r_star);
    N_VDestroy(q);
    N_VDestroy(d);
    N_VDestroy(v);
    N_VDestroy(p);
    N_VDestroyVectorArray(r, 2);
    N_VDestroy(u);
    N_VDestroy(vtemp1);
    return(NULL);
  }

  vtemp3 = NULL;
  vtemp3 = N_VClone(vec_tmpl);
  if (vtemp3 == NULL) {
    N_VDestroy(r_star);
    N_VDestroy(q);
    N_VDestroy(d);
    N_VDestroy(v);
    N_VDestroy(p);
    N_VDestroyVectorArray(r, 2);
    N_VDestroy(u);
    N_VDestroy(vtemp1);
    N_VDestroy(vtemp2);
    return(NULL);
  }

  /* Allocate memory for SptfqmrMemRec */
  mem = NULL;
  mem = (SptfqmrMem) malloc(sizeof(SptfqmrMemRec));
  if (mem == NULL) {
    N_VDestroy(r_star);
    N_VDestroy(q);
    N_VDestroy(d);
    N_VDestroy(v);
    N_VDestroy(p);
    N_VDestroyVectorArray(r, 2);
    N_VDestroy(u);
    N_VDestroy(vtemp1);
    N_VDestroy(vtemp2);
    N_VDestroy(vtemp3);
    return(NULL);
  }

  /* Intialize SptfqmrMemRec data structure */
  mem->l_max  = l_max;
  mem->r_star = r_star;
  mem->q      = q;
  mem->d      = d;
  mem->v      = v;
  mem->p      = p;
  mem->r      = r;
  mem->u      = u;
  mem->vtemp1 = vtemp1;
  mem->vtemp2 = vtemp2;
  mem->vtemp3 = vtemp3;

  /* Return pointer to SPTFQMR memory block */
  return(mem);
}

#define l_max  (mem->l_max)
#define r_star (mem->r_star)
#define q_     (mem->q)
#define d_     (mem->d)
#define v_     (mem->v)
#define p_     (mem->p)
#define r_     (mem->r)
#define u_     (mem->u)
#define vtemp1 (mem->vtemp1)
#define vtemp2 (mem->vtemp2)
#define vtemp3 (mem->vtemp3)

/*
 * -----------------------------------------------------------------
 * Function : SptfqmrSolve
 * -----------------------------------------------------------------
 */

int SptfqmrSolve(SptfqmrMem mem, void *A_data, N_Vector x, N_Vector b,
		 int pretype, realtype delta, void *P_data, N_Vector sx,
		 N_Vector sb, ATimesFn atimes, PSolveFn psolve,
		 realtype *res_norm, int *nli, int *nps)
{
  realtype alpha, tau, eta, beta, c, sigma, v_bar, omega;
  realtype rho[2];
  realtype r_init_norm, r_curr_norm;
  realtype temp_val;
  booleantype preOnLeft, preOnRight, scale_x, scale_b, converged;
  booleantype b_ok;
  int n, m, ier;

  /* Exit immediately if memory pointer is NULL */
  if (mem == NULL) return(SPTFQMR_MEM_NULL);

  temp_val = r_curr_norm = -ONE;  /* Initialize to avoid compiler warnings */

  *nli = *nps = 0;    /* Initialize counters */
  converged = FALSE;  /* Initialize convergence flag */
  b_ok = FALSE;

  if ((pretype != PREC_LEFT)  &&
      (pretype != PREC_RIGHT) &&
      (pretype != PREC_BOTH)) pretype = PREC_NONE;

  preOnLeft  = ((pretype == PREC_BOTH) || (pretype == PREC_LEFT));
  preOnRight = ((pretype == PREC_BOTH) || (pretype == PREC_RIGHT));

  scale_x = (sx != NULL);
  scale_b = (sb != NULL);

  /* Set r_star to initial (unscaled) residual r_star = r_0 = b - A*x_0 */
  /* NOTE: if x == 0 then just set residual to b and continue */
  if (N_VDotProd(x, x) == ZERO) N_VScale(ONE, b, r_star);
  else {
    ier = atimes(A_data, x, r_star);
    if (ier != 0)
      return((ier < 0) ? SPTFQMR_ATIMES_FAIL_UNREC : SPTFQMR_ATIMES_FAIL_REC);
    N_VLinearSum(ONE, b, -ONE, r_star, r_star);
  }

  /* Apply left preconditioner and b-scaling to r_star (or really just r_0) */
  if (preOnLeft) {
    ier = psolve(P_data, r_star, vtemp1, PREC_LEFT);
    (*nps)++;
    if (ier != 0)
      return((ier < 0) ? SPTFQMR_PSOLVE_FAIL_UNREC : SPTFQMR_PSOLVE_FAIL_REC);
  }
  else N_VScale(ONE, r_star, vtemp1);
  if (scale_b) N_VProd(sb, vtemp1, r_star);
  else N_VScale(ONE, vtemp1, r_star);

  /* Initialize rho[0] */
  /* NOTE: initialized here to reduce number of computations - avoid need
           to compute r_star^T*r_star twice, and avoid needlessly squaring
           values */
  rho[0] = N_VDotProd(r_star, r_star);

  /* Compute norm of initial residual (r_0) to see if we really need
     to do anything */
  *res_norm = r_init_norm = RSqrt(rho[0]);
  if (r_init_norm <= delta) return(SPTFQMR_SUCCESS);

  /* Set v_ = A*r_0 (preconditioned and scaled) */
  if (scale_x) N_VDiv(r_star, sx, vtemp1);
  else N_VScale(ONE, r_star, vtemp1);
  if (preOnRight) {
    N_VScale(ONE, vtemp1, v_);
    ier = psolve(P_data, v_, vtemp1, PREC_RIGHT);
    (*nps)++;
    if (ier != 0) return((ier < 0) ? SPTFQMR_PSOLVE_FAIL_UNREC : SPTFQMR_PSOLVE_FAIL_REC);
  }
  ier = atimes(A_data, vtemp1, v_);
  if (ier != 0)
    return((ier < 0) ? SPTFQMR_ATIMES_FAIL_UNREC : SPTFQMR_ATIMES_FAIL_REC);
  if (preOnLeft) {
    ier = psolve(P_data, v_, vtemp1, PREC_LEFT);
    (*nps)++;
    if (ier != 0) return((ier < 0) ? SPTFQMR_PSOLVE_FAIL_UNREC : SPTFQMR_PSOLVE_FAIL_REC);
  }
  else N_VScale(ONE, v_, vtemp1);
  if (scale_b) N_VProd(sb, vtemp1, v_);
  else N_VScale(ONE, vtemp1, v_);

  /* Initialize remaining variables */
  N_VScale(ONE, r_star, r_[0]);
  N_VScale(ONE, r_star, u_);
  N_VScale(ONE, r_star, p_);
  N_VConst(ZERO, d_);

  tau = r_init_norm;
  v_bar = eta = ZERO;

  /* START outer loop */
  for (n = 0; n < l_max; ++n) {

    /* Increment linear iteration counter */
    (*nli)++;

    /* sigma = r_star^T*v_ */
    sigma = N_VDotProd(r_star, v_);

    /* alpha = rho[0]/sigma */
    alpha = rho[0]/sigma;

    /* q_ = u_-alpha*v_ */
    N_VLinearSum(ONE, u_, -alpha, v_, q_);

    /* r_[1] = r_[0]-alpha*A*(u_+q_) */
    N_VLinearSum(ONE, u_, ONE, q_, r_[1]);
    if (scale_x) N_VDiv(r_[1], sx, r_[1]);
    if (preOnRight) {
      N_VScale(ONE, r_[1], vtemp1);
      ier = psolve(P_data, vtemp1, r_[1], PREC_RIGHT);
      (*nps)++;
      if (ier != 0) return((ier < 0) ? SPTFQMR_PSOLVE_FAIL_UNREC : SPTFQMR_PSOLVE_FAIL_REC);
    }
    ier = atimes(A_data, r_[1], vtemp1);
    if (ier != 0)
      return((ier < 0) ? SPTFQMR_ATIMES_FAIL_UNREC : SPTFQMR_ATIMES_FAIL_REC);
    if (preOnLeft) {
      ier = psolve(P_data, vtemp1, r_[1], PREC_LEFT);
      (*nps)++;
      if (ier != 0) return((ier < 0) ? SPTFQMR_PSOLVE_FAIL_UNREC : SPTFQMR_PSOLVE_FAIL_REC);
    }
    else N_VScale(ONE, vtemp1, r_[1]);
    if (scale_b) N_VProd(sb, r_[1], vtemp1);
    else N_VScale(ONE, r_[1], vtemp1);
    N_VLinearSum(ONE, r_[0], -alpha, vtemp1, r_[1]);

    /* START inner loop */
    for (m = 0; m < 2; ++m) {

      /* d_ = [*]+(v_bar^2*eta/alpha)*d_ */
      /* NOTES:
       *   (1) [*] = u_ if m == 0, and q_ if m == 1
       *   (2) using temp_val reduces the number of required computations
       *       if the inner loop is executed twice
       */
      if (m == 0) {
	temp_val = RSqrt(N_VDotProd(r_[1], r_[1]));
	omega = RSqrt(RSqrt(N_VDotProd(r_[0], r_[0]))*temp_val);
	N_VLinearSum(ONE, u_, SQR(v_bar)*eta/alpha, d_, d_);
      }
      else {
	omega = temp_val;
	N_VLinearSum(ONE, q_, SQR(v_bar)*eta/alpha, d_, d_);
      }

      /* v_bar = omega/tau */
      v_bar = omega/tau;

      /* c = (1+v_bar^2)^(-1/2) */
      c = ONE / RSqrt(ONE+SQR(v_bar));

      /* tau = tau*v_bar*c */
      tau = tau*v_bar*c;

      /* eta = c^2*alpha */
      eta = SQR(c)*alpha;

      /* x = x+eta*d_ */
      N_VLinearSum(ONE, x, eta, d_, x);

      /* Check for convergence... */
      /* NOTE: just use approximation to norm of residual, if possible */
      *res_norm = r_curr_norm = tau*RSqrt(m+1);

      /* Exit inner loop if iteration has converged based upon approximation
	 to norm of current residual */
      if (r_curr_norm <= delta) {
	converged = TRUE;
	break;
      }

      /* Decide if actual norm of residual vector should be computed */
      /* NOTES:
       *   (1) if r_curr_norm > delta, then check if actual residual norm
       *       is OK (recall we first compute an approximation)
       *   (2) if r_curr_norm >= r_init_norm and m == 1 and n == l_max, then
       *       compute actual residual norm to see if the iteration can be
       *       saved
       *   (3) the scaled and preconditioned right-hand side of the given
       *       linear system (denoted by b) is only computed once, and the
       *       result is stored in vtemp3 so it can be reused - reduces the
       *       number of psovles if using left preconditioning
       */
      if ((r_curr_norm > delta) ||
	  (r_curr_norm >= r_init_norm && m == 1 && n == l_max)) {

	/* Compute norm of residual ||b-A*x||_2 (preconditioned and scaled) */
	if (scale_x) N_VDiv(x, sx, vtemp1);
	else N_VScale(ONE, x, vtemp1);
	if (preOnRight) {
	  ier = psolve(P_data, vtemp1, vtemp2, PREC_RIGHT);
	  (*nps)++;
	  if (ier != 0) return((ier < 0) ? SPTFQMR_PSOLVE_FAIL_UNREC : SPTFQMR_PSOLVE_FAIL_UNREC);
	  N_VScale(ONE, vtemp2, vtemp1);
	}
	ier = atimes(A_data, vtemp1, vtemp2);
        if (ier != 0)
          return((ier < 0) ? SPTFQMR_ATIMES_FAIL_UNREC : SPTFQMR_ATIMES_FAIL_REC);
	if (preOnLeft) {
	  ier = psolve(P_data, vtemp2, vtemp1, PREC_LEFT);
	  (*nps)++;
	  if (ier != 0) return((ier < 0) ? SPTFQMR_PSOLVE_FAIL_UNREC : SPTFQMR_PSOLVE_FAIL_REC);
	}
	else N_VScale(ONE, vtemp2, vtemp1);
	if (scale_b) N_VProd(sb, vtemp1, vtemp2);
	else N_VScale(ONE, vtemp1, vtemp2);
	/* Only precondition and scale b once (result saved for reuse) */
	if (!b_ok) {
	  b_ok = TRUE;
	  if (preOnLeft) {
	    ier = psolve(P_data, b, vtemp3, PREC_LEFT);
	    (*nps)++;
	    if (ier != 0) return((ier < 0) ? SPTFQMR_PSOLVE_FAIL_UNREC : SPTFQMR_PSOLVE_FAIL_REC);
	  }
	  else N_VScale(ONE, b, vtemp3);
	  if (scale_b) N_VProd(sb, vtemp3, vtemp3);
	}
	N_VLinearSum(ONE, vtemp3, -ONE, vtemp2, vtemp1);
	*res_norm = r_curr_norm = RSqrt(N_VDotProd(vtemp1, vtemp1));

	/* Exit inner loop if inequality condition is satisfied 
	   (meaning exit if we have converged) */
	if (r_curr_norm <= delta) {
	  converged = TRUE;
	  break;
	}

      }

    }  /* END inner loop */

    /* If converged, then exit outer loop as well */
    if (converged == TRUE) break;

    /* rho[1] = r_star^T*r_[1] */
    rho[1] = N_VDotProd(r_star, r_[1]);

    /* beta = rho[1]/rho[0] */
    beta = rho[1]/rho[0];

    /* u_ = r_[1]+beta*q_ */
    N_VLinearSum(ONE, r_[1], beta, q_, u_);

    /* p_ = u_+beta*(q_+beta*p_) */
    N_VLinearSum(beta, q_, SQR(beta), p_, p_);
    N_VLinearSum(ONE, u_, ONE, p_, p_);

    /* v_ = A*p_ */
    if (scale_x) N_VDiv(p_, sx, vtemp1);
    else N_VScale(ONE, p_, vtemp1);
    if (preOnRight) {
      N_VScale(ONE, vtemp1, v_);
      ier = psolve(P_data, v_, vtemp1, PREC_RIGHT);
      (*nps)++;
      if (ier != 0) return((ier < 0) ? SPTFQMR_PSOLVE_FAIL_UNREC : SPTFQMR_PSOLVE_FAIL_REC);
    }
    ier = atimes(A_data, vtemp1, v_);
    if (ier != 0)
      return((ier < 0) ? SPTFQMR_ATIMES_FAIL_UNREC : SPTFQMR_ATIMES_FAIL_REC);
    if (preOnLeft) {
      ier = psolve(P_data, v_, vtemp1, PREC_LEFT);
      (*nps)++;
      if (ier != 0) return((ier < 0) ? SPTFQMR_PSOLVE_FAIL_UNREC : SPTFQMR_PSOLVE_FAIL_REC);
    }
    else N_VScale(ONE, v_, vtemp1);
    if (scale_b) N_VProd(sb, vtemp1, v_);
    else N_VScale(ONE, vtemp1, v_);

    /* Shift variable values */
    /* NOTE: reduces storage requirements */
    N_VScale(ONE, r_[1], r_[0]);
    rho[0] = rho[1];

  }  /* END outer loop */

  /* Determine return value */
  /* If iteration converged or residual was reduced, then return current iterate (x) */
  if ((converged == TRUE) || (r_curr_norm < r_init_norm)) {
    if (scale_x) N_VDiv(x, sx, x);
    if (preOnRight) {
      ier = psolve(P_data, x, vtemp1, PREC_RIGHT);
      (*nps)++;
      if (ier != 0) return((ier < 0) ? SPTFQMR_PSOLVE_FAIL_UNREC : SPTFQMR_PSOLVE_FAIL_UNREC);
      N_VScale(ONE, vtemp1, x);
    }
    if (converged == TRUE) return(SPTFQMR_SUCCESS);
    else return(SPTFQMR_RES_REDUCED);
  }
  /* Otherwise, return error code */
  else return(SPTFQMR_CONV_FAIL);
}

/*
 * -----------------------------------------------------------------
 * Function : SptfqmrFree
 * -----------------------------------------------------------------
 */

void SptfqmrFree(SptfqmrMem mem)
{

  if (mem == NULL) return;

  N_VDestroy(r_star);
  N_VDestroy(q_);
  N_VDestroy(d_);
  N_VDestroy(v_);
  N_VDestroy(p_);
  N_VDestroyVectorArray(r_, 2);
  N_VDestroy(u_);
  N_VDestroy(vtemp1);
  N_VDestroy(vtemp2);
  N_VDestroy(vtemp3);

  free(mem); mem = NULL;
}
```
