## Supplementary scripts for "A multi-scale *in silico* mouse model for insulin resistance and humanoid type 2 diabetes": convertNNCa2NANNCaSBPD.m

```
function [output] = convertNNCa2NANNCaSBPD(input)
% convertNNCa2NANNCaSBPD: complicated name for simple function. This
% function is called by a MEX simulation function when returning
% non-numeric initial conditions. input is a cell-array with string
% entries, defining the initial conditions. some of the strings only
% contain numeric values. this function here will convert these from
% strings to numbers. 

% Information:
% ============
% Copyright (C) 2005-2013 Henning Schmidt,
%
% This program is free software; you can redistribute it and/or modify
% it under the terms of the GNU General Public License as published by
% the Free Software Foundation; either version 2 of the License, or
% (at your option) any later version.
% 
% This program is distributed in the hope that it will be useful,
% but WITHOUT ANY WARRANTY; without even the implied warranty of
% MERCHANTABILITY or FITNESS FOR A PARTICULAR PURPOSE.  See the
% GNU General Public License for more details.
% 
% You should have received a copy of the GNU General Public License
% along with this program; if not, write to the Free Software
% Foundation, Inc., 59 Temple Place, Suite 330, Boston, MA  02111-1307  
% USA.

output = {};
for k=1:length(input),
    test = str2double(input{k});
    if isnan(test),
        output{k} = input{k};
    else
        output{k} = test;
    end
end
```
