## Supplementary scripts for "A multi-scale *in silico* mouse model for insulin resistance and humanoid type 2 diabetes": getparamindicesSBPD.m

```
function [output] = getparamindicesSBPD(model,parameters)
% getparamindicesSBPD: Determines a vector of model parameter indices, the 
% order corresponding to the parameters argument.
% 
% USAGE:
% ======
% [output] = getparamindicesSBPD(model,parameters)
% 
% model: SBmodel, ODE file model, or MEX simulation model
% parameters: Cell-array of parameter names
% 
% OUTPUT:
% =======
% output: Vector of parameter indices

% Information:
% ============
% Copyright (C) 2005-2013 Henning Schmidt,
%
% This program is free software; you can redistribute it and/or modify
% it under the terms of the GNU General Public License as published by
% the Free Software Foundation; either version 2 of the License, or
% (at your option) any later version.
% 
% This program is distributed in the hope that it will be useful,
% but WITHOUT ANY WARRANTY; without even the implied warranty of
% MERCHANTABILITY or FITNESS FOR A PARTICULAR PURPOSE.  See the
% GNU General Public License for more details.
% 
% You should have received a copy of the GNU General Public License
% along with this program; if not, write to the Free Software
% Foundation, Inc., 59 Temple Place, Suite 330, Boston, MA  02111-1307  
% USA.


% check if parameter names is not a cell array
if ischar(parameters),
    parameters = {parameters};
end
% Get all parameter names in the order as they are stored in the model
allnames = SBparameters(model);
% Get the indices
output = zeros(1,length(parameters));
for k = 1:length(parameters),
    % get the index
    index = strmatchSB(parameters{k},allnames,'exact');
    % check ... if error then the current parameter does not exist in the model
    if isempty(index),
        error('Parameter ''%s'' does not exist in the model.',parameters{k});
    end
    output(k) = index;
end
return
```
