## Supplementary scripts for "A multi-scale *in silico* mouse model for insulin resistance and humanoid type 2 diabetes": makeinicondvectorSBPD.m

```
function [icvector] = makeinicondvectorSBPD(model,states,statevalues)
% makeinicondvectorSBPD: The mex simulation functions take a full initial
% condition vector as an input. In order to construct such a vector while
% changing some but possibly not all IC values this function can be used.
%
% This function does handle also non-numeric initial conditions. The
% default state vector is determined based on the given model. Then the
% user provided changes are added as given.
%
% USAGE:
% ======
% [icvector] = makeinicondvectorSBPD(model,states,statevalues)
% 
% model: SBmodel, ODE file model, or MEX file model
% states: Cell-array of state names for which to change the ICs
% statevalues: Vector with the corresponding values for the ICs of the states.
% 
% OUTPUT:
% =======
% icvector: Constructed initial condition vector

% Information:
% ============
% Copyright (C) 2005-2013 Henning Schmidt,
%
% This program is free software; you can redistribute it and/or modify
% it under the terms of the GNU General Public License as published by
% the Free Software Foundation; either version 2 of the License, or
% (at your option) any later version.
% 
% This program is distributed in the hope that it will be useful,
% but WITHOUT ANY WARRANTY; without even the implied warranty of
% MERCHANTABILITY or FITNESS FOR A PARTICULAR PURPOSE.  See the
% GNU General Public License for more details.
% 
% You should have received a copy of the GNU General Public License
% along with this program; if not, write to the Free Software
% Foundation, Inc., 59 Temple Place, Suite 330, Boston, MA  02111-1307  
% USA.


% Convert to cell-array if necessary
if ischar(states),
    states = {states};
end
% State input arguments need to have same length ...
if length(states) ~= length(statevalues),
    error('Different numbers of states and statevalues.');
end
% Get the initialconditions from the model
icvector = SBcalcICvector(model);
% Get all state names to check against the given state names
allstates = SBstates(model);
% Update the icvector with given values
for k = 1:length(states),
    index = strmatchSB(states{k},allstates,'exact');
    if isempty(index),
        error(sprintf('Statename ''%s'' is no state in the model.',states{k}));
    end
    icvector(index) = statevalues(k);
end
return
```
