## Supplementary scripts for "A multi-scale *in silico* mouse model for insulin resistance and humanoid type 2 diabetes": makeparamvecSBPD.m

```
function [paramvector] = makeparamvecSBPD(model,parameters,parametervalues)
% makeparamvecSBPD: The mex simulation functions take a full parameter value
% vector as an input. In order to construct such a vector while changing 
% some but possibly not all parameter values this function can be used.
% 
% USAGE:
% ======
% [paramvector] = makeparamvecSBPD(model,parameters,parametervalues)
% 
% model: SBmodel, ODE file model, or MEX file model
% parameters: Cell-array of parameter names for which to change the parametervalues
% parametervalues: Vector with the corresponding values for the parameters 
% 
% OUTPUT:
% =======
% paramvector: Constructed initial condition vector

% Information:
% ============
% Copyright (C) 2005-2013 Henning Schmidt,
%
% This program is free software; you can redistribute it and/or modify
% it under the terms of the GNU General Public License as published by
% the Free Software Foundation; either version 2 of the License, or
% (at your option) any later version.
% 
% This program is distributed in the hope that it will be useful,
% but WITHOUT ANY WARRANTY; without even the implied warranty of
% MERCHANTABILITY or FITNESS FOR A PARTICULAR PURPOSE.  See the
% GNU General Public License for more details.
% 
% You should have received a copy of the GNU General Public License
% along with this program; if not, write to the Free Software
% Foundation, Inc., 59 Temple Place, Suite 330, Boston, MA  02111-1307  
% USA.

% Convert to cell-array if necessary
if ischar(parameters),
    parameters = {parameters};
end
% Parameter input arguments need to have same length ...
if length(parameters) ~= length(parametervalues),
    error('Different numbers of parameters and parametervalues.');
end
% Get the parametervalues from the model
[allParams,paramvector] = SBparameters(model);
% Update the paramvector with given values
for k = 1:length(parameters),
    index = strmatchSB(parameters{k},allParams,'exact');
    if isempty(index),
        error(sprintf('Parameter name ''%s'' is no parameter in the model.',parameters{k}));
    end
    paramvector(index) = parametervalues(k);
end
return
```
