## Supplementary scripts for "A multi-scale *in silico* mouse model for insulin resistance and humanoid type 2 diabetes": makeTempMEXmodelSBPD.m

```
function [MEXmodel, MEXmodelfullpath] = makeTempMEXmodelSBPD(model)
% makeTempMEXmodelSBPD: This function can be used to create a temporary 
% MEX simulation function. Temporary means that it is created in the 
% systems temp directory and that no name can be chosen, but that a unique
% temporary filename is automatically chosen and returned.
% 
% USAGE:
% ======
% [MEXmodel, MEXmodelfullpath] = makeTempMEXmodelSBPD(model)
% 
% model: SBmodel to convert to a temporary MEX simulation function
% 
% OUTPUT ARGUMENTS:
% =================
% MEXmodel: name of the MEX simulation function (filename without .mex extension)
% MEXmodelfullpath: complete path to the created MEX simulation function (including mex extension)
%                   This information can be used to delete the function as
%                   soon as it is not needed anymore.

% Information:
% ============
% Copyright (C) 2005-2013 Henning Schmidt,
%
% This program is free software; you can redistribute it and/or modify
% it under the terms of the GNU General Public License as published by
% the Free Software Foundation; either version 2 of the License, or
% (at your option) any later version.
% 
% This program is distributed in the hope that it will be useful,
% but WITHOUT ANY WARRANTY; without even the implied warranty of
% MERCHANTABILITY or FITNESS FOR A PARTICULAR PURPOSE.  See the
% GNU General Public License for more details.
% 
% You should have received a copy of the GNU General Public License
% along with this program; if not, write to the Free Software
% Foundation, Inc., 59 Temple Place, Suite 330, Boston, MA  02111-1307  
% USA.

%%%%%%%%%%%%%%%%%%%%%%%%%%%%%%%%%%%%%%%%%%%%%%%
% TEMPDIR 
%%%%%%%%%%%%%%%%%%%%%%%%%%%%%%%%%%%%%%%%%%%%%%%
addpath(tempdirSB);   % add it to the path
returndir = pwd;      % save the current dir to return to it
cd(tempdirSB);        % change to tempdir

%%%%%%%%%%%%%%%%%%%%%%%%%%%%%%%%%%%%%%%%%%%%%%%
% CREATE THE MEX SIMULATION FUNCTION
%%%%%%%%%%%%%%%%%%%%%%%%%%%%%%%%%%%%%%%%%%%%%%%
% Unfortunately this strange while loop is required,
% since the compilation of MEX files using MinGW sometimes 
% produces invalid MEX files. It is so seldom that it does not matter
% to have this loop here. And after all a 3x speed increase in simulation
% is worth a lot :). Why does Mathworks use such a sh...y compiler?
notOk = 1;
warning off;
while notOk,
    try
        % GET TEMP FILE NAME
        tempfilefullpath = tempnameSB;
        [pathstr,tempfilename,ext] = fileparts(tempfilefullpath);
        SBPDmakeMEXmodel(model,tempfilename);
        feval(tempfilename,'parameters');
        notOk = 0;
    catch
        if ~isempty(strfind(lasterr,'Invalid MEX-file')),
            % disp('Nu isses wieder passiert');
            clear mex;
            delete([tempfilename,'.',mexext]);
            delete([tempfilename,'.c']);
            delete([tempfilename,'.h']);
            notOk = 1;
        elseif ~isempty(strfind(lasterr,'Error using ==> mex')),
            error(sprintf('MEX compilation of model failed. Please check the model syntax.\nThe output of the compiler above will guide you to the error.\ntp******.c is the .c file that showed the error.\nOpen it using ">> edit tp******.c" and look at the indicated lines.\nDo not correct it in the .c file but in the corresponding model\n(or experiment) description.'));
        else
            cd(returndir);
            error(lasterr); 
        end            
    end
end
warning on;

%%%%%%%%%%%%%%%%%%%%%%%%%%%%%%%%%%%%%%%%%%%%%%%
% RETURN TO THE PREVIOUS DIRECTORY AND CREATE OUTPUT
%%%%%%%%%%%%%%%%%%%%%%%%%%%%%%%%%%%%%%%%%%%%%%%
rehash;
cd(returndir);
MEXmodel = tempfilename;
MEXmodelfullpath = sprintf('%s.%s',tempfilefullpath,mexext);
```
