## Supplementary scripts for "A multi-scale *in silico* mouse model for insulin resistance and humanoid type 2 diabetes": mexcompileSBPD.m

```
function [] = mexcompileSBPD(filename)
% mexcompileSBPD: Compiles an SBmodel that has been converted to a mexfile.h 
% and mexfile.c and links it with the CVODE integrator from the SUNDIALS
% suite (http://www.llnl.gov/CASC/sundials/). 
% 
% USAGE:
% ======
% [] = mexcompileSBPD(mexfile)
%
% mexfile: Basename of the model files (mexfile.c and mexfile.h)  
    
% Information:
% ============
% Copyright (C) 2005-2013 Henning Schmidt,
%
% This program is free software; you can redistribute it and/or modify
% it under the terms of the GNU General Public License as published by
% the Free Software Foundation; either version 2 of the License, or
% (at your option) any later version.
% 
% This program is distributed in the hope that it will be useful,
% but WITHOUT ANY WARRANTY; without even the implied warranty of
% MERCHANTABILITY or FITNESS FOR A PARTICULAR PURPOSE.  See the
% GNU General Public License for more details.
% 
% You should have received a copy of the GNU General Public License
% along with this program; if not, write to the Free Software
% Foundation, Inc., 59 Temple Place, Suite 330, Boston, MA  02111-1307  
% USA.

if ~isunix,
    libpath = which('CVODEmex25.lib');
else
    libpath = which('CVODEmex25.a');
end
includefolder = fileparts(which('mexmathaddon.h'));
% do compilation
eval(sprintf('mex -O -I''%s'' %s.c ''%s'';',includefolder,filename,libpath));
```
