## Supplementary scripts for "A multi-scale *in silico* mouse model for insulin resistance and humanoid type 2 diabetes": updateparamvecSBPD.m

```
function [newparamvector] = updateparamvecSBPD(paramnames, oldparamvector, pnames, pvalues)
% updateparamvecSBPD: The mex simulation functions take a full parameter value
% vector as an input. In order to construct such a vector from an existing 
% parameter vector while changing some but possibly not all parameter
% values this function can be used. 
% 
% USAGE:
% ======
% [newparamvector] = updateparamvecSBPD(paramnames, oldparamvector, pnames, pvalues)
% 
% paramnames: Cell-array of all model parameter names (in the order as they
%   appear in the model)
% oldparamvector: Full parameter vector before the change
% pnames: Cell-array of parameter names for which to change the
%   parametervalues 
% pvalues: Vector with parameter values for the parameters to be changed
%   (order defined by pnames)
% 
% OUTPUT:
% =======
% newparamvector: Constructed new parameter vector with changed parameters

% Information:
% ============
% Copyright (C) 2005-2013 Henning Schmidt,
%
% This program is free software; you can redistribute it and/or modify
% it under the terms of the GNU General Public License as published by
% the Free Software Foundation; either version 2 of the License, or
% (at your option) any later version.
% 
% This program is distributed in the hope that it will be useful,
% but WITHOUT ANY WARRANTY; without even the implied warranty of
% MERCHANTABILITY or FITNESS FOR A PARTICULAR PURPOSE.  See the
% GNU General Public License for more details.
% 
% You should have received a copy of the GNU General Public License
% along with this program; if not, write to the Free Software
% Foundation, Inc., 59 Temple Place, Suite 330, Boston, MA  02111-1307  
% USA.

% Convert to cell-array if necessary
if ischar(pnames),
    pnames = {pnames};
end
% Parameter input arguments need to have same length ...
if length(pnames) ~= length(pvalues),
    error('Different numbers of parameters and parametervalues to be changed.');
end

% Update the paramvector with given values
newparamvector = oldparamvector;
for k = 1:length(pnames),
    index = strmatchSB(pnames{k}, paramnames, 'exact');
    if isempty(index),
        disp(sprintf('Parameter name ''%s'' is no parameter in the model.',pnames{k}));
    end
    newparamvector(index) = pvalues(k);
end
return
```
