## Supplementary scripts for "A multi-scale *in silico* mouse model for insulin resistance and humanoid type 2 diabetes": SBPDmakeMEXmodel.m

```
function [] = SBPDmakeMEXmodel(model,varargin)
% SBPDmakeMEXmodel: This function converts an SBmodel to an executable
% C-code
% MEX model and links it with the CVODE integrator from SUNDIALS.
% 
% USAGE:
% ======
% [] = SBPDmakeMEXmodel(model)
% [] = SBPDmakeMEXmodel(model, MEXfile)
% [] = SBPDmakeMEXmodel(model, MEXfile, doNOTcompileFlag)
%
% model: SBmodel
% MEXfile: the name of the MEX file (default: models name)
% doNOTcompileFlag: =0 (default): create executable MEX simulation file 
%                   =1: only create the source code but do not compile
%
% The calling syntax of the MEX simulation function is the following:
%           
%   output = MEXfile()                     % returns vector with initial conditions
%   output = MEXfile('states')             % returns cell array with state names
%   output = MEXfile('parameters')         % returns cell array with parameter names
%   output = MEXfile('parametervalues')    % returns vector with parameter values
%   output = MEXfile(timevector)
%   output = MEXfile(timevector, ICs)
%   output = MEXfile(timevector, ICs, param)
%   output = MEXfile(timevector, ICs, param, OPTIONS)
%   
%   INPUT ARGUMENTS TO 'MEXfile':
%   timevector: simulation time vector
%   ICs: full initial condition vector
%   param: full parametervalue vector
%   OPTIONS: structure with OPTIONS for the integrator
%       OPTIONS.showIntegratorStats: =0 (off), =1 shows integrator
%               statistics in the MATLAB console window
%       OPTIONS.method:             'stiff' or 'nonstiff' (default: 'stiff')
%       OPTIONS.abstol:             abs tolerance (default: 1e-6)
%       OPTIONS.reltol:             rel tolerance (default: 1e-6)
%       OPTIONS.minstep:            min step-size integrator (default: 0)
%       OPTIONS.maxstep:            max step-size integrator (default: inf)
%       OPTIONS.maxnumsteps:        max number of steps between two output
%                                   points (default: 100000)
%       OPTIONS.maxerrtestfails:    maximum number of error test failures
%                                   permitted in attempting one step
%                                   (default: 50)
%       OPTIONS.maxorder:           maximum order of the linear multistep
%                                   method (default: 5 BDF, 12 ADAMS)
%       OPTIONS.maxconvfails:       maximum number of nonlinear solver convergence 
%                                   failures permitted during one step
%                                   (default: 10)
%       OPTIONS.initstep:           initial step size to be attempted
%                                   (default: 0)
%       OPTIONS.maxnonlineariter:   maximum number of nonlinear solver
%                                   iterations permitted per step 
%                                   (default: 3)
%       OPTIONS.xdotcalc: =0: do integration, =1: return RHS of ODEs for
%                         given state and parameter values. Time information
%                         is neglected and it is assumed that time=0.
    
% Information:
% ============
% Copyright (C) 2005-2013 Henning Schmidt,
%
% This program is free software; you can redistribute it and/or modify
% it under the terms of the GNU General Public License as published by
% the Free Software Foundation; either version 2 of the License, or
% (at your option) any later version.
% 
% This program is distributed in the hope that it will be useful,
% but WITHOUT ANY WARRANTY; without even the implied warranty of
% MERCHANTABILITY or FITNESS FOR A PARTICULAR PURPOSE.  See the
% GNU General Public License for more details.
% 
% You should have received a copy of the GNU General Public License
% along with this program; if not, write to the Free Software
% Foundation, Inc., 59 Temple Place, Suite 330, Boston, MA  02111-1307  
% USA.

global MAX_NRPERROW 
MAX_NRPERROW = 20;  % Max. number of elements per rows

%%%%%%%%%%%%%%%%%%%%%%%%%%%%%%%%%%%%%%%%%%%%%%%
% GENERATE DELAY BASE NAME
%%%%%%%%%%%%%%%%%%%%%%%%%%%%%%%%%%%%%%%%%%%%%%%
global delaycount delaybase
delaycount = 0;
[dummy,delaybase] = fileparts(tempname);
delaybase = char([double('delaybase_') double(delaybase(1:8))]);

%%%%%%%%%%%%%%%%%%%%%%%%%%%%%%%%%%%%%%%%%%%%%%%
% GENERATE interpcseSB, etc. counters
%%%%%%%%%%%%%%%%%%%%%%%%%%%%%%%%%%%%%%%%%%%%%%%
global interpcseSBcount interpcseSlopeSBcount
interpcseSBcount = 0;
interpcseSlopeSBcount = 0;

%%%%%%%%%%%%%%%%%%%%%%%%%%%%%%%%%%%%%%%%%%%%%%%
% CHECK THE MODEL
%%%%%%%%%%%%%%%%%%%%%%%%%%%%%%%%%%%%%%%%%%%%%%%
if ~strcmp('SBmodel',class(model)),
    error('Function only defined for SBmodels.');
end
if ~isempty(SBfunctionsMATLAB(model)),
    error('Matlab functions present in model. Conversion to MEX simulation file not possible.');
end

%%%%%%%%%%%%%%%%%%%%%%%%%%%%%%%%%%%%%%%%%%%%%%%
% ADD PIECEWISE TRIGGERS AS EVENTS
%%%%%%%%%%%%%%%%%%%%%%%%%%%%%%%%%%%%%%%%%%%%%%%
model = addpiecewiseeventsSB(model);

%%%%%%%%%%%%%%%%%%%%%%%%%%%%%%%%%%%%%%%%%%%%%%%
% CHECK IF DELAY PRESENT ... THEN ERROR
%%%%%%%%%%%%%%%%%%%%%%%%%%%%%%%%%%%%%%%%%%%%%%%
if usedelaySB(model),
    error('The model contains delays. This is not supported for MEX file simulation.');
end

%%%%%%%%%%%%%%%%%%%%%%%%%%%%%%%%%%%%%%%%%%%%%%%
% GET THE UNDERLYING MODEL STRUCTURE
%%%%%%%%%%%%%%%%%%%%%%%%%%%%%%%%%%%%%%%%%%%%%%%
modelstruct = SBstruct(model);

%%%%%%%%%%%%%%%%%%%%%%%%%%%%%%%%%%%%%%%%%%%%%%%
% DEFAULT ARGUMENTS
%%%%%%%%%%%%%%%%%%%%%%%%%%%%%%%%%%%%%%%%%%%%%%%
MEXfile = regexprep(modelstruct.name,'[\W]','');  % name of model as default MEX function name
doNOTcompileFlag = 0;         % do compile per default

%%%%%%%%%%%%%%%%%%%%%%%%%%%%%%%%%%%%%%%%%%%%%%%
% TAKE CAR OF VARIABLE INPUT ARGUMENTS
%%%%%%%%%%%%%%%%%%%%%%%%%%%%%%%%%%%%%%%%%%%%%%%
if nargin < 1 || nargin > 3,
    error('Incorrect number of input arguments.');
end
if nargin == 2,
    MEXfile = varargin{1};
elseif nargin == 3,
    MEXfile = varargin{1};
    doNOTcompileFlag = varargin{2};
end

%%%%%%%%%%%%%%%%%%%%%%%%%%%%%%%%%%%%%%%%%%%%%%%
% HANDLE AN EVENTUAL FULL PATH (GET PATH AND THE NAME OF THE FUNCTION)
%%%%%%%%%%%%%%%%%%%%%%%%%%%%%%%%%%%%%%%%%%%%%%%
[MEXfunctionPath,MEXfunctionName] = fileparts(MEXfile);
filenameH = fullfile(MEXfunctionPath,strcat(MEXfunctionName,'.h'));
filenameC = fullfile(MEXfunctionPath,strcat(MEXfunctionName,'.c'));

%%%%%%%%%%%%%%%%%%%%%%%%%%%%%%%%%%%%%%%%%%%%%%%
% GET THE MODELS ELEMENTS
%%%%%%%%%%%%%%%%%%%%%%%%%%%%%%%%%%%%%%%%%%%%%%%
[staten,stateode,stateICs] = SBstates(model);
[parametern, parameterv] = SBparameters(model);
[variablen, variablef] = SBvariables(model);
[reactionn, reactionf] = SBreactions(model);
[functionn,functionf,functiona] = SBfunctions(model);
[eventn,eventt,eventv,eventf] = SBevents(model);

%%%%%%%%%%%%%%%%%%%%%%%%%%%%%%%%%%%%%%%%%%%%%%%
% GET NUMBER OF THE MODELS ELEMENTS
%%%%%%%%%%%%%%%%%%%%%%%%%%%%%%%%%%%%%%%%%%%%%%%
NRSTATES = length(staten);
NRPARAMETERS = length(parametern);
NRVARIABLES = length(variablen);
NRREACTIONS = length(reactionn);
NRFUNCTIONS = length(functionn);
NREVENTS = length(eventn);

%%%%%%%%%%%%%%%%%%%%%%%%%%%%%%%%%%%%%%%%%%%%%%%
% EXCHANGE SPECIAL NAMES
%%%%%%%%%%%%%%%%%%%%%%%%%%%%%%%%%%%%%%%%%%%%%%%
% Certain reserved element names might have to be exchanged. Example:
% "default", which is an often used name for the root compartment in SBML,
% but it is a reserved name in C.
staten = exchangeNames(staten);
parametern = exchangeNames(parametern); 
variablen = exchangeNames(variablen);
reactionn = exchangeNames(reactionn);
functionn = exchangeNames(functionn);
eventn = exchangeNames(eventn); 
eventt = exchangeNames(eventt);
for k=1:length(eventn),
    eventv{k} = exchangeNames(eventv{k});
end
functiona = exchangeNames(functiona);

%%%%%%%%%%%%%%%%%%%%%%%%%%%%%%%%%%%%%%%%%%%%%%%
% DEAL WITH THE FORMULAS
%%%%%%%%%%%%%%%%%%%%%%%%%%%%%%%%%%%%%%%%%%%%%%%
% For C the double representation needs to be made (1->1.0) ...
% Furthermore, several other things need to be fixed.
stateode = dealFormulas(stateode);
variablef = dealFormulas(variablef); 
reactionf = dealFormulas(reactionf); 
functionf = dealFormulas(functionf); 
eventt = dealFormulas(eventt);
for k = 1:length(eventf),
    eventf{k} = dealFormulas(eventf{k});
end

%%%%%%%%%%%%%%%%%%%%%%%%%%%%%%%%%%%%%%%%%%%%%%%%%%%%%%%%%%%%%%%%%%%%%%%%%%
% WRITE THE MODEL HEADER FILE
%%%%%%%%%%%%%%%%%%%%%%%%%%%%%%%%%%%%%%%%%%%%%%%%%%%%%%%%%%%%%%%%%%%%%%%%%%
fid = fopen(filenameH,'w');
fprintf(fid,'#include "mex.h"\n\n');
fprintf(fid,'const int NRSTATES = %d;\n',NRSTATES);
fprintf(fid,'const int NRPARAMETERS = %d;\n',NRPARAMETERS);
fprintf(fid,'const int NRVARIABLES = %d;\n',NRVARIABLES);
fprintf(fid,'const int NRREACTIONS = %d;\n',NRREACTIONS);
fprintf(fid,'const int NREVENTS = %d;\n',NREVENTS);
fprintf(fid,'\n');
% add flag for numeric or non-numeric ICs
if hasonlynumericICsSB(model),
    fprintf(fid,'const int hasOnlyNumericICs = 1;\n');
    outputHeaderData(fid,'defaultICs_num',NRSTATES,stateICs)
    fprintf(fid,'char *defaultICs_nonnum[1];\n');    
else
    fprintf(fid,'const int hasOnlyNumericICs = 0;\n');
    fprintf(fid,'double defaultICs_num[1];\n');    
    outputHeaderData(fid,'defaultICs_nonnum',NRSTATES,stateICs)
end
fprintf(fid,'\n');
if isempty(parameterv), parameterv = []; end
outputHeaderData(fid,'defaultParam',NRPARAMETERS,parameterv)
outputHeaderData(fid,'stateNames',NRSTATES,staten)
outputHeaderData(fid,'parameterNames',NRPARAMETERS,parametern)
outputHeaderData(fid,'variableNames',NRVARIABLES,variablen)
outputHeaderData(fid,'reactionNames',NRREACTIONS,reactionn)
outputHeaderData(fid,'eventNames',NREVENTS,eventn)
fprintf(fid,'\n');
fprintf(fid,'void model(double time, double *stateVector, double *DDTvector, ParamData *paramdataPtr, int DOflag, double *variableVector, double *reactionVector, double *gout, int *eventVector);\n');
fprintf(fid,'void calc_ic_model(double *icVector, ParamData *paramdataPtr);\n\n');
fprintf(fid,'void CVODEmex25(int nlhs, mxArray *plhs[], int nrhs, const mxArray *prhs[]);\n');
fprintf(fid,'void mexFunction(int nlhs, mxArray *plhs[], int nrhs, const mxArray *prhs[])\n');
fprintf(fid,'{\n    CVODEmex25(nlhs, plhs, nrhs, prhs);\n}\n');
fclose(fid);

%%%%%%%%%%%%%%%%%%%%%%%%%%%%%%%%%%%%%%%%%%%%%%%
% WRITE THE MODEL C FILE
%%%%%%%%%%%%%%%%%%%%%%%%%%%%%%%%%%%%%%%%%%%%%%%
fid = fopen(filenameC,'w');
fprintf(fid,'#include "stddef.h"\n');
fprintf(fid,'#include "stdarg.h"\n');
fprintf(fid,'#include "math.h"\n');
fprintf(fid,'#include "CVODEmex25.h"\n');
fprintf(fid,'#include "%s"\n',strcat(MEXfunctionName,'.h'));
fprintf(fid,'#include "mexsplineaddon.h"\n');
fprintf(fid,'#include "mexmathaddon.h"\n');
fprintf(fid,'#include "kineticformulas.h"\n\n');
fprintf(fid,'double time;\n\n');
% First define the functions
for k = 1:NRFUNCTIONS,
    % write declaration
    fprintf(fid,'static double %s(',functionn{k});
    % write arguments
    arguments = explodePCSB(functiona{k},','); for k2 = 1:length(arguments), if k2 < length(arguments), fprintf(fid,'double %s,',arguments{k2}); else fprintf(fid,'double %s)\n',arguments{k2}); end; end
    % write forumla and return
    fprintf(fid,'{\n'); fprintf(fid,'    return %s;\n',functionf{k}); fprintf(fid,'}\n'); fprintf(fid,'\n');
end
% Define the model function
fprintf(fid,'void model(double time_local, double *stateVector, double *DDTvector, ParamData *paramdataPtr, int DOflag, double *variableVector, double *reactionVector, double *gout, int *eventVector)\n');
fprintf(fid,'{\n');
outputDeclarationData(fid,NRSTATES,staten)
outputDeclarationData(fid,NRPARAMETERS,parametern)
outputDeclarationData(fid,NRVARIABLES,variablen)
outputDeclarationData(fid,NRREACTIONS,reactionn)
eventassignn = {};
for k=1:NREVENTS,
    eventassignformulak = eventf{k};
    for k2=1:length(eventassignformulak),
        eventassignn{end+1} = sprintf('eventassign_%d_%d',k,k2);
    end
end
outputDeclarationData(fid,length(eventassignn),eventassignn)
fprintf(fid,'\n');
fprintf(fid,'    time = time_local;\n');
fprintf(fid,'\n');
for k=1:NRSTATES, fprintf(fid,'    %s = stateVector[%d];\n',staten{k},k-1); end
for k=1:NRPARAMETERS, fprintf(fid,'    %s = paramdataPtr->parametervector[%d]; /* %g */\n',parametern{k},k-1,parameterv(k)); end
%for k=1:NRVARIABLES, fprintf(fid,'    %s = %s;\n',variablen{k},variablef{k}); end
%for k=1:NRREACTIONS, fprintf(fid,'    %s = %s;\n',reactionn{k},reactionf{k}); end
for k=1:NRVARIABLES, writeOutFormulasConvertPiecewise(fid,variablen{k},variablef{k},sprintf('\t')); end
for k=1:NRREACTIONS, writeOutFormulasConvertPiecewise(fid,reactionn{k},reactionf{k},sprintf('\t')); end
for k=1:NREVENTS,
    eventassignformulak = eventf{k};
    for k2=1:length(eventassignformulak),
        namevar = sprintf('eventassign_%d_%d',k,k2);
        fprintf(fid,'    %s = %s;\n',namevar,eventassignformulak{k2}); 
    end
end
fprintf(fid,'    if (DOflag == DOFLAG_DDT) {\n');
%for k=1:NRSTATES, fprintf(fid,'        DDTvector[%d] = %s;\n',k-1,stateode{k}); end
for k=1:NRSTATES, writeOutFormulasConvertPiecewise(fid,sprintf('\tDDTvector[%d]',k-1),stateode{k},sprintf('\t\t')); end
fprintf(fid,'    } else if (DOflag == DOFLAG_VARREAC) {\n');
for k=1:NRVARIABLES, fprintf(fid,'        variableVector[%d] = %s;\n',k-1,variablen{k}); end
for k=1:NRREACTIONS, fprintf(fid,'        reactionVector[%d] = %s;\n',k-1,reactionn{k}); end
fprintf(fid,'    } else if (DOflag == DOFLAG_EVENTS) {\n');
for k = 1:NREVENTS,
    tExpr = getTriggerExpression(eventt{k});
    fprintf(fid,'        gout[%d] = %s;\n',k-1,tExpr);
end
fprintf(fid,'    } else if (DOflag == DOFLAG_EVENTASSIGN) {\n');
for k = 1:NREVENTS,
    fprintf(fid,'        if (eventVector[%d] == 1 && gout[%d] < 0) {\n',k-1,k-1);
    fprintf(fid,'            DDTvector[0] = 1;\n');
    vars = eventv{k};
    for k2 = 1:length(vars),
        index = strmatchSB(vars{k2},staten,'exact')-1;
        if ~isempty(index),
            fprintf(fid,'            stateVector[%d] = eventassign_%d_%d;\n',index,k,k2);
        end
        index = strmatchSB(vars{k2},parametern,'exact')-1;
        if ~isempty(index),
            fprintf(fid,'            paramdataPtr->parametervector[%d] = eventassign_%d_%d;\n',index,k,k2);
        end
    end
    fprintf(fid,'        }\n');
end
fprintf(fid,'    }\n');
fprintf(fid,'}\n');
fprintf(fid,'\n');
fprintf(fid,'\n');
fprintf(fid,'/* Function for initial condition calculation */\n');
fprintf(fid,'void calc_ic_model(double *icVector, ParamData *paramdataPtr)\n');
fprintf(fid,'{\n');
outputDeclarationData(fid,NRSTATES,staten)
outputDeclarationData(fid,NRPARAMETERS,parametern)
for k=1:NRPARAMETERS, fprintf(fid,'    %s = paramdataPtr->parametervector[%d]; /* %g */\n',parametern{k},k-1,parameterv(k)); end
modelics = SBinitialconditions(model);
for k = 1:NRSTATES,
    if hasonlynumericICsSB(model),
        value = modelics(k);
    else
        value = modelics{k};
    end
    if isnumeric(value),
        value = sprintf('%g',value);
    end
    stateIC = dealFormulas({value});
    fprintf(fid,'    %s = %s;\n',staten{k},stateIC{1});
end
for k = 1:NRSTATES,
    fprintf(fid,'    icVector[%d] = %s;\n',k-1,staten{k});
end
fprintf(fid,'}\n');
fprintf(fid,'\n');
fclose(fid);

%%%%%%%%%%%%%%%%%%%%%%%%%%%%%%%%%%%%%%%%%%%%%%%
% COMPILE THE MEX MODEL
%%%%%%%%%%%%%%%%%%%%%%%%%%%%%%%%%%%%%%%%%%%%%%%
if ~doNOTcompileFlag,
    mexcompileSBPD(MEXfunctionName);
    delete(filenameC);
    delete(filenameH);
end

clear delaycount delaybase interpcseSBcount interpcseSlopeSBcount
return

%%%%%%%%%%%%%%%%%%%%%%%%%%%%%%%%%%%%%%%%%%%%%%%
% WRITING OUT FORMULAS AND REPLACING PIECEWISESB BY IF THEN ELSE
%%%%%%%%%%%%%%%%%%%%%%%%%%%%%%%%%%%%%%%%%%%%%%%
function [] = writeOutFormulasConvertPiecewise(fid,lhs,rhs,pretext)
% check if rhs starts with a piecewiseSB expression
rhs = strtrim(rhs);
% Search for both the normal and the T0 type of piecewise expressions
index1 = strfind(rhs,'piecewiseSB(');
index2 = strfind(rhs,'piecewiseT0SB(');
index = [index1(:)', index2(:)'];
if isempty(index),
    % not a piecewise expression
    fprintf(fid,'    %s = %s;\n',lhs,rhs); 
    return
elseif length(index) > 1,
    % more than one piecewise expression
    fprintf(fid,'    %s = %s;\n',lhs,rhs); 
    return
elseif index ~= 1,
    % does not start with a piecewise expression
    fprintf(fid,'    %s = %s;\n',lhs,rhs); 
    return
end
% At this point the rhs has a single piecewiseSB or piecewiseT0SB expression and starts with
% it. We only need to check if after the piecewiseSB expression there are
% other components in the rhs.
parOpen = 1;
if ~isempty(index1),
    indexStart = length('piecewiseSB(')+1; 
else
    indexStart = length('piecewiseT0SB(')+1; 
end    
indexEnd   = indexStart;
while parOpen ~= 0,
    indexEnd = indexEnd + 1;
    if rhs(indexEnd) == '(',
        parOpen = parOpen + 1;
    elseif rhs(indexEnd) == ')',
        parOpen = parOpen - 1;
    end
end
if ~isempty(rhs(indexEnd+1:end)),
    % piecewiseSB expression is not the single component on the RHS => not if then else
    fprintf(fid,'    %s = %s;\n',lhs,rhs); 
    return
end
% Finally we are left with only the rhs which only contain a single piecewise expression
% that easily can be represented in terms of if else if else if else ...
% Get only the arguments as comma separated list
arguments = rhs(indexStart:indexEnd-1);
% Explode to get the terms
terms = explodePCSB(arguments);
% First term gives the number of arguments. Needs to be an odd number,
% because otherwise the default value is missing and this is not good :)
% We still do not use the number ... since the length-1 gives the same
% result.
n = length(terms)-1;
% Check if n is odd 
if 2*round(n/2) == n,
    error('All piecewise expressions need an odd number of arguments to specify also a default value.');
end
% Construct the if statement and write it out
terms = terms(2:end);
for k=2:2:n-1,
    if k==2,
        fprintf(fid,'%sif (%s) {\n',pretext,terms{k});
    else
        fprintf(fid,'%s} else if (%s) {\n',pretext,terms{k});
    end
    fprintf(fid,'%s\t%s = %s;\n',pretext,lhs,terms{k-1});
end
fprintf(fid,'%s} else {\n',pretext);
fprintf(fid,'%s\t%s = %s;\n%s}\n',pretext,lhs,terms{n},pretext);
return


%%%%%%%%%%%%%%%%%%%%%%%%%%%%%%%%%%%%%%%%%%%%%%%
% DETERMINE THE TRIGGER EXPRESSION
%%%%%%%%%%%%%%%%%%%%%%%%%%%%%%%%%%%%%%%%%%%%%%%
function [tExpr] = getTriggerExpression(data)
tExpr = sprintf('%s-0.5',data);
return

%%%%%%%%%%%%%%%%%%%%%%%%%%%%%%%%%%%%%%%%%%%%%%%
% OUTPUT HEADER DATA
%%%%%%%%%%%%%%%%%%%%%%%%%%%%%%%%%%%%%%%%%%%%%%%
function [] = outputHeaderData(fid,name,NR,data)
global MAX_NRPERROW 
if NR == 0,
    if iscell(data),
        fprintf(fid,'char *%s[1];\n',name);
    else
        fprintf(fid,'double %s[1];\n',name);
    end
else
    if iscell(data),
        fprintf(fid,'char *%s[%d] = {\n\t',name,NR);
    else
        fprintf(fid,'double %s[%d] = {\n\t',name,NR);
    end        
    nrperrow = 0;
    for k=1:NR,
        if nrperrow == MAX_NRPERROW, fprintf(fid,'\n\t'); nrperrow = 0; end
        if iscell(data),   
            if ~isnumeric(data{k}),
                if k<NR, fprintf(fid,'"%s",',data{k}); else fprintf(fid,'"%s"',data{k}); end
            else
                % handling numeric entries in cell-arrays: happening if
                % non-numeric ICs
                if k<NR, fprintf(fid,'"%g",',data{k}); else fprintf(fid,'"%g"',data{k}); end
            end
        else
            if k<NR, fprintf(fid,'%g,',data(k)); else fprintf(fid,'%g',data(k)); end
        end
        nrperrow = nrperrow + 1;
    end
    fprintf(fid,'};\n');
end

%%%%%%%%%%%%%%%%%%%%%%%%%%%%%%%%%%%%%%%%%%%%%%%
% OUTPUT DECLARATION DATA
%%%%%%%%%%%%%%%%%%%%%%%%%%%%%%%%%%%%%%%%%%%%%%%
function [] = outputDeclarationData(fid,NR,data)
global MAX_NRPERROW 
nrperrow = 0;
for k=1:NR,
    if nrperrow==0, fprintf(fid,'    double '); end
    if k<NR && nrperrow<MAX_NRPERROW-1, fprintf(fid,'%s,',data{k}); else fprintf(fid,'%s',data{k}); end
    nrperrow = nrperrow + 1;
    if nrperrow == MAX_NRPERROW, fprintf(fid,';\n'); nrperrow = 0; end
end
if nrperrow ~= 0, fprintf(fid,';\n'); end
return

%%%%%%%%%%%%%%%%%%%%%%%%%%%%%%%%%%%%%%%%%%%%%%%%%%%%%%%%%%%%%%%%%%%%%%%%%%
% EXCHANGE THE NAMES
%%%%%%%%%%%%%%%%%%%%%%%%%%%%%%%%%%%%%%%%%%%%%%%%%%%%%%%%%%%%%%%%%%%%%%%%%%
function [names] = exchangeNames(names)
if isempty(names), return; end
% do not allow "default" being used as component name!!!
if ~isempty(strmatchSB('default',names,'exact')),
    error(sprintf('The name "default" is not allowed to be used in an SBmodel when you want\nto create MEX simulation functions. The reason is that "default" is a reserved\n"C" word. Of course this is the case also for all other reserved "C" words.\nPlease rename the "default" component in your model with another name!\n'));
end
oldText = {}; 
newText = {};
names = regexprep(names,oldText,newText);
return

%%%%%%%%%%%%%%%%%%%%%%%%%%%%%%%%%%%%%%%%%%%%%%%%%%%%%%%%%%%%%%%%%%%%%%%%%%
% HANDLE INTERP1SB (LOOKUP TABLE W/ LINEAR INTERPOLATION)
%%%%%%%%%%%%%%%%%%%%%%%%%%%%%%%%%%%%%%%%%%%%%%%%%%%%%%%%%%%%%%%%%%%%%%%%%%
function [textnew] = exchangeInterp1SB(text)
textnew = text;
indexstart = strfind(text,'interp1SB(')+length('interp1SB(');
if length(indexstart) > 1,
    error('The ''interp1SB'' function is only allowed to be present once in each formula.');
end
if ~isempty(indexstart),
    % interp1SB found ... handle it
    % cut out the content using the parentheses
    % Get the end index of the k-th statement (closing parentheses
    % belonging to the opening)
    % count parentheses
    pc = 1;
    cstart = indexstart;
    cend = cstart;
    while pc ~= 0,
        cend = cend + 1;
        if textnew(cend) == '(',
            pc = pc+1;
        elseif textnew(cend) == ')',
            pc = pc-1;
        end
    end
    indexend = cend-1;
    indexafter = indexend+1;
    % indexstart/indexend identify the content in the parentheses to be
    % processed and replaced
    textinside = textnew(indexstart:indexend);
    terms = explodePCSB(textinside,',',{'(','['},{')',']'});
    % fine so far. We need now to make sure that the elements in the
    % vectors are separated using commata (otherwise big problem)!
    xtermstring = terms{1};
    ytermstring = terms{2};
    % remove parentheses
    xtermstring = xtermstring(2:end-1);
    ytermstring = ytermstring(2:end-1);
    % get single elements
    xtermelements = explodePCSB(xtermstring);
    ytermelements = explodePCSB(ytermstring);
    if length(xtermelements) == 1,
        error(sprintf('The elements of the lookup table using the interp1SB function need to be separated by commata!\nAre you sure you have done that correctly?'));
    end
    if length(xtermelements) ~= length(ytermelements),
        error('x and y arguments for interp1SB function do not have same number of elements.');
    end
    pwText = sprintf('%s,lt(%s,%s),',ytermelements{1},terms{3},xtermelements{1});
    for k=1:length(xtermelements)-1,
        pwText = sprintf('%s(%s-(%s))/(%s-(%s))*(%s-(%s))+(%s)',pwText,ytermelements{k+1},ytermelements{k},xtermelements{k+1},xtermelements{k},terms{3},xtermelements{k},ytermelements{k});
        if k<length(xtermelements)-1,
            pwText = sprintf('%s,andSB(lt(%s,%s),ge(%s,%s)),',pwText,terms{3},xtermelements{k+1},terms{3},xtermelements{k});
        end
    end
    pwText = sprintf('%s,andSB(lt(%s,%s),ge(%s,%s)),%s',pwText,terms{3},xtermelements{end},terms{3},xtermelements{end-1},ytermelements{end});
    textnew = [text(1:indexstart-1) pwText text(indexend+1:end)];
    textnew = strrep(textnew,'interp1SB','piecewiseSB');
end
return

%%%%%%%%%%%%%%%%%%%%%%%%%%%%%%%%%%%%%%%%%%%%%%%%%%%%%%%%%%%%%%%%%%%%%%%%%%
% HANDLE INTERP0SB (LOOKUP TABLE W/ ZERO-ORDER INTERPOLATION)
%%%%%%%%%%%%%%%%%%%%%%%%%%%%%%%%%%%%%%%%%%%%%%%%%%%%%%%%%%%%%%%%%%%%%%%%%%
function [textnew] = exchangeInterp0SB(text)
textnew = text;
indexstart = strfind(text,'interp0SB(')+length('interp0SB(');
if length(indexstart) > 1,
    error('The ''interp0SB'' function is only allowed to be present once in each formula.');
end
if ~isempty(indexstart),
    % interp0SB found ... handle it
    % cut out the content using the parentheses
    % Get the end index of the k-th statement (closing parentheses
    % belonging to the opening)
    % count parentheses
    pc = 1;
    cstart = indexstart;
    cend = cstart;
    while pc ~= 0,
        cend = cend + 1;
        if textnew(cend) == '(',
            pc = pc+1;
        elseif textnew(cend) == ')',
            pc = pc-1;
        end
    end
    indexend = cend-1;
    indexafter = indexend+1;
    % indexstart/indexend identify the content in the parentheses to be
    % processed and replaced
    textinside = textnew(indexstart:indexend);
    terms = explodePCSB(textinside,',',{'(','['},{')',']'});
    % fine so far. We need now to make sure that the elements in the
    % vectors are separated using commata (otherwise big problem)!
    xtermstring = terms{1};
    ytermstring = terms{2};
    % remove parentheses
    xtermstring = xtermstring(2:end-1);
    ytermstring = ytermstring(2:end-1);
    % get single elements
    xtermelements = explodePCSB(xtermstring);
    ytermelements = explodePCSB(ytermstring);
    if length(xtermelements) == 1,
        error(sprintf('The elements of the lookup table using the interp0SB function need to be separated by commata!\nAre you sure you have done that correctly?'));
    end
    if length(xtermelements) == 2,
        error('The interp0SB function requires at least 3 points on the x and y axis.');
    end
    if length(xtermelements) ~= length(ytermelements),
        error('x and y arguments for interp0SB function do not have same number of elements.');
    end
    pwText = sprintf('%s,lt(%s,%s),',ytermelements{1},terms{3},xtermelements{2});
    for k=2:length(xtermelements)-1,
        pwText = sprintf('%s(%s)',pwText,ytermelements{k});
        if k<length(xtermelements)-1,
            pwText = sprintf('%s,andSB(lt(%s,%s),ge(%s,%s)),',pwText,terms{3},xtermelements{k+1},terms{3},xtermelements{k});
        end
    end
    pwText = sprintf('%s,andSB(lt(%s,%s),ge(%s,%s)),%s',pwText,terms{3},xtermelements{end},terms{3},xtermelements{end-1},ytermelements{end});
    textnew = [text(1:indexstart-1) pwText text(indexend+1:end)];
    textnew = strrep(textnew,'interp0SB','piecewiseSB');
end
return

%%%%%%%%%%%%%%%%%%%%%%%%%%%%%%%%%%%%%%%%%%%%%%%%%%%%%%%%%%%%%%%%%%%%%%%%%%
% HANDLE INTERPCSSB (LOOKUP TABLE W/ CUBIC SPLINE INTERPOLATION)
%%%%%%%%%%%%%%%%%%%%%%%%%%%%%%%%%%%%%%%%%%%%%%%%%%%%%%%%%%%%%%%%%%%%%%%%%%
function [textnew] = exchangeInterpcsSB(text)
textnew = text;
indexbefore = strfind(text,'interpcsSB(');
indexstart = indexbefore+length('interpcsSB(');
if length(indexstart) > 1,
    error('The ''interpcsSB'' function is only allowed to be present once in each formula.');
end
if ~isempty(indexstart),
    % interpcsSB found ... handle it
    % cut out the content using the parentheses
    % Get the end index of the k-th statement (closing parentheses
    % belonging to the opening)
    % count parentheses
    pc = 1;
    cstart = indexstart;
    cend = cstart;
    while pc ~= 0,
        cend = cend + 1;
        if textnew(cend) == '(',
            pc = pc+1;
        elseif textnew(cend) == ')',
            pc = pc-1;
        end
    end
    indexend = cend-1;
    indexafter = indexend+1;
    % indexstart/indexend identify the content in the parentheses to be
    % processed and replaced
    textinside = textnew(indexstart:indexend);
    terms = explodePCSB(textinside,',',{'(','['},{')',']'});
    if length(terms) ~= 3, 
        error('interpcsSB function has wrong number of input arguments.');
    end
    % fine so far. We need now to make sure that the elements in the
    % vectors are separated using commata (otherwise big problem)!
    xtermstring = terms{1};
    ytermstring = terms{2};
    % remove parentheses
    xtermstring = xtermstring(2:end-1);
    ytermstring = ytermstring(2:end-1);
    % get single elements
    xtermelements = explodePCSB(xtermstring);
    ytermelements = explodePCSB(ytermstring);
    if length(xtermelements) == 1,
        error(sprintf('The elements of the lookup table using the interpcsSB function need to be separated by commata!\nAre you sure you have done that correctly?'));
    end
    if length(xtermelements) ~= length(ytermelements),
        error('x and y arguments for interpcsSB function do not have same number of elements.');
    end
    newexpr = sprintf('%d,%s',length(xtermelements),terms{3});
    newexpr = strcat('interpcsSB(',newexpr,',',xtermstring,',',ytermstring,')');
    textnew = strcat(text(1:indexbefore-1),newexpr,text(indexend+2:end));
end
return

%%%%%%%%%%%%%%%%%%%%%%%%%%%%%%%%%%%%%%%%%%%%%%%%%%%%%%%%%%%%%%%%%%%%%%%%%%
% HANDLE INTERPCSeSB (LOOKUP TABLE W/ CUBIC SPLINE INTERPOLATION and
% possible endpoint-slope definition
%%%%%%%%%%%%%%%%%%%%%%%%%%%%%%%%%%%%%%%%%%%%%%%%%%%%%%%%%%%%%%%%%%%%%%%%%%
function [textnew] = exchangeInterpcseSB(text)
global interpcseSBcount

% Determine the number of required replacements (number of interpcseSB functions)
nrRep = length(strfind(text,'interpcseSB('));
% Create new text string from input string
textnew = text;

% Replace on after the other expression sequentially
% if interpcseSB not present then nothing is done ... simple
for k=1:nrRep,
    % Find start of interpcseSB in modified string
    indexbefore = strfind(textnew,'interpcseSB(');
    % Get the start index for modification for the k-th modification
    indexbefore = indexbefore(k);
    indexstart = indexbefore+length('interpcseSB(');
    % Get the end index of the k-th statement (closing parentheses
    % belonging to the opening)
    % count parentheses
    pc = 1;
    cstart = indexstart;
    cend = cstart;
    while pc ~= 0,
        cend = cend + 1;
        if textnew(cend) == '(',
            pc = pc+1;
        elseif textnew(cend) == ')',
            pc = pc-1;
        end
    end
    indexend = cend-1;
    indexafter = indexend+1;
    % indexstart/indexend identify the content in the parentheses to be
    % processed and replaced
    textinside = textnew(indexstart:indexend);
    terms = explodePCSB(textinside,',',{'(','['},{')',']'});
    nterms = length(terms);
    if nterms ~= 3 && nterms ~= 5, 
        error('interpcseSB function has wrong number of input arguments.');
    end
    % fine so far. We need now to make sure that the elements in the
    % vectors are separated using commata (otherwise big problem)!
    xtermstring = terms{1};
    ytermstring = terms{2};
    % remove parentheses
    xtermstring = xtermstring(2:end-1);
    ytermstring = ytermstring(2:end-1);
    % get single elements
    xtermelements = explodePCSB(xtermstring);
    ytermelements = explodePCSB(ytermstring);
    if length(xtermelements) == 1,
        error(sprintf('The elements of the lookup table using the interpcseSB function need to be separated by commata!\nAre you sure you have done that correctly?'));
    end
    if length(xtermelements) ~= length(ytermelements),
        error('x and y arguments for interpcseSB function do not have same number of elements.');
    end
    % construct the text to replace with
    if nterms == 3,
        reptextnew = [sprintf('interpcseSB(%d,%d',interpcseSBcount,length(xtermelements)),',',terms{3} ',',xtermstring,',',ytermstring,')'];
    else
        reptextnew = [sprintf('interpcseSB(%d,%d',interpcseSBcount,length(xtermelements)),',',terms{3},',',terms{4},',',terms{5} ',',xtermstring,',',ytermstring,')'];
    end
    % get the text to replace
    reptextold = textnew(indexbefore:indexafter);
    % do a simple strrep operation
    textnew = strrep(textnew,reptextold,reptextnew);
    % increment the interpcseSB counter
    interpcseSBcount = interpcseSBcount + 1;
end
return

%%%%%%%%%%%%%%%%%%%%%%%%%%%%%%%%%%%%%%%%%%%%%%%%%%%%%%%%%%%%%%%%%%%%%%%%%%
% HANDLE INTERPCSeSlopeSB (LOOKUP TABLE W/ CUBIC SPLINE INTERPOLATION and
% possible endpoint-slope definition - DERIVATIVE)
%%%%%%%%%%%%%%%%%%%%%%%%%%%%%%%%%%%%%%%%%%%%%%%%%%%%%%%%%%%%%%%%%%%%%%%%%%
function [textnew] = exchangeInterpcseSlopeSB(text)
global interpcseSlopeSBcount

% Determine the number of required replacements (number of interpcseSlopeSB functions)
nrRep = length(strfind(text,'interpcseSlopeSB('));
% Create new text string from input string
textnew = text;

% Replace on after the other expression sequentially
% if interpcseSlopeSB not present then nothing is done ... simple
for k=1:nrRep,
    % Find start of interpcseSlopeSB in modified string
    indexbefore = strfind(textnew,'interpcseSlopeSB(');
    % Get the start index for modification for the k-th modification
    indexbefore = indexbefore(k);
    indexstart = indexbefore+length('interpcseSlopeSB(');
    % Get the end index of the k-th statement (closing parentheses
    % belonging to the opening)
    % count parentheses
    pc = 1;
    cstart = indexstart;
    cend = cstart;
    while pc ~= 0,
        cend = cend + 1;
        if textnew(cend) == '(',
            pc = pc+1;
        elseif textnew(cend) == ')',
            pc = pc-1;
        end
    end
    indexend = cend-1;
    indexafter = indexend+1;
    % indexstart/indexend identify the content in the parentheses to be
    % processed and replaced
    textinside = textnew(indexstart:indexend);
    terms = explodePCSB(textinside,',',{'(','['},{')',']'});
    nterms = length(terms);
    if nterms ~= 3 && nterms ~= 5, 
        error('interpcseSlopeSB function has wrong number of input arguments.');
    end
    % fine so far. We need now to make sure that the elements in the
    % vectors are separated using commata (otherwise big problem)!
    xtermstring = terms{1};
    ytermstring = terms{2};
    % remove parentheses
    xtermstring = xtermstring(2:end-1);
    ytermstring = ytermstring(2:end-1);
    % get single elements
    xtermelements = explodePCSB(xtermstring);
    ytermelements = explodePCSB(ytermstring);
    if length(xtermelements) == 1,
        error(sprintf('The elements of the lookup table using the interpcseSlopeSB function need to be separated by commata!\nAre you sure you have done that correctly?'));
    end
    if length(xtermelements) ~= length(ytermelements),
        error('x and y arguments for interpcseSlopeSB function do not have same number of elements.');
    end
    % construct the text to replace with
    if nterms == 3,
        reptextnew = [sprintf('interpcseSlopeSB(%d,%d',interpcseSlopeSBcount,length(xtermelements)),',',terms{3} ',',xtermstring,',',ytermstring,')'];
    else
        reptextnew = [sprintf('interpcseSlopeSB(%d,%d',interpcseSlopeSBcount,length(xtermelements)),',',terms{3},',',terms{4},',',terms{5} ',',xtermstring,',',ytermstring,')'];
    end
    % get the text to replace
    reptextold = textnew(indexbefore:indexafter);
    % do a simple strrep operation
    textnew = strrep(textnew,reptextold,reptextnew);
    % increment the interpcseSlopeSB counter
    interpcseSlopeSBcount = interpcseSlopeSBcount + 1;
end
return

%%%%%%%%%%%%%%%%%%%%%%%%%%%%%%%%%%%%%%%%%%%%%%%%%%%%%%%%%%%%%%%%%%%%%%%%%%
% HANDLE INTERPCSexSB (LOOKUP TABLE W/ CUBIC SPLINE INTERPOLATION and
% possible endpoint-slope definition
%%%%%%%%%%%%%%%%%%%%%%%%%%%%%%%%%%%%%%%%%%%%%%%%%%%%%%%%%%%%%%%%%%%%%%%%%%
function [textnew] = exchangeInterpcsexSB(text)
% Determine the number of required replacements (number of interpcsexSB functions)
nrRep = length(strfind(text,'interpcsexSB('));
% Create new text string from input string
textnew = text;

% Replace on after the other expression sequentially
% if interpcsexSB not present then nothing is done ... simple
for k=1:nrRep,
    % Find start of interpcseSB in modified string
    indexbefore = strfind(textnew,'interpcsexSB(');
    % Get the start index for modification for the k-th modification
    indexbefore = indexbefore(k);
    indexstart = indexbefore+length('interpcsexSB(');
    % Get the end index of the k-th statement (closing parentheses
    % belonging to the opening)
    % count parentheses
    pc = 1;
    cstart = indexstart;
    cend = cstart;
    while pc ~= 0,
        cend = cend + 1;
        if textnew(cend) == '(',
            pc = pc+1;
        elseif textnew(cend) == ')',
            pc = pc-1;
        end
    end
    indexend = cend-1;
    indexafter = indexend+1;
    % indexstart/indexend identify the content in the parentheses to be
    % processed and replaced
    textinside = textnew(indexstart:indexend);
    terms = explodePCSB(textinside,',',{'(','['},{')',']'});
    nterms = length(terms);
    if nterms ~= 3 && nterms ~= 5, 
        error('interpcsexSB function has wrong number of input arguments.');
    end
    % fine so far. We need now to make sure that the elements in the
    % vectors are separated using commata (otherwise big problem)!
    xtermstring = terms{1};
    ytermstring = terms{2};
    % remove parentheses
    xtermstring = xtermstring(2:end-1);
    ytermstring = ytermstring(2:end-1);
    % get single elements
    xtermelements = explodePCSB(xtermstring);
    ytermelements = explodePCSB(ytermstring);
    if length(xtermelements) == 1,
        error(sprintf('The elements of the lookup table using the interpcsexSB function need to be separated by commata!\nAre you sure you have done that correctly?'));
    end
    if length(xtermelements) ~= length(ytermelements),
        error('x and y arguments for interpcsexSB function do not have same number of elements.');
    end
    % construct the text to replace with
    if nterms == 3,
        reptextnew = [sprintf('interpcsexSB(%d',length(xtermelements)) ',',xtermstring,',',ytermstring,',',terms{3},')'];
    else
        reptextnew = [sprintf('interpcsexSB(%d',length(xtermelements)),',',terms{4},',',terms{5} ',',xtermstring,',',ytermstring,',',terms{3},')'];
    end
    % get the text to replace
    reptextold = textnew(indexbefore:indexafter);
    % do a simple strrep operation
    textnew = strrep(textnew,reptextold,reptextnew);
end
return

%%%%%%%%%%%%%%%%%%%%%%%%%%%%%%%%%%%%%%%%%%%%%%%%%%%%%%%%%%%%%%%%%%%%%%%%%%
% HANDLE INTERPCSexSB (LOOKUP TABLE W/ CUBIC SPLINE INTERPOLATION and
% possible endpoint-slope definition
%%%%%%%%%%%%%%%%%%%%%%%%%%%%%%%%%%%%%%%%%%%%%%%%%%%%%%%%%%%%%%%%%%%%%%%%%%
function [textnew] = exchangeInterpcsexSlopeSB(text)
% Determine the number of required replacements (number of interpcsexSlopeSB functions)
nrRep = length(strfind(text,'interpcsexSlopeSB('));
% Create new text string from input string
textnew = text;

% Replace on after the other expression sequentially
% if interpcsexSlopeSB not present then nothing is done ... simple
for k=1:nrRep,
    % Find start of interpcsexSlopeSB in modified string
    indexbefore = strfind(textnew,'interpcsexSlopeSB(');
    % Get the start index for modification for the k-th modification
    indexbefore = indexbefore(k);
    indexstart = indexbefore+length('interpcsexSlopeSB(');
    % Get the end index of the k-th statement (closing parentheses
    % belonging to the opening)
    % count parentheses
    pc = 1;
    cstart = indexstart;
    cend = cstart;
    while pc ~= 0,
        cend = cend + 1;
        if textnew(cend) == '(',
            pc = pc+1;
        elseif textnew(cend) == ')',
            pc = pc-1;
        end
    end
    indexend = cend-1;
    indexafter = indexend+1;
    % indexstart/indexend identify the content in the parentheses to be
    % processed and replaced
    textinside = textnew(indexstart:indexend);
    terms = explodePCSB(textinside,',',{'(','['},{')',']'});
    nterms = length(terms);
    if nterms ~= 3 && nterms ~= 5, 
        error('interpcsexSlopeSB function has wrong number of input arguments.');
    end
    % fine so far. We need now to make sure that the elements in the
    % vectors are separated using commata (otherwise big problem)!
    xtermstring = terms{1};
    ytermstring = terms{2};
    % remove parentheses
    xtermstring = xtermstring(2:end-1);
    ytermstring = ytermstring(2:end-1);
    % get single elements
    xtermelements = explodePCSB(xtermstring);
    ytermelements = explodePCSB(ytermstring);
    if length(xtermelements) == 1,
        error(sprintf('The elements of the lookup table using the interpcsexSlopeSB function need to be separated by commata!\nAre you sure you have done that correctly?'));
    end
    if length(xtermelements) ~= length(ytermelements),
        error('x and y arguments for interpcsexSlopeSB function do not have same number of elements.');
    end
    % construct the text to replace with
    if nterms == 3,
        reptextnew = [sprintf('interpcsexSlopeSB(%d',length(xtermelements)) ',',xtermstring,',',ytermstring,',',terms{3},')'];
    else
        reptextnew = [sprintf('interpcsexSlopeSB(%d',length(xtermelements)),',',terms{4},',',terms{5} ',',xtermstring,',',ytermstring,',',terms{3},')'];
    end
    % get the text to replace
    reptextold = textnew(indexbefore:indexafter);
    % do a simple strrep operation
    textnew = strrep(textnew,reptextold,reptextnew);
end
return

%%%%%%%%%%%%%%%%%%%%%%%%%%%%%%%%%%%%%%%%%%%%%%%%%%%%%%%%%%%%%%%%%%%%%%%%%%
% DEAL WITH THE FORMULAS
%%%%%%%%%%%%%%%%%%%%%%%%%%%%%%%%%%%%%%%%%%%%%%%%%%%%%%%%%%%%%%%%%%%%%%%%%%
function [newformulaArray] = dealFormulas(formulaArray)
if isempty(formulaArray),
    newformulaArray = formulaArray;
    return
end

% replace names first (and we do it once more after the handling of the
% formulas ... at least for power that is necessary)
oldElements = {'\<nthroot\>','\<and\>','\<or\>','\<power\>','\<abs\>'};
newElements = {'nthrootSB','andSB','orSB','pow','absSB'};
formulaArray = regexprep(formulaArray,oldElements,newElements);

% handle calls with variable number of input arguments (these get the
% number of input arguments as first (additional) argument).
checkElements = {'\<indexmaxSB\>','\<minSB\>','\<maxSB\>','\<andSB\>','\<orSB\>','\<piecewiseSB\>','\<interpcsSB\>','\<interpcseSB\>','\<interpcsexSB\>','\<piecewiseT0SB\>','\<interpcseSlopeSB\>','\<interpcsexSlopeSB\>'};
for k0=1:length(formulaArray),
    formula = formulaArray{k0};
    % handle interp1SB -> piecewiseSB
    formula = exchangeInterp1SB(formula);
    % handle interp0SB -> piecewiseSB
    formula = exchangeInterp0SB(formula);
    % handle interpcsSB: changing the syntax
    formula = exchangeInterpcsSB(formula);
    % handle interpcseSB: changing the syntax
    formula = exchangeInterpcseSB(formula);
    % handle interpcseSlopeSB: changing the syntax
    formula = exchangeInterpcseSlopeSB(formula);
    % handle interpcsexSB: changing the syntax
    formula = exchangeInterpcsexSB(formula);
    % handle interpcsexSlopeSB: changing the syntax
    formula = exchangeInterpcsexSlopeSB(formula);
    % handle the power operator
    formula = convertPowerOperator(formula);
    % handle delay operator
%    [formula] = processDelayCallSB(formula);
    % fix the c notation of doubles
    formula = regexprep(formula,'((?<!\d*[eE][-+])(?<!\.)\<\d*\>(?!\.))','$1.0');
    % delete "." at end
    formula = regexprep(formula,'([\w])+[.]{1}([\W])','$1$2');
    formula = regexprep(formula,'([\W])+[.]{1}([\W])','$1$2');
    % Add number of variable input arguments to function calls as first
    % input argument (for the functions, defined above)
    % Add number of variable input arguments to function calls as first
    % input argument (for the functions, defined above)
    for k1=1:length(checkElements),
        index = regexp(formula,checkElements{k1});
        if ~isempty(index),
            for k2 = 1:length(index),
                indexStart = index(k2)+length(checkElements{k1})-4;
                indexEnd = indexStart;
                parOpen = 1;
                while parOpen ~= 0,
                    indexEnd = indexEnd + 1;
                    if formula(indexEnd) == '(',
                        parOpen = parOpen + 1;
                    elseif formula(indexEnd) == ')',
                        parOpen = parOpen - 1;
                    end
                end
                oldarguments = formula(indexStart+1:indexEnd-1);
                newargstring = sprintf('%d,%s',length(explodePCSB(oldarguments,',')),oldarguments);
                command = checkElements{k1};
                oldrep  = [command(3:end-2) '(' oldarguments ')'];
                newrep = [command(3:end-2) '(' newargstring ')'];
                formula = strrep(formula,oldrep,newrep);
                index = index + length(newargstring)-length(oldarguments);
            end
        end
    end
    formulaArray{k0} = formula;
end

% replace names again
oldElements = {'\<nthroot\>','\<and\>','\<or\>','\<power\>','\<abs\>'};
newElements = {'nthrootSB','andSB','orSB','pow','absSB'};
formulaArray = regexprep(formulaArray,oldElements,newElements);

% ready
newformulaArray = formulaArray;
return

%%%%%%%%%%%%%%%%%%%%%%%%%%%%%%%%%%%%%%%%%%%%%%%%%%%%%%%%%%%%%%%%%%%%%%%%%%
% HANDLE THE POWER OPERATOR
%%%%%%%%%%%%%%%%%%%%%%%%%%%%%%%%%%%%%%%%%%%%%%%%%%%%%%%%%%%%%%%%%%%%%%%%%%
function [formula] = convertPowerOperator(formula)
% remove whitespaces from formula
formula = regexprep(formula,'\s','');
% first fix the simple problem
formula = regexprep(formula,'([\w]+[.]?[\w]*)\^([\w]+[.]?[\w]*)','power($1,$2)');
% then do the more complicated stuff
indices = strfind(formula,'^');
while ~isempty(indices),
    index = indices(1);
    formula1 = strtrim(formula(1:index-1));
    formula2 = strtrim(formula(index+1:end));
    % check formula1 from the right
    firstargument = regexp(formula1,'([\w]+[.]?[\w]*)$','match');
    if isempty(firstargument),
        % check if last character is a closing parenthesis
        if formula1(end) ~= ')',
            error(sprintf('Error in formula: %s',formula));
        end
        % count parentheses
        pc = 1; 
        cend = length(formula1);
        cstart = cend;
        while pc ~= 0,
            cstart = cstart - 1;
            if formula1(cstart) == ')',
                pc = pc+1;
            elseif formula1(cstart) == '(',
                pc = pc-1;
            end
        end
        firstargument = formula1(cstart+1:cend-1);
    else
        firstargument = firstargument{1};
        cstart = length(formula1)-length(firstargument)+1; 
    end
    cendfirst = cstart;
    % check formula2 from the left
    secondargument = regexp(formula2,'^([\w]+[.]?[\w]*)','match');
    if isempty(secondargument),
        % check if first character is an opening parenthesis
        if formula2(1) ~= '(',
            error(sprintf('Error in formula: %s',formula));
        end
        % count parentheses
        pc = 1; 
        cstart = 1;
        cend = cstart;
        while pc ~= 0,
            cend = cend + 1;
            if formula2(cend) == '(',
                pc = pc+1;
            elseif formula2(cend) == ')',
                pc = pc-1;
            end
        end
        secondargument = formula2(cstart+1:cend-1);
    else
        secondargument = secondargument{1};    
        cend = length(secondargument);
    end
    cstartsecond = cend;
    % construct power expression
    powerexp = sprintf('pow(%s,%s)',firstargument,secondargument);
    % construct new formula
    formula = [formula1(1:cendfirst-1) powerexp formula2(cstartsecond+1:end)];
    % get new indices for '^' character
    indices = strfind(formula,'^');
end
return

% function [formula] = processDelayCallSB(formula)
% global delaycount delaybase
% count = 1;
% while 1,
%     index = strfind(formula,'delaySB(');
%     if length(index) < count,
%         break;
%     end
%     indexstart = index(count)+length('delaySB(');
%     indexend = indexstart;
%     
%     % search the end of the delay argument definition
%     parOpen = 1;
%     while parOpen ~= 0,
%         if formula(indexend) == '(',
%             parOpen = parOpen + 1;
%         elseif formula(indexend) == ')',
%             parOpen = parOpen - 1;
%         end
%         indexend = indexend + 1;
%     end
%     % check if the delaybasename has to be changed
%     if length(index) > 1,
%         delayname = [delaybase '_' sprintf('%d', delaycount) '_' sprintf('%d', count)];
%     else
%         delayname = [delaybase '_' sprintf('%d', delaycount)];
%     end
%     % add info to delaySB call
%     firstpart = formula(1:indexend-2);
%     lastpart = formula(indexend-1:end);
%     middlepart = sprintf(',time,"%s"',delayname);
%     formula = char([double(firstpart) double(middlepart) double(lastpart)]);
%     % increase counters
%     count = count + 1;
%     delaycount = delaycount + 1;
% end
% return
```
