## Appendix: Model description for "A multi-scale *in silico* mouse model for insulin resistance and humanoid type 2 diabetes"

**Supplementary Appendix**

**Simonsson et al, 2021**

### **1. Model description: the multi-level and multi-timescale model**

The complete model is comprised of four different sub-models describing metabolic control on the three different levels (Fig 1A-B). This section of the appendix goes through the equations in each of these sub-models, one-by-one, and also highlights the novel model interconnections that were required to create the multi-level structure and the new insulin resistance description.

**I. Body-composition model**

The body-composition model developed by Guo *et al*. (1) was used to describe the weight-change part of our multi-level dataset. The model (Fig. 2B) divides body mass into two compartments with one state each: fat mass ($FM$) and fat-free mass ($FFM$). As can be seen in Fig 2B, body mass is governed by the balance between energy intake ($\mathrm{EI}$) and expenditure ($\mathrm{EE}$), according to:

$\rho_{FM}\left( \frac{dFM}{dt} \right)+\rho_{FFM}\left( \frac{dFFM}{dt} \right)=EI-EE$	(Eq.1)

where $\rho_{FM}$ and $\rho_{FFM}$ are the energy densities for $FM$and $FFM$, respectively. The energy density is deﬁned as amount of energy stored per unit mass (kcal/g) and is higher for $FM$ than for $FFM$. More specifically, the EI is the daily energy intake rate and is a set value defined in kcal/day. The EE is the energy expenditure rate (kcal/day) defined as:

$EE=\frac{K+\beta\cdot\Delta EI + \left( \gamma_{FFM}+\lambda\right)\cdot FFM+\left( \gamma_{FM}+\lambda\right)\cdot FM+ \eta_{FFM}\cdot\alpha\cdot g\cdot EI+\eta_{FM}\cdot g\cdot EI}{1+ \eta_{FM}\cdot g+\eta_{FFM}\cdot\alpha\cdot g}$	(Eq. 2A)

$\Delta EI=EI-\mathrm{EI}_{\mathrm{standard}}$	(Eq. 2B)

where $K$ represents a thermogenesis parameter; the $\beta$ parameter scales the $\Delta EI$ to effect on thermogenesis; $\Delta EI$ represents the change of EI compared to a standard low-fat diet, here denoted $\mathrm{EI}_{\mathrm{standard}} (Eq. 2B)$; $\gamma_{\mathrm{FFM}}$ and $\gamma_{\mathrm{FM}}$ are parameters representing the metabolic rate for *FFM* and *FM* respectively; the parameter $\lambda$ represents the physical activity; $\eta_{FFM}$ and $\eta_{FM}$ are parameters representing synthesis of fat-free mass and fat mass for *FFM* and *FM* respectively; $\alpha$ is an empirical function, a Forbes curve relating changes in *FFM* and *FM;* and $g$ is a variable used to simplify the EE equation. The variable $g$ is defined as:

$g=\frac{1}{\alpha\cdot\rho_{FFM}\cdot\rho_{FM}}$	(Eq. 3)

The empirical function α, (1, 2), that describes the relationship between changes of *FFM* and *FM* is defined as:

$\alpha=\frac{dFFM}{dFM}$ = $\alpha_{c}+ \alpha_{d}+e^{\alpha_{k}\cdot FM}$	(Eq.4)

where $\alpha_{c}$, $\alpha_{d}$ are $\alpha_{k}$ empirically estimated constants. We used the values for male C57BL/6J measured by *Gao et al* (1)*.* Given $\alpha$, Eq. 4 can be simpliﬁed and rearranged as:

$\frac{dFFM}{dt}=\frac{\alpha}{\rho_{FFM}\alpha+\rho_{FM}}(EI-EE)$ $, FFM\left( 0 \right)=fraction_{FFM}\cdot BW\left( 0 \right)\left[ g \right]$	(Eq.5)

$\frac{dFM}{dt}=\frac{1}{\rho_{FFM}\alpha+\rho_{FM}}(EI-EE)$ $, FM\left( 0 \right)=fraction_{FM}\cdot BW$(0) [g]	(Eq.6)

where the initial conditions ($FM\left( 0 \right)$and$FFM\left( 0 \right)$) were set to estimated fat and fat-free mass from the body weight of the mice included in our own experiments; and where the variables $fraction_{FM}$ and $fraction_{FFM}$ describes fat and fat-free mass divided by the body weight, respectively. The mean composition between fat- and fat-free mass was based on estimates done experimentally (3): $fraction_{FM}=0.11$ and $fraction_{FFM}=0.89$. Similarly, the value of $EI$ was also set to match our own experiments by using the measured values (9–11 kcal/day for chow diet and 16–18 kcal/day for HFD).

Finally, some comments on the model parameters. All parameters for the body weight model, except the physical activity parameter ($\lambda)$ were kept constant, and not included in the parameter estimation. The values were taken from Gao *et al.* (2009) and can be seen in Table 1.

**Table 1:** *Parameters of the body-composition model. The parameters presented in the first part of the table were kept constant and are refence values taken from Gao et al (2009). The lower part of the table presents the parameters that were included in the parameter estimation.*

| Constants | | |
| --- | --- | --- |
| Name | **Value** | **Unit** |
| $\boldsymbol{K}$ | 2.1 | $kcal\cdot g^{-1}$ |
| $\boldsymbol{\beta}$ | 0.4 |  |
| $\mathbf{EI}_{\mathbf{standard}}$ | 10 | $kcal\cdot{day}^{-1}$ |
| $\boldsymbol{\gamma}_{\boldsymbol{FFM}}$ | 0.15 | $kcal\cdot g^{-1}\cdot day^{-1}$ |
| $\boldsymbol{\gamma}_{\mathbf{FM}}$ | 0.03 | $kcal\cdot g^{-1}\cdot day^{-1}$ |
| $\boldsymbol{\eta}_{\boldsymbol{FFM}}$ | 0.23 | $kcal\cdot g^{-1}$ |
| $\boldsymbol{\eta}_{\boldsymbol{FM}}$ | 0.18 | $kcal\cdot g^{-1}$ |
| $\boldsymbol{\rho}_{\boldsymbol{FFM}}$ | 1.8 | $kcal\cdot g^{-1}$ |
| $\boldsymbol{\rho}_{\boldsymbol{FM}}$ | 9.4 | $kcal\cdot g^{-1}$ |
| $\boldsymbol{\alpha}_{\mathbf{c}}$ | 0.1 |  |
| $\boldsymbol{\alpha}_{\mathbf{d}}$ | 0.00019 |  |
| $\boldsymbol{\alpha}_{\mathbf{k}}$ | 0.45 |  |
| Estimated parameters | | |
| Name | $\boldsymbol{\theta}_{\mathbf{best}}$ | **Unit** |
| $\boldsymbol{\lambda}_{\boldsymbol{chow2w}}$ | 0.166 | $kcal\cdot g^{-1}$ |
| $\boldsymbol{\lambda}_{\boldsymbol{HFD2w}}$ | 0.162 | $kcal\cdot g^{-1}$ |
| $\boldsymbol{\lambda}_{\boldsymbol{chow8w}}$ | 0.145 | $kcal\cdot g^{-1}$ |
| $\boldsymbol{\lambda}_{\boldsymbol{HFD8w}}$ | 0.179 | $kcal\cdot g^{-1}$ |
| $\mathbf{EI}_{\mathbf{chow2w}}$ | 10.069 | $kcal$ |
| $\mathbf{EI}_{\mathbf{chow8w}}$ | 11.000 | $kcal$ |
| $\mathbf{EI}_{\mathbf{chow8w}}$ | 14.999 | $kcal$ |

To find agreement to the training data we estimated the physical activity parameter, $\lambda,$as also done in Gennemark *et al (**4**).* An $\lambda$ was estimated for each individual diet in the training data set, to allow for difference in experimental conditions. The value of $\lambda$ for each diet is presented in Table 1. We also estimated the energy intake for the 2 and 8 chow diets as well as the 8-week HFD intervention. An assumption that was made was that we used the mean energy intake during all diets. However, HFD mice has a higher energy intake during the initial time period and then becomes isocaloric in respect to the chow controls. Thus, it was not possible to use the measured mean energy intake value for the 8-week HFD intervention, as it would have been much to low compared to the mean for the 2-week HFD and 2-week switch diet interventions, and it was instead estimated.

**II. Glucose and insulin regulation model**

The glucose and insulin regulation model developed by Alskär *et al*. (5) was used as a basis to describe plasma glucose and insulin data. The structure of our modified model can be seen in Fig. 2C. Our own data included plasma glucose values during an intraperitoneal glucose tolerance test (IPGTT) at varying time-points and diets, as well as the fasting insulin level over the course of the experiment. Note that the Alskär *et al*. model was originally developed using intra-venous GTT plasma glucose data. To model our own IPGTT data, a state representing glucose in an intraperitoneal compartment, $G_{ip}$, was added to the model. This state is equal to zero prior to the glucose dose of the IPGTT and assigned the glucose dose (in moles) at the time of the glucose dose. As can be seen in Fig 2D, the ensuing dynamics is only given by an export out from the intraperitoneal compartment, since there normally is no glucose transported to this compartment, except in the exceptional situation of an injection of glucose. In other words, the $G_{ip}$ derivative is defined as

$\frac{dG_{ip}}{dt}= -{{k_{i}}_{p}G}_{ip}$	(Eq.7)

where *k_ip_* governs the rate at which the injected glucose load enters the blood stream, which is the central glucose compartment $G_{c}$. As can be seen in Fig 2D, the amount of glucose in the central plasma compartment is determined by several terms, representing some of the major influxes and effluxes, such as glucose utilization and endogenous production. More specifically, the equations are:

$\frac{dG_{c}}{dt}= EGP + Q_{VP}\cdot G_{P}- Q_{VG}\cdot G_{C}- U + G_{ip}\cdot k_{ip} , G_{c}\left( 0 \right)=G_{SS}\cdot V_{G}[mmol]$	(Eq.8A)

$Q_{VG}=\frac{Q}{V_{G}}$	(Eq.8B)

$Q_{VP}=\frac{Q}{V_{P}}$	(Eq.8C)

where $EGP$is the endogenous glucose production; $U$ is a big expression which represents the glucose utilization; $G_{P}$ denotes the amount of peripheral glucose; $Q_{VG}$is a variable describing the rate of glucose moving from plasma to peripheral tissue; $Q_{VP}$ is the variable describing the rate for glucose returning to plasma from peripheral tissue; Q is the rate constant of diffusion between $G_{c}$ and $G_{P}$; $V_{G}$is the plasma glucose volume of distribution and $V_{P}$ is peripheral glucose volume of distribution; and where $G_{SS}$ is a parameter that represents the initial concentration of glucose in the central compartment, $G_{c}\left( 0 \right)$. As previously described, plasma glucose can freely diffuse into peripheral tissue, which glucose level is described by the state $G_{P}$. The ordinary differential equation (ODE) for $G_{p}$ is given by fluxes already introduced, i.e.

$\frac{dG_{P}}{dt}= - Q_{VP} \cdot G_{P}+ Q_{VG}\cdot G_{C}$, $G_{P}\left( 0 \right)=G_{SS}\cdot V_{P}[mmol]$	(Eq.9)

The endogenous production of glucose is represented by the EGP, which describes glucose output from liver and kidney. The EGP depends on $G_{SS}$, the basal value of plasma insulin $I_{SS}$, and the rate parameters for insulin independent $k_{CLG}$ and dependent $k_{CLGI}$glucose utilization. The variable denoted ${f_{IR}}_{EGP}$ describes insulin resistance progression and is explained in more detail in section IV (Eq. 64). The EGP also depend on the amount of plasma glucose through the variable $G_{EG}$, which represents the effect of plasma glucose on EGP. These dependences together means that EGP is governed by the following expression:

$EGP=G_{SS}{\cdot G}_{EG}\left( k_{CLG}+k_{CLGI}\cdot I_{SS} \right)\cdot{f_{IR}}_{EGP}$	(Eq.10)

The variable $G_{EG}$ is given by:

$G_{EG}= \left( \frac{G_{E2}}{G_{SS}} \right)^{GPRG}$	(Eq.11)

where the parameter $GPRG$ is the exponent and $G_{E2}$ is a model state which represents functional plasma glucose in the liver. The $GPRG$ parameter is always negative to ensure the negative effect glucose has on the EGP. Note that Eq. 11 involves a ratio, to normalize $G_{E2}$ with $G_{SS}$. The state $G_{E2}$ represent the plasma glucose that effects the liver, and its ODE is written in the following way:

$\frac{dG_{E2}}{dt}=k_{G_{E2}}\left( \frac{G_{C}}{V_{G}}-G_{E2} \right), G_{E2}\left( 0 \right)=G_{SS} [mmol]$	(Eq.12)

where $k_{G_{E2}}$ is the rate parameter of glucose diffusion between liver and plasma.

The final not yet mentioned term in Eq. 8 governs the utilization of glucose. This term is denoted U and is divided into insulin independent ($U_{ii}$) and insulin-dependent ($U_{id}$) glucose utilization, i.e.:

$U=U_{ii}+U_{id}$	(Eq.13)

The insulin independent glucose utilization ($U_{ii}$) only varies with the amount of glucose present in plasma, and is written as:

$U_{ii}=k_{CLG}\frac{G_{C}}{V_{G}}$	(Eq.14)

where $k_{CLG}$ is the rate of insulin independent glucose utilization. The $U_{ii}$ represents passive uptake, e.g., glucose uptake by the brain. To be able to create some of the main multi-level connections, Alskär’s original single-term insulin dependent glucose utilization ($U_{id}$) is now divided into three terms, representing the relative uptake from the three major tissues - adipose ($u_{a})$, muscle ($u_{m}$) and hepatic tissue ($u_{h}$) - and is thus written as:

$U_{id}=u_{a}+u_{m}+u_{h}$	(Eq.15)

The expression for adipose glucose uptake ($u_{a})$is closely connected to the intracellular insulin signaling model, and is described in section III*,* Eq. 22. The expression for glucose uptake in muscle is written as:

$u_{m}=\frac{kCLGIm\cdot\frac{FFM}{FFM_{0}}\cdot I_{E}\cdot\frac{G_{C}}{V_{G}}}{f_{IR_{CLGI}}}$	(Eq.16)

where $kCLGIm$ is the rate constant for glucose uptake by muscle tissue; $\frac{FFM}{FFM_{0}}$ is the increase in fat-free mass; $I_{E}$ is the model state representing central plasma insulin and $f_{IR_{CLGI}}$ is a model variable inducing the effect of insulin resistance on the insulin dependent glucose uptake (in muscle and hepatic tissue), described in more detail in section IV (Eq. 66). Similarly, the expression for glucose uptake in the liver ($u_{h})$is written as

$u_{h}=\frac{kCLGIh\cdot I_{E}\cdot\frac{G_{C}}{V_{G}}}{f_{IR_{CLGI}}}$	(Eq.17)

where $kCLGIh$ is the rate constant for glucose uptake by hepatic tissue. The key difference from the muscle expression is that liver expression has no dependency on fat free mass, i.e. we assume that the functional liver volume is constant.

We will now go through the model states associated with plasma insulin. The acting insulin, $I_{E}$, represent the insulin that has diffused into tissue. The acting insulin $I_{E}$ has the following ODE:

$\frac{dI_{E}}{dt}=k_{I_{E}}( \frac{I_{C}}{V_{I}}-I_{E} )$, $I_{E}\left( 0 \right)= I_{SS}$	(Eq.18)

where $k_{I_{E}}$ is the rate of which insulin enters tissue; $I_{C}$ is a model state representing central plasma insulin; $V_{I}$ is the volume of distribution of plasma insulin; and $I_{SS}$ is the baseline plasma insulin level. The derivative for the central plasma insulin is defined as:

$\frac{dI_{c}}{dt}=I_{sec}-\frac{{k_{CLI}\cdot I}_{C}}{V_{I}}$ , $I_{C}\left( 0 \right)=$ $I_{SS}$*$V_{I}$	(Eq.19)

where the rate parameter $k_{CLI}$ governs the insulin clearance in circulation and the $I_{sec}$ is a model variable describing insulin secretion from pancreas into plasma. $I_{sec}$ is defined as:

$I_{sec}= \frac{I_{SS}\cdot k_{CLI}\cdot{f_{InsRes}}_{I_{sec}}\cdot G_{Eins}}{T2d_{effect}}$	(Eq.20)

where $G_{Eins}$ is a model variable representing the effect of glucose on insulin secretion; the variable ${f_{InsRes}}_{I_{sec}}$ describes the effect of insulin resistance on the insulin secretion and is described in section IV (Eq. 65) and the $T2d_{effect}$ variable introduces the effects of T2D into the system, described in detail in section V (Eq 69). The parameters $k_{CLI}$ and $I_{SS}$ determine the baseline secretion of insulin. The $G_{Eins}$ variable is a power function dependent on the glucose acting on the pancreas $G_{E1}$, and is written as:

$G_{Eins}=\left( \frac{G_{E1}}{G_{SS}} \right)^{IPRG}$	(Eq.21)

where the parameter $IPRG$is an exponent that determines the sensitivity of the insulin secretion in response to glucose. Note that $G_{E1}$ is normalized with the $G_{SS}$ parameter. As described, $G_{E1}$ represent the plasma glucose that acts on the pancreas, and its dynamics is given by the following ODE:

$\frac{dG_{E1}}{dt}=k_{G_{E1}}\cdot\left( \frac{G_{C}}{V_{G}}-G_{E1} \right)$	$, G_{E1}\left( 0 \right)=G_{SS}$	(Eq.22)

where $k_{G_{E1}}$ is the rate constant governing the diffusion on plasma glucose to the pancreas.

Finally, some comments on the scaling of the parameters in our modified Alskär model. We used the model parameters values presented in Alskär *et al.* as a start guesses (θ_0_) for the parameter estimation. The parameters were also allometrically scaled to mice as presented in Alskär *et al.* using the allometric equation given by:

$P_{mice}=P_{human}\cdot\left( \frac{BW_{mice}}{BW_{human}} \right)^{b}$	(Eq.23)

where $P_{mice}$ is approximated parameter value in mice; $P_{human}$ is the parameter value in humans; $BW_{mice}$ is the mice body weight; $BW_{human}$ is the human bodyweight; and *b* is the allometric exponent. The human body weight used was 70 kg and the mice body weight used was 23 g. The alometric exponents vary depending on what the parameter represents. In Alskär *et al.,* the following values were presented: when scaling parameters representing clearance *b* was set 0.75, for parameters representing volume *b* was set to 1, and for rate parameters *b* was set to -0.25. Also, in Alskär *et al.,* the rate parameters were expressed in change per minute, which we have changed to per hour. In Table 3 a parameter comparison is presented, showing the allometrically scaled parameters for mice used by Alskär *et al.* and the parameters which we used as our estimation start guess (θ_0_).

**Table 2:** *Table showing the model parameters for the glucose and insulin model allosteric scaled to mice in the same way as described in Alskär et al. Also, the same values but converted into rate of change per hour were instead used as the parameter estimation start guess θ_0._ The last column of the table shows the model parameter set which yielded the best agreement to the training data.*

| Name | Alskär *et al.* | θ_0_ | $\boldsymbol{\theta}_{\mathbf{best}}$ |
| --- | --- | --- | --- |
| *Q* | 0.001078 ($L\cdot\min^{-1})$ | 0.064721($L\cdot h^{-1}$) | 0.0388($L\cdot h^{-1}$) |
| *k_IE_* | 0.158206 ($\min^{-1}$) | 9.492342($h^{-1}$) | 5.9375 ($h^{-1}$) |
| *k_GE1_* | 0.214654 ($\min^{-1}$) | 12.879281($h^{-1}$) | 11.6564($h^{-1}$) |
| *k_GE2_* | 0.425595 ($\min^{-1}$) | 25.535738($h^{-1}$) | 14.3337($h^{-1}$) |
| *CL_GI_* | 5.71414e-06($L\cdot\min^{-1})$ | 3.428489e-04($L\cdot h^{-1}$) | 3.429e-05($L\cdot h^{-1}$) |
| *CL_G_* | 0.000363($L\cdot\min^{-1})$ | 0.0217862($L\cdot h^{-1}$) | 0.0022($L\cdot h^{-1}$) |
| *CL_I_* | 0.002977($L\cdot\min^{-1})$ | 0.178641($L\cdot h^{-1}$) | 0.2436($L\cdot h^{-1}$) |
| *IPRG* | 1.42 | 1.42 | 1.845 |
| *GPRG* | -2.79 | -2.79 | -3.90 |
| *V_G_* | 0.003065 (L) | 0.003065 (L) | - |
| *V_P_* | 0.0028125(L) | 0.0028125(L) | - |
| *V_I_* | 0.002001 (L) | 0.002001 (L) | - |
| *k_ip_* | - | - | 1.8513($L\cdot h^{-1}$) |
| *Gss* | - | 5.85$(mmol\cdot L^{-1})$ | 6.0712$(mmol\cdot L^{-1})$ |
| *Iss* | - | 68.8($pmol\cdot L^{-1})$ | 68.168($pmol\cdot L^{-1})$ |

During parameter estimation all parameters can vary at most 40% from these values, with some exceptions. The parameters for glucose clearance had to be allowed more freedom and could vary 100% from the initial start guess (**θ_0_**), for the model to able to fit data. The parameter set best describing the training dataset can be found in Table 3. The parameters representing the different volumes of distribution (V_G_ , V_P_ and V_I_) were held constant and were not included in the estimation.

**III. The adipocyte intracellular insulin signaling model**

Our previously published adipocyte insulin signaling model describes adipose glucose uptake in a mechanistic manner (6). The model input is the acting insulin $I_{E}$ (Eq.18). Upon insulin stimulation of the insulin receptor (IR), a signaling cascade is triggered which leads to the translocation of GLUT4 into the plasma membrane. The ODE for the concentration of IR, denoted $IR$, is given by the three arrows going in and out of IR in Fig. 2E, i.e.:

$\frac{dIR}{dt}= -ik1\cdot IR\left( I_{e}\cdot kscale \right)-ik1basal\cdot IR+ikR\cdot IRi$	(Eq.24)

$$IR\left( 0 \right)= 9.8696 [A. u]$$

where $ik1$ is the rate parameter governing the insulin dependent phosphorylation of the receptor; $ik1basal$ is the rate parameter governing the basal phosphorylation of the receptors; $ikR$ is the rate parameter for the re-surfacing of receptors after internalization; where $kscale$ is the conversion of acting insulin, defined for organ-level dynamics, into nM, defined for intracellular dynamics; and where $IRi$ represents the concentration of internalized IR. The model state$IRp$ represents the concentration of phosphorylated receptors and the ODE is defined as

$\frac{dIRp}{dt}= ik1\cdot IR1\left( I_{e}\cdot kscale \right)+ik1basal\cdot IR-ik2\cdot IRp$	(Eq.25)

$$IR_{p}\left( 0 \right)= 0.0269364 [A. u]$$

where $ik2$ is the rate of receptor internalization. The internalized $(IRi$) receptor ODE is given by:

$\frac{dIRi}{dt}=ik2\cdot IRp- ikR\cdot IRi$	(Eq.26)

$$IR_{i}\left( 0 \right)= 0.103462 [A. u]$$

The phosphorylated receptor, $IRp,$ initiates signaling downstream. This downstream cascade has two major pathways: the pathway via insulin receptor substrate ($IRS$) and a pathway located close to plasma membrane near the IR which activity is represented by the state *PMR*. In the model, the *IRS* pathway is more complex compared to the PMR pathway (which almost directly regulates the GLUT4 translocation). The most established pathway via $IRS$ includes the phosphorylation of multiple sites of protein kinase B (PKB), which occurs downstream IRS activation. The derivatives of $IRS$are written as

$\frac{dIRS}{dt}=-\frac{ik3\cdot IRp\cdot IRS}{1+ikf\cdot Xp}+ikm3\cdot IRSp+ k_{basal}\left( \frac{1}{{f_{InsRes}}_{IRS}} -IRS \right)$	(Eq.27)

$$IRS\left( 0 \right)= 9.99981 [A. u]$$

$\frac{dIRSp}{dt}=- ikm3\cdot IRSp+\frac{ik3\cdot IRp\cdot IRS}{1+ikf\cdot Xp}$	(Eq.28)

$$IRS_{p}\left( 0 \right)= 0.000189 [A. u]$$

where $ik3$ is the rate of phosphorylation; $IRSp$ is the model state representing the phosphorylated form of IRS; $Xp$ represents the active form of an unknown protein; $ikf$ is a parameter governing the inhibition of the same unknown protein; $ikm3$ is the rate parameter governing the dephosphorylation of IRS; $k_{basal}$ governs the rate of production and degradation of *IRS* and ${f_{InsRes}}_{IRS}$ is a variable representing the effect of insulin resistance on IRS expression, described in detail in section IV (Eq. 67). The first term in (Eq. 27) represents the $IRp$ dependent rate of phosphorylation of IRS. The IRS phosphorylation has a negative feedback loop, which have been investigated in our previous publication (6). As is already alluded to, when IRS becomes phosphorylated, it facilitates phosphorylation of an unknown protein, $X$. The phosphorylated form of the unknown protein *Xp* then inhibits the phosphorylation of IRS. The ODEs of the unknown protein are defined as

$\frac{dX}{dt}=-ik4\cdot IRSp\cdot X+ikm4\cdot Xp$	(Eq.29)

$X\left( 0 \right)= 9.89417 [A. u]$

$\frac{dXp}{dt}=ik4\cdot IRSp\cdot X-ikm4\cdot Xp$	(Eq.30)

$$X_{P}\left( 0 \right)=0.105826 [A. u]$$

where, $ik4$ is the rate parameter governing phosphorylation and $ikm4$ is the rate parameter governing deactivation.

Phosphorylated IRS ($IRSp)$ facilitates downstream signaling. $IRSp$ facilitates the activation of the mTOR Complex 2 ($mTorc2)$. The ODEs of $mTorc2$ are written as:

$\frac{dmTorc2}{dt}=-mTorc2\cdot kmt1\cdot IRSp+mTorc2a\cdot kmt2$	(Eq.31)

$$mTorc2\left( 0 \right)= 1 [A. u]$$

$\frac{dmTorc2a}{dt}=-mTorc2\cdot kmt2+ mTorc2\cdot kmt1\cdot IRSp$	(Eq.32)

$$mTorc2a\left( 0 \right)= 0 [A. u]$$

where $kmt1$ is the rate parameter governing the activation; $kmt2$ is the parameter for the rate of deactivation and $mTorc2a$ is the model state representing the active form of the complex. After activation, $mTorc2a$ then facilitates phosphorylation of the S473 site in the PKB/Akt complex. The signaling from $IRS$ also initiates the transition of membrane-bound phosphatidylinositol (4,5)-bisphosphate ($PIP2$) to phosphatidylinositol 3,4,5-trisphosphate ($PIP3$) with ODEs written as

$\frac{dPIP2}{dt}=-PIP2\cdot IRSp\cdot kpi1+PIP3\cdot kpi2$	(Eq.33)

$$PIP2\left( 0 \right)= 10 [A. u]$$

$\frac{dPIP3}{dt}=-PIP3\cdot kpi2+PIP2\cdot IRSp\cdot kpi1$	(Eq.34)

$$PIP3\left( 0 \right)= 0.001 [A. u]$$

Here, $kpi1$ is the rate parameter governing transition from PIP2 to PIP3 and $kpi2$ governs the rate backward transition from PIP3 to PIP2. Thereafter, $PIP3$ activates phosphoinositide-dependent protein kinase-1 ($PDK1$) with ODEs defined as

$\frac{dPDK1}{dt}=-PDK1\cdot PIP3\cdot kpd1+PDK1a\cdot kpd2$	(Eq.35)

$$PDK1\left( 0 \right)= 1 [A. u]$$

$\frac{dPDK1a}{dt}=-PDK1a\cdot kpd2+PDK1\cdot PIP3\cdot kpd1$	(Eq.36)

$$PDK1a\left( 0 \right)= 0.01 [A. u]$$

where $kpd1$ is the rate parameter governing activation of $PDK1$ by $PIP3$, and where $kpd2$ governs the rate of deactivation of $PDK1$. Both $PDK1$a and $mTorc2a$ help facilitate phosphorylation of site T308 and S473 on $PKB/Akt$. The model describes four different states of PKB/Akt, representing all combinations of the non-phosphorylated and phosphorylated states of the two phosphorylation sites. The non-phosphorylated form of $PKB/Akt$ is described by a series of reactions, which just as for all other equations above, are explicitly depicted in Fig. 2E. Using standard mass action kinetics these reactions become the following equation:

$\frac{dPKB}{dt}=-PKB\cdot k3081\cdot PDK1a+{PKB}_{308}\cdot k3082-PKB\cdot k4731\cdot mTorc2a+{PKB}_{473}\cdot k4732$	(Eq.37)

$$PKB\left( 0 \right)= 6 [A. u]$$

where $k3081$ is the rate parameter governing the PDK1 dependent phosphorylation of PKB at site 308; the rate parameter $k3082$determines the rate of dephosphorylation of${PKB}_{308}$; the rate parameter $k4731$ governs the rate of $mTorc2$ dependent phosphorylation of PKB at site 473; the rate parameter $k4732$ governs the rate of the dephosphorylation of ${PKB}_{473}$ and ${PKB}_{473}$ and ${PKB}_{308}$ are model states representing phosphorylated PKB at different sites. The derivative for ${PKB}_{308}$ is written as

$\frac{dPKB_{308}}{dt}=PKB\cdot k3081\cdot PDK1a-{PKB}_{308}\cdot k308-{PKB}_{308}\cdot k308473\cdot mTorc2a+{PKB}_{308473}\cdot k34$(Eq.38)

$$PKB_{308}\left( 0 \right)= 0 [A. u]$$

where $k308473$ is the rate parameter governing the phosphorylation of site S473 for ${PKB}_{308}$ ; $k34$is the rate parameter governing the dephosphorylation of site S473 on PKB phosphorylated at both sites and ${PKB}_{308473}$ is PKB phosphorylated at both sites. Similarly, the derivative for ${PKB}_{473}$ is written as

$\frac{dPKB_{473}}{dt}=PKB\cdot k4731\cdot mTorc2a-{PKB}_{473}\cdot k4732-{PKB}_{473}\cdot k473308\cdot PDK1a+{PKB}_{308473}\cdot k43$ (Eq.39)

$$PKB_{473}\left( 0 \right)= 0.07 [A. u]$$

where $k473308$is the rate parameter governing the phosphorylation of site T308 for ${PKB}_{473}$ and $k43$is the rate parameter governing the dephosphorylation of site T308 on PKB phosphorylated at both sites. The derivative for PKB phosphorylated on both sites is defined as:

$\frac{dPKB_{308473}}{dt}={PKB}_{308}\cdot k308473\cdot mTorc2a+{PKB}_{473}\cdot k473308\cdot PDK1a -{PKB}_{308473}\cdot k34-{PKB}_{308473}\cdot k43$ (Eq.40)

$$PKB_{308473}\left( 0 \right)= 0 [A. u]$$

The different states of $PKB/Akt$ also have different signaling actions. However, $PKB/Akt$ with phosphorylation on site T308 is the only active version of PKB in the model and is defined as:

$Total_{PKB_{308}}= dPKB_{308473}$ + $PKB_{308}$	(Eq.41)

This total activated form of PKB facilitates the activity of two pathways. The first such downstream activation is to facilitate the activation of an AS160-independent pathway downstream of PKB. AS160 denotes $PKB/Akt$ substrate of 160 kDa. This AS160-independent activation step is referred to as $PKBd$, to symbolize that it is dependent on PKB (in contrast to PMR, which is independent of both AS160 and PKB). The ODEs for the state $PKBd$, and its active form $PKBda$ are defined by:

$\frac{dPKBd}{dt}=PKBda\cdot k_{pI2t2} -PKBd\cdot k_{pI1t2}\cdot Total_{PKB_{308}}$	(Eq.42)

$$PKBd\left( 0 \right)= 1 [A. u]$$

$\frac{dPKBda}{dt}=$	$PKBd\cdot k_{pI1t2}\cdot Total_{PKB_{308}}- PKBda\cdot k_{pI2t2}$	(Eq.43)

$$PKBda\left( 0 \right)= 0.01 [A. u]$$

where the rate parameter $k_{pI1t2}$ governs the activation of the pathway and the deactivation of the active pathway form is governed by the rate parameter $k_{pI2t2}$. The second signaling action of the active PKB is to facilitate phosphorylation on the $AS160$ which is given by the following equations:

$\frac{dAS160}{dt}=AS160p\cdot k_{as2} -AS160\cdot k_{as1}\cdot Total_{PKB_{308}}$	(Eq.44)

$$AS160\left( 0 \right)= 2.5 [A. u]$$

$\frac{dAS160p}{dt}=AS160\cdot k_{as1}\cdot Total_{PKB_{308}}-AS160p\cdot k_{as2}$	(Eq.45)

$$AS160p\left( 0 \right)= 0.1 [A. u]$$

where $k_{as2}$ is the rate parameter governing the rate of dephosphorylation of AS160 ; $k_{as1}$ is the parameter governing the rate of phosphorylation of AS160 and $AS160p$ is the model state representing phosphorylated $AS160$.

AS160 interacts with the small G-protein Rab, and in the non-insulin stimulated state, AS160, through its GTPase-activating protein (GAP) domain, keeps maintaining the inactive GDP-bound form of Rab. When AS160 becomes phosphorylated, the GAP activity becomes inhibited, resulting in an increase in the active form of Rab (RABGTP). The ODEs for *RAB* are defined as:

$\frac{dRABGTP}{dt}=RABGDP\cdot k_{gka1} -RABGTP\cdot k_{gka2}\cdot AS160$	(Eq.46)

$$RABGTP\left( 0 \right)= 0 [A. u]$$

$\frac{d\mathrm{RABGDP}}{dt}=RABGTP\cdot k_{gka2}\cdot AS160-RABGDP\cdot k_{gka1}$	(Eq.47)

$$RABGDP\left( 0 \right)= 5 [A. u]$$

where $k_{gka1}$is the rate parameter governing $RABGDP$ moving to the active GTP bound and $k_{gka2}$ is the rate parameter for RABGTP moving into de inactive GDP bound form. The active form RAB-GTP interacts with the GLUT4 translocation in various ways.

Going back to the activation of the insulin receptor and the second major signaling pathway which is started already by IR. In other words, the second alternative pathway describes a plasma membrane-located pathway near the IR, denoted “PMR”. This pathway describes the insulin-induced effect of plasma membrane activity associated with changes in the rate of GLUT4 movement from GLUT4 clusters to monomer form in the membrane. The PMR pathway also inhibits the rate of endocytosis of GLUT4 cluster into endosomes. The ODEs for the PMR pathway are defined as:

$\frac{dPMR}{dt}=PMRa\cdot k_{gk12} -PMR\cdot k_{gk11}\cdot\frac{IRp}{f_{IR_{PMR}}}$	(Eq.48)

$$PMR\left( 0 \right)= 1 [A. u]$$

$\frac{dPMRa}{dt}=PMR\cdot k_{gk11}\cdot\frac{IRp}{f_{IR_{PMR}}}-PMRa\cdot k_{gk12}$	(Eq.49)

$$PMRa\left( 0 \right)= 0 [A. u]$$

where$k_{gk11}$is the rate parameter governing the pathway activation; $k_{gk12}$ is the rate parameter for deactivation; $PMRa$ is the model state representing the active form of the pathway and $f_{IR_{PMR}}$ is a variable representing the effect of insulin resistance on PMR activity, described in detail in section IV (Eq. 68).

The $PMR$ and the two *IRS* signaling pathways are the three ways of regulating the GLUT4 translocation in the model. The GLUT4 translocation in the model describes the movement between stages of either GLUT4 storage in vesicles or active membrane bound form. In the model, the stored form of GLUT4 is either described by the ratio of GLUT4 (compared to the populations of the other GLUT4 states) in the endosome (denoted *C3* in the model, Eq. 50) or the amount of GLUT4 stored in vesicles (C0, Eq. 51).

$C3=1-C0-C1-C2$	(Eq.50)

The state C0 is defined by the influx and efflux to the GLUT4 vesicle storage and the derivative of *C0* can be written as

$\frac{dC0}{dt}=gk3\cdot C3-gv1-gv2$	(Eq.51)

$$C0\left( 0 \right)= 0.75 [A. u]$$

where the $gk3$ rate parameter governs the influx of GLUT4 from the endosome compartment (C3); $gv1$ represent the transport of GLUT4 from the vesicles storage to membrane bound active GLUT4 monomers and $gv2$represent the transport from GLUT4 vesicle storage (C0) to active membrane bound GLUT4 clusters. The $gv1$ variable can be written as

$$gv1=gk1bas\cdot C0\cdot RABGTP\cdot PKBda\cdot PMRa$$

(Eq.52)

where, $gk1bas$ is the rate parameter governing the GLUT4 transport to membrane bound GLUT4 monomers. The transport represented by $gv1,$is facilitated by all major insulin signaling pathways in the model; RABGTP, the $PKBd$pathway and the plasma membrane located pathway $PMR$. The $gv2$ variable is defined as

$$gv2=gk2bas\cdot C0\cdot RABGTP$$

(Eq.53)

where, $gk2bas$ is the rate parameter governing the GLUT4 transport to membrane bound GLUT4 clusters. The transport is dependent on the active form of RAB-GTP$.$ The model states describing membrane bound GLUT4 represent the level of GLUT4 protein in monomers (C1) or clusters (C2). The C1 derivative can be written as

$\frac{dC1}{dt}=gv1-gvc+gvr$	(Eq.54)

$$C1\left( 0 \right)= 0.1 [A. u]$$

where $gvc$ represents the transport of GLUT4 from monomer GLUT4 into clusters and $gvr$ which represents the conversion from GLUT4 clusters into monomers. The transport rate in $gvc$ is dependent on the rate parameter $gkc$, as well as the $C1$ and $C2$, and is written as

$$gvc=gkc\cdot C1\cdot C2$$

(Eq.55)

The transport rate in $gvr$is dependent on the rate parameter $gkr$, as well as the $C2$ and $PMRa$, and is written as

$$gvr=gkr\cdot C2\cdot PMRa$$

(Eq.56)

Here the transport is dependent on active PMR pathway as well as the amount of GLUT4 clusters readily available. The ODE for membrane bound GLUT4 cluster, *C2*, can be written as

$$\frac{dC2}{dt}=gv2+gvc-gvr-gve$$

(Eq.57)

$$C2\left( 0 \right)= 0.1 [A. u]$$

Where, $gve$ represents the translocation of GLUT4 cluster back to the endosome storage and the three first terms have already been previously described (Eq.49, 51, and 52). The $gve$ translocation is dependent on the amount available cluster and is also inhibited by the PMR pathway. The *gve* can be written as:

$gve=C2\cdot\frac{gke}{1+PMRa\cdot gn1}$	(Eq.58)

where, the parameter $gn1$ governs the inhibition made by the PMR pathway and the transport rate is governed by the $gke$ parameter.

Using the model states describing the active membrane bound forms (C1 and C2) of GLUT4, the adipocyte glucose uptake ($u_{a})$ is described in the following way:

$u_{a}=\frac{G_{c}}{V_{G}}\cdot\frac{FM}{FM_{0}}\cdot kGLUTprop\cdot(C1+C2)$	(Eq.59)

where, the parameter $kGLUTprop$ scales the value of GLUT4, so that the computed value of $u_{a}$is in a realistic range for the adipose glucose uptake. The scaling is needed because the intracellular model is trained by *in vitro* data from isolated adipocytes. Note that the adipose tissue glucose uptake also is dependent on the concentration of glucose in plasma ($\frac{G_{c}}{V_{G}}$) as well as the change in fat mass ($FM$) (which are specified further in Sections I and II above).

Finally, some comments on the model parameter values for the intracellular model. The parameters shown in Table 5, were taken from Bergqvist *et al.* and were kept constant and no included in the parameter estimation. This was done to keep the model dynamics in line with what was presented in Bergqvist *et al.,* and also because the lack of intracellular data included in our dataset. However, Bergqvist *et al.* trained the model using data of primary adipocytes from rats (our data is on mice), also for much larger doses of insulin compared to our data. To account for these differences, we included the parameter governing the phosphorylation rate of the IR, *ik1,* in the parameter, values are shown in Table 6.

**Table 3*:*** *Table showing the model parameters for the intracellular insulin signaling model from Bergqvist et al. The parameters listed under constants was not included in the parameter estimation and the values were kept the same as in Bergqvist etl al. (2017). The parameters listed in estimation were included in the optimization. The presented parameter values correspond to the parameter-set that yielded the simulation with the lowest cost.*

| Constants | | |
| --- | --- | --- |
| *Name* | **Value** | **Unit** |
| *ik1basal* | 0.0043 | ${min}^{-1}$ |
| *ik2* | 1.9332 | ${min}^{-1}$ |
| *ikR* | 0.7932 | ${min}^{-1}$ |
| *ik3* | 0.1243 | ${min}^{-1}$ |
| *ikm3* | 19.6444 | ${min}^{-1}$ |
| *ik4* | 79.0576 | ${min}^{-1}$ |
| *ikm4* | 1.1018 | ${min}^{-1}$ |
| *ikf* | 73.8741 | ${min}^{-1}$ |
| *kpI1t2* | 7.4089 | ${min}^{-1}$ |
| *kpI2t2* | 39.6374 | ${min}^{-1}$ |
| *kpi1* | 50.3746 | ${min}^{-1}$ |
| *kpi2* | 3.7679 | ${min}^{-1}$ |
| *kpd1* | 2.7857 | ${min}^{-1}$ |
| *kpd2* | 0.5348 | ${min}^{-1}$ |
| *k4731* | 67.1693 | ${min}^{-1}$ |
| *k3081* | 6.2902 | ${min}^{-1}$ |
| *k3082* | 78.8825 | ${min}^{-1}$ |
| *k43* | 42.8235 | ${min}^{-1}$ |
| *k473308* | 73.2418 | ${min}^{-1}$ |
| *k308473* | 79.5252 | ${min}^{-1}$ |
| *k4732* | 30.8146 | ${min}^{-1}$ |
| *k34* | 64.2238 | ${min}^{-1}$ |
| *as1* | 57.3653 | ${min}^{-1}$ |
| *as2* | 33.3260 | ${min}^{-1}$ |
| *gkr* | 0.4528 | ${min}^{-1}$ |
| *gk3* | 0.2919 | ${min}^{-1}$ |
| *gke* | 1.3427 | ${min}^{-1}$ |
| *gkc* | 1.4054 | ${min}^{-1}$ |
| *gk1bas* | 43.2974 | ${min}^{-1}$ |
| *gk2bas* | 3.8534 | ${min}^{-1}$ |
| *gn1* | 5.4502 | ${min}^{-1}$ |
| *gka1* | 0.0818 | ${min}^{-1}$ |
| *gka2* | 24.4059 | ${min}^{-1}$ |
| *kmt1* | 23.6746 | ${min}^{-1}$ |
| *kmt2* | 24.9801 | ${min}^{-1}$ |
| *gkl1* | 70.3705 | ${min}^{-1}$ |
| *gkl2* | 2.9805 | ${min}^{-1}$ |
| Estimated parameters | | |
| Name | $\boldsymbol{\theta}_{\mathbf{best}}$ | **Unit** |
| *kbasalIRS* | 627.3036 | ${min}^{-1}$ |
| *kGluProp* | 0.0047 |  |
| *ik1* | 2.1541e-4 | ${min}^{-1}$ |

To allow for some freedom in the model adipose glucose uptake (Eq. 59), we also included the scaling parameter *kGluProp* into the parameter estimation. Also, the newly added parameter determining IRS expression, *kbasalIRS,* was included in the estimation. Most of the equations so far have come more or less directly from existing models, except for a few additions concerning e.g. the impact of insulin resistance. Let us now look at the equations for the insulin resistance as such.

**IV. The new insulin resistance model**

To describe the effects of insulin resistance in the system, five scaling-functions were created to either increase or decrease key rates in the model. The input to the scaling functions is a new variable ($x_{IR}$) that combines the relative increase in fat-mass $\frac{FM}{FM_{0}}$. with the time spent on HFD. The time spent on HFD is described using the model state $t_{HFD}$, written as:

$\frac{dt_{HFD}}{dt}=\left\{ \begin{aligned} cf>0.7\to0 \\ cf<0.7\to1 \end{aligned} t_{HFD}\left( 0 \right)=0 \right.$	(Eq.61)

where $cf$is an input constant describing the carbohydrate fraction of the energy intake. For the chow diets, $cf$ was set to 0.7 and for HFD, $cf$, was set to 0.2, thus $t_{HFD}$ only increased when the diet was defined as an HFD. The new variable, $x_{IR}$, is defined as

$x_{IR}=\frac{FM}{FM_{0}}\cdot t_{HFD}$	(Eq.62)

As mentioned, the new variable $x_{IR}$ was used as an input for the insulin resistance functions. One of the functions is a linear regression function (Eq. 63a) and four are log-linear regression functions (Eq. 63b), written as:

$f\left( x_{IR} \right)=1+k\cdot x_{IR}$	*where,* $x_{IR}=t_{HFD}\cdot\frac{FM}{FM_{0}}$	(Eq.63a)

$f\left( x_{IR} \right)=1+k\cdot log(x_{IR})$	*where,* $x_{IR}=t_{HFD}\cdot\frac{FM}{FM_{0}}$	(Eq.63b)

where *k* is the slope that was estimated as a free parameter allowing for adjustment of each scaling function to the insulin resistance in data. We will now detail each of the affected rates in our phenomenological description of insulin resistance progression.

*a) Endogenous glucose production*

The EGP rate was allowed to change with increasing insulin resistance. This was done to be able to phenomenologically describe the higher baseline level of fasting glucose seen in our data (Figure 5A-D). The scaling function ${f_{IR}}_{EGP}$ describes the increase in the glucose production with developing insulin resistance and is given by

${f_{IR}}_{EGP}(x)=1+k_{EGP}\cdot log(x_{IR}),$	(Eq.64)

where $k_{EGP}$ represents the slope and $x_{IR}$ is the function input. Here we use a log-regression function to allow for a sharper increase in the fasting plasma glucose levels during the first days of HFD, which we see in our data (Fig 5A-D). The slope, $k_{EGP}$, determines how fast and how much the EGP is affected during the progression of insulin resistance.

*b) Insulin secretion*

The second rate that was allowed to change with insulin resistance was the basal insulin secretion (Eq. 20). This was done to be able to describe the high fasting plasma insulin levels seen in our data (Figure 4D). Hyperinsulinemia has been shown to be connected to insulin resistance and beta-cell dysfunction. In mice, increased insulin secretion has been correlated to insulin resistance and HFD (7). This is also seen in our own data (Fig 4C). The scaling function used to describe phenomenologically is a linear regression function.

${f_{InsRes}}_{I_{sec}}(x)=1+k_{Isec}\cdot\log\left( x_{IR} \right),$	(Eq. 65)

were the slope, $k_{Isec}$ governs the increased effect of insulin resistance on the rate of insulin secretion with prolonged HFD and $x_{IR}$ is the function input.

*c) Hepatic and muscle glucose utilization*

The third rate impacted by insulin resistance is the insulin dependent glucose utilization, UID. Insulin resistance is characterized by the inability of tissue (muscle and adipose) to properly respond to insulin stimulation (8). This is one of the main mechanisms behind hyperglycemia. In mice, HFD is correlated with lower levels of skeletal muscle glucose uptake (9). The hepatic glucose uptake is indirectly stimulated by insulin. However, it has been shown that hepatic glucose uptake is lowered in individuals with T2D (10). In the model, we describe the tissue (muscle and hepatic) dysfunction by gradually lowering the glucose uptake rate to phenomenologically describe the effects of increasing insulin resistance. For the muscle and hepatic tissue, we use the same scaling function.

$f_{IR_{CLGI}}(x)=1+k_{CLGI}\cdot x_{IR}$	(Eq.66)

Here,$k_{CLGI}$ is the slope and $x_{IR}$ is the function input. We use a linear-regression function to allow for a gradual change in plasma glucose dynamics and fasting levels, which we see in our data (Fig 5 A-D). The slope $k_{CLGI}$ governs the effect of insulin resistance on the glucose uptake from muscle and hepatic tissue. To describe the effects of insulin resistance on adipose tissue we had to modify the insulin intracellular signal model.

*d) Adipose glucose utilization - IRS and PMR*

The last two modifications to account for lower adipose glucose utilization following insulin resistance. Two key signaling intermediates, directly downstream IR, were affected by separate scaling functions with the goal to phenomenologically described the blunted signaling. The first affected rate is the production of IRS (Eq. 27-28) which was scaled with the log-linear-regression function$:$

${f_{InsRes}}_{IRS}\left( x \right)=1+k_{IRS}\cdot\log\left( x_{IR} \right)$	(Eq.67)

Here,$k_{IRS}$ is the slope and $x_{IR}$ is the function input. Data from Hansson *et al.* (11), show that the total IRS1 expression is decreased with prolonged durations of HFD and increased insulin resistance, which could contribute to blunted insulin signaling. To describe this in the model we added a production and degradation of the IRS protein in the adipocyte insulin signaling model (Eq. 27-28). Using the scaling function, the production of IRS is downregulated with increase in insulin resistance. Also, the mTORC2 model state was moved to be downstream IRS (Eq. 27-28), and to our knowledge this is not in conflict with any known data.

The second scaling function, ${f_{InsRes}}_{PMA}$, (Eq. 30) affecting adipose glucose utilization is implemented on the other main arm of insulin signaling in the model of the PMR pathway to blunt the GLUT4 translocation (Eq. 48-49). The scaling function is written as:

${f_{InsRes}}_{PMR}(x)=1+k_{PMA}\cdot\log\left( x\_IR \right)$	(Eq.68)

where,$k_{IRS}$ is the slope and $x_{IR}$ is the function input. Being on a prolonged HFD will result in a decrease of plasma membrane activation, and subsequent decrease of GLUT4 translocation and adipose glucose uptake (Eq. 59).

Finally, some comments on the model parameter values for the insulin resistance model. All the slope parameters were included in the parameter estimation. As this was new parameters in the model the parameter start guess (*θ_0_*) was arbitrary. In Table 4 we show the parameter set which yielded simulation corresponding the lowest cost-value.

**Table 4:** *The estimated parameter values for the insulin resistance functions. These values come from the parameter-set which produced the simulation corresponding to the lowest cost values for all training data. These parameters are dimensionless.*

| Name | $\boldsymbol{\theta}_{\mathbf{best}}$ |
| --- | --- |
| $\boldsymbol{k}_{\boldsymbol{EGP}}$ | 1.3599 |
| $\boldsymbol{k}_{\boldsymbol{Isec}}$ | 0.5150 |
| $\boldsymbol{k}_{\boldsymbol{CLGI}}$ | 0.3147 |
| $\boldsymbol{k}_{\boldsymbol{IRS}}$ | 0.6423 |
| $\boldsymbol{k}_{\boldsymbol{PMA}}$ | 99.82 |

**V. Implementation of HFD-induced T2D response**

The final new thing added to the model, was an ability to describe the human T2D response in our resulting *in silico* mouse model. To this end, we made some model variables capable of inducing loss in beta-cell mass. This was done by adding variables representing build-up of beta-cell damage and failure and their effect on insulin secretion. The model states responsible for inducing beta-cell failure were pancreatic stress, $P_{stress}$ and the subsequent beta-cell failure, $Beta_{failure}$. The $P_{stress}$ ODE is defined as:

$\frac{dP_{stress}}{dt} =\left\{ \begin{aligned} G_{c}<T_{fi}\to0 \\ G_{c}>T_{fi}\to kp_{stress} \end{aligned} P_{stress}\left( 0 \right)=0 \right.$	(Eq.69)

where, $T_{fi}$ is a constant representing the set threshold for plasma glucose for when pancreatic stress begins and $kp_{stress}$ is the constant rate at which the $P_{stress}$ derivative will increase after the plasma glucose reach the set threshold. The beta-cell failure is dependent on the $P_{stress}$ state to reach a certain value, which is represented by the constant $T_{PNR}$ (threshold representing the point of no return into T2D). When this happens the ${Beta}_{failure}$ derivative will start to increase with a constant set by the ${kbeta}_{failure}$ parameter. The ${Beta}_{failure}$ ODE is written as:

$\frac{d{Beta}_{failure}}{dt} =\left\{ \begin{aligned} P_{stress}<T_{PNR}\to0 \\ P_{stress}>T_{PNR}\to{kbeta}_{failure} \end{aligned} {Beta}_{failure}\left( 0 \right)=0 \right.$	(Eq.70)

The cumulative effect of both these model states is denoted the $T2D_{effect}$which is defined as:

${T2D}_{effect}=Beta_{failure}\cdot P_{stress}$	(Eq.71)

Let us now go through these equations. We used the fasting plasma glucose level as the main driving force behind the increase in the pancreatic stress variable, $P_{Stress}$ (Eq. 67). Here, $T_{fi}$ is the threshold value of fasting glucose when the system is deemed to have reached a sufficient level of insulin resistance to progress into T2D, and $P_{Stress}$ starts to increase. In isolation, increasing $P_{stress}$ only results in a blunted insulin secretion. To get a decline in insulin secretion, representing loss in beta-cell mass, the ${Beta}_{failure}$ state is also needed. When $P_{stress}$ reaches the threshold value of $T_{PNR}$, representing the point of no return in T2D development, the $Beta_{failure}$ starts to increase. The two variables, $P_{Stress}$ and ${Beta}_{failure}$, form the factor denoted the ${T2D}_{effect}$. This variable is then inserted into Eq. 20 to inhibit the fasting insulin secretion.

Finally, some comments on the model parameters. The parameters for this part of the model were manually changed to tune the T2D progression to produce the simulations shown in Fig. 6. The parameters values used can be seen in Table 5. Note that these parameters can be set to simulate any sought-after behavior in the T2D progression.

**Table 5:** *Parameters for the HFD-induced T2D response in the in silico humanoid mouse. These values were manually tuned to produce the desired T2D, and they can be set to produce any sought-after progression into the T2D state.*

| Name | Value | Unit |
| --- | --- | --- |
| $\boldsymbol{T}_{\boldsymbol{fi}}$ | 7.5 | $mmol$ |
| $\boldsymbol{k}\boldsymbol{p}_{\boldsymbol{sress}}$ | 0.15 | ${day}^{-1}$ |
| $\boldsymbol{T}_{\boldsymbol{PNR}}$ | 1.5 | - |
| $\boldsymbol{kbeta}_{\boldsymbol{failure}}$ | 0.25 | ${day}^{-1}$ |

### **2. *Adhoc* requirements for glucose uptake in the objective function**

To not allow any unphysiological behavior, *adhoc* requirements were added to the objective function to limit the behavior of the glucose uptake. These *adhoc* requirements were written to limit the possibility of finding solutions (parameter-sets) that would yield unsatisfactory qualitative behavior, defined as: 1) the insulin dependent uptake being smaller than the insulin independent uptake during IPGTT’s 2) the insulin independent uptake being unphysiologically small, 3) limit the size of the adipose tissue uptake (relative to the other tissues) and 4) the adipose tissue uptake can’t be unphysiologically small. These *adhoc* requirements were added to three simulations: IPGTT after two-week chow (Fig 4A), IPGTT after two-week HFD (Fig 4D), as well as IPGTT after six-week HFD. In practice this was done by calculating the area under curve (AUC) for each glucose uptake, which was then compared relative to each other. These requirements were written as four statements which was enforced with punishment to the cost-value if broken. The requirements regarding the insulin dependent and independent uptake were written as:

1) $Adhoc_{Uid}=if AUC\left( Uid \right)< AUC\left( Uii \right)$	$cost_{adhoc}=cost_{achoc}+max(0,AUC\left( Uii \right)- AUC\left( Uid \right) )+\mathcal{F}_{\chi2}^{\mathrm{cdf}-inv}\left( 0.95,35 \right)$

2) $Adhoc_{Uii}=if 0.05\cdot AUC\left( Uid \right)> AUC\left( Uii \right)$

The requirements regarding the adipose tissue glucose uptake was written as:

3) $Adhoc_{UidA\_high}=if 0.15\cdot AUC\left( UidM \right)< AUC\left( UidA \right)$

4) $Adhoc_{UidA\_low}=if 0.05\cdot AUC\left( UidM \right)> AUC\left( UidA \right)$

In Fig SA1 we show the corresponding uptake for all threes simulations.

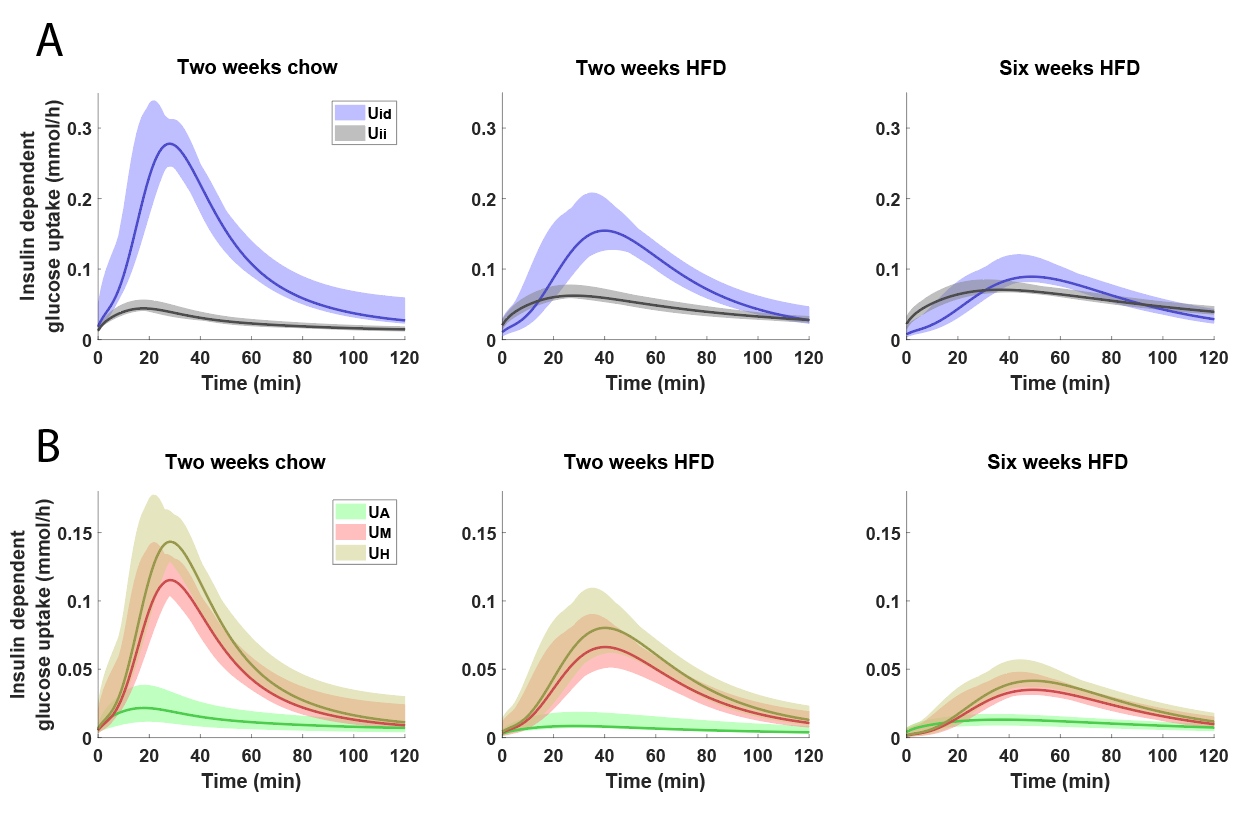

***Figure SA1:*** *Simulations concerning various aspects of the glucose uptake during an IPGTT at the end of the three different diet schemes: two weeks chow (left), two weeks HFD (middle), and six weeks HFD (right).* ***A)*** *Model simulations (lines) with uncertainty (shaded areas) shown for the insulin dependent glucose uptake (blue) and insulin independent glucose uptake (grey).* ***B)*** *Model simulations (lines) with uncertainty (shaded areas) shown for the insulin dependent glucose uptake in each tissue; adipose- (light green), muscle- (red) and hepatic-tissue (brown).*

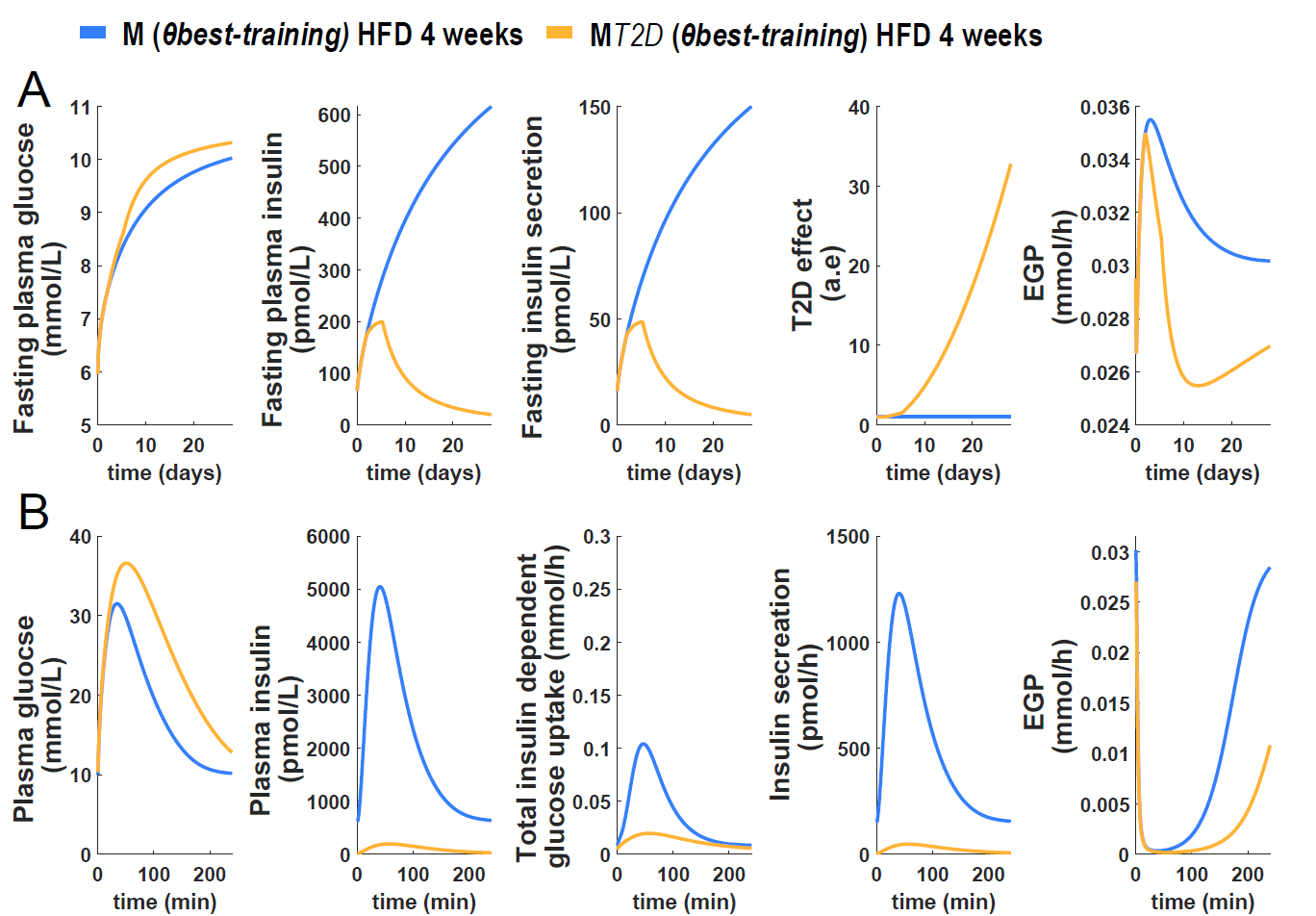

***Figure SA2:*** *Model simulation for all states relevant to the progression of insulin resistance into T2D, and the subsequent short-term response during an IPGTT.* *These simulations correspond to Fig 6 in the main file and shows more model states.* ***A)*** *Simulations showing the long-term changes in several model states: plasma glucose, plasma insulin, insulin secretion, and EGP. Also shown is the change in the variable “T2D effect” which is the sum of the pancreatic stress and the beta-cell failure variables.* ***B)*** *Simulations showing the short-term response to an IPGTT in several states as predicted in the end of the HFD scheme.*
